# Memory by a thousand rules: Automated discovery of multi-type plasticity rules reveals variety & degeneracy at the heart of learning

**DOI:** 10.1101/2025.05.28.656584

**Authors:** Basile Confavreux, Zoe P. M. Harrington, Maciej Kania, Poornima Ramesh, Anastasia N. Krouglova, Panos A. Bozelos, Jakob H. Macke, Andrew M. Saxe, Pedro J. Gonçalves, Tim P. Vogels

## Abstract

Synaptic plasticity is the basis of learning and memory, but the link between synaptic changes and neural function remains elusive. Here, we used automated search algorithms to obtain thousands of strikingly diverse quadruplets of excitatory(E)-to-E, E-to-inhibitory(I), IE, and II plasticity rules, cooperating to stabilize recurrent spiking networks. Despite the fact that quadruplets were selected for homeostasis, more than 90% of them performed well in simple and more difficult memory tasks such as novelty detection, contextual novelty and sequence replay. Co-activity was crucial, i.e., most rules failed in isolation. Our purely local, unsupervised plasticity rules could also help solve computer games such as pong. Our work showcases automated discovery augmenting human intuition to find en masse solutions for high dimensional problems.

## I. Introduction

How do millions of heterogeneous synapses coordinate to form memories? Direct measurements of multiple synapses remain most difficult^1,2^, and theoretical work has focused on single-rule models^3–8, c.f.9–12^, leaving the joint impact of multiple concurrent rules largely uncharted. Indeed, synaptic plasticity rules—typically expressed as weight changes dependent on pre- and post-synaptic activity—are difficult to tune by hand^9,10,13^, as they comprise many parameters and their solution spaces become unruly very quickly. Meta-learning approaches, in which plasticity rules are generated and evaluated by automated methods, have been gaining traction to explore more systematically multiple coactive rules^12,14–26^, but only for small networks or simple functions. Here, we used simulation-based inference^12,27^ to meta-learn thousands of co-active quadruplets of plasticity rules that produce stable and biologically plausible network activity. Strikingly, most of these rule quadruplets also supported various forms of memory acquisition when stimulated accordingly. Without fine-tuning, we could identify families of rule quadruplets for various experimentally observed behaviors such as familiarity—or novelty—detection, contextual novelty, and replay, with memory lifetimes from seconds to hours. In some, but not all cases, rule quadruplets with similar network functions had notable trends in their plasticity parameters, hinting at what features of each learning rule bestowed a particular function. We also found that most individual rules were unstable in isolation, and only functioned successfully as part of their quadruplet, providing a possible explanation for why plasticity is so difficult to probe experimentally. Finally, we showed how these unsupervised, co-active rules could be a stepping stone towards more complex computations, such as helping a computer model play the game of pong.

## II. Results

### Meta-learning co-active plasticity rules at scale

Recent work uncovered a space of co-active EE, EI, IE and II plasticity rule quadruplets which could produce stable and robust activity for a few minutes in recurrent spiking networks of excitatory and inhibitory neurons^12^ (Fig. 1A). We wondered if these rules could support neural functions beyond stability. To investigate, we considered such networks in which synapses from neuron *i* to *j* underwent spike-timing dependent plasticity, STDP^1,3^, with additional pre- or post-only updates^12^:

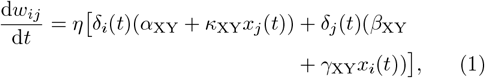

where *η* is the learning rate and variables *x*_*i*_(*t*) and *x*_*j*_(*t*) are low-pass filters of the pre- and post-synaptic spike trains with time constants 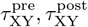, with (X, Y) ∈ {E, I}. Plasticity updates at each synapse type depended on 6 parameters 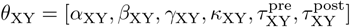, resulting in 24 plasticity parameters across the four synapse types. We visualized plasticity rule quadruplets using a classic representation of weight changes as a function of the time lag between a pre- and a post-synaptic spike^1–3^ (Fig. 1B).

**Fig. 1.**
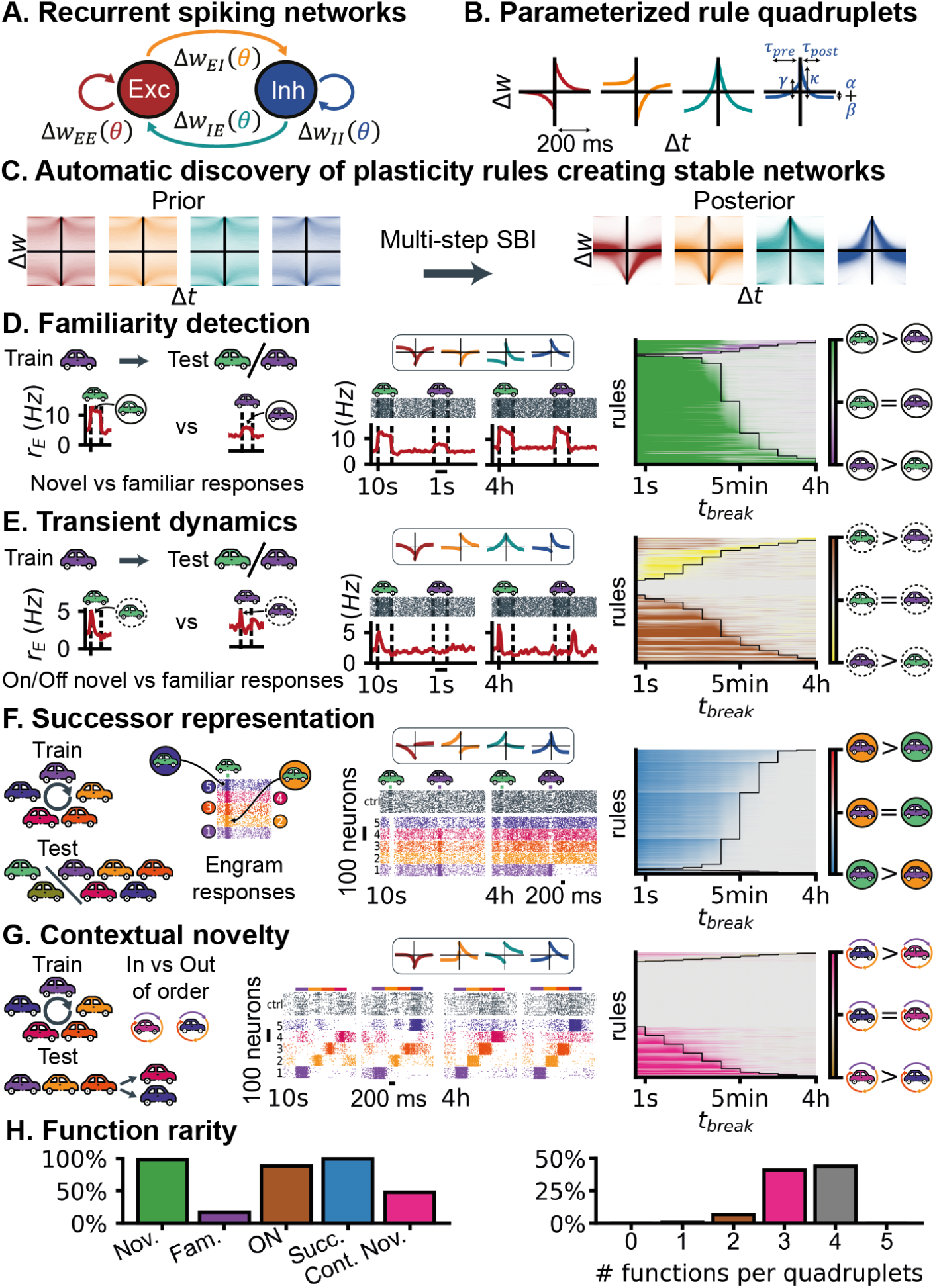
Rule quadruplets meta-learned for stability elicit memories. **A**: Recurrent spiking network with 4096 excitatory and 1024 inhibitory neurons. **B**: Each recurrent synapse type (EE, EI, IE and II) has its own, STDP-like, plasticity rule. Weight change as a function of the time-lag between a pre- and a post-synaptic spike. **C**: Obtaining the stability manifold starting from a uniform prior. **D**: Familiarity detection task. Middle: example rule quadruplet in this task. Top: pre-post protocol of the rule quadruplet, like in B. Bottom: raster plot of 250 excitatory neurons, and population firing rate. Dotted vertical lines denote onset and offset of the stimuli presentations. Right: evaluation of network function in the familiarity task for each meta-learned rule tested. The black line denotes the last timestep at which the rule quadruplet displayed a significant difference between novel and familiar. **E**: Same as D but now reading out memories from ON responses. **F**: Sequence learning task: a network is trained for 50s on a loop of 5 stimuli. During testing, each training stimulus is presented independently, as well as two novel stimuli. All stimuli are non-overlapping. **G**: Contextual novelty task, comparing network population responses when familiar stimuli are presented in vs out of order. **H**: Left: percentage of rule quadruplets from the 2,500 sampled from the stability manifold that have significant responses for least at one break time in the task shown above. Right: distribution of the number of functions per rule quadruplet.

To meta-learn plasticity rules, we combined filter-based simulation-based inference, fSBI^12^, with a multi-fidelity simulation-based inference approach^27^. This combination of methods dramatically reduced the compute required to explore the 24-dimensional plasticity parameter space and allowed us to numerically infer a manifold of varied co-active learning rules with hours-long stability, the stability manifold (Fig. 1C and Fig. Supp. 1).

### Memory as a byproduct of network stabilization

We sampled 2,500 rule quadruplets from the stability manifold and embedded them in networks performing a range of memory tasks. We began with a familiarity detection task that consisted of a single 10s high-frequency input stimulus to a subset of excitatory neurons, followed by a break period—during which the network received only background inputs—ranging from 1s to 4h (Fig. 1D, left). After the break, we compared the network response to the familiar stimulus and to a novel stimulus (Fig. 1D, middle).

Immediately after the initial stimulus, we observed significant differences (Student t-test, *p <* 0.05) between the response to novel and familiar stimuli for 94% of the sampled quadruplets, i.e., a memory of the initial stimulus was expressed. Memory lifetime, the time until a memory could not be recalled with significant firing rate differences, varied across rule quadruplets. After a 4h break, the fraction of networks still expressing a memory dropped to 7% (Fig. 1D).

When considering only transient responses to stimulus onset and offset (so-called “ON” or “OFF” responses^28^), we found that some quadruplets created networks with rich transient dynamics that were absent from the naïve networks (Fig. Supp. 5). In some cases, these transient responses were significantly different for novel and familiar stimuli, and could serve as a readout signal for familiarity or novelty (Fig. 1E).

We also played looping sequences of 5 different inputs that stimulated 5 non-overlapping groups of excitatory neurons in the network (Fig. 1F). After the break period, we tested if the presentation of any one stimulus would elicit “successor representations” in the other, non-stimulated groups. 98% of meta-learned rules produced various signatures of sequence learning after 10s, gradually dropping to 3% after 4h (Fig. 1F).

We tested for contextual novelty—also called omission novelty^29^, or error prediction in predictive coding frameworks^30^—by presenting a set of stimuli out of order compared to the training sequence (Fig. 1G). This task was more difficult than regular familiarity detection as all test stimuli were encountered during training, only their ordering during the test phase was novel or familiar. 23% of meta-learned rules produced networks that could detect this more subtle form of novelty after 10s, and memories were more short-lived across the board (Fig. 1G).

Our meta-learning approach generated thousands of co-active, unsupervised rule quadruplets producing complex network functions despite having been selected solely for network stability. Only one of the 2,500 quadruplets sampled from the stability manifold did not respond to any memory task according to our predefined metrics (Methods, Fig. 1H).

### Experimental predictions

To discover plasticity rules consistent with more specific experimental observations, we zoomed in on sub-regions of the stability manifold by performing additional rounds of inference (see Methods). We chose several published datasets for which the underlying plasticity rules are unknown, or have been subject to previous theoretical predictions^2,29,31–35^.

We first focused on two subfamilies of rule quadruplets that reproduced the trends observed in two experimental studies on novelty^32^, and familiarity detection^31^ (Fig. 2B-D). For novelty detection, we considered rule quadruplets as plausible if they robustly produced stronger network responses for novel stimuli than for familiar stimuli, with a memory lifetime of at least 4h. Plausible novelty-detecting quadruplets revealed a striking diversity of shapes, i.e. a number of degenerate solutions (Fig. 2B). In contrast, familiarity detection was much less common within the stability manifold (as suggested by Fig. 1D). Interestingly, all quadruplets which satisfied the criteria of familiarity detection switched from novelty detection early in the simulation (Fig. 2B). No obvious commonalities could be observed in the shapes of all plausible familiar-detecting quadruplets.

**Fig. 2.**
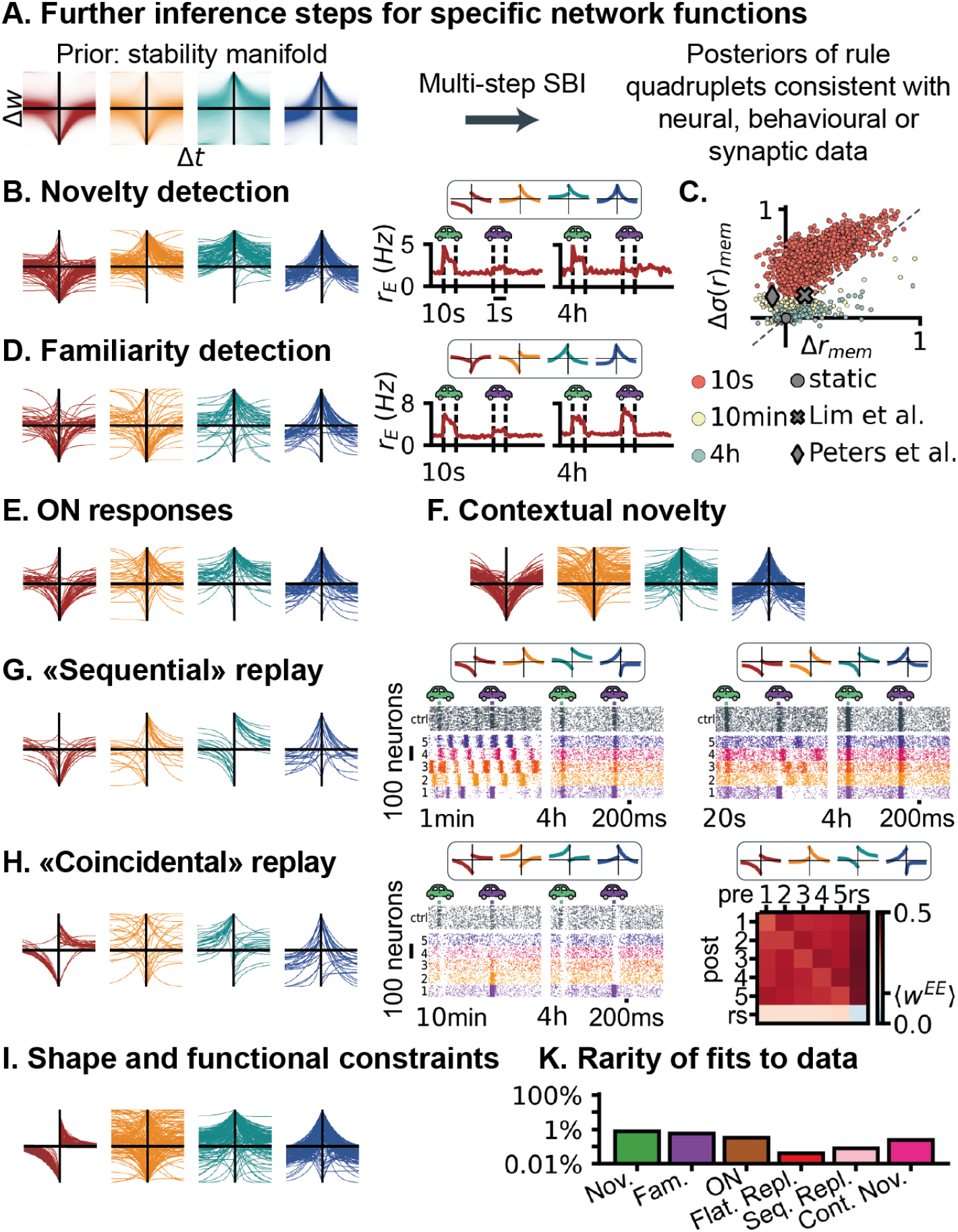
Isolating subfamilies of rule quadruplets consistent with data. **A**: Using the stability manifold as a prior, new posteriors with specific network functions are obtained with the same inference pipeline as in Fig. 1 (see methods). **B**: Left: Overlay of the shapes of quadruplets that robustly elicit long-term novelty detection, tested on 4h-simulations; Middle: Firing rate of an example quadruplet undergoing the familiarity task; Right: Response of the 2,500 meta-learned quadruplets used in Fig. 1 on the familiarity task, with difference in mean (x-axis) and variance (y-axis) in the network response to familiar and novel stimuli, static networks (with no plasticity), data from^31,32^. **C**-**H**: Same as B, but selecting for robust transient familiarity/transient contextual novelty/replay-like events/sequential memory without temporal unfolding of the memory at recall/robust ON responses (see methods for the criteria to define the function of quadruplets). Example quadruplets are shown, as well as the network dynamics that they elicit (similar format as Fig. 1F). Bottom right: EE matrix of a quadruplet exhibiting spontaneous replay. Weights are grouped by which engram their pre- and post-synaptic (excitatory) neurons belong to (see methods for engram definition). ‘rs’ stands for the rest of neurons not participating in any of the 5 familiar engrams. **I**: Selecting quadruplets based on the network function as well as their shape (condition on the plasticity parameter values). Here, selecting stable quadruplets with EE rules that are similar to the STDP rule^1–3^. **K**: Considering only the 2,500 quadruplets sampled from the stability manifold (and not from the subsequent posteriors with specific network functions), proportion of rules that exhibit each function shown above (see methods for selection criteria).

Plausible quadruplets for contextual novelty^29^ appeared to constrain the shape of the EE rule to symmetric anti-Hebbian rules, similar to what has been observed experimentally^36,37^ (Fig. 2F). Note however, that many stable quadruplets with symmetric anti-Hebbian EE rule did not elicit contextual novelty detection.

A few, rare quadruplets (*<* 0.1%, Fig. 2K) could imprint sequential memories into a network. We expanded on this subregion of the stability manifold to obtain two classes of solutions. Some quadruplets created networks with spontaneous or triggered, forward or backward sequential dynamics during the recall phase (Fig. 2G), reminiscent of hippocampal replay^33,35^. Other quadruplets could recall the training sequence without any sequential, replay information (Fig. 2H, “coincidental” replay). Quadruplets from each class could be distinguished by their shapes: coincidental quadruplets comprised mostly asymmetric EE rules—reminiscent of STDP^2^—and asymmetric IE rules with pre-before-post potentiation (Fig. 2H). Conversely, replay rules consisted mostly of symmetric anti-Hebbian EE and asymmetric IE rules, in this case with post-before-pre potentiation (Fig. 2G).

Meta-learned rules could also be compared directly with published data from *ex vivo* patch clamp recordings. For example, the oft-assumed unstable^13,38^ STDP EE rule observed in cortex^1,2^ appeared stable in many quadruplet configurations (Fig. 2I), suggesting that stable Hebbian learning may have to depend on co-activity of plasticity rules across different synapse types.

### Illuminating plasticity-function relationships

We wanted to further investigate the inter-dependencies between co-active plasticity rules. We had observed that stable rule quadruplets could take a plethora of shapes (Fig. 3A), with no clear, linear, low-dimensional macro structure in the parameter space (Fig. 3B, and Supplementary Material).

**Fig. 3.**
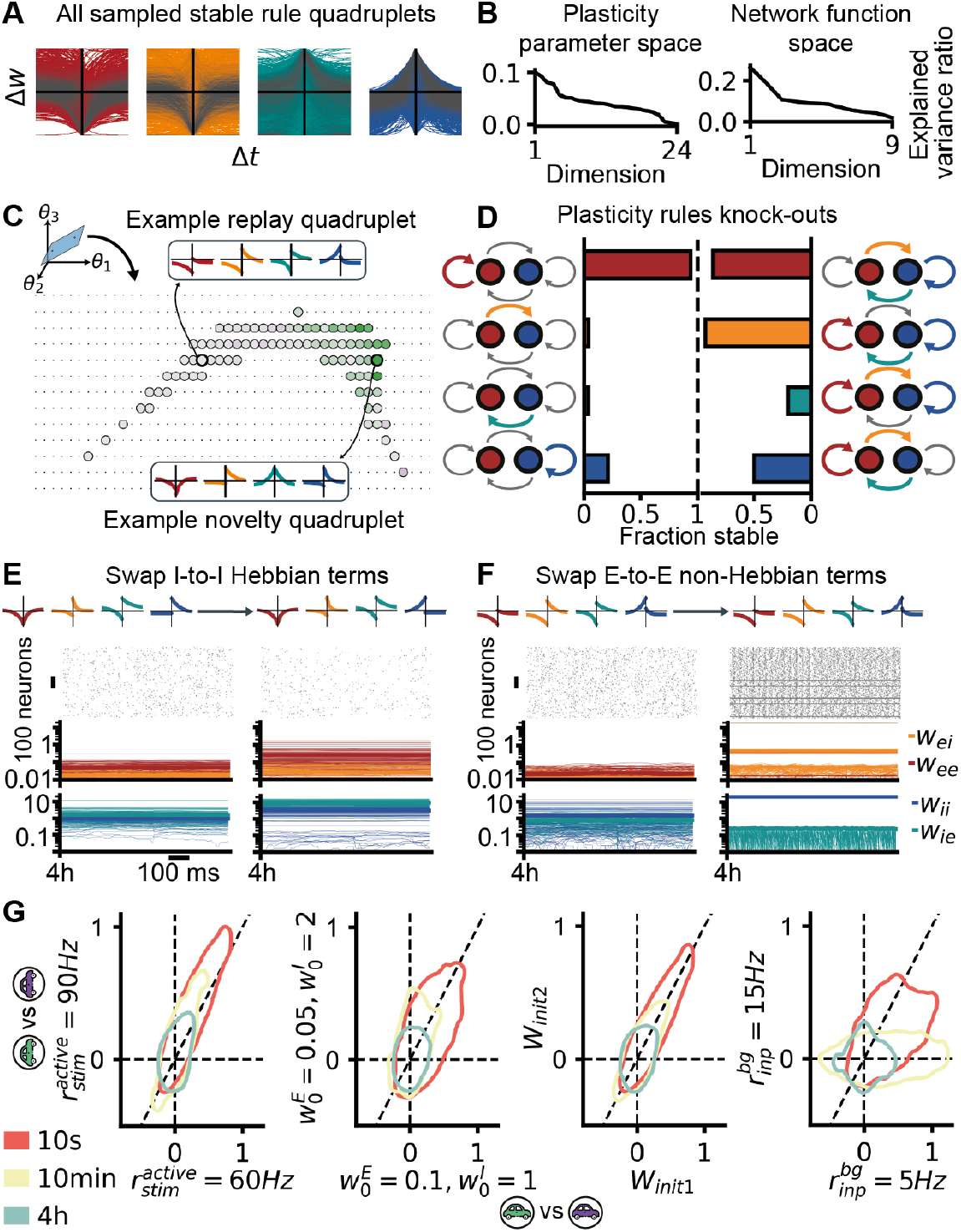
Analysis of the stability manifold. **A**: Histogram of the shapes of the 2,500 rule quadruplets stable for at least 4h used in Fig. 1. **B**: Left: Principal Component Analysis (PCA) performed on the plasticity parameters from the quadruplets shown in A, variance explained by each dimension. Right: PCA performed on the network responses elicited by the quadruplets in the tasks shown in Fig. 1. **C**: Two quadruplets from the stability manifold (shown in Fig. 1E and Fig. 2G) in plasticity parameter space shown with strong black borders. A chosen random plane containing the two quadruplets and simulated quadruplets sampled across this plane. Small black dots denote unstable quadruplets. Larger dots denote stable quadruplets and the color represents their response after 10 minutes on the familiarity task (same color bar as Fig. 1D). **D**: Left: Considering the 2,500 stable quadruplets used in Fig.1, simulating networks with only one of the 4 rules active (“triple KO”), and computing the fraction of these networks that were stable. Right: Same when removing a single rule from the stable quadruplets (“single KO”). Black connections are static (KO), colored connections are plastic. **E**: Raster plot and evolution of 100 weights of each synapse type and their means in bold, for two rule quadruplets. The left one is stable (see Fig. 1G), the right one has the Hebbian terms of the II rule swapped (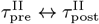 and 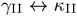) and has unrealistically high inhibitory weights. **F**: Same as B for two other quadruplets. the left one is stable (see Fig. 3), the right one has the non-Hebbian parameters of the EE rule swapped (*α*_EE_ *↔ β*_EE_, note this cannot be seen on the pre-post protocol plots). **G**: For each of the 2,500 stable quadruplets used in Fig.1, comparisons of two simulations with changes to one task parameter at a time. From left to right: Firing rate of input neurons that are part of a stimulus pattern while that pattern is active, initial network connectivity (strength of initialization and random initialization), initial network connectivity (random initialization seed only) and general tonic input firing rate received by all recurrent neurons in the network. The circles denote 50% of the mass, the color indicates different break times.

To visualize the micro structure of the stability manifold, we sampled rules from a plane with two stable quadruplets of dissimilar shape and functions, one quadruplet elicited transient replay (Fig. 3G), the other detected novelty for at least 4 hours (Fig. 3B). Our analysis revealed a likely-continuous, non-linear manifold surface of stable rule quadruplets, with smooth functional variations (Fig. 3C).

Next, we studied how individual rules contributed to the stability of a quadruplet. We compared the network dynamics and function of single rules in isolation (“triple KO”), or rule triplets with one deactivated rule in the quadruplet (“single KO”, see Methods and Fig. 3D). With the exception of many isolated EE rules, the majority of isolated rules proved unstable (Fig. 1D). Curiously, removing an unstable rule from a quadruplet often destabilized the remaining triplet of rules. For instance, many rule triplets without active (intrinsically unstable) IE or II plasticity failed to support stable network dynamics (Fig. 3D). Further, stable triplets and singlets exhibited altered and typically diminished function (Fig. Supp. 10). Our findings suggest that unstable rules can be “rescued”, and even utilized by co-activity, thereby broadening the space of stable parameter combinations compared to single rule cases.

Having established that the stability of rule quadruplets relied on complex parameter inter-dependencies across and within synapse types, we wondered whether mean-field theory—a rate-based description of spiking plasticity rules between independent neurons^39^— was able to capture these observations. We devised a perturbation to the rule quadruplets that would not affect their mean-field description: swapping the Hebbian parameters of one individual rule at a time: 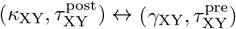 (see Fig. Supp. 12). This corresponds to swapping the response to pre-post vs. post-pre spike pairs. We found that such swaps destabilized the quadruplets around 40% of the time, typically because the weights of the affected rule would grow out of physiologically reasonable bounds within the 4h of simulation (Fig. 3E). We concluded that the simplified rate description of our quadruplets was not sufficient to predict stability.

Similarly to swapping terms that did not change a rule’s mean-field description, we could also swap non-Hebbian parameters *α* and *β* to explicitly change the mean-field description, but not the shape of the rule with regards to the pre-post protocol. Such swaps destabilized over 95% of tested quadruplets (Fig. 3F), highlighting the insufficiency of the pre-post protocol representation, as well.

We also tested the robustness of our quadruplets with regard to the parameters of the networks in which they acted (Fig. 3G). Most simulation parameters, such as stimulus strength and weight initialization (Fig. 2G) did not affect stability or function. Some, e.g., background drive, impacted the performance of most rules, sometimes in interesting ways (Fig. 4A). For example, one of our quadruplets only memorized familiar stimuli that were presented with low background rates, but not with high rates (Fig. 4A), reminiscent of previously reported effects of neuro-modulation on memory acquisition^40,41^.

**Fig. 4.**
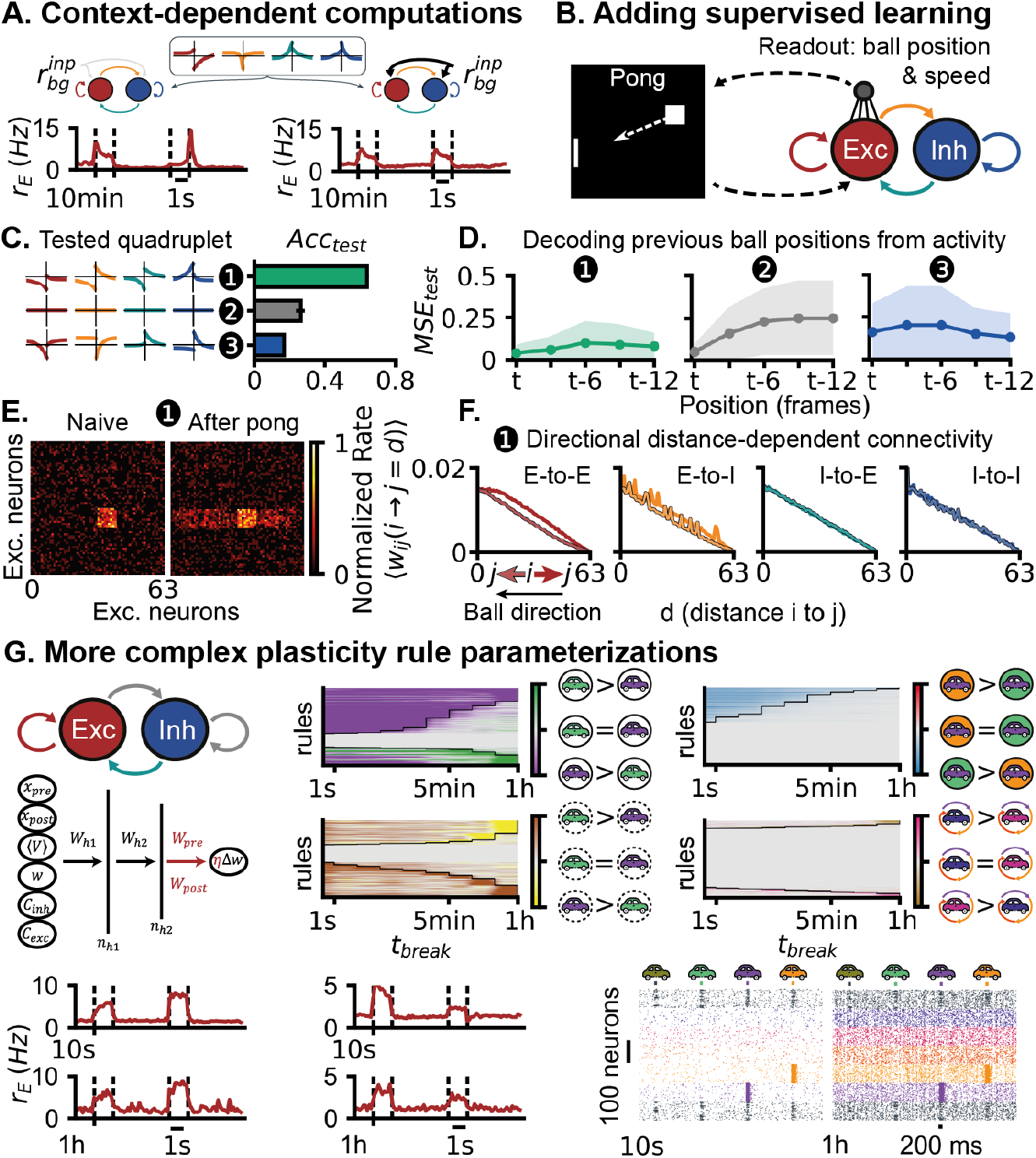
Beyond unsupervised learning in isolation: additional rules and network functions. **A**: Example quadruplet from the stability manifold with different responses on the familiarity task depending on the tonic input background firing rate (see Fig. 3G). **B**: Learning linear readouts on top of a plastic recurrent network playing pong. **C**: Validation set accuracy at pong for three rule quadruplets. Quadruplet 1 is shown in Fig. 2G and elicits sequential replay, quadruplet 2 corresponds to no plasticity (static network), quadruplet 3 elicits no function on any of the memory tasks. **D**: Given the network activity at time *t* (with the ball at position (*x, y*)), accuracy of three networks with the three rule quadruplets described above at predicting current and previous ball positions. **E**: Firing rates of recurrent excitatory neurons of the network evolving with quadruplet 1, during presentation of the same ball position before and after the network as been exposed to several pong sequences. **F**: Weight distribution of the network evolving with quadruplet 1 after minutes of exposure to pong. Synapses are sorted by their distance and direction along the x-axis (axis along which the ball moves). **G**: Same tasks as in Fig. 1, but considering different plasticity rule parameterization, MLP-based (see methods). Note that only the EE and IE synapses are plastic in this case. Bottom: Three example doublets, two undergoing the familiarity task and one the sequential task.

### More complex tasks and rules

The rules quadruplets we have meta-learned act purely locally. We wondered whether such local plasticity could help with learning supervised tasks. Towards this goal, we added readout units to the spiking network, whose inputs were trained using supervised learning^42,43^. We then assessed if the readout units performed better in combination with plastic (vs. static) upstream networks. First, we trained the readouts to report on familiar stimuli with a specified firing rate (Fig. Supp. 18). Plastic networks outperformed a static control network, as long as familiar and novel stimuli elicited different responses that could be mapped onto the desired output signal (Fig. Supp. 18). We then turned to a more difficult task, i.e., the Atari game of pong (Fig. 4B) that has been used to demonstrate learning capabilities of biological and artificial systems^44,45^. We tested if some quadruplets could produce networks from which the location and direction-of-motion of the ball could be inferred. Two quadruplets— those which had displayed sequential learning capabilities (Fig. 2G,H)—outperformed static networks substantially (Fig. 2G,H). Interestingly, these quadruplets created place-field-like plasticity (Fig. 4E,^46,47^), as well as directional, distance-dependent connectivity (i.e. aligned vs. against ball direction, Fig. 4F). Quadruplets that exhibited no significant responses in any memory task performed similar to, or worse than a static network (Fig. 4C,D).

Finally, we tested a higher-dimensional rule parameterization, in which synaptic changes in the spiking network were computed by a feedforward neural network, based on additional variables beyond spike pairs: post-synaptic membrane potential, activity at neighboring synapses, spike triplets and the synaptic weight^12^. Stable quadruplets from this manifold also performed successfully many of the above mentioned tasks, again as a byproduct of network stabilization (Fig. 4G). Unlike for the polynomial implementation, the most frequent function was transient familiarity detection (Fig. 4G) instead of novelty detection (Fig. 1D).

## III. Discussion

Our findings challenge the prevailing view that complex cognitive functions require intricately designed plasticity rules. Instead, we demonstrate that memory in its many forms may emerge spontaneously when simple plasticity rules work together to maintain network stability. Our discovery has three major implications: First, it explains why experimental studies of single plasticity rules often yield contradictory results. Second, it suggests that evolution may have discovered memory through the simpler route of ensuring stable neural dynamics. Third, it provides a new framework for understanding learning disorders as failures of plasticity coordination rather than single-rule deficits.

A key ingredient in stable and functional dynamics was co-activity (Fig. 3), by producing negative feedback cycles that counteracted the destabilizing tendencies of Hebbian learning and thus increased the size of the solution manifold. Such interactions might spell doom for the success of studying single rules in isolation, as has been the standard in the field. Our finding on co-activity also provides a new perspective on the computational advantages of the cell type diversity seen in the brain^48^ that may broaden the repertoire of possible network computations.

The plasticity rules we considered are likely an over-simplification of the mechanisms taking place in the brain. We did not consider slower mechanisms such as behavioral time scale plasticity^8^, homeostatic plasticity, consolidation and sleep. Different memory functions may appear more naturally for some types of plasticity rules than others (Fig. 4G). Moreover, plasticity rules themselves have been shown to change across time, for instance as a result of neuro-modulation^49,50^. The stable rule quadruplets discovered in this study can be thought as basic building blocks towards making more complex and realistic plasticity processes (Fig. 4A).

Though it is clear that the brain implements some form of reward signal for learning^51^, a good part of the brain’s ongoing plasticity may be unsupervised^52^. We provide *in silico* backing to this hypothesis, with complex network functions pertaining to memory emerging from simple, co-active and unsupervised rules. As our approach was limited to unsupervised learning—our plasticity rules did not incorporate error signals or top-down feedback—we added a readout trained with supervised learning and showed that we could use the network for reservoir computing^53,54^ to solve real-world tasks, such as pong (Fig. 4).

In summary, we have demonstrated the viability of algorithms for automated discovery of synaptic plasticity rules, and show that relatively simple, unsupervised rules can elicit complex network functions through cooperation. Our manifold may serve as an open resource for designing closed-loop plasticity experiments. More generally, our work suggests that simple parts can work together and interact with each other to produce something greater than the sum of their parts.

## IV. Methods

Unless mentioned otherwise, all networks in this study were simulated using Auryn^55^. All network simulations were run on the ISTA HPC cluster (913 cpu cores). The simulations for pong were run in python (Brian2 and Brian2CUDA^56,57^). All the code for running the networks and reproducing the analysis is available on Github https://github.com/VogelsLab/dataSBI. Numpy^58^, Pytorch^59^, Pandas^60^, Matplotlib^61^, Scipy^62^, sklearn^63^ and the sbi libraries^64^ were used for the analysis.

### Network model

Throughout the study, we considered recurrent spiking networks with *N*_E_ = 4096 excitatory neurons and *N*_I_ = 1024 inhibitory neurons (leaky-integrate and fire point neurons with variable threshold, AMPA and NMDA currents, and conductance-based synapses). These networks were similar to previous work^10,12^, except for the inputs that depended on the task considered. The membrane potential dynamics of neuron *j* (excitatory or inhibitory) followed:

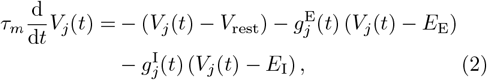

where E stands for excitation and I for inhibition, *τ*_*m*_ = 20 ms, *V*_rest_ = −70 mV, *E*_E_ = 0 mV and *E*_I_ = −80 mV.

A post-synaptic spike was emitted whenever the membrane potential *V*_*j*_(*t*) crossed a threshold 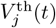, with an instantaneous reset to *V*_reset_ = −70 mV. This threshold 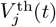 was incremented by 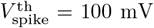 every time neuron *j* spiked and otherwise decayed following:

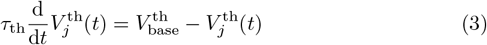

with 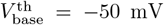. The excitatory and inhibitory conductances, *g*^E^ and *g*^I^ evolved such that

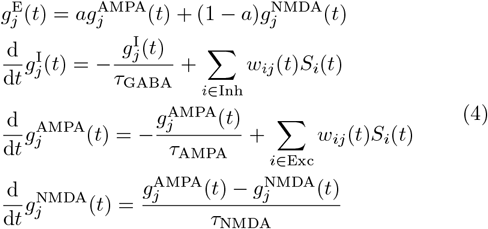

with *w*_*ij*_(*t*) the connection strength between neurons *i* and *j* (unitless), *a* = 0.23 (unitless), *τ*_GABA_ = 10 ms, *τ*_AMPA_ = 5 ms, *τ*_NMDA_ = 100 ms, 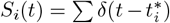 the spike train of pre-synaptic neuron *i*, where 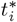 denotes the spike times of neuron *k*, and *δ* the Dirac delta.

Network initializations: Unless mentioned otherwise, all networks were initialized with random sparse connectivity (10%), with 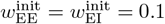 and 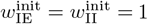.

The inputs received by the network depended on the exact task, which are described in subsequent sections.

### Rule quadruplets parameterization

The parameterization of synaptic plasticity, initially defined in^21^, captured first order Hebbian spike-triggered updates, the weight from neuron *i* of type X ∈ (E, I) to neuron *j* of type Y ∈ (E, I) evolved such that:

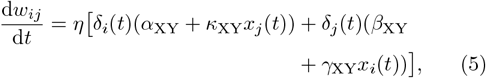

with *η* = 0.01 a fixed learning rate, 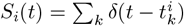 the spike train of neuron *i, δ* the Dirac delta function to denote the presence of a pre (post)-synaptic spike at time *t*. The synaptic traces *x*_*i*_ and *x*_*j*_ are low-pass filters of the activity of pre-synaptic neuron *i* and post-synaptic neuron *j*, with time constants *τ*_pre_ and *τ*_post_, such that:

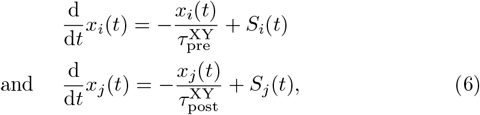

Overall, this search space comprised 6 tunable plasticity parameters per synapse type 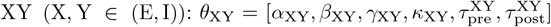, for a total of 24 plasticity parameters across all four synapse types.

Note that all weights in the network were capped at all times, in the [0, *w*_max_] range, with *w*_max_ = 20.

The MLP rules used in Fig. 4 were defined as in previous work^12^.

Note that there were no other plasticity mechanisms in these networks such as synaptic normalization, homeostatic plasticity or short-term plasticity.

### Inferring rule quadruplets: fSBI and MF-NPE

We got posterior distributions over plasticity rule quadruplets (parameterized with 24 parameters *θ* = [*θ*_EE_, *θ*_EI_, *θ*_IE_, *θ*_II_]; more details in section above) that created stable network dynamics for at least 4h, by combining two simulation-based inference (SBI) methods: filter Simulation-Based Inference (fSBI)^12^ and Multi-Fidelity Neural Posterior Estimation (MF-NPE)^27^. We proceeded in the following way:

1. Define the prior over rule quadruplets: In line with previous work^12^, the prior was a uniform distribution, such that *α*_XY_, *β*_XY_, *γ*_XY_, *κ*_XY_ ∼ 𝒰[−2, 2] and 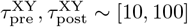ms.
2. Obtain 2min-stable quadruplets with fSBI: This step was performed in prior work^12^. In that study, fSBI was used to gradually filter out rule quadruplets that created unstable network dynamics after 2 minutes (rather than 4h). On this shorter task, rule quadruplets were deemed stable if they passed a number of conditions on the activity and weight dynamics^21^. The result was a neural density estimator that approximated a posterior over plasticity parameters that generated stable quadruplets with high probability (≈50%). In total, 110k stable quadruplets were obtained with this method. The stability of each quadruplet in practice was verified with numerical simulations.
3. Obtain 4h-stable quadruplets with MF-NPE: We used the 110k 2min-stable quadruplets to pre-train a posterior density estimator *p*(*θ*|*r*_exc_(2min)), with *r*_exc_(2min) the firing rate of the excitatory population after 2 minutes. We then fine-tuned the density estimator on a subset of these 110k quadruplets that led to stable networks for at least 4h, *p*(*θ*|*r*_exc_(4h)) (2,500 quadruplets, see Fig. Supp. 1). The 4h-stability criteria are defined below. Finally, we sampled 1,000 rule quadruplets from this posterior, conditioning on low firing rates (*r*_exc_(4h) ∼ 𝒰 [2 − 10]Hz). 49.2% of the posterior samples led to stable dynamics of the spiking networks when simulated for 4h.

#### Criteria for 4h-stability

In this study, a network was classified as stable in the 4h simulations if:

- Firing rate condition: *r*_exc_(*t*) ∈ [0.1, 100] for all *t <* 4h, with *r*_exc_(*t*) the firing rate of the excitatory population computed over a 10s window.
- Irregularity condition: CV_ISI_(*t*) *>* 0.7 for all *t <* 4h, with CV_ISI_(*t*) the coefficient of variation of the inter spike interval computed over a 10s window averaged across all excitatory neurons.
- Final weight values: ⟨*w*_EE_(4h)⟩ *<* 0.5, ⟨*w*_EI_(4h)⟩ *<* 0.5, ⟨*w*_IE_(4h)⟩ *<* 5, ⟨*w*_II_(4h)⟩ *<* 5 with *w*_EE_(4h) (resp. *w*_EI_(4h), *w*_IE_(4h), *w*_II_(4h) the average EE weight (resp. EI, IE, II weight) after the 4h simulation.
- Final weight distribution: *w*_blow_(4h) *<* 0.1, with *w*_blow_(4*h*) the fraction of synaptic weights that reached 0 or *w*_max_ (maximum over all 4 synapse types). This condition effectively enforced that no more than 10% of the weights of each synapse types had “blown up”, i.e., had reached extreme values.

Note that these criteria were less stringent than for 2 minutes, as many metrics on the 2min-task were designed to detect networks that would be unstable in the future. In our hands, these conditions which were added on top of the pre-screening performed in previous work^12^ were enough to constrain the network dynamics to plausible regimes. Note that these criteria for stability were established with a focus on cortex, and that in the future one may want to impose different conditions to better fit other brain regions.

#### Posteriors for specific network functions

To infer plasticity rules that created networks able to solve specific memory tasks, we applied a similar procedure as for the 4h-stability posterior. However, we did not have any indication of memory performance for the samples obtained from the 2min pre-screening. However, we had access to the memory performance of the 2,500 4h-stable quadruplets, and used these values to assign memory performance to the 110k quadruplets, using a nearest-neighbor algorithm. Several posteriors were obtained this way: *p*(*θ*|Δ*r*_mem_(10min)), *p*(*θ*|ON_nov_(10min)), 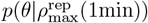and 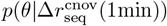 (see Fig. Supp. 13). Refer to the following sections for the definition of these metrics of network responses. We then sampled rule quadruplets from these posteriors conditioned on metric values reminiscent of experimental observations. In practice, this step increased the probability of sampling quadruplets with desired properties compared to the 4h-stable posterior (Fig. Supp. 13E). However, all quadruplets were verified numerically before being used in further analyses.

### Familiarity detection task

#### Input connectivity

The connectivity from *N*_inp→E_ = 10000 input neurons to excitatory recurrent neurons was receptive-field-like, similar to previous work^10^ (Fig. Supp. 2). For each recurrent excitatory neuron, we selected a random input neuron as the center of the circular receptive field of radius 8. The connections from neurons of this circular patch of input neurons to the considered recurrent neuron was *w*_inp_ = 0.075, and 0 to all other input neurons. The inhibitory neurons received inputs from *N*_inp→I_ = 4096 Poisson neurons with weight *w*_inp_ and similar receptive field connectivity than for the excitatory population. However, inhibitory neurons only received background inputs and no specific stimulus patterns. All input neurons fired at a baseline firing rate of 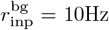 at all time, unless the input was part of an active input pattern (e.g. an active familiar stimulus).

#### Task

After a burn-out period with only background inputs of duration *l*_pre-train_ = 30s, the network was presented with one training stimulus (the familiar stimulus) for *l*_train_ = 10s. A break period with only background inputs followed, which could last *t*_b_ ∈ [1s, 10s, 20s, 1min, 2min, 5min, 10min, 30min, 1h, 4h]. A test session then began, during which a novel stimulus was presented to the network for 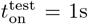 followed by 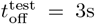 of background inputs. The same was then done with the familiar stimulus. Plasticity was turned on throughout the task. To avoid any biases due to the ordering of stimulus presentations (novel before familiar), the network state was saved at the beginning of each test session and loaded before presenting the familiar stimuli.

#### Familiar and novel stimulus design

For each stimulus pattern (familiar or novel), 500 adjacent input neurons were selected. While the pattern was active, the participating input neurons fired at 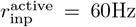 instead of 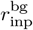. There was no overlap between the stimulus patterns.

### Evaluation of network response

The firing rate of the excitatory population was computed during the 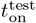 presentation of the familiar (*r*_fam_(*t*_b_)) and novel (*r*_nov_(*t*_b_)) stimuli. We then computed the relative preference of the network for the novel stimulus:

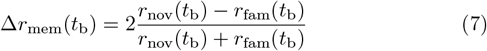

In Fig. 1, each rule quadruplet was tested *N*_trials_ = 5 times. Each trial involved a different draw of the connectivity matrices and the input Poisson spike trains. A rule quadruplet was deemed responsive at break time *t*_b_ if it passed student T-test comparing 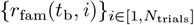and 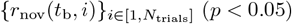.

Other metrics to evaluate memory recall on this task can be designed, for instance the transient dynamics (see next section for ON and OFF responses), or by taking into account the standard deviation of the neuron activities (see Fig. Supp. 4E):

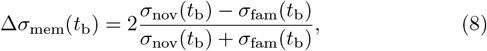

with *σ*_nov_(*t*_b_) = *σ*({*r*_nov,*i*_(*t*_b_)}_*i*∈E_) (resp. *σ*_fam_(*t*_b_) = *σ*({*r*_fam,*i*_(*t*_b_)} _*i*∈E_)) the standard deviation of the firing rates of excitatory neurons computed during presentation of the novel (resp. familiar) stimulus.

Note that we also checked that the simulation respected the stability criteria detailed in the above paragraphs.

### Transient dynamics task

This task was similar to the familiarity detection task described above, only the scoring metrics differed. We computed ON and OFF responses for familiar and novel stimuli presentations:

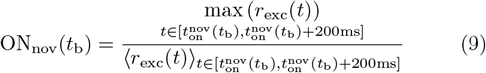

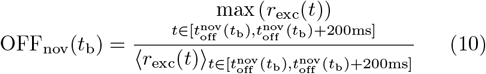

with *t*_on_(*t*_b_) (resp. *t*_off_(*t*_b_)) are the time of stimulus onset (resp. offset) of the novel stimulus after break period *t*_b_.

We computed the relative preference for novel stimuli when reading out memories from ON responses:

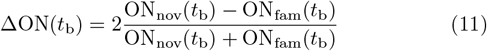

Other metrics were designed to quantify the quadruplet responses on this task. For instance, memory could also be read out from the OFF responses (see Fig. Supp. 5):

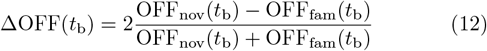

We observed that ON responses had less variance than the OFF responses (Fig. Supp. 5C), and thus favored the ON response metric.

### Sequential task

#### Input connectivity

The excitatory neurons in the network received *N*_inp→E_ = 11025 inputs from Poisson neurons firing at 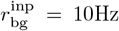. When a stimulus was active, a subset of the input neurons increased their firing rate to 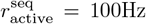 (see Fig. Supp. 3). The connectivity from input neurons to excitatory and inhibitory neurons was receptive-field-like (similar to the familiarity task, see Fig. Supp. 2). The inhibitory neurons received inputs from *N*_inp→I_ = 4096 Poisson neurons with *w*_inp_ and similar receptive field connectivity than for the excitatory population. However, inhibitory neurons only received background inputs 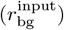 and no specific stimulus patterns.

#### Task

After a burn-out period 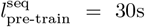 with only background inputs, the network was trained for 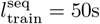 on a loop of 5 stimuli (the “familiar” stimuli). After a break period ranging from 1s to 4h with only background inputs, each training stimulus was presented independently for 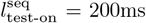, as well as two novel stimuli.

#### Familiar and novel stimuli design

All stimuli were non-overlapping (Fig. Supp. 3). To isolate as best as possible the effects of plasticity, the stimuli were designed such that a static network did not create any memories on this task (Fig. Supp. 3).

#### Engram neurons definition

Engram neurons were defined for every stimulus pattern (novel and familiar), at the beginning of the task. With plasticity turned off, we presented every stimulus to the network for 1s and collected the firing rates of each neuron during the whole stimulus presentation. The top 1/7th most active neurons were defined to be engram neurons for that stimulus.

#### Successor representation metric

This metric is used in Fig. 1F. We define 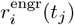 the firing rate of engram *i* during presentation of the stimulus *j* as the population firing rate computed over engram neurons for pattern *i* during 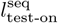 from stimulus onset. Then, 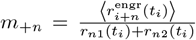. In particular, *m*_+1_ is plotted in Fig. 1F.

In Fig. 1, rule quadruplets were deemed responsive at a break duration *t*_b_ for the *m* + 1 metric if they passed a Student T-test comparing 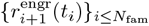 and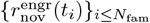.

### Metrics to evaluate replay

- 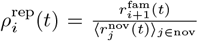: During the presentation of stimulus *i*, ratio of the firing rate of the engram of next stimulus *i* + 1 in the sequence, compared to the firing rate of the engram associated to a novel stimulus.
- 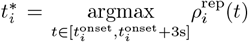 the time at which the ratio above was maximal, i.e. when the next stimulus was maximally activated following presentation of a given stimulus.
- 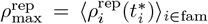 the maximum ratios averaged across all familiar stimuli.
- 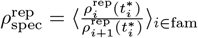, a metric to check if the maximal activation of the next stimulus is concurrent with activation of 2 stimuli in the future.
- 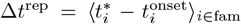 the time at which the maximal reactivation takes place.

In our hands, it was important for some metrics to compare engram firing rates to other engram firing rates, even if they were engrams associated to novel stimuli, as the process of selecting neurons to be part of an engram induced a bias on their overall activity.

### Contextual novelty task

This task was similar to the sequential task described above, only the test periods differed. After each break duration, we presented three groups of sequences of four stimuli to the network (see Fig. Supp. 9). The first group of sequences was fully familiar: Fam1-Fam2-Fam3-Fam4; Fam2-Fam3-Fam4-Fam5; Fam3-Fam4-Fam5-Fam1 etc. with each stimulus being on for 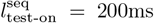 and no down time between stimuli presentations. The second group was composed of sequences ending with a novel stimulus: Fam1-Fam2-Fam3-Nov1; Fam2-Fam3-Fam4-Nov2; Fam3-Fam4-Fam5-Nov1 etc. The third group was composed of sequences ending in a out-of-order familiar stimulus: a contextual novel stimulus: Fam1-Fam2-Fam3-Fam5; Fam2-Fam3-Fam4-Fam1; Fam3-Fam4-Fam5-Fam2 etc. To assess the network responses to these stimuli, we define 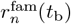 the firing rate of the excitatory population computed during the fourth stimulus in the *n*^th^ familiar sequence, and similarly 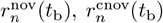. From these, we defined two metrics for this memory task:

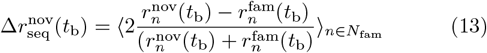

and

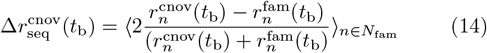

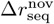 can be seen as a generalization of Δ*r*_mem_ used in the familiarity task, but for the case of multiple familiar inputs (see Fig. Supp. 9). 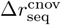 is the relative preference of the network to the contextually novel stimulus compared to the familiar one.

### Criteria for quadruplets to be responsive to a task

In this study, we used two criteria for each network function.

#### Loose criteria

In Fig. 1, we use permissive criteria to label a rule quadruplet as responsive to a task. These criteria involves averaging over several trials.

- Novelty detection: A rule was deemed able to perform novelty detection if at any of the break times a significant difference between *r*_nov_(*t*_b_ and *r*_fam_(*t*_b_ with *r*_nov_(*t*_b_) ≥ *r*_fam_(*t*_b_) (see section on familiarity task for the statistical test).
- Familiarity detection: Same as above but with *r*_nov_(*t*_b_) ≤ *r*_fam_(*t*_b_)
- Transient dynamics: The condition was ON_nov_(*t*_b_) *>* 1.5 for all break after 5min. Note that here, we are only assessing whether a quadruplets generates notable transient dynamics and not whether it read out any memories.
- Successor representation: A rule was deemed able to be responsive to the sequential task if at any of the break times a significant difference between 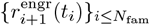 and 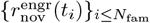 could be detected (see section on sequential task for the statistical test).
- Contextual novelty: A rule was deemed responsive if at any of the break times there was a significant difference between 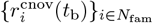 and 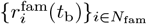.

These criteria only applied to the 2,500 quadruplets initially sampled from the stability manifold. For the other quadruplets sampled from more specialized posteriors, we employed the criteria below.

#### Tight criteria

In Fig. 2, we use more stringent criteria to qualify rule quadruplets as consistent with experimental observations:

- Novelty detection: ∀*t*_b_ ≥ 2min, Δ*r*_mem_.(*t*_b_) *>* 0.1.
- Familiarity detection: Δ*r*_mem_(*t*_b_) *<* −0.1 for 3 consecutive break durations.
- Transient dynamics: ∀*t*_b_ ≥ 2min, ON_nov_(*t*_b_) *>* 1.8.
- Flat replay: 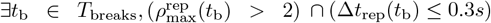.
- Sequential replay: 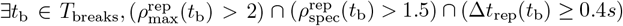.
- Contextual novelty: ∀*t*_b_ ≤ 10min, 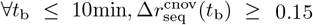.

Note that these criteria above are relatively arbitrary and were chosen to ensure that the network responses were qualitatively like experimental data and significant. The exact stringency of the criteria was tuned to select the best ≈ 100 rule quadruplets in the dataset. Due to the compute constraint, in this second part, we only tested rule quadruplets on a single trial (i.e. only the 2,500 base quadruplets were tested on all tasks on several trials).

#### Knock-outs

We conducted two types of knock-out (KO) experiments: single KOs, where plasticity was removed from one connection type at a time, and triple KOs, where plasticity was removed from three connection types simultaneously, leaving only one connection type plastic. For both KO types, we tested all possible combinations across the four connection types, for all 2,500 stable rule quadruplets tested in Fig. 1. Knocking out the plasticity rule of type XY was defined as setting *η*_XY_ = 0 in equation (1) after a burn-in period of *l*_pre-train_ = 30s, during which only background inputs were present. This burn-in allowed synaptic weights to evolve from their default values, ensuring that KO weights were not overly dependent on our choice of initialization weights. The networks then underwent the familiarity detection task as described previously. Simulations were performed across all *t*_break_ values up to 4h, with all break times listed. Networks were then classified as stable or unstable using the stability criteria defined previously.

### Reservoir computing and pong

#### Familiarity detection task

A dataset of 2,500 simulations with the stable rule quadruplets used in Fig. 1 undergoing the familiarity task was prepared. During each simulation, spiking activity was recorded from all 5120 recurrent neurons. Simulations were conducted using the Brian2 simulator^56^. One readout unit (leaky-integrate-and-fire dynamics) was trained to perform the familiarity task receiving 55s of spike trains from the dataset as input. To show that we could obtain a readout of the network that was not bound to the preference induced by the quadruplet, the task was either to increase (resp. decrease) activity when a novel (resp. familiar) stimulus had been shown (Fig. Supp. 18). In a similar fashion to previous work^53^, the readout neuron received spike trains from 5,120 inputs and underwent training using the surrogate gradient descent approach^42^. The loss function was defined as the mean squared error between the firing rate of the output unit and the expected firing rate (in the case of novelty detection: 50Hz for novel and 0Hz for familiar; and vice versa for familiarity detection). 80% of the dataset was used for training and the remaining 20% for the validation phase. Accuracy was calculated as: 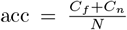, where *C*_*f*_ is the number of correct familiar responses, *C*_*n*_ is the number of correct novel responses, and *N* is the total number of validation trials. Maximum accuracy was calculated as the highest achievable accuracy for a threshold separating familiar and novel responses between 0 and 50Hz. To avoid an input location bias, during the training session, locations of familiar and novel inputs were regularly swapped between two different populations of neurons. During the validation phase, novel and familiar inputs were provided to the neurons that had never been exposed to them during training.

#### Pong – Offline

To simulate the game environment, we started from existing code training electro-active polymer hydrogels to play pong^65^. The game field was a 640×640 pixels square field and the ball was a 90 × 90 pixel square (Fig. Supp. 19). The ball was initialized on the right side of the field in a random direction (an angle between 109^◦^ and 253^◦^) and traveled across the field at a constant speed. No paddle was included in these simulations.

Following the idea of training a neural culture to play pong^45^, 4096 excitatory neurons were arranged spatially onto a 64 × 64 grid. Neurons received external inputs that modeled the position of a ball during the simulated games, i.e. neurons at the location of the ball would receive increased excitatory inputs compared to baseline. Each ball position excited a 9 × 9 square of recurrent neurons for 200 ms. Simulations were conducted using the Brian2CUDA simulator^57^. Every 200ms, the firing rate of each neuron was calculated to match separate stimulation periods and ball movement. To facilitate the position readout from the network activity, the firing rate was normalized to the highest firing rate during one simulation, and the ball position coordinates were normalized to the range (−1,1).

To decode the current and previous positions of the ball from the network activity, we trained readout weights from the recurrent network with a dataset composed of 2,000 pong games for training and 1,000 for validation. Since trials were independent, no position prediction was allowed using network activity from more than one trial. The readout comprised one linear layer with input size *N*_E_ = 4096 and 4 linear output units. These four output neurons modeled the decoded x and y coordinates of the current and previous ball positions. The loss function was defined as follows:

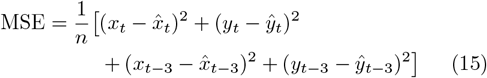

with *n* is the number of output units, *x*_*t*_, *y*_*t*_ are the true x and y coordinates of the ball at the current time step 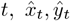 are the predicted x and y coordinates at time *t, x*_*t*−3_, *y*_*t*−3_ are the true x and y coordinates of the ball at the previous time step 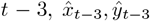 are the predicted x and y coordinates at time *t* −3. We did not train the recurrent network to take actions from these decoded values. Instead, using the decoded position and direction of the ball, a straight line was fitted to the direction of the ball, and the paddle movement was simulated accordingly, to intersect with the ball. The paddle size was equal to the size of the ball. The accuracy was implemented by comparing the ball position with the paddle. If both overlapped, the agent scored a point. The score was calculated as the number of successful trials over the total number of trials.

#### Place field calculation

Place field experiment involved the same game environment as pong (offline version). The simulation of the network activity consisted of showing solely one ball trajectory 200 times during one simulation. A single ball position that was part of the training trajectory was shown for 2s, first at the beginning of the simulation (“naive network” in Fig. 4E) and 30s after the last trajectory was shown (“after pong” in Fig. 4E). The weights were extracted for each connection type and sorted by their direction and distance along the x-axis of the 64×64 grid network. For each neuron in the grid, we calculated the distribution of weights of the connections sent to the neurons on the left side (right-to-left direction) vs the weights of the connections sent to the neurons on the right side (left-to-right direction). The summed weights were plotted against the distance on the x-axis of the grid.

#### Pong - Real-time

This game version extends the offline pong to the real-time prediction of the ball direction and control of the paddle. The game used the same environment as described in the section above. To decrease the computations needed to train the readout model, a model trained offline was used to predict the ball position in real-time. Real-time pong followed the same protocol as its offline version: ball position was tracked during the game and immediately provided as a sensory input to the network. Next, the activity of the network was normalized and sent to the readout which predicted the direction of the ball by fitting a linear equation to two predicted positions. As a result, the paddle was controlled. Real-time pong allowed for changing the velocity of the ball and initializing the ball in different directions than during the offline game. Two different versions of readout models for the network with static or plastic connections were verified (Table III). First, the mean score per trial and the maximum score for the games were measured with models that were trained offline and validated in real-time (no fine-tuning). Second, the same statistics were measured for models that underwent offline training and a small real-time fine-tuning for 50 trials before validating them. This was done to leverage any possible differences related to a real-time setup such as different ball trajectories or speed.

## Acknowledgments

We would like to thank Tim Behrens, Henning Sprekeler, Nicolas Brunel, David Scheinberg, Andrew Peters, Matteo Carandini, Maneesh Sahani, Nicoleta Condruz, Chaitanya Chintaluri and Douglas Feitosa Tomé for useful discussions, as well as Stefano Elefante and Alois Schlögl for their help with deploying simulations on the ISTA cluster.

## Author contributions

BC, PR, PG, JHM and TV designed the study, BC, MK and ZH ran and analyzed the simulations. BC and TV wrote the manuscript, with help from all coauthors. PB created the online supplementary material.

## Data and materials availability

All the code for running the networks and reproducing the analysis is available on Github https://github.com/VogelsLab/dataSBI.

