## Supplementary material for "Memory by a thousand rules: Automated discovery of multi-type plasticity rules reveals variety & degeneracy at the heart of learning": DataSBI_bioRxiv_AllSupp.pdf

#### Appendixes

##### Linear regression of network function from plasticity parameters

To assess the relationship between plasticity rule quadruplet parameterization and emergent network firing rates, we performed multiple linear regressions using Ordinary Least Squares (OLS) implemented in scikit-learn (Python). Each regression model included 24 independent variables representing the parameters of the rule quadruplet, namely $\theta_{\mathrm{XY}} = [\alpha_{\mathrm{XY}}, \beta_{\mathrm{XY}}, \gamma_{\mathrm{XY}}, \kappa_{\mathrm{XY}}, \tau_{\mathrm{pre}}^{\mathrm{XY}}, \tau_{\mathrm{post}}^{\mathrm{XY}}]$ for synapse types X,Y ($\mathrm{X}, \mathrm{Y} \in (\mathrm{E}, \mathrm{I})$), as defined in Eq. (1). All parameters were z-scored prior to analysis to standardize their scales. We included parameters of the 2,500 rule quadruplets producing stable network dynamics (used in Fig. 1). The dependent variables, assessed in three separate regressions, were three metrics evaluated at $t_{\mathrm{break}} = 10\mathrm{s}$: excitatory neuron population firing rate ($r_{\mathrm{exc}}(t_{\mathrm{break}})$), inhibitory neuron population firing rate ($r_{\mathrm{inh}}(t_{\mathrm{break}})$), and relative novelty preference ($\Delta r_{\mathrm{mem}}(t_{\mathrm{break}})$) defined for the familiarity detection task in Eq. 7. For clarity, we refer to these metrics without the explicit break time index in the remainder of the analysis. The dataset for each regression model was split into a training set (80%) and a test set (20%) to evaluate the performance of the model. If a specific firing rate metric was undefined for a given rule quadruplet, the corresponding 24 parameters were excluded from the regression analysis involving the dependent variable. The regression results for each dependent variable are summarized in Table (I), and the values of significant coefficients with a p-value below 0.05 are listed in Table (II). The linear regression models for the excitatory neuron population firing rate ($r_{\mathrm{exc}}$) and inhibitory neuron population firing rate ($r_{\mathrm{inh}}$) performed well, with $r_{\mathrm{exc}}$ achieving an R-squared of 0.712 and an adjusted R-squared of 0.708 (F-statistic $= 182.8$, $p < 0.001$), and $r_{\mathrm{inh}}$ showing a moderate fit with an R-squared of 0.662 and an adjusted R-squared of 0.657 (F-statistic $= 131.9$, $p < 0.001$). These results indicate a reasonable degree of explanatory power for both models, consistent with expectations given the model complexity. The adjusted R-squared values, being close to the R-squared values (0.708 vs. 0.712 for $\mathrm{r}_{\mathrm{exc}}$, and 0.657 vs. 0.662 for $r_{\mathrm{inh}}$, suggest that these models are not overfitting despite including 24 predictors. In contrast, the model for relative novelty preference ($\Delta r_{\mathrm{mem}}$) exhibited a weak fit, with an R-squared of 0.112 and an adjusted R-squared of 0.096 ($F$-statistic $= 6.993$, $p < 0.001$), suggesting that novelty preference is challenging to predict using the quadruplet parameterization in a linear regression framework. These results highlight that, while the plasticity parameters effectively predict population firing rates, they struggle to capture more complex metrics, assessing network function, such as the relative preference for the novel stimulus.

##### Online supporting material

Given the wealth of rules proposed in this study, we provide a semi-automated analysis of every rule quadruplet in our dataset. The resulting online supplementary material can be found at *https://plasticity.vogelslab.org*

| Dependent Variable | R-squared | Adj. R-squared | F-statistic (p-value) |
|---|---|---|---|
| $r_{\text{exc}}$ | 0.712 | 0.708 | $F(24, 1796) = 182.8$ ($< 0.001$) |
| $r_{\text{inh}}$ | 0.662 | 0.657 | $F(24, 1638) = 131.9$ ($< 0.001$) |
| $\Delta r_{\text{mem}}$ | 0.112 | 0.096 | $F(24, 1359) = 6.993$ ($< 0.001$) |

**Table I. Summary of regression results for the three dependent variables.**

| Dependent Variable | Significant Coefficients |
|---|---|
| $r_{\text{exc}}$ | $\tau_{\text{pre}_{\text{EE}}} = -0.3545$, $\tau_{\text{post}_{\text{EE}}} = -0.1659$, $\alpha_{\text{EE}} = 1.8891$, $\beta_{\text{EE}} = 1.9163$, $\gamma_{\text{EE}} = 0.5152$, $\kappa_{\text{EE}} = 0.4954$, $\tau_{\text{pre}_{\text{IE}}} = -1.1067$, $\tau_{\text{post}_{\text{IE}}} = -0.1345$, $\alpha_{\text{IE}} = -0.8065$, $\beta_{\text{IE}} = -1.0850$, $\gamma_{\text{IE}} = -0.2594$, $\kappa_{\text{IE}} = -0.2614$, $\tau_{\text{pre}_{\text{II}}} = -0.1264$, $\tau_{\text{post}_{\text{II}}} = -0.1006$, $\alpha_{\text{II}} = -0.7869$, $\beta_{\text{II}} = -0.8533$, $\gamma_{\text{II}} = -0.1280$, $\kappa_{\text{II}} = -0.2760$ |
| $r_{\text{inh}}$ | $\tau_{\text{pre}_{\text{EE}}} = -0.0799$, $\alpha_{\text{EE}} = 0.3457$, $\beta_{\text{EE}} = 0.3136$, $\gamma_{\text{EE}} = 0.0729$, $\kappa_{\text{EE}} = 0.0858$, $\tau_{\text{pre}_{\text{EI}}} = -0.0505$, $\tau_{\text{post}_{\text{EI}}} = -0.0392$, $\alpha_{\text{EI}} = 0.4016$, $\beta_{\text{EI}} = 0.3102$, $\gamma_{\text{EI}} = 0.1242$, $\kappa_{\text{EI}} = 0.1442$, $\alpha_{\text{IE}} = -0.1177$, $\beta_{\text{IE}} = -0.1802$, $\gamma_{\text{IE}} = -0.0430$, $\kappa_{\text{IE}} = -0.0380$, $\tau_{\text{pre}_{\text{II}}} = -0.3108$, $\tau_{\text{post}_{\text{II}}} = -0.2137$, $\alpha_{\text{II}} = -1.7196$, $\beta_{\text{II}} = -1.8886$, $\gamma_{\text{II}} = -0.2735$, $\kappa_{\text{II}} = -0.4103$ |
| $\Delta r_{\text{mem}}$ | $\alpha_{\text{EE}} = -0.0156$, $\beta_{\text{EE}} = 0.0115$, $\tau_{\text{pre}_{\text{EI}}} = -0.0058$, $\gamma_{\text{EI}} = -0.0174$, $\kappa_{\text{EI}} = 0.2018$, $\tau_{\text{post}_{\text{IE}}} = -0.0073$, $\alpha_{\text{IE}} = -0.0202$, $\beta_{\text{IE}} = -0.0190$, $\gamma_{\text{IE}} = -0.0062$, $\kappa_{\text{IE}} = -0.0096$ |

**Table II. Significant regression coefficients for each dependent variable ($p < 0.05$).**

| **Validation Scores** # Trials: 1000 | **Plastic $\theta_1$** | **Plastic $\theta_2$** | **Plastic $\theta_3$** | **Static** |
|---|---|---|---|---|
| Offline: Accuracy | 0.65 | 0.57 | 0.17 | 0.27 |
| Real-time w/o tuning: Mean Score per Trial | 0.65 | 1.00 | 0.37 | 0.56 |
| Real-time w/o tuning: Max Score | 8 | 14 | 6 | 4 |
| Real-time with tuning: Mean Score per Trial | 0.79 | 1.19 | 0.76 | 0.43 |
| Real-time with tuning: Max Score | 10 | 16 | 11 | 5 |

**Table III. Summary of pong game validation scores for plastic and static networks during offline and real-time trials.**

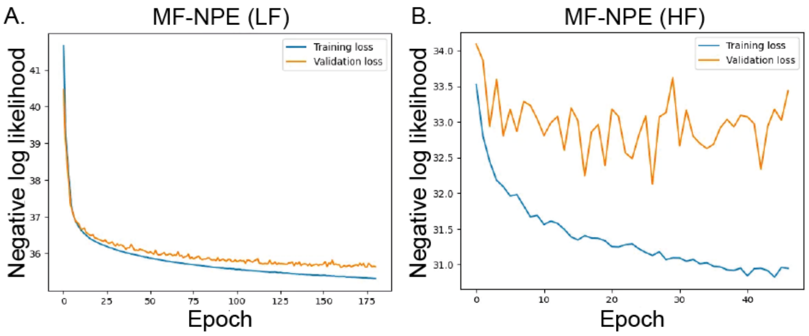


**Fig. Supp. 1. Example of fSBI/MF-NPE training. A**: Training and validation loss for training a posterior on the low fidelity samples (100k rule quadruplets stable for 2min obtained with fSBI[12]). The summary statistic used for that posterior was the firing rate of the excitatory population. **B**: Training and validation loss of the second part of the posterior training, using the high fidelity samples (2,500 rule quadruplets tested to be stable after 4h).

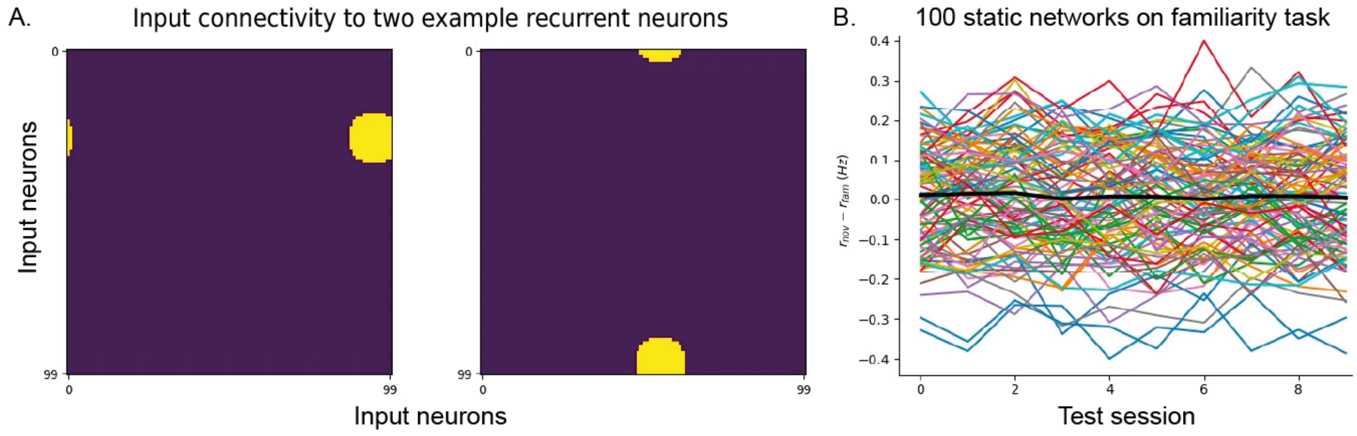


**Fig. Supp. 2. Familiarity task design.** **A**: Two example connectivity matrices from the input neurons (arranged in 2D) to example recurrent excitatory neurons. Yellow denotes a non-zero weight. **B**: Difference of firing rate between the novel and familiar network response over test sessions (break periods ranging from 1s to 4h), shown for 100 randomly initialized static networks, average firing rate for each test session over networks is shown in black.

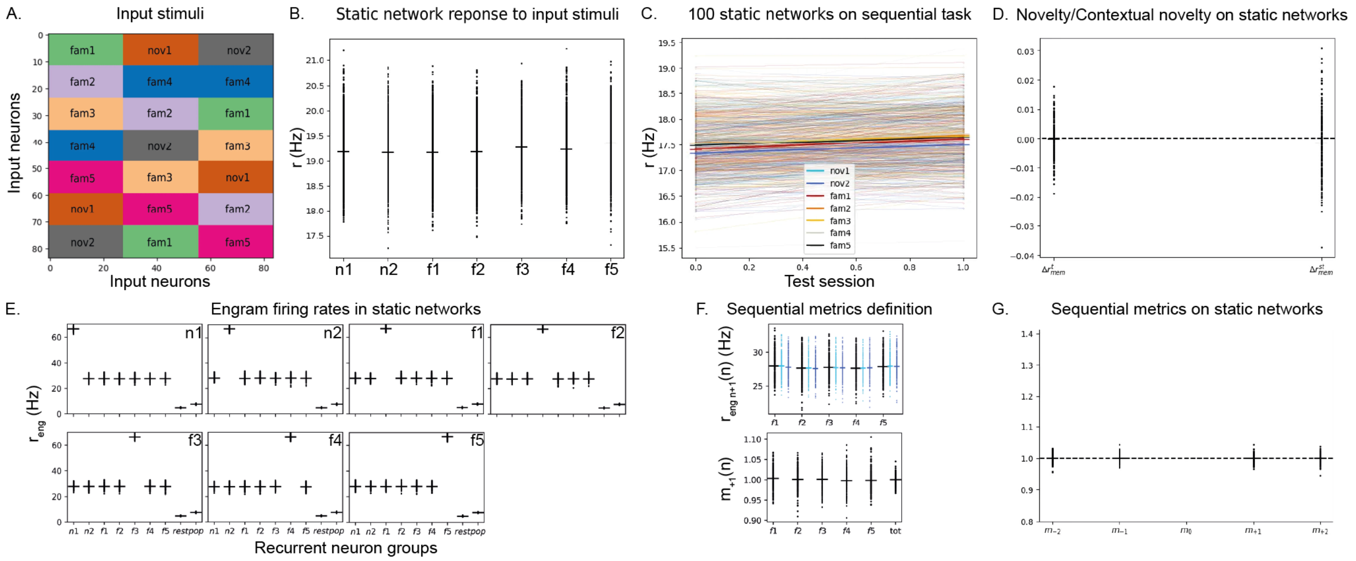


**Fig. Supp. 3. Sequential task design.** **A**: Arrangement of the 7 input patterns used in the task (5 used for training–familiar–, 2 only presented in testing–novel–. **B**: Population firing rate in response to stimulation of a static network with the 7 input stimuli shown in A (after a random break period of 1s to 4h, firing rates computed during a 1s presentation of each input stimulus). Results averaged over 100 randomly initialized static networks. **C**: Population firing rate across test sessions for 100 static networks in response to each input stimulus. Average for each stimulus shown in bold. **D**: Metrics for the contextual novelty task computed on 100 static networks, see methods. **E**: Firing rate of each engram (group of recurrent excitatory neurons, see definition in methods) during presentation of the input stimuli. "Rest" denotes all the excitatory neurons not part of any familiar engrams, "pop" the firing rate of the entire population. **F**: Top: firing rate of the engram n+1 while input stimulus n was presented (familiar 2 engram during familiar 3 presentation / familiar 1 engram during familiar 5 presentation), for various engram sizes (5% in black, 10% light blue and 14% dark blue). Bottom: $m_{+}1$ metric computed for each stimuli, on the right averaged over all stimuli, as described in methods. Done in static networks. **G**: all the $m$ metrics for successor representation (see methods). $m_0$ is a large value, not shown here.

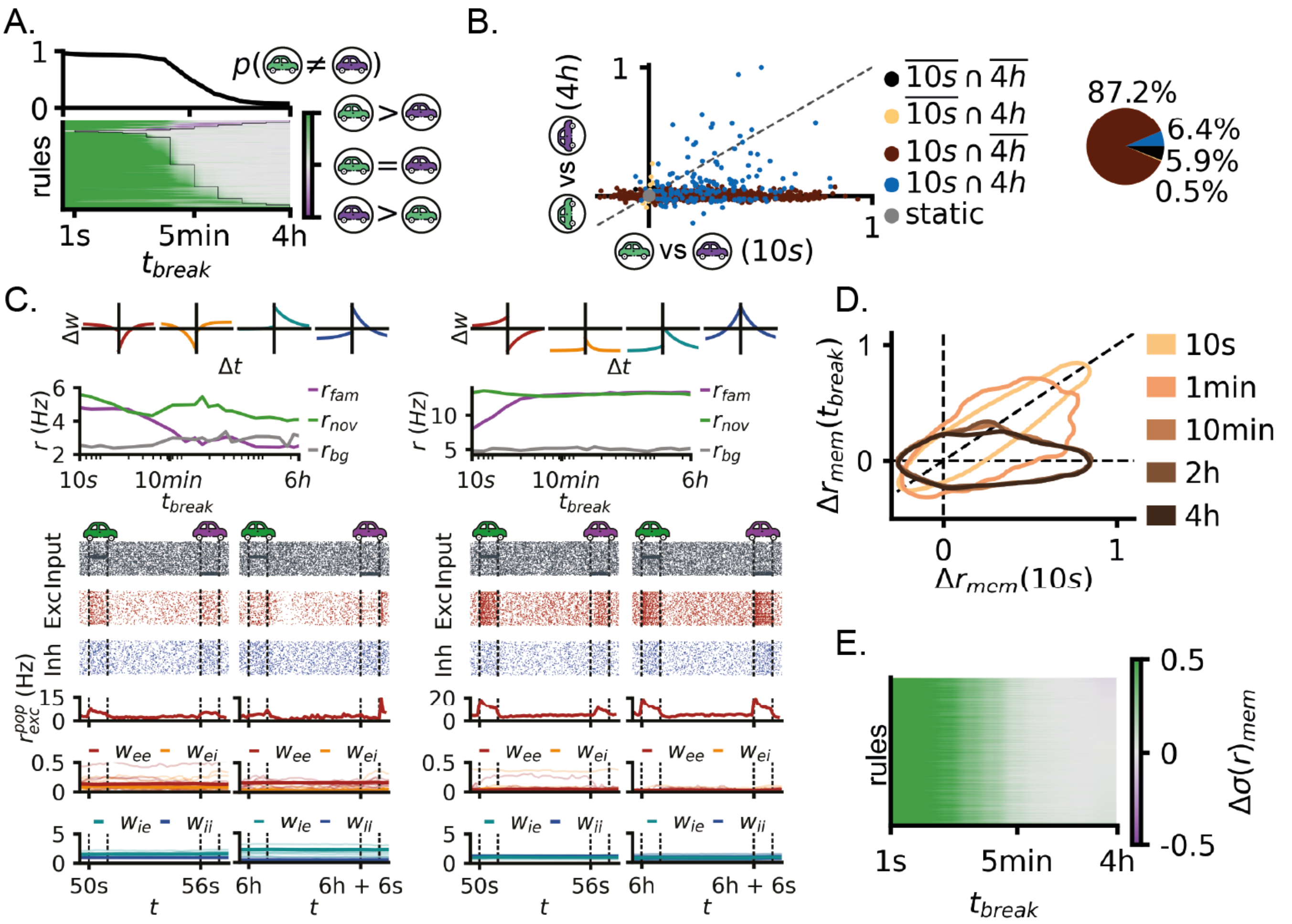

**Fig. Supp. 4. Additional analysis of the familiarity detection task.** **A**: 2,500 stable quadruplets from the stability manifold (the same as in Fig. 1) going through the familiarity task. Fraction of quadruplets that elicit significantly different familiar vs novel responses as a function of the break time. **B**: Relative novel preference ($\Delta r_{\text{mem}}$, see methods) after 4h vs after 10s for of the 2,500 stable quadruplets. Quadruplets are colored depending on the significance of their novel vs familiar responses at 10s and 4h (e.g. blue dots are quadruplets with significant responses after 10s and 4h. The proportion of such quadruplets is shown with the same colors on the right. **C**: Dynamics of networks evolving with 2 rule quadruplets from the stability manifold. From top to bottom: shape of the quadruplets in the pre-post protocol; population firing rate of the network during background inputs/familiar stimulus/ novel stimulus active for different break times; raster plot of input/excitatory/inhibitory neurons over two test sessions. Times of presentation of novel (resp. familiar) stimulus is shown by the green (resp. purple) cartoon car; firing rate of the excitatory population; weights of each synapse type, means shown in bold. **D**: Relative novel preference for the 2,500 stable quadruplets across break times, the 50% mass contour is shown. **E**: Relative novel preference computed from the standard deviation of individual firing rates (instead of the mean, see methods).

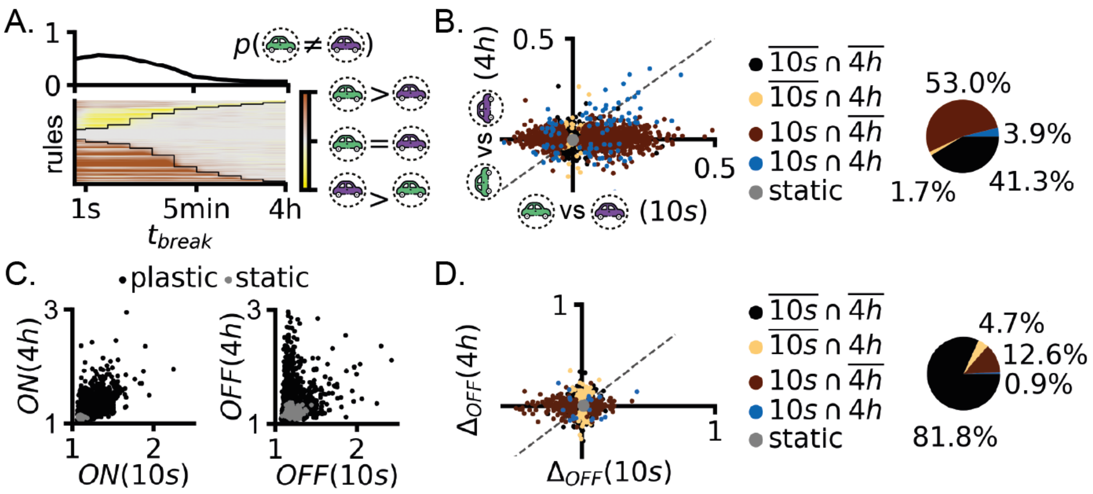


**Fig. Supp. 5. Additional analysis of the transient amplification task.** **A**: 2,500 stable quadruplets from the stability manifold (the same as in Fig. 1) going through the familiarity task. Fraction of quadruplets that elicit significantly different familiar vs novel responses as a function of the break time, as read out by the $\Delta\text{ON}(t_{\text{break}})$ metric. **B**: Relative novel preference computed using ON responses ($\Delta\text{ON}(t_{\text{break}})$, see methods) after 4h vs after 10s for of the 2,500 stable quadruplets. Quadruplets are colored depending on the significance of their novel vs familiar responses at 10s and 4h (e.g. blue dots are quadruplets with significant responses after 10s and 4h). The proportion of such quadruplets is shown with the same colors on the right. **C**: Right: ON responses to a novel stimulus for different break periods. Each dot denotes one of the 2,500 stable quadruplets. Left: same but for the OFF responses. **D**: Same as B, but reading out memories using OFF responses.

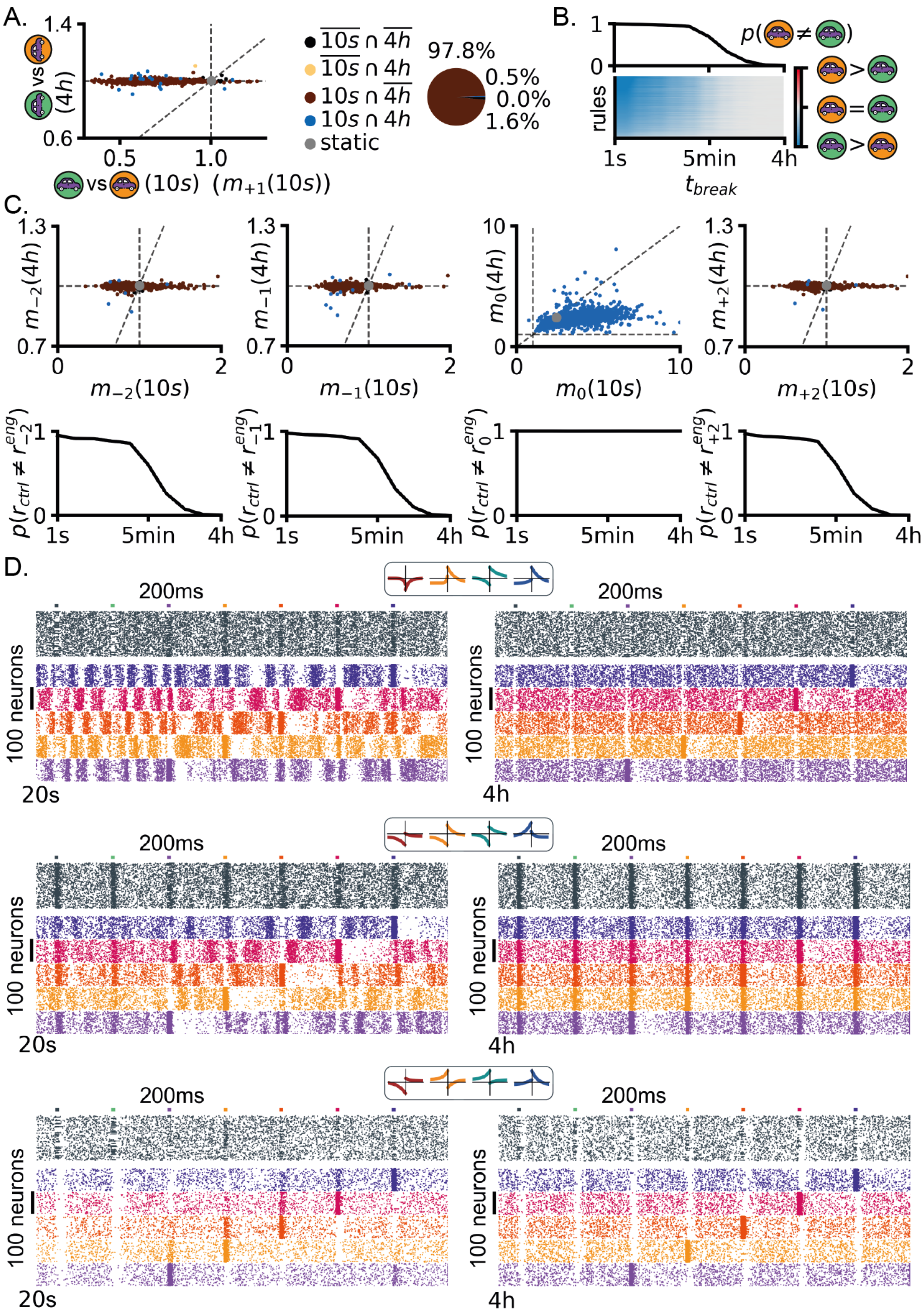

**Fig. Supp. 6. Additional analysis of the sequential task.** **A**: Successor representation metric ($m_{+1}(t_{\text{break}})$, see methods) after 4h vs after 10s for of the 2,500 stable quadruplets. Quadruplets are colored depending on the significance of their novel vs familiar responses at 10s and 4h. **B**: 2,500 stable quadruplets from the stability manifold (the same as in Fig. 1) going through the sequential task. Fraction of quadruplets that elicit significantly different responses of the next engram in the training sequence (compared to the one presented) vs control neurons (see methods). **C**: Same as A and B, but for the other successor representation metrics (e.g. $m_{-1}$ for the previous stimulus in the training sequence). **D**: Raster plots grouped by engrams over the entire test session, for the same three example rules as in Fig. 1, with the same color code and notations.

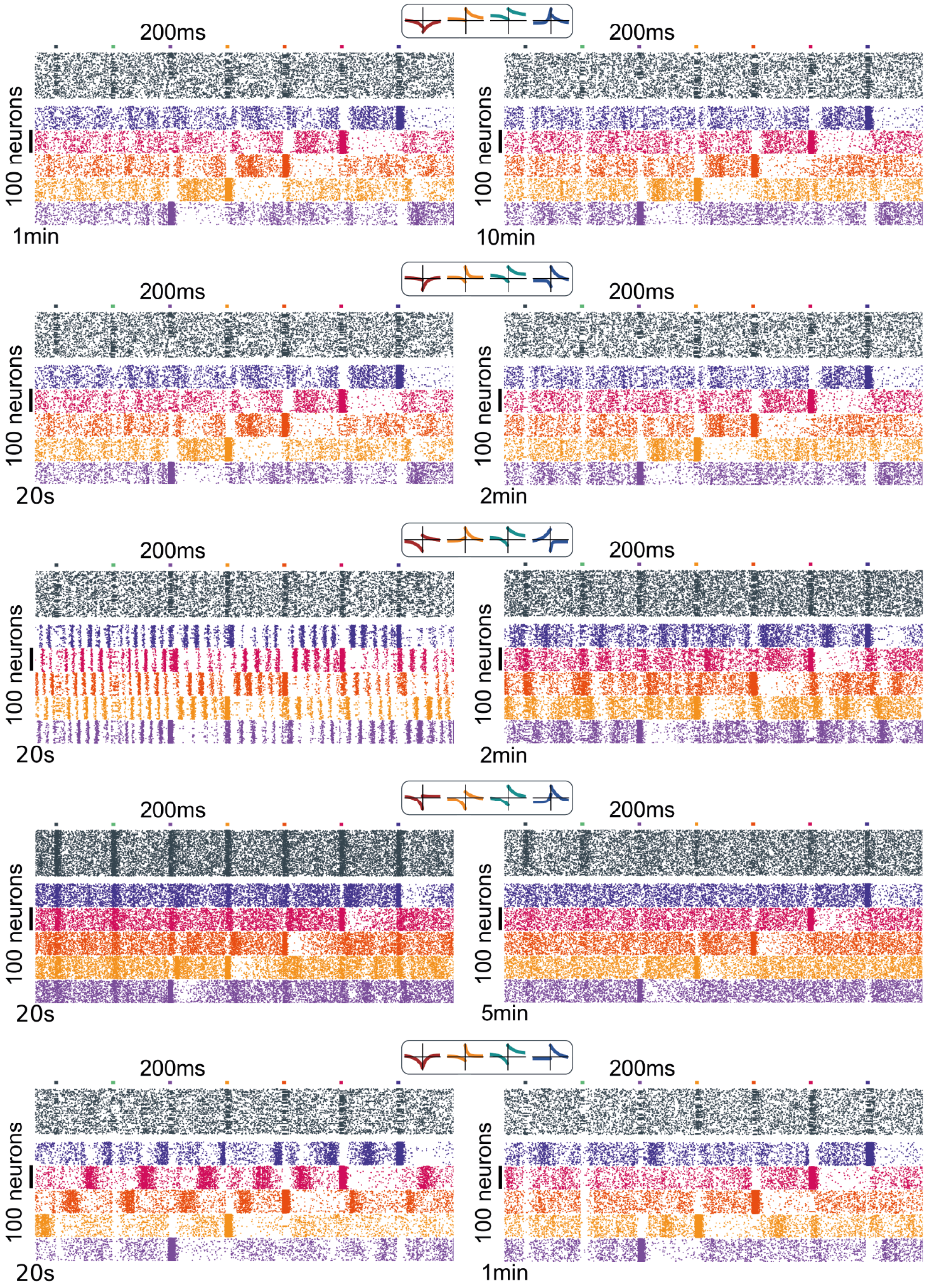


**Fig. Supp. 7. Additional example sequential rules.** Raster plots grouped by engrams over the entire test session, for a few example rule quadruplets, with the same color code and notations as in Fig. 1.

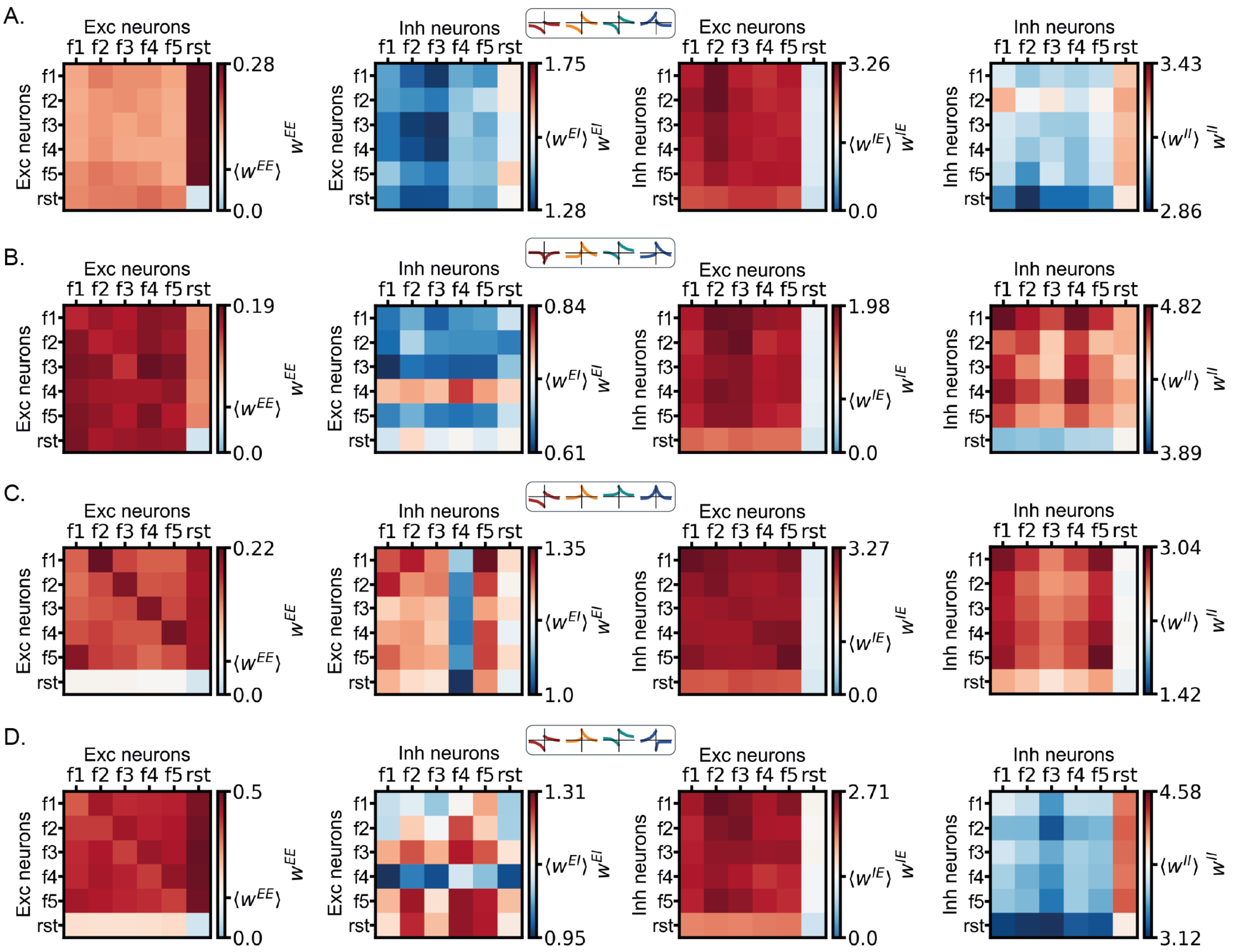


**Fig. Supp. 8. Weight matrices in the sequential task. A-D:** Weight matrices extracted directly after the end of training for three rule quadruplets. Weights were grouped by which engrams their post and pre-synaptic neurons belonged to (see methods for definition of engrams). "rst" denotes all remaining neurons not participating in any familiar engram. Weights were forced to be positive, but the color bars are centered on the mean weight of a given matrix.

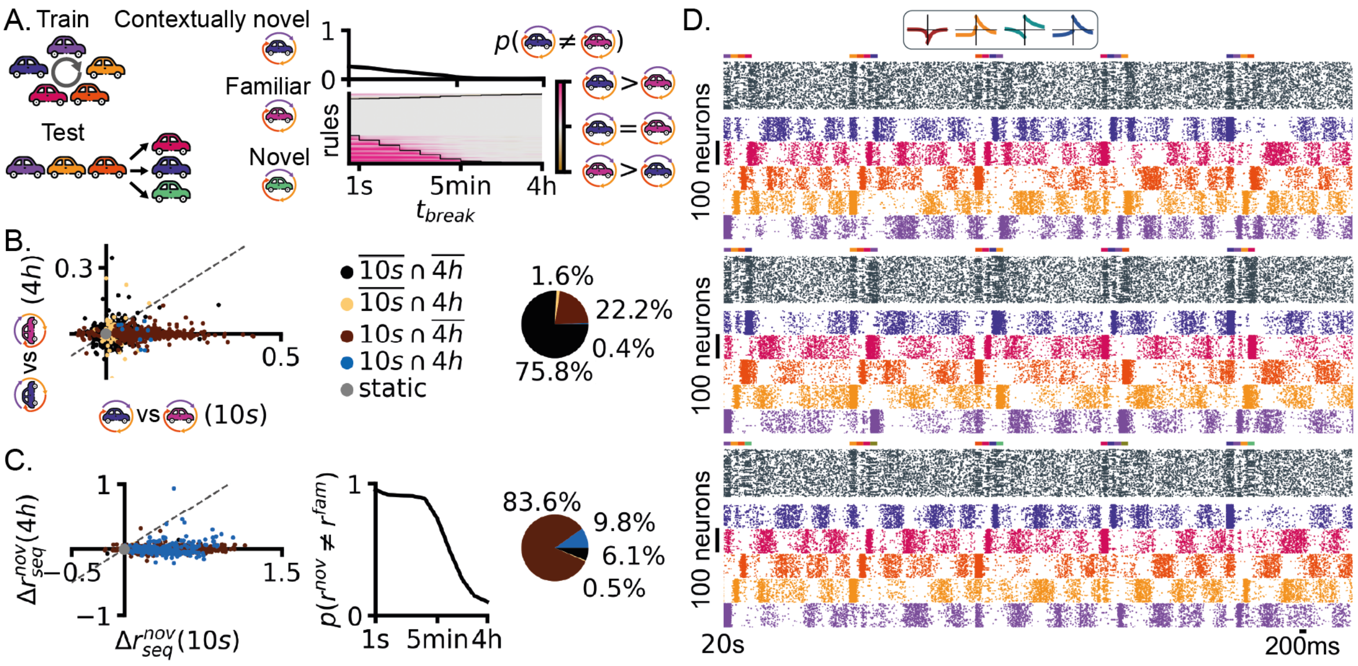


**Fig. Supp. 9. Additional analysis of the contextual novelty detection task.** **A**: Left: Description of the task. Right: 2,500 stable quadruplets from the stability manifold (the same as in Fig. 1) going through the contextual novelty task. Fraction of quadruplets that elicit significantly different familiar vs contextually novel responses as a function of the break time, as read out by the $\Delta r^{\mathrm{cnov}}_{\mathrm{seq}}(t_{\mathrm{break}})$ metric (see methods). **B**: Relative contextual novel preference ($\Delta r^{\mathrm{cnov}}_{\mathrm{seq}}(t_{\mathrm{break}})$, see methods) after 4h vs after 10s for of the 2,500 stable quadruplets. **C**: Same as B, but for $\Delta r^{\mathrm{nov}}_{\mathrm{seq}}(t_{\mathrm{break}})$, the relative novelty preference computed in the sequential task. **D**: Raster plots grouped by engrams over the entire test session of the contextual novelty task, for the same three example rules as in Fig. 1, with the same color code and notations. Top row shows the familiar sequences, middle the contextually novel and bottom the novel sequences.

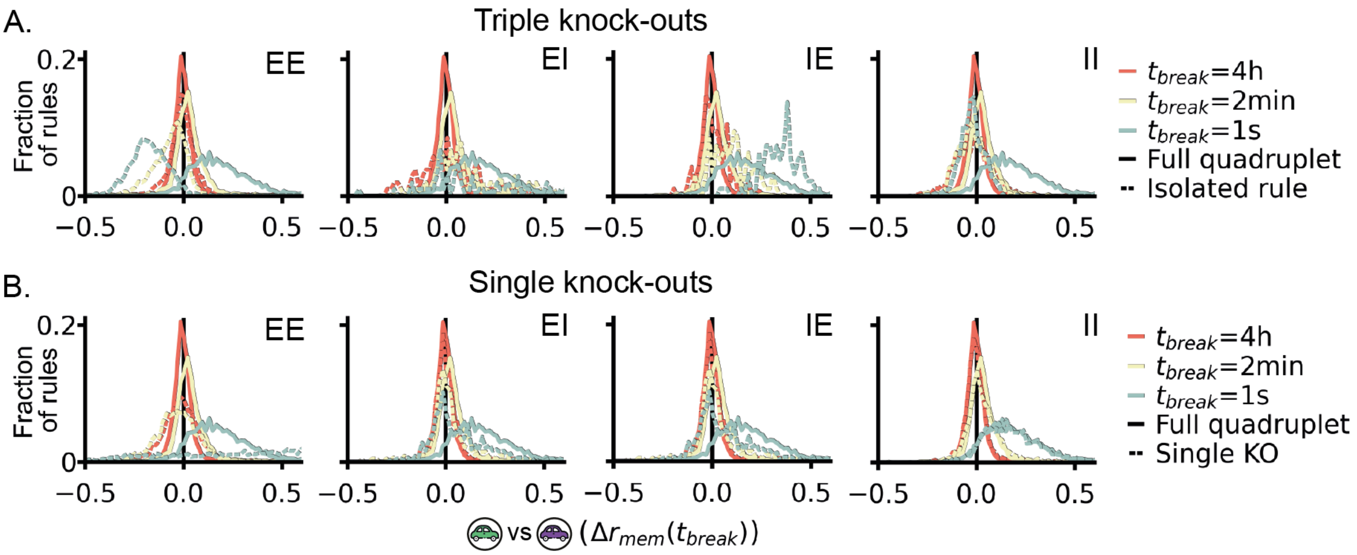


**Fig. Supp. 10. Additional analysis of the knock-outs experiments.** **A:** For the triple knock-outs (3 out of 4 plasticity rules within a stable quadruplets are removed when the task starts, see methods), among the stable resulting quadruplets, distribution of their relative novel preference for different break times. The distribution for all stable quadruplets at the same break points is shown for reference. Left to right: isolated stable EE rules (resp. EI, IE and II). **B**: Same as A but for the single knock-outs (only one of the four co-active rules is removed).

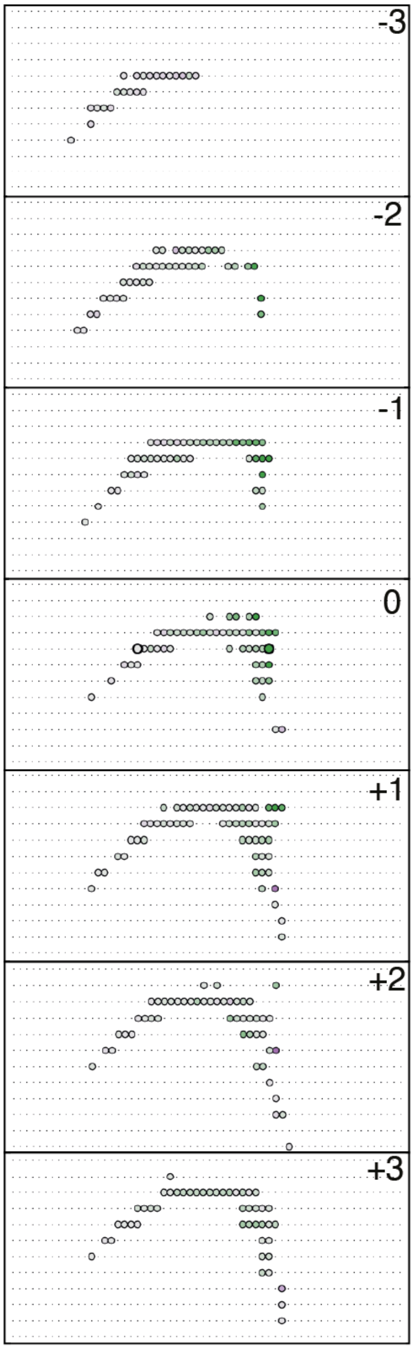


**Fig. Supp. 11. Additional analysis of stability manifold shape.** Similar to Fig. 3C, sampling quadruplets using two quadruplets from the stability manifold (the same as in Fig. 3C). The randomly chosen plane is different than in the main figure, and 7 parallel sheets are generated, on which quadruplets are sampled on a grid. Black dots denote unstable quadruplets. Larger dots denote stable quadruplets and the color represents their response after 10 minutes on the familiarity task (same colorbar as Fig. 1D).

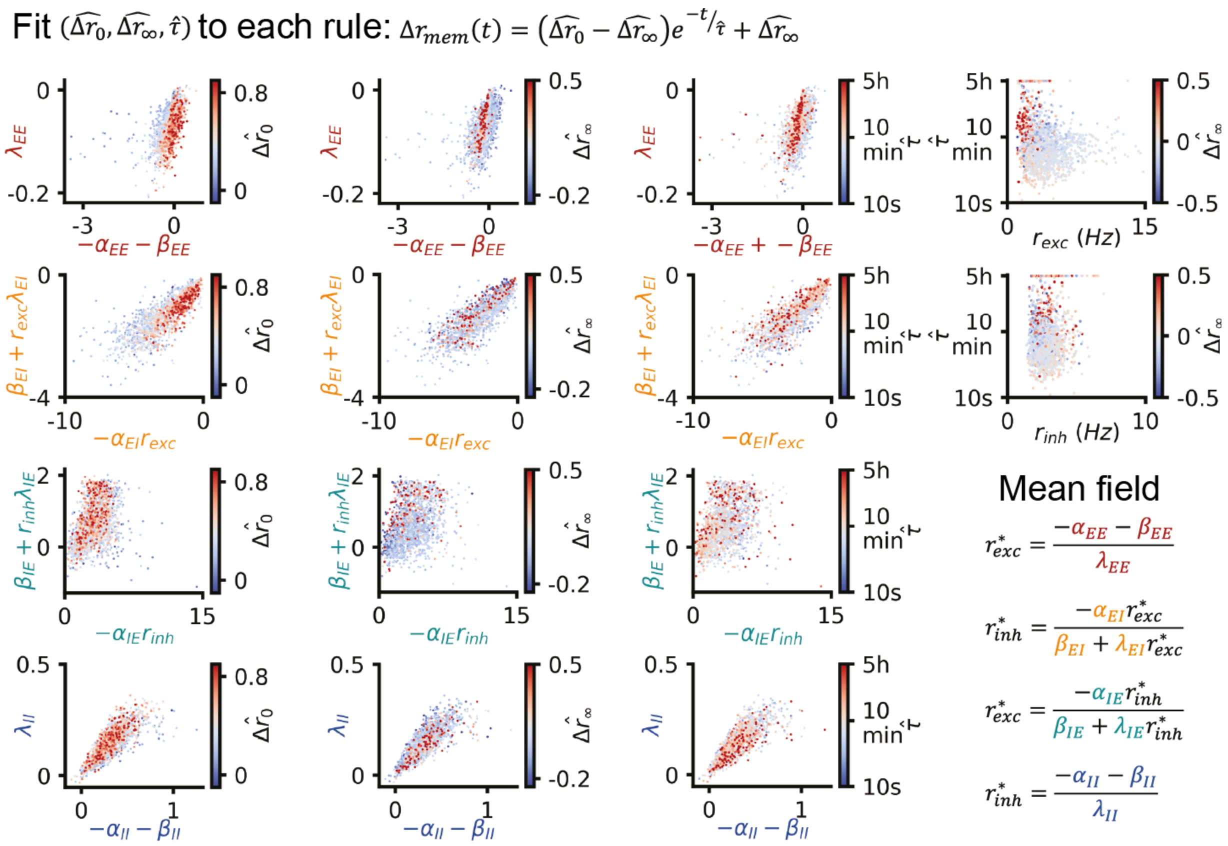


**Fig. Supp. 12. Predicting function from plasticity parameters.** For the 2,500 stable quadruplets from Fig. 1, we fit their responses to the familiarity task–$\{\Delta r_{\mathrm{mem}}(t_{\mathrm{break}})\}_{t_{\mathrm{break}} \in [1\mathrm{s}, 4\mathrm{h}]}$–to an exponential model with 3 parameters for each quadruplet. We then plot two plasticity parameters of each stable quadruplet and color by the the value of one fitted parameter.

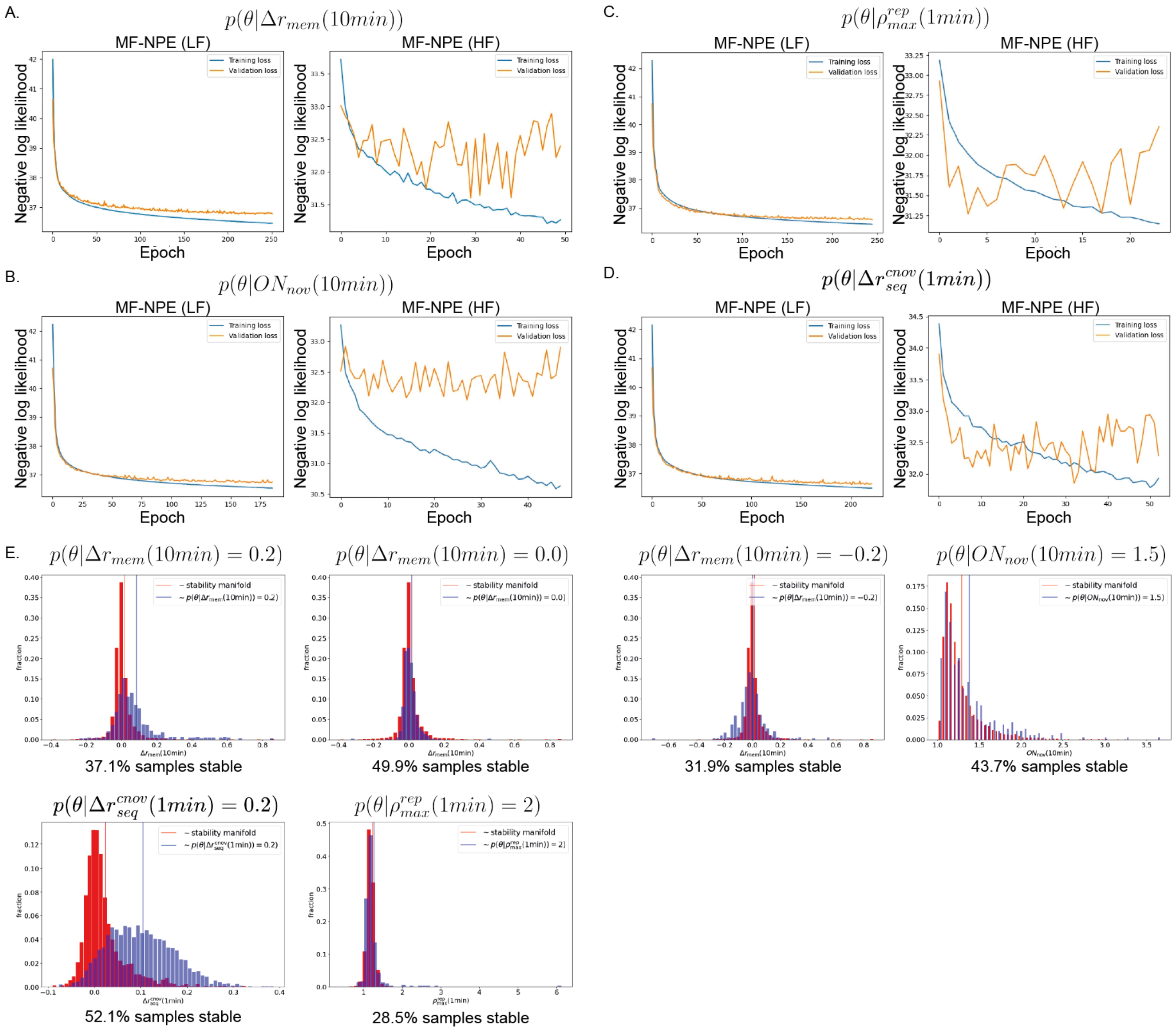


**Fig. Supp. 13. Posteriors on specific network functions. A-D**: Left: Training and validation loss for training a posterior on the low-fidelity samples (110k rule quadruplets stable for 2min obtained with fSBI[12]). Right: Training and validation loss of the second part of the posterior training, using the high fidelity samples (2,500 rule quadruplets tested to be stable after 4h). **E**: For each posterior, we sample 1000 quadruplets conditioned on a specific network metric value. Red: histogram of that metric value within the stability manifold, blue: among the 1000 samples.

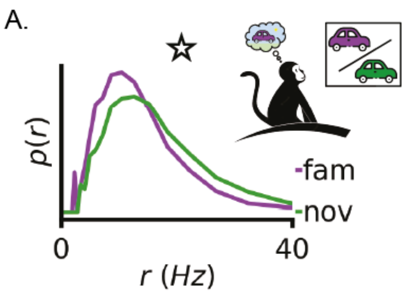


**Fig. Supp. 14. Experimental data. A**: Data[32,67], histogram of recorded neuron responses to familiar (purple) and novel (green) stimuli, recorded in ITC of monkeys. Novel stimuli elicit neural responses with higher mean and variance than familiar stimuli.

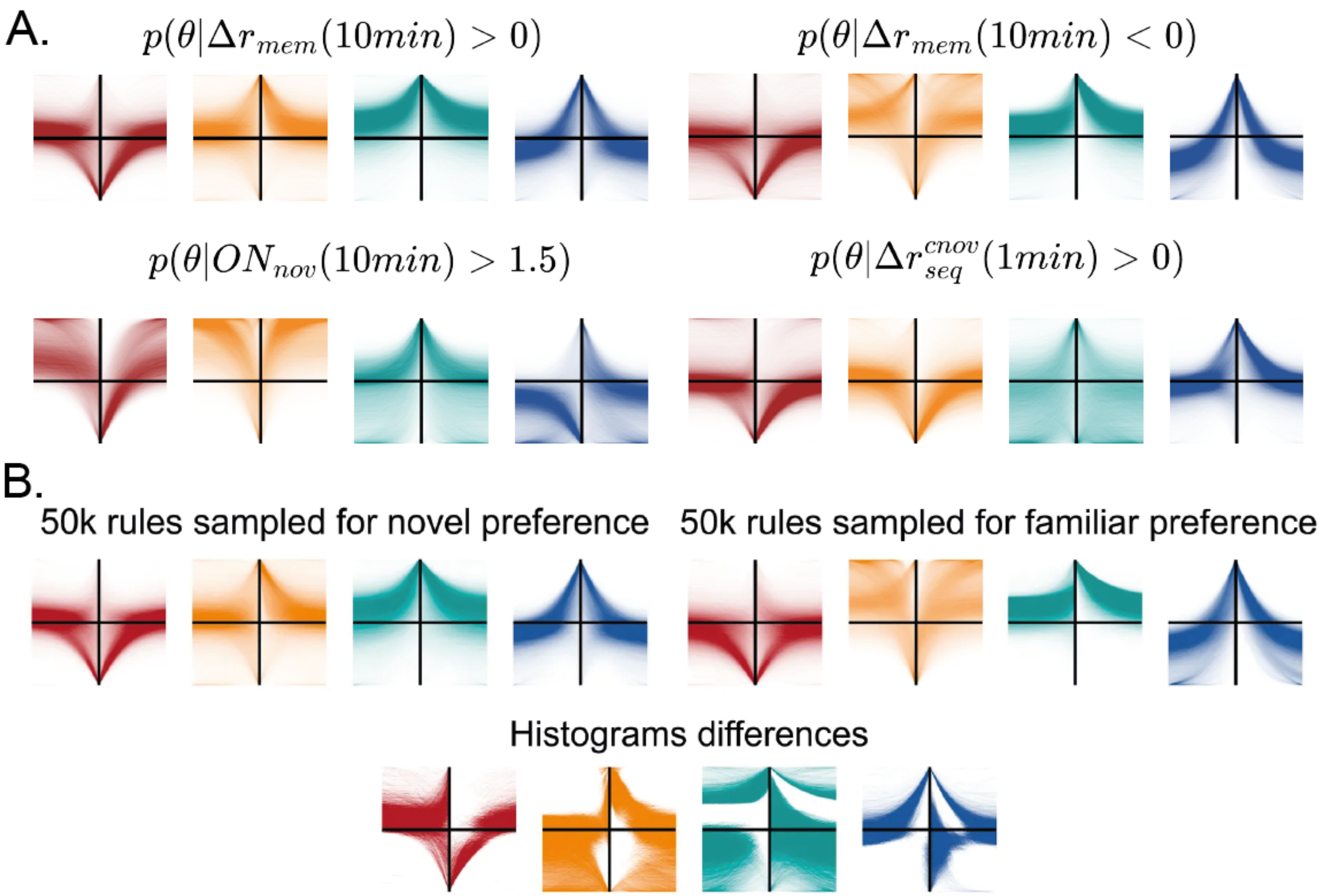


**Fig. Supp. 15. Analysis of the functional posteriors. A**: Sampling 50k rule quadruplets from four posteriors on several memory metrics. The 50k rules are represented as a histogram in pre-post protocol space. **B**: Taking the difference between two pre-post protocol histograms, showing in which parts of the pre-post protocol rule quadruplets differed most.

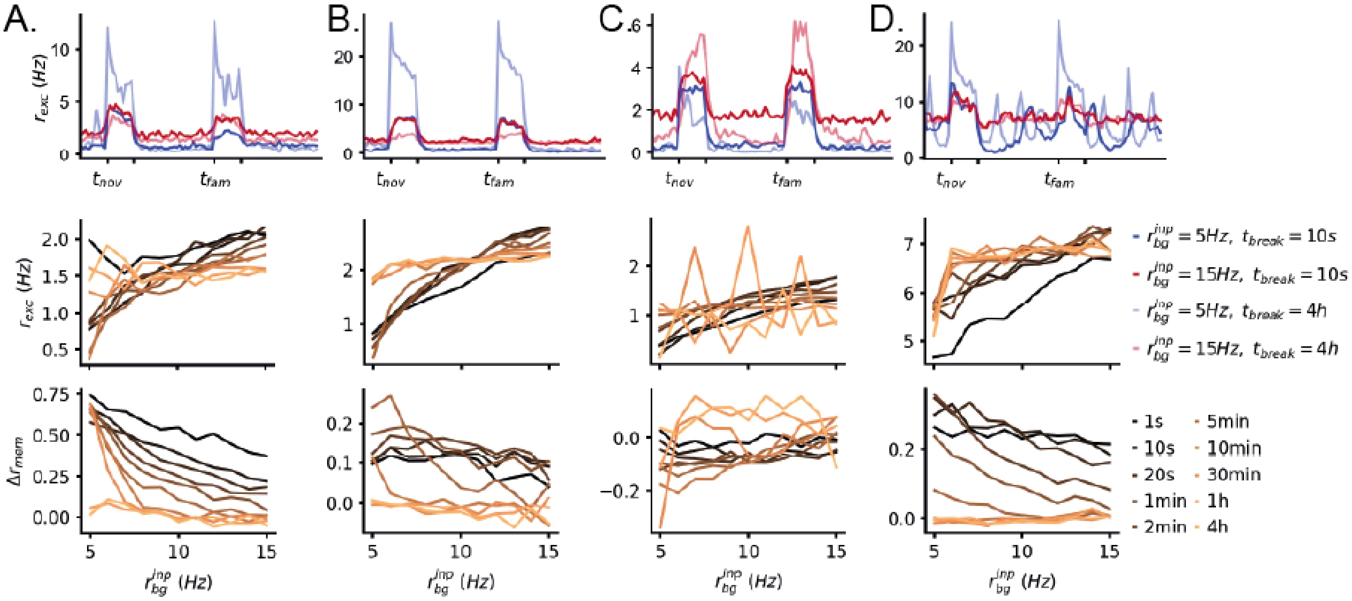


**Fig. Supp. 16. Dependency on background input rate. A**: One stable quadruplet simulated for a range of background input rates $r^{\text{bg}}_{\text{inp}} \in [5, 15]$Hz. Top: Excitatory (red) and inhibitory (blue) population firing rate during a test session of the familiarity task. Each stimulus presentation lasts 1s. Middle: evolution of the firing rate of the excitatory population with background inputs. Top: same but with the relative novel preference for different break periods. **B-D**: Same as A but for three other quadruplets

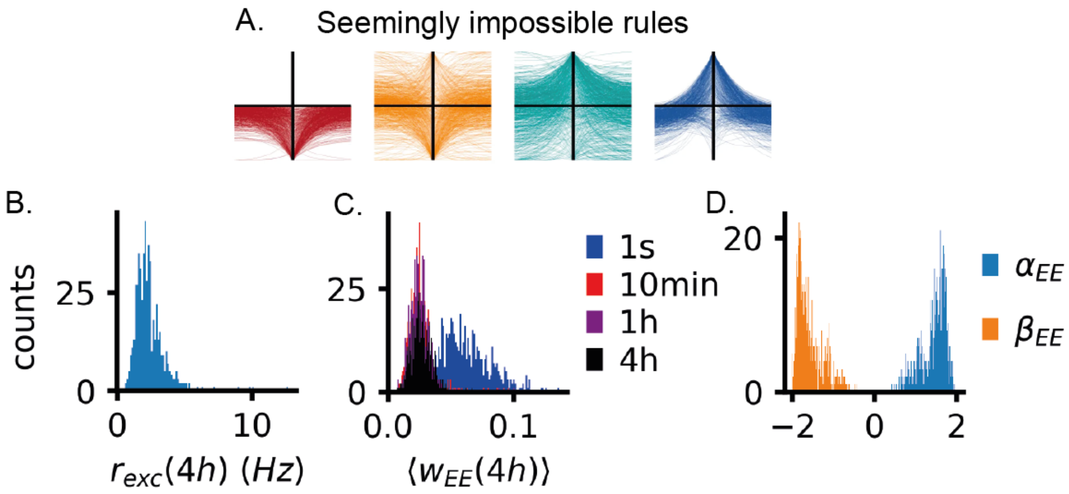


**Fig. Supp. 17. Analysis of meta-learned EE rules with counter intuitive shapes.** **A**: Histogram of quadruplet shapes that belong to the stability manifold, but which never trigger any potentiation on the pre-post protocol. **B**: Histogram of firing rate of networks evolving with these "negative EE" quadruplets. **C**: Histogram of the mean EE weights at different points in a network simulation in these networks. The weights are not going to zero, suggesting that these rules can potentiate the EE weights in practice. **D**: Histogram of the non-Hebbian EE parameters of these negative EE quadruplets. All of them have pre-only potentiation and post-only depression, which may explain why such rules can still undergo potentiation in practice.

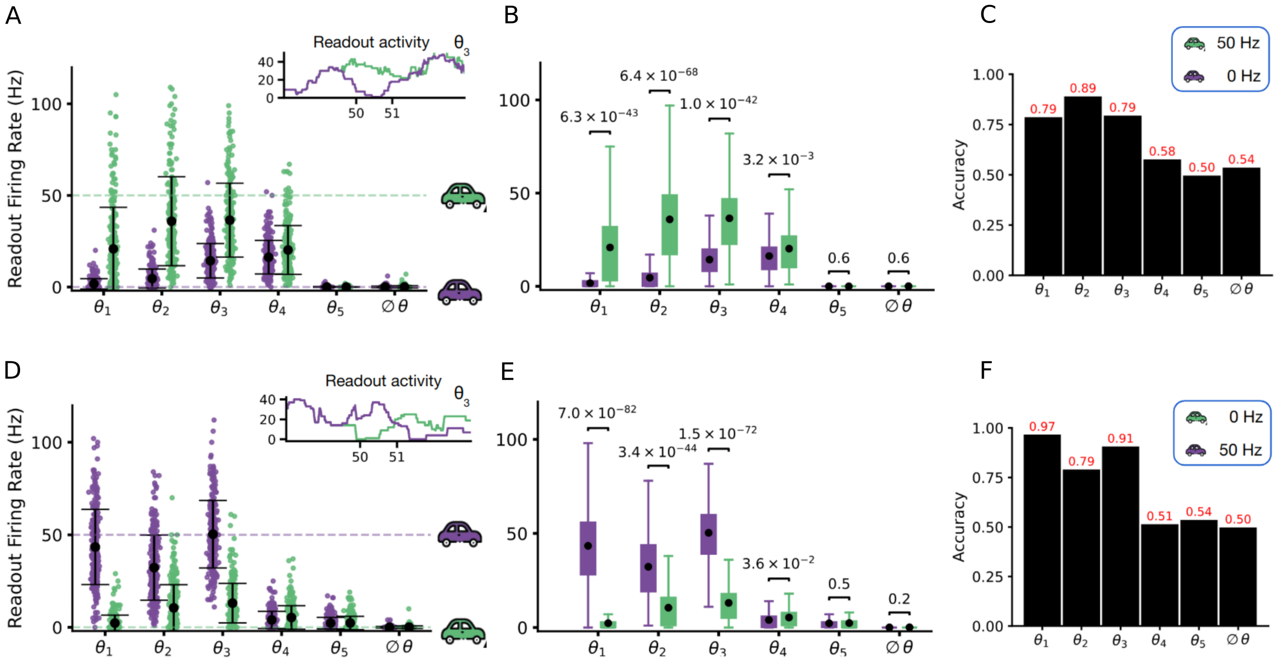


**Fig. Supp. 18. Reservoir computing 1: Novelty detection.** As a first, simple task to train a readout from the plastic recurrent networks, we trained networks to do novelty detection with desired firing population rates in response to novel and familiar stimuli. **A**: Firing rate of the readout LIF neuron in the trials with novel (green) and familiar (purple) stimuli for five different plasticity rules and one static network. Inset: Example of activity of the LIF neuron when a novel or familiar stimulus was presented. **B**: Firing rate of the readout LIF neuron in the trials with novel (green) and familiar (purple) stimuli with statistical p-values. **C**: Maximum accuracy obtained for every network. Inset: target firing rate of the neuron for two different stimuli. **D, E, F**: The same analysis for the reversed target firing rate conditions.

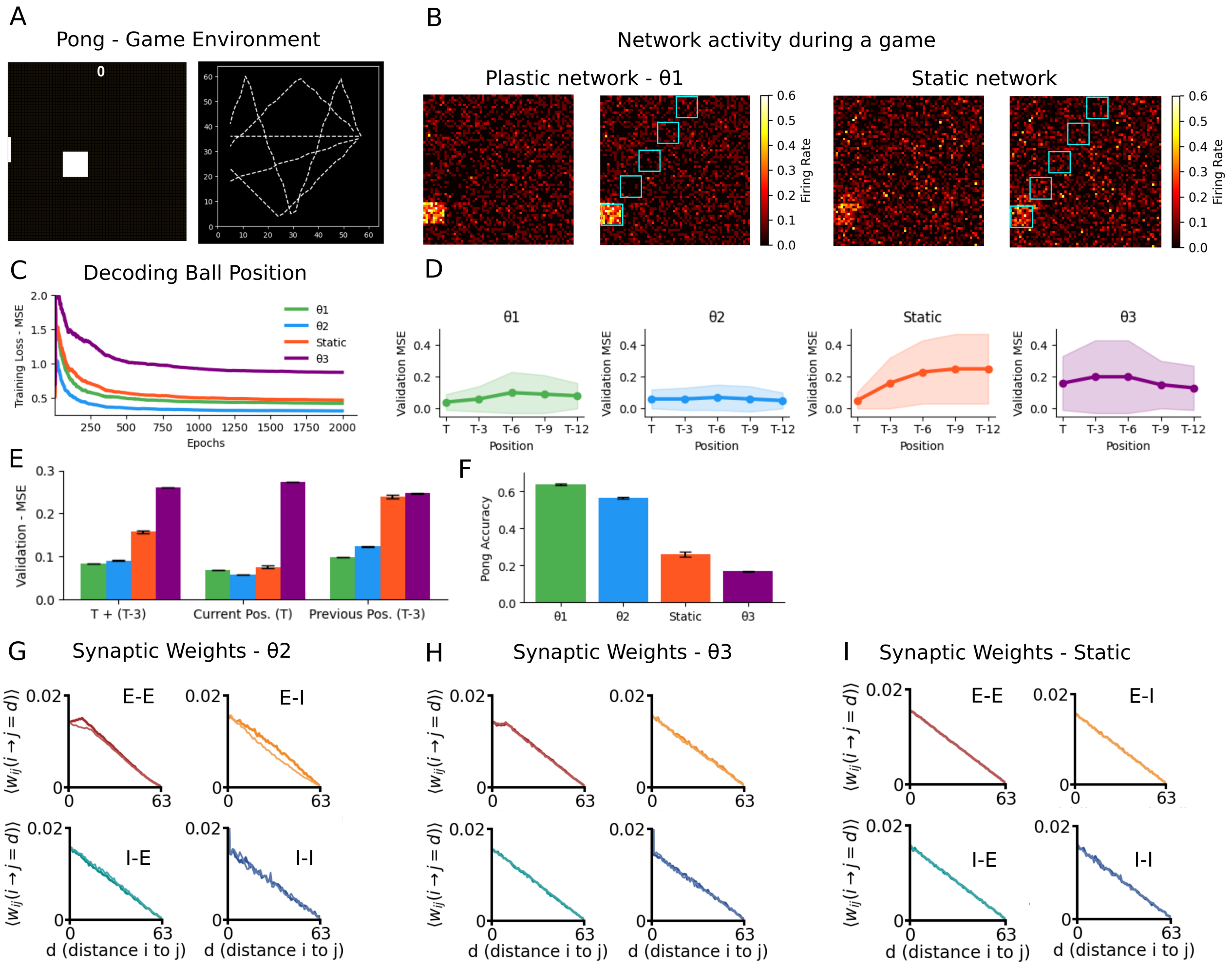


**Fig. Supp. 19. Reservoir computing 2: Pong A**: Game environment. Left: a screenshot of the game. Right: example of 5 different trajectories of the ball. **B**: Firing rate of the network during a game for a plastic (left) and a static (right) network. Blue squares show the current and four previous positions of the ball. **C**: Decoding current and previous positions of the ball. Training loss for 2000 trials (epochs). **D**: Validation loss for three plastic networks and one static network for decoding current and four previous positions. **E**: Decoding the ball's position to predict the ball's direction during a pong game. Validation loss for decoding current and last previous position. **F**: Pong accuracy as the number of successful trials divided by the size of the validation phase. **G, H, I**: Synaptic weights distributions for two plastic networks and one static network.

Supplementary Figure 20: Documentation of all sampled rule quadruplets.

**Performance scores** are shown as percentages compared to the best quadruplet within the dataset for a given task. Significant task responses are displayed in green, non-significant changes in black.

**Rule shapes** are a visualization of the individual rules of a quadruplet (EE, EI, IE, II). Given a pair of pre- and post-synaptic spikes with a given temporal lag (x-axis), efficacy change of a synapse (y-axis).

The **star tally** conveys overall task performance: ⭐ exceptional quadruplet for the task (criteria from Fig. 2), ⚪ quadruplet with at least one significant response over the break period. The symbols also appear beside each task and are ordered as: Novelty, Transience, Replay and Omission.

**ID** of the rule quadruplet.

**Figure** in which the rule quadruplet features.

The **novelty** panel shows each quadruplet's performance in the novelty detection task (Fig. 1D), after a short (20s, ⏳) or long (4h, ⌛) break with only background activity.

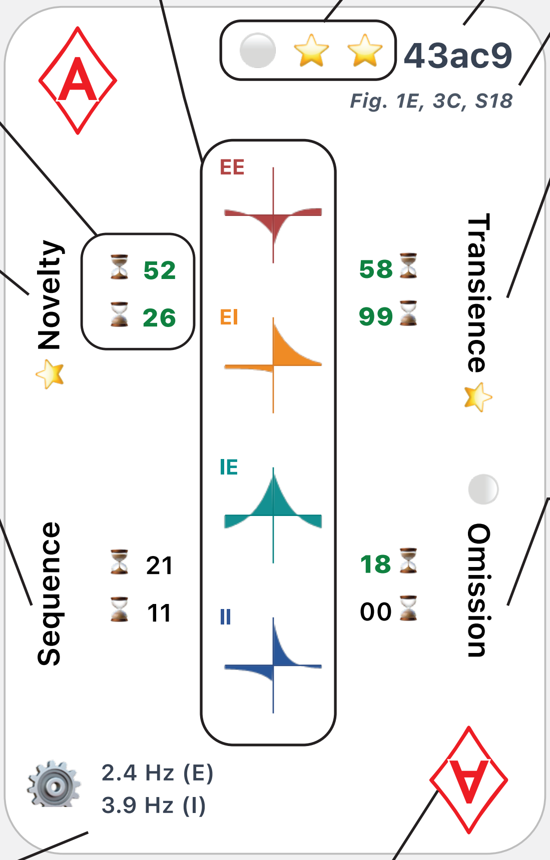


The **transience** panel highlights each quadruplet's ability to produce ON responses (transient dynamics at stimulus onset) in the novelty detection task (Fig. 1E), after a short (20s, ⏳) or long (4h, ⌛) break.

The **sequence** panel shows each quadruplet's performance on the sequential task (Fig. 1F). The score assesses the quadruplet's ability to recall the training sequence after a short (20s, ⏳) or long (10min, ⌛) break.

The **omission** panel shows each quadruplet's performance on the contextual novelty task (Fig. 1G), also known as omission novelty, recall the training sequence after a short (20s, ⏳) or long (10min, ⌛) break.

The **baseline firing rate** (⚙️) corresponds to the excitatory and inhibitory population firing rates (in Hz) after 4h of simulation.

The **suit** in the diagonal corners displays the conventional rank (A–2) and suit (♥ ♠ ♦ ♣) used to sort rules into physical decks.These assignments were made based on which of the four functions the quadruplet at hand responded most to.

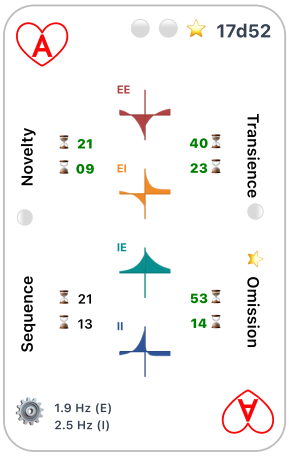
17d52
Novelty
21
09
Transience
40
23
EE
EI
IE
II
Sequence
21
13
Omission
53
14
1.9 Hz (E)
2.5 Hz (I)

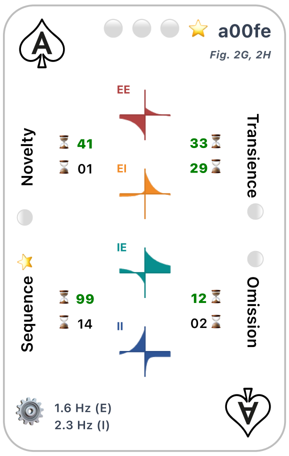
a00fe
Fig. 2G, 2H
Novelty
41
01
Transience
33
29
EE
EI
IE
II
Sequence
99
14
Omission
12
02
1.6 Hz (E)
2.3 Hz (I)

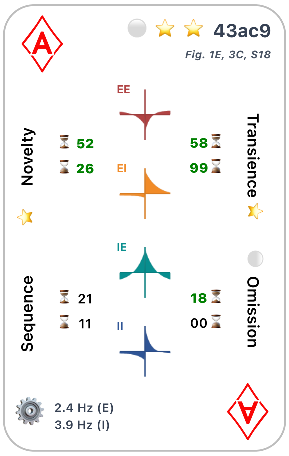
43ac9
Fig. 1E, 3C, S18
Novelty
52
26
Transience
58
99
EE
EI
IE
II
Sequence
21
11
Omission
18
00
2.4 Hz (E)
3.9 Hz (I)

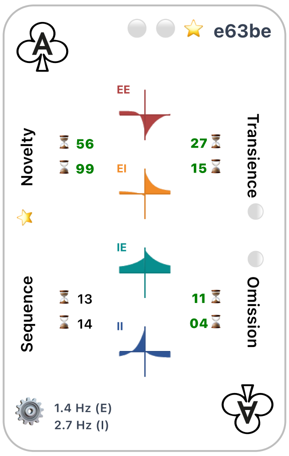
e63be
Novelty
56
99
Transience
27
15
EE
EI
IE
II
Sequence
13
14
Omission
11
04
1.4 Hz (E)
2.7 Hz (I)

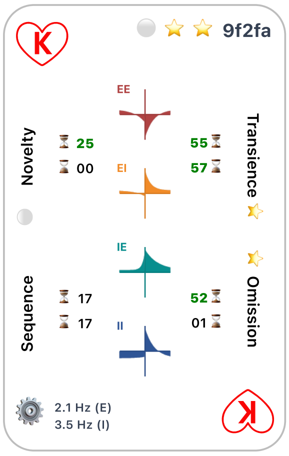
9f2fa
Novelty
25
00
Transience
55
57
EE
EI
IE
II
Sequence
17
17
Omission
52
01
2.1 Hz (E)
3.5 Hz (I)

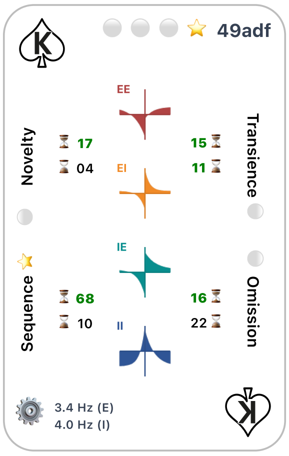
49adf
Novelty
17
04
Transience
15
11
EE
EI
IE
II
Sequence
68
10
Omission
16
22
3.4 Hz (E)
4.0 Hz (I)

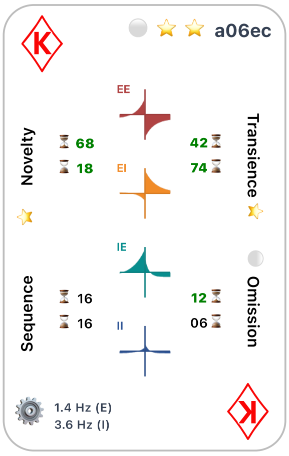
a06ec
Novelty
68
18
Transience
42
74
EE
EI
IE
II
Sequence
16
16
Omission
12
06
1.4 Hz (E)
3.6 Hz (I)

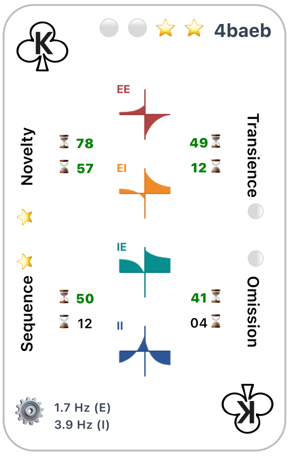
4baeb
Novelty
78
57
Transience
49
12
EE
EI
IE
II
Sequence
50
12
Omission
41
04
1.7 Hz (E)
3.9 Hz (I)

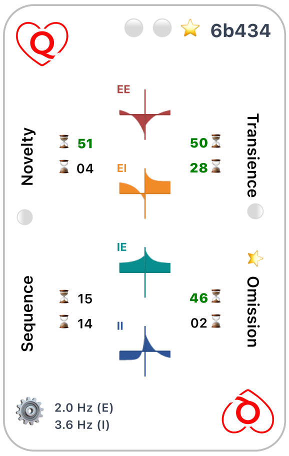
6b434
Novelty
51
04
Transience
50
28
EE
EI
IE
II
Sequence
15
14
Omission
46
02
2.0 Hz (E)
3.6 Hz (I)

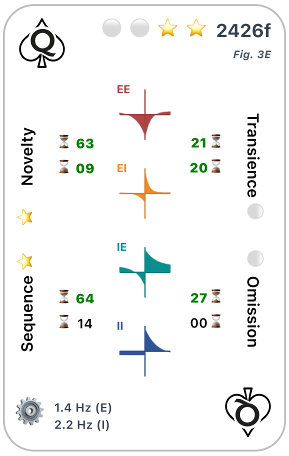
2426f
Fig. 3E
Novelty
63
09
Transience
21
20
EE
EI
IE
II
Sequence
64
14
Omission
27
00
1.4 Hz (E)
2.2 Hz (I)

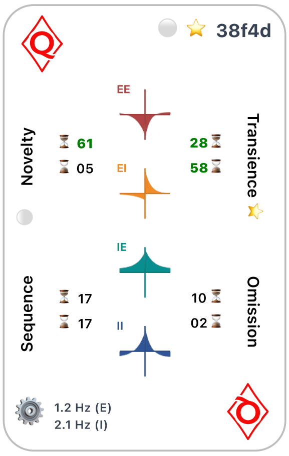
38f4d
Novelty
61
05
Transience
28
58
EE
EI
IE
II
Sequence
17
17
Omission
10
02
1.2 Hz (E)
2.1 Hz (I)

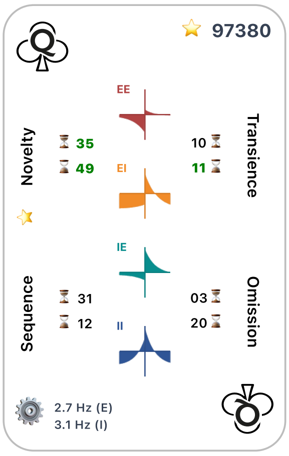
97380
Novelty
35
49
Transience
10
11
EE
EI
IE
II
Sequence
31
12
Omission
03
20
2.7 Hz (E)
3.1 Hz (I)

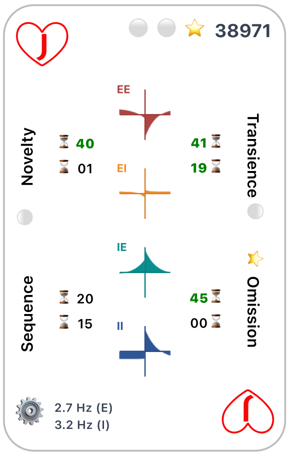
38971
Novelty
40
01
Transience
41
19
EE
EI
IE
II
Sequence
20
15
Omission
45
00
2.7 Hz (E)
3.2 Hz (I)

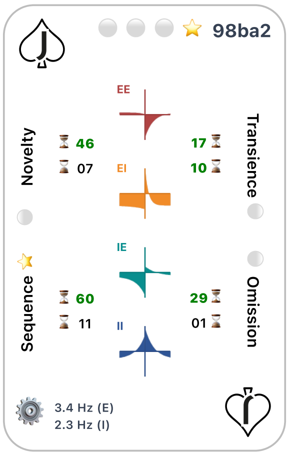
98ba2
Novelty
46
07
Transience
17
10
EE
EI
IE
II
Sequence
60
11
Omission
29
01
3.4 Hz (E)
2.3 Hz (I)

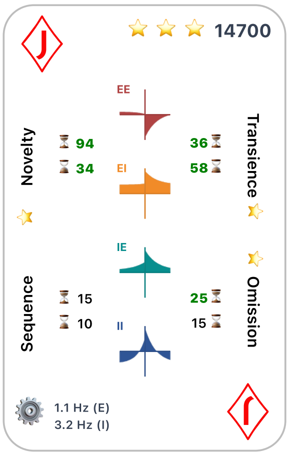
14700
Novelty
94
34
Transience
36
58
EE
EI
IE
II
Sequence
15
10
Omission
25
15
1.1 Hz (E)
3.2 Hz (I)

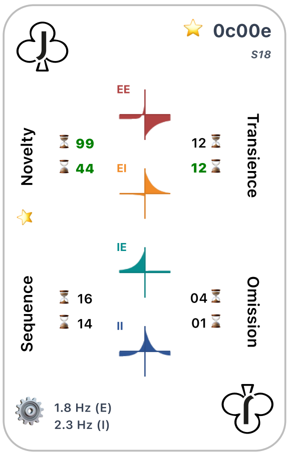
0c00e
S18
Novelty
99
44
Transience
12
12
EE
EI
IE
II
Sequence
16
14
Omission
04
01
1.8 Hz (E)
2.3 Hz (I)

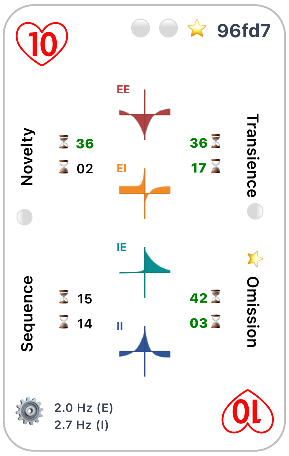
10
96fd7
EE
Novelty
36
02
36
17
Transience
EI
IE
Sequence
15
14
42
03
Omission
II
2.0 Hz (E)
2.7 Hz (I)

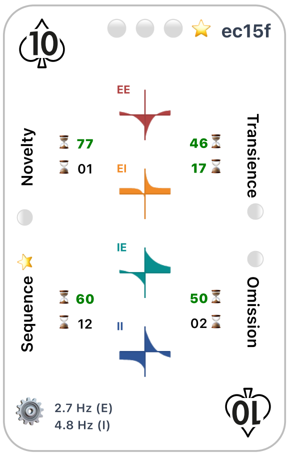
10
ec15f
EE
Novelty
77
01
46
17
Transience
EI
IE
Sequence
60
12
50
02
Omission
II
2.7 Hz (E)
4.8 Hz (I)

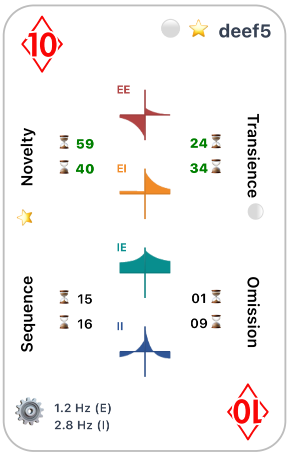
10
deef5
EE
Novelty
59
40
24
34
Transience
EI
IE
Sequence
15
16
01
09
Omission
II
1.2 Hz (E)
2.8 Hz (I)

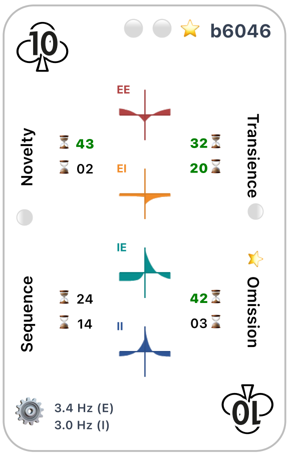
10
b6046
EE
Novelty
43
02
32
20
Transience
EI
IE
Sequence
24
14
42
03
Omission
II
3.4 Hz (E)
3.0 Hz (I)

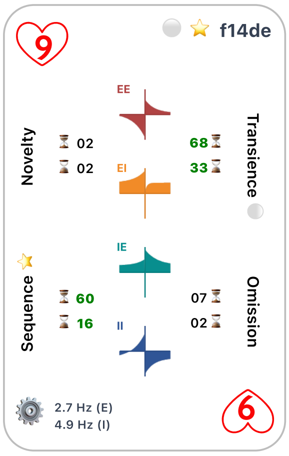
9
f14de
EE
Novelty
02
02
68
33
Transience
EI
IE
Sequence
60
16
07
02
Omission
II
2.7 Hz (E)
4.9 Hz (I)

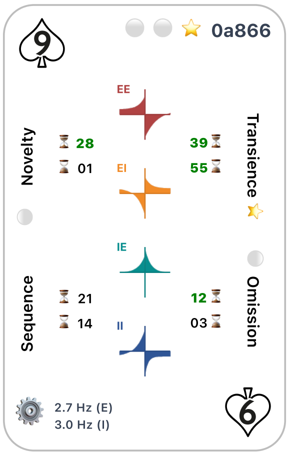
9
0a866
EE
Novelty
28
01
39
55
Transience
EI
IE
Sequence
21
14
12
03
Omission
II
2.7 Hz (E)
3.0 Hz (I)

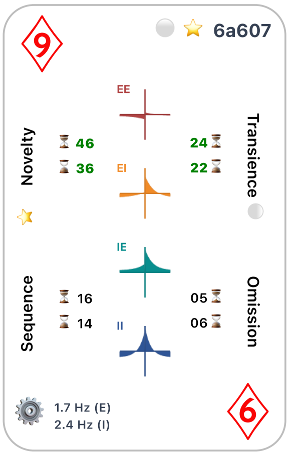
9
6a607
EE
Novelty
46
36
24
22
Transience
EI
IE
Sequence
16
14
05
06
Omission
II
1.7 Hz (E)
2.4 Hz (I)

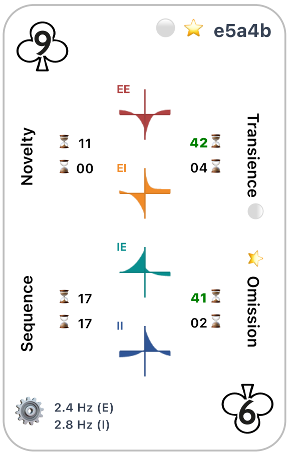
9
e5a4b
EE
Novelty
11
00
42
04
Transience
EI
IE
Sequence
17
17
41
02
Omission
II
2.4 Hz (E)
2.8 Hz (I)

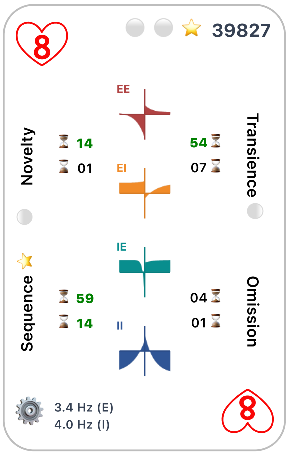
8
39827
EE
Novelty
14
01
54
07
Transience
EI
IE
Sequence
59
14
04
01
Omission
II
3.4 Hz (E)
4.0 Hz (I)

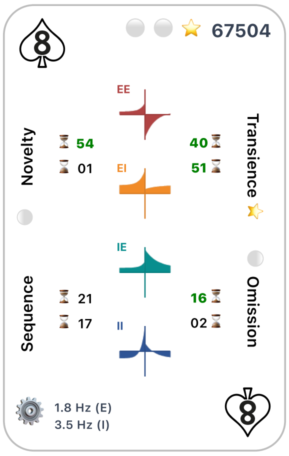
8
67504
EE
Novelty
54
01
40
51
Transience
EI
IE
Sequence
21
17
16
02
Omission
II
1.8 Hz (E)
3.5 Hz (I)

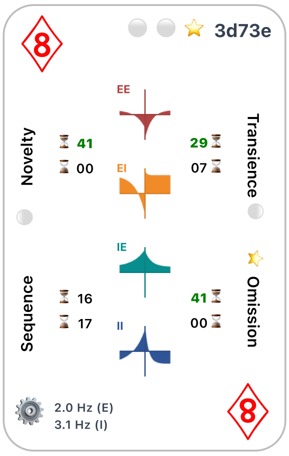
8
3d73e
EE
Novelty
41
00
29
07
Transience
EI
IE
Sequence
16
17
41
00
Omission
II
2.0 Hz (E)
3.1 Hz (I)

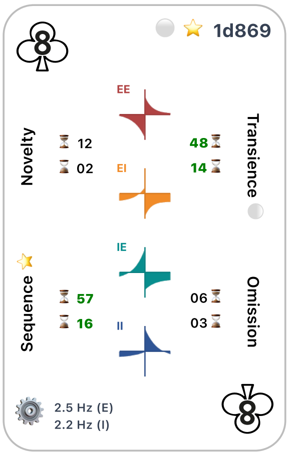
8
1d869
EE
Novelty
12
02
48
14
Transience
EI
IE
Sequence
57
16
06
03
Omission
II
2.5 Hz (E)
2.2 Hz (I)

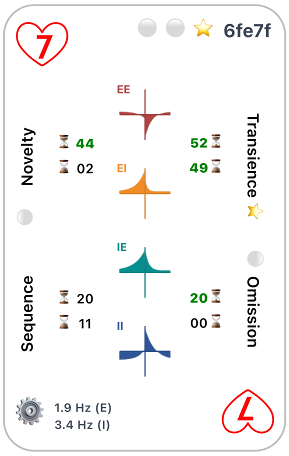
7
6fe7f
EE
Novelty
44
02
52
49
Transience
EI
IE
Sequence
20
11
20
00
Omission
II
1.9 Hz (E)
3.4 Hz (I)

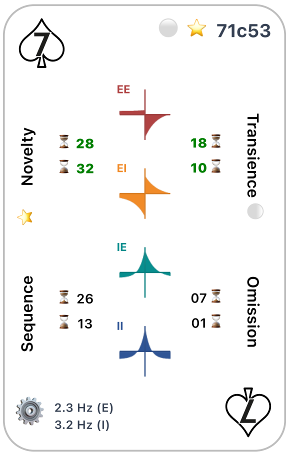
7
71c53
EE
Novelty
28
32
18
10
Transience
EI
IE
Sequence
26
13
07
01
Omission
II
2.3 Hz (E)
3.2 Hz (I)

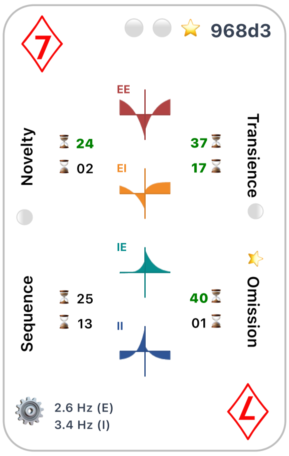
7
968d3
EE
Novelty
24
02
37
17
Transience
EI
IE
Sequence
25
13
40
01
Omission
II
2.6 Hz (E)
3.4 Hz (I)

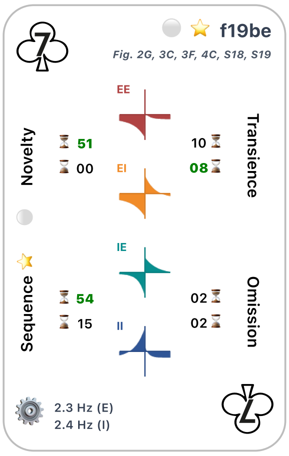
7
f19be
Fig. 2G, 3C, 3F, 4C, S18, S19
EE
Novelty
51
00
10
08
Transience
EI
IE
Sequence
54
15
02
02
Omission
II
2.3 Hz (E)
2.4 Hz (I)

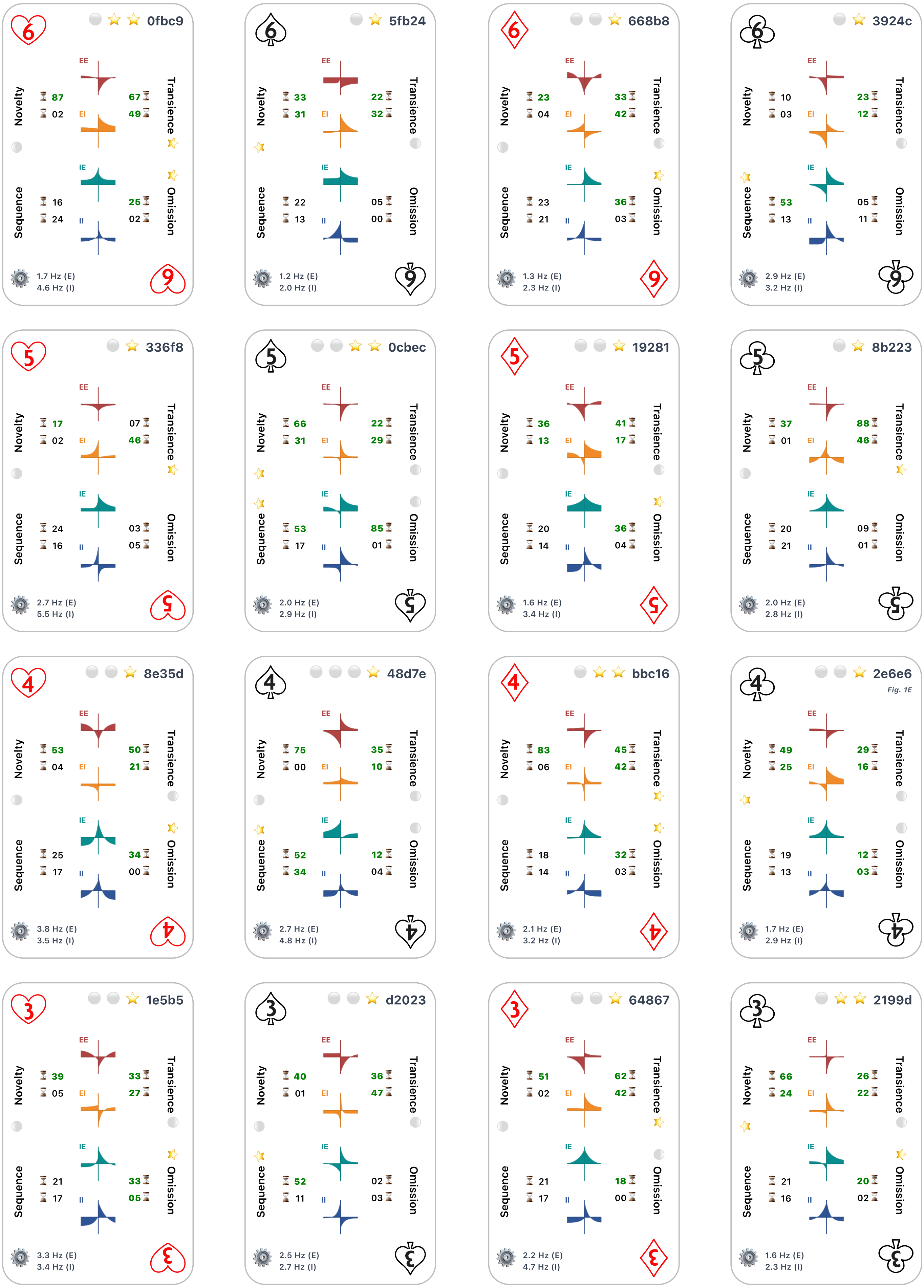
6 0fbc9
EE EI IE II
Novelty 87 02 67 49 Transience
Sequence 16 24 25 02 Omission
1.7 Hz (E) 4.6 Hz (I)
9
6 5fb24
Novelty 33 31 22 32 Transience
Sequence 22 13 05 00 Omission
1.2 Hz (E) 2.0 Hz (I)
9
6 668b8
Novelty 23 04 33 42 Transience
Sequence 23 21 36 03 Omission
1.3 Hz (E) 2.3 Hz (I)
9
6 3924c
Novelty 10 03 23 12 Transience
Sequence 53 13 05 11 Omission
2.9 Hz (E) 3.2 Hz (I)
9
5 336f8
Novelty 17 02 07 46 Transience
Sequence 24 16 03 05 Omission
2.7 Hz (E) 5.5 Hz (I)
5
5 0cbec
Novelty 66 31 22 29 Transience
Sequence 53 17 85 01 Omission
2.0 Hz (E) 2.9 Hz (I)
5
5 19281
Novelty 36 13 41 17 Transience
Sequence 20 14 36 04 Omission
1.6 Hz (E) 3.4 Hz (I)
5
5 8b223
Novelty 37 01 88 46 Transience
Sequence 20 21 09 01 Omission
2.0 Hz (E) 2.8 Hz (I)
5
4 8e35d
Novelty 53 04 50 21 Transience
Sequence 25 17 34 00 Omission
3.8 Hz (E) 3.5 Hz (I)
4
4 48d7e
Novelty 75 00 35 10 Transience
Sequence 52 34 12 04 Omission
2.7 Hz (E) 4.8 Hz (I)
4
4 bbc16
Novelty 83 06 45 42 Transience
Sequence 18 14 32 03 Omission
2.1 Hz (E) 3.2 Hz (I)
4
4 2e6e6
Fig. 1E
Novelty 49 25 29 16 Transience
Sequence 19 13 12 03 Omission
1.7 Hz (E) 2.9 Hz (I)
4
3 1e5b5
Novelty 39 05 33 27 Transience
Sequence 21 17 33 05 Omission
3.3 Hz (E) 3.4 Hz (I)
3
3 d2023
Novelty 40 01 36 47 Transience
Sequence 52 11 02 03 Omission
2.5 Hz (E) 2.7 Hz (I)
3
3 64867
Novelty 51 02 62 42 Transience
Sequence 21 17 18 00 Omission
2.2 Hz (E) 4.7 Hz (I)
3
3 2199d
Novelty 66 24 26 22 Transience
Sequence 21 16 20 02 Omission
1.6 Hz (E) 2.3 Hz (I)
3

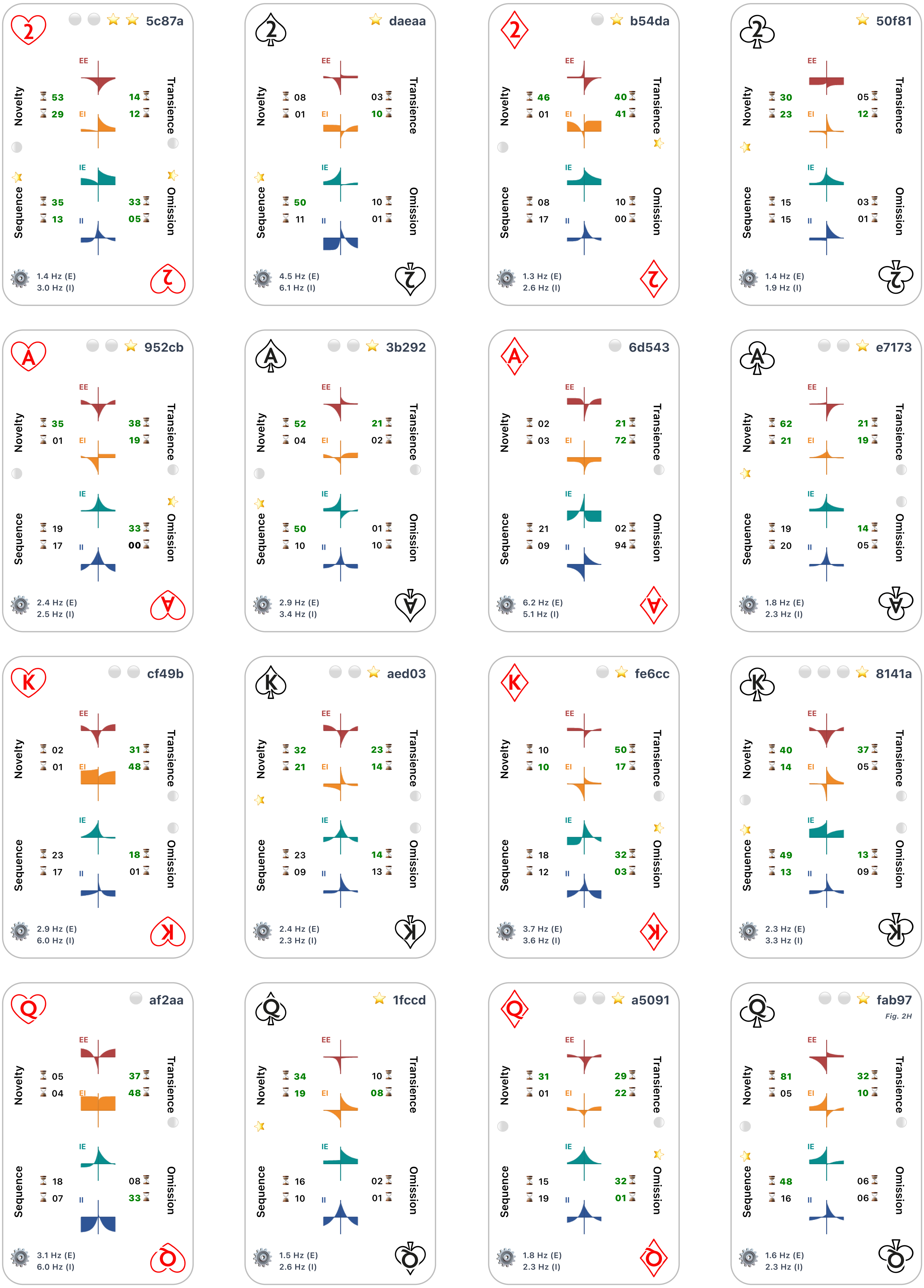

2 5c87a Novelty 53 29 Transience 14 12 Sequence 35 13 Omission 33 05 EE EI IE II 1.4 Hz (E) 3.0 Hz (I)
2 daeaa Novelty 08 01 Transience 03 10 Sequence 50 11 Omission 10 01 EE EI IE II 4.5 Hz (E) 6.1 Hz (I)
2 b54da Novelty 46 01 Transience 40 41 Sequence 08 17 Omission 10 00 EE EI IE II 1.3 Hz (E) 2.6 Hz (I)
2 50f81 Novelty 30 23 Transience 05 12 Sequence 15 15 Omission 03 01 EE EI IE II 1.4 Hz (E) 1.9 Hz (I)
A 952cb Novelty 35 01 Transience 38 19 Sequence 19 17 Omission 33 00 EE EI IE II 2.4 Hz (E) 2.5 Hz (I)
A 3b292 Novelty 52 04 Transience 21 02 Sequence 50 10 Omission 01 10 EE EI IE II 2.9 Hz (E) 3.4 Hz (I)
A 6d543 Novelty 02 03 Transience 21 72 Sequence 21 09 Omission 02 94 EE EI IE II 6.2 Hz (E) 5.1 Hz (I)
A e7173 Novelty 62 21 Transience 21 19 Sequence 19 20 Omission 14 05 EE EI IE II 1.8 Hz (E) 2.3 Hz (I)
K cf49b Novelty 02 01 Transience 31 48 Sequence 23 17 Omission 18 01 EE EI IE II 2.9 Hz (E) 6.0 Hz (I)
K aed03 Novelty 32 21 Transience 23 14 Sequence 23 09 Omission 14 13 EE EI IE II 2.4 Hz (E) 2.3 Hz (I)
K fe6cc Novelty 10 10 Transience 50 17 Sequence 18 12 Omission 32 03 EE EI IE II 3.7 Hz (E) 3.6 Hz (I)
K 8141a Novelty 40 14 Transience 37 05 Sequence 49 13 Omission 13 09 EE EI IE II 2.3 Hz (E) 3.3 Hz (I)
Q af2aa Novelty 05 04 Transience 37 48 Sequence 18 07 Omission 08 33 EE EI IE II 3.1 Hz (E) 6.0 Hz (I)
Q 1fccd Novelty 34 19 Transience 10 08 Sequence 16 10 Omission 02 01 EE EI IE II 1.5 Hz (E) 2.6 Hz (I)
Q a5091 Novelty 31 01 Transience 29 22 Sequence 15 19 Omission 32 01 EE EI IE II 1.8 Hz (E) 2.3 Hz (I)
Q fab97 Fig. 2H Novelty 81 05 Transience 32 10 Sequence 48 16 Omission 06 06 EE EI IE II 1.6 Hz (E) 2.3 Hz (I)

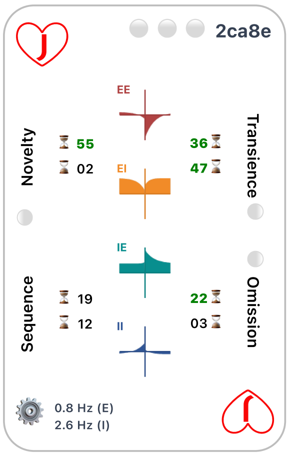
2ca8e
EE
EI
IE
II
Novelty 55 02
Transience 36 47
Sequence 19 12
Omission 22 03
0.8 Hz (E)
2.6 Hz (I)

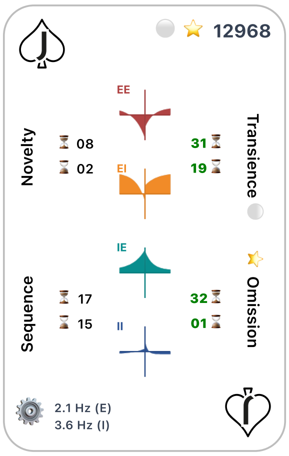
12968
EE
EI
IE
II
Novelty 08 02
Transience 31 19
Sequence 17 15
Omission 32 01
2.1 Hz (E)
3.6 Hz (I)

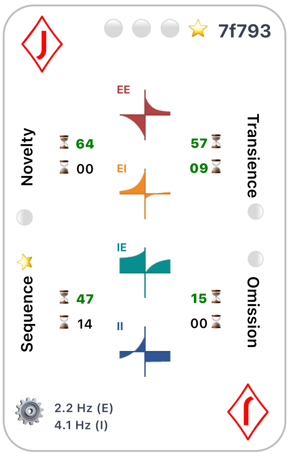
7f793
EE
EI
IE
II
Novelty 64 00
Transience 57 09
Sequence 47 14
Omission 15 00
2.2 Hz (E)
4.1 Hz (I)

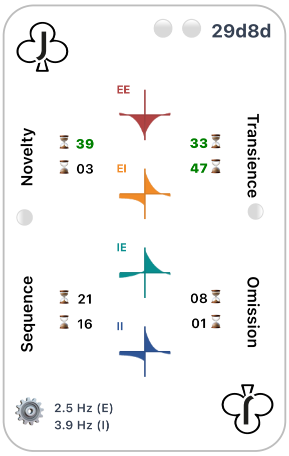
29d8d
EE
EI
IE
II
Novelty 39 03
Transience 33 47
Sequence 21 16
Omission 08 01
2.5 Hz (E)
3.9 Hz (I)

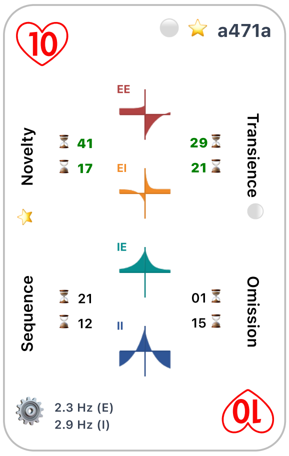
a471a
EE
EI
IE
II
Novelty 41 17
Transience 29 21
Sequence 21 12
Omission 01 15
2.3 Hz (E)
2.9 Hz (I)

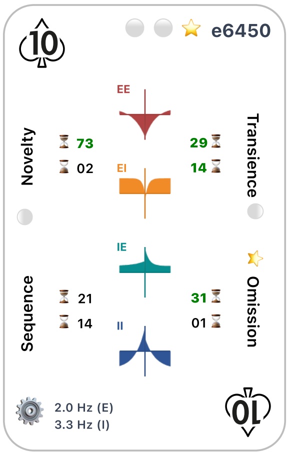
e6450
EE
EI
IE
II
Novelty 73 02
Transience 29 14
Sequence 21 14
Omission 31 01
2.0 Hz (E)
3.3 Hz (I)

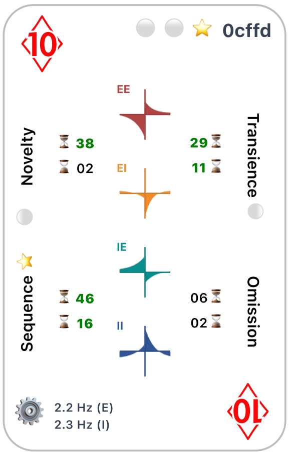
0cffd
EE
EI
IE
II
Novelty 38 02
Transience 29 11
Sequence 46 16
Omission 06 02
2.2 Hz (E)
2.3 Hz (I)

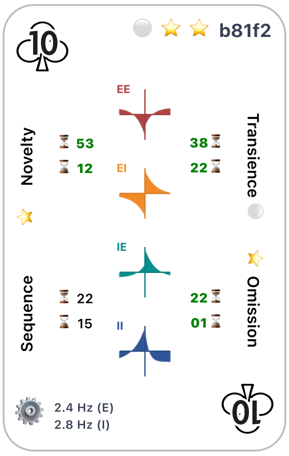
b81f2
EE
EI
IE
II
Novelty 53 12
Transience 38 22
Sequence 22 15
Omission 22 01
2.4 Hz (E)
2.8 Hz (I)

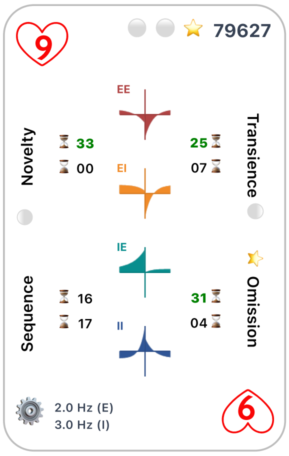
79627
EE
EI
IE
II
Novelty 33 00
Transience 25 07
Sequence 16 17
Omission 31 04
2.0 Hz (E)
3.0 Hz (I)

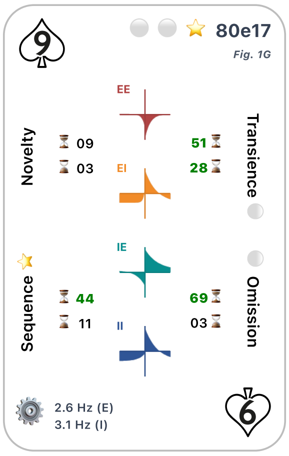
80e17
Fig. 1G
EE
EI
IE
II
Novelty 09 03
Transience 51 28
Sequence 44 11
Omission 69 03
2.6 Hz (E)
3.1 Hz (I)

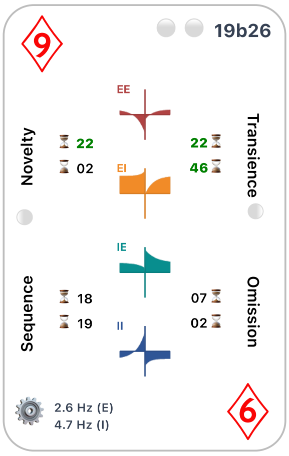
19b26
EE
EI
IE
II
Novelty 22 02
Transience 22 46
Sequence 18 19
Omission 07 02
2.6 Hz (E)
4.7 Hz (I)

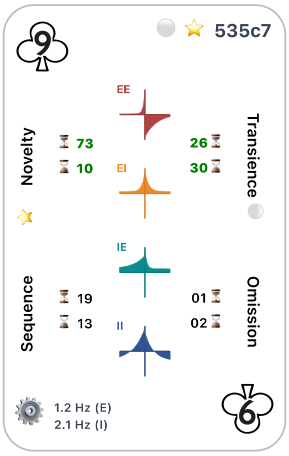
535c7
EE
EI
IE
II
Novelty 73 10
Transience 26 30
Sequence 19 13
Omission 01 02
1.2 Hz (E)
2.1 Hz (I)

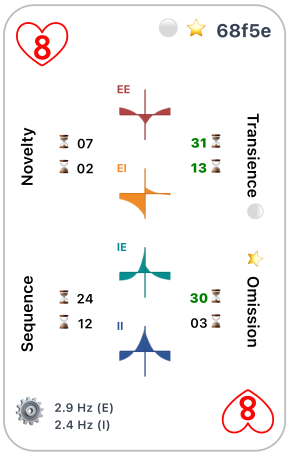
68f5e
EE
EI
IE
II
Novelty 07 02
Transience 31 13
Sequence 24 12
Omission 30 03
2.9 Hz (E)
2.4 Hz (I)

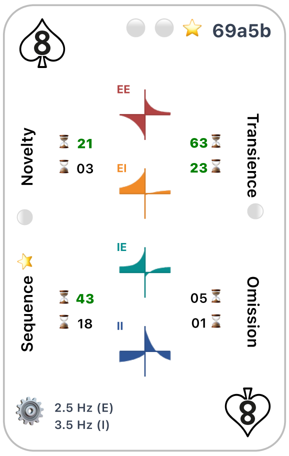
69a5b
EE
EI
IE
II
Novelty 21 03
Transience 63 23
Sequence 43 18
Omission 05 01
2.5 Hz (E)
3.5 Hz (I)

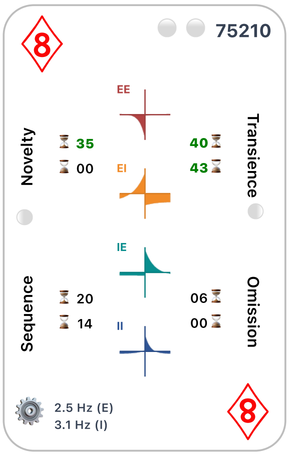
75210
EE
EI
IE
II
Novelty 35 00
Transience 40 43
Sequence 20 14
Omission 06 00
2.5 Hz (E)
3.1 Hz (I)

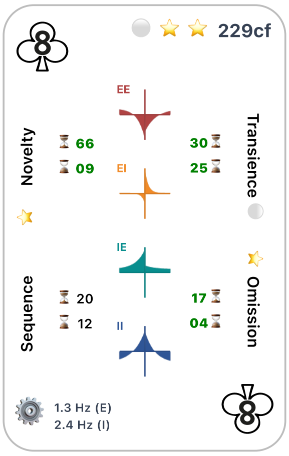
229cf
EE
EI
IE
II
Novelty 66 09
Transience 30 25
Sequence 20 12
Omission 17 04
1.3 Hz (E)
2.4 Hz (I)

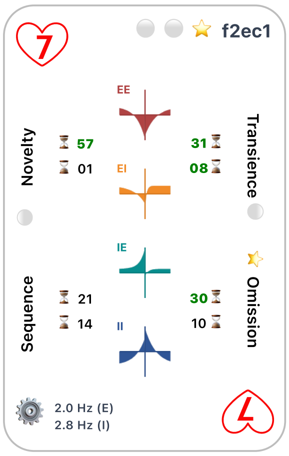
f2ec1
EE
EI
IE
II
Novelty
57
01
Transience
31
08
Sequence
21
14
Omission
30
10
2.0 Hz (E)
2.8 Hz (I)

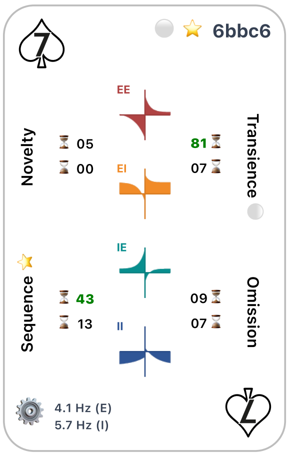
6bbc6
EE
EI
IE
II
Novelty
05
00
Transience
81
07
Sequence
43
13
Omission
09
07
4.1 Hz (E)
5.7 Hz (I)

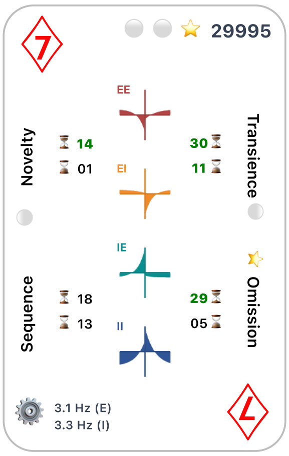
29995
EE
EI
IE
II
Novelty
14
01
Transience
30
11
Sequence
18
13
Omission
29
05
3.1 Hz (E)
3.3 Hz (I)

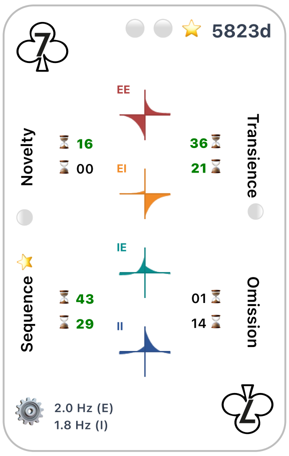
5823d
EE
EI
IE
II
Novelty
16
00
Transience
36
21
Sequence
43
29
Omission
01
14
2.0 Hz (E)
1.8 Hz (I)

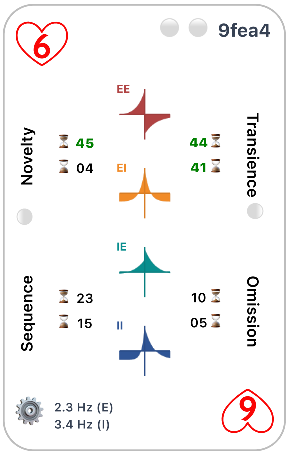
9fea4
EE
EI
IE
II
Novelty
45
04
Transience
44
41
Sequence
23
15
Omission
10
05
2.3 Hz (E)
3.4 Hz (I)

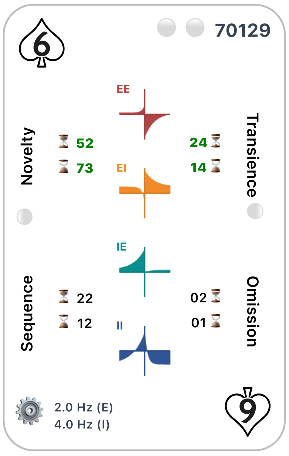
70129
EE
EI
IE
II
Novelty
52
73
Transience
24
14
Sequence
22
12
Omission
02
01
2.0 Hz (E)
4.0 Hz (I)

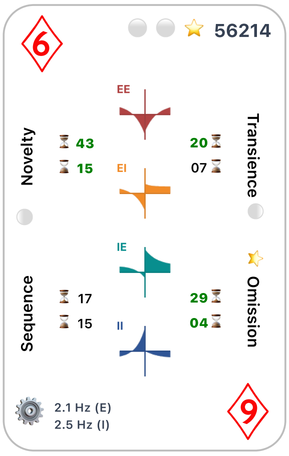
56214
EE
EI
IE
II
Novelty
43
15
Transience
20
07
Sequence
17
15
Omission
29
04
2.1 Hz (E)
2.5 Hz (I)

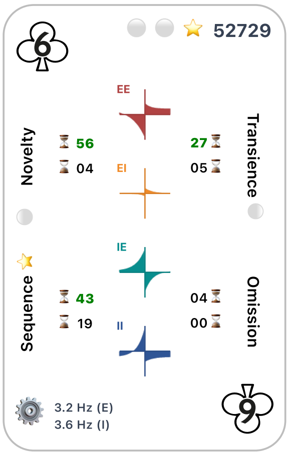
52729
EE
EI
IE
II
Novelty
56
04
Transience
27
05
Sequence
43
19
Omission
04
00
3.2 Hz (E)
3.6 Hz (I)

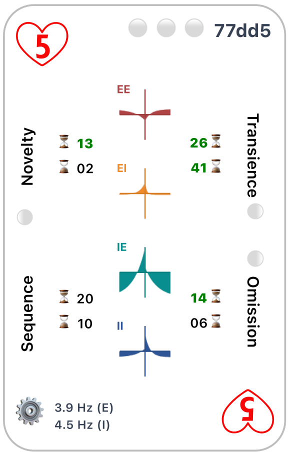
77dd5
EE
EI
IE
II
Novelty
13
02
Transience
26
41
Sequence
20
10
Omission
14
06
3.9 Hz (E)
4.5 Hz (I)

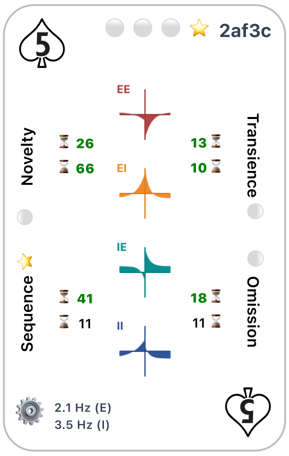
2af3c
EE
EI
IE
II
Novelty
26
66
Transience
13
10
Sequence
41
11
Omission
18
11
2.1 Hz (E)
3.5 Hz (I)

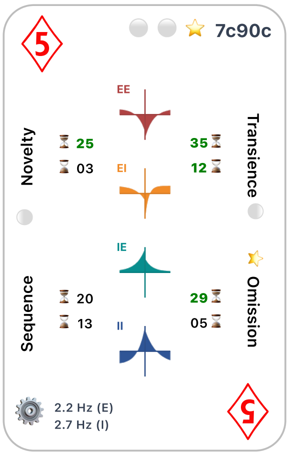
7c90c
EE
EI
IE
II
Novelty
25
03
Transience
35
12
Sequence
20
13
Omission
29
05
2.2 Hz (E)
2.7 Hz (I)

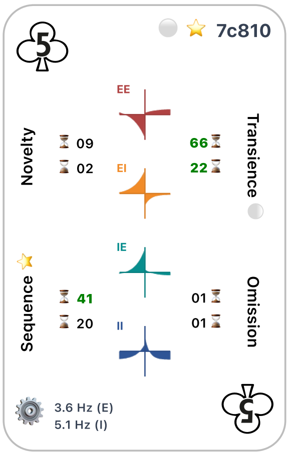
7c810
EE
EI
IE
II
Novelty
09
02
Transience
66
22
Sequence
41
20
Omission
01
01
3.6 Hz (E)
5.1 Hz (I)

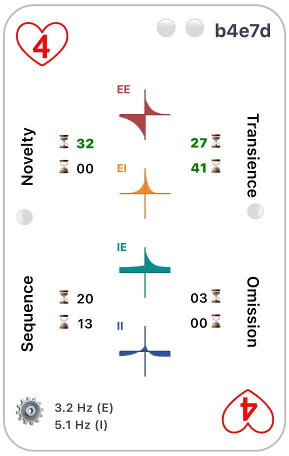
b4e7d
EE
EI
IE
II
Novelty
32
00
Transience
27
41
Sequence
20
13
Omission
03
00
3.2 Hz (E)
5.1 Hz (I)

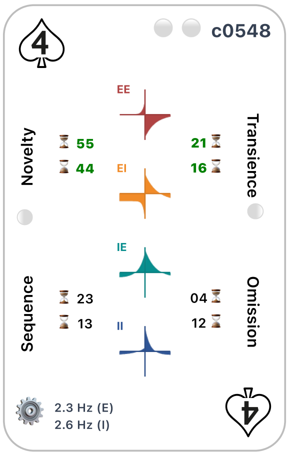
c0548
EE
EI
IE
II
Novelty
55
44
Transience
21
16
Sequence
23
13
Omission
04
12
2.3 Hz (E)
2.6 Hz (I)

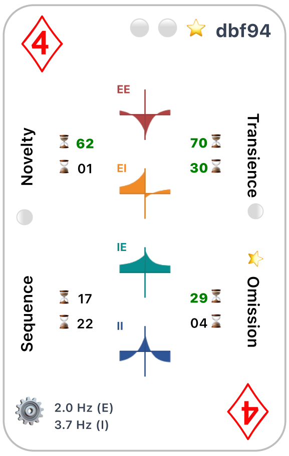
dbf94
EE
EI
IE
II
Novelty
62
01
Transience
70
30
Sequence
17
22
Omission
29
04
2.0 Hz (E)
3.7 Hz (I)

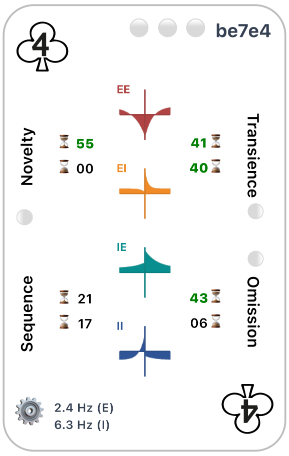
be7e4
EE
EI
IE
II
Novelty
55
00
Transience
41
40
Sequence
21
17
Omission
43
06
2.4 Hz (E)
6.3 Hz (I)

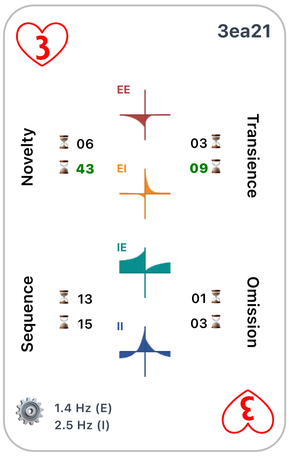
3
3ea21
Novelty
06
43
EE
EI
03
09
Transience
Sequence
13
15
IE
II
01
03
Omission
1.4 Hz (E)
2.5 Hz (I)
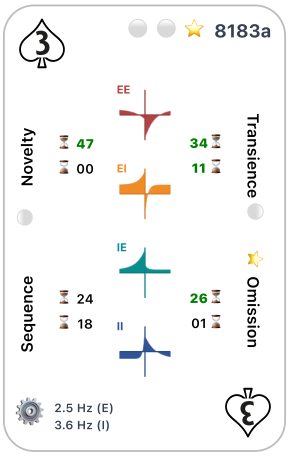
3
8183a
Novelty
47
00
EE
EI
34
11
Transience
Sequence
24
18
IE
II
26
01
Omission
2.5 Hz (E)
3.6 Hz (I)
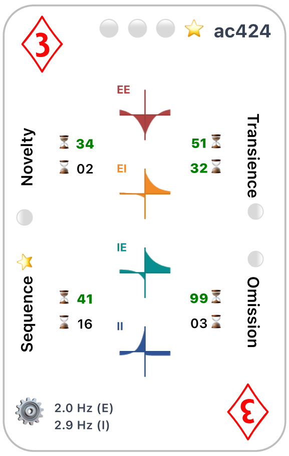
3
ac424
Novelty
34
02
EE
EI
51
32
Transience
Sequence
41
16
IE
II
99
03
Omission
2.0 Hz (E)
2.9 Hz (I)
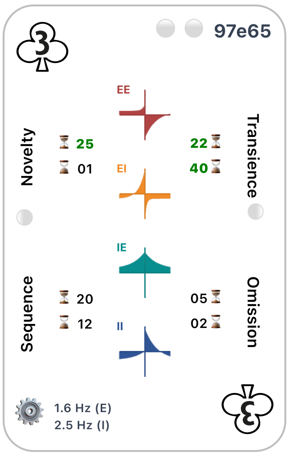
3
97e65
Novelty
25
01
EE
EI
22
40
Transience
Sequence
20
12
IE
II
05
02
Omission
1.6 Hz (E)
2.5 Hz (I)
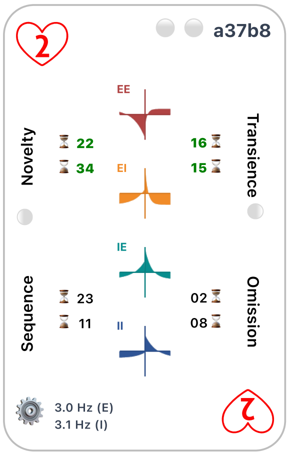
2
a37b8
Novelty
22
34
EE
EI
16
15
Transience
Sequence
23
11
IE
II
02
08
Omission
3.0 Hz (E)
3.1 Hz (I)
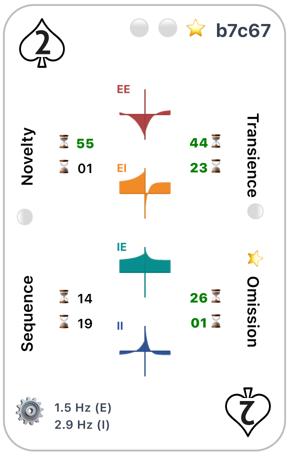
2
b7c67
Novelty
55
01
EE
EI
44
23
Transience
Sequence
14
19
IE
II
26
01
Omission
1.5 Hz (E)
2.9 Hz (I)
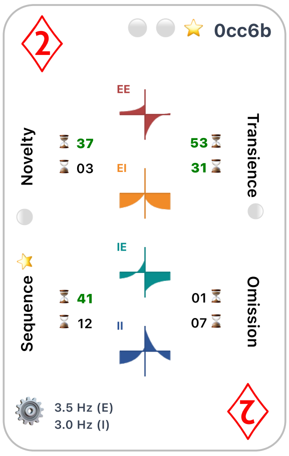
2
0cc6b
Novelty
37
03
EE
EI
53
31
Transience
Sequence
41
12
IE
II
01
07
Omission
3.5 Hz (E)
3.0 Hz (I)
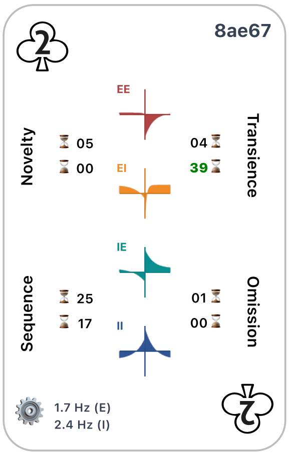
2
8ae67
Novelty
05
00
EE
EI
04
39
Transience
Sequence
25
17
IE
II
01
00
Omission
1.7 Hz (E)
2.4 Hz (I)
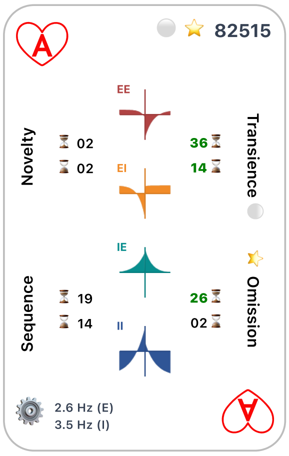
A
82515
Novelty
02
02
EE
EI
36
14
Transience
Sequence
19
14
IE
II
26
02
Omission
2.6 Hz (E)
3.5 Hz (I)
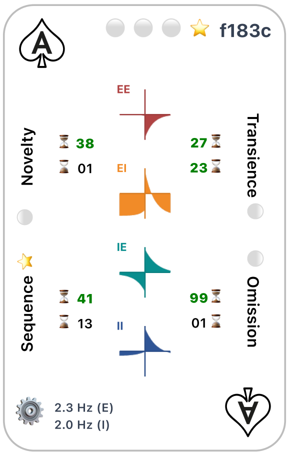
A
f183c
Novelty
38
01
EE
EI
27
23
Transience
Sequence
41
13
IE
II
99
01
Omission
2.3 Hz (E)
2.0 Hz (I)
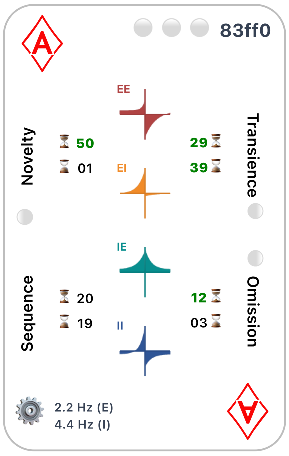
A
83ff0
Novelty
50
01
EE
EI
29
39
Transience
Sequence
20
19
IE
II
12
03
Omission
2.2 Hz (E)
4.4 Hz (I)
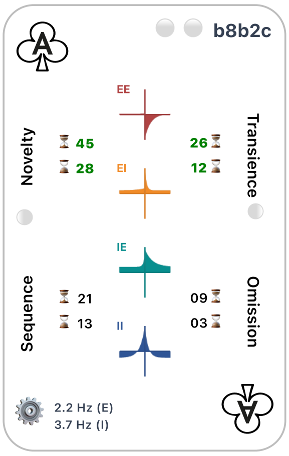
A
b8b2c
Novelty
45
28
EE
EI
26
12
Transience
Sequence
21
13
IE
II
09
03
Omission
2.2 Hz (E)
3.7 Hz (I)
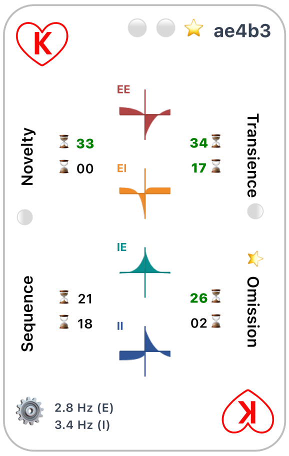
K
ae4b3
Novelty
33
00
EE
EI
34
17
Transience
Sequence
21
18
IE
II
26
02
Omission
2.8 Hz (E)
3.4 Hz (I)
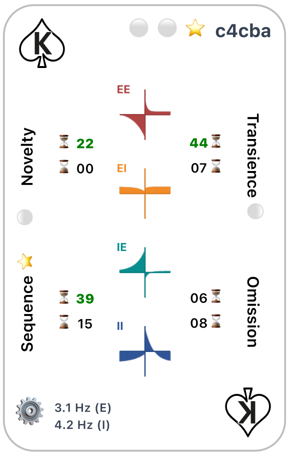
K
c4cba
Novelty
22
00
EE
EI
44
07
Transience
Sequence
39
15
IE
II
06
08
Omission
3.1 Hz (E)
4.2 Hz (I)
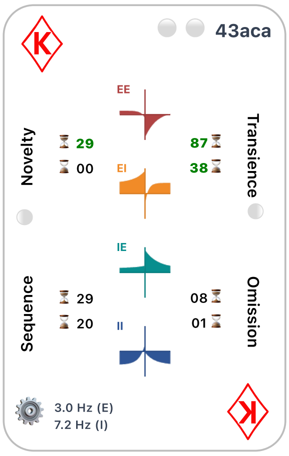
K
43aca
Novelty
29
00
EE
EI
87
38
Transience
Sequence
29
20
IE
II
08
01
Omission
3.0 Hz (E)
7.2 Hz (I)
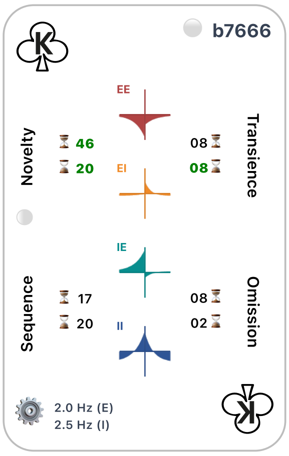
K
b7666
Novelty
46
20
EE
EI
08
08
Transience
Sequence
17
20
IE
II
08
02
Omission
2.0 Hz (E)
2.5 Hz (I)

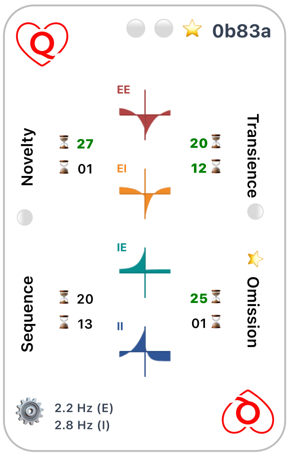
Q
0b83a
EE
EI
IE
II
Novelty 27 01
Transience 20 12
Sequence 20 13
Omission 25 01
2.2 Hz (E)
2.8 Hz (I)

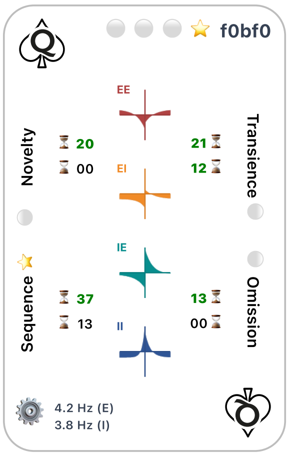
Q
f0bf0
EE
EI
IE
II
Novelty 20 00
Transience 21 12
Sequence 37 13
Omission 13 00
4.2 Hz (E)
3.8 Hz (I)

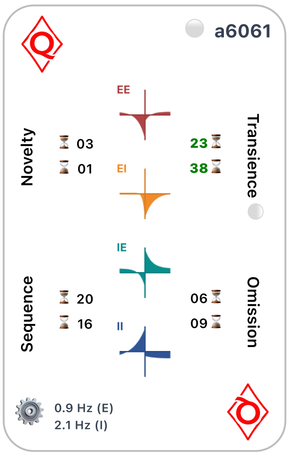
Q
a6061
EE
EI
IE
II
Novelty 03 01
Transience 23 38
Sequence 20 16
Omission 06 09
0.9 Hz (E)
2.1 Hz (I)

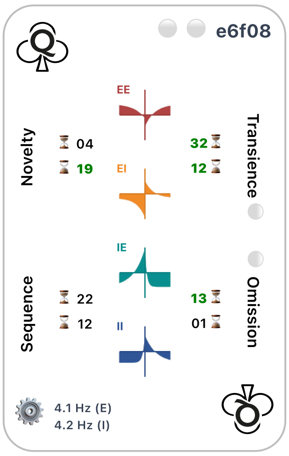
Q
e6f08
EE
EI
IE
II
Novelty 04 19
Transience 32 12
Sequence 22 12
Omission 13 01
4.1 Hz (E)
4.2 Hz (I)

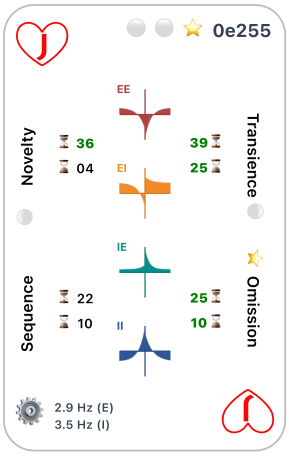
J
0e255
EE
EI
IE
II
Novelty 36 04
Transience 39 25
Sequence 22 10
Omission 25 10
2.9 Hz (E)
3.5 Hz (I)

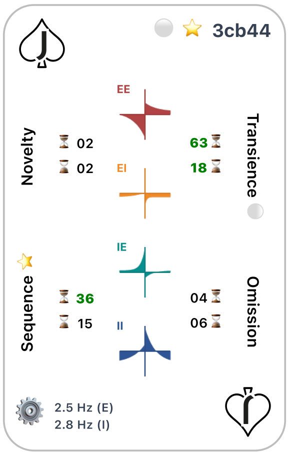
J
3cb44
EE
EI
IE
II
Novelty 02 02
Transience 63 18
Sequence 36 15
Omission 04 06
2.5 Hz (E)
2.8 Hz (I)

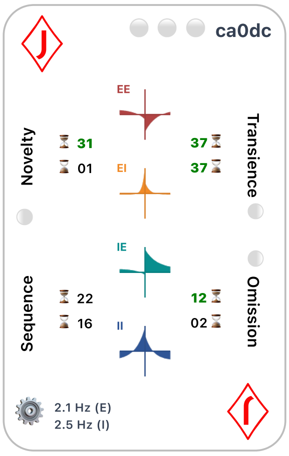
J
ca0dc
EE
EI
IE
II
Novelty 31 01
Transience 37 37
Sequence 22 16
Omission 12 02
2.1 Hz (E)
2.5 Hz (I)

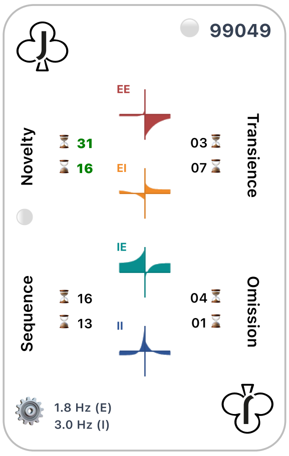
J
99049
EE
EI
IE
II
Novelty 31 16
Transience 03 07
Sequence 16 13
Omission 04 01
1.8 Hz (E)
3.0 Hz (I)

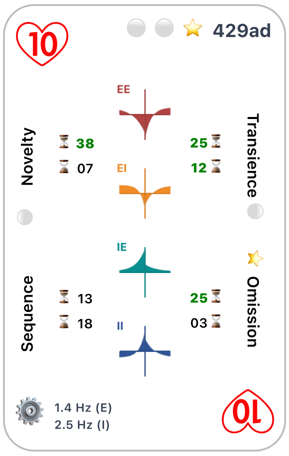
10
429ad
EE
EI
IE
II
Novelty 38 07
Transience 25 12
Sequence 13 18
Omission 25 03
1.4 Hz (E)
2.5 Hz (I)

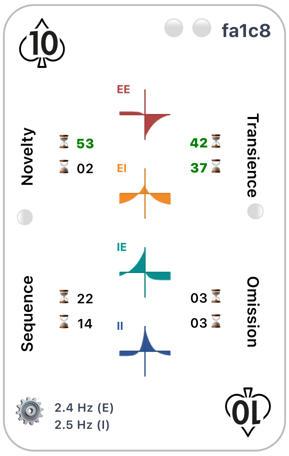
10
fa1c8
EE
EI
IE
II
Novelty 53 02
Transience 42 37
Sequence 22 14
Omission 03 03
2.4 Hz (E)
2.5 Hz (I)

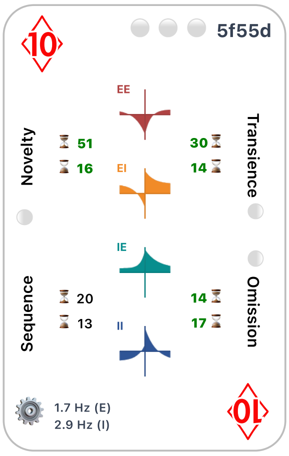
10
5f55d
EE
EI
IE
II
Novelty 51 16
Transience 30 14
Sequence 20 13
Omission 14 17
1.7 Hz (E)
2.9 Hz (I)

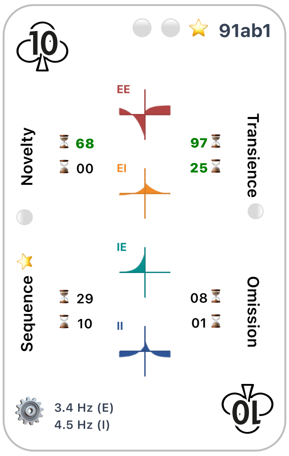
10
91ab1
EE
EI
IE
II
Novelty 68 00
Transience 97 25
Sequence 29 10
Omission 08 01
3.4 Hz (E)
4.5 Hz (I)

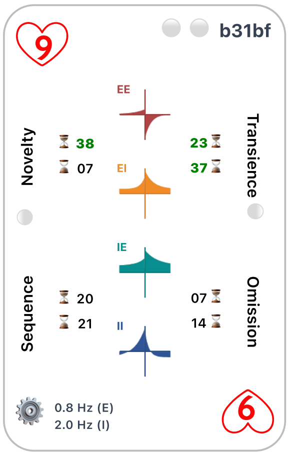
9
b31bf
EE
EI
IE
II
Novelty 38 07
Transience 23 37
Sequence 20 21
Omission 07 14
0.8 Hz (E)
2.0 Hz (I)

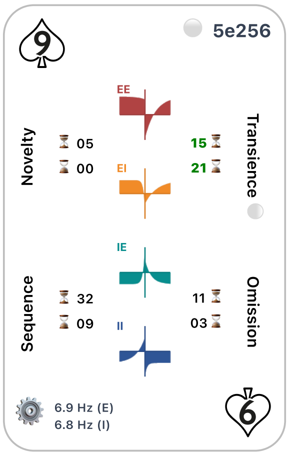
9
5e256
EE
EI
IE
II
Novelty 05 00
Transience 15 21
Sequence 32 09
Omission 11 03
6.9 Hz (E)
6.8 Hz (I)

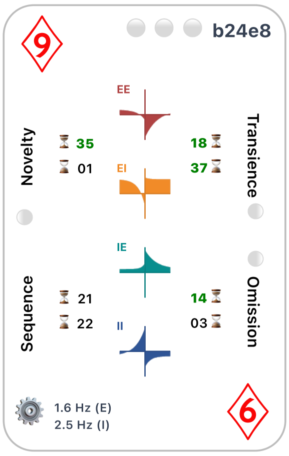
9
b24e8
EE
EI
IE
II
Novelty 35 01
Transience 18 37
Sequence 21 22
Omission 14 03
1.6 Hz (E)
2.5 Hz (I)

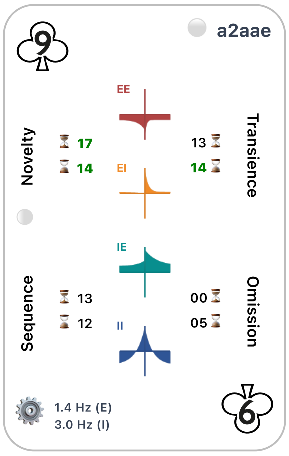
9
a2aae
EE
EI
IE
II
Novelty 17 14
Transience 13 14
Sequence 13 12
Omission 00 05
1.4 Hz (E)
3.0 Hz (I)

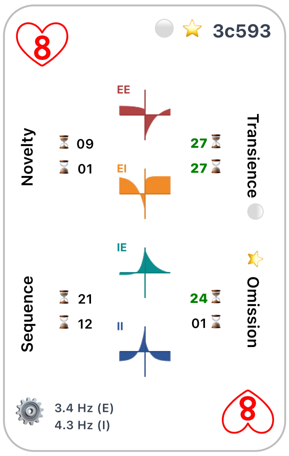
8
3c593
EE
Novelty
09
01
27
27
Transience
EI
IE
Sequence
21
12
24
01
Omission
II
3.4 Hz (E)
4.3 Hz (I)
8
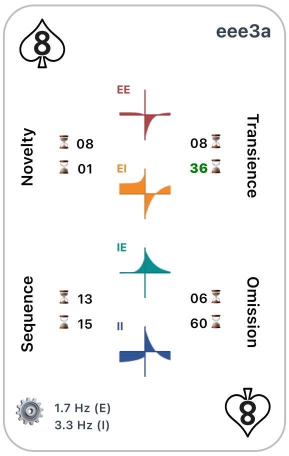
8
eee3a
EE
Novelty
08
01
08
36
Transience
EI
IE
Sequence
13
15
06
60
Omission
II
1.7 Hz (E)
3.3 Hz (I)
8
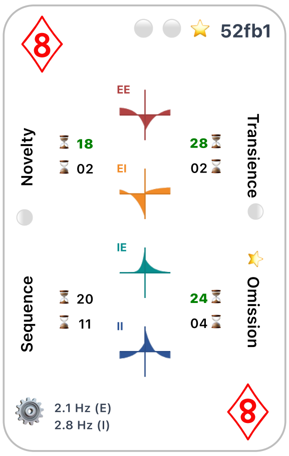
8
52fb1
EE
Novelty
18
02
28
02
Transience
EI
IE
Sequence
20
11
24
04
Omission
II
2.1 Hz (E)
2.8 Hz (I)
8
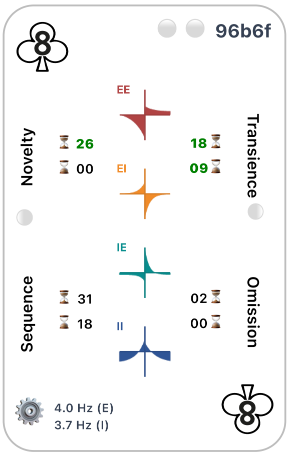
8
96b6f
EE
Novelty
26
00
18
09
Transience
EI
IE
Sequence
31
18
02
00
Omission
II
4.0 Hz (E)
3.7 Hz (I)
8
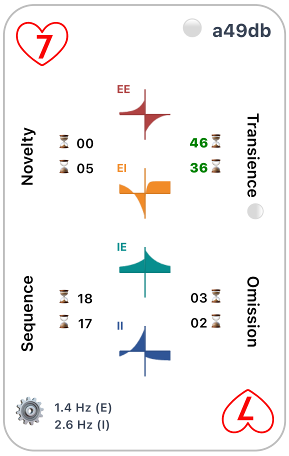
7
a49db
EE
Novelty
00
05
46
36
Transience
EI
IE
Sequence
18
17
03
02
Omission
II
1.4 Hz (E)
2.6 Hz (I)
7
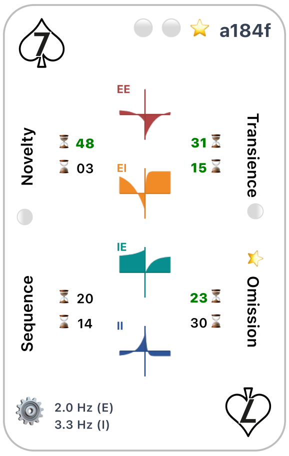
7
a184f
EE
Novelty
48
03
31
15
Transience
EI
IE
Sequence
20
14
23
30
Omission
II
2.0 Hz (E)
3.3 Hz (I)
7
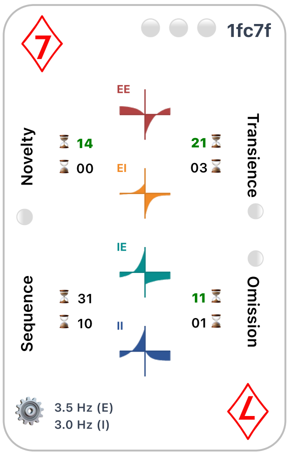
7
1fc7f
EE
Novelty
14
00
21
03
Transience
EI
IE
Sequence
31
10
11
01
Omission
II
3.5 Hz (E)
3.0 Hz (I)
7
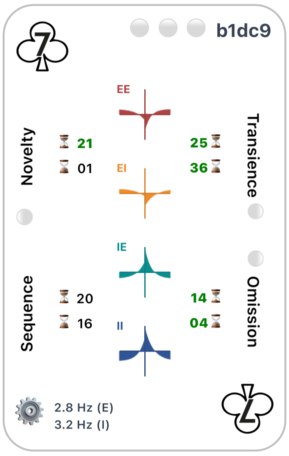
7
b1dc9
EE
Novelty
21
01
25
36
Transience
EI
IE
Sequence
20
16
14
04
Omission
II
2.8 Hz (E)
3.2 Hz (I)
7
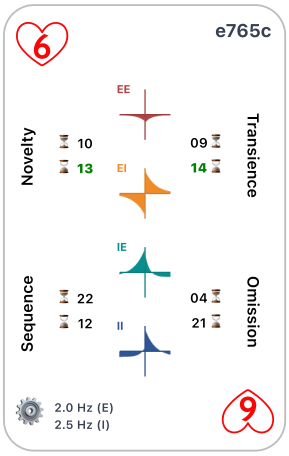
6
e765c
EE
Novelty
10
13
09
14
Transience
EI
IE
Sequence
22
12
04
21
Omission
II
2.0 Hz (E)
2.5 Hz (I)
6
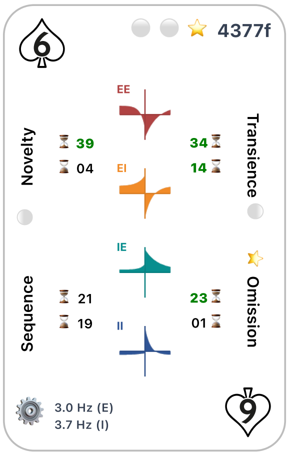
6
4377f
EE
Novelty
39
04
34
14
Transience
EI
IE
Sequence
21
19
23
01
Omission
II
3.0 Hz (E)
3.7 Hz (I)
6
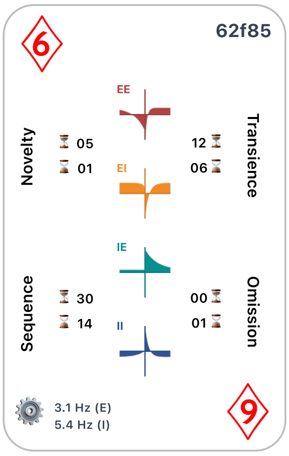
6
62f85
EE
Novelty
05
01
12
06
Transience
EI
IE
Sequence
30
14
00
01
Omission
II
3.1 Hz (E)
5.4 Hz (I)
6
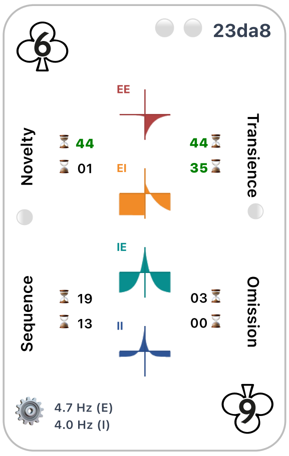
6
23da8
EE
Novelty
44
01
44
35
Transience
EI
IE
Sequence
19
13
03
00
Omission
II
4.7 Hz (E)
4.0 Hz (I)
6
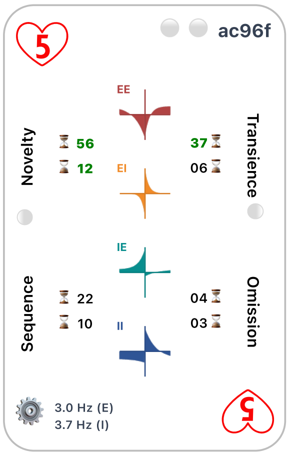
5
ac96f
EE
Novelty
56
12
37
06
Transience
EI
IE
Sequence
22
10
04
03
Omission
II
3.0 Hz (E)
3.7 Hz (I)
5
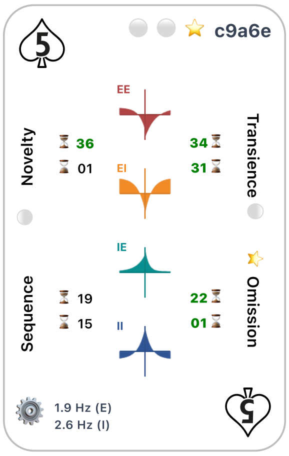
5
c9a6e
EE
Novelty
36
01
34
31
Transience
EI
IE
Sequence
19
15
22
01
Omission
II
1.9 Hz (E)
2.6 Hz (I)
5
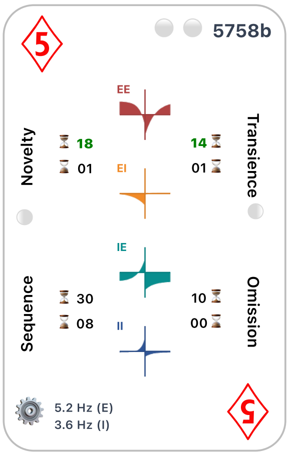
5
5758b
EE
Novelty
18
01
14
01
Transience
EI
IE
Sequence
30
08
10
00
Omission
II
5.2 Hz (E)
3.6 Hz (I)
5
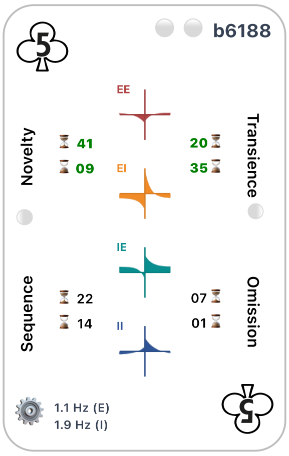
5
b6188
EE
Novelty
41
09
20
35
Transience
EI
IE
Sequence
22
14
07
01
Omission
II
1.1 Hz (E)
1.9 Hz (I)
5

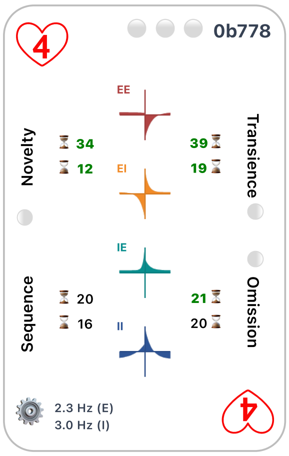
4
0b778
EE
EI
IE
II
Novelty
34
12
Transience
39
19
Sequence
20
16
Omission
21
20
2.3 Hz (E)
3.0 Hz (I)
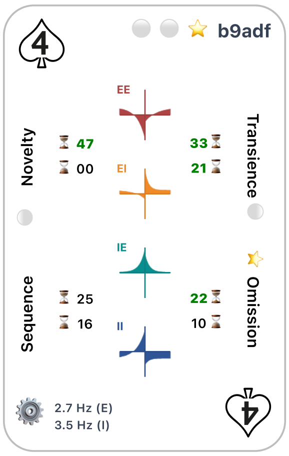
4
b9adf
EE
EI
IE
II
Novelty
47
00
Transience
33
21
Sequence
25
16
Omission
22
10
2.7 Hz (E)
3.5 Hz (I)
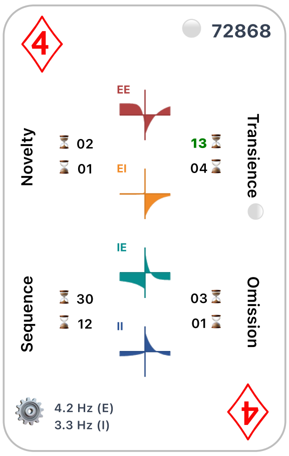
4
72868
EE
EI
IE
II
Novelty
02
01
Transience
13
04
Sequence
30
12
Omission
03
01
4.2 Hz (E)
3.3 Hz (I)
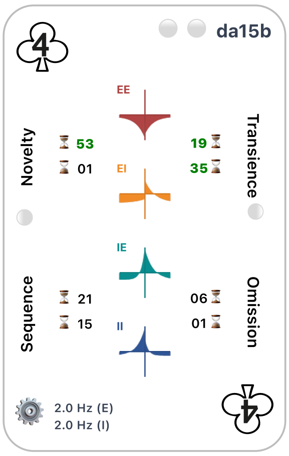
4
da15b
EE
EI
IE
II
Novelty
53
01
Transience
19
35
Sequence
21
15
Omission
06
01
2.0 Hz (E)
2.0 Hz (I)
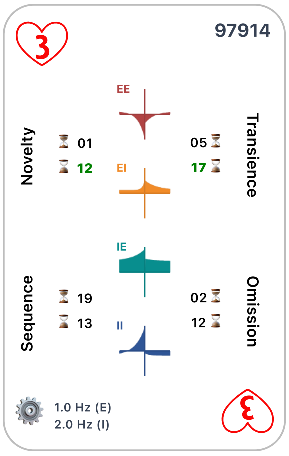
3
97914
EE
EI
IE
II
Novelty
01
12
Transience
05
17
Sequence
19
13
Omission
02
12
1.0 Hz (E)
2.0 Hz (I)
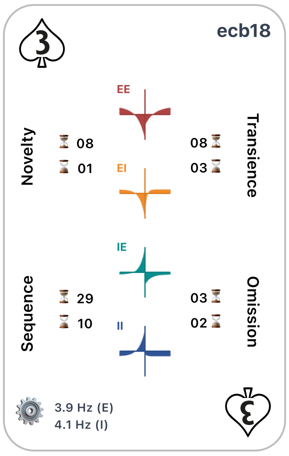
3
ecb18
EE
EI
IE
II
Novelty
08
01
Transience
08
03
Sequence
29
10
Omission
03
02
3.9 Hz (E)
4.1 Hz (I)
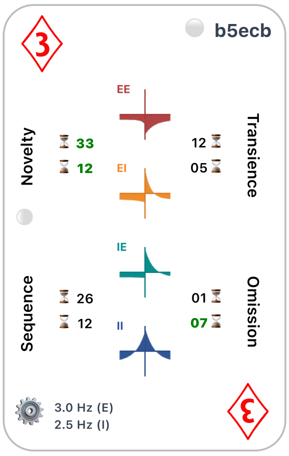
3
b5ecb
EE
EI
IE
II
Novelty
33
12
Transience
12
05
Sequence
26
12
Omission
01
07
3.0 Hz (E)
2.5 Hz (I)
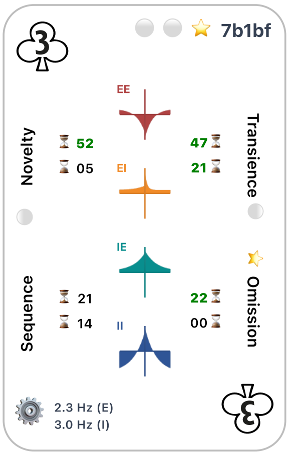
3
7b1bf
EE
EI
IE
II
Novelty
52
05
Transience
47
21
Sequence
21
14
Omission
22
00
2.3 Hz (E)
3.0 Hz (I)
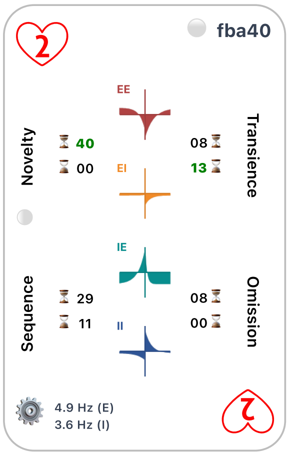
2
fba40
EE
EI
IE
II
Novelty
40
00
Transience
08
13
Sequence
29
11
Omission
08
00
4.9 Hz (E)
3.6 Hz (I)
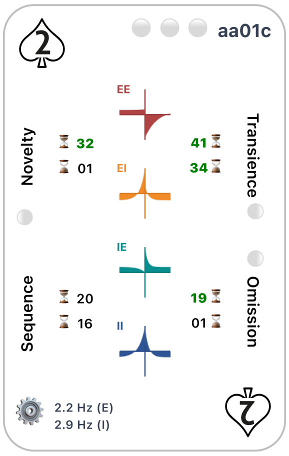
2
aa01c
EE
EI
IE
II
Novelty
32
01
Transience
41
34
Sequence
20
16
Omission
19
01
2.2 Hz (E)
2.9 Hz (I)
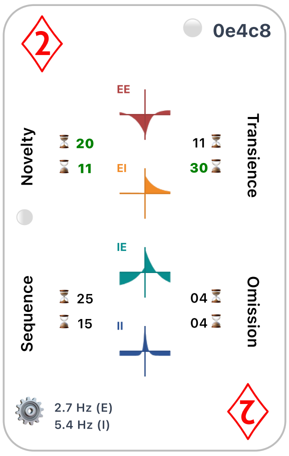
2
0e4c8
EE
EI
IE
II
Novelty
20
11
Transience
11
30
Sequence
25
15
Omission
04
04
2.7 Hz (E)
5.4 Hz (I)
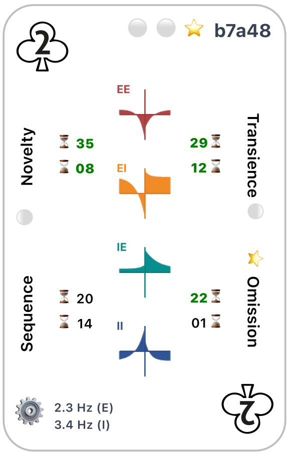
2
b7a48
EE
EI
IE
II
Novelty
35
08
Transience
29
12
Sequence
20
14
Omission
22
01
2.3 Hz (E)
3.4 Hz (I)
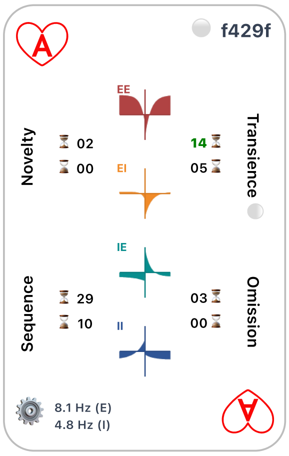
A
f429f
EE
EI
IE
II
Novelty
02
00
Transience
14
05
Sequence
29
10
Omission
03
00
8.1 Hz (E)
4.8 Hz (I)
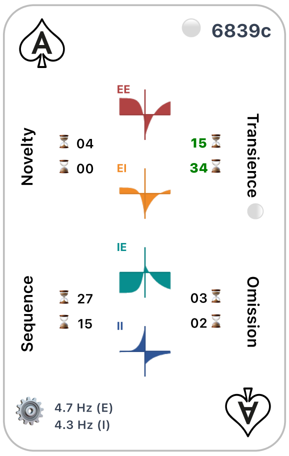
A
6839c
EE
EI
IE
II
Novelty
04
00
Transience
15
34
Sequence
27
15
Omission
03
02
4.7 Hz (E)
4.3 Hz (I)
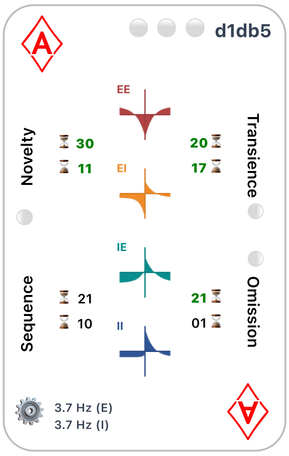
A
d1db5
EE
EI
IE
II
Novelty
30
11
Transience
20
17
Sequence
21
10
Omission
21
01
3.7 Hz (E)
3.7 Hz (I)
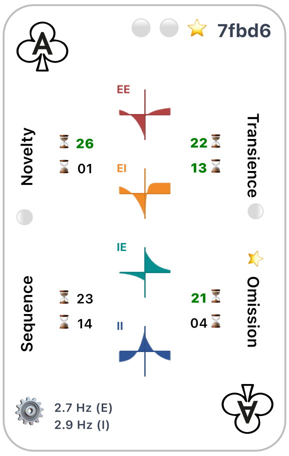
A
7fbd6
EE
EI
IE
II
Novelty
26
01
Transience
22
13
Sequence
23
14
Omission
21
04
2.7 Hz (E)
2.9 Hz (I)

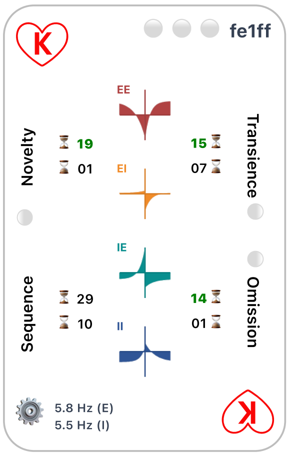
fe1ff
Novelty
19
01
EE
EI
15
07
Transience
IE
II
Sequence
29
10
14
01
Omission
5.8 Hz (E)
5.5 Hz (I)

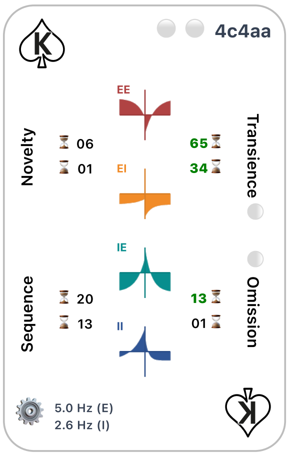
4c4aa
Novelty
06
01
EE
EI
65
34
Transience
IE
II
Sequence
20
13
13
01
Omission
5.0 Hz (E)
2.6 Hz (I)

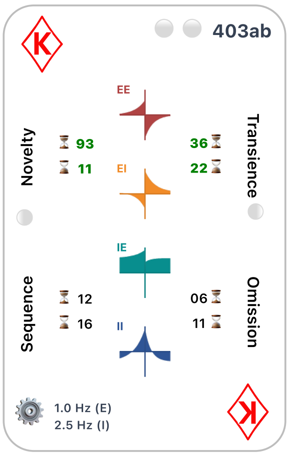
403ab
Novelty
93
11
EE
EI
36
22
Transience
IE
II
Sequence
12
16
06
11
Omission
1.0 Hz (E)
2.5 Hz (I)

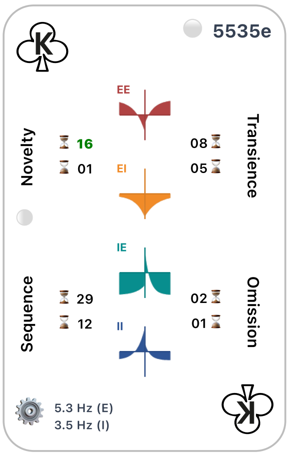
5535e
Novelty
16
01
EE
EI
08
05
Transience
IE
II
Sequence
29
12
02
01
Omission
5.3 Hz (E)
3.5 Hz (I)

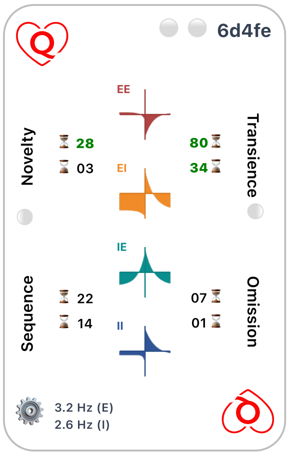
6d4fe
Novelty
28
03
EE
EI
80
34
Transience
IE
II
Sequence
22
14
07
01
Omission
3.2 Hz (E)
2.6 Hz (I)

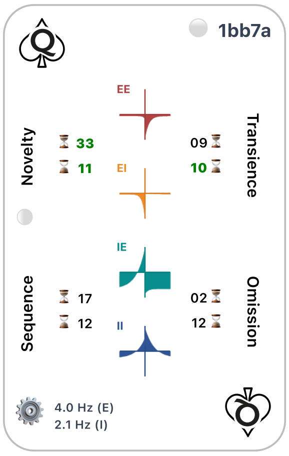
1bb7a
Novelty
33
11
EE
EI
09
10
Transience
IE
II
Sequence
17
12
02
12
Omission
4.0 Hz (E)
2.1 Hz (I)

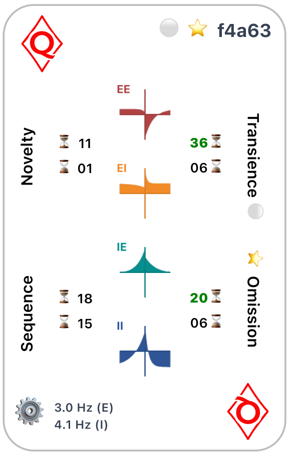
f4a63
Novelty
11
01
EE
EI
36
06
Transience
IE
II
Sequence
18
15
20
06
Omission
3.0 Hz (E)
4.1 Hz (I)

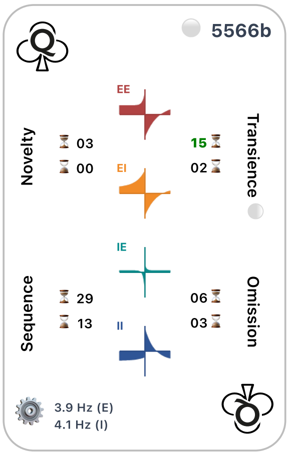
5566b
Novelty
03
00
EE
EI
15
02
Transience
IE
II
Sequence
29
13
06
03
Omission
3.9 Hz (E)
4.1 Hz (I)

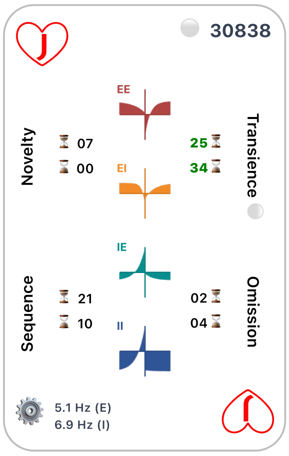
30838
Novelty
07
00
EE
EI
25
34
Transience
IE
II
Sequence
21
10
02
04
Omission
5.1 Hz (E)
6.9 Hz (I)

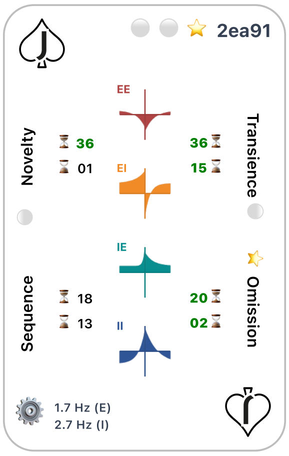
2ea91
Novelty
36
01
EE
EI
36
15
Transience
IE
II
Sequence
18
13
20
02
Omission
1.7 Hz (E)
2.7 Hz (I)

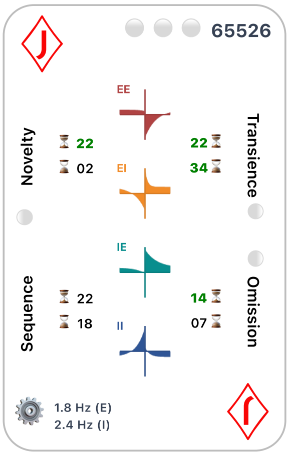
65526
Novelty
22
02
EE
EI
22
34
Transience
IE
II
Sequence
22
18
14
07
Omission
1.8 Hz (E)
2.4 Hz (I)

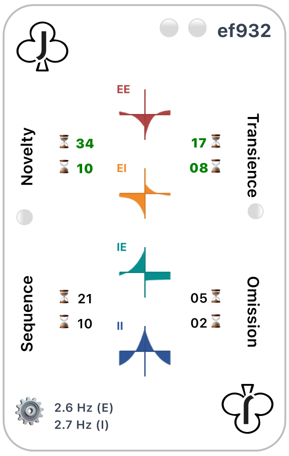
ef932
Novelty
34
10
EE
EI
17
08
Transience
IE
II
Sequence
21
10
05
02
Omission
2.6 Hz (E)
2.7 Hz (I)

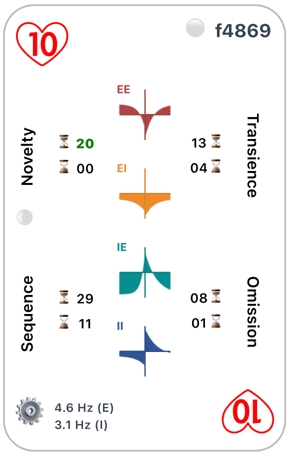
f4869
Novelty
20
00
EE
EI
13
04
Transience
IE
II
Sequence
29
11
08
01
Omission
4.6 Hz (E)
3.1 Hz (I)

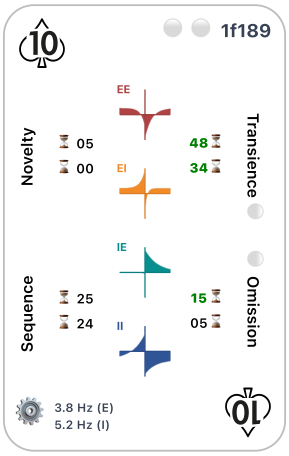
1f189
Novelty
05
00
EE
EI
48
34
Transience
IE
II
Sequence
25
24
15
05
Omission
3.8 Hz (E)
5.2 Hz (I)

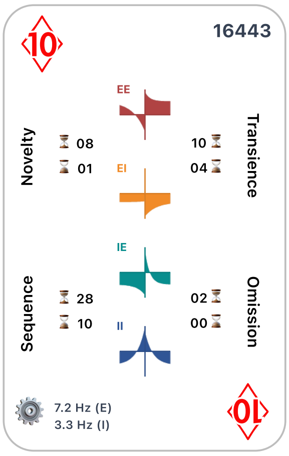
16443
Novelty
08
01
EE
EI
10
04
Transience
IE
II
Sequence
28
10
02
00
Omission
7.2 Hz (E)
3.3 Hz (I)

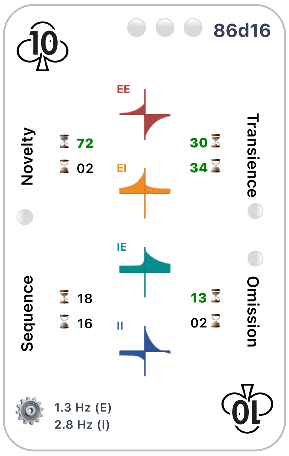
86d16
Novelty
72
02
EE
EI
30
34
Transience
IE
II
Sequence
18
16
13
02
Omission
1.3 Hz (E)
2.8 Hz (I)

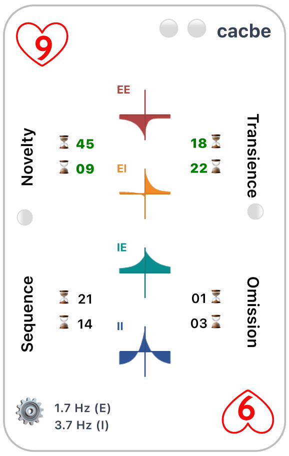
9
cacbe
EE
EI
IE
II
Novelty
45
09
Transience
18
22
Sequence
21
14
Omission
01
03
1.7 Hz (E)
3.7 Hz (I)
6

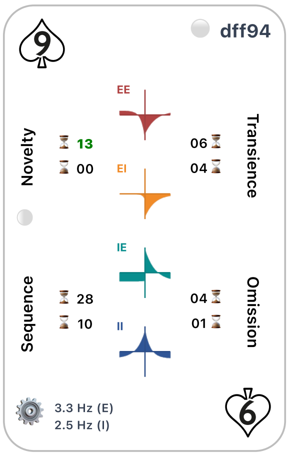
9
dff94
EE
EI
IE
II
Novelty
13
00
Transience
06
04
Sequence
28
10
Omission
04
01
3.3 Hz (E)
2.5 Hz (I)
6

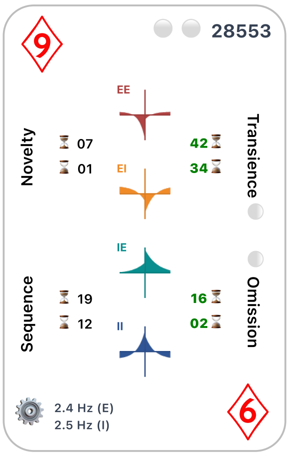
9
28553
EE
EI
IE
II
Novelty
07
01
Transience
42
34
Sequence
19
12
Omission
16
02
2.4 Hz (E)
2.5 Hz (I)
6

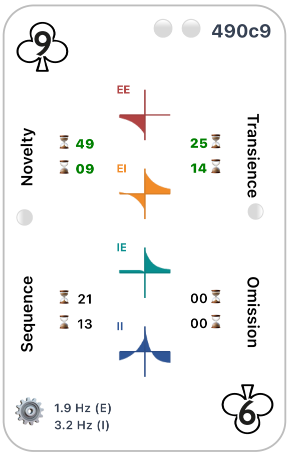
9
490c9
EE
EI
IE
II
Novelty
49
09
Transience
25
14
Sequence
21
13
Omission
00
00
1.9 Hz (E)
3.2 Hz (I)
6

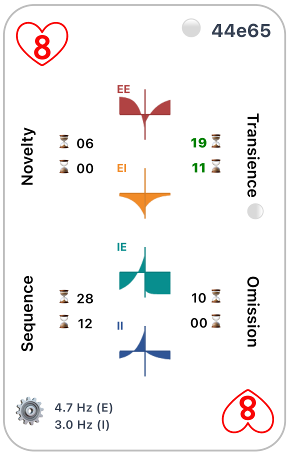
8
44e65
EE
EI
IE
II
Novelty
06
00
Transience
19
11
Sequence
28
12
Omission
10
00
4.7 Hz (E)
3.0 Hz (I)
8

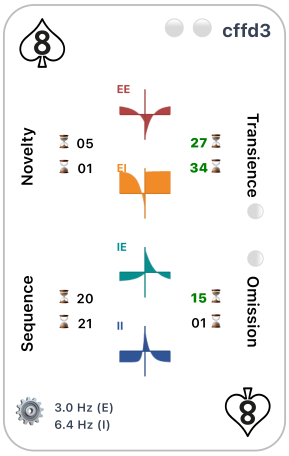
8
cffd3
EE
EI
IE
II
Novelty
05
01
Transience
27
34
Sequence
20
21
Omission
15
01
3.0 Hz (E)
6.4 Hz (I)
8

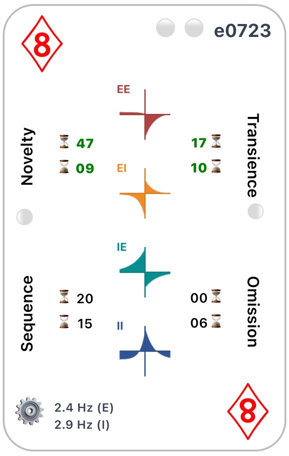
8
e0723
EE
EI
IE
II
Novelty
47
09
Transience
17
10
Sequence
20
15
Omission
00
06
2.4 Hz (E)
2.9 Hz (I)
8

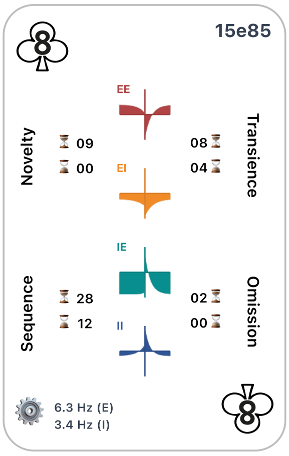
8
15e85
EE
EI
IE
II
Novelty
09
00
Transience
08
04
Sequence
28
12
Omission
02
00
6.3 Hz (E)
3.4 Hz (I)
8

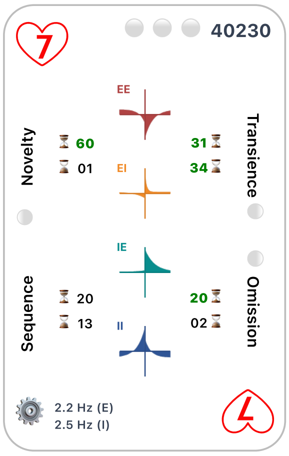
7
40230
EE
EI
IE
II
Novelty
60
01
Transience
31
34
Sequence
20
13
Omission
20
02
2.2 Hz (E)
2.5 Hz (I)
7

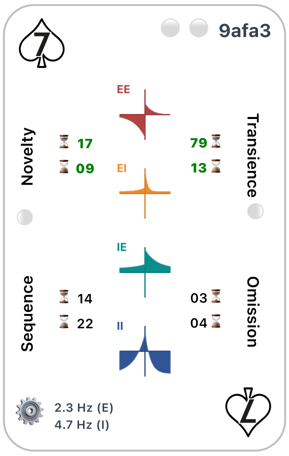
7
9afa3
EE
EI
IE
II
Novelty
17
09
Transience
79
13
Sequence
14
22
Omission
03
04
2.3 Hz (E)
4.7 Hz (I)
7

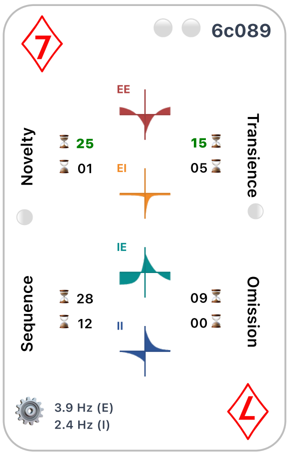
7
6c089
EE
EI
IE
II
Novelty
25
01
Transience
15
05
Sequence
28
12
Omission
09
00
3.9 Hz (E)
2.4 Hz (I)
7

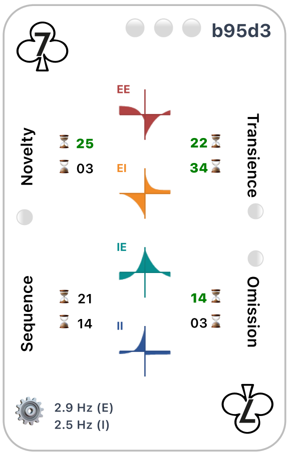
7
b95d3
EE
EI
IE
II
Novelty
25
03
Transience
22
34
Sequence
21
14
Omission
14
03
2.9 Hz (E)
2.5 Hz (I)
7

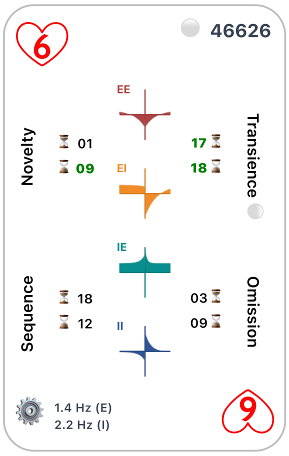
6
46626
EE
EI
IE
II
Novelty
01
09
Transience
17
18
Sequence
18
12
Omission
03
09
1.4 Hz (E)
2.2 Hz (I)
9

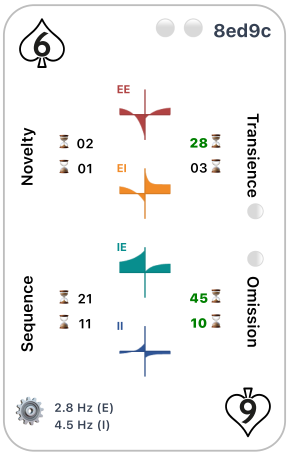
6
8ed9c
EE
EI
IE
II
Novelty
02
01
Transience
28
03
Sequence
21
11
Omission
45
10
2.8 Hz (E)
4.5 Hz (I)
9

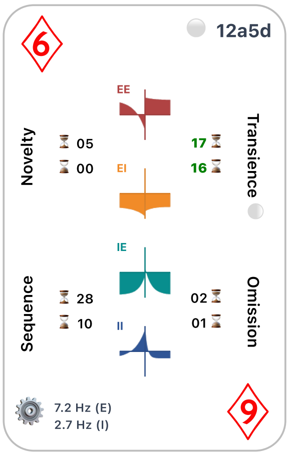
6
12a5d
EE
EI
IE
II
Novelty
05
00
Transience
17
16
Sequence
28
10
Omission
02
01
7.2 Hz (E)
2.7 Hz (I)
9

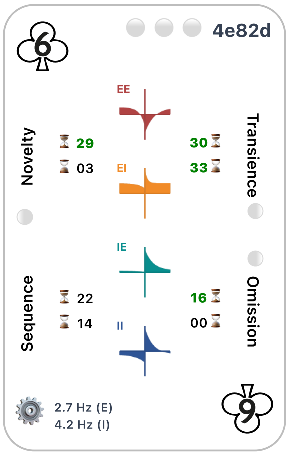
6
4e82d
EE
EI
IE
II
Novelty
29
03
Transience
30
33
Sequence
22
14
Omission
16
00
2.7 Hz (E)
4.2 Hz (I)
9

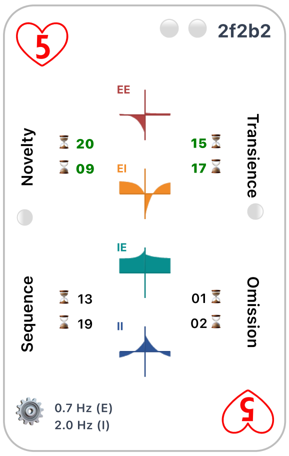
5
2f2b2
Novelty
20
09
EE
EI
15
17
Transience
Sequence
13
19
IE
II
01
02
Omission
0.7 Hz (E)
2.0 Hz (I)

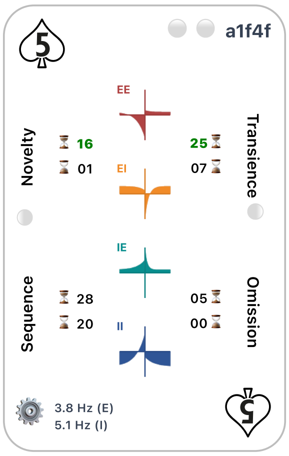
5
a1f4f
Novelty
16
01
EE
EI
25
07
Transience
Sequence
28
20
IE
II
05
00
Omission
3.8 Hz (E)
5.1 Hz (I)

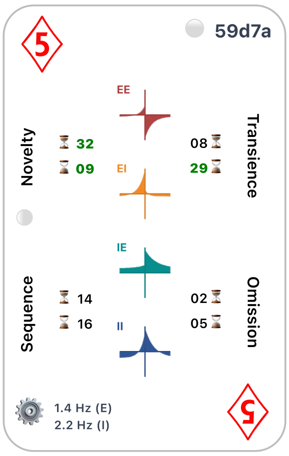
5
59d7a
Novelty
32
09
EE
EI
08
29
Transience
Sequence
14
16
IE
II
02
05
Omission
1.4 Hz (E)
2.2 Hz (I)

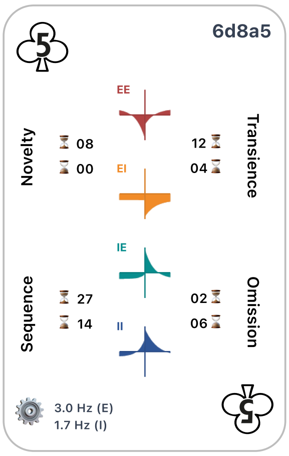
5
6d8a5
Novelty
08
00
EE
EI
12
04
Transience
Sequence
27
14
IE
II
02
06
Omission
3.0 Hz (E)
1.7 Hz (I)

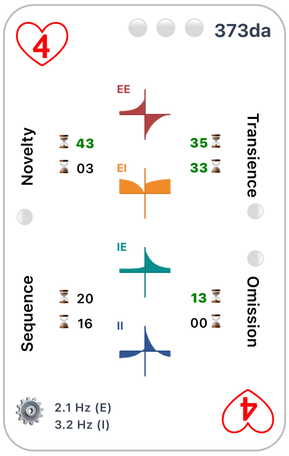
4
373da
Novelty
43
03
EE
EI
35
33
Transience
Sequence
20
16
IE
II
13
00
Omission
2.1 Hz (E)
3.2 Hz (I)

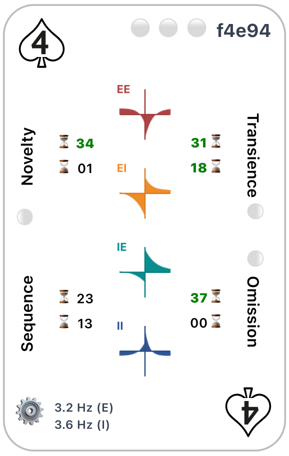
4
f4e94
Novelty
34
01
EE
EI
31
18
Transience
Sequence
23
13
IE
II
37
00
Omission
3.2 Hz (E)
3.6 Hz (I)

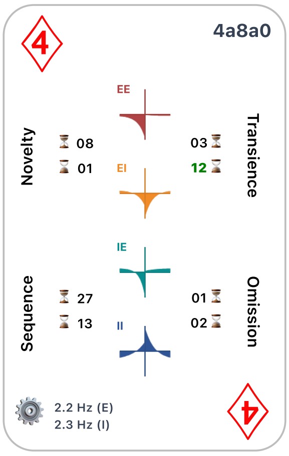
4
4a8a0
Novelty
08
01
EE
EI
03
12
Transience
Sequence
27
13
IE
II
01
02
Omission
2.2 Hz (E)
2.3 Hz (I)

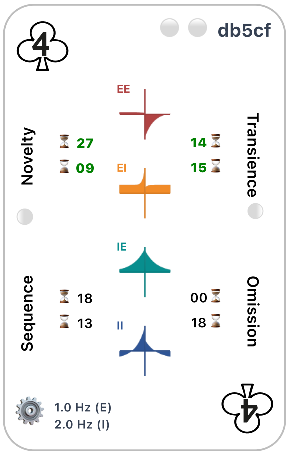
4
db5cf
Novelty
27
09
EE
EI
14
15
Transience
Sequence
18
13
IE
II
00
18
Omission
1.0 Hz (E)
2.0 Hz (I)

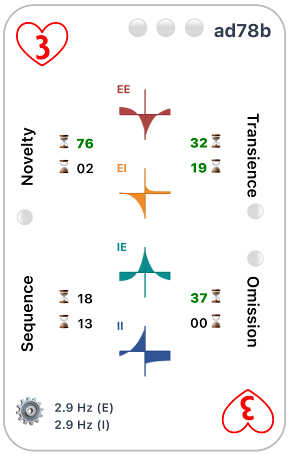
3
ad78b
Novelty
76
02
EE
EI
32
19
Transience
Sequence
18
13
IE
II
37
00
Omission
2.9 Hz (E)
2.9 Hz (I)

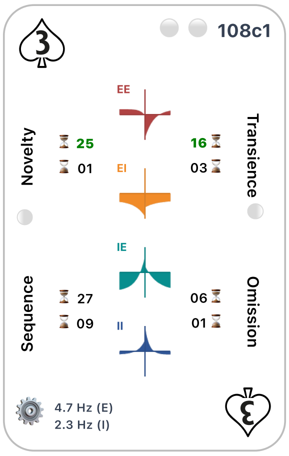
3
108c1
Novelty
25
01
EE
EI
16
03
Transience
Sequence
27
09
IE
II
06
01
Omission
4.7 Hz (E)
2.3 Hz (I)

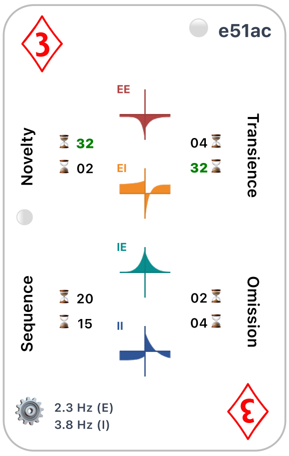
3
e51ac
Novelty
32
02
EE
EI
04
32
Transience
Sequence
20
15
IE
II
02
04
Omission
2.3 Hz (E)
3.8 Hz (I)

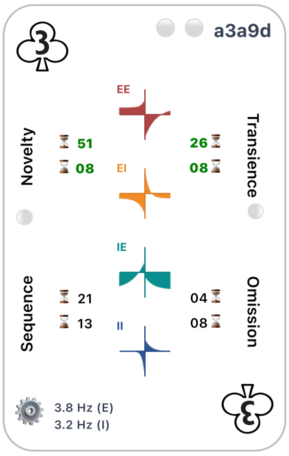
3
a3a9d
Novelty
51
08
EE
EI
26
08
Transience
Sequence
21
13
IE
II
04
08
Omission
3.8 Hz (E)
3.2 Hz (I)

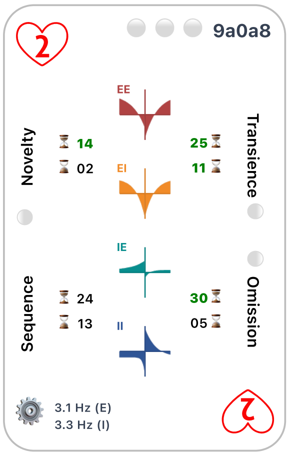
2
9a0a8
Novelty
14
02
EE
EI
25
11
Transience
Sequence
24
13
IE
II
30
05
Omission
3.1 Hz (E)
3.3 Hz (I)

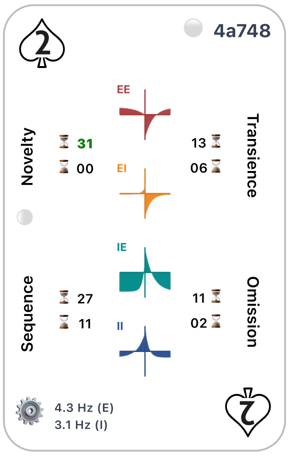
2
4a748
Novelty
31
00
EE
EI
13
06
Transience
Sequence
27
11
IE
II
11
02
Omission
4.3 Hz (E)
3.1 Hz (I)

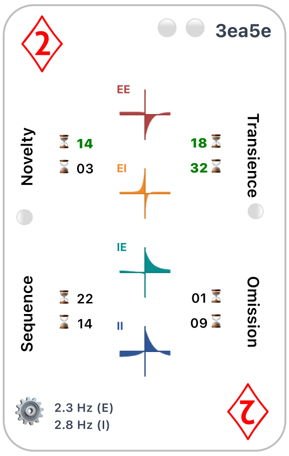
2
3ea5e
Novelty
14
03
EE
EI
18
32
Transience
Sequence
22
14
IE
II
01
09
Omission
2.3 Hz (E)
2.8 Hz (I)

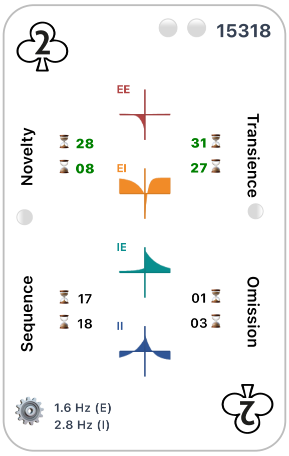
2
15318
Novelty
28
08
EE
EI
31
27
Transience
Sequence
17
18
IE
II
01
03
Omission
1.6 Hz (E)
2.8 Hz (I)

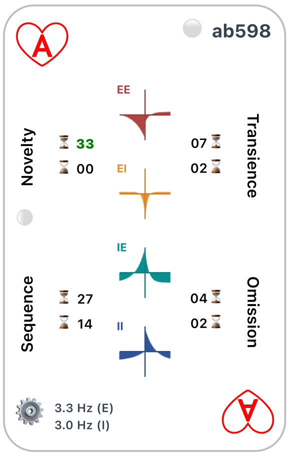
A
ab598
Novelty 33 00
Transience 07 02
Sequence 27 14
Omission 04 02
EE EI IE II
3.3 Hz (E)
3.0 Hz (I)

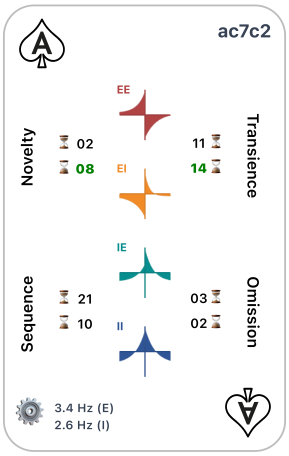
A
ac7c2
Novelty 02 08
Transience 11 14
Sequence 21 10
Omission 03 02
EE EI IE II
3.4 Hz (E)
2.6 Hz (I)

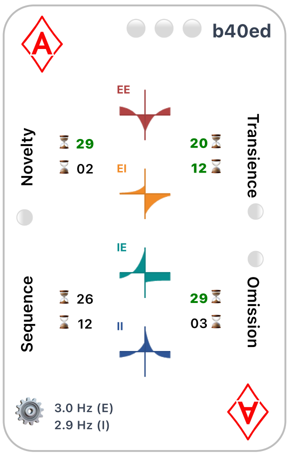
A
b40ed
Novelty 29 02
Transience 20 12
Sequence 26 12
Omission 29 03
EE EI IE II
3.0 Hz (E)
2.9 Hz (I)

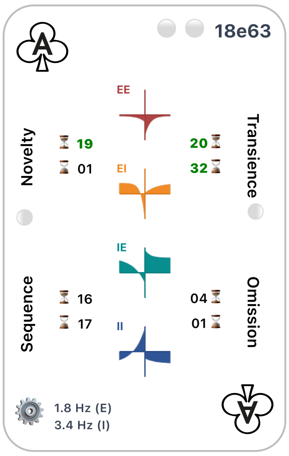
A
18e63
Novelty 19 01
Transience 20 32
Sequence 16 17
Omission 04 01
EE EI IE II
1.8 Hz (E)
3.4 Hz (I)

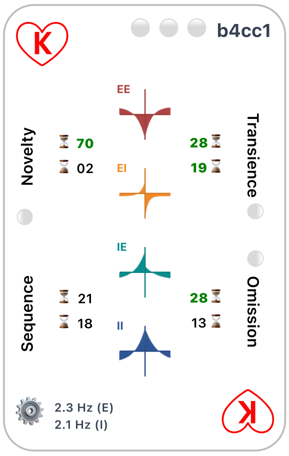
K
b4cc1
Novelty 70 02
Transience 28 19
Sequence 21 18
Omission 28 13
EE EI IE II
2.3 Hz (E)
2.1 Hz (I)

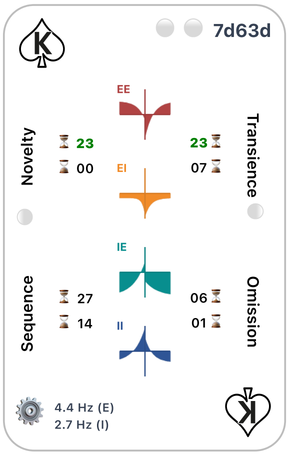
K
7d63d
Novelty 23 00
Transience 23 07
Sequence 27 14
Omission 06 01
EE EI IE II
4.4 Hz (E)
2.7 Hz (I)

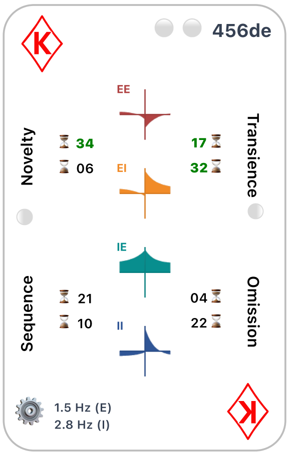
K
456de
Novelty 34 06
Transience 17 32
Sequence 21 10
Omission 04 22
EE EI IE II
1.5 Hz (E)
2.8 Hz (I)

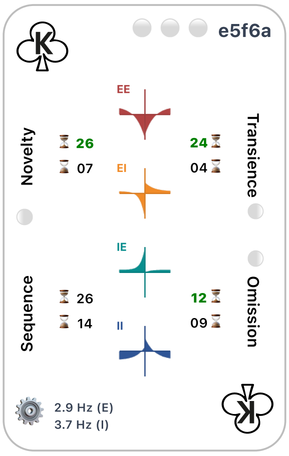
K
e5f6a
Novelty 26 07
Transience 24 04
Sequence 26 14
Omission 12 09
EE EI IE II
2.9 Hz (E)
3.7 Hz (I)

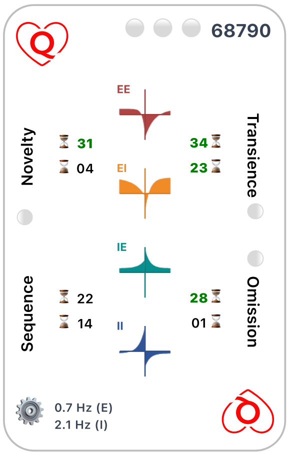
Q
68790
Novelty 31 04
Transience 34 23
Sequence 22 14
Omission 28 01
EE EI IE II
0.7 Hz (E)
2.1 Hz (I)

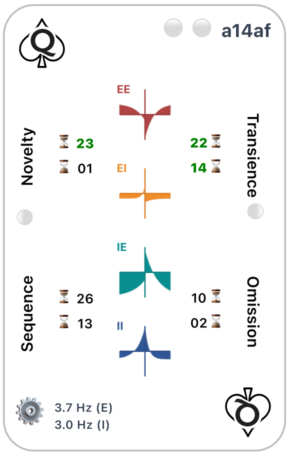
Q
a14af
Novelty 23 01
Transience 22 14
Sequence 26 13
Omission 10 02
EE EI IE II
3.7 Hz (E)
3.0 Hz (I)

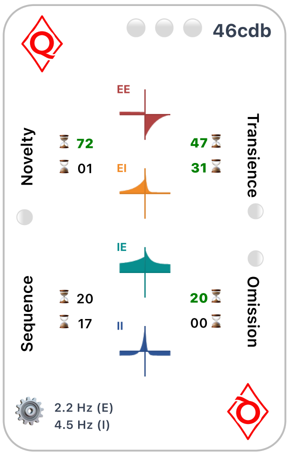
Q
46cdb
Novelty 72 01
Transience 47 31
Sequence 20 17
Omission 20 00
EE EI IE II
2.2 Hz (E)
4.5 Hz (I)

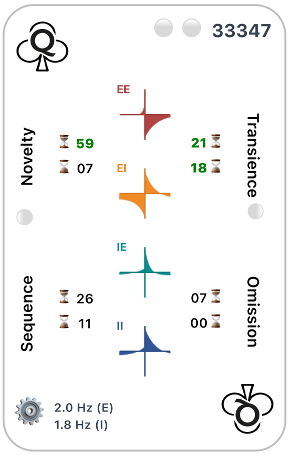
Q
33347
Novelty 59 07
Transience 21 18
Sequence 26 11
Omission 07 00
EE EI IE II
2.0 Hz (E)
1.8 Hz (I)

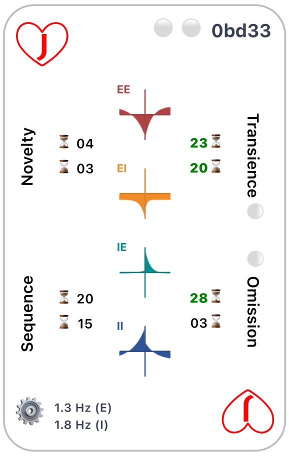
J
0bd33
Novelty 04 03
Transience 23 20
Sequence 20 15
Omission 28 03
EE EI IE II
1.3 Hz (E)
1.8 Hz (I)

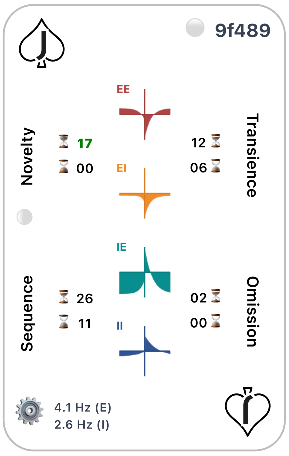
J
9f489
Novelty 17 00
Transience 12 06
Sequence 26 11
Omission 02 00
EE EI IE II
4.1 Hz (E)
2.6 Hz (I)

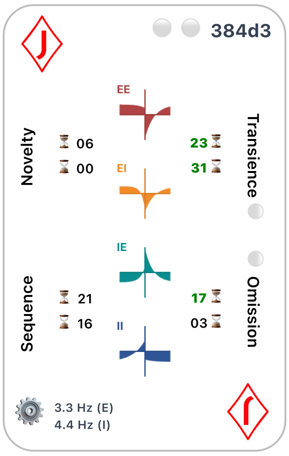
J
384d3
Novelty 06 00
Transience 23 31
Sequence 21 16
Omission 17 03
EE EI IE II
3.3 Hz (E)
4.4 Hz (I)

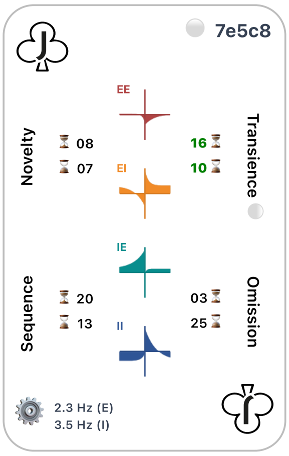
J
7e5c8
Novelty 08 07
Transience 16 10
Sequence 20 13
Omission 03 25
EE EI IE II
2.3 Hz (E)
3.5 Hz (I)

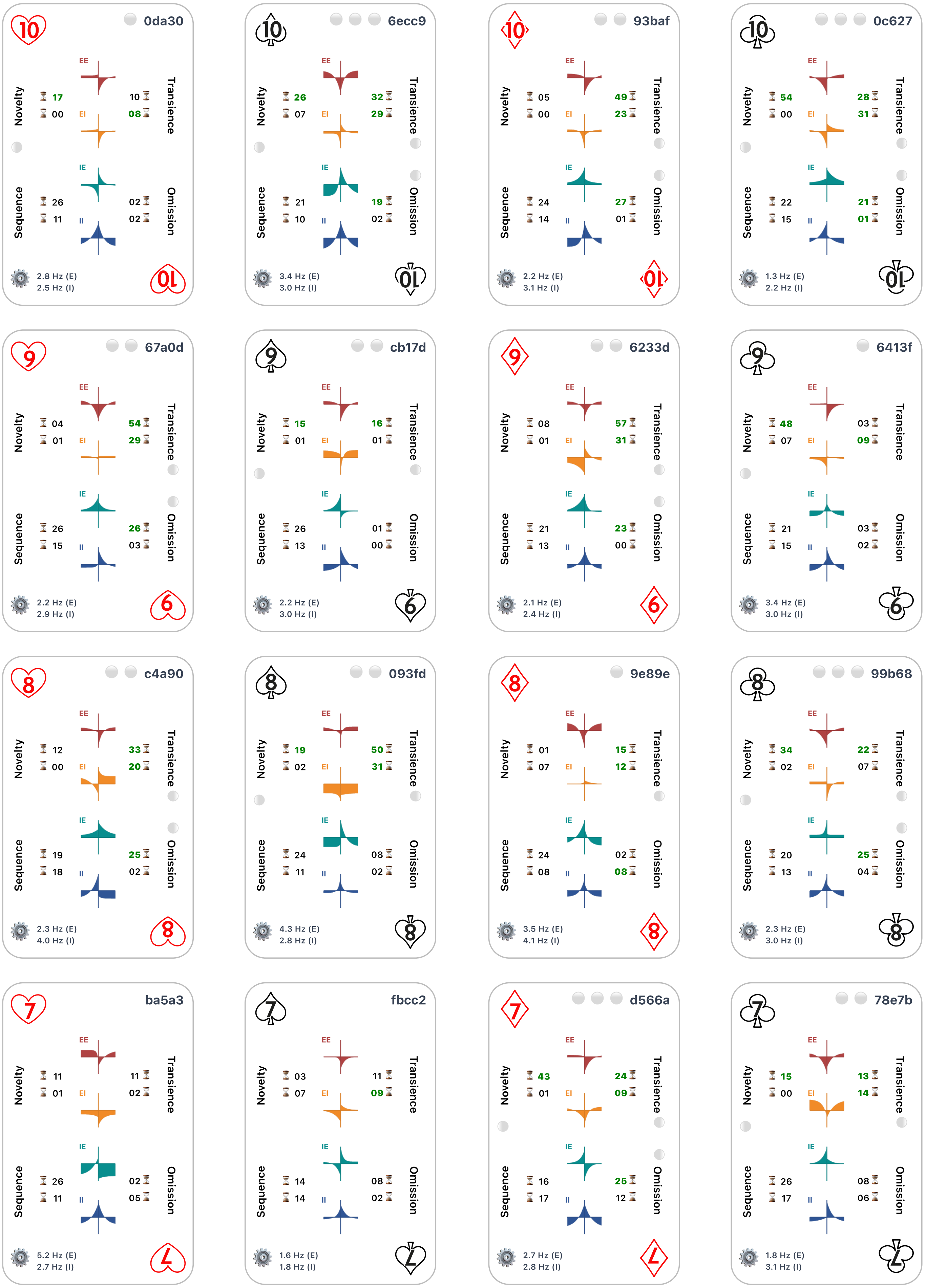

10 0da30 Novelty 17 00 Transience 10 08 Sequence 26 11 Omission 02 02 EE EI IE II 2.8 Hz (E) 2.5 Hz (I)
10 6ecc9 Novelty 26 07 Transience 32 29 Sequence 21 10 Omission 19 02 EE EI IE II 3.4 Hz (E) 3.0 Hz (I)
10 93baf Novelty 05 00 Transience 49 23 Sequence 24 14 Omission 27 01 EE EI IE II 2.2 Hz (E) 3.1 Hz (I)
10 0c627 Novelty 54 00 Transience 28 31 Sequence 22 15 Omission 21 01 EE EI IE II 1.3 Hz (E) 2.2 Hz (I)
9 67a0d Novelty 04 01 Transience 54 29 Sequence 26 15 Omission 26 03 EE EI IE II 2.2 Hz (E) 2.9 Hz (I)
9 cb17d Novelty 15 01 Transience 16 01 Sequence 26 13 Omission 01 00 EE EI IE II 2.2 Hz (E) 3.0 Hz (I)
9 6233d Novelty 08 01 Transience 57 31 Sequence 21 13 Omission 23 00 EE EI IE II 2.1 Hz (E) 2.4 Hz (I)
9 6413f Novelty 48 07 Transience 03 09 Sequence 21 15 Omission 03 02 EE EI IE II 3.4 Hz (E) 3.0 Hz (I)
8 c4a90 Novelty 12 00 Transience 33 20 Sequence 19 18 Omission 25 02 EE EI IE II 2.3 Hz (E) 4.0 Hz (I)
8 093fd Novelty 19 02 Transience 50 31 Sequence 24 11 Omission 08 02 EE EI IE II 4.3 Hz (E) 2.8 Hz (I)
8 9e89e Novelty 01 07 Transience 15 12 Sequence 24 08 Omission 02 08 EE EI IE II 3.5 Hz (E) 4.1 Hz (I)
8 99b68 Novelty 34 02 Transience 22 07 Sequence 20 13 Omission 25 04 EE EI IE II 2.3 Hz (E) 3.0 Hz (I)
7 ba5a3 Novelty 11 01 Transience 11 02 Sequence 26 11 Omission 02 05 EE EI IE II 5.2 Hz (E) 2.7 Hz (I)
7 fbcc2 Novelty 03 07 Transience 11 09 Sequence 14 14 Omission 08 02 EE EI IE II 1.6 Hz (E) 1.8 Hz (I)
7 d566a Novelty 43 01 Transience 24 09 Sequence 16 17 Omission 25 12 EE EI IE II 2.7 Hz (E) 2.8 Hz (I)
7 78e7b Novelty 15 00 Transience 13 14 Sequence 26 17 Omission 08 06 EE EI IE II 1.8 Hz (E) 3.1 Hz (I)

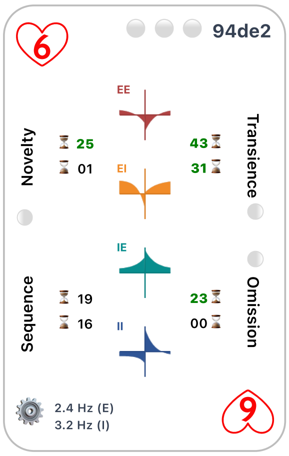
6
94de2
EE
EI
IE
II
Novelty
25
01
43
31
Transience
Sequence
19
16
23
00
Omission
2.4 Hz (E)
3.2 Hz (I)
9

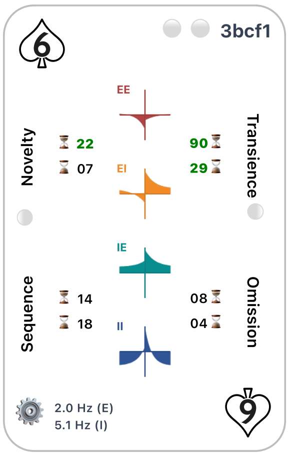
6
3bcf1
EE
EI
IE
II
Novelty
22
07
90
29
Transience
Sequence
14
18
08
04
Omission
2.0 Hz (E)
5.1 Hz (I)
9

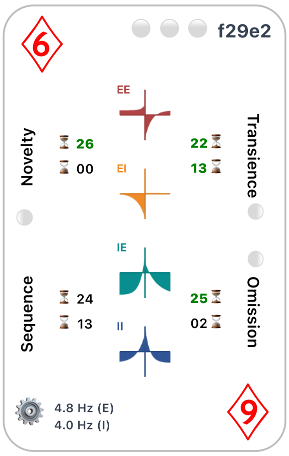
6
f29e2
EE
EI
IE
II
Novelty
26
00
22
13
Transience
Sequence
24
13
25
02
Omission
4.8 Hz (E)
4.0 Hz (I)
9

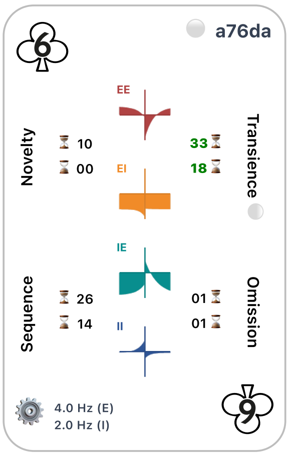
6
a76da
EE
EI
IE
II
Novelty
10
00
33
18
Transience
Sequence
26
14
01
01
Omission
4.0 Hz (E)
2.0 Hz (I)
9

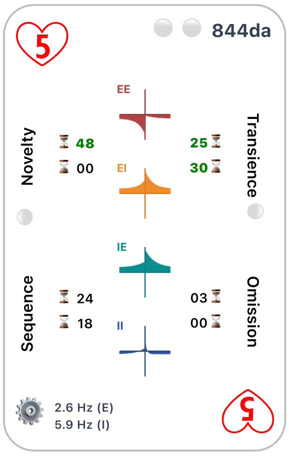
5
844da
EE
EI
IE
II
Novelty
48
00
25
30
Transience
Sequence
24
18
03
00
Omission
2.6 Hz (E)
5.9 Hz (I)
5

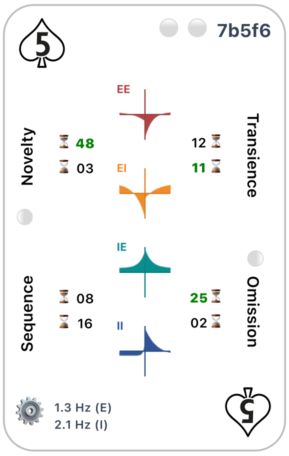
5
7b5f6
EE
EI
IE
II
Novelty
48
03
12
11
Transience
Sequence
08
16
25
02
Omission
1.3 Hz (E)
2.1 Hz (I)
5

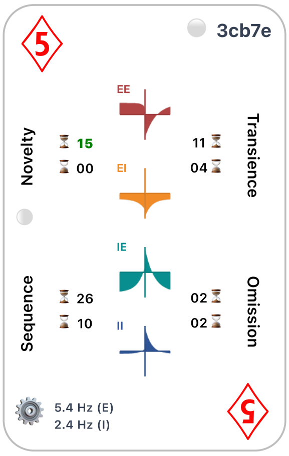
5
3cb7e
EE
EI
IE
II
Novelty
15
00
11
04
Transience
Sequence
26
10
02
02
Omission
5.4 Hz (E)
2.4 Hz (I)
5

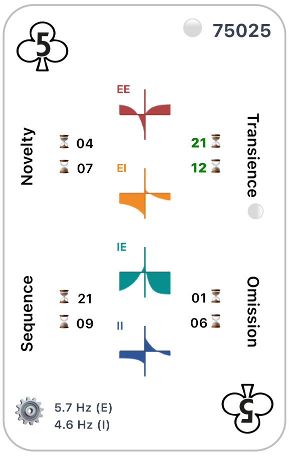
5
75025
EE
EI
IE
II
Novelty
04
07
21
12
Transience
Sequence
21
09
01
06
Omission
5.7 Hz (E)
4.6 Hz (I)
5

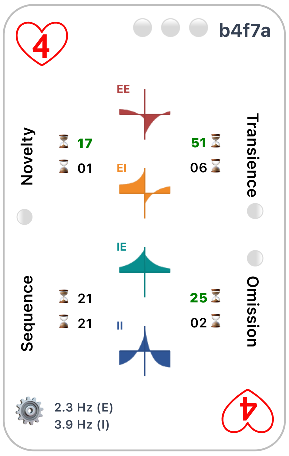
4
b4f7a
EE
EI
IE
II
Novelty
17
01
51
06
Transience
Sequence
21
21
25
02
Omission
2.3 Hz (E)
3.9 Hz (I)
4

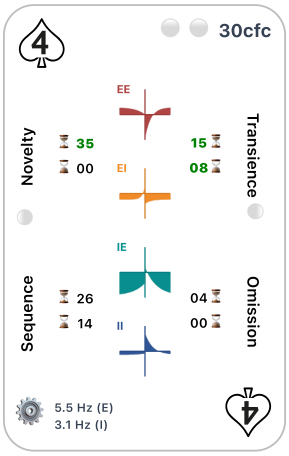
4
30cfc
EE
EI
IE
II
Novelty
35
00
15
08
Transience
Sequence
26
14
04
00
Omission
5.5 Hz (E)
3.1 Hz (I)
4

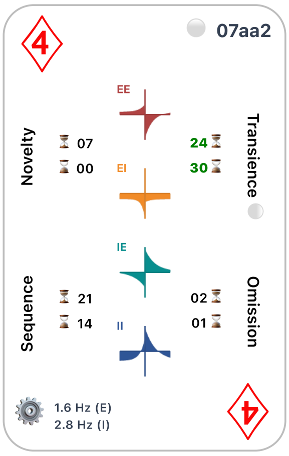
4
07aa2
EE
EI
IE
II
Novelty
07
00
24
30
Transience
Sequence
21
14
02
01
Omission
1.6 Hz (E)
2.8 Hz (I)
4

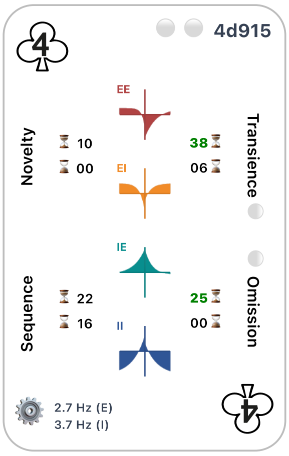
4
4d915
EE
EI
IE
II
Novelty
10
00
38
06
Transience
Sequence
22
16
25
00
Omission
2.7 Hz (E)
3.7 Hz (I)
4

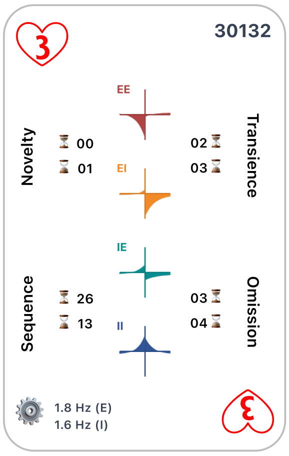
3
30132
EE
EI
IE
II
Novelty
00
01
02
03
Transience
Sequence
26
13
03
04
Omission
1.8 Hz (E)
1.6 Hz (I)
3

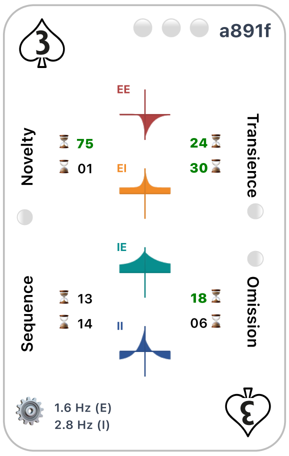
3
a891f
EE
EI
IE
II
Novelty
75
01
24
30
Transience
Sequence
13
14
18
06
Omission
1.6 Hz (E)
2.8 Hz (I)
3

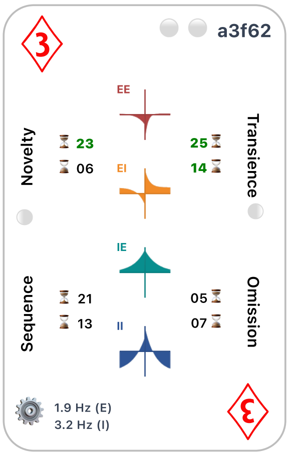
3
a3f62
EE
EI
IE
II
Novelty
23
06
25
14
Transience
Sequence
21
13
05
07
Omission
1.9 Hz (E)
3.2 Hz (I)
3

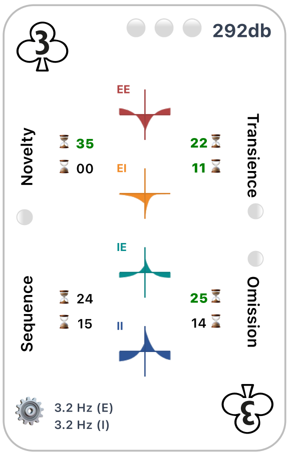
3
292db
EE
EI
IE
II
Novelty
35
00
22
11
Transience
Sequence
24
15
25
14
Omission
3.2 Hz (E)
3.2 Hz (I)
3

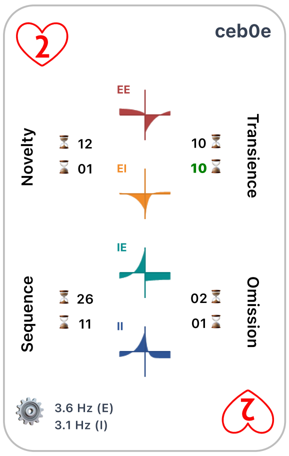
2
ceb0e
EE
EI
IE
II
Novelty
12
01
Transience
10
10
Sequence
26
11
Omission
02
01
3.6 Hz (E)
3.1 Hz (I)

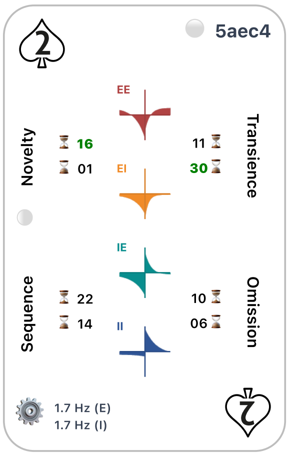
2
5aec4
EE
EI
IE
II
Novelty
16
01
Transience
11
30
Sequence
22
14
Omission
10
06
1.7 Hz (E)
1.7 Hz (I)

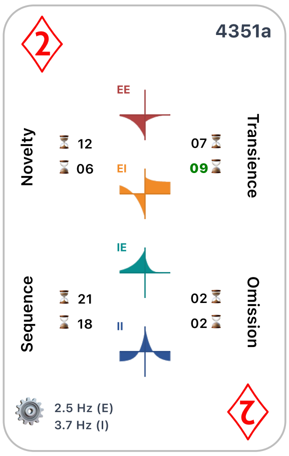
2
4351a
EE
EI
IE
II
Novelty
12
06
Transience
07
09
Sequence
21
18
Omission
02
02
2.5 Hz (E)
3.7 Hz (I)

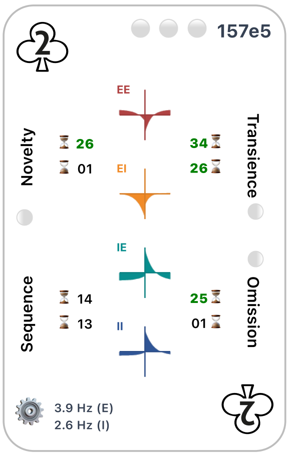
2
157e5
EE
EI
IE
II
Novelty
26
01
Transience
34
26
Sequence
14
13
Omission
25
01
3.9 Hz (E)
2.6 Hz (I)

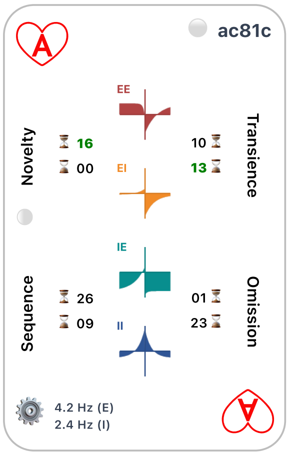
A
ac81c
EE
EI
IE
II
Novelty
16
00
Transience
10
13
Sequence
26
09
Omission
01
23
4.2 Hz (E)
2.4 Hz (I)

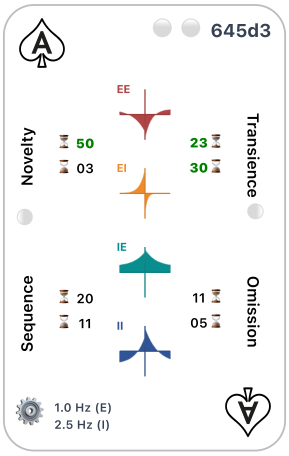
A
645d3
EE
EI
IE
II
Novelty
50
03
Transience
23
30
Sequence
20
11
Omission
11
05
1.0 Hz (E)
2.5 Hz (I)

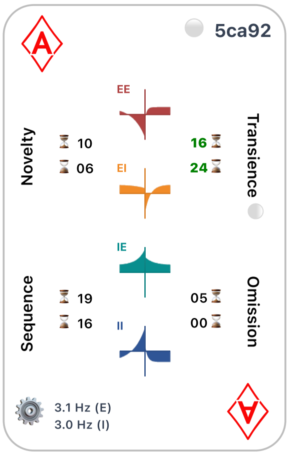
A
5ca92
EE
EI
IE
II
Novelty
10
06
Transience
16
24
Sequence
19
16
Omission
05
00
3.1 Hz (E)
3.0 Hz (I)

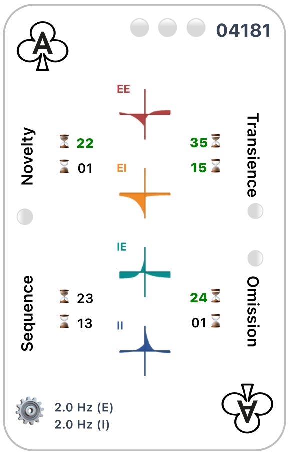
A
04181
EE
EI
IE
II
Novelty
22
01
Transience
35
15
Sequence
23
13
Omission
24
01
2.0 Hz (E)
2.0 Hz (I)

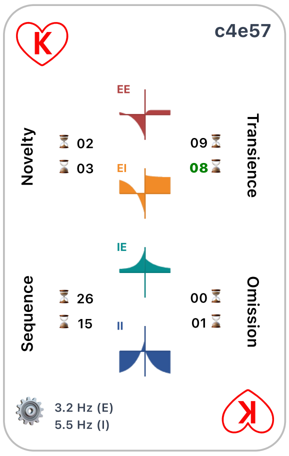
K
c4e57
EE
EI
IE
II
Novelty
02
03
Transience
09
08
Sequence
26
15
Omission
00
01
3.2 Hz (E)
5.5 Hz (I)

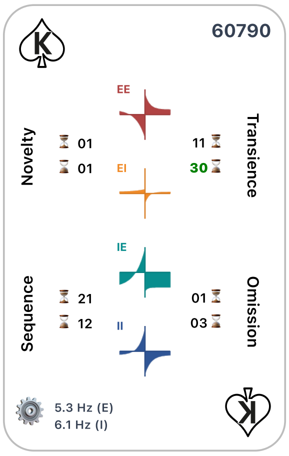
K
60790
EE
EI
IE
II
Novelty
01
01
Transience
11
30
Sequence
21
12
Omission
01
03
5.3 Hz (E)
6.1 Hz (I)

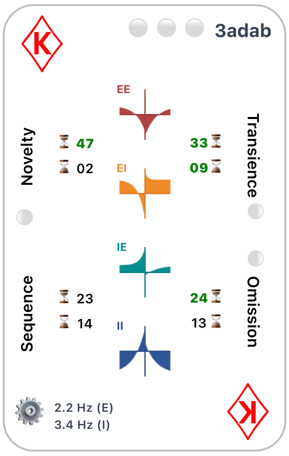
K
3adab
EE
EI
IE
II
Novelty
47
02
Transience
33
09
Sequence
23
14
Omission
24
13
2.2 Hz (E)
3.4 Hz (I)

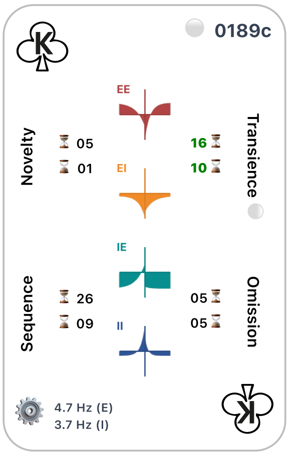
K
0189c
EE
EI
IE
II
Novelty
05
01
Transience
16
10
Sequence
26
09
Omission
05
05
4.7 Hz (E)
3.7 Hz (I)

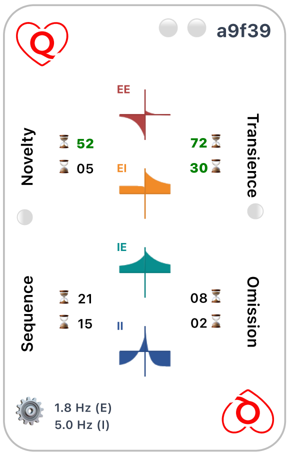
Q
a9f39
EE
EI
IE
II
Novelty
52
05
Transience
72
30
Sequence
21
15
Omission
08
02
1.8 Hz (E)
5.0 Hz (I)

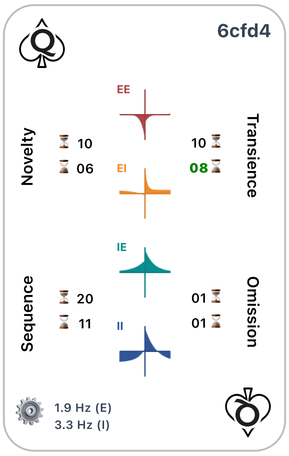
Q
6cfd4
EE
EI
IE
II
Novelty
10
06
Transience
10
08
Sequence
20
11
Omission
01
01
1.9 Hz (E)
3.3 Hz (I)

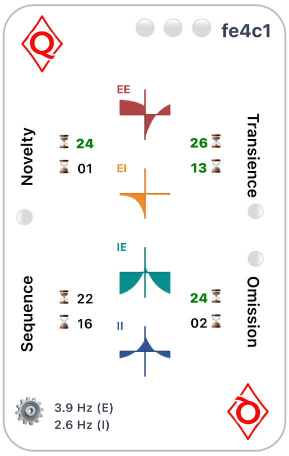
Q
fe4c1
EE
EI
IE
II
Novelty
24
01
Transience
26
13
Sequence
22
16
Omission
24
02
3.9 Hz (E)
2.6 Hz (I)

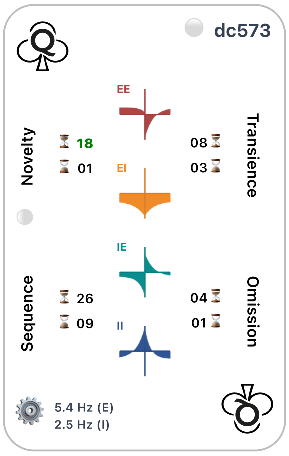
Q
dc573
EE
EI
IE
II
Novelty
18
01
Transience
08
03
Sequence
26
09
Omission
04
01
5.4 Hz (E)
2.5 Hz (I)

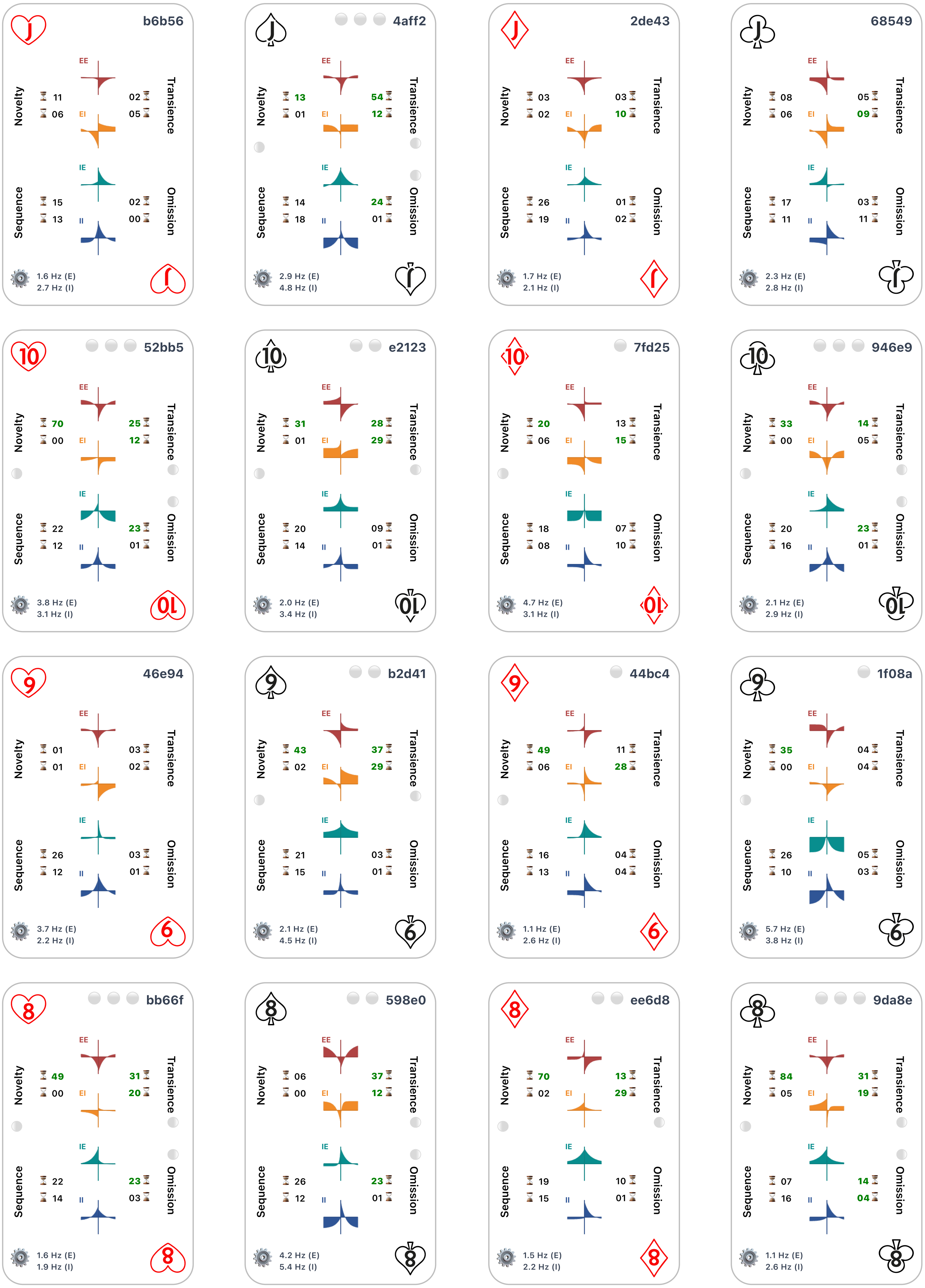
b6b56
Novelty 11 06 · Transience 02 05 · Sequence 15 13 · Omission 02 00
EE EI IE II
1.6 Hz (E) 2.7 Hz (I)
4aff2
Novelty 13 01 · Transience 54 12 · Sequence 14 18 · Omission 24 01
EE EI IE II
2.9 Hz (E) 4.8 Hz (I)
2de43
Novelty 03 02 · Transience 03 10 · Sequence 26 19 · Omission 01 02
EE EI IE II
1.7 Hz (E) 2.1 Hz (I)
68549
Novelty 08 06 · Transience 05 09 · Sequence 17 11 · Omission 03 11
EE EI IE II
2.3 Hz (E) 2.8 Hz (I)
52bb5
Novelty 70 00 · Transience 25 12 · Sequence 22 12 · Omission 23 01
EE EI IE II
3.8 Hz (E) 3.1 Hz (I)
e2123
Novelty 31 01 · Transience 28 29 · Sequence 20 14 · Omission 09 01
EE EI IE II
2.0 Hz (E) 3.4 Hz (I)
7fd25
Novelty 20 06 · Transience 13 15 · Sequence 18 08 · Omission 07 10
EE EI IE II
4.7 Hz (E) 3.1 Hz (I)
946e9
Novelty 33 00 · Transience 14 05 · Sequence 20 16 · Omission 23 01
EE EI IE II
2.1 Hz (E) 2.9 Hz (I)
46e94
Novelty 01 01 · Transience 03 02 · Sequence 26 12 · Omission 03 01
EE EI IE II
3.7 Hz (E) 2.2 Hz (I)
b2d41
Novelty 43 02 · Transience 37 29 · Sequence 21 15 · Omission 03 01
EE EI IE II
2.1 Hz (E) 4.5 Hz (I)
44bc4
Novelty 49 06 · Transience 11 28 · Sequence 16 13 · Omission 04 04
EE EI IE II
1.1 Hz (E) 2.6 Hz (I)
1f08a
Novelty 35 00 · Transience 04 04 · Sequence 26 10 · Omission 05 03
EE EI IE II
5.7 Hz (E) 3.8 Hz (I)
bb66f
Novelty 49 00 · Transience 31 20 · Sequence 22 14 · Omission 23 03
EE EI IE II
1.6 Hz (E) 1.9 Hz (I)
598e0
Novelty 06 00 · Transience 37 12 · Sequence 26 12 · Omission 23 01
EE EI IE II
4.2 Hz (E) 5.4 Hz (I)
ee6d8
Novelty 70 02 · Transience 13 29 · Sequence 19 15 · Omission 10 01
EE EI IE II
1.5 Hz (E) 2.2 Hz (I)
9da8e
Novelty 84 05 · Transience 31 19 · Sequence 07 16 · Omission 14 04
EE EI IE II
1.1 Hz (E) 2.6 Hz (I)

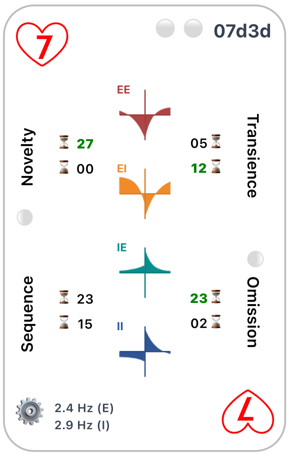
07d3d
Novelty
27
00
Transience
05
12
EE
EI
IE
II
Sequence
23
15
Omission
23
02
2.4 Hz (E)
2.9 Hz (I)

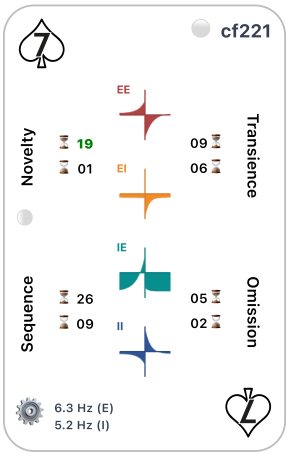
cf221
Novelty
19
01
Transience
09
06
EE
EI
IE
II
Sequence
26
09
Omission
05
02
6.3 Hz (E)
5.2 Hz (I)

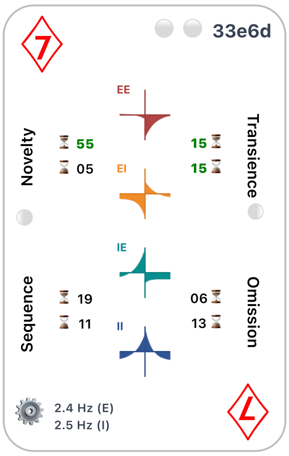
33e6d
Novelty
55
05
Transience
15
15
EE
EI
IE
II
Sequence
19
11
Omission
06
13
2.4 Hz (E)
2.5 Hz (I)

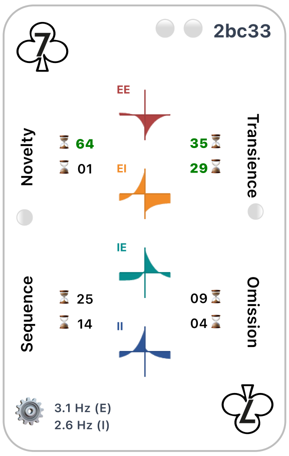
2bc33
Novelty
64
01
Transience
35
29
EE
EI
IE
II
Sequence
25
14
Omission
09
04
3.1 Hz (E)
2.6 Hz (I)

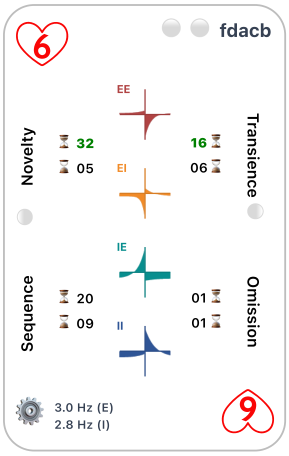
fdacb
Novelty
32
05
Transience
16
06
EE
EI
IE
II
Sequence
20
09
Omission
01
01
3.0 Hz (E)
2.8 Hz (I)

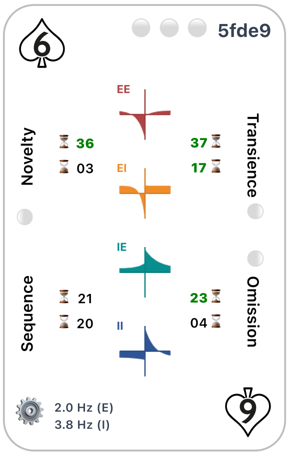
5fde9
Novelty
36
03
Transience
37
17
EE
EI
IE
II
Sequence
21
20
Omission
23
04
2.0 Hz (E)
3.8 Hz (I)

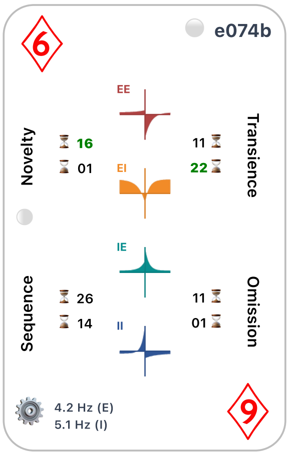
e074b
Novelty
16
01
Transience
11
22
EE
EI
IE
II
Sequence
26
14
Omission
11
01
4.2 Hz (E)
5.1 Hz (I)

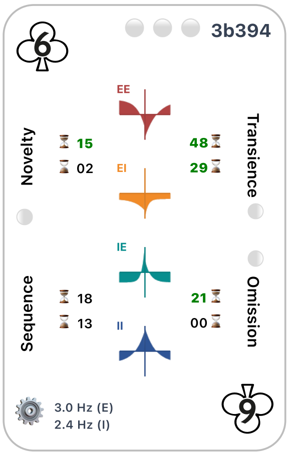
3b394
Novelty
15
02
Transience
48
29
EE
EI
IE
II
Sequence
18
13
Omission
21
00
3.0 Hz (E)
2.4 Hz (I)

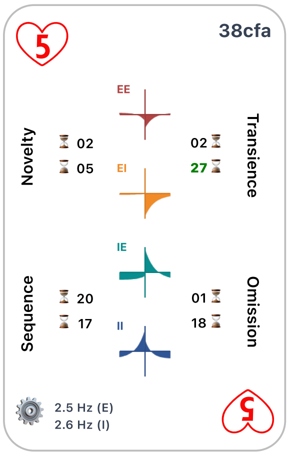
38cfa
Novelty
02
05
Transience
02
27
EE
EI
IE
II
Sequence
20
17
Omission
01
18
2.5 Hz (E)
2.6 Hz (I)

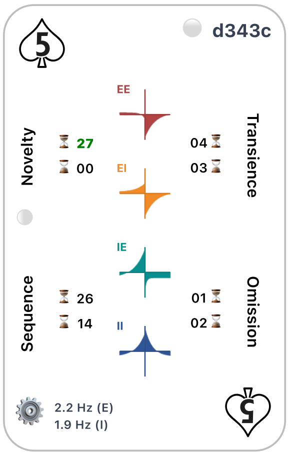
d343c
Novelty
27
00
Transience
04
03
EE
EI
IE
II
Sequence
26
14
Omission
01
02
2.2 Hz (E)
1.9 Hz (I)

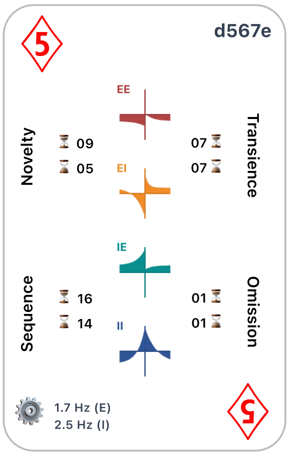
d567e
Novelty
09
05
Transience
07
07
EE
EI
IE
II
Sequence
16
14
Omission
01
01
1.7 Hz (E)
2.5 Hz (I)

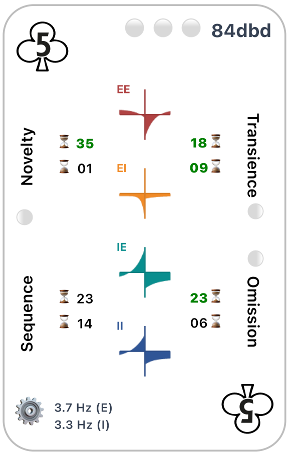
84dbd
Novelty
35
01
Transience
18
09
EE
EI
IE
II
Sequence
23
14
Omission
23
06
3.7 Hz (E)
3.3 Hz (I)

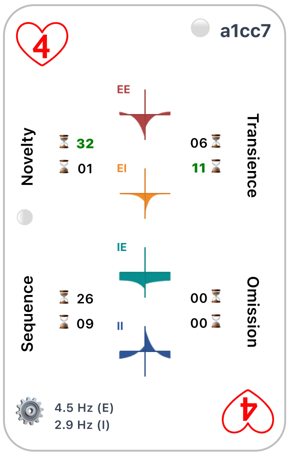
a1cc7
Novelty
32
01
Transience
06
11
EE
EI
IE
II
Sequence
26
09
Omission
00
00
4.5 Hz (E)
2.9 Hz (I)

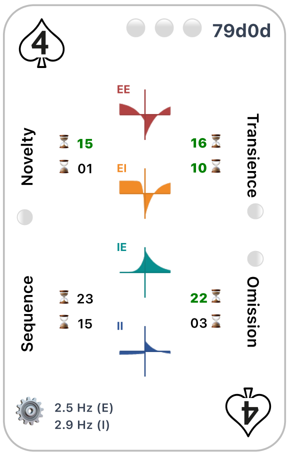
79d0d
Novelty
15
01
Transience
16
10
EE
EI
IE
II
Sequence
23
15
Omission
22
03
2.5 Hz (E)
2.9 Hz (I)

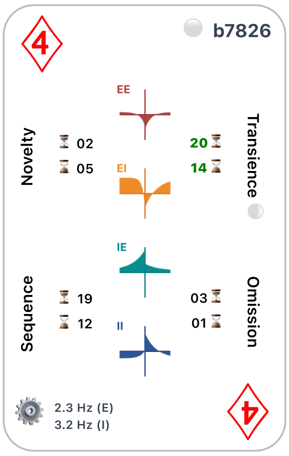
b7826
Novelty
02
05
Transience
20
14
EE
EI
IE
II
Sequence
19
12
Omission
03
01
2.3 Hz (E)
3.2 Hz (I)

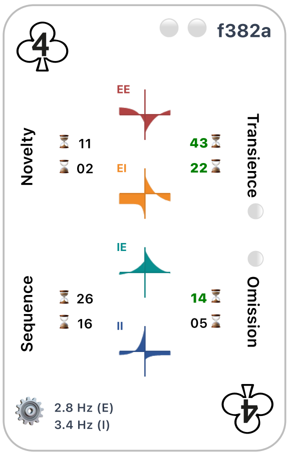
f382a
Novelty
11
02
Transience
43
22
EE
EI
IE
II
Sequence
26
16
Omission
14
05
2.8 Hz (E)
3.4 Hz (I)

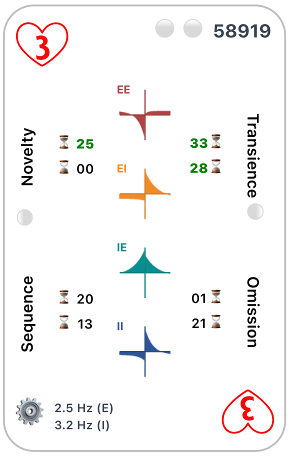
3
58919
Novelty
25
00
EE
EI
33
28
Transience
Sequence
20
13
IE
II
01
21
Omission
2.5 Hz (E)
3.2 Hz (I)
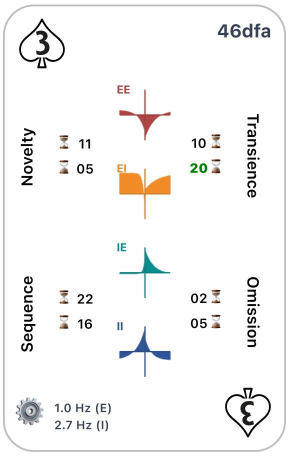
3
46dfa
Novelty
11
05
EE
EI
10
20
Transience
Sequence
22
16
IE
II
02
05
Omission
1.0 Hz (E)
2.7 Hz (I)
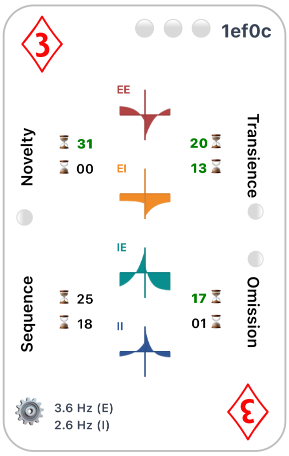
3
1ef0c
Novelty
31
00
EE
EI
20
13
Transience
Sequence
25
18
IE
II
17
01
Omission
3.6 Hz (E)
2.6 Hz (I)
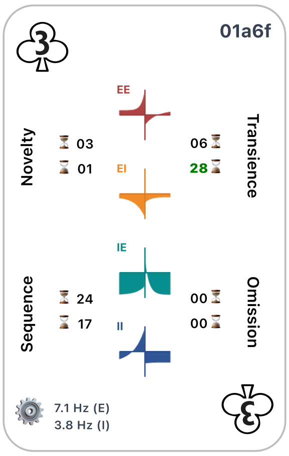
3
01a6f
Novelty
03
01
EE
EI
06
28
Transience
Sequence
24
17
IE
II
00
00
Omission
7.1 Hz (E)
3.8 Hz (I)
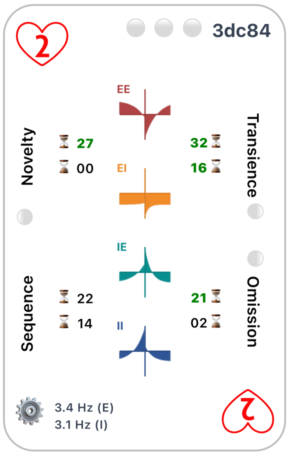
2
3dc84
Novelty
27
00
EE
EI
32
16
Transience
Sequence
22
14
IE
II
21
02
Omission
3.4 Hz (E)
3.1 Hz (I)
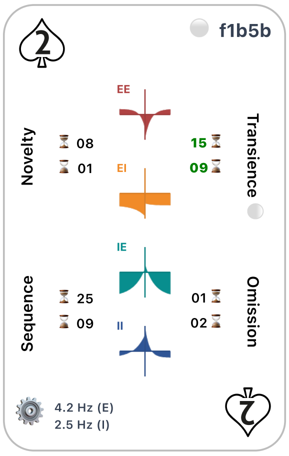
2
f1b5b
Novelty
08
01
EE
EI
15
09
Transience
Sequence
25
09
IE
II
01
02
Omission
4.2 Hz (E)
2.5 Hz (I)
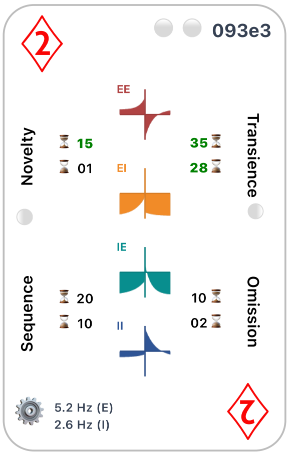
2
093e3
Novelty
15
01
EE
EI
35
28
Transience
Sequence
20
10
IE
II
10
02
Omission
5.2 Hz (E)
2.6 Hz (I)
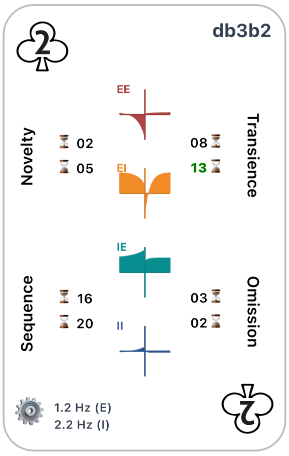
2
db3b2
Novelty
02
05
EE
EI
08
13
Transience
Sequence
16
20
IE
II
03
02
Omission
1.2 Hz (E)
2.2 Hz (I)
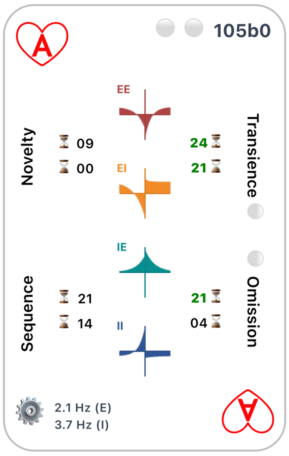
A
105b0
Novelty
09
00
EE
EI
24
21
Transience
Sequence
21
14
IE
II
21
04
Omission
2.1 Hz (E)
3.7 Hz (I)
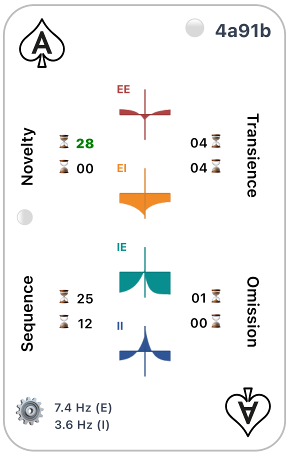
A
4a91b
Novelty
28
00
EE
EI
04
04
Transience
Sequence
25
12
IE
II
01
00
Omission
7.4 Hz (E)
3.6 Hz (I)
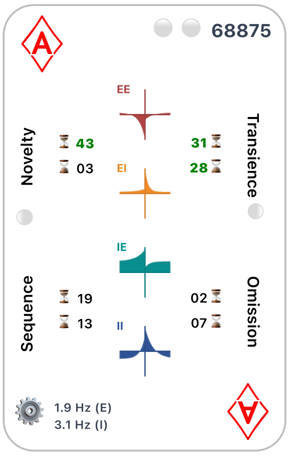
A
68875
Novelty
43
03
EE
EI
31
28
Transience
Sequence
19
13
IE
II
02
07
Omission
1.9 Hz (E)
3.1 Hz (I)
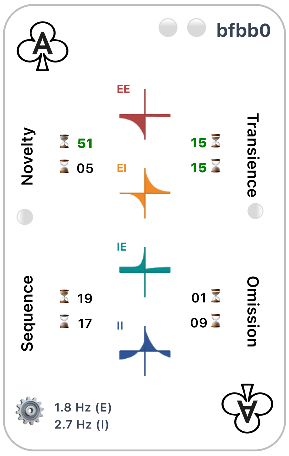
A
bfbb0
Novelty
51
05
EE
EI
15
15
Transience
Sequence
19
17
IE
II
01
09
Omission
1.8 Hz (E)
2.7 Hz (I)
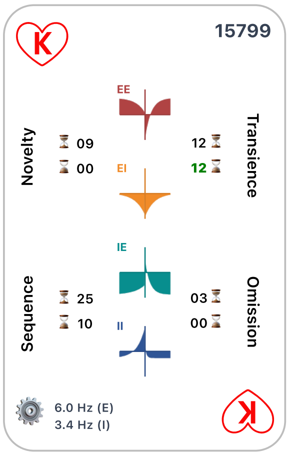
K
15799
Novelty
09
00
EE
EI
12
12
Transience
Sequence
25
10
IE
II
03
00
Omission
6.0 Hz (E)
3.4 Hz (I)
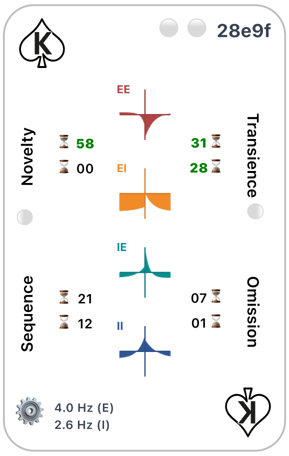
K
28e9f
Novelty
58
00
EE
EI
31
28
Transience
Sequence
21
12
IE
II
07
01
Omission
4.0 Hz (E)
2.6 Hz (I)
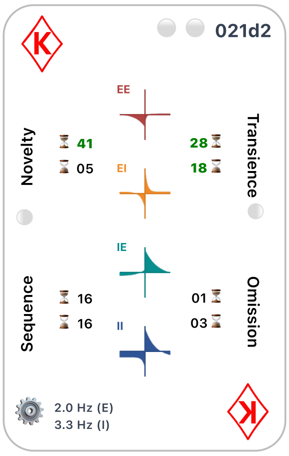
K
021d2
Novelty
41
05
EE
EI
28
18
Transience
Sequence
16
16
IE
II
01
03
Omission
2.0 Hz (E)
3.3 Hz (I)
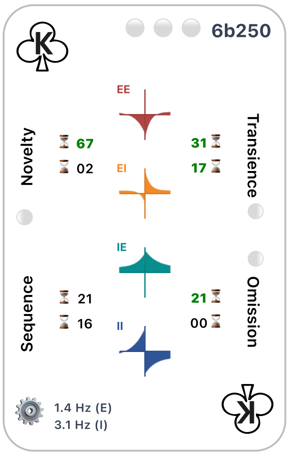
K
6b250
Novelty
67
02
EE
EI
31
17
Transience
Sequence
21
16
IE
II
21
00
Omission
1.4 Hz (E)
3.1 Hz (I)

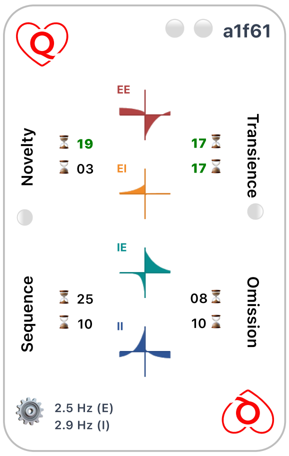
Q
a1f61
EE
EI
IE
II
Novelty
19
03
17
17
Transience
Sequence
25
10
08
10
Omission
2.5 Hz (E)
2.9 Hz (I)

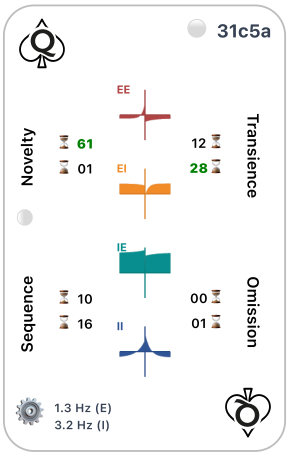
Q
31c5a
EE
EI
IE
II
Novelty
61
01
12
28
Transience
Sequence
10
16
00
01
Omission
1.3 Hz (E)
3.2 Hz (I)

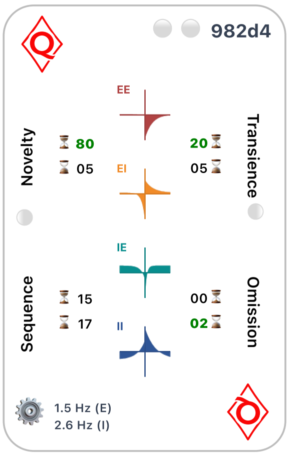
Q
982d4
EE
EI
IE
II
Novelty
80
05
20
05
Transience
Sequence
15
17
00
02
Omission
1.5 Hz (E)
2.6 Hz (I)

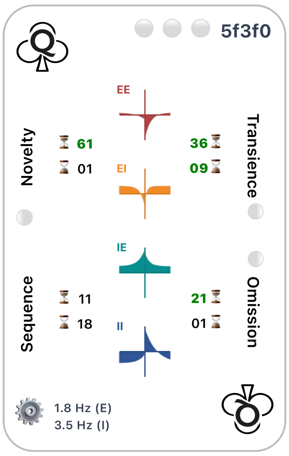
Q
5f3f0
EE
EI
IE
II
Novelty
61
01
36
09
Transience
Sequence
11
18
21
01
Omission
1.8 Hz (E)
3.5 Hz (I)

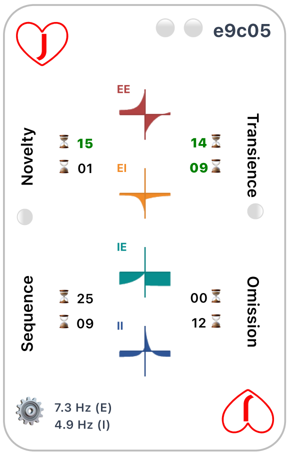
J
e9c05
EE
EI
IE
II
Novelty
15
01
14
09
Transience
Sequence
25
09
00
12
Omission
7.3 Hz (E)
4.9 Hz (I)

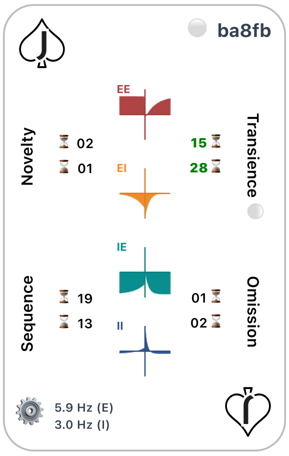
J
ba8fb
EE
EI
IE
II
Novelty
02
01
15
28
Transience
Sequence
19
13
01
02
Omission
5.9 Hz (E)
3.0 Hz (I)

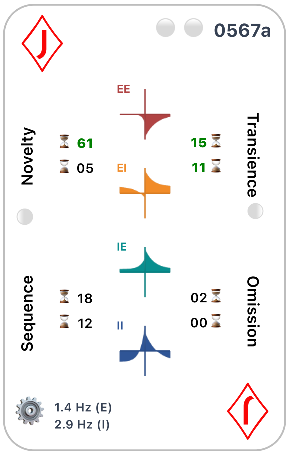
J
0567a
EE
EI
IE
II
Novelty
61
05
15
11
Transience
Sequence
18
12
02
00
Omission
1.4 Hz (E)
2.9 Hz (I)

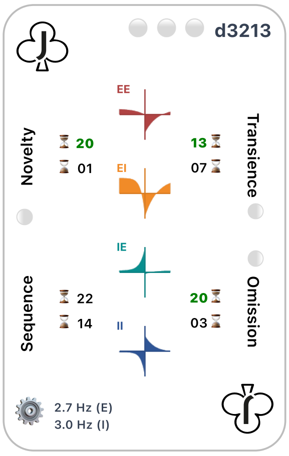
J
d3213
EE
EI
IE
II
Novelty
20
01
13
07
Transience
Sequence
22
14
20
03
Omission
2.7 Hz (E)
3.0 Hz (I)

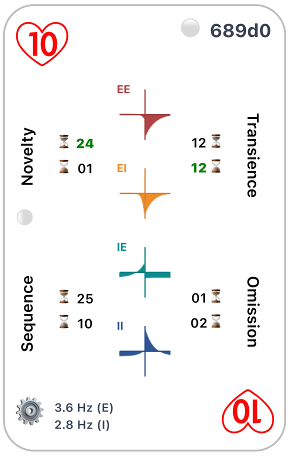
10
689d0
EE
EI
IE
II
Novelty
24
01
12
12
Transience
Sequence
25
10
01
02
Omission
3.6 Hz (E)
2.8 Hz (I)

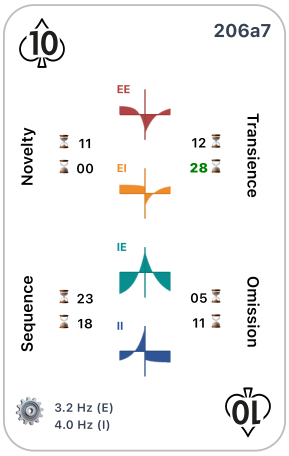
10
206a7
EE
EI
IE
II
Novelty
11
00
12
28
Transience
Sequence
23
18
05
11
Omission
3.2 Hz (E)
4.0 Hz (I)

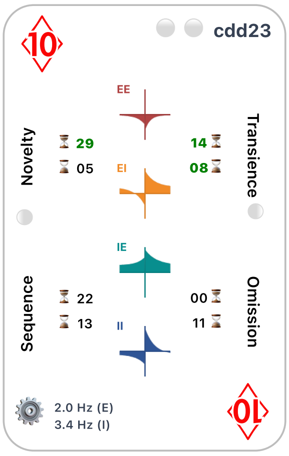
10
cdd23
EE
EI
IE
II
Novelty
29
05
14
08
Transience
Sequence
22
13
00
11
Omission
2.0 Hz (E)
3.4 Hz (I)

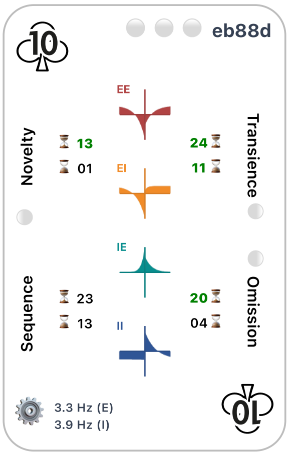
10
eb88d
EE
EI
IE
II
Novelty
13
01
24
11
Transience
Sequence
23
13
20
04
Omission
3.3 Hz (E)
3.9 Hz (I)

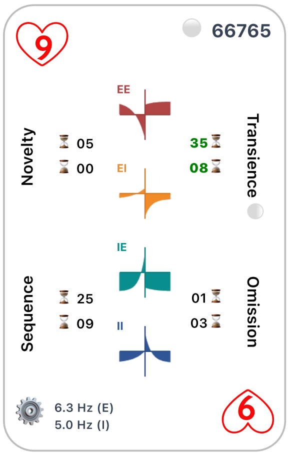
9
66765
EE
EI
IE
II
Novelty
05
00
35
08
Transience
Sequence
25
09
01
03
Omission
6.3 Hz (E)
5.0 Hz (I)

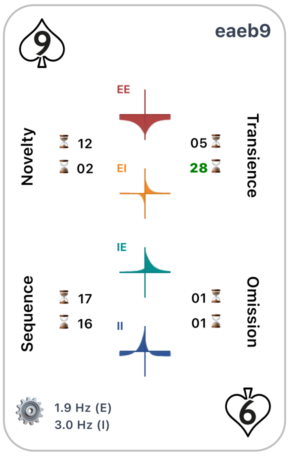
9
eaeb9
EE
EI
IE
II
Novelty
12
02
05
28
Transience
Sequence
17
16
01
01
Omission
1.9 Hz (E)
3.0 Hz (I)

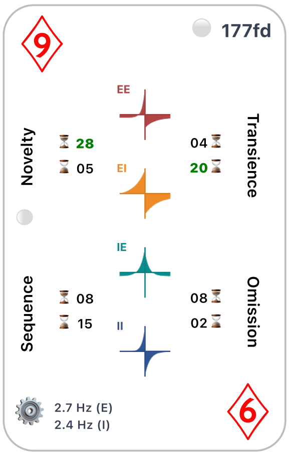
9
177fd
EE
EI
IE
II
Novelty
28
05
04
20
Transience
Sequence
08
15
08
02
Omission
2.7 Hz (E)
2.4 Hz (I)

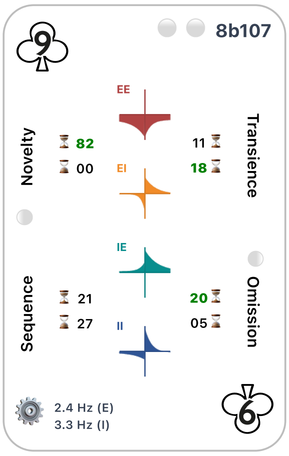
9
8b107
EE
EI
IE
II
Novelty
82
00
11
18
Transience
Sequence
21
27
20
05
Omission
2.4 Hz (E)
3.3 Hz (I)

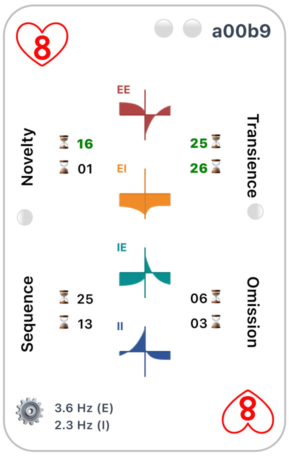
8
a00b9
Novelty
16
01
EE
EI
25
26
Transience
Sequence
25
13
IE
II
06
03
Omission
3.6 Hz (E)
2.3 Hz (I)
8
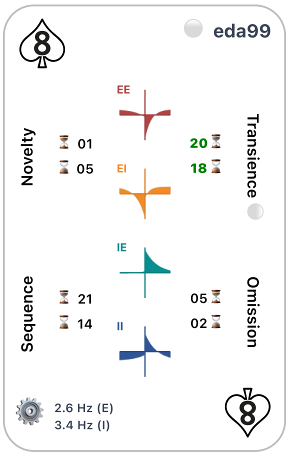
8
eda99
Novelty
01
05
EE
EI
20
18
Transience
Sequence
21
14
IE
II
05
02
Omission
2.6 Hz (E)
3.4 Hz (I)
8
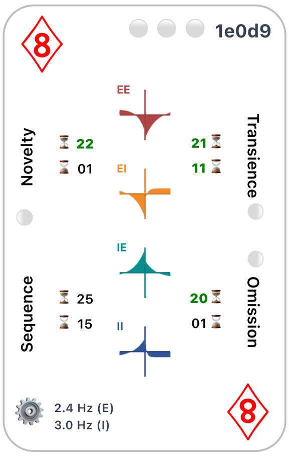
8
1e0d9
Novelty
22
01
EE
EI
21
11
Transience
Sequence
25
15
IE
II
20
01
Omission
2.4 Hz (E)
3.0 Hz (I)
8
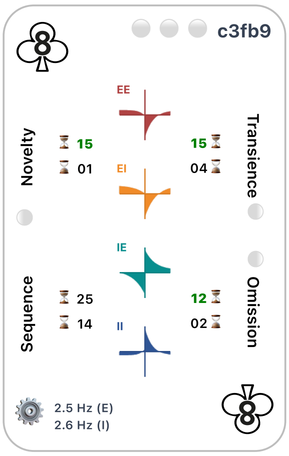
8
c3fb9
Novelty
15
01
EE
EI
15
04
Transience
Sequence
25
14
IE
II
12
02
Omission
2.5 Hz (E)
2.6 Hz (I)
8
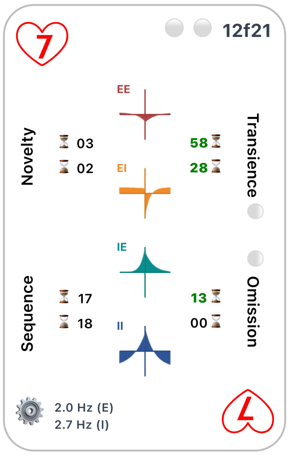
7
12f21
Novelty
03
02
EE
EI
58
28
Transience
Sequence
17
18
IE
II
13
00
Omission
2.0 Hz (E)
2.7 Hz (I)
7
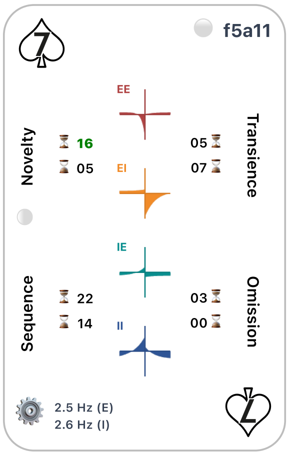
7
f5a11
Novelty
16
05
EE
EI
05
07
Transience
Sequence
22
14
IE
II
03
00
Omission
2.5 Hz (E)
2.6 Hz (I)
7
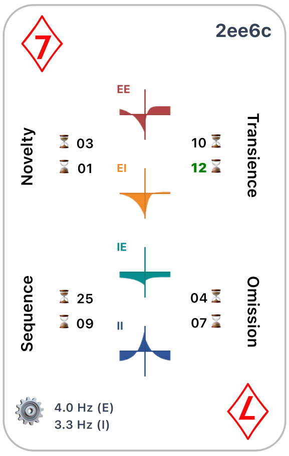
7
2ee6c
Novelty
03
01
EE
EI
10
12
Transience
Sequence
25
09
IE
II
04
07
Omission
4.0 Hz (E)
3.3 Hz (I)
7
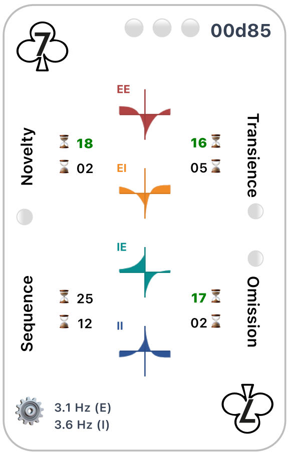
7
00d85
Novelty
18
02
EE
EI
16
05
Transience
Sequence
25
12
IE
II
17
02
Omission
3.1 Hz (E)
3.6 Hz (I)
7
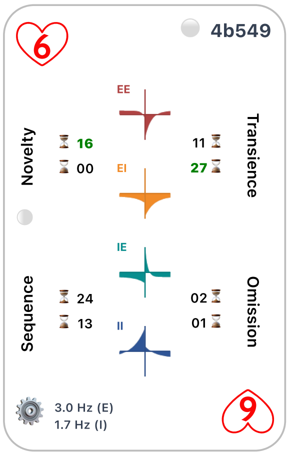
6
4b549
Novelty
16
00
EE
EI
11
27
Transience
Sequence
24
13
IE
II
02
01
Omission
3.0 Hz (E)
1.7 Hz (I)
9
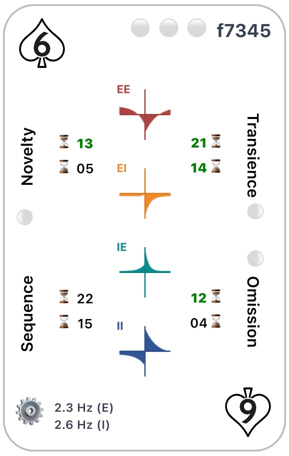
6
f7345
Novelty
13
05
EE
EI
21
14
Transience
Sequence
22
15
IE
II
12
04
Omission
2.3 Hz (E)
2.6 Hz (I)
9
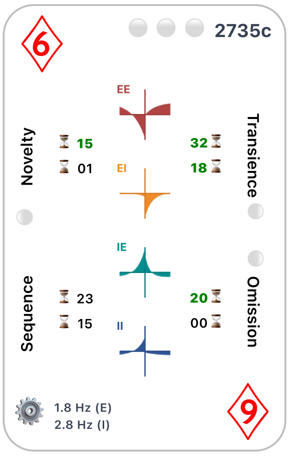
6
2735c
Novelty
15
01
EE
EI
32
18
Transience
Sequence
23
15
IE
II
20
00
Omission
1.8 Hz (E)
2.8 Hz (I)
9
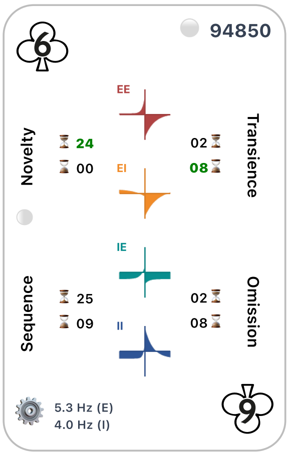
6
94850
Novelty
24
00
EE
EI
02
08
Transience
Sequence
25
09
IE
II
02
08
Omission
5.3 Hz (E)
4.0 Hz (I)
9
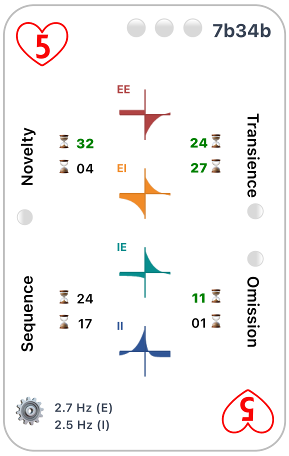
5
7b34b
Novelty
32
04
EE
EI
24
27
Transience
Sequence
24
17
IE
II
11
01
Omission
2.7 Hz (E)
2.5 Hz (I)
5
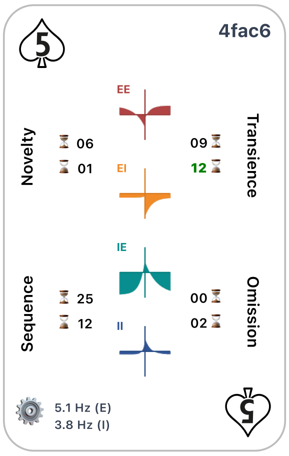
5
4fac6
Novelty
06
01
EE
EI
09
12
Transience
Sequence
25
12
IE
II
00
02
Omission
5.1 Hz (E)
3.8 Hz (I)
5
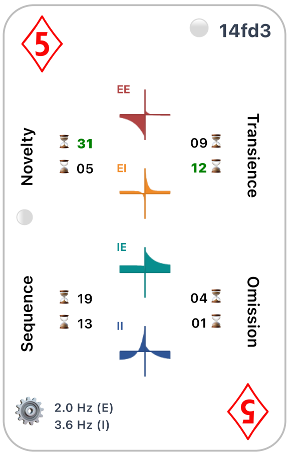
5
14fd3
Novelty
31
05
EE
EI
09
12
Transience
Sequence
19
13
IE
II
04
01
Omission
2.0 Hz (E)
3.6 Hz (I)
5
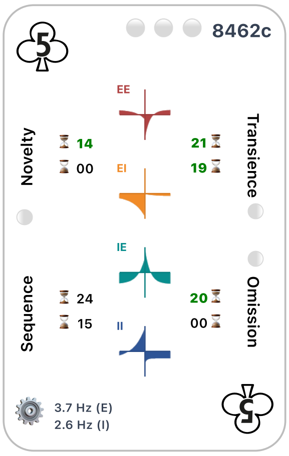
5
8462c
Novelty
14
00
EE
EI
21
19
Transience
Sequence
24
15
IE
II
20
00
Omission
3.7 Hz (E)
2.6 Hz (I)
5

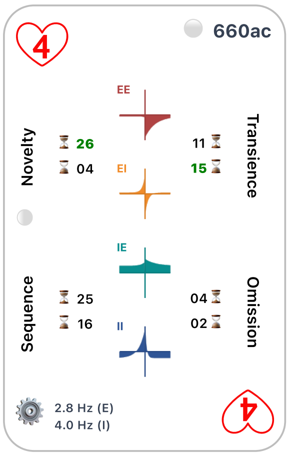
4
660ac
EE
Novelty
26
04
11
15
Transience
EI
IE
Sequence
25
16
04
02
Omission
II
2.8 Hz (E)
4.0 Hz (I)

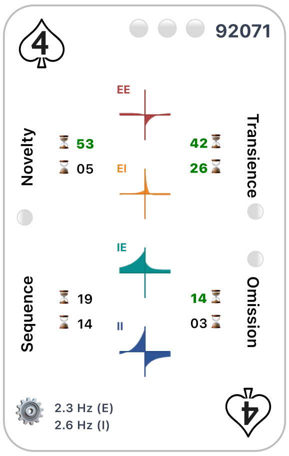
4
92071
EE
Novelty
53
05
42
26
Transience
EI
IE
Sequence
19
14
14
03
Omission
II
2.3 Hz (E)
2.6 Hz (I)

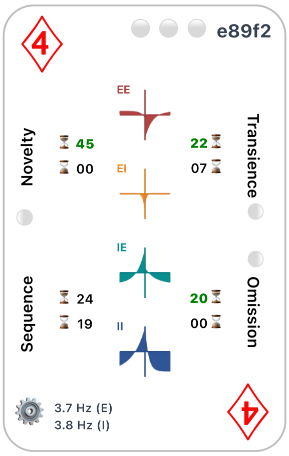
4
e89f2
EE
Novelty
45
00
22
07
Transience
EI
IE
Sequence
24
19
20
00
Omission
II
3.7 Hz (E)
3.8 Hz (I)

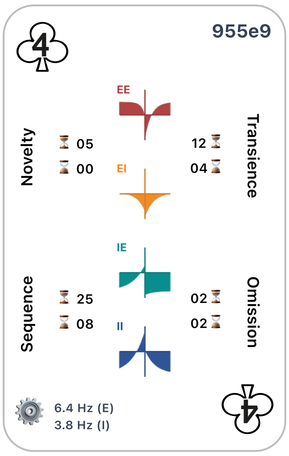
4
955e9
EE
Novelty
05
00
12
04
Transience
EI
IE
Sequence
25
08
02
02
Omission
II
6.4 Hz (E)
3.8 Hz (I)

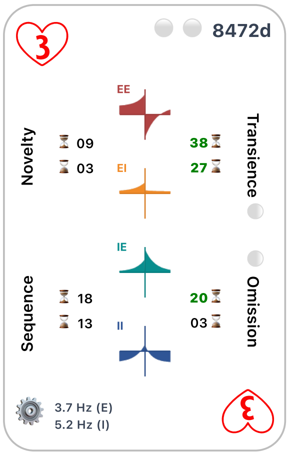
3
8472d
EE
Novelty
09
03
38
27
Transience
EI
IE
Sequence
18
13
20
03
Omission
II
3.7 Hz (E)
5.2 Hz (I)

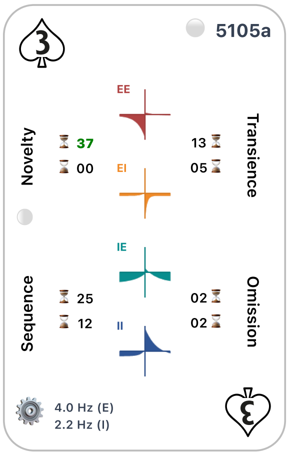
3
5105a
EE
Novelty
37
00
13
05
Transience
EI
IE
Sequence
25
12
02
02
Omission
II
4.0 Hz (E)
2.2 Hz (I)

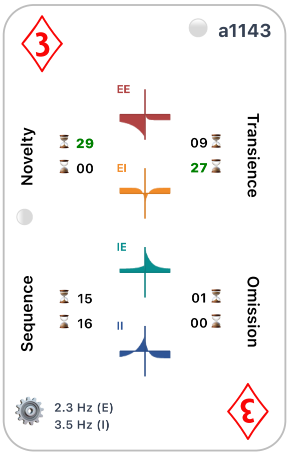
3
a1143
EE
Novelty
29
00
09
27
Transience
EI
IE
Sequence
15
16
01
00
Omission
II
2.3 Hz (E)
3.5 Hz (I)

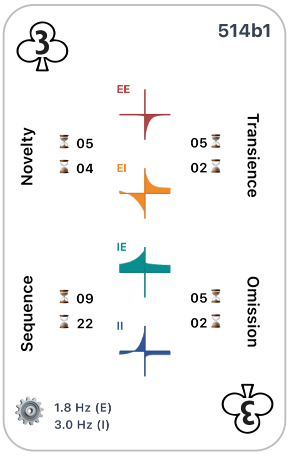
3
514b1
EE
Novelty
05
04
05
02
Transience
EI
IE
Sequence
09
22
05
02
Omission
II
1.8 Hz (E)
3.0 Hz (I)

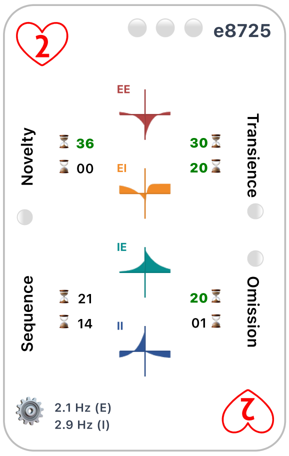
2
e8725
EE
Novelty
36
00
30
20
Transience
EI
IE
Sequence
21
14
20
01
Omission
II
2.1 Hz (E)
2.9 Hz (I)

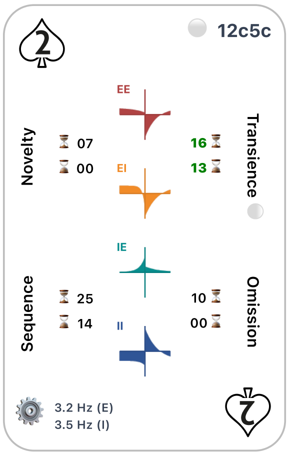
2
12c5c
EE
Novelty
07
00
16
13
Transience
EI
IE
Sequence
25
14
10
00
Omission
II
3.2 Hz (E)
3.5 Hz (I)

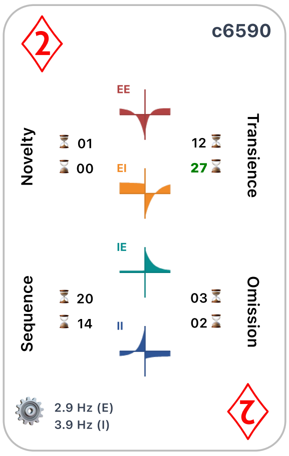
2
c6590
EE
Novelty
01
00
12
27
Transience
EI
IE
Sequence
20
14
03
02
Omission
II
2.9 Hz (E)
3.9 Hz (I)

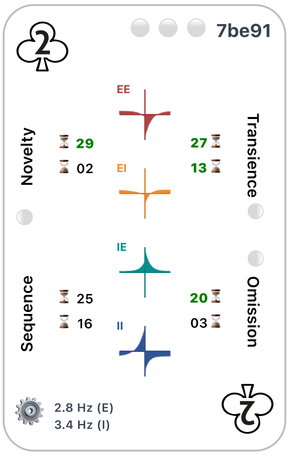
2
7be91
EE
Novelty
29
02
27
13
Transience
EI
IE
Sequence
25
16
20
03
Omission
II
2.8 Hz (E)
3.4 Hz (I)

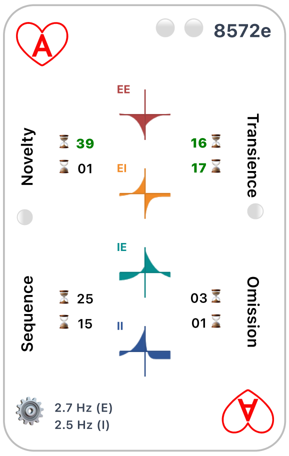
A
8572e
EE
Novelty
39
01
16
17
Transience
EI
IE
Sequence
25
15
03
01
Omission
II
2.7 Hz (E)
2.5 Hz (I)

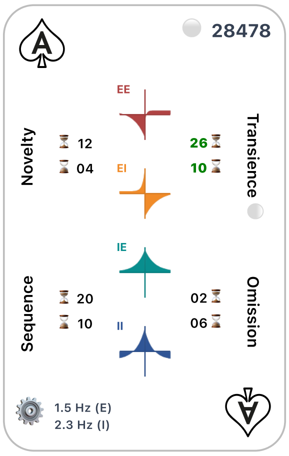
A
28478
EE
Novelty
12
04
26
10
Transience
EI
IE
Sequence
20
10
02
06
Omission
II
1.5 Hz (E)
2.3 Hz (I)

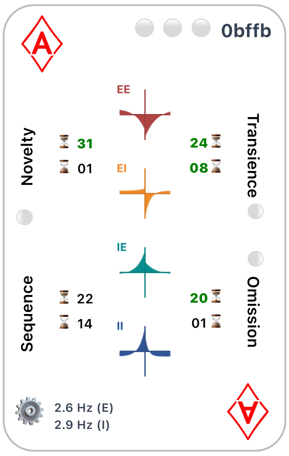
A
0bffb
EE
Novelty
31
01
24
08
Transience
EI
IE
Sequence
22
14
20
01
Omission
II
2.6 Hz (E)
2.9 Hz (I)

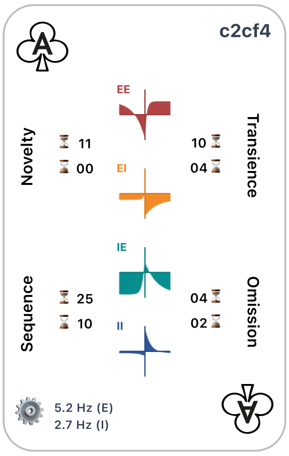
A
c2cf4
EE
Novelty
11
00
10
04
Transience
EI
IE
Sequence
25
10
04
02
Omission
II
5.2 Hz (E)
2.7 Hz (I)

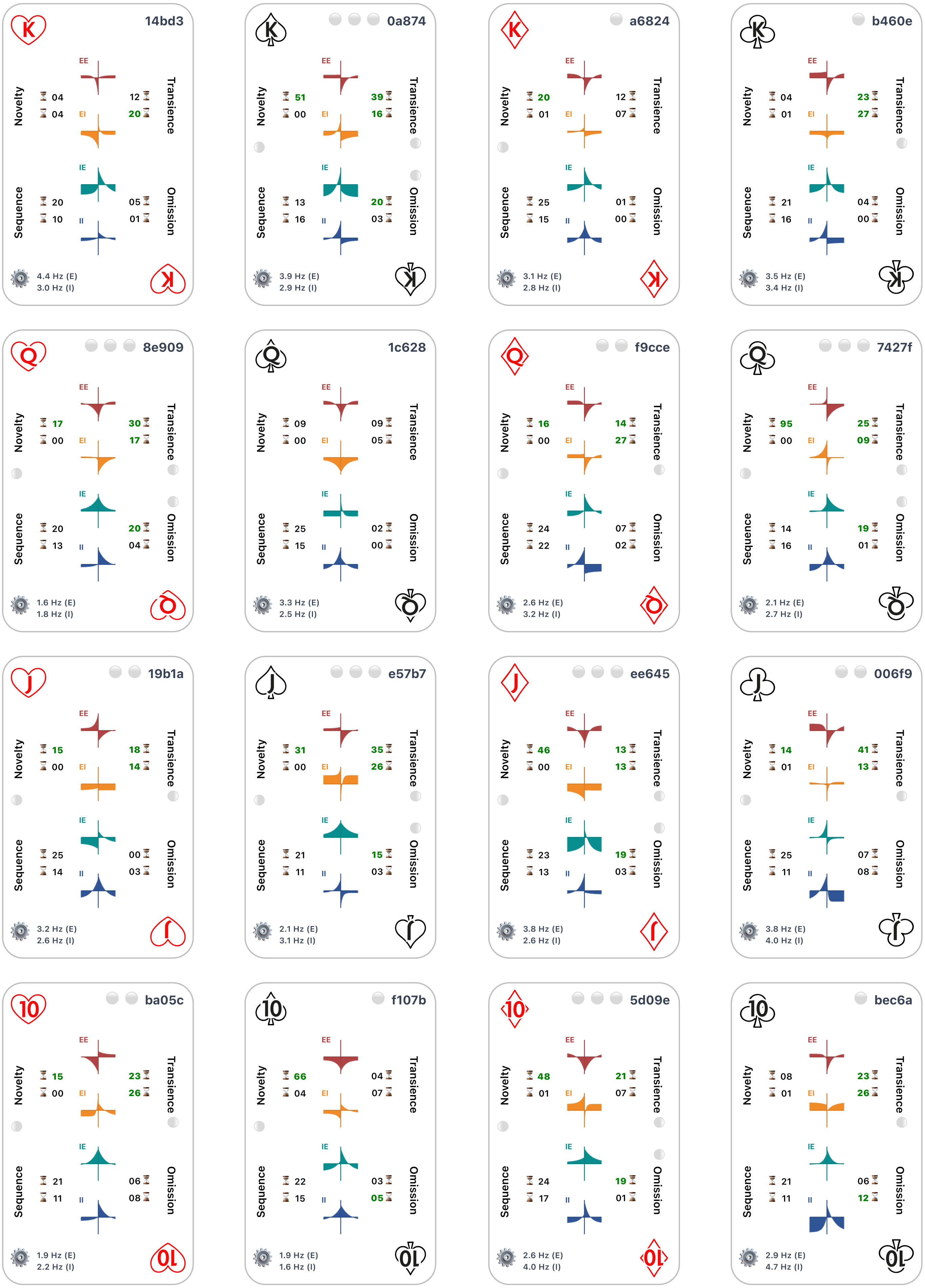
K 14bd3 — Novelty 04 04 — Transience 12 20 — Sequence 20 10 — Omission 05 01 — EE EI IE II — 4.4 Hz (E) 3.0 Hz (I)
K 0a874 — Novelty 51 00 — Transience 39 16 — Sequence 13 16 — Omission 20 03 — EE EI IE II — 3.9 Hz (E) 2.9 Hz (I)
K a6824 — Novelty 20 01 — Transience 12 07 — Sequence 25 15 — Omission 01 00 — EE EI IE II — 3.1 Hz (E) 2.8 Hz (I)
K b460e — Novelty 04 01 — Transience 23 27 — Sequence 21 16 — Omission 04 00 — EE EI IE II — 3.5 Hz (E) 3.4 Hz (I)
Q 8e909 — Novelty 17 00 — Transience 30 17 — Sequence 20 13 — Omission 20 04 — EE EI IE II — 1.6 Hz (E) 1.8 Hz (I)
Q 1c628 — Novelty 09 00 — Transience 09 05 — Sequence 25 15 — Omission 02 00 — EE EI IE II — 3.3 Hz (E) 2.5 Hz (I)
Q f9cce — Novelty 16 00 — Transience 14 27 — Sequence 24 22 — Omission 07 02 — EE EI IE II — 2.6 Hz (E) 3.2 Hz (I)
Q 7427f — Novelty 95 00 — Transience 25 09 — Sequence 14 16 — Omission 19 01 — EE EI IE II — 2.1 Hz (E) 2.7 Hz (I)
J 19b1a — Novelty 15 00 — Transience 18 14 — Sequence 25 14 — Omission 00 03 — EE EI IE II — 3.2 Hz (E) 2.6 Hz (I)
J e57b7 — Novelty 31 00 — Transience 35 26 — Sequence 21 11 — Omission 15 03 — EE EI IE II — 2.1 Hz (E) 3.1 Hz (I)
J ee645 — Novelty 46 00 — Transience 13 13 — Sequence 23 13 — Omission 19 03 — EE EI IE II — 3.8 Hz (E) 2.6 Hz (I)
J 006f9 — Novelty 14 01 — Transience 41 13 — Sequence 25 11 — Omission 07 08 — EE EI IE II — 3.8 Hz (E) 4.0 Hz (I)
10 ba05c — Novelty 15 00 — Transience 23 26 — Sequence 21 11 — Omission 06 08 — EE EI IE II — 1.9 Hz (E) 2.2 Hz (I)
10 f107b — Novelty 66 04 — Transience 04 07 — Sequence 22 15 — Omission 03 05 — EE EI IE II — 1.9 Hz (E) 1.6 Hz (I)
10 5d09e — Novelty 48 01 — Transience 21 07 — Sequence 24 17 — Omission 19 01 — EE EI IE II — 2.6 Hz (E) 4.0 Hz (I)
10 bec6a — Novelty 08 01 — Transience 23 26 — Sequence 21 11 — Omission 06 12 — EE EI IE II — 2.9 Hz (E) 4.7 Hz (I)

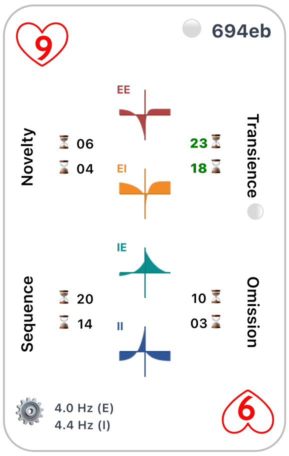
9
694eb
EE
Novelty
06
04
23
18
Transience
EI
IE
Sequence
20
14
10
03
Omission
II
4.0 Hz (E)
4.4 Hz (I)
6

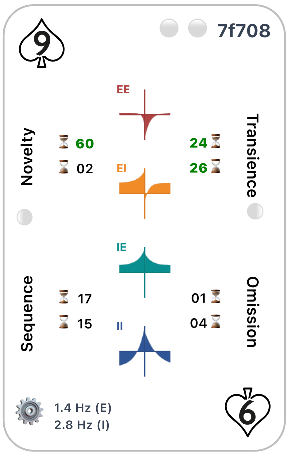
9
7f708
EE
Novelty
60
02
24
26
Transience
EI
IE
Sequence
17
15
01
04
Omission
II
1.4 Hz (E)
2.8 Hz (I)
6

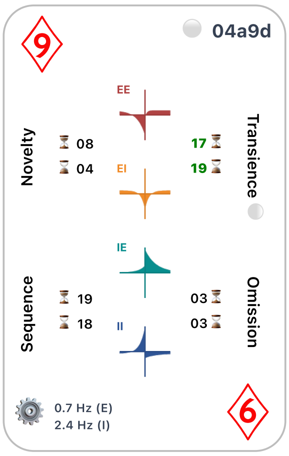
9
04a9d
EE
Novelty
08
04
17
19
Transience
EI
IE
Sequence
19
18
03
03
Omission
II
0.7 Hz (E)
2.4 Hz (I)
6

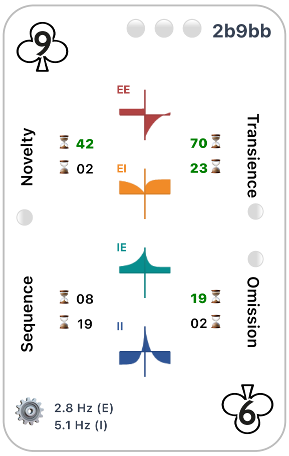
9
2b9bb
EE
Novelty
42
02
70
23
Transience
EI
IE
Sequence
08
19
19
02
Omission
II
2.8 Hz (E)
5.1 Hz (I)
6

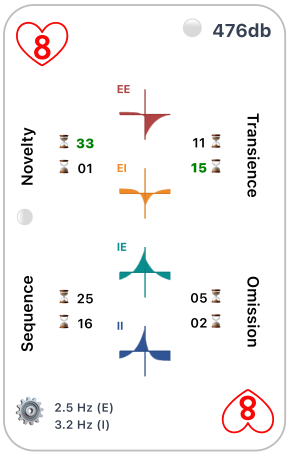
8
476db
EE
Novelty
33
01
11
15
Transience
EI
IE
Sequence
25
16
05
02
Omission
II
2.5 Hz (E)
3.2 Hz (I)
8

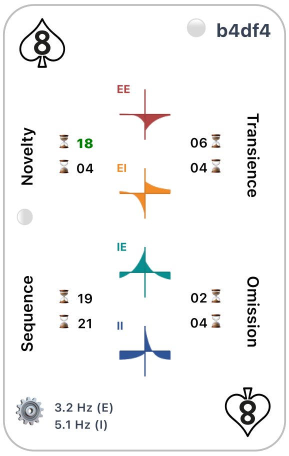
8
b4df4
EE
Novelty
18
04
06
04
Transience
EI
IE
Sequence
19
21
02
04
Omission
II
3.2 Hz (E)
5.1 Hz (I)
8

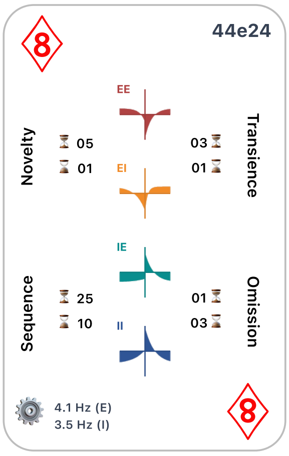
8
44e24
EE
Novelty
05
01
03
01
Transience
EI
IE
Sequence
25
10
01
03
Omission
II
4.1 Hz (E)
3.5 Hz (I)
8

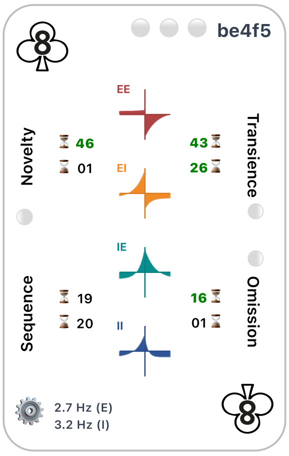
8
be4f5
EE
Novelty
46
01
43
26
Transience
EI
IE
Sequence
19
20
16
01
Omission
II
2.7 Hz (E)
3.2 Hz (I)
8

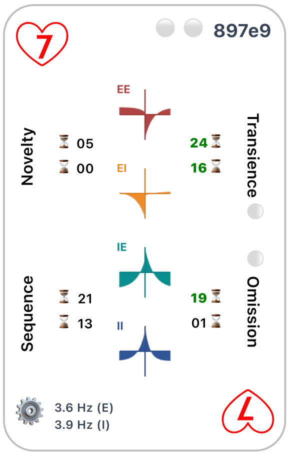
7
897e9
EE
Novelty
05
00
24
16
Transience
EI
IE
Sequence
21
13
19
01
Omission
II
3.6 Hz (E)
3.9 Hz (I)
7

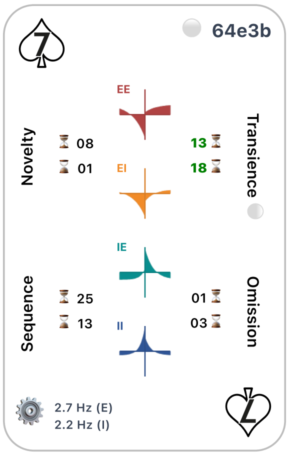
7
64e3b
EE
Novelty
08
01
13
18
Transience
EI
IE
Sequence
25
13
01
03
Omission
II
2.7 Hz (E)
2.2 Hz (I)
7

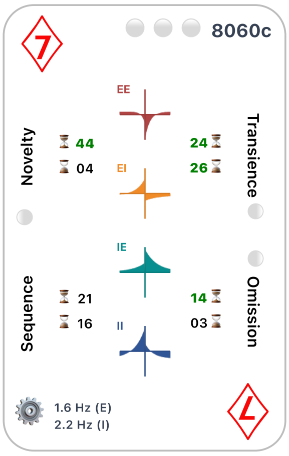
7
8060c
EE
Novelty
44
04
24
26
Transience
EI
IE
Sequence
21
16
14
03
Omission
II
1.6 Hz (E)
2.2 Hz (I)
7

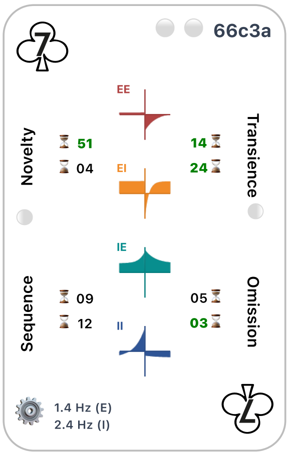
7
66c3a
EE
Novelty
51
04
14
24
Transience
EI
IE
Sequence
09
12
05
03
Omission
II
1.4 Hz (E)
2.4 Hz (I)
7

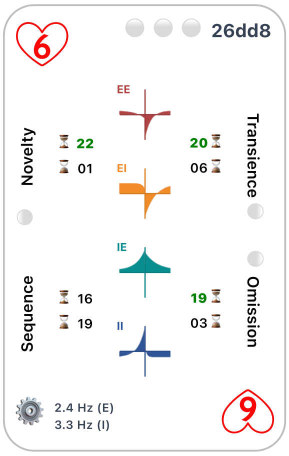
6
26dd8
EE
Novelty
22
01
20
06
Transience
EI
IE
Sequence
16
19
19
03
Omission
II
2.4 Hz (E)
3.3 Hz (I)
9

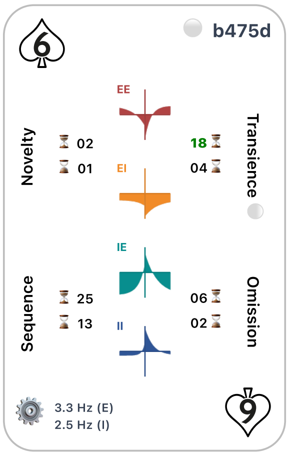
6
b475d
EE
Novelty
02
01
18
04
Transience
EI
IE
Sequence
25
13
06
02
Omission
II
3.3 Hz (E)
2.5 Hz (I)
9

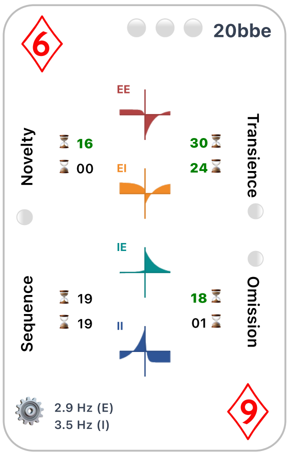
6
20bbe
EE
Novelty
16
00
30
24
Transience
EI
IE
Sequence
19
19
18
01
Omission
II
2.9 Hz (E)
3.5 Hz (I)
9

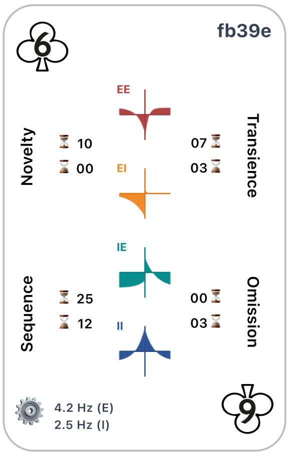
6
fb39e
EE
Novelty
10
00
07
03
Transience
EI
IE
Sequence
25
12
00
03
Omission
II
4.2 Hz (E)
2.5 Hz (I)
9

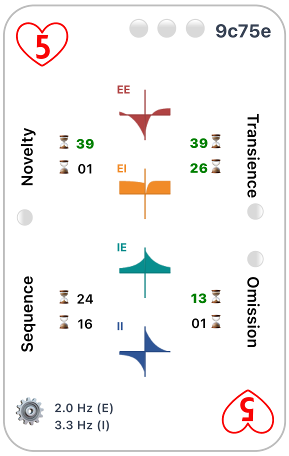
5
9c75e
EE
EI
IE
II
Novelty
39
01
Transience
39
26
Sequence
24
16
Omission
13
01
2.0 Hz (E)
3.3 Hz (I)

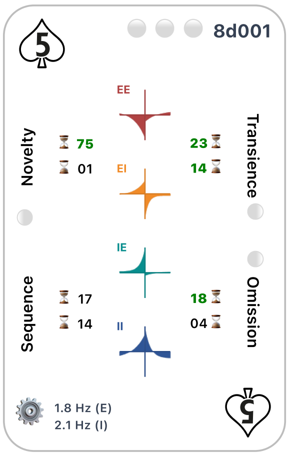
5
8d001
EE
EI
IE
II
Novelty
75
01
Transience
23
14
Sequence
17
14
Omission
18
04
1.8 Hz (E)
2.1 Hz (I)

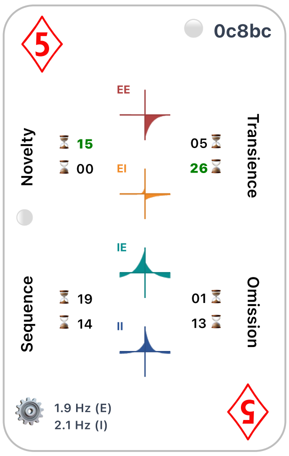
5
0c8bc
EE
EI
IE
II
Novelty
15
00
Transience
05
26
Sequence
19
14
Omission
01
13
1.9 Hz (E)
2.1 Hz (I)

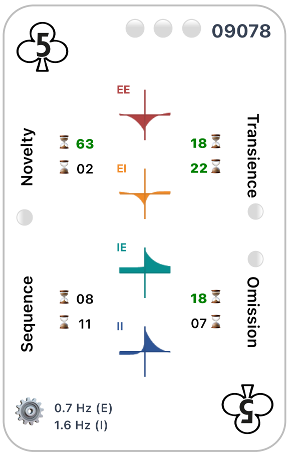
5
09078
EE
EI
IE
II
Novelty
63
02
Transience
18
22
Sequence
08
11
Omission
18
07
0.7 Hz (E)
1.6 Hz (I)

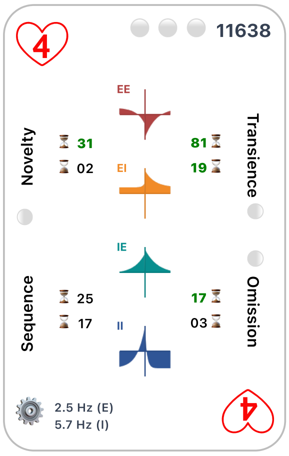
4
11638
EE
EI
IE
II
Novelty
31
02
Transience
81
19
Sequence
25
17
Omission
17
03
2.5 Hz (E)
5.7 Hz (I)

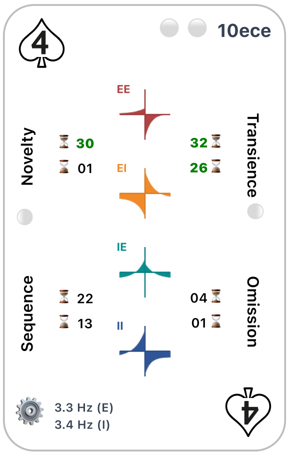
4
10ece
EE
EI
IE
II
Novelty
30
01
Transience
32
26
Sequence
22
13
Omission
04
01
3.3 Hz (E)
3.4 Hz (I)

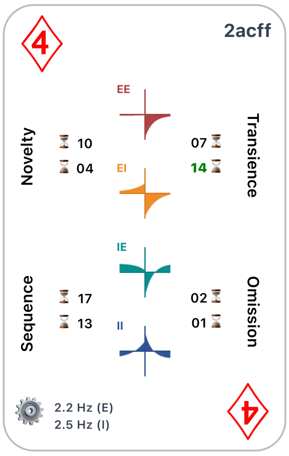
4
2acff
EE
EI
IE
II
Novelty
10
04
Transience
07
14
Sequence
17
13
Omission
02
01
2.2 Hz (E)
2.5 Hz (I)

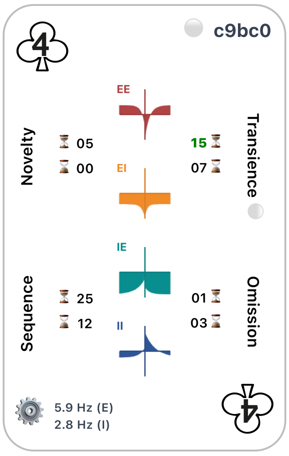
4
c9bc0
EE
EI
IE
II
Novelty
05
00
Transience
15
07
Sequence
25
12
Omission
01
03
5.9 Hz (E)
2.8 Hz (I)

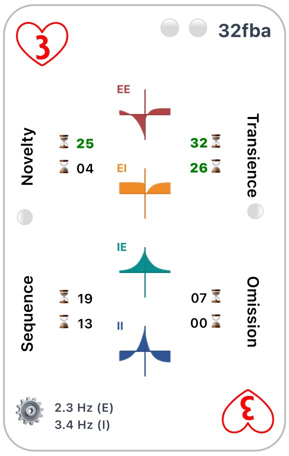
3
32fba
EE
EI
IE
II
Novelty
25
04
Transience
32
26
Sequence
19
13
Omission
07
00
2.3 Hz (E)
3.4 Hz (I)

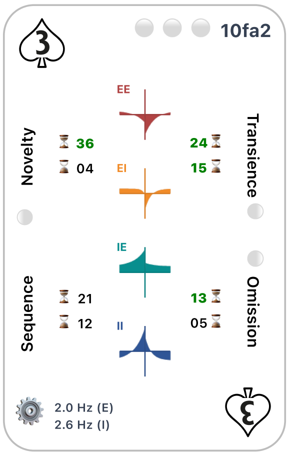
3
10fa2
EE
EI
IE
II
Novelty
36
04
Transience
24
15
Sequence
21
12
Omission
13
05
2.0 Hz (E)
2.6 Hz (I)

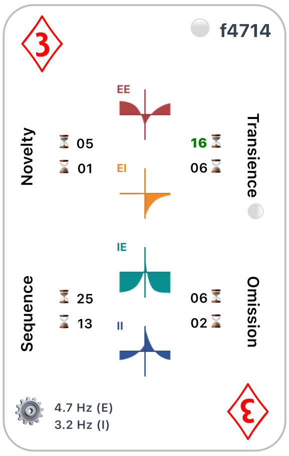
3
f4714
EE
EI
IE
II
Novelty
05
01
Transience
16
06
Sequence
25
13
Omission
06
02
4.7 Hz (E)
3.2 Hz (I)

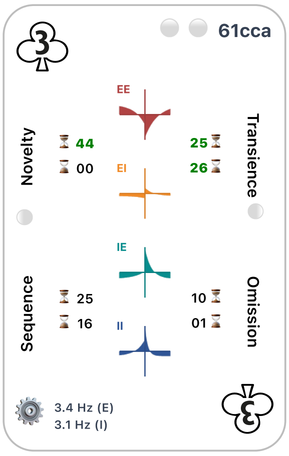
3
61cca
EE
EI
IE
II
Novelty
44
00
Transience
25
26
Sequence
25
16
Omission
10
01
3.4 Hz (E)
3.1 Hz (I)

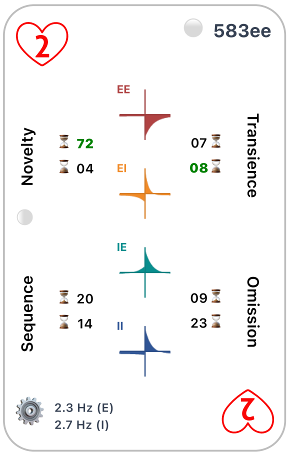
2
583ee
EE
EI
IE
II
Novelty
72
04
Transience
07
08
Sequence
20
14
Omission
09
23
2.3 Hz (E)
2.7 Hz (I)

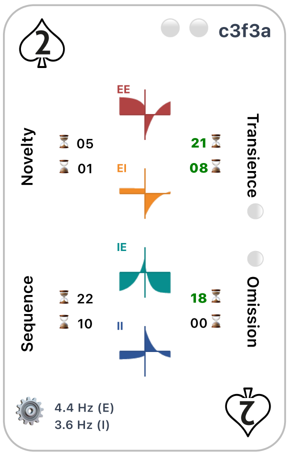
2
c3f3a
EE
EI
IE
II
Novelty
05
01
Transience
21
08
Sequence
22
10
Omission
18
00
4.4 Hz (E)
3.6 Hz (I)

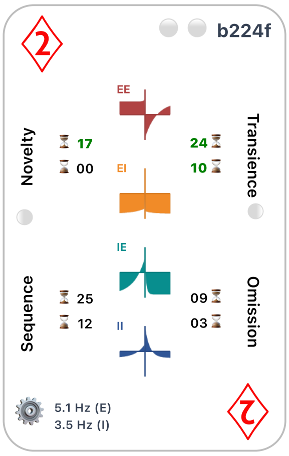
2
b224f
EE
EI
IE
II
Novelty
17
00
Transience
24
10
Sequence
25
12
Omission
09
03
5.1 Hz (E)
3.5 Hz (I)

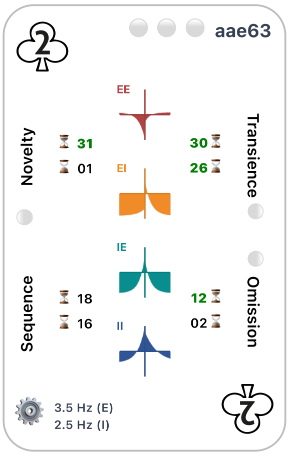
2
aae63
EE
EI
IE
II
Novelty
31
01
Transience
30
26
Sequence
18
16
Omission
12
02
3.5 Hz (E)
2.5 Hz (I)

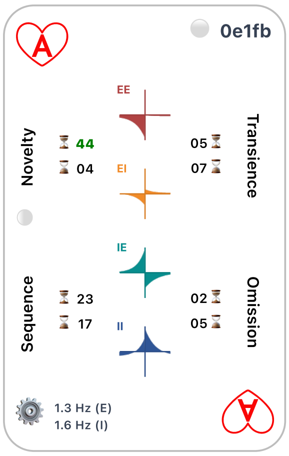
0e1fb
Novelty
44
04
EE
EI
05
07
Transience
Sequence
23
17
IE
II
02
05
Omission
1.3 Hz (E)
1.6 Hz (I)

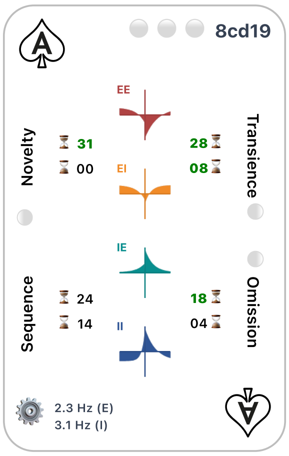
8cd19
Novelty
31
00
EE
EI
28
08
Transience
Sequence
24
14
IE
II
18
04
Omission
2.3 Hz (E)
3.1 Hz (I)

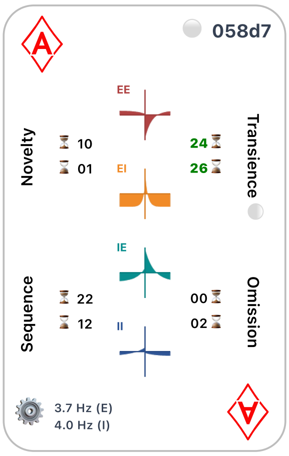
058d7
Novelty
10
01
EE
EI
24
26
Transience
Sequence
22
12
IE
II
00
02
Omission
3.7 Hz (E)
4.0 Hz (I)

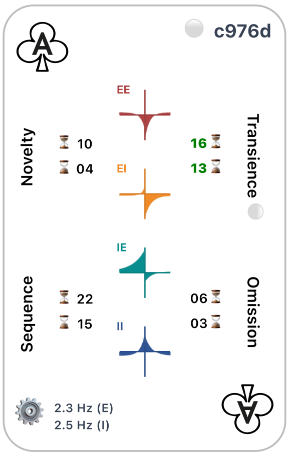
c976d
Novelty
10
04
EE
EI
16
13
Transience
Sequence
22
15
IE
II
06
03
Omission
2.3 Hz (E)
2.5 Hz (I)

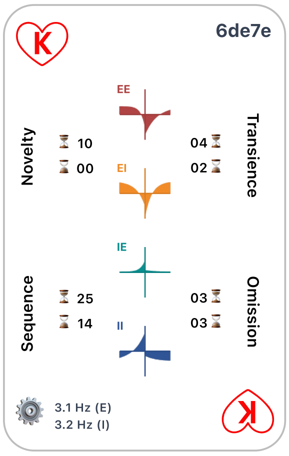
6de7e
Novelty
10
00
EE
EI
04
02
Transience
Sequence
25
14
IE
II
03
03
Omission
3.1 Hz (E)
3.2 Hz (I)

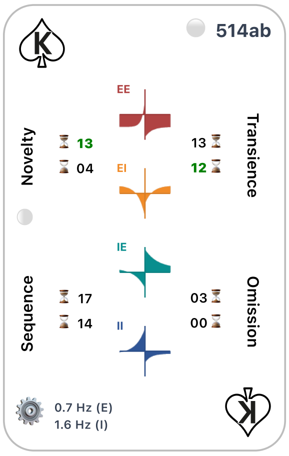
514ab
Novelty
13
04
EE
EI
13
12
Transience
Sequence
17
14
IE
II
03
00
Omission
0.7 Hz (E)
1.6 Hz (I)

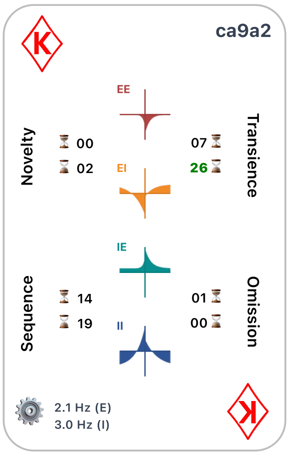
ca9a2
Novelty
00
02
EE
EI
07
26
Transience
Sequence
14
19
IE
II
01
00
Omission
2.1 Hz (E)
3.0 Hz (I)

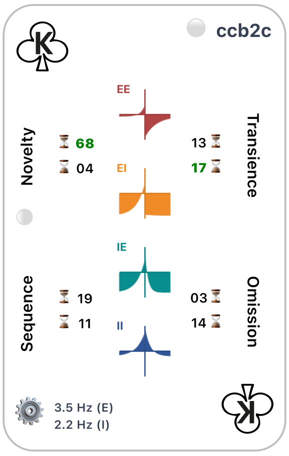
ccb2c
Novelty
68
04
EE
EI
13
17
Transience
Sequence
19
11
IE
II
03
14
Omission
3.5 Hz (E)
2.2 Hz (I)

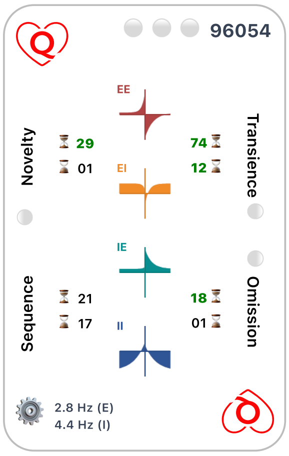
96054
Novelty
29
01
EE
EI
74
12
Transience
Sequence
21
17
IE
II
18
01
Omission
2.8 Hz (E)
4.4 Hz (I)

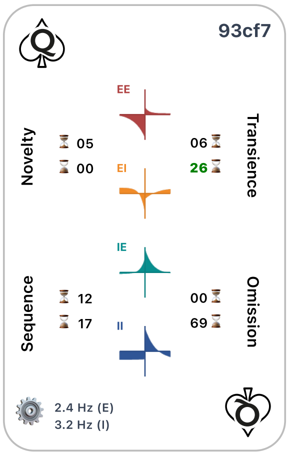
93cf7
Novelty
05
00
EE
EI
06
26
Transience
Sequence
12
17
IE
II
00
69
Omission
2.4 Hz (E)
3.2 Hz (I)

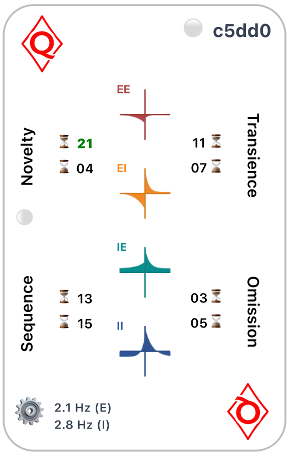
c5dd0
Novelty
21
04
EE
EI
11
07
Transience
Sequence
13
15
IE
II
03
05
Omission
2.1 Hz (E)
2.8 Hz (I)

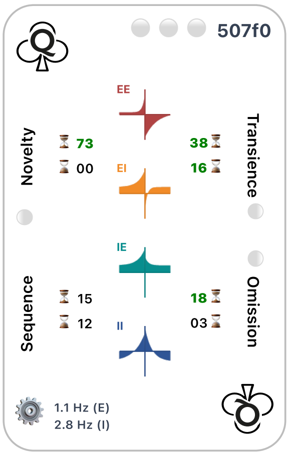
507f0
Novelty
73
00
EE
EI
38
16
Transience
Sequence
15
12
IE
II
18
03
Omission
1.1 Hz (E)
2.8 Hz (I)

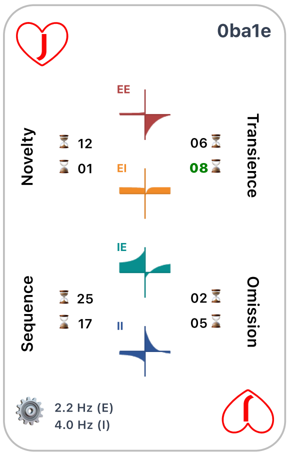
0ba1e
Novelty
12
01
EE
EI
06
08
Transience
Sequence
25
17
IE
II
02
05
Omission
2.2 Hz (E)
4.0 Hz (I)

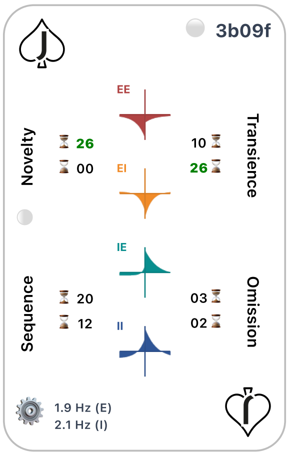
3b09f
Novelty
26
00
EE
EI
10
26
Transience
Sequence
20
12
IE
II
03
02
Omission
1.9 Hz (E)
2.1 Hz (I)

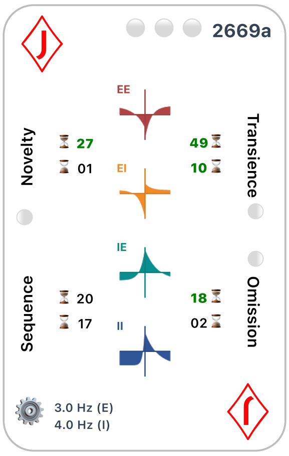
2669a
Novelty
27
01
EE
EI
49
10
Transience
Sequence
20
17
IE
II
18
02
Omission
3.0 Hz (E)
4.0 Hz (I)

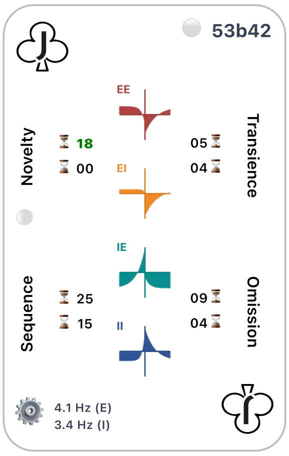
53b42
Novelty
18
00
EE
EI
05
04
Transience
Sequence
25
15
IE
II
09
04
Omission
4.1 Hz (E)
3.4 Hz (I)

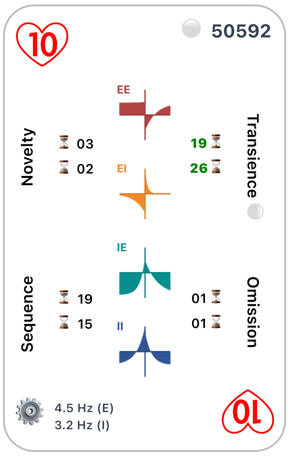
10
50592
EE
EI
IE
II
Novelty 03 02
Transience 19 26
Sequence 19 15
Omission 01 01
4.5 Hz (E)
3.2 Hz (I)
10

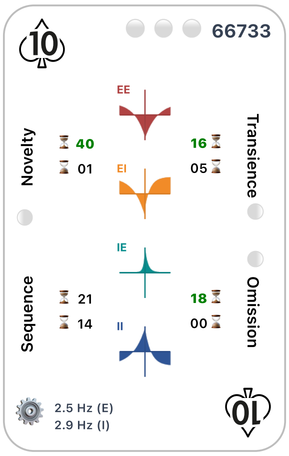
10
66733
EE
EI
IE
II
Novelty 40 01
Transience 16 05
Sequence 21 14
Omission 18 00
2.5 Hz (E)
2.9 Hz (I)
10

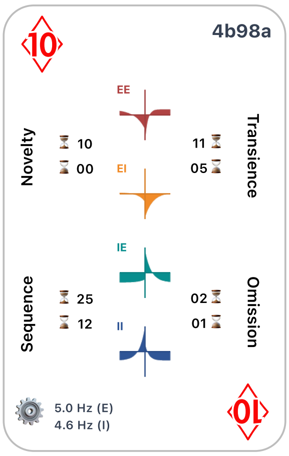
10
4b98a
EE
EI
IE
II
Novelty 10 00
Transience 11 05
Sequence 25 12
Omission 02 01
5.0 Hz (E)
4.6 Hz (I)
10

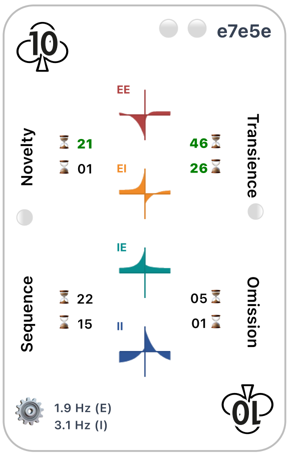
10
e7e5e
EE
EI
IE
II
Novelty 21 01
Transience 46 26
Sequence 22 15
Omission 05 01
1.9 Hz (E)
3.1 Hz (I)
10

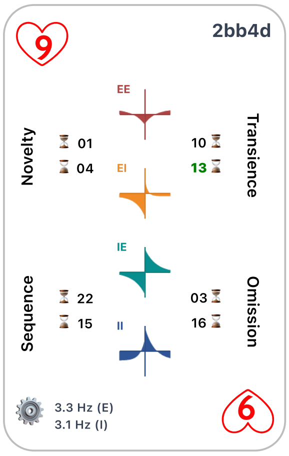
9
2bb4d
EE
EI
IE
II
Novelty 01 04
Transience 10 13
Sequence 22 15
Omission 03 16
3.3 Hz (E)
3.1 Hz (I)
9

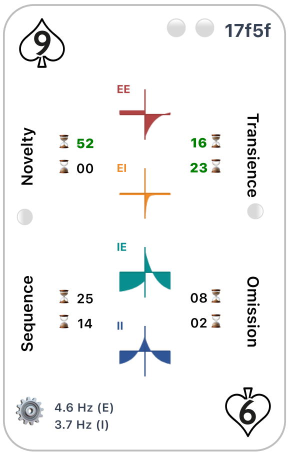
9
17f5f
EE
EI
IE
II
Novelty 52 00
Transience 16 23
Sequence 25 14
Omission 08 02
4.6 Hz (E)
3.7 Hz (I)
9

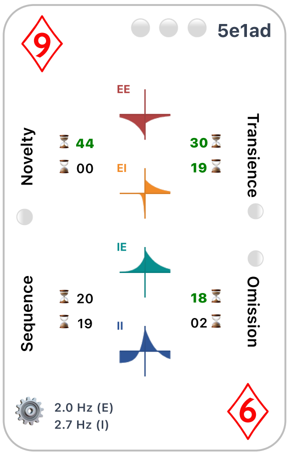
9
5e1ad
EE
EI
IE
II
Novelty 44 00
Transience 30 19
Sequence 20 19
Omission 18 02
2.0 Hz (E)
2.7 Hz (I)
9

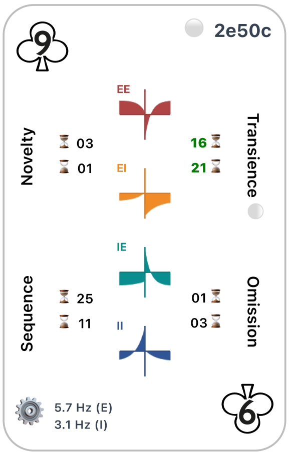
9
2e50c
EE
EI
IE
II
Novelty 03 01
Transience 16 21
Sequence 25 11
Omission 01 03
5.7 Hz (E)
3.1 Hz (I)
9

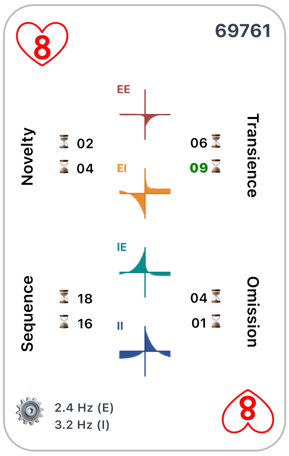
8
69761
EE
EI
IE
II
Novelty 02 04
Transience 06 09
Sequence 18 16
Omission 04 01
2.4 Hz (E)
3.2 Hz (I)
8

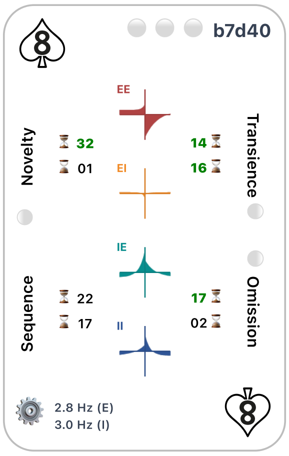
8
b7d40
EE
EI
IE
II
Novelty 32 01
Transience 14 16
Sequence 22 17
Omission 17 02
2.8 Hz (E)
3.0 Hz (I)
8

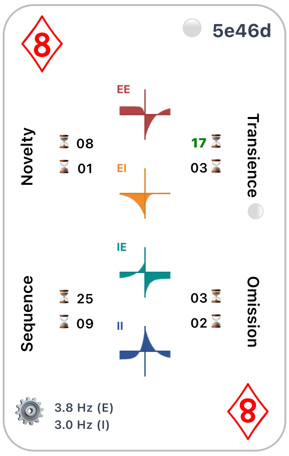
8
5e46d
EE
EI
IE
II
Novelty 08 01
Transience 17 03
Sequence 25 09
Omission 03 02
3.8 Hz (E)
3.0 Hz (I)
8

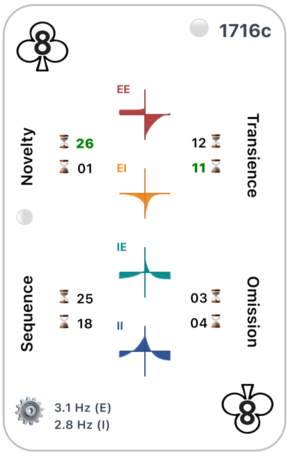
8
1716c
EE
EI
IE
II
Novelty 26 01
Transience 12 11
Sequence 25 18
Omission 03 04
3.1 Hz (E)
2.8 Hz (I)
8

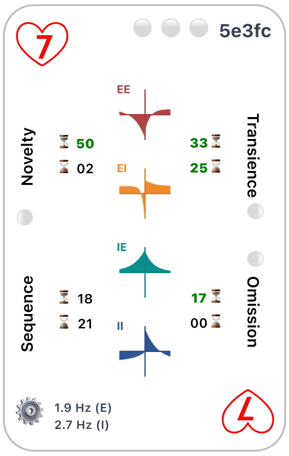
7
5e3fc
EE
EI
IE
II
Novelty 50 02
Transience 33 25
Sequence 18 21
Omission 17 00
1.9 Hz (E)
2.7 Hz (I)
7

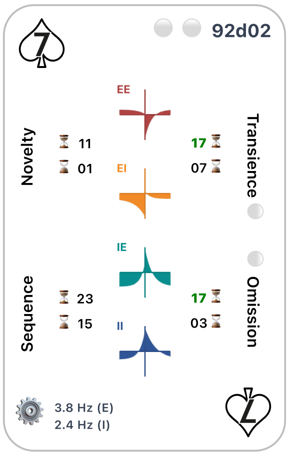
7
92d02
EE
EI
IE
II
Novelty 11 01
Transience 17 07
Sequence 23 15
Omission 17 03
3.8 Hz (E)
2.4 Hz (I)
7

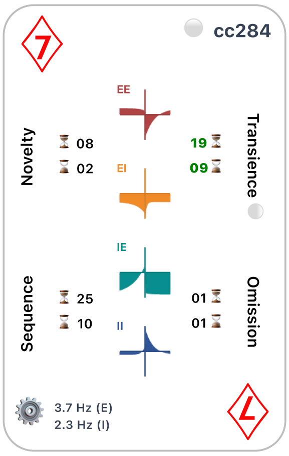
7
cc284
EE
EI
IE
II
Novelty 08 02
Transience 19 09
Sequence 25 10
Omission 01 01
3.7 Hz (E)
2.3 Hz (I)
7

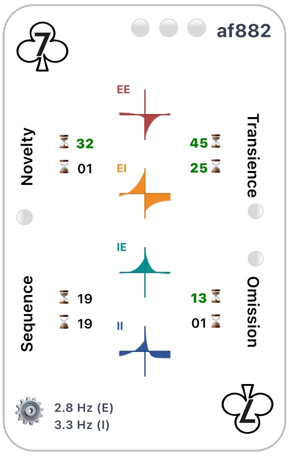
7
af882
EE
EI
IE
II
Novelty 32 01
Transience 45 25
Sequence 19 19
Omission 13 01
2.8 Hz (E)
3.3 Hz (I)
7

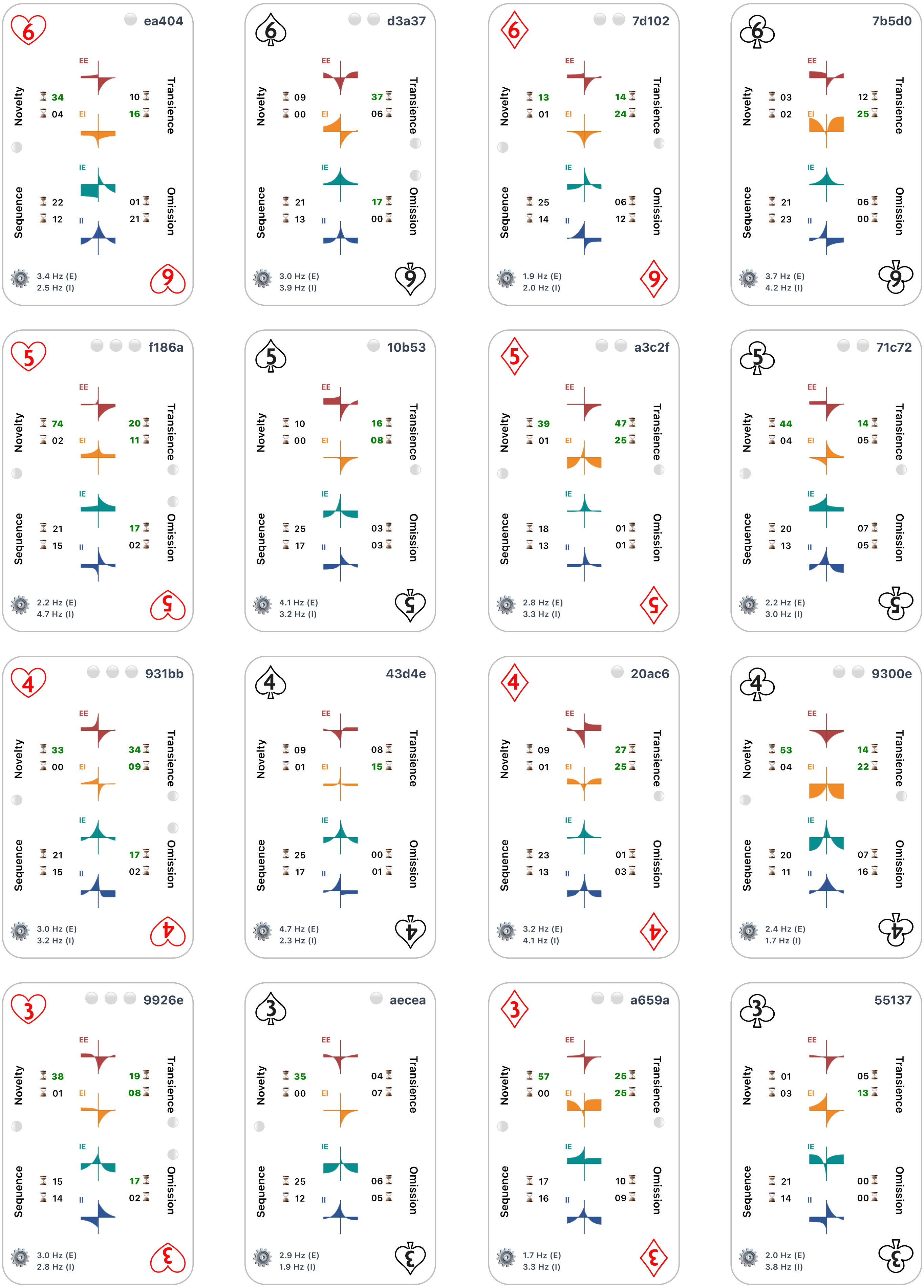
6 ea404
EE EI IE II
Novelty 34 04
Transience 10 16
Sequence 22 12
Omission 01 21
3.4 Hz (E)
2.5 Hz (I)
6 d3a37
Novelty 09 00
Transience 37 06
Sequence 21 13
Omission 17 00
3.0 Hz (E)
3.9 Hz (I)
6 7d102
Novelty 13 01
Transience 14 24
Sequence 25 14
Omission 06 12
1.9 Hz (E)
2.0 Hz (I)
6 7b5d0
Novelty 03 02
Transience 12 25
Sequence 21 23
Omission 06 00
3.7 Hz (E)
4.2 Hz (I)
5 f186a
Novelty 74 02
Transience 20 11
Sequence 21 15
Omission 17 02
2.2 Hz (E)
4.7 Hz (I)
5 10b53
Novelty 10 00
Transience 16 08
Sequence 25 17
Omission 03 03
4.1 Hz (E)
3.2 Hz (I)
5 a3c2f
Novelty 39 01
Transience 47 25
Sequence 18 13
Omission 01 01
2.8 Hz (E)
3.3 Hz (I)
5 71c72
Novelty 44 04
Transience 14 05
Sequence 20 13
Omission 07 05
2.2 Hz (E)
3.0 Hz (I)
4 931bb
Novelty 33 00
Transience 34 09
Sequence 21 15
Omission 17 02
3.0 Hz (E)
3.2 Hz (I)
4 43d4e
Novelty 09 01
Transience 08 15
Sequence 25 17
Omission 00 01
4.7 Hz (E)
2.3 Hz (I)
4 20ac6
Novelty 09 01
Transience 27 25
Sequence 23 13
Omission 01 03
3.2 Hz (E)
4.1 Hz (I)
4 9300e
Novelty 53 04
Transience 14 22
Sequence 20 11
Omission 07 16
2.4 Hz (E)
1.7 Hz (I)
3 9926e
Novelty 38 01
Transience 19 08
Sequence 15 14
Omission 17 02
3.0 Hz (E)
2.8 Hz (I)
3 aecea
Novelty 35 00
Transience 04 07
Sequence 25 12
Omission 06 05
2.9 Hz (E)
1.9 Hz (I)
3 a659a
Novelty 57 00
Transience 25 25
Sequence 17 16
Omission 10 09
1.7 Hz (E)
3.3 Hz (I)
3 55137
Novelty 01 03
Transience 05 13
Sequence 21 14
Omission 00 00
2.0 Hz (E)
3.8 Hz (I)

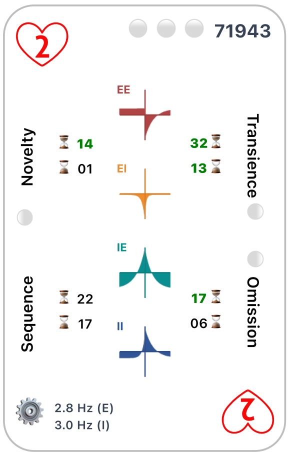
2
71943
EE
Novelty
14
01
32
13
Transience
EI
IE
Sequence
22
17
17
06
Omission
II
2.8 Hz (E)
3.0 Hz (I)

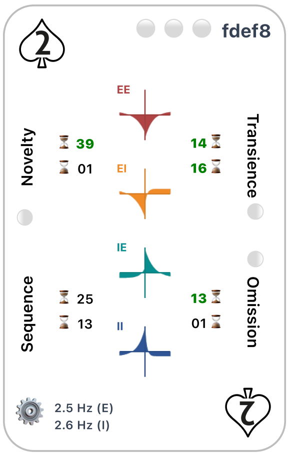
2
fdef8
EE
Novelty
39
01
14
16
Transience
EI
IE
Sequence
25
13
13
01
Omission
II
2.5 Hz (E)
2.6 Hz (I)

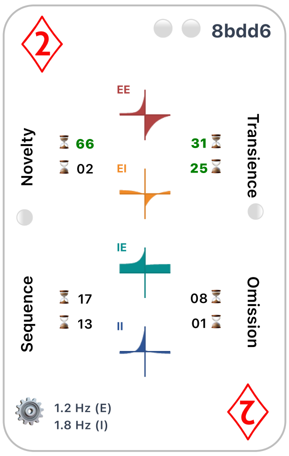
2
8bdd6
EE
Novelty
66
02
31
25
Transience
EI
IE
Sequence
17
13
08
01
Omission
II
1.2 Hz (E)
1.8 Hz (I)

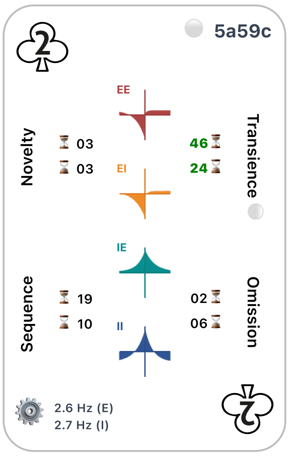
2
5a59c
EE
Novelty
03
03
46
24
Transience
EI
IE
Sequence
19
10
02
06
Omission
II
2.6 Hz (E)
2.7 Hz (I)

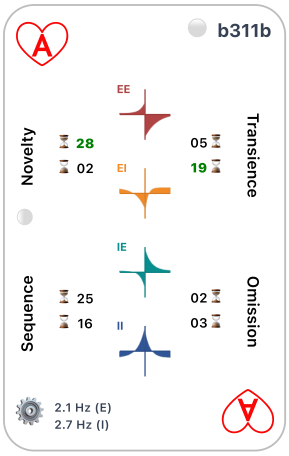
A
b311b
EE
Novelty
28
02
05
19
Transience
EI
IE
Sequence
25
16
02
03
Omission
II
2.1 Hz (E)
2.7 Hz (I)

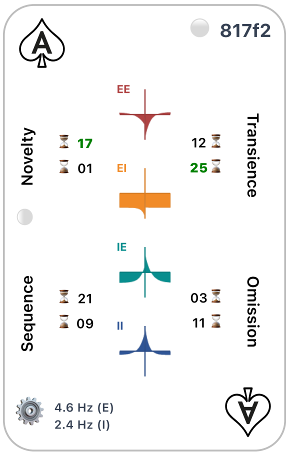
A
817f2
EE
Novelty
17
01
12
25
Transience
EI
IE
Sequence
21
09
03
11
Omission
II
4.6 Hz (E)
2.4 Hz (I)

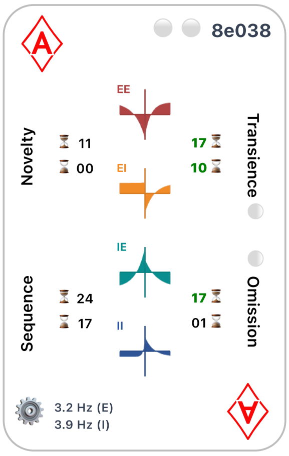
A
8e038
EE
Novelty
11
00
17
10
Transience
EI
IE
Sequence
24
17
17
01
Omission
II
3.2 Hz (E)
3.9 Hz (I)

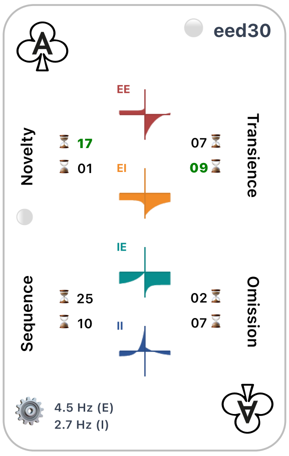
A
eed30
EE
Novelty
17
01
07
09
Transience
EI
IE
Sequence
25
10
02
07
Omission
II
4.5 Hz (E)
2.7 Hz (I)

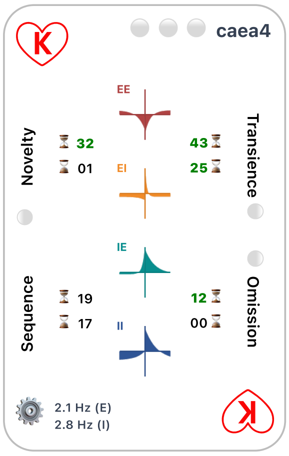
K
caea4
EE
Novelty
32
01
43
25
Transience
EI
IE
Sequence
19
17
12
00
Omission
II
2.1 Hz (E)
2.8 Hz (I)

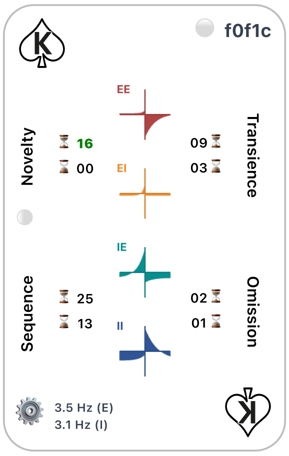
K
f0f1c
EE
Novelty
16
00
09
03
Transience
EI
IE
Sequence
25
13
02
01
Omission
II
3.5 Hz (E)
3.1 Hz (I)

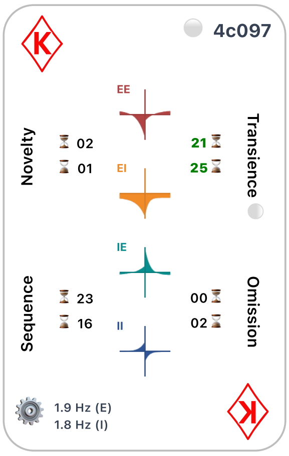
K
4c097
EE
Novelty
02
01
21
25
Transience
EI
IE
Sequence
23
16
00
02
Omission
II
1.9 Hz (E)
1.8 Hz (I)

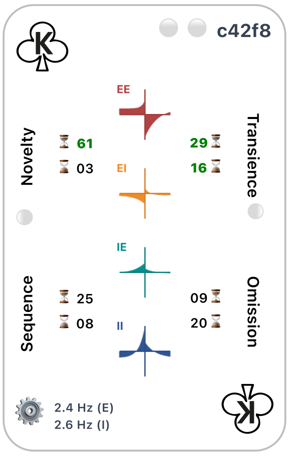
K
c42f8
EE
Novelty
61
03
29
16
Transience
EI
IE
Sequence
25
08
09
20
Omission
II
2.4 Hz (E)
2.6 Hz (I)

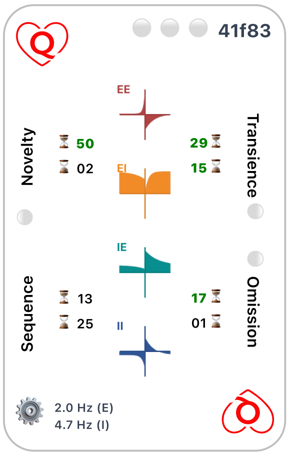
Q
41f83
EE
Novelty
50
02
29
15
Transience
EI
IE
Sequence
13
25
17
01
Omission
II
2.0 Hz (E)
4.7 Hz (I)

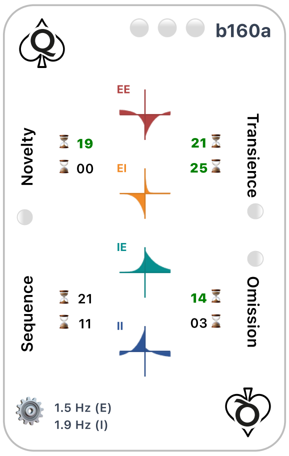
Q
b160a
EE
Novelty
19
00
21
25
Transience
EI
IE
Sequence
21
11
14
03
Omission
II
1.5 Hz (E)
1.9 Hz (I)

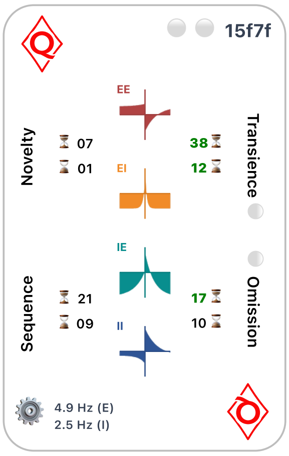
Q
15f7f
EE
Novelty
07
01
38
12
Transience
EI
IE
Sequence
21
09
17
10
Omission
II
4.9 Hz (E)
2.5 Hz (I)

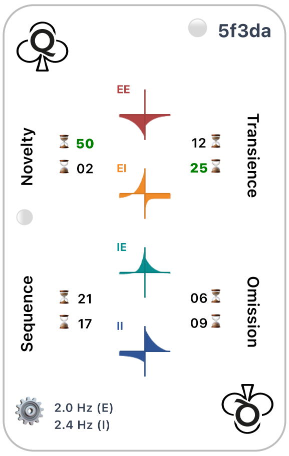
Q
5f3da
EE
Novelty
50
02
12
25
Transience
EI
IE
Sequence
21
17
06
09
Omission
II
2.0 Hz (E)
2.4 Hz (I)

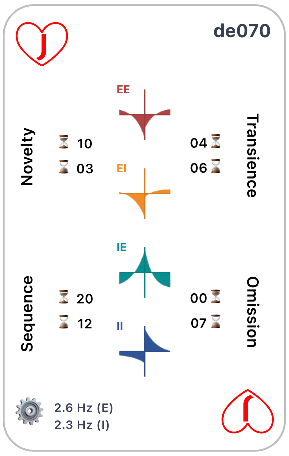
J
de070
EE
Novelty
10
03
04
06
Transience
EI
IE
Sequence
20
12
00
07
Omission
II
2.6 Hz (E)
2.3 Hz (I)

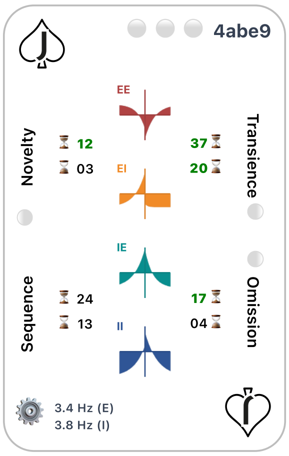
J
4abe9
EE
Novelty
12
03
37
20
Transience
EI
IE
Sequence
24
13
17
04
Omission
II
3.4 Hz (E)
3.8 Hz (I)

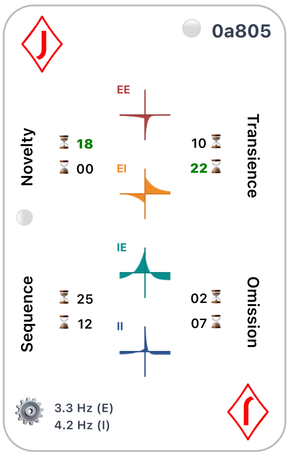
J
0a805
EE
Novelty
18
00
10
22
Transience
EI
IE
Sequence
25
12
02
07
Omission
II
3.3 Hz (E)
4.2 Hz (I)

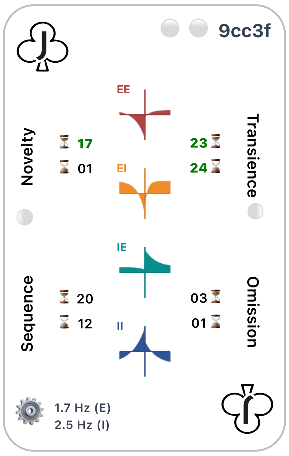
J
9cc3f
EE
Novelty
17
01
23
24
Transience
EI
IE
Sequence
20
12
03
01
Omission
II
1.7 Hz (E)
2.5 Hz (I)

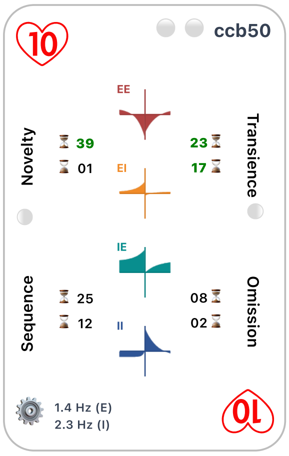
10
ccb50
EE
Novelty
39
01
23
17
Transience
EI
IE
Sequence
25
12
08
02
Omission
II
1.4 Hz (E)
2.3 Hz (I)

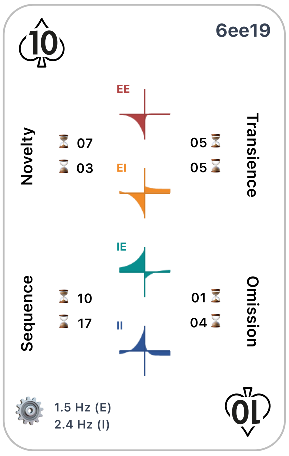
10
6ee19
EE
Novelty
07
03
05
05
Transience
EI
IE
Sequence
10
17
01
04
Omission
II
1.5 Hz (E)
2.4 Hz (I)

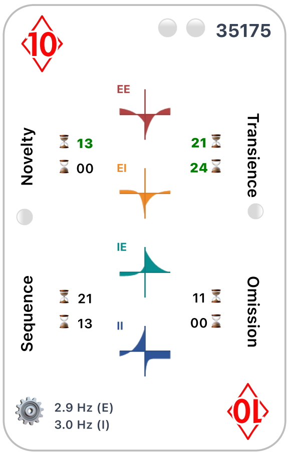
10
35175
EE
Novelty
13
00
21
24
Transience
EI
IE
Sequence
21
13
11
00
Omission
II
2.9 Hz (E)
3.0 Hz (I)

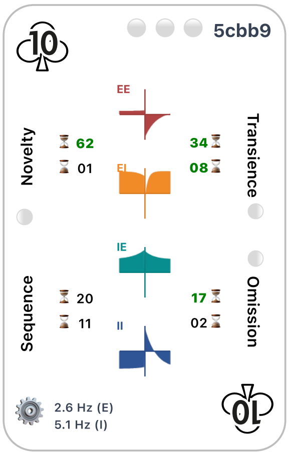
10
5cbb9
EE
Novelty
62
01
34
08
Transience
EI
IE
Sequence
20
11
17
02
Omission
II
2.6 Hz (E)
5.1 Hz (I)

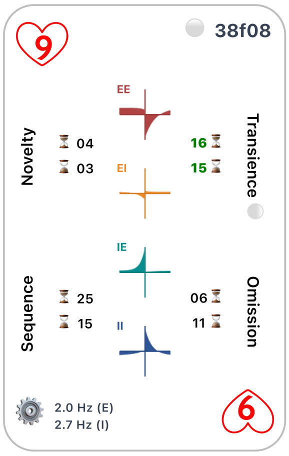
9
38f08
EE
Novelty
04
03
16
15
Transience
EI
IE
Sequence
25
15
06
11
Omission
II
2.0 Hz (E)
2.7 Hz (I)

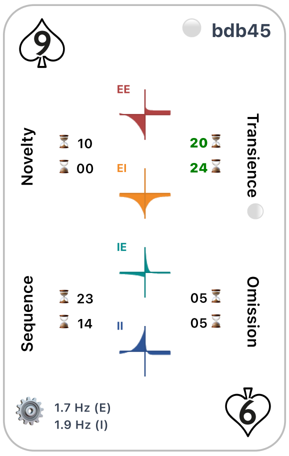
9
bdb45
EE
Novelty
10
00
20
24
Transience
EI
IE
Sequence
23
14
05
05
Omission
II
1.7 Hz (E)
1.9 Hz (I)

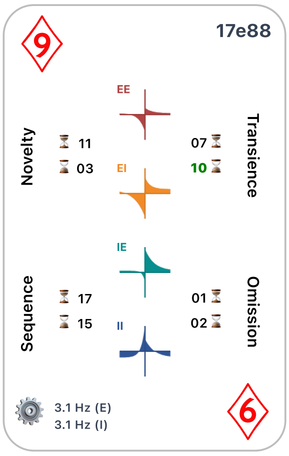
9
17e88
EE
Novelty
11
03
07
10
Transience
EI
IE
Sequence
17
15
01
02
Omission
II
3.1 Hz (E)
3.1 Hz (I)

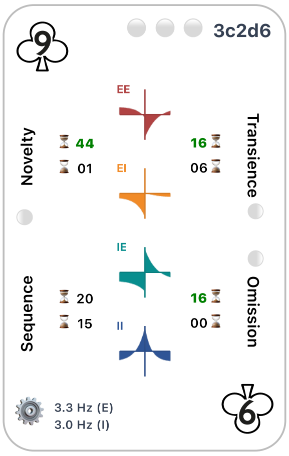
9
3c2d6
EE
Novelty
44
01
16
06
Transience
EI
IE
Sequence
20
15
16
00
Omission
II
3.3 Hz (E)
3.0 Hz (I)

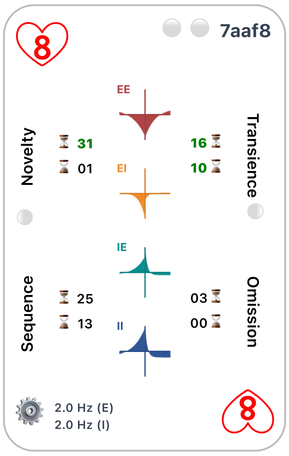
8
7aaf8
EE
Novelty
31
01
16
10
Transience
EI
IE
Sequence
25
13
03
00
Omission
II
2.0 Hz (E)
2.0 Hz (I)

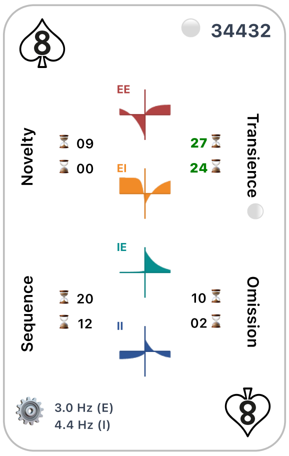
8
34432
EE
Novelty
09
00
27
24
Transience
EI
IE
Sequence
20
12
10
02
Omission
II
3.0 Hz (E)
4.4 Hz (I)

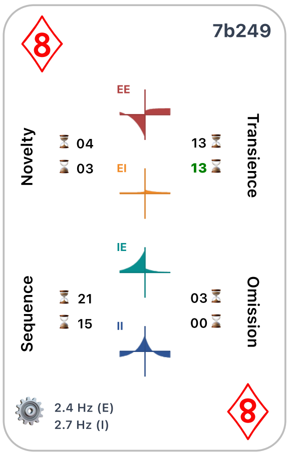
8
7b249
EE
Novelty
04
03
13
13
Transience
EI
IE
Sequence
21
15
03
00
Omission
II
2.4 Hz (E)
2.7 Hz (I)

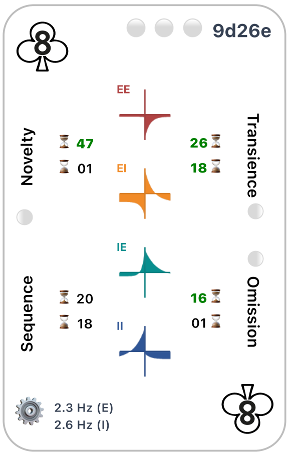
8
9d26e
EE
Novelty
47
01
26
18
Transience
EI
IE
Sequence
20
18
16
01
Omission
II
2.3 Hz (E)
2.6 Hz (I)

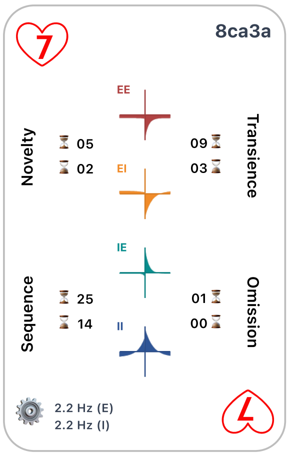
7
8ca3a
EE
Novelty
05
02
09
03
Transience
EI
IE
Sequence
25
14
01
00
Omission
II
2.2 Hz (E)
2.2 Hz (I)

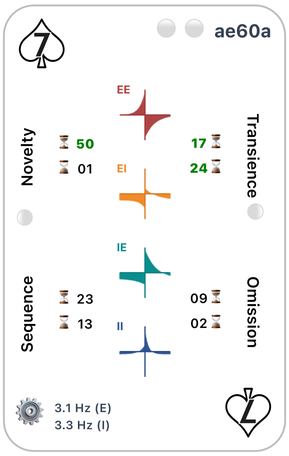
7
ae60a
EE
Novelty
50
01
17
24
Transience
EI
IE
Sequence
23
13
09
02
Omission
II
3.1 Hz (E)
3.3 Hz (I)

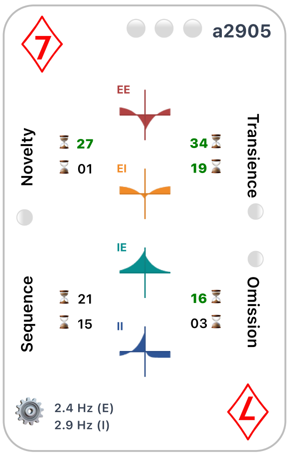
7
a2905
EE
Novelty
27
01
34
19
Transience
EI
IE
Sequence
21
15
16
03
Omission
II
2.4 Hz (E)
2.9 Hz (I)

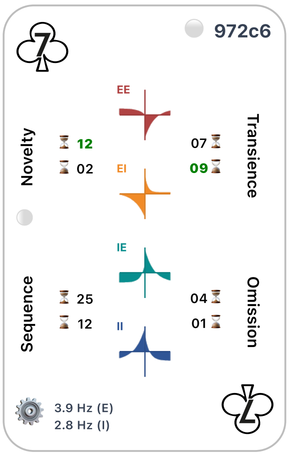
7
972c6
EE
Novelty
12
02
07
09
Transience
EI
IE
Sequence
25
12
04
01
Omission
II
3.9 Hz (E)
2.8 Hz (I)

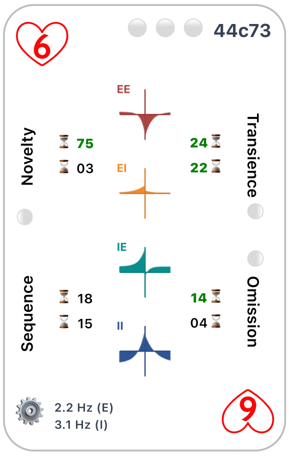
6
44c73
EE
Novelty
75
03
24
22
Transience
EI
IE
Sequence
18
15
14
04
Omission
II
2.2 Hz (E)
3.1 Hz (I)

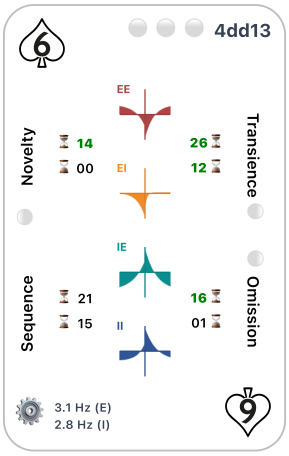
6
4dd13
EE
Novelty
14
00
26
12
Transience
EI
IE
Sequence
21
15
16
01
Omission
II
3.1 Hz (E)
2.8 Hz (I)

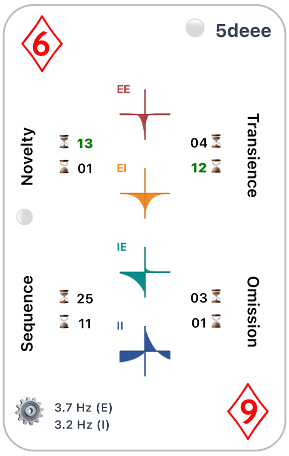
6
5deee
EE
Novelty
13
01
04
12
Transience
EI
IE
Sequence
25
11
03
01
Omission
II
3.7 Hz (E)
3.2 Hz (I)

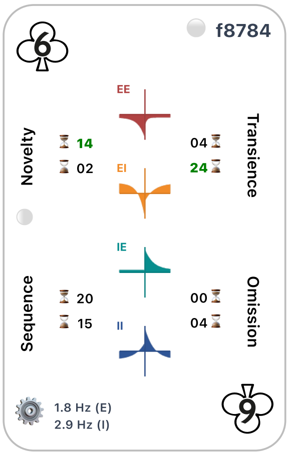
6
f8784
EE
Novelty
14
02
04
24
Transience
EI
IE
Sequence
20
15
00
04
Omission
II
1.8 Hz (E)
2.9 Hz (I)

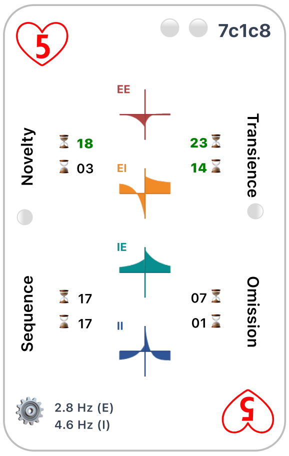
5
7c1c8
EE
Novelty
18
03
23
14
Transience
EI
IE
Sequence
17
17
07
01
Omission
II
2.8 Hz (E)
4.6 Hz (I)

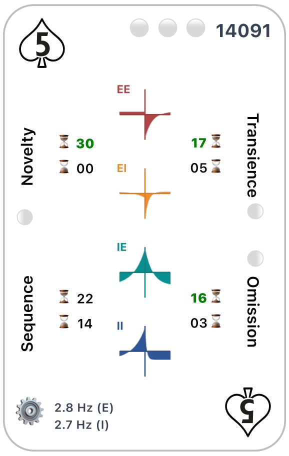
5
14091
EE
Novelty
30
00
17
05
Transience
EI
IE
Sequence
22
14
16
03
Omission
II
2.8 Hz (E)
2.7 Hz (I)

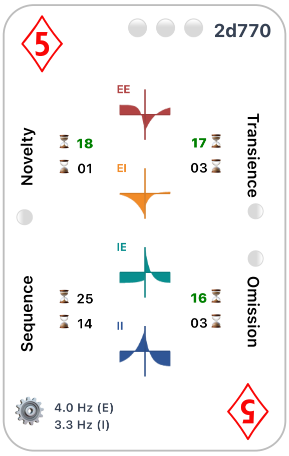
5
2d770
EE
Novelty
18
01
17
03
Transience
EI
IE
Sequence
25
14
16
03
Omission
II
4.0 Hz (E)
3.3 Hz (I)

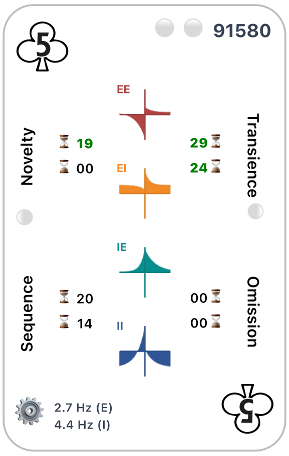
5
91580
EE
Novelty
19
00
29
24
Transience
EI
IE
Sequence
20
14
00
00
Omission
II
2.7 Hz (E)
4.4 Hz (I)

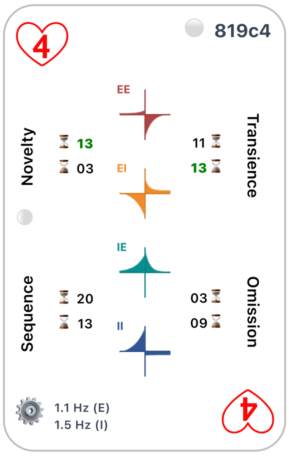
4
819c4
EE
Novelty
13
03
11
13
Transience
EI
IE
Sequence
20
13
03
09
Omission
II
1.1 Hz (E)
1.5 Hz (I)

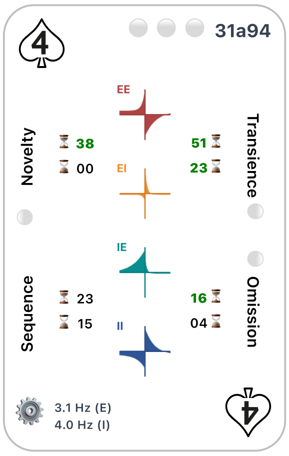
4
31a94
EE
Novelty
38
00
51
23
Transience
EI
IE
Sequence
23
15
16
04
Omission
II
3.1 Hz (E)
4.0 Hz (I)

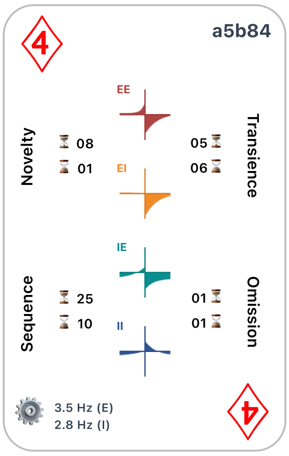
4
a5b84
EE
Novelty
08
01
05
06
Transience
EI
IE
Sequence
25
10
01
01
Omission
II
3.5 Hz (E)
2.8 Hz (I)

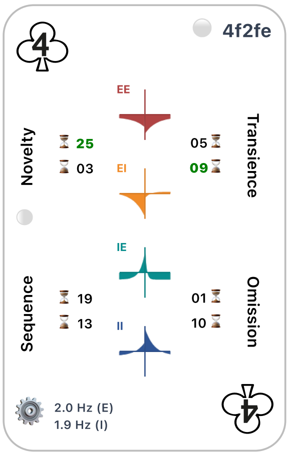
4
4f2fe
EE
Novelty
25
03
05
09
Transience
EI
IE
Sequence
19
13
01
10
Omission
II
2.0 Hz (E)
1.9 Hz (I)

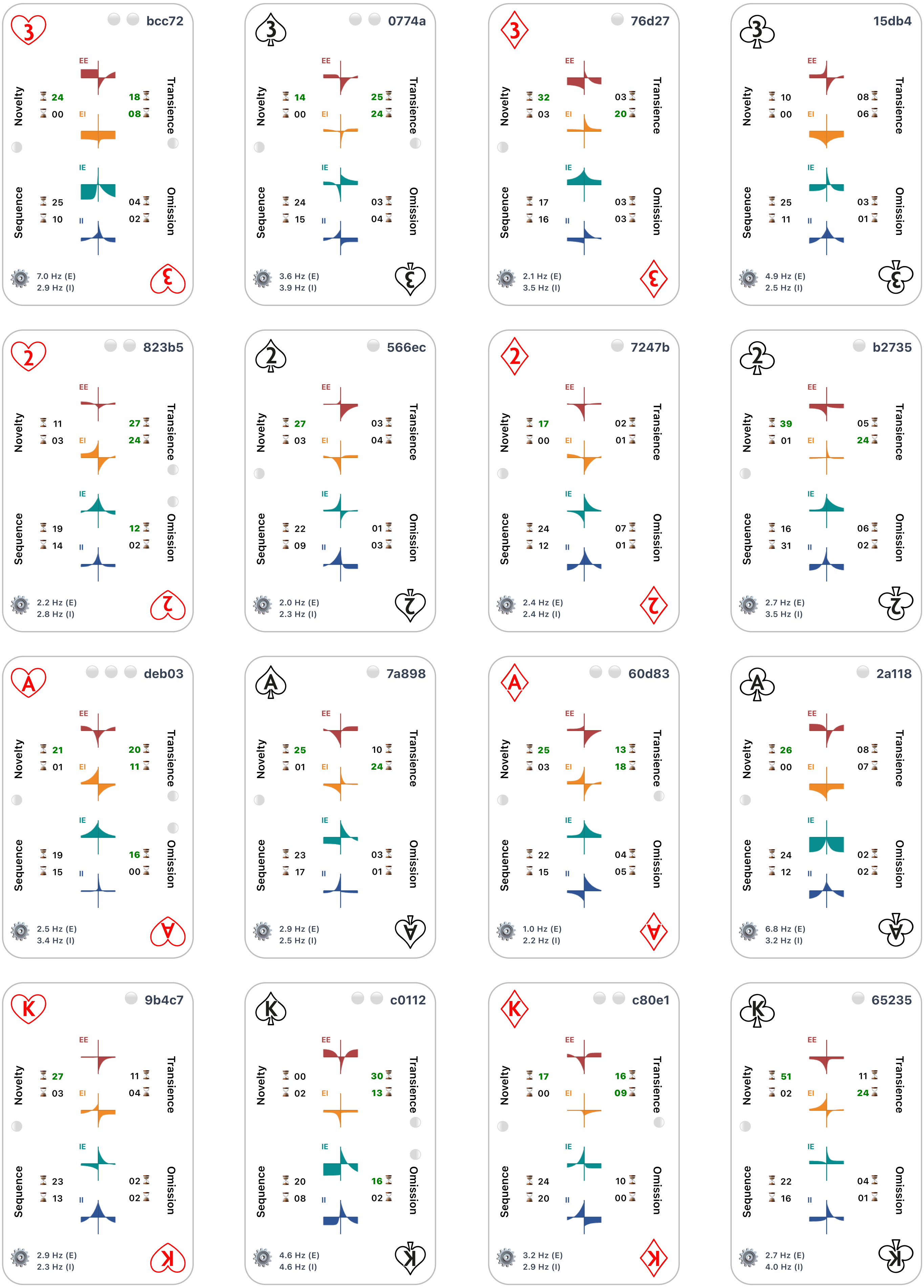
3 bcc72
EE EI IE II
Novelty 24 00
Transience 18 08
Sequence 25 10
Omission 04 02
7.0 Hz (E)
2.9 Hz (I)
3 0774a
EE EI IE II
Novelty 14 00
Transience 25 24
Sequence 24 15
Omission 03 04
3.6 Hz (E)
3.9 Hz (I)
3 76d27
EE EI IE II
Novelty 32 03
Transience 03 20
Sequence 17 16
Omission 03 03
2.1 Hz (E)
3.5 Hz (I)
3 15db4
EE EI IE II
Novelty 10 00
Transience 08 06
Sequence 25 11
Omission 03 01
4.9 Hz (E)
2.5 Hz (I)
2 823b5
EE EI IE II
Novelty 11 03
Transience 27 24
Sequence 19 14
Omission 12 02
2.2 Hz (E)
2.8 Hz (I)
2 566ec
EE EI IE II
Novelty 27 03
Transience 03 04
Sequence 22 09
Omission 01 03
2.0 Hz (E)
2.3 Hz (I)
2 7247b
EE EI IE II
Novelty 17 00
Transience 02 01
Sequence 24 12
Omission 07 01
2.4 Hz (E)
2.4 Hz (I)
2 b2735
EE EI IE II
Novelty 39 01
Transience 05 24
Sequence 16 31
Omission 06 02
2.7 Hz (E)
3.5 Hz (I)
A deb03
EE EI IE II
Novelty 21 01
Transience 20 11
Sequence 19 15
Omission 16 00
2.5 Hz (E)
3.4 Hz (I)
A 7a898
EE EI IE II
Novelty 25 01
Transience 10 24
Sequence 23 17
Omission 03 01
2.9 Hz (E)
2.5 Hz (I)
A 60d83
EE EI IE II
Novelty 25 03
Transience 13 18
Sequence 22 15
Omission 04 05
1.0 Hz (E)
2.2 Hz (I)
A 2a118
EE EI IE II
Novelty 26 00
Transience 08 07
Sequence 24 12
Omission 02 02
6.8 Hz (E)
3.2 Hz (I)
K 9b4c7
EE EI IE II
Novelty 27 03
Transience 11 04
Sequence 23 13
Omission 02 02
2.9 Hz (E)
2.3 Hz (I)
K c0112
EE EI IE II
Novelty 00 02
Transience 30 13
Sequence 20 08
Omission 16 02
4.6 Hz (E)
4.6 Hz (I)
K c80e1
EE EI IE II
Novelty 17 00
Transience 16 09
Sequence 24 20
Omission 10 00
3.2 Hz (E)
2.9 Hz (I)
K 65235
EE EI IE II
Novelty 51 02
Transience 11 24
Sequence 22 16
Omission 04 01
2.7 Hz (E)
4.0 Hz (I)

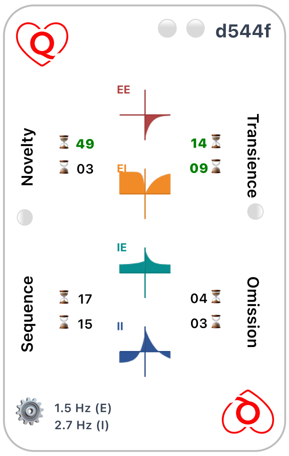
d544f
Novelty 49 03
Transience 14 09
Sequence 17 15
Omission 04 03
EE EI IE II
1.5 Hz (E)
2.7 Hz (I)

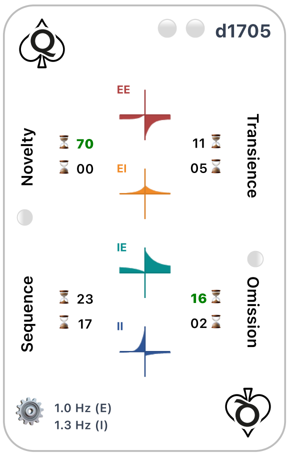
d1705
Novelty 70 00
Transience 11 05
Sequence 23 17
Omission 16 02
EE EI IE II
1.0 Hz (E)
1.3 Hz (I)

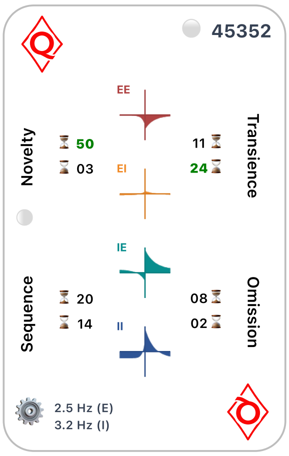
45352
Novelty 50 03
Transience 11 24
Sequence 20 14
Omission 08 02
EE EI IE II
2.5 Hz (E)
3.2 Hz (I)

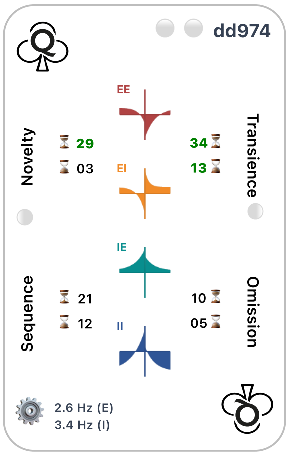
dd974
Novelty 29 03
Transience 34 13
Sequence 21 12
Omission 10 05
EE EI IE II
2.6 Hz (E)
3.4 Hz (I)

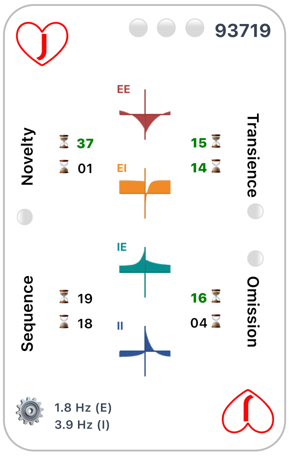
93719
Novelty 37 01
Transience 15 14
Sequence 19 18
Omission 16 04
EE EI IE II
1.8 Hz (E)
3.9 Hz (I)

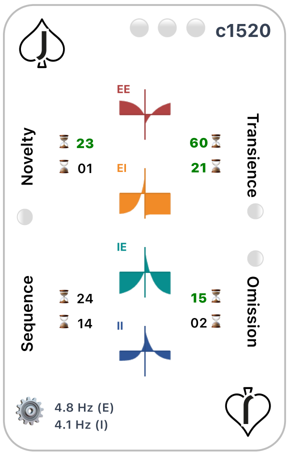
c1520
Novelty 23 01
Transience 60 21
Sequence 24 14
Omission 15 02
EE EI IE II
4.8 Hz (E)
4.1 Hz (I)

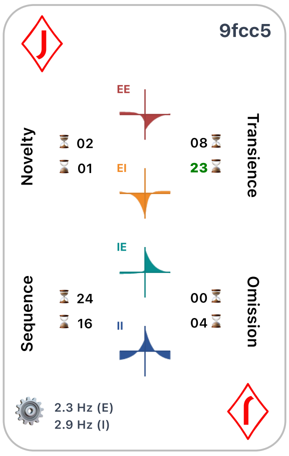
9fcc5
Novelty 02 01
Transience 08 23
Sequence 24 16
Omission 00 04
EE EI IE II
2.3 Hz (E)
2.9 Hz (I)

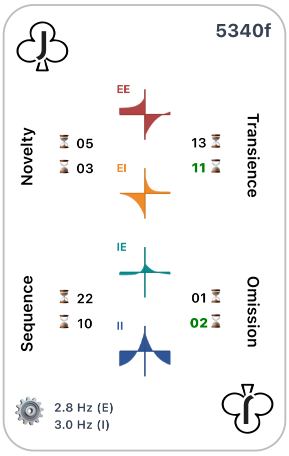
5340f
Novelty 05 03
Transience 13 11
Sequence 22 10
Omission 01 02
EE EI IE II
2.8 Hz (E)
3.0 Hz (I)

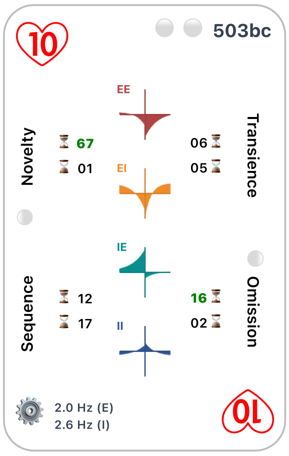
503bc
Novelty 67 01
Transience 06 05
Sequence 12 17
Omission 16 02
EE EI IE II
2.0 Hz (E)
2.6 Hz (I)

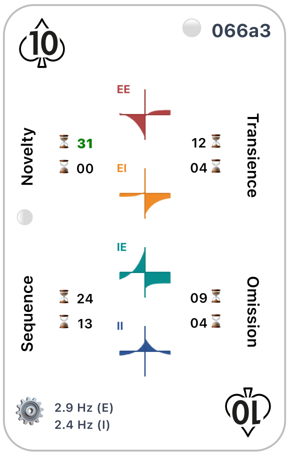
066a3
Novelty 31 00
Transience 12 04
Sequence 24 13
Omission 09 04
EE EI IE II
2.9 Hz (E)
2.4 Hz (I)

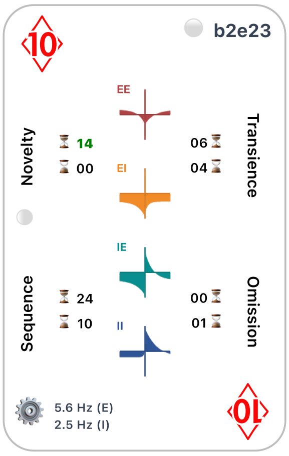
b2e23
Novelty 14 00
Transience 06 04
Sequence 24 10
Omission 00 01
EE EI IE II
5.6 Hz (E)
2.5 Hz (I)

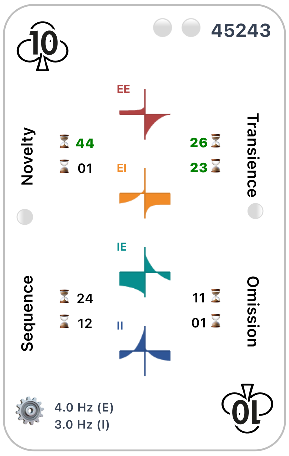
45243
Novelty 44 01
Transience 26 23
Sequence 24 12
Omission 11 01
EE EI IE II
4.0 Hz (E)
3.0 Hz (I)

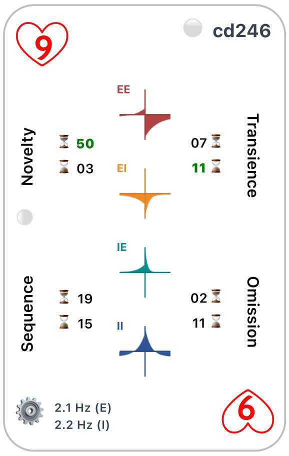
cd246
Novelty 50 03
Transience 07 11
Sequence 19 15
Omission 02 11
EE EI IE II
2.1 Hz (E)
2.2 Hz (I)

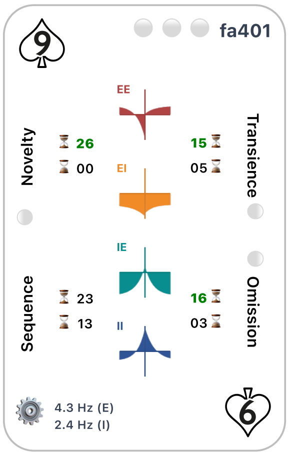
fa401
Novelty 26 00
Transience 15 05
Sequence 23 13
Omission 16 03
EE EI IE II
4.3 Hz (E)
2.4 Hz (I)

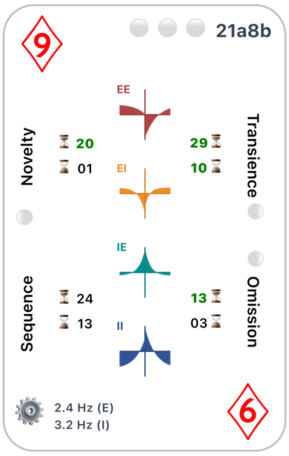
21a8b
Novelty 20 01
Transience 29 10
Sequence 24 13
Omission 13 03
EE EI IE II
2.4 Hz (E)
3.2 Hz (I)

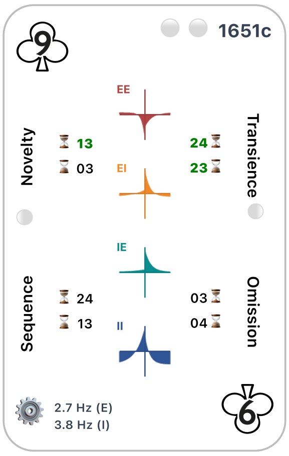
1651c
Novelty 13 03
Transience 24 23
Sequence 24 13
Omission 03 04
EE EI IE II
2.7 Hz (E)
3.8 Hz (I)

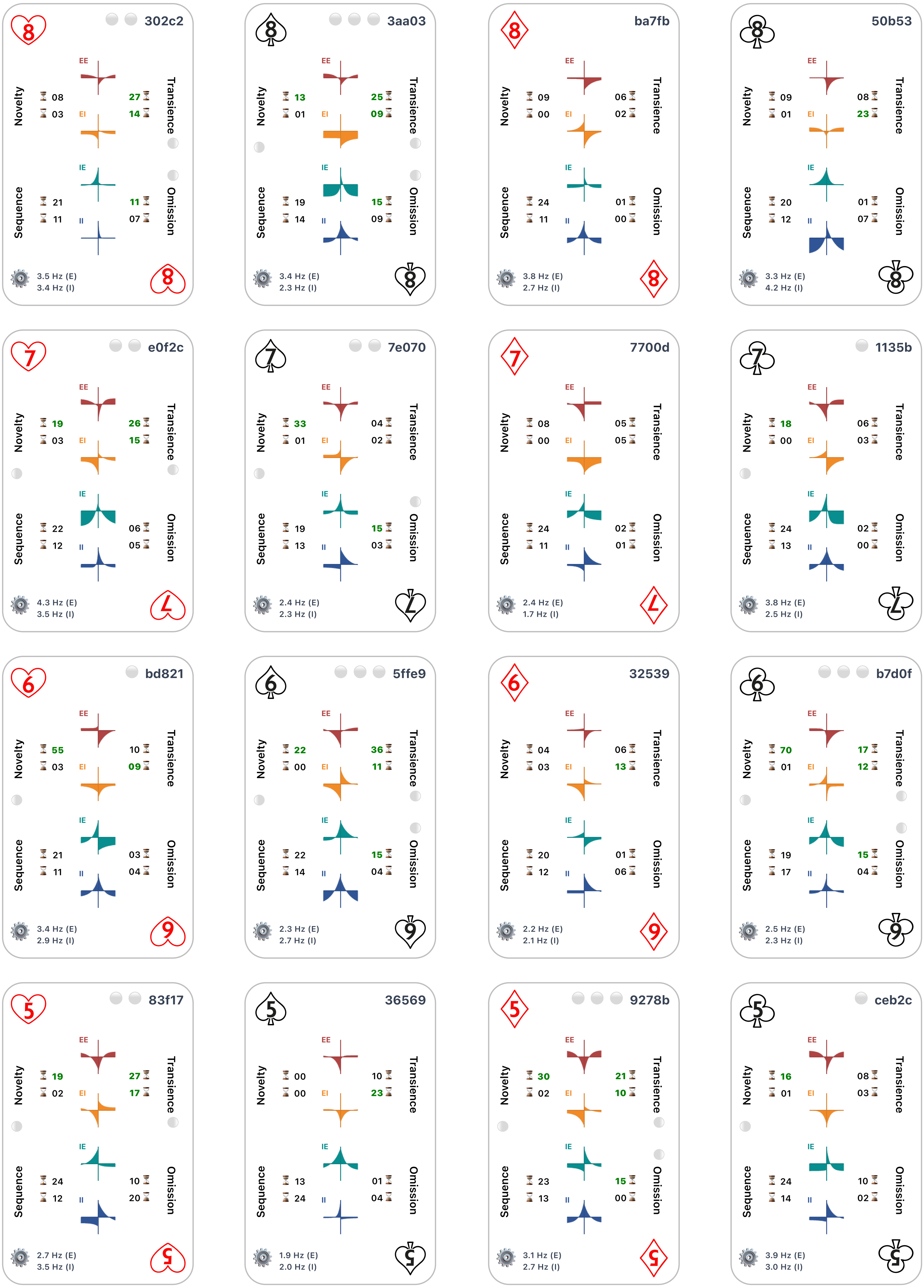

8
302c2
Novelty
08
03
27
14
Transience
EE
EI
IE
II
Sequence
21
11
11
07
Omission
3.5 Hz (E)
3.4 Hz (I)
8
3aa03
Novelty
13
01
25
09
Transience
Sequence
19
14
15
09
Omission
3.4 Hz (E)
2.3 Hz (I)
8
ba7fb
Novelty
09
00
06
02
Transience
Sequence
24
11
01
00
Omission
3.8 Hz (E)
2.7 Hz (I)
8
50b53
Novelty
09
01
08
23
Transience
Sequence
20
12
01
07
Omission
3.3 Hz (E)
4.2 Hz (I)
7
e0f2c
Novelty
19
03
26
15
Transience
Sequence
22
12
06
05
Omission
4.3 Hz (E)
3.5 Hz (I)
7
7e070
Novelty
33
01
04
02
Transience
Sequence
19
13
15
03
Omission
2.4 Hz (E)
2.3 Hz (I)
7
7700d
Novelty
08
00
05
05
Transience
Sequence
24
11
02
01
Omission
2.4 Hz (E)
1.7 Hz (I)
7
1135b
Novelty
18
00
06
03
Transience
Sequence
24
13
02
00
Omission
3.8 Hz (E)
2.5 Hz (I)
6
bd821
Novelty
55
03
10
09
Transience
Sequence
21
11
03
04
Omission
3.4 Hz (E)
2.9 Hz (I)
6
5ffe9
Novelty
22
00
36
11
Transience
Sequence
22
14
15
04
Omission
2.3 Hz (E)
2.7 Hz (I)
6
32539
Novelty
04
03
06
13
Transience
Sequence
20
12
01
06
Omission
2.2 Hz (E)
2.1 Hz (I)
6
b7d0f
Novelty
70
01
17
12
Transience
Sequence
19
17
15
04
Omission
2.5 Hz (E)
2.3 Hz (I)
5
83f17
Novelty
19
02
27
17
Transience
Sequence
24
12
10
20
Omission
2.7 Hz (E)
3.5 Hz (I)
5
36569
Novelty
00
00
10
23
Transience
Sequence
13
24
01
04
Omission
1.9 Hz (E)
2.0 Hz (I)
5
9278b
Novelty
30
02
21
10
Transience
Sequence
23
13
15
00
Omission
3.1 Hz (E)
2.7 Hz (I)
5
ceb2c
Novelty
16
01
08
03
Transience
Sequence
24
14
10
02
Omission
3.9 Hz (E)
3.0 Hz (I)

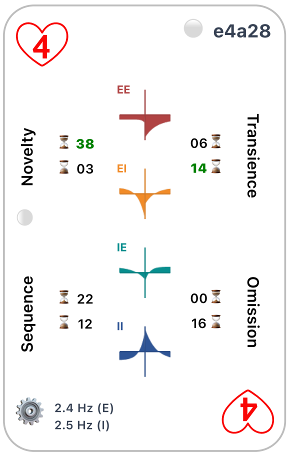
4
e4a28
EE
Novelty
38
03
06
14
Transience
EI
IE
Sequence
22
12
00
16
Omission
II
2.4 Hz (E)
2.5 Hz (I)

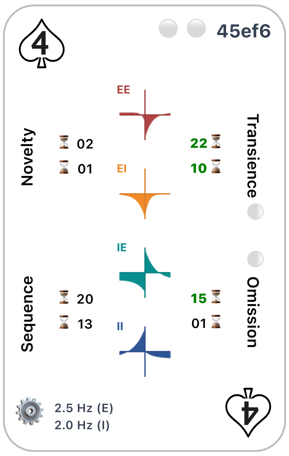
4
45ef6
EE
Novelty
02
01
22
10
Transience
EI
IE
Sequence
20
13
15
01
Omission
II
2.5 Hz (E)
2.0 Hz (I)

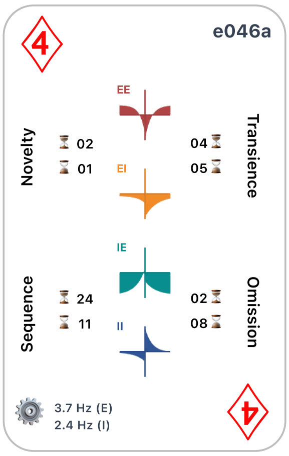
4
e046a
EE
Novelty
02
01
04
05
Transience
EI
IE
Sequence
24
11
02
08
Omission
II
3.7 Hz (E)
2.4 Hz (I)

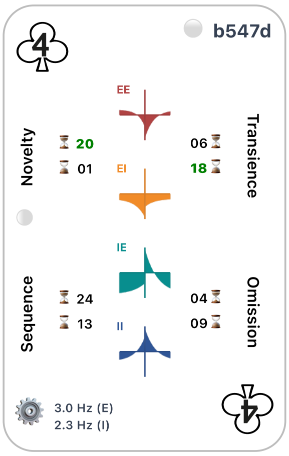
4
b547d
EE
Novelty
20
01
06
18
Transience
EI
IE
Sequence
24
13
04
09
Omission
II
3.0 Hz (E)
2.3 Hz (I)

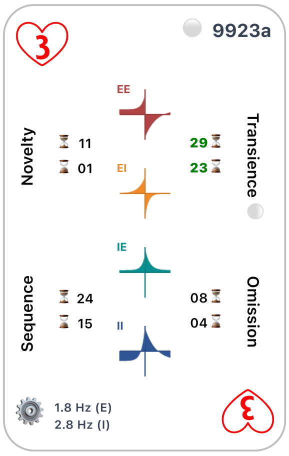
3
9923a
EE
Novelty
11
01
29
23
Transience
EI
IE
Sequence
24
15
08
04
Omission
II
1.8 Hz (E)
2.8 Hz (I)

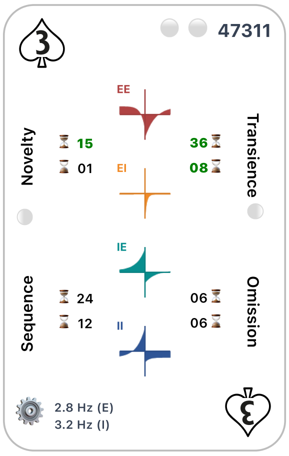
3
47311
EE
Novelty
15
01
36
08
Transience
EI
IE
Sequence
24
12
06
06
Omission
II
2.8 Hz (E)
3.2 Hz (I)

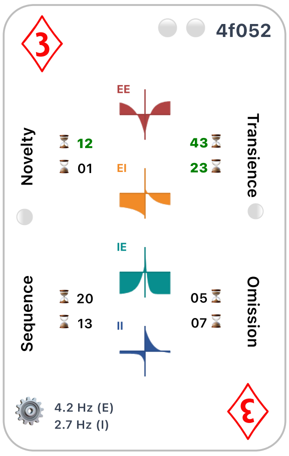
3
4f052
EE
Novelty
12
01
43
23
Transience
EI
IE
Sequence
20
13
05
07
Omission
II
4.2 Hz (E)
2.7 Hz (I)

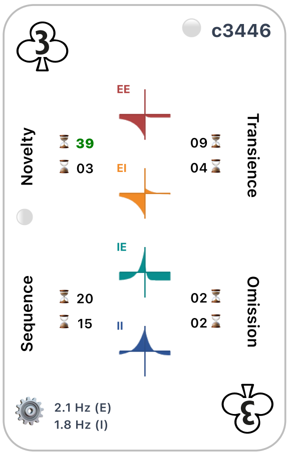
3
c3446
EE
Novelty
39
03
09
04
Transience
EI
IE
Sequence
20
15
02
02
Omission
II
2.1 Hz (E)
1.8 Hz (I)

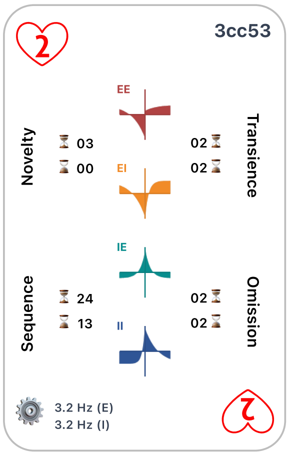
2
3cc53
EE
Novelty
03
00
02
02
Transience
EI
IE
Sequence
24
13
02
02
Omission
II
3.2 Hz (E)
3.2 Hz (I)

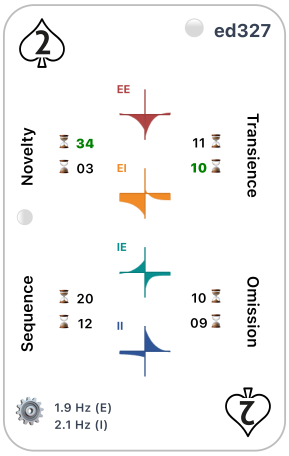
2
ed327
EE
Novelty
34
03
11
10
Transience
EI
IE
Sequence
20
12
10
09
Omission
II
1.9 Hz (E)
2.1 Hz (I)

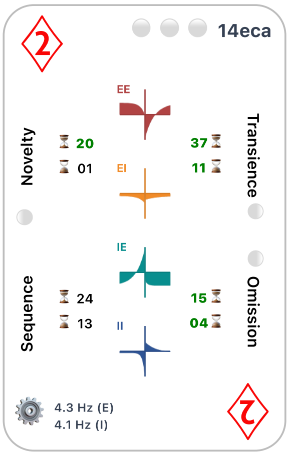
2
14eca
EE
Novelty
20
01
37
11
Transience
EI
IE
Sequence
24
13
15
04
Omission
II
4.3 Hz (E)
4.1 Hz (I)

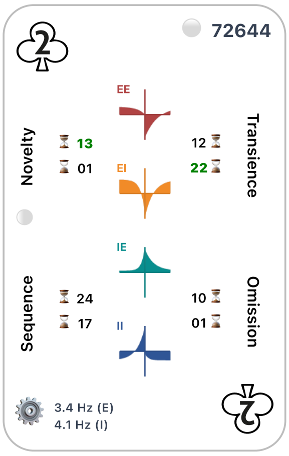
2
72644
EE
Novelty
13
01
12
22
Transience
EI
IE
Sequence
24
17
10
01
Omission
II
3.4 Hz (E)
4.1 Hz (I)

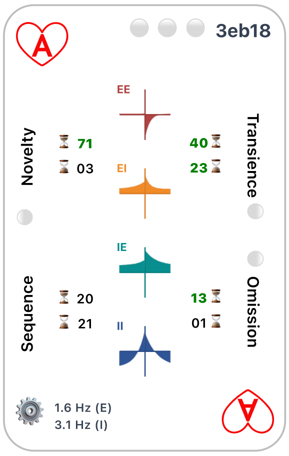
A
3eb18
EE
Novelty
71
03
40
23
Transience
EI
IE
Sequence
20
21
13
01
Omission
II
1.6 Hz (E)
3.1 Hz (I)

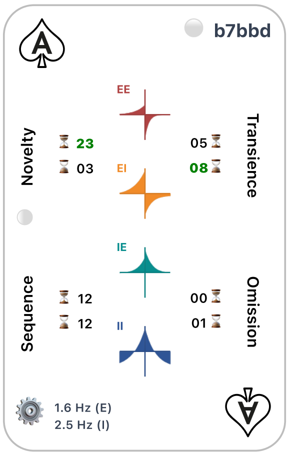
A
b7bbd
EE
Novelty
23
03
05
08
Transience
EI
IE
Sequence
12
12
00
01
Omission
II
1.6 Hz (E)
2.5 Hz (I)

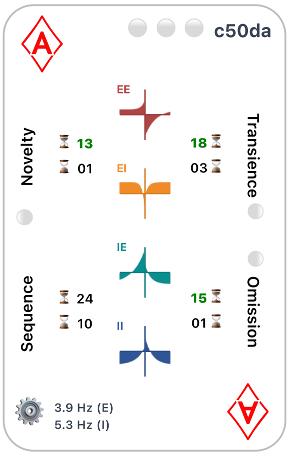
A
c50da
EE
Novelty
13
01
18
03
Transience
EI
IE
Sequence
24
10
15
01
Omission
II
3.9 Hz (E)
5.3 Hz (I)

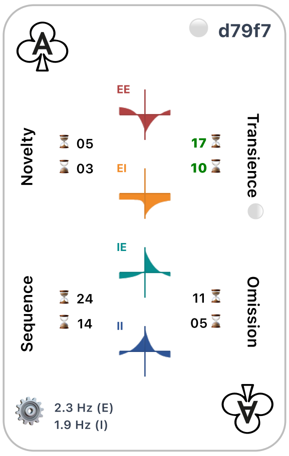
A
d79f7
EE
Novelty
05
03
17
10
Transience
EI
IE
Sequence
24
14
11
05
Omission
II
2.3 Hz (E)
1.9 Hz (I)

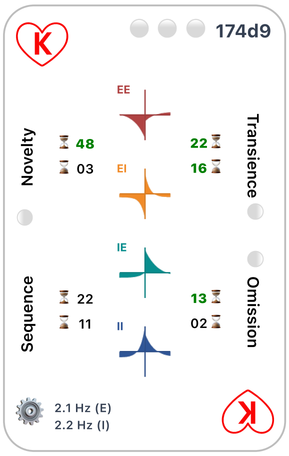
K
174d9
EE
Novelty
48
03
EI
22
16
Transience
IE
Sequence
22
11
II
13
02
Omission
2.1 Hz (E)
2.2 Hz (I)

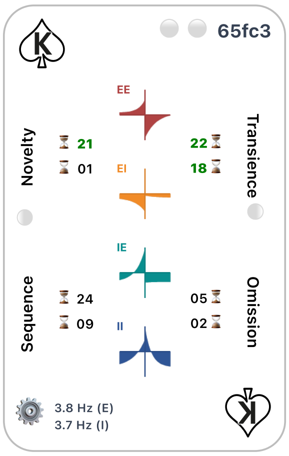
K
65fc3
EE
Novelty
21
01
EI
22
18
Transience
IE
Sequence
24
09
II
05
02
Omission
3.8 Hz (E)
3.7 Hz (I)

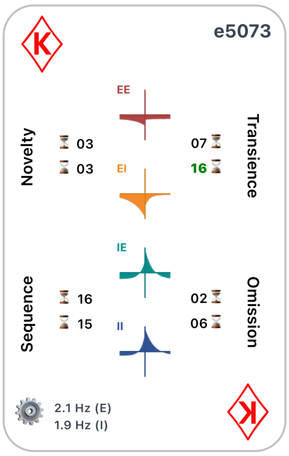
K
e5073
EE
Novelty
03
03
EI
07
16
Transience
IE
Sequence
16
15
II
02
06
Omission
2.1 Hz (E)
1.9 Hz (I)

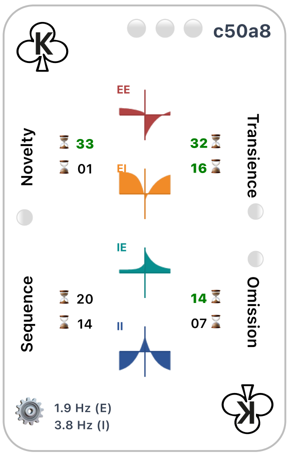
K
c50a8
EE
Novelty
33
01
EI
32
16
Transience
IE
Sequence
20
14
II
14
07
Omission
1.9 Hz (E)
3.8 Hz (I)

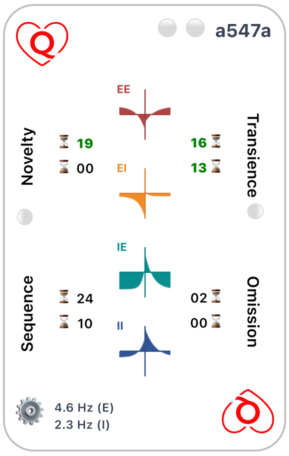
Q
a547a
EE
Novelty
19
00
EI
16
13
Transience
IE
Sequence
24
10
II
02
00
Omission
4.6 Hz (E)
2.3 Hz (I)

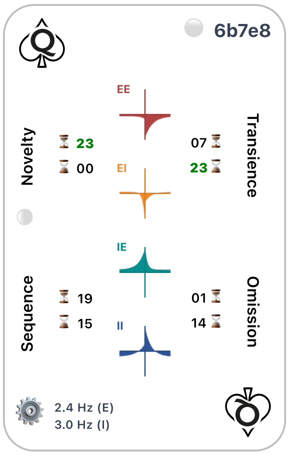
Q
6b7e8
EE
Novelty
23
00
EI
07
23
Transience
IE
Sequence
19
15
II
01
14
Omission
2.4 Hz (E)
3.0 Hz (I)

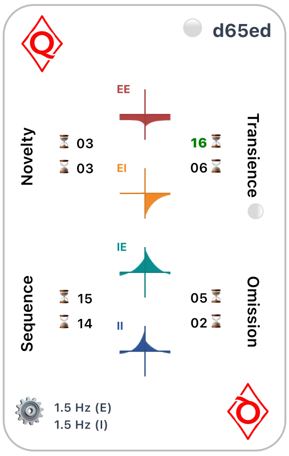
Q
d65ed
EE
Novelty
03
03
EI
16
06
Transience
IE
Sequence
15
14
II
05
02
Omission
1.5 Hz (E)
1.5 Hz (I)

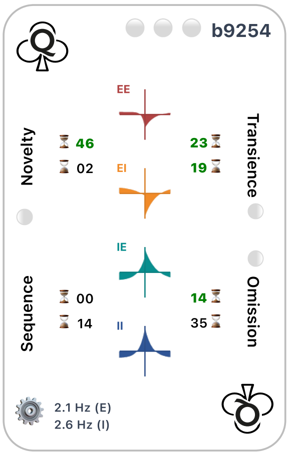
Q
b9254
EE
Novelty
46
02
EI
23
19
Transience
IE
Sequence
00
14
II
14
35
Omission
2.1 Hz (E)
2.6 Hz (I)

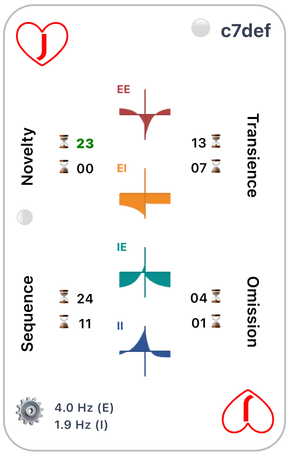
J
c7def
EE
Novelty
23
00
EI
13
07
Transience
IE
Sequence
24
11
II
04
01
Omission
4.0 Hz (E)
1.9 Hz (I)

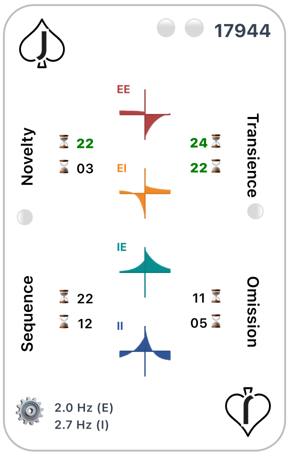
J
17944
EE
Novelty
22
03
EI
24
22
Transience
IE
Sequence
22
12
II
11
05
Omission
2.0 Hz (E)
2.7 Hz (I)

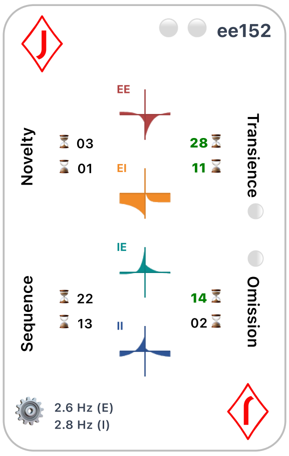
J
ee152
EE
Novelty
03
01
EI
28
11
Transience
IE
Sequence
22
13
II
14
02
Omission
2.6 Hz (E)
2.8 Hz (I)

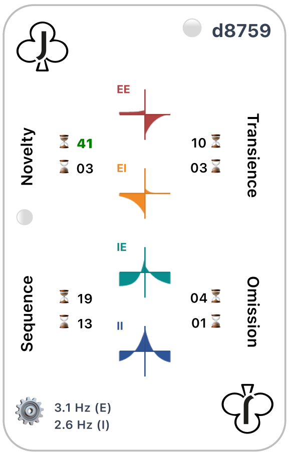
J
d8759
EE
Novelty
41
03
EI
10
03
Transience
IE
Sequence
19
13
II
04
01
Omission
3.1 Hz (E)
2.6 Hz (I)

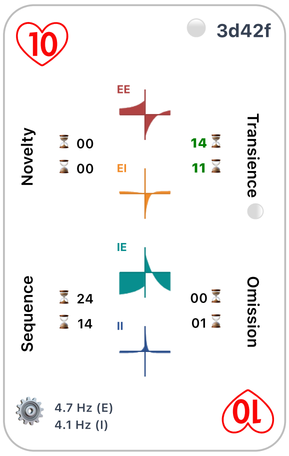
10
3d42f
EE
Novelty
00
00
EI
14
11
Transience
IE
Sequence
24
14
II
00
01
Omission
4.7 Hz (E)
4.1 Hz (I)

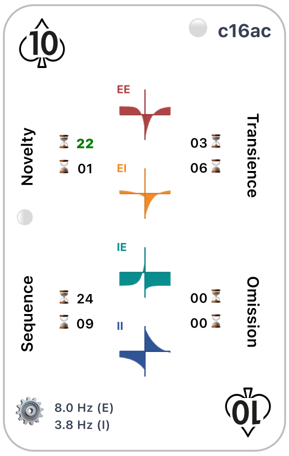
10
c16ac
EE
Novelty
22
01
EI
03
06
Transience
IE
Sequence
24
09
II
00
00
Omission
8.0 Hz (E)
3.8 Hz (I)

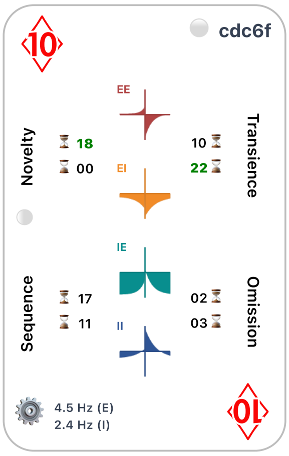
10
cdc6f
EE
Novelty
18
00
EI
10
22
Transience
IE
Sequence
17
11
II
02
03
Omission
4.5 Hz (E)
2.4 Hz (I)

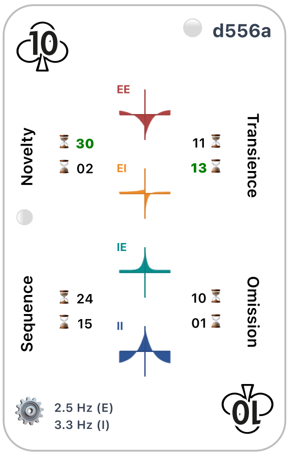
10
d556a
EE
Novelty
30
02
EI
11
13
Transience
IE
Sequence
24
15
II
10
01
Omission
2.5 Hz (E)
3.3 Hz (I)

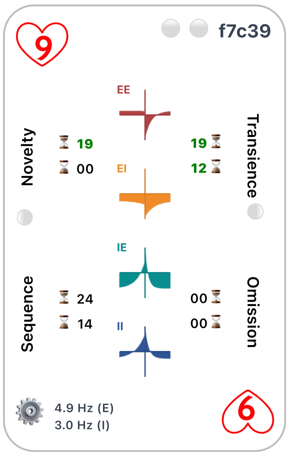
9
f7c39
EE
EI
IE
II
Novelty
19
00
Transience
19
12
Sequence
24
14
Omission
00
00
4.9 Hz (E)
3.0 Hz (I)
6

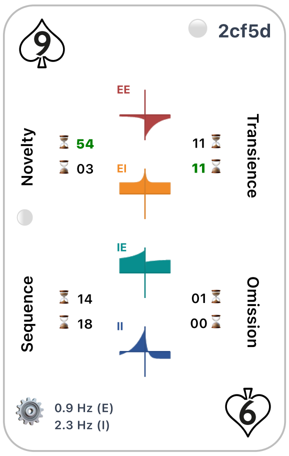
9
2cf5d
EE
EI
IE
II
Novelty
54
03
Transience
11
11
Sequence
14
18
Omission
01
00
0.9 Hz (E)
2.3 Hz (I)
6

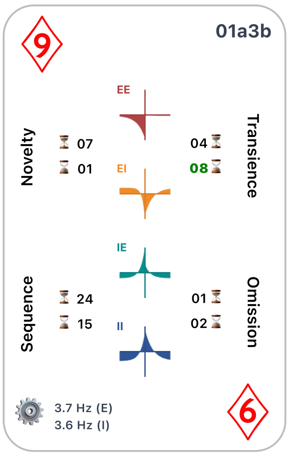
9
01a3b
EE
EI
IE
II
Novelty
07
01
Transience
04
08
Sequence
24
15
Omission
01
02
3.7 Hz (E)
3.6 Hz (I)
6

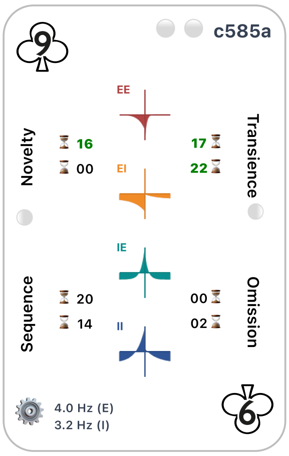
9
c585a
EE
EI
IE
II
Novelty
16
00
Transience
17
22
Sequence
20
14
Omission
00
02
4.0 Hz (E)
3.2 Hz (I)
6

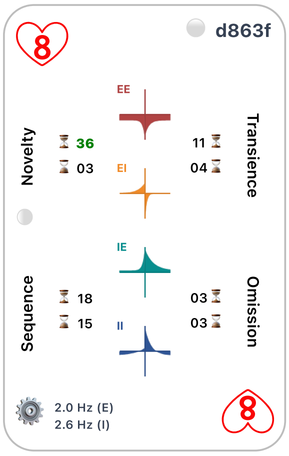
8
d863f
EE
EI
IE
II
Novelty
36
03
Transience
11
04
Sequence
18
15
Omission
03
03
2.0 Hz (E)
2.6 Hz (I)
8

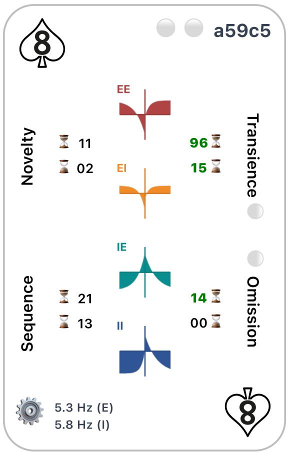
8
a59c5
EE
EI
IE
II
Novelty
11
02
Transience
96
15
Sequence
21
13
Omission
14
00
5.3 Hz (E)
5.8 Hz (I)
8

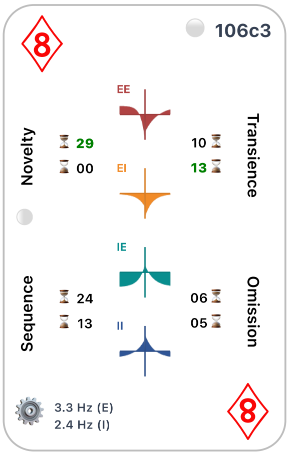
8
106c3
EE
EI
IE
II
Novelty
29
00
Transience
10
13
Sequence
24
13
Omission
06
05
3.3 Hz (E)
2.4 Hz (I)
8

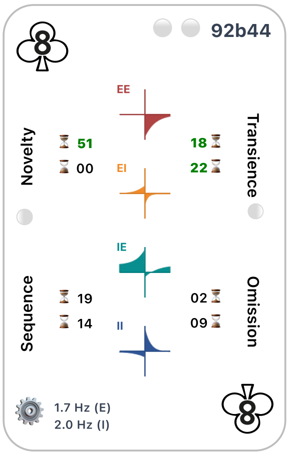
8
92b44
EE
EI
IE
II
Novelty
51
00
Transience
18
22
Sequence
19
14
Omission
02
09
1.7 Hz (E)
2.0 Hz (I)
8

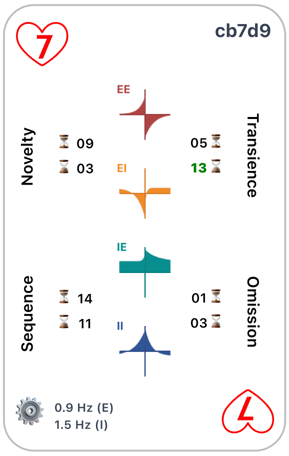
7
cb7d9
EE
EI
IE
II
Novelty
09
03
Transience
05
13
Sequence
14
11
Omission
01
03
0.9 Hz (E)
1.5 Hz (I)
7

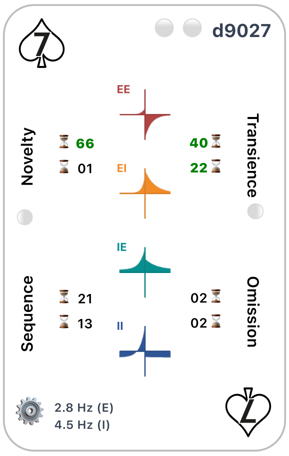
7
d9027
EE
EI
IE
II
Novelty
66
01
Transience
40
22
Sequence
21
13
Omission
02
02
2.8 Hz (E)
4.5 Hz (I)
7

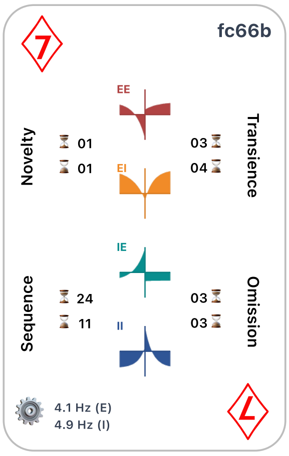
7
fc66b
EE
EI
IE
II
Novelty
01
01
Transience
03
04
Sequence
24
11
Omission
03
03
4.1 Hz (E)
4.9 Hz (I)
7

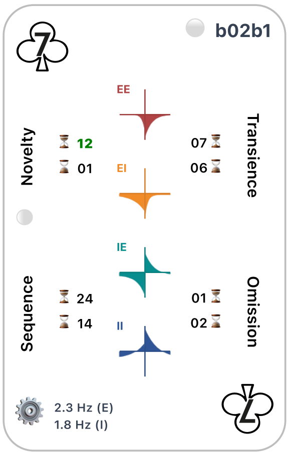
7
b02b1
EE
EI
IE
II
Novelty
12
01
Transience
07
06
Sequence
24
14
Omission
01
02
2.3 Hz (E)
1.8 Hz (I)
7

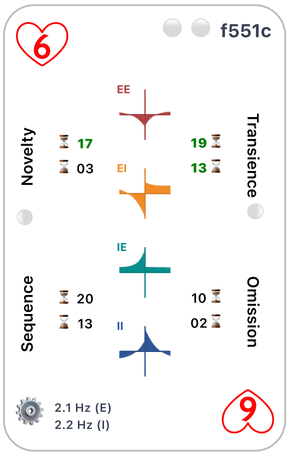
6
f551c
EE
EI
IE
II
Novelty
17
03
Transience
19
13
Sequence
20
13
Omission
10
02
2.1 Hz (E)
2.2 Hz (I)
9

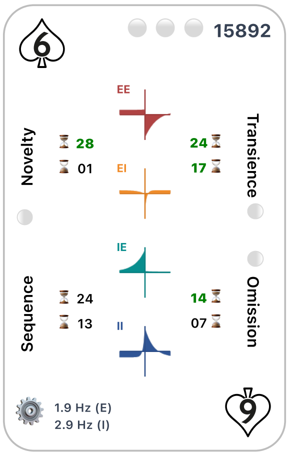
6
15892
EE
EI
IE
II
Novelty
28
01
Transience
24
17
Sequence
24
13
Omission
14
07
1.9 Hz (E)
2.9 Hz (I)
9

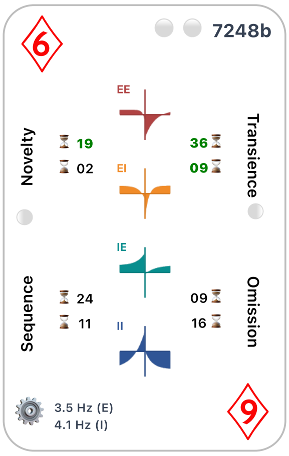
6
7248b
EE
EI
IE
II
Novelty
19
02
Transience
36
09
Sequence
24
11
Omission
09
16
3.5 Hz (E)
4.1 Hz (I)
9

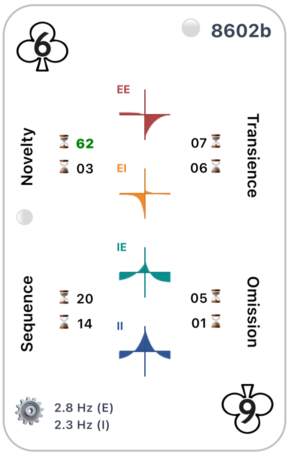
6
8602b
EE
EI
IE
II
Novelty
62
03
Transience
07
06
Sequence
20
14
Omission
05
01
2.8 Hz (E)
2.3 Hz (I)
9

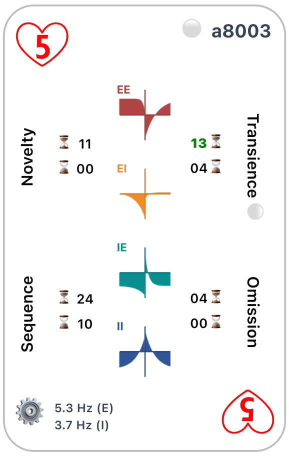
5
a8003
EE
EI
IE
II
Novelty 11 00
Transience 13 04
Sequence 24 10
Omission 04 00
5.3 Hz (E)
3.7 Hz (I)

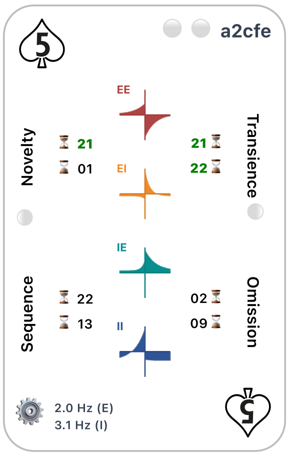
5
a2cfe
EE
EI
IE
II
Novelty 21 01
Transience 21 22
Sequence 22 13
Omission 02 09
2.0 Hz (E)
3.1 Hz (I)

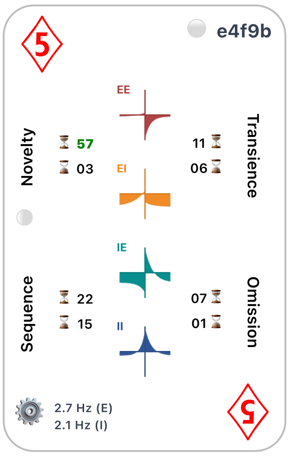
5
e4f9b
EE
EI
IE
II
Novelty 57 03
Transience 11 06
Sequence 22 15
Omission 07 01
2.7 Hz (E)
2.1 Hz (I)

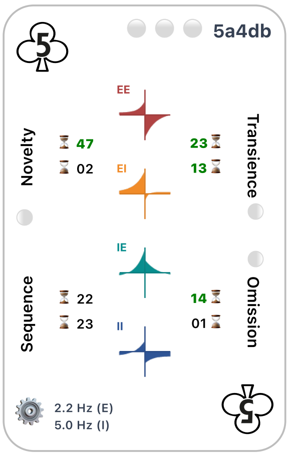
5
5a4db
EE
EI
IE
II
Novelty 47 02
Transience 23 13
Sequence 22 23
Omission 14 01
2.2 Hz (E)
5.0 Hz (I)

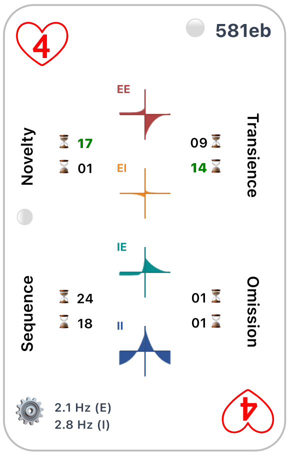
4
581eb
EE
EI
IE
II
Novelty 17 01
Transience 09 14
Sequence 24 18
Omission 01 01
2.1 Hz (E)
2.8 Hz (I)

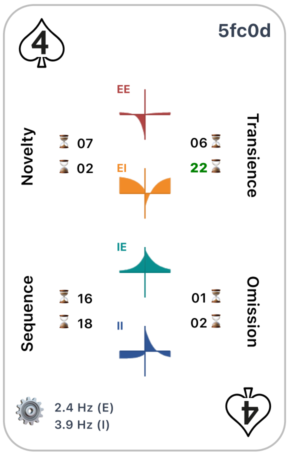
4
5fc0d
EE
EI
IE
II
Novelty 07 02
Transience 06 22
Sequence 16 18
Omission 01 02
2.4 Hz (E)
3.9 Hz (I)

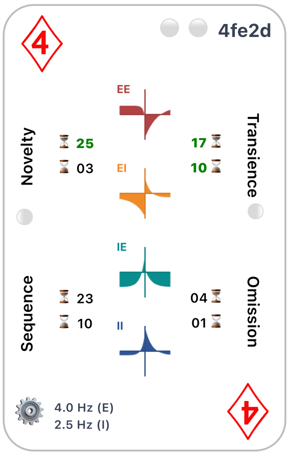
4
4fe2d
EE
EI
IE
II
Novelty 25 03
Transience 17 10
Sequence 23 10
Omission 04 01
4.0 Hz (E)
2.5 Hz (I)

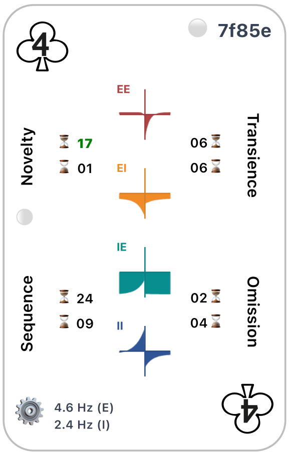
4
7f85e
EE
EI
IE
II
Novelty 17 01
Transience 06 06
Sequence 24 09
Omission 02 04
4.6 Hz (E)
2.4 Hz (I)

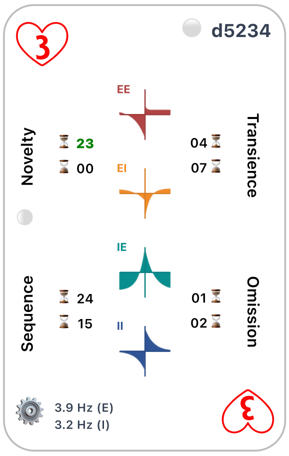
3
d5234
EE
EI
IE
II
Novelty 23 00
Transience 04 07
Sequence 24 15
Omission 01 02
3.9 Hz (E)
3.2 Hz (I)

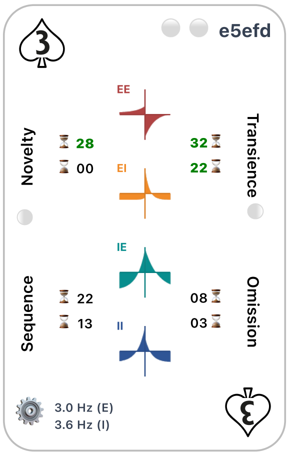
3
e5efd
EE
EI
IE
II
Novelty 28 00
Transience 32 22
Sequence 22 13
Omission 08 03
3.0 Hz (E)
3.6 Hz (I)

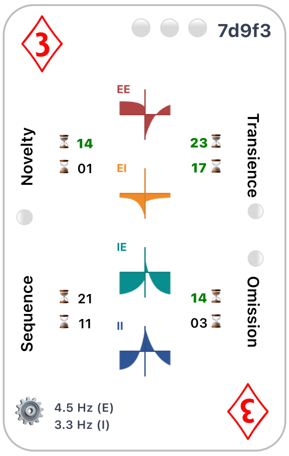
3
7d9f3
EE
EI
IE
II
Novelty 14 01
Transience 23 17
Sequence 21 11
Omission 14 03
4.5 Hz (E)
3.3 Hz (I)

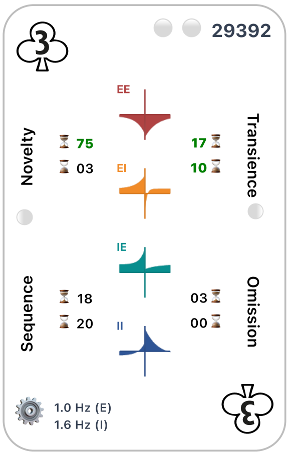
3
29392
EE
EI
IE
II
Novelty 75 03
Transience 17 10
Sequence 18 20
Omission 03 00
1.0 Hz (E)
1.6 Hz (I)

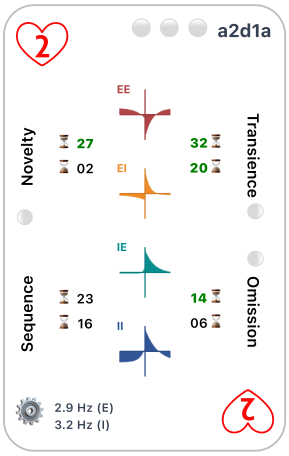
2
a2d1a
EE
EI
IE
II
Novelty 27 02
Transience 32 20
Sequence 23 16
Omission 14 06
2.9 Hz (E)
3.2 Hz (I)

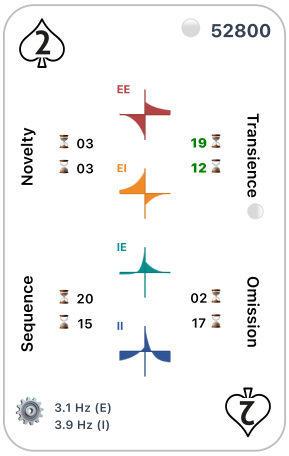
2
52800
EE
EI
IE
II
Novelty 03 03
Transience 19 12
Sequence 20 15
Omission 02 17
3.1 Hz (E)
3.9 Hz (I)

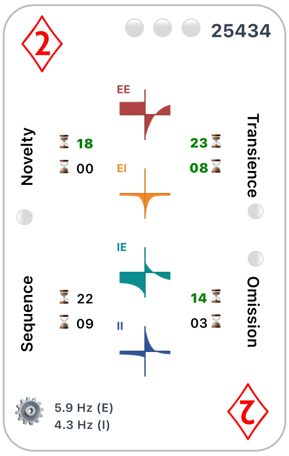
2
25434
EE
EI
IE
II
Novelty 18 00
Transience 23 08
Sequence 22 09
Omission 14 03
5.9 Hz (E)
4.3 Hz (I)

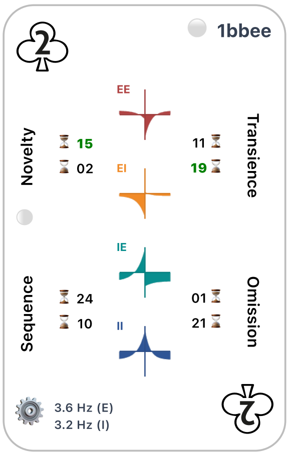
2
1bbee
EE
EI
IE
II
Novelty 15 02
Transience 11 19
Sequence 24 10
Omission 01 21
3.6 Hz (E)
3.2 Hz (I)

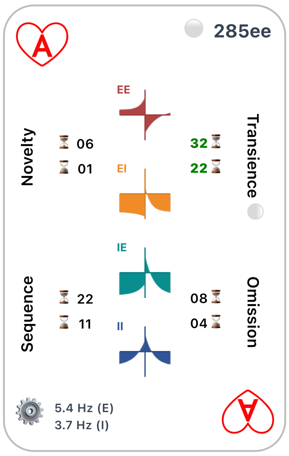
A
285ee
EE
Novelty
06
01
32
22
Transience
EI
IE
Sequence
22
11
08
04
Omission
II
5.4 Hz (E)
3.7 Hz (I)

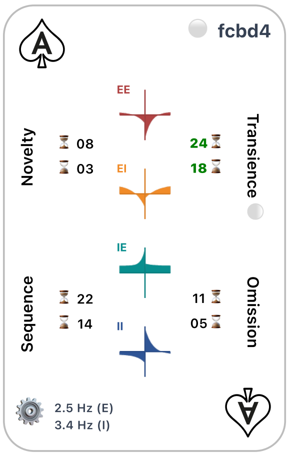
A
fcbd4
EE
Novelty
08
03
24
18
Transience
EI
IE
Sequence
22
14
11
05
Omission
II
2.5 Hz (E)
3.4 Hz (I)

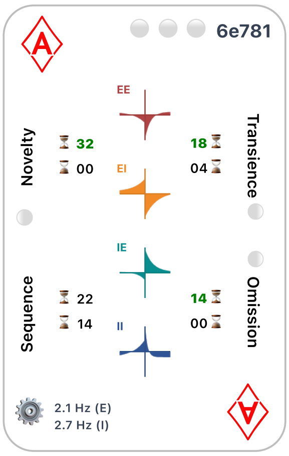
A
6e781
EE
Novelty
32
00
18
04
Transience
EI
IE
Sequence
22
14
14
00
Omission
II
2.1 Hz (E)
2.7 Hz (I)

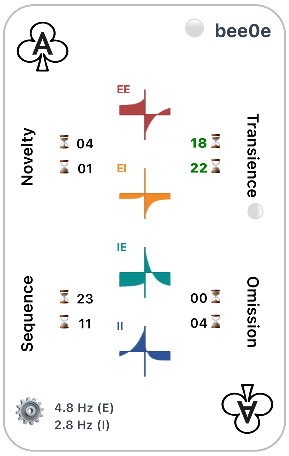
A
bee0e
EE
Novelty
04
01
18
22
Transience
EI
IE
Sequence
23
11
00
04
Omission
II
4.8 Hz (E)
2.8 Hz (I)

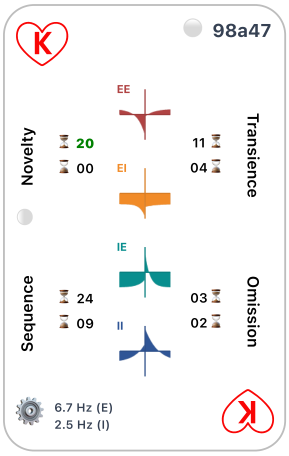
K
98a47
EE
Novelty
20
00
11
04
Transience
EI
IE
Sequence
24
09
03
02
Omission
II
6.7 Hz (E)
2.5 Hz (I)

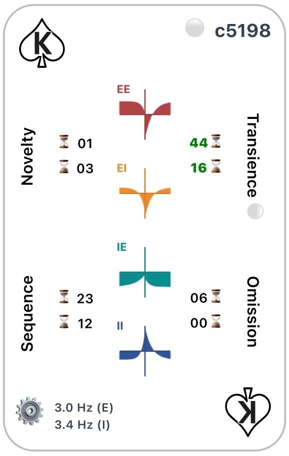
K
c5198
EE
Novelty
01
03
44
16
Transience
EI
IE
Sequence
23
12
06
00
Omission
II
3.0 Hz (E)
3.4 Hz (I)

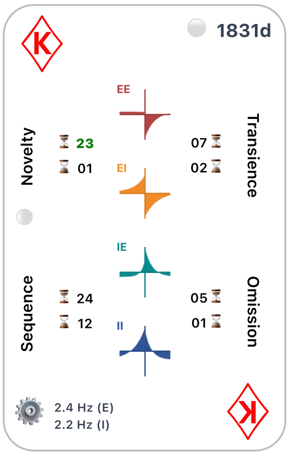
K
1831d
EE
Novelty
23
01
07
02
Transience
EI
IE
Sequence
24
12
05
01
Omission
II
2.4 Hz (E)
2.2 Hz (I)

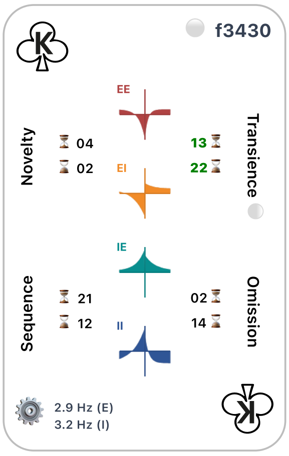
K
f3430
EE
Novelty
04
02
13
22
Transience
EI
IE
Sequence
21
12
02
14
Omission
II
2.9 Hz (E)
3.2 Hz (I)

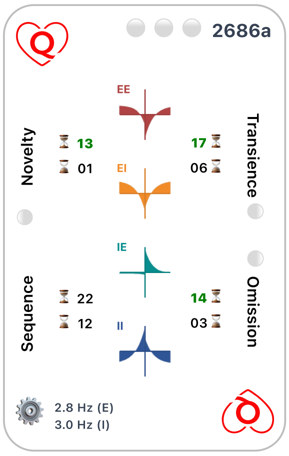
Q
2686a
EE
Novelty
13
01
17
06
Transience
EI
IE
Sequence
22
12
14
03
Omission
II
2.8 Hz (E)
3.0 Hz (I)

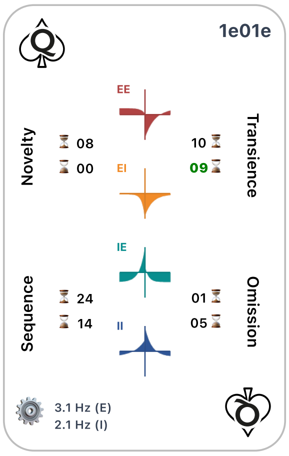
Q
1e01e
EE
Novelty
08
00
10
09
Transience
EI
IE
Sequence
24
14
01
05
Omission
II
3.1 Hz (E)
2.1 Hz (I)

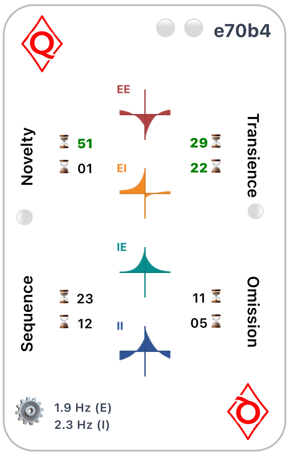
Q
e70b4
EE
Novelty
51
01
29
22
Transience
EI
IE
Sequence
23
12
11
05
Omission
II
1.9 Hz (E)
2.3 Hz (I)

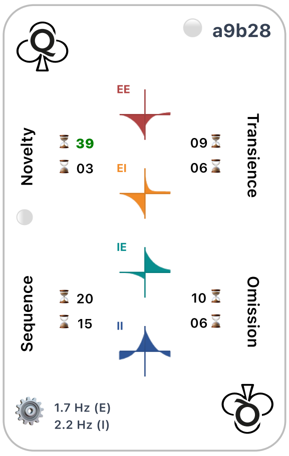
Q
a9b28
EE
Novelty
39
03
09
06
Transience
EI
IE
Sequence
20
15
10
06
Omission
II
1.7 Hz (E)
2.2 Hz (I)

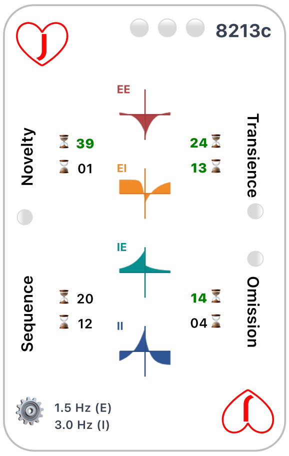
J
8213c
EE
Novelty
39
01
24
13
Transience
EI
IE
Sequence
20
12
14
04
Omission
II
1.5 Hz (E)
3.0 Hz (I)

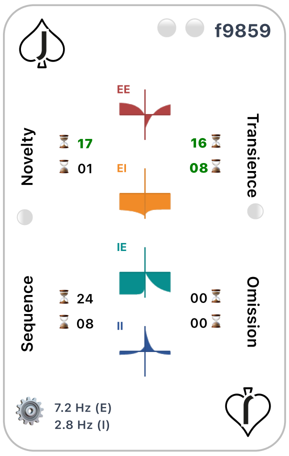
J
f9859
EE
Novelty
17
01
16
08
Transience
EI
IE
Sequence
24
08
00
00
Omission
II
7.2 Hz (E)
2.8 Hz (I)

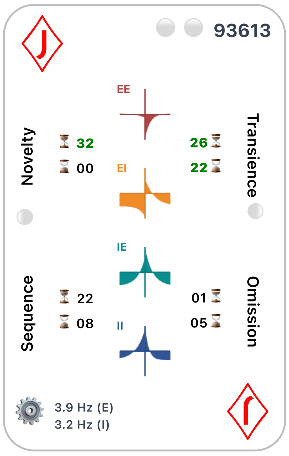
J
93613
EE
Novelty
32
00
26
22
Transience
EI
IE
Sequence
22
08
01
05
Omission
II
3.9 Hz (E)
3.2 Hz (I)

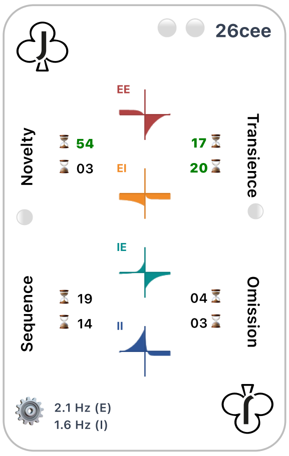
J
26cee
EE
Novelty
54
03
17
20
Transience
EI
IE
Sequence
19
14
04
03
Omission
II
2.1 Hz (E)
1.6 Hz (I)

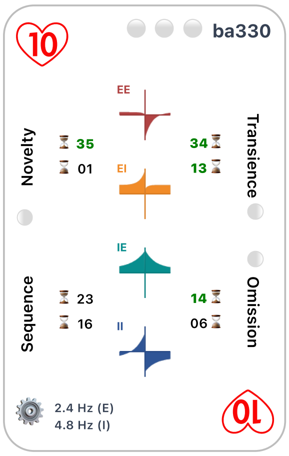
10
ba330
EE
EI
IE
II
Novelty
35
01
Transience
34
13
Sequence
23
16
Omission
14
06
2.4 Hz (E)
4.8 Hz (I)

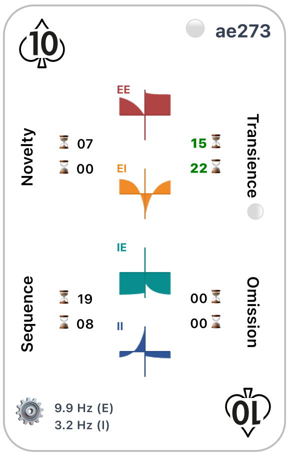
10
ae273
EE
EI
IE
II
Novelty
07
00
Transience
15
22
Sequence
19
08
Omission
00
00
9.9 Hz (E)
3.2 Hz (I)

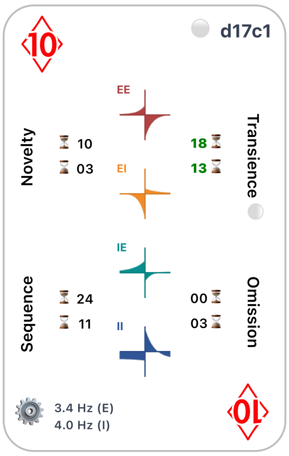
10
d17c1
EE
EI
IE
II
Novelty
10
03
Transience
18
13
Sequence
24
11
Omission
00
03
3.4 Hz (E)
4.0 Hz (I)

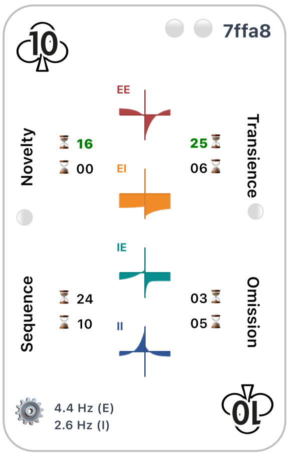
10
7ffa8
EE
EI
IE
II
Novelty
16
00
Transience
25
06
Sequence
24
10
Omission
03
05
4.4 Hz (E)
2.6 Hz (I)

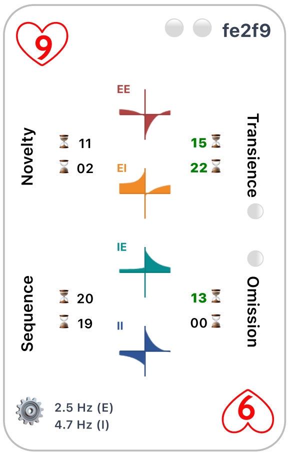
9
fe2f9
EE
EI
IE
II
Novelty
11
02
Transience
15
22
Sequence
20
19
Omission
13
00
2.5 Hz (E)
4.7 Hz (I)

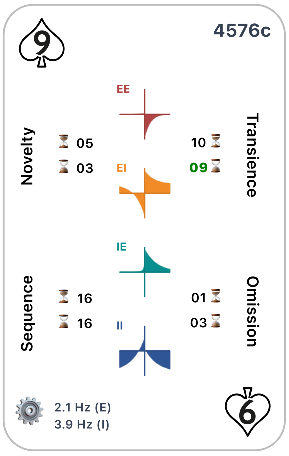
9
4576c
EE
EI
IE
II
Novelty
05
03
Transience
10
09
Sequence
16
16
Omission
01
03
2.1 Hz (E)
3.9 Hz (I)

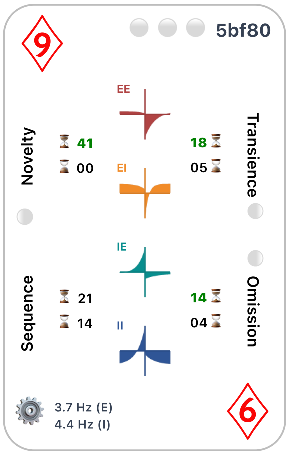
9
5bf80
EE
EI
IE
II
Novelty
41
00
Transience
18
05
Sequence
21
14
Omission
14
04
3.7 Hz (E)
4.4 Hz (I)

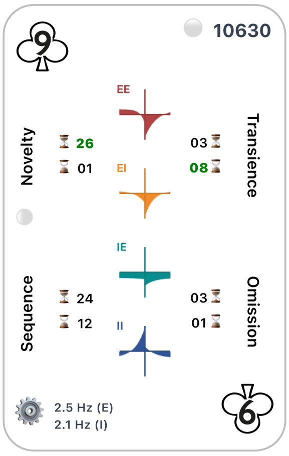
9
10630
EE
EI
IE
II
Novelty
26
01
Transience
03
08
Sequence
24
12
Omission
03
01
2.5 Hz (E)
2.1 Hz (I)

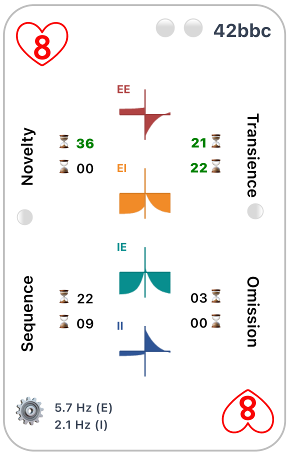
8
42bbc
EE
EI
IE
II
Novelty
36
00
Transience
21
22
Sequence
22
09
Omission
03
00
5.7 Hz (E)
2.1 Hz (I)

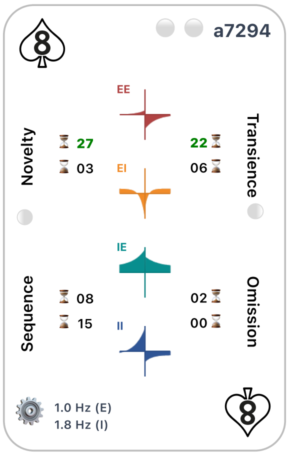
8
a7294
EE
EI
IE
II
Novelty
27
03
Transience
22
06
Sequence
08
15
Omission
02
00
1.0 Hz (E)
1.8 Hz (I)

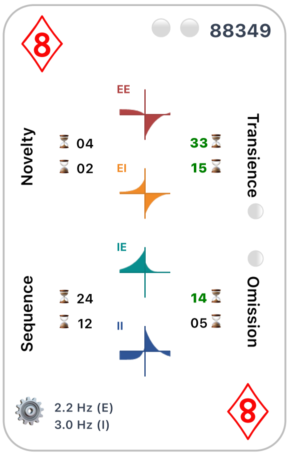
8
88349
EE
EI
IE
II
Novelty
04
02
Transience
33
15
Sequence
24
12
Omission
14
05
2.2 Hz (E)
3.0 Hz (I)

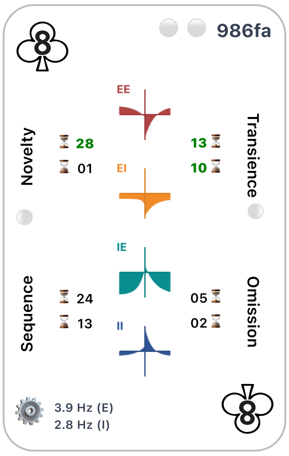
8
986fa
EE
EI
IE
II
Novelty
28
01
Transience
13
10
Sequence
24
13
Omission
05
02
3.9 Hz (E)
2.8 Hz (I)

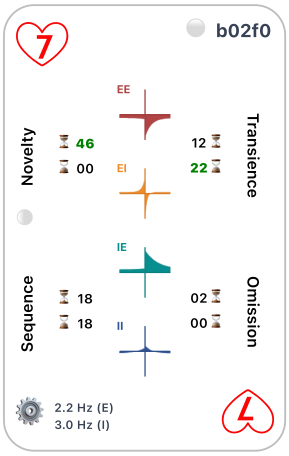
7
b02f0
EE
EI
IE
II
Novelty
46
00
Transience
12
22
Sequence
18
18
Omission
02
00
2.2 Hz (E)
3.0 Hz (I)

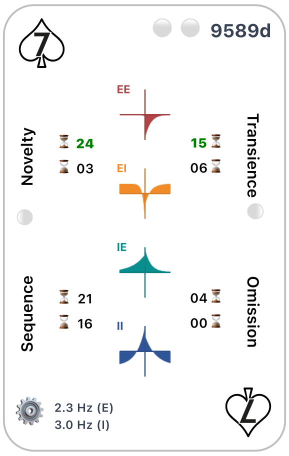
7
9589d
EE
EI
IE
II
Novelty
24
03
Transience
15
06
Sequence
21
16
Omission
04
00
2.3 Hz (E)
3.0 Hz (I)

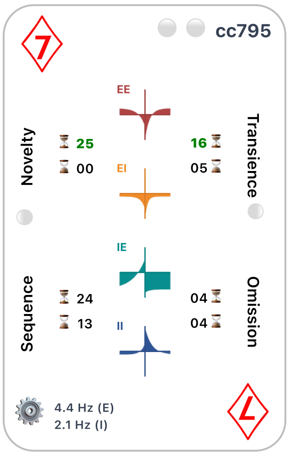
7
cc795
EE
EI
IE
II
Novelty
25
00
Transience
16
05
Sequence
24
13
Omission
04
04
4.4 Hz (E)
2.1 Hz (I)

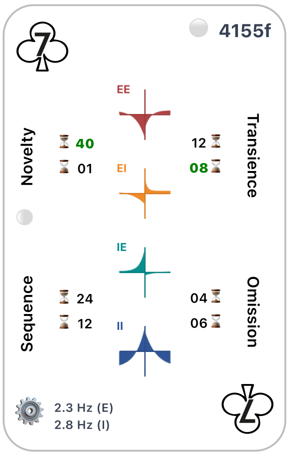
7
4155f
EE
EI
IE
II
Novelty
40
01
Transience
12
08
Sequence
24
12
Omission
04
06
2.3 Hz (E)
2.8 Hz (I)

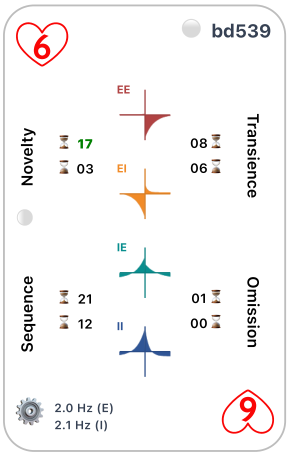
6
bd539
Novelty 17 03
Transience 08 06
Sequence 21 12
Omission 01 00
EE EI IE II
2.0 Hz (E)
2.1 Hz (I)
9

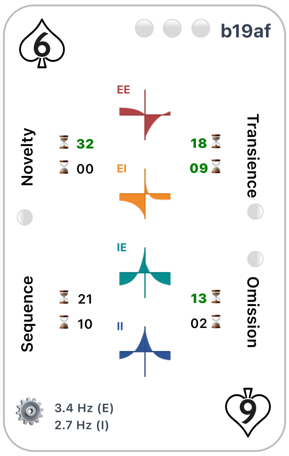
6
b19af
Novelty 32 00
Transience 18 09
Sequence 21 10
Omission 13 02
EE EI IE II
3.4 Hz (E)
2.7 Hz (I)
9

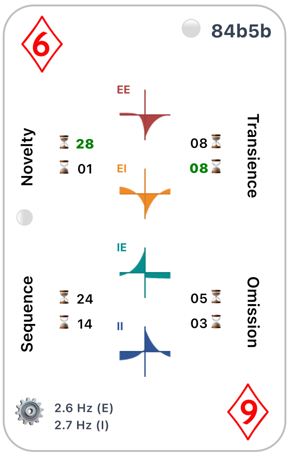
6
84b5b
Novelty 28 01
Transience 08 08
Sequence 24 14
Omission 05 03
EE EI IE II
2.6 Hz (E)
2.7 Hz (I)
9

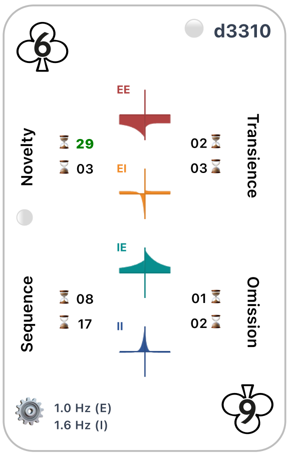
6
d3310
Novelty 29 03
Transience 02 03
Sequence 08 17
Omission 01 02
EE EI IE II
1.0 Hz (E)
1.6 Hz (I)
9

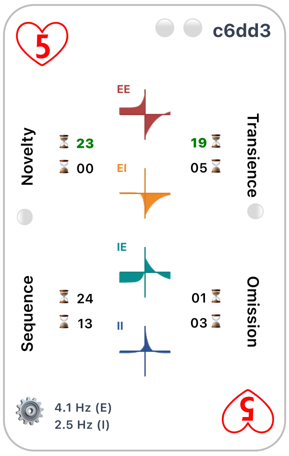
5
c6dd3
Novelty 23 00
Transience 19 05
Sequence 24 13
Omission 01 03
EE EI IE II
4.1 Hz (E)
2.5 Hz (I)
5

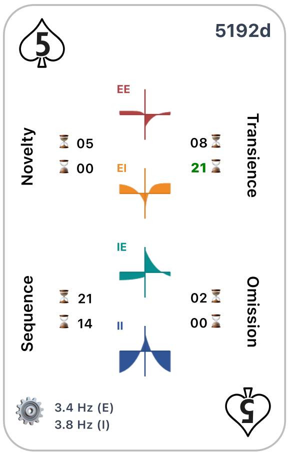
5
5192d
Novelty 05 00
Transience 08 21
Sequence 21 14
Omission 02 00
EE EI IE II
3.4 Hz (E)
3.8 Hz (I)
5

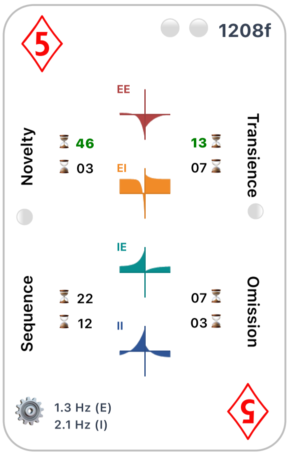
5
1208f
Novelty 46 03
Transience 13 07
Sequence 22 12
Omission 07 03
EE EI IE II
1.3 Hz (E)
2.1 Hz (I)
5

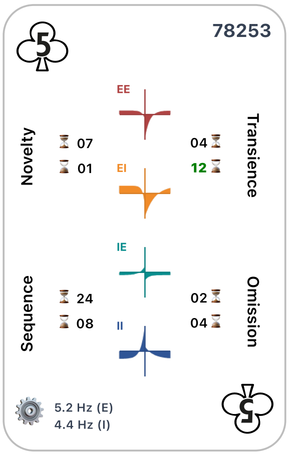
5
78253
Novelty 07 01
Transience 04 12
Sequence 24 08
Omission 02 04
EE EI IE II
5.2 Hz (E)
4.4 Hz (I)
5

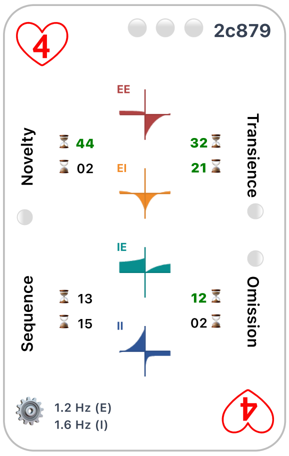
4
2c879
Novelty 44 02
Transience 32 21
Sequence 13 15
Omission 12 02
EE EI IE II
1.2 Hz (E)
1.6 Hz (I)
4

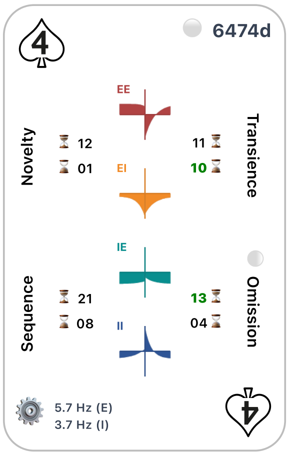
4
6474d
Novelty 12 01
Transience 11 10
Sequence 21 08
Omission 13 04
EE EI IE II
5.7 Hz (E)
3.7 Hz (I)
4

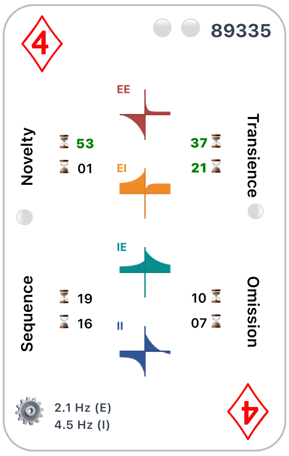
4
89335
Novelty 53 01
Transience 37 21
Sequence 19 16
Omission 10 07
EE EI IE II
2.1 Hz (E)
4.5 Hz (I)
4

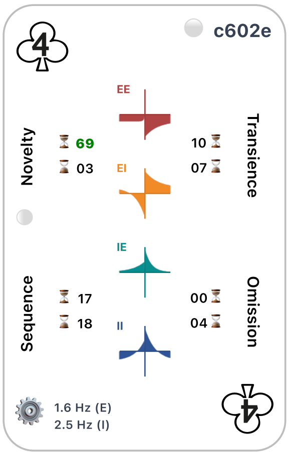
4
c602e
Novelty 69 03
Transience 10 07
Sequence 17 18
Omission 00 04
EE EI IE II
1.6 Hz (E)
2.5 Hz (I)
4

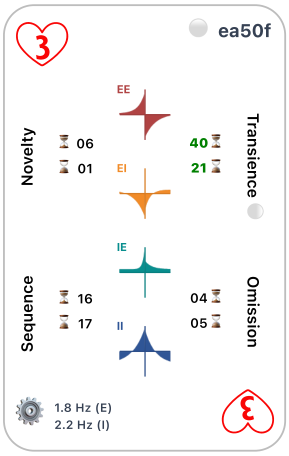
3
ea50f
Novelty 06 01
Transience 40 21
Sequence 16 17
Omission 04 05
EE EI IE II
1.8 Hz (E)
2.2 Hz (I)
3

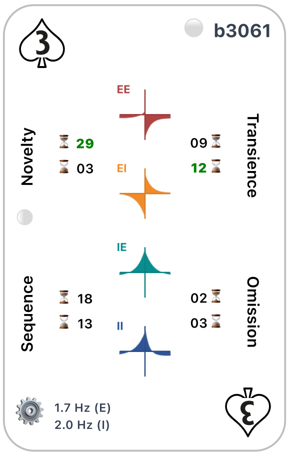
3
b3061
Novelty 29 03
Transience 09 12
Sequence 18 13
Omission 02 03
EE EI IE II
1.7 Hz (E)
2.0 Hz (I)
3

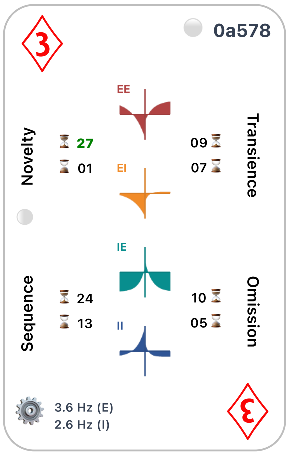
3
0a578
Novelty 27 01
Transience 09 07
Sequence 24 13
Omission 10 05
EE EI IE II
3.6 Hz (E)
2.6 Hz (I)
3

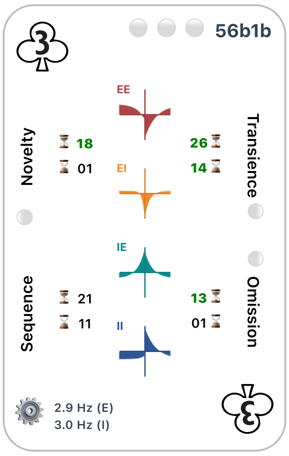
3
56b1b
Novelty 18 01
Transience 26 14
Sequence 21 11
Omission 13 01
EE EI IE II
2.9 Hz (E)
3.0 Hz (I)
3

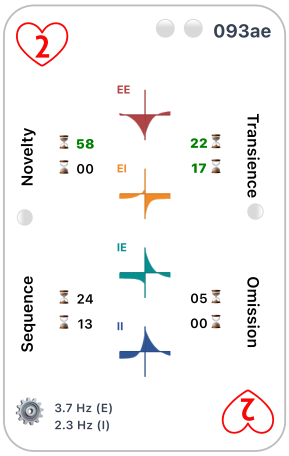
093ae
EE
Novelty
58
00
22
17
Transience
EI
IE
Sequence
24
13
05
00
Omission
II
3.7 Hz (E)
2.3 Hz (I)

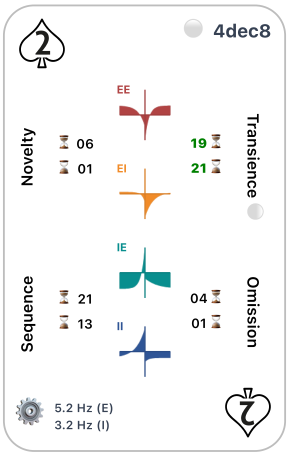
4dec8
EE
Novelty
06
01
19
21
Transience
EI
IE
Sequence
21
13
04
01
Omission
II
5.2 Hz (E)
3.2 Hz (I)

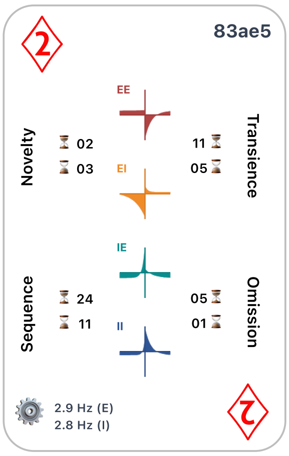
83ae5
EE
Novelty
02
03
11
05
Transience
EI
IE
Sequence
24
11
05
01
Omission
II
2.9 Hz (E)
2.8 Hz (I)

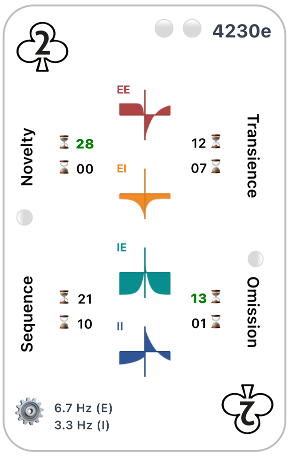
4230e
EE
Novelty
28
00
12
07
Transience
EI
IE
Sequence
21
10
13
01
Omission
II
6.7 Hz (E)
3.3 Hz (I)

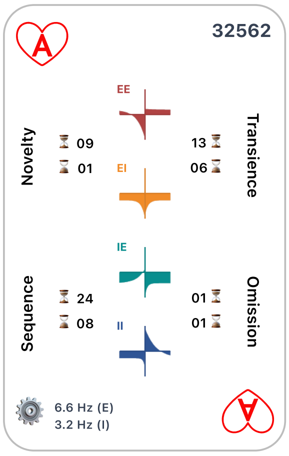
32562
EE
Novelty
09
01
13
06
Transience
EI
IE
Sequence
24
08
01
01
Omission
II
6.6 Hz (E)
3.2 Hz (I)

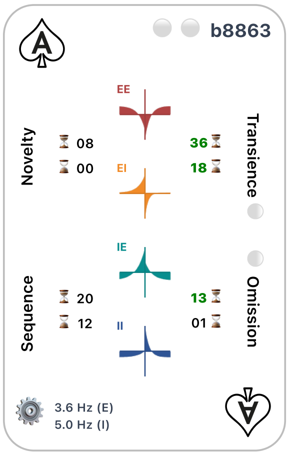
b8863
EE
Novelty
08
00
36
18
Transience
EI
IE
Sequence
20
12
13
01
Omission
II
3.6 Hz (E)
5.0 Hz (I)

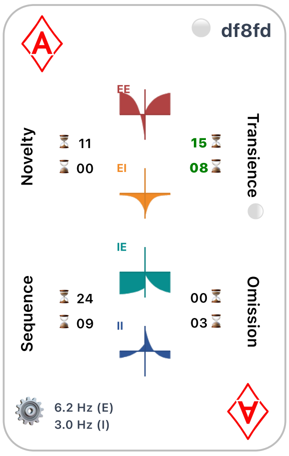
df8fd
EE
Novelty
11
00
15
08
Transience
EI
IE
Sequence
24
09
00
03
Omission
II
6.2 Hz (E)
3.0 Hz (I)

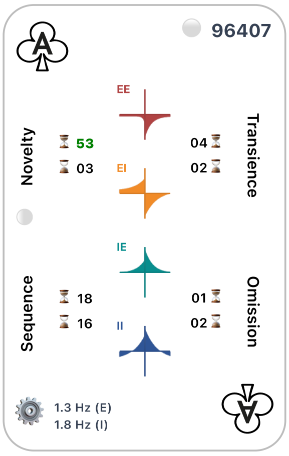
96407
EE
Novelty
53
03
04
02
Transience
EI
IE
Sequence
18
16
01
02
Omission
II
1.3 Hz (E)
1.8 Hz (I)

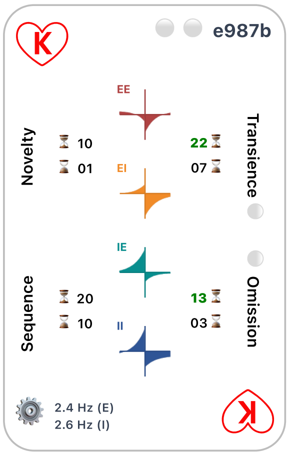
e987b
EE
Novelty
10
01
22
07
Transience
EI
IE
Sequence
20
10
13
03
Omission
II
2.4 Hz (E)
2.6 Hz (I)

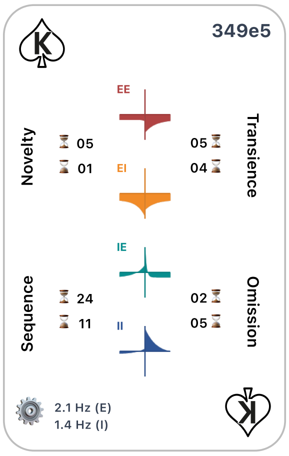
349e5
EE
Novelty
05
01
05
04
Transience
EI
IE
Sequence
24
11
02
05
Omission
II
2.1 Hz (E)
1.4 Hz (I)

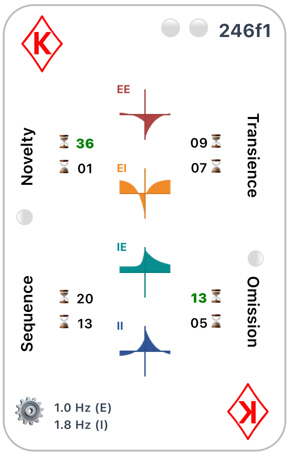
246f1
EE
Novelty
36
01
09
07
Transience
EI
IE
Sequence
20
13
13
05
Omission
II
1.0 Hz (E)
1.8 Hz (I)

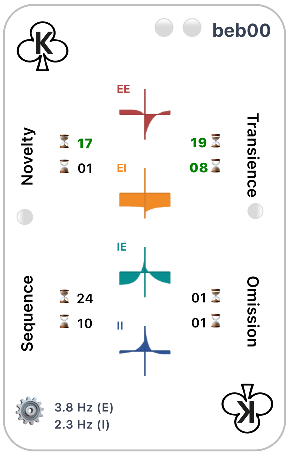
beb00
EE
Novelty
17
01
19
08
Transience
EI
IE
Sequence
24
10
01
01
Omission
II
3.8 Hz (E)
2.3 Hz (I)

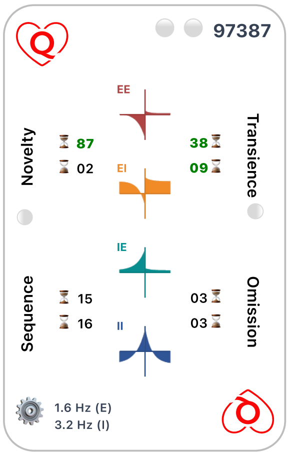
97387
EE
Novelty
87
02
38
09
Transience
EI
IE
Sequence
15
16
03
03
Omission
II
1.6 Hz (E)
3.2 Hz (I)

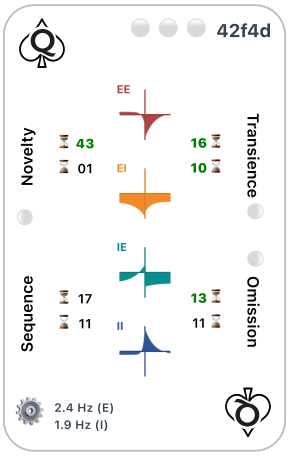
42f4d
EE
Novelty
43
01
16
10
Transience
EI
IE
Sequence
17
11
13
11
Omission
II
2.4 Hz (E)
1.9 Hz (I)

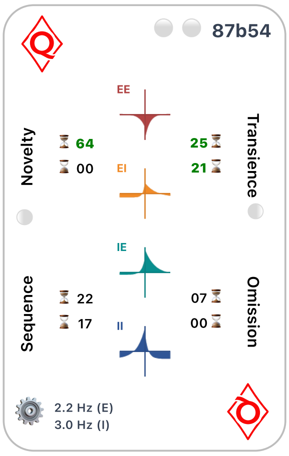
87b54
EE
Novelty
64
00
25
21
Transience
EI
IE
Sequence
22
17
07
00
Omission
II
2.2 Hz (E)
3.0 Hz (I)

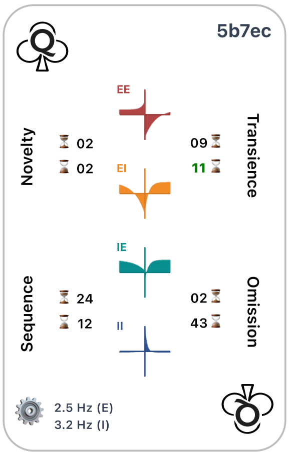
5b7ec
EE
Novelty
02
02
09
11
Transience
EI
IE
Sequence
24
12
02
43
Omission
II
2.5 Hz (E)
3.2 Hz (I)

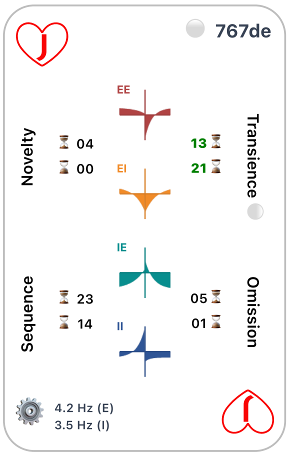
767de
Novelty
04
00
Transience
13
21
EE
EI
IE
II
Sequence
23
14
Omission
05
01
4.2 Hz (E)
3.5 Hz (I)
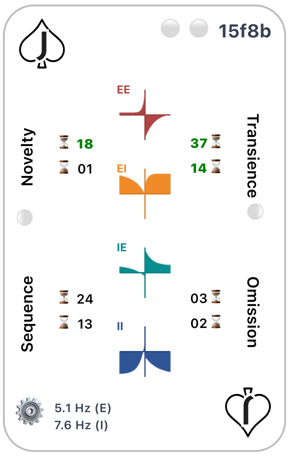
15f8b
Novelty
18
01
Transience
37
14
EE
EI
IE
II
Sequence
24
13
Omission
03
02
5.1 Hz (E)
7.6 Hz (I)
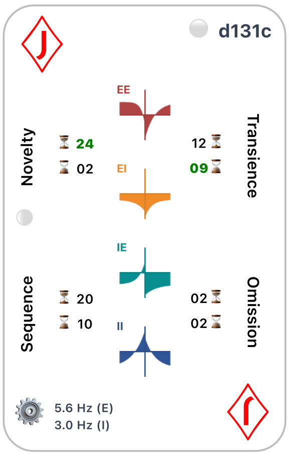
d131c
Novelty
24
02
Transience
12
09
EE
EI
IE
II
Sequence
20
10
Omission
02
02
5.6 Hz (E)
3.0 Hz (I)
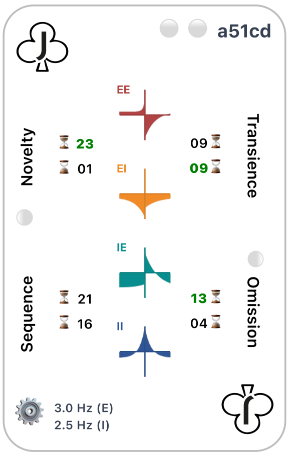
a51cd
Novelty
23
01
Transience
09
09
EE
EI
IE
II
Sequence
21
16
Omission
13
04
3.0 Hz (E)
2.5 Hz (I)
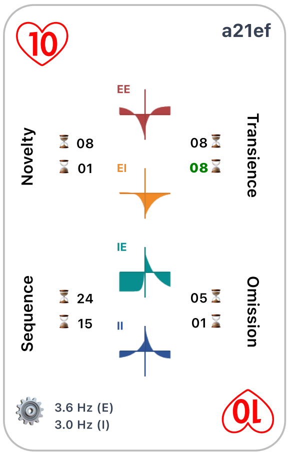
a21ef
Novelty
08
01
Transience
08
08
EE
EI
IE
II
Sequence
24
15
Omission
05
01
3.6 Hz (E)
3.0 Hz (I)
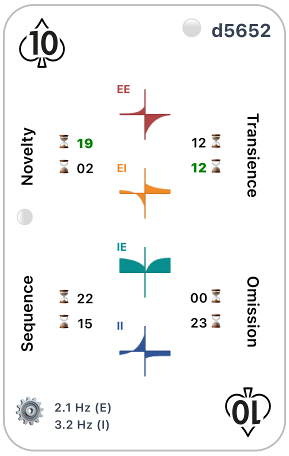
d5652
Novelty
19
02
Transience
12
12
EE
EI
IE
II
Sequence
22
15
Omission
00
23
2.1 Hz (E)
3.2 Hz (I)
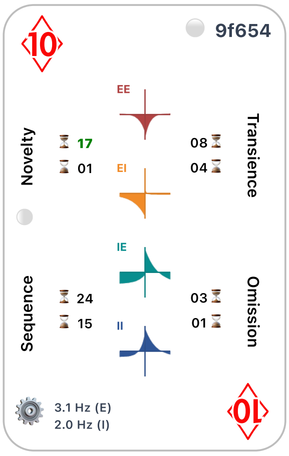
9f654
Novelty
17
01
Transience
08
04
EE
EI
IE
II
Sequence
24
15
Omission
03
01
3.1 Hz (E)
2.0 Hz (I)
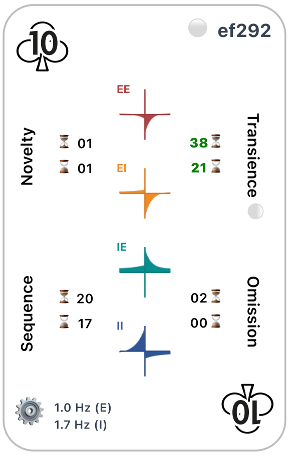
ef292
Novelty
01
01
Transience
38
21
EE
EI
IE
II
Sequence
20
17
Omission
02
00
1.0 Hz (E)
1.7 Hz (I)
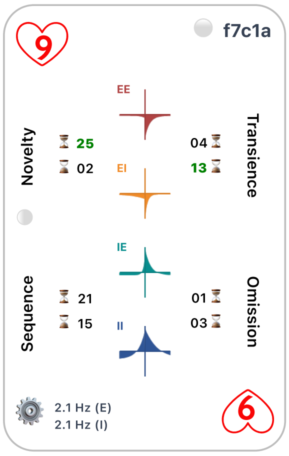
f7c1a
Novelty
25
02
Transience
04
13
EE
EI
IE
II
Sequence
21
15
Omission
01
03
2.1 Hz (E)
2.1 Hz (I)
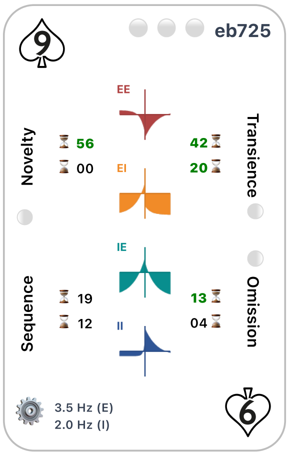
eb725
Novelty
56
00
Transience
42
20
EE
EI
IE
II
Sequence
19
12
Omission
13
04
3.5 Hz (E)
2.0 Hz (I)
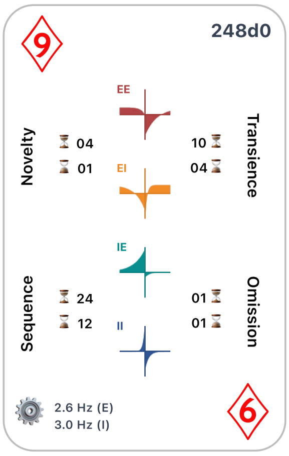
248d0
Novelty
04
01
Transience
10
04
EE
EI
IE
II
Sequence
24
12
Omission
01
01
2.6 Hz (E)
3.0 Hz (I)
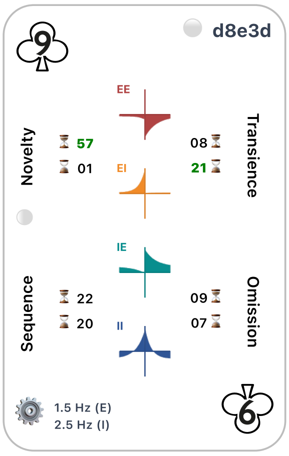
d8e3d
Novelty
57
01
Transience
08
21
EE
EI
IE
II
Sequence
22
20
Omission
09
07
1.5 Hz (E)
2.5 Hz (I)
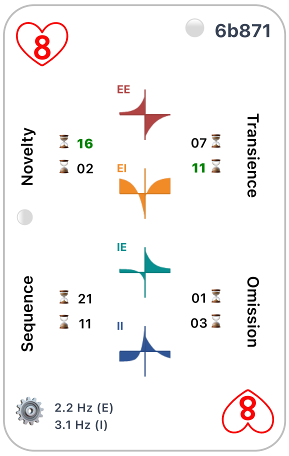
6b871
Novelty
16
02
Transience
07
11
EE
EI
IE
II
Sequence
21
11
Omission
01
03
2.2 Hz (E)
3.1 Hz (I)
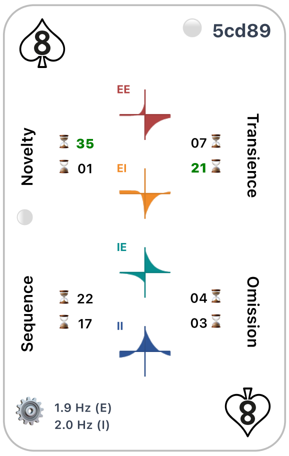
5cd89
Novelty
35
01
Transience
07
21
EE
EI
IE
II
Sequence
22
17
Omission
04
03
1.9 Hz (E)
2.0 Hz (I)
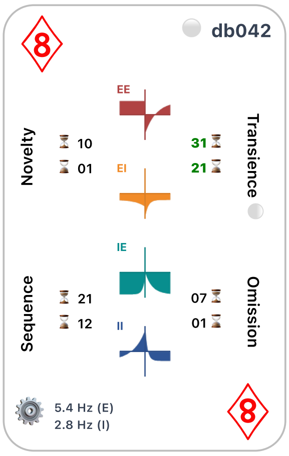
db042
Novelty
10
01
Transience
31
21
EE
EI
IE
II
Sequence
21
12
Omission
07
01
5.4 Hz (E)
2.8 Hz (I)
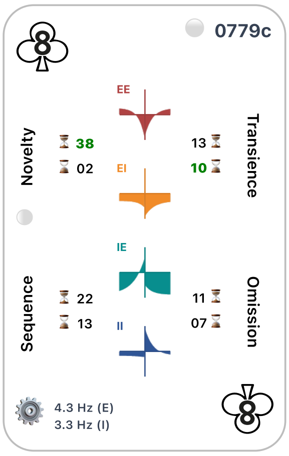
0779c
Novelty
38
02
Transience
13
10
EE
EI
IE
II
Sequence
22
13
Omission
11
07
4.3 Hz (E)
3.3 Hz (I)

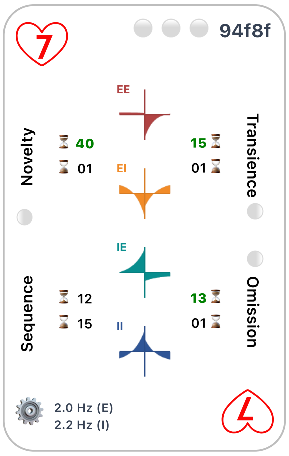
7
94f8f
EE
EI
IE
II
Novelty
40
01
Transience
15
01
Sequence
12
15
Omission
13
01
2.0 Hz (E)
2.2 Hz (I)

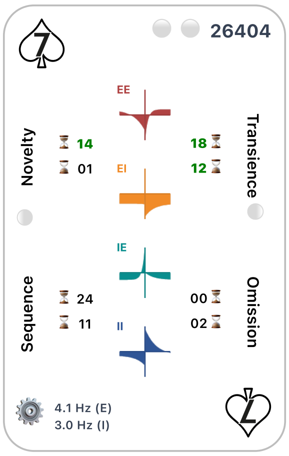
7
26404
EE
EI
IE
II
Novelty
14
01
Transience
18
12
Sequence
24
11
Omission
00
02
4.1 Hz (E)
3.0 Hz (I)

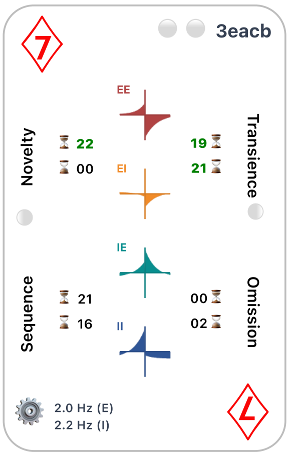
7
3eacb
EE
EI
IE
II
Novelty
22
00
Transience
19
21
Sequence
21
16
Omission
00
02
2.0 Hz (E)
2.2 Hz (I)

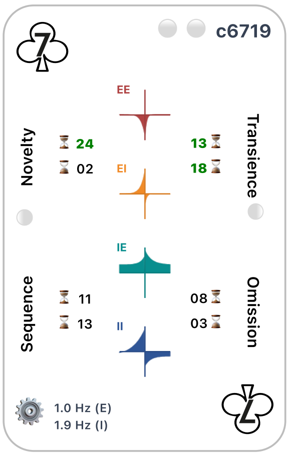
7
c6719
EE
EI
IE
II
Novelty
24
02
Transience
13
18
Sequence
11
13
Omission
08
03
1.0 Hz (E)
1.9 Hz (I)

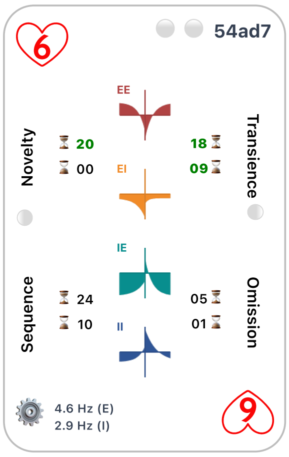
6
54ad7
EE
EI
IE
II
Novelty
20
00
Transience
18
09
Sequence
24
10
Omission
05
01
4.6 Hz (E)
2.9 Hz (I)

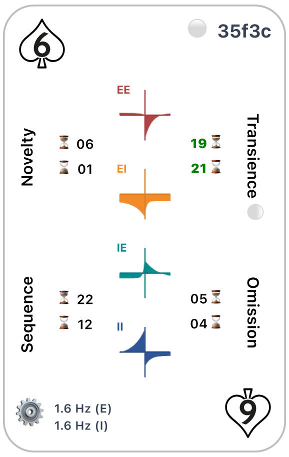
6
35f3c
EE
EI
IE
II
Novelty
06
01
Transience
19
21
Sequence
22
12
Omission
05
04
1.6 Hz (E)
1.6 Hz (I)

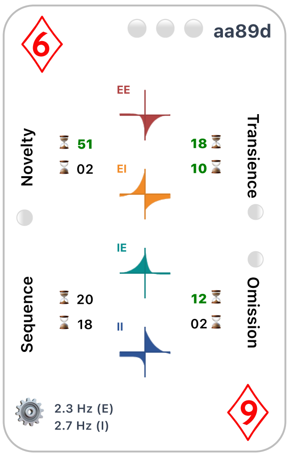
6
aa89d
EE
EI
IE
II
Novelty
51
02
Transience
18
10
Sequence
20
18
Omission
12
02
2.3 Hz (E)
2.7 Hz (I)

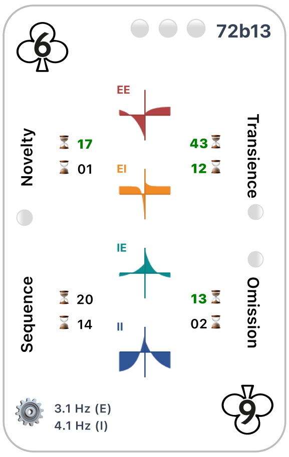
6
72b13
EE
EI
IE
II
Novelty
17
01
Transience
43
12
Sequence
20
14
Omission
13
02
3.1 Hz (E)
4.1 Hz (I)

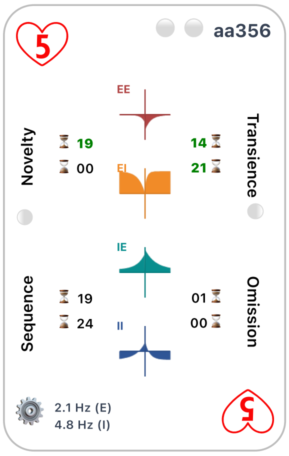
5
aa356
EE
EI
IE
II
Novelty
19
00
Transience
14
21
Sequence
19
24
Omission
01
00
2.1 Hz (E)
4.8 Hz (I)

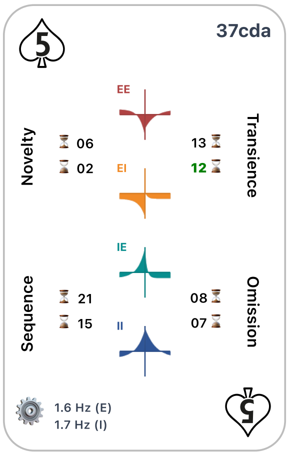
5
37cda
EE
EI
IE
II
Novelty
06
02
Transience
13
12
Sequence
21
15
Omission
08
07
1.6 Hz (E)
1.7 Hz (I)

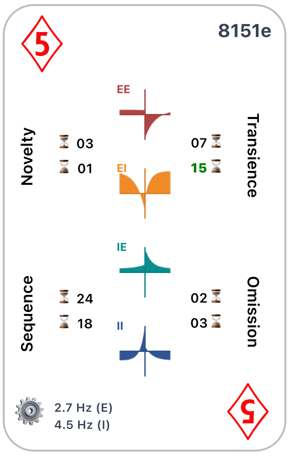
5
8151e
EE
EI
IE
II
Novelty
03
01
Transience
07
15
Sequence
24
18
Omission
02
03
2.7 Hz (E)
4.5 Hz (I)

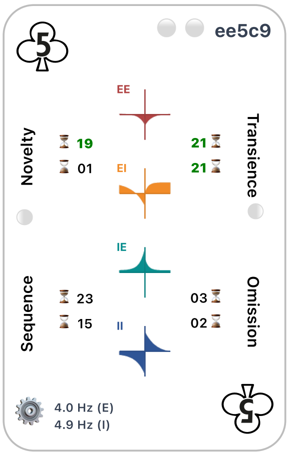
5
ee5c9
EE
EI
IE
II
Novelty
19
01
Transience
21
21
Sequence
23
15
Omission
03
02
4.0 Hz (E)
4.9 Hz (I)

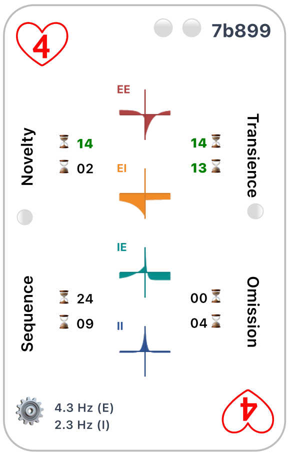
4
7b899
EE
EI
IE
II
Novelty
14
02
Transience
14
13
Sequence
24
09
Omission
00
04
4.3 Hz (E)
2.3 Hz (I)

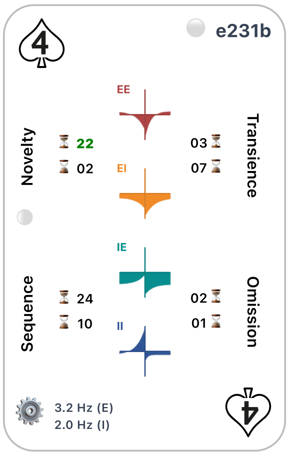
4
e231b
EE
EI
IE
II
Novelty
22
02
Transience
03
07
Sequence
24
10
Omission
02
01
3.2 Hz (E)
2.0 Hz (I)

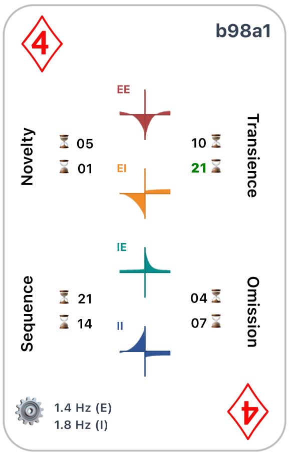
4
b98a1
EE
EI
IE
II
Novelty
05
01
Transience
10
21
Sequence
21
14
Omission
04
07
1.4 Hz (E)
1.8 Hz (I)

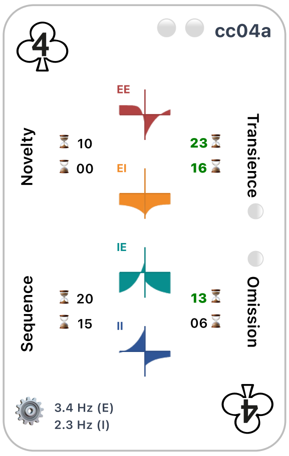
4
cc04a
EE
EI
IE
II
Novelty
10
00
Transience
23
16
Sequence
20
15
Omission
13
06
3.4 Hz (E)
2.3 Hz (I)

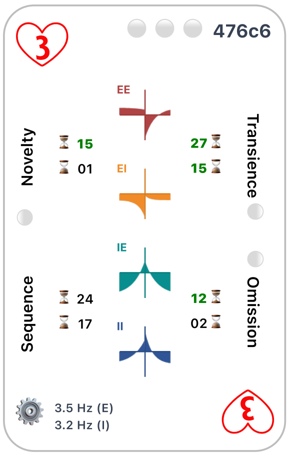
3
476c6
EE
EI
IE
II
Novelty 15 01
Transience 27 15
Sequence 24 17
Omission 12 02
3.5 Hz (E)
3.2 Hz (I)

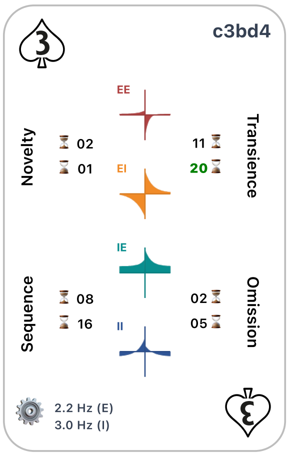
3
c3bd4
EE
EI
IE
II
Novelty 02 01
Transience 11 20
Sequence 08 16
Omission 02 05
2.2 Hz (E)
3.0 Hz (I)

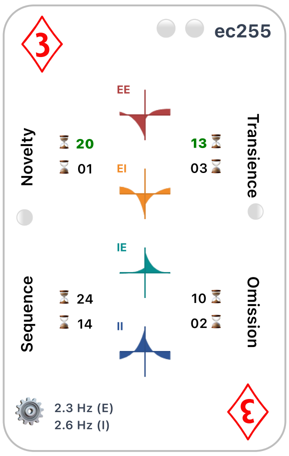
3
ec255
EE
EI
IE
II
Novelty 20 01
Transience 13 03
Sequence 24 14
Omission 10 02
2.3 Hz (E)
2.6 Hz (I)

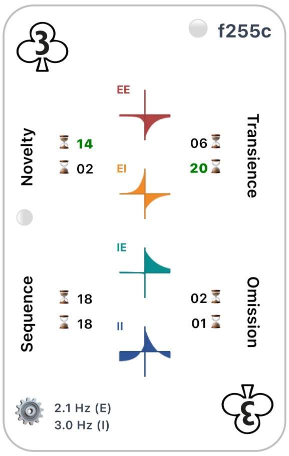
3
f255c
EE
EI
IE
II
Novelty 14 02
Transience 06 20
Sequence 18 18
Omission 02 01
2.1 Hz (E)
3.0 Hz (I)

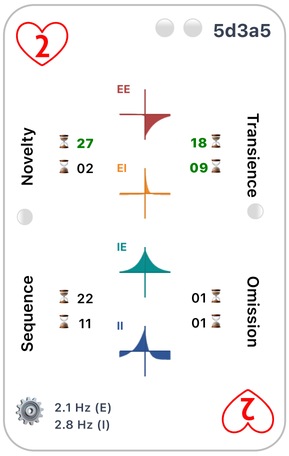
2
5d3a5
EE
EI
IE
II
Novelty 27 02
Transience 18 09
Sequence 22 11
Omission 01 01
2.1 Hz (E)
2.8 Hz (I)

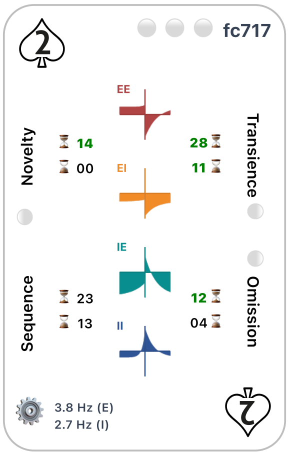
2
fc717
EE
EI
IE
II
Novelty 14 00
Transience 28 11
Sequence 23 13
Omission 12 04
3.8 Hz (E)
2.7 Hz (I)

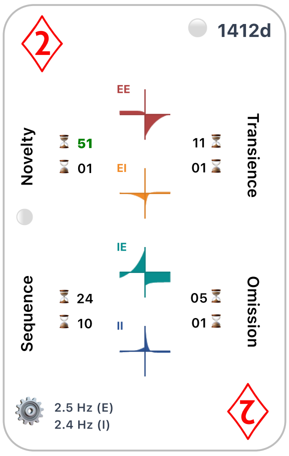
2
1412d
EE
EI
IE
II
Novelty 51 01
Transience 11 01
Sequence 24 10
Omission 05 01
2.5 Hz (E)
2.4 Hz (I)

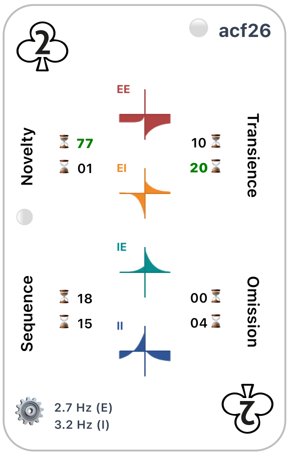
2
acf26
EE
EI
IE
II
Novelty 77 01
Transience 10 20
Sequence 18 15
Omission 00 04
2.7 Hz (E)
3.2 Hz (I)

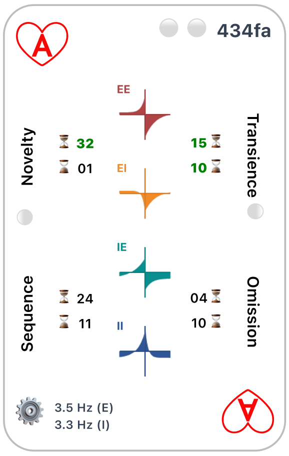
A
434fa
EE
EI
IE
II
Novelty 32 01
Transience 15 10
Sequence 24 11
Omission 04 10
3.5 Hz (E)
3.3 Hz (I)

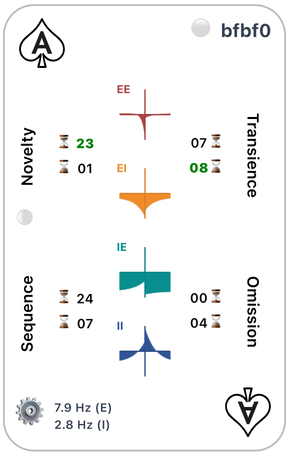
A
bfbf0
EE
EI
IE
II
Novelty 23 01
Transience 07 08
Sequence 24 07
Omission 00 04
7.9 Hz (E)
2.8 Hz (I)

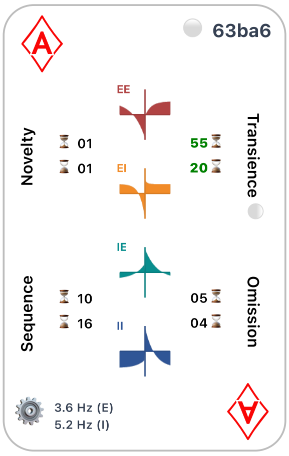
A
63ba6
EE
EI
IE
II
Novelty 01 01
Transience 55 20
Sequence 10 16
Omission 05 04
3.6 Hz (E)
5.2 Hz (I)

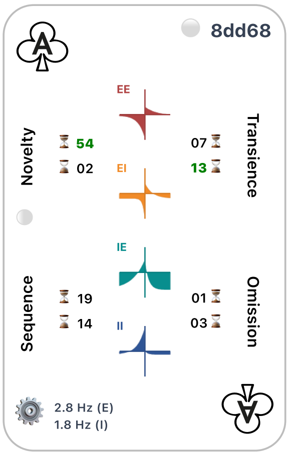
A
8dd68
EE
EI
IE
II
Novelty 54 02
Transience 07 13
Sequence 19 14
Omission 01 03
2.8 Hz (E)
1.8 Hz (I)

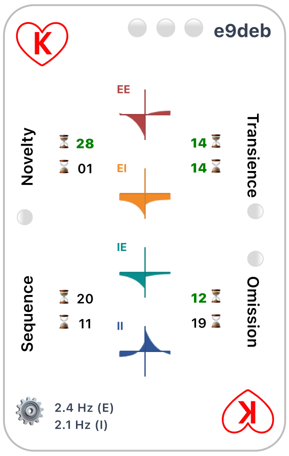
K
e9deb
EE
EI
IE
II
Novelty 28 01
Transience 14 14
Sequence 20 11
Omission 12 19
2.4 Hz (E)
2.1 Hz (I)

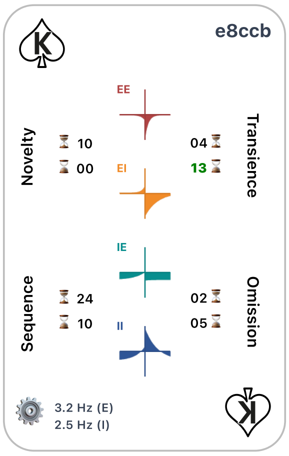
K
e8ccb
EE
EI
IE
II
Novelty 10 00
Transience 04 13
Sequence 24 10
Omission 02 05
3.2 Hz (E)
2.5 Hz (I)

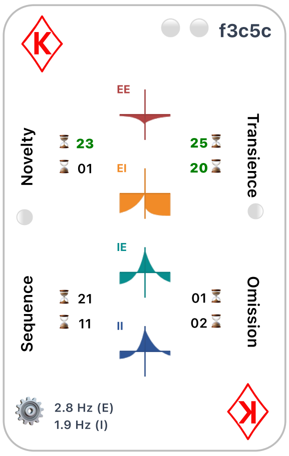
K
f3c5c
EE
EI
IE
II
Novelty 23 01
Transience 25 20
Sequence 21 11
Omission 01 02
2.8 Hz (E)
1.9 Hz (I)

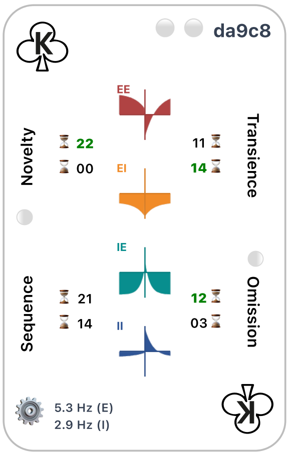
K
da9c8
EE
EI
IE
II
Novelty 22 00
Transience 11 14
Sequence 21 14
Omission 12 03
5.3 Hz (E)
2.9 Hz (I)

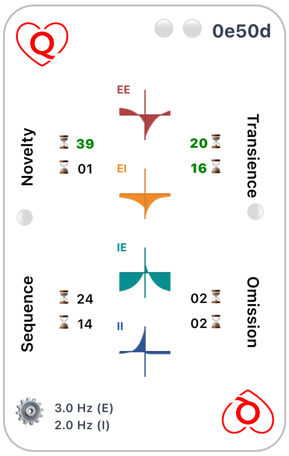
Q
0e50d
EE
EI
IE
II
Novelty
39
01
Transience
20
16
Sequence
24
14
Omission
02
02
3.0 Hz (E)
2.0 Hz (I)

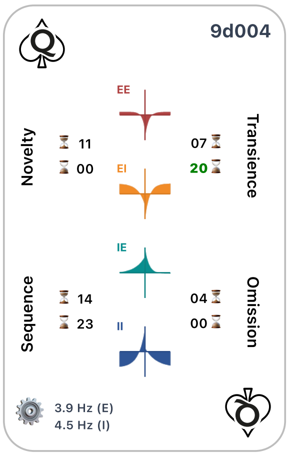
Q
9d004
EE
EI
IE
II
Novelty
11
00
Transience
07
20
Sequence
14
23
Omission
04
00
3.9 Hz (E)
4.5 Hz (I)

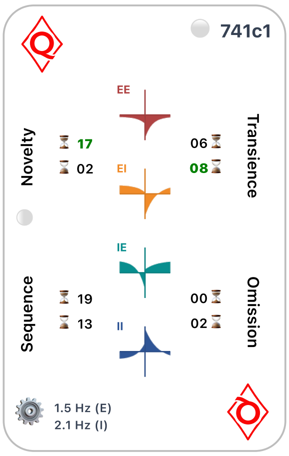
Q
741c1
EE
EI
IE
II
Novelty
17
02
Transience
06
08
Sequence
19
13
Omission
00
02
1.5 Hz (E)
2.1 Hz (I)

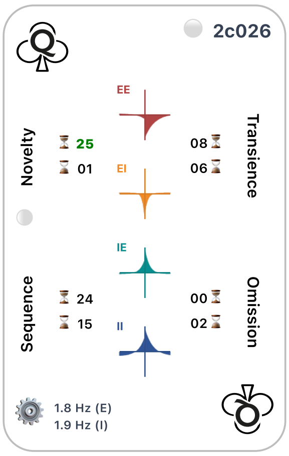
Q
2c026
EE
EI
IE
II
Novelty
25
01
Transience
08
06
Sequence
24
15
Omission
00
02
1.8 Hz (E)
1.9 Hz (I)

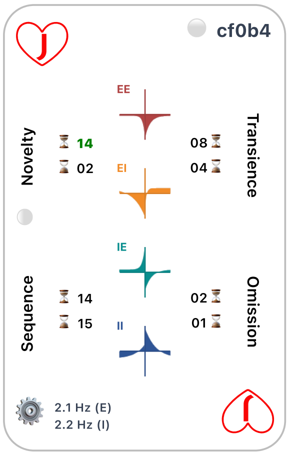
J
cf0b4
EE
EI
IE
II
Novelty
14
02
Transience
08
04
Sequence
14
15
Omission
02
01
2.1 Hz (E)
2.2 Hz (I)

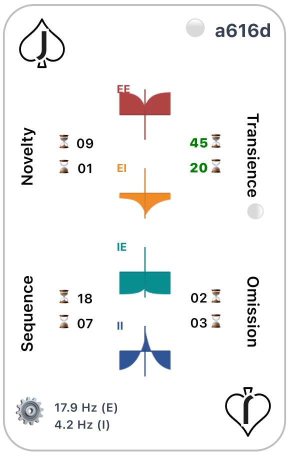
J
a616d
EE
EI
IE
II
Novelty
09
01
Transience
45
20
Sequence
18
07
Omission
02
03
17.9 Hz (E)
4.2 Hz (I)

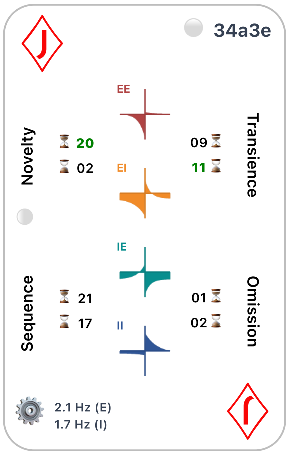
J
34a3e
EE
EI
IE
II
Novelty
20
02
Transience
09
11
Sequence
21
17
Omission
01
02
2.1 Hz (E)
1.7 Hz (I)

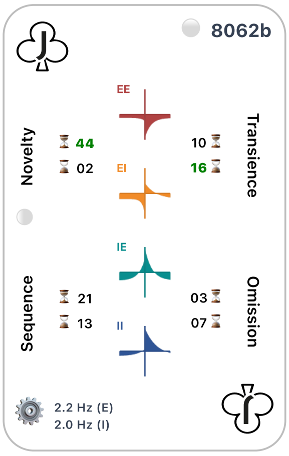
J
8062b
EE
EI
IE
II
Novelty
44
02
Transience
10
16
Sequence
21
13
Omission
03
07
2.2 Hz (E)
2.0 Hz (I)

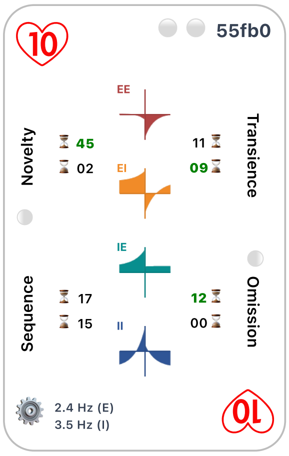
10
55fb0
EE
EI
IE
II
Novelty
45
02
Transience
11
09
Sequence
17
15
Omission
12
00
2.4 Hz (E)
3.5 Hz (I)

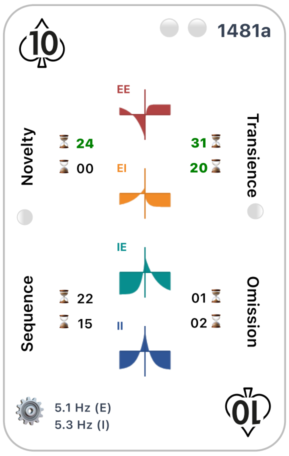
10
1481a
EE
EI
IE
II
Novelty
24
00
Transience
31
20
Sequence
22
15
Omission
01
02
5.1 Hz (E)
5.3 Hz (I)

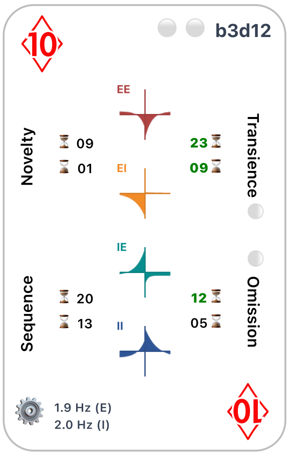
10
b3d12
EE
EI
IE
II
Novelty
09
01
Transience
23
09
Sequence
20
13
Omission
12
05
1.9 Hz (E)
2.0 Hz (I)

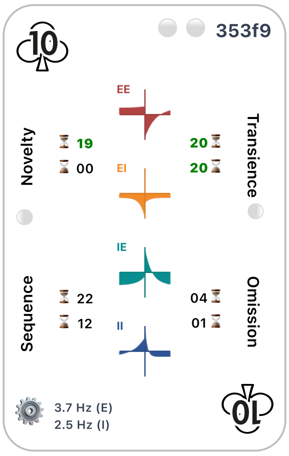
10
353f9
EE
EI
IE
II
Novelty
19
00
Transience
20
20
Sequence
22
12
Omission
04
01
3.7 Hz (E)
2.5 Hz (I)

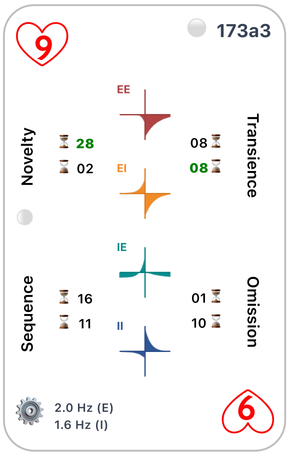
9
173a3
EE
EI
IE
II
Novelty
28
02
Transience
08
08
Sequence
16
11
Omission
01
10
2.0 Hz (E)
1.6 Hz (I)

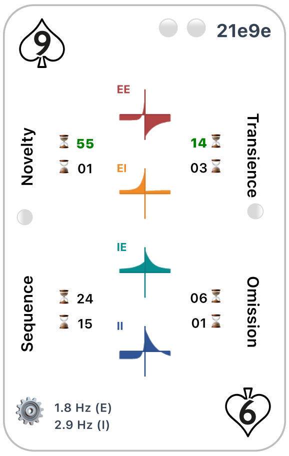
9
21e9e
EE
EI
IE
II
Novelty
55
01
Transience
14
03
Sequence
24
15
Omission
06
01
1.8 Hz (E)
2.9 Hz (I)

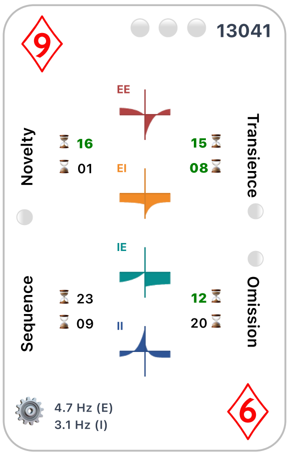
9
13041
EE
EI
IE
II
Novelty
16
01
Transience
15
08
Sequence
23
09
Omission
12
20
4.7 Hz (E)
3.1 Hz (I)

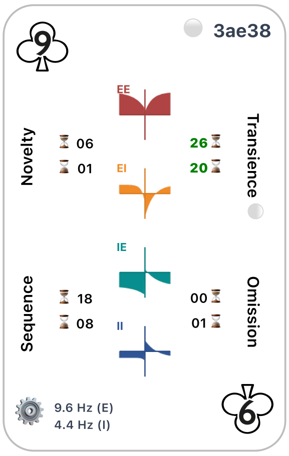
9
3ae38
EE
EI
IE
II
Novelty
06
01
Transience
26
20
Sequence
18
08
Omission
00
01
9.6 Hz (E)
4.4 Hz (I)

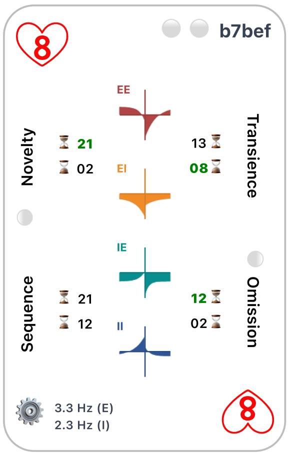
8
b7bef
EE
EI
IE
II
Novelty
21
02
13
08
Transience
Sequence
21
12
12
02
Omission
3.3 Hz (E)
2.3 Hz (I)

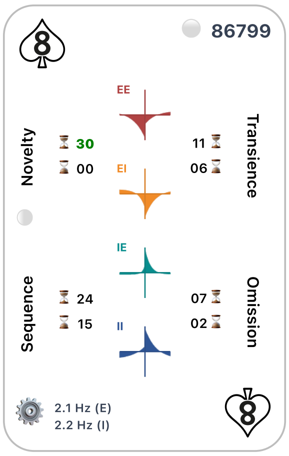
8
86799
EE
EI
IE
II
Novelty
30
00
11
06
Transience
Sequence
24
15
07
02
Omission
2.1 Hz (E)
2.2 Hz (I)

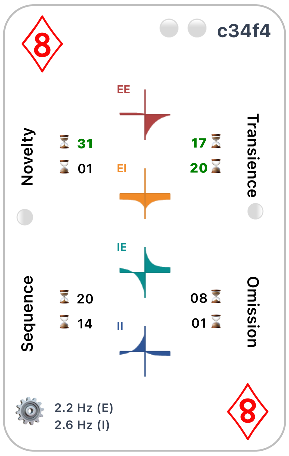
8
c34f4
EE
EI
IE
II
Novelty
31
01
17
20
Transience
Sequence
20
14
08
01
Omission
2.2 Hz (E)
2.6 Hz (I)

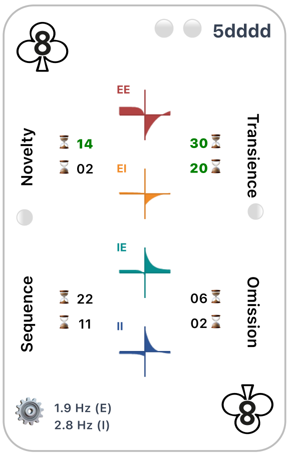
8
5dddd
EE
EI
IE
II
Novelty
14
02
30
20
Transience
Sequence
22
11
06
02
Omission
1.9 Hz (E)
2.8 Hz (I)

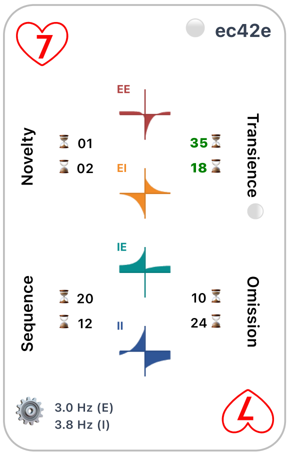
7
ec42e
EE
EI
IE
II
Novelty
01
02
35
18
Transience
Sequence
20
12
10
24
Omission
3.0 Hz (E)
3.8 Hz (I)

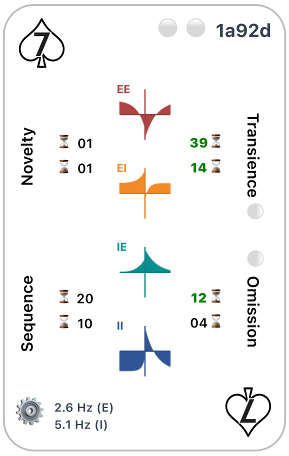
7
1a92d
EE
EI
IE
II
Novelty
01
01
39
14
Transience
Sequence
20
10
12
04
Omission
2.6 Hz (E)
5.1 Hz (I)

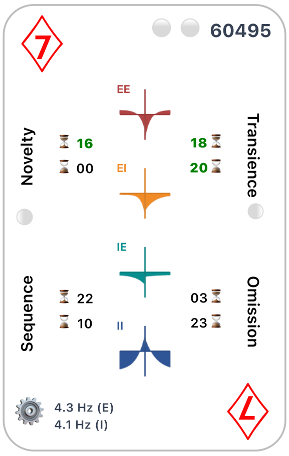
7
60495
EE
EI
IE
II
Novelty
16
00
18
20
Transience
Sequence
22
10
03
23
Omission
4.3 Hz (E)
4.1 Hz (I)

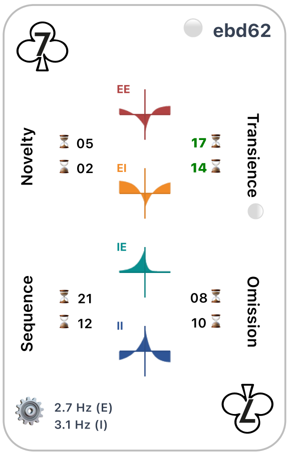
7
ebd62
EE
EI
IE
II
Novelty
05
02
17
14
Transience
Sequence
21
12
08
10
Omission
2.7 Hz (E)
3.1 Hz (I)

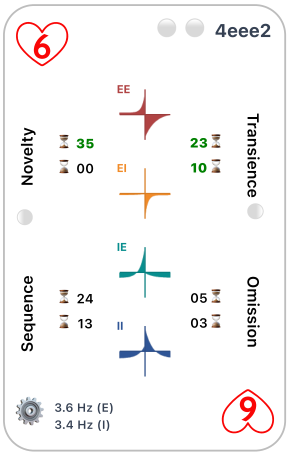
6
4eee2
EE
EI
IE
II
Novelty
35
00
23
10
Transience
Sequence
24
13
05
03
Omission
3.6 Hz (E)
3.4 Hz (I)

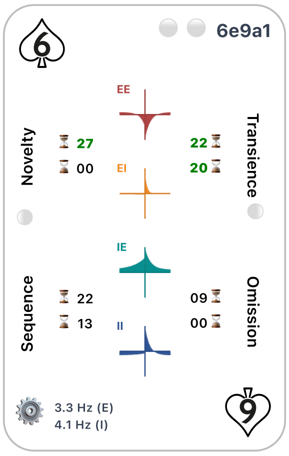
6
6e9a1
EE
EI
IE
II
Novelty
27
00
22
20
Transience
Sequence
22
13
09
00
Omission
3.3 Hz (E)
4.1 Hz (I)

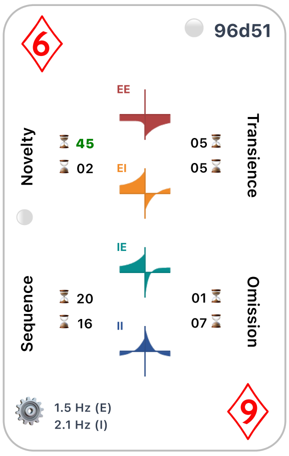
6
96d51
EE
EI
IE
II
Novelty
45
02
05
05
Transience
Sequence
20
16
01
07
Omission
1.5 Hz (E)
2.1 Hz (I)

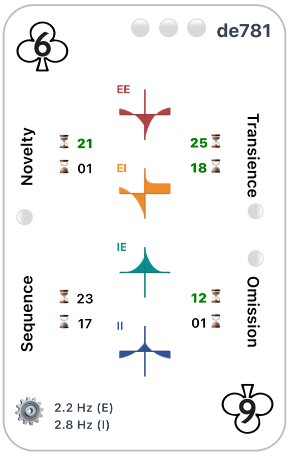
6
de781
EE
EI
IE
II
Novelty
21
01
25
18
Transience
Sequence
23
17
12
01
Omission
2.2 Hz (E)
2.8 Hz (I)

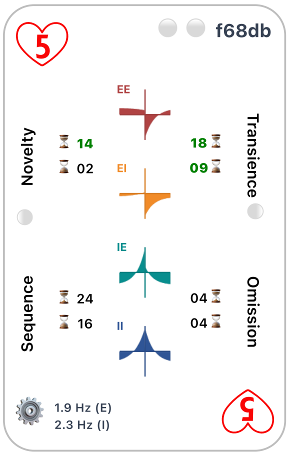
5
f68db
EE
EI
IE
II
Novelty
14
02
18
09
Transience
Sequence
24
16
04
04
Omission
1.9 Hz (E)
2.3 Hz (I)

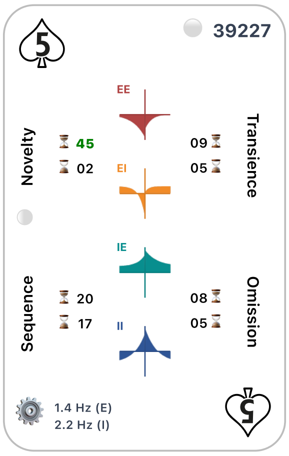
5
39227
EE
EI
IE
II
Novelty
45
02
09
05
Transience
Sequence
20
17
08
05
Omission
1.4 Hz (E)
2.2 Hz (I)

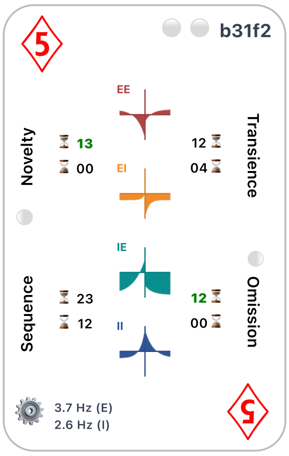
5
b31f2
EE
EI
IE
II
Novelty
13
00
12
04
Transience
Sequence
23
12
12
00
Omission
3.7 Hz (E)
2.6 Hz (I)

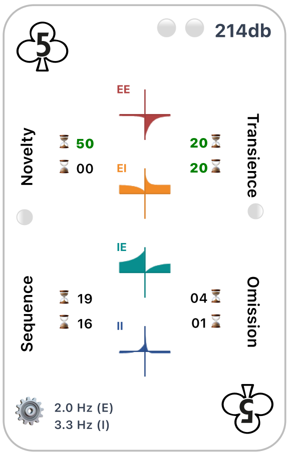
5
214db
EE
EI
IE
II
Novelty
50
00
20
20
Transience
Sequence
19
16
04
01
Omission
2.0 Hz (E)
3.3 Hz (I)

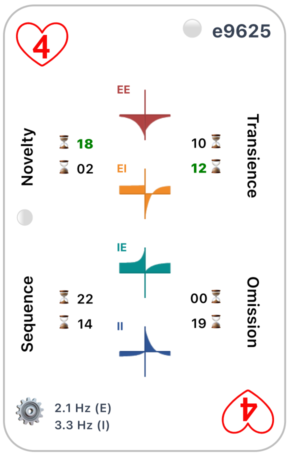
4
e9625
EE
Novelty
18
02
10
12
Transience
EI
IE
Sequence
22
14
00
19
Omission
II
2.1 Hz (E)
3.3 Hz (I)

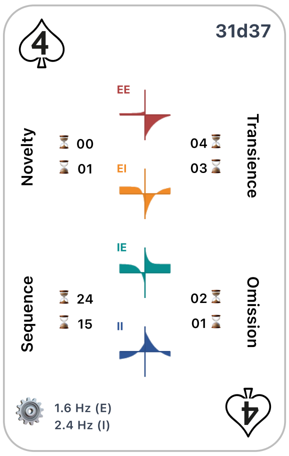
4
31d37
EE
Novelty
00
01
04
03
Transience
EI
IE
Sequence
24
15
02
01
Omission
II
1.6 Hz (E)
2.4 Hz (I)

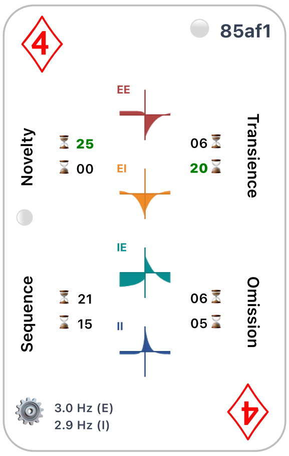
4
85af1
EE
Novelty
25
00
06
20
Transience
EI
IE
Sequence
21
15
06
05
Omission
II
3.0 Hz (E)
2.9 Hz (I)

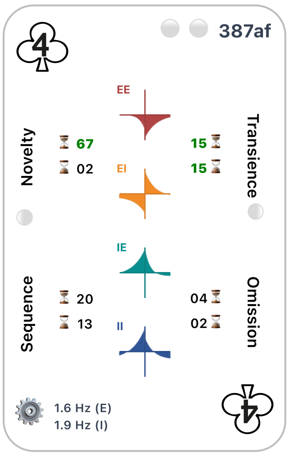
4
387af
EE
Novelty
67
02
15
15
Transience
EI
IE
Sequence
20
13
04
02
Omission
II
1.6 Hz (E)
1.9 Hz (I)

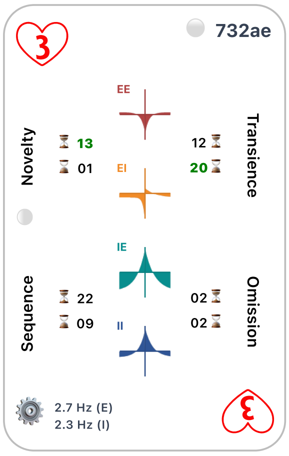
3
732ae
EE
Novelty
13
01
12
20
Transience
EI
IE
Sequence
22
09
02
02
Omission
II
2.7 Hz (E)
2.3 Hz (I)

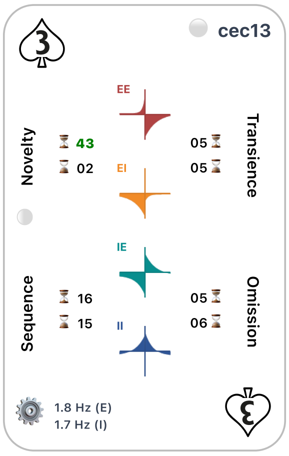
3
cec13
EE
Novelty
43
02
05
05
Transience
EI
IE
Sequence
16
15
05
06
Omission
II
1.8 Hz (E)
1.7 Hz (I)

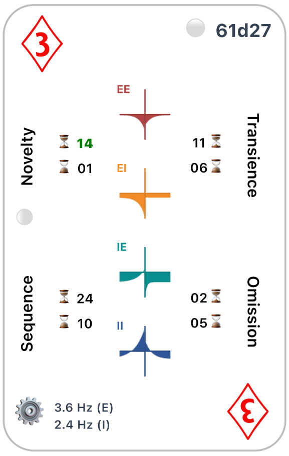
3
61d27
EE
Novelty
14
01
11
06
Transience
EI
IE
Sequence
24
10
02
05
Omission
II
3.6 Hz (E)
2.4 Hz (I)

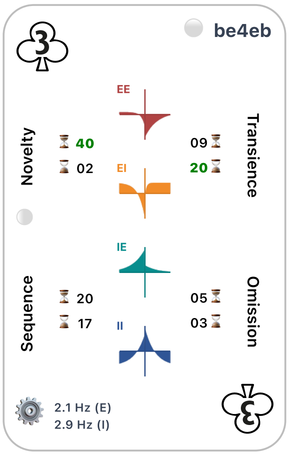
3
be4eb
EE
Novelty
40
02
09
20
Transience
EI
IE
Sequence
20
17
05
03
Omission
II
2.1 Hz (E)
2.9 Hz (I)

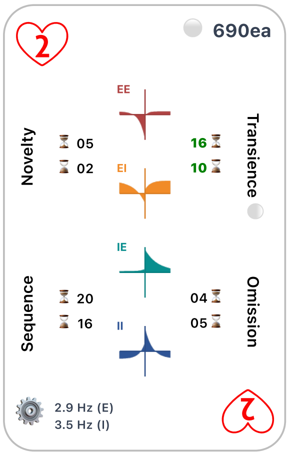
2
690ea
EE
Novelty
05
02
16
10
Transience
EI
IE
Sequence
20
16
04
05
Omission
II
2.9 Hz (E)
3.5 Hz (I)

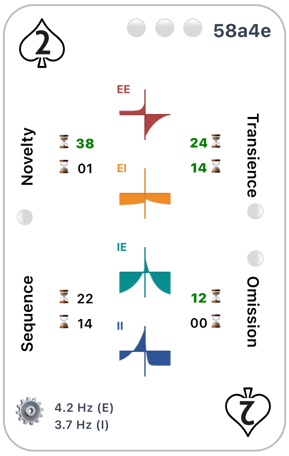
2
58a4e
EE
Novelty
38
01
24
14
Transience
EI
IE
Sequence
22
14
12
00
Omission
II
4.2 Hz (E)
3.7 Hz (I)

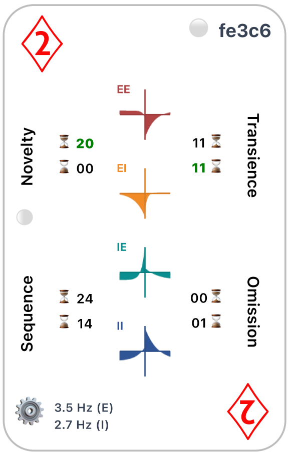
2
fe3c6
EE
Novelty
20
00
11
11
Transience
EI
IE
Sequence
24
14
00
01
Omission
II
3.5 Hz (E)
2.7 Hz (I)

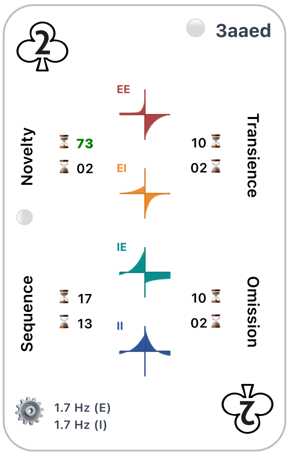
2
3aaed
EE
Novelty
73
02
10
02
Transience
EI
IE
Sequence
17
13
10
02
Omission
II
1.7 Hz (E)
1.7 Hz (I)

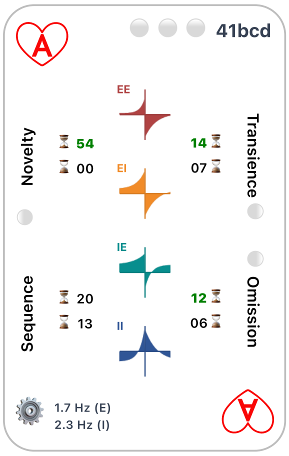
A
41bcd
EE
Novelty
54
00
14
07
Transience
EI
IE
Sequence
20
13
12
06
Omission
II
1.7 Hz (E)
2.3 Hz (I)

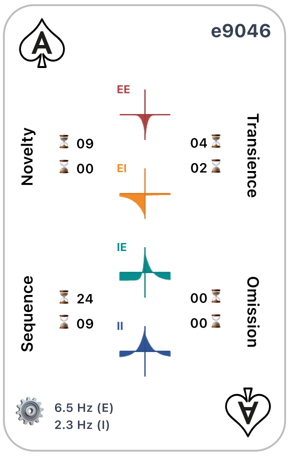
A
e9046
EE
Novelty
09
00
04
02
Transience
EI
IE
Sequence
24
09
00
00
Omission
II
6.5 Hz (E)
2.3 Hz (I)

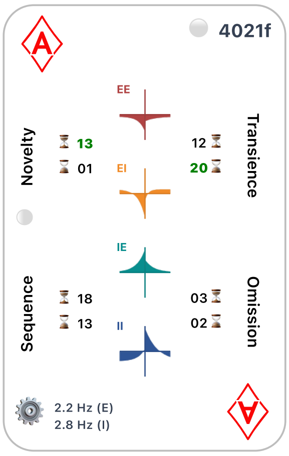
A
4021f
EE
Novelty
13
01
12
20
Transience
EI
IE
Sequence
18
13
03
02
Omission
II
2.2 Hz (E)
2.8 Hz (I)

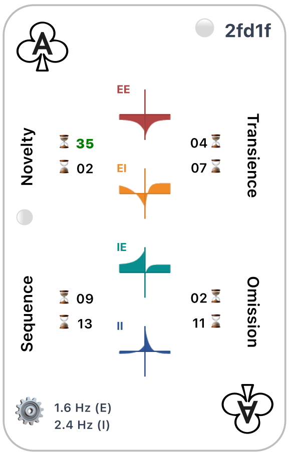
A
2fd1f
EE
Novelty
35
02
04
07
Transience
EI
IE
Sequence
09
13
02
11
Omission
II
1.6 Hz (E)
2.4 Hz (I)

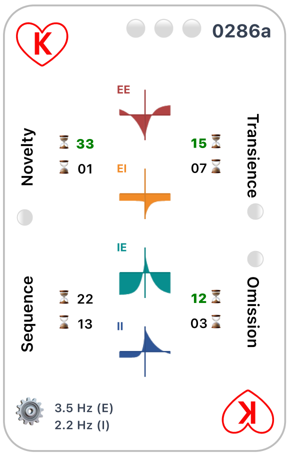
K
0286a
EE
Novelty
33
01
Transience
15
07
EI
IE
Sequence
22
13
Omission
12
03
II
3.5 Hz (E)
2.2 Hz (I)

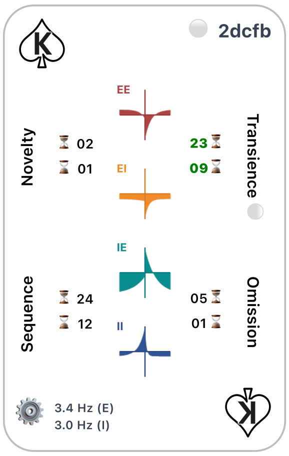
K
2dcfb
EE
Novelty
02
01
Transience
23
09
EI
IE
Sequence
24
12
Omission
05
01
II
3.4 Hz (E)
3.0 Hz (I)

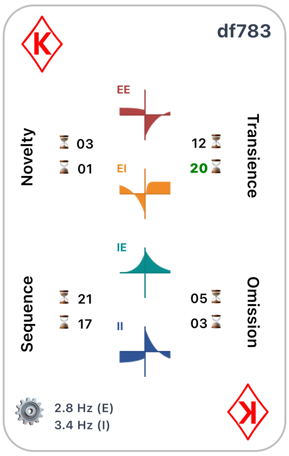
K
df783
EE
Novelty
03
01
Transience
12
20
EI
IE
Sequence
21
17
Omission
05
03
II
2.8 Hz (E)
3.4 Hz (I)

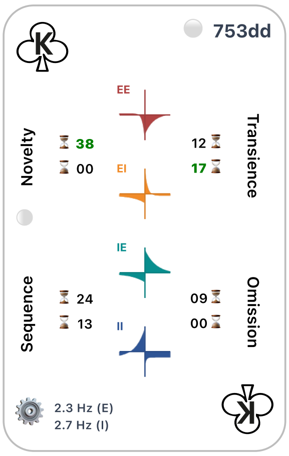
K
753dd
EE
Novelty
38
00
Transience
12
17
EI
IE
Sequence
24
13
Omission
09
00
II
2.3 Hz (E)
2.7 Hz (I)

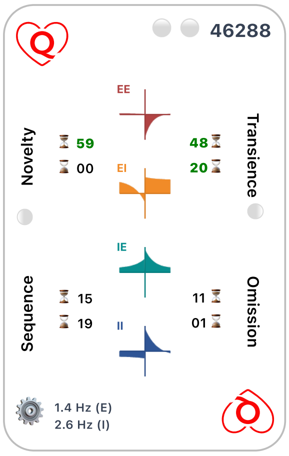
Q
46288
EE
Novelty
59
00
Transience
48
20
EI
IE
Sequence
15
19
Omission
11
01
II
1.4 Hz (E)
2.6 Hz (I)

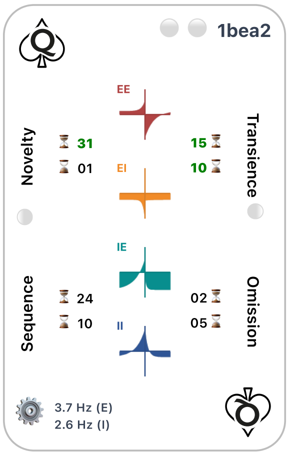
Q
1bea2
EE
Novelty
31
01
Transience
15
10
EI
IE
Sequence
24
10
Omission
02
05
II
3.7 Hz (E)
2.6 Hz (I)

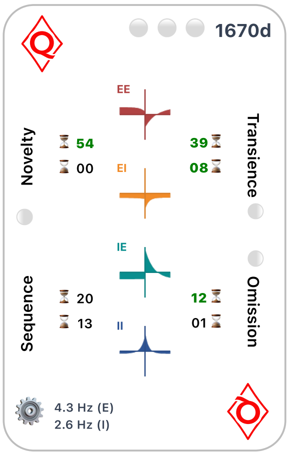
Q
1670d
EE
Novelty
54
00
Transience
39
08
EI
IE
Sequence
20
13
Omission
12
01
II
4.3 Hz (E)
2.6 Hz (I)

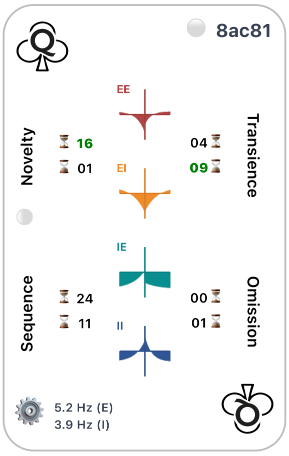
Q
8ac81
EE
Novelty
16
01
Transience
04
09
EI
IE
Sequence
24
11
Omission
00
01
II
5.2 Hz (E)
3.9 Hz (I)

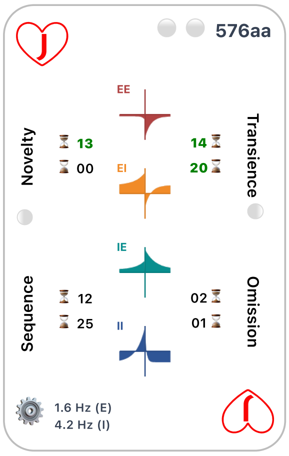
J
576aa
EE
Novelty
13
00
Transience
14
20
EI
IE
Sequence
12
25
Omission
02
01
II
1.6 Hz (E)
4.2 Hz (I)

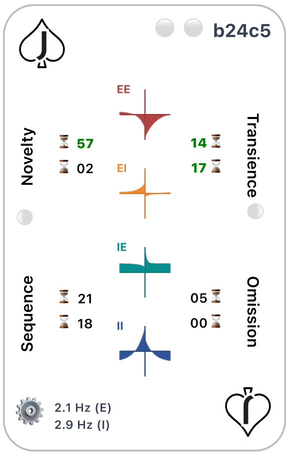
J
b24c5
EE
Novelty
57
02
Transience
14
17
EI
IE
Sequence
21
18
Omission
05
00
II
2.1 Hz (E)
2.9 Hz (I)

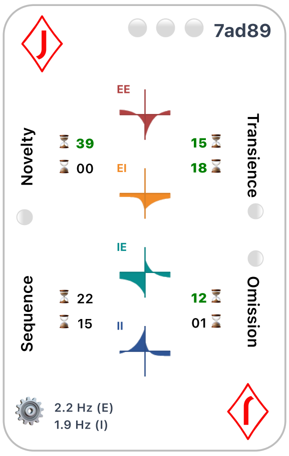
J
7ad89
EE
Novelty
39
00
Transience
15
18
EI
IE
Sequence
22
15
Omission
12
01
II
2.2 Hz (E)
1.9 Hz (I)

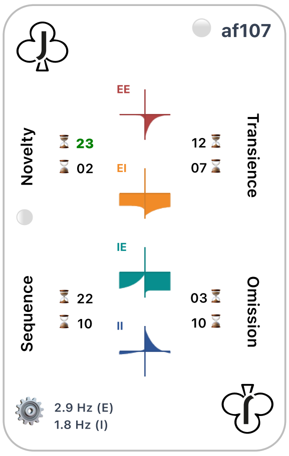
J
af107
EE
Novelty
23
02
Transience
12
07
EI
IE
Sequence
22
10
Omission
03
10
II
2.9 Hz (E)
1.8 Hz (I)

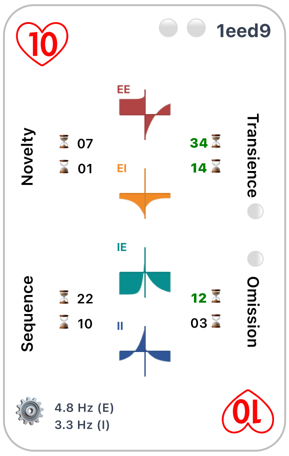
10
1eed9
EE
Novelty
07
01
Transience
34
14
EI
IE
Sequence
22
10
Omission
12
03
II
4.8 Hz (E)
3.3 Hz (I)

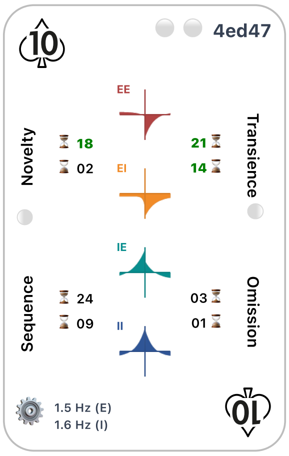
10
4ed47
EE
Novelty
18
02
Transience
21
14
EI
IE
Sequence
24
09
Omission
03
01
II
1.5 Hz (E)
1.6 Hz (I)

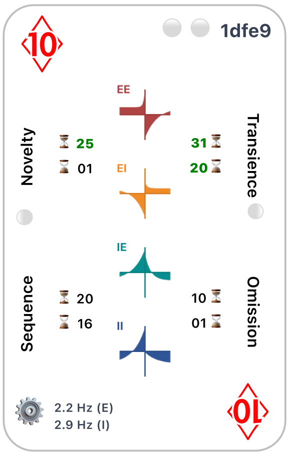
10
1dfe9
EE
Novelty
25
01
Transience
31
20
EI
IE
Sequence
20
16
Omission
10
01
II
2.2 Hz (E)
2.9 Hz (I)

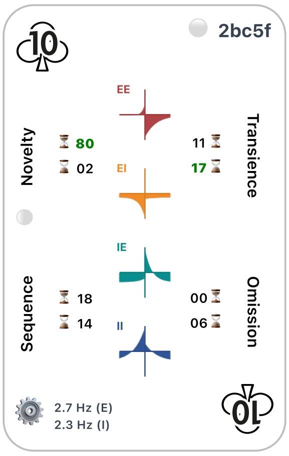
10
2bc5f
EE
Novelty
80
02
Transience
11
17
EI
IE
Sequence
18
14
Omission
00
06
II
2.7 Hz (E)
2.3 Hz (I)

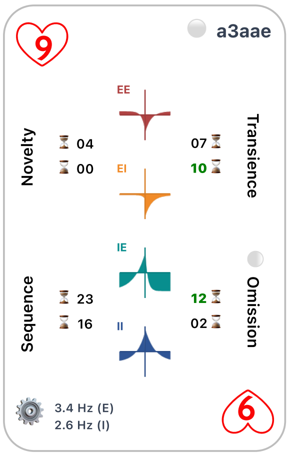
9
a3aae
EE
EI
IE
II
Novelty
04
00
07
10
Transience
Sequence
23
16
12
02
Omission
3.4 Hz (E)
2.6 Hz (I)
6

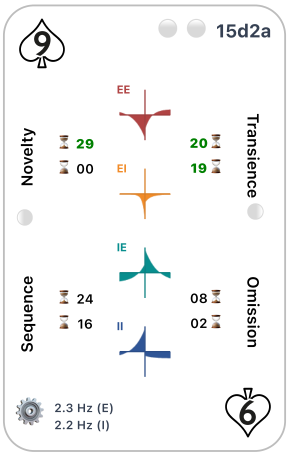
9
15d2a
EE
EI
IE
II
Novelty
29
00
20
19
Transience
Sequence
24
16
08
02
Omission
2.3 Hz (E)
2.2 Hz (I)
6

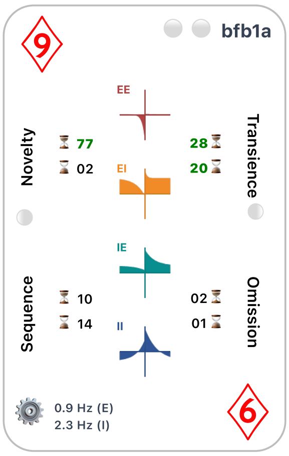
9
bfb1a
EE
EI
IE
II
Novelty
77
02
28
20
Transience
Sequence
10
14
02
01
Omission
0.9 Hz (E)
2.3 Hz (I)
6

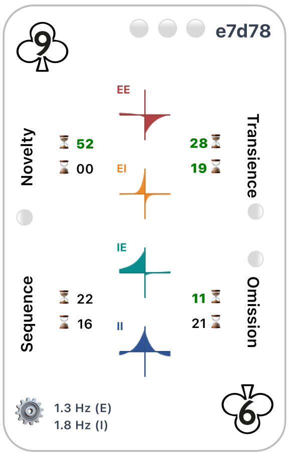
9
e7d78
EE
EI
IE
II
Novelty
52
00
28
19
Transience
Sequence
22
16
11
21
Omission
1.3 Hz (E)
1.8 Hz (I)
6

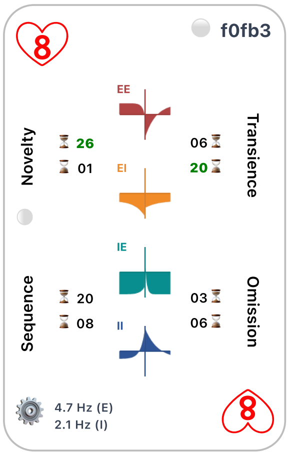
8
f0fb3
EE
EI
IE
II
Novelty
26
01
06
20
Transience
Sequence
20
08
03
06
Omission
4.7 Hz (E)
2.1 Hz (I)
8

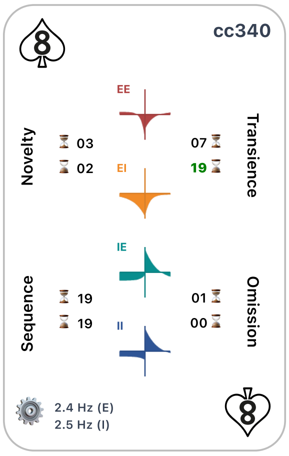
8
cc340
EE
EI
IE
II
Novelty
03
02
07
19
Transience
Sequence
19
19
01
00
Omission
2.4 Hz (E)
2.5 Hz (I)
8

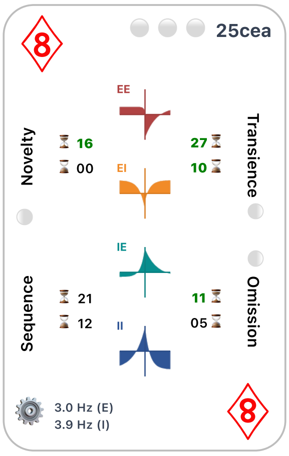
8
25cea
EE
EI
IE
II
Novelty
16
00
27
10
Transience
Sequence
21
12
11
05
Omission
3.0 Hz (E)
3.9 Hz (I)
8

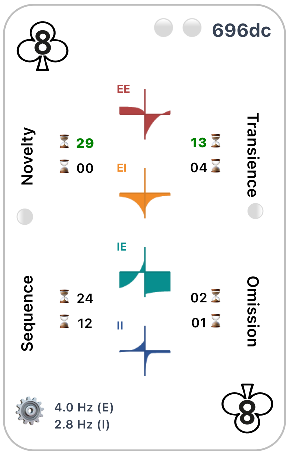
8
696dc
EE
EI
IE
II
Novelty
29
00
13
04
Transience
Sequence
24
12
02
01
Omission
4.0 Hz (E)
2.8 Hz (I)
8

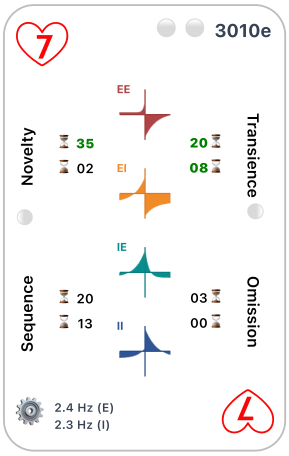
7
3010e
EE
EI
IE
II
Novelty
35
02
20
08
Transience
Sequence
20
13
03
00
Omission
2.4 Hz (E)
2.3 Hz (I)
7

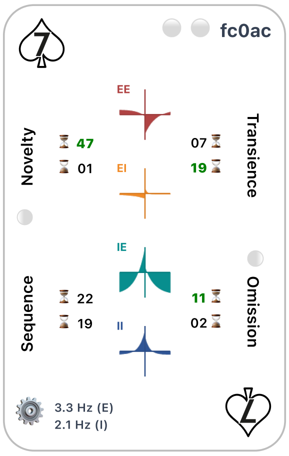
7
fc0ac
EE
EI
IE
II
Novelty
47
01
07
19
Transience
Sequence
22
19
11
02
Omission
3.3 Hz (E)
2.1 Hz (I)
7

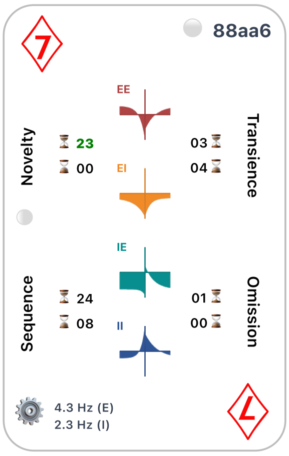
7
88aa6
EE
EI
IE
II
Novelty
23
00
03
04
Transience
Sequence
24
08
01
00
Omission
4.3 Hz (E)
2.3 Hz (I)
7

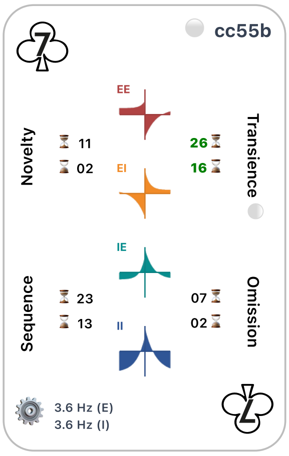
7
cc55b
EE
EI
IE
II
Novelty
11
02
26
16
Transience
Sequence
23
13
07
02
Omission
3.6 Hz (E)
3.6 Hz (I)
7

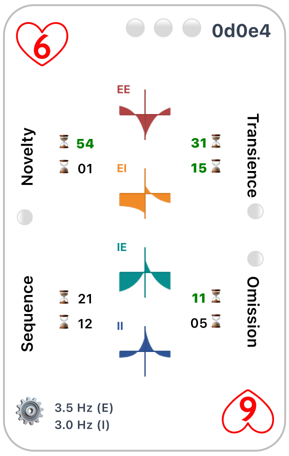
6
0d0e4
EE
EI
IE
II
Novelty
54
01
31
15
Transience
Sequence
21
12
11
05
Omission
3.5 Hz (E)
3.0 Hz (I)
9

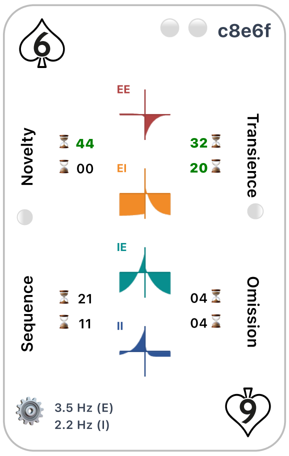
6
c8e6f
EE
EI
IE
II
Novelty
44
00
32
20
Transience
Sequence
21
11
04
04
Omission
3.5 Hz (E)
2.2 Hz (I)
9

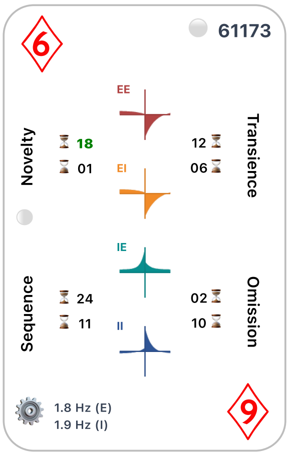
6
61173
EE
EI
IE
II
Novelty
18
01
12
06
Transience
Sequence
24
11
02
10
Omission
1.8 Hz (E)
1.9 Hz (I)
9

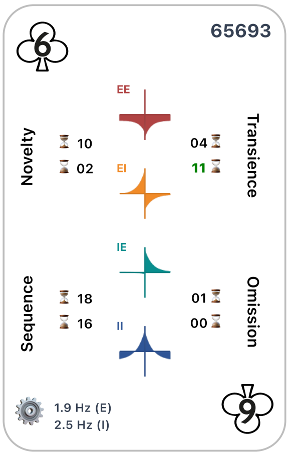
6
65693
EE
EI
IE
II
Novelty
10
02
04
11
Transience
Sequence
18
16
01
00
Omission
1.9 Hz (E)
2.5 Hz (I)
9

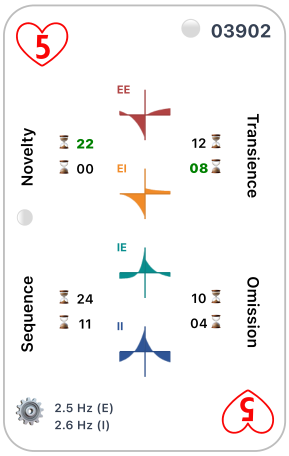
5
03902
EE
EI
IE
II
Novelty
22
00
Transience
12
08
Sequence
24
11
Omission
10
04
2.5 Hz (E)
2.6 Hz (I)

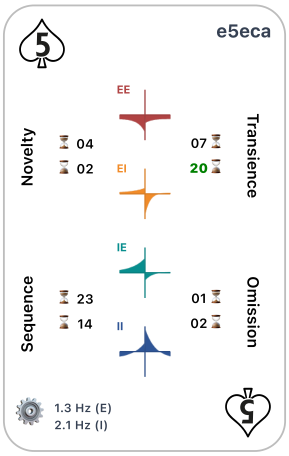
5
e5eca
EE
EI
IE
II
Novelty
04
02
Transience
07
20
Sequence
23
14
Omission
01
02
1.3 Hz (E)
2.1 Hz (I)

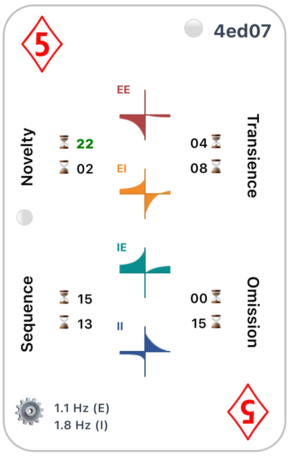
5
4ed07
EE
EI
IE
II
Novelty
22
02
Transience
04
08
Sequence
15
13
Omission
00
15
1.1 Hz (E)
1.8 Hz (I)

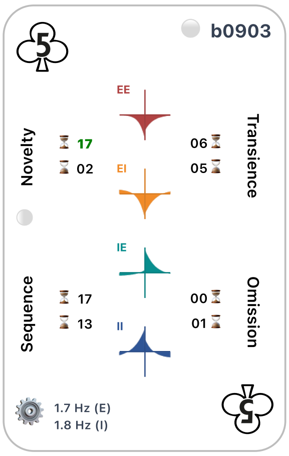
5
b0903
EE
EI
IE
II
Novelty
17
02
Transience
06
05
Sequence
17
13
Omission
00
01
1.7 Hz (E)
1.8 Hz (I)

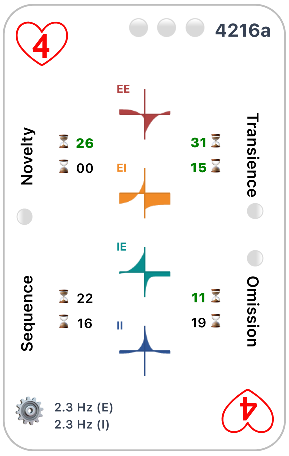
4
4216a
EE
EI
IE
II
Novelty
26
00
Transience
31
15
Sequence
22
16
Omission
11
19
2.3 Hz (E)
2.3 Hz (I)

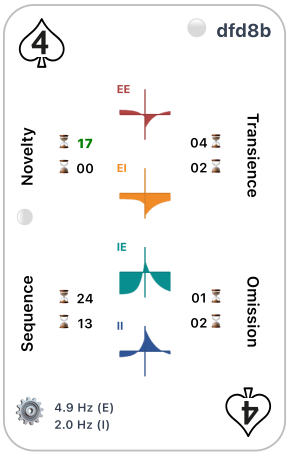
4
dfd8b
EE
EI
IE
II
Novelty
17
00
Transience
04
02
Sequence
24
13
Omission
01
02
4.9 Hz (E)
2.0 Hz (I)

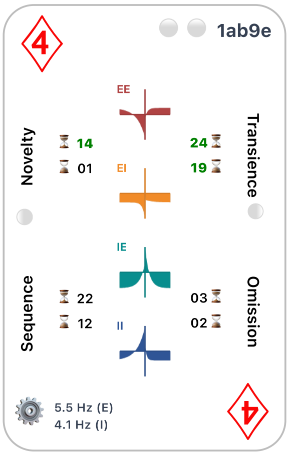
4
1ab9e
EE
EI
IE
II
Novelty
14
01
Transience
24
19
Sequence
22
12
Omission
03
02
5.5 Hz (E)
4.1 Hz (I)

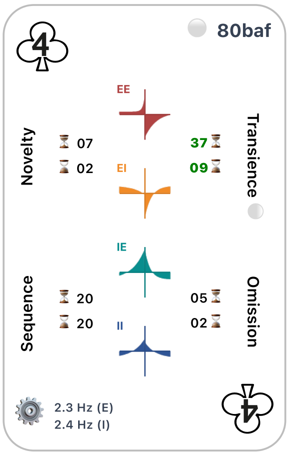
4
80baf
EE
EI
IE
II
Novelty
07
02
Transience
37
09
Sequence
20
20
Omission
05
02
2.3 Hz (E)
2.4 Hz (I)

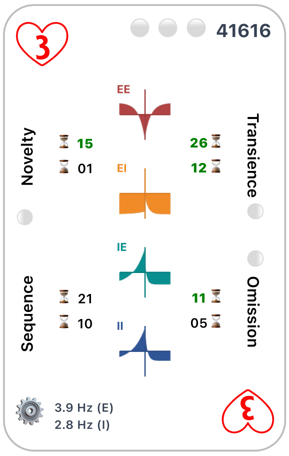
3
41616
EE
EI
IE
II
Novelty
15
01
Transience
26
12
Sequence
21
10
Omission
11
05
3.9 Hz (E)
2.8 Hz (I)

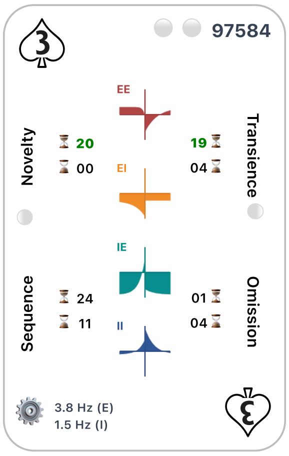
3
97584
EE
EI
IE
II
Novelty
20
00
Transience
19
04
Sequence
24
11
Omission
01
04
3.8 Hz (E)
1.5 Hz (I)

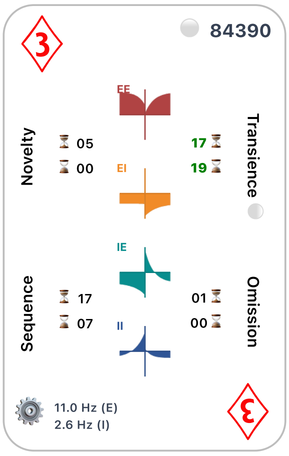
3
84390
EE
EI
IE
II
Novelty
05
00
Transience
17
19
Sequence
17
07
Omission
01
00
11.0 Hz (E)
2.6 Hz (I)

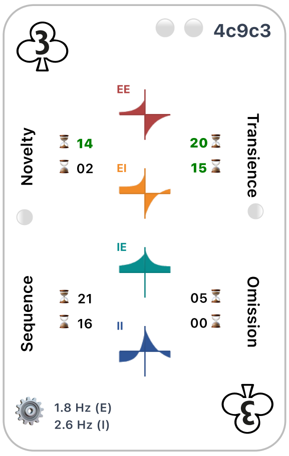
3
4c9c3
EE
EI
IE
II
Novelty
14
02
Transience
20
15
Sequence
21
16
Omission
05
00
1.8 Hz (E)
2.6 Hz (I)

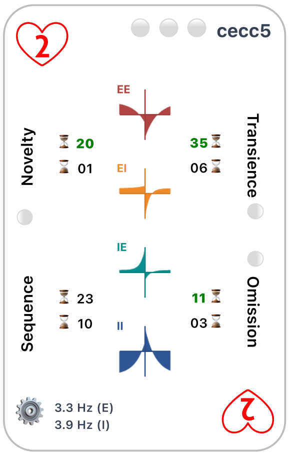
2
cecc5
EE
EI
IE
II
Novelty
20
01
Transience
35
06
Sequence
23
10
Omission
11
03
3.3 Hz (E)
3.9 Hz (I)

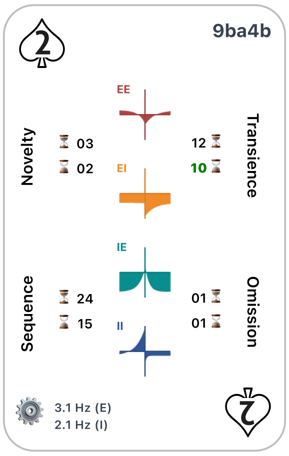
2
9ba4b
EE
EI
IE
II
Novelty
03
02
Transience
12
10
Sequence
24
15
Omission
01
01
3.1 Hz (E)
2.1 Hz (I)

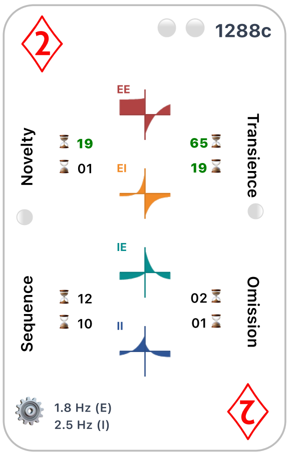
2
1288c
EE
EI
IE
II
Novelty
19
01
Transience
65
19
Sequence
12
10
Omission
02
01
1.8 Hz (E)
2.5 Hz (I)

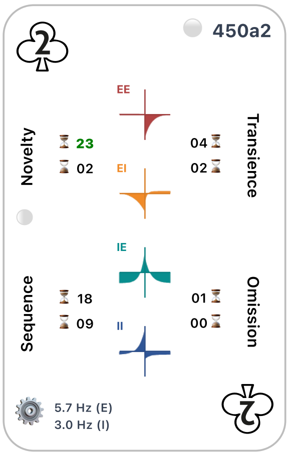
2
450a2
EE
EI
IE
II
Novelty
23
02
Transience
04
02
Sequence
18
09
Omission
01
00
5.7 Hz (E)
3.0 Hz (I)

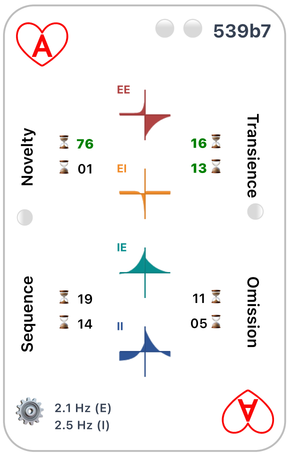
539b7
A
EE
EI
IE
II
Novelty
76
01
Transience
16
13
Sequence
19
14
Omission
11
05
2.1 Hz (E)
2.5 Hz (I)

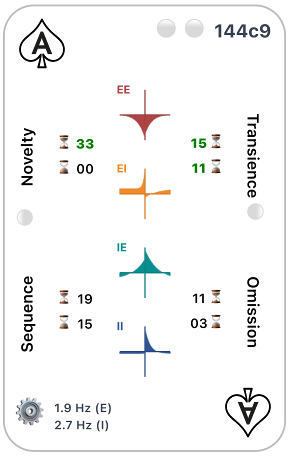
144c9
A
EE
EI
IE
II
Novelty
33
00
Transience
15
11
Sequence
19
15
Omission
11
03
1.9 Hz (E)
2.7 Hz (I)

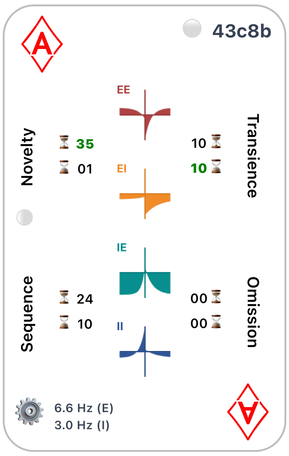
43c8b
A
EE
EI
IE
II
Novelty
35
01
Transience
10
10
Sequence
24
10
Omission
00
00
6.6 Hz (E)
3.0 Hz (I)

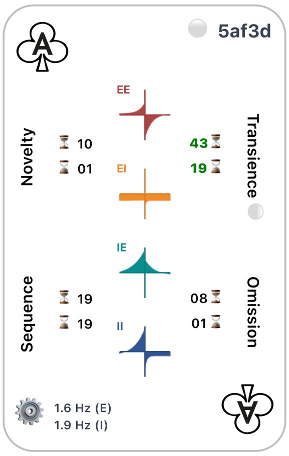
5af3d
A
EE
EI
IE
II
Novelty
10
01
Transience
43
19
Sequence
19
19
Omission
08
01
1.6 Hz (E)
1.9 Hz (I)

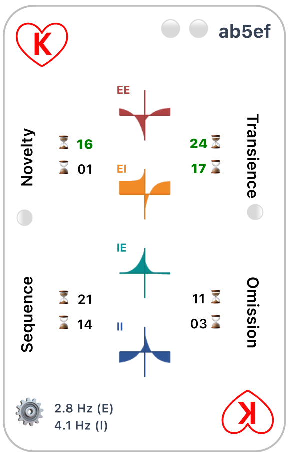
ab5ef
K
EE
EI
IE
II
Novelty
16
01
Transience
24
17
Sequence
21
14
Omission
11
03
2.8 Hz (E)
4.1 Hz (I)

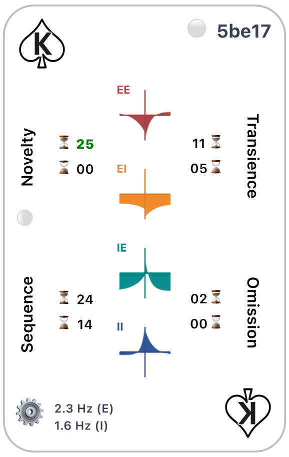
5be17
K
EE
EI
IE
II
Novelty
25
00
Transience
11
05
Sequence
24
14
Omission
02
00
2.3 Hz (E)
1.6 Hz (I)

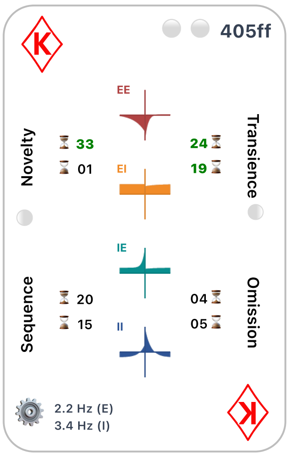
405ff
K
EE
EI
IE
II
Novelty
33
01
Transience
24
19
Sequence
20
15
Omission
04
05
2.2 Hz (E)
3.4 Hz (I)

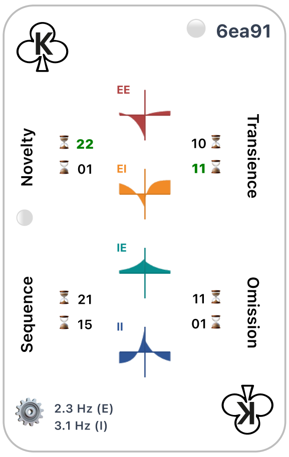
6ea91
K
EE
EI
IE
II
Novelty
22
01
Transience
10
11
Sequence
21
15
Omission
11
01
2.3 Hz (E)
3.1 Hz (I)

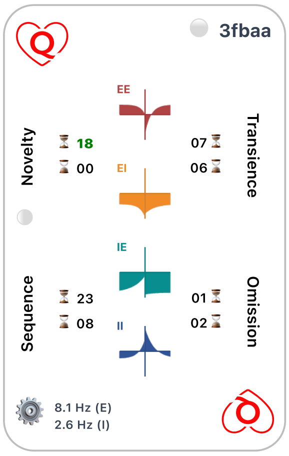
3fbaa
Q
EE
EI
IE
II
Novelty
18
00
Transience
07
06
Sequence
23
08
Omission
01
02
8.1 Hz (E)
2.6 Hz (I)

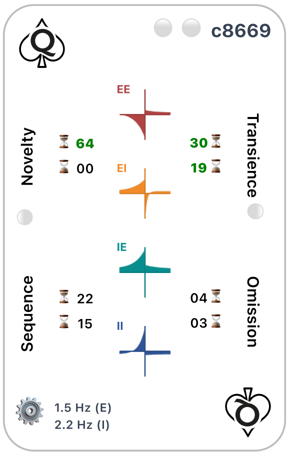
c8669
Q
EE
EI
IE
II
Novelty
64
00
Transience
30
19
Sequence
22
15
Omission
04
03
1.5 Hz (E)
2.2 Hz (I)

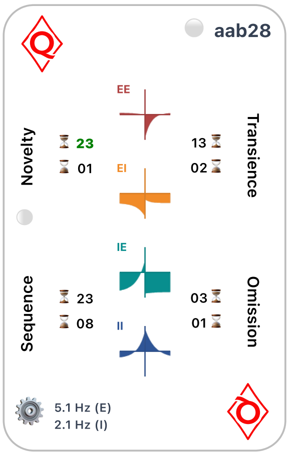
aab28
Q
EE
EI
IE
II
Novelty
23
01
Transience
13
02
Sequence
23
08
Omission
03
01
5.1 Hz (E)
2.1 Hz (I)

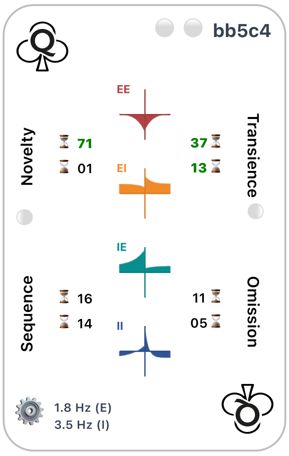
bb5c4
Q
EE
EI
IE
II
Novelty
71
01
Transience
37
13
Sequence
16
14
Omission
11
05
1.8 Hz (E)
3.5 Hz (I)

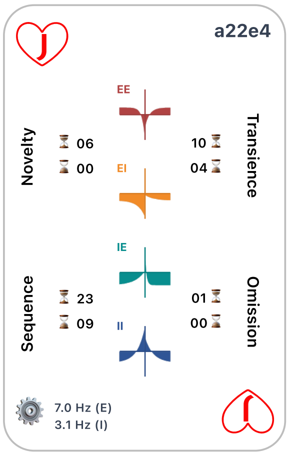
a22e4
J
EE
EI
IE
II
Novelty
06
00
Transience
10
04
Sequence
23
09
Omission
01
00
7.0 Hz (E)
3.1 Hz (I)

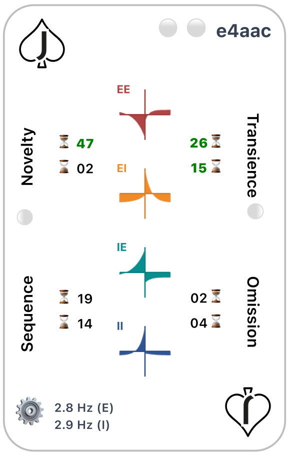
e4aac
J
EE
EI
IE
II
Novelty
47
02
Transience
26
15
Sequence
19
14
Omission
02
04
2.8 Hz (E)
2.9 Hz (I)

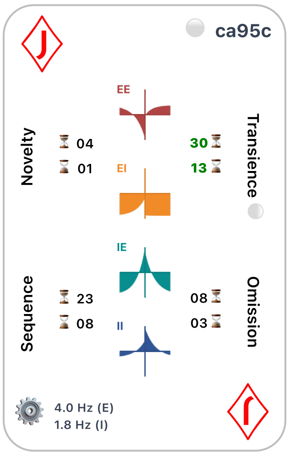
ca95c
J
EE
EI
IE
II
Novelty
04
01
Transience
30
13
Sequence
23
08
Omission
08
03
4.0 Hz (E)
1.8 Hz (I)

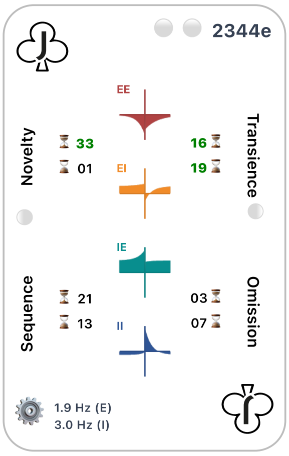
2344e
J
EE
EI
IE
II
Novelty
33
01
Transience
16
19
Sequence
21
13
Omission
03
07
1.9 Hz (E)
3.0 Hz (I)

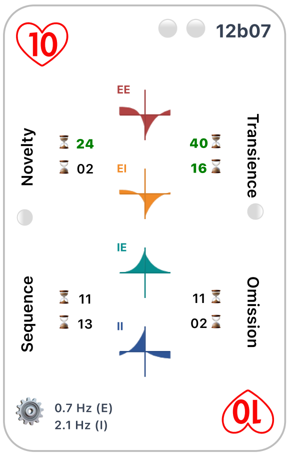
10
12b07
Novelty
24
02
Transience
40
16
EE
EI
IE
II
Sequence
11
13
Omission
11
02
0.7 Hz (E)
2.1 Hz (I)

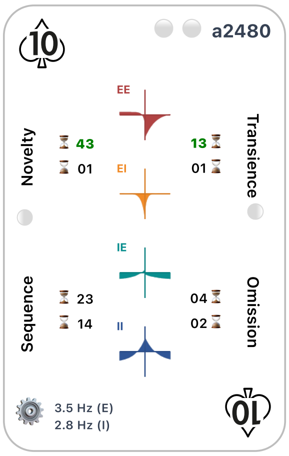
10
a2480
Novelty
43
01
Transience
13
01
EE
EI
IE
II
Sequence
23
14
Omission
04
02
3.5 Hz (E)
2.8 Hz (I)

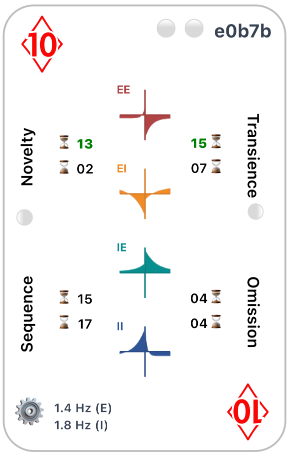
10
e0b7b
Novelty
13
02
Transience
15
07
EE
EI
IE
II
Sequence
15
17
Omission
04
04
1.4 Hz (E)
1.8 Hz (I)

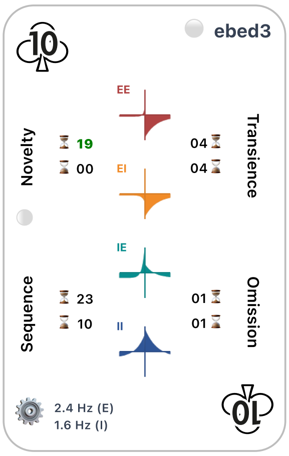
10
ebed3
Novelty
19
00
Transience
04
04
EE
EI
IE
II
Sequence
23
10
Omission
01
01
2.4 Hz (E)
1.6 Hz (I)

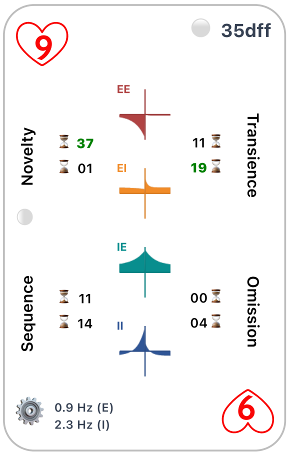
9
35dff
Novelty
37
01
Transience
11
19
EE
EI
IE
II
Sequence
11
14
Omission
00
04
0.9 Hz (E)
2.3 Hz (I)

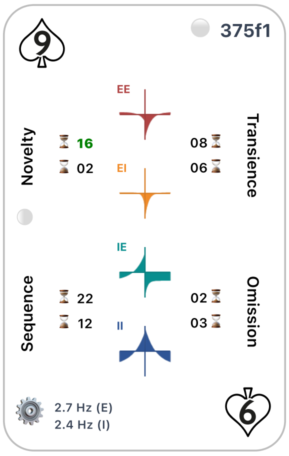
9
375f1
Novelty
16
02
Transience
08
06
EE
EI
IE
II
Sequence
22
12
Omission
02
03
2.7 Hz (E)
2.4 Hz (I)

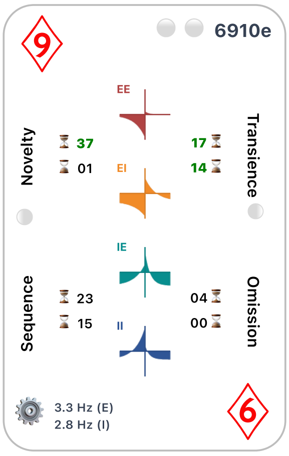
9
6910e
Novelty
37
01
Transience
17
14
EE
EI
IE
II
Sequence
23
15
Omission
04
00
3.3 Hz (E)
2.8 Hz (I)

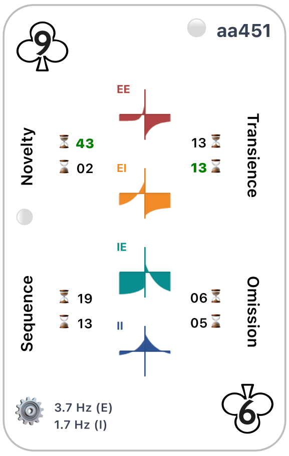
9
aa451
Novelty
43
02
Transience
13
13
EE
EI
IE
II
Sequence
19
13
Omission
06
05
3.7 Hz (E)
1.7 Hz (I)

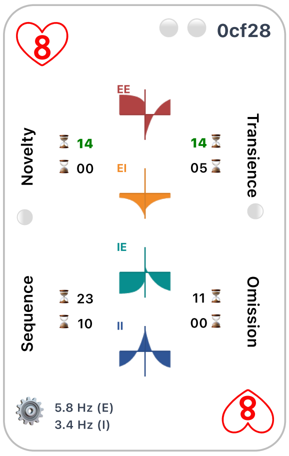
8
0cf28
Novelty
14
00
Transience
14
05
EE
EI
IE
II
Sequence
23
10
Omission
11
00
5.8 Hz (E)
3.4 Hz (I)

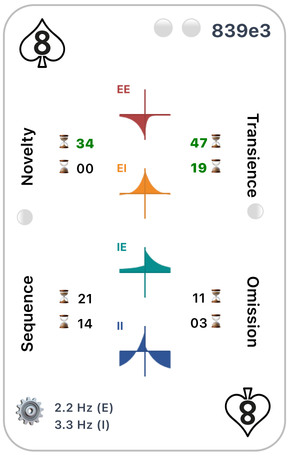
8
839e3
Novelty
34
00
Transience
47
19
EE
EI
IE
II
Sequence
21
14
Omission
11
03
2.2 Hz (E)
3.3 Hz (I)

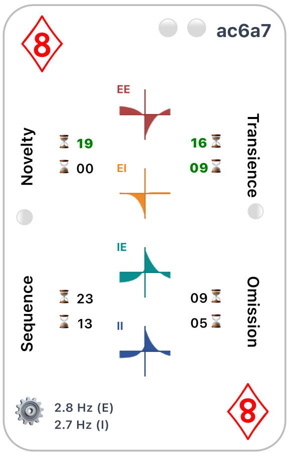
8
ac6a7
Novelty
19
00
Transience
16
09
EE
EI
IE
II
Sequence
23
13
Omission
09
05
2.8 Hz (E)
2.7 Hz (I)

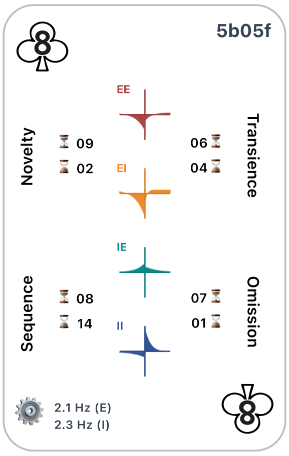
8
5b05f
Novelty
09
02
Transience
06
04
EE
EI
IE
II
Sequence
08
14
Omission
07
01
2.1 Hz (E)
2.3 Hz (I)

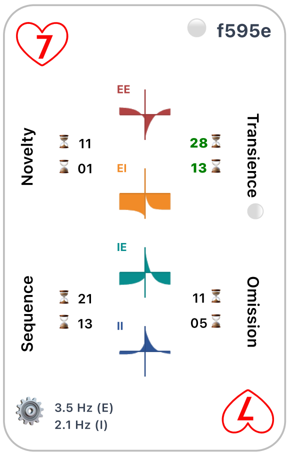
7
f595e
Novelty
11
01
Transience
28
13
EE
EI
IE
II
Sequence
21
13
Omission
11
05
3.5 Hz (E)
2.1 Hz (I)

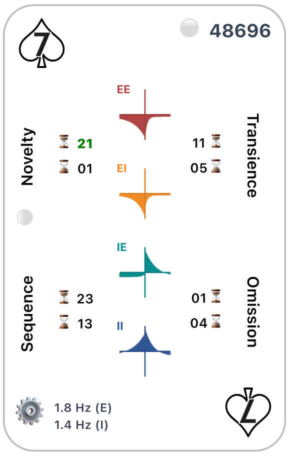
7
48696
Novelty
21
01
Transience
11
05
EE
EI
IE
II
Sequence
23
13
Omission
01
04
1.8 Hz (E)
1.4 Hz (I)

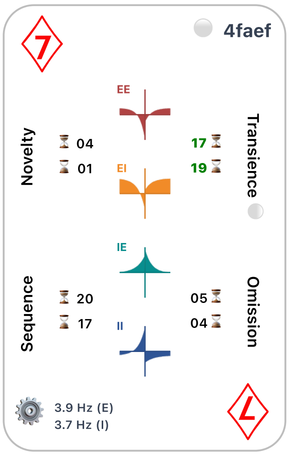
7
4faef
Novelty
04
01
Transience
17
19
EE
EI
IE
II
Sequence
20
17
Omission
05
04
3.9 Hz (E)
3.7 Hz (I)

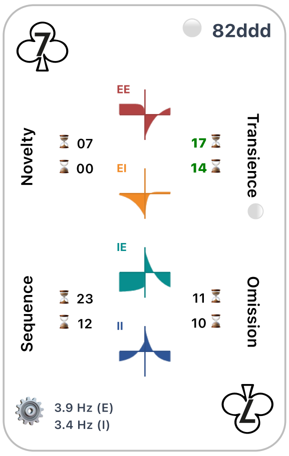
7
82ddd
Novelty
07
00
Transience
17
14
EE
EI
IE
II
Sequence
23
12
Omission
11
10
3.9 Hz (E)
3.4 Hz (I)

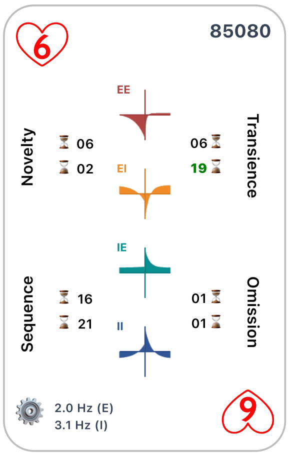
6
85080
EE
Novelty
06
02
06
19
Transience
EI
IE
Sequence
16
21
01
01
Omission
II
2.0 Hz (E)
3.1 Hz (I)
9

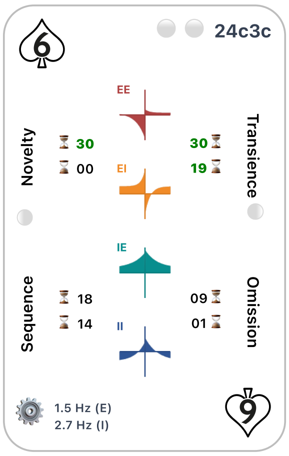
6
24c3c
EE
Novelty
30
00
30
19
Transience
EI
IE
Sequence
18
14
09
01
Omission
II
1.5 Hz (E)
2.7 Hz (I)
9

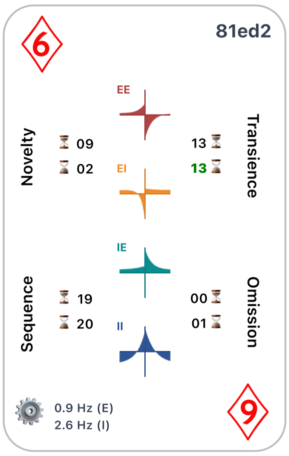
6
81ed2
EE
Novelty
09
02
13
13
Transience
EI
IE
Sequence
19
20
00
01
Omission
II
0.9 Hz (E)
2.6 Hz (I)
9

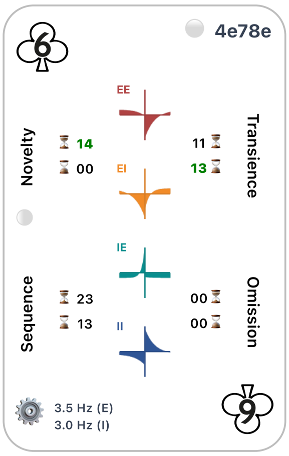
6
4e78e
EE
Novelty
14
00
11
13
Transience
EI
IE
Sequence
23
13
00
00
Omission
II
3.5 Hz (E)
3.0 Hz (I)
9

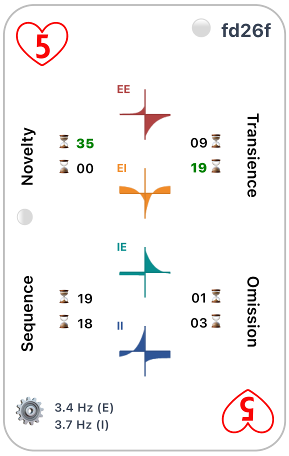
5
fd26f
EE
Novelty
35
00
09
19
Transience
EI
IE
Sequence
19
18
01
03
Omission
II
3.4 Hz (E)
3.7 Hz (I)
5

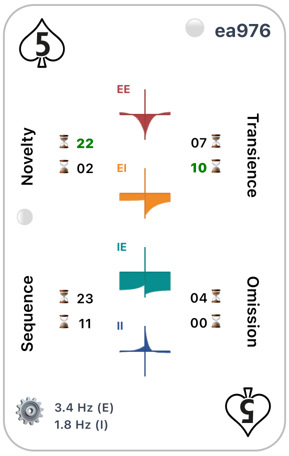
5
ea976
EE
Novelty
22
02
07
10
Transience
EI
IE
Sequence
23
11
04
00
Omission
II
3.4 Hz (E)
1.8 Hz (I)
5

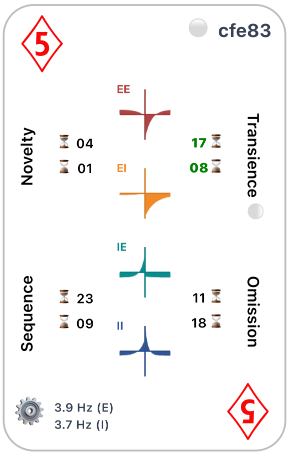
5
cfe83
EE
Novelty
04
01
17
08
Transience
EI
IE
Sequence
23
09
11
18
Omission
II
3.9 Hz (E)
3.7 Hz (I)
5

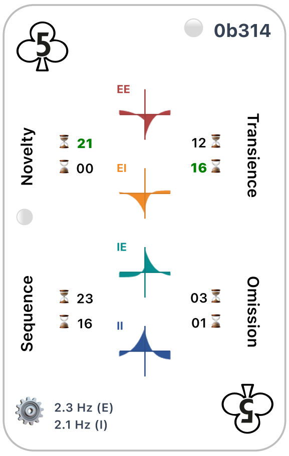
5
0b314
EE
Novelty
21
00
12
16
Transience
EI
IE
Sequence
23
16
03
01
Omission
II
2.3 Hz (E)
2.1 Hz (I)
5

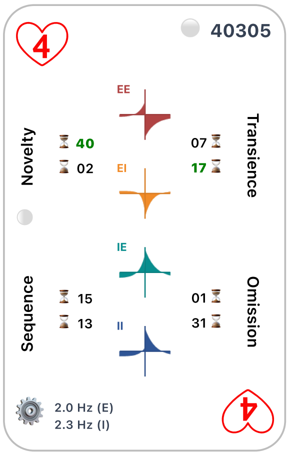
4
40305
EE
Novelty
40
02
07
17
Transience
EI
IE
Sequence
15
13
01
31
Omission
II
2.0 Hz (E)
2.3 Hz (I)
4

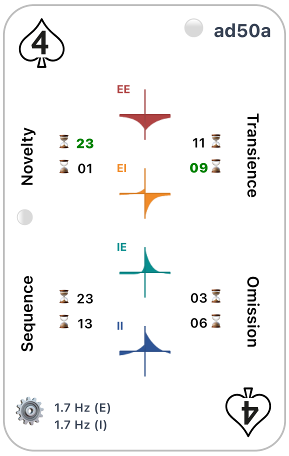
4
ad50a
EE
Novelty
23
01
11
09
Transience
EI
IE
Sequence
23
13
03
06
Omission
II
1.7 Hz (E)
1.7 Hz (I)
4

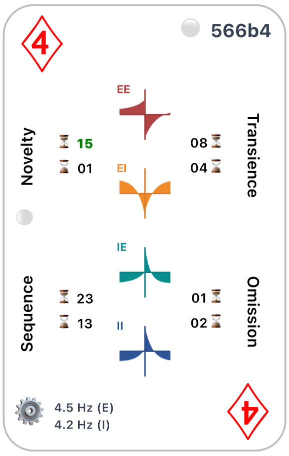
4
566b4
EE
Novelty
15
01
08
04
Transience
EI
IE
Sequence
23
13
01
02
Omission
II
4.5 Hz (E)
4.2 Hz (I)
4

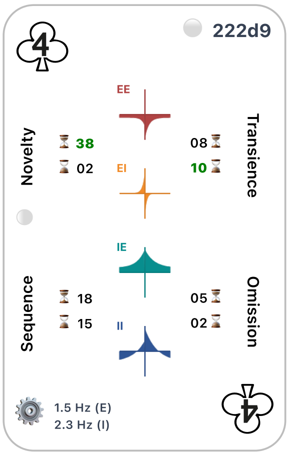
4
222d9
EE
Novelty
38
02
08
10
Transience
EI
IE
Sequence
18
15
05
02
Omission
II
1.5 Hz (E)
2.3 Hz (I)
4

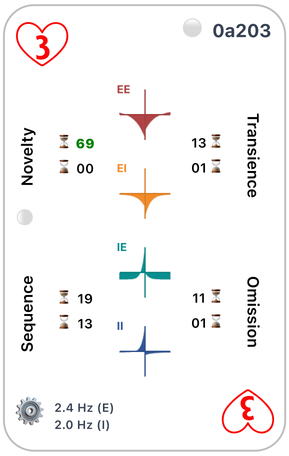
3
0a203
EE
Novelty
69
00
13
01
Transience
EI
IE
Sequence
19
13
11
01
Omission
II
2.4 Hz (E)
2.0 Hz (I)
3

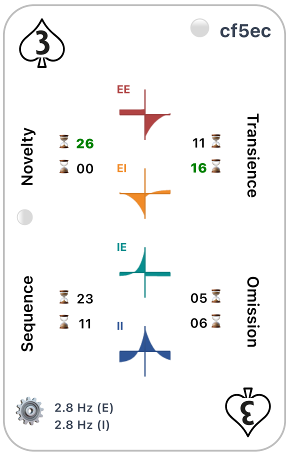
3
cf5ec
EE
Novelty
26
00
11
16
Transience
EI
IE
Sequence
23
11
05
06
Omission
II
2.8 Hz (E)
2.8 Hz (I)
3

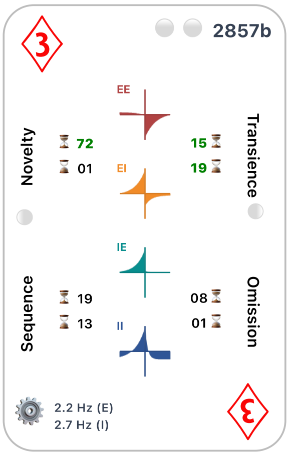
3
2857b
EE
Novelty
72
01
15
19
Transience
EI
IE
Sequence
19
13
08
01
Omission
II
2.2 Hz (E)
2.7 Hz (I)
3

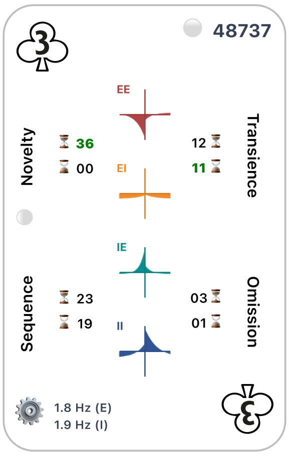
3
48737
EE
Novelty
36
00
12
11
Transience
EI
IE
Sequence
23
19
03
01
Omission
II
1.8 Hz (E)
1.9 Hz (I)
3

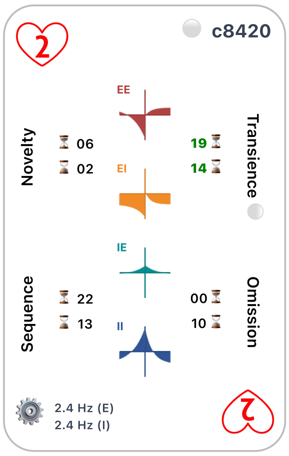
2
c8420
EE
EI
IE
II
Novelty
06
02
19
14
Transience
Sequence
22
13
00
10
Omission
2.4 Hz (E)
2.4 Hz (I)

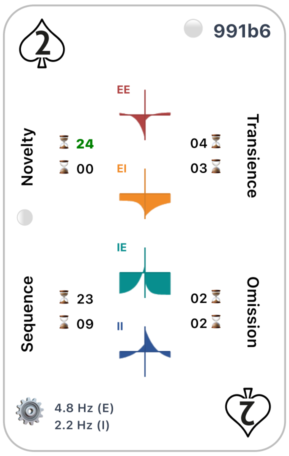
2
991b6
EE
EI
IE
II
Novelty
24
00
04
03
Transience
Sequence
23
09
02
02
Omission
4.8 Hz (E)
2.2 Hz (I)

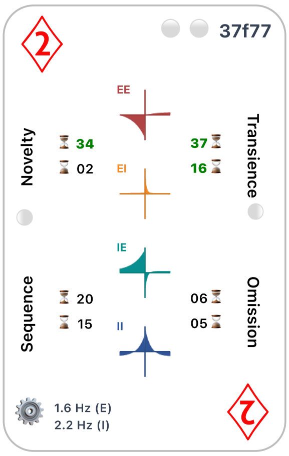
2
37f77
EE
EI
IE
II
Novelty
34
02
37
16
Transience
Sequence
20
15
06
05
Omission
1.6 Hz (E)
2.2 Hz (I)

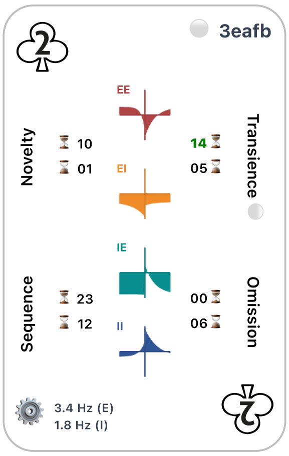
2
3eafb
EE
EI
IE
II
Novelty
10
01
14
05
Transience
Sequence
23
12
00
06
Omission
3.4 Hz (E)
1.8 Hz (I)

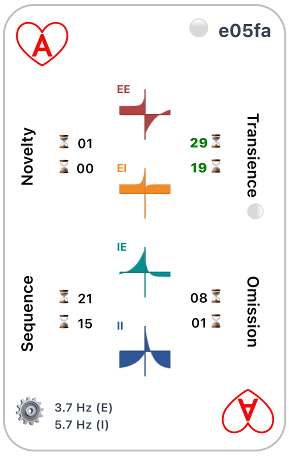
A
e05fa
EE
EI
IE
II
Novelty
01
00
29
19
Transience
Sequence
21
15
08
01
Omission
3.7 Hz (E)
5.7 Hz (I)

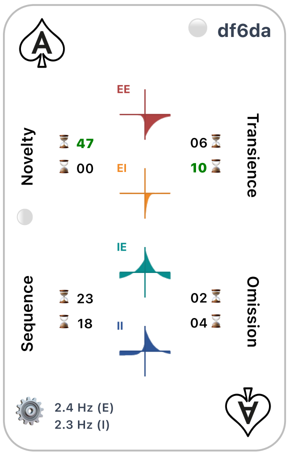
A
df6da
EE
EI
IE
II
Novelty
47
00
06
10
Transience
Sequence
23
18
02
04
Omission
2.4 Hz (E)
2.3 Hz (I)

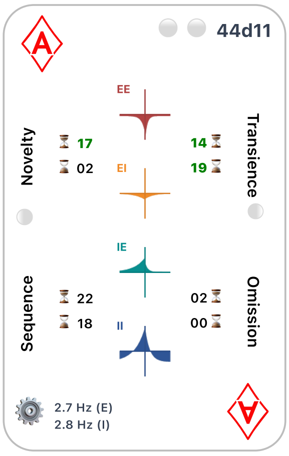
A
44d11
EE
EI
IE
II
Novelty
17
02
14
19
Transience
Sequence
22
18
02
00
Omission
2.7 Hz (E)
2.8 Hz (I)

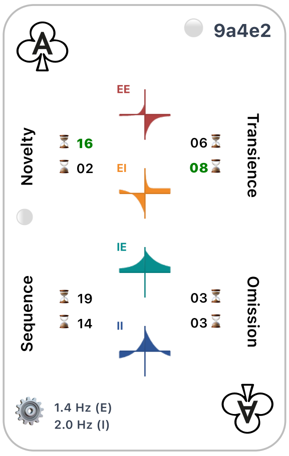
A
9a4e2
EE
EI
IE
II
Novelty
16
02
06
08
Transience
Sequence
19
14
03
03
Omission
1.4 Hz (E)
2.0 Hz (I)

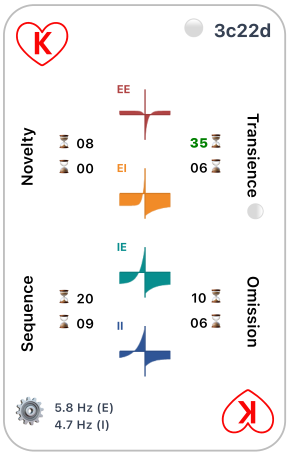
K
3c22d
EE
EI
IE
II
Novelty
08
00
35
06
Transience
Sequence
20
09
10
06
Omission
5.8 Hz (E)
4.7 Hz (I)

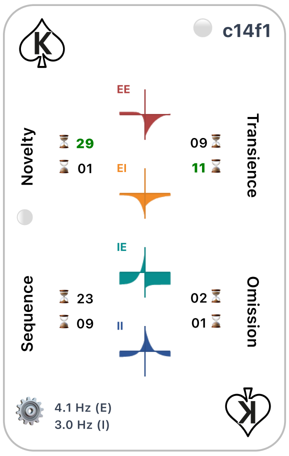
K
c14f1
EE
EI
IE
II
Novelty
29
01
09
11
Transience
Sequence
23
09
02
01
Omission
4.1 Hz (E)
3.0 Hz (I)

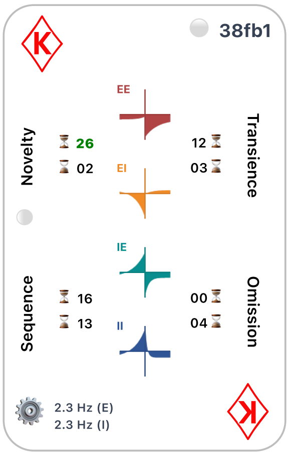
K
38fb1
EE
EI
IE
II
Novelty
26
02
12
03
Transience
Sequence
16
13
00
04
Omission
2.3 Hz (E)
2.3 Hz (I)

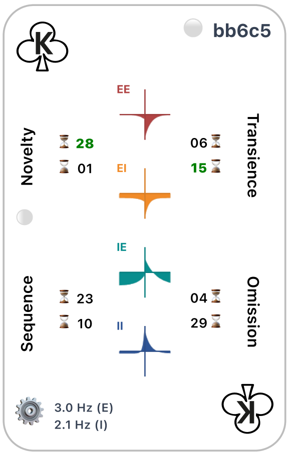
K
bb6c5
EE
EI
IE
II
Novelty
28
01
06
15
Transience
Sequence
23
10
04
29
Omission
3.0 Hz (E)
2.1 Hz (I)

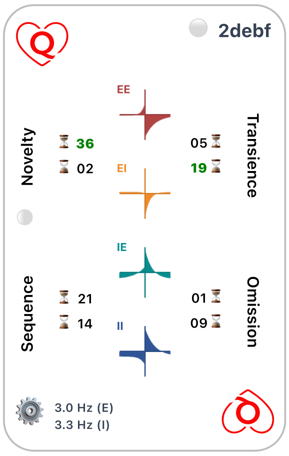
Q
2debf
EE
EI
IE
II
Novelty
36
02
05
19
Transience
Sequence
21
14
01
09
Omission
3.0 Hz (E)
3.3 Hz (I)

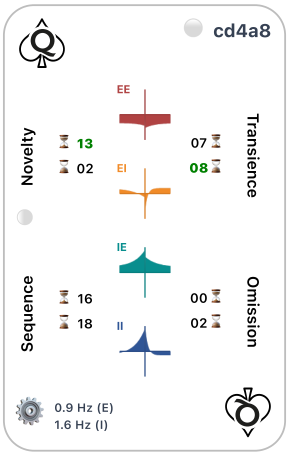
Q
cd4a8
EE
EI
IE
II
Novelty
13
02
07
08
Transience
Sequence
16
18
00
02
Omission
0.9 Hz (E)
1.6 Hz (I)

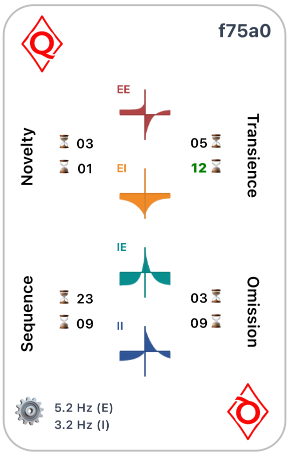
Q
f75a0
EE
EI
IE
II
Novelty
03
01
05
12
Transience
Sequence
23
09
03
09
Omission
5.2 Hz (E)
3.2 Hz (I)

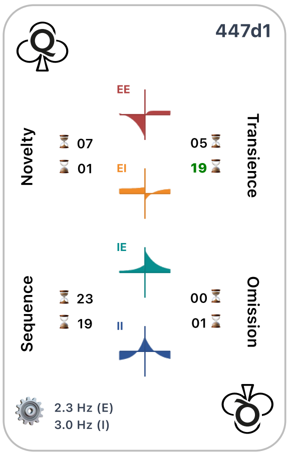
Q
447d1
EE
EI
IE
II
Novelty
07
01
05
19
Transience
Sequence
23
19
00
01
Omission
2.3 Hz (E)
3.0 Hz (I)

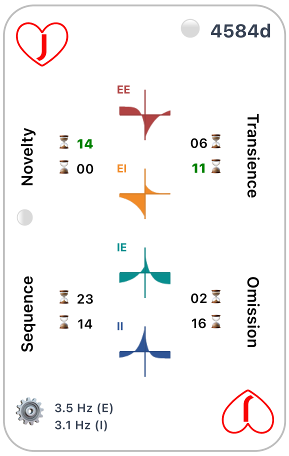
J
4584d
EE
EI
IE
II
Novelty 14 00
Transience 06 11
Sequence 23 14
Omission 02 16
3.5 Hz (E)
3.1 Hz (I)

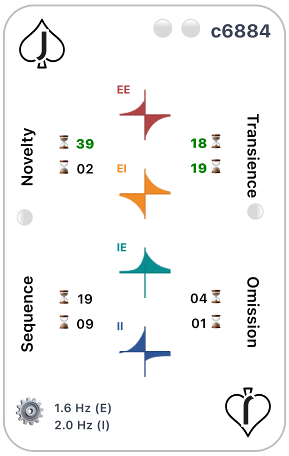
J
c6884
EE
EI
IE
II
Novelty 39 02
Transience 18 19
Sequence 19 09
Omission 04 01
1.6 Hz (E)
2.0 Hz (I)

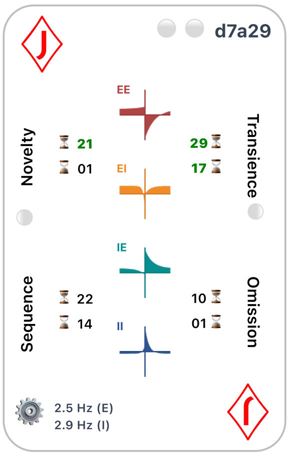
J
d7a29
EE
EI
IE
II
Novelty 21 01
Transience 29 17
Sequence 22 14
Omission 10 01
2.5 Hz (E)
2.9 Hz (I)

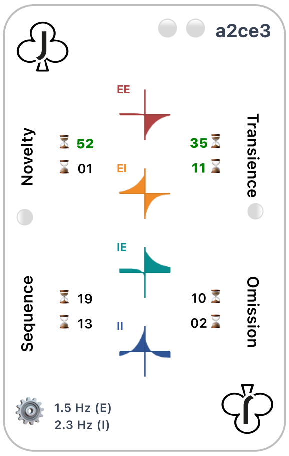
J
a2ce3
EE
EI
IE
II
Novelty 52 01
Transience 35 11
Sequence 19 13
Omission 10 02
1.5 Hz (E)
2.3 Hz (I)

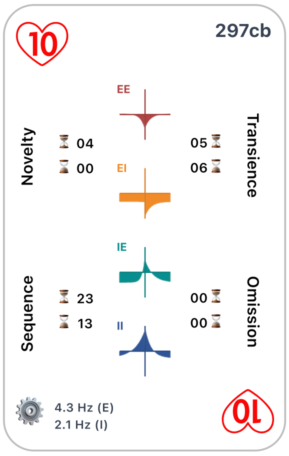
10
297cb
EE
EI
IE
II
Novelty 04 00
Transience 05 06
Sequence 23 13
Omission 00 00
4.3 Hz (E)
2.1 Hz (I)

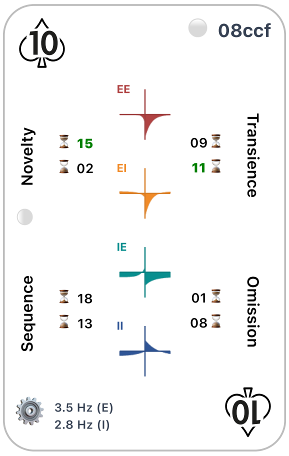
10
08ccf
EE
EI
IE
II
Novelty 15 02
Transience 09 11
Sequence 18 13
Omission 01 08
3.5 Hz (E)
2.8 Hz (I)

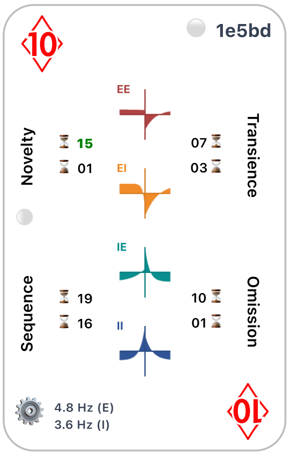
10
1e5bd
EE
EI
IE
II
Novelty 15 01
Transience 07 03
Sequence 19 16
Omission 10 01
4.8 Hz (E)
3.6 Hz (I)

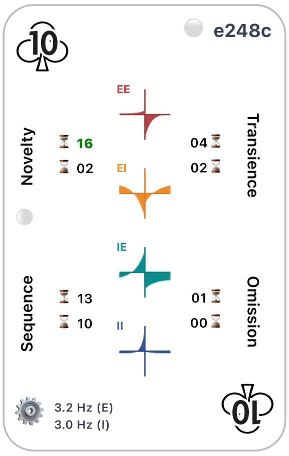
10
e248c
EE
EI
IE
II
Novelty 16 02
Transience 04 02
Sequence 13 10
Omission 01 00
3.2 Hz (E)
3.0 Hz (I)

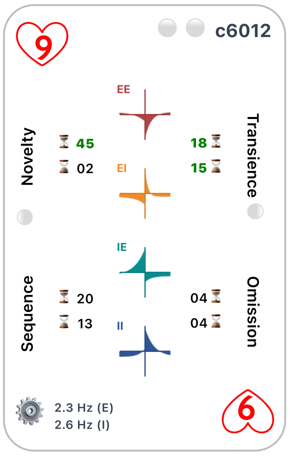
9
c6012
EE
EI
IE
II
Novelty 45 02
Transience 18 15
Sequence 20 13
Omission 04 04
2.3 Hz (E)
2.6 Hz (I)

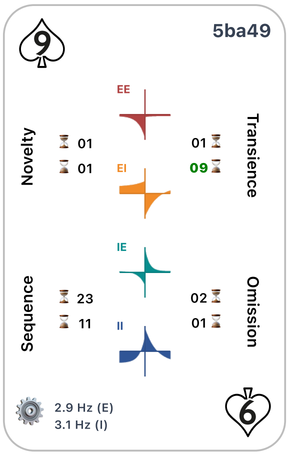
9
5ba49
EE
EI
IE
II
Novelty 01 01
Transience 01 09
Sequence 23 11
Omission 02 01
2.9 Hz (E)
3.1 Hz (I)

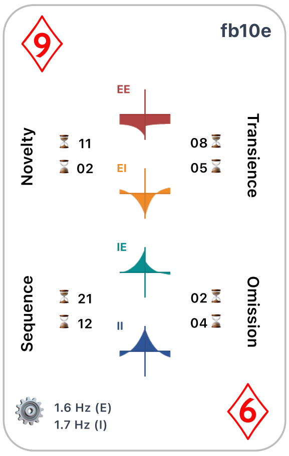
9
fb10e
EE
EI
IE
II
Novelty 11 02
Transience 08 05
Sequence 21 12
Omission 02 04
1.6 Hz (E)
1.7 Hz (I)

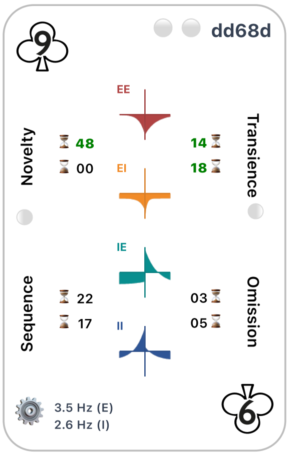
9
dd68d
EE
EI
IE
II
Novelty 48 00
Transience 14 18
Sequence 22 17
Omission 03 05
3.5 Hz (E)
2.6 Hz (I)

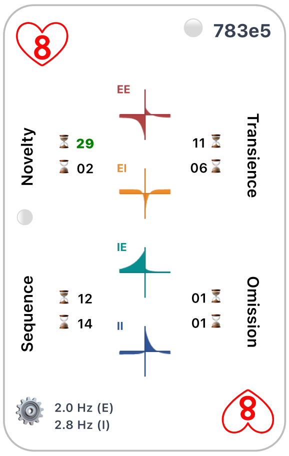
8
783e5
EE
EI
IE
II
Novelty 29 02
Transience 11 06
Sequence 12 14
Omission 01 01
2.0 Hz (E)
2.8 Hz (I)

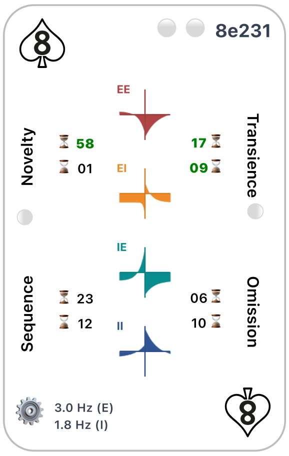
8
8e231
EE
EI
IE
II
Novelty 58 01
Transience 17 09
Sequence 23 12
Omission 06 10
3.0 Hz (E)
1.8 Hz (I)

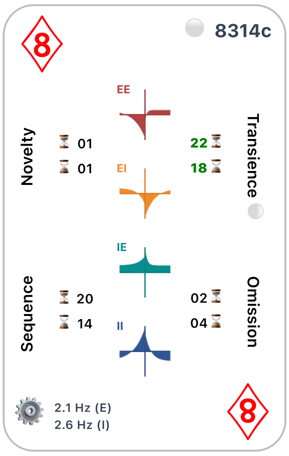
8
8314c
EE
EI
IE
II
Novelty 01 01
Transience 22 18
Sequence 20 14
Omission 02 04
2.1 Hz (E)
2.6 Hz (I)

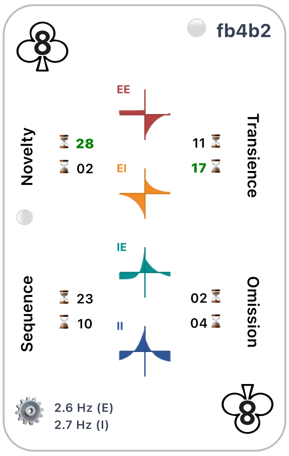
8
fb4b2
EE
EI
IE
II
Novelty 28 02
Transience 11 17
Sequence 23 10
Omission 02 04
2.6 Hz (E)
2.7 Hz (I)

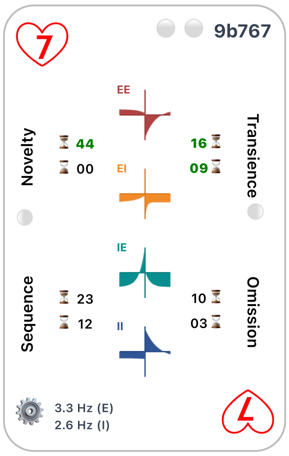
7
9b767
EE
EI
IE
II
Novelty
44
00
Transience
16
09
Sequence
23
12
Omission
10
03
3.3 Hz (E)
2.6 Hz (I)
7

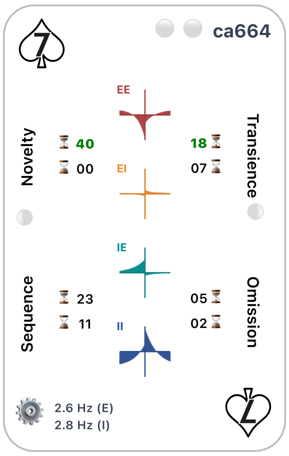
7
ca664
EE
EI
IE
II
Novelty
40
00
Transience
18
07
Sequence
23
11
Omission
05
02
2.6 Hz (E)
2.8 Hz (I)
7

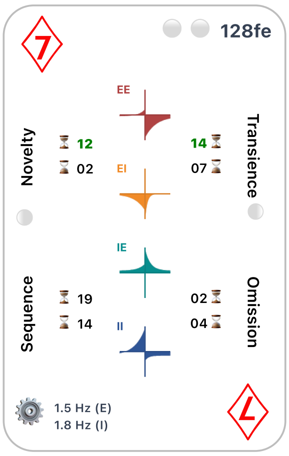
7
128fe
EE
EI
IE
II
Novelty
12
02
Transience
14
07
Sequence
19
14
Omission
02
04
1.5 Hz (E)
1.8 Hz (I)
7

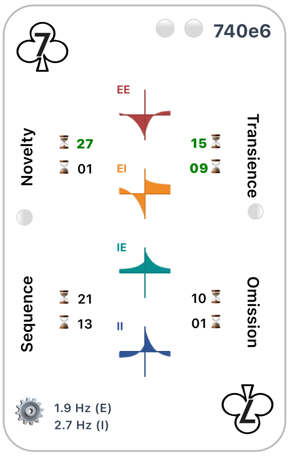
7
740e6
EE
EI
IE
II
Novelty
27
01
Transience
15
09
Sequence
21
13
Omission
10
01
1.9 Hz (E)
2.7 Hz (I)
7

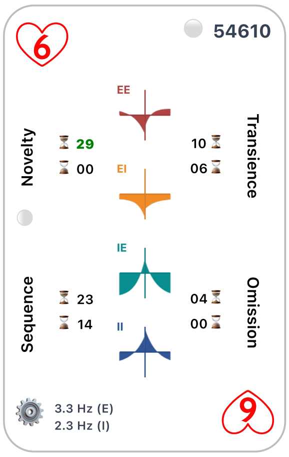
6
54610
EE
EI
IE
II
Novelty
29
00
Transience
10
06
Sequence
23
14
Omission
04
00
3.3 Hz (E)
2.3 Hz (I)
6

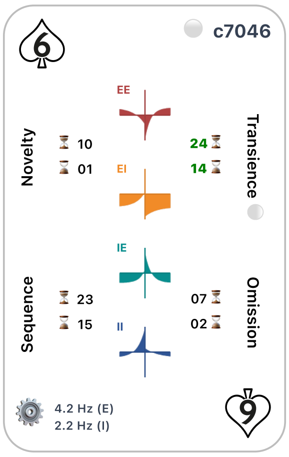
6
c7046
EE
EI
IE
II
Novelty
10
01
Transience
24
14
Sequence
23
15
Omission
07
02
4.2 Hz (E)
2.2 Hz (I)
6

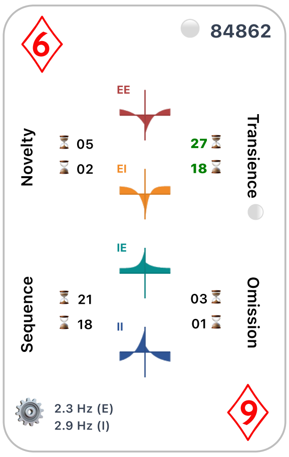
6
84862
EE
EI
IE
II
Novelty
05
02
Transience
27
18
Sequence
21
18
Omission
03
01
2.3 Hz (E)
2.9 Hz (I)
6

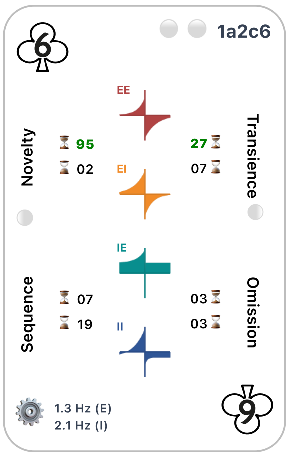
6
1a2c6
EE
EI
IE
II
Novelty
95
02
Transience
27
07
Sequence
07
19
Omission
03
03
1.3 Hz (E)
2.1 Hz (I)
6

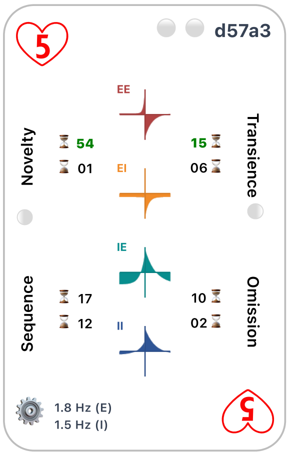
5
d57a3
EE
EI
IE
II
Novelty
54
01
Transience
15
06
Sequence
17
12
Omission
10
02
1.8 Hz (E)
1.5 Hz (I)
5

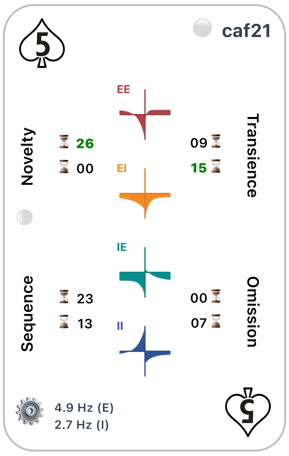
5
caf21
EE
EI
IE
II
Novelty
26
00
Transience
09
15
Sequence
23
13
Omission
00
07
4.9 Hz (E)
2.7 Hz (I)
5

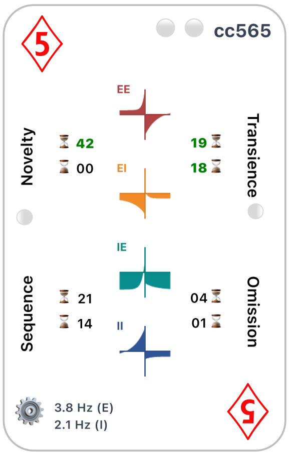
5
cc565
EE
EI
IE
II
Novelty
42
00
Transience
19
18
Sequence
21
14
Omission
04
01
3.8 Hz (E)
2.1 Hz (I)
5

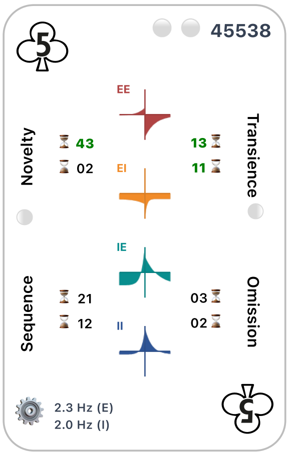
5
45538
EE
EI
IE
II
Novelty
43
02
Transience
13
11
Sequence
21
12
Omission
03
02
2.3 Hz (E)
2.0 Hz (I)
5

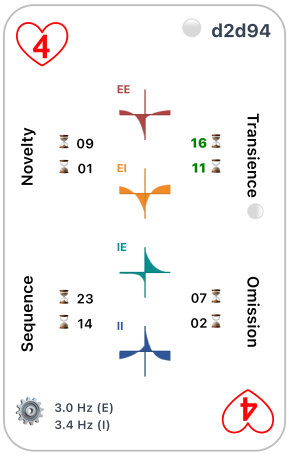
4
d2d94
EE
EI
IE
II
Novelty
09
01
Transience
16
11
Sequence
23
14
Omission
07
02
3.0 Hz (E)
3.4 Hz (I)
4

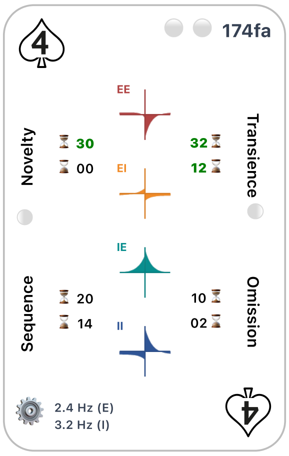
4
174fa
EE
EI
IE
II
Novelty
30
00
Transience
32
12
Sequence
20
14
Omission
10
02
2.4 Hz (E)
3.2 Hz (I)
4

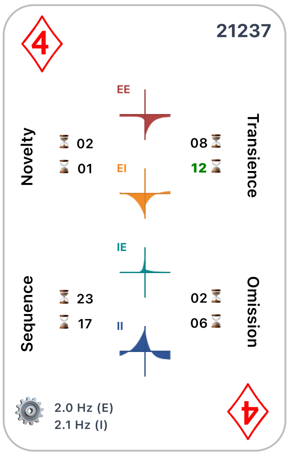
4
21237
EE
EI
IE
II
Novelty
02
01
Transience
08
12
Sequence
23
17
Omission
02
06
2.0 Hz (E)
2.1 Hz (I)
4

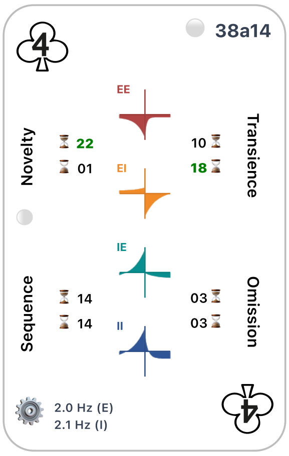
4
38a14
EE
EI
IE
II
Novelty
22
01
Transience
10
18
Sequence
14
14
Omission
03
03
2.0 Hz (E)
2.1 Hz (I)
4

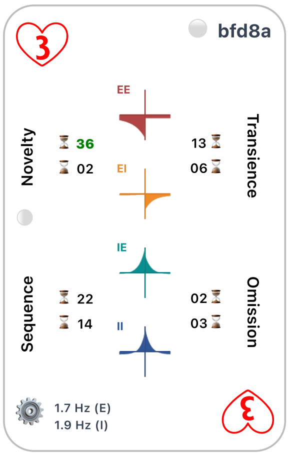
3
bfd8a
EE
EI
IE
II
Novelty
36
02
Transience
13
06
Sequence
22
14
Omission
02
03
1.7 Hz (E)
1.9 Hz (I)

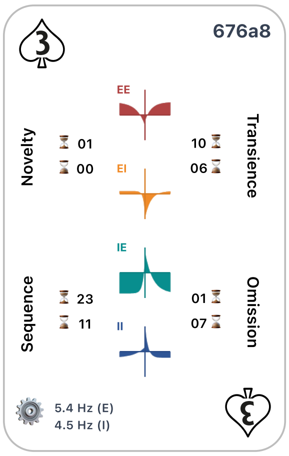
3
676a8
EE
EI
IE
II
Novelty
01
00
Transience
10
06
Sequence
23
11
Omission
01
07
5.4 Hz (E)
4.5 Hz (I)

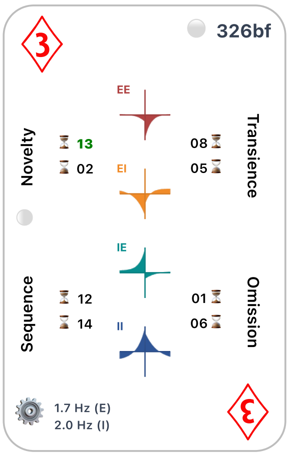
3
326bf
EE
EI
IE
II
Novelty
13
02
Transience
08
05
Sequence
12
14
Omission
01
06
1.7 Hz (E)
2.0 Hz (I)

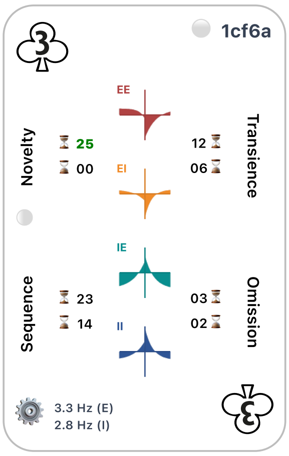
3
1cf6a
EE
EI
IE
II
Novelty
25
00
Transience
12
06
Sequence
23
14
Omission
03
02
3.3 Hz (E)
2.8 Hz (I)

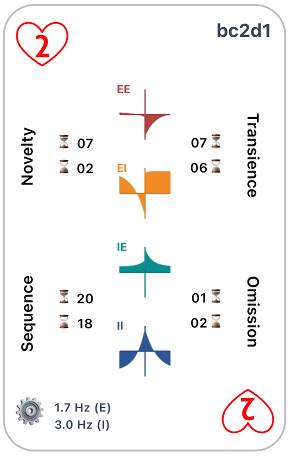
2
bc2d1
EE
EI
IE
II
Novelty
07
02
Transience
07
06
Sequence
20
18
Omission
01
02
1.7 Hz (E)
3.0 Hz (I)

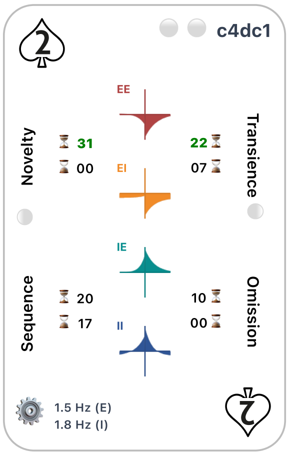
2
c4dc1
EE
EI
IE
II
Novelty
31
00
Transience
22
07
Sequence
20
17
Omission
10
00
1.5 Hz (E)
1.8 Hz (I)

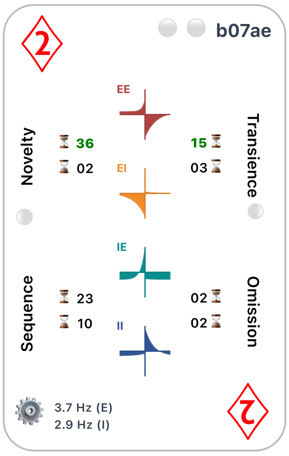
2
b07ae
EE
EI
IE
II
Novelty
36
02
Transience
15
03
Sequence
23
10
Omission
02
02
3.7 Hz (E)
2.9 Hz (I)

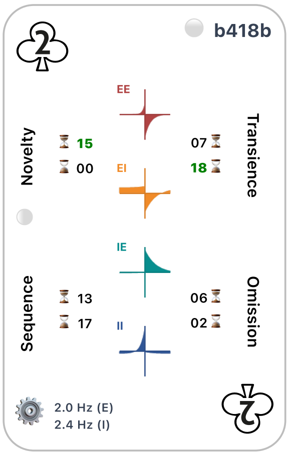
2
b418b
EE
EI
IE
II
Novelty
15
00
Transience
07
18
Sequence
13
17
Omission
06
02
2.0 Hz (E)
2.4 Hz (I)

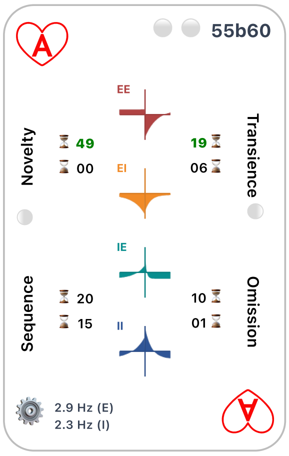
A
55b60
EE
EI
IE
II
Novelty
49
00
Transience
19
06
Sequence
20
15
Omission
10
01
2.9 Hz (E)
2.3 Hz (I)

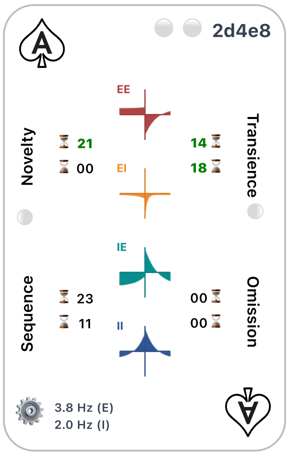
A
2d4e8
EE
EI
IE
II
Novelty
21
00
Transience
14
18
Sequence
23
11
Omission
00
00
3.8 Hz (E)
2.0 Hz (I)

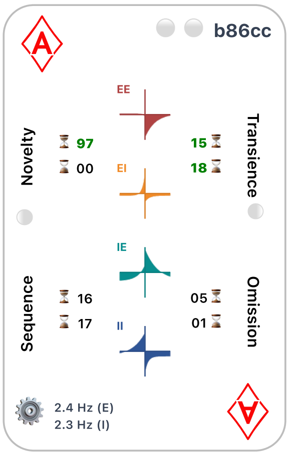
A
b86cc
EE
EI
IE
II
Novelty
97
00
Transience
15
18
Sequence
16
17
Omission
05
01
2.4 Hz (E)
2.3 Hz (I)

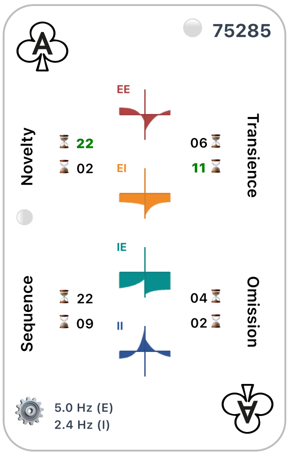
A
75285
EE
EI
IE
II
Novelty
22
02
Transience
06
11
Sequence
22
09
Omission
04
02
5.0 Hz (E)
2.4 Hz (I)

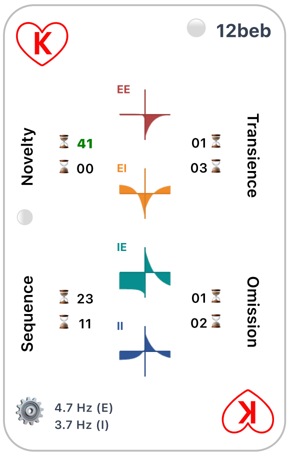
K
12beb
EE
EI
IE
II
Novelty
41
00
Transience
01
03
Sequence
23
11
Omission
01
02
4.7 Hz (E)
3.7 Hz (I)

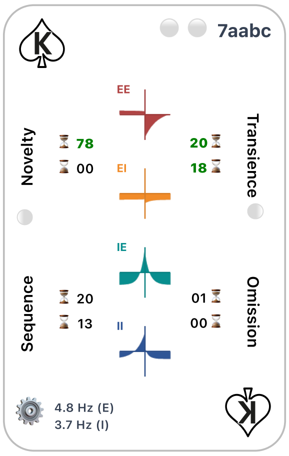
K
7aabc
EE
EI
IE
II
Novelty
78
00
Transience
20
18
Sequence
20
13
Omission
01
00
4.8 Hz (E)
3.7 Hz (I)

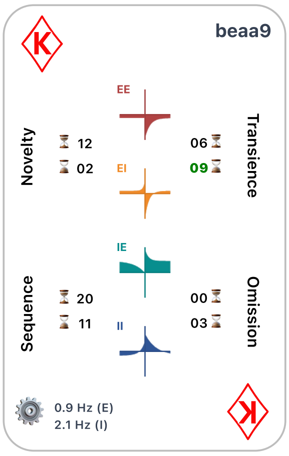
K
beaa9
EE
EI
IE
II
Novelty
12
02
Transience
06
09
Sequence
20
11
Omission
00
03
0.9 Hz (E)
2.1 Hz (I)

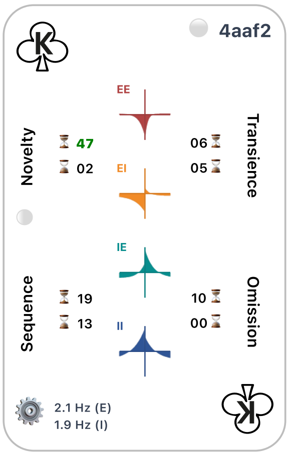
K
4aaf2
EE
EI
IE
II
Novelty
47
02
Transience
06
05
Sequence
19
13
Omission
10
00
2.1 Hz (E)
1.9 Hz (I)

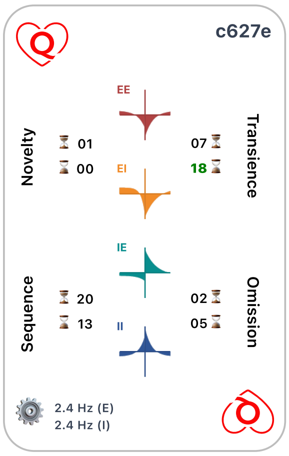
Q
c627e
EE
EI
IE
II
Novelty 01 00
Transience 07 18
Sequence 20 13
Omission 02 05
2.4 Hz (E)
2.4 Hz (I)

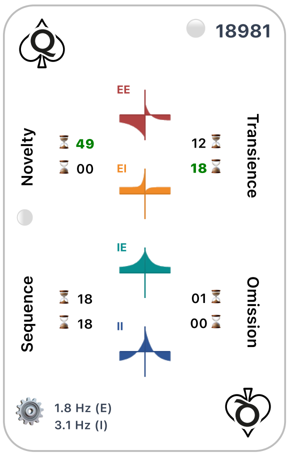
Q
18981
EE
EI
IE
II
Novelty 49 00
Transience 12 18
Sequence 18 18
Omission 01 00
1.8 Hz (E)
3.1 Hz (I)

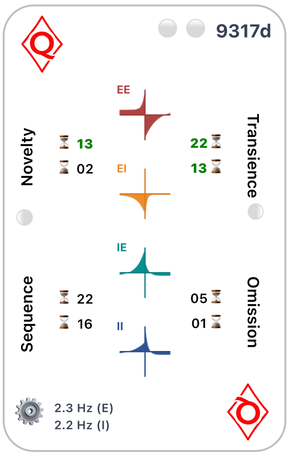
Q
9317d
EE
EI
IE
II
Novelty 13 02
Transience 22 13
Sequence 22 16
Omission 05 01
2.3 Hz (E)
2.2 Hz (I)

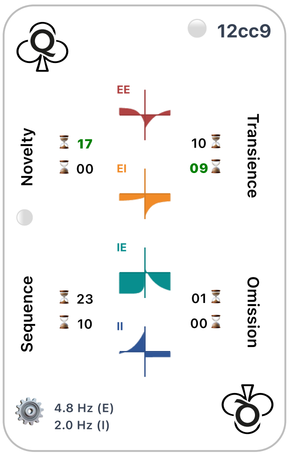
Q
12cc9
EE
EI
IE
II
Novelty 17 00
Transience 10 09
Sequence 23 10
Omission 01 00
4.8 Hz (E)
2.0 Hz (I)

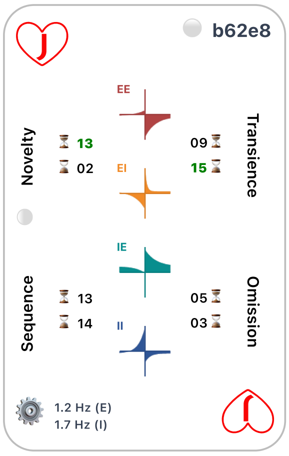
J
b62e8
EE
EI
IE
II
Novelty 13 02
Transience 09 15
Sequence 13 14
Omission 05 03
1.2 Hz (E)
1.7 Hz (I)

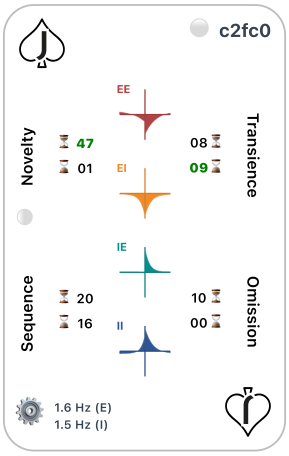
J
c2fc0
EE
EI
IE
II
Novelty 47 01
Transience 08 09
Sequence 20 16
Omission 10 00
1.6 Hz (E)
1.5 Hz (I)

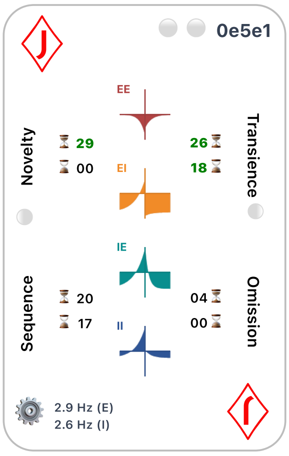
J
0e5e1
EE
EI
IE
II
Novelty 29 00
Transience 26 18
Sequence 20 17
Omission 04 00
2.9 Hz (E)
2.6 Hz (I)

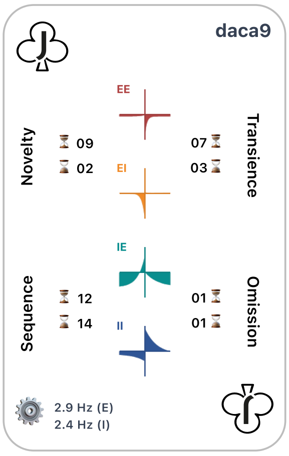
J
daca9
EE
EI
IE
II
Novelty 09 02
Transience 07 03
Sequence 12 14
Omission 01 01
2.9 Hz (E)
2.4 Hz (I)

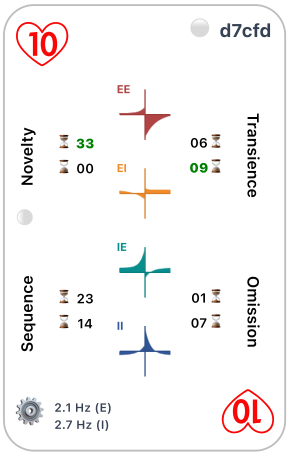
10
d7cfd
EE
EI
IE
II
Novelty 33 00
Transience 06 09
Sequence 23 14
Omission 01 07
2.1 Hz (E)
2.7 Hz (I)

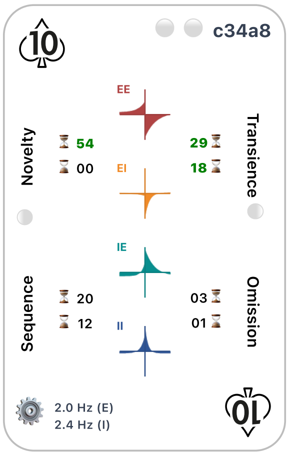
10
c34a8
EE
EI
IE
II
Novelty 54 00
Transience 29 18
Sequence 20 12
Omission 03 01
2.0 Hz (E)
2.4 Hz (I)

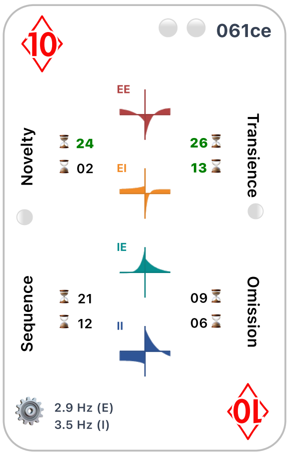
10
061ce
EE
EI
IE
II
Novelty 24 02
Transience 26 13
Sequence 21 12
Omission 09 06
2.9 Hz (E)
3.5 Hz (I)

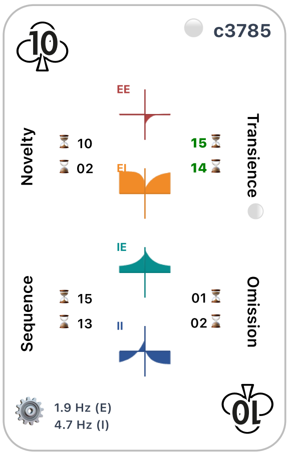
10
c3785
EE
EI
IE
II
Novelty 10 02
Transience 15 14
Sequence 15 13
Omission 01 02
1.9 Hz (E)
4.7 Hz (I)

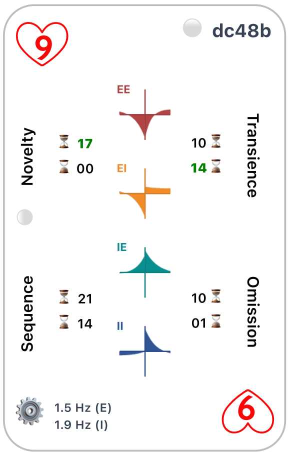
9
dc48b
EE
EI
IE
II
Novelty 17 00
Transience 10 14
Sequence 21 14
Omission 10 01
1.5 Hz (E)
1.9 Hz (I)

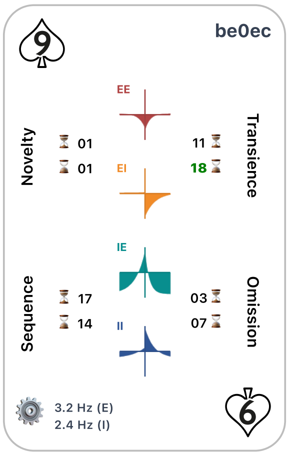
9
be0ec
EE
EI
IE
II
Novelty 01 01
Transience 11 18
Sequence 17 14
Omission 03 07
3.2 Hz (E)
2.4 Hz (I)

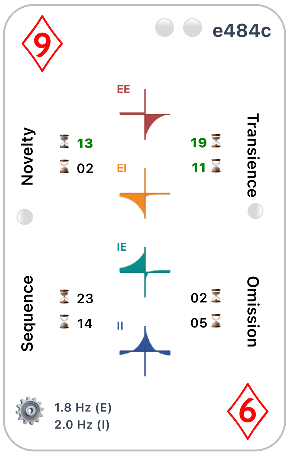
9
e484c
EE
EI
IE
II
Novelty 13 02
Transience 19 11
Sequence 23 14
Omission 02 05
1.8 Hz (E)
2.0 Hz (I)

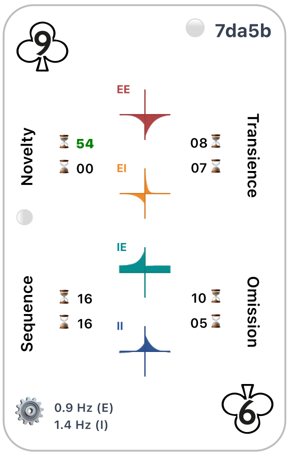
9
7da5b
EE
EI
IE
II
Novelty 54 00
Transience 08 07
Sequence 16 16
Omission 10 05
0.9 Hz (E)
1.4 Hz (I)

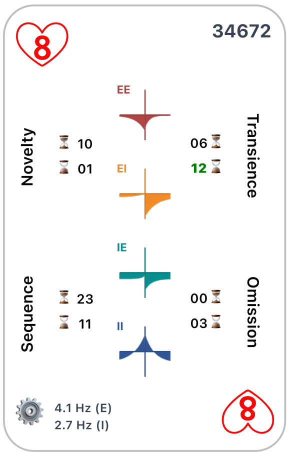
8
34672
EE
EI
IE
II
Novelty 10 01
Transience 06 12
Sequence 23 11
Omission 00 03
4.1 Hz (E)
2.7 Hz (I)

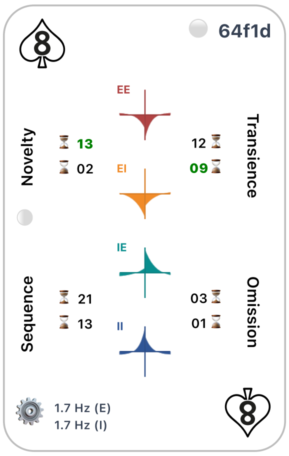
8
64f1d
EE
EI
IE
II
Novelty 13 02
Transience 12 09
Sequence 21 13
Omission 03 01
1.7 Hz (E)
1.7 Hz (I)

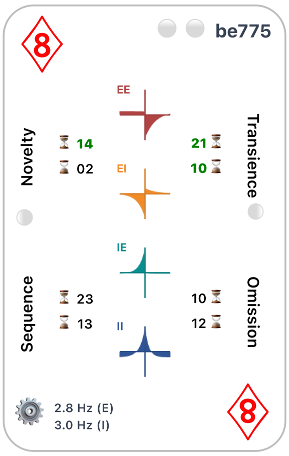
8
be775
EE
EI
IE
II
Novelty 14 02
Transience 21 10
Sequence 23 13
Omission 10 12
2.8 Hz (E)
3.0 Hz (I)

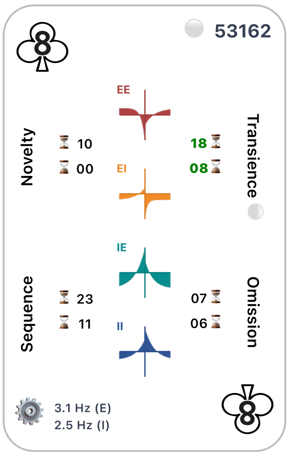
8
53162
EE
EI
IE
II
Novelty 10 00
Transience 18 08
Sequence 23 11
Omission 07 06
3.1 Hz (E)
2.5 Hz (I)

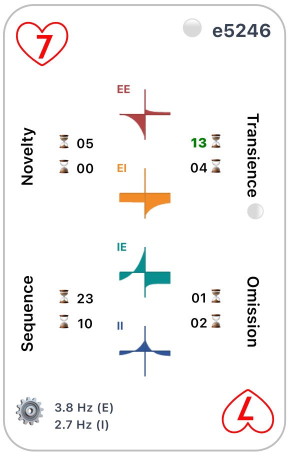
7
e5246
EE
EI
IE
II
Novelty 05 00
Transience 13 04
Sequence 23 10
Omission 01 02
3.8 Hz (E)
2.7 Hz (I)

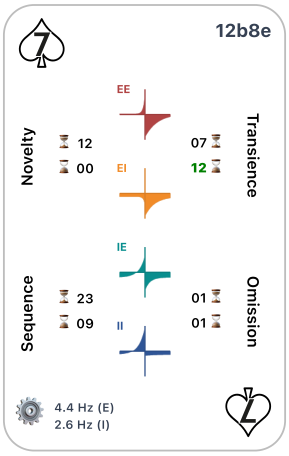
7
12b8e
EE
EI
IE
II
Novelty 12 00
Transience 07 12
Sequence 23 09
Omission 01 01
4.4 Hz (E)
2.6 Hz (I)

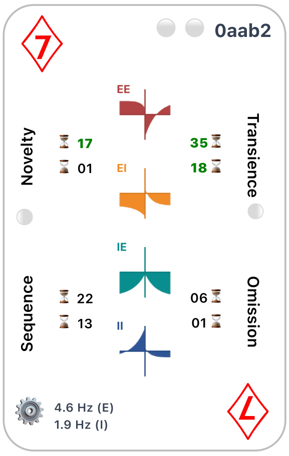
7
0aab2
EE
EI
IE
II
Novelty 17 01
Transience 35 18
Sequence 22 13
Omission 06 01
4.6 Hz (E)
1.9 Hz (I)

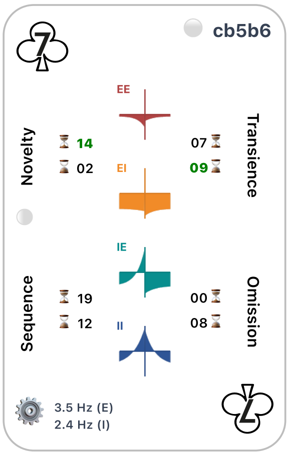
7
cb5b6
EE
EI
IE
II
Novelty 14 02
Transience 07 09
Sequence 19 12
Omission 00 08
3.5 Hz (E)
2.4 Hz (I)

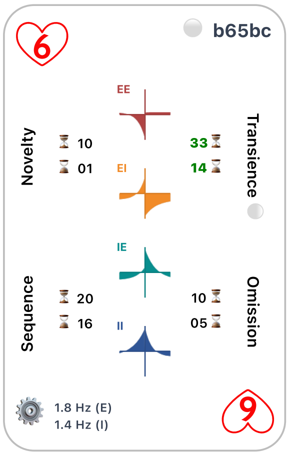
6
b65bc
EE
EI
IE
II
Novelty 10 01
Transience 33 14
Sequence 20 16
Omission 10 05
1.8 Hz (E)
1.4 Hz (I)

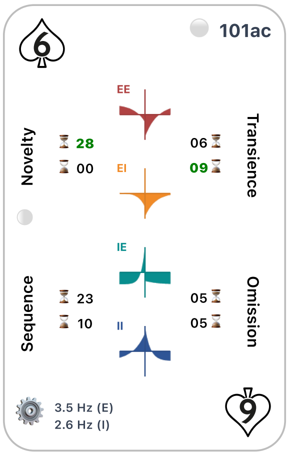
6
101ac
EE
EI
IE
II
Novelty 28 00
Transience 06 09
Sequence 23 10
Omission 05 05
3.5 Hz (E)
2.6 Hz (I)

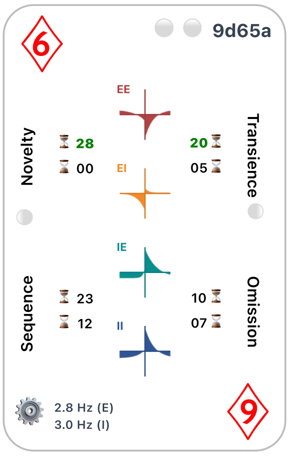
6
9d65a
EE
EI
IE
II
Novelty 28 00
Transience 20 05
Sequence 23 12
Omission 10 07
2.8 Hz (E)
3.0 Hz (I)

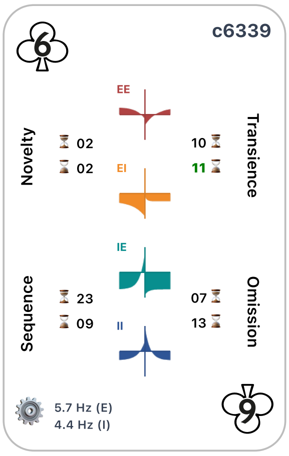
6
c6339
EE
EI
IE
II
Novelty 02 02
Transience 10 11
Sequence 23 09
Omission 07 13
5.7 Hz (E)
4.4 Hz (I)

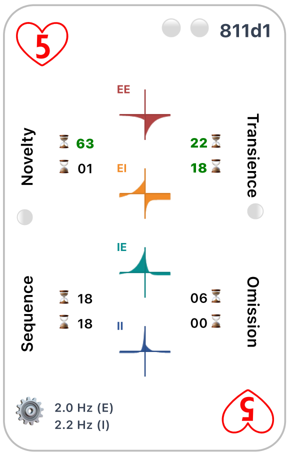
5
811d1
EE
EI
IE
II
Novelty 63 01
Transience 22 18
Sequence 18 18
Omission 06 00
2.0 Hz (E)
2.2 Hz (I)

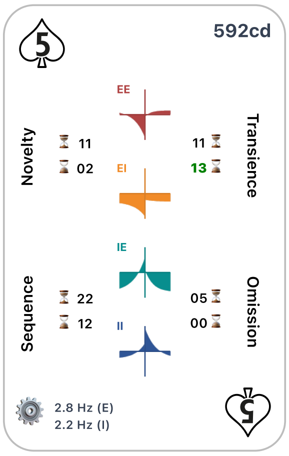
5
592cd
EE
EI
IE
II
Novelty 11 02
Transience 11 13
Sequence 22 12
Omission 05 00
2.8 Hz (E)
2.2 Hz (I)

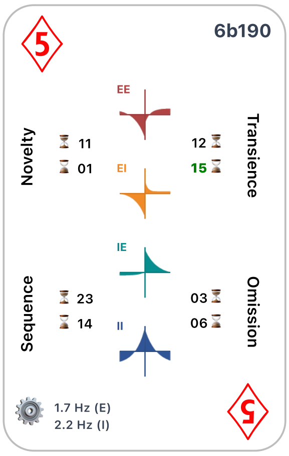
5
6b190
EE
EI
IE
II
Novelty 11 01
Transience 12 15
Sequence 23 14
Omission 03 06
1.7 Hz (E)
2.2 Hz (I)

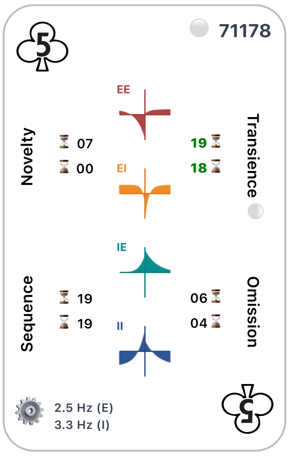
5
71178
EE
EI
IE
II
Novelty 07 00
Transience 19 18
Sequence 19 19
Omission 06 04
2.5 Hz (E)
3.3 Hz (I)

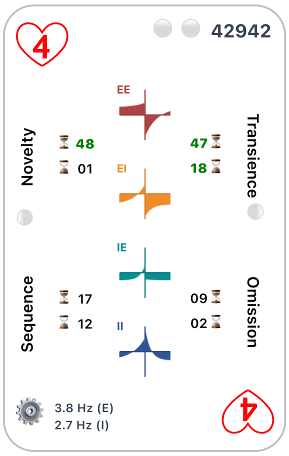
4
42942
EE
EI
IE
II
Novelty 48 01
Transience 47 18
Sequence 17 12
Omission 09 02
3.8 Hz (E)
2.7 Hz (I)

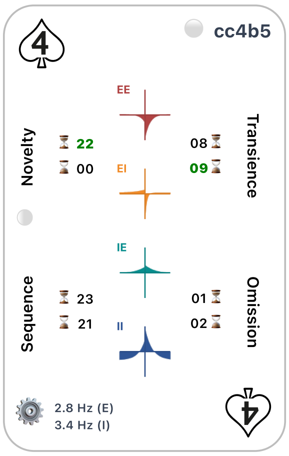
4
cc4b5
EE
EI
IE
II
Novelty 22 00
Transience 08 09
Sequence 23 21
Omission 01 02
2.8 Hz (E)
3.4 Hz (I)

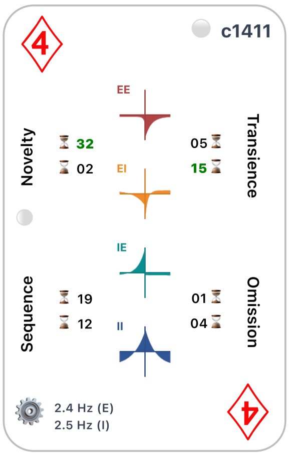
4
c1411
EE
EI
IE
II
Novelty 32 02
Transience 05 15
Sequence 19 12
Omission 01 04
2.4 Hz (E)
2.5 Hz (I)

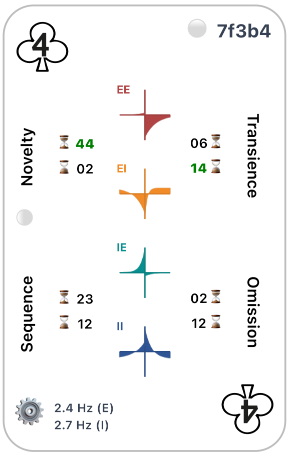
4
7f3b4
EE
EI
IE
II
Novelty 44 02
Transience 06 14
Sequence 23 12
Omission 02 12
2.4 Hz (E)
2.7 Hz (I)

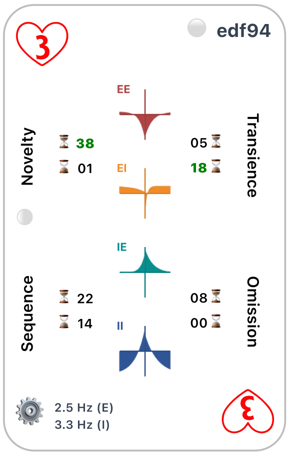
3
edf94
EE
EI
IE
II
Novelty 38 01
Transience 05 18
Sequence 22 14
Omission 08 00
2.5 Hz (E)
3.3 Hz (I)

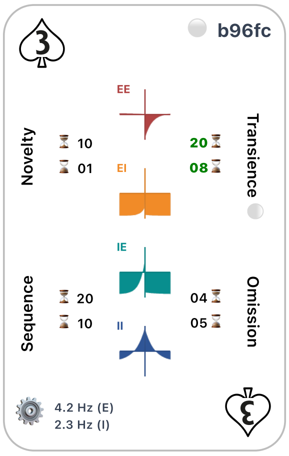
3
b96fc
EE
EI
IE
II
Novelty 10 01
Transience 20 08
Sequence 20 10
Omission 04 05
4.2 Hz (E)
2.3 Hz (I)

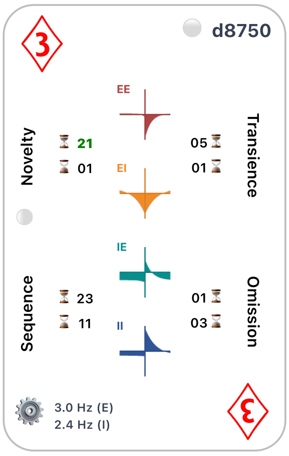
3
d8750
EE
EI
IE
II
Novelty 21 01
Transience 05 01
Sequence 23 11
Omission 01 03
3.0 Hz (E)
2.4 Hz (I)

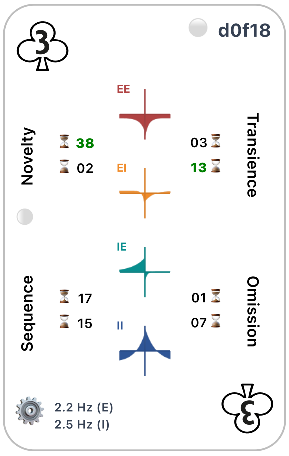
3
d0f18
EE
EI
IE
II
Novelty 38 02
Transience 03 13
Sequence 17 15
Omission 01 07
2.2 Hz (E)
2.5 Hz (I)

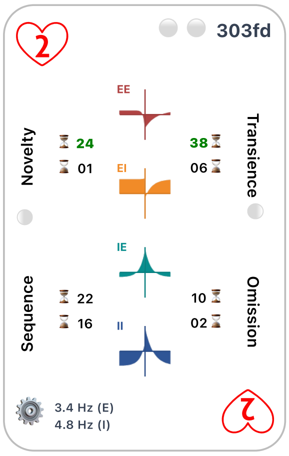
2
303fd
EE
EI
IE
II
Novelty 24 01
Transience 38 06
Sequence 22 16
Omission 10 02
3.4 Hz (E)
4.8 Hz (I)

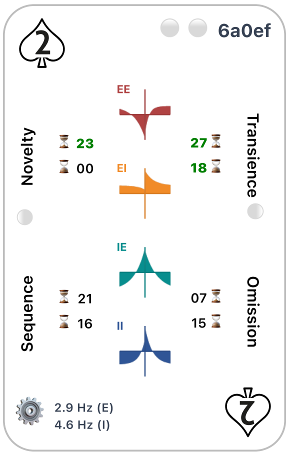
2
6a0ef
EE
EI
IE
II
Novelty 23 00
Transience 27 18
Sequence 21 16
Omission 07 15
2.9 Hz (E)
4.6 Hz (I)

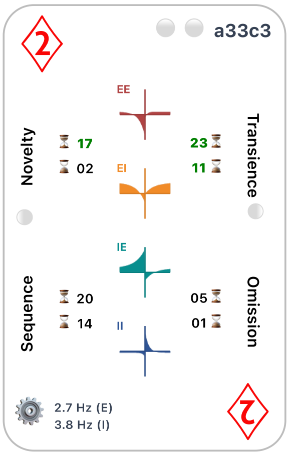
2
a33c3
EE
EI
IE
II
Novelty 17 02
Transience 23 11
Sequence 20 14
Omission 05 01
2.7 Hz (E)
3.8 Hz (I)

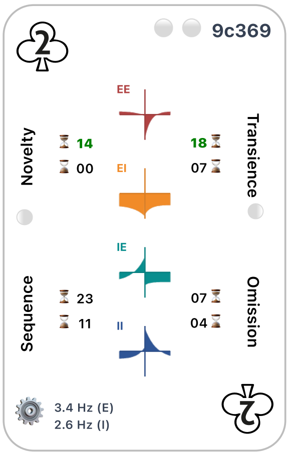
2
9c369
EE
EI
IE
II
Novelty 14 00
Transience 18 07
Sequence 23 11
Omission 07 04
3.4 Hz (E)
2.6 Hz (I)

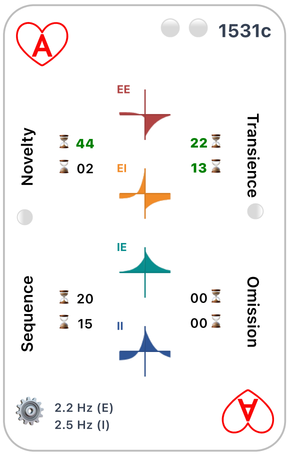
A
1531c
EE
EI
IE
II
Novelty 44 02
Transience 22 13
Sequence 20 15
Omission 00 00
2.2 Hz (E)
2.5 Hz (I)

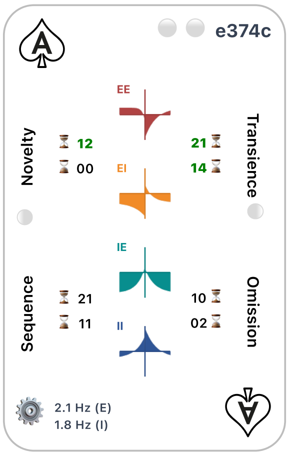
A
e374c
EE
EI
IE
II
Novelty 12 00
Transience 21 14
Sequence 21 11
Omission 10 02
2.1 Hz (E)
1.8 Hz (I)

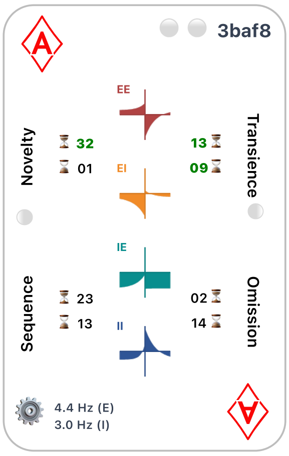
A
3baf8
EE
EI
IE
II
Novelty 32 01
Transience 13 09
Sequence 23 13
Omission 02 14
4.4 Hz (E)
3.0 Hz (I)

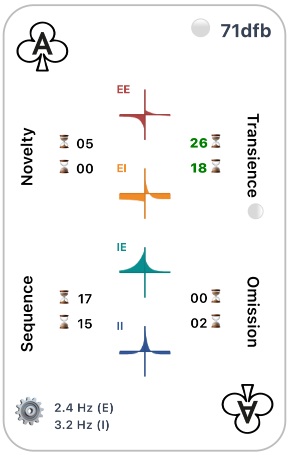
A
71dfb
EE
EI
IE
II
Novelty 05 00
Transience 26 18
Sequence 17 15
Omission 00 02
2.4 Hz (E)
3.2 Hz (I)

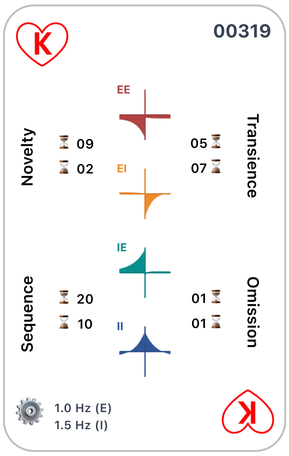
K
00319
EE
Novelty
09
02
05
07
Transience
EI
IE
Sequence
20
10
01
01
Omission
II
1.0 Hz (E)
1.5 Hz (I)

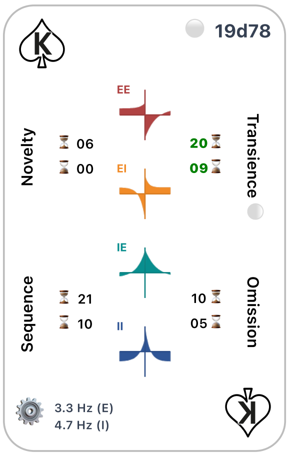
K
19d78
EE
Novelty
06
00
20
09
Transience
EI
IE
Sequence
21
10
10
05
Omission
II
3.3 Hz (E)
4.7 Hz (I)

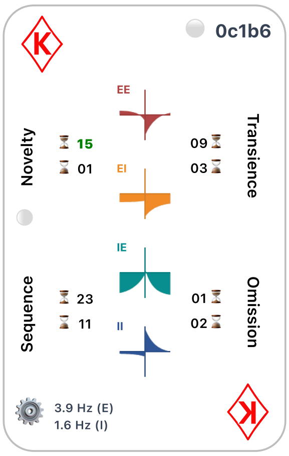
K
0c1b6
EE
Novelty
15
01
09
03
Transience
EI
IE
Sequence
23
11
01
02
Omission
II
3.9 Hz (E)
1.6 Hz (I)

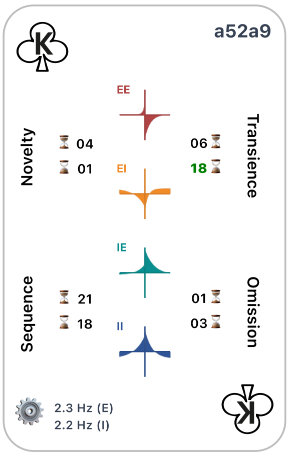
K
a52a9
EE
Novelty
04
01
06
18
Transience
EI
IE
Sequence
21
18
01
03
Omission
II
2.3 Hz (E)
2.2 Hz (I)

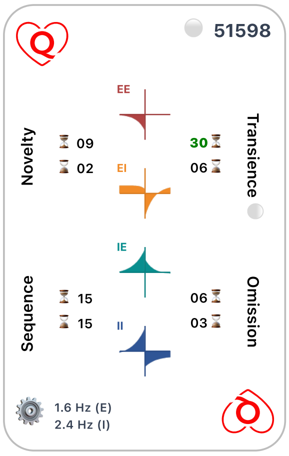
Q
51598
EE
Novelty
09
02
30
06
Transience
EI
IE
Sequence
15
15
06
03
Omission
II
1.6 Hz (E)
2.4 Hz (I)

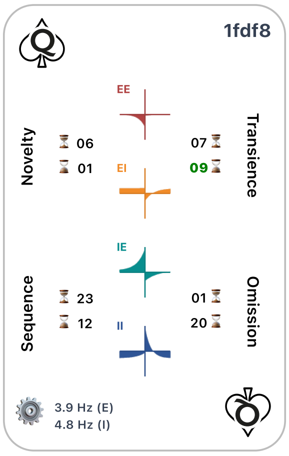
Q
1fdf8
EE
Novelty
06
01
07
09
Transience
EI
IE
Sequence
23
12
01
20
Omission
II
3.9 Hz (E)
4.8 Hz (I)

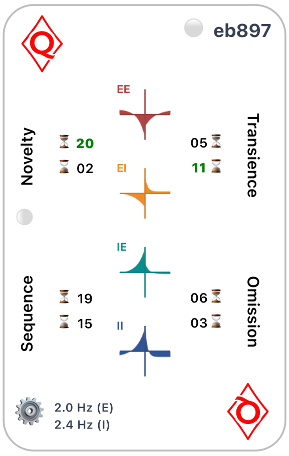
Q
eb897
EE
Novelty
20
02
05
11
Transience
EI
IE
Sequence
19
15
06
03
Omission
II
2.0 Hz (E)
2.4 Hz (I)

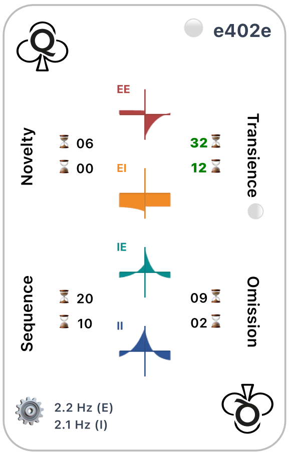
Q
e402e
EE
Novelty
06
00
32
12
Transience
EI
IE
Sequence
20
10
09
02
Omission
II
2.2 Hz (E)
2.1 Hz (I)

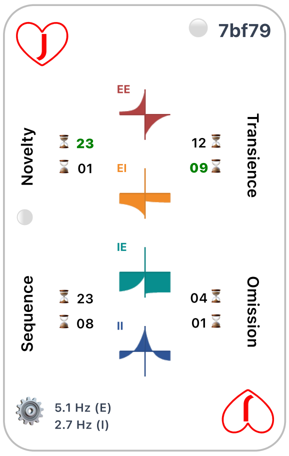
J
7bf79
EE
Novelty
23
01
12
09
Transience
EI
IE
Sequence
23
08
04
01
Omission
II
5.1 Hz (E)
2.7 Hz (I)

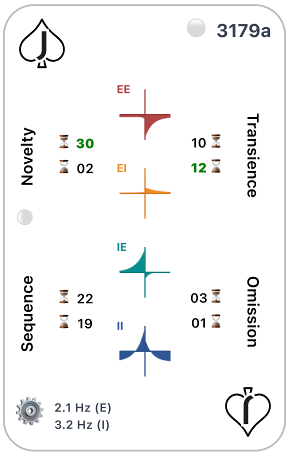
J
3179a
EE
Novelty
30
02
10
12
Transience
EI
IE
Sequence
22
19
03
01
Omission
II
2.1 Hz (E)
3.2 Hz (I)

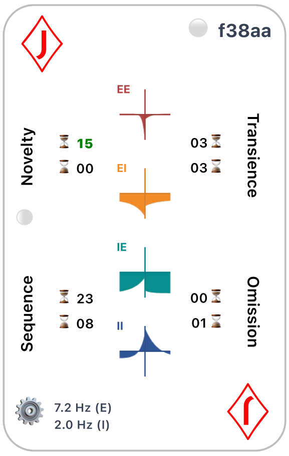
J
f38aa
EE
Novelty
15
00
03
03
Transience
EI
IE
Sequence
23
08
00
01
Omission
II
7.2 Hz (E)
2.0 Hz (I)

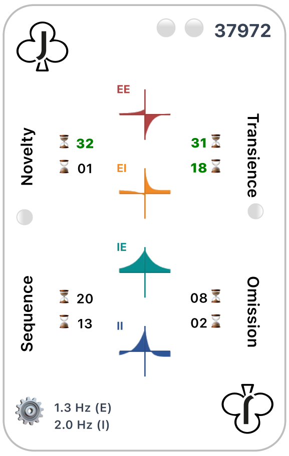
J
37972
EE
Novelty
32
01
31
18
Transience
EI
IE
Sequence
20
13
08
02
Omission
II
1.3 Hz (E)
2.0 Hz (I)

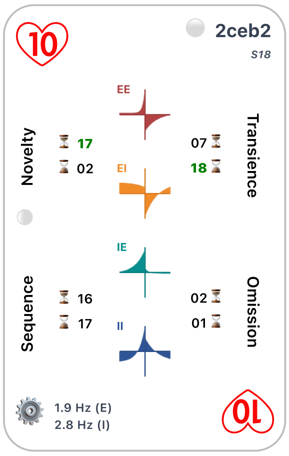
10
2ceb2
S18
EE
Novelty
17
02
07
18
Transience
EI
IE
Sequence
16
17
02
01
Omission
II
1.9 Hz (E)
2.8 Hz (I)

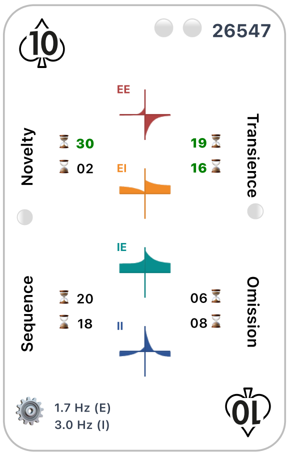
10
26547
EE
Novelty
30
02
19
16
Transience
EI
IE
Sequence
20
18
06
08
Omission
II
1.7 Hz (E)
3.0 Hz (I)

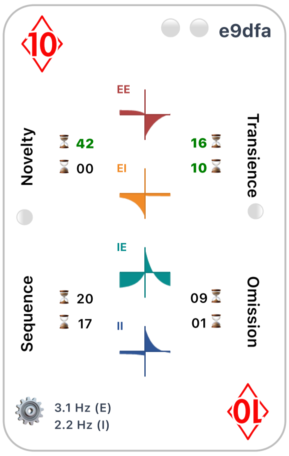
10
e9dfa
EE
Novelty
42
00
16
10
Transience
EI
IE
Sequence
20
17
09
01
Omission
II
3.1 Hz (E)
2.2 Hz (I)

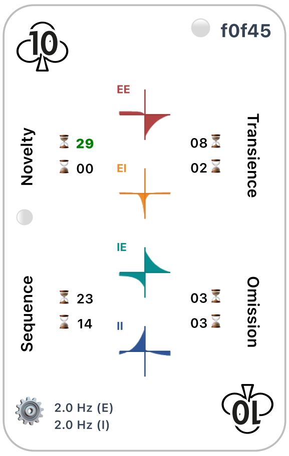
10
f0f45
EE
Novelty
29
00
08
02
Transience
EI
IE
Sequence
23
14
03
03
Omission
II
2.0 Hz (E)
2.0 Hz (I)

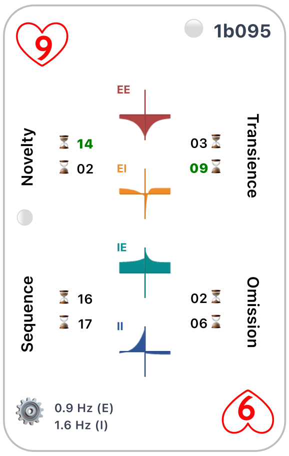
9
1b095
EE
EI
IE
II
Novelty
14
02
Transience
03
09
Sequence
16
17
Omission
02
06
0.9 Hz (E)
1.6 Hz (I)

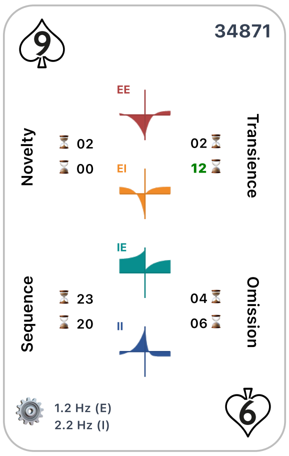
9
34871
EE
EI
IE
II
Novelty
02
00
Transience
02
12
Sequence
23
20
Omission
04
06
1.2 Hz (E)
2.2 Hz (I)

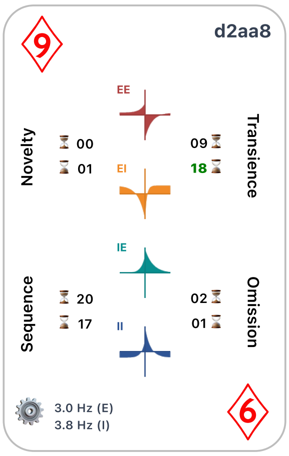
9
d2aa8
EE
EI
IE
II
Novelty
00
01
Transience
09
18
Sequence
20
17
Omission
02
01
3.0 Hz (E)
3.8 Hz (I)

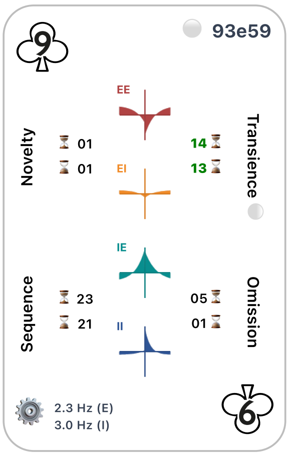
9
93e59
EE
EI
IE
II
Novelty
01
01
Transience
14
13
Sequence
23
21
Omission
05
01
2.3 Hz (E)
3.0 Hz (I)

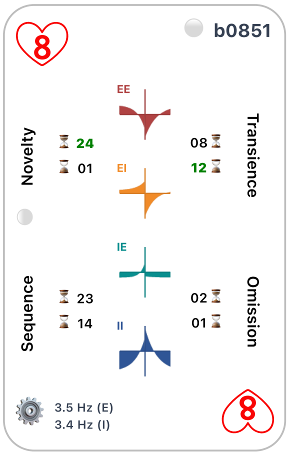
8
b0851
EE
EI
IE
II
Novelty
24
01
Transience
08
12
Sequence
23
14
Omission
02
01
3.5 Hz (E)
3.4 Hz (I)

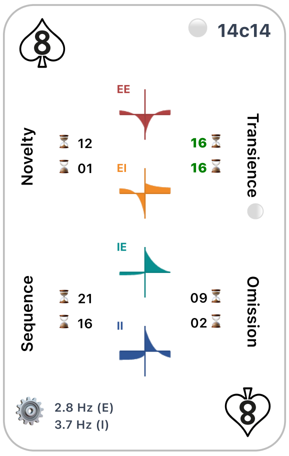
8
14c14
EE
EI
IE
II
Novelty
12
01
Transience
16
16
Sequence
21
16
Omission
09
02
2.8 Hz (E)
3.7 Hz (I)

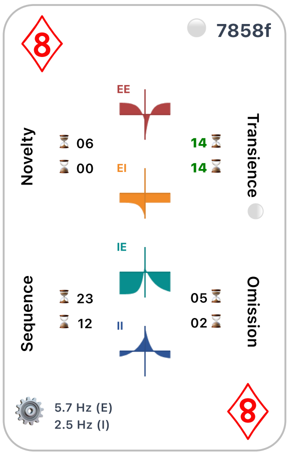
8
7858f
EE
EI
IE
II
Novelty
06
00
Transience
14
14
Sequence
23
12
Omission
05
02
5.7 Hz (E)
2.5 Hz (I)

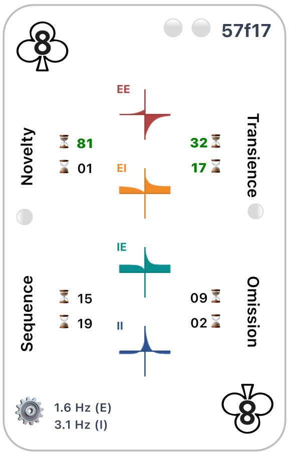
8
57f17
EE
EI
IE
II
Novelty
81
01
Transience
32
17
Sequence
15
19
Omission
09
02
1.6 Hz (E)
3.1 Hz (I)

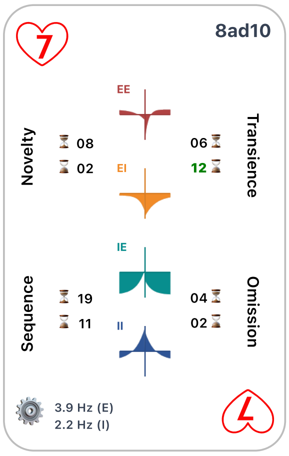
7
8ad10
EE
EI
IE
II
Novelty
08
02
Transience
06
12
Sequence
19
11
Omission
04
02
3.9 Hz (E)
2.2 Hz (I)

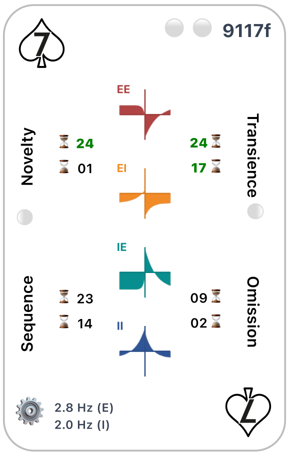
7
9117f
EE
EI
IE
II
Novelty
24
01
Transience
24
17
Sequence
23
14
Omission
09
02
2.8 Hz (E)
2.0 Hz (I)

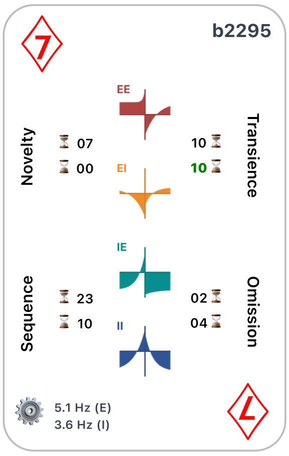
7
b2295
EE
EI
IE
II
Novelty
07
00
Transience
10
10
Sequence
23
10
Omission
02
04
5.1 Hz (E)
3.6 Hz (I)

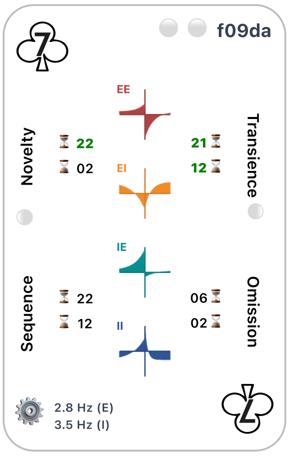
7
f09da
EE
EI
IE
II
Novelty
22
02
Transience
21
12
Sequence
22
12
Omission
06
02
2.8 Hz (E)
3.5 Hz (I)

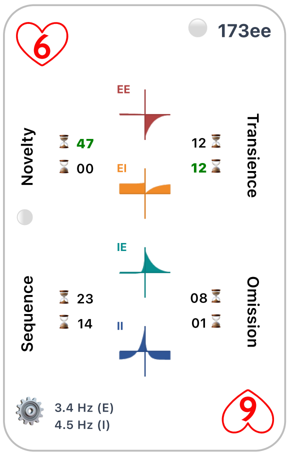
6
173ee
EE
EI
IE
II
Novelty
47
00
Transience
12
12
Sequence
23
14
Omission
08
01
3.4 Hz (E)
4.5 Hz (I)

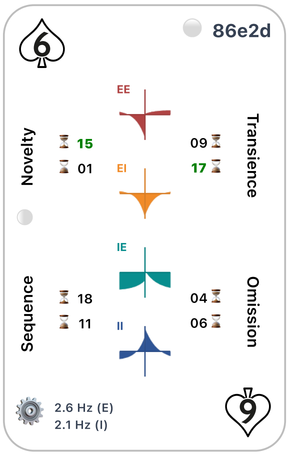
6
86e2d
EE
EI
IE
II
Novelty
15
01
Transience
09
17
Sequence
18
11
Omission
04
06
2.6 Hz (E)
2.1 Hz (I)

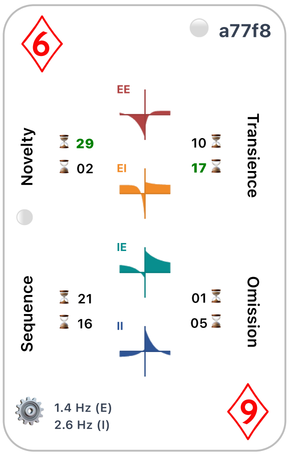
6
a77f8
EE
EI
IE
II
Novelty
29
02
Transience
10
17
Sequence
21
16
Omission
01
05
1.4 Hz (E)
2.6 Hz (I)

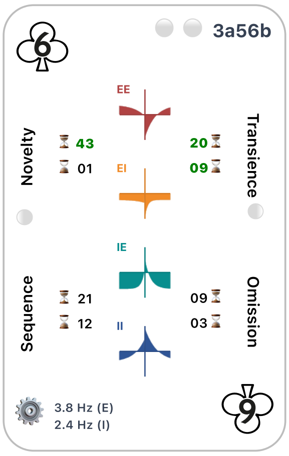
6
3a56b
EE
EI
IE
II
Novelty
43
01
Transience
20
09
Sequence
21
12
Omission
09
03
3.8 Hz (E)
2.4 Hz (I)

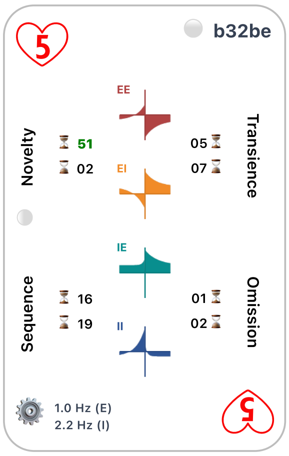
5
b32ebe
EE
Novelty
51
02
05
07
Transience
EI
IE
Sequence
16
19
01
02
Omission
II
1.0 Hz (E)
2.2 Hz (I)

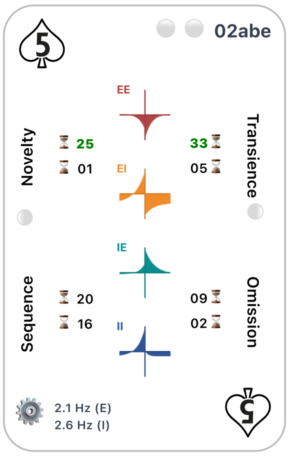
5
02abe
EE
Novelty
25
01
33
05
Transience
EI
IE
Sequence
20
16
09
02
Omission
II
2.1 Hz (E)
2.6 Hz (I)

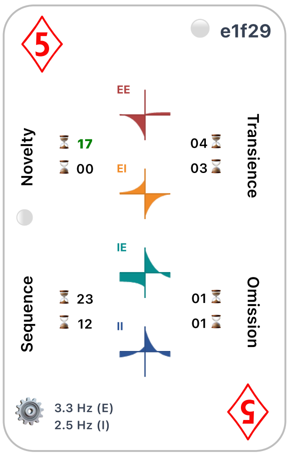
5
e1f29
EE
Novelty
17
00
04
03
Transience
EI
IE
Sequence
23
12
01
01
Omission
II
3.3 Hz (E)
2.5 Hz (I)

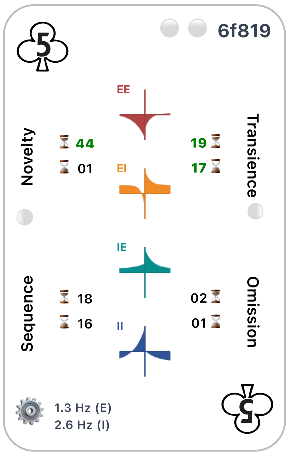
5
6f819
EE
Novelty
44
01
19
17
Transience
EI
IE
Sequence
18
16
02
01
Omission
II
1.3 Hz (E)
2.6 Hz (I)

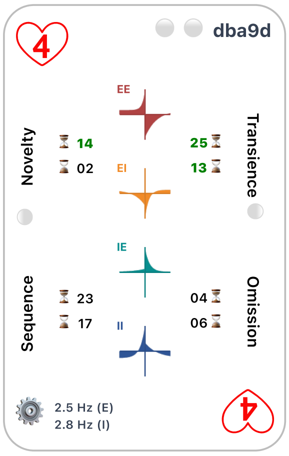
4
dba9d
EE
Novelty
14
02
25
13
Transience
EI
IE
Sequence
23
17
04
06
Omission
II
2.5 Hz (E)
2.8 Hz (I)

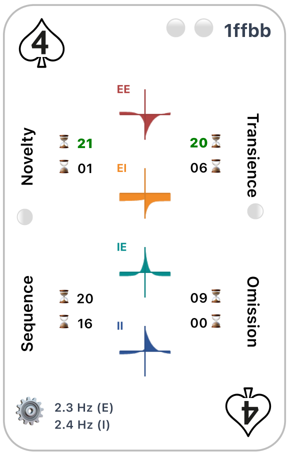
4
1ffbb
EE
Novelty
21
01
20
06
Transience
EI
IE
Sequence
20
16
09
00
Omission
II
2.3 Hz (E)
2.4 Hz (I)

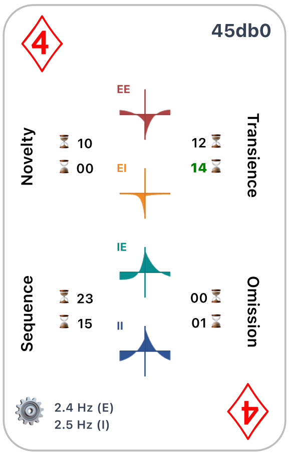
4
45db0
EE
Novelty
10
00
12
14
Transience
EI
IE
Sequence
23
15
00
01
Omission
II
2.4 Hz (E)
2.5 Hz (I)

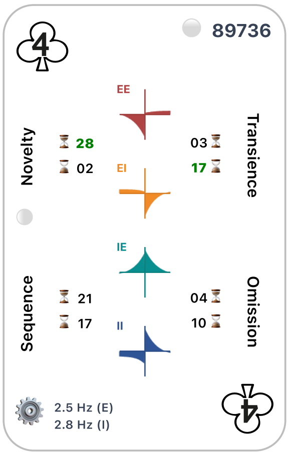
4
89736
EE
Novelty
28
02
03
17
Transience
EI
IE
Sequence
21
17
04
10
Omission
II
2.5 Hz (E)
2.8 Hz (I)

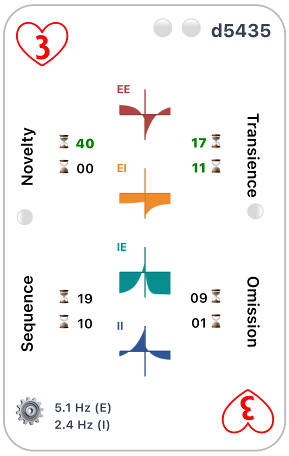
3
d5435
EE
Novelty
40
00
17
11
Transience
EI
IE
Sequence
19
10
09
01
Omission
II
5.1 Hz (E)
2.4 Hz (I)

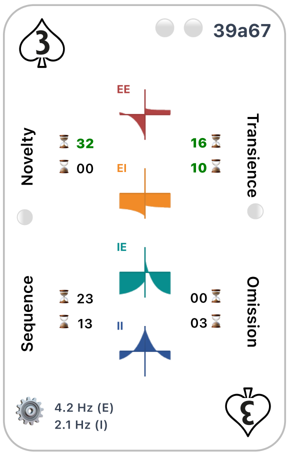
3
39a67
EE
Novelty
32
00
16
10
Transience
EI
IE
Sequence
23
13
00
03
Omission
II
4.2 Hz (E)
2.1 Hz (I)

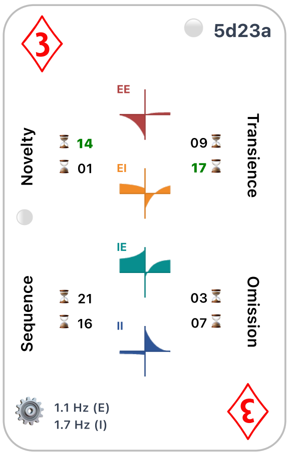
3
5d23a
EE
Novelty
14
01
09
17
Transience
EI
IE
Sequence
21
16
03
07
Omission
II
1.1 Hz (E)
1.7 Hz (I)

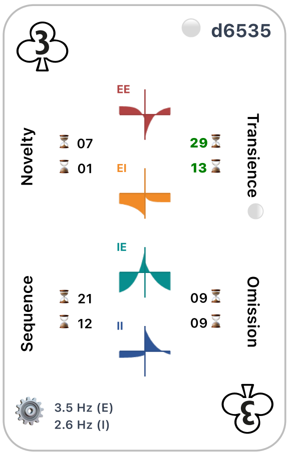
3
d6535
EE
Novelty
07
01
29
13
Transience
EI
IE
Sequence
21
12
09
09
Omission
II
3.5 Hz (E)
2.6 Hz (I)

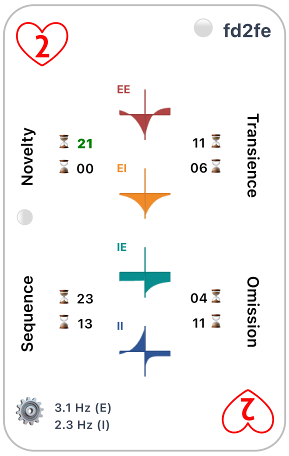
2
fd2fe
EE
Novelty
21
00
11
06
Transience
EI
IE
Sequence
23
13
04
11
Omission
II
3.1 Hz (E)
2.3 Hz (I)

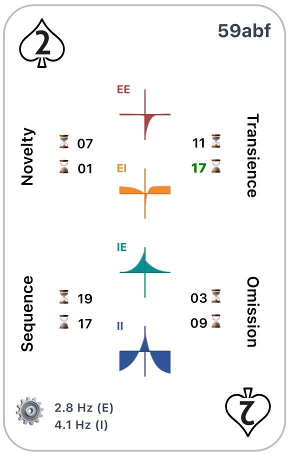
2
59abf
EE
Novelty
07
01
11
17
Transience
EI
IE
Sequence
19
17
03
09
Omission
II
2.8 Hz (E)
4.1 Hz (I)

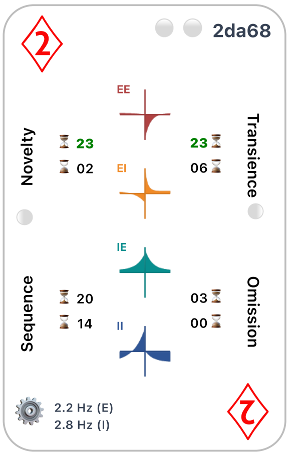
2
2da68
EE
Novelty
23
02
23
06
Transience
EI
IE
Sequence
20
14
03
00
Omission
II
2.2 Hz (E)
2.8 Hz (I)

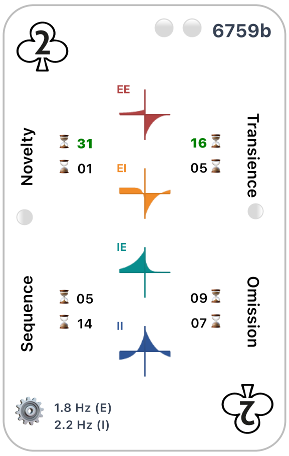
2
6759b
EE
Novelty
31
01
16
05
Transience
EI
IE
Sequence
05
14
09
07
Omission
II
1.8 Hz (E)
2.2 Hz (I)

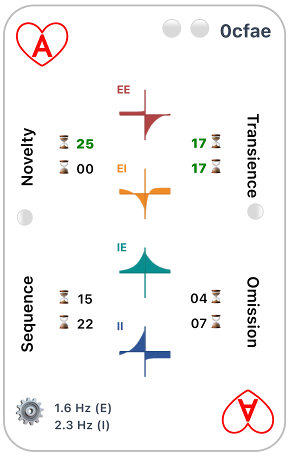
A
0cfae
Novelty
25
00
Transience
17
17
EE
EI
IE
II
Sequence
15
22
Omission
04
07
1.6 Hz (E)
2.3 Hz (I)

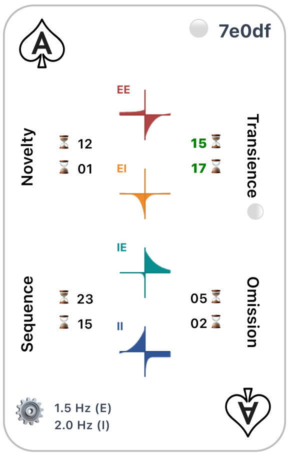
A
7e0df
Novelty
12
01
Transience
15
17
EE
EI
IE
II
Sequence
23
15
Omission
05
02
1.5 Hz (E)
2.0 Hz (I)

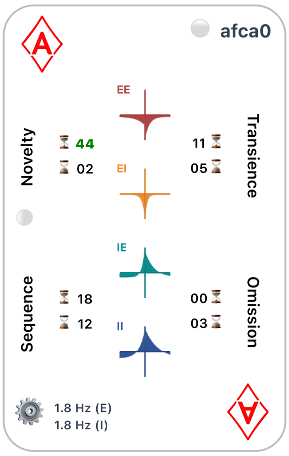
A
afca0
Novelty
44
02
Transience
11
05
EE
EI
IE
II
Sequence
18
12
Omission
00
03
1.8 Hz (E)
1.8 Hz (I)

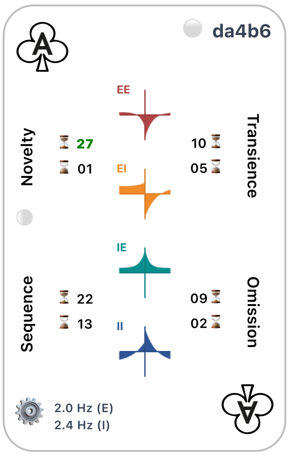
A
da4b6
Novelty
27
01
Transience
10
05
EE
EI
IE
II
Sequence
22
13
Omission
09
02
2.0 Hz (E)
2.4 Hz (I)

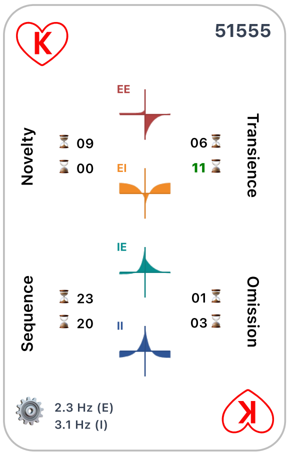
K
51555
Novelty
09
00
Transience
06
11
EE
EI
IE
II
Sequence
23
20
Omission
01
03
2.3 Hz (E)
3.1 Hz (I)

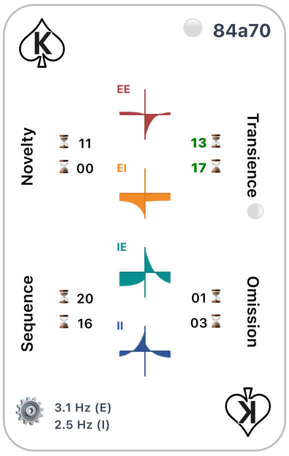
K
84a70
Novelty
11
00
Transience
13
17
EE
EI
IE
II
Sequence
20
16
Omission
01
03
3.1 Hz (E)
2.5 Hz (I)

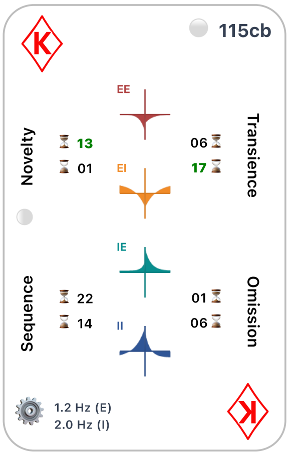
K
115cb
Novelty
13
01
Transience
06
17
EE
EI
IE
II
Sequence
22
14
Omission
01
06
1.2 Hz (E)
2.0 Hz (I)

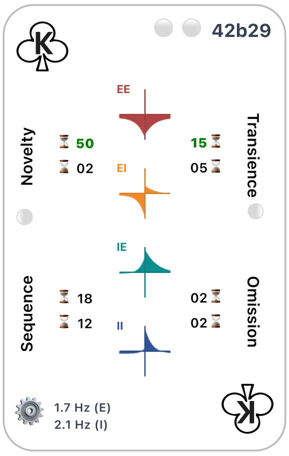
K
42b29
Novelty
50
02
Transience
15
05
EE
EI
IE
II
Sequence
18
12
Omission
02
02
1.7 Hz (E)
2.1 Hz (I)

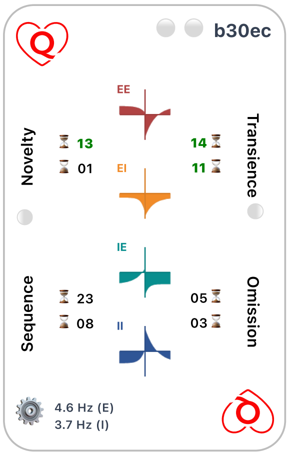
Q
b30ec
Novelty
13
01
Transience
14
11
EE
EI
IE
II
Sequence
23
08
Omission
05
03
4.6 Hz (E)
3.7 Hz (I)

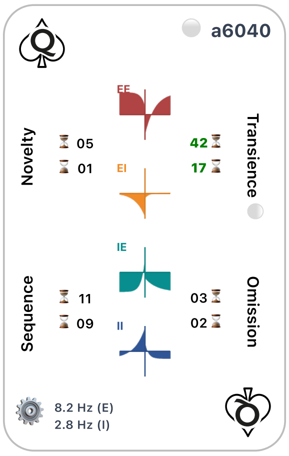
Q
a6040
Novelty
05
01
Transience
42
17
EE
EI
IE
II
Sequence
11
09
Omission
03
02
8.2 Hz (E)
2.8 Hz (I)

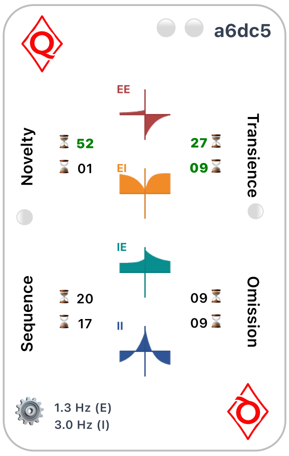
Q
a6dc5
Novelty
52
01
Transience
27
09
EE
EI
IE
II
Sequence
20
17
Omission
09
09
1.3 Hz (E)
3.0 Hz (I)

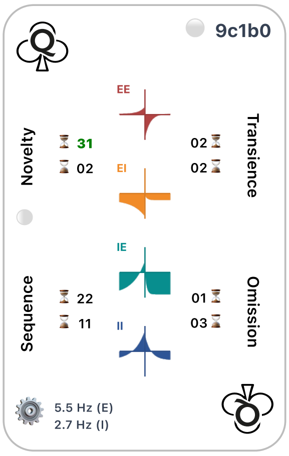
Q
9c1b0
Novelty
31
02
Transience
02
02
EE
EI
IE
II
Sequence
22
11
Omission
01
03
5.5 Hz (E)
2.7 Hz (I)

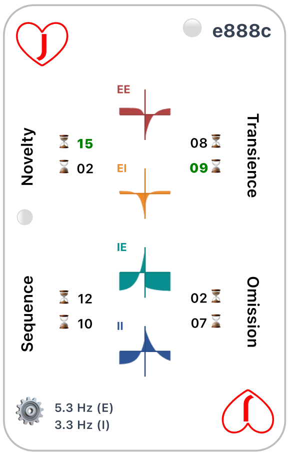
J
e888c
Novelty
15
02
Transience
08
09
EE
EI
IE
II
Sequence
12
10
Omission
02
07
5.3 Hz (E)
3.3 Hz (I)

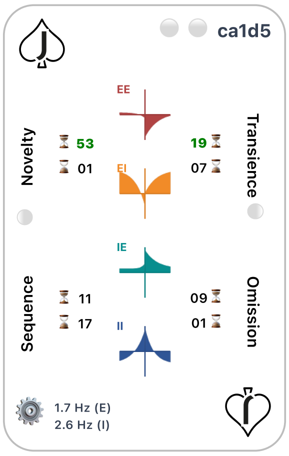
J
ca1d5
Novelty
53
01
Transience
19
07
EE
EI
IE
II
Sequence
11
17
Omission
09
01
1.7 Hz (E)
2.6 Hz (I)

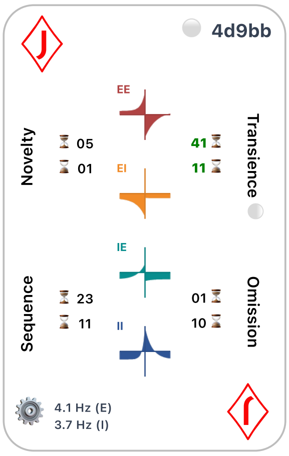
J
4d9bb
Novelty
05
01
Transience
41
11
EE
EI
IE
II
Sequence
23
11
Omission
01
10
4.1 Hz (E)
3.7 Hz (I)

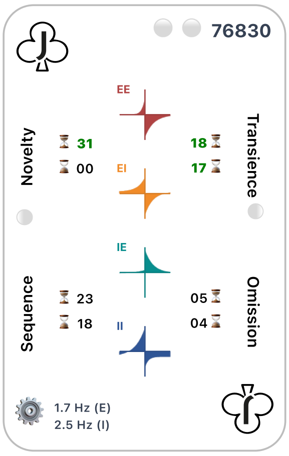
J
76830
Novelty
31
00
Transience
18
17
EE
EI
IE
II
Sequence
23
18
Omission
05
04
1.7 Hz (E)
2.5 Hz (I)

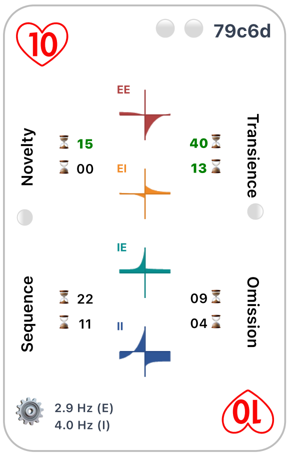
10
79c6d
EE
EI
IE
II
Novelty
15
00
Transience
40
13
Sequence
22
11
Omission
09
04
2.9 Hz (E)
4.0 Hz (I)

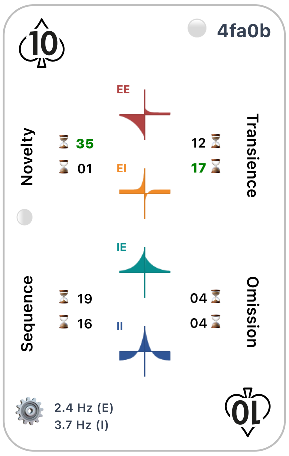
10
4fa0b
EE
EI
IE
II
Novelty
35
01
Transience
12
17
Sequence
19
16
Omission
04
04
2.4 Hz (E)
3.7 Hz (I)

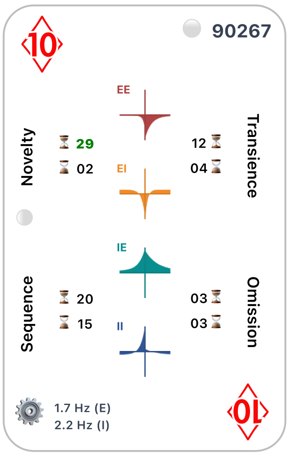
10
90267
EE
EI
IE
II
Novelty
29
02
Transience
12
04
Sequence
20
15
Omission
03
03
1.7 Hz (E)
2.2 Hz (I)

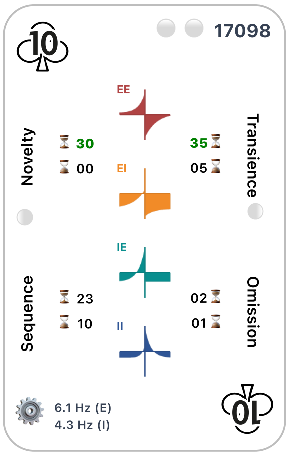
10
17098
EE
EI
IE
II
Novelty
30
00
Transience
35
05
Sequence
23
10
Omission
02
01
6.1 Hz (E)
4.3 Hz (I)

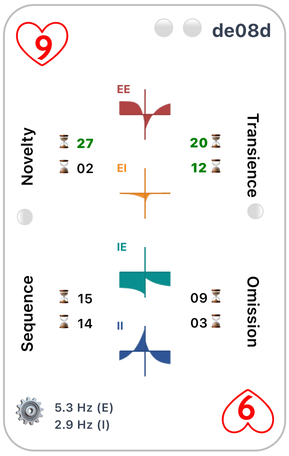
9
de08d
EE
EI
IE
II
Novelty
27
02
Transience
20
12
Sequence
15
14
Omission
09
03
5.3 Hz (E)
2.9 Hz (I)

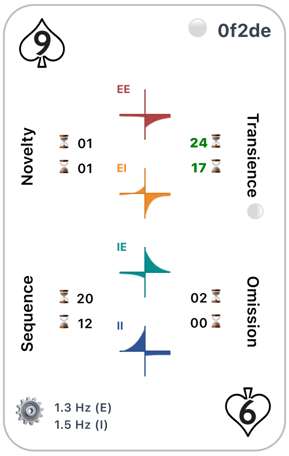
9
0f2de
EE
EI
IE
II
Novelty
01
01
Transience
24
17
Sequence
20
12
Omission
02
00
1.3 Hz (E)
1.5 Hz (I)

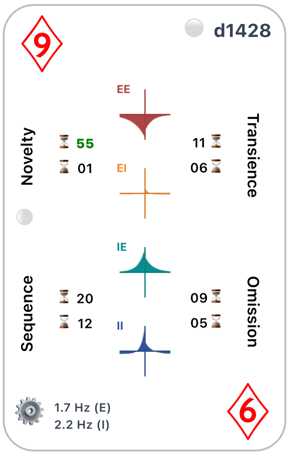
9
d1428
EE
EI
IE
II
Novelty
55
01
Transience
11
06
Sequence
20
12
Omission
09
05
1.7 Hz (E)
2.2 Hz (I)

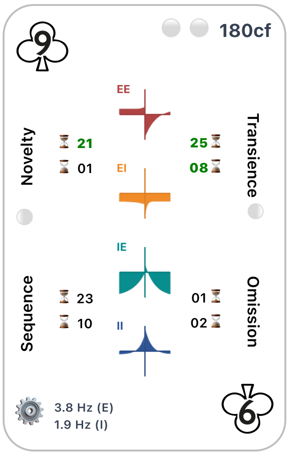
9
180cf
EE
EI
IE
II
Novelty
21
01
Transience
25
08
Sequence
23
10
Omission
01
02
3.8 Hz (E)
1.9 Hz (I)

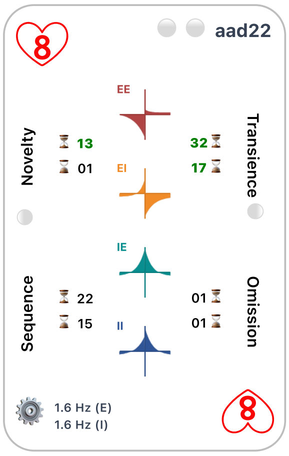
8
aad22
EE
EI
IE
II
Novelty
13
01
Transience
32
17
Sequence
22
15
Omission
01
01
1.6 Hz (E)
1.6 Hz (I)

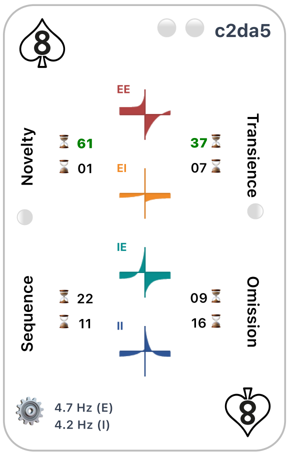
8
c2da5
EE
EI
IE
II
Novelty
61
01
Transience
37
07
Sequence
22
11
Omission
09
16
4.7 Hz (E)
4.2 Hz (I)

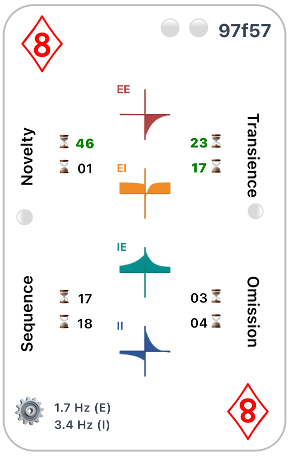
8
97f57
EE
EI
IE
II
Novelty
46
01
Transience
23
17
Sequence
17
18
Omission
03
04
1.7 Hz (E)
3.4 Hz (I)

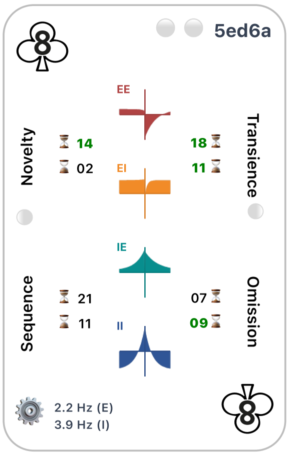
8
5ed6a
EE
EI
IE
II
Novelty
14
02
Transience
18
11
Sequence
21
11
Omission
07
09
2.2 Hz (E)
3.9 Hz (I)

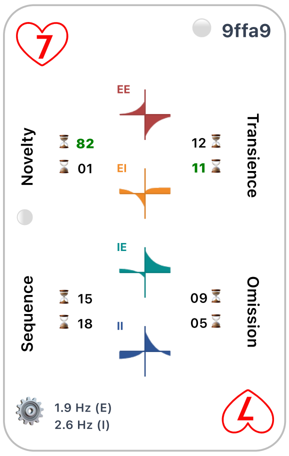
7
9ffa9
EE
EI
IE
II
Novelty
82
01
Transience
12
11
Sequence
15
18
Omission
09
05
1.9 Hz (E)
2.6 Hz (I)

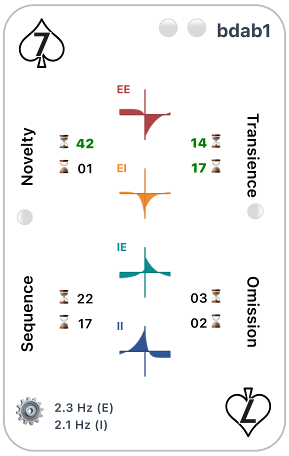
7
bdab1
EE
EI
IE
II
Novelty
42
01
Transience
14
17
Sequence
22
17
Omission
03
02
2.3 Hz (E)
2.1 Hz (I)

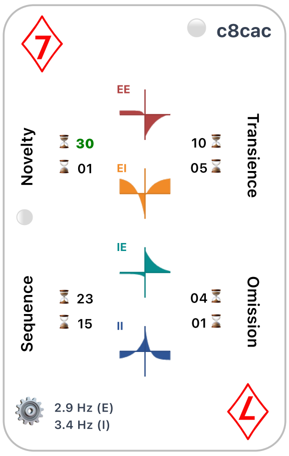
7
c8cac
EE
EI
IE
II
Novelty
30
01
Transience
10
05
Sequence
23
15
Omission
04
01
2.9 Hz (E)
3.4 Hz (I)

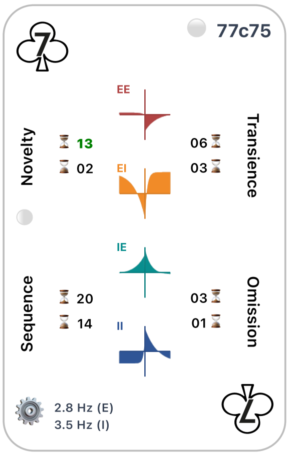
7
77c75
EE
EI
IE
II
Novelty
13
02
Transience
06
03
Sequence
20
14
Omission
03
01
2.8 Hz (E)
3.5 Hz (I)

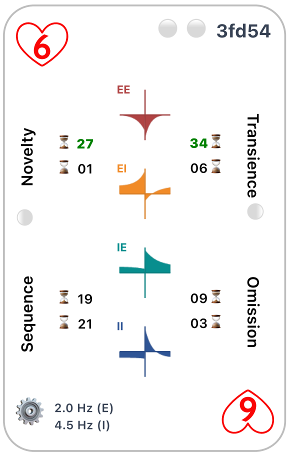
6
3fd54
EE
EI
IE
II
Novelty
27
01
Transience
34
06
Sequence
19
21
Omission
09
03
2.0 Hz (E)
4.5 Hz (I)
9

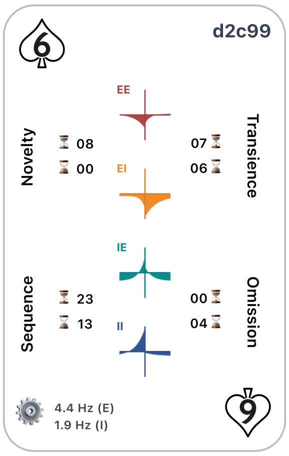
6
d2c99
EE
EI
IE
II
Novelty
08
00
Transience
07
06
Sequence
23
13
Omission
00
04
4.4 Hz (E)
1.9 Hz (I)
9

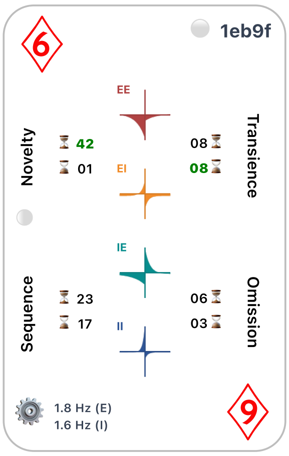
6
1eb9f
EE
EI
IE
II
Novelty
42
01
Transience
08
08
Sequence
23
17
Omission
06
03
1.8 Hz (E)
1.6 Hz (I)
9

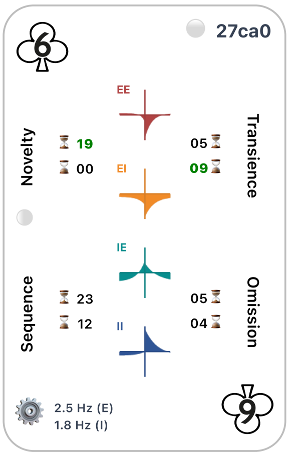
6
27ca0
EE
EI
IE
II
Novelty
19
00
Transience
05
09
Sequence
23
12
Omission
05
04
2.5 Hz (E)
1.8 Hz (I)
9

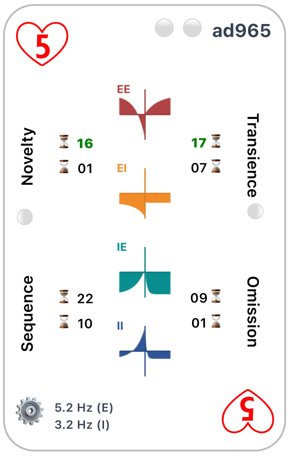
5
ad965
EE
EI
IE
II
Novelty
16
01
Transience
17
07
Sequence
22
10
Omission
09
01
5.2 Hz (E)
3.2 Hz (I)
5

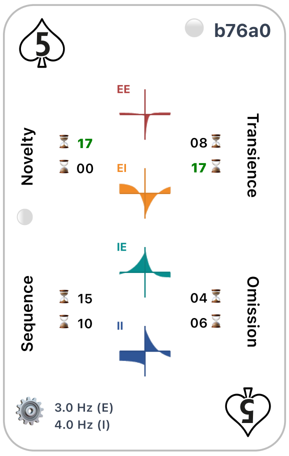
5
b76a0
EE
EI
IE
II
Novelty
17
00
Transience
08
17
Sequence
15
10
Omission
04
06
3.0 Hz (E)
4.0 Hz (I)
5

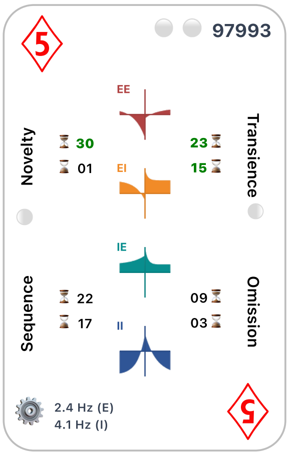
5
97993
EE
EI
IE
II
Novelty
30
01
Transience
23
15
Sequence
22
17
Omission
09
03
2.4 Hz (E)
4.1 Hz (I)
5

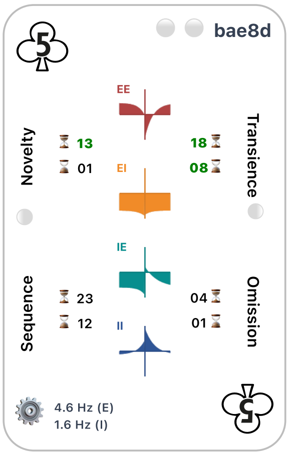
5
bae8d
EE
EI
IE
II
Novelty
13
01
Transience
18
08
Sequence
23
12
Omission
04
01
4.6 Hz (E)
1.6 Hz (I)
5

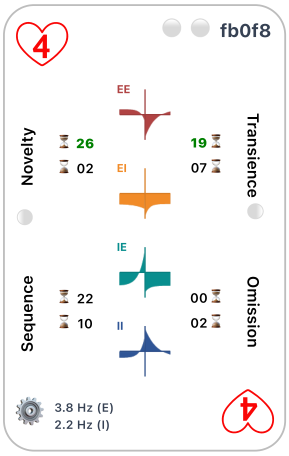
4
fb0f8
EE
EI
IE
II
Novelty
26
02
Transience
19
07
Sequence
22
10
Omission
00
02
3.8 Hz (E)
2.2 Hz (I)
4

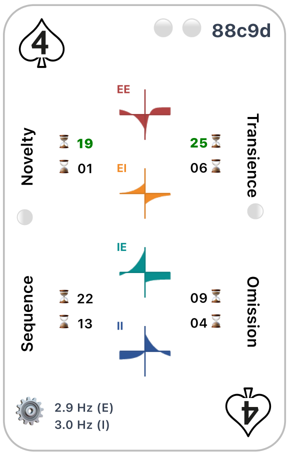
4
88c9d
EE
EI
IE
II
Novelty
19
01
Transience
25
06
Sequence
22
13
Omission
09
04
2.9 Hz (E)
3.0 Hz (I)
4

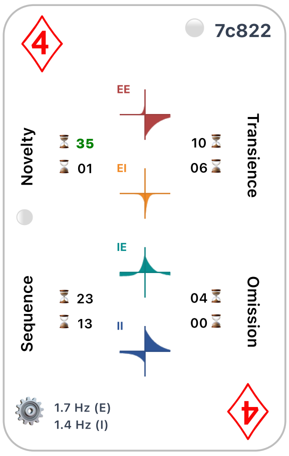
4
7c822
EE
EI
IE
II
Novelty
35
01
Transience
10
06
Sequence
23
13
Omission
04
00
1.7 Hz (E)
1.4 Hz (I)
4

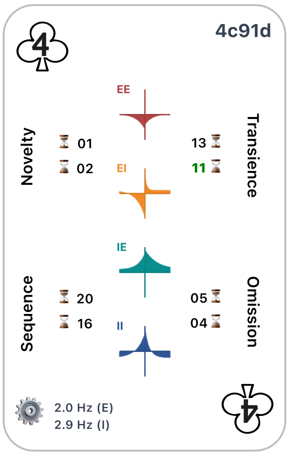
4
4c91d
EE
EI
IE
II
Novelty
01
02
Transience
13
11
Sequence
20
16
Omission
05
04
2.0 Hz (E)
2.9 Hz (I)
4

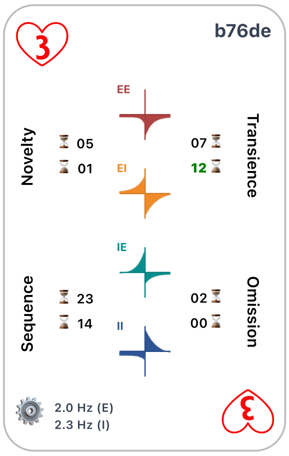
3
b76de
EE
EI
IE
II
Novelty
05
01
Transience
07
12
Sequence
23
14
Omission
02
00
2.0 Hz (E)
2.3 Hz (I)
3

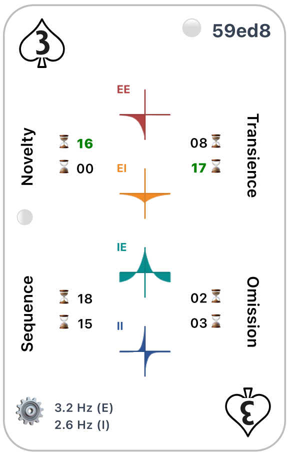
3
59ed8
EE
EI
IE
II
Novelty
16
00
Transience
08
17
Sequence
18
15
Omission
02
03
3.2 Hz (E)
2.6 Hz (I)
3

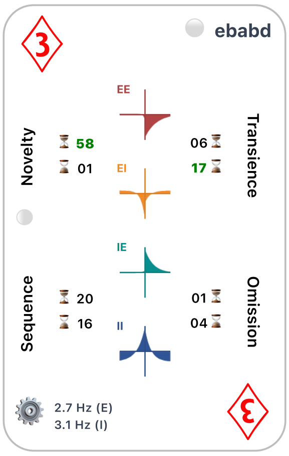
3
ebabd
EE
EI
IE
II
Novelty
58
01
Transience
06
17
Sequence
20
16
Omission
01
04
2.7 Hz (E)
3.1 Hz (I)
3

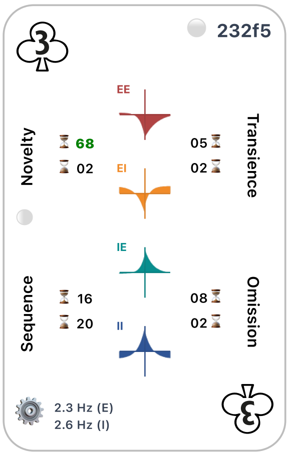
3
232f5
EE
EI
IE
II
Novelty
68
02
Transience
05
02
Sequence
16
20
Omission
08
02
2.3 Hz (E)
2.6 Hz (I)
3

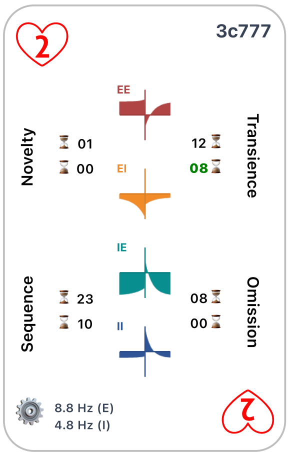
2
3c777
Novelty
01
00
EE
EI
12
08
Transience
IE
II
Sequence
23
10
08
00
Omission
8.8 Hz (E)
4.8 Hz (I)

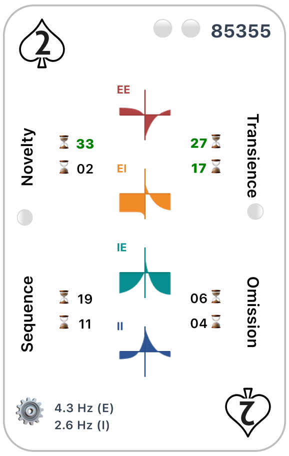
2
85355
Novelty
33
02
EE
EI
27
17
Transience
IE
II
Sequence
19
11
06
04
Omission
4.3 Hz (E)
2.6 Hz (I)

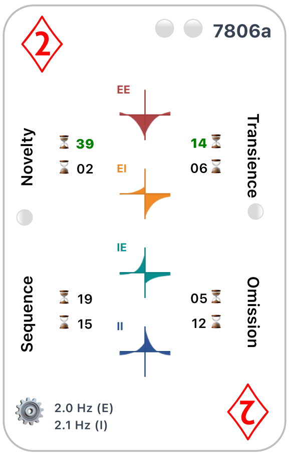
2
7806a
Novelty
39
02
EE
EI
14
06
Transience
IE
II
Sequence
19
15
05
12
Omission
2.0 Hz (E)
2.1 Hz (I)

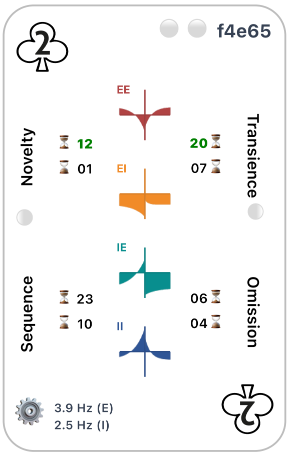
2
f4e65
Novelty
12
01
EE
EI
20
07
Transience
IE
II
Sequence
23
10
06
04
Omission
3.9 Hz (E)
2.5 Hz (I)

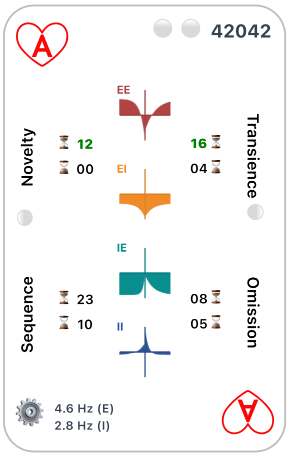
A
42042
Novelty
12
00
EE
EI
16
04
Transience
IE
II
Sequence
23
10
08
05
Omission
4.6 Hz (E)
2.8 Hz (I)

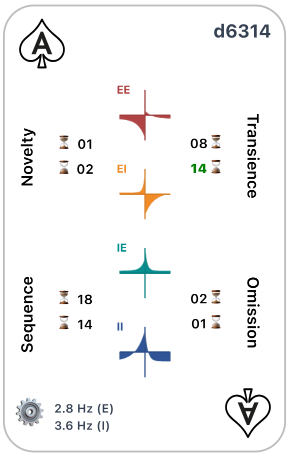
A
d6314
Novelty
01
02
EE
EI
08
14
Transience
IE
II
Sequence
18
14
02
01
Omission
2.8 Hz (E)
3.6 Hz (I)

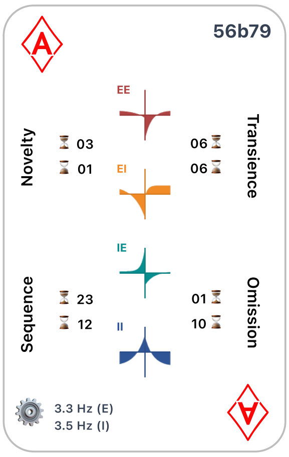
A
56b79
Novelty
03
01
EE
EI
06
06
Transience
IE
II
Sequence
23
12
01
10
Omission
3.3 Hz (E)
3.5 Hz (I)

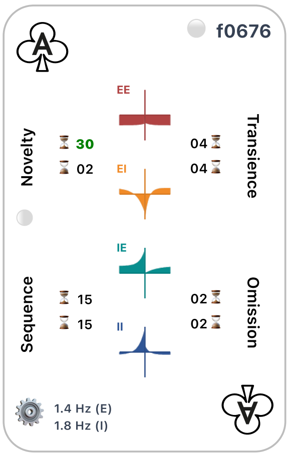
A
f0676
Novelty
30
02
EE
EI
04
04
Transience
IE
II
Sequence
15
15
02
02
Omission
1.4 Hz (E)
1.8 Hz (I)

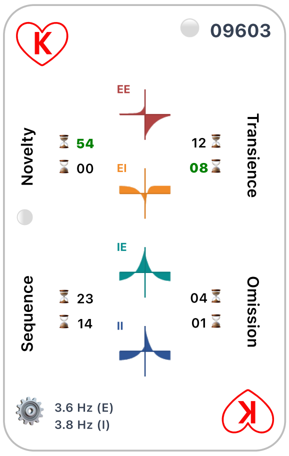
K
09603
Novelty
54
00
EE
EI
12
08
Transience
IE
II
Sequence
23
14
04
01
Omission
3.6 Hz (E)
3.8 Hz (I)

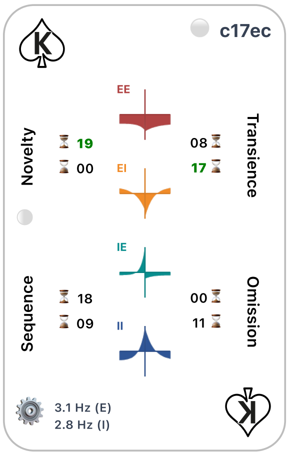
K
c17ec
Novelty
19
00
EE
EI
08
17
Transience
IE
II
Sequence
18
09
00
11
Omission
3.1 Hz (E)
2.8 Hz (I)

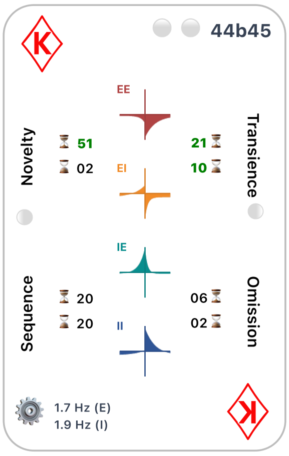
K
44b45
Novelty
51
02
EE
EI
21
10
Transience
IE
II
Sequence
20
20
06
02
Omission
1.7 Hz (E)
1.9 Hz (I)

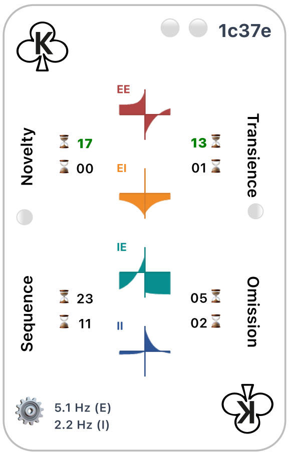
K
1c37e
Novelty
17
00
EE
EI
13
01
Transience
IE
II
Sequence
23
11
05
02
Omission
5.1 Hz (E)
2.2 Hz (I)

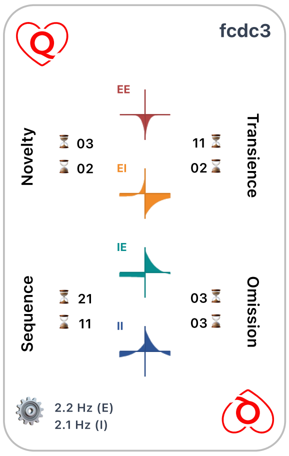
Q
fcdc3
Novelty
03
02
EE
EI
11
02
Transience
IE
II
Sequence
21
11
03
03
Omission
2.2 Hz (E)
2.1 Hz (I)

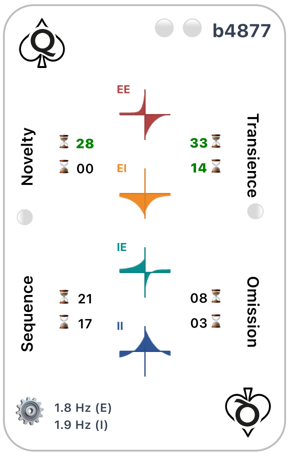
Q
b4877
Novelty
28
00
EE
EI
33
14
Transience
IE
II
Sequence
21
17
08
03
Omission
1.8 Hz (E)
1.9 Hz (I)

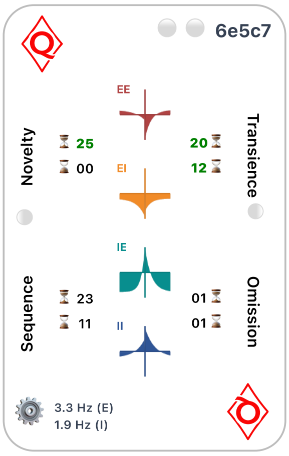
Q
6e5c7
Novelty
25
00
EE
EI
20
12
Transience
IE
II
Sequence
23
11
01
01
Omission
3.3 Hz (E)
1.9 Hz (I)

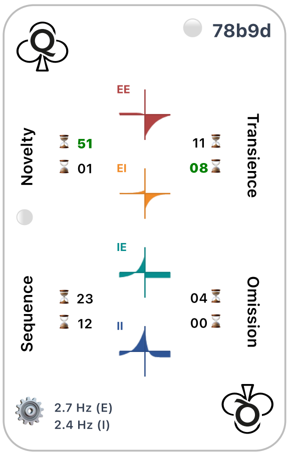
Q
78b9d
Novelty
51
01
EE
EI
11
08
Transience
IE
II
Sequence
23
12
04
00
Omission
2.7 Hz (E)
2.4 Hz (I)

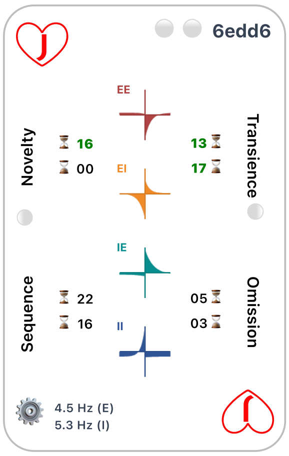
J
6edd6
EE
EI
IE
II
Novelty 16 00
Transience 13 17
Sequence 22 16
Omission 05 03
4.5 Hz (E)
5.3 Hz (I)

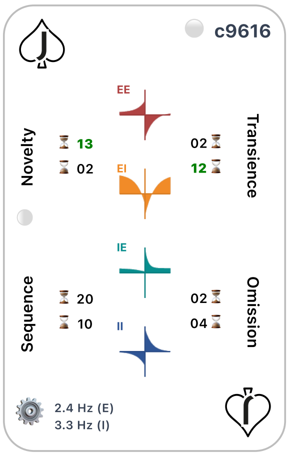
J
c9616
EE
EI
IE
II
Novelty 13 02
Transience 02 12
Sequence 20 10
Omission 02 04
2.4 Hz (E)
3.3 Hz (I)

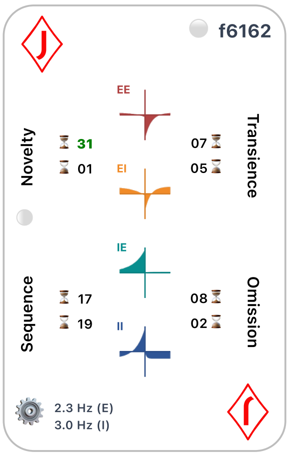
J
f6162
EE
EI
IE
II
Novelty 31 01
Transience 07 05
Sequence 17 19
Omission 08 02
2.3 Hz (E)
3.0 Hz (I)

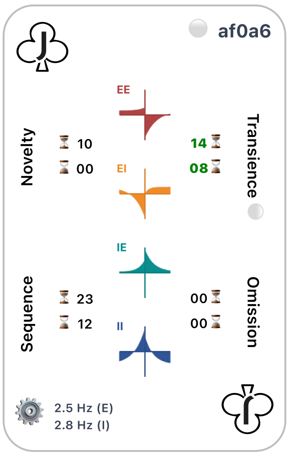
J
af0a6
EE
EI
IE
II
Novelty 10 00
Transience 14 08
Sequence 23 12
Omission 00 00
2.5 Hz (E)
2.8 Hz (I)

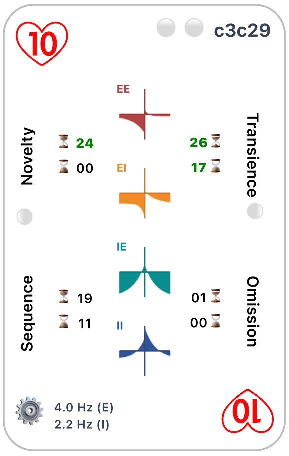
10
c3c29
EE
EI
IE
II
Novelty 24 00
Transience 26 17
Sequence 19 11
Omission 01 00
4.0 Hz (E)
2.2 Hz (I)

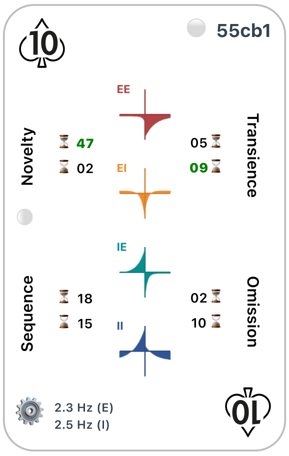
10
55cb1
EE
EI
IE
II
Novelty 47 02
Transience 05 09
Sequence 18 15
Omission 02 10
2.3 Hz (E)
2.5 Hz (I)

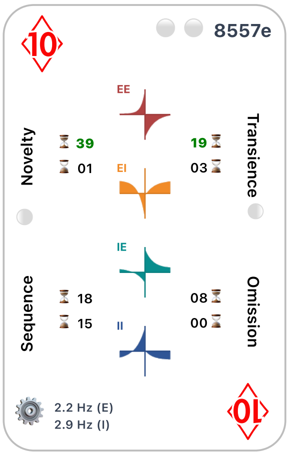
10
8557e
EE
EI
IE
II
Novelty 39 01
Transience 19 03
Sequence 18 15
Omission 08 00
2.2 Hz (E)
2.9 Hz (I)

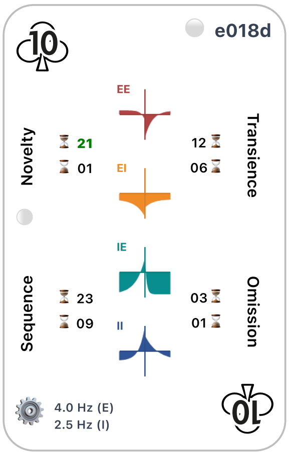
10
e018d
EE
EI
IE
II
Novelty 21 01
Transience 12 06
Sequence 23 09
Omission 03 01
4.0 Hz (E)
2.5 Hz (I)

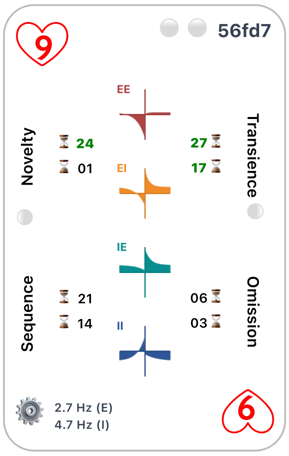
9
56fd7
EE
EI
IE
II
Novelty 24 01
Transience 27 17
Sequence 21 14
Omission 06 03
2.7 Hz (E)
4.7 Hz (I)

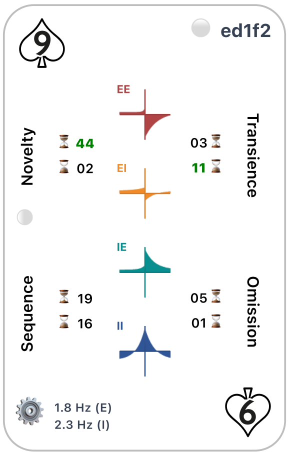
9
ed1f2
EE
EI
IE
II
Novelty 44 02
Transience 03 11
Sequence 19 16
Omission 05 01
1.8 Hz (E)
2.3 Hz (I)

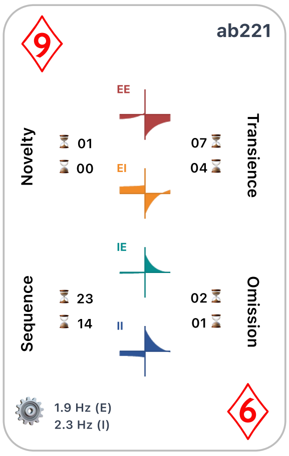
9
ab221
EE
EI
IE
II
Novelty 01 00
Transience 07 04
Sequence 23 14
Omission 02 01
1.9 Hz (E)
2.3 Hz (I)

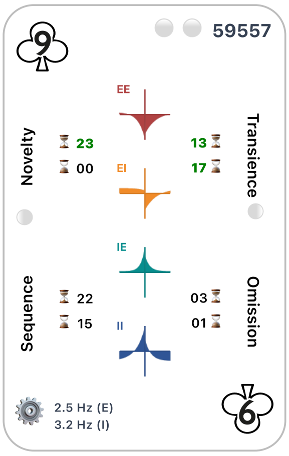
9
59557
EE
EI
IE
II
Novelty 23 00
Transience 13 17
Sequence 22 15
Omission 03 01
2.5 Hz (E)
3.2 Hz (I)

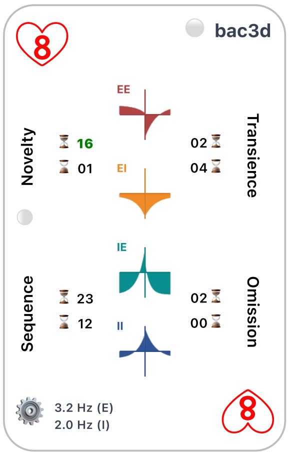
8
bac3d
EE
EI
IE
II
Novelty 16 01
Transience 02 04
Sequence 23 12
Omission 02 00
3.2 Hz (E)
2.0 Hz (I)

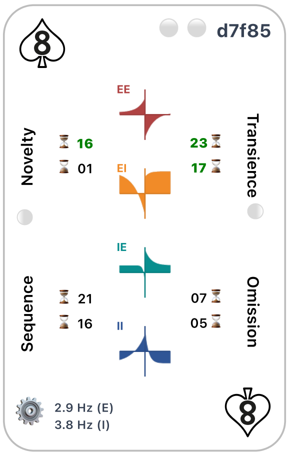
8
d7f85
EE
EI
IE
II
Novelty 16 01
Transience 23 17
Sequence 21 16
Omission 07 05
2.9 Hz (E)
3.8 Hz (I)

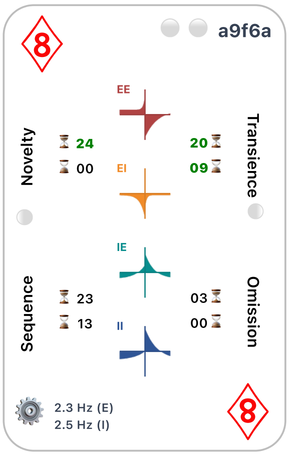
8
a9f6a
EE
EI
IE
II
Novelty 24 00
Transience 20 09
Sequence 23 13
Omission 03 00
2.3 Hz (E)
2.5 Hz (I)

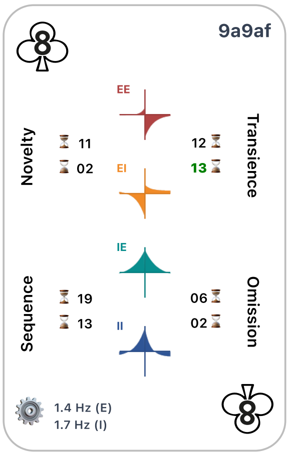
8
9a9af
EE
EI
IE
II
Novelty 11 02
Transience 12 13
Sequence 19 13
Omission 06 02
1.4 Hz (E)
1.7 Hz (I)

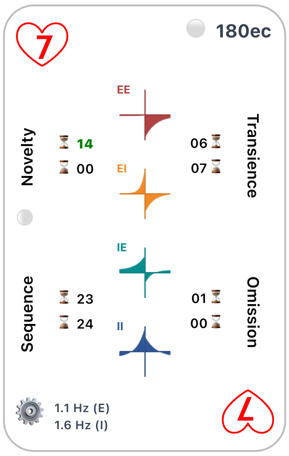
7
180ec
EE
Novelty
14
00
06
07
Transience
EI
IE
Sequence
23
24
01
00
Omission
II
1.1 Hz (E)
1.6 Hz (I)
7

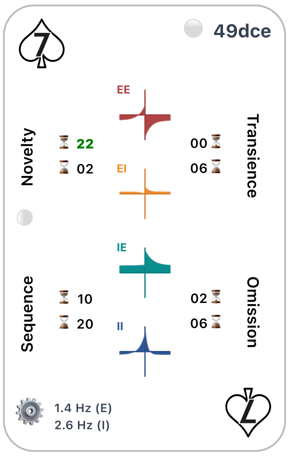
7
49dce
EE
Novelty
22
02
00
06
Transience
EI
IE
Sequence
10
20
02
06
Omission
II
1.4 Hz (E)
2.6 Hz (I)
7

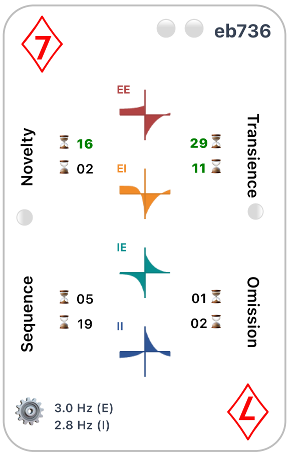
7
eb736
EE
Novelty
16
02
29
11
Transience
EI
IE
Sequence
05
19
01
02
Omission
II
3.0 Hz (E)
2.8 Hz (I)
7

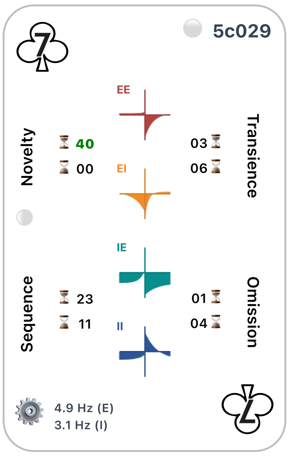
7
5c029
EE
Novelty
40
00
03
06
Transience
EI
IE
Sequence
23
11
01
04
Omission
II
4.9 Hz (E)
3.1 Hz (I)
7

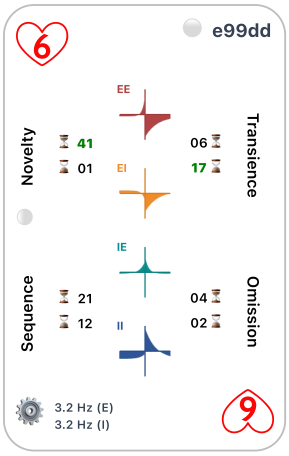
6
e99dd
EE
Novelty
41
01
06
17
Transience
EI
IE
Sequence
21
12
04
02
Omission
II
3.2 Hz (E)
3.2 Hz (I)
9

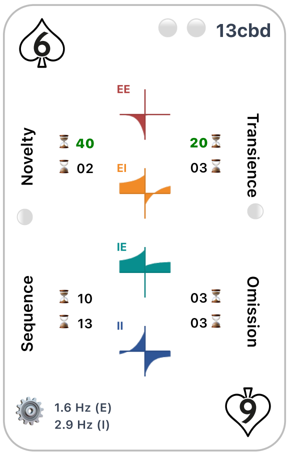
6
13cbd
EE
Novelty
40
02
20
03
Transience
EI
IE
Sequence
10
13
03
03
Omission
II
1.6 Hz (E)
2.9 Hz (I)
9

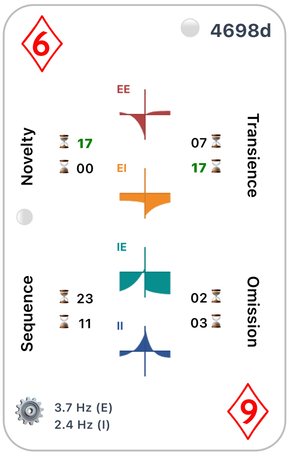
6
4698d
EE
Novelty
17
00
07
17
Transience
EI
IE
Sequence
23
11
02
03
Omission
II
3.7 Hz (E)
2.4 Hz (I)
9

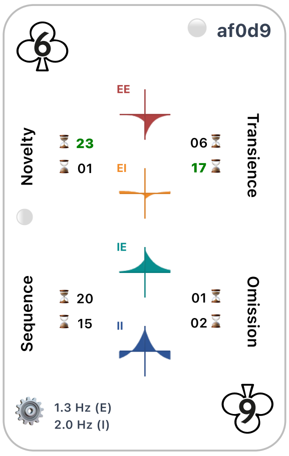
6
af0d9
EE
Novelty
23
01
06
17
Transience
EI
IE
Sequence
20
15
01
02
Omission
II
1.3 Hz (E)
2.0 Hz (I)
9

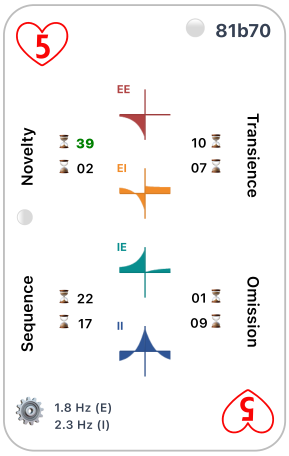
5
81b70
EE
Novelty
39
02
10
07
Transience
EI
IE
Sequence
22
17
01
09
Omission
II
1.8 Hz (E)
2.3 Hz (I)
5

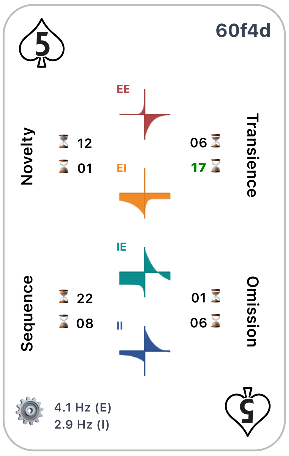
5
60f4d
EE
Novelty
12
01
06
17
Transience
EI
IE
Sequence
22
08
01
06
Omission
II
4.1 Hz (E)
2.9 Hz (I)
5

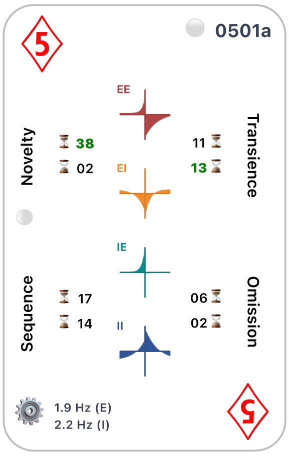
5
0501a
EE
Novelty
38
02
11
13
Transience
EI
IE
Sequence
17
14
06
02
Omission
II
1.9 Hz (E)
2.2 Hz (I)
5

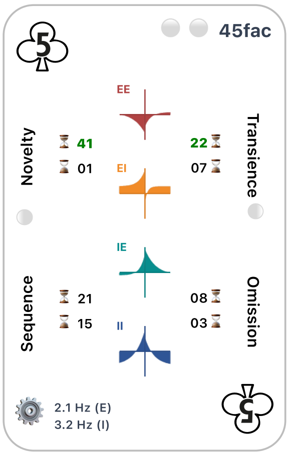
5
45fac
EE
Novelty
41
01
22
07
Transience
EI
IE
Sequence
21
15
08
03
Omission
II
2.1 Hz (E)
3.2 Hz (I)
5

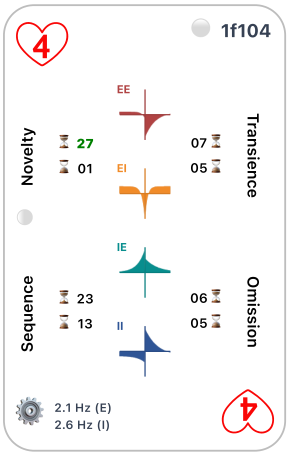
4
1f104
EE
Novelty
27
01
07
05
Transience
EI
IE
Sequence
23
13
06
05
Omission
II
2.1 Hz (E)
2.6 Hz (I)
4

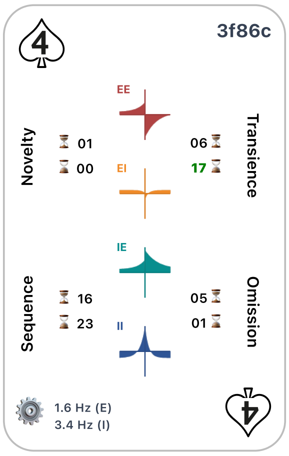
4
3f86c
EE
Novelty
01
00
06
17
Transience
EI
IE
Sequence
16
23
05
01
Omission
II
1.6 Hz (E)
3.4 Hz (I)
4

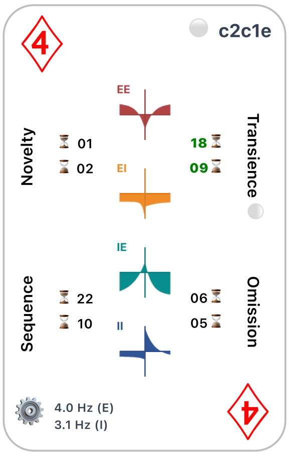
4
c2c1e
EE
Novelty
01
02
18
09
Transience
EI
IE
Sequence
22
10
06
05
Omission
II
4.0 Hz (E)
3.1 Hz (I)
4

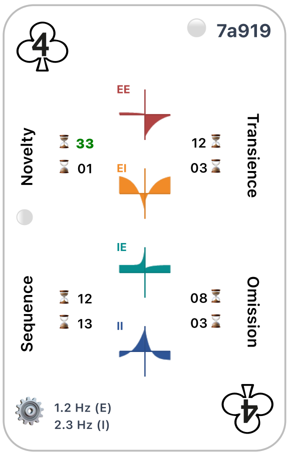
4
7a919
EE
Novelty
33
01
12
03
Transience
EI
IE
Sequence
12
13
08
03
Omission
II
1.2 Hz (E)
2.3 Hz (I)
4

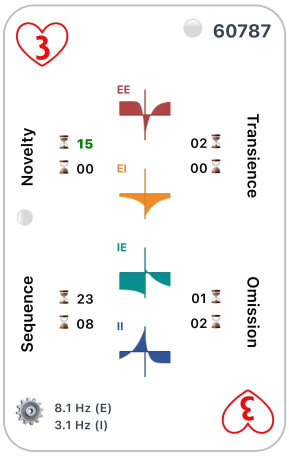
3
60787
EE
EI
IE
II
Novelty 15 00
Transience 02 00
Sequence 23 08
Omission 01 02
8.1 Hz (E)
3.1 Hz (I)

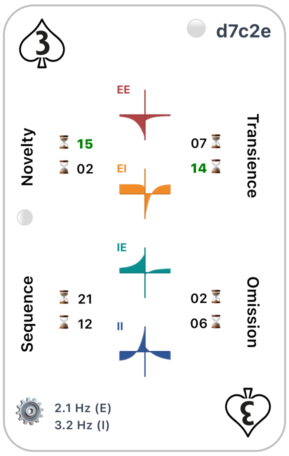
3
d7c2e
EE
EI
IE
II
Novelty 15 02
Transience 07 14
Sequence 21 12
Omission 02 06
2.1 Hz (E)
3.2 Hz (I)

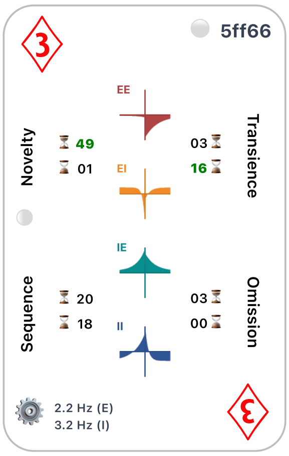
3
5ff66
EE
EI
IE
II
Novelty 49 01
Transience 03 16
Sequence 20 18
Omission 03 00
2.2 Hz (E)
3.2 Hz (I)

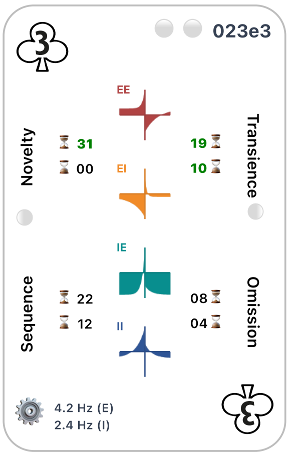
3
023e3
EE
EI
IE
II
Novelty 31 00
Transience 19 10
Sequence 22 12
Omission 08 04
4.2 Hz (E)
2.4 Hz (I)

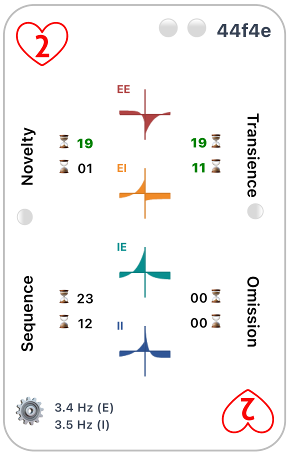
2
44f4e
EE
EI
IE
II
Novelty 19 01
Transience 19 11
Sequence 23 12
Omission 00 00
3.4 Hz (E)
3.5 Hz (I)

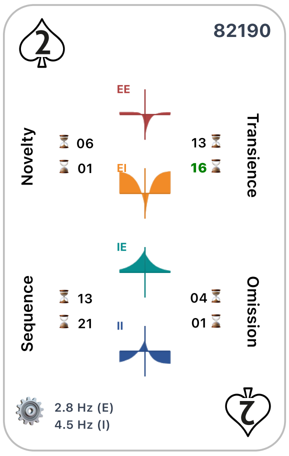
2
82190
EE
EI
IE
II
Novelty 06 01
Transience 13 16
Sequence 13 21
Omission 04 01
2.8 Hz (E)
4.5 Hz (I)

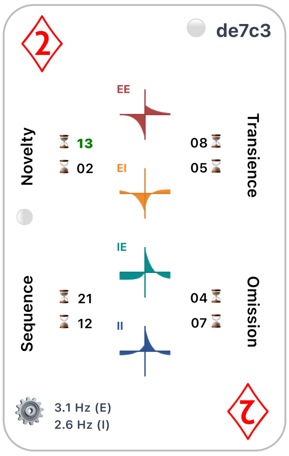
2
de7c3
EE
EI
IE
II
Novelty 13 02
Transience 08 05
Sequence 21 12
Omission 04 07
3.1 Hz (E)
2.6 Hz (I)

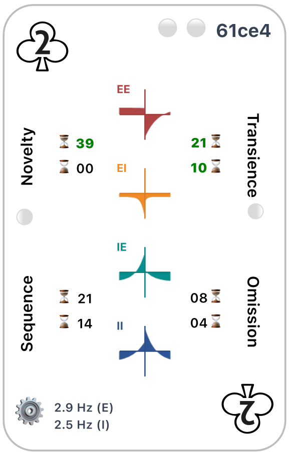
2
61ce4
EE
EI
IE
II
Novelty 39 00
Transience 21 10
Sequence 21 14
Omission 08 04
2.9 Hz (E)
2.5 Hz (I)

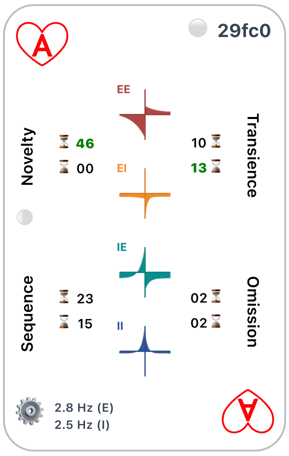
A
29fc0
EE
EI
IE
II
Novelty 46 00
Transience 10 13
Sequence 23 15
Omission 02 02
2.8 Hz (E)
2.5 Hz (I)

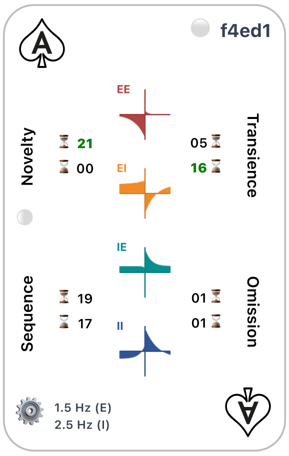
A
f4ed1
EE
EI
IE
II
Novelty 21 00
Transience 05 16
Sequence 19 17
Omission 01 01
1.5 Hz (E)
2.5 Hz (I)

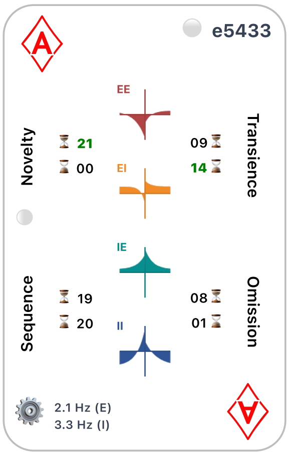
A
e5433
EE
EI
IE
II
Novelty 21 00
Transience 09 14
Sequence 19 20
Omission 08 01
2.1 Hz (E)
3.3 Hz (I)

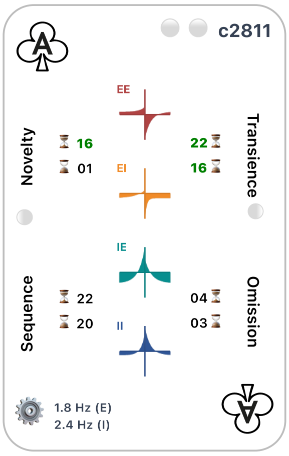
A
c2811
EE
EI
IE
II
Novelty 16 01
Transience 22 16
Sequence 22 20
Omission 04 03
1.8 Hz (E)
2.4 Hz (I)

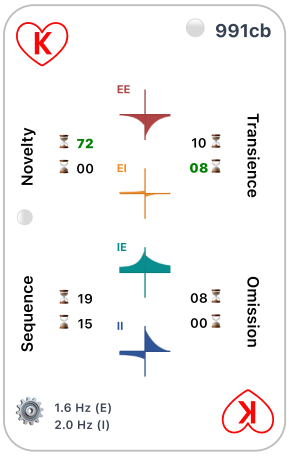
K
991cb
EE
EI
IE
II
Novelty 72 00
Transience 10 08
Sequence 19 15
Omission 08 00
1.6 Hz (E)
2.0 Hz (I)

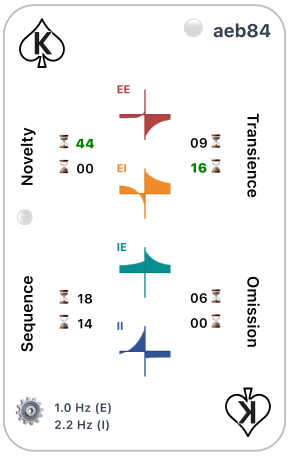
K
aeb84
EE
EI
IE
II
Novelty 44 00
Transience 09 16
Sequence 18 14
Omission 06 00
1.0 Hz (E)
2.2 Hz (I)

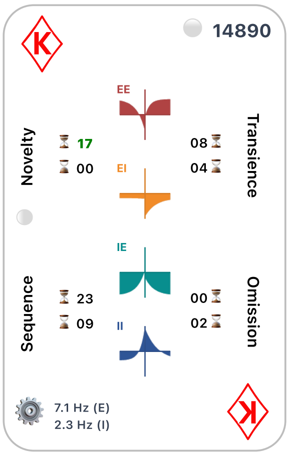
K
14890
EE
EI
IE
II
Novelty 17 00
Transience 08 04
Sequence 23 09
Omission 00 02
7.1 Hz (E)
2.3 Hz (I)

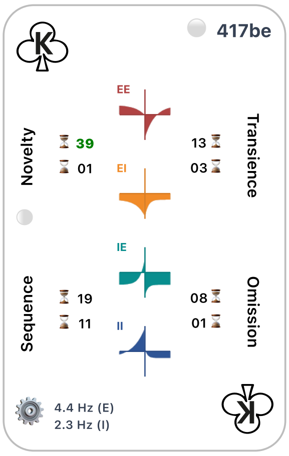
K
417be
EE
EI
IE
II
Novelty 39 01
Transience 13 03
Sequence 19 11
Omission 08 01
4.4 Hz (E)
2.3 Hz (I)

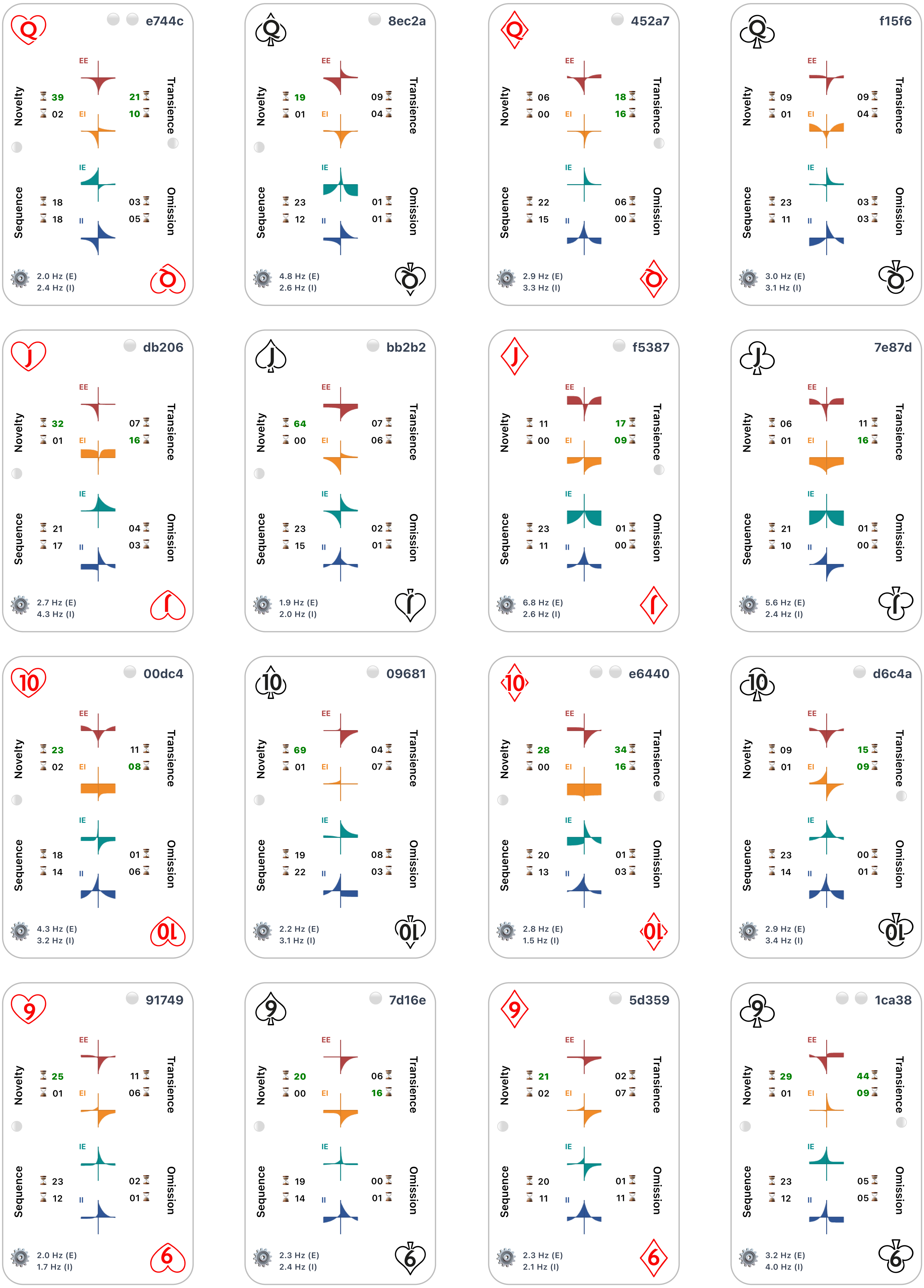
Q e744c
EE EI IE II
Novelty 39 02 — Transience 21 10
Sequence 18 18 — Omission 03 05
2.0 Hz (E) 2.4 Hz (I)
Q 8ec2a
Novelty 19 01 — Transience 09 04
Sequence 23 12 — Omission 01 01
4.8 Hz (E) 2.6 Hz (I)
Q 452a7
Novelty 06 00 — Transience 18 16
Sequence 22 15 — Omission 06 00
2.9 Hz (E) 3.3 Hz (I)
Q f15f6
Novelty 09 01 — Transience 09 04
Sequence 23 11 — Omission 03 03
3.0 Hz (E) 3.1 Hz (I)
J db206
Novelty 32 01 — Transience 07 16
Sequence 21 17 — Omission 04 03
2.7 Hz (E) 4.3 Hz (I)
J bb2b2
Novelty 64 00 — Transience 07 06
Sequence 23 15 — Omission 02 01
1.9 Hz (E) 2.0 Hz (I)
J f5387
Novelty 11 00 — Transience 17 09
Sequence 23 11 — Omission 01 00
6.8 Hz (E) 2.6 Hz (I)
J 7e87d
Novelty 06 01 — Transience 11 16
Sequence 21 10 — Omission 01 00
5.6 Hz (E) 2.4 Hz (I)
10 00dc4
Novelty 23 02 — Transience 11 08
Sequence 18 14 — Omission 01 06
4.3 Hz (E) 3.2 Hz (I)
10 09681
Novelty 69 01 — Transience 04 07
Sequence 19 22 — Omission 08 03
2.2 Hz (E) 3.1 Hz (I)
10 e6440
Novelty 28 00 — Transience 34 16
Sequence 20 13 — Omission 01 03
2.8 Hz (E) 1.5 Hz (I)
10 d6c4a
Novelty 09 01 — Transience 15 09
Sequence 23 14 — Omission 00 01
2.9 Hz (E) 3.4 Hz (I)
9 91749
Novelty 25 01 — Transience 11 06
Sequence 23 12 — Omission 02 01
2.0 Hz (E) 1.7 Hz (I)
9 7d16e
Novelty 20 00 — Transience 06 16
Sequence 19 14 — Omission 00 01
2.3 Hz (E) 2.4 Hz (I)
9 5d359
Novelty 21 02 — Transience 02 07
Sequence 20 11 — Omission 01 11
2.3 Hz (E) 2.1 Hz (I)
9 1ca38
Novelty 29 01 — Transience 44 09
Sequence 23 12 — Omission 05 05
3.2 Hz (E) 4.0 Hz (I)

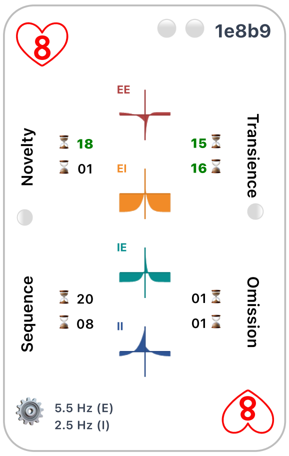
8
1e8b9
EE
Novelty
18
01
15
16
Transience
EI
IE
Sequence
20
08
01
01
Omission
II
5.5 Hz (E)
2.5 Hz (I)

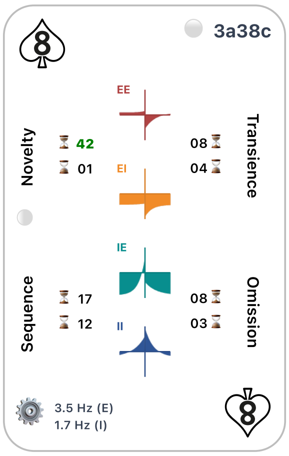
8
3a38c
EE
Novelty
42
01
08
04
Transience
EI
IE
Sequence
17
12
08
03
Omission
II
3.5 Hz (E)
1.7 Hz (I)

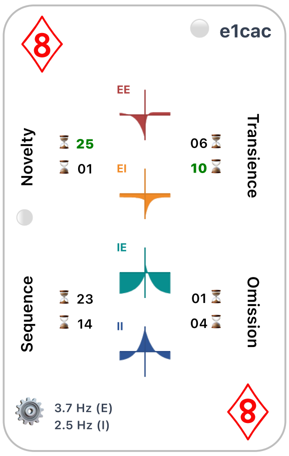
8
e1cac
EE
Novelty
25
01
06
10
Transience
EI
IE
Sequence
23
14
01
04
Omission
II
3.7 Hz (E)
2.5 Hz (I)

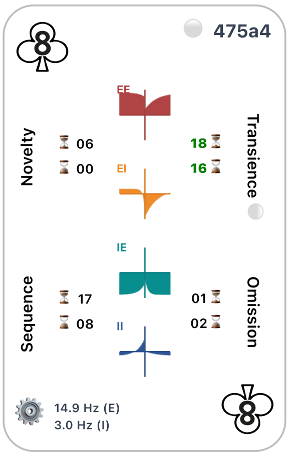
8
475a4
EE
Novelty
06
00
18
16
Transience
EI
IE
Sequence
17
08
01
02
Omission
II
14.9 Hz (E)
3.0 Hz (I)

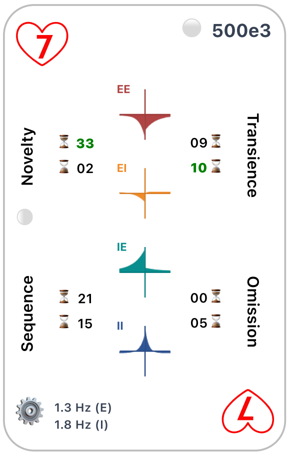
7
500e3
EE
Novelty
33
02
09
10
Transience
EI
IE
Sequence
21
15
00
05
Omission
II
1.3 Hz (E)
1.8 Hz (I)

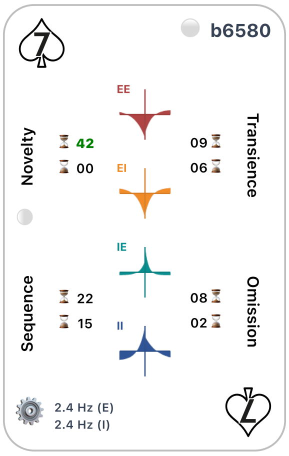
7
b6580
EE
Novelty
42
00
09
06
Transience
EI
IE
Sequence
22
15
08
02
Omission
II
2.4 Hz (E)
2.4 Hz (I)

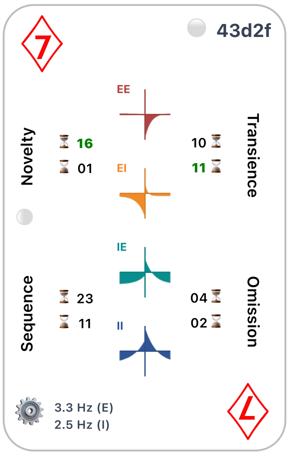
7
43d2f
EE
Novelty
16
01
10
11
Transience
EI
IE
Sequence
23
11
04
02
Omission
II
3.3 Hz (E)
2.5 Hz (I)

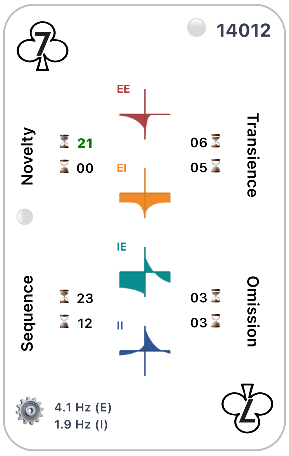
7
14012
EE
Novelty
21
00
06
05
Transience
EI
IE
Sequence
23
12
03
03
Omission
II
4.1 Hz (E)
1.9 Hz (I)

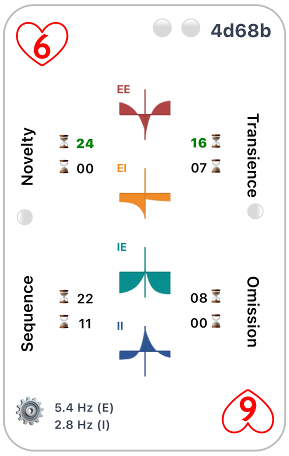
6
4d68b
EE
Novelty
24
00
16
07
Transience
EI
IE
Sequence
22
11
08
00
Omission
II
5.4 Hz (E)
2.8 Hz (I)

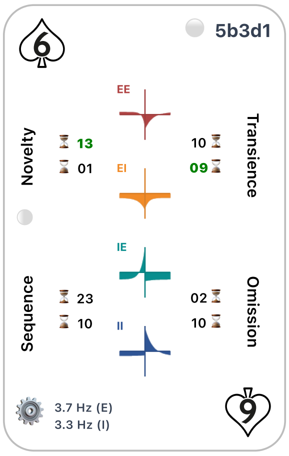
6
5b3d1
EE
Novelty
13
01
10
09
Transience
EI
IE
Sequence
23
10
02
10
Omission
II
3.7 Hz (E)
3.3 Hz (I)

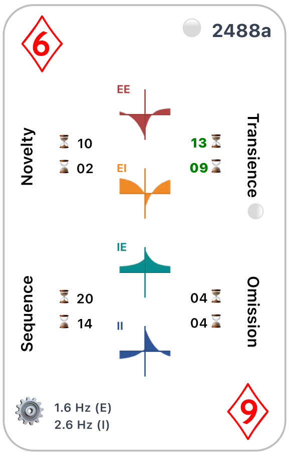
6
2488a
EE
Novelty
10
02
13
09
Transience
EI
IE
Sequence
20
14
04
04
Omission
II
1.6 Hz (E)
2.6 Hz (I)

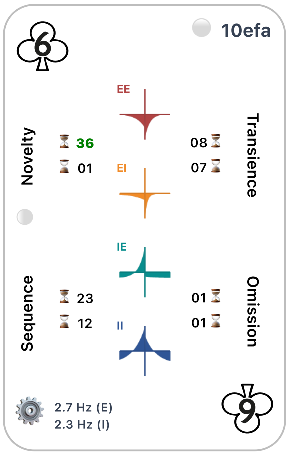
6
10efa
EE
Novelty
36
01
08
07
Transience
EI
IE
Sequence
23
12
01
01
Omission
II
2.7 Hz (E)
2.3 Hz (I)

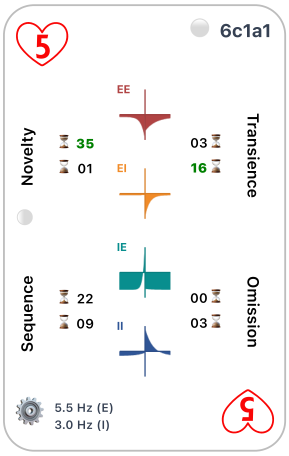
5
6c1a1
EE
Novelty
35
01
03
16
Transience
EI
IE
Sequence
22
09
00
03
Omission
II
5.5 Hz (E)
3.0 Hz (I)

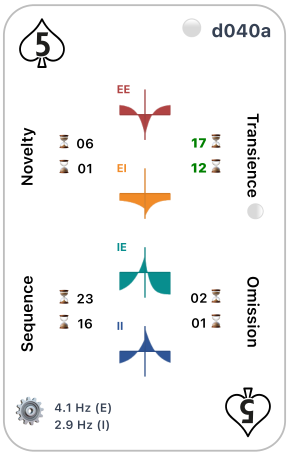
5
d040a
EE
Novelty
06
01
17
12
Transience
EI
IE
Sequence
23
16
02
01
Omission
II
4.1 Hz (E)
2.9 Hz (I)

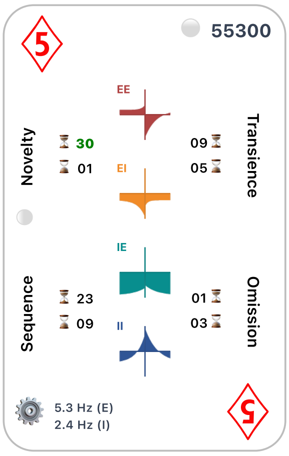
5
55300
EE
Novelty
30
01
09
05
Transience
EI
IE
Sequence
23
09
01
03
Omission
II
5.3 Hz (E)
2.4 Hz (I)

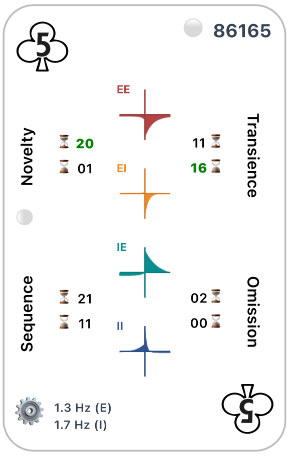
5
86165
EE
Novelty
20
01
11
16
Transience
EI
IE
Sequence
21
11
02
00
Omission
II
1.3 Hz (E)
1.7 Hz (I)

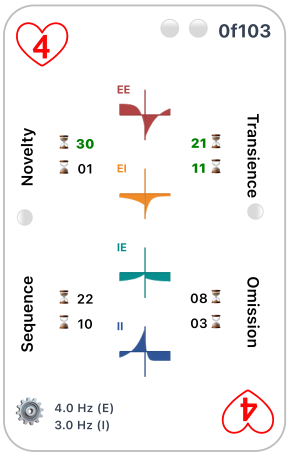
4
0f103
EE
EI
IE
II
Novelty
Transience
Sequence
Omission
30
01
21
11
22
10
08
03
4.0 Hz (E)
3.0 Hz (I)

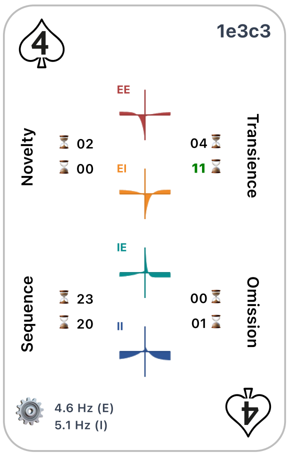
4
1e3c3
EE
EI
IE
II
Novelty
Transience
Sequence
Omission
02
00
04
11
23
20
00
01
4.6 Hz (E)
5.1 Hz (I)

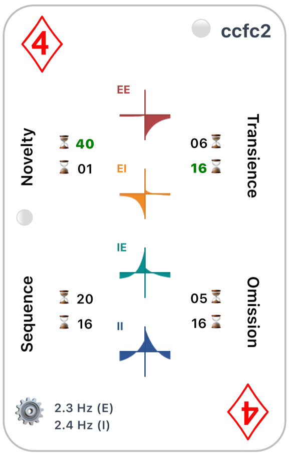
4
ccfc2
EE
EI
IE
II
Novelty
Transience
Sequence
Omission
40
01
06
16
20
16
05
16
2.3 Hz (E)
2.4 Hz (I)

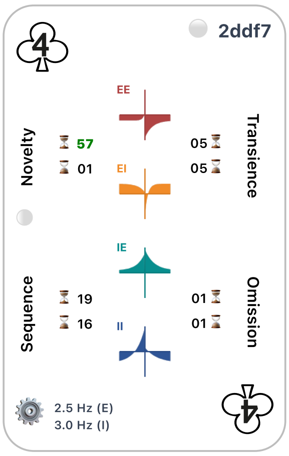
4
2ddf7
EE
EI
IE
II
Novelty
Transience
Sequence
Omission
57
01
05
05
19
16
01
01
2.5 Hz (E)
3.0 Hz (I)

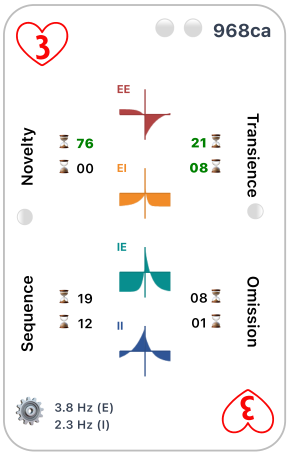
3
968ca
EE
EI
IE
II
Novelty
Transience
Sequence
Omission
76
00
21
08
19
12
08
01
3.8 Hz (E)
2.3 Hz (I)

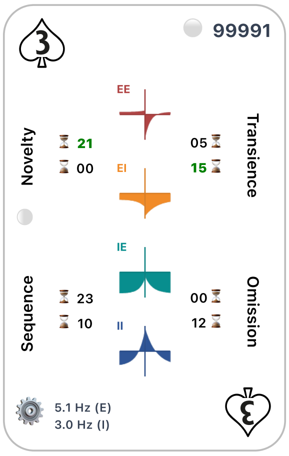
3
99991
EE
EI
IE
II
Novelty
Transience
Sequence
Omission
21
00
05
15
23
10
00
12
5.1 Hz (E)
3.0 Hz (I)

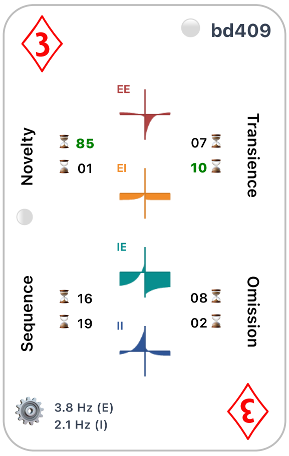
3
bd409
EE
EI
IE
II
Novelty
Transience
Sequence
Omission
85
01
07
10
16
19
08
02
3.8 Hz (E)
2.1 Hz (I)

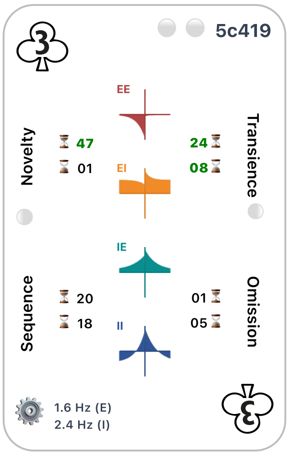
3
5c419
EE
EI
IE
II
Novelty
Transience
Sequence
Omission
47
01
24
08
20
18
01
05
1.6 Hz (E)
2.4 Hz (I)

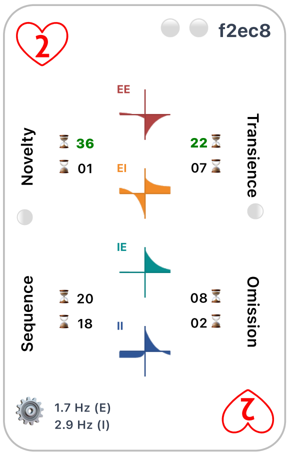
2
f2ec8
EE
EI
IE
II
Novelty
Transience
Sequence
Omission
36
01
22
07
20
18
08
02
1.7 Hz (E)
2.9 Hz (I)

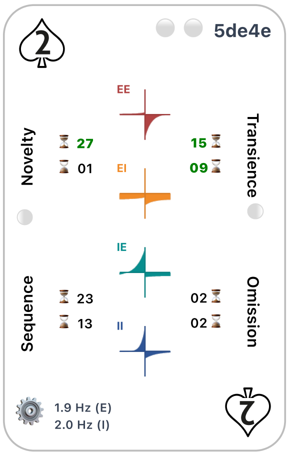
2
5de4e
EE
EI
IE
II
Novelty
Transience
Sequence
Omission
27
01
15
09
23
13
02
02
1.9 Hz (E)
2.0 Hz (I)

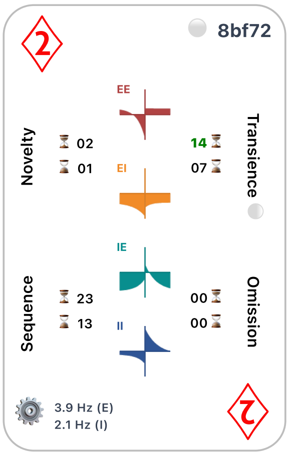
2
8bf72
EE
EI
IE
II
Novelty
Transience
Sequence
Omission
02
01
14
07
23
13
00
00
3.9 Hz (E)
2.1 Hz (I)

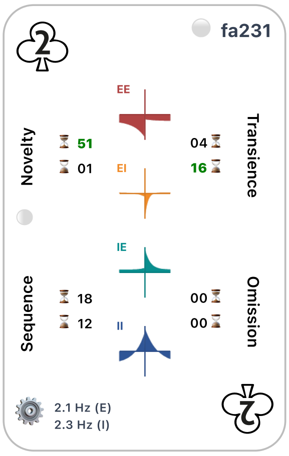
2
fa231
EE
EI
IE
II
Novelty
Transience
Sequence
Omission
51
01
04
16
18
12
00
00
2.1 Hz (E)
2.3 Hz (I)

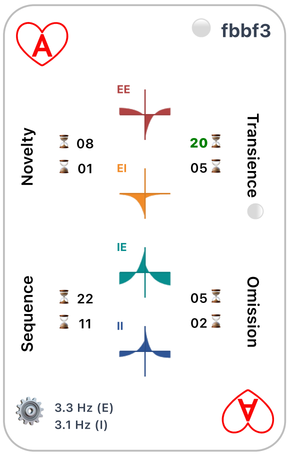
A
fbbf3
EE
EI
IE
II
Novelty
Transience
Sequence
Omission
08
01
20
05
22
11
05
02
3.3 Hz (E)
3.1 Hz (I)

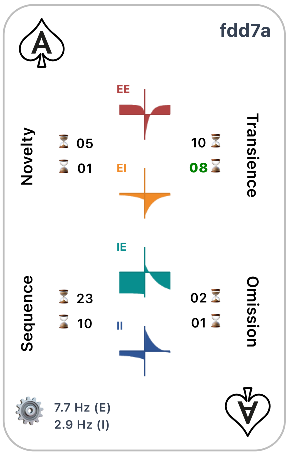
A
fdd7a
EE
EI
IE
II
Novelty
Transience
Sequence
Omission
05
01
10
08
23
10
02
01
7.7 Hz (E)
2.9 Hz (I)

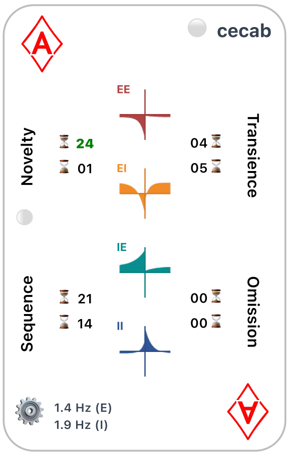
A
cecab
EE
EI
IE
II
Novelty
Transience
Sequence
Omission
24
01
04
05
21
14
00
00
1.4 Hz (E)
1.9 Hz (I)

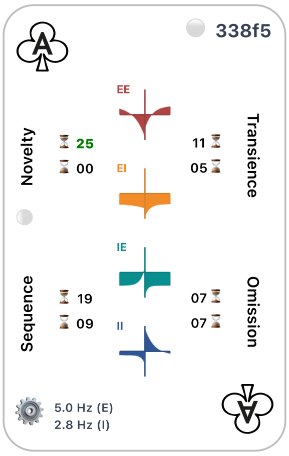
A
338f5
EE
EI
IE
II
Novelty
Transience
Sequence
Omission
25
00
11
05
19
09
07
07
5.0 Hz (E)
2.8 Hz (I)

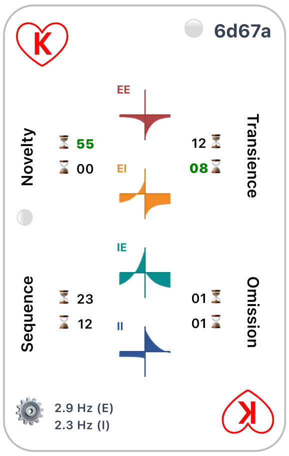
K
6d67a
EE
EI
IE
II
Novelty
55
00
Transience
12
08
Sequence
23
12
Omission
01
01
2.9 Hz (E)
2.3 Hz (I)

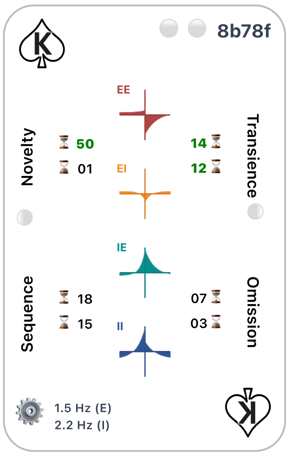
K
8b78f
EE
EI
IE
II
Novelty
50
01
Transience
14
12
Sequence
18
15
Omission
07
03
1.5 Hz (E)
2.2 Hz (I)

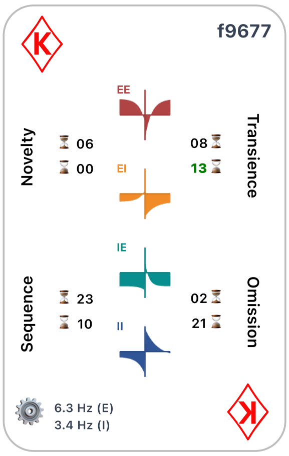
K
f9677
EE
EI
IE
II
Novelty
06
00
Transience
08
13
Sequence
23
10
Omission
02
21
6.3 Hz (E)
3.4 Hz (I)

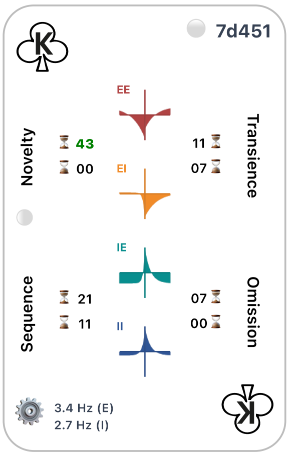
K
7d451
EE
EI
IE
II
Novelty
43
00
Transience
11
07
Sequence
21
11
Omission
07
00
3.4 Hz (E)
2.7 Hz (I)

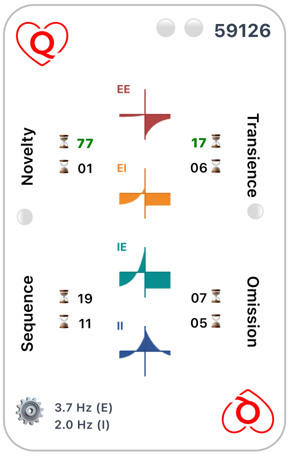
Q
59126
EE
EI
IE
II
Novelty
77
01
Transience
17
06
Sequence
19
11
Omission
07
05
3.7 Hz (E)
2.0 Hz (I)

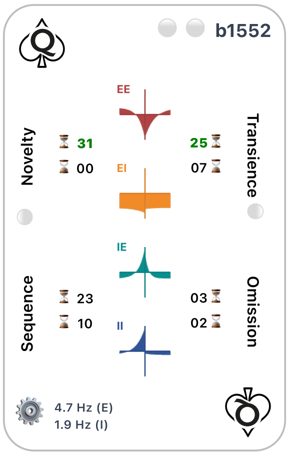
Q
b1552
EE
EI
IE
II
Novelty
31
00
Transience
25
07
Sequence
23
10
Omission
03
02
4.7 Hz (E)
1.9 Hz (I)

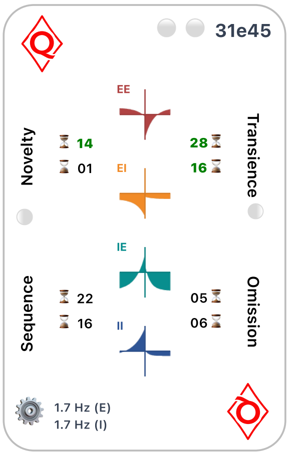
Q
31e45
EE
EI
IE
II
Novelty
14
01
Transience
28
16
Sequence
22
16
Omission
05
06
1.7 Hz (E)
1.7 Hz (I)

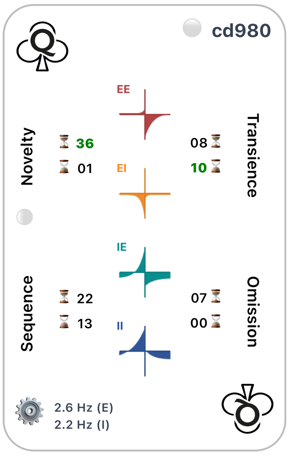
Q
cd980
EE
EI
IE
II
Novelty
36
01
Transience
08
10
Sequence
22
13
Omission
07
00
2.6 Hz (E)
2.2 Hz (I)

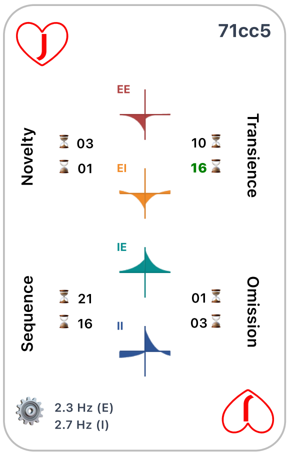
J
71cc5
EE
EI
IE
II
Novelty
03
01
Transience
10
16
Sequence
21
16
Omission
01
03
2.3 Hz (E)
2.7 Hz (I)

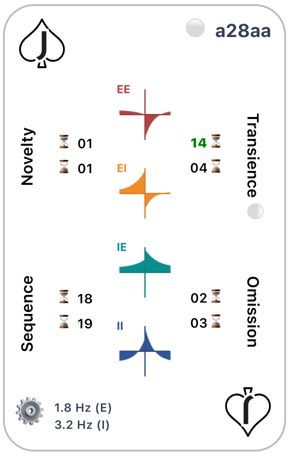
J
a28aa
EE
EI
IE
II
Novelty
01
01
Transience
14
04
Sequence
18
19
Omission
02
03
1.8 Hz (E)
3.2 Hz (I)

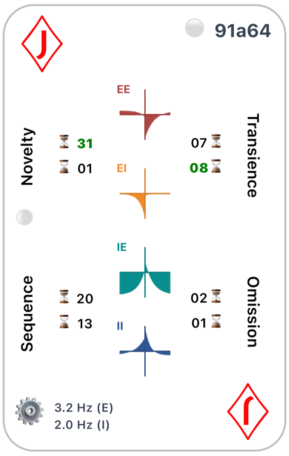
J
91a64
EE
EI
IE
II
Novelty
31
01
Transience
07
08
Sequence
20
13
Omission
02
01
3.2 Hz (E)
2.0 Hz (I)

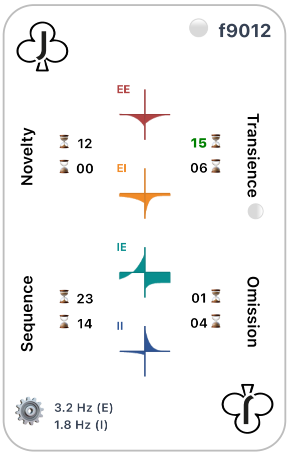
J
f9012
EE
EI
IE
II
Novelty
12
00
Transience
15
06
Sequence
23
14
Omission
01
04
3.2 Hz (E)
1.8 Hz (I)

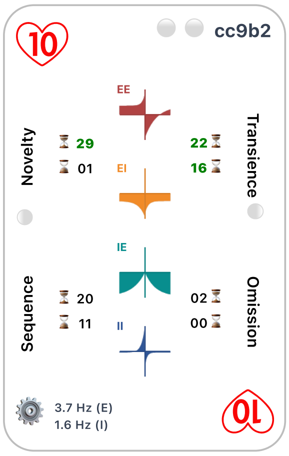
10
cc9b2
EE
EI
IE
II
Novelty
29
01
Transience
22
16
Sequence
20
11
Omission
02
00
3.7 Hz (E)
1.6 Hz (I)

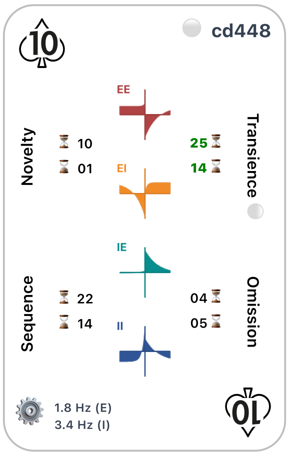
10
cd448
EE
EI
IE
II
Novelty
10
01
Transience
25
14
Sequence
22
14
Omission
04
05
1.8 Hz (E)
3.4 Hz (I)

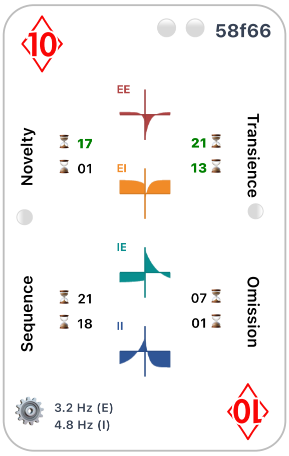
10
58f66
EE
EI
IE
II
Novelty
17
01
Transience
21
13
Sequence
21
18
Omission
07
01
3.2 Hz (E)
4.8 Hz (I)

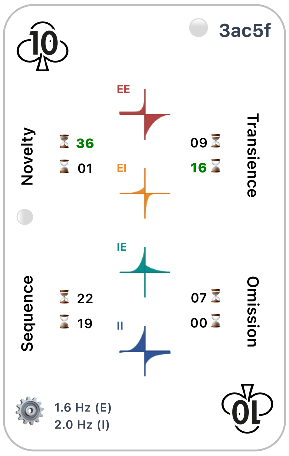
10
3ac5f
EE
EI
IE
II
Novelty
36
01
Transience
09
16
Sequence
22
19
Omission
07
00
1.6 Hz (E)
2.0 Hz (I)

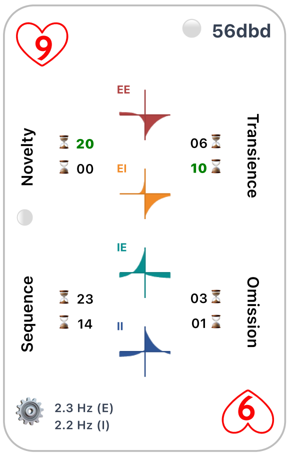
56dbd
EE
EI
IE
II
Novelty
20
00
Transience
06
10
Sequence
23
14
Omission
03
01
2.3 Hz (E)
2.2 Hz (I)

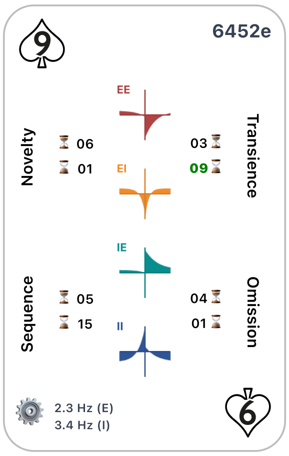
6452e
EE
EI
IE
II
Novelty
06
01
Transience
03
09
Sequence
05
15
Omission
04
01
2.3 Hz (E)
3.4 Hz (I)

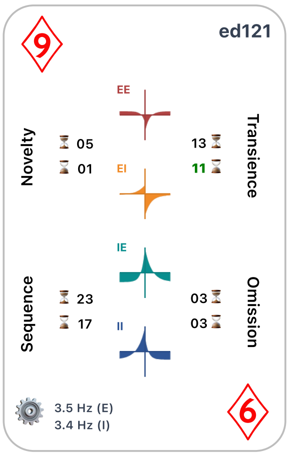
ed121
EE
EI
IE
II
Novelty
05
01
Transience
13
11
Sequence
23
17
Omission
03
03
3.5 Hz (E)
3.4 Hz (I)

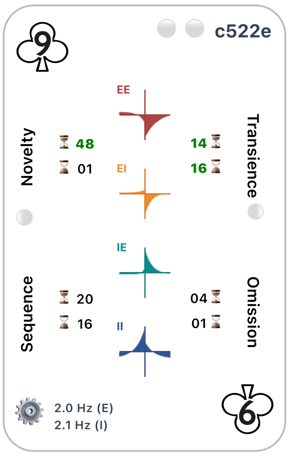
c522e
EE
EI
IE
II
Novelty
48
01
Transience
14
16
Sequence
20
16
Omission
04
01
2.0 Hz (E)
2.1 Hz (I)

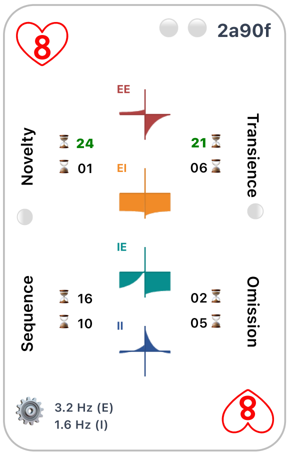
2a90f
EE
EI
IE
II
Novelty
24
01
Transience
21
06
Sequence
16
10
Omission
02
05
3.2 Hz (E)
1.6 Hz (I)

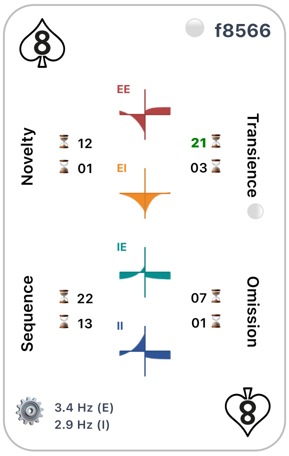
f8566
EE
EI
IE
II
Novelty
12
01
Transience
21
03
Sequence
22
13
Omission
07
01
3.4 Hz (E)
2.9 Hz (I)

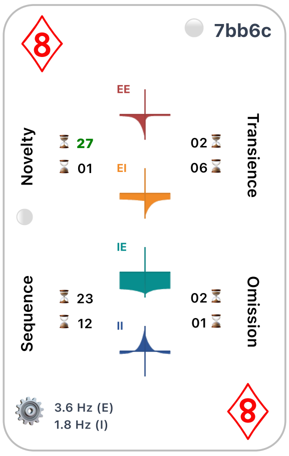
7bb6c
EE
EI
IE
II
Novelty
27
01
Transience
02
06
Sequence
23
12
Omission
02
01
3.6 Hz (E)
1.8 Hz (I)

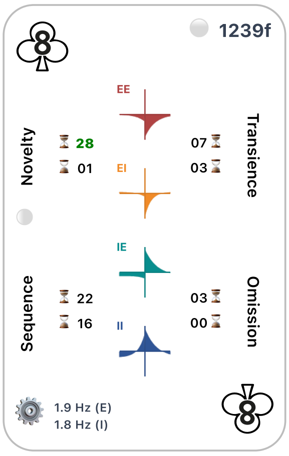
1239f
EE
EI
IE
II
Novelty
28
01
Transience
07
03
Sequence
22
16
Omission
03
00
1.9 Hz (E)
1.8 Hz (I)

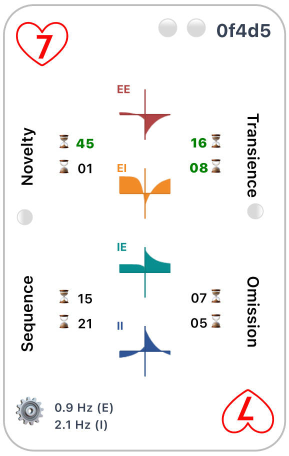
0f4d5
EE
EI
IE
II
Novelty
45
01
Transience
16
08
Sequence
15
21
Omission
07
05
0.9 Hz (E)
2.1 Hz (I)

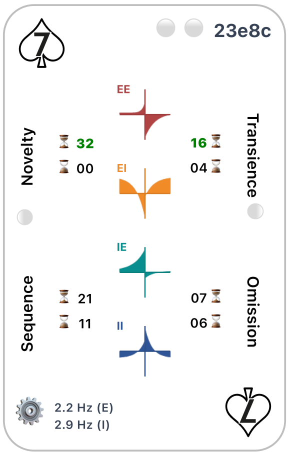
23e8c
EE
EI
IE
II
Novelty
32
00
Transience
16
04
Sequence
21
11
Omission
07
06
2.2 Hz (E)
2.9 Hz (I)

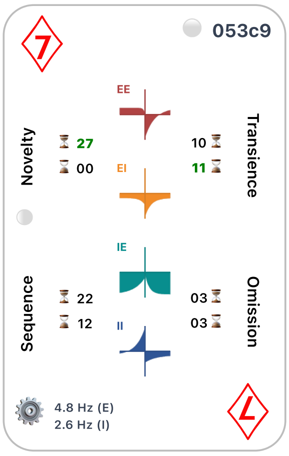
053c9
EE
EI
IE
II
Novelty
27
00
Transience
10
11
Sequence
22
12
Omission
03
03
4.8 Hz (E)
2.6 Hz (I)

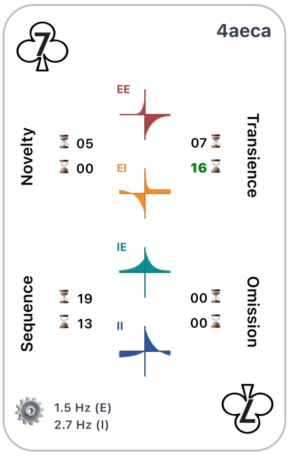
4aeca
EE
EI
IE
II
Novelty
05
00
Transience
07
16
Sequence
19
13
Omission
00
00
1.5 Hz (E)
2.7 Hz (I)

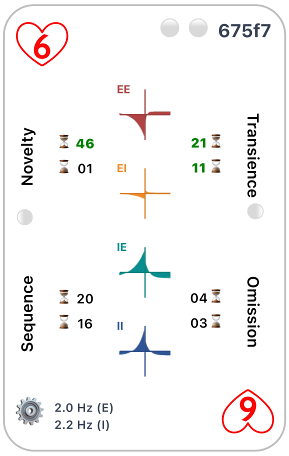
675f7
EE
EI
IE
II
Novelty
46
01
Transience
21
11
Sequence
20
16
Omission
04
03
2.0 Hz (E)
2.2 Hz (I)

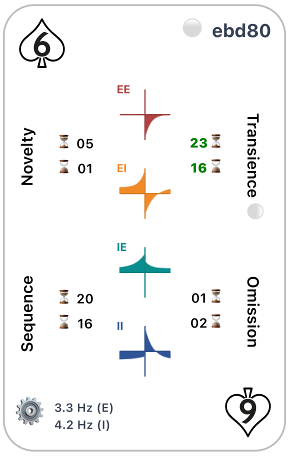
ebd80
EE
EI
IE
II
Novelty
05
01
Transience
23
16
Sequence
20
16
Omission
01
02
3.3 Hz (E)
4.2 Hz (I)

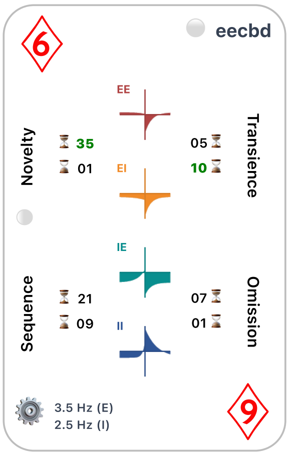
eecbd
EE
EI
IE
II
Novelty
35
01
Transience
05
10
Sequence
21
09
Omission
07
01
3.5 Hz (E)
2.5 Hz (I)

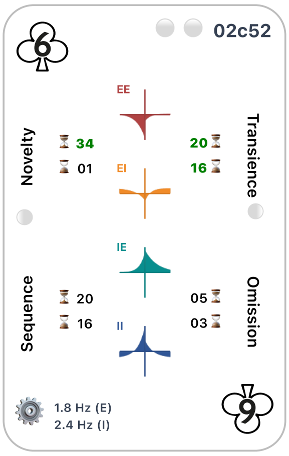
02c52
EE
EI
IE
II
Novelty
34
01
Transience
20
16
Sequence
20
16
Omission
05
03
1.8 Hz (E)
2.4 Hz (I)

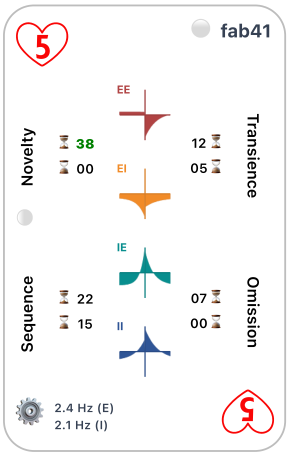
5
fab41
EE
Novelty
38
00
12
05
Transience
EI
IE
Sequence
22
15
07
00
Omission
II
2.4 Hz (E)
2.1 Hz (I)

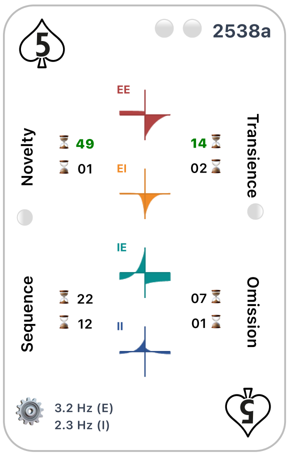
5
2538a
EE
Novelty
49
01
14
02
Transience
EI
IE
Sequence
22
12
07
01
Omission
II
3.2 Hz (E)
2.3 Hz (I)

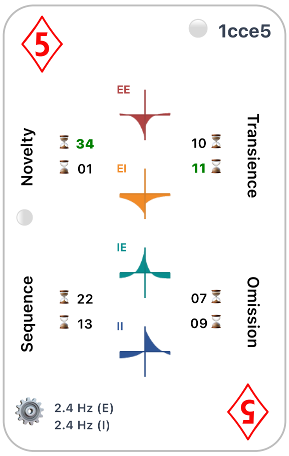
5
1cce5
EE
Novelty
34
01
10
11
Transience
EI
IE
Sequence
22
13
07
09
Omission
II
2.4 Hz (E)
2.4 Hz (I)

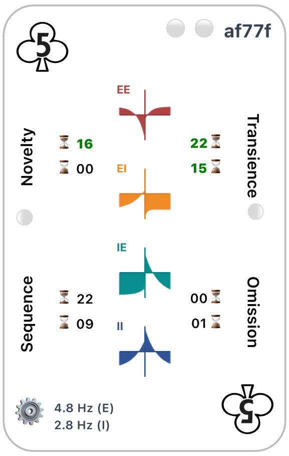
5
af77f
EE
Novelty
16
00
22
15
Transience
EI
IE
Sequence
22
09
00
01
Omission
II
4.8 Hz (E)
2.8 Hz (I)

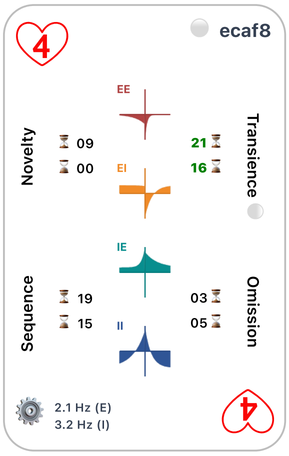
4
ecaf8
EE
Novelty
09
00
21
16
Transience
EI
IE
Sequence
19
15
03
05
Omission
II
2.1 Hz (E)
3.2 Hz (I)

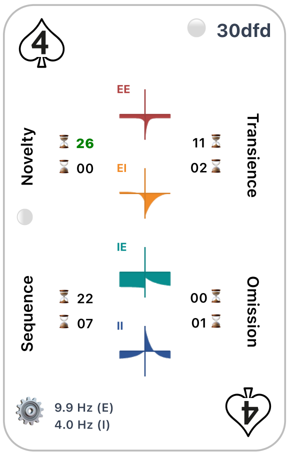
4
30dfd
EE
Novelty
26
00
11
02
Transience
EI
IE
Sequence
22
07
00
01
Omission
II
9.9 Hz (E)
4.0 Hz (I)

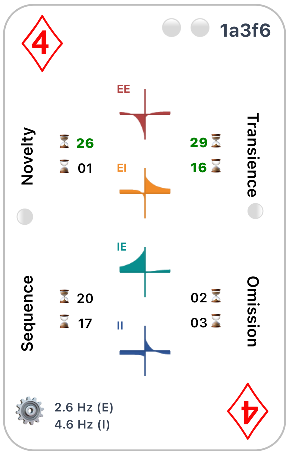
4
1a3f6
EE
Novelty
26
01
29
16
Transience
EI
IE
Sequence
20
17
02
03
Omission
II
2.6 Hz (E)
4.6 Hz (I)

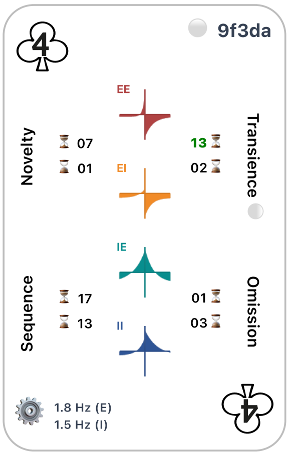
4
9f3da
EE
Novelty
07
01
13
02
Transience
EI
IE
Sequence
17
13
01
03
Omission
II
1.8 Hz (E)
1.5 Hz (I)

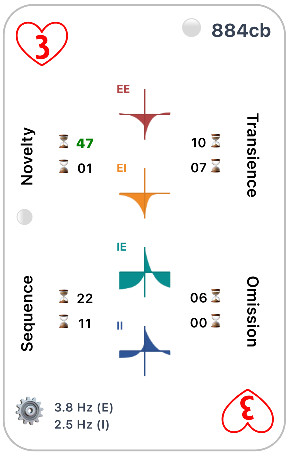
3
884cb
EE
Novelty
47
01
10
07
Transience
EI
IE
Sequence
22
11
06
00
Omission
II
3.8 Hz (E)
2.5 Hz (I)

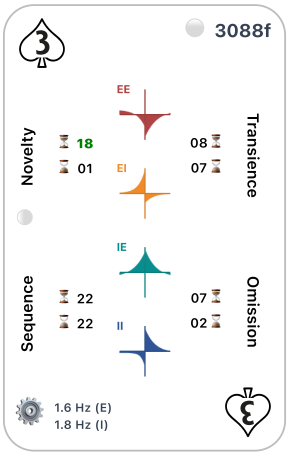
3
3088f
EE
Novelty
18
01
08
07
Transience
EI
IE
Sequence
22
22
07
02
Omission
II
1.6 Hz (E)
1.8 Hz (I)

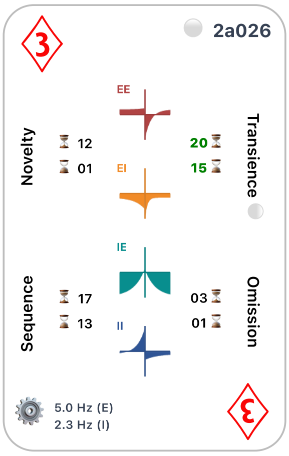
3
2a026
EE
Novelty
12
01
20
15
Transience
EI
IE
Sequence
17
13
03
01
Omission
II
5.0 Hz (E)
2.3 Hz (I)

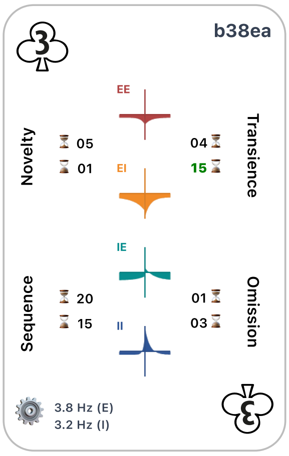
3
b38ea
EE
Novelty
05
01
04
15
Transience
EI
IE
Sequence
20
15
01
03
Omission
II
3.8 Hz (E)
3.2 Hz (I)

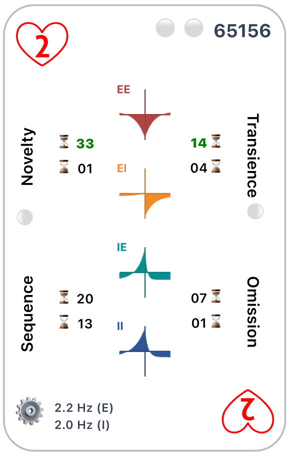
2
65156
EE
Novelty
33
01
14
04
Transience
EI
IE
Sequence
20
13
07
01
Omission
II
2.2 Hz (E)
2.0 Hz (I)

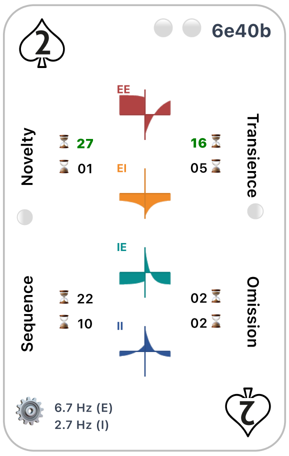
2
6e40b
EE
Novelty
27
01
16
05
Transience
EI
IE
Sequence
22
10
02
02
Omission
II
6.7 Hz (E)
2.7 Hz (I)

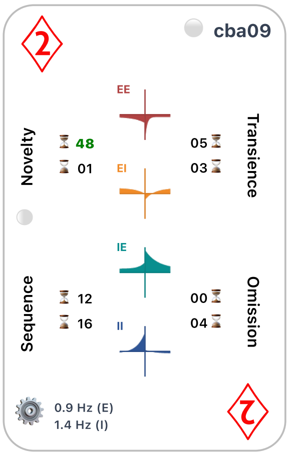
2
cba09
EE
Novelty
48
01
05
03
Transience
EI
IE
Sequence
12
16
00
04
Omission
II
0.9 Hz (E)
1.4 Hz (I)

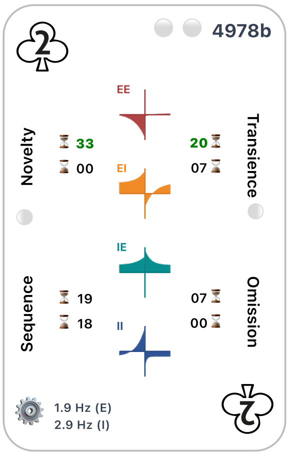
2
4978b
EE
Novelty
33
00
20
07
Transience
EI
IE
Sequence
19
18
07
00
Omission
II
1.9 Hz (E)
2.9 Hz (I)

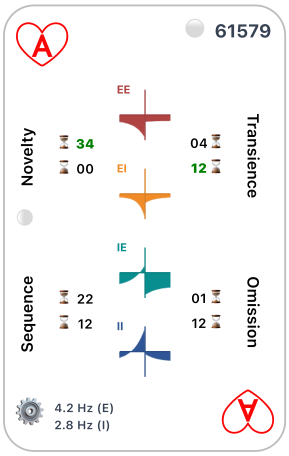
A
61579
EE
EI
IE
II
Novelty
34
00
04
12
Transience
Sequence
22
12
01
12
Omission
4.2 Hz (E)
2.8 Hz (I)

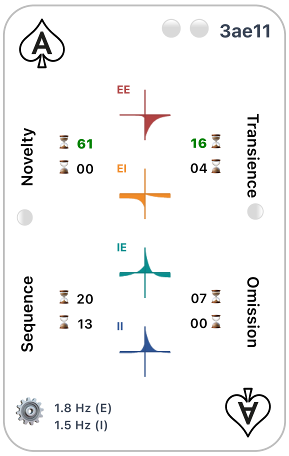
A
3ae11
EE
EI
IE
II
Novelty
61
00
16
04
Transience
Sequence
20
13
07
00
Omission
1.8 Hz (E)
1.5 Hz (I)

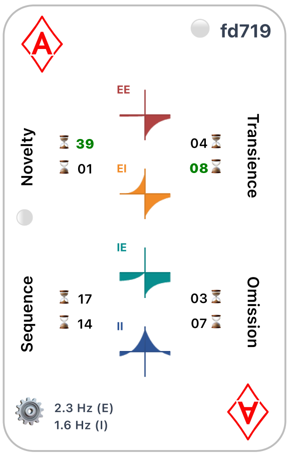
A
fd719
EE
EI
IE
II
Novelty
39
01
04
08
Transience
Sequence
17
14
03
07
Omission
2.3 Hz (E)
1.6 Hz (I)

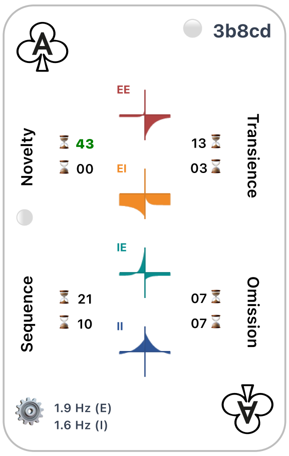
A
3b8cd
EE
EI
IE
II
Novelty
43
00
13
03
Transience
Sequence
21
10
07
07
Omission
1.9 Hz (E)
1.6 Hz (I)

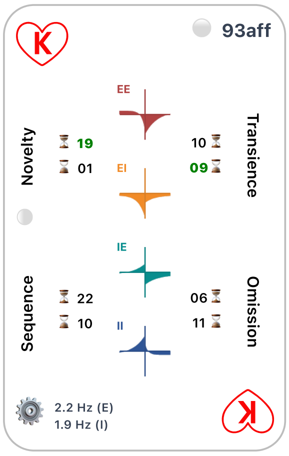
K
93aff
EE
EI
IE
II
Novelty
19
01
10
09
Transience
Sequence
22
10
06
11
Omission
2.2 Hz (E)
1.9 Hz (I)

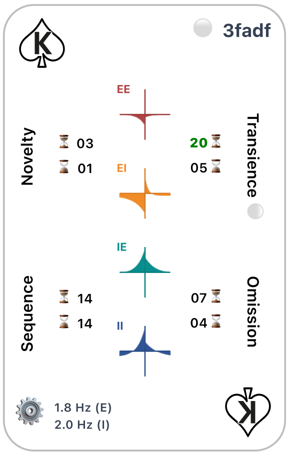
K
3fadf
EE
EI
IE
II
Novelty
03
01
20
05
Transience
Sequence
14
14
07
04
Omission
1.8 Hz (E)
2.0 Hz (I)

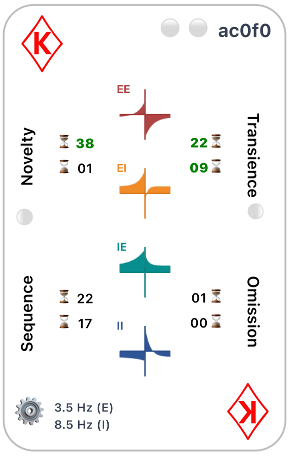
K
ac0f0
EE
EI
IE
II
Novelty
38
01
22
09
Transience
Sequence
22
17
01
00
Omission
3.5 Hz (E)
8.5 Hz (I)

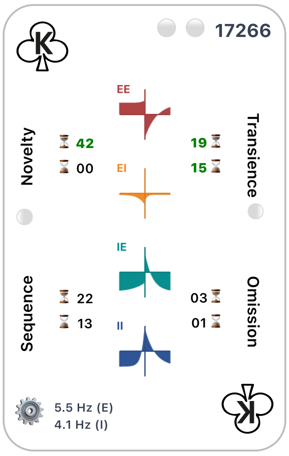
K
17266
EE
EI
IE
II
Novelty
42
00
19
15
Transience
Sequence
22
13
03
01
Omission
5.5 Hz (E)
4.1 Hz (I)

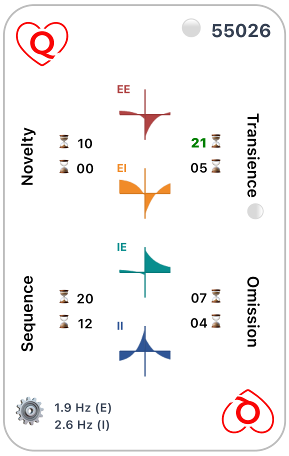
Q
55026
EE
EI
IE
II
Novelty
10
00
21
05
Transience
Sequence
20
12
07
04
Omission
1.9 Hz (E)
2.6 Hz (I)

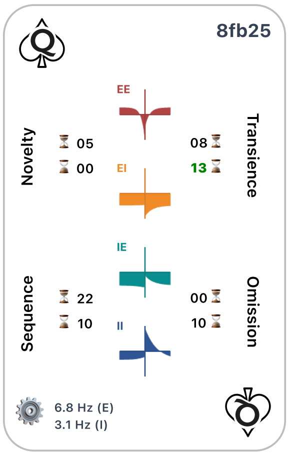
Q
8fb25
EE
EI
IE
II
Novelty
05
00
08
13
Transience
Sequence
22
10
00
10
Omission
6.8 Hz (E)
3.1 Hz (I)

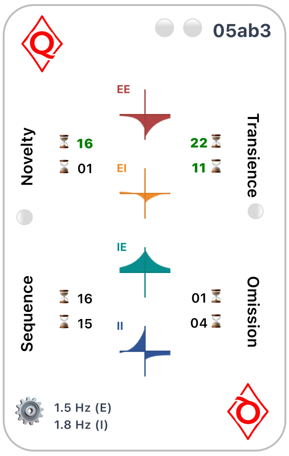
Q
05ab3
EE
EI
IE
II
Novelty
16
01
22
11
Transience
Sequence
16
15
01
04
Omission
1.5 Hz (E)
1.8 Hz (I)

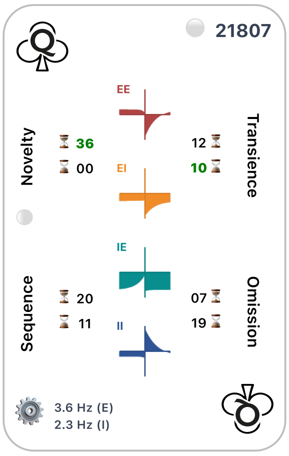
Q
21807
EE
EI
IE
II
Novelty
36
00
12
10
Transience
Sequence
20
11
07
19
Omission
3.6 Hz (E)
2.3 Hz (I)

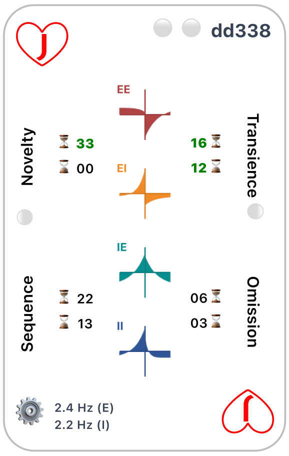
J
dd338
EE
EI
IE
II
Novelty
33
00
16
12
Transience
Sequence
22
13
06
03
Omission
2.4 Hz (E)
2.2 Hz (I)

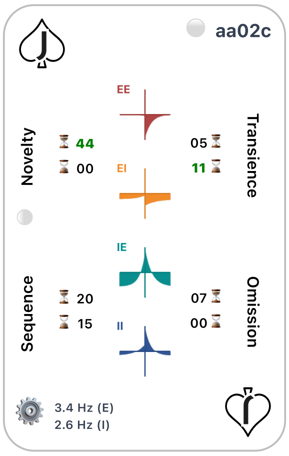
J
aa02c
EE
EI
IE
II
Novelty
44
00
05
11
Transience
Sequence
20
15
07
00
Omission
3.4 Hz (E)
2.6 Hz (I)

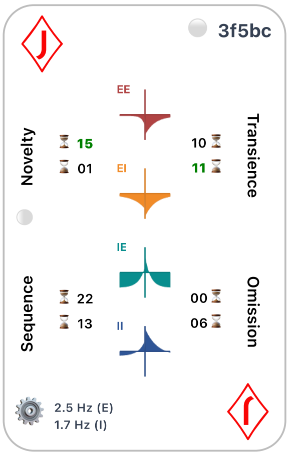
J
3f5bc
EE
EI
IE
II
Novelty
15
01
10
11
Transience
Sequence
22
13
00
06
Omission
2.5 Hz (E)
1.7 Hz (I)

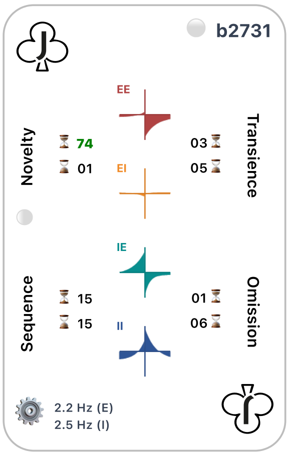
J
b2731
EE
EI
IE
II
Novelty
74
01
03
05
Transience
Sequence
15
15
01
06
Omission
2.2 Hz (E)
2.5 Hz (I)

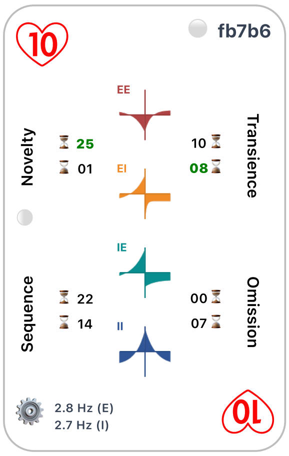
10
fb7b6
Novelty
25
01
Transience
10
08
EE
EI
IE
II
Sequence
22
14
Omission
00
07
2.8 Hz (E)
2.7 Hz (I)

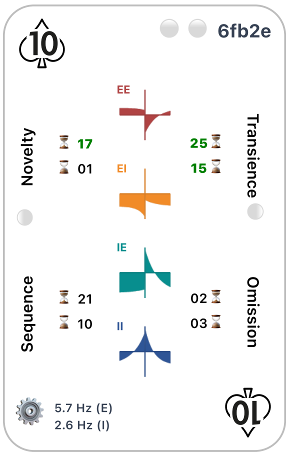
10
6fb2e
Novelty
17
01
Transience
25
15
EE
EI
IE
II
Sequence
21
10
Omission
02
03
5.7 Hz (E)
2.6 Hz (I)

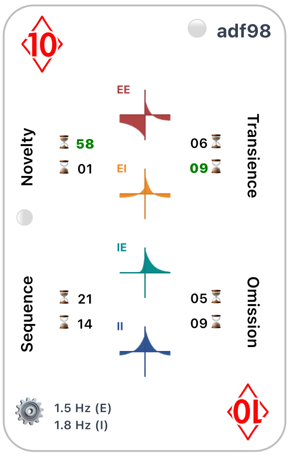
10
adf98
Novelty
58
01
Transience
06
09
EE
EI
IE
II
Sequence
21
14
Omission
05
09
1.5 Hz (E)
1.8 Hz (I)

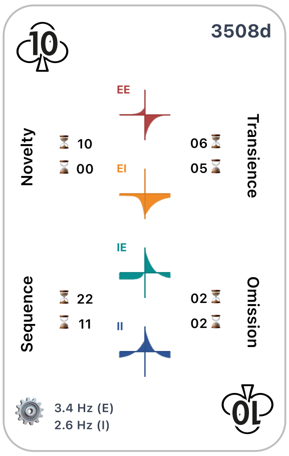
10
3508d
Novelty
10
00
Transience
06
05
EE
EI
IE
II
Sequence
22
11
Omission
02
02
3.4 Hz (E)
2.6 Hz (I)

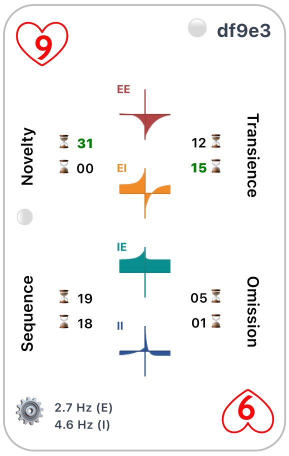
9
df9e3
Novelty
31
00
Transience
12
15
EE
EI
IE
II
Sequence
19
18
Omission
05
01
2.7 Hz (E)
4.6 Hz (I)

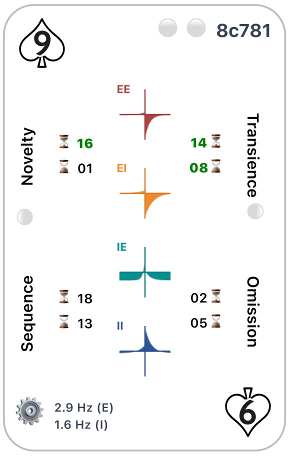
9
8c781
Novelty
16
01
Transience
14
08
EE
EI
IE
II
Sequence
18
13
Omission
02
05
2.9 Hz (E)
1.6 Hz (I)

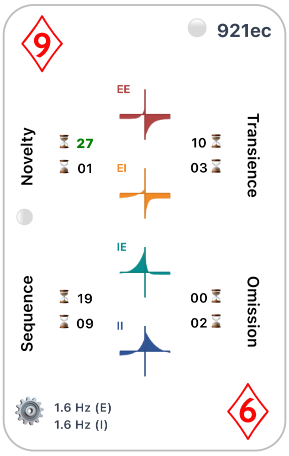
9
921ec
Novelty
27
01
Transience
10
03
EE
EI
IE
II
Sequence
19
09
Omission
00
02
1.6 Hz (E)
1.6 Hz (I)

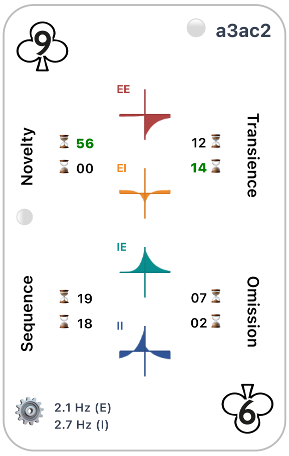
9
a3ac2
Novelty
56
00
Transience
12
14
EE
EI
IE
II
Sequence
19
18
Omission
07
02
2.1 Hz (E)
2.7 Hz (I)

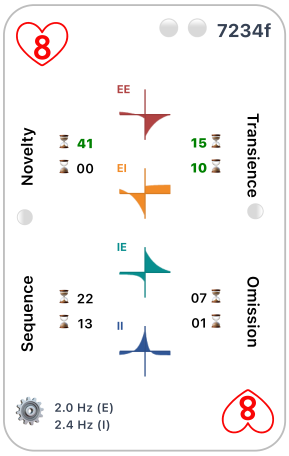
8
7234f
Novelty
41
00
Transience
15
10
EE
EI
IE
II
Sequence
22
13
Omission
07
01
2.0 Hz (E)
2.4 Hz (I)

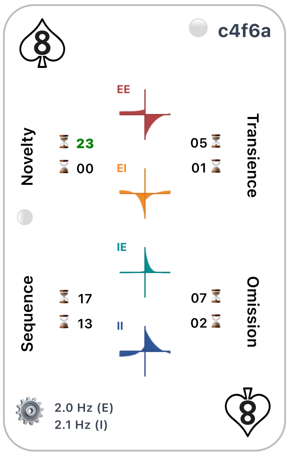
8
c4f6a
Novelty
23
00
Transience
05
01
EE
EI
IE
II
Sequence
17
13
Omission
07
02
2.0 Hz (E)
2.1 Hz (I)

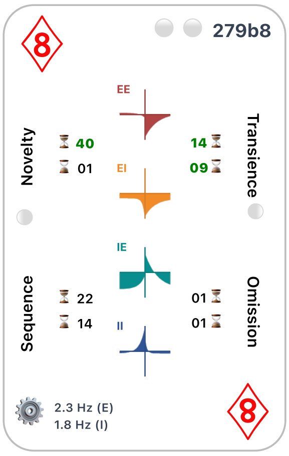
8
279b8
Novelty
40
01
Transience
14
09
EE
EI
IE
II
Sequence
22
14
Omission
01
01
2.3 Hz (E)
1.8 Hz (I)

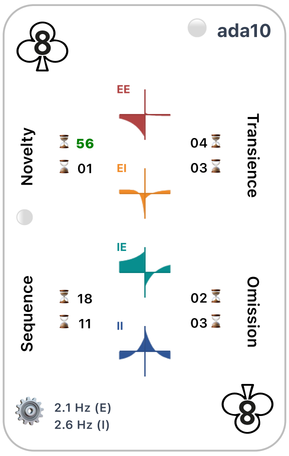
8
ada10
Novelty
56
01
Transience
04
03
EE
EI
IE
II
Sequence
18
11
Omission
02
03
2.1 Hz (E)
2.6 Hz (I)

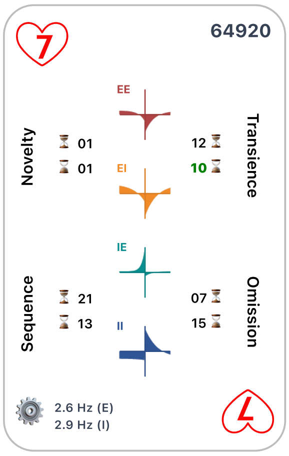
7
64920
Novelty
01
01
Transience
12
10
EE
EI
IE
II
Sequence
21
13
Omission
07
15
2.6 Hz (E)
2.9 Hz (I)

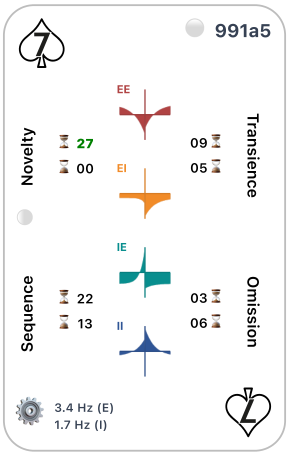
7
991a5
Novelty
27
00
Transience
09
05
EE
EI
IE
II
Sequence
22
13
Omission
03
06
3.4 Hz (E)
1.7 Hz (I)

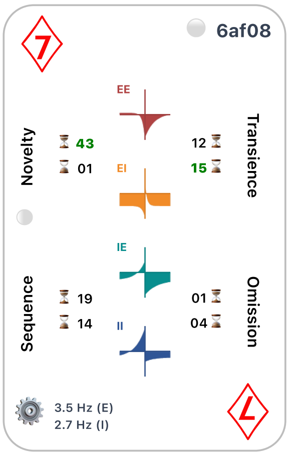
7
6af08
Novelty
43
01
Transience
12
15
EE
EI
IE
II
Sequence
19
14
Omission
01
04
3.5 Hz (E)
2.7 Hz (I)

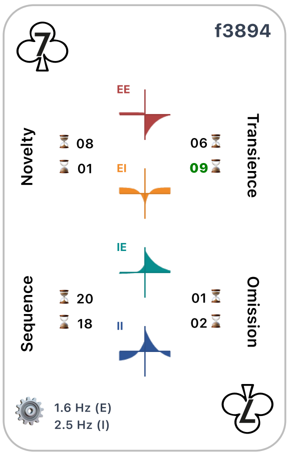
7
f3894
Novelty
08
01
Transience
06
09
EE
EI
IE
II
Sequence
20
18
Omission
01
02
1.6 Hz (E)
2.5 Hz (I)

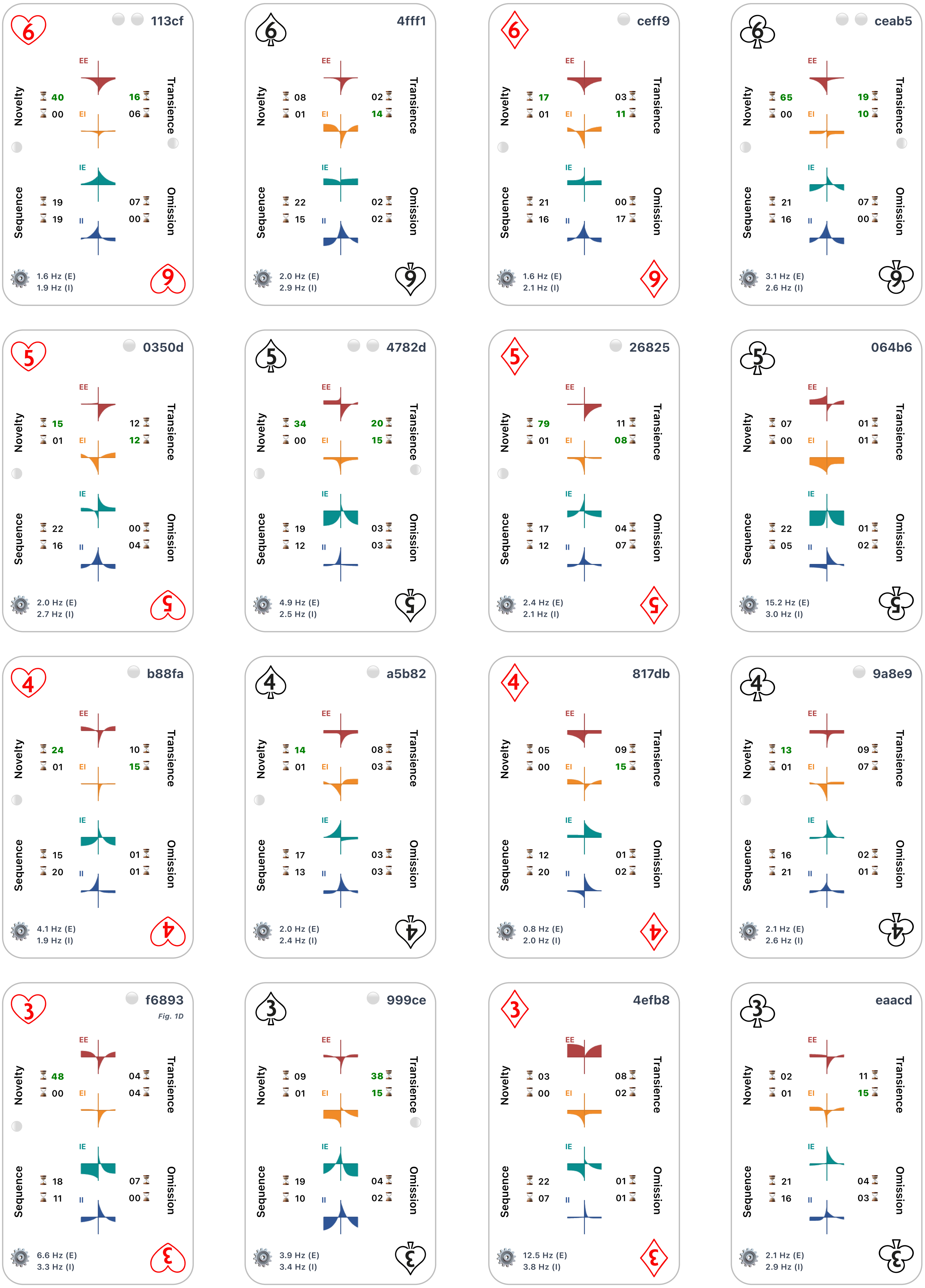

113cf
Novelty 40 00
Transience 16 06
Sequence 19 19
Omission 07 00
1.6 Hz (E) 1.9 Hz (I)
4fff1
Novelty 08 01
Transience 02 14
Sequence 22 15
Omission 02 02
2.0 Hz (E) 2.9 Hz (I)
ceff9
Novelty 17 01
Transience 03 11
Sequence 21 16
Omission 00 17
1.6 Hz (E) 2.1 Hz (I)
ceab5
Novelty 65 00
Transience 19 10
Sequence 21 16
Omission 07 00
3.1 Hz (E) 2.6 Hz (I)
0350d
Novelty 15 01
Transience 12 12
Sequence 22 16
Omission 00 04
2.0 Hz (E) 2.7 Hz (I)
4782d
Novelty 34 00
Transience 20 15
Sequence 19 12
Omission 03 03
4.9 Hz (E) 2.5 Hz (I)
26825
Novelty 79 01
Transience 11 08
Sequence 17 12
Omission 04 07
2.4 Hz (E) 2.1 Hz (I)
064b6
Novelty 07 00
Transience 01 01
Sequence 22 05
Omission 01 02
15.2 Hz (E) 3.0 Hz (I)
b88fa
Novelty 24 01
Transience 10 15
Sequence 15 20
Omission 01 01
4.1 Hz (E) 1.9 Hz (I)
a5b82
Novelty 14 01
Transience 08 03
Sequence 17 13
Omission 03 03
2.0 Hz (E) 2.4 Hz (I)
817db
Novelty 05 00
Transience 09 15
Sequence 12 20
Omission 01 02
0.8 Hz (E) 2.0 Hz (I)
9a8e9
Novelty 13 01
Transience 09 07
Sequence 16 21
Omission 02 01
2.1 Hz (E) 2.6 Hz (I)
f6893
Fig. 1D
Novelty 48 00
Transience 04 04
Sequence 18 11
Omission 07 00
6.6 Hz (E) 3.3 Hz (I)
999ce
Novelty 09 01
Transience 38 15
Sequence 19 10
Omission 04 02
3.9 Hz (E) 3.4 Hz (I)
4efb8
Novelty 03 00
Transience 08 02
Sequence 22 07
Omission 01 01
12.5 Hz (E) 3.8 Hz (I)
eaacd
Novelty 02 01
Transience 11 15
Sequence 21 16
Omission 04 03
2.1 Hz (E) 2.9 Hz (I)

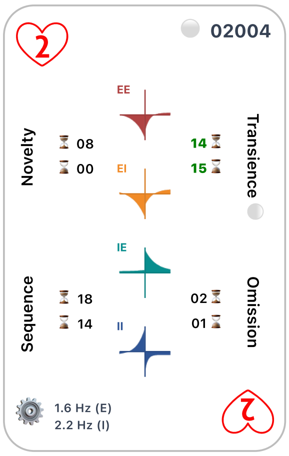
2
02004
EE
EI
IE
II
Novelty
08
00
Transience
14
15
Sequence
18
14
Omission
02
01
1.6 Hz (E)
2.2 Hz (I)

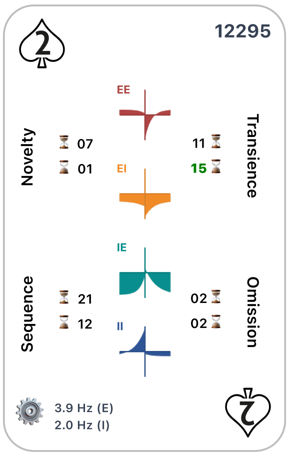
2
12295
EE
EI
IE
II
Novelty
07
01
Transience
11
15
Sequence
21
12
Omission
02
02
3.9 Hz (E)
2.0 Hz (I)

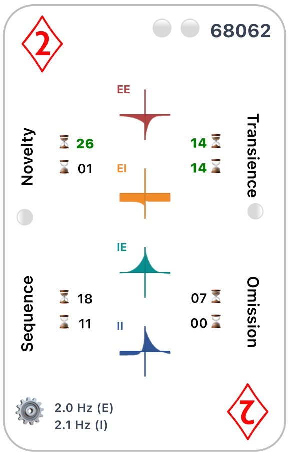
2
68062
EE
EI
IE
II
Novelty
26
01
Transience
14
14
Sequence
18
11
Omission
07
00
2.0 Hz (E)
2.1 Hz (I)

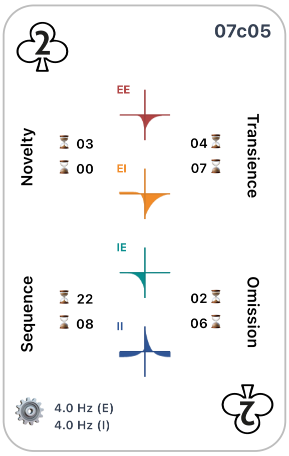
2
07c05
EE
EI
IE
II
Novelty
03
00
Transience
04
07
Sequence
22
08
Omission
02
06
4.0 Hz (E)
4.0 Hz (I)

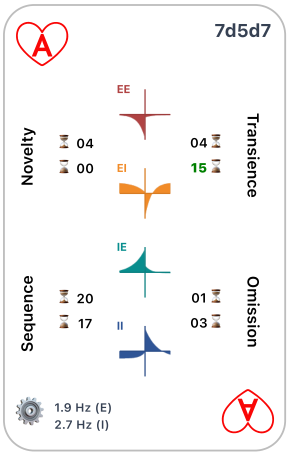
A
7d5d7
EE
EI
IE
II
Novelty
04
00
Transience
04
15
Sequence
20
17
Omission
01
03
1.9 Hz (E)
2.7 Hz (I)

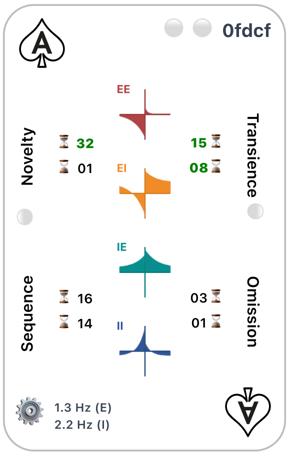
A
0fdcf
EE
EI
IE
II
Novelty
32
01
Transience
15
08
Sequence
16
14
Omission
03
01
1.3 Hz (E)
2.2 Hz (I)

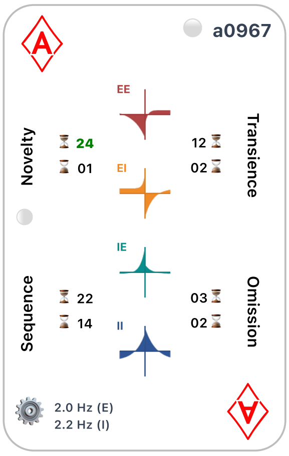
A
a0967
EE
EI
IE
II
Novelty
24
01
Transience
12
02
Sequence
22
14
Omission
03
02
2.0 Hz (E)
2.2 Hz (I)

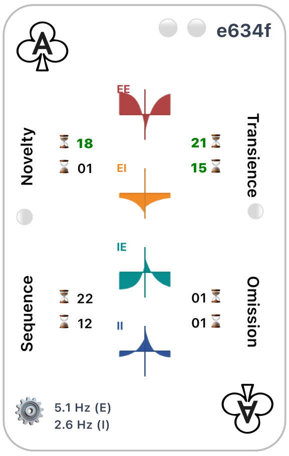
A
e634f
EE
EI
IE
II
Novelty
18
01
Transience
21
15
Sequence
22
12
Omission
01
01
5.1 Hz (E)
2.6 Hz (I)

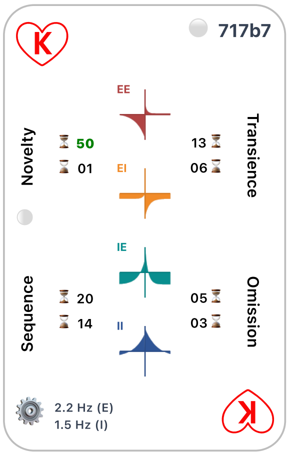
K
717b7
EE
EI
IE
II
Novelty
50
01
Transience
13
06
Sequence
20
14
Omission
05
03
2.2 Hz (E)
1.5 Hz (I)

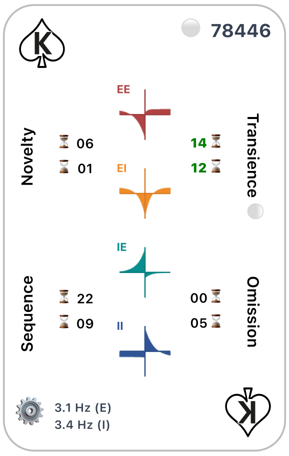
K
78446
EE
EI
IE
II
Novelty
06
01
Transience
14
12
Sequence
22
09
Omission
00
05
3.1 Hz (E)
3.4 Hz (I)

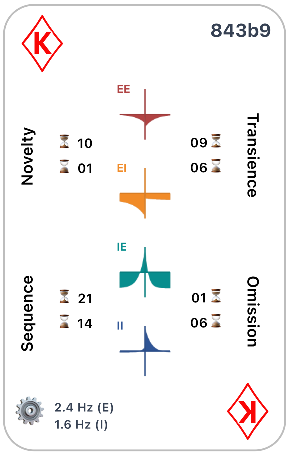
K
843b9
EE
EI
IE
II
Novelty
10
01
Transience
09
06
Sequence
21
14
Omission
01
06
2.4 Hz (E)
1.6 Hz (I)

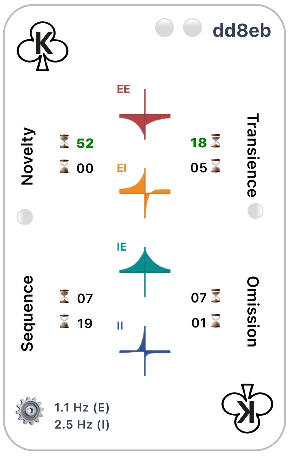
K
dd8eb
EE
EI
IE
II
Novelty
52
00
Transience
18
05
Sequence
07
19
Omission
07
01
1.1 Hz (E)
2.5 Hz (I)

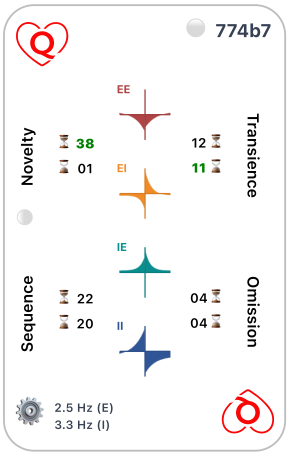
Q
774b7
EE
EI
IE
II
Novelty
38
01
Transience
12
11
Sequence
22
20
Omission
04
04
2.5 Hz (E)
3.3 Hz (I)

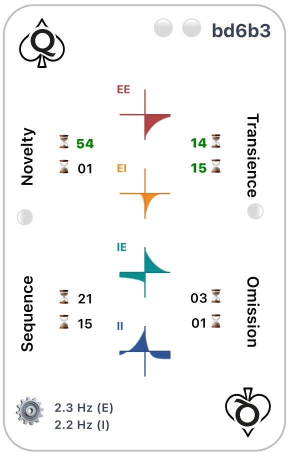
Q
bd6b3
EE
EI
IE
II
Novelty
54
01
Transience
14
15
Sequence
21
15
Omission
03
01
2.3 Hz (E)
2.2 Hz (I)

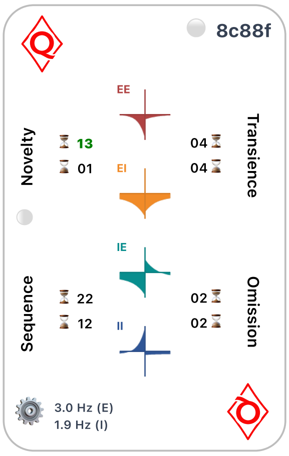
Q
8c88f
EE
EI
IE
II
Novelty
13
01
Transience
04
04
Sequence
22
12
Omission
02
02
3.0 Hz (E)
1.9 Hz (I)

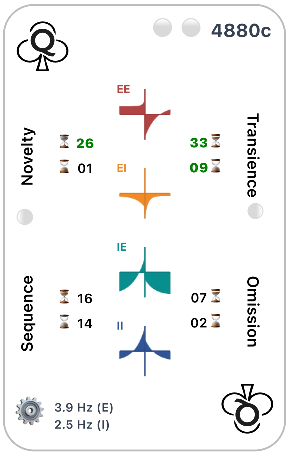
Q
4880c
EE
EI
IE
II
Novelty
26
01
Transience
33
09
Sequence
16
14
Omission
07
02
3.9 Hz (E)
2.5 Hz (I)

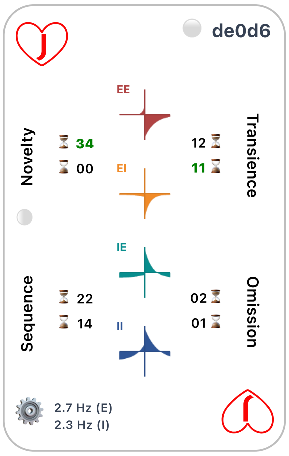
J
de0d6
EE
Novelty
34
00
12
11
Transience
EI
IE
Sequence
22
14
02
01
Omission
II
2.7 Hz (E)
2.3 Hz (I)
J

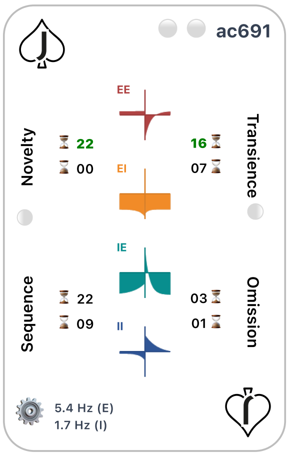
J
ac691
EE
Novelty
22
00
16
07
Transience
EI
IE
Sequence
22
09
03
01
Omission
II
5.4 Hz (E)
1.7 Hz (I)
J

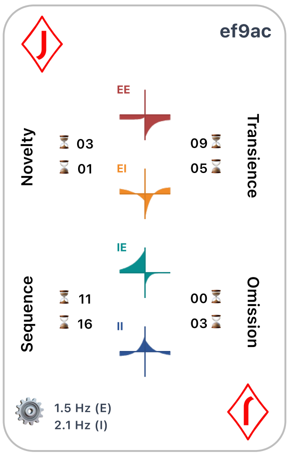
J
ef9ac
EE
Novelty
03
01
09
05
Transience
EI
IE
Sequence
11
16
00
03
Omission
II
1.5 Hz (E)
2.1 Hz (I)
J

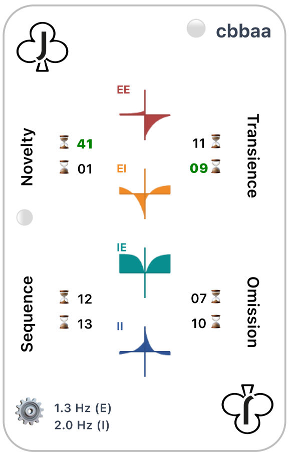
J
cbbaa
EE
Novelty
41
01
11
09
Transience
EI
IE
Sequence
12
13
07
10
Omission
II
1.3 Hz (E)
2.0 Hz (I)
J

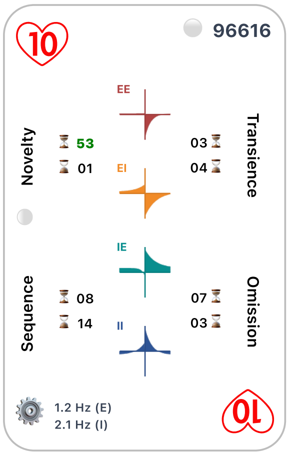
10
96616
EE
Novelty
53
01
03
04
Transience
EI
IE
Sequence
08
14
07
03
Omission
II
1.2 Hz (E)
2.1 Hz (I)
10

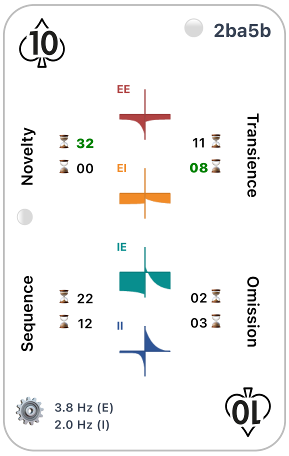
10
2ba5b
EE
Novelty
32
00
11
08
Transience
EI
IE
Sequence
22
12
02
03
Omission
II
3.8 Hz (E)
2.0 Hz (I)
10

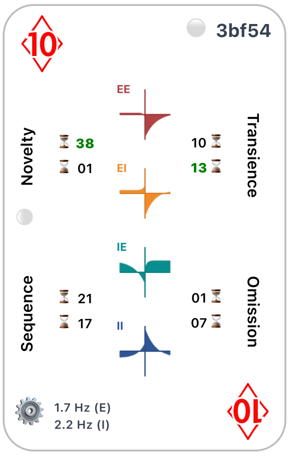
10
3bf54
EE
Novelty
38
01
10
13
Transience
EI
IE
Sequence
21
17
01
07
Omission
II
1.7 Hz (E)
2.2 Hz (I)
10

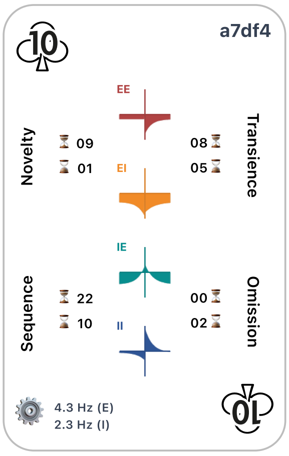
10
a7df4
EE
Novelty
09
01
08
05
Transience
EI
IE
Sequence
22
10
00
02
Omission
II
4.3 Hz (E)
2.3 Hz (I)
10

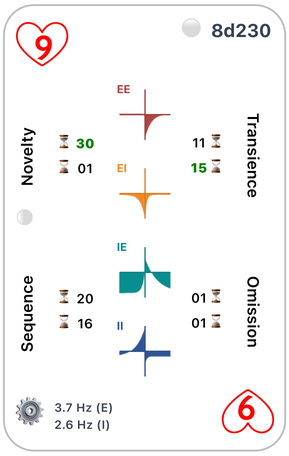
9
8d230
EE
Novelty
30
01
11
15
Transience
EI
IE
Sequence
20
16
01
01
Omission
II
3.7 Hz (E)
2.6 Hz (I)
9

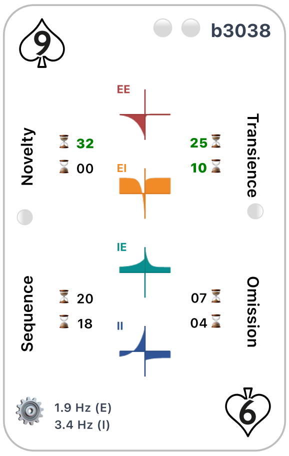
9
b3038
EE
Novelty
32
00
25
10
Transience
EI
IE
Sequence
20
18
07
04
Omission
II
1.9 Hz (E)
3.4 Hz (I)
9

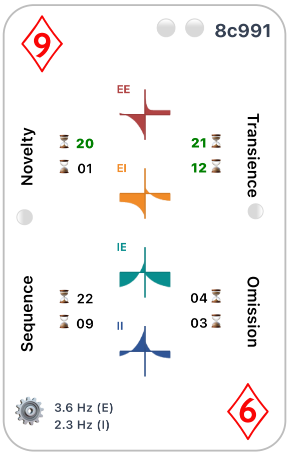
9
8c991
EE
Novelty
20
01
21
12
Transience
EI
IE
Sequence
22
09
04
03
Omission
II
3.6 Hz (E)
2.3 Hz (I)
9

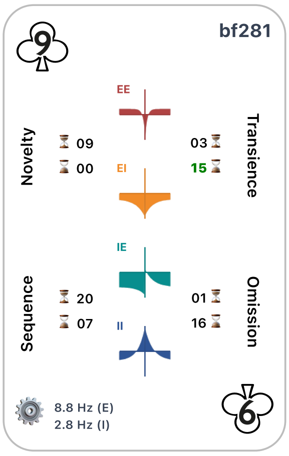
9
bf281
EE
Novelty
09
00
03
15
Transience
EI
IE
Sequence
20
07
01
16
Omission
II
8.8 Hz (E)
2.8 Hz (I)
9

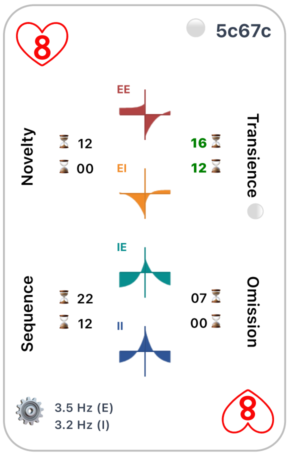
8
5c67c
EE
Novelty
12
00
16
12
Transience
EI
IE
Sequence
22
12
07
00
Omission
II
3.5 Hz (E)
3.2 Hz (I)
8

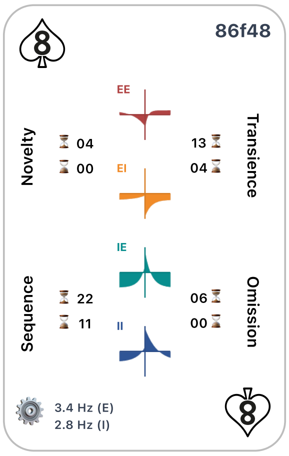
8
86f48
EE
Novelty
04
00
13
04
Transience
EI
IE
Sequence
22
11
06
00
Omission
II
3.4 Hz (E)
2.8 Hz (I)
8

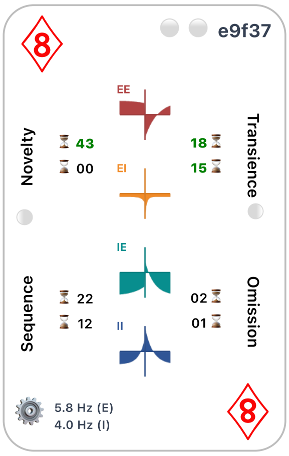
8
e9f37
EE
Novelty
43
00
18
15
Transience
EI
IE
Sequence
22
12
02
01
Omission
II
5.8 Hz (E)
4.0 Hz (I)
8

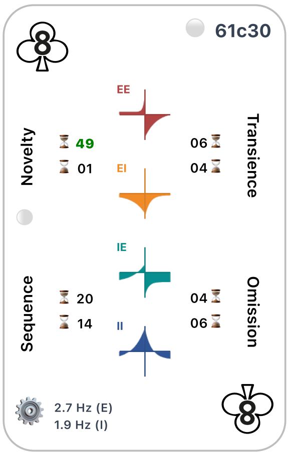
8
61c30
EE
Novelty
49
01
06
04
Transience
EI
IE
Sequence
20
14
04
06
Omission
II
2.7 Hz (E)
1.9 Hz (I)
8

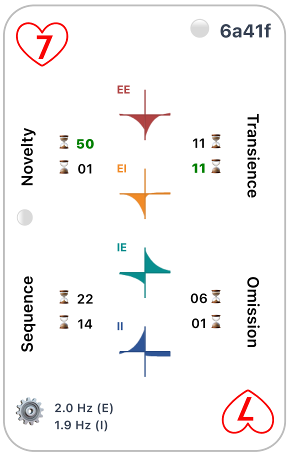
7
6a41f
Novelty
50
01
Transience
11
11
Sequence
22
14
Omission
06
01
EE
EI
IE
II
2.0 Hz (E)
1.9 Hz (I)

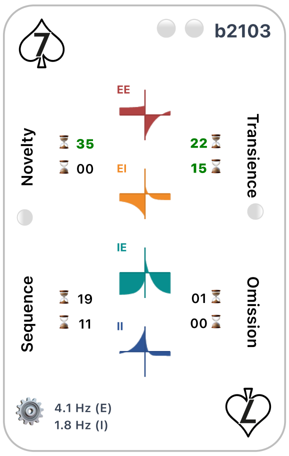
7
b2103
Novelty
35
00
Transience
22
15
Sequence
19
11
Omission
01
00
EE
EI
IE
II
4.1 Hz (E)
1.8 Hz (I)

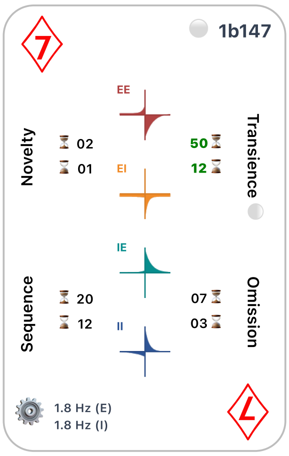
7
1b147
Novelty
02
01
Transience
50
12
Sequence
20
12
Omission
07
03
EE
EI
IE
II
1.8 Hz (E)
1.8 Hz (I)

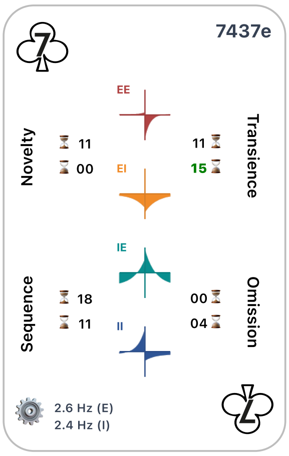
7
7437e
Novelty
11
00
Transience
11
15
Sequence
18
11
Omission
00
04
EE
EI
IE
II
2.6 Hz (E)
2.4 Hz (I)

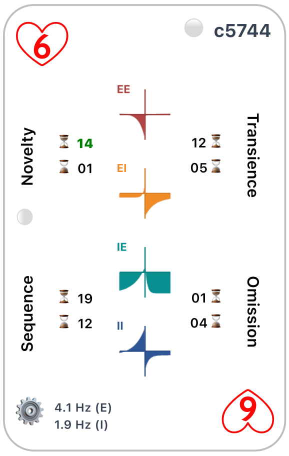
6
c5744
Novelty
14
01
Transience
12
05
Sequence
19
12
Omission
01
04
EE
EI
IE
II
4.1 Hz (E)
1.9 Hz (I)

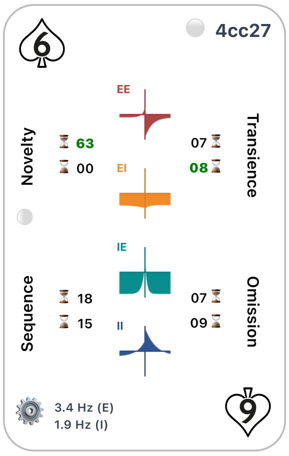
6
4cc27
Novelty
63
00
Transience
07
08
Sequence
18
15
Omission
07
09
EE
EI
IE
II
3.4 Hz (E)
1.9 Hz (I)

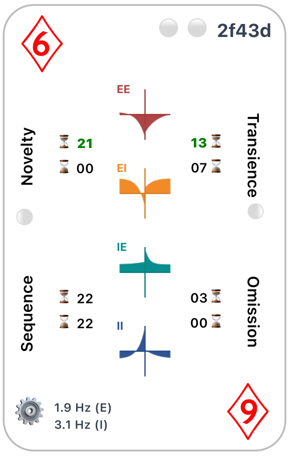
6
2f43d
Novelty
21
00
Transience
13
07
Sequence
22
22
Omission
03
00
EE
EI
IE
II
1.9 Hz (E)
3.1 Hz (I)

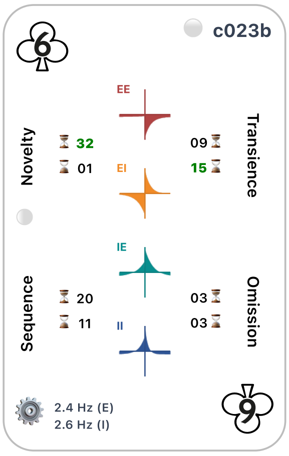
6
c023b
Novelty
32
01
Transience
09
15
Sequence
20
11
Omission
03
03
EE
EI
IE
II
2.4 Hz (E)
2.6 Hz (I)

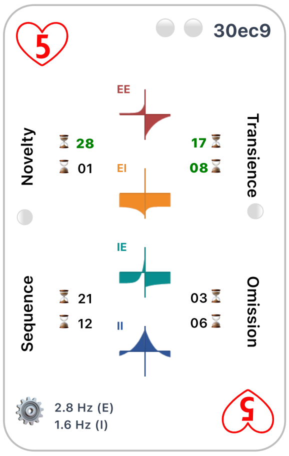
5
30ec9
Novelty
28
01
Transience
17
08
Sequence
21
12
Omission
03
06
EE
EI
IE
II
2.8 Hz (E)
1.6 Hz (I)

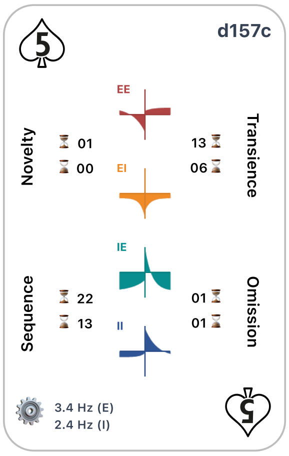
5
d157c
Novelty
01
00
Transience
13
06
Sequence
22
13
Omission
01
01
EE
EI
IE
II
3.4 Hz (E)
2.4 Hz (I)

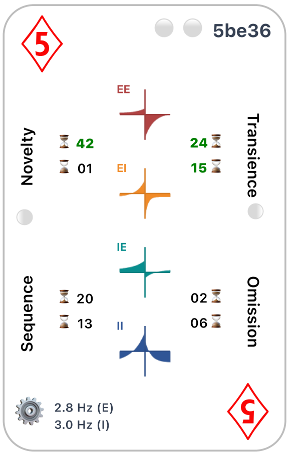
5
5be36
Novelty
42
01
Transience
24
15
Sequence
20
13
Omission
02
06
EE
EI
IE
II
2.8 Hz (E)
3.0 Hz (I)

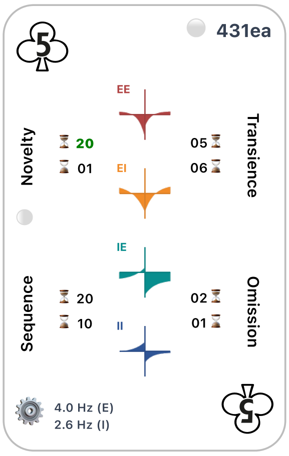
5
431ea
Novelty
20
01
Transience
05
06
Sequence
20
10
Omission
02
01
EE
EI
IE
II
4.0 Hz (E)
2.6 Hz (I)

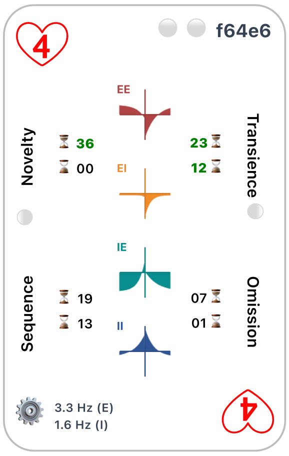
4
f64e6
Novelty
36
00
Transience
23
12
Sequence
19
13
Omission
07
01
EE
EI
IE
II
3.3 Hz (E)
1.6 Hz (I)

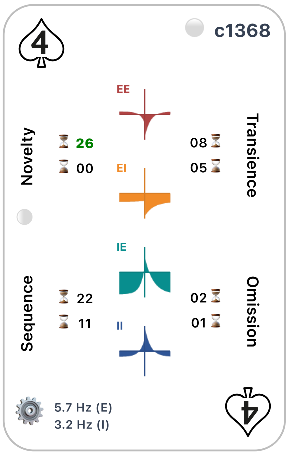
4
c1368
Novelty
26
00
Transience
08
05
Sequence
22
11
Omission
02
01
EE
EI
IE
II
5.7 Hz (E)
3.2 Hz (I)

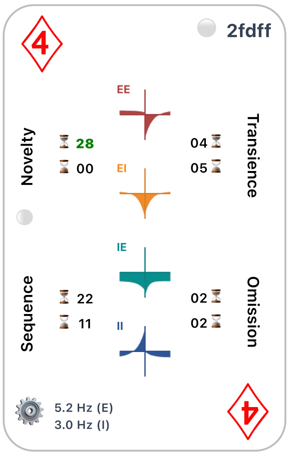
4
2fdff
Novelty
28
00
Transience
04
05
Sequence
22
11
Omission
02
02
EE
EI
IE
II
5.2 Hz (E)
3.0 Hz (I)

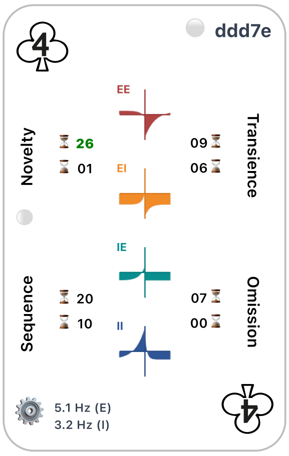
4
ddd7e
Novelty
26
01
Transience
09
06
Sequence
20
10
Omission
07
00
EE
EI
IE
II
5.1 Hz (E)
3.2 Hz (I)

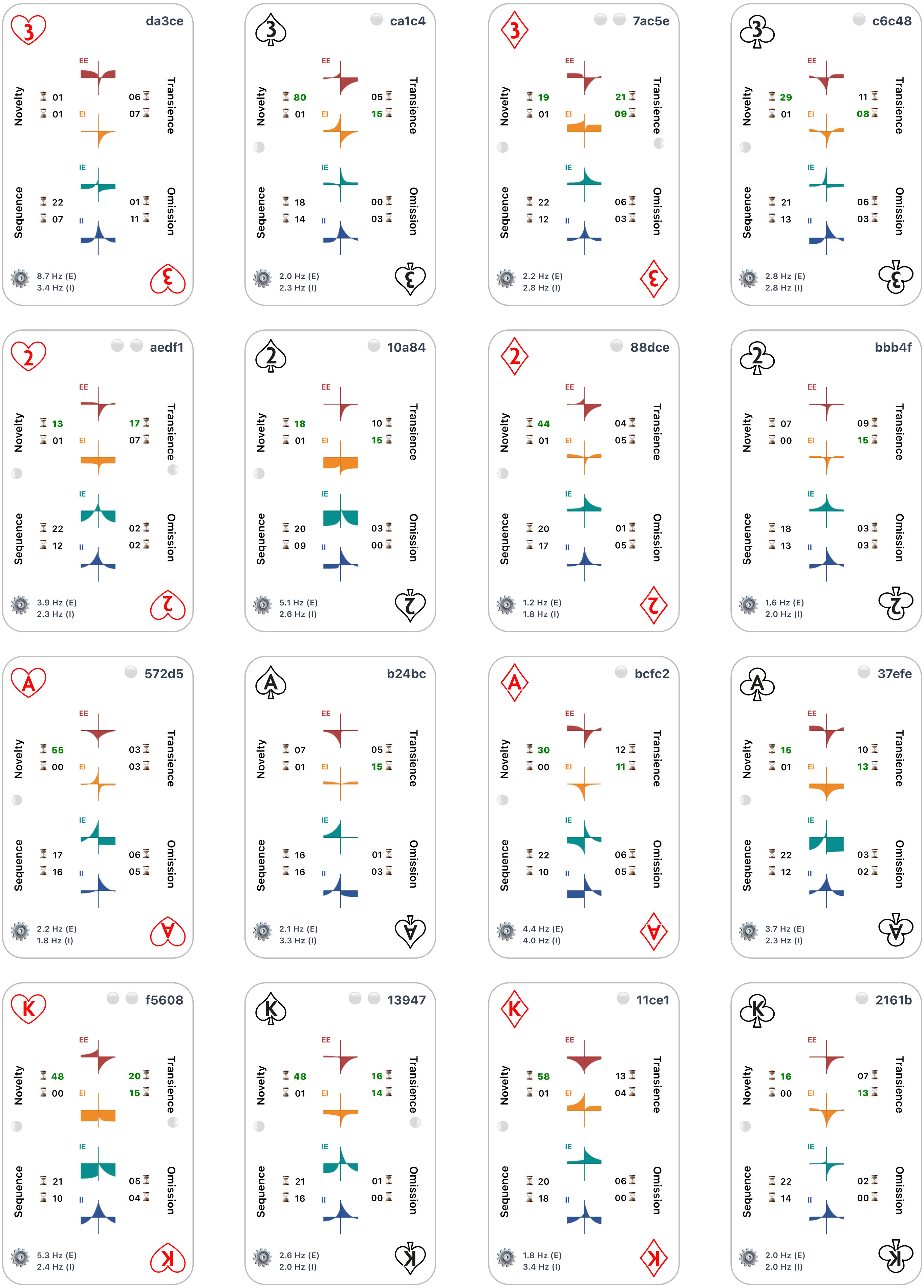
3 da3ce
Novelty 01 01
Transience 06 07
Sequence 22 07
Omission 01 11
EE EI IE II
8.7 Hz (E) 3.4 Hz (I)
3 ca1c4
Novelty 80 01
Transience 05 15
Sequence 18 14
Omission 00 03
EE EI IE II
2.0 Hz (E) 2.3 Hz (I)
3 7ac5e
Novelty 19 01
Transience 21 09
Sequence 22 12
Omission 06 03
EE EI IE II
2.2 Hz (E) 2.8 Hz (I)
3 c6c48
Novelty 29 01
Transience 11 08
Sequence 21 13
Omission 06 03
EE EI IE II
2.8 Hz (E) 2.8 Hz (I)
2 aedf1
Novelty 13 01
Transience 17 07
Sequence 22 12
Omission 02 02
EE EI IE II
3.9 Hz (E) 2.3 Hz (I)
2 10a84
Novelty 18 01
Transience 10 15
Sequence 20 09
Omission 03 00
EE EI IE II
5.1 Hz (E) 2.6 Hz (I)
2 88dce
Novelty 44 01
Transience 04 05
Sequence 20 17
Omission 01 05
EE EI IE II
1.2 Hz (E) 1.8 Hz (I)
2 bbb4f
Novelty 07 00
Transience 09 15
Sequence 18 13
Omission 03 03
EE EI IE II
1.6 Hz (E) 2.0 Hz (I)
A 572d5
Novelty 55 00
Transience 03 03
Sequence 17 16
Omission 06 05
EE EI IE II
2.2 Hz (E) 1.8 Hz (I)
A b24bc
Novelty 07 01
Transience 05 15
Sequence 16 16
Omission 01 03
EE EI IE II
2.1 Hz (E) 3.3 Hz (I)
A bcfc2
Novelty 30 00
Transience 12 11
Sequence 22 10
Omission 06 05
EE EI IE II
4.4 Hz (E) 4.0 Hz (I)
A 37efe
Novelty 15 01
Transience 10 13
Sequence 22 12
Omission 03 02
EE EI IE II
3.7 Hz (E) 2.3 Hz (I)
K f5608
Novelty 48 00
Transience 20 15
Sequence 21 10
Omission 05 04
EE EI IE II
5.3 Hz (E) 2.4 Hz (I)
K 13947
Novelty 48 01
Transience 16 14
Sequence 21 16
Omission 01 00
EE EI IE II
2.6 Hz (E) 2.0 Hz (I)
K 11ce1
Novelty 58 01
Transience 13 04
Sequence 20 18
Omission 06 00
EE EI IE II
1.8 Hz (E) 3.4 Hz (I)
K 2161b
Novelty 16 00
Transience 07 13
Sequence 22 14
Omission 02 00
EE EI IE II
2.0 Hz (E) 2.0 Hz (I)

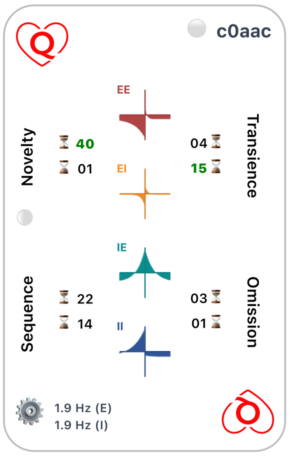
Q
c0aac
EE
Novelty
40
01
EI
04
15
Transience
IE
Sequence
22
14
II
03
01
Omission
1.9 Hz (E)
1.9 Hz (I)

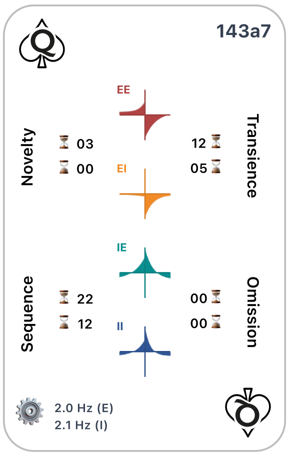
Q
143a7
EE
Novelty
03
00
EI
12
05
Transience
IE
Sequence
22
12
II
00
00
Omission
2.0 Hz (E)
2.1 Hz (I)

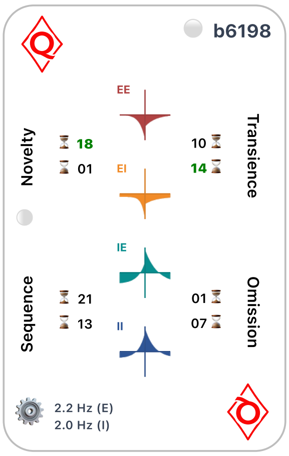
Q
b6198
EE
Novelty
18
01
EI
10
14
Transience
IE
Sequence
21
13
II
01
07
Omission
2.2 Hz (E)
2.0 Hz (I)

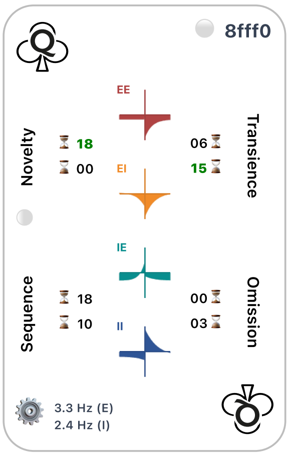
Q
8fff0
EE
Novelty
18
00
EI
06
15
Transience
IE
Sequence
18
10
II
00
03
Omission
3.3 Hz (E)
2.4 Hz (I)

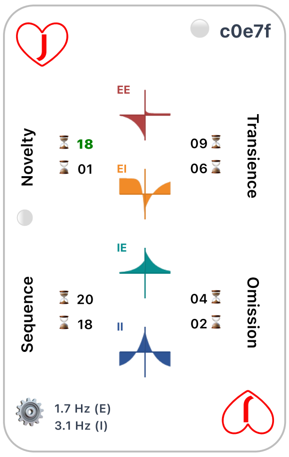
J
c0e7f
EE
Novelty
18
01
EI
09
06
Transience
IE
Sequence
20
18
II
04
02
Omission
1.7 Hz (E)
3.1 Hz (I)

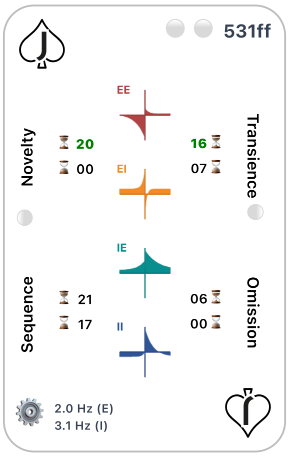
J
531ff
EE
Novelty
20
00
EI
16
07
Transience
IE
Sequence
21
17
II
06
00
Omission
2.0 Hz (E)
3.1 Hz (I)

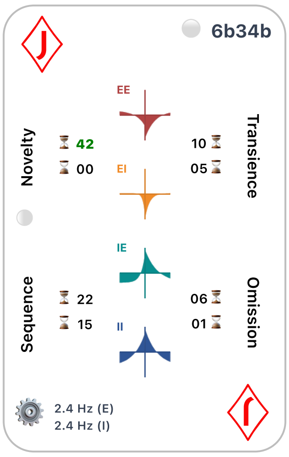
J
6b34b
EE
Novelty
42
00
EI
10
05
Transience
IE
Sequence
22
15
II
06
01
Omission
2.4 Hz (E)
2.4 Hz (I)

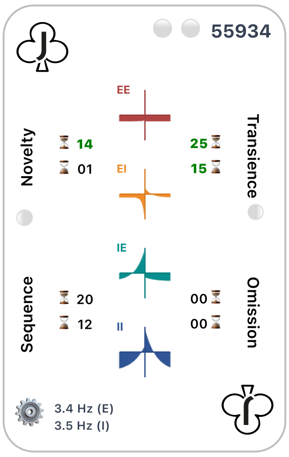
J
55934
EE
Novelty
14
01
EI
25
15
Transience
IE
Sequence
20
12
II
00
00
Omission
3.4 Hz (E)
3.5 Hz (I)

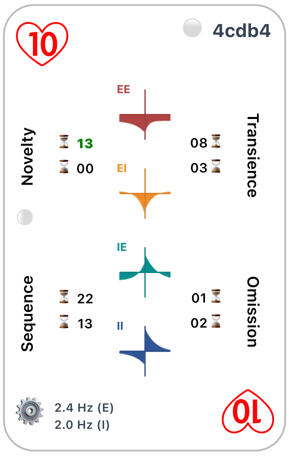
10
4cdb4
EE
Novelty
13
00
EI
08
03
Transience
IE
Sequence
22
13
II
01
02
Omission
2.4 Hz (E)
2.0 Hz (I)

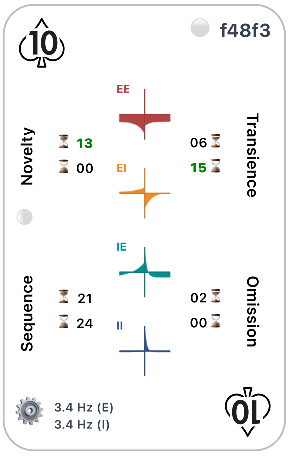
10
f48f3
EE
Novelty
13
00
EI
06
15
Transience
IE
Sequence
21
24
II
02
00
Omission
3.4 Hz (E)
3.4 Hz (I)

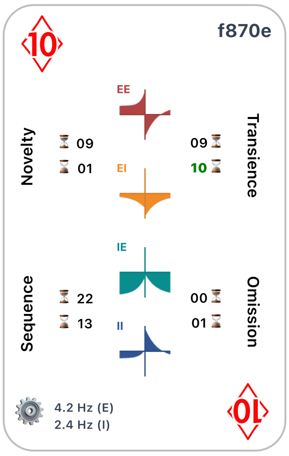
10
f870e
EE
Novelty
09
01
EI
09
10
Transience
IE
Sequence
22
13
II
00
01
Omission
4.2 Hz (E)
2.4 Hz (I)

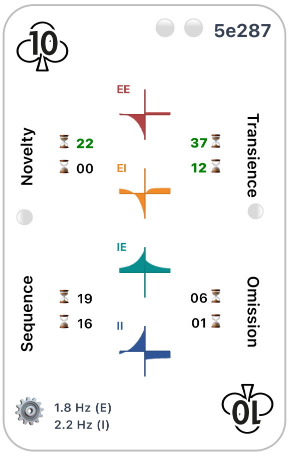
10
5e287
EE
Novelty
22
00
EI
37
12
Transience
IE
Sequence
19
16
II
06
01
Omission
1.8 Hz (E)
2.2 Hz (I)

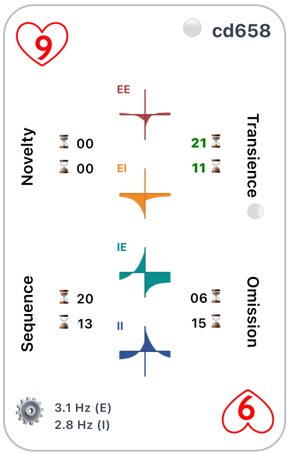
9
cd658
EE
Novelty
00
00
EI
21
11
Transience
IE
Sequence
20
13
II
06
15
Omission
3.1 Hz (E)
2.8 Hz (I)

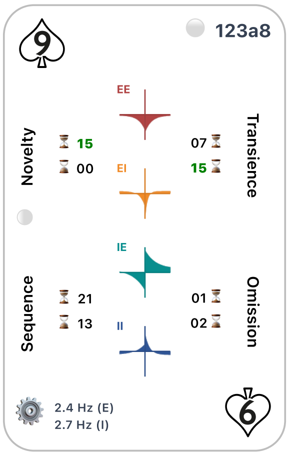
9
123a8
EE
Novelty
15
00
EI
07
15
Transience
IE
Sequence
21
13
II
01
02
Omission
2.4 Hz (E)
2.7 Hz (I)

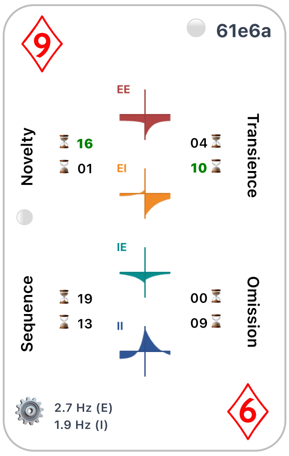
9
61e6a
EE
Novelty
16
01
EI
04
10
Transience
IE
Sequence
19
13
II
00
09
Omission
2.7 Hz (E)
1.9 Hz (I)

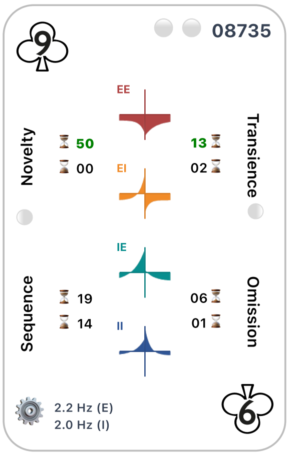
9
08735
EE
Novelty
50
00
EI
13
02
Transience
IE
Sequence
19
14
II
06
01
Omission
2.2 Hz (E)
2.0 Hz (I)

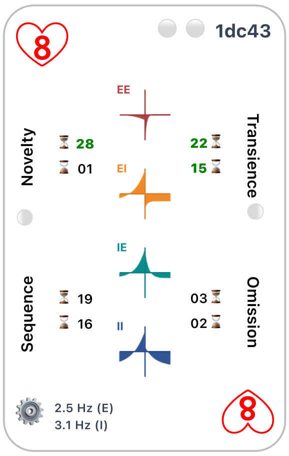
8
1dc43
EE
EI
IE
II
Novelty
28
01
Transience
22
15
Sequence
19
16
Omission
03
02
2.5 Hz (E)
3.1 Hz (I)

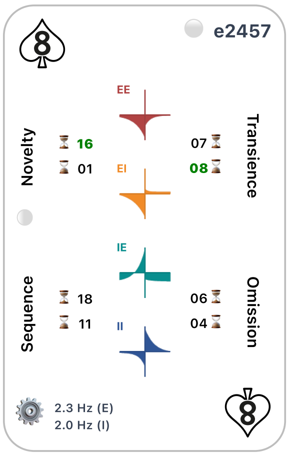
8
e2457
EE
EI
IE
II
Novelty
16
01
Transience
07
08
Sequence
18
11
Omission
06
04
2.3 Hz (E)
2.0 Hz (I)

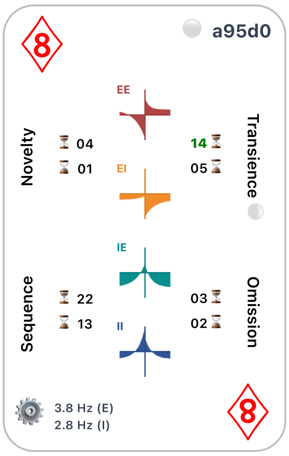
8
a95d0
EE
EI
IE
II
Novelty
04
01
Transience
14
05
Sequence
22
13
Omission
03
02
3.8 Hz (E)
2.8 Hz (I)

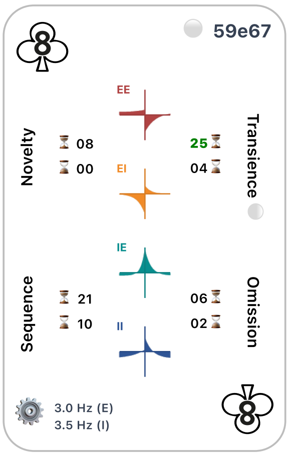
8
59e67
EE
EI
IE
II
Novelty
08
00
Transience
25
04
Sequence
21
10
Omission
06
02
3.0 Hz (E)
3.5 Hz (I)

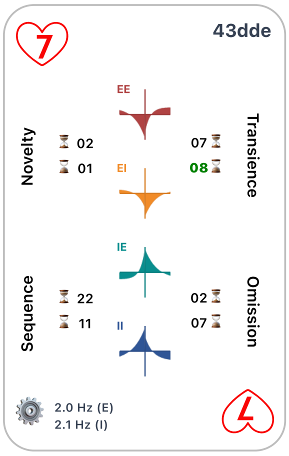
7
43dde
EE
EI
IE
II
Novelty
02
01
Transience
07
08
Sequence
22
11
Omission
02
07
2.0 Hz (E)
2.1 Hz (I)

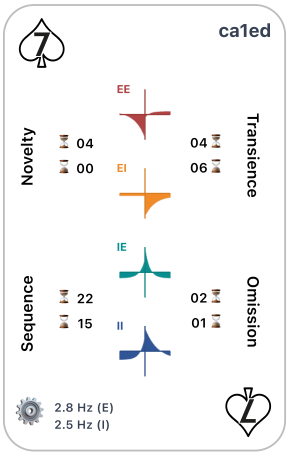
7
ca1ed
EE
EI
IE
II
Novelty
04
00
Transience
04
06
Sequence
22
15
Omission
02
01
2.8 Hz (E)
2.5 Hz (I)

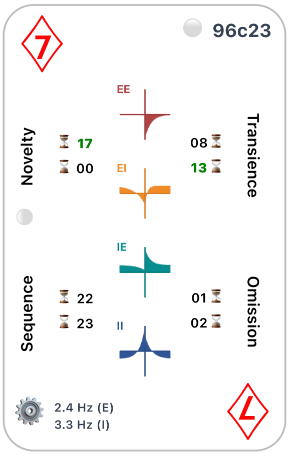
7
96c23
EE
EI
IE
II
Novelty
17
00
Transience
08
13
Sequence
22
23
Omission
01
02
2.4 Hz (E)
3.3 Hz (I)

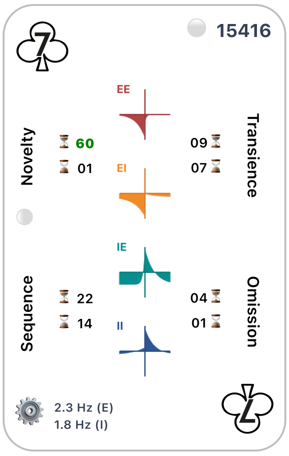
7
15416
EE
EI
IE
II
Novelty
60
01
Transience
09
07
Sequence
22
14
Omission
04
01
2.3 Hz (E)
1.8 Hz (I)

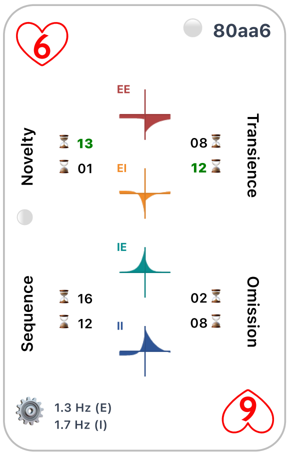
6
80aa6
EE
EI
IE
II
Novelty
13
01
Transience
08
12
Sequence
16
12
Omission
02
08
1.3 Hz (E)
1.7 Hz (I)

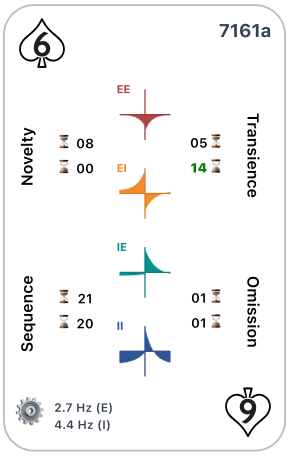
6
7161a
EE
EI
IE
II
Novelty
08
00
Transience
05
14
Sequence
21
20
Omission
01
01
2.7 Hz (E)
4.4 Hz (I)

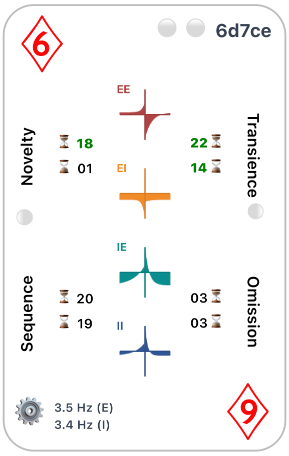
6
6d7ce
EE
EI
IE
II
Novelty
18
01
Transience
22
14
Sequence
20
19
Omission
03
03
3.5 Hz (E)
3.4 Hz (I)

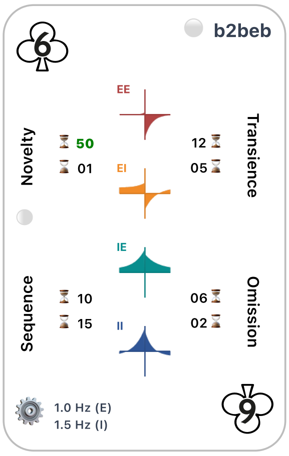
6
b2beb
EE
EI
IE
II
Novelty
50
01
Transience
12
05
Sequence
10
15
Omission
06
02
1.0 Hz (E)
1.5 Hz (I)

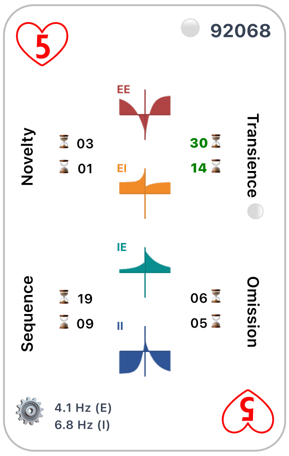
5
92068
EE
EI
IE
II
Novelty
03
01
Transience
30
14
Sequence
19
09
Omission
06
05
4.1 Hz (E)
6.8 Hz (I)

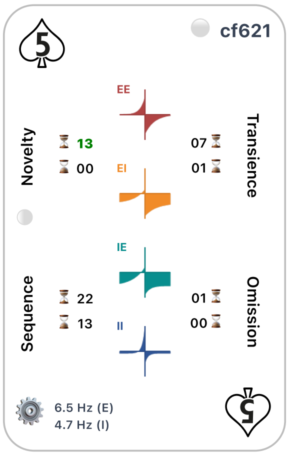
5
cf621
EE
EI
IE
II
Novelty
13
00
Transience
07
01
Sequence
22
13
Omission
01
00
6.5 Hz (E)
4.7 Hz (I)

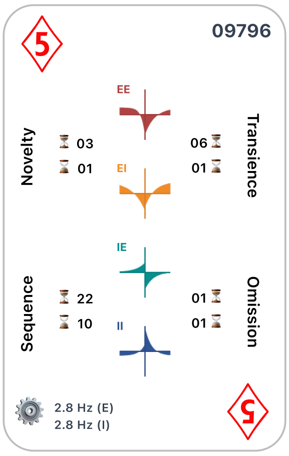
5
09796
EE
EI
IE
II
Novelty
03
01
Transience
06
01
Sequence
22
10
Omission
01
01
2.8 Hz (E)
2.8 Hz (I)

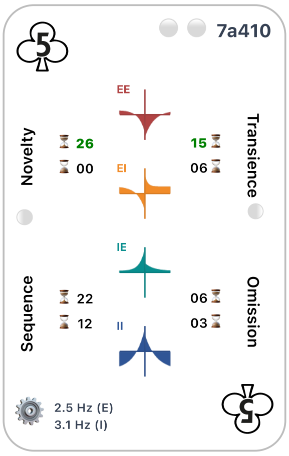
5
7a410
EE
EI
IE
II
Novelty
26
00
Transience
15
06
Sequence
22
12
Omission
06
03
2.5 Hz (E)
3.1 Hz (I)

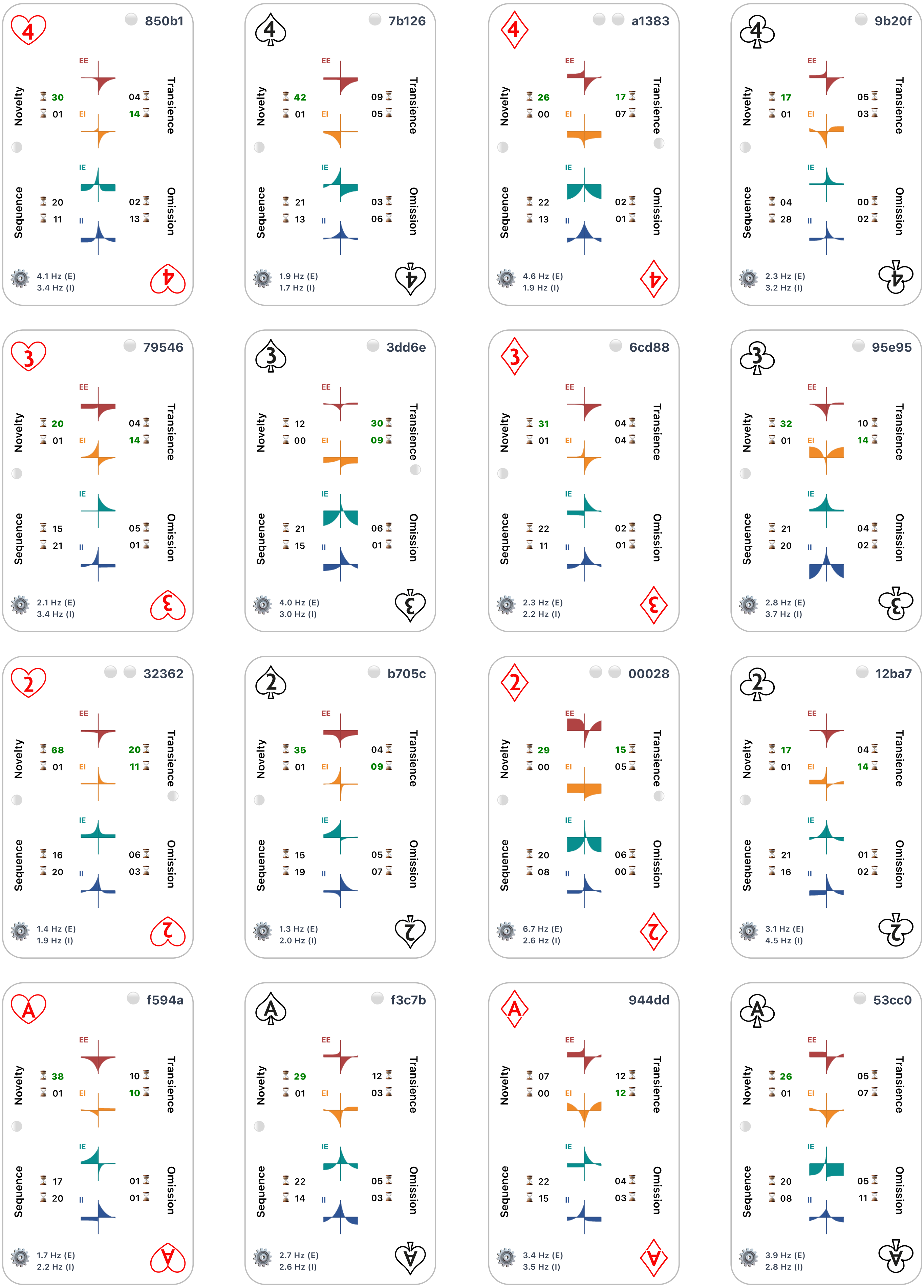
4
850b1
EE
EI
IE
II
Novelty 30 01
Transience 04 14
Sequence 20 11
Omission 02 13
4.1 Hz (E)
3.4 Hz (I)
4
7b126
EE
EI
IE
II
Novelty 42 01
Transience 09 05
Sequence 21 13
Omission 03 06
1.9 Hz (E)
1.7 Hz (I)
4
a1383
EE
EI
IE
II
Novelty 26 00
Transience 17 07
Sequence 22 13
Omission 02 01
4.6 Hz (E)
1.9 Hz (I)
4
9b20f
EE
EI
IE
II
Novelty 17 01
Transience 05 03
Sequence 04 28
Omission 00 02
2.3 Hz (E)
3.2 Hz (I)
3
79546
EE
EI
IE
II
Novelty 20 01
Transience 04 14
Sequence 15 21
Omission 05 01
2.1 Hz (E)
3.4 Hz (I)
3
3dd6e
EE
EI
IE
II
Novelty 12 00
Transience 30 09
Sequence 21 15
Omission 06 01
4.0 Hz (E)
3.0 Hz (I)
3
6cd88
EE
EI
IE
II
Novelty 31 01
Transience 04 04
Sequence 22 11
Omission 02 01
2.3 Hz (E)
2.2 Hz (I)
3
95e95
EE
EI
IE
II
Novelty 32 01
Transience 10 14
Sequence 21 20
Omission 04 02
2.8 Hz (E)
3.7 Hz (I)
2
32362
EE
EI
IE
II
Novelty 68 01
Transience 20 11
Sequence 16 20
Omission 06 03
1.4 Hz (E)
1.9 Hz (I)
2
b705c
EE
EI
IE
II
Novelty 35 01
Transience 04 09
Sequence 15 19
Omission 05 07
1.3 Hz (E)
2.0 Hz (I)
2
00028
EE
EI
IE
II
Novelty 29 00
Transience 15 05
Sequence 20 08
Omission 06 00
6.7 Hz (E)
2.6 Hz (I)
2
12ba7
EE
EI
IE
II
Novelty 17 01
Transience 04 14
Sequence 21 16
Omission 01 02
3.1 Hz (E)
4.5 Hz (I)
A
f594a
EE
EI
IE
II
Novelty 38 01
Transience 10 10
Sequence 17 20
Omission 01 01
1.7 Hz (E)
2.2 Hz (I)
A
f3c7b
EE
EI
IE
II
Novelty 29 01
Transience 12 03
Sequence 22 14
Omission 05 03
2.7 Hz (E)
2.6 Hz (I)
A
944dd
EE
EI
IE
II
Novelty 07 00
Transience 12 12
Sequence 22 15
Omission 04 03
3.4 Hz (E)
3.5 Hz (I)
A
53cc0
EE
EI
IE
II
Novelty 26 01
Transience 05 07
Sequence 20 08
Omission 05 11
3.9 Hz (E)
2.8 Hz (I)

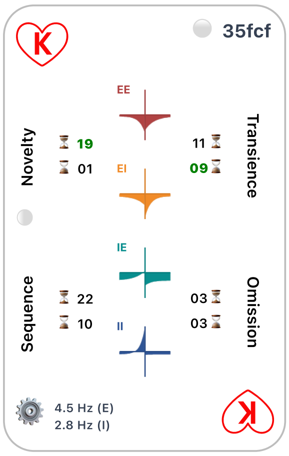
35fcf
EE
EI
IE
II
Novelty
19
01
Transience
11
09
Sequence
22
10
Omission
03
03
4.5 Hz (E)
2.8 Hz (I)

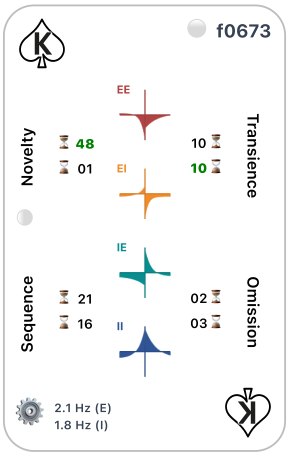
f0673
EE
EI
IE
II
Novelty
48
01
Transience
10
10
Sequence
21
16
Omission
02
03
2.1 Hz (E)
1.8 Hz (I)

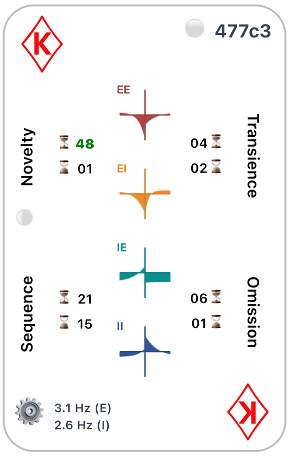
477c3
EE
EI
IE
II
Novelty
48
01
Transience
04
02
Sequence
21
15
Omission
06
01
3.1 Hz (E)
2.6 Hz (I)

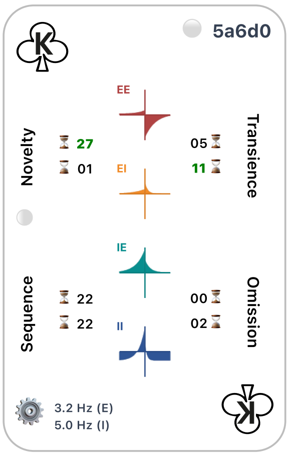
5a6d0
EE
EI
IE
II
Novelty
27
01
Transience
05
11
Sequence
22
22
Omission
00
02
3.2 Hz (E)
5.0 Hz (I)

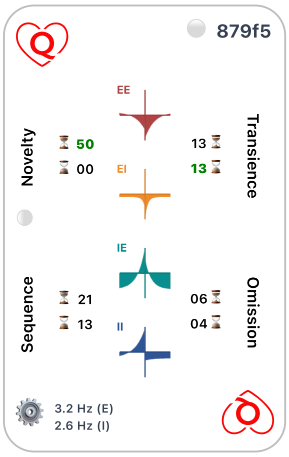
879f5
EE
EI
IE
II
Novelty
50
00
Transience
13
13
Sequence
21
13
Omission
06
04
3.2 Hz (E)
2.6 Hz (I)

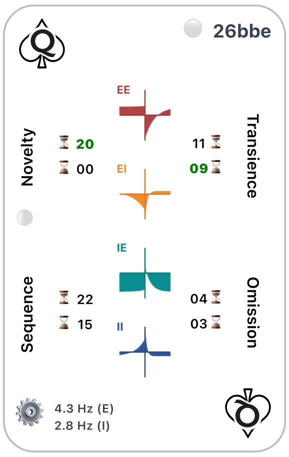
26bbe
EE
EI
IE
II
Novelty
20
00
Transience
11
09
Sequence
22
15
Omission
04
03
4.3 Hz (E)
2.8 Hz (I)

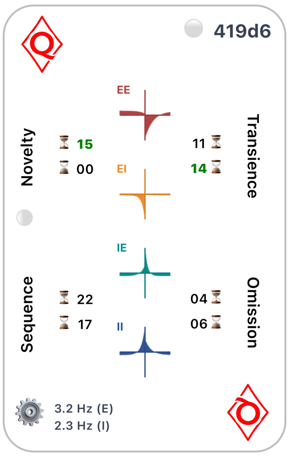
419d6
EE
EI
IE
II
Novelty
15
00
Transience
11
14
Sequence
22
17
Omission
04
06
3.2 Hz (E)
2.3 Hz (I)

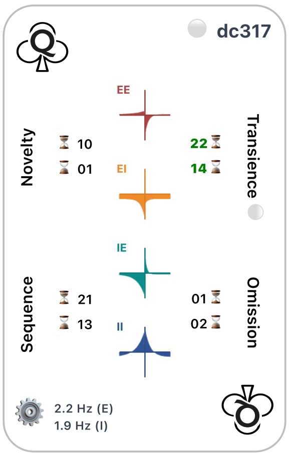
dc317
EE
EI
IE
II
Novelty
10
01
Transience
22
14
Sequence
21
13
Omission
01
02
2.2 Hz (E)
1.9 Hz (I)

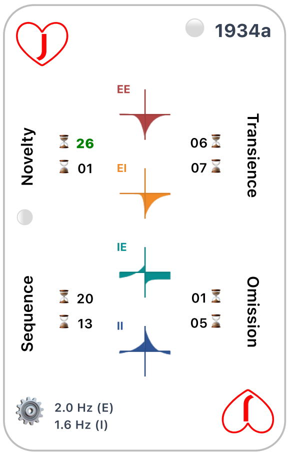
1934a
EE
EI
IE
II
Novelty
26
01
Transience
06
07
Sequence
20
13
Omission
01
05
2.0 Hz (E)
1.6 Hz (I)

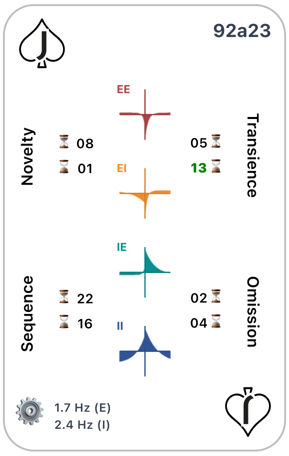
92a23
EE
EI
IE
II
Novelty
08
01
Transience
05
13
Sequence
22
16
Omission
02
04
1.7 Hz (E)
2.4 Hz (I)

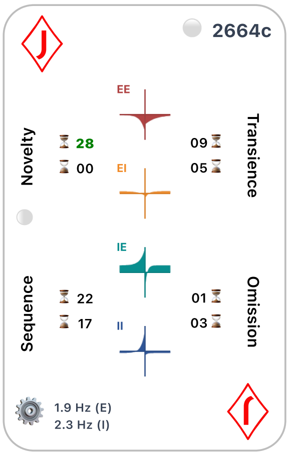
2664c
EE
EI
IE
II
Novelty
28
00
Transience
09
05
Sequence
22
17
Omission
01
03
1.9 Hz (E)
2.3 Hz (I)

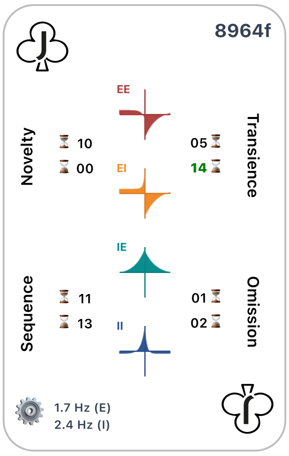
8964f
EE
EI
IE
II
Novelty
10
00
Transience
05
14
Sequence
11
13
Omission
01
02
1.7 Hz (E)
2.4 Hz (I)

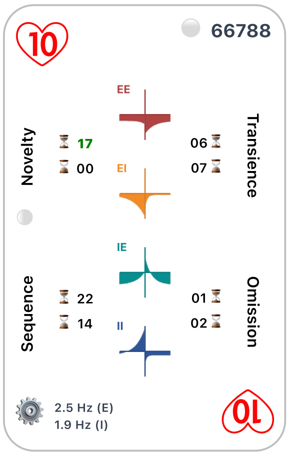
66788
EE
EI
IE
II
Novelty
17
00
Transience
06
07
Sequence
22
14
Omission
01
02
2.5 Hz (E)
1.9 Hz (I)

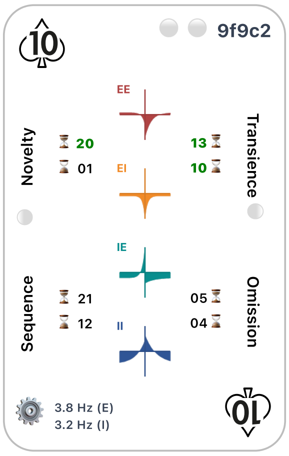
9f9c2
EE
EI
IE
II
Novelty
20
01
Transience
13
10
Sequence
21
12
Omission
05
04
3.8 Hz (E)
3.2 Hz (I)

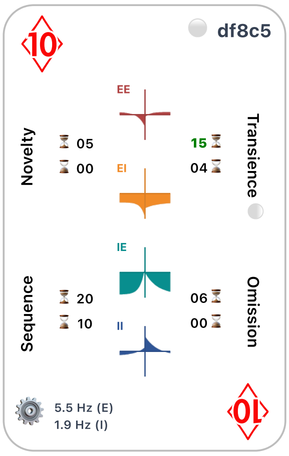
df8c5
EE
EI
IE
II
Novelty
05
00
Transience
15
04
Sequence
20
10
Omission
06
00
5.5 Hz (E)
1.9 Hz (I)

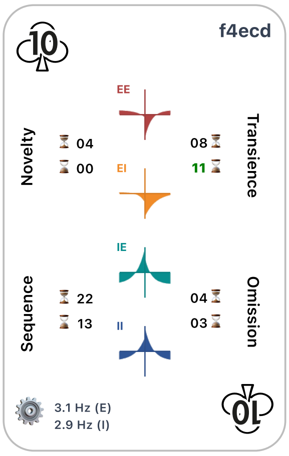
f4ecd
EE
EI
IE
II
Novelty
04
00
Transience
08
11
Sequence
22
13
Omission
04
03
3.1 Hz (E)
2.9 Hz (I)

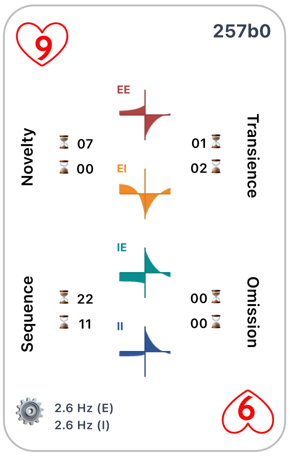
257b0
EE
EI
IE
II
Novelty 07 00
Transience 01 02
Sequence 22 11
Omission 00 00
2.6 Hz (E)
2.6 Hz (I)

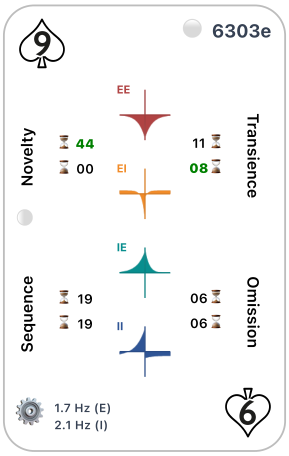
6303e
EE
EI
IE
II
Novelty 44 00
Transience 11 08
Sequence 19 19
Omission 06 06
1.7 Hz (E)
2.1 Hz (I)

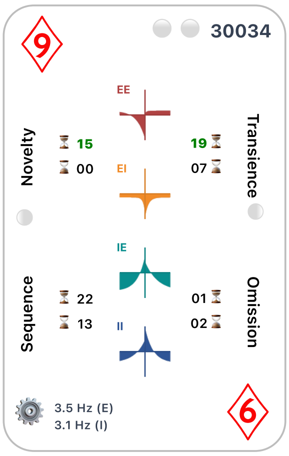
30034
EE
EI
IE
II
Novelty 15 00
Transience 19 07
Sequence 22 13
Omission 01 02
3.5 Hz (E)
3.1 Hz (I)

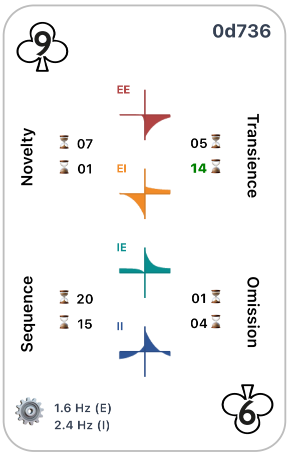
0d736
EE
EI
IE
II
Novelty 07 01
Transience 05 14
Sequence 20 15
Omission 01 04
1.6 Hz (E)
2.4 Hz (I)

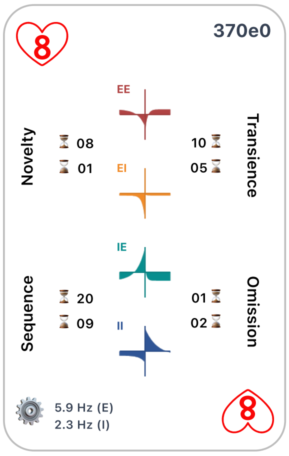
370e0
EE
EI
IE
II
Novelty 08 01
Transience 10 05
Sequence 20 09
Omission 01 02
5.9 Hz (E)
2.3 Hz (I)

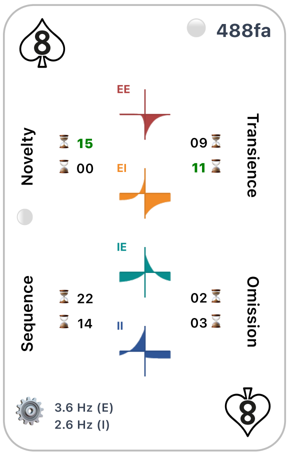
488fa
EE
EI
IE
II
Novelty 15 00
Transience 09 11
Sequence 22 14
Omission 02 03
3.6 Hz (E)
2.6 Hz (I)

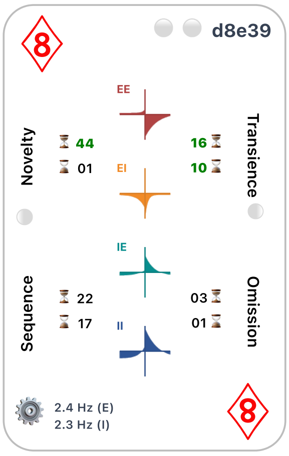
d8e39
EE
EI
IE
II
Novelty 44 01
Transience 16 10
Sequence 22 17
Omission 03 01
2.4 Hz (E)
2.3 Hz (I)

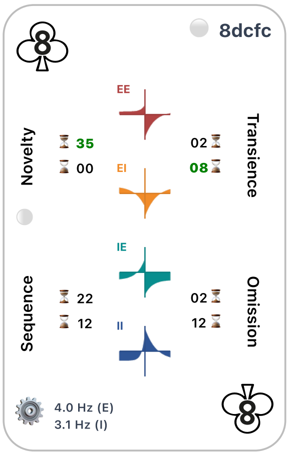
8dcfc
EE
EI
IE
II
Novelty 35 00
Transience 02 08
Sequence 22 12
Omission 02 12
4.0 Hz (E)
3.1 Hz (I)

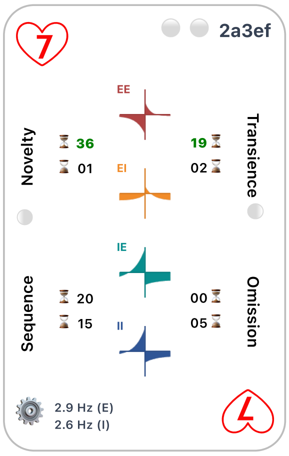
2a3ef
EE
EI
IE
II
Novelty 36 01
Transience 19 02
Sequence 20 15
Omission 00 05
2.9 Hz (E)
2.6 Hz (I)

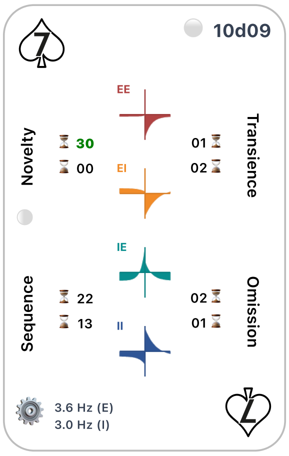
10d09
EE
EI
IE
II
Novelty 30 00
Transience 01 02
Sequence 22 13
Omission 02 01
3.6 Hz (E)
3.0 Hz (I)

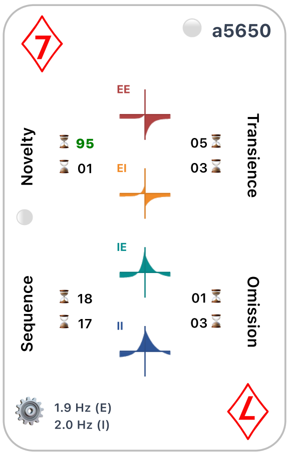
a5650
EE
EI
IE
II
Novelty 95 01
Transience 05 03
Sequence 18 17
Omission 01 03
1.9 Hz (E)
2.0 Hz (I)

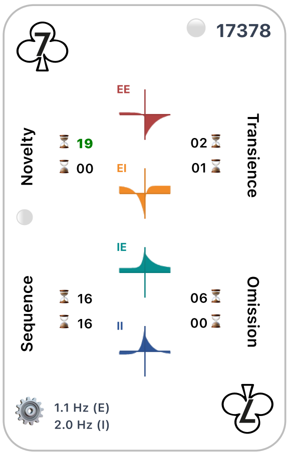
17378
EE
EI
IE
II
Novelty 19 00
Transience 02 01
Sequence 16 16
Omission 06 00
1.1 Hz (E)
2.0 Hz (I)

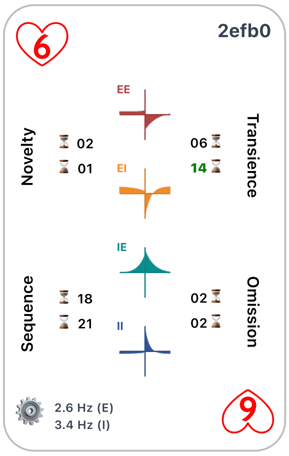
2efb0
EE
EI
IE
II
Novelty 02 01
Transience 06 14
Sequence 18 21
Omission 02 02
2.6 Hz (E)
3.4 Hz (I)

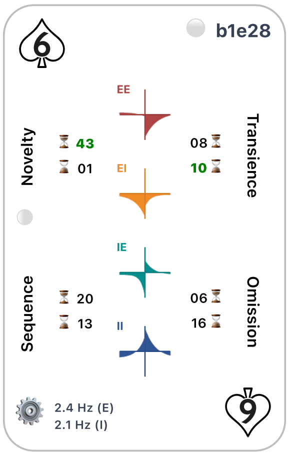
b1e28
EE
EI
IE
II
Novelty 43 01
Transience 08 10
Sequence 20 13
Omission 06 16
2.4 Hz (E)
2.1 Hz (I)

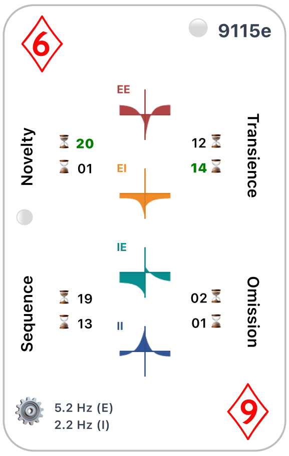
9115e
EE
EI
IE
II
Novelty 20 01
Transience 12 14
Sequence 19 13
Omission 02 01
5.2 Hz (E)
2.2 Hz (I)

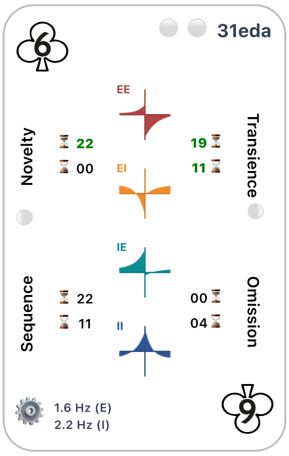
31eda
EE
EI
IE
II
Novelty 22 00
Transience 19 11
Sequence 22 11
Omission 00 04
1.6 Hz (E)
2.2 Hz (I)

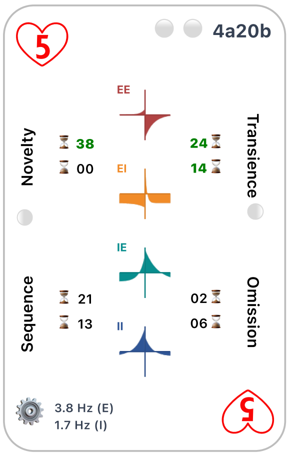
5
4a20b
EE
EI
IE
II
Novelty
38
00
Transience
24
14
Sequence
21
13
Omission
02
06
3.8 Hz (E)
1.7 Hz (I)

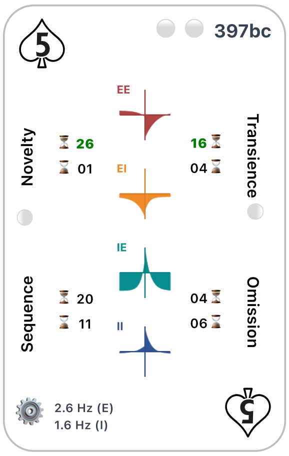
5
397bc
EE
EI
IE
II
Novelty
26
01
Transience
16
04
Sequence
20
11
Omission
04
06
2.6 Hz (E)
1.6 Hz (I)

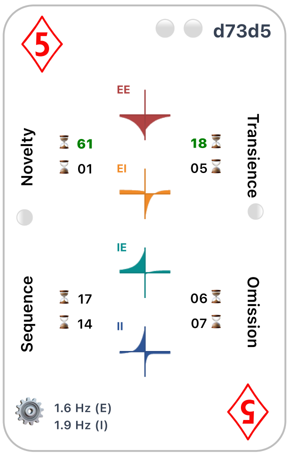
5
d73d5
EE
EI
IE
II
Novelty
61
01
Transience
18
05
Sequence
17
14
Omission
06
07
1.6 Hz (E)
1.9 Hz (I)

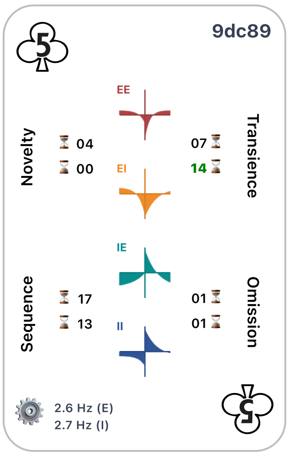
5
9dc89
EE
EI
IE
II
Novelty
04
00
Transience
07
14
Sequence
17
13
Omission
01
01
2.6 Hz (E)
2.7 Hz (I)

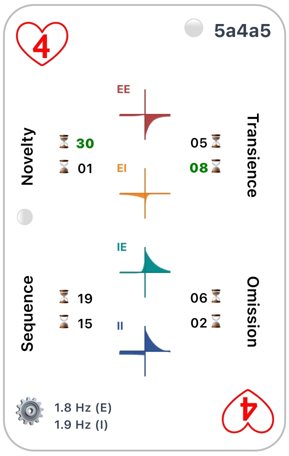
4
5a4a5
EE
EI
IE
II
Novelty
30
01
Transience
05
08
Sequence
19
15
Omission
06
02
1.8 Hz (E)
1.9 Hz (I)

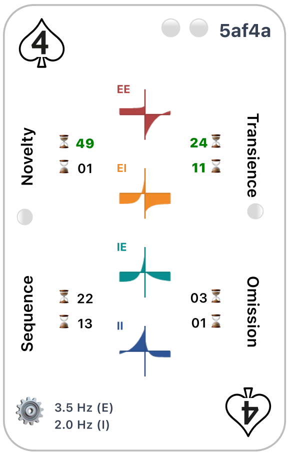
4
5af4a
EE
EI
IE
II
Novelty
49
01
Transience
24
11
Sequence
22
13
Omission
03
01
3.5 Hz (E)
2.0 Hz (I)

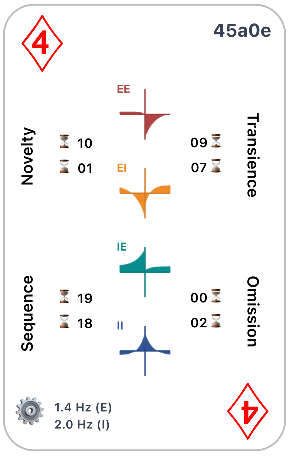
4
45a0e
EE
EI
IE
II
Novelty
10
01
Transience
09
07
Sequence
19
18
Omission
00
02
1.4 Hz (E)
2.0 Hz (I)

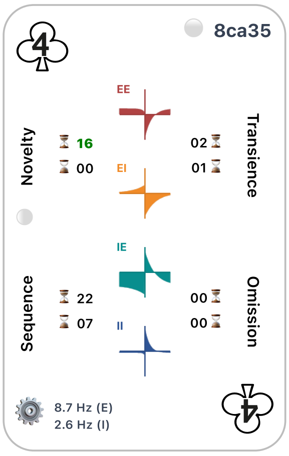
4
8ca35
EE
EI
IE
II
Novelty
16
00
Transience
02
01
Sequence
22
07
Omission
00
00
8.7 Hz (E)
2.6 Hz (I)

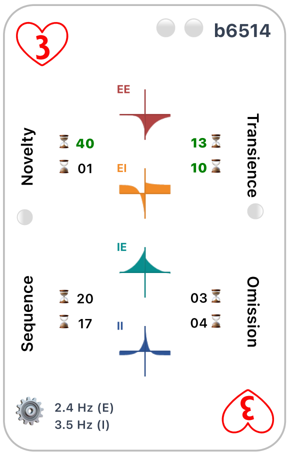
3
b6514
EE
EI
IE
II
Novelty
40
01
Transience
13
10
Sequence
20
17
Omission
03
04
2.4 Hz (E)
3.5 Hz (I)

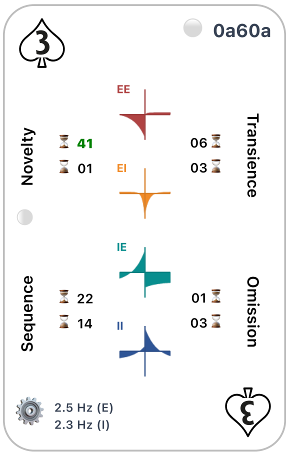
3
0a60a
EE
EI
IE
II
Novelty
41
01
Transience
06
03
Sequence
22
14
Omission
01
03
2.5 Hz (E)
2.3 Hz (I)

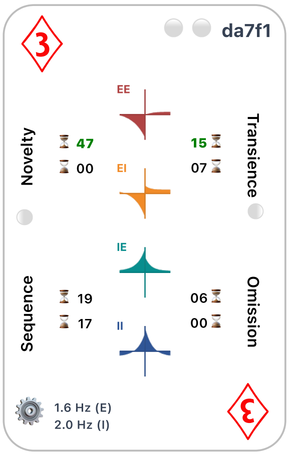
3
da7f1
EE
EI
IE
II
Novelty
47
00
Transience
15
07
Sequence
19
17
Omission
06
00
1.6 Hz (E)
2.0 Hz (I)

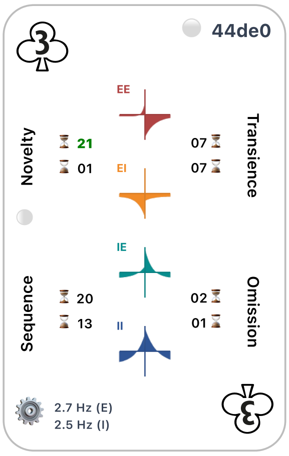
3
44de0
EE
EI
IE
II
Novelty
21
01
Transience
07
07
Sequence
20
13
Omission
02
01
2.7 Hz (E)
2.5 Hz (I)

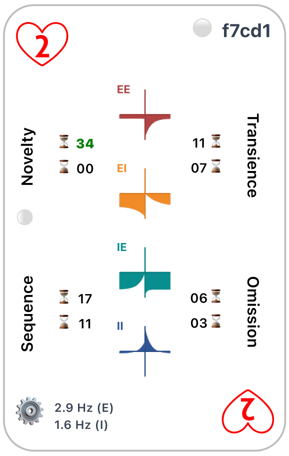
2
f7cd1
EE
EI
IE
II
Novelty
34
00
Transience
11
07
Sequence
17
11
Omission
06
03
2.9 Hz (E)
1.6 Hz (I)

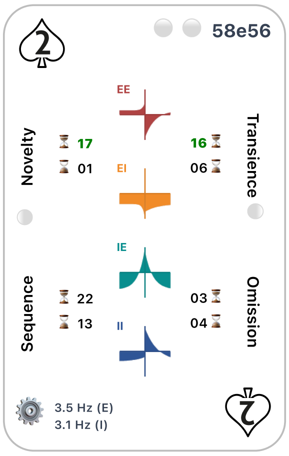
2
58e56
EE
EI
IE
II
Novelty
17
01
Transience
16
06
Sequence
22
13
Omission
03
04
3.5 Hz (E)
3.1 Hz (I)

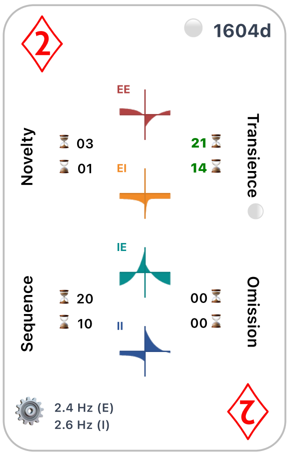
2
1604d
EE
EI
IE
II
Novelty
03
01
Transience
21
14
Sequence
20
10
Omission
00
00
2.4 Hz (E)
2.6 Hz (I)

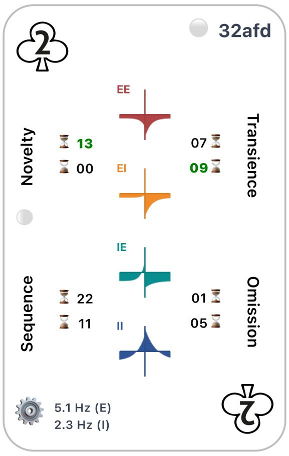
2
32afd
EE
EI
IE
II
Novelty
13
00
Transience
07
09
Sequence
22
11
Omission
01
05
5.1 Hz (E)
2.3 Hz (I)

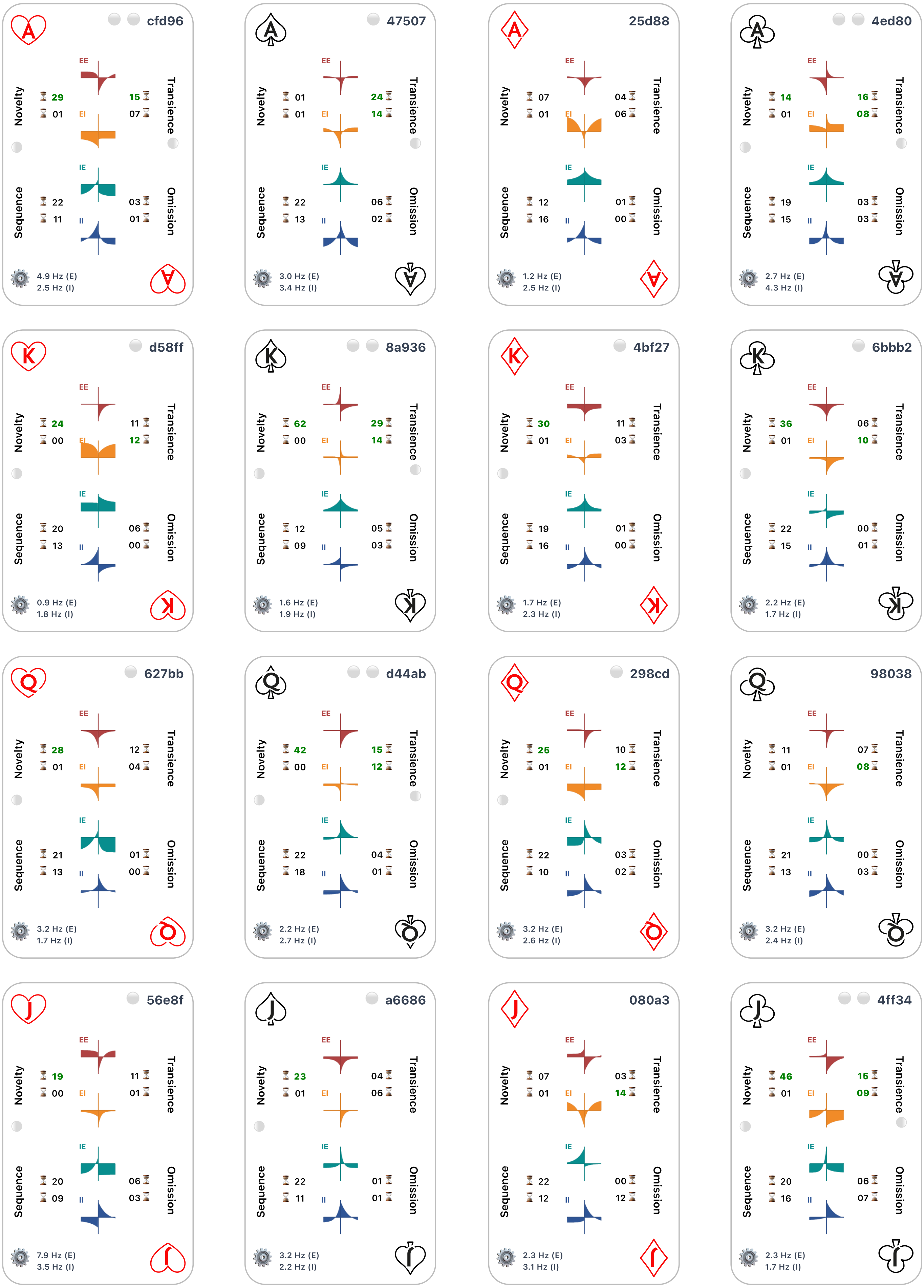
A cfd96
Novelty 29 01
Transience 15 07
Sequence 22 11
Omission 03 01
EE EI IE II
4.9 Hz (E) 2.5 Hz (I)
A 47507
Novelty 01 01
Transience 24 14
Sequence 22 13
Omission 06 02
EE EI IE II
3.0 Hz (E) 3.4 Hz (I)
A 25d88
Novelty 07 01
Transience 04 06
Sequence 12 16
Omission 01 00
EE EI IE II
1.2 Hz (E) 2.5 Hz (I)
A 4ed80
Novelty 14 01
Transience 16 08
Sequence 19 15
Omission 03 03
EE EI IE II
2.7 Hz (E) 4.3 Hz (I)
K d58ff
Novelty 24 00
Transience 11 12
Sequence 20 13
Omission 06 00
EE EI IE II
0.9 Hz (E) 1.8 Hz (I)
K 8a936
Novelty 62 00
Transience 29 14
Sequence 12 09
Omission 05 03
EE EI IE II
1.6 Hz (E) 1.9 Hz (I)
K 4bf27
Novelty 30 01
Transience 11 03
Sequence 19 16
Omission 01 00
EE EI IE II
1.7 Hz (E) 2.3 Hz (I)
K 6bbb2
Novelty 36 01
Transience 06 10
Sequence 22 15
Omission 00 01
EE EI IE II
2.2 Hz (E) 1.7 Hz (I)
Q 627bb
Novelty 28 01
Transience 12 04
Sequence 21 13
Omission 01 00
EE EI IE II
3.2 Hz (E) 1.7 Hz (I)
Q d44ab
Novelty 42 00
Transience 15 12
Sequence 22 18
Omission 04 01
EE EI IE II
2.2 Hz (E) 2.7 Hz (I)
Q 298cd
Novelty 25 01
Transience 10 12
Sequence 22 10
Omission 03 02
EE EI IE II
3.2 Hz (E) 2.6 Hz (I)
Q 98038
Novelty 11 01
Transience 07 08
Sequence 21 13
Omission 00 03
EE EI IE II
3.2 Hz (E) 2.4 Hz (I)
J 56e8f
Novelty 19 00
Transience 11 01
Sequence 20 09
Omission 06 03
EE EI IE II
7.9 Hz (E) 3.5 Hz (I)
J a6686
Novelty 23 01
Transience 04 06
Sequence 22 11
Omission 01 01
EE EI IE II
3.2 Hz (E) 2.2 Hz (I)
J 080a3
Novelty 07 01
Transience 03 14
Sequence 22 12
Omission 00 12
EE EI IE II
2.3 Hz (E) 3.1 Hz (I)
J 4ff34
Novelty 46 01
Transience 15 09
Sequence 20 16
Omission 06 07
EE EI IE II
2.3 Hz (E) 1.7 Hz (I)

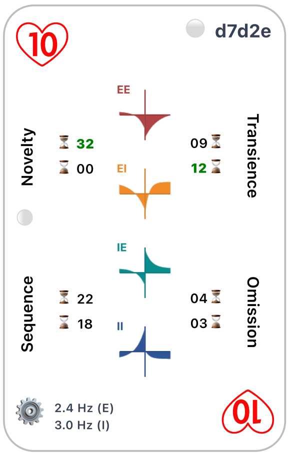
10
d7d2e
EE
Novelty
32
00
09
12
Transience
EI
IE
Sequence
22
18
04
03
Omission
II
2.4 Hz (E)
3.0 Hz (I)
10

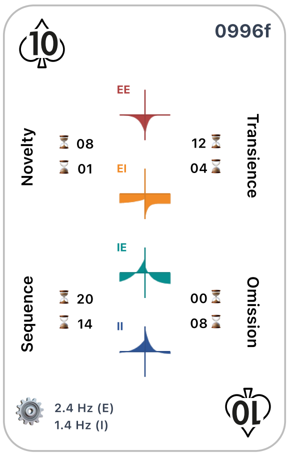
10
0996f
EE
Novelty
08
01
12
04
Transience
EI
IE
Sequence
20
14
00
08
Omission
II
2.4 Hz (E)
1.4 Hz (I)
10

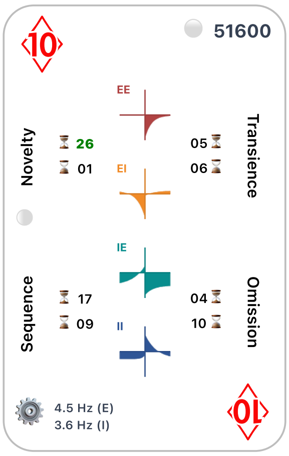
10
51600
EE
Novelty
26
01
05
06
Transience
EI
IE
Sequence
17
09
04
10
Omission
II
4.5 Hz (E)
3.6 Hz (I)
10

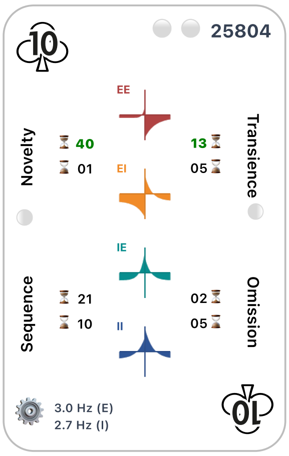
10
25804
EE
Novelty
40
01
13
05
Transience
EI
IE
Sequence
21
10
02
05
Omission
II
3.0 Hz (E)
2.7 Hz (I)
10

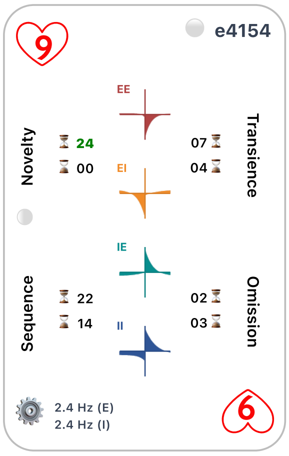
9
e4154
EE
Novelty
24
00
07
04
Transience
EI
IE
Sequence
22
14
02
03
Omission
II
2.4 Hz (E)
2.4 Hz (I)
9

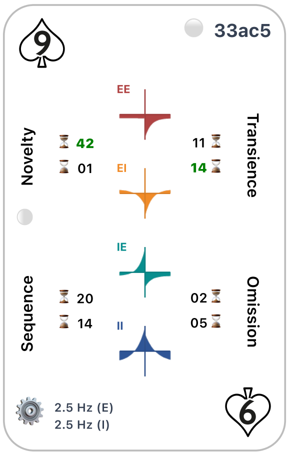
9
33ac5
EE
Novelty
42
01
11
14
Transience
EI
IE
Sequence
20
14
02
05
Omission
II
2.5 Hz (E)
2.5 Hz (I)
9

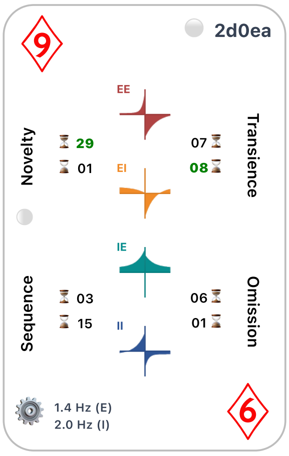
9
2d0ea
EE
Novelty
29
01
07
08
Transience
EI
IE
Sequence
03
15
06
01
Omission
II
1.4 Hz (E)
2.0 Hz (I)
9

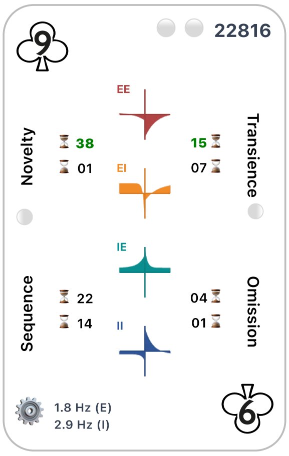
9
22816
EE
Novelty
38
01
15
07
Transience
EI
IE
Sequence
22
14
04
01
Omission
II
1.8 Hz (E)
2.9 Hz (I)
9

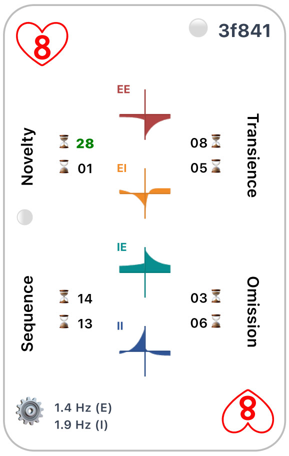
8
3f841
EE
Novelty
28
01
08
05
Transience
EI
IE
Sequence
14
13
03
06
Omission
II
1.4 Hz (E)
1.9 Hz (I)
8

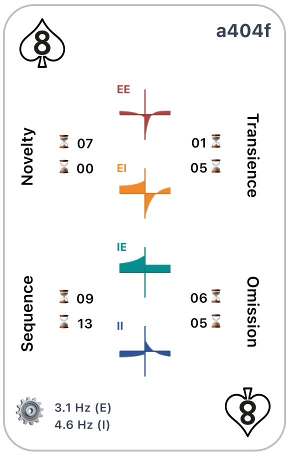
8
a404f
EE
Novelty
07
00
01
05
Transience
EI
IE
Sequence
09
13
06
05
Omission
II
3.1 Hz (E)
4.6 Hz (I)
8

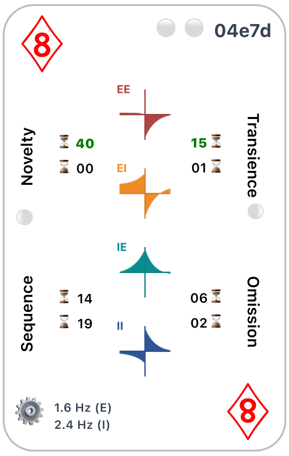
8
04e7d
EE
Novelty
40
00
15
01
Transience
EI
IE
Sequence
14
19
06
02
Omission
II
1.6 Hz (E)
2.4 Hz (I)
8

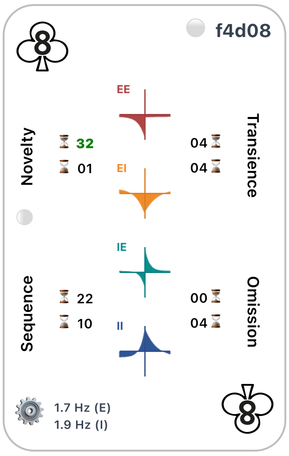
8
f4d08
EE
Novelty
32
01
04
04
Transience
EI
IE
Sequence
22
10
00
04
Omission
II
1.7 Hz (E)
1.9 Hz (I)
8

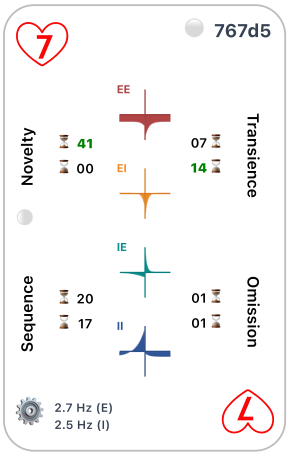
7
767d5
EE
Novelty
41
00
07
14
Transience
EI
IE
Sequence
20
17
01
01
Omission
II
2.7 Hz (E)
2.5 Hz (I)
7

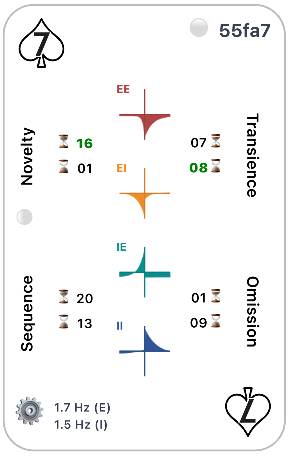
7
55fa7
EE
Novelty
16
01
07
08
Transience
EI
IE
Sequence
20
13
01
09
Omission
II
1.7 Hz (E)
1.5 Hz (I)
7

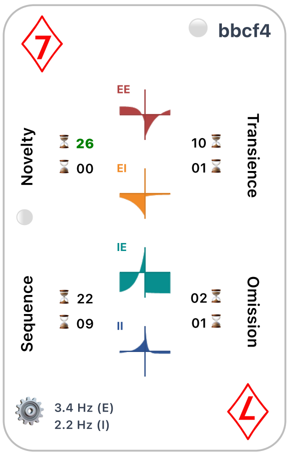
7
bbcf4
EE
Novelty
26
00
10
01
Transience
EI
IE
Sequence
22
09
02
01
Omission
II
3.4 Hz (E)
2.2 Hz (I)
7

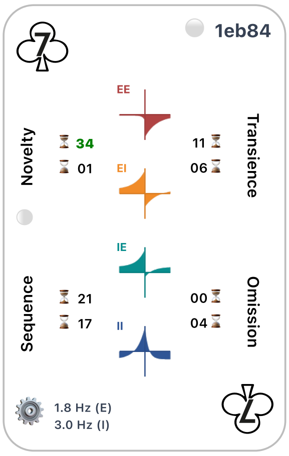
7
1eb84
EE
Novelty
34
01
11
06
Transience
EI
IE
Sequence
21
17
00
04
Omission
II
1.8 Hz (E)
3.0 Hz (I)
7

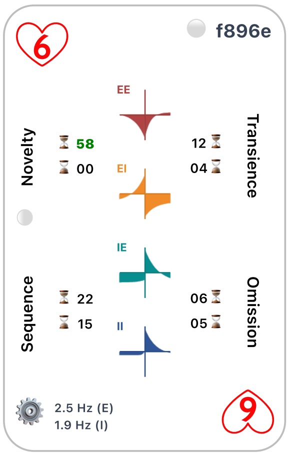
6
f896e
EE
EI
IE
II
Novelty
58
00
Transience
12
04
Sequence
22
15
Omission
06
05
2.5 Hz (E)
1.9 Hz (I)
9

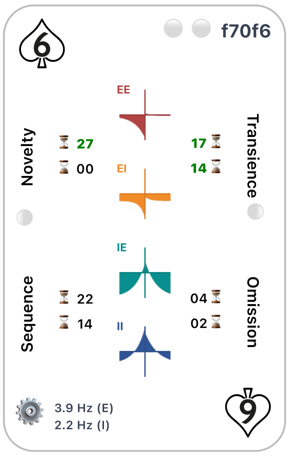
6
f70f6
EE
EI
IE
II
Novelty
27
00
Transience
17
14
Sequence
22
14
Omission
04
02
3.9 Hz (E)
2.2 Hz (I)
9

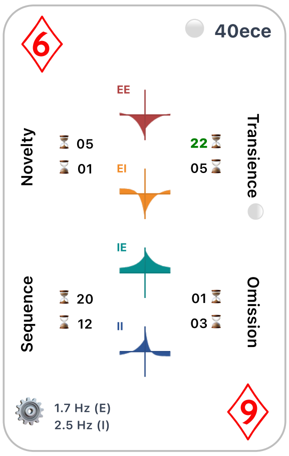
6
40ece
EE
EI
IE
II
Novelty
05
01
Transience
22
05
Sequence
20
12
Omission
01
03
1.7 Hz (E)
2.5 Hz (I)
9

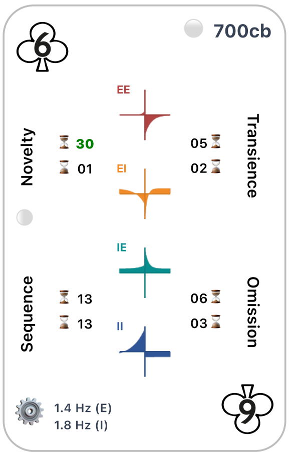
6
700cb
EE
EI
IE
II
Novelty
30
01
Transience
05
02
Sequence
13
13
Omission
06
03
1.4 Hz (E)
1.8 Hz (I)
9

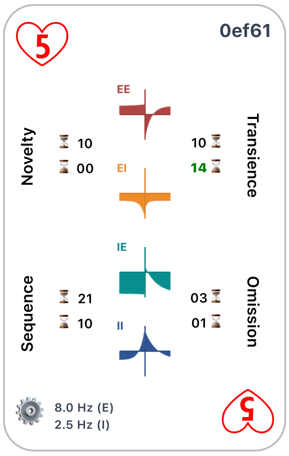
5
0ef61
EE
EI
IE
II
Novelty
10
00
Transience
10
14
Sequence
21
10
Omission
03
01
8.0 Hz (E)
2.5 Hz (I)
5

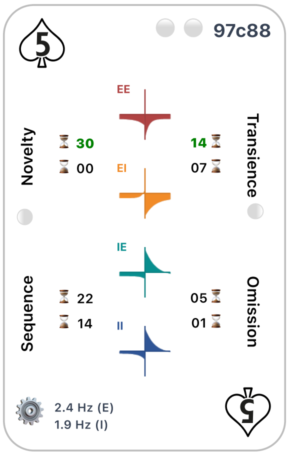
5
97c88
EE
EI
IE
II
Novelty
30
00
Transience
14
07
Sequence
22
14
Omission
05
01
2.4 Hz (E)
1.9 Hz (I)
5

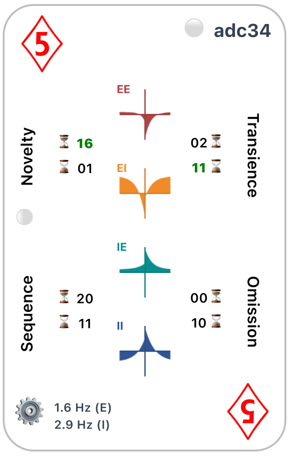
5
adc34
EE
EI
IE
II
Novelty
16
01
Transience
02
11
Sequence
20
11
Omission
00
10
1.6 Hz (E)
2.9 Hz (I)
5

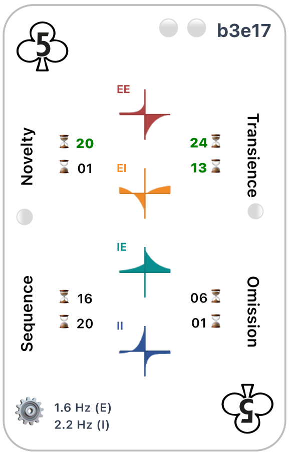
5
b3e17
EE
EI
IE
II
Novelty
20
01
Transience
24
13
Sequence
16
20
Omission
06
01
1.6 Hz (E)
2.2 Hz (I)
5

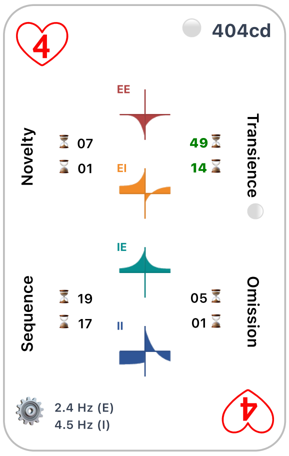
4
404cd
EE
EI
IE
II
Novelty
07
01
Transience
49
14
Sequence
19
17
Omission
05
01
2.4 Hz (E)
4.5 Hz (I)
4

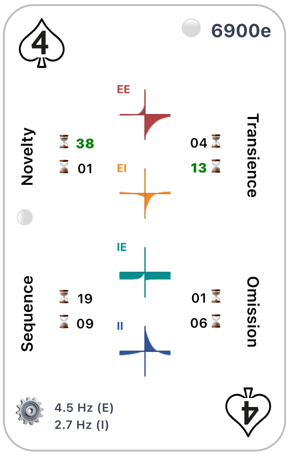
4
6900e
EE
EI
IE
II
Novelty
38
01
Transience
04
13
Sequence
19
09
Omission
01
06
4.5 Hz (E)
2.7 Hz (I)
4

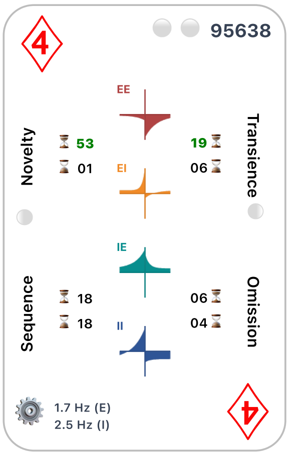
4
95638
EE
EI
IE
II
Novelty
53
01
Transience
19
06
Sequence
18
18
Omission
06
04
1.7 Hz (E)
2.5 Hz (I)
4

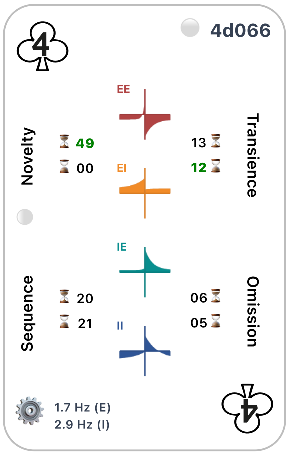
4
4d066
EE
EI
IE
II
Novelty
49
00
Transience
13
12
Sequence
20
21
Omission
06
05
1.7 Hz (E)
2.9 Hz (I)
4

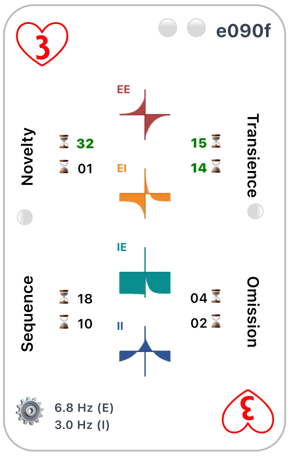
3
e090f
EE
EI
IE
II
Novelty
32
01
Transience
15
14
Sequence
18
10
Omission
04
02
6.8 Hz (E)
3.0 Hz (I)
3

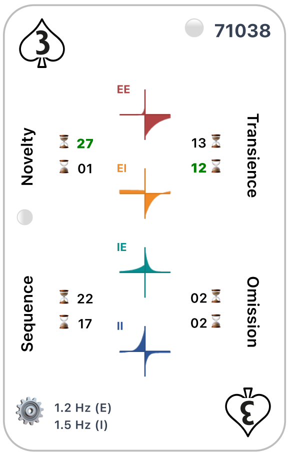
3
71038
EE
EI
IE
II
Novelty
27
01
Transience
13
12
Sequence
22
17
Omission
02
02
1.2 Hz (E)
1.5 Hz (I)
3

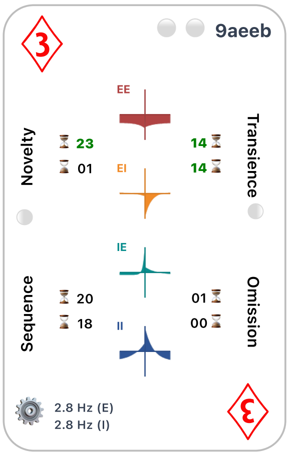
3
9aeeb
EE
EI
IE
II
Novelty
23
01
Transience
14
14
Sequence
20
18
Omission
01
00
2.8 Hz (E)
2.8 Hz (I)
3

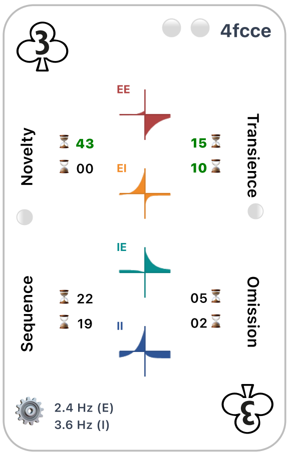
3
4fcce
EE
EI
IE
II
Novelty
43
00
Transience
15
10
Sequence
22
19
Omission
05
02
2.4 Hz (E)
3.6 Hz (I)
3

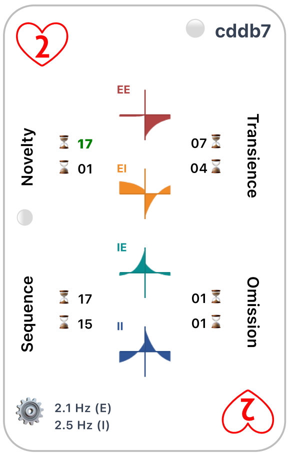
cddb7
EE
EI
IE
II
Novelty 17 01
Transience 07 04
Sequence 17 15
Omission 01 01
2.1 Hz (E)
2.5 Hz (I)

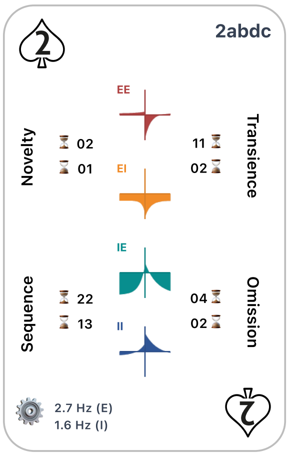
2abdc
EE
EI
IE
II
Novelty 02 01
Transience 11 02
Sequence 22 13
Omission 04 02
2.7 Hz (E)
1.6 Hz (I)

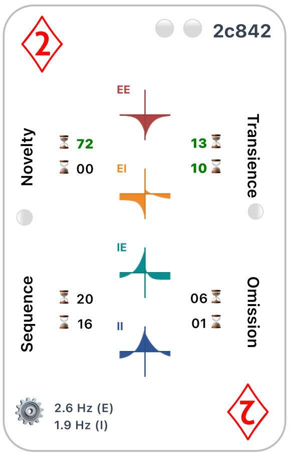
2c842
EE
EI
IE
II
Novelty 72 00
Transience 13 10
Sequence 20 16
Omission 06 01
2.6 Hz (E)
1.9 Hz (I)

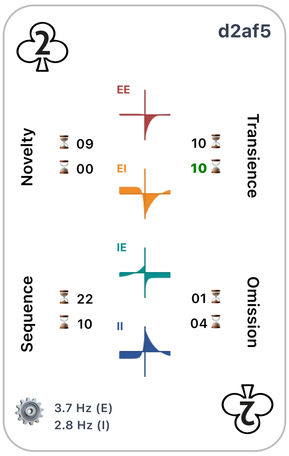
d2af5
EE
EI
IE
II
Novelty 09 00
Transience 10 10
Sequence 22 10
Omission 01 04
3.7 Hz (E)
2.8 Hz (I)

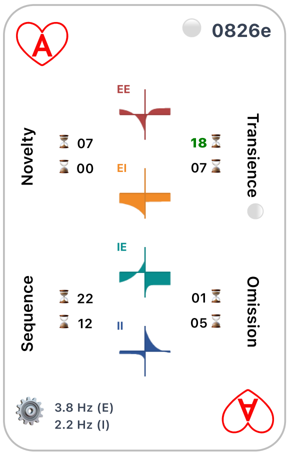
0826e
EE
EI
IE
II
Novelty 07 00
Transience 18 07
Sequence 22 12
Omission 01 05
3.8 Hz (E)
2.2 Hz (I)

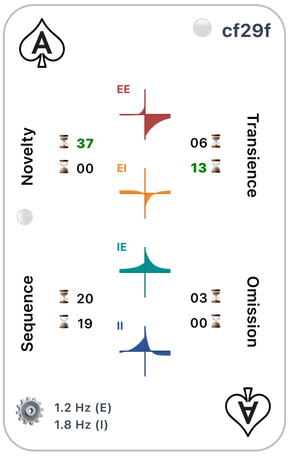
cf29f
EE
EI
IE
II
Novelty 37 00
Transience 06 13
Sequence 20 19
Omission 03 00
1.2 Hz (E)
1.8 Hz (I)

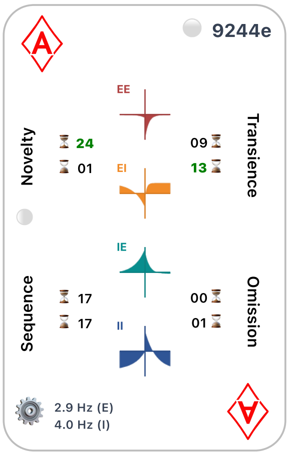
9244e
EE
EI
IE
II
Novelty 24 01
Transience 09 13
Sequence 17 17
Omission 00 01
2.9 Hz (E)
4.0 Hz (I)

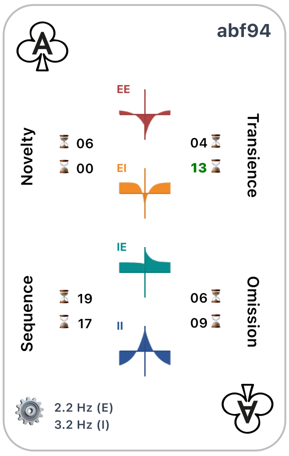
abf94
EE
EI
IE
II
Novelty 06 00
Transience 04 13
Sequence 19 17
Omission 06 09
2.2 Hz (E)
3.2 Hz (I)

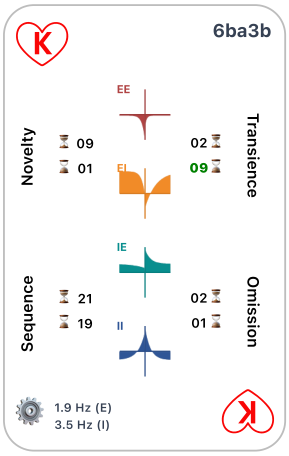
6ba3b
EE
EI
IE
II
Novelty 09 01
Transience 02 09
Sequence 21 19
Omission 02 01
1.9 Hz (E)
3.5 Hz (I)

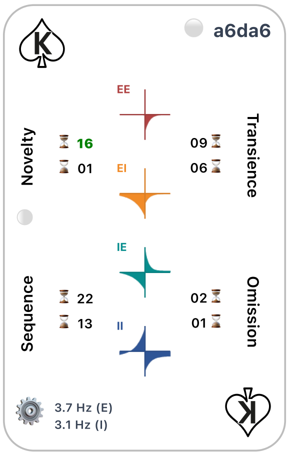
a6da6
EE
EI
IE
II
Novelty 16 01
Transience 09 06
Sequence 22 13
Omission 02 01
3.7 Hz (E)
3.1 Hz (I)

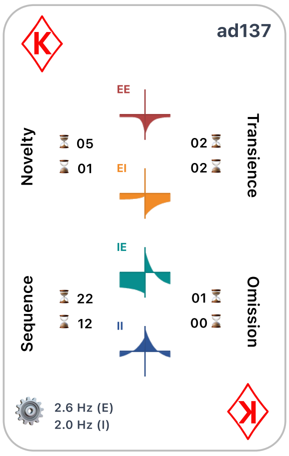
ad137
EE
EI
IE
II
Novelty 05 01
Transience 02 02
Sequence 22 12
Omission 01 00
2.6 Hz (E)
2.0 Hz (I)

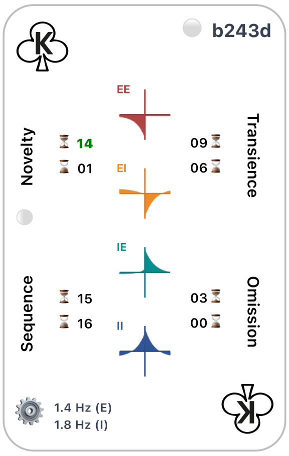
b243d
EE
EI
IE
II
Novelty 14 01
Transience 09 06
Sequence 15 16
Omission 03 00
1.4 Hz (E)
1.8 Hz (I)

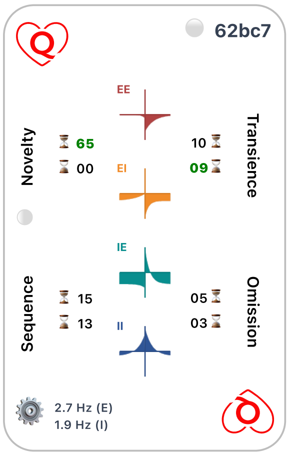
62bc7
EE
EI
IE
II
Novelty 65 00
Transience 10 09
Sequence 15 13
Omission 05 03
2.7 Hz (E)
1.9 Hz (I)

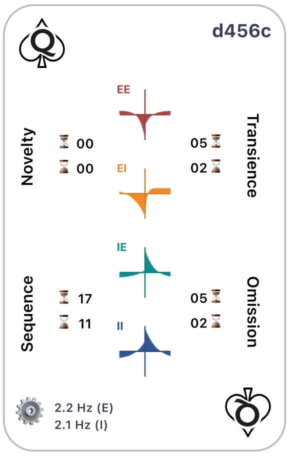
d456c
EE
EI
IE
II
Novelty 00 00
Transience 05 02
Sequence 17 11
Omission 05 02
2.2 Hz (E)
2.1 Hz (I)

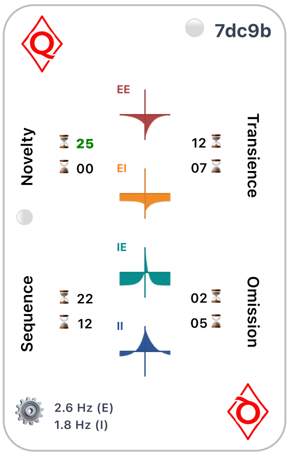
7dc9b
EE
EI
IE
II
Novelty 25 00
Transience 12 07
Sequence 22 12
Omission 02 05
2.6 Hz (E)
1.8 Hz (I)

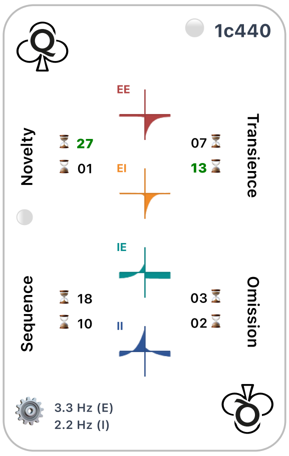
1c440
EE
EI
IE
II
Novelty 27 01
Transience 07 13
Sequence 18 10
Omission 03 02
3.3 Hz (E)
2.2 Hz (I)

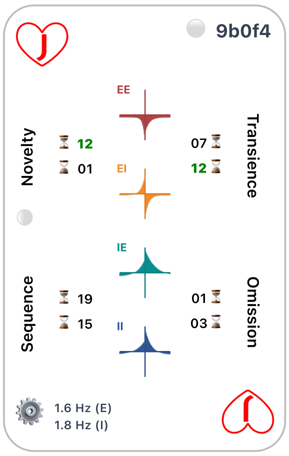
J
9b0f4
EE
Novelty 12 01
Transience 07 12
EI
IE
Sequence 19 15
Omission 01 03
II
1.6 Hz (E)
1.8 Hz (I)

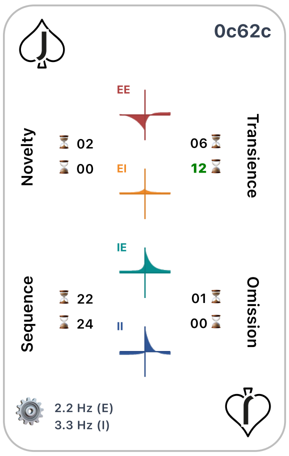
J
0c62c
EE
Novelty 02 00
Transience 06 12
EI
IE
Sequence 22 24
Omission 01 00
II
2.2 Hz (E)
3.3 Hz (I)

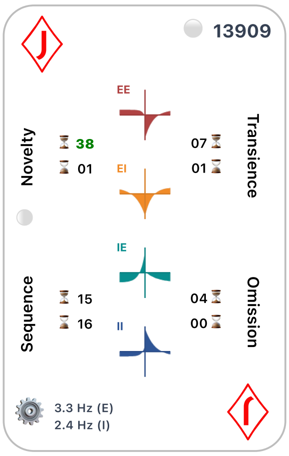
J
13909
EE
Novelty 38 01
Transience 07 01
EI
IE
Sequence 15 16
Omission 04 00
II
3.3 Hz (E)
2.4 Hz (I)

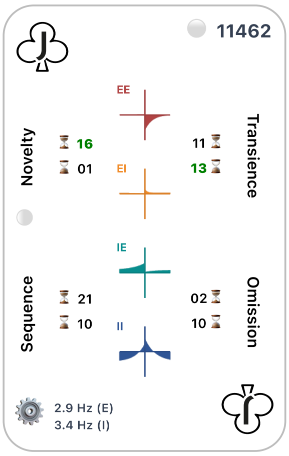
J
11462
EE
Novelty 16 01
Transience 11 13
EI
IE
Sequence 21 10
Omission 02 10
II
2.9 Hz (E)
3.4 Hz (I)

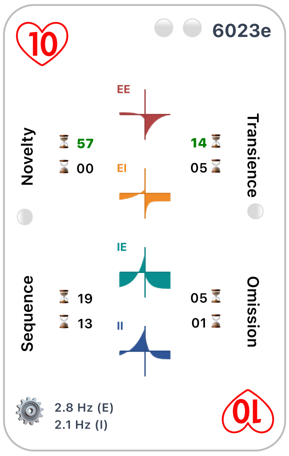
10
6023e
EE
Novelty 57 00
Transience 14 05
EI
IE
Sequence 19 13
Omission 05 01
II
2.8 Hz (E)
2.1 Hz (I)

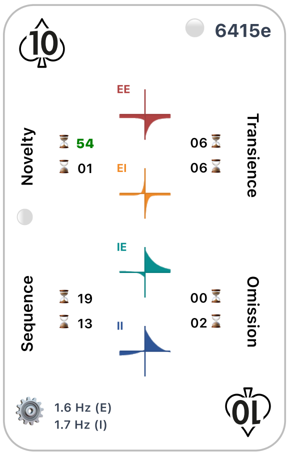
10
6415e
EE
Novelty 54 01
Transience 06 06
EI
IE
Sequence 19 13
Omission 00 02
II
1.6 Hz (E)
1.7 Hz (I)

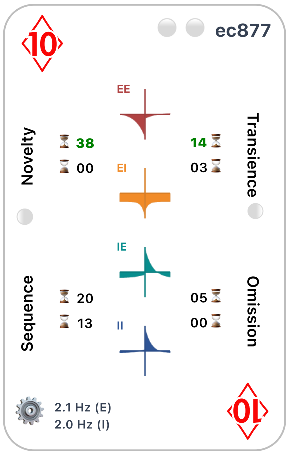
10
ec877
EE
Novelty 38 00
Transience 14 03
EI
IE
Sequence 20 13
Omission 05 00
II
2.1 Hz (E)
2.0 Hz (I)

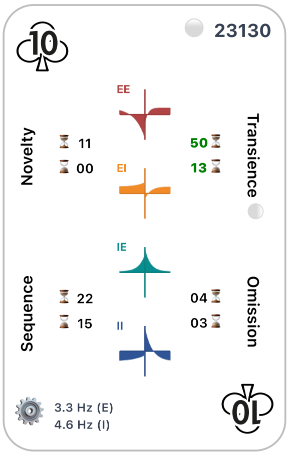
10
23130
EE
Novelty 11 00
Transience 50 13
EI
IE
Sequence 22 15
Omission 04 03
II
3.3 Hz (E)
4.6 Hz (I)

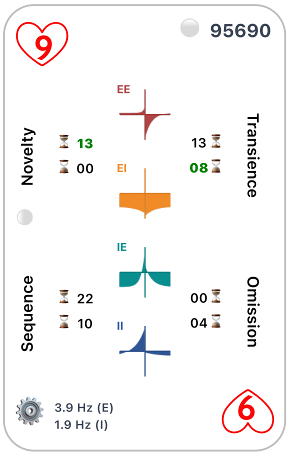
9
95690
EE
Novelty 13 00
Transience 13 08
EI
IE
Sequence 22 10
Omission 00 04
II
3.9 Hz (E)
1.9 Hz (I)

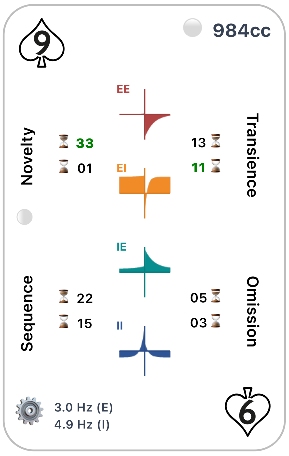
9
984cc
EE
Novelty 33 01
Transience 13 11
EI
IE
Sequence 22 15
Omission 05 03
II
3.0 Hz (E)
4.9 Hz (I)

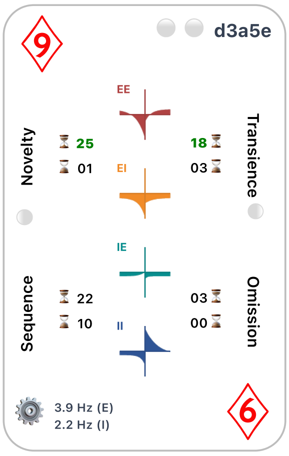
9
d3a5e
EE
Novelty 25 01
Transience 18 03
EI
IE
Sequence 22 10
Omission 03 00
II
3.9 Hz (E)
2.2 Hz (I)

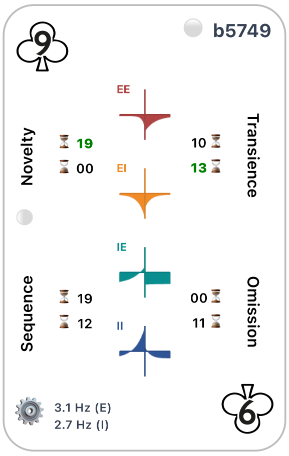
9
b5749
EE
Novelty 19 00
Transience 10 13
EI
IE
Sequence 19 12
Omission 00 11
II
3.1 Hz (E)
2.7 Hz (I)

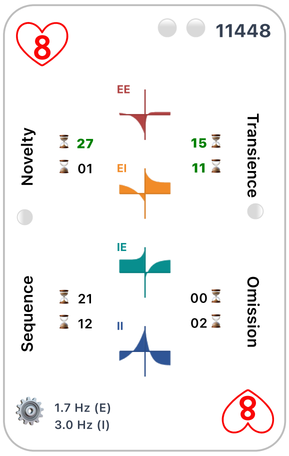
8
11448
EE
Novelty 27 01
Transience 15 11
EI
IE
Sequence 21 12
Omission 00 02
II
1.7 Hz (E)
3.0 Hz (I)

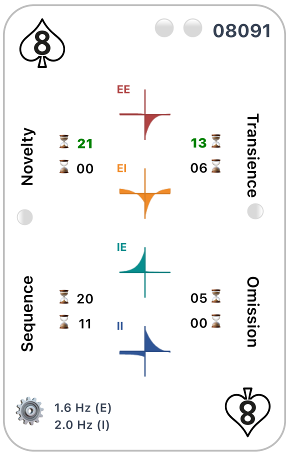
8
08091
EE
Novelty 21 00
Transience 13 06
EI
IE
Sequence 20 11
Omission 05 00
II
1.6 Hz (E)
2.0 Hz (I)

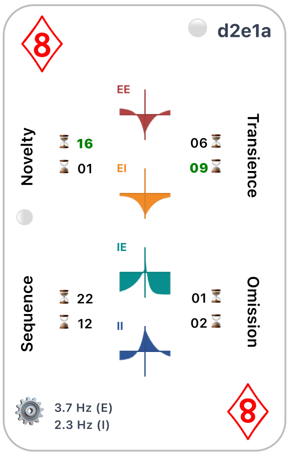
8
d2e1a
EE
Novelty 16 01
Transience 06 09
EI
IE
Sequence 22 12
Omission 01 02
II
3.7 Hz (E)
2.3 Hz (I)

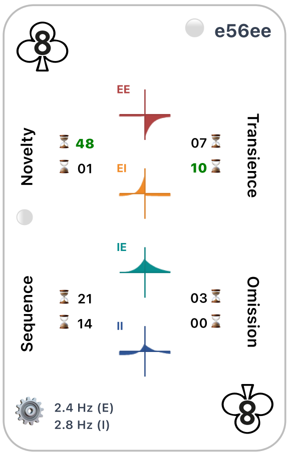
8
e56ee
EE
Novelty 48 01
Transience 07 10
EI
IE
Sequence 21 14
Omission 03 00
II
2.4 Hz (E)
2.8 Hz (I)

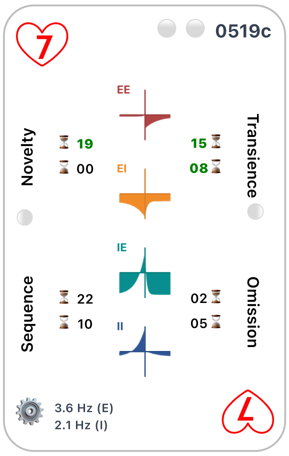
7
0519c
EE
EI
IE
II
Novelty
19
00
Transience
15
08
Sequence
22
10
Omission
02
05
3.6 Hz (E)
2.1 Hz (I)

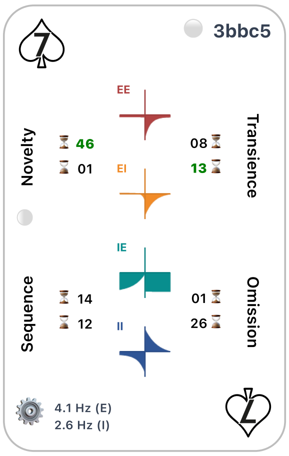
7
3bbc5
EE
EI
IE
II
Novelty
46
01
Transience
08
13
Sequence
14
12
Omission
01
26
4.1 Hz (E)
2.6 Hz (I)

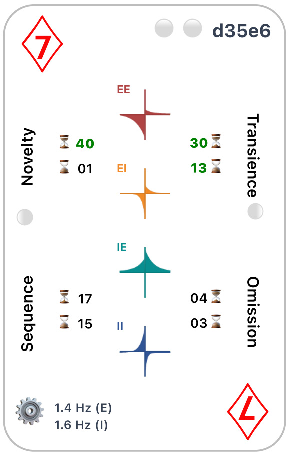
7
d35e6
EE
EI
IE
II
Novelty
40
01
Transience
30
13
Sequence
17
15
Omission
04
03
1.4 Hz (E)
1.6 Hz (I)

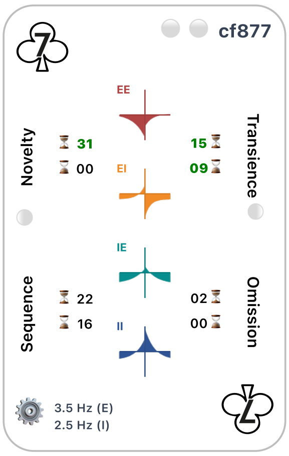
7
cf877
EE
EI
IE
II
Novelty
31
00
Transience
15
09
Sequence
22
16
Omission
02
00
3.5 Hz (E)
2.5 Hz (I)

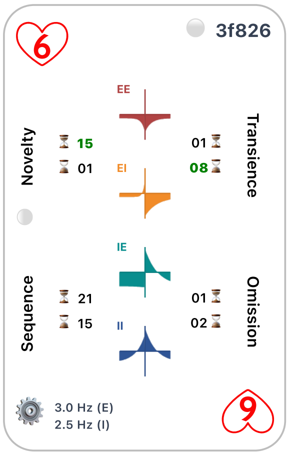
6
3f826
EE
EI
IE
II
Novelty
15
01
Transience
01
08
Sequence
21
15
Omission
01
02
3.0 Hz (E)
2.5 Hz (I)

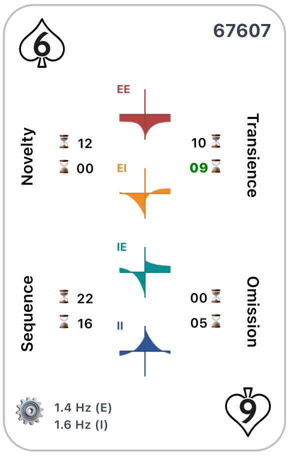
6
67607
EE
EI
IE
II
Novelty
12
00
Transience
10
09
Sequence
22
16
Omission
00
05
1.4 Hz (E)
1.6 Hz (I)

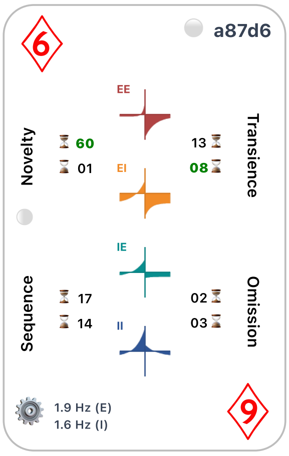
6
a87d6
EE
EI
IE
II
Novelty
60
01
Transience
13
08
Sequence
17
14
Omission
02
03
1.9 Hz (E)
1.6 Hz (I)

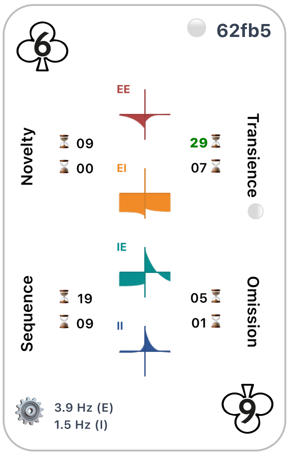
6
62fb5
EE
EI
IE
II
Novelty
09
00
Transience
29
07
Sequence
19
09
Omission
05
01
3.9 Hz (E)
1.5 Hz (I)

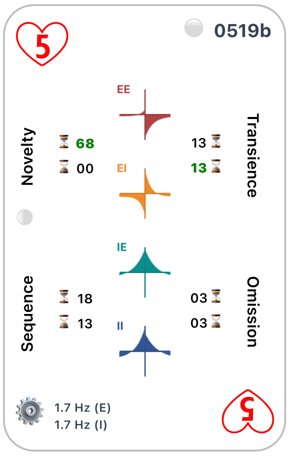
5
0519b
EE
EI
IE
II
Novelty
68
00
Transience
13
13
Sequence
18
13
Omission
03
03
1.7 Hz (E)
1.7 Hz (I)

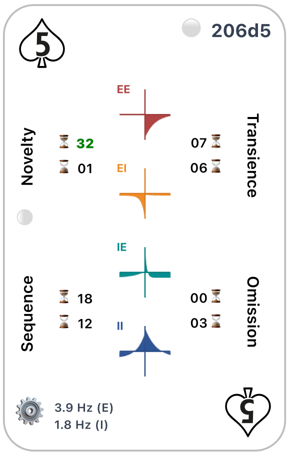
5
206d5
EE
EI
IE
II
Novelty
32
01
Transience
07
06
Sequence
18
12
Omission
00
03
3.9 Hz (E)
1.8 Hz (I)

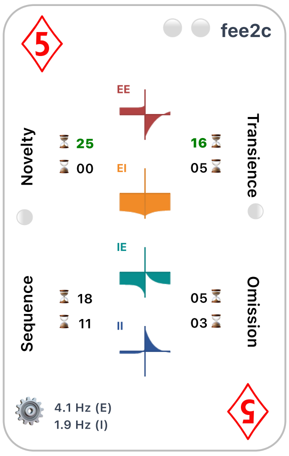
5
fee2c
EE
EI
IE
II
Novelty
25
00
Transience
16
05
Sequence
18
11
Omission
05
03
4.1 Hz (E)
1.9 Hz (I)

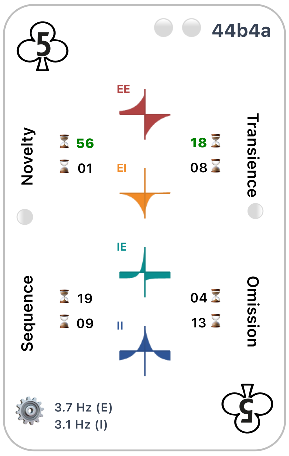
5
44b4a
EE
EI
IE
II
Novelty
56
01
Transience
18
08
Sequence
19
09
Omission
04
13
3.7 Hz (E)
3.1 Hz (I)

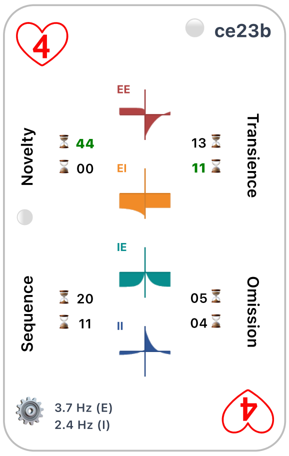
4
ce23b
EE
EI
IE
II
Novelty
44
00
Transience
13
11
Sequence
20
11
Omission
05
04
3.7 Hz (E)
2.4 Hz (I)

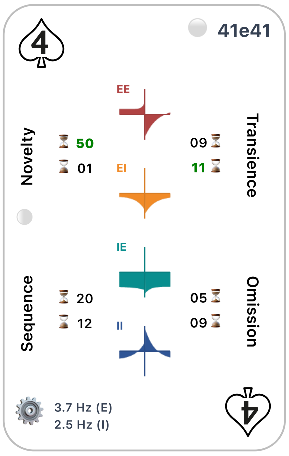
4
41e41
EE
EI
IE
II
Novelty
50
01
Transience
09
11
Sequence
20
12
Omission
05
09
3.7 Hz (E)
2.5 Hz (I)

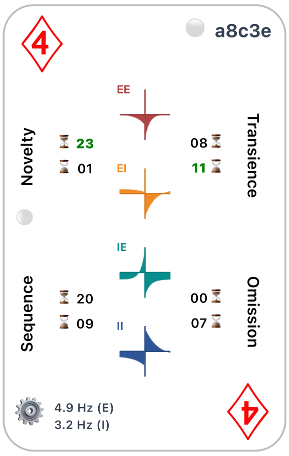
4
a8c3e
EE
EI
IE
II
Novelty
23
01
Transience
08
11
Sequence
20
09
Omission
00
07
4.9 Hz (E)
3.2 Hz (I)

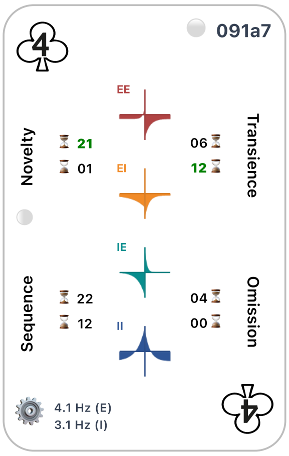
4
091a7
EE
EI
IE
II
Novelty
21
01
Transience
06
12
Sequence
22
12
Omission
04
00
4.1 Hz (E)
3.1 Hz (I)

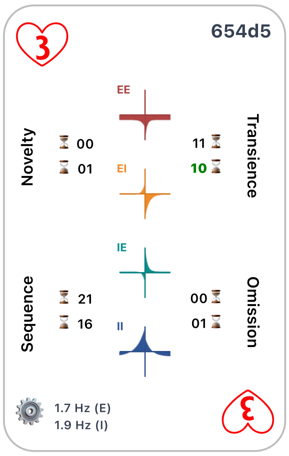
3
654d5
Novelty 00 01
Transience 11 10
EE EI IE II
Sequence 21 16
Omission 00 01
1.7 Hz (E)
1.9 Hz (I)
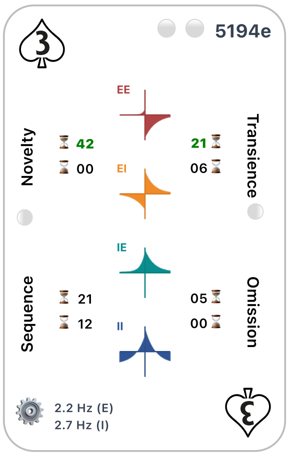
3
5194e
Novelty 42 00
Transience 21 06
EE EI IE II
Sequence 21 12
Omission 05 00
2.2 Hz (E)
2.7 Hz (I)
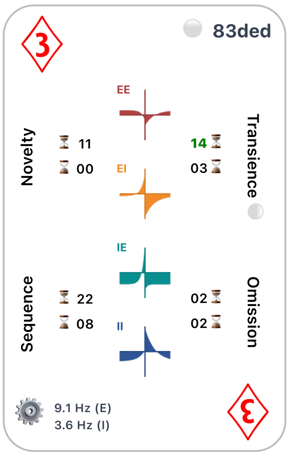
3
83ded
Novelty 11 00
Transience 14 03
EE EI IE II
Sequence 22 08
Omission 02 02
9.1 Hz (E)
3.6 Hz (I)
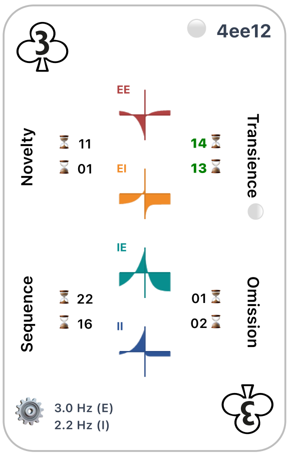
3
4ee12
Novelty 11 01
Transience 14 13
EE EI IE II
Sequence 22 16
Omission 01 02
3.0 Hz (E)
2.2 Hz (I)
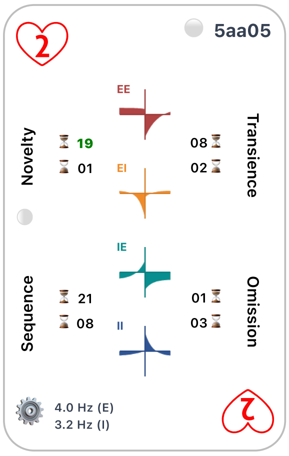
2
5aa05
Novelty 19 01
Transience 08 02
EE EI IE II
Sequence 21 08
Omission 01 03
4.0 Hz (E)
3.2 Hz (I)
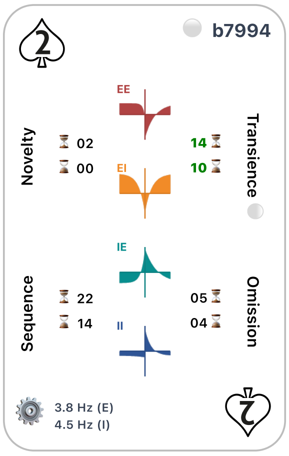
2
b7994
Novelty 02 00
Transience 14 10
EE EI IE II
Sequence 22 14
Omission 05 04
3.8 Hz (E)
4.5 Hz (I)
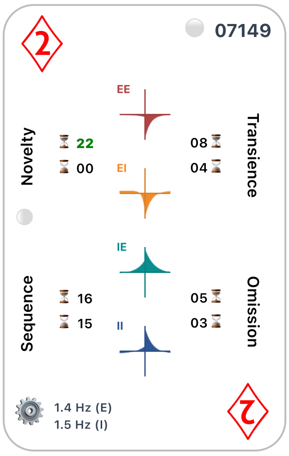
2
07149
Novelty 22 00
Transience 08 04
EE EI IE II
Sequence 16 15
Omission 05 03
1.4 Hz (E)
1.5 Hz (I)
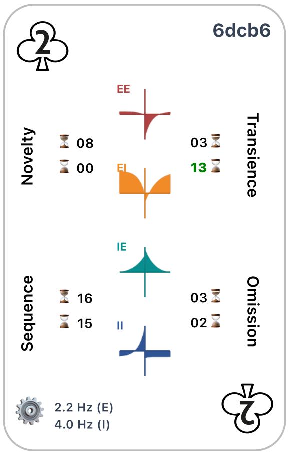
2
6dcb6
Novelty 08 00
Transience 03 13
EE EI IE II
Sequence 16 15
Omission 03 02
2.2 Hz (E)
4.0 Hz (I)
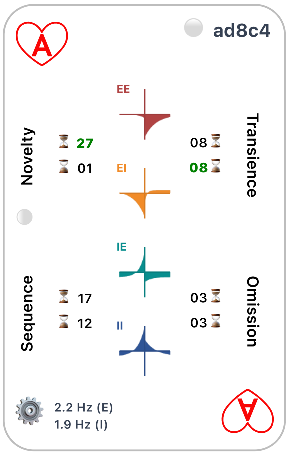
A
ad8c4
Novelty 27 01
Transience 08 08
EE EI IE II
Sequence 17 12
Omission 03 03
2.2 Hz (E)
1.9 Hz (I)
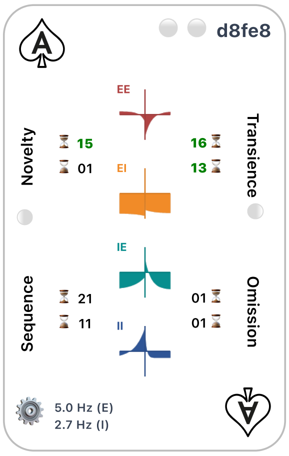
A
d8fe8
Novelty 15 01
Transience 16 13
EE EI IE II
Sequence 21 11
Omission 01 01
5.0 Hz (E)
2.7 Hz (I)
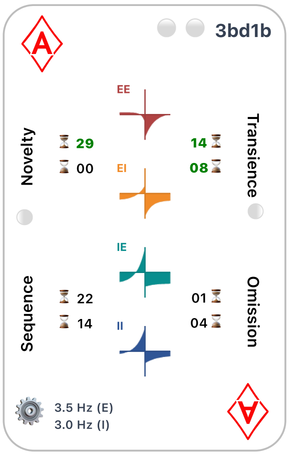
A
3bd1b
Novelty 29 00
Transience 14 08
EE EI IE II
Sequence 22 14
Omission 01 04
3.5 Hz (E)
3.0 Hz (I)
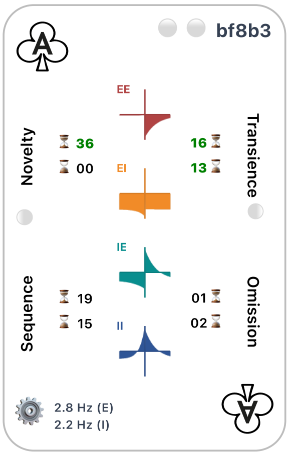
A
bf8b3
Novelty 36 00
Transience 16 13
EE EI IE II
Sequence 19 15
Omission 01 02
2.8 Hz (E)
2.2 Hz (I)
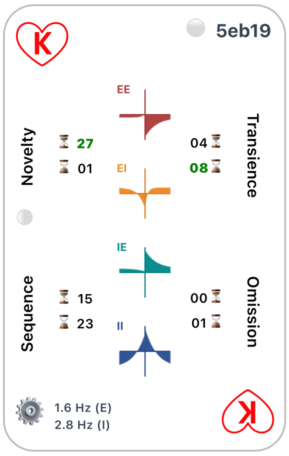
K
5eb19
Novelty 27 01
Transience 04 08
EE EI IE II
Sequence 15 23
Omission 00 01
1.6 Hz (E)
2.8 Hz (I)
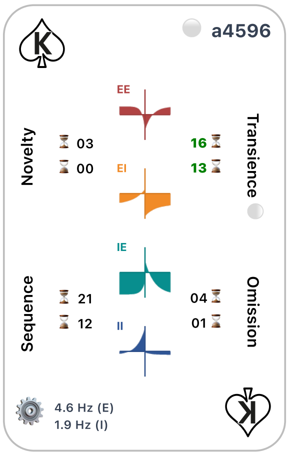
K
a4596
Novelty 03 00
Transience 16 13
EE EI IE II
Sequence 21 12
Omission 04 01
4.6 Hz (E)
1.9 Hz (I)
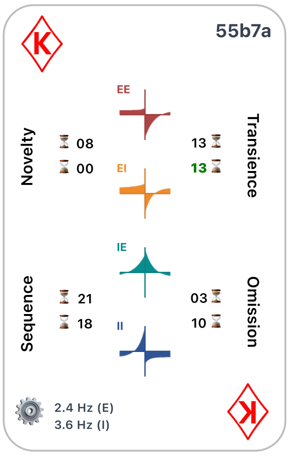
K
55b7a
Novelty 08 00
Transience 13 13
EE EI IE II
Sequence 21 18
Omission 03 10
2.4 Hz (E)
3.6 Hz (I)
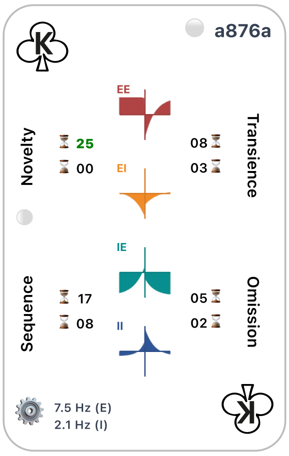
K
a876a
Novelty 25 00
Transience 08 03
EE EI IE II
Sequence 17 08
Omission 05 02
7.5 Hz (E)
2.1 Hz (I)

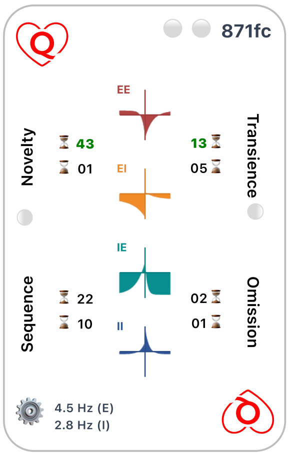
Q
871fc
Novelty 43 01
Transience 13 05
Sequence 22 10
Omission 02 01
EE EI IE II
4.5 Hz (E)
2.8 Hz (I)

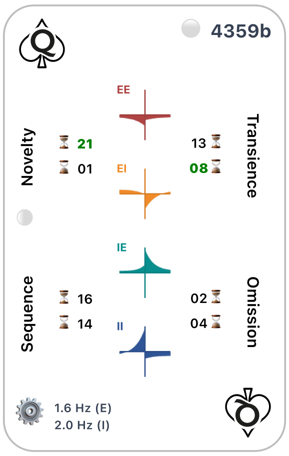
Q
4359b
Novelty 21 01
Transience 13 08
Sequence 16 14
Omission 02 04
EE EI IE II
1.6 Hz (E)
2.0 Hz (I)

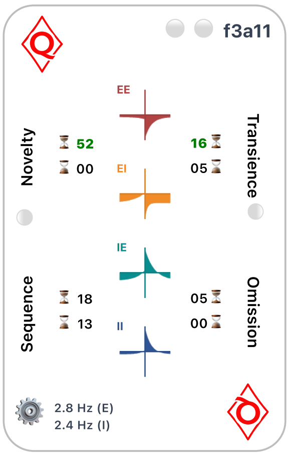
Q
f3a11
Novelty 52 00
Transience 16 05
Sequence 18 13
Omission 05 00
EE EI IE II
2.8 Hz (E)
2.4 Hz (I)

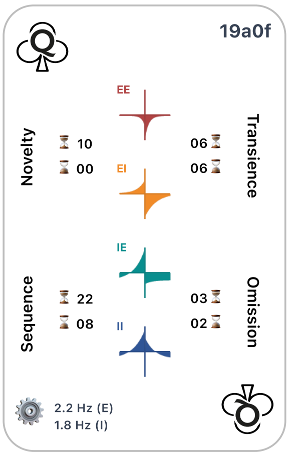
Q
19a0f
Novelty 10 00
Transience 06 06
Sequence 22 08
Omission 03 02
EE EI IE II
2.2 Hz (E)
1.8 Hz (I)

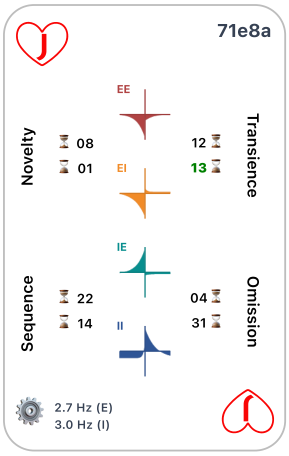
J
71e8a
Novelty 08 01
Transience 12 13
Sequence 22 14
Omission 04 31
EE EI IE II
2.7 Hz (E)
3.0 Hz (I)

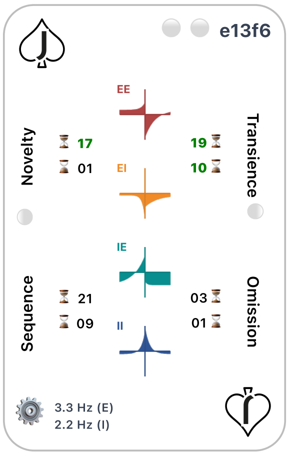
J
e13f6
Novelty 17 01
Transience 19 10
Sequence 21 09
Omission 03 01
EE EI IE II
3.3 Hz (E)
2.2 Hz (I)

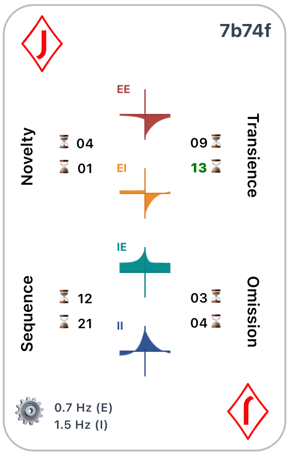
J
7b74f
Novelty 04 01
Transience 09 13
Sequence 12 21
Omission 03 04
EE EI IE II
0.7 Hz (E)
1.5 Hz (I)

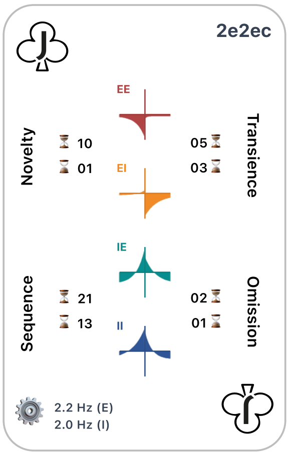
J
2e2ec
Novelty 10 01
Transience 05 03
Sequence 21 13
Omission 02 01
EE EI IE II
2.2 Hz (E)
2.0 Hz (I)

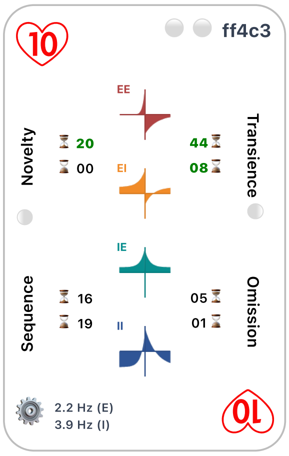
10
ff4c3
Novelty 20 00
Transience 44 08
Sequence 16 19
Omission 05 01
EE EI IE II
2.2 Hz (E)
3.9 Hz (I)

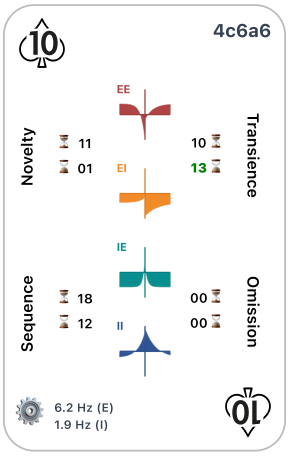
10
4c6a6
Novelty 11 01
Transience 10 13
Sequence 18 12
Omission 00 00
EE EI IE II
6.2 Hz (E)
1.9 Hz (I)

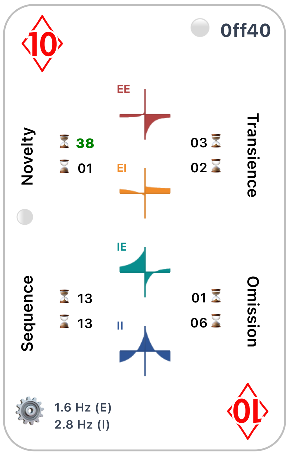
10
0ff40
Novelty 38 01
Transience 03 02
Sequence 13 13
Omission 01 06
EE EI IE II
1.6 Hz (E)
2.8 Hz (I)

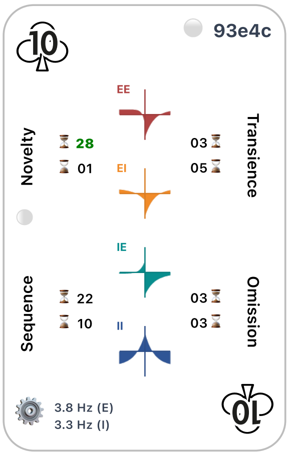
10
93e4c
Novelty 28 01
Transience 03 05
Sequence 22 10
Omission 03 03
EE EI IE II
3.8 Hz (E)
3.3 Hz (I)

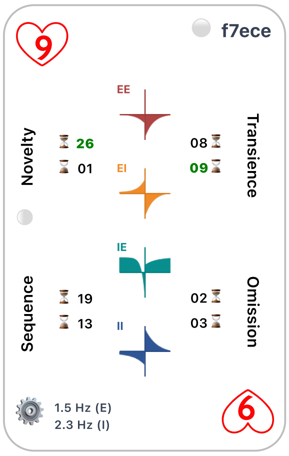
9
f7ece
Novelty 26 01
Transience 08 09
Sequence 19 13
Omission 02 03
EE EI IE II
1.5 Hz (E)
2.3 Hz (I)

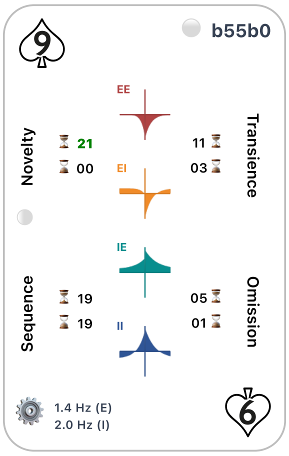
9
b55b0
Novelty 21 00
Transience 11 03
Sequence 19 19
Omission 05 01
EE EI IE II
1.4 Hz (E)
2.0 Hz (I)

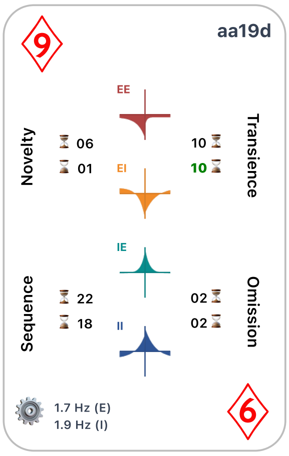
9
aa19d
Novelty 06 01
Transience 10 10
Sequence 22 18
Omission 02 02
EE EI IE II
1.7 Hz (E)
1.9 Hz (I)

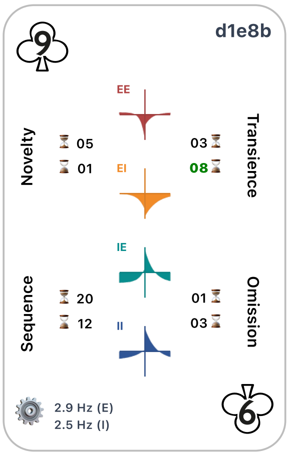
9
d1e8b
Novelty 05 01
Transience 03 08
Sequence 20 12
Omission 01 03
EE EI IE II
2.9 Hz (E)
2.5 Hz (I)

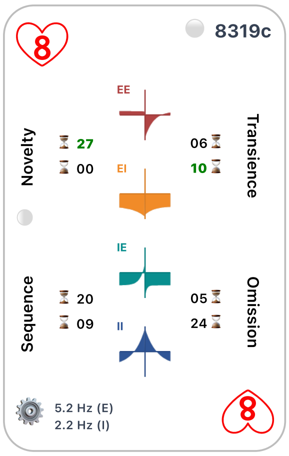
8
8319c
EE
EI
IE
II
Novelty
27
00
Transience
06
10
Sequence
20
09
Omission
05
24
5.2 Hz (E)
2.2 Hz (I)
8

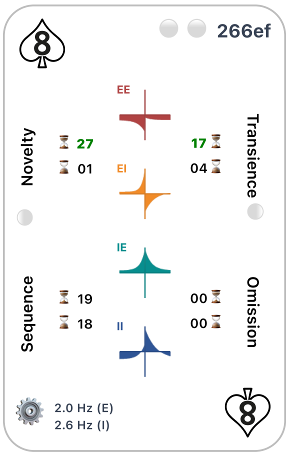
8
266ef
EE
EI
IE
II
Novelty
27
01
Transience
17
04
Sequence
19
18
Omission
00
00
2.0 Hz (E)
2.6 Hz (I)
8

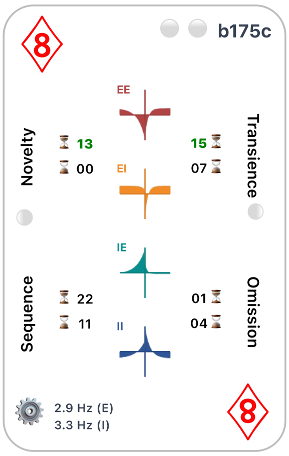
8
b175c
EE
EI
IE
II
Novelty
13
00
Transience
15
07
Sequence
22
11
Omission
01
04
2.9 Hz (E)
3.3 Hz (I)
8

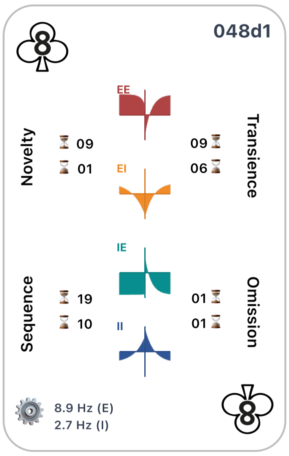
8
048d1
EE
EI
IE
II
Novelty
09
01
Transience
09
06
Sequence
19
10
Omission
01
01
8.9 Hz (E)
2.7 Hz (I)
8

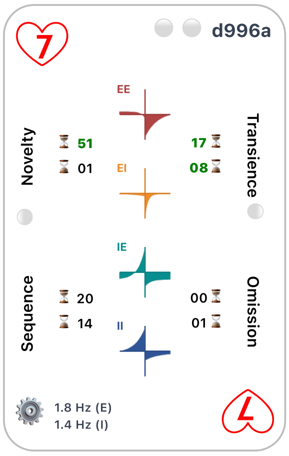
7
d996a
EE
EI
IE
II
Novelty
51
01
Transience
17
08
Sequence
20
14
Omission
00
01
1.8 Hz (E)
1.4 Hz (I)
7

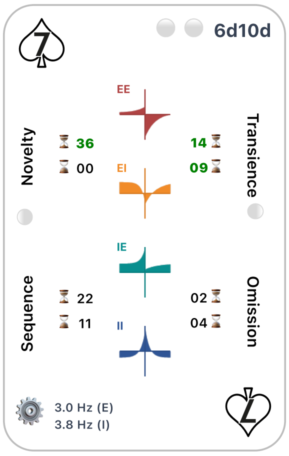
7
6d10d
EE
EI
IE
II
Novelty
36
00
Transience
14
09
Sequence
22
11
Omission
02
04
3.0 Hz (E)
3.8 Hz (I)
7

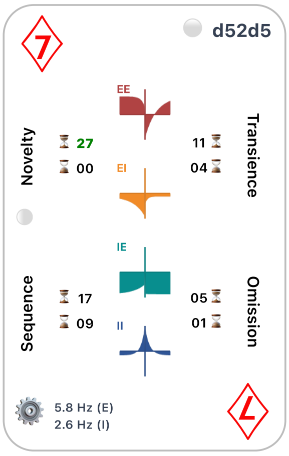
7
d52d5
EE
EI
IE
II
Novelty
27
00
Transience
11
04
Sequence
17
09
Omission
05
01
5.8 Hz (E)
2.6 Hz (I)
7

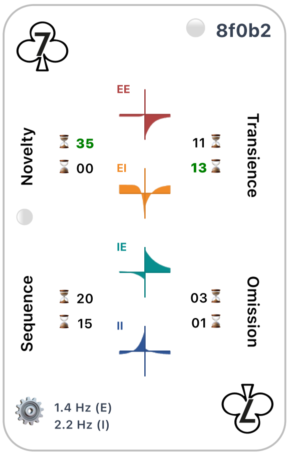
7
8f0b2
EE
EI
IE
II
Novelty
35
00
Transience
11
13
Sequence
20
15
Omission
03
01
1.4 Hz (E)
2.2 Hz (I)
7

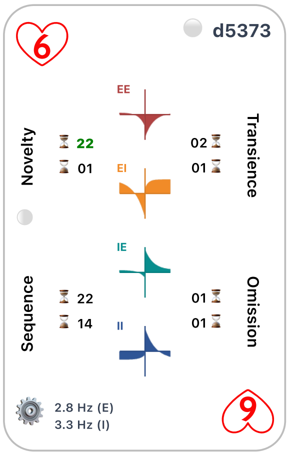
6
d5373
EE
EI
IE
II
Novelty
22
01
Transience
02
01
Sequence
22
14
Omission
01
01
2.8 Hz (E)
3.3 Hz (I)
6

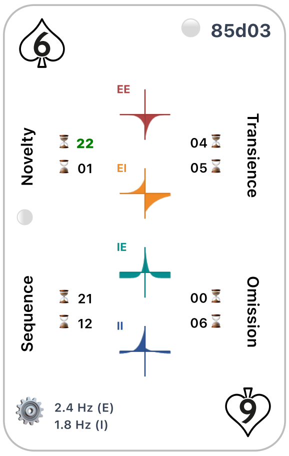
6
85d03
EE
EI
IE
II
Novelty
22
01
Transience
04
05
Sequence
21
12
Omission
00
06
2.4 Hz (E)
1.8 Hz (I)
6

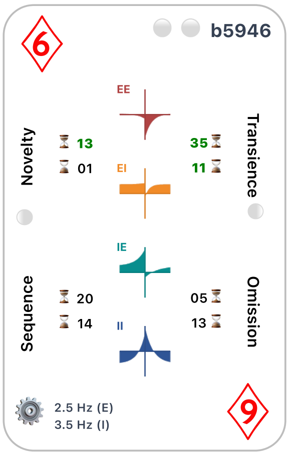
6
b5946
EE
EI
IE
II
Novelty
13
01
Transience
35
11
Sequence
20
14
Omission
05
13
2.5 Hz (E)
3.5 Hz (I)
6

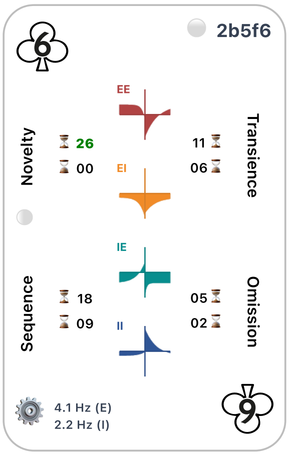
6
2b5f6
EE
EI
IE
II
Novelty
26
00
Transience
11
06
Sequence
18
09
Omission
05
02
4.1 Hz (E)
2.2 Hz (I)
6

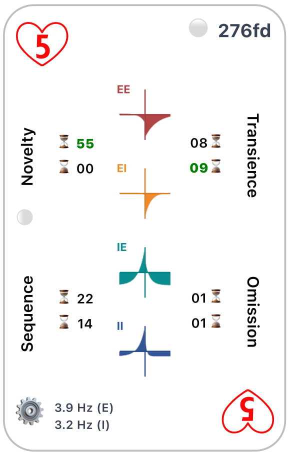
5
276fd
EE
EI
IE
II
Novelty
55
00
Transience
08
09
Sequence
22
14
Omission
01
01
3.9 Hz (E)
3.2 Hz (I)
5

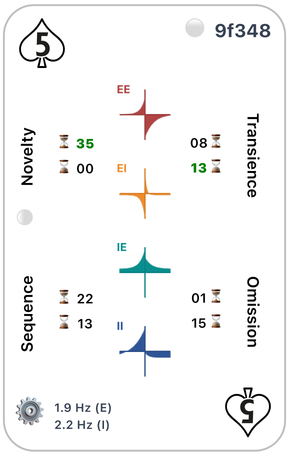
5
9f348
EE
EI
IE
II
Novelty
35
00
Transience
08
13
Sequence
22
13
Omission
01
15
1.9 Hz (E)
2.2 Hz (I)
5

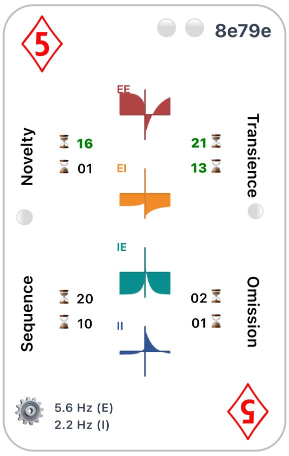
5
8e79e
EE
EI
IE
II
Novelty
16
01
Transience
21
13
Sequence
20
10
Omission
02
01
5.6 Hz (E)
2.2 Hz (I)
5

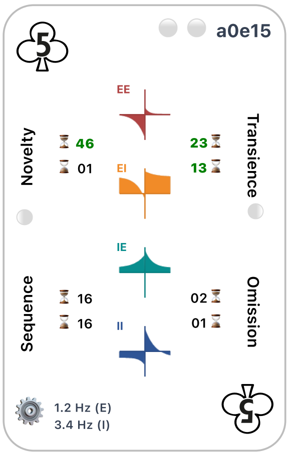
5
a0e15
EE
EI
IE
II
Novelty
46
01
Transience
23
13
Sequence
16
16
Omission
02
01
1.2 Hz (E)
3.4 Hz (I)
5

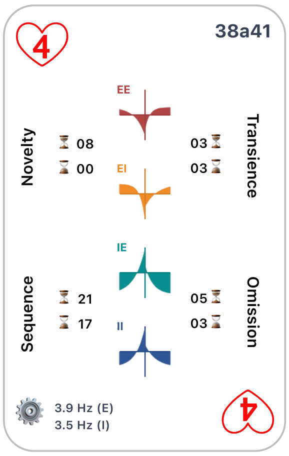
4
38a41
EE
Novelty
08
00
03
03
Transience
EI
IE
Sequence
21
17
05
03
Omission
II
3.9 Hz (E)
3.5 Hz (I)

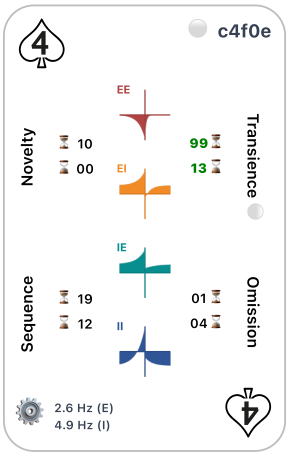
4
c4f0e
EE
Novelty
10
00
99
13
Transience
EI
IE
Sequence
19
12
01
04
Omission
II
2.6 Hz (E)
4.9 Hz (I)

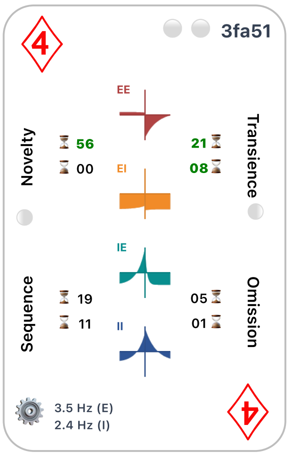
4
3fa51
EE
Novelty
56
00
21
08
Transience
EI
IE
Sequence
19
11
05
01
Omission
II
3.5 Hz (E)
2.4 Hz (I)

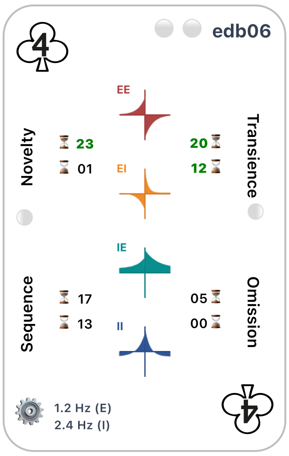
4
edb06
EE
Novelty
23
01
20
12
Transience
EI
IE
Sequence
17
13
05
00
Omission
II
1.2 Hz (E)
2.4 Hz (I)

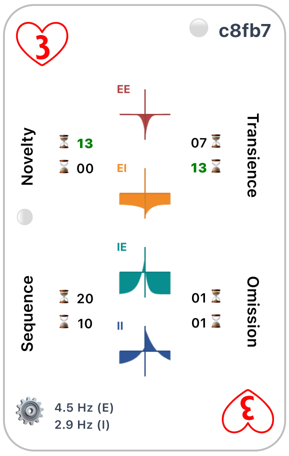
3
c8fb7
EE
Novelty
13
00
07
13
Transience
EI
IE
Sequence
20
10
01
01
Omission
II
4.5 Hz (E)
2.9 Hz (I)

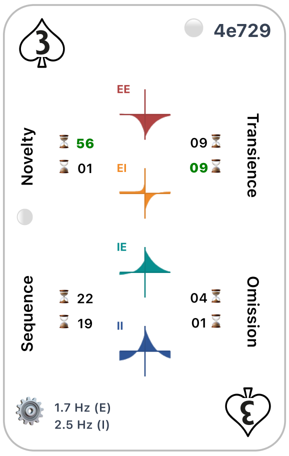
3
4e729
EE
Novelty
56
01
09
09
Transience
EI
IE
Sequence
22
19
04
01
Omission
II
1.7 Hz (E)
2.5 Hz (I)

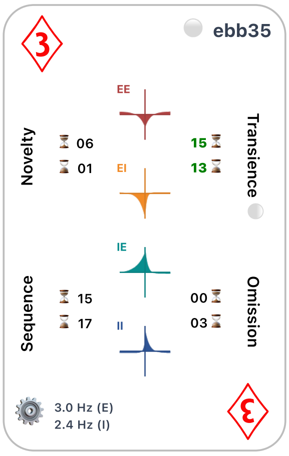
3
ebb35
EE
Novelty
06
01
15
13
Transience
EI
IE
Sequence
15
17
00
03
Omission
II
3.0 Hz (E)
2.4 Hz (I)

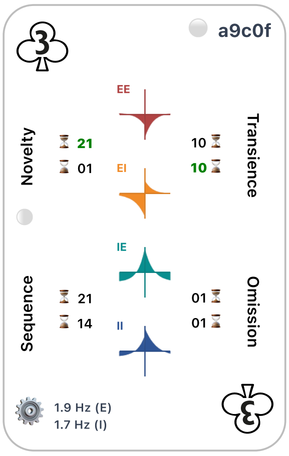
3
a9c0f
EE
Novelty
21
01
10
10
Transience
EI
IE
Sequence
21
14
01
01
Omission
II
1.9 Hz (E)
1.7 Hz (I)

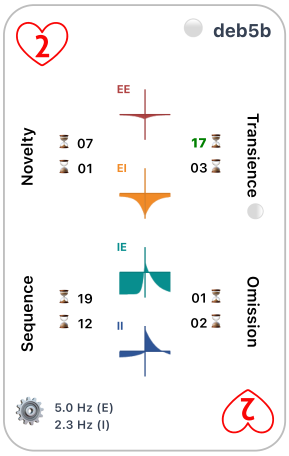
2
deb5b
EE
Novelty
07
01
17
03
Transience
EI
IE
Sequence
19
12
01
02
Omission
II
5.0 Hz (E)
2.3 Hz (I)

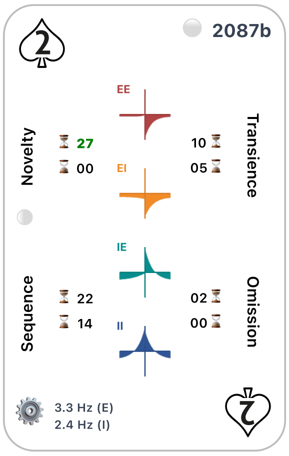
2
2087b
EE
Novelty
27
00
10
05
Transience
EI
IE
Sequence
22
14
02
00
Omission
II
3.3 Hz (E)
2.4 Hz (I)

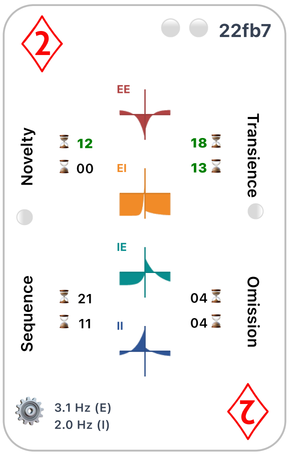
2
22fb7
EE
Novelty
12
00
18
13
Transience
EI
IE
Sequence
21
11
04
04
Omission
II
3.1 Hz (E)
2.0 Hz (I)

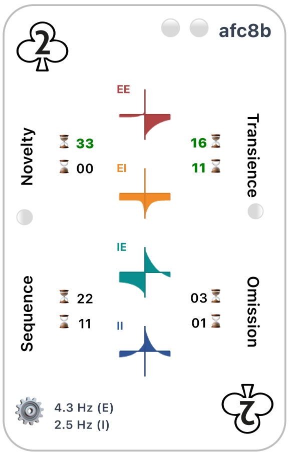
2
afc8b
EE
Novelty
33
00
16
11
Transience
EI
IE
Sequence
22
11
03
01
Omission
II
4.3 Hz (E)
2.5 Hz (I)

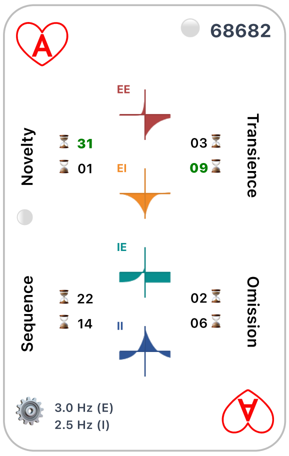
A
68682
EE
Novelty
31
01
03
09
Transience
EI
IE
Sequence
22
14
02
06
Omission
II
3.0 Hz (E)
2.5 Hz (I)

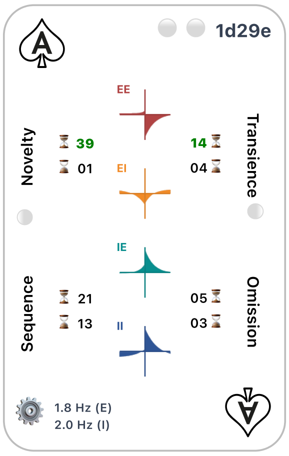
A
1d29e
EE
Novelty
39
01
14
04
Transience
EI
IE
Sequence
21
13
05
03
Omission
II
1.8 Hz (E)
2.0 Hz (I)

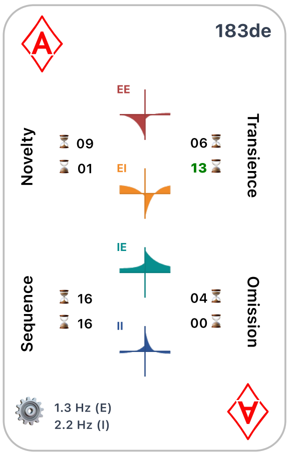
A
183de
EE
Novelty
09
01
06
13
Transience
EI
IE
Sequence
16
16
04
00
Omission
II
1.3 Hz (E)
2.2 Hz (I)

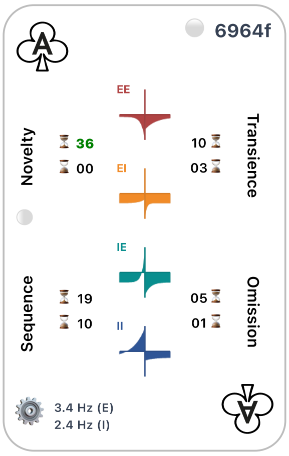
A
6964f
EE
Novelty
36
00
10
03
Transience
EI
IE
Sequence
19
10
05
01
Omission
II
3.4 Hz (E)
2.4 Hz (I)

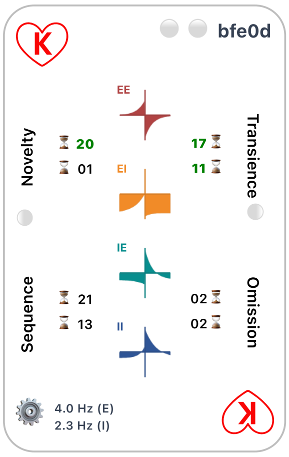
K
bfe0d
Novelty
20
01
EE
EI
17
11
Transience
Sequence
21
13
IE
II
02
02
Omission
4.0 Hz (E)
2.3 Hz (I)

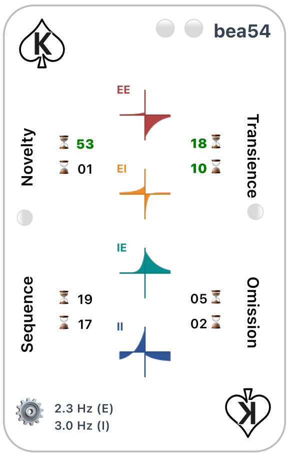
K
bea54
Novelty
53
01
EE
EI
18
10
Transience
Sequence
19
17
IE
II
05
02
Omission
2.3 Hz (E)
3.0 Hz (I)

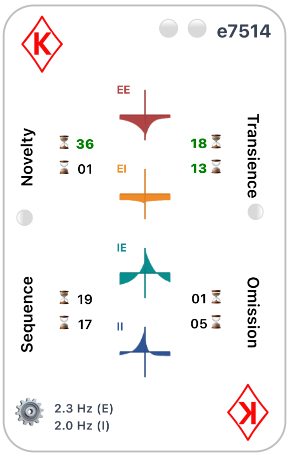
K
e7514
Novelty
36
01
EE
EI
18
13
Transience
Sequence
19
17
IE
II
01
05
Omission
2.3 Hz (E)
2.0 Hz (I)

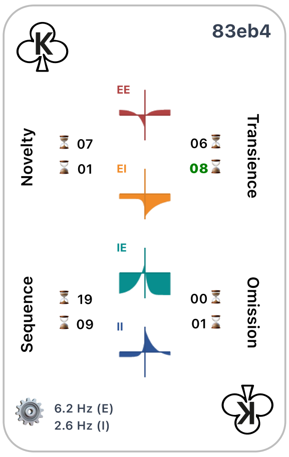
K
83eb4
Novelty
07
01
EE
EI
06
08
Transience
Sequence
19
09
IE
II
00
01
Omission
6.2 Hz (E)
2.6 Hz (I)

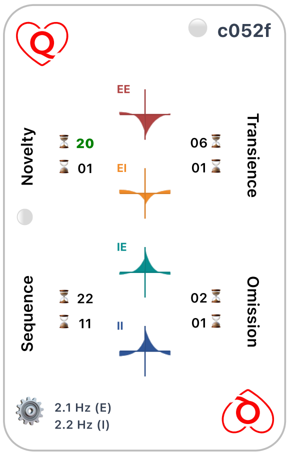
Q
c052f
Novelty
20
01
EE
EI
06
01
Transience
Sequence
22
11
IE
II
02
01
Omission
2.1 Hz (E)
2.2 Hz (I)

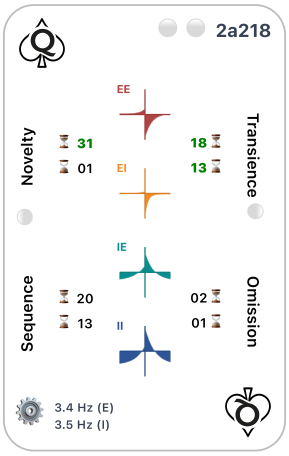
Q
2a218
Novelty
31
01
EE
EI
18
13
Transience
Sequence
20
13
IE
II
02
01
Omission
3.4 Hz (E)
3.5 Hz (I)

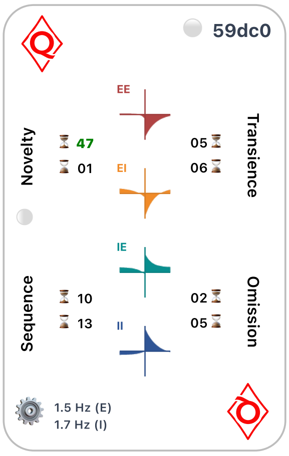
Q
59dc0
Novelty
47
01
EE
EI
05
06
Transience
Sequence
10
13
IE
II
02
05
Omission
1.5 Hz (E)
1.7 Hz (I)

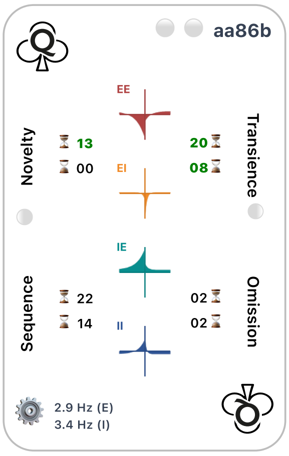
Q
aa86b
Novelty
13
00
EE
EI
20
08
Transience
Sequence
22
14
IE
II
02
02
Omission
2.9 Hz (E)
3.4 Hz (I)

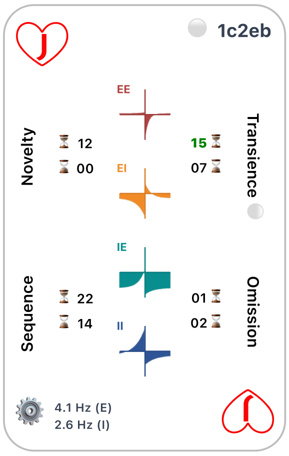
J
1c2eb
Novelty
12
00
EE
EI
15
07
Transience
Sequence
22
14
IE
II
01
02
Omission
4.1 Hz (E)
2.6 Hz (I)

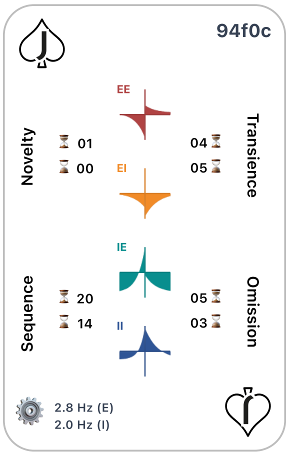
J
94f0c
Novelty
01
00
EE
EI
04
05
Transience
Sequence
20
14
IE
II
05
03
Omission
2.8 Hz (E)
2.0 Hz (I)

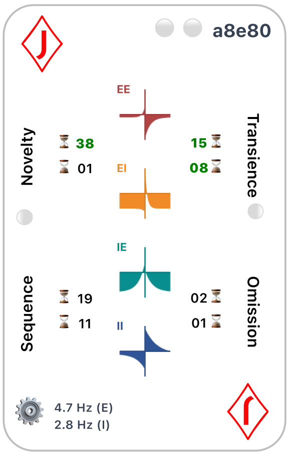
J
a8e80
Novelty
38
01
EE
EI
15
08
Transience
Sequence
19
11
IE
II
02
01
Omission
4.7 Hz (E)
2.8 Hz (I)

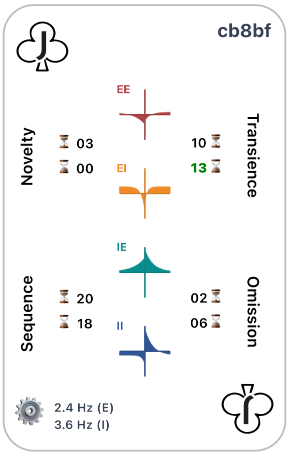
J
cb8bf
Novelty
03
00
EE
EI
10
13
Transience
Sequence
20
18
IE
II
02
06
Omission
2.4 Hz (E)
3.6 Hz (I)

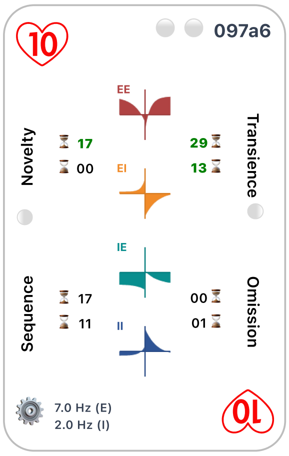
10
097a6
Novelty
17
00
EE
EI
29
13
Transience
Sequence
17
11
IE
II
00
01
Omission
7.0 Hz (E)
2.0 Hz (I)

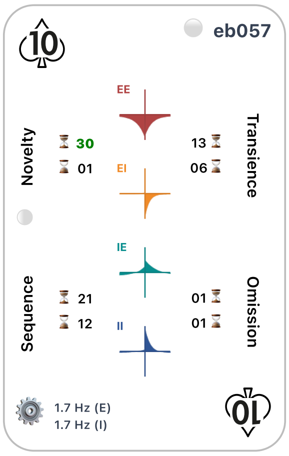
10
eb057
Novelty
30
01
EE
EI
13
06
Transience
Sequence
21
12
IE
II
01
01
Omission
1.7 Hz (E)
1.7 Hz (I)

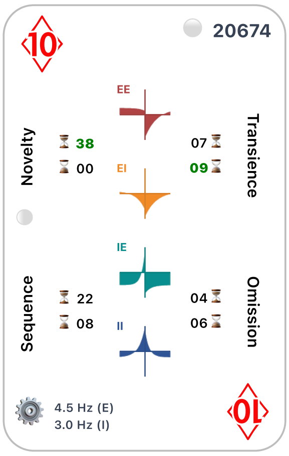
10
20674
Novelty
38
00
EE
EI
07
09
Transience
Sequence
22
08
IE
II
04
06
Omission
4.5 Hz (E)
3.0 Hz (I)

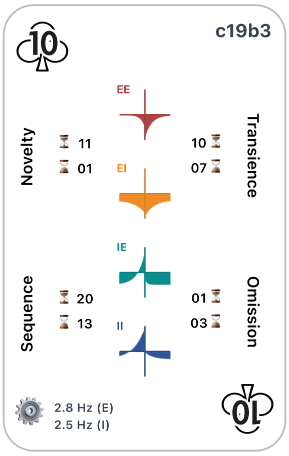
10
c19b3
Novelty
11
01
EE
EI
10
07
Transience
Sequence
20
13
IE
II
01
03
Omission
2.8 Hz (E)
2.5 Hz (I)

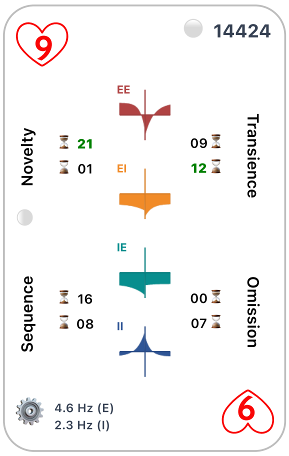
9
14424
EE
Novelty
21
01
EI
09
12
Transience
IE
Sequence
16
08
II
00
07
Omission
4.6 Hz (E)
2.3 Hz (I)
6

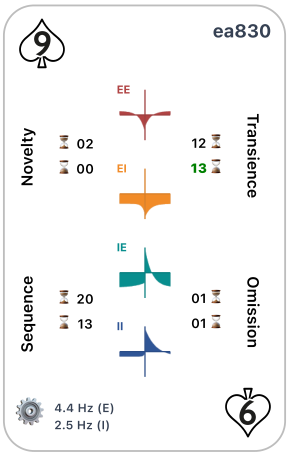
9
ea830
EE
Novelty
02
00
EI
12
13
Transience
IE
Sequence
20
13
II
01
01
Omission
4.4 Hz (E)
2.5 Hz (I)
6

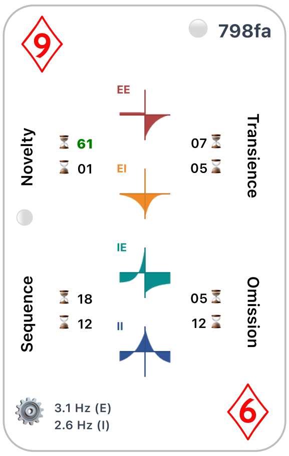
9
798fa
EE
Novelty
61
01
EI
07
05
Transience
IE
Sequence
18
12
II
05
12
Omission
3.1 Hz (E)
2.6 Hz (I)
6

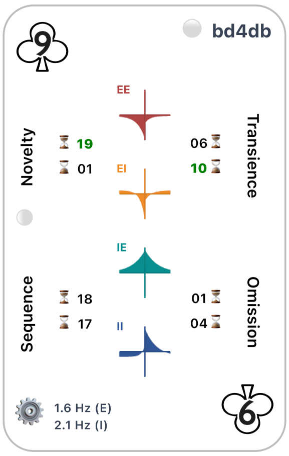
9
bd4db
EE
Novelty
19
01
EI
06
10
Transience
IE
Sequence
18
17
II
01
04
Omission
1.6 Hz (E)
2.1 Hz (I)
6

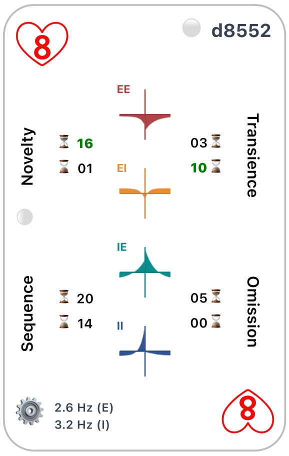
8
d8552
EE
Novelty
16
01
EI
03
10
Transience
IE
Sequence
20
14
II
05
00
Omission
2.6 Hz (E)
3.2 Hz (I)
8

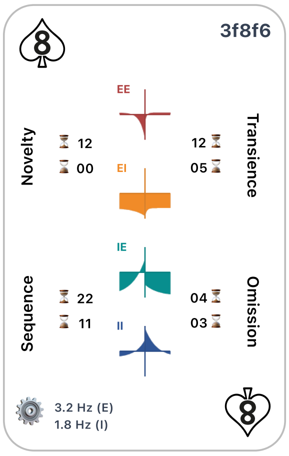
8
3f8f6
EE
Novelty
12
00
EI
12
05
Transience
IE
Sequence
22
11
II
04
03
Omission
3.2 Hz (E)
1.8 Hz (I)
8

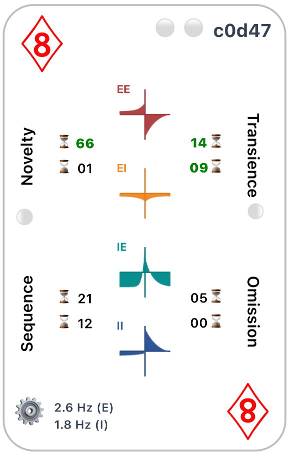
8
c0d47
EE
Novelty
66
01
EI
14
09
Transience
IE
Sequence
21
12
II
05
00
Omission
2.6 Hz (E)
1.8 Hz (I)
8

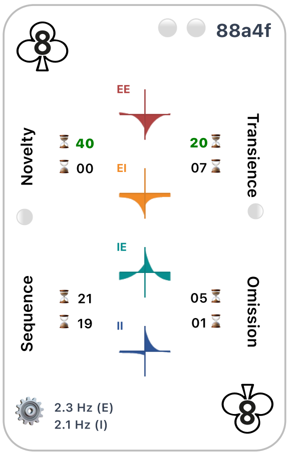
8
88a4f
EE
Novelty
40
00
EI
20
07
Transience
IE
Sequence
21
19
II
05
01
Omission
2.3 Hz (E)
2.1 Hz (I)
8

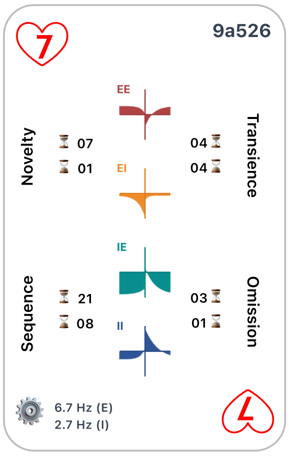
7
9a526
EE
Novelty
07
01
EI
04
04
Transience
IE
Sequence
21
08
II
03
01
Omission
6.7 Hz (E)
2.7 Hz (I)
7

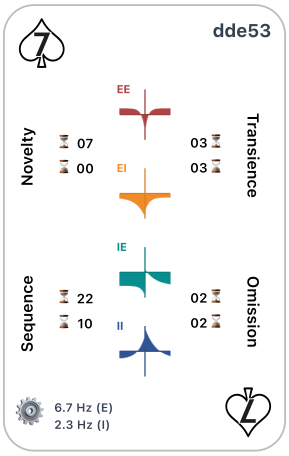
7
dde53
EE
Novelty
07
00
EI
03
03
Transience
IE
Sequence
22
10
II
02
02
Omission
6.7 Hz (E)
2.3 Hz (I)
7

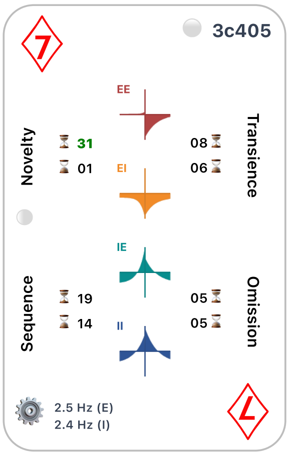
7
3c405
EE
Novelty
31
01
EI
08
06
Transience
IE
Sequence
19
14
II
05
05
Omission
2.5 Hz (E)
2.4 Hz (I)
7

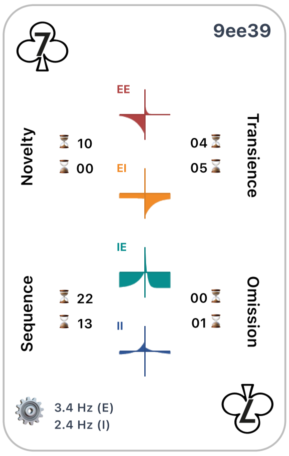
7
9ee39
EE
Novelty
10
00
EI
04
05
Transience
IE
Sequence
22
13
II
00
01
Omission
3.4 Hz (E)
2.4 Hz (I)
7

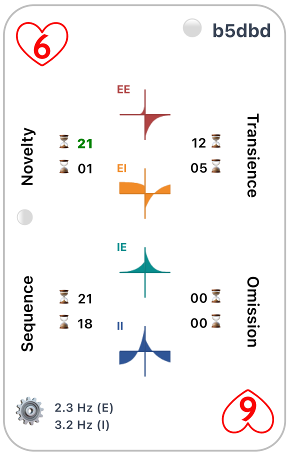
6
b5dbd
EE
Novelty
21
01
EI
12
05
Transience
IE
Sequence
21
18
II
00
00
Omission
2.3 Hz (E)
3.2 Hz (I)
9

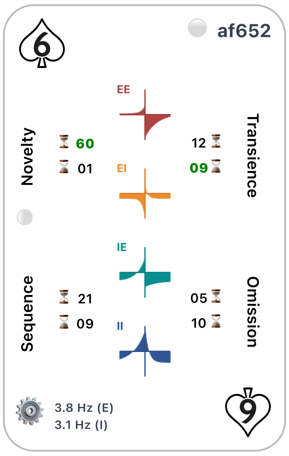
6
af652
EE
Novelty
60
01
EI
12
09
Transience
IE
Sequence
21
09
II
05
10
Omission
3.8 Hz (E)
3.1 Hz (I)
9

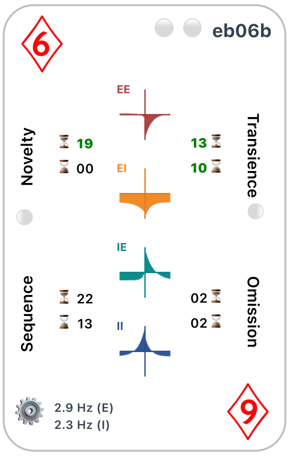
6
eb06b
EE
Novelty
19
00
EI
13
10
Transience
IE
Sequence
22
13
II
02
02
Omission
2.9 Hz (E)
2.3 Hz (I)
9

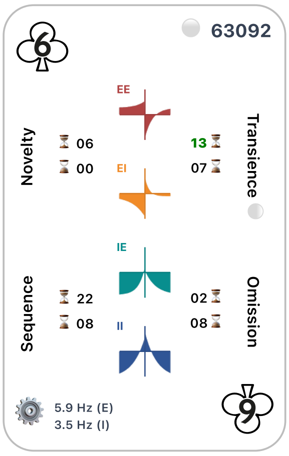
6
63092
EE
Novelty
06
00
EI
13
07
Transience
IE
Sequence
22
08
II
02
08
Omission
5.9 Hz (E)
3.5 Hz (I)
9

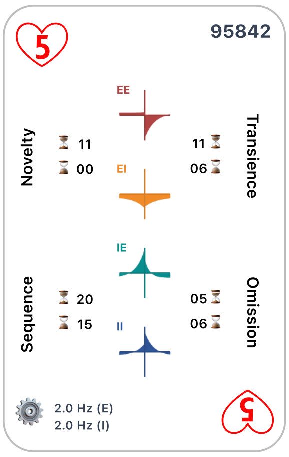
5
95842
EE
EI
IE
II
Novelty 11 00
Transience 11 06
Sequence 20 15
Omission 05 06
2.0 Hz (E)
2.0 Hz (I)

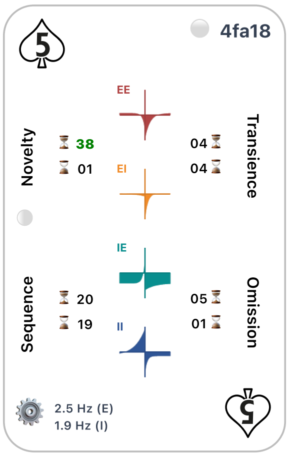
5
4fa18
EE
EI
IE
II
Novelty 38 01
Transience 04 04
Sequence 20 19
Omission 05 01
2.5 Hz (E)
1.9 Hz (I)

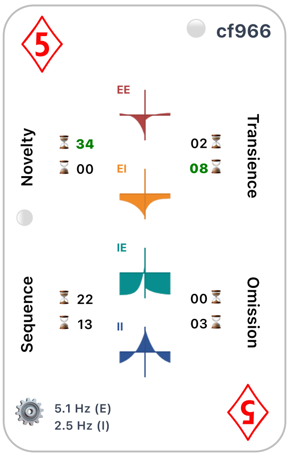
5
cf966
EE
EI
IE
II
Novelty 34 00
Transience 02 08
Sequence 22 13
Omission 00 03
5.1 Hz (E)
2.5 Hz (I)

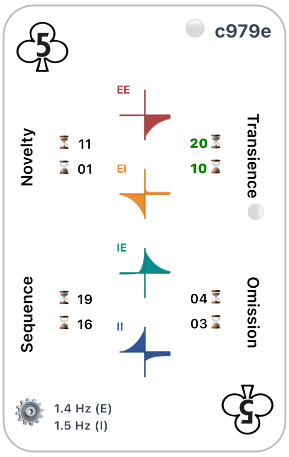
5
c979e
EE
EI
IE
II
Novelty 11 01
Transience 20 10
Sequence 19 16
Omission 04 03
1.4 Hz (E)
1.5 Hz (I)

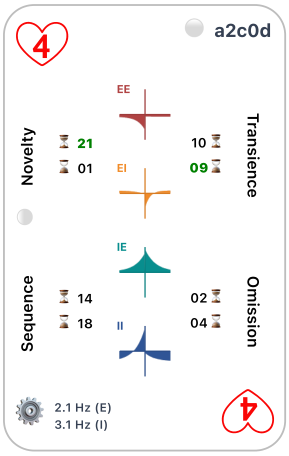
4
a2c0d
EE
EI
IE
II
Novelty 21 01
Transience 10 09
Sequence 14 18
Omission 02 04
2.1 Hz (E)
3.1 Hz (I)

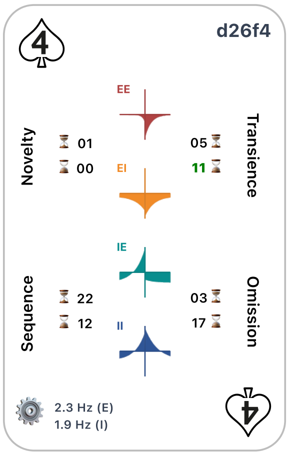
4
d26f4
EE
EI
IE
II
Novelty 01 00
Transience 05 11
Sequence 22 12
Omission 03 17
2.3 Hz (E)
1.9 Hz (I)

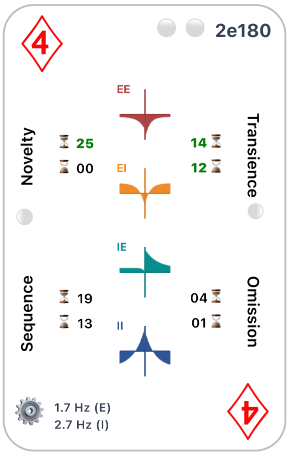
4
2e180
EE
EI
IE
II
Novelty 25 00
Transience 14 12
Sequence 19 13
Omission 04 01
1.7 Hz (E)
2.7 Hz (I)

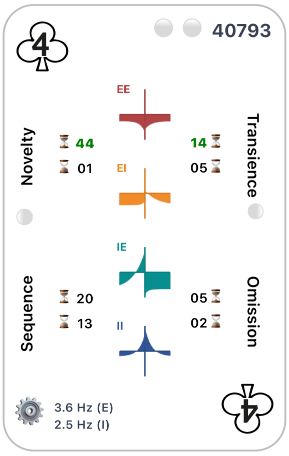
4
40793
EE
EI
IE
II
Novelty 44 01
Transience 14 05
Sequence 20 13
Omission 05 02
3.6 Hz (E)
2.5 Hz (I)

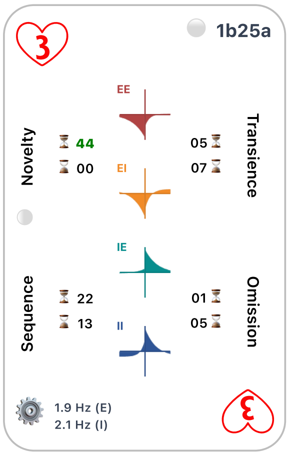
3
1b25a
EE
EI
IE
II
Novelty 44 00
Transience 05 07
Sequence 22 13
Omission 01 05
1.9 Hz (E)
2.1 Hz (I)

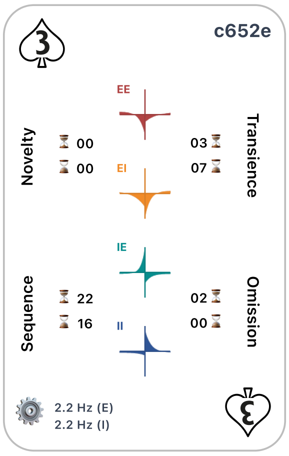
3
c652e
EE
EI
IE
II
Novelty 00 00
Transience 03 07
Sequence 22 16
Omission 02 00
2.2 Hz (E)
2.2 Hz (I)

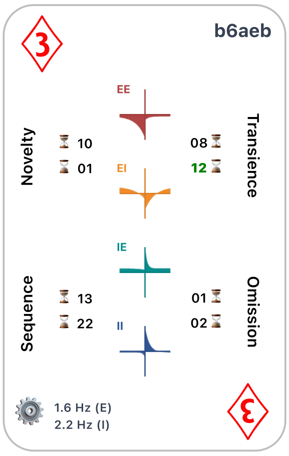
3
b6aeb
EE
EI
IE
II
Novelty 10 01
Transience 08 12
Sequence 13 22
Omission 01 02
1.6 Hz (E)
2.2 Hz (I)

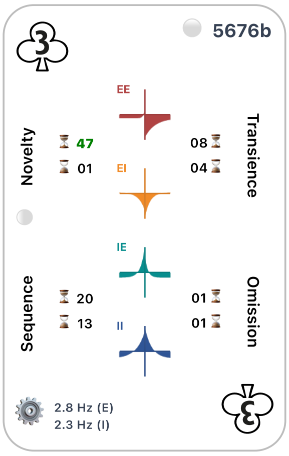
3
5676b
EE
EI
IE
II
Novelty 47 01
Transience 08 04
Sequence 20 13
Omission 01 01
2.8 Hz (E)
2.3 Hz (I)

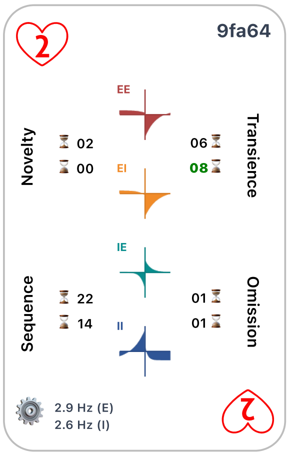
2
9fa64
EE
EI
IE
II
Novelty 02 00
Transience 06 08
Sequence 22 14
Omission 01 01
2.9 Hz (E)
2.6 Hz (I)

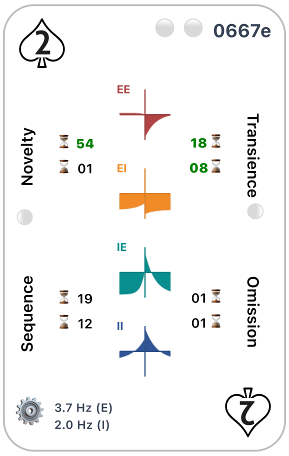
2
0667e
EE
EI
IE
II
Novelty 54 01
Transience 18 08
Sequence 19 12
Omission 01 01
3.7 Hz (E)
2.0 Hz (I)

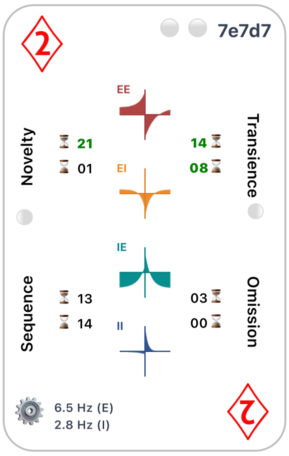
2
7e7d7
EE
EI
IE
II
Novelty 21 01
Transience 14 08
Sequence 13 14
Omission 03 00
6.5 Hz (E)
2.8 Hz (I)

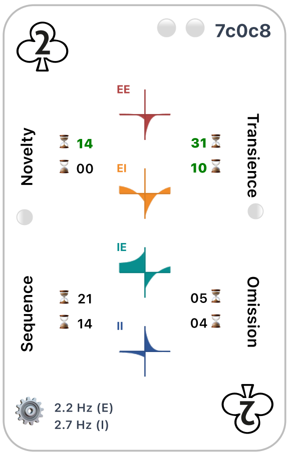
2
7c0c8
EE
EI
IE
II
Novelty 14 00
Transience 31 10
Sequence 21 14
Omission 05 04
2.2 Hz (E)
2.7 Hz (I)

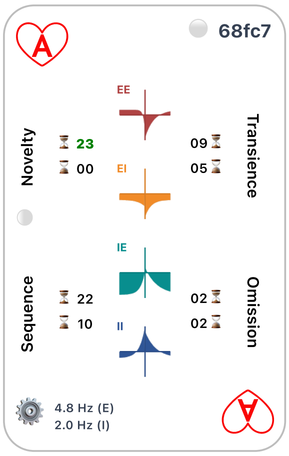
A
68fc7
EE
Novelty
23
00
09
05
Transience
EI
IE
Sequence
22
10
02
02
Omission
II
4.8 Hz (E)
2.0 Hz (I)

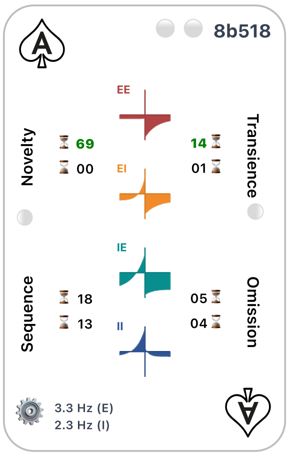
A
8b518
EE
Novelty
69
00
14
01
Transience
EI
IE
Sequence
18
13
05
04
Omission
II
3.3 Hz (E)
2.3 Hz (I)

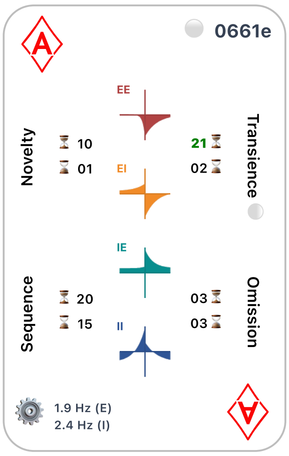
A
0661e
EE
Novelty
10
01
21
02
Transience
EI
IE
Sequence
20
15
03
03
Omission
II
1.9 Hz (E)
2.4 Hz (I)

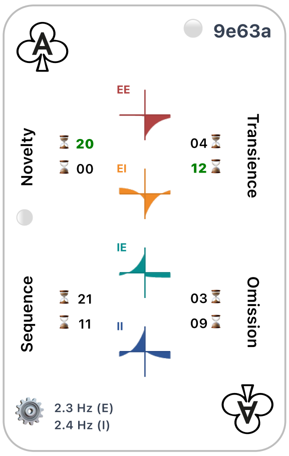
A
9e63a
EE
Novelty
20
00
04
12
Transience
EI
IE
Sequence
21
11
03
09
Omission
II
2.3 Hz (E)
2.4 Hz (I)

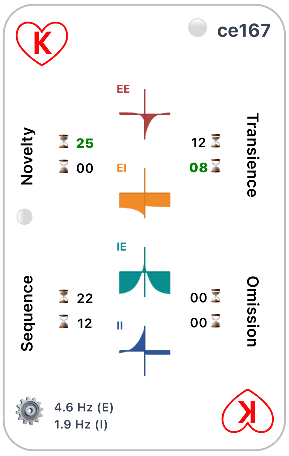
K
ce167
EE
Novelty
25
00
12
08
Transience
EI
IE
Sequence
22
12
00
00
Omission
II
4.6 Hz (E)
1.9 Hz (I)

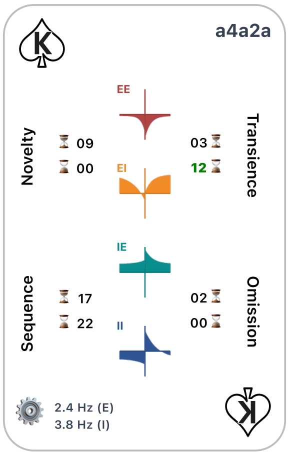
K
a4a2a
EE
Novelty
09
00
03
12
Transience
EI
IE
Sequence
17
22
02
00
Omission
II
2.4 Hz (E)
3.8 Hz (I)

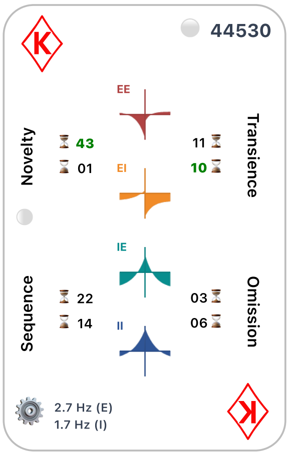
K
44530
EE
Novelty
43
01
11
10
Transience
EI
IE
Sequence
22
14
03
06
Omission
II
2.7 Hz (E)
1.7 Hz (I)

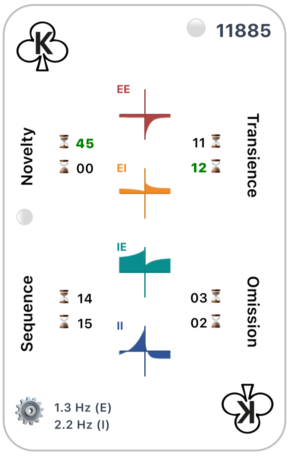
K
11885
EE
Novelty
45
00
11
12
Transience
EI
IE
Sequence
14
15
03
02
Omission
II
1.3 Hz (E)
2.2 Hz (I)

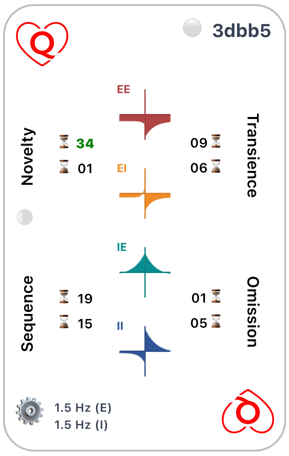
Q
3dbb5
EE
Novelty
34
01
09
06
Transience
EI
IE
Sequence
19
15
01
05
Omission
II
1.5 Hz (E)
1.5 Hz (I)

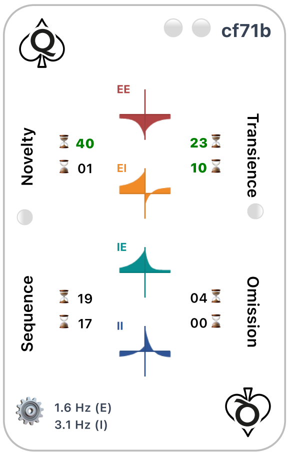
Q
cf71b
EE
Novelty
40
01
23
10
Transience
EI
IE
Sequence
19
17
04
00
Omission
II
1.6 Hz (E)
3.1 Hz (I)

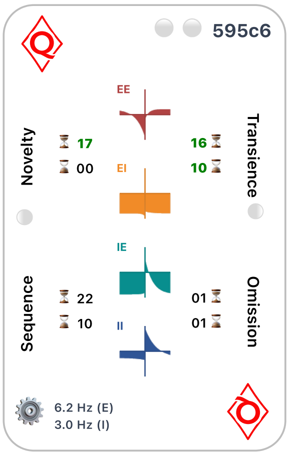
Q
595c6
EE
Novelty
17
00
16
10
Transience
EI
IE
Sequence
22
10
01
01
Omission
II
6.2 Hz (E)
3.0 Hz (I)

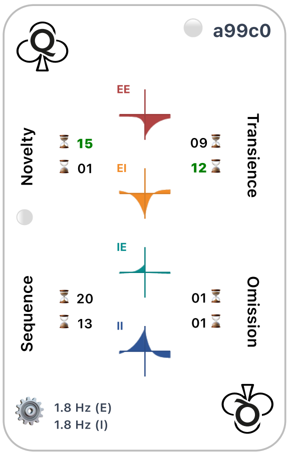
Q
a99c0
EE
Novelty
15
01
09
12
Transience
EI
IE
Sequence
20
13
01
01
Omission
II
1.8 Hz (E)
1.8 Hz (I)

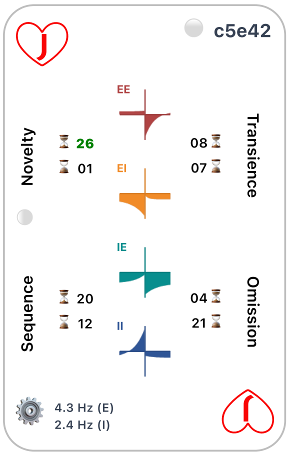
J
c5e42
EE
Novelty
26
01
08
07
Transience
EI
IE
Sequence
20
12
04
21
Omission
II
4.3 Hz (E)
2.4 Hz (I)

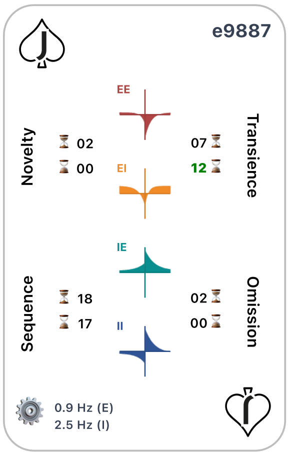
J
e9887
EE
Novelty
02
00
07
12
Transience
EI
IE
Sequence
18
17
02
00
Omission
II
0.9 Hz (E)
2.5 Hz (I)

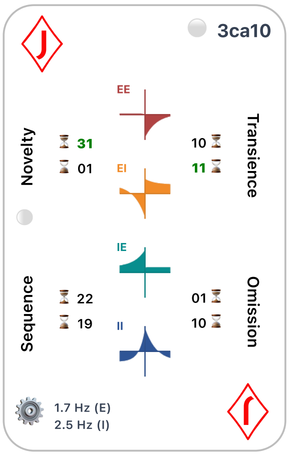
J
3ca10
EE
Novelty
31
01
10
11
Transience
EI
IE
Sequence
22
19
01
10
Omission
II
1.7 Hz (E)
2.5 Hz (I)

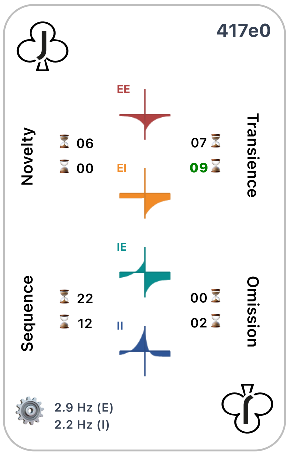
J
417e0
EE
Novelty
06
00
07
09
Transience
EI
IE
Sequence
22
12
00
02
Omission
II
2.9 Hz (E)
2.2 Hz (I)

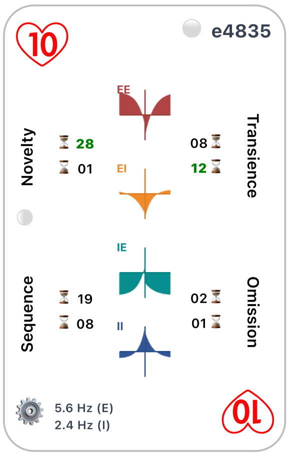
10
e4835
EE
EI
IE
II
Novelty
28
01
Transience
08
12
Sequence
19
08
Omission
02
01
5.6 Hz (E)
2.4 Hz (I)

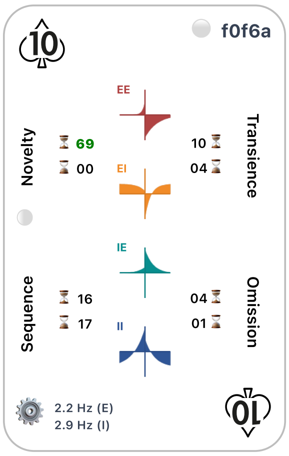
10
f0f6a
EE
EI
IE
II
Novelty
69
00
Transience
10
04
Sequence
16
17
Omission
04
01
2.2 Hz (E)
2.9 Hz (I)

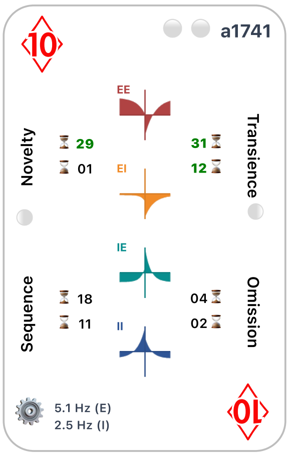
10
a1741
EE
EI
IE
II
Novelty
29
01
Transience
31
12
Sequence
18
11
Omission
04
02
5.1 Hz (E)
2.5 Hz (I)

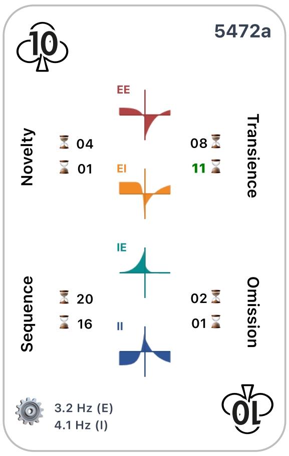
10
5472a
EE
EI
IE
II
Novelty
04
01
Transience
08
11
Sequence
20
16
Omission
02
01
3.2 Hz (E)
4.1 Hz (I)

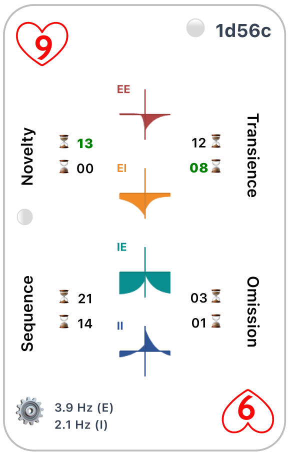
9
1d56c
EE
EI
IE
II
Novelty
13
00
Transience
12
08
Sequence
21
14
Omission
03
01
3.9 Hz (E)
2.1 Hz (I)

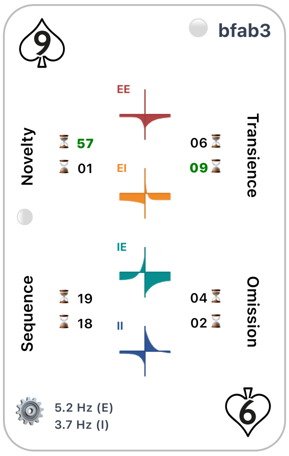
9
bfab3
EE
EI
IE
II
Novelty
57
01
Transience
06
09
Sequence
19
18
Omission
04
02
5.2 Hz (E)
3.7 Hz (I)

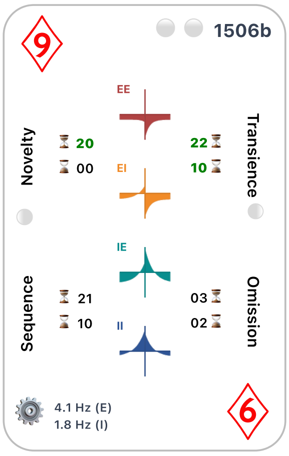
9
1506b
EE
EI
IE
II
Novelty
20
00
Transience
22
10
Sequence
21
10
Omission
03
02
4.1 Hz (E)
1.8 Hz (I)

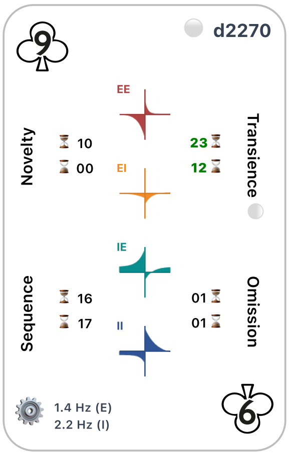
9
d2270
EE
EI
IE
II
Novelty
10
00
Transience
23
12
Sequence
16
17
Omission
01
01
1.4 Hz (E)
2.2 Hz (I)

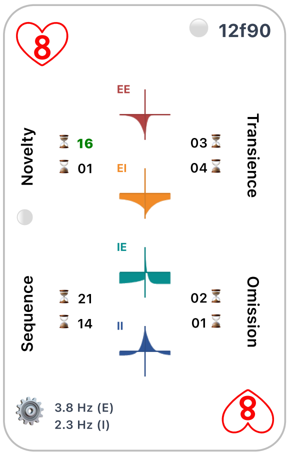
8
12f90
EE
EI
IE
II
Novelty
16
01
Transience
03
04
Sequence
21
14
Omission
02
01
3.8 Hz (E)
2.3 Hz (I)

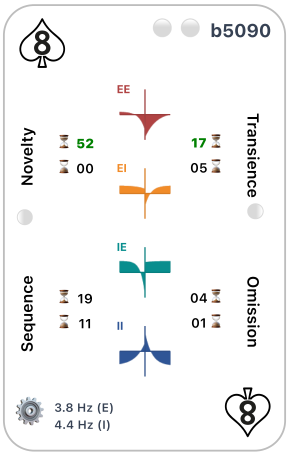
8
b5090
EE
EI
IE
II
Novelty
52
00
Transience
17
05
Sequence
19
11
Omission
04
01
3.8 Hz (E)
4.4 Hz (I)

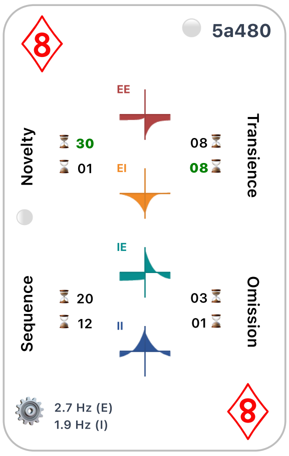
8
5a480
EE
EI
IE
II
Novelty
30
01
Transience
08
08
Sequence
20
12
Omission
03
01
2.7 Hz (E)
1.9 Hz (I)

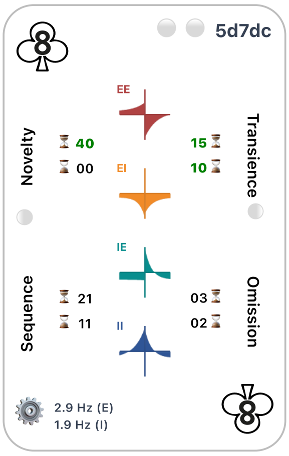
8
5d7dc
EE
EI
IE
II
Novelty
40
00
Transience
15
10
Sequence
21
11
Omission
03
02
2.9 Hz (E)
1.9 Hz (I)

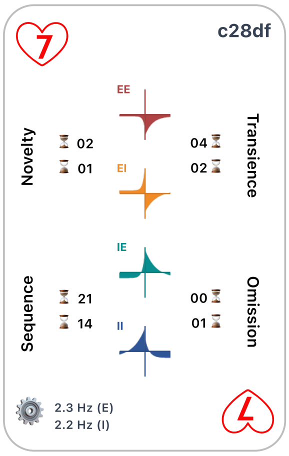
7
c28df
EE
EI
IE
II
Novelty
02
01
Transience
04
02
Sequence
21
14
Omission
00
01
2.3 Hz (E)
2.2 Hz (I)

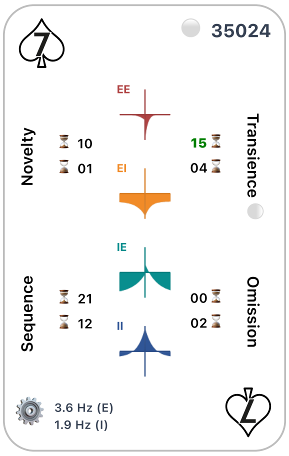
7
35024
EE
EI
IE
II
Novelty
10
01
Transience
15
04
Sequence
21
12
Omission
00
02
3.6 Hz (E)
1.9 Hz (I)

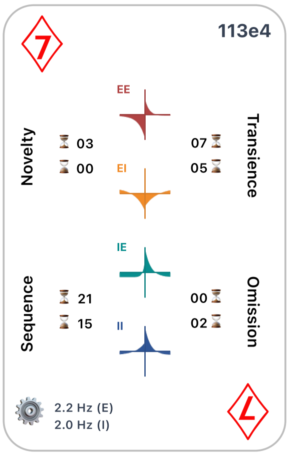
7
113e4
EE
EI
IE
II
Novelty
03
00
Transience
07
05
Sequence
21
15
Omission
00
02
2.2 Hz (E)
2.0 Hz (I)

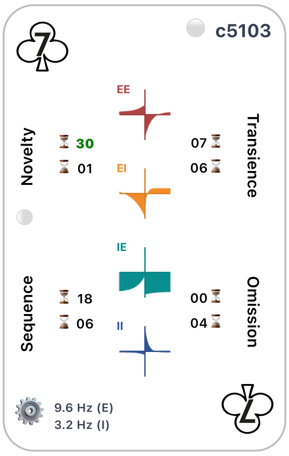
7
c5103
EE
EI
IE
II
Novelty
30
01
Transience
07
06
Sequence
18
06
Omission
00
04
9.6 Hz (E)
3.2 Hz (I)

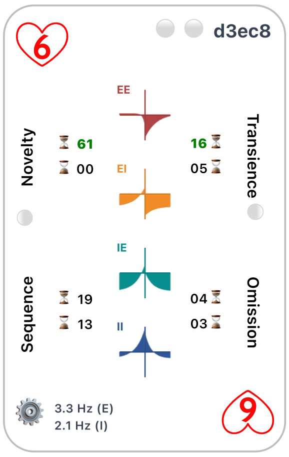
6
d3ec8
EE
EI
IE
II
Novelty
61
00
Transience
16
05
Sequence
19
13
Omission
04
03
3.3 Hz (E)
2.1 Hz (I)

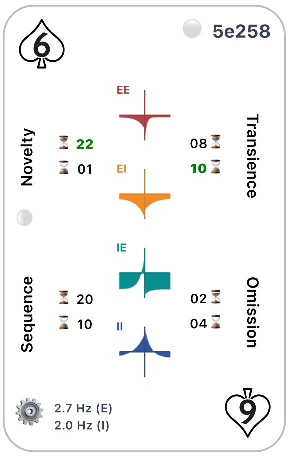
6
5e258
EE
EI
IE
II
Novelty
22
01
Transience
08
10
Sequence
20
10
Omission
02
04
2.7 Hz (E)
2.0 Hz (I)

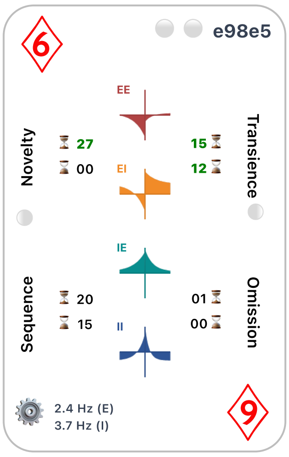
6
e98e5
EE
EI
IE
II
Novelty
27
00
Transience
15
12
Sequence
20
15
Omission
01
00
2.4 Hz (E)
3.7 Hz (I)

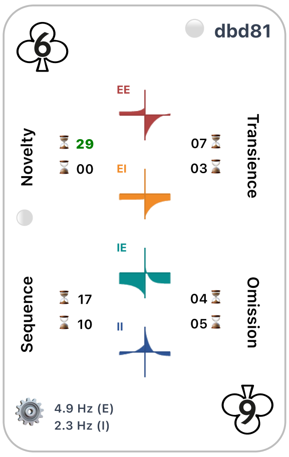
6
dbd81
EE
EI
IE
II
Novelty
29
00
Transience
07
03
Sequence
17
10
Omission
04
05
4.9 Hz (E)
2.3 Hz (I)

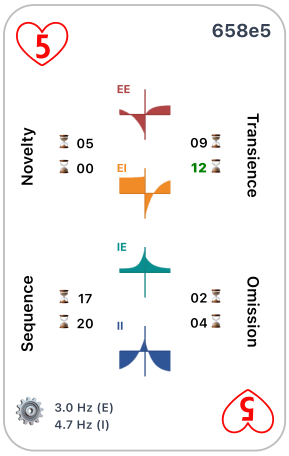
5
658e5
EE
EI
IE
II
Novelty
05
00
Transience
09
12
Sequence
17
20
Omission
02
04
3.0 Hz (E)
4.7 Hz (I)

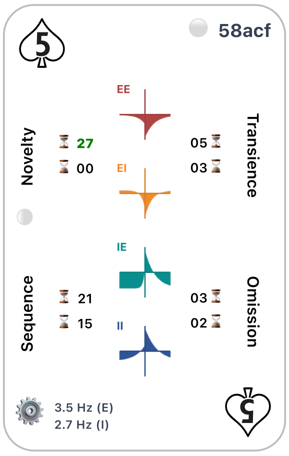
5
58acf
EE
EI
IE
II
Novelty
27
00
Transience
05
03
Sequence
21
15
Omission
03
02
3.5 Hz (E)
2.7 Hz (I)

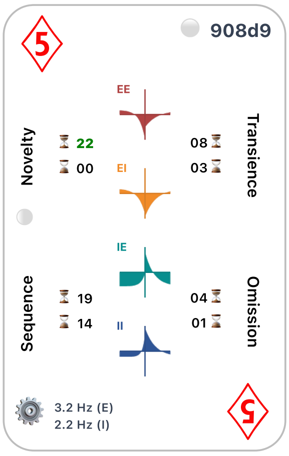
5
908d9
EE
EI
IE
II
Novelty
22
00
Transience
08
03
Sequence
19
14
Omission
04
01
3.2 Hz (E)
2.2 Hz (I)

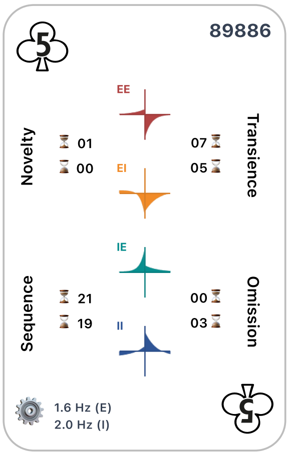
5
89886
EE
EI
IE
II
Novelty
01
00
Transience
07
05
Sequence
21
19
Omission
00
03
1.6 Hz (E)
2.0 Hz (I)

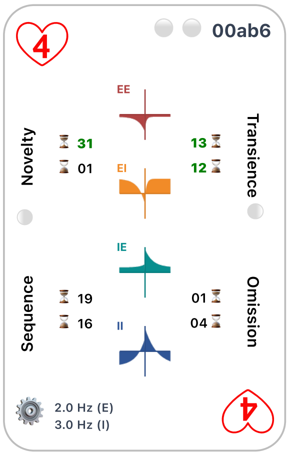
4
00ab6
EE
EI
IE
II
Novelty
31
01
Transience
13
12
Sequence
19
16
Omission
01
04
2.0 Hz (E)
3.0 Hz (I)

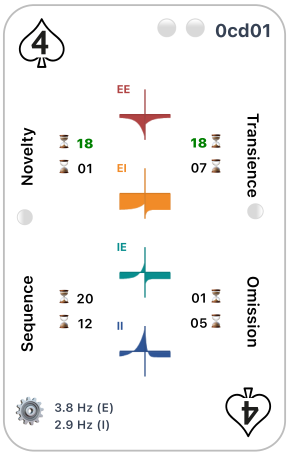
4
0cd01
EE
EI
IE
II
Novelty
18
01
Transience
18
07
Sequence
20
12
Omission
01
05
3.8 Hz (E)
2.9 Hz (I)

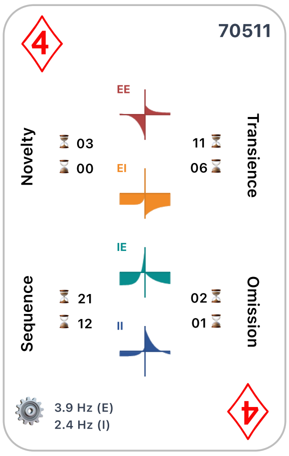
4
70511
EE
EI
IE
II
Novelty
03
00
Transience
11
06
Sequence
21
12
Omission
02
01
3.9 Hz (E)
2.4 Hz (I)

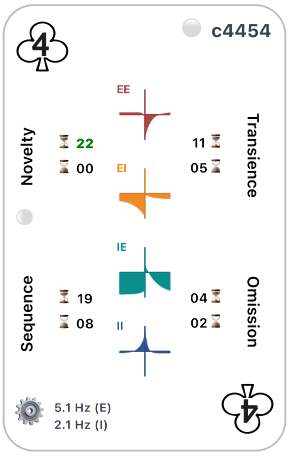
4
c4454
EE
EI
IE
II
Novelty
22
00
Transience
11
05
Sequence
19
08
Omission
04
02
5.1 Hz (E)
2.1 Hz (I)

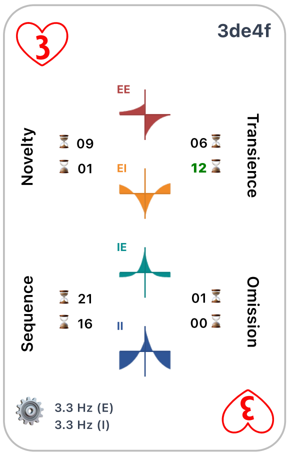
3
3de4f
EE
EI
IE
II
Novelty
09
01
Transience
06
12
Sequence
21
16
Omission
01
00
3.3 Hz (E)
3.3 Hz (I)

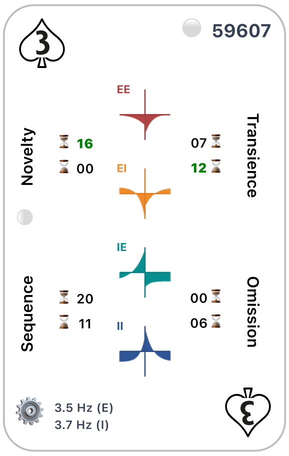
3
59607
EE
EI
IE
II
Novelty
16
00
Transience
07
12
Sequence
20
11
Omission
00
06
3.5 Hz (E)
3.7 Hz (I)

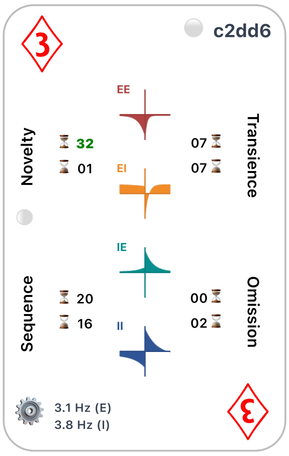
3
c2dd6
EE
EI
IE
II
Novelty
32
01
Transience
07
07
Sequence
20
16
Omission
00
02
3.1 Hz (E)
3.8 Hz (I)

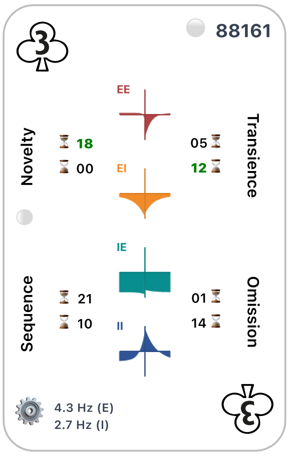
3
88161
EE
EI
IE
II
Novelty
18
00
Transience
05
12
Sequence
21
10
Omission
01
14
4.3 Hz (E)
2.7 Hz (I)

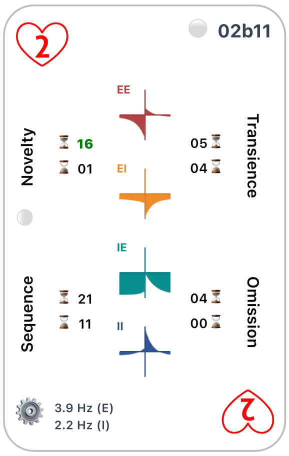
2
02b11
Novelty
16
01
Transience
05
04
EE
EI
IE
II
Sequence
21
11
Omission
04
00
3.9 Hz (E)
2.2 Hz (I)

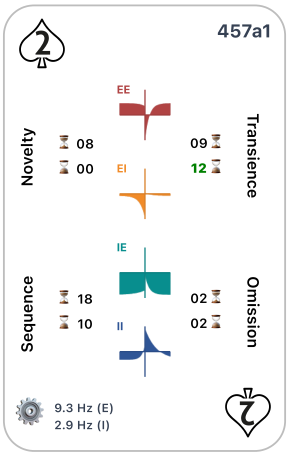
2
457a1
Novelty
08
00
Transience
09
12
EE
EI
IE
II
Sequence
18
10
Omission
02
02
9.3 Hz (E)
2.9 Hz (I)

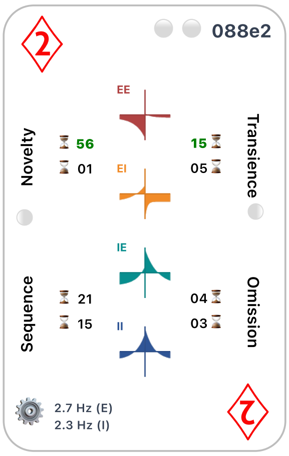
2
088e2
Novelty
56
01
Transience
15
05
EE
EI
IE
II
Sequence
21
15
Omission
04
03
2.7 Hz (E)
2.3 Hz (I)

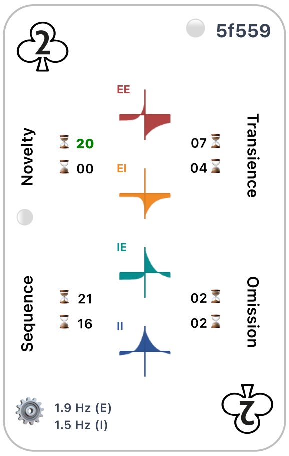
2
5f559
Novelty
20
00
Transience
07
04
EE
EI
IE
II
Sequence
21
16
Omission
02
02
1.9 Hz (E)
1.5 Hz (I)

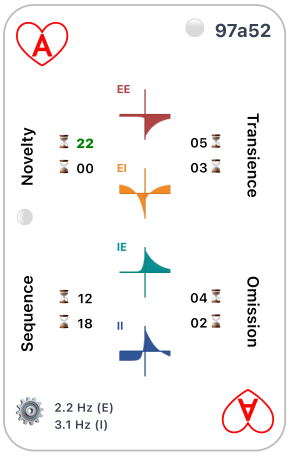
A
97a52
Novelty
22
00
Transience
05
03
EE
EI
IE
II
Sequence
12
18
Omission
04
02
2.2 Hz (E)
3.1 Hz (I)

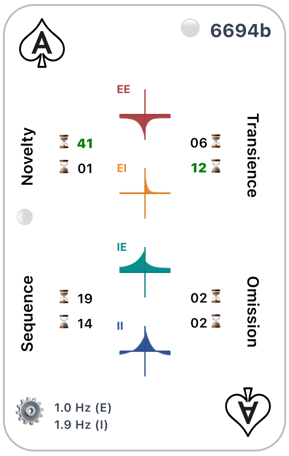
A
6694b
Novelty
41
01
Transience
06
12
EE
EI
IE
II
Sequence
19
14
Omission
02
02
1.0 Hz (E)
1.9 Hz (I)

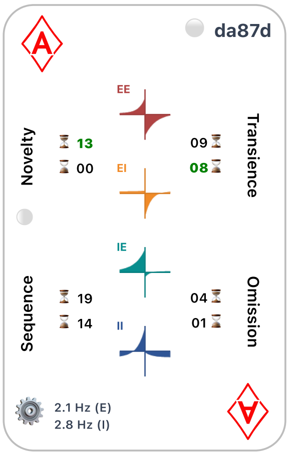
A
da87d
Novelty
13
00
Transience
09
08
EE
EI
IE
II
Sequence
19
14
Omission
04
01
2.1 Hz (E)
2.8 Hz (I)

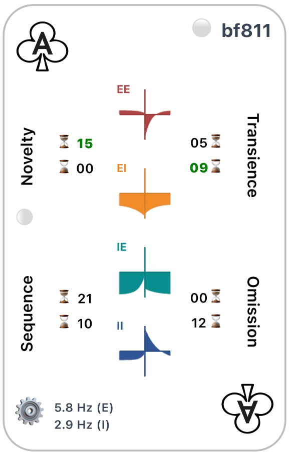
A
bf811
Novelty
15
00
Transience
05
09
EE
EI
IE
II
Sequence
21
10
Omission
00
12
5.8 Hz (E)
2.9 Hz (I)

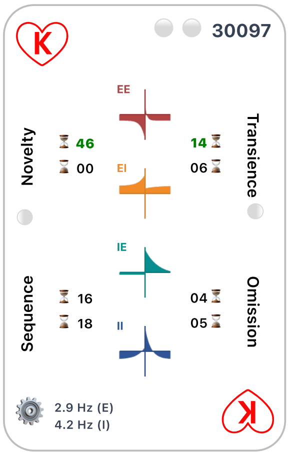
K
30097
Novelty
46
00
Transience
14
06
EE
EI
IE
II
Sequence
16
18
Omission
04
05
2.9 Hz (E)
4.2 Hz (I)

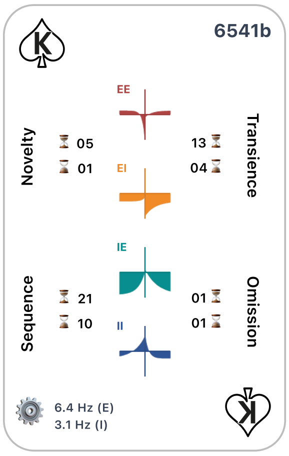
K
6541b
Novelty
05
01
Transience
13
04
EE
EI
IE
II
Sequence
21
10
Omission
01
01
6.4 Hz (E)
3.1 Hz (I)

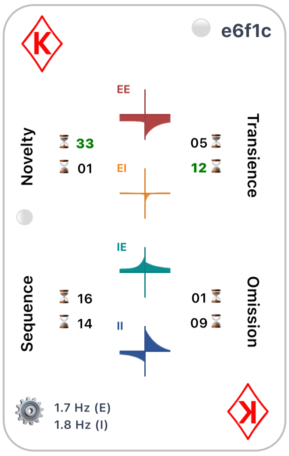
K
e6f1c
Novelty
33
01
Transience
05
12
EE
EI
IE
II
Sequence
16
14
Omission
01
09
1.7 Hz (E)
1.8 Hz (I)

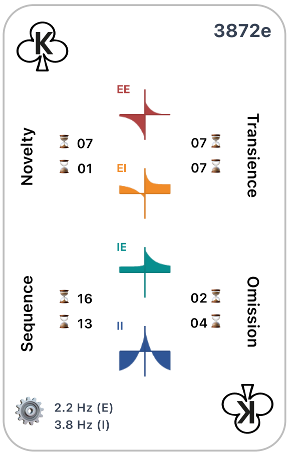
K
3872e
Novelty
07
01
Transience
07
07
EE
EI
IE
II
Sequence
16
13
Omission
02
04
2.2 Hz (E)
3.8 Hz (I)

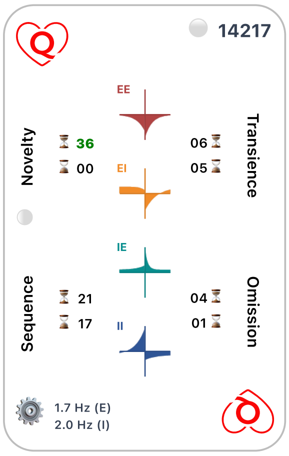
Q
14217
Novelty
36
00
Transience
06
05
EE
EI
IE
II
Sequence
21
17
Omission
04
01
1.7 Hz (E)
2.0 Hz (I)

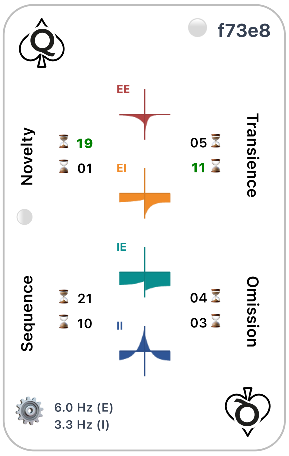
Q
f73e8
Novelty
19
01
Transience
05
11
EE
EI
IE
II
Sequence
21
10
Omission
04
03
6.0 Hz (E)
3.3 Hz (I)

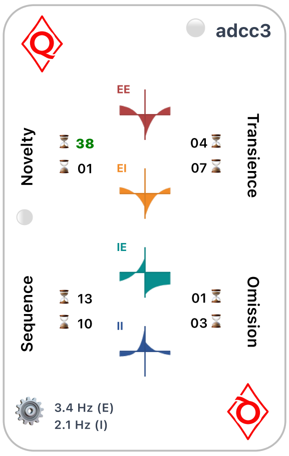
Q
adcc3
Novelty
38
01
Transience
04
07
EE
EI
IE
II
Sequence
13
10
Omission
01
03
3.4 Hz (E)
2.1 Hz (I)

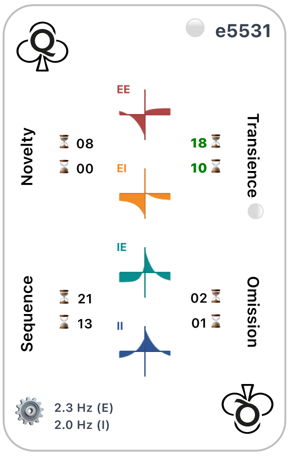
Q
e5531
Novelty
08
00
Transience
18
10
EE
EI
IE
II
Sequence
21
13
Omission
02
01
2.3 Hz (E)
2.0 Hz (I)

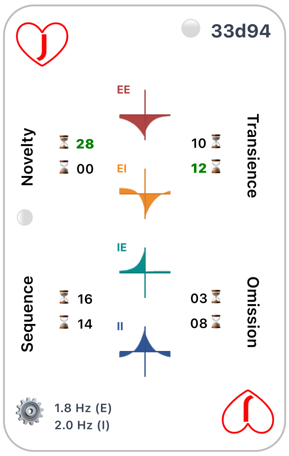
J
33d94
EE
EI
IE
II
Novelty
28
00
Transience
10
12
Sequence
16
14
Omission
03
08
1.8 Hz (E)
2.0 Hz (I)

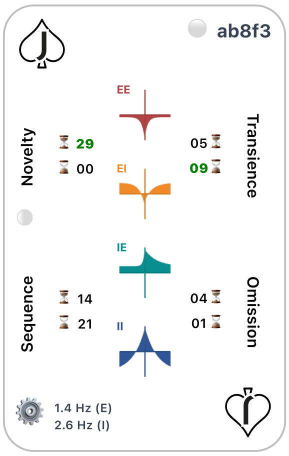
J
ab8f3
EE
EI
IE
II
Novelty
29
00
Transience
05
09
Sequence
14
21
Omission
04
01
1.4 Hz (E)
2.6 Hz (I)

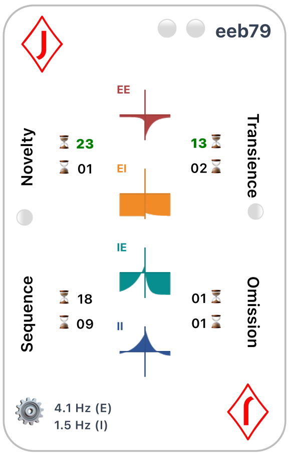
J
eeb79
EE
EI
IE
II
Novelty
23
01
Transience
13
02
Sequence
18
09
Omission
01
01
4.1 Hz (E)
1.5 Hz (I)

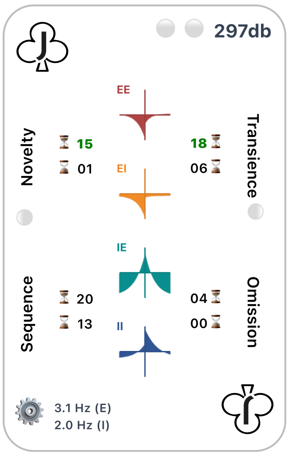
J
297db
EE
EI
IE
II
Novelty
15
01
Transience
18
06
Sequence
20
13
Omission
04
00
3.1 Hz (E)
2.0 Hz (I)

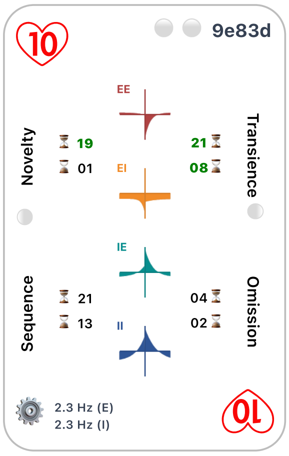
10
9e83d
EE
EI
IE
II
Novelty
19
01
Transience
21
08
Sequence
21
13
Omission
04
02
2.3 Hz (E)
2.3 Hz (I)

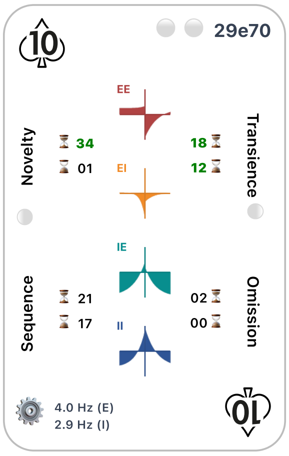
10
29e70
EE
EI
IE
II
Novelty
34
01
Transience
18
12
Sequence
21
17
Omission
02
00
4.0 Hz (E)
2.9 Hz (I)

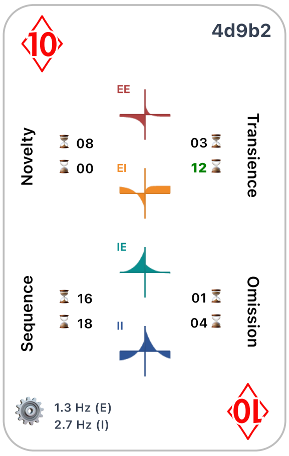
10
4d9b2
EE
EI
IE
II
Novelty
08
00
Transience
03
12
Sequence
16
18
Omission
01
04
1.3 Hz (E)
2.7 Hz (I)

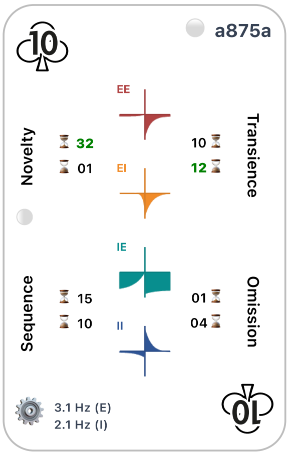
10
a875a
EE
EI
IE
II
Novelty
32
01
Transience
10
12
Sequence
15
10
Omission
01
04
3.1 Hz (E)
2.1 Hz (I)

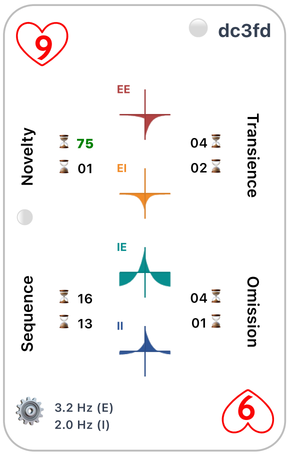
9
dc3fd
EE
EI
IE
II
Novelty
75
01
Transience
04
02
Sequence
16
13
Omission
04
01
3.2 Hz (E)
2.0 Hz (I)

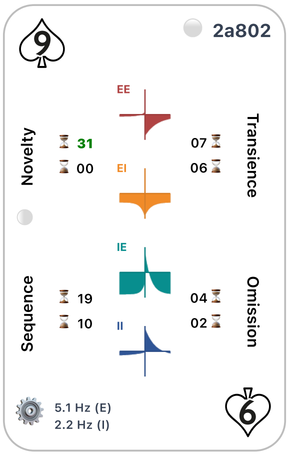
9
2a802
EE
EI
IE
II
Novelty
31
00
Transience
07
06
Sequence
19
10
Omission
04
02
5.1 Hz (E)
2.2 Hz (I)

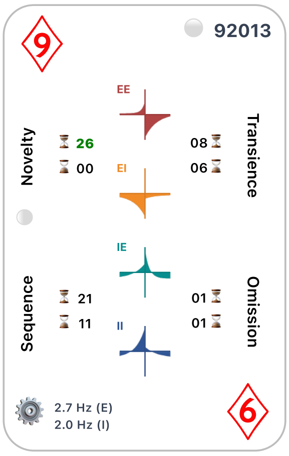
9
92013
EE
EI
IE
II
Novelty
26
00
Transience
08
06
Sequence
21
11
Omission
01
01
2.7 Hz (E)
2.0 Hz (I)

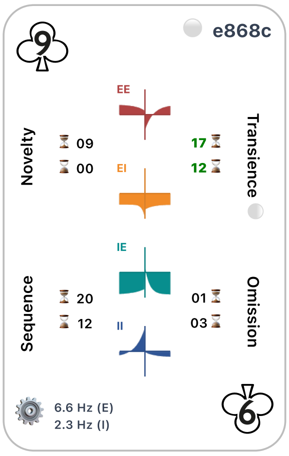
9
e868c
EE
EI
IE
II
Novelty
09
00
Transience
17
12
Sequence
20
12
Omission
01
03
6.6 Hz (E)
2.3 Hz (I)

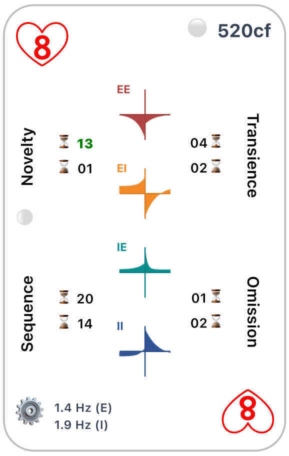
8
520cf
EE
EI
IE
II
Novelty
13
01
Transience
04
02
Sequence
20
14
Omission
01
02
1.4 Hz (E)
1.9 Hz (I)

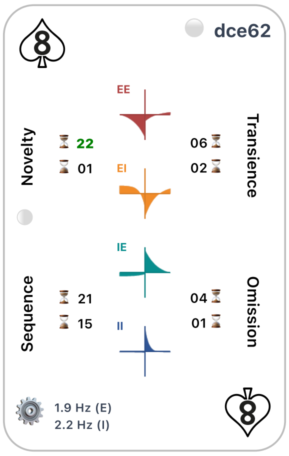
8
dce62
EE
EI
IE
II
Novelty
22
01
Transience
06
02
Sequence
21
15
Omission
04
01
1.9 Hz (E)
2.2 Hz (I)

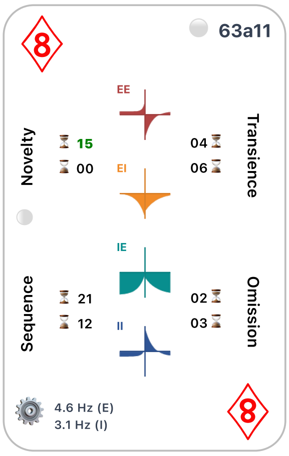
8
63a11
EE
EI
IE
II
Novelty
15
00
Transience
04
06
Sequence
21
12
Omission
02
03
4.6 Hz (E)
3.1 Hz (I)

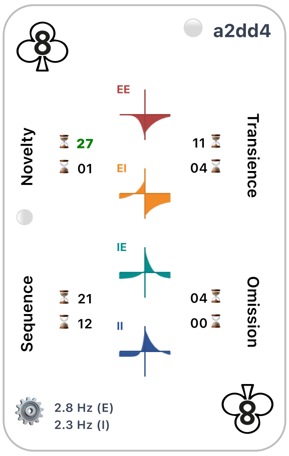
8
a2dd4
EE
EI
IE
II
Novelty
27
01
Transience
11
04
Sequence
21
12
Omission
04
00
2.8 Hz (E)
2.3 Hz (I)

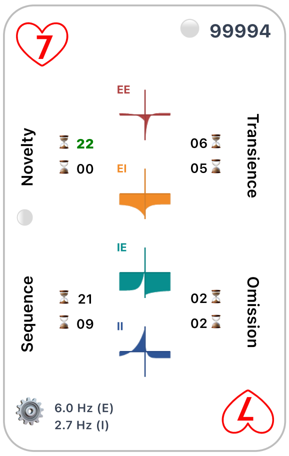
7
99994
EE
EI
IE
II
Novelty
22
00
Transience
06
05
Sequence
21
09
Omission
02
02
6.0 Hz (E)
2.7 Hz (I)
7

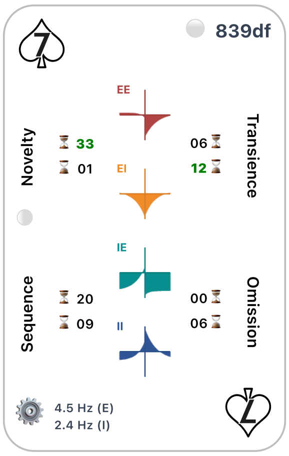
7
839df
EE
EI
IE
II
Novelty
33
01
Transience
06
12
Sequence
20
09
Omission
00
06
4.5 Hz (E)
2.4 Hz (I)
7

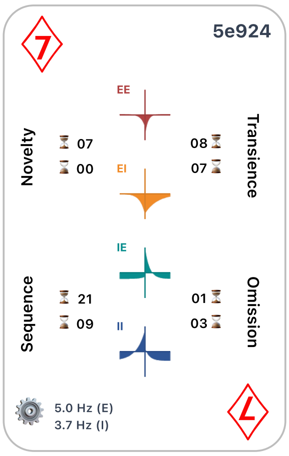
7
5e924
EE
EI
IE
II
Novelty
07
00
Transience
08
07
Sequence
21
09
Omission
01
03
5.0 Hz (E)
3.7 Hz (I)
7

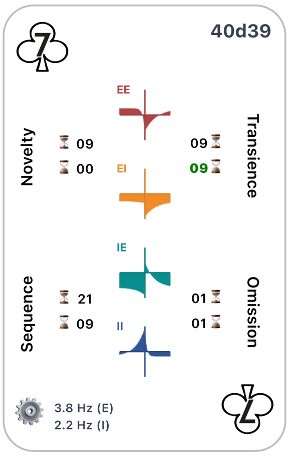
7
40d39
EE
EI
IE
II
Novelty
09
00
Transience
09
09
Sequence
21
09
Omission
01
01
3.8 Hz (E)
2.2 Hz (I)
7

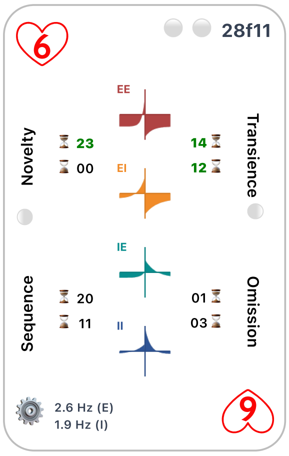
6
28f11
EE
EI
IE
II
Novelty
23
00
Transience
14
12
Sequence
20
11
Omission
01
03
2.6 Hz (E)
1.9 Hz (I)
6

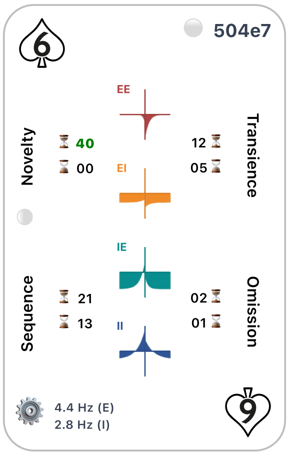
6
504e7
EE
EI
IE
II
Novelty
40
00
Transience
12
05
Sequence
21
13
Omission
02
01
4.4 Hz (E)
2.8 Hz (I)
6

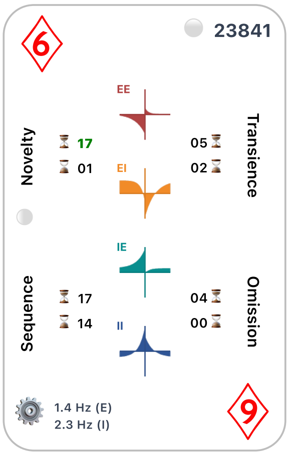
6
23841
EE
EI
IE
II
Novelty
17
01
Transience
05
02
Sequence
17
14
Omission
04
00
1.4 Hz (E)
2.3 Hz (I)
6

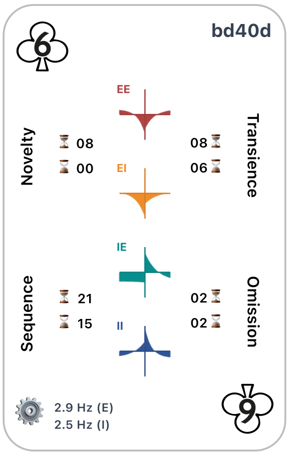
6
bd40d
EE
EI
IE
II
Novelty
08
00
Transience
08
06
Sequence
21
15
Omission
02
02
2.9 Hz (E)
2.5 Hz (I)
6

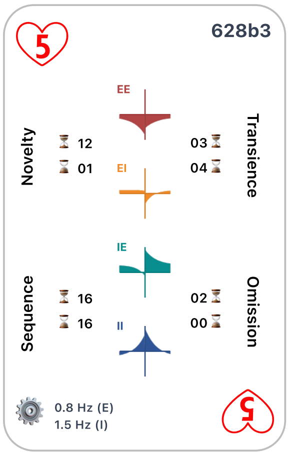
5
628b3
EE
EI
IE
II
Novelty
12
01
Transience
03
04
Sequence
16
16
Omission
02
00
0.8 Hz (E)
1.5 Hz (I)
5

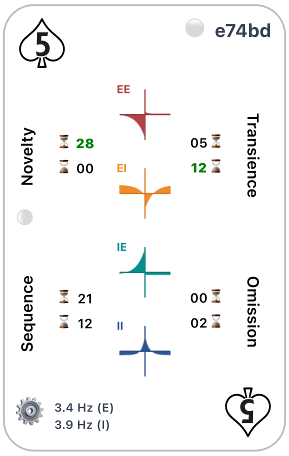
5
e74bd
EE
EI
IE
II
Novelty
28
00
Transience
05
12
Sequence
21
12
Omission
00
02
3.4 Hz (E)
3.9 Hz (I)
5

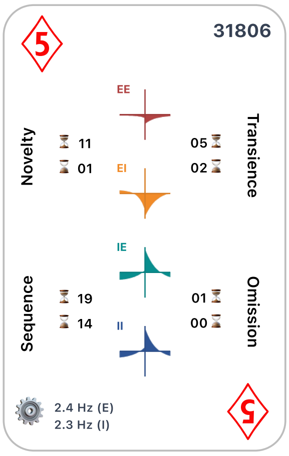
5
31806
EE
EI
IE
II
Novelty
11
01
Transience
05
02
Sequence
19
14
Omission
01
00
2.4 Hz (E)
2.3 Hz (I)
5

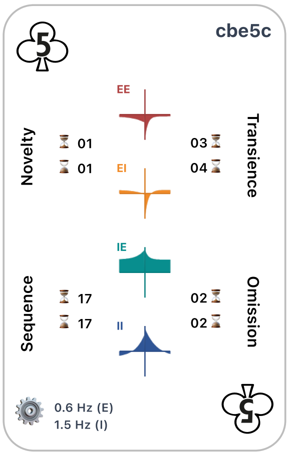
5
cbe5c
EE
EI
IE
II
Novelty
01
01
Transience
03
04
Sequence
17
17
Omission
02
02
0.6 Hz (E)
1.5 Hz (I)
5

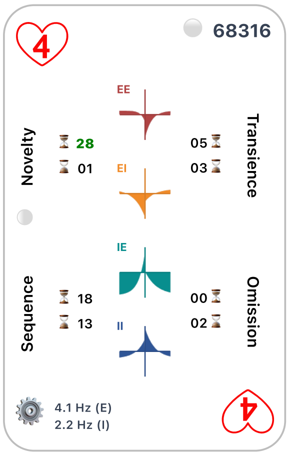
4
68316
EE
EI
IE
II
Novelty
28
01
Transience
05
03
Sequence
18
13
Omission
00
02
4.1 Hz (E)
2.2 Hz (I)
4

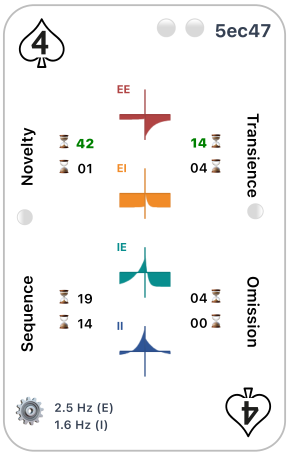
4
5ec47
EE
EI
IE
II
Novelty
42
01
Transience
14
04
Sequence
19
14
Omission
04
00
2.5 Hz (E)
1.6 Hz (I)
4

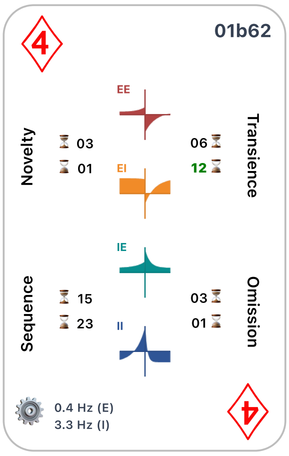
4
01b62
EE
EI
IE
II
Novelty
03
01
Transience
06
12
Sequence
15
23
Omission
03
01
0.4 Hz (E)
3.3 Hz (I)
4

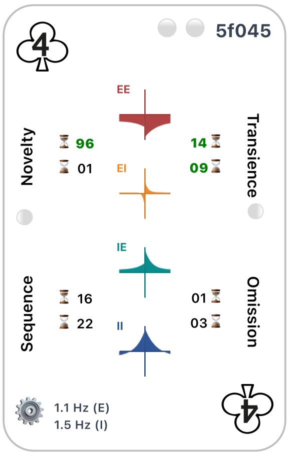
4
5f045
EE
EI
IE
II
Novelty
96
01
Transience
14
09
Sequence
16
22
Omission
01
03
1.1 Hz (E)
1.5 Hz (I)
4

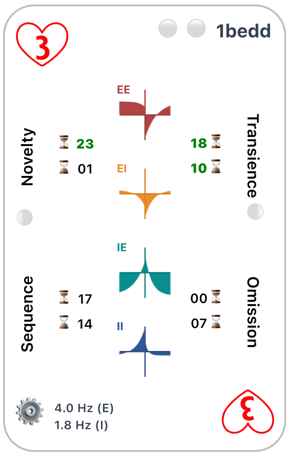
3
1bedd
Novelty
23
01
EE
EI
18
10
Transience
Sequence
17
14
IE
II
00
07
Omission
4.0 Hz (E)
1.8 Hz (I)
3

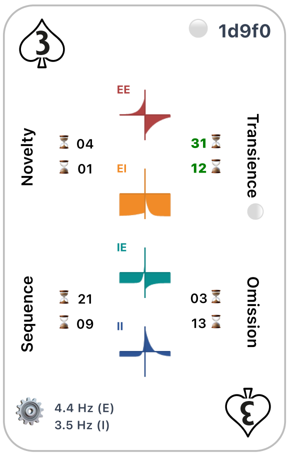
3
1d9f0
Novelty
04
01
EE
EI
31
12
Transience
Sequence
21
09
IE
II
03
13
Omission
4.4 Hz (E)
3.5 Hz (I)
3

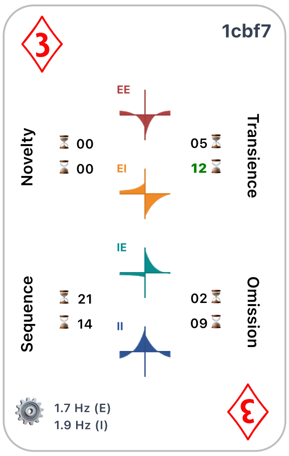
3
1cbf7
Novelty
00
00
EE
EI
05
12
Transience
Sequence
21
14
IE
II
02
09
Omission
1.7 Hz (E)
1.9 Hz (I)
3

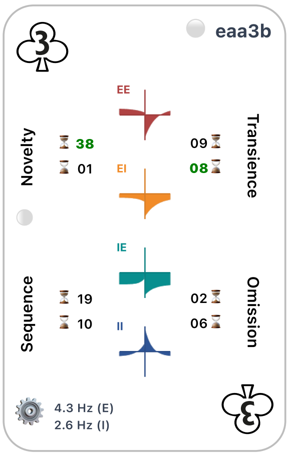
3
eaa3b
Novelty
38
01
EE
EI
09
08
Transience
Sequence
19
10
IE
II
02
06
Omission
4.3 Hz (E)
2.6 Hz (I)
3

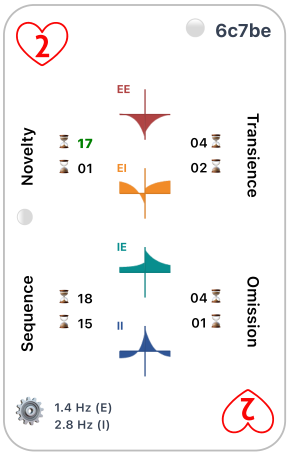
2
6c7be
Novelty
17
01
EE
EI
04
02
Transience
Sequence
18
15
IE
II
04
01
Omission
1.4 Hz (E)
2.8 Hz (I)
2

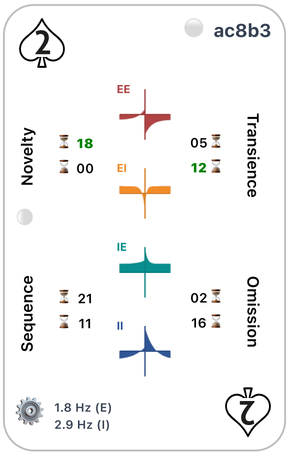
2
ac8b3
Novelty
18
00
EE
EI
05
12
Transience
Sequence
21
11
IE
II
02
16
Omission
1.8 Hz (E)
2.9 Hz (I)
2

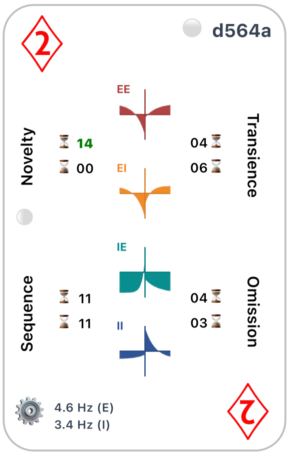
2
d564a
Novelty
14
00
EE
EI
04
06
Transience
Sequence
11
11
IE
II
04
03
Omission
4.6 Hz (E)
3.4 Hz (I)
2

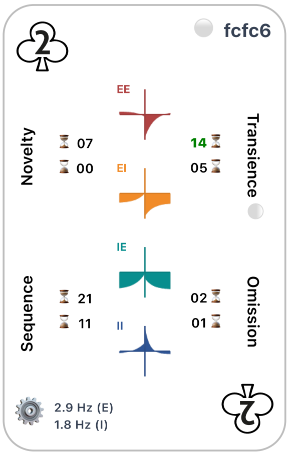
2
fcfc6
Novelty
07
00
EE
EI
14
05
Transience
Sequence
21
11
IE
II
02
01
Omission
2.9 Hz (E)
1.8 Hz (I)
2

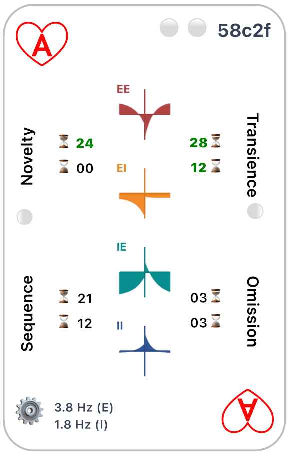
A
58c2f
Novelty
24
00
EE
EI
28
12
Transience
Sequence
21
12
IE
II
03
03
Omission
3.8 Hz (E)
1.8 Hz (I)
A

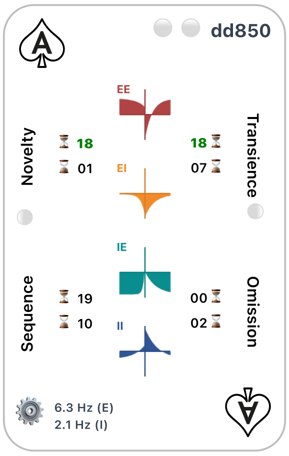
A
dd850
Novelty
18
01
EE
EI
18
07
Transience
Sequence
19
10
IE
II
00
02
Omission
6.3 Hz (E)
2.1 Hz (I)
A

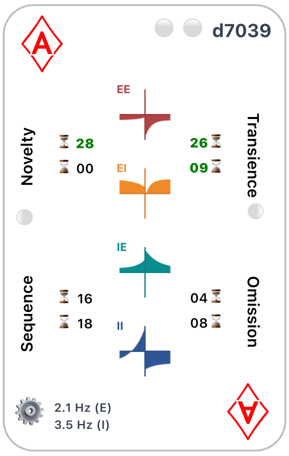
A
d7039
Novelty
28
00
EE
EI
26
09
Transience
Sequence
16
18
IE
II
04
08
Omission
2.1 Hz (E)
3.5 Hz (I)
A

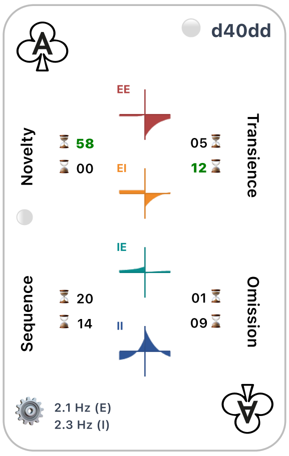
A
d40dd
Novelty
58
00
EE
EI
05
12
Transience
Sequence
20
14
IE
II
01
09
Omission
2.1 Hz (E)
2.3 Hz (I)
A

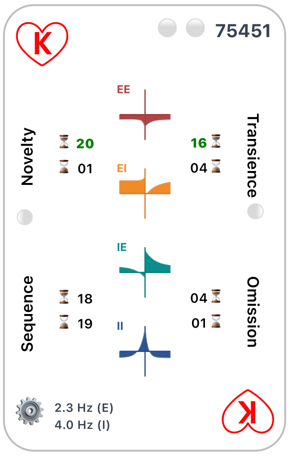
K
75451
Novelty
20
01
EE
EI
16
04
Transience
Sequence
18
19
IE
II
04
01
Omission
2.3 Hz (E)
4.0 Hz (I)
K

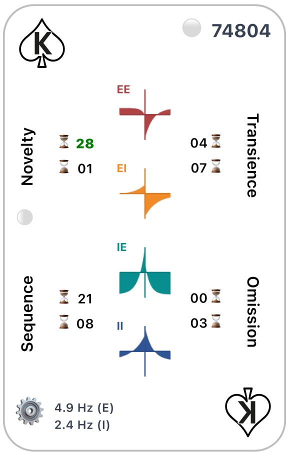
K
74804
Novelty
28
01
EE
EI
04
07
Transience
Sequence
21
08
IE
II
00
03
Omission
4.9 Hz (E)
2.4 Hz (I)
K

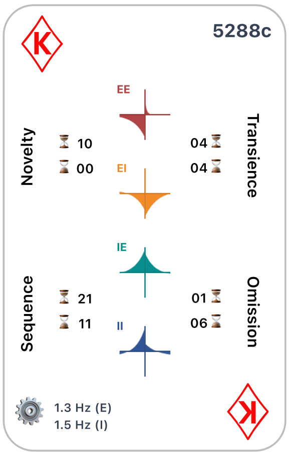
K
5288c
Novelty
10
00
EE
EI
04
04
Transience
Sequence
21
11
IE
II
01
06
Omission
1.3 Hz (E)
1.5 Hz (I)
K

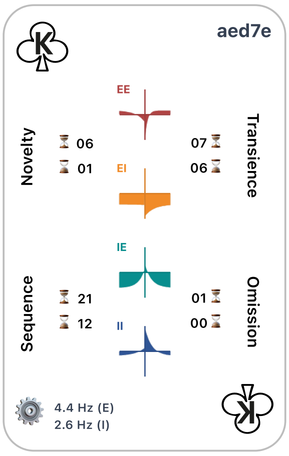
K
aed7e
Novelty
06
01
EE
EI
07
06
Transience
Sequence
21
12
IE
II
01
00
Omission
4.4 Hz (E)
2.6 Hz (I)
K

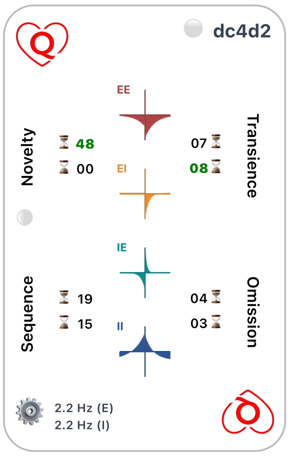
dc4d2
EE
EI
IE
II
Novelty
48
00
Transience
07
08
Sequence
19
15
Omission
04
03
2.2 Hz (E)
2.2 Hz (I)

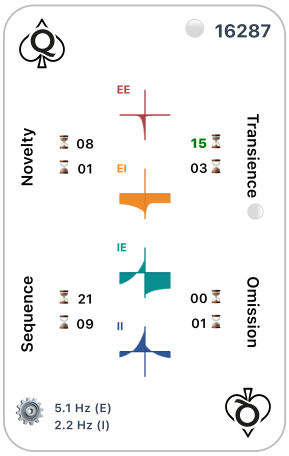
16287
EE
EI
IE
II
Novelty
08
01
Transience
15
03
Sequence
21
09
Omission
00
01
5.1 Hz (E)
2.2 Hz (I)

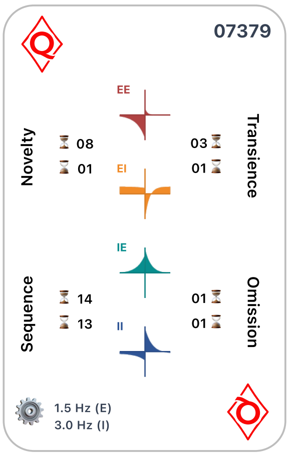
07379
EE
EI
IE
II
Novelty
08
01
Transience
03
01
Sequence
14
13
Omission
01
01
1.5 Hz (E)
3.0 Hz (I)

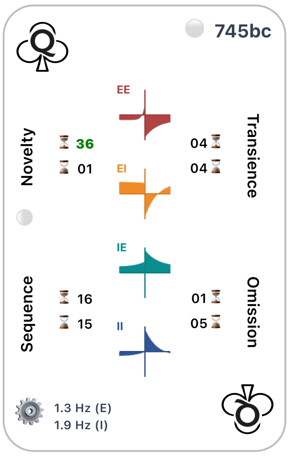
745bc
EE
EI
IE
II
Novelty
36
01
Transience
04
04
Sequence
16
15
Omission
01
05
1.3 Hz (E)
1.9 Hz (I)

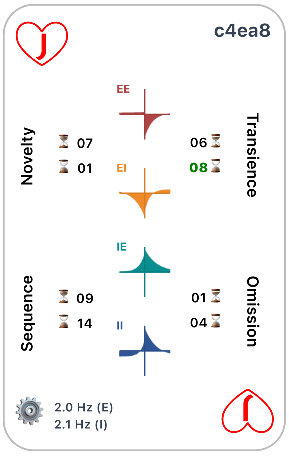
c4ea8
EE
EI
IE
II
Novelty
07
01
Transience
06
08
Sequence
09
14
Omission
01
04
2.0 Hz (E)
2.1 Hz (I)

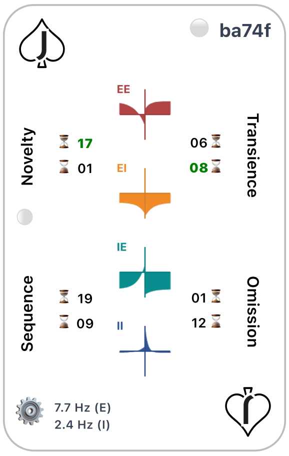
ba74f
EE
EI
IE
II
Novelty
17
01
Transience
06
08
Sequence
19
09
Omission
01
12
7.7 Hz (E)
2.4 Hz (I)

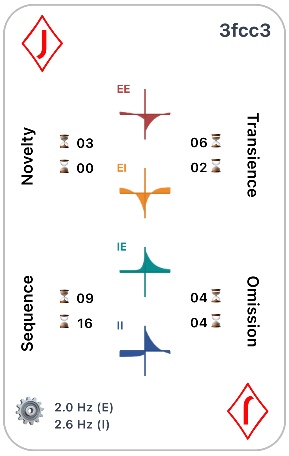
3fcc3
EE
EI
IE
II
Novelty
03
00
Transience
06
02
Sequence
09
16
Omission
04
04
2.0 Hz (E)
2.6 Hz (I)

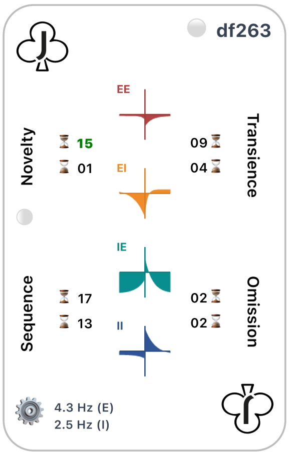
df263
EE
EI
IE
II
Novelty
15
01
Transience
09
04
Sequence
17
13
Omission
02
02
4.3 Hz (E)
2.5 Hz (I)

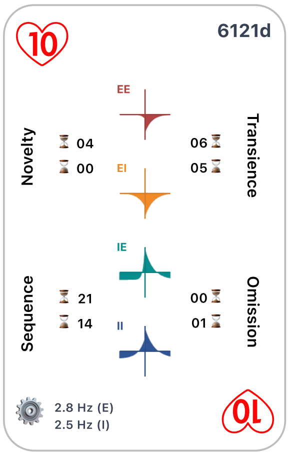
6121d
EE
EI
IE
II
Novelty
04
00
Transience
06
05
Sequence
21
14
Omission
00
01
2.8 Hz (E)
2.5 Hz (I)

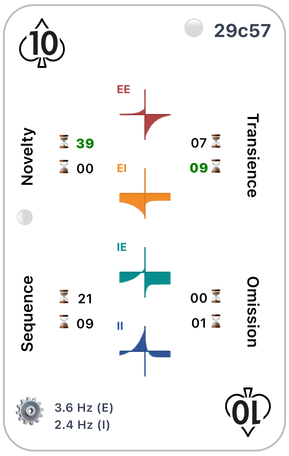
29c57
EE
EI
IE
II
Novelty
39
00
Transience
07
09
Sequence
21
09
Omission
00
01
3.6 Hz (E)
2.4 Hz (I)

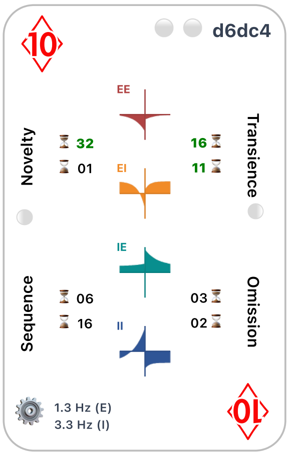
d6dc4
EE
EI
IE
II
Novelty
32
01
Transience
16
11
Sequence
06
16
Omission
03
02
1.3 Hz (E)
3.3 Hz (I)

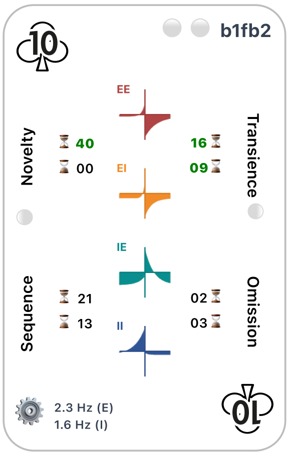
b1fb2
EE
EI
IE
II
Novelty
40
00
Transience
16
09
Sequence
21
13
Omission
02
03
2.3 Hz (E)
1.6 Hz (I)

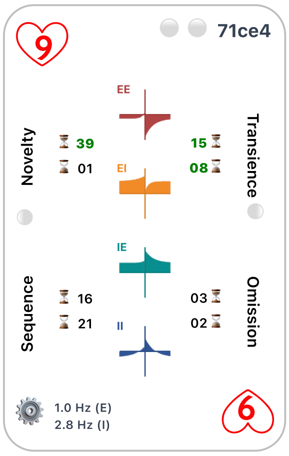
71ce4
EE
EI
IE
II
Novelty
39
01
Transience
15
08
Sequence
16
21
Omission
03
02
1.0 Hz (E)
2.8 Hz (I)

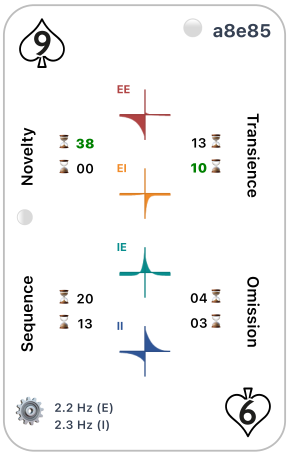
a8e85
EE
EI
IE
II
Novelty
38
00
Transience
13
10
Sequence
20
13
Omission
04
03
2.2 Hz (E)
2.3 Hz (I)

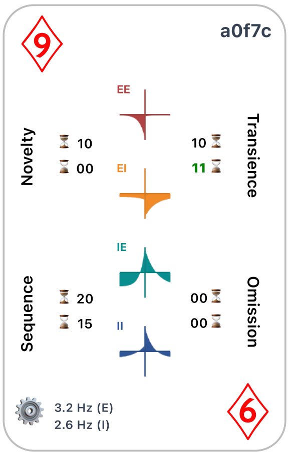
a0f7c
EE
EI
IE
II
Novelty
10
00
Transience
10
11
Sequence
20
15
Omission
00
00
3.2 Hz (E)
2.6 Hz (I)

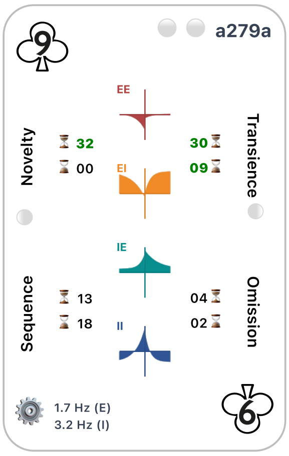
a279a
EE
EI
IE
II
Novelty
32
00
Transience
30
09
Sequence
13
18
Omission
04
02
1.7 Hz (E)
3.2 Hz (I)

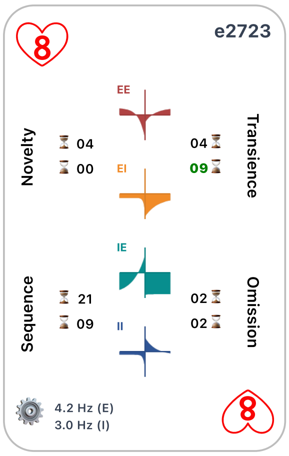
8
e2723
Novelty 04 00
Transience 04 09
EE
EI
IE
II
Sequence 21 09
Omission 02 02
4.2 Hz (E)
3.0 Hz (I)

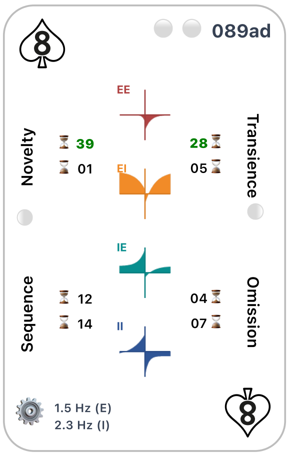
8
089ad
Novelty 39 01
Transience 28 05
EE
EI
IE
II
Sequence 12 14
Omission 04 07
1.5 Hz (E)
2.3 Hz (I)

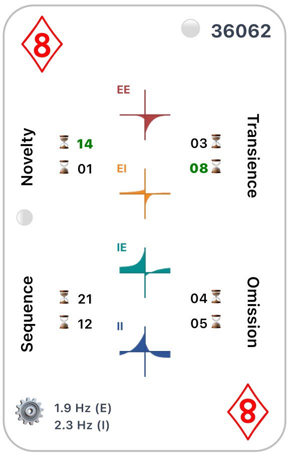
8
36062
Novelty 14 01
Transience 03 08
EE
EI
IE
II
Sequence 21 12
Omission 04 05
1.9 Hz (E)
2.3 Hz (I)

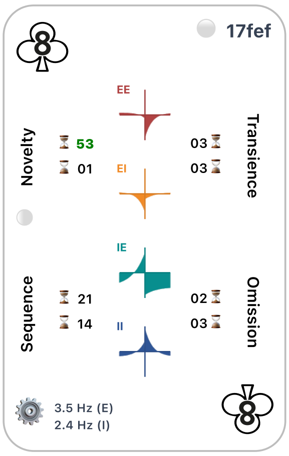
8
17fef
Novelty 53 01
Transience 03 03
EE
EI
IE
II
Sequence 21 14
Omission 02 03
3.5 Hz (E)
2.4 Hz (I)

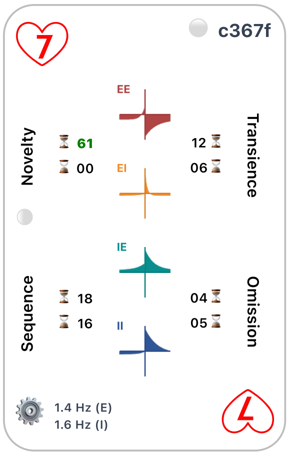
7
c367f
Novelty 61 00
Transience 12 06
EE
EI
IE
II
Sequence 18 16
Omission 04 05
1.4 Hz (E)
1.6 Hz (I)

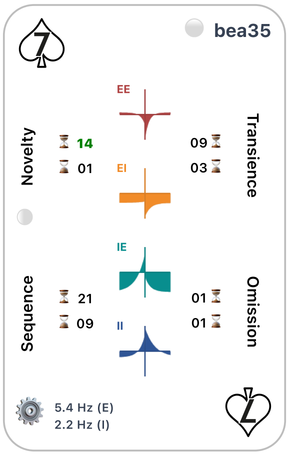
7
bea35
Novelty 14 01
Transience 09 03
EE
EI
IE
II
Sequence 21 09
Omission 01 01
5.4 Hz (E)
2.2 Hz (I)

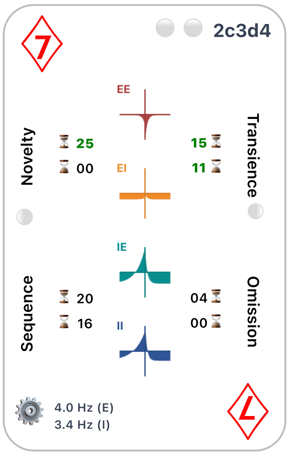
7
2c3d4
Novelty 25 00
Transience 15 11
EE
EI
IE
II
Sequence 20 16
Omission 04 00
4.0 Hz (E)
3.4 Hz (I)

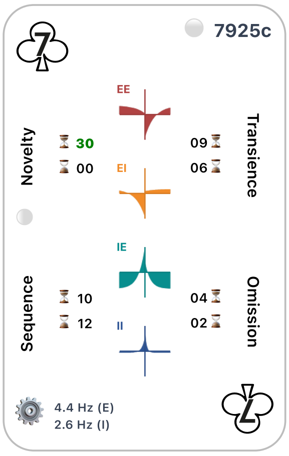
7
7925c
Novelty 30 00
Transience 09 06
EE
EI
IE
II
Sequence 10 12
Omission 04 02
4.4 Hz (E)
2.6 Hz (I)

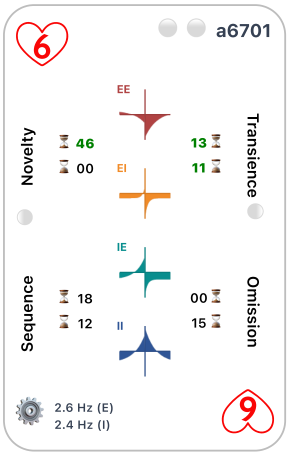
6
a6701
Novelty 46 00
Transience 13 11
EE
EI
IE
II
Sequence 18 12
Omission 00 15
2.6 Hz (E)
2.4 Hz (I)

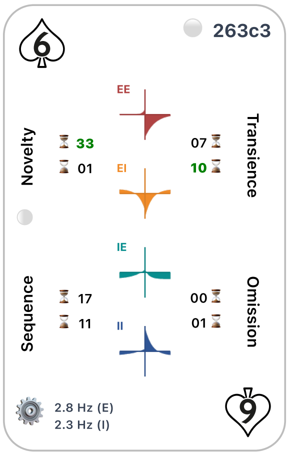
6
263c3
Novelty 33 01
Transience 07 10
EE
EI
IE
II
Sequence 17 11
Omission 00 01
2.8 Hz (E)
2.3 Hz (I)

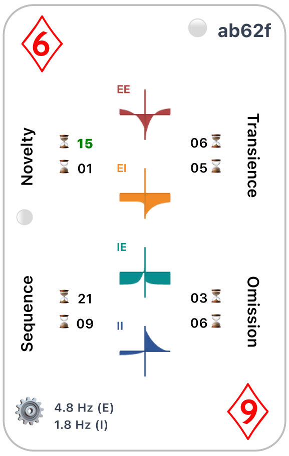
6
ab62f
Novelty 15 01
Transience 06 05
EE
EI
IE
II
Sequence 21 09
Omission 03 06
4.8 Hz (E)
1.8 Hz (I)

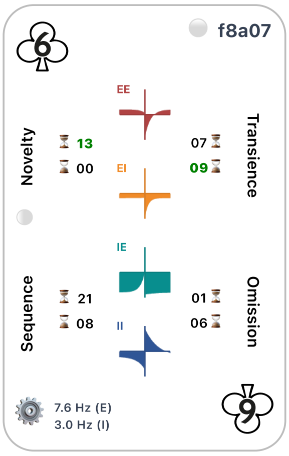
6
f8a07
Novelty 13 00
Transience 07 09
EE
EI
IE
II
Sequence 21 08
Omission 01 06
7.6 Hz (E)
3.0 Hz (I)

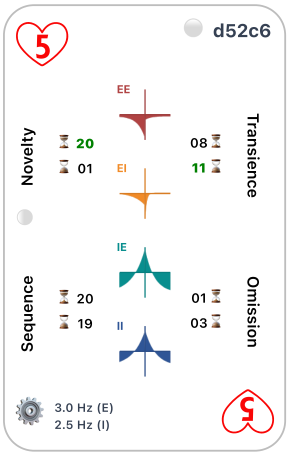
5
d52c6
Novelty 20 01
Transience 08 11
EE
EI
IE
II
Sequence 20 19
Omission 01 03
3.0 Hz (E)
2.5 Hz (I)

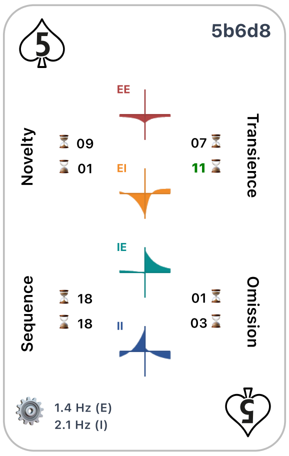
5
5b6d8
Novelty 09 01
Transience 07 11
EE
EI
IE
II
Sequence 18 18
Omission 01 03
1.4 Hz (E)
2.1 Hz (I)

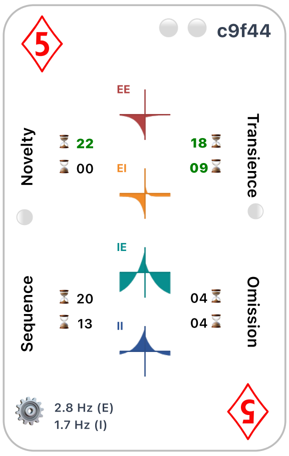
5
c9f44
Novelty 22 00
Transience 18 09
EE
EI
IE
II
Sequence 20 13
Omission 04 04
2.8 Hz (E)
1.7 Hz (I)

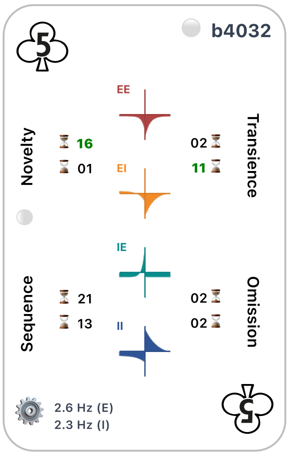
5
b4032
Novelty 16 01
Transience 02 11
EE
EI
IE
II
Sequence 21 13
Omission 02 02
2.6 Hz (E)
2.3 Hz (I)

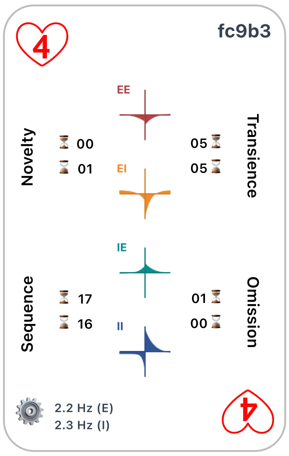
4
fc9b3
Novelty 00 01
Transience 05 05
Sequence 17 16
Omission 01 00
EE EI IE II
2.2 Hz (E)
2.3 Hz (I)

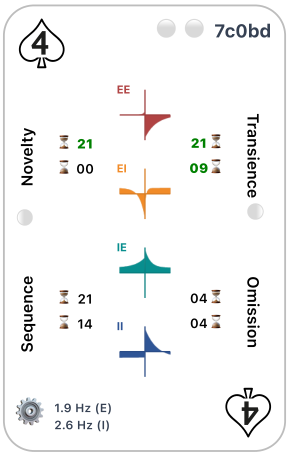
4
7c0bd
Novelty 21 00
Transience 21 09
Sequence 21 14
Omission 04 04
EE EI IE II
1.9 Hz (E)
2.6 Hz (I)

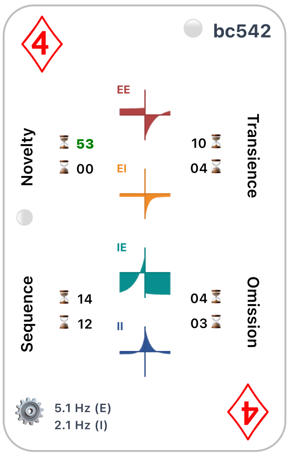
4
bc542
Novelty 53 00
Transience 10 04
Sequence 14 12
Omission 04 03
EE EI IE II
5.1 Hz (E)
2.1 Hz (I)

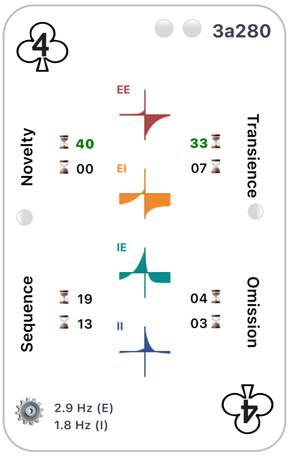
4
3a280
Novelty 40 00
Transience 33 07
Sequence 19 13
Omission 04 03
EE EI IE II
2.9 Hz (E)
1.8 Hz (I)

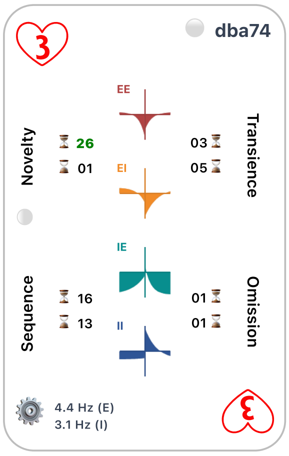
3
dba74
Novelty 26 01
Transience 03 05
Sequence 16 13
Omission 01 01
EE EI IE II
4.4 Hz (E)
3.1 Hz (I)

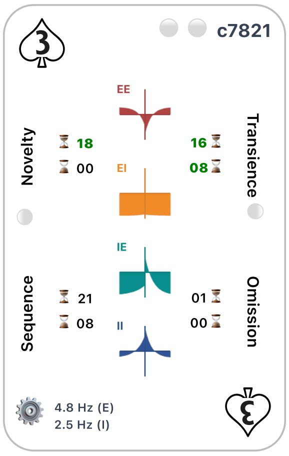
3
c7821
Novelty 18 00
Transience 16 08
Sequence 21 08
Omission 01 00
EE EI IE II
4.8 Hz (E)
2.5 Hz (I)

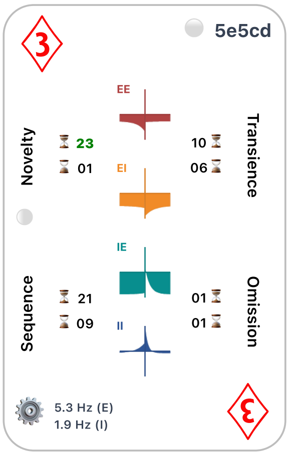
3
5e5cd
Novelty 23 01
Transience 10 06
Sequence 21 09
Omission 01 01
EE EI IE II
5.3 Hz (E)
1.9 Hz (I)

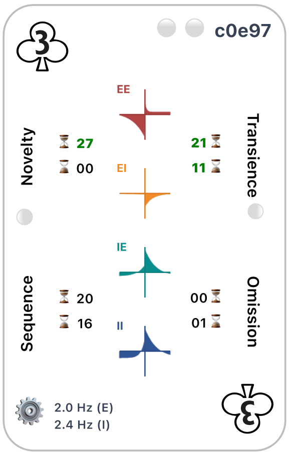
3
c0e97
Novelty 27 00
Transience 21 11
Sequence 20 16
Omission 00 01
EE EI IE II
2.0 Hz (E)
2.4 Hz (I)

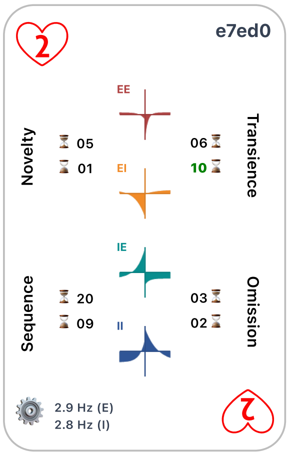
2
e7ed0
Novelty 05 01
Transience 06 10
Sequence 20 09
Omission 03 02
EE EI IE II
2.9 Hz (E)
2.8 Hz (I)

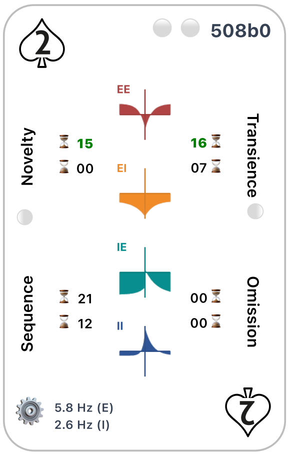
2
508b0
Novelty 15 00
Transience 16 07
Sequence 21 12
Omission 00 00
EE EI IE II
5.8 Hz (E)
2.6 Hz (I)

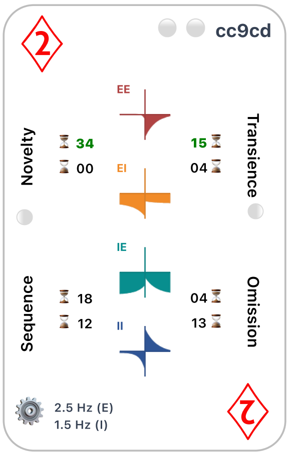
2
cc9cd
Novelty 34 00
Transience 15 04
Sequence 18 12
Omission 04 13
EE EI IE II
2.5 Hz (E)
1.5 Hz (I)

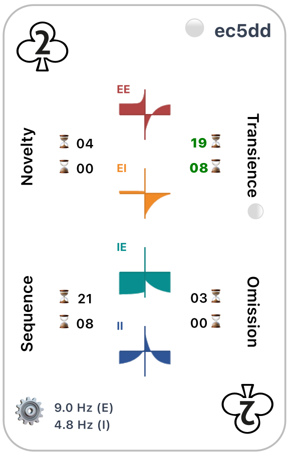
2
ec5dd
Novelty 04 00
Transience 19 08
Sequence 21 08
Omission 03 00
EE EI IE II
9.0 Hz (E)
4.8 Hz (I)

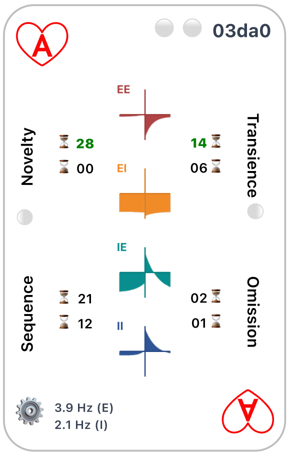
A
03da0
Novelty 28 00
Transience 14 06
Sequence 21 12
Omission 02 01
EE EI IE II
3.9 Hz (E)
2.1 Hz (I)

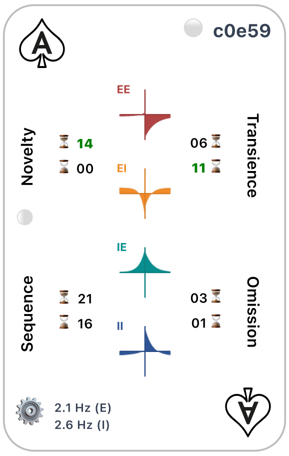
A
c0e59
Novelty 14 00
Transience 06 11
Sequence 21 16
Omission 03 01
EE EI IE II
2.1 Hz (E)
2.6 Hz (I)

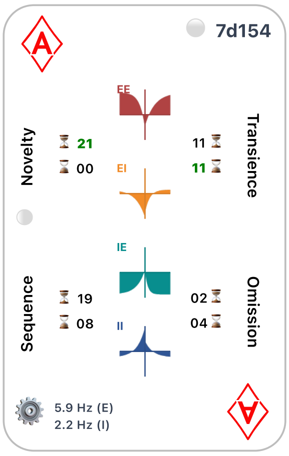
A
7d154
Novelty 21 00
Transience 11 11
Sequence 19 08
Omission 02 04
EE EI IE II
5.9 Hz (E)
2.2 Hz (I)

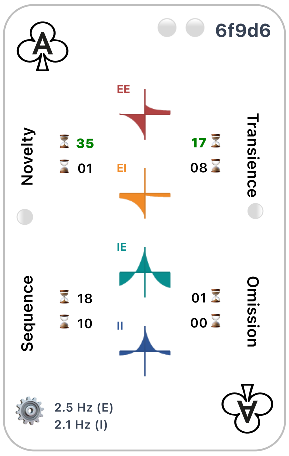
A
6f9d6
Novelty 35 01
Transience 17 08
Sequence 18 10
Omission 01 00
EE EI IE II
2.5 Hz (E)
2.1 Hz (I)

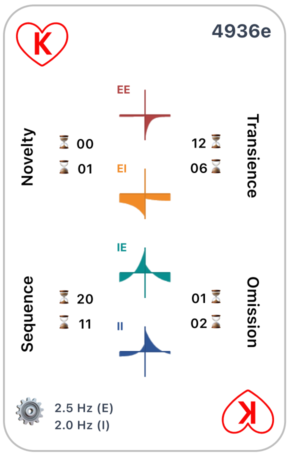
K
4936e
EE
EI
IE
II
Novelty 00 01
Transience 12 06
Sequence 20 11
Omission 01 02
2.5 Hz (E)
2.0 Hz (I)

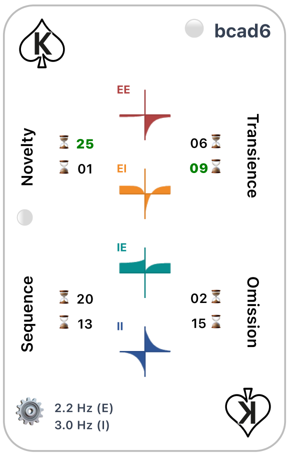
K
bcad6
EE
EI
IE
II
Novelty 25 01
Transience 06 09
Sequence 20 13
Omission 02 15
2.2 Hz (E)
3.0 Hz (I)

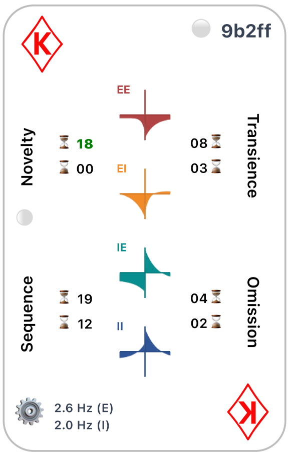
K
9b2ff
EE
EI
IE
II
Novelty 18 00
Transience 08 03
Sequence 19 12
Omission 04 02
2.6 Hz (E)
2.0 Hz (I)

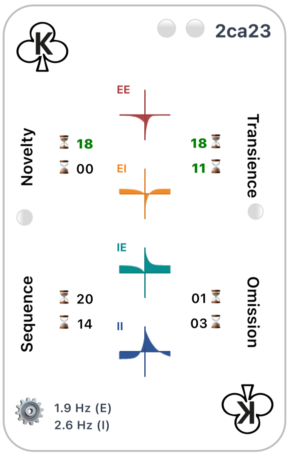
K
2ca23
EE
EI
IE
II
Novelty 18 00
Transience 18 11
Sequence 20 14
Omission 01 03
1.9 Hz (E)
2.6 Hz (I)

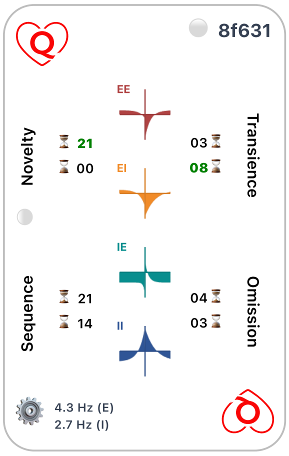
Q
8f631
EE
EI
IE
II
Novelty 21 00
Transience 03 08
Sequence 21 14
Omission 04 03
4.3 Hz (E)
2.7 Hz (I)

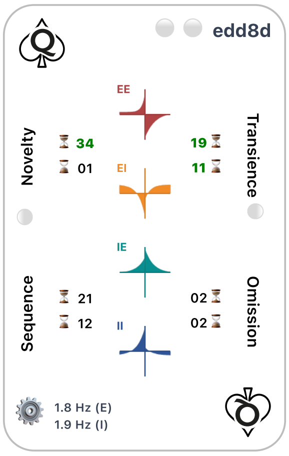
Q
edd8d
EE
EI
IE
II
Novelty 34 01
Transience 19 11
Sequence 21 12
Omission 02 02
1.8 Hz (E)
1.9 Hz (I)

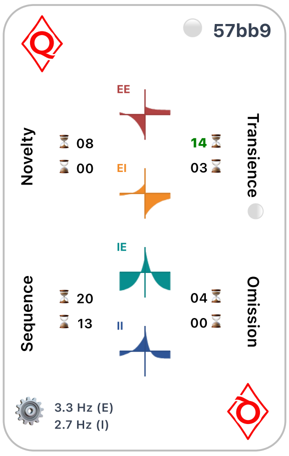
Q
57bb9
EE
EI
IE
II
Novelty 08 00
Transience 14 03
Sequence 20 13
Omission 04 00
3.3 Hz (E)
2.7 Hz (I)

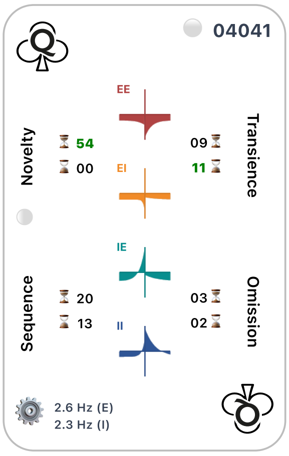
Q
04041
EE
EI
IE
II
Novelty 54 00
Transience 09 11
Sequence 20 13
Omission 03 02
2.6 Hz (E)
2.3 Hz (I)

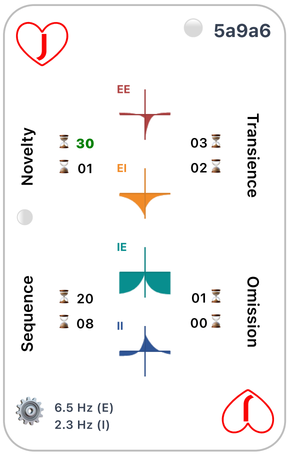
J
5a9a6
EE
EI
IE
II
Novelty 30 01
Transience 03 02
Sequence 20 08
Omission 01 00
6.5 Hz (E)
2.3 Hz (I)

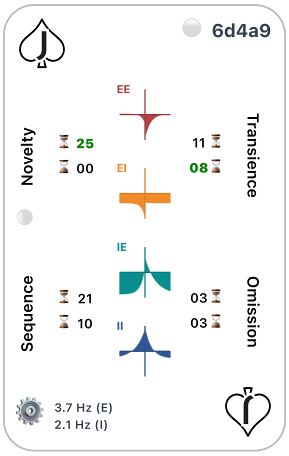
J
6d4a9
EE
EI
IE
II
Novelty 25 00
Transience 11 08
Sequence 21 10
Omission 03 03
3.7 Hz (E)
2.1 Hz (I)

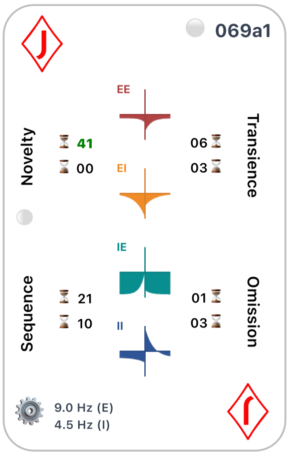
J
069a1
EE
EI
IE
II
Novelty 41 00
Transience 06 03
Sequence 21 10
Omission 01 03
9.0 Hz (E)
4.5 Hz (I)

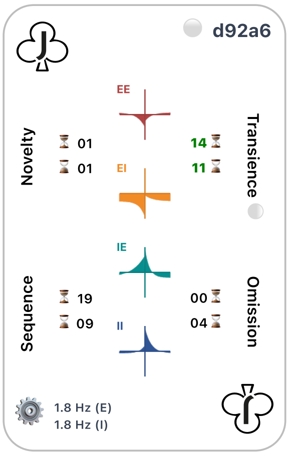
J
d92a6
EE
EI
IE
II
Novelty 01 01
Transience 14 11
Sequence 19 09
Omission 00 04
1.8 Hz (E)
1.8 Hz (I)

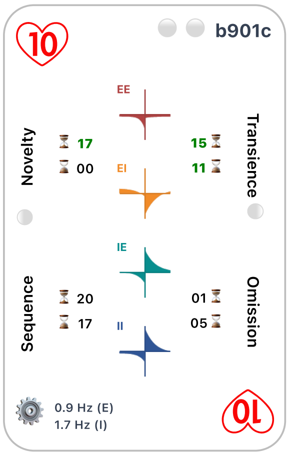
10
b901c
EE
EI
IE
II
Novelty 17 00
Transience 15 11
Sequence 20 17
Omission 01 05
0.9 Hz (E)
1.7 Hz (I)

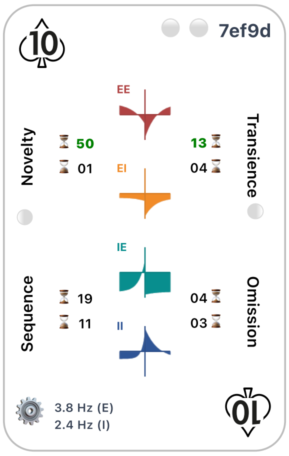
10
7ef9d
EE
EI
IE
II
Novelty 50 01
Transience 13 04
Sequence 19 11
Omission 04 03
3.8 Hz (E)
2.4 Hz (I)

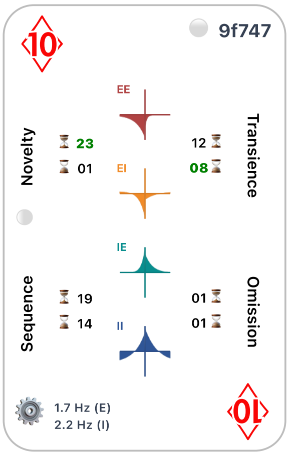
10
9f747
EE
EI
IE
II
Novelty 23 01
Transience 12 08
Sequence 19 14
Omission 01 01
1.7 Hz (E)
2.2 Hz (I)

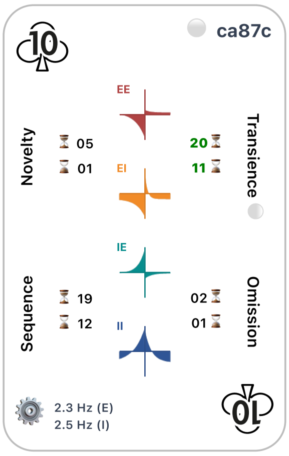
10
ca87c
EE
EI
IE
II
Novelty 05 01
Transience 20 11
Sequence 19 12
Omission 02 01
2.3 Hz (E)
2.5 Hz (I)

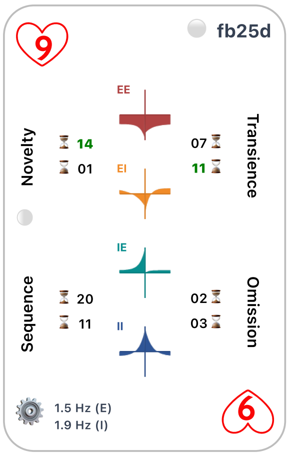
9
fb25d
EE
EI
IE
II
Novelty 14 01
Transience 07 11
Sequence 20 11
Omission 02 03
1.5 Hz (E)
1.9 Hz (I)

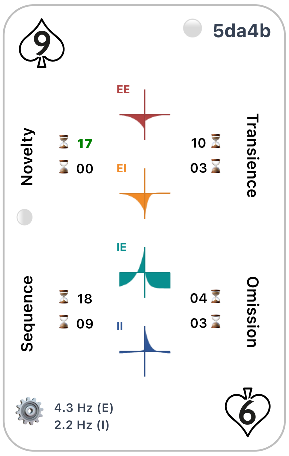
9
5da4b
EE
EI
IE
II
Novelty 17 00
Transience 10 03
Sequence 18 09
Omission 04 03
4.3 Hz (E)
2.2 Hz (I)

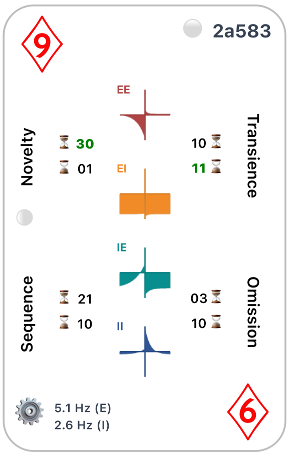
9
2a583
EE
EI
IE
II
Novelty 30 01
Transience 10 11
Sequence 21 10
Omission 03 10
5.1 Hz (E)
2.6 Hz (I)

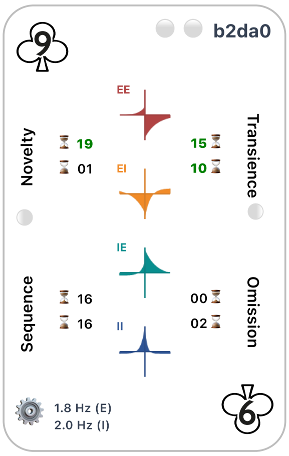
9
b2da0
EE
EI
IE
II
Novelty 19 01
Transience 15 10
Sequence 16 16
Omission 00 02
1.8 Hz (E)
2.0 Hz (I)

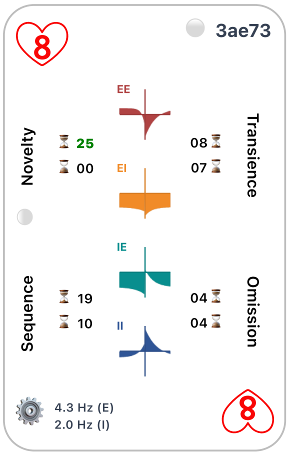
8
3ae73
EE
EI
IE
II
Novelty 25 00
Transience 08 07
Sequence 19 10
Omission 04 04
4.3 Hz (E)
2.0 Hz (I)

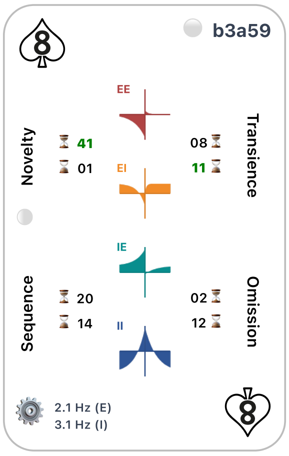
8
b3a59
EE
EI
IE
II
Novelty 41 01
Transience 08 11
Sequence 20 14
Omission 02 12
2.1 Hz (E)
3.1 Hz (I)

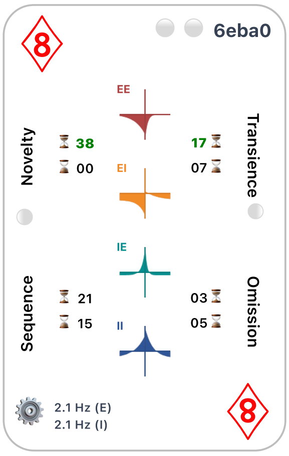
8
6eba0
EE
EI
IE
II
Novelty 38 00
Transience 17 07
Sequence 21 15
Omission 03 05
2.1 Hz (E)
2.1 Hz (I)

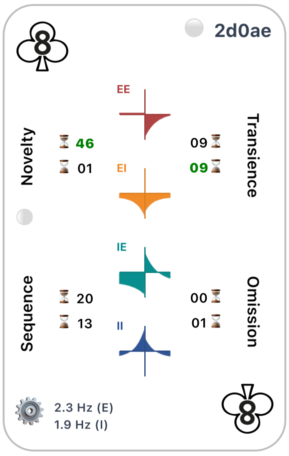
8
2d0ae
EE
EI
IE
II
Novelty 46 01
Transience 09 09
Sequence 20 13
Omission 00 01
2.3 Hz (E)
1.9 Hz (I)

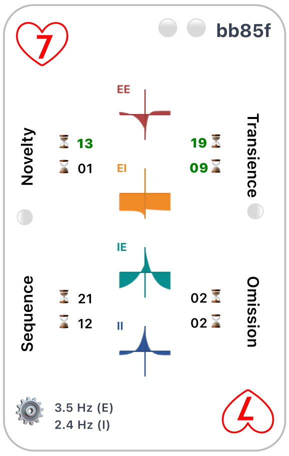
7
bb85f
EE
EI
IE
II
Novelty 13 01
Transience 19 09
Sequence 21 12
Omission 02 02
3.5 Hz (E)
2.4 Hz (I)

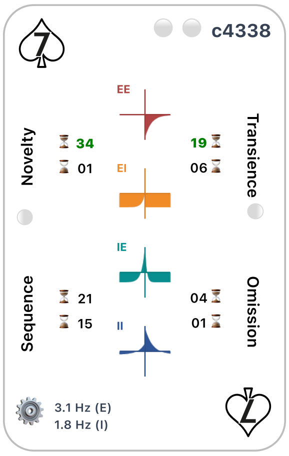
7
c4338
EE
EI
IE
II
Novelty 34 01
Transience 19 06
Sequence 21 15
Omission 04 01
3.1 Hz (E)
1.8 Hz (I)

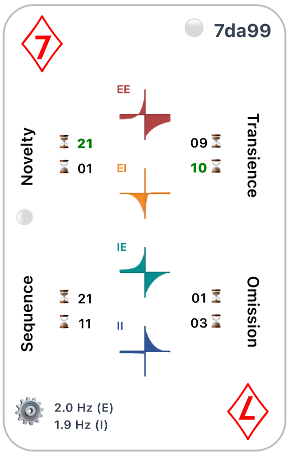
7
7da99
EE
EI
IE
II
Novelty 21 01
Transience 09 10
Sequence 21 11
Omission 01 03
2.0 Hz (E)
1.9 Hz (I)

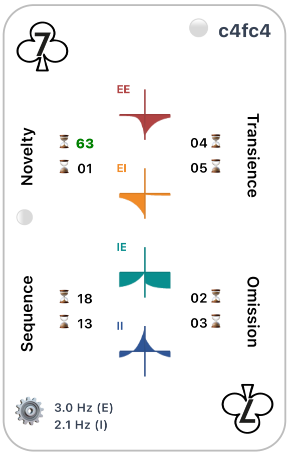
7
c4fc4
EE
EI
IE
II
Novelty 63 01
Transience 04 05
Sequence 18 13
Omission 02 03
3.0 Hz (E)
2.1 Hz (I)

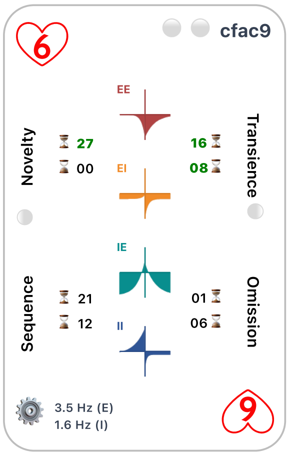
6
cfac9
EE
EI
IE
II
Novelty 27 00
Transience 16 08
Sequence 21 12
Omission 01 06
3.5 Hz (E)
1.6 Hz (I)

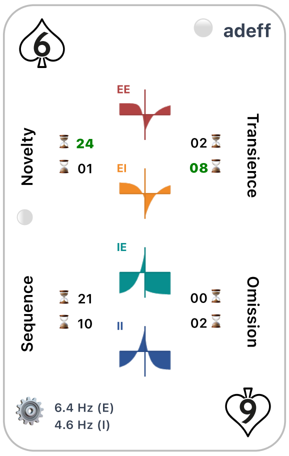
6
adeff
EE
EI
IE
II
Novelty 24 01
Transience 02 08
Sequence 21 10
Omission 00 02
6.4 Hz (E)
4.6 Hz (I)

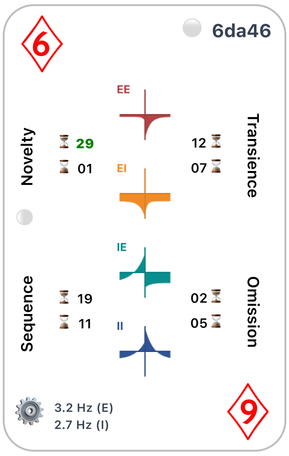
6
6da46
EE
EI
IE
II
Novelty 29 01
Transience 12 07
Sequence 19 11
Omission 02 05
3.2 Hz (E)
2.7 Hz (I)

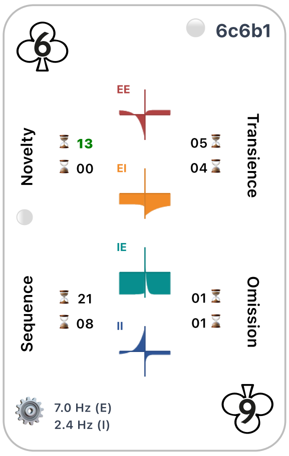
6
6c6b1
EE
EI
IE
II
Novelty 13 00
Transience 05 04
Sequence 21 08
Omission 01 01
7.0 Hz (E)
2.4 Hz (I)

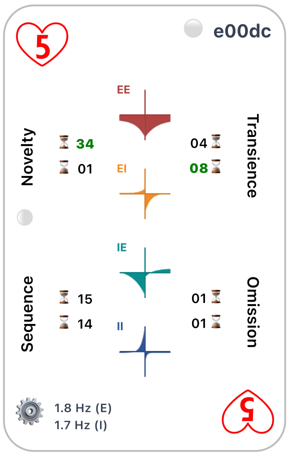
5
e00dc
EE
Novelty
34
01
04
08
Transience
EI
IE
Sequence
15
14
01
01
Omission
II
1.8 Hz (E)
1.7 Hz (I)

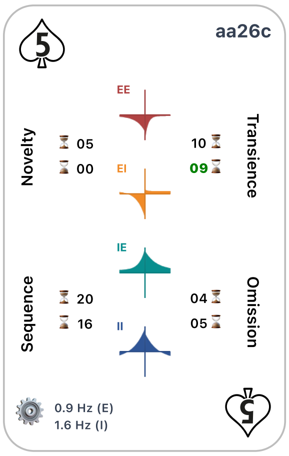
5
aa26c
EE
Novelty
05
00
10
09
Transience
EI
IE
Sequence
20
16
04
05
Omission
II
0.9 Hz (E)
1.6 Hz (I)

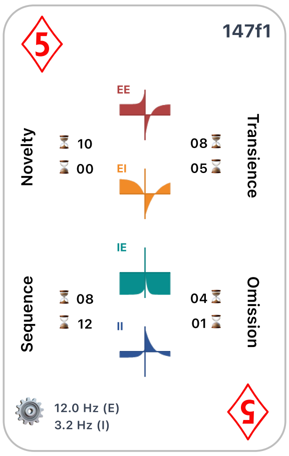
5
147f1
EE
Novelty
10
00
08
05
Transience
EI
IE
Sequence
08
12
04
01
Omission
II
12.0 Hz (E)
3.2 Hz (I)

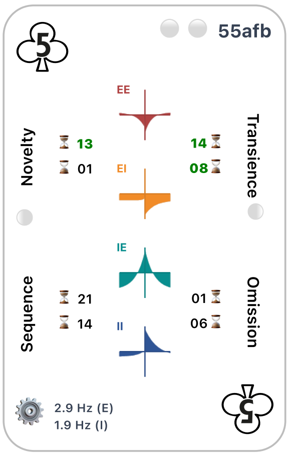
5
55afb
EE
Novelty
13
01
14
08
Transience
EI
IE
Sequence
21
14
01
06
Omission
II
2.9 Hz (E)
1.9 Hz (I)

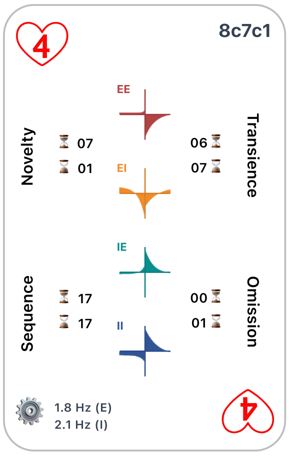
4
8c7c1
EE
Novelty
07
01
06
07
Transience
EI
IE
Sequence
17
17
00
01
Omission
II
1.8 Hz (E)
2.1 Hz (I)

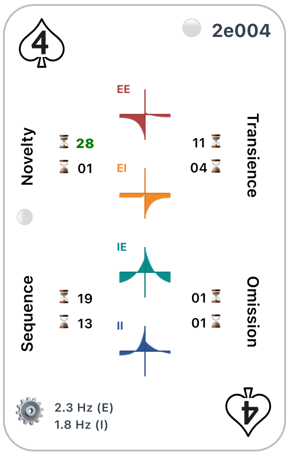
4
2e004
EE
Novelty
28
01
11
04
Transience
EI
IE
Sequence
19
13
01
01
Omission
II
2.3 Hz (E)
1.8 Hz (I)

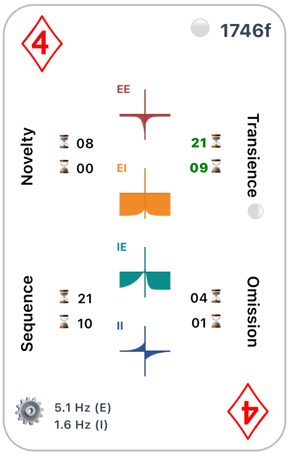
4
1746f
EE
Novelty
08
00
21
09
Transience
EI
IE
Sequence
21
10
04
01
Omission
II
5.1 Hz (E)
1.6 Hz (I)

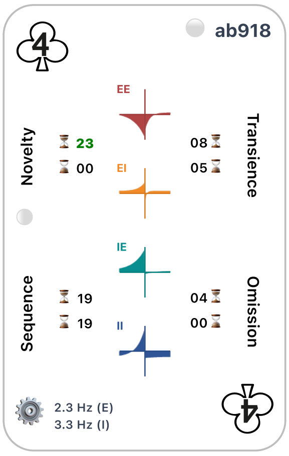
4
ab918
EE
Novelty
23
00
08
05
Transience
EI
IE
Sequence
19
19
04
00
Omission
II
2.3 Hz (E)
3.3 Hz (I)

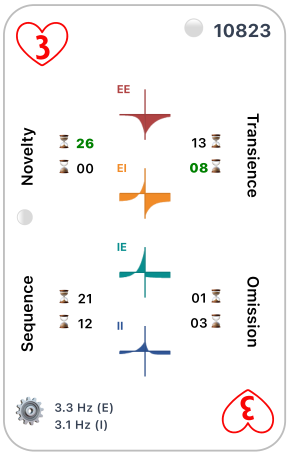
3
10823
EE
Novelty
26
00
13
08
Transience
EI
IE
Sequence
21
12
01
03
Omission
II
3.3 Hz (E)
3.1 Hz (I)

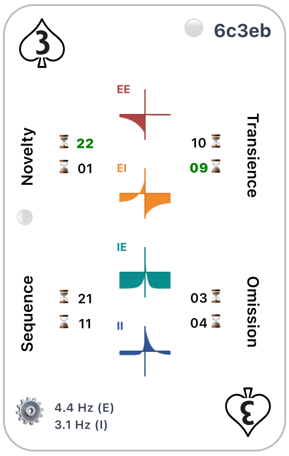
3
6c3eb
EE
Novelty
22
01
10
09
Transience
EI
IE
Sequence
21
11
03
04
Omission
II
4.4 Hz (E)
3.1 Hz (I)

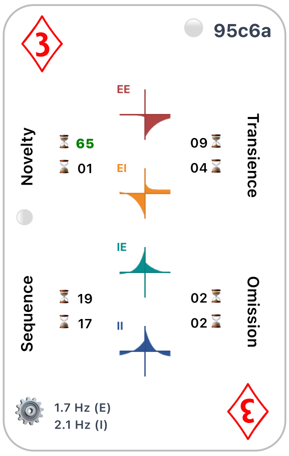
3
95c6a
EE
Novelty
65
01
09
04
Transience
EI
IE
Sequence
19
17
02
02
Omission
II
1.7 Hz (E)
2.1 Hz (I)

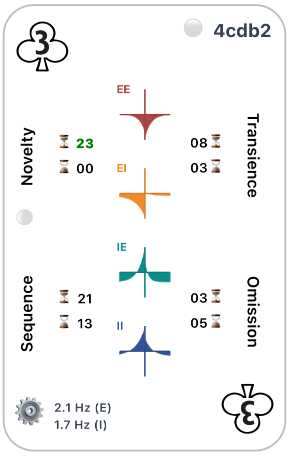
3
4cdb2
EE
Novelty
23
00
08
03
Transience
EI
IE
Sequence
21
13
03
05
Omission
II
2.1 Hz (E)
1.7 Hz (I)

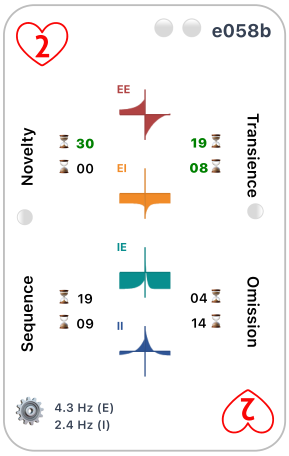
2
e058b
EE
Novelty
30
00
19
08
Transience
EI
IE
Sequence
19
09
04
14
Omission
II
4.3 Hz (E)
2.4 Hz (I)

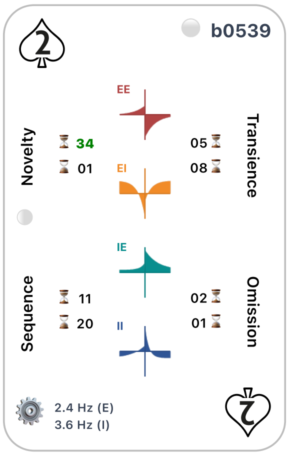
2
b0539
EE
Novelty
34
01
05
08
Transience
EI
IE
Sequence
11
20
02
01
Omission
II
2.4 Hz (E)
3.6 Hz (I)

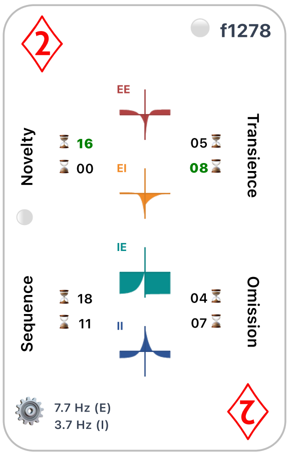
2
f1278
EE
Novelty
16
00
05
08
Transience
EI
IE
Sequence
18
11
04
07
Omission
II
7.7 Hz (E)
3.7 Hz (I)

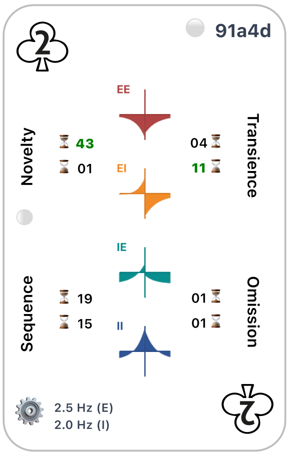
2
91a4d
EE
Novelty
43
01
04
11
Transience
EI
IE
Sequence
19
15
01
01
Omission
II
2.5 Hz (E)
2.0 Hz (I)

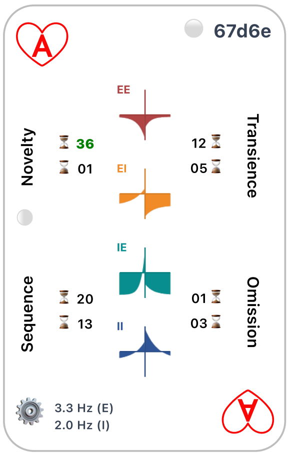
A
67d6e
EE
Novelty
36
01
12
05
Transience
EI
IE
Sequence
20
13
01
03
Omission
II
3.3 Hz (E)
2.0 Hz (I)

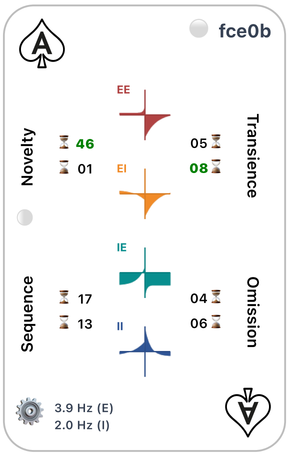
A
fce0b
EE
Novelty
46
01
05
08
Transience
EI
IE
Sequence
17
13
04
06
Omission
II
3.9 Hz (E)
2.0 Hz (I)

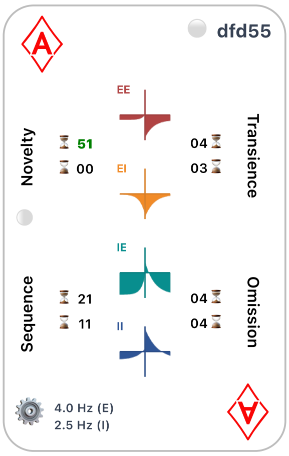
A
dfd55
EE
Novelty
51
00
04
03
Transience
EI
IE
Sequence
21
11
04
04
Omission
II
4.0 Hz (E)
2.5 Hz (I)

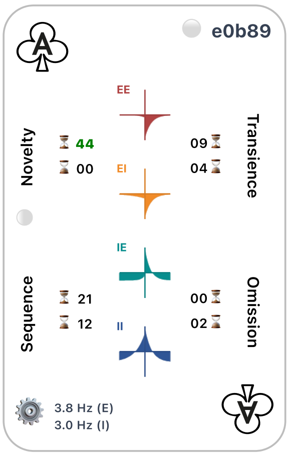
A
e0b89
EE
Novelty
44
00
09
04
Transience
EI
IE
Sequence
21
12
00
02
Omission
II
3.8 Hz (E)
3.0 Hz (I)

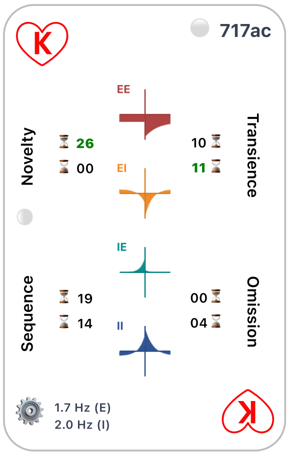
K
717ac
EE
Novelty
26
00
10
11
Transience
EI
IE
Sequence
19
14
00
04
Omission
II
1.7 Hz (E)
2.0 Hz (I)

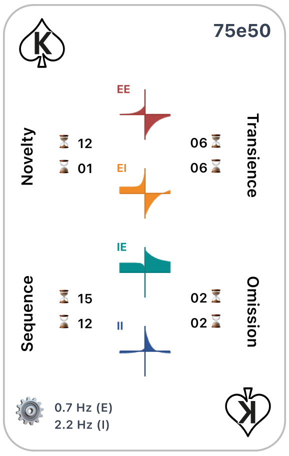
K
75e50
EE
Novelty
12
01
06
06
Transience
EI
IE
Sequence
15
12
02
02
Omission
II
0.7 Hz (E)
2.2 Hz (I)

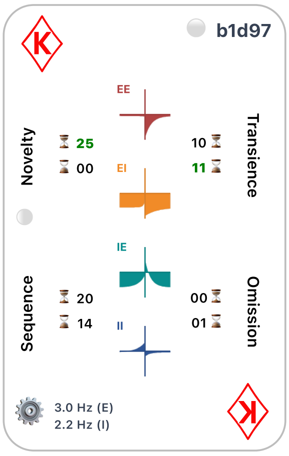
K
b1d97
EE
Novelty
25
00
10
11
Transience
EI
IE
Sequence
20
14
00
01
Omission
II
3.0 Hz (E)
2.2 Hz (I)

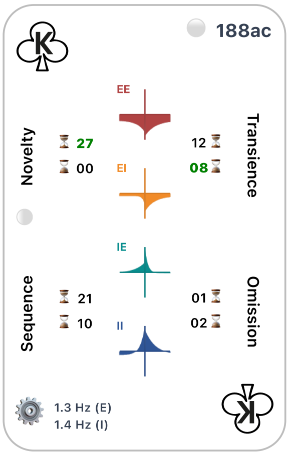
K
188ac
EE
Novelty
27
00
12
08
Transience
EI
IE
Sequence
21
10
01
02
Omission
II
1.3 Hz (E)
1.4 Hz (I)

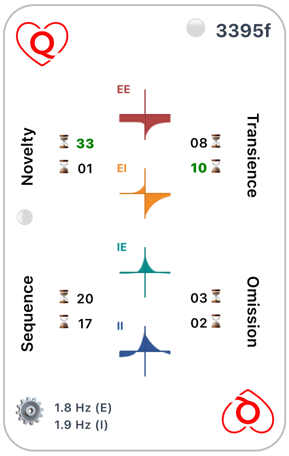
Q
3395f
EE
Novelty
33
01
08
10
Transience
EI
IE
Sequence
20
17
03
02
Omission
II
1.8 Hz (E)
1.9 Hz (I)

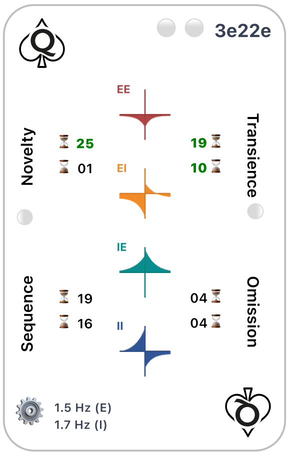
Q
3e22e
EE
Novelty
25
01
19
10
Transience
EI
IE
Sequence
19
16
04
04
Omission
II
1.5 Hz (E)
1.7 Hz (I)

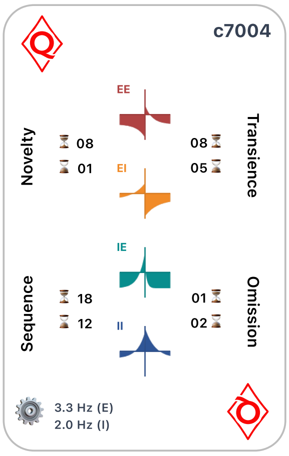
Q
c7004
EE
Novelty
08
01
08
05
Transience
EI
IE
Sequence
18
12
01
02
Omission
II
3.3 Hz (E)
2.0 Hz (I)

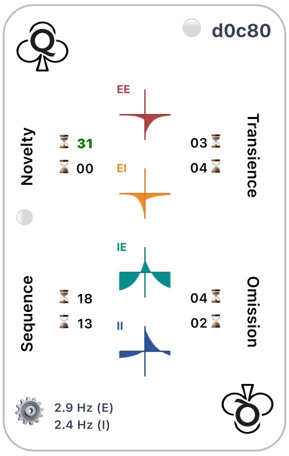
Q
d0c80
EE
Novelty
31
00
03
04
Transience
EI
IE
Sequence
18
13
04
02
Omission
II
2.9 Hz (E)
2.4 Hz (I)

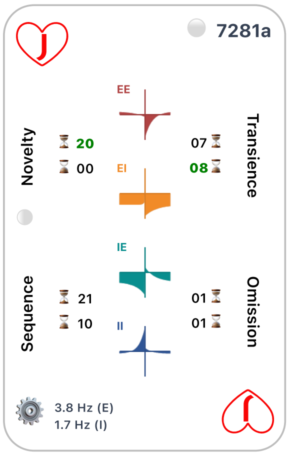
J
7281a
EE
Novelty
20
00
07
08
Transience
EI
IE
Sequence
21
10
01
01
Omission
II
3.8 Hz (E)
1.7 Hz (I)

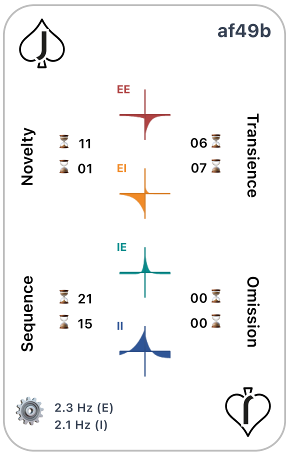
J
af49b
EE
Novelty
11
01
06
07
Transience
EI
IE
Sequence
21
15
00
00
Omission
II
2.3 Hz (E)
2.1 Hz (I)

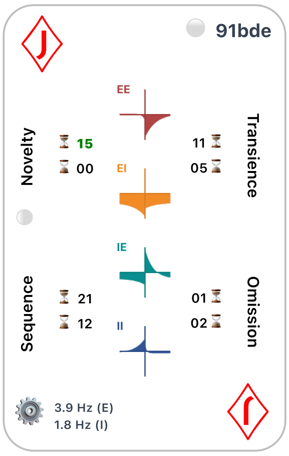
J
91bde
EE
Novelty
15
00
11
05
Transience
EI
IE
Sequence
21
12
01
02
Omission
II
3.9 Hz (E)
1.8 Hz (I)

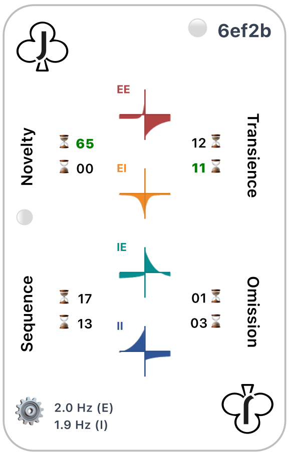
J
6ef2b
EE
Novelty
65
00
12
11
Transience
EI
IE
Sequence
17
13
01
03
Omission
II
2.0 Hz (E)
1.9 Hz (I)

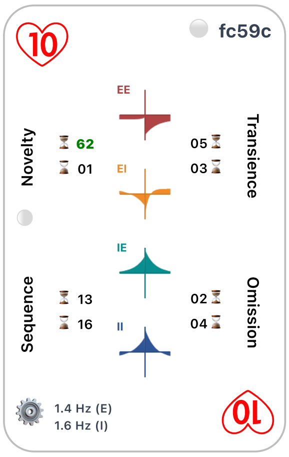
10
fc59c
EE
EI
IE
II
Novelty
62
01
Transience
05
03
Sequence
13
16
Omission
02
04
1.4 Hz (E)
1.6 Hz (I)

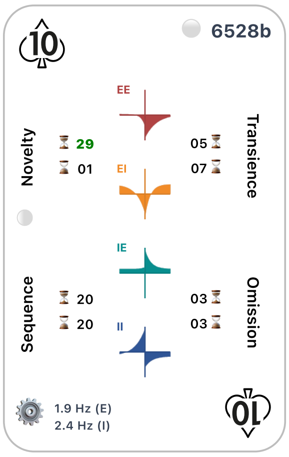
10
6528b
EE
EI
IE
II
Novelty
29
01
Transience
05
07
Sequence
20
20
Omission
03
03
1.9 Hz (E)
2.4 Hz (I)

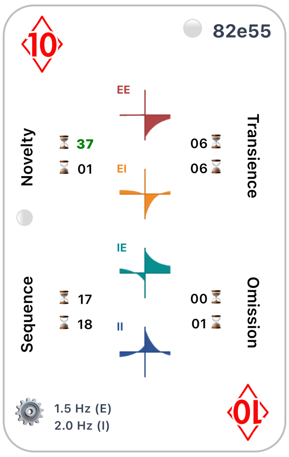
10
82e55
EE
EI
IE
II
Novelty
37
01
Transience
06
06
Sequence
17
18
Omission
00
01
1.5 Hz (E)
2.0 Hz (I)

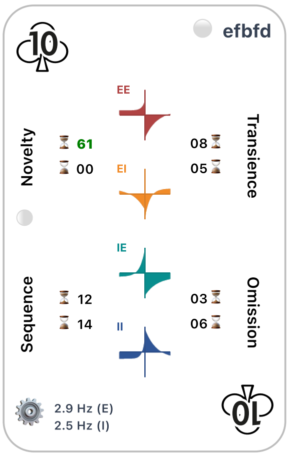
10
efbfd
EE
EI
IE
II
Novelty
61
00
Transience
08
05
Sequence
12
14
Omission
03
06
2.9 Hz (E)
2.5 Hz (I)

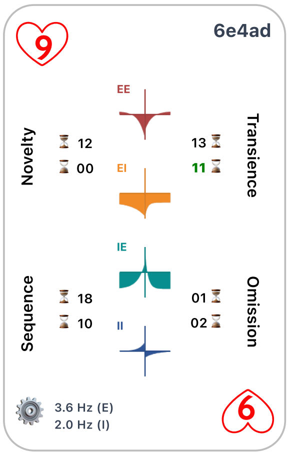
9
6e4ad
EE
EI
IE
II
Novelty
12
00
Transience
13
11
Sequence
18
10
Omission
01
02
3.6 Hz (E)
2.0 Hz (I)

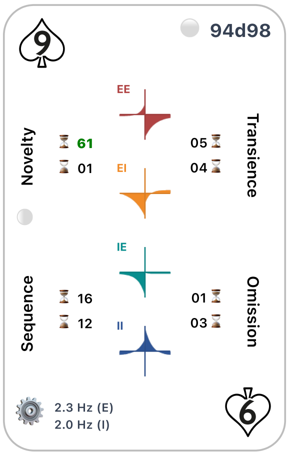
9
94d98
EE
EI
IE
II
Novelty
61
01
Transience
05
04
Sequence
16
12
Omission
01
03
2.3 Hz (E)
2.0 Hz (I)

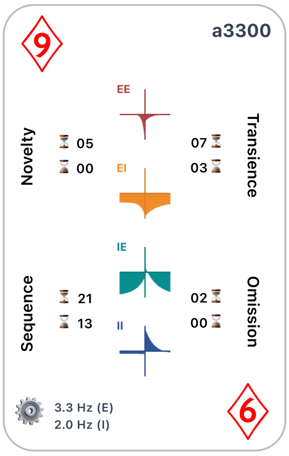
9
a3300
EE
EI
IE
II
Novelty
05
00
Transience
07
03
Sequence
21
13
Omission
02
00
3.3 Hz (E)
2.0 Hz (I)

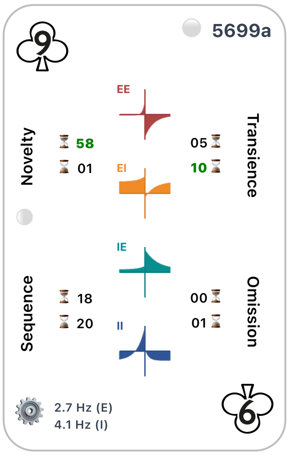
9
5699a
EE
EI
IE
II
Novelty
58
01
Transience
05
10
Sequence
18
20
Omission
00
01
2.7 Hz (E)
4.1 Hz (I)

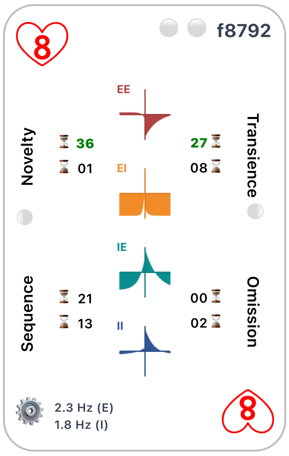
8
f8792
EE
EI
IE
II
Novelty
36
01
Transience
27
08
Sequence
21
13
Omission
00
02
2.3 Hz (E)
1.8 Hz (I)

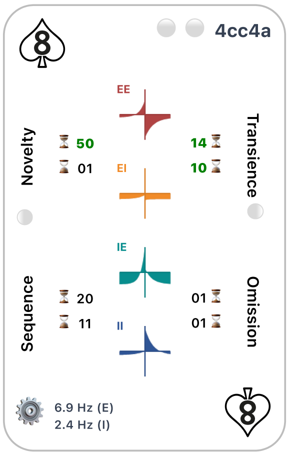
8
4cc4a
EE
EI
IE
II
Novelty
50
01
Transience
14
10
Sequence
20
11
Omission
01
01
6.9 Hz (E)
2.4 Hz (I)

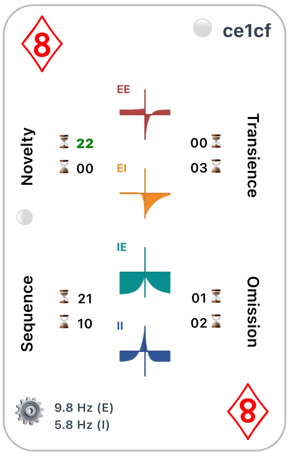
8
ce1cf
EE
EI
IE
II
Novelty
22
00
Transience
00
03
Sequence
21
10
Omission
01
02
9.8 Hz (E)
5.8 Hz (I)

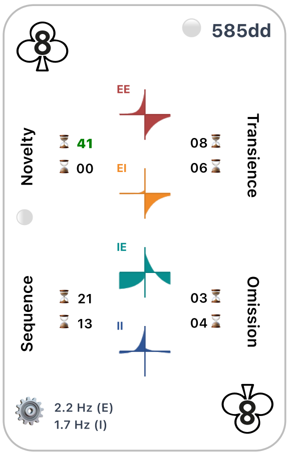
8
585dd
EE
EI
IE
II
Novelty
41
00
Transience
08
06
Sequence
21
13
Omission
03
04
2.2 Hz (E)
1.7 Hz (I)

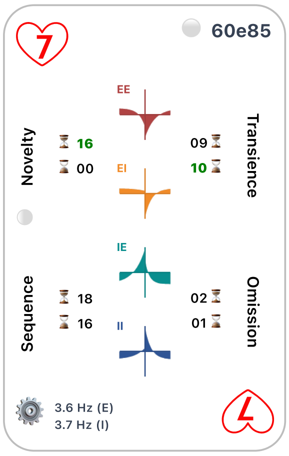
7
60e85
EE
EI
IE
II
Novelty
16
00
Transience
09
10
Sequence
18
16
Omission
02
01
3.6 Hz (E)
3.7 Hz (I)

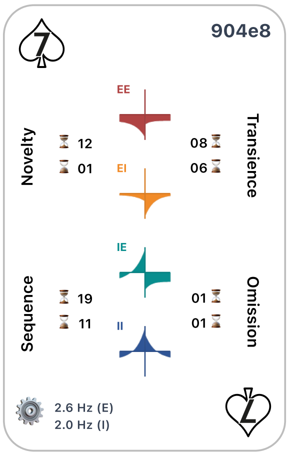
7
904e8
EE
EI
IE
II
Novelty
12
01
Transience
08
06
Sequence
19
11
Omission
01
01
2.6 Hz (E)
2.0 Hz (I)

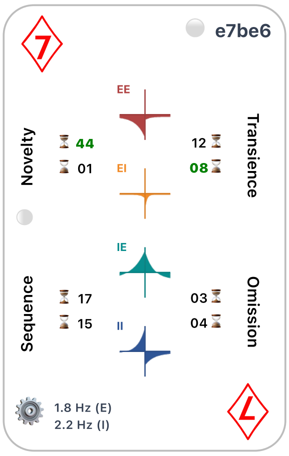
7
e7be6
EE
EI
IE
II
Novelty
44
01
Transience
12
08
Sequence
17
15
Omission
03
04
1.8 Hz (E)
2.2 Hz (I)

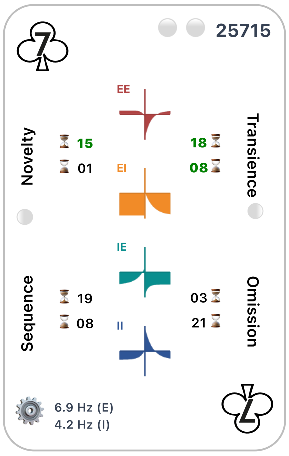
7
25715
EE
EI
IE
II
Novelty
15
01
Transience
18
08
Sequence
19
08
Omission
03
21
6.9 Hz (E)
4.2 Hz (I)

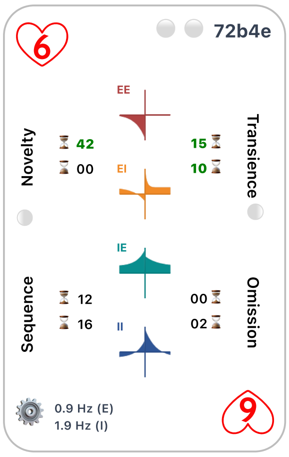
6
72b4e
EE
Novelty
42
00
15
10
Transience
EI
IE
Sequence
12
16
00
02
Omission
II
0.9 Hz (E)
1.9 Hz (I)

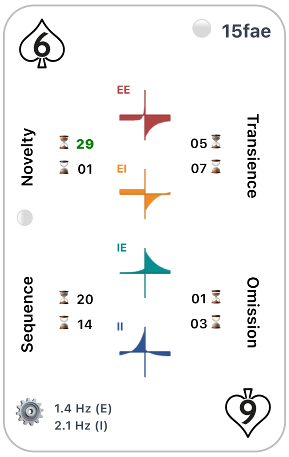
6
15fae
EE
Novelty
29
01
05
07
Transience
EI
IE
Sequence
20
14
01
03
Omission
II
1.4 Hz (E)
2.1 Hz (I)

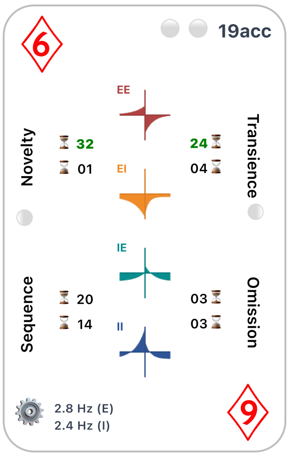
6
19acc
EE
Novelty
32
01
24
04
Transience
EI
IE
Sequence
20
14
03
03
Omission
II
2.8 Hz (E)
2.4 Hz (I)

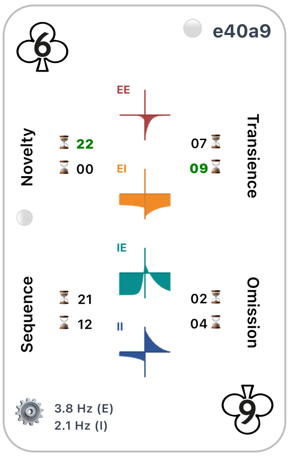
6
e40a9
EE
Novelty
22
00
07
09
Transience
EI
IE
Sequence
21
12
02
04
Omission
II
3.8 Hz (E)
2.1 Hz (I)

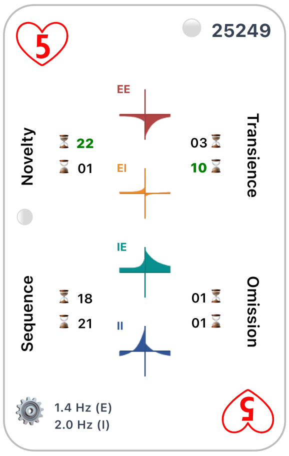
5
25249
EE
Novelty
22
01
03
10
Transience
EI
IE
Sequence
18
21
01
01
Omission
II
1.4 Hz (E)
2.0 Hz (I)

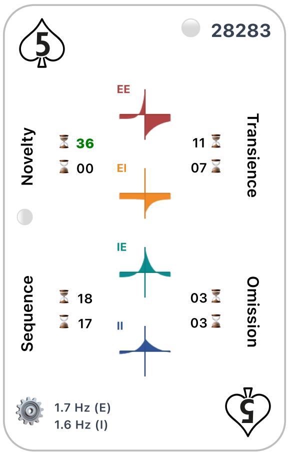
5
28283
EE
Novelty
36
00
11
07
Transience
EI
IE
Sequence
18
17
03
03
Omission
II
1.7 Hz (E)
1.6 Hz (I)

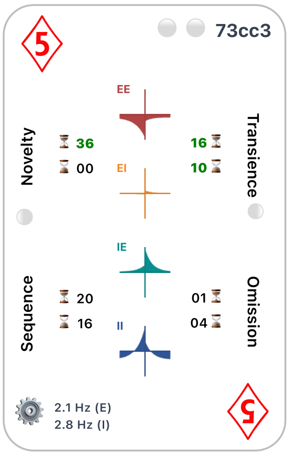
5
73cc3
EE
Novelty
36
00
16
10
Transience
EI
IE
Sequence
20
16
01
04
Omission
II
2.1 Hz (E)
2.8 Hz (I)

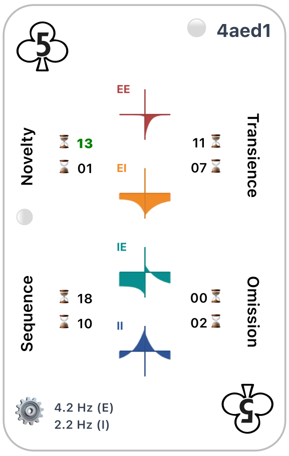
5
4aed1
EE
Novelty
13
01
11
07
Transience
EI
IE
Sequence
18
10
00
02
Omission
II
4.2 Hz (E)
2.2 Hz (I)

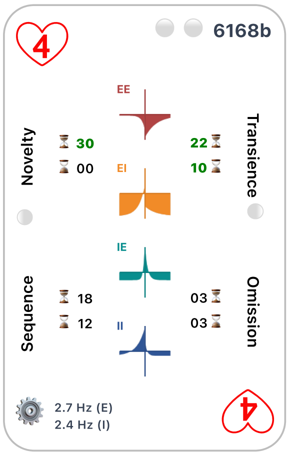
4
6168b
EE
Novelty
30
00
22
10
Transience
EI
IE
Sequence
18
12
03
03
Omission
II
2.7 Hz (E)
2.4 Hz (I)

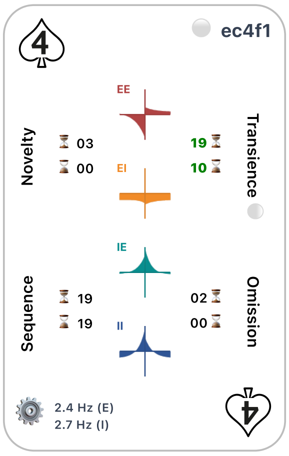
4
ec4f1
EE
Novelty
03
00
19
10
Transience
EI
IE
Sequence
19
19
02
00
Omission
II
2.4 Hz (E)
2.7 Hz (I)

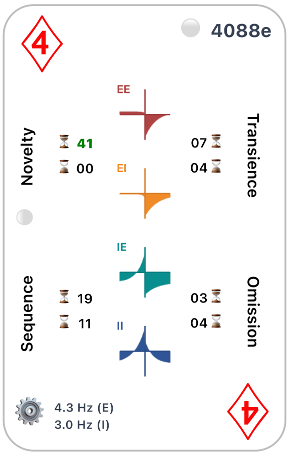
4
4088e
EE
Novelty
41
00
07
04
Transience
EI
IE
Sequence
19
11
03
04
Omission
II
4.3 Hz (E)
3.0 Hz (I)

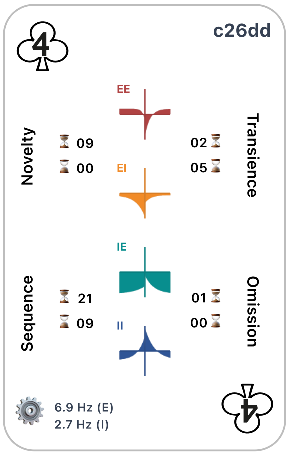
4
c26dd
EE
Novelty
09
00
02
05
Transience
EI
IE
Sequence
21
09
01
00
Omission
II
6.9 Hz (E)
2.7 Hz (I)

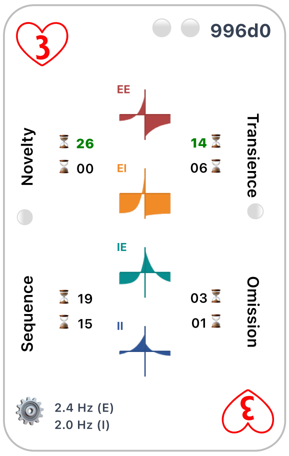
3
996d0
EE
Novelty
26
00
14
06
Transience
EI
IE
Sequence
19
15
03
01
Omission
II
2.4 Hz (E)
2.0 Hz (I)

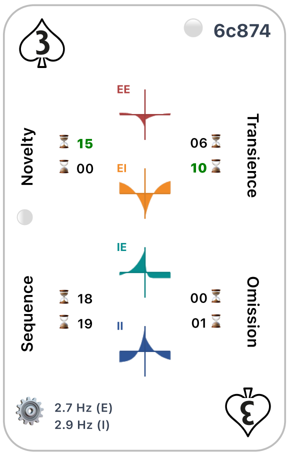
3
6c874
EE
Novelty
15
00
06
10
Transience
EI
IE
Sequence
18
19
00
01
Omission
II
2.7 Hz (E)
2.9 Hz (I)

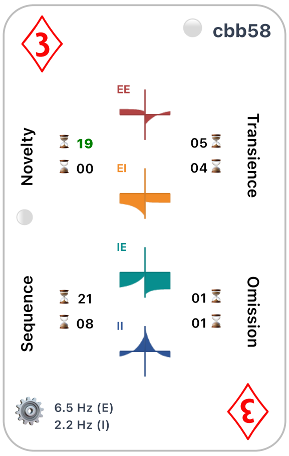
3
cbb58
EE
Novelty
19
00
05
04
Transience
EI
IE
Sequence
21
08
01
01
Omission
II
6.5 Hz (E)
2.2 Hz (I)

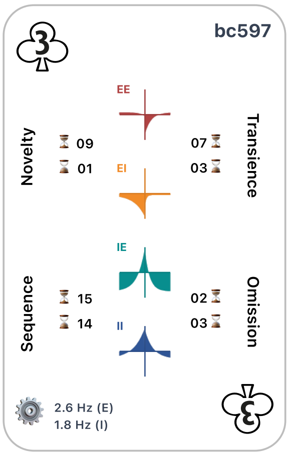
3
bc597
EE
Novelty
09
01
07
03
Transience
EI
IE
Sequence
15
14
02
03
Omission
II
2.6 Hz (E)
1.8 Hz (I)

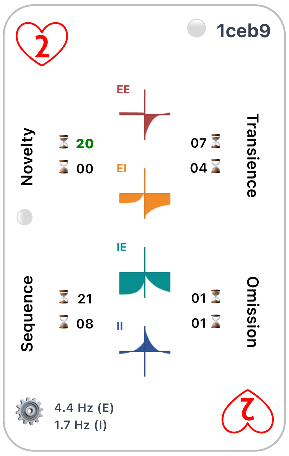
2
1ceb9
EE
Novelty
20
00
07
04
Transience
EI
IE
Sequence
21
08
01
01
Omission
II
4.4 Hz (E)
1.7 Hz (I)

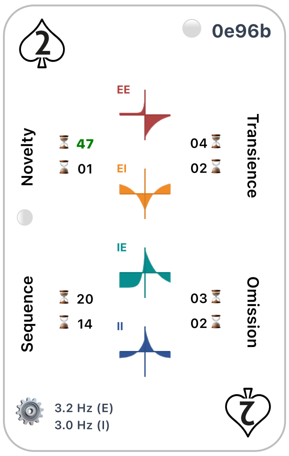
2
0e96b
EE
Novelty
47
01
04
02
Transience
EI
IE
Sequence
20
14
03
02
Omission
II
3.2 Hz (E)
3.0 Hz (I)

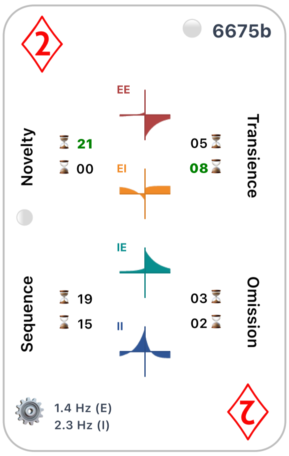
2
6675b
EE
Novelty
21
00
05
08
Transience
EI
IE
Sequence
19
15
03
02
Omission
II
1.4 Hz (E)
2.3 Hz (I)

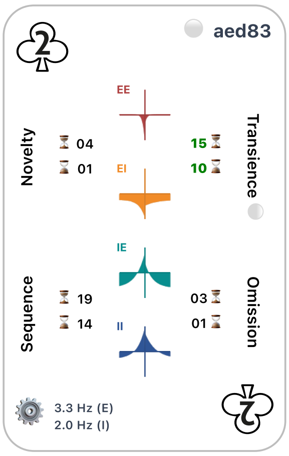
2
aed83
EE
Novelty
04
01
15
10
Transience
EI
IE
Sequence
19
14
03
01
Omission
II
3.3 Hz (E)
2.0 Hz (I)

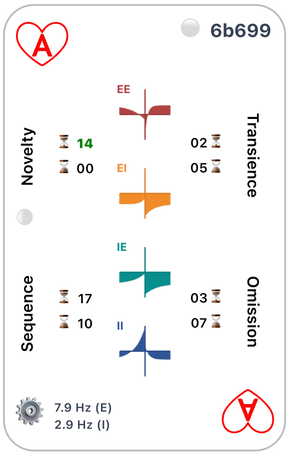
A
6b699
EE
Novelty
14
00
02
05
Transience
EI
IE
Sequence
17
10
03
07
Omission
II
7.9 Hz (E)
2.9 Hz (I)

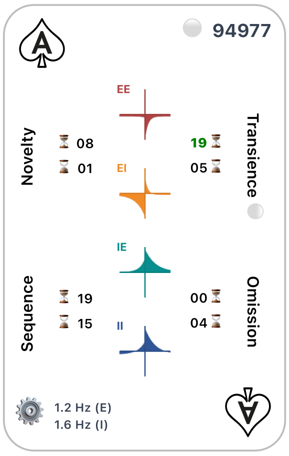
A
94977
EE
Novelty
08
01
19
05
Transience
EI
IE
Sequence
19
15
00
04
Omission
II
1.2 Hz (E)
1.6 Hz (I)

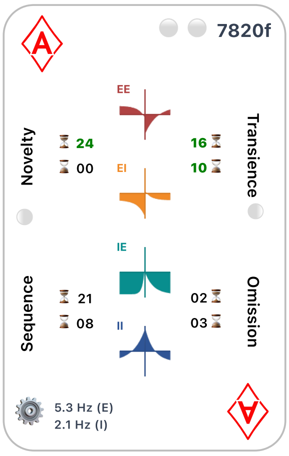
A
7820f
EE
Novelty
24
00
16
10
Transience
EI
IE
Sequence
21
08
02
03
Omission
II
5.3 Hz (E)
2.1 Hz (I)

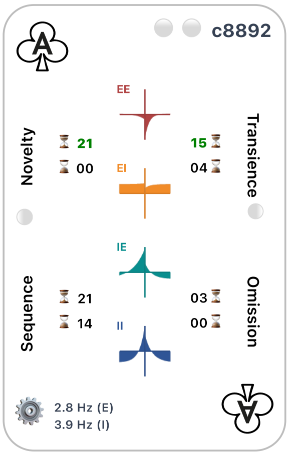
A
c8892
EE
Novelty
21
00
15
04
Transience
EI
IE
Sequence
21
14
03
00
Omission
II
2.8 Hz (E)
3.9 Hz (I)

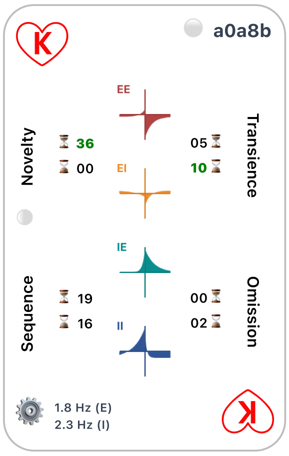
K
a0a8b
EE
Novelty
36
00
05
10
Transience
EI
IE
Sequence
19
16
00
02
Omission
II
1.8 Hz (E)
2.3 Hz (I)

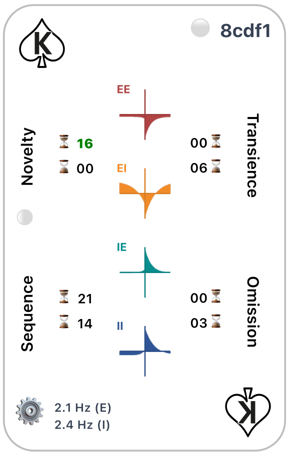
K
8cdf1
EE
Novelty
16
00
00
06
Transience
EI
IE
Sequence
21
14
00
03
Omission
II
2.1 Hz (E)
2.4 Hz (I)

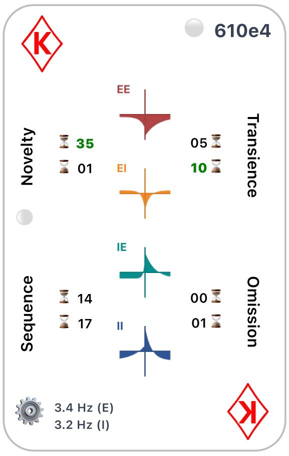
K
610e4
EE
Novelty
35
01
05
10
Transience
EI
IE
Sequence
14
17
00
01
Omission
II
3.4 Hz (E)
3.2 Hz (I)

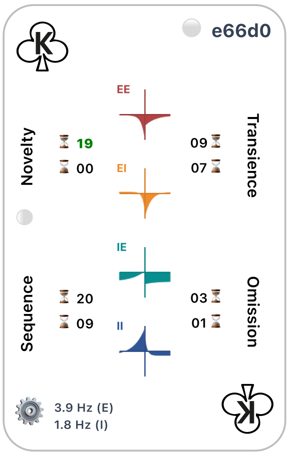
K
e66d0
EE
Novelty
19
00
09
07
Transience
EI
IE
Sequence
20
09
03
01
Omission
II
3.9 Hz (E)
1.8 Hz (I)

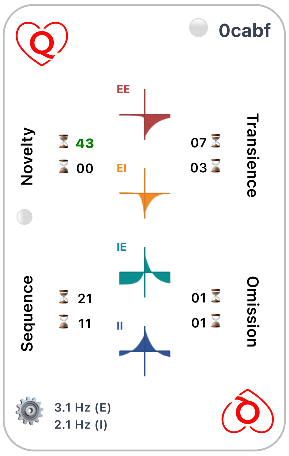
Q
0cabf
EE
Novelty
43
00
07
03
Transience
EI
IE
Sequence
21
11
01
01
Omission
II
3.1 Hz (E)
2.1 Hz (I)

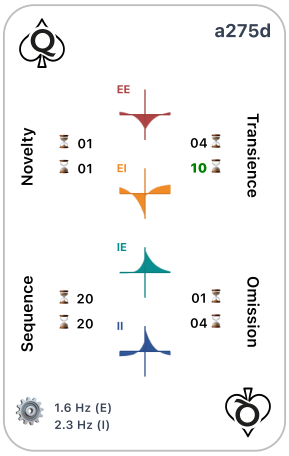
Q
a275d
EE
Novelty
01
01
04
10
Transience
EI
IE
Sequence
20
20
01
04
Omission
II
1.6 Hz (E)
2.3 Hz (I)

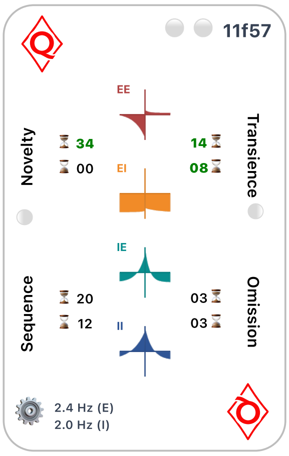
Q
11f57
EE
Novelty
34
00
14
08
Transience
EI
IE
Sequence
20
12
03
03
Omission
II
2.4 Hz (E)
2.0 Hz (I)

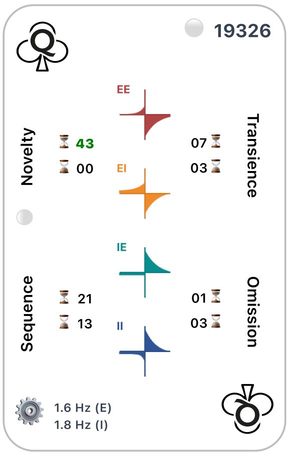
Q
19326
EE
Novelty
43
00
07
03
Transience
EI
IE
Sequence
21
13
01
03
Omission
II
1.6 Hz (E)
1.8 Hz (I)

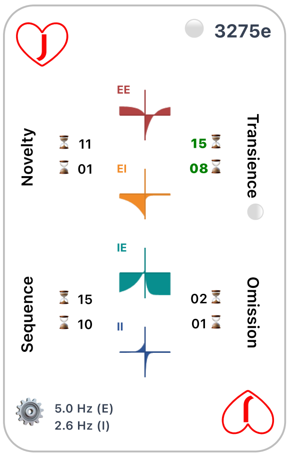
J
3275e
EE
EI
IE
II
Novelty
11
01
Transience
15
08
Sequence
15
10
Omission
02
01
5.0 Hz (E)
2.6 Hz (I)

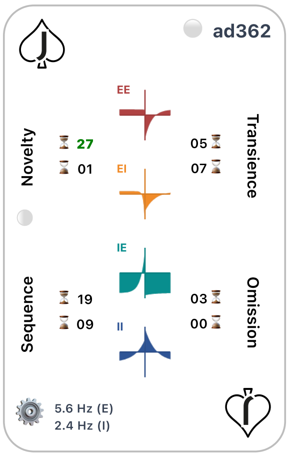
J
ad362
EE
EI
IE
II
Novelty
27
01
Transience
05
07
Sequence
19
09
Omission
03
00
5.6 Hz (E)
2.4 Hz (I)

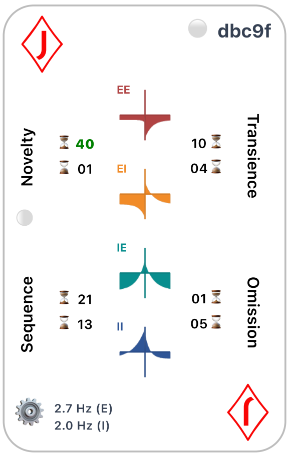
J
dbc9f
EE
EI
IE
II
Novelty
40
01
Transience
10
04
Sequence
21
13
Omission
01
05
2.7 Hz (E)
2.0 Hz (I)

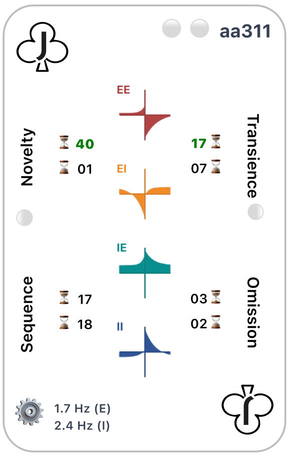
J
aa311
EE
EI
IE
II
Novelty
40
01
Transience
17
07
Sequence
17
18
Omission
03
02
1.7 Hz (E)
2.4 Hz (I)

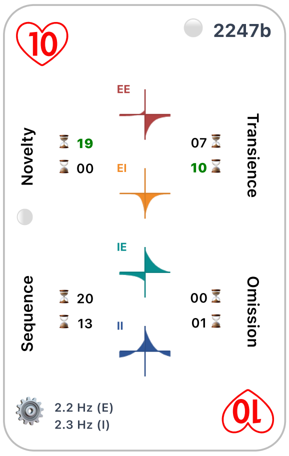
10
2247b
EE
EI
IE
II
Novelty
19
00
Transience
07
10
Sequence
20
13
Omission
00
01
2.2 Hz (E)
2.3 Hz (I)

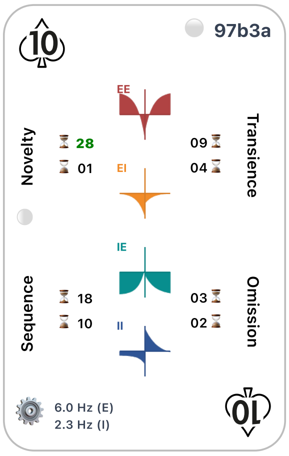
10
97b3a
EE
EI
IE
II
Novelty
28
01
Transience
09
04
Sequence
18
10
Omission
03
02
6.0 Hz (E)
2.3 Hz (I)

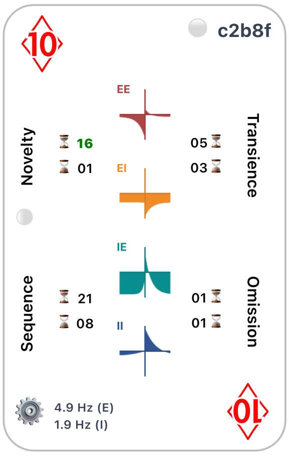
10
c2b8f
EE
EI
IE
II
Novelty
16
01
Transience
05
03
Sequence
21
08
Omission
01
01
4.9 Hz (E)
1.9 Hz (I)

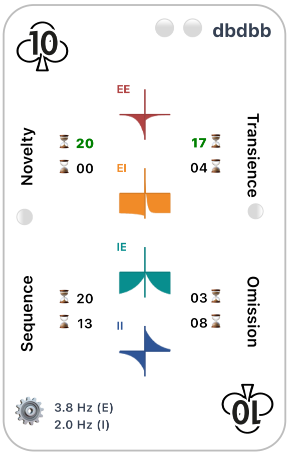
10
dbdbb
EE
EI
IE
II
Novelty
20
00
Transience
17
04
Sequence
20
13
Omission
03
08
3.8 Hz (E)
2.0 Hz (I)

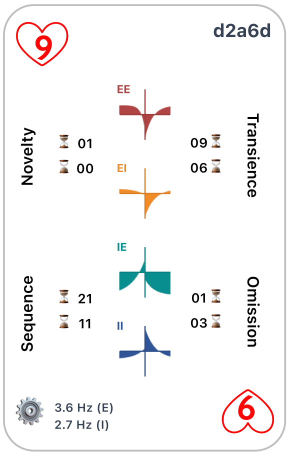
9
d2a6d
EE
EI
IE
II
Novelty
01
00
Transience
09
06
Sequence
21
11
Omission
01
03
3.6 Hz (E)
2.7 Hz (I)

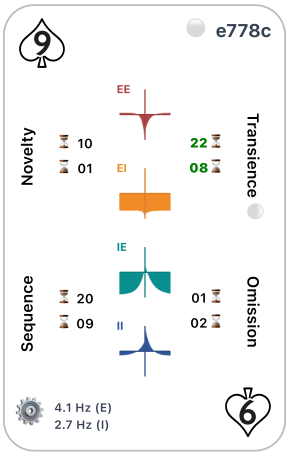
9
e778c
EE
EI
IE
II
Novelty
10
01
Transience
22
08
Sequence
20
09
Omission
01
02
4.1 Hz (E)
2.7 Hz (I)

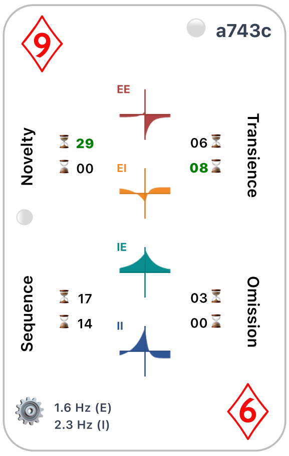
9
a743c
EE
EI
IE
II
Novelty
29
00
Transience
06
08
Sequence
17
14
Omission
03
00
1.6 Hz (E)
2.3 Hz (I)

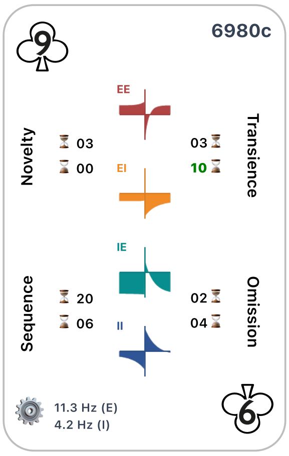
9
6980c
EE
EI
IE
II
Novelty
03
00
Transience
03
10
Sequence
20
06
Omission
02
04
11.3 Hz (E)
4.2 Hz (I)

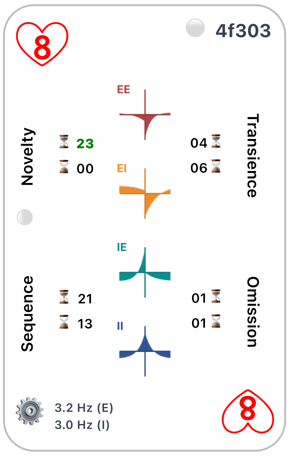
8
4f303
EE
EI
IE
II
Novelty
23
00
Transience
04
06
Sequence
21
13
Omission
01
01
3.2 Hz (E)
3.0 Hz (I)

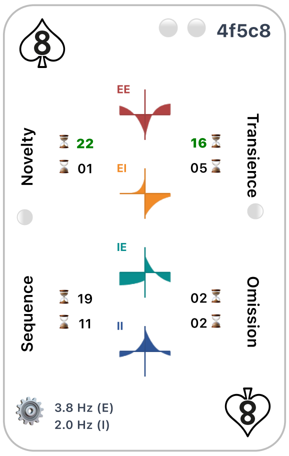
8
4f5c8
EE
EI
IE
II
Novelty
22
01
Transience
16
05
Sequence
19
11
Omission
02
02
3.8 Hz (E)
2.0 Hz (I)

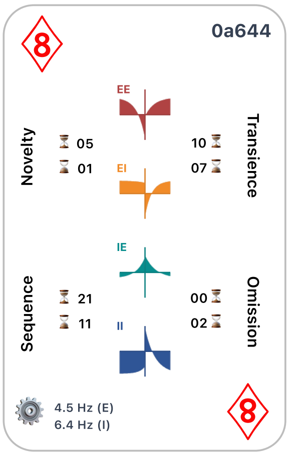
8
0a644
EE
EI
IE
II
Novelty
05
01
Transience
10
07
Sequence
21
11
Omission
00
02
4.5 Hz (E)
6.4 Hz (I)

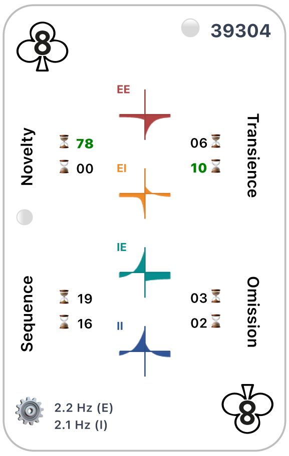
8
39304
EE
EI
IE
II
Novelty
78
00
Transience
06
10
Sequence
19
16
Omission
03
02
2.2 Hz (E)
2.1 Hz (I)

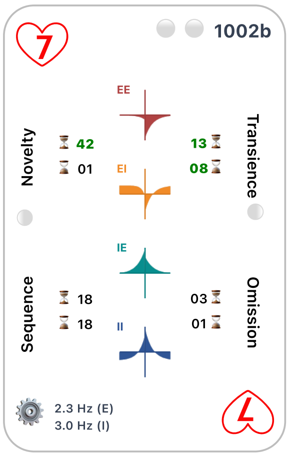
7
1002b
Novelty
42
01
EE
EI
13
08
Transience
Sequence
18
18
IE
II
03
01
Omission
2.3 Hz (E)
3.0 Hz (I)

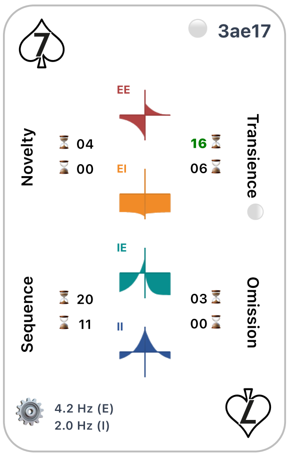
7
3ae17
Novelty
04
00
EE
EI
16
06
Transience
Sequence
20
11
IE
II
03
00
Omission
4.2 Hz (E)
2.0 Hz (I)

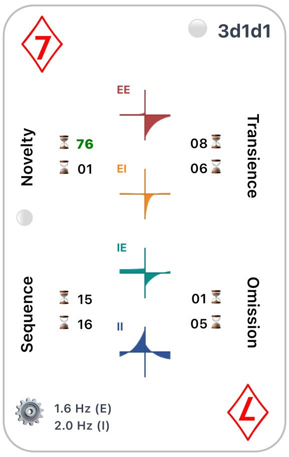
7
3d1d1
Novelty
76
01
EE
EI
08
06
Transience
Sequence
15
16
IE
II
01
05
Omission
1.6 Hz (E)
2.0 Hz (I)

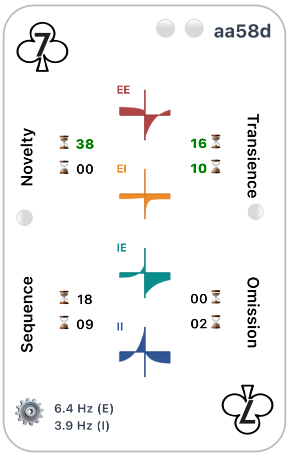
7
aa58d
Novelty
38
00
EE
EI
16
10
Transience
Sequence
18
09
IE
II
00
02
Omission
6.4 Hz (E)
3.9 Hz (I)

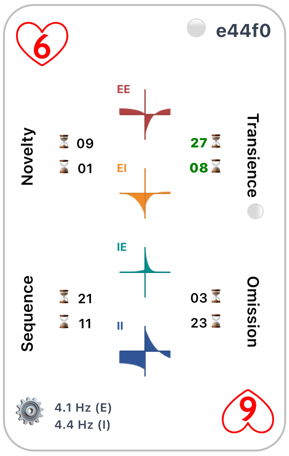
6
e44f0
Novelty
09
01
EE
EI
27
08
Transience
Sequence
21
11
IE
II
03
23
Omission
4.1 Hz (E)
4.4 Hz (I)

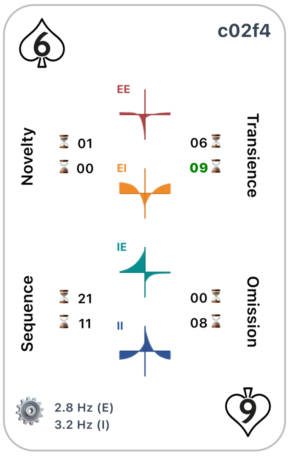
6
c02f4
Novelty
01
00
EE
EI
06
09
Transience
Sequence
21
11
IE
II
00
08
Omission
2.8 Hz (E)
3.2 Hz (I)

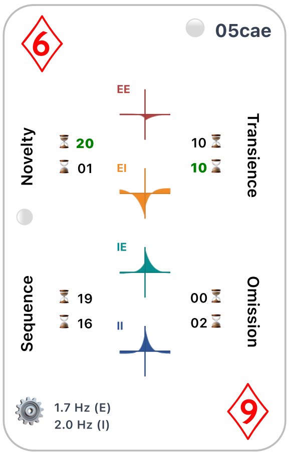
6
05cae
Novelty
20
01
EE
EI
10
10
Transience
Sequence
19
16
IE
II
00
02
Omission
1.7 Hz (E)
2.0 Hz (I)

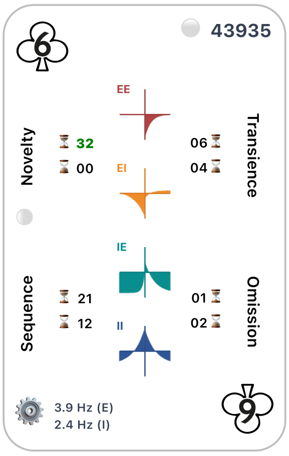
6
43935
Novelty
32
00
EE
EI
06
04
Transience
Sequence
21
12
IE
II
01
02
Omission
3.9 Hz (E)
2.4 Hz (I)

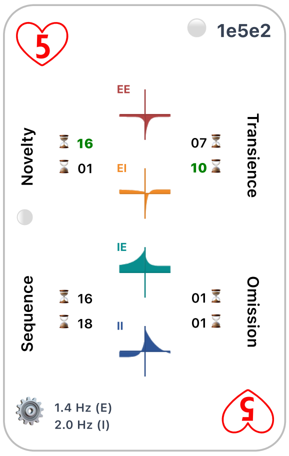
5
1e5e2
Novelty
16
01
EE
EI
07
10
Transience
Sequence
16
18
IE
II
01
01
Omission
1.4 Hz (E)
2.0 Hz (I)

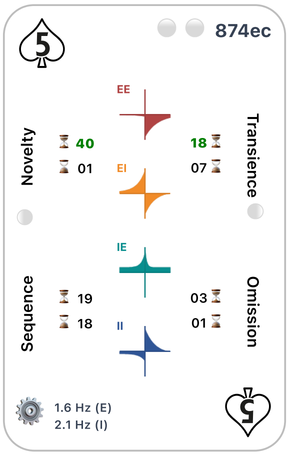
5
874ec
Novelty
40
01
EE
EI
18
07
Transience
Sequence
19
18
IE
II
03
01
Omission
1.6 Hz (E)
2.1 Hz (I)

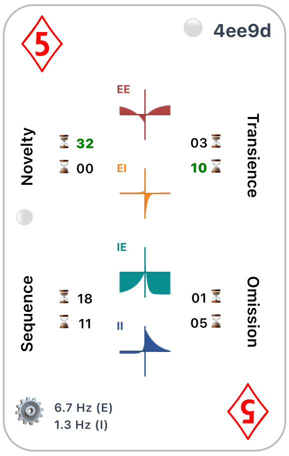
5
4ee9d
Novelty
32
00
EE
EI
03
10
Transience
Sequence
18
11
IE
II
01
05
Omission
6.7 Hz (E)
1.3 Hz (I)

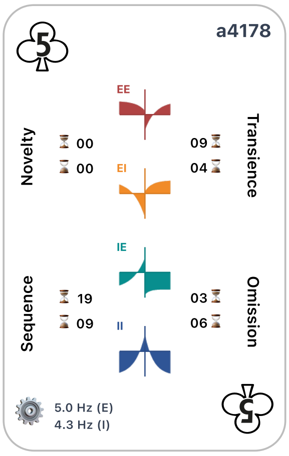
5
a4178
Novelty
00
00
EE
EI
09
04
Transience
Sequence
19
09
IE
II
03
06
Omission
5.0 Hz (E)
4.3 Hz (I)

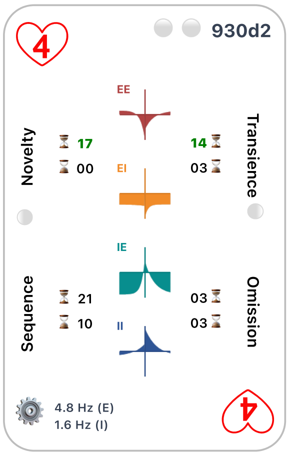
4
930d2
Novelty
17
00
EE
EI
14
03
Transience
Sequence
21
10
IE
II
03
03
Omission
4.8 Hz (E)
1.6 Hz (I)

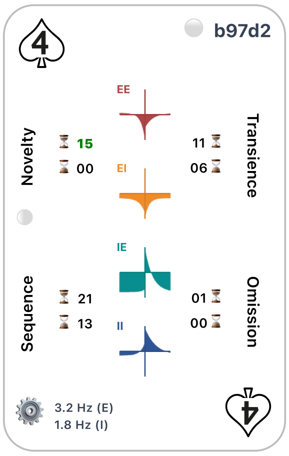
4
b97d2
Novelty
15
00
EE
EI
11
06
Transience
Sequence
21
13
IE
II
01
00
Omission
3.2 Hz (E)
1.8 Hz (I)

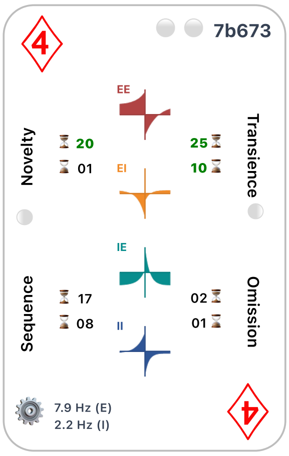
4
7b673
Novelty
20
01
EE
EI
25
10
Transience
Sequence
17
08
IE
II
02
01
Omission
7.9 Hz (E)
2.2 Hz (I)

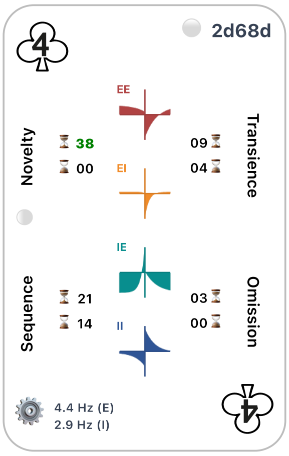
4
2d68d
Novelty
38
00
EE
EI
09
04
Transience
Sequence
21
14
IE
II
03
00
Omission
4.4 Hz (E)
2.9 Hz (I)

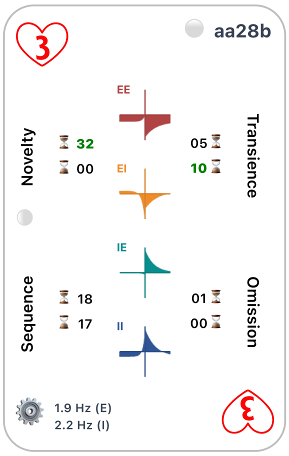
3
aa28b
Novelty
32
00
Transience
05
10
EE
EI
IE
II
Sequence
18
17
Omission
01
00
1.9 Hz (E)
2.2 Hz (I)

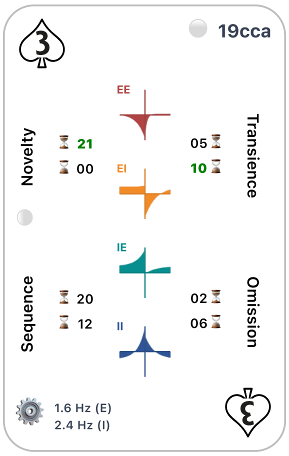
3
19cca
Novelty
21
00
Transience
05
10
EE
EI
IE
II
Sequence
20
12
Omission
02
06
1.6 Hz (E)
2.4 Hz (I)

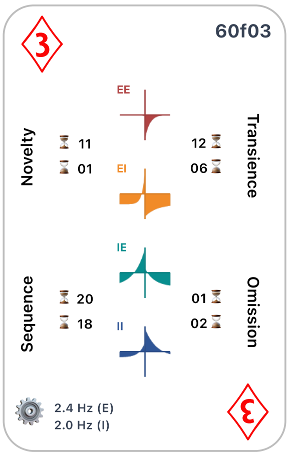
3
60f03
Novelty
11
01
Transience
12
06
EE
EI
IE
II
Sequence
20
18
Omission
01
02
2.4 Hz (E)
2.0 Hz (I)

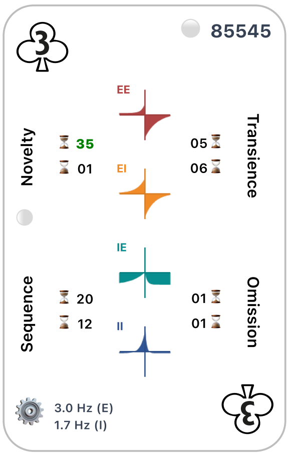
3
85545
Novelty
35
01
Transience
05
06
EE
EI
IE
II
Sequence
20
12
Omission
01
01
3.0 Hz (E)
1.7 Hz (I)

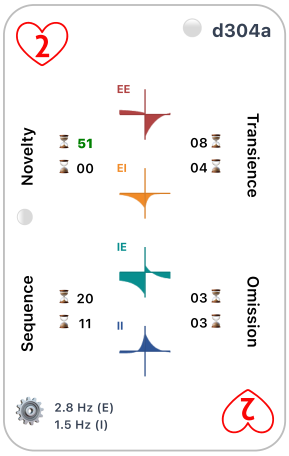
2
d304a
Novelty
51
00
Transience
08
04
EE
EI
IE
II
Sequence
20
11
Omission
03
03
2.8 Hz (E)
1.5 Hz (I)

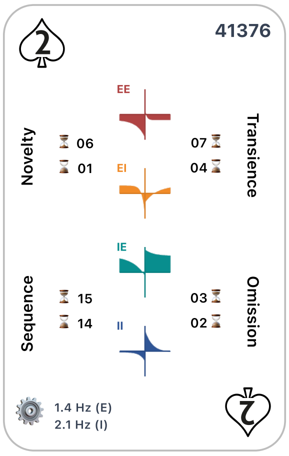
2
41376
Novelty
06
01
Transience
07
04
EE
EI
IE
II
Sequence
15
14
Omission
03
02
1.4 Hz (E)
2.1 Hz (I)

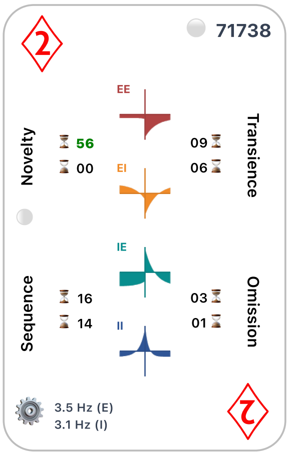
2
71738
Novelty
56
00
Transience
09
06
EE
EI
IE
II
Sequence
16
14
Omission
03
01
3.5 Hz (E)
3.1 Hz (I)

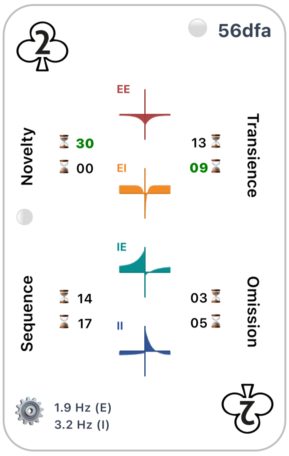
2
56dfa
Novelty
30
00
Transience
13
09
EE
EI
IE
II
Sequence
14
17
Omission
03
05
1.9 Hz (E)
3.2 Hz (I)

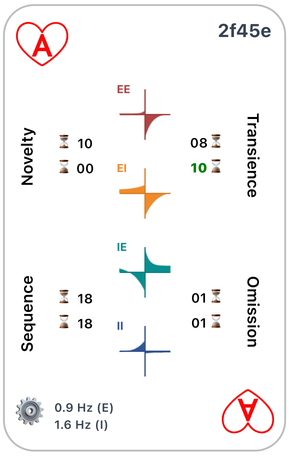
A
2f45e
Novelty
10
00
Transience
08
10
EE
EI
IE
II
Sequence
18
18
Omission
01
01
0.9 Hz (E)
1.6 Hz (I)

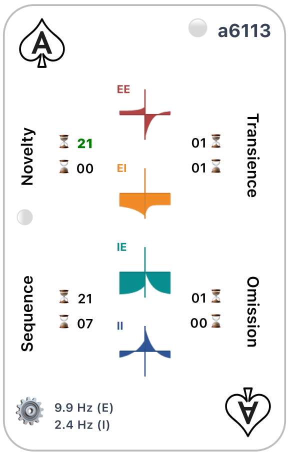
A
a6113
Novelty
21
00
Transience
01
01
EE
EI
IE
II
Sequence
21
07
Omission
01
00
9.9 Hz (E)
2.4 Hz (I)

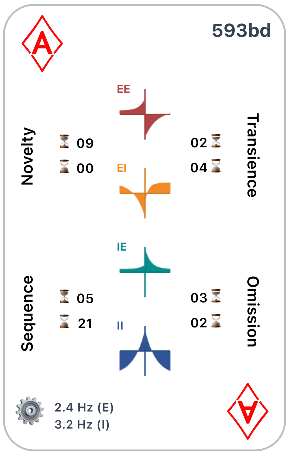
A
593bd
Novelty
09
00
Transience
02
04
EE
EI
IE
II
Sequence
05
21
Omission
03
02
2.4 Hz (E)
3.2 Hz (I)

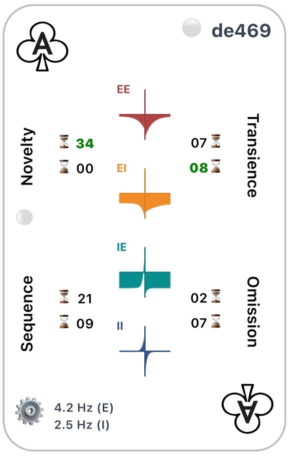
A
de469
Novelty
34
00
Transience
07
08
EE
EI
IE
II
Sequence
21
09
Omission
02
07
4.2 Hz (E)
2.5 Hz (I)

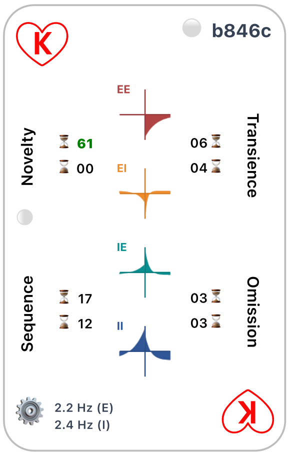
K
b846c
Novelty
61
00
Transience
06
04
EE
EI
IE
II
Sequence
17
12
Omission
03
03
2.2 Hz (E)
2.4 Hz (I)

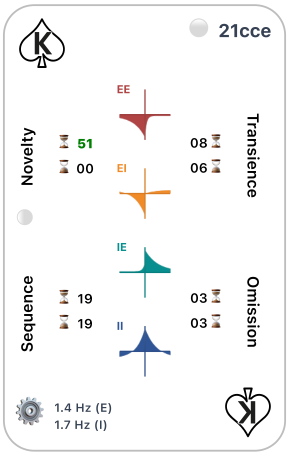
K
21cce
Novelty
51
00
Transience
08
06
EE
EI
IE
II
Sequence
19
19
Omission
03
03
1.4 Hz (E)
1.7 Hz (I)

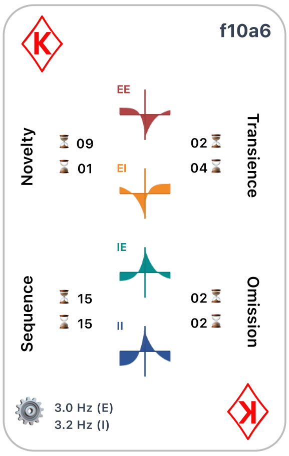
K
f10a6
Novelty
09
01
Transience
02
04
EE
EI
IE
II
Sequence
15
15
Omission
02
02
3.0 Hz (E)
3.2 Hz (I)

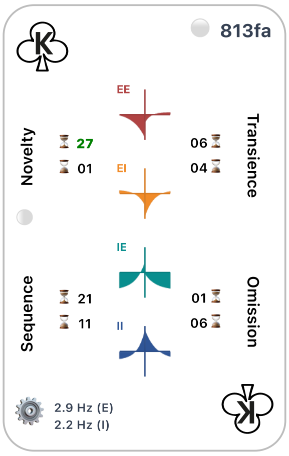
K
813fa
Novelty
27
01
Transience
06
04
EE
EI
IE
II
Sequence
21
11
Omission
01
06
2.9 Hz (E)
2.2 Hz (I)

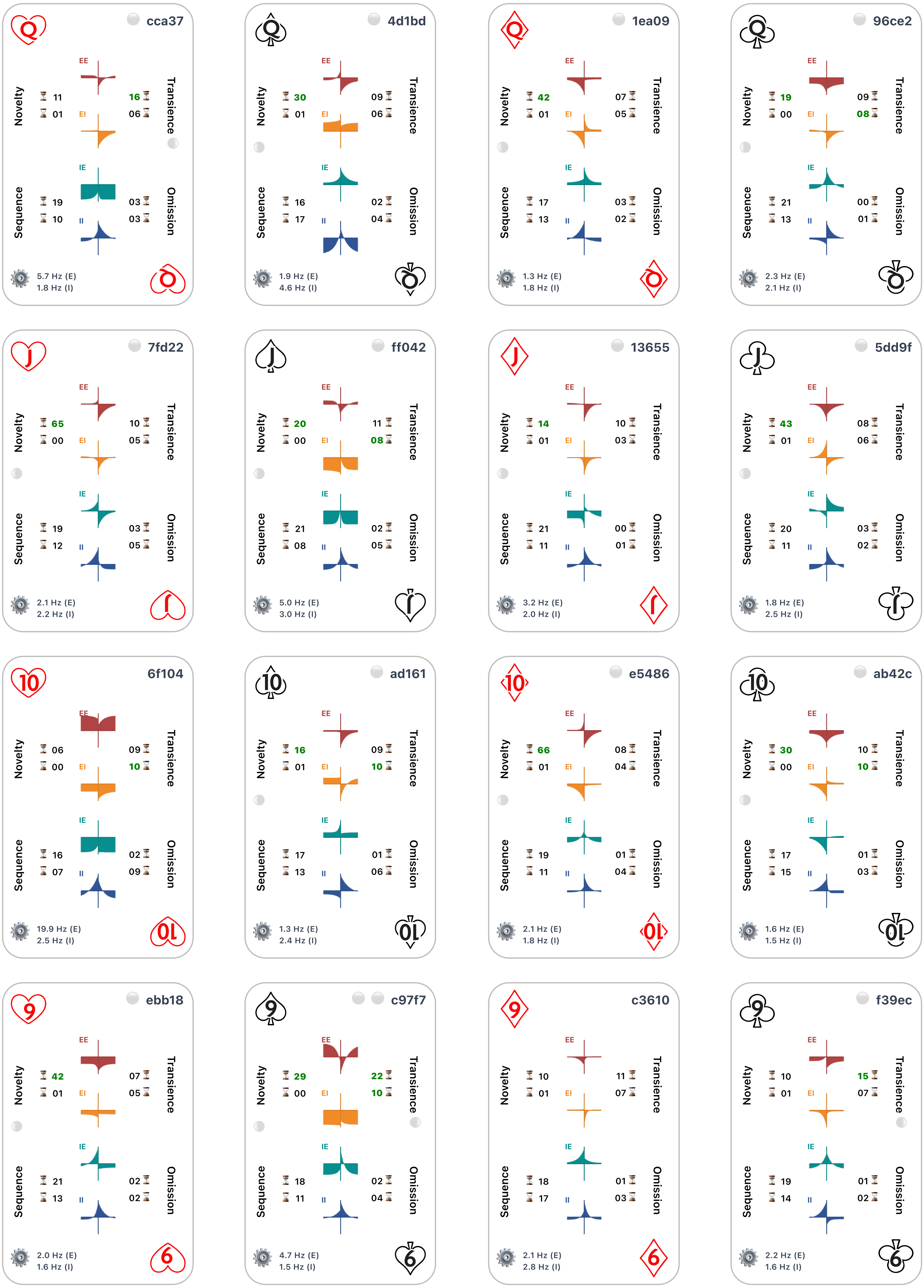
Q cca37
EE EI IE II
Novelty 11 01
Transience 16 06
Sequence 19 10
Omission 03 03
5.7 Hz (E) 1.8 Hz (I)
Q 4d1bd
Novelty 30 01
Transience 09 06
Sequence 16 17
Omission 02 04
1.9 Hz (E) 4.6 Hz (I)
Q 1ea09
Novelty 42 01
Transience 07 05
Sequence 17 13
Omission 03 02
1.3 Hz (E) 1.8 Hz (I)
Q 96ce2
Novelty 19 00
Transience 09 08
Sequence 21 13
Omission 00 01
2.3 Hz (E) 2.1 Hz (I)
J 7fd22
Novelty 65 00
Transience 10 05
Sequence 19 12
Omission 03 05
2.1 Hz (E) 2.2 Hz (I)
J ff042
Novelty 20 00
Transience 11 08
Sequence 21 08
Omission 02 05
5.0 Hz (E) 3.0 Hz (I)
J 13655
Novelty 14 01
Transience 10 03
Sequence 21 11
Omission 00 01
3.2 Hz (E) 2.0 Hz (I)
J 5dd9f
Novelty 43 01
Transience 08 06
Sequence 20 11
Omission 03 02
1.8 Hz (E) 2.5 Hz (I)
10 6f104
Novelty 06 00
Transience 09 10
Sequence 16 07
Omission 02 09
19.9 Hz (E) 2.5 Hz (I)
10 ad161
Novelty 16 01
Transience 09 10
Sequence 17 13
Omission 01 06
1.3 Hz (E) 2.4 Hz (I)
10 e5486
Novelty 66 01
Transience 08 04
Sequence 19 11
Omission 01 04
2.1 Hz (E) 1.8 Hz (I)
10 ab42c
Novelty 30 00
Transience 10 10
Sequence 17 15
Omission 01 03
1.6 Hz (E) 1.5 Hz (I)
9 ebb18
Novelty 42 01
Transience 07 05
Sequence 21 13
Omission 02 02
2.0 Hz (E) 1.6 Hz (I)
9 c97f7
Novelty 29 00
Transience 22 10
Sequence 18 11
Omission 02 04
4.7 Hz (E) 1.5 Hz (I)
9 c3610
Novelty 10 01
Transience 11 07
Sequence 18 17
Omission 01 03
2.1 Hz (E) 2.8 Hz (I)
9 f39ec
Novelty 10 01
Transience 15 07
Sequence 19 14
Omission 01 02
2.2 Hz (E) 1.6 Hz (I)

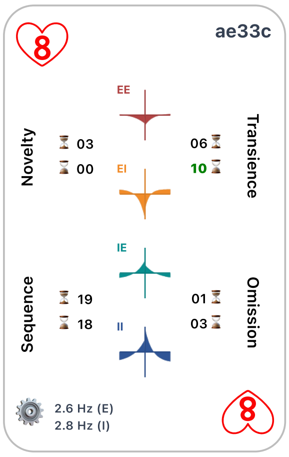
8
ae33c
EE
Novelty
03
00
06
10
Transience
EI
IE
Sequence
19
18
01
03
Omission
II
2.6 Hz (E)
2.8 Hz (I)

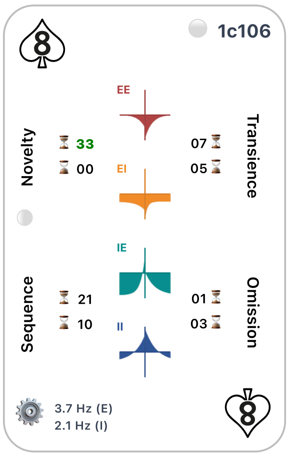
8
1c106
EE
Novelty
33
00
07
05
Transience
EI
IE
Sequence
21
10
01
03
Omission
II
3.7 Hz (E)
2.1 Hz (I)

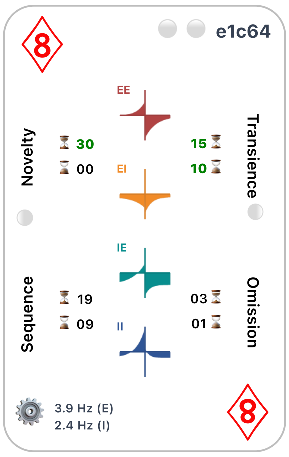
8
e1c64
EE
Novelty
30
00
15
10
Transience
EI
IE
Sequence
19
09
03
01
Omission
II
3.9 Hz (E)
2.4 Hz (I)

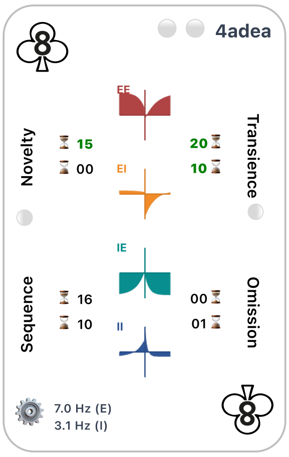
8
4adea
EE
Novelty
15
00
20
10
Transience
EI
IE
Sequence
16
10
00
01
Omission
II
7.0 Hz (E)
3.1 Hz (I)

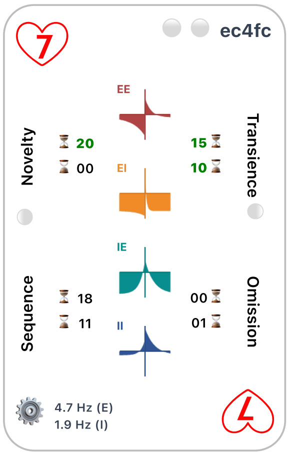
7
ec4fc
EE
Novelty
20
00
15
10
Transience
EI
IE
Sequence
18
11
00
01
Omission
II
4.7 Hz (E)
1.9 Hz (I)

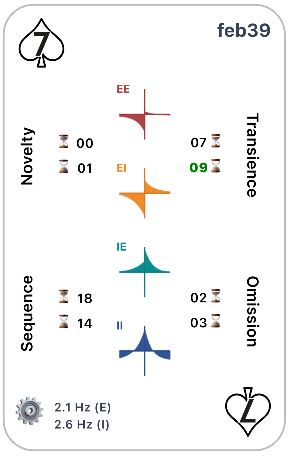
7
feb39
EE
Novelty
00
01
07
09
Transience
EI
IE
Sequence
18
14
02
03
Omission
II
2.1 Hz (E)
2.6 Hz (I)

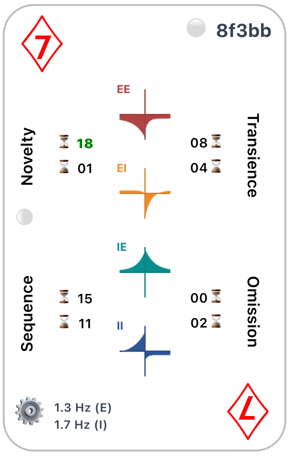
7
8f3bb
EE
Novelty
18
01
08
04
Transience
EI
IE
Sequence
15
11
00
02
Omission
II
1.3 Hz (E)
1.7 Hz (I)

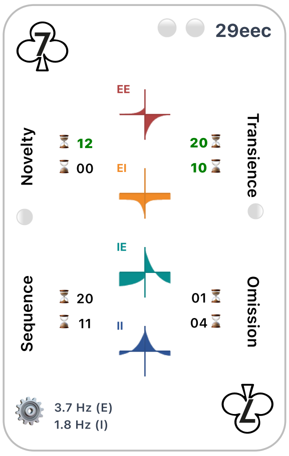
7
29eec
EE
Novelty
12
00
20
10
Transience
EI
IE
Sequence
20
11
01
04
Omission
II
3.7 Hz (E)
1.8 Hz (I)

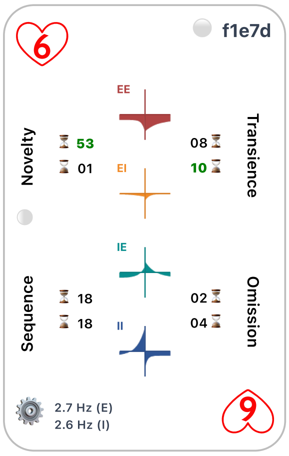
6
f1e7d
EE
Novelty
53
01
08
10
Transience
EI
IE
Sequence
18
18
02
04
Omission
II
2.7 Hz (E)
2.6 Hz (I)

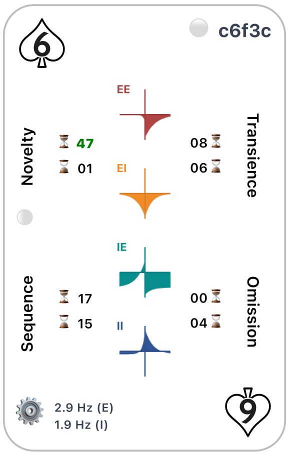
6
c6f3c
EE
Novelty
47
01
08
06
Transience
EI
IE
Sequence
17
15
00
04
Omission
II
2.9 Hz (E)
1.9 Hz (I)

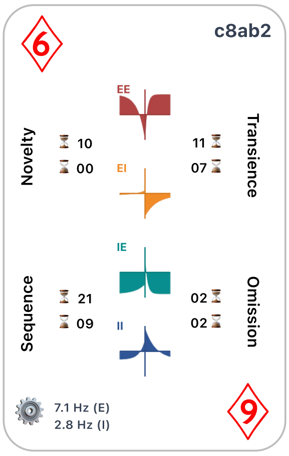
6
c8ab2
EE
Novelty
10
00
11
07
Transience
EI
IE
Sequence
21
09
02
02
Omission
II
7.1 Hz (E)
2.8 Hz (I)

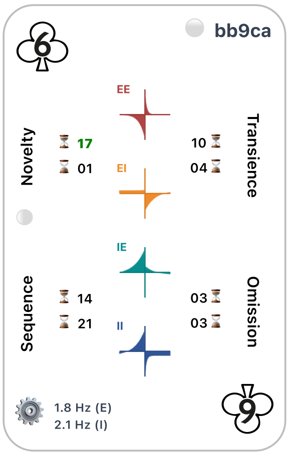
6
bb9ca
EE
Novelty
17
01
10
04
Transience
EI
IE
Sequence
14
21
03
03
Omission
II
1.8 Hz (E)
2.1 Hz (I)

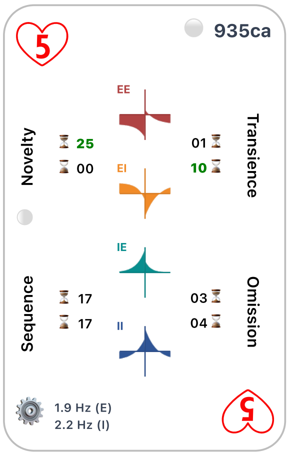
5
935ca
EE
Novelty
25
00
01
10
Transience
EI
IE
Sequence
17
17
03
04
Omission
II
1.9 Hz (E)
2.2 Hz (I)

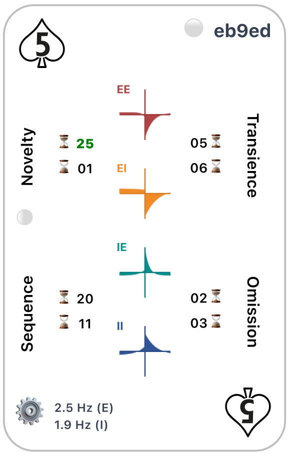
5
eb9ed
EE
Novelty
25
01
05
06
Transience
EI
IE
Sequence
20
11
02
03
Omission
II
2.5 Hz (E)
1.9 Hz (I)

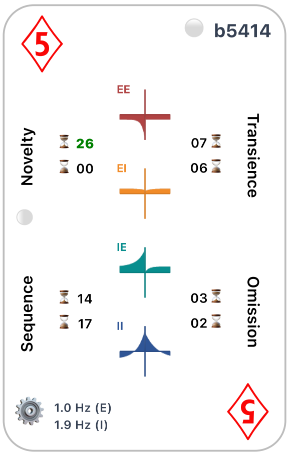
5
b5414
EE
Novelty
26
00
07
06
Transience
EI
IE
Sequence
14
17
03
02
Omission
II
1.0 Hz (E)
1.9 Hz (I)

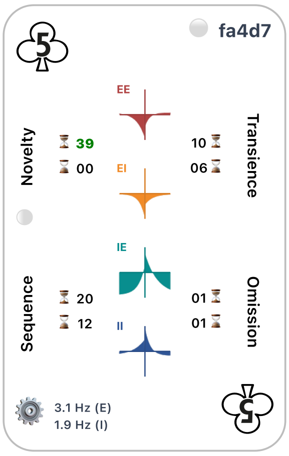
5
fa4d7
EE
Novelty
39
00
10
06
Transience
EI
IE
Sequence
20
12
01
01
Omission
II
3.1 Hz (E)
1.9 Hz (I)

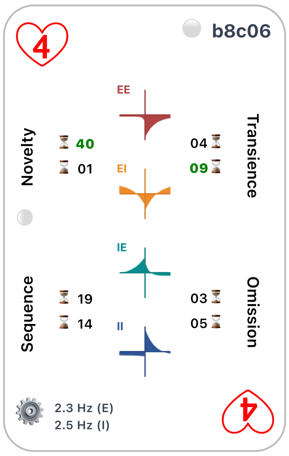
4
b8c06
EE
Novelty 40 01
04 09 Transience
EI
IE
Sequence 19 14
03 05 Omission
II
2.3 Hz (E)
2.5 Hz (I)

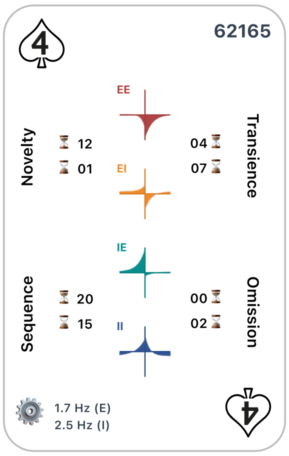
4
62165
EE
Novelty 12 01
04 07 Transience
EI
IE
Sequence 20 15
00 02 Omission
II
1.7 Hz (E)
2.5 Hz (I)

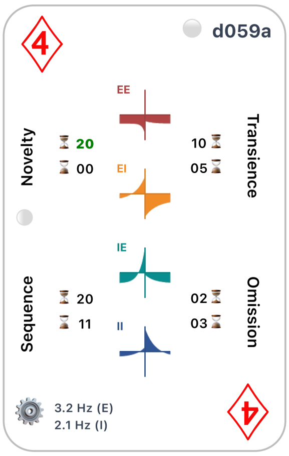
4
d059a
EE
Novelty 20 00
10 05 Transience
EI
IE
Sequence 20 11
02 03 Omission
II
3.2 Hz (E)
2.1 Hz (I)

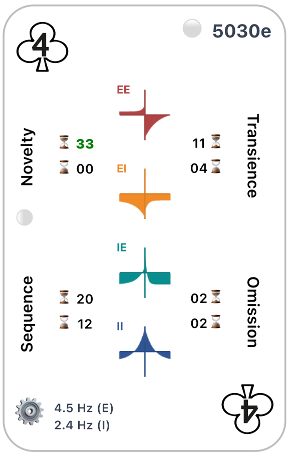
4
5030e
EE
Novelty 33 00
11 04 Transience
EI
IE
Sequence 20 12
02 02 Omission
II
4.5 Hz (E)
2.4 Hz (I)

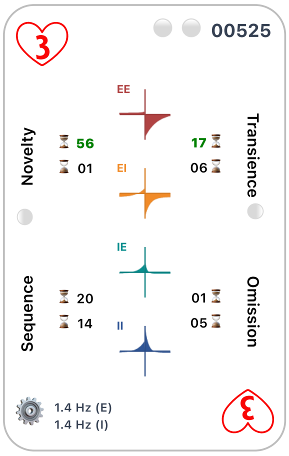
3
00525
EE
Novelty 56 01
17 06 Transience
EI
IE
Sequence 20 14
01 05 Omission
II
1.4 Hz (E)
1.4 Hz (I)

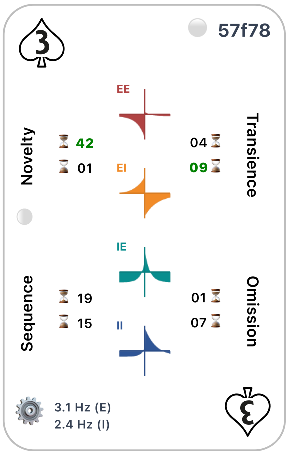
3
57f78
EE
Novelty 42 01
04 09 Transience
EI
IE
Sequence 19 15
01 07 Omission
II
3.1 Hz (E)
2.4 Hz (I)

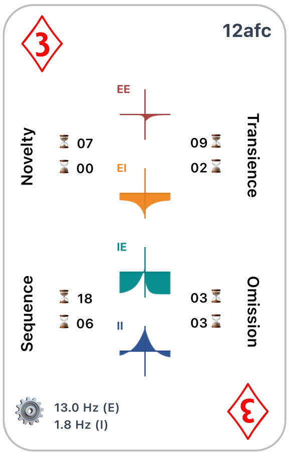
3
12afc
EE
Novelty 07 00
09 02 Transience
EI
IE
Sequence 18 06
03 03 Omission
II
13.0 Hz (E)
1.8 Hz (I)

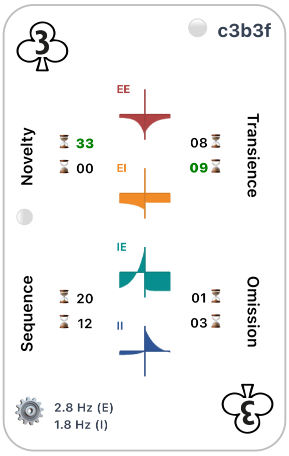
3
c3b3f
EE
Novelty 33 00
08 09 Transience
EI
IE
Sequence 20 12
01 03 Omission
II
2.8 Hz (E)
1.8 Hz (I)

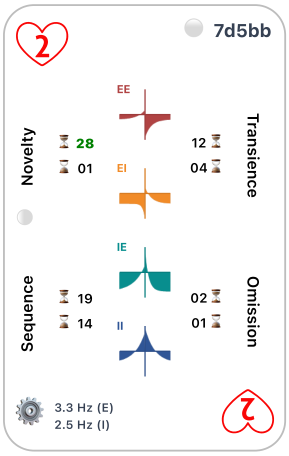
2
7d5bb
EE
Novelty 28 01
12 04 Transience
EI
IE
Sequence 19 14
02 01 Omission
II
3.3 Hz (E)
2.5 Hz (I)

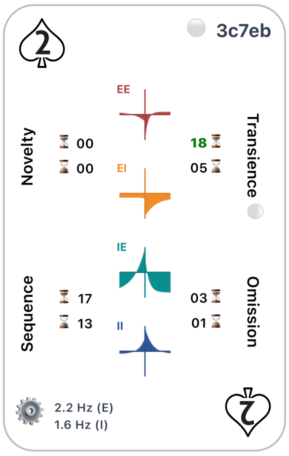
2
3c7eb
EE
Novelty 00 00
18 05 Transience
EI
IE
Sequence 17 13
03 01 Omission
II
2.2 Hz (E)
1.6 Hz (I)

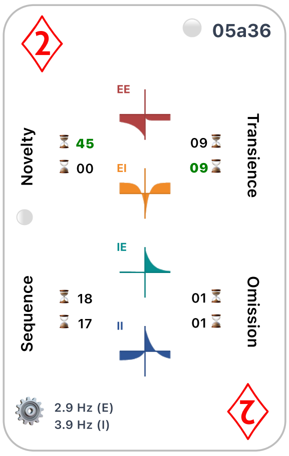
2
05a36
EE
Novelty 45 00
09 09 Transience
EI
IE
Sequence 18 17
01 01 Omission
II
2.9 Hz (E)
3.9 Hz (I)

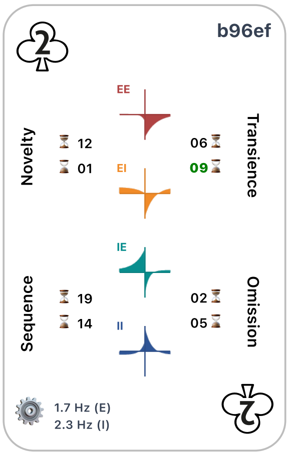
2
b96ef
EE
Novelty 12 01
06 09 Transience
EI
IE
Sequence 19 14
02 05 Omission
II
1.7 Hz (E)
2.3 Hz (I)

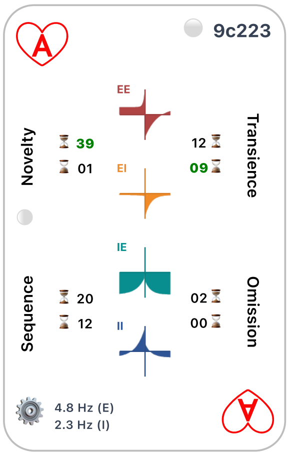
A
9c223
EE
Novelty 39 01
12 09 Transience
EI
IE
Sequence 20 12
02 00 Omission
II
4.8 Hz (E)
2.3 Hz (I)

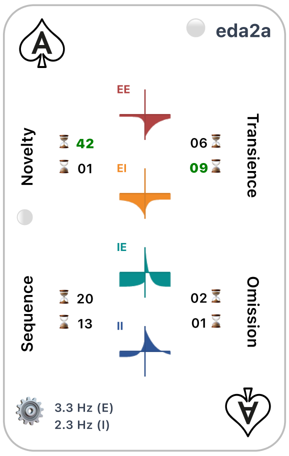
A
eda2a
EE
Novelty 42 01
06 09 Transience
EI
IE
Sequence 20 13
02 01 Omission
II
3.3 Hz (E)
2.3 Hz (I)

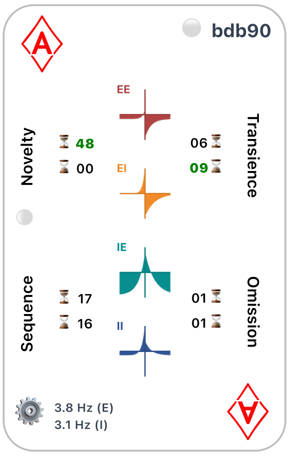
A
bdb90
EE
Novelty 48 00
06 09 Transience
EI
IE
Sequence 17 16
01 01 Omission
II
3.8 Hz (E)
3.1 Hz (I)

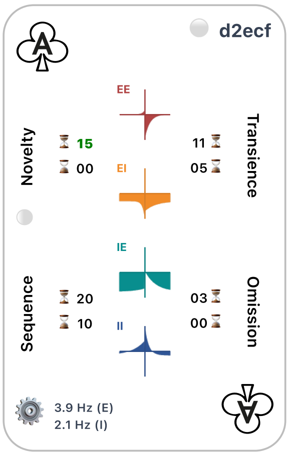
A
d2ecf
EE
Novelty 15 00
11 05 Transience
EI
IE
Sequence 20 10
03 00 Omission
II
3.9 Hz (E)
2.1 Hz (I)

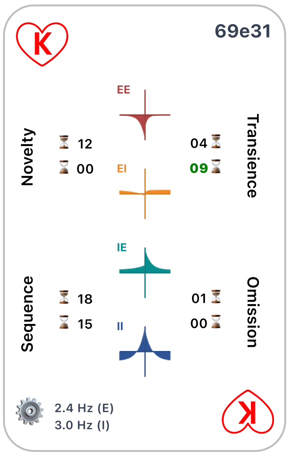
K
69e31
EE
EI
IE
II
Novelty
12
00
Transience
04
09
Sequence
18
15
Omission
01
00
2.4 Hz (E)
3.0 Hz (I)

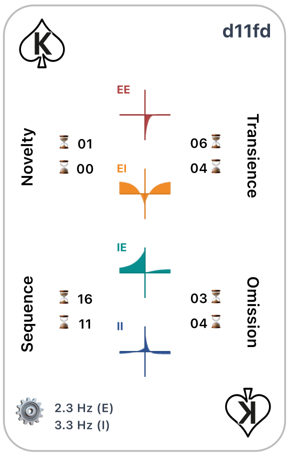
K
d11fd
EE
EI
IE
II
Novelty
01
00
Transience
06
04
Sequence
16
11
Omission
03
04
2.3 Hz (E)
3.3 Hz (I)

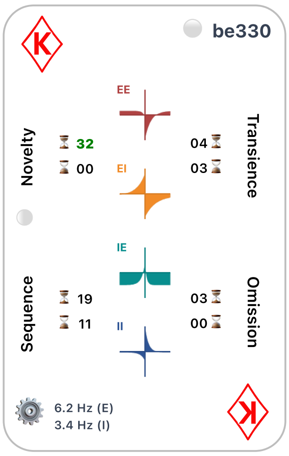
K
be330
EE
EI
IE
II
Novelty
32
00
Transience
04
03
Sequence
19
11
Omission
03
00
6.2 Hz (E)
3.4 Hz (I)

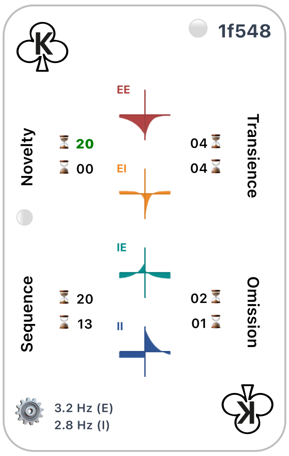
K
1f548
EE
EI
IE
II
Novelty
20
00
Transience
04
04
Sequence
20
13
Omission
02
01
3.2 Hz (E)
2.8 Hz (I)

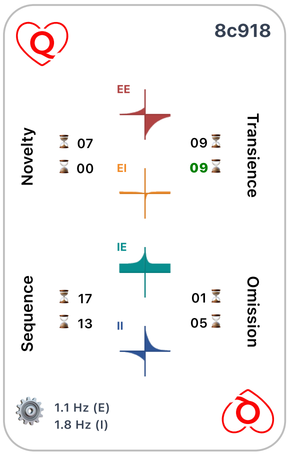
Q
8c918
EE
EI
IE
II
Novelty
07
00
Transience
09
09
Sequence
17
13
Omission
01
05
1.1 Hz (E)
1.8 Hz (I)

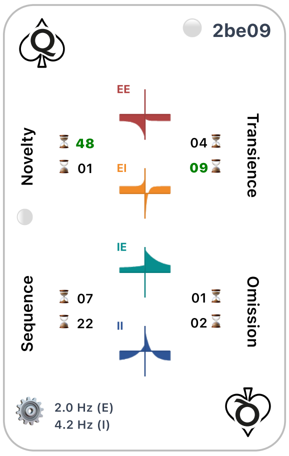
Q
2be09
EE
EI
IE
II
Novelty
48
01
Transience
04
09
Sequence
07
22
Omission
01
02
2.0 Hz (E)
4.2 Hz (I)

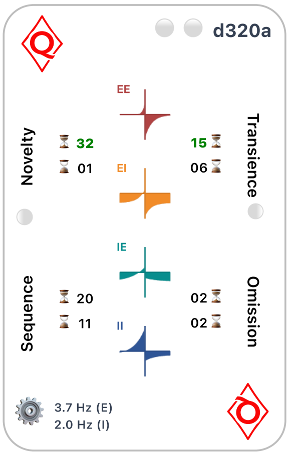
Q
d320a
EE
EI
IE
II
Novelty
32
01
Transience
15
06
Sequence
20
11
Omission
02
02
3.7 Hz (E)
2.0 Hz (I)

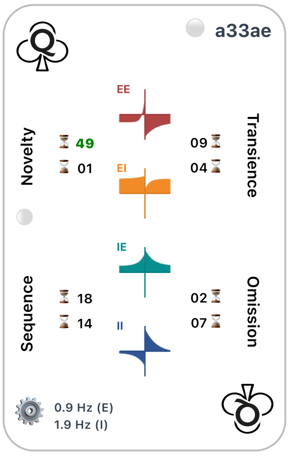
Q
a33ae
EE
EI
IE
II
Novelty
49
01
Transience
09
04
Sequence
18
14
Omission
02
07
0.9 Hz (E)
1.9 Hz (I)

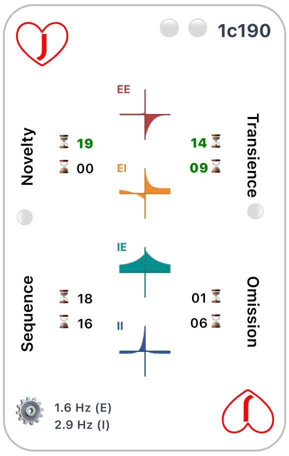
J
1c190
EE
EI
IE
II
Novelty
19
00
Transience
14
09
Sequence
18
16
Omission
01
06
1.6 Hz (E)
2.9 Hz (I)

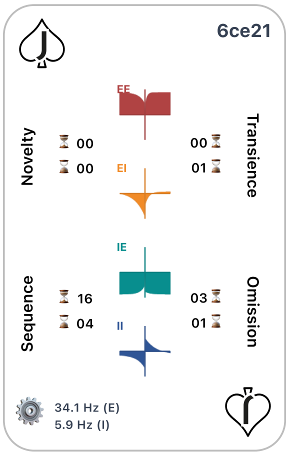
J
6ce21
EE
EI
IE
II
Novelty
00
00
Transience
00
01
Sequence
16
04
Omission
03
01
34.1 Hz (E)
5.9 Hz (I)

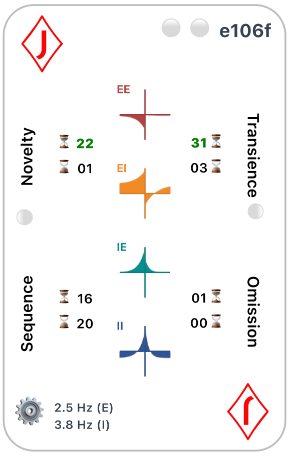
J
e106f
EE
EI
IE
II
Novelty
22
01
Transience
31
03
Sequence
16
20
Omission
01
00
2.5 Hz (E)
3.8 Hz (I)

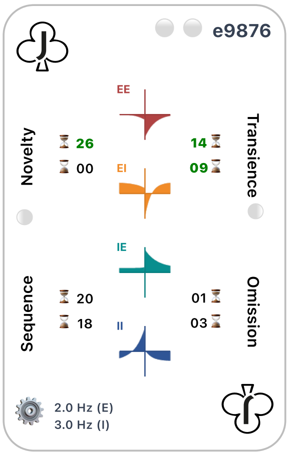
J
e9876
EE
EI
IE
II
Novelty
26
00
Transience
14
09
Sequence
20
18
Omission
01
03
2.0 Hz (E)
3.0 Hz (I)

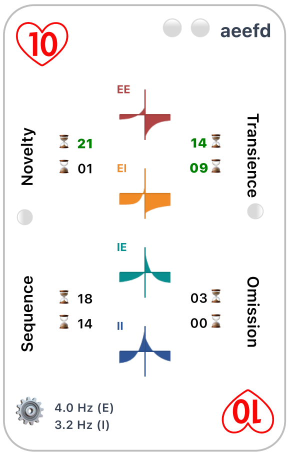
10
aeefd
EE
EI
IE
II
Novelty
21
01
Transience
14
09
Sequence
18
14
Omission
03
00
4.0 Hz (E)
3.2 Hz (I)

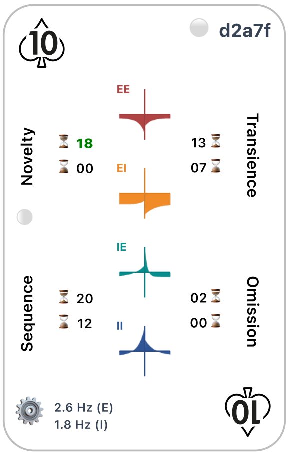
10
d2a7f
EE
EI
IE
II
Novelty
18
00
Transience
13
07
Sequence
20
12
Omission
02
00
2.6 Hz (E)
1.8 Hz (I)

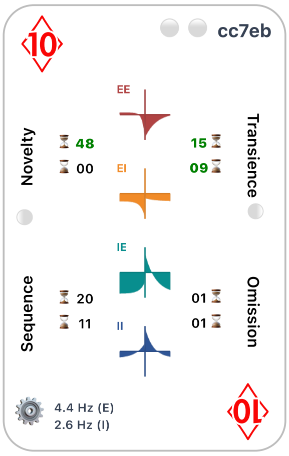
10
cc7eb
EE
EI
IE
II
Novelty
48
00
Transience
15
09
Sequence
20
11
Omission
01
01
4.4 Hz (E)
2.6 Hz (I)

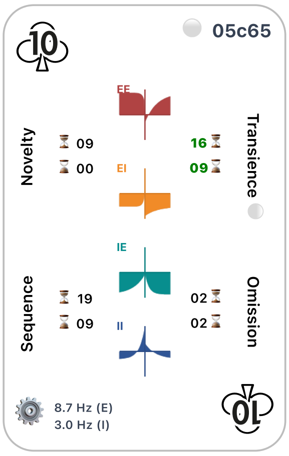
10
05c65
EE
EI
IE
II
Novelty
09
00
Transience
16
09
Sequence
19
09
Omission
02
02
8.7 Hz (E)
3.0 Hz (I)

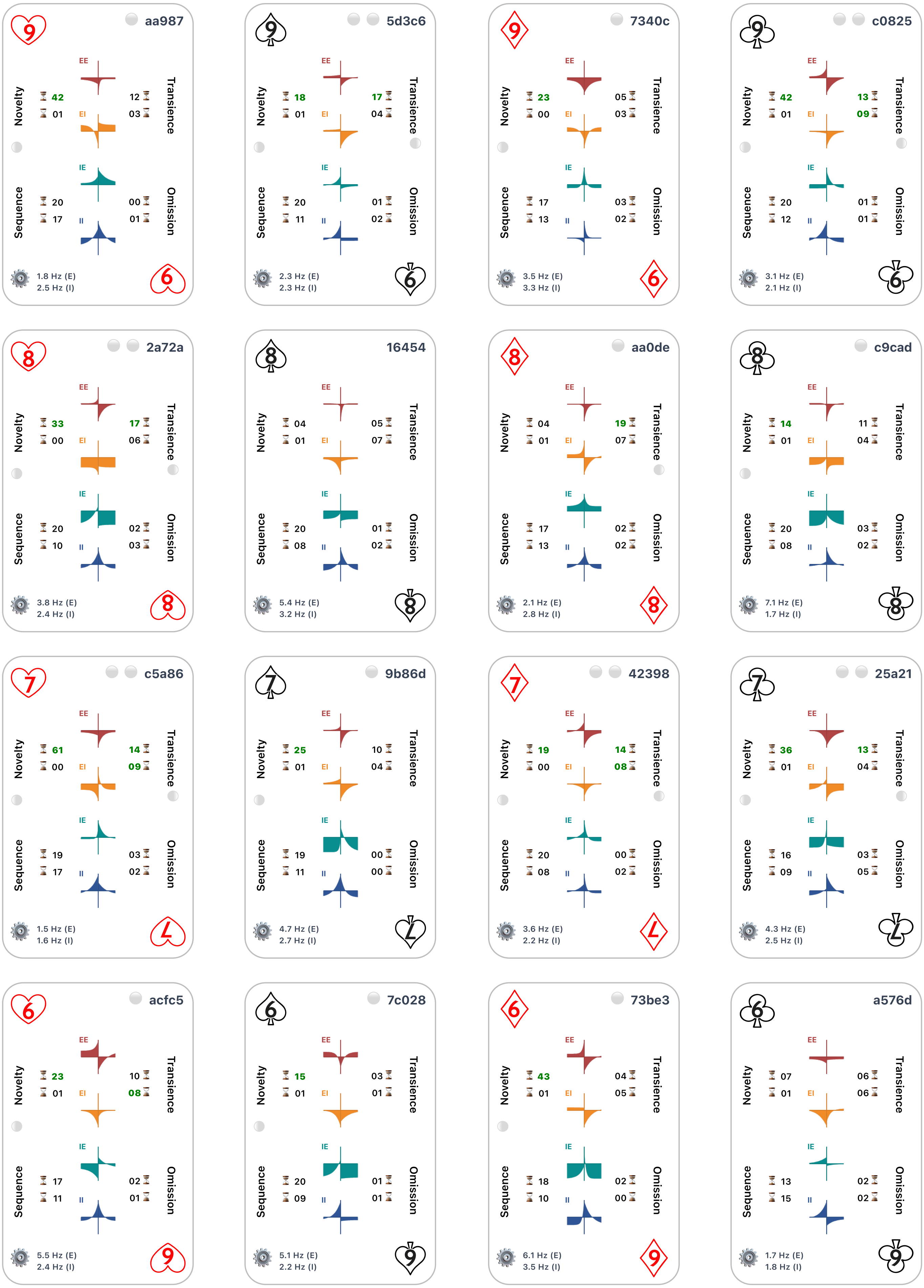
9 aa987
EE EI IE II
Novelty 42 01
Transience 12 03
Sequence 20 17
Omission 00 01
1.8 Hz (E) 2.5 Hz (I)
6
9 5d3c6
EE EI IE II
Novelty 18 01
Transience 17 04
Sequence 20 11
Omission 01 02
2.3 Hz (E) 2.3 Hz (I)
6
9 7340c
EE EI IE II
Novelty 23 00
Transience 05 03
Sequence 17 13
Omission 03 02
3.5 Hz (E) 3.3 Hz (I)
6
9 c0825
EE EI IE II
Novelty 42 01
Transience 13 09
Sequence 20 12
Omission 01 01
3.1 Hz (E) 2.1 Hz (I)
6
8 2a72a
EE EI IE II
Novelty 33 00
Transience 17 06
Sequence 20 10
Omission 02 03
3.8 Hz (E) 2.4 Hz (I)
8
8 16454
EE EI IE II
Novelty 04 01
Transience 05 07
Sequence 20 08
Omission 01 02
5.4 Hz (E) 3.2 Hz (I)
8
8 aa0de
EE EI IE II
Novelty 04 01
Transience 19 07
Sequence 17 13
Omission 02 02
2.1 Hz (E) 2.8 Hz (I)
8
8 c9cad
EE EI IE II
Novelty 14 01
Transience 11 04
Sequence 20 08
Omission 03 02
7.1 Hz (E) 1.7 Hz (I)
8
7 c5a86
EE EI IE II
Novelty 61 00
Transience 14 09
Sequence 19 17
Omission 03 02
1.5 Hz (E) 1.6 Hz (I)
7
7 9b86d
EE EI IE II
Novelty 25 01
Transience 10 04
Sequence 19 11
Omission 00 00
4.7 Hz (E) 2.7 Hz (I)
7
7 42398
EE EI IE II
Novelty 19 00
Transience 14 08
Sequence 20 08
Omission 00 02
3.6 Hz (E) 2.2 Hz (I)
7
7 25a21
EE EI IE II
Novelty 36 01
Transience 13 04
Sequence 16 09
Omission 03 05
4.3 Hz (E) 2.5 Hz (I)
7
6 acfc5
EE EI IE II
Novelty 23 01
Transience 10 08
Sequence 17 11
Omission 02 01
5.5 Hz (E) 2.4 Hz (I)
9
6 7c028
EE EI IE II
Novelty 15 01
Transience 03 01
Sequence 20 09
Omission 01 01
5.1 Hz (E) 2.2 Hz (I)
9
6 73be3
EE EI IE II
Novelty 43 01
Transience 04 05
Sequence 18 10
Omission 02 00
6.1 Hz (E) 3.5 Hz (I)
9
6 a576d
EE EI IE II
Novelty 07 01
Transience 06 06
Sequence 13 15
Omission 02 02
1.7 Hz (E) 1.8 Hz (I)
9

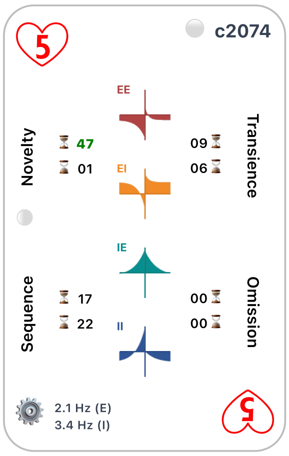
5
c2074
EE
Novelty
47
01
09
06
Transience
EI
IE
Sequence
17
22
00
00
Omission
II
2.1 Hz (E)
3.4 Hz (I)

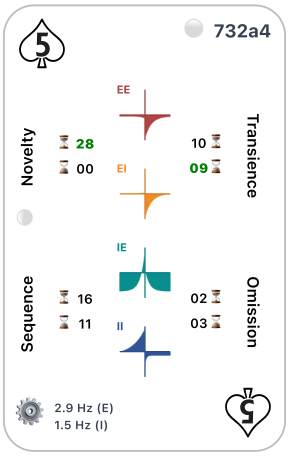
5
732a4
EE
Novelty
28
00
10
09
Transience
EI
IE
Sequence
16
11
02
03
Omission
II
2.9 Hz (E)
1.5 Hz (I)

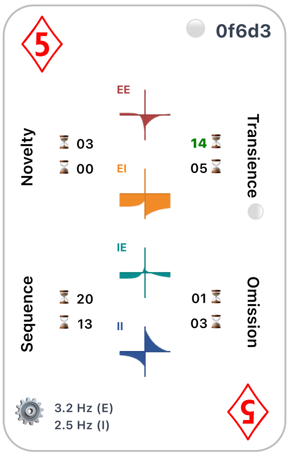
5
0f6d3
EE
Novelty
03
00
14
05
Transience
EI
IE
Sequence
20
13
01
03
Omission
II
3.2 Hz (E)
2.5 Hz (I)

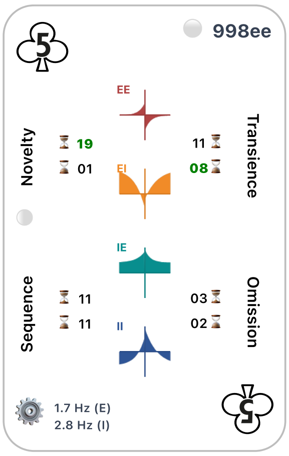
5
998ee
EE
Novelty
19
01
11
08
Transience
EI
IE
Sequence
11
11
03
02
Omission
II
1.7 Hz (E)
2.8 Hz (I)

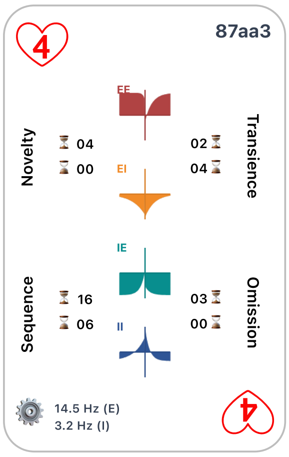
4
87aa3
EE
Novelty
04
00
02
04
Transience
EI
IE
Sequence
16
06
03
00
Omission
II
14.5 Hz (E)
3.2 Hz (I)

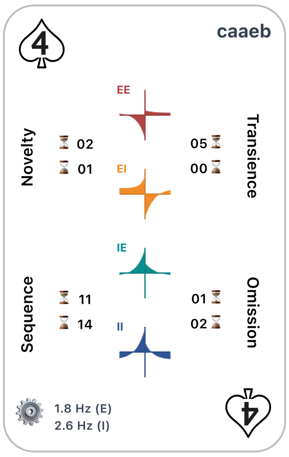
4
caaeb
EE
Novelty
02
01
05
00
Transience
EI
IE
Sequence
11
14
01
02
Omission
II
1.8 Hz (E)
2.6 Hz (I)

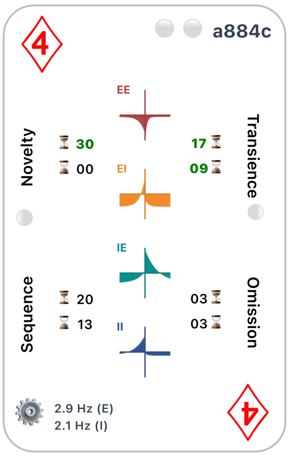
4
a884c
EE
Novelty
30
00
17
09
Transience
EI
IE
Sequence
20
13
03
03
Omission
II
2.9 Hz (E)
2.1 Hz (I)

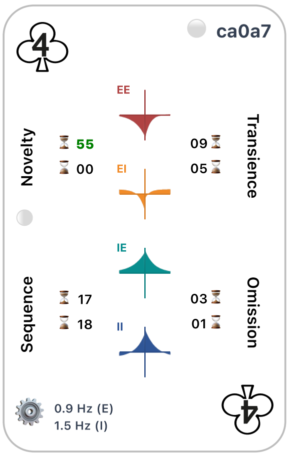
4
ca0a7
EE
Novelty
55
00
09
05
Transience
EI
IE
Sequence
17
18
03
01
Omission
II
0.9 Hz (E)
1.5 Hz (I)

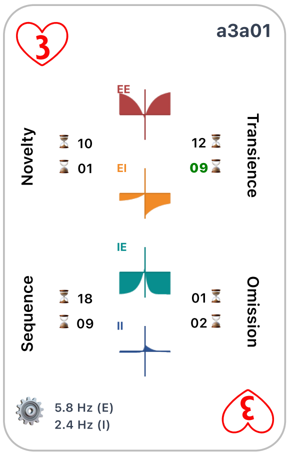
3
a3a01
EE
Novelty
10
01
12
09
Transience
EI
IE
Sequence
18
09
01
02
Omission
II
5.8 Hz (E)
2.4 Hz (I)

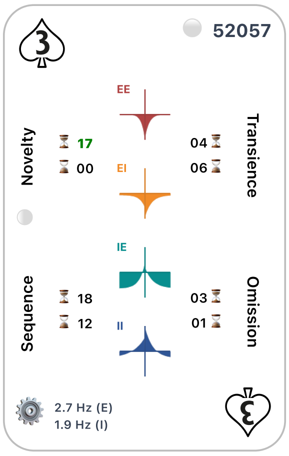
3
52057
EE
Novelty
17
00
04
06
Transience
EI
IE
Sequence
18
12
03
01
Omission
II
2.7 Hz (E)
1.9 Hz (I)

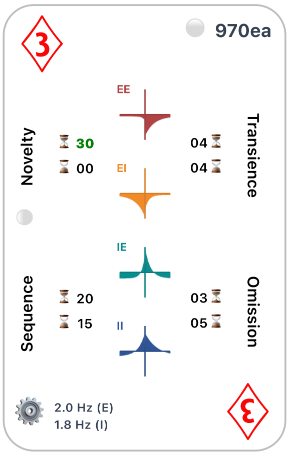
3
970ea
EE
Novelty
30
00
04
04
Transience
EI
IE
Sequence
20
15
03
05
Omission
II
2.0 Hz (E)
1.8 Hz (I)

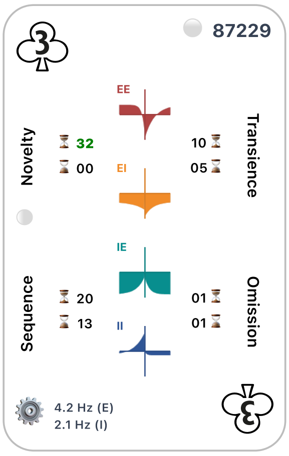
3
87229
EE
Novelty
32
00
10
05
Transience
EI
IE
Sequence
20
13
01
01
Omission
II
4.2 Hz (E)
2.1 Hz (I)

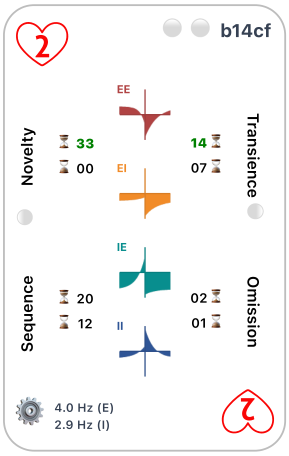
2
b14cf
EE
Novelty
33
00
14
07
Transience
EI
IE
Sequence
20
12
02
01
Omission
II
4.0 Hz (E)
2.9 Hz (I)

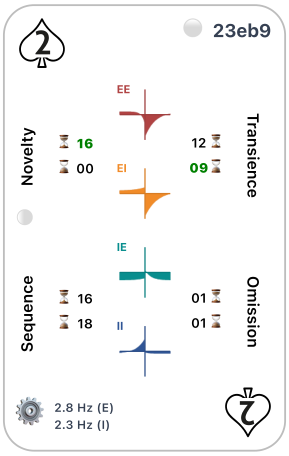
2
23eb9
EE
Novelty
16
00
12
09
Transience
EI
IE
Sequence
16
18
01
01
Omission
II
2.8 Hz (E)
2.3 Hz (I)

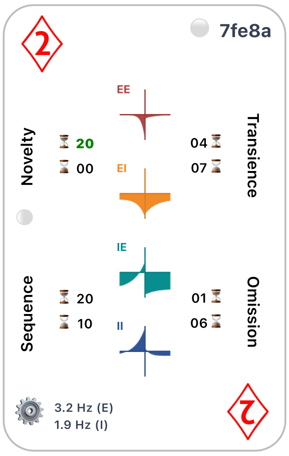
2
7fe8a
EE
Novelty
20
00
04
07
Transience
EI
IE
Sequence
20
10
01
06
Omission
II
3.2 Hz (E)
1.9 Hz (I)

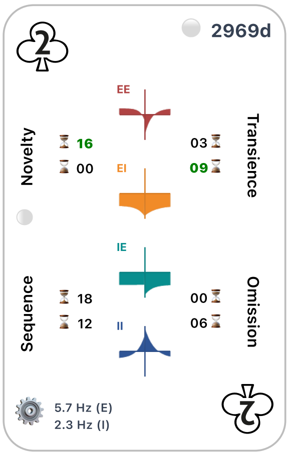
2
2969d
EE
Novelty
16
00
03
09
Transience
EI
IE
Sequence
18
12
00
06
Omission
II
5.7 Hz (E)
2.3 Hz (I)

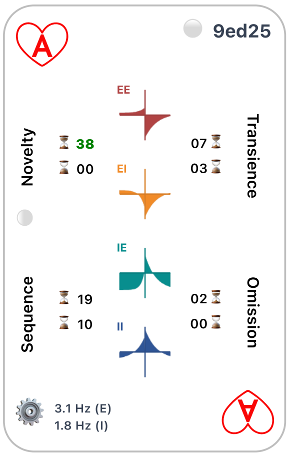
9ed25
EE
Novelty
38
00
07
03
Transience
EI
IE
Sequence
19
10
02
00
Omission
II
3.1 Hz (E)
1.8 Hz (I)

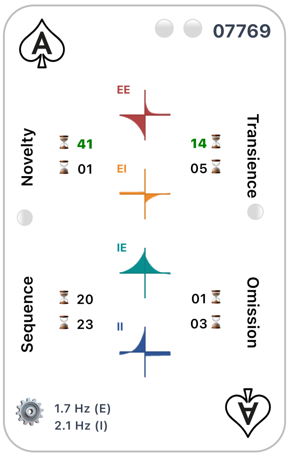
07769
EE
Novelty
41
01
14
05
Transience
EI
IE
Sequence
20
23
01
03
Omission
II
1.7 Hz (E)
2.1 Hz (I)

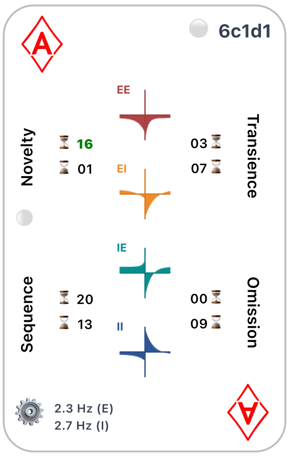
6c1d1
EE
Novelty
16
01
03
07
Transience
EI
IE
Sequence
20
13
00
09
Omission
II
2.3 Hz (E)
2.7 Hz (I)

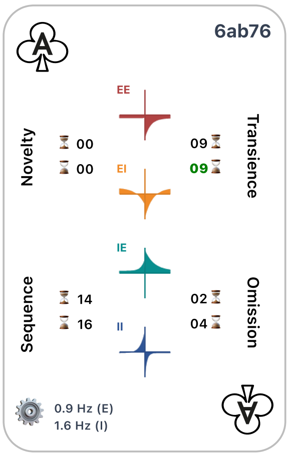
6ab76
EE
Novelty
00
00
09
09
Transience
EI
IE
Sequence
14
16
02
04
Omission
II
0.9 Hz (E)
1.6 Hz (I)

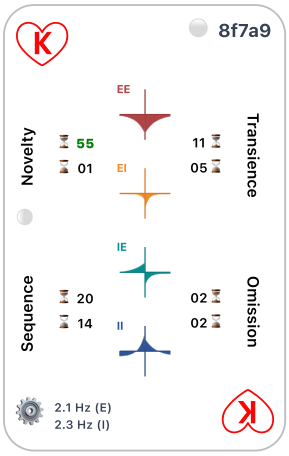
8f7a9
EE
Novelty
55
01
11
05
Transience
EI
IE
Sequence
20
14
02
02
Omission
II
2.1 Hz (E)
2.3 Hz (I)

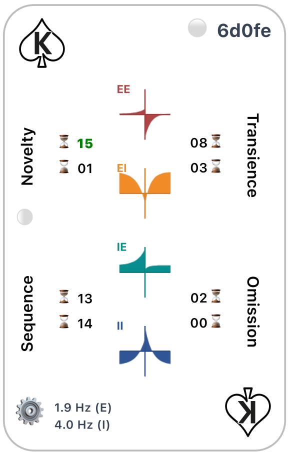
6d0fe
EE
Novelty
15
01
08
03
Transience
EI
IE
Sequence
13
14
02
00
Omission
II
1.9 Hz (E)
4.0 Hz (I)

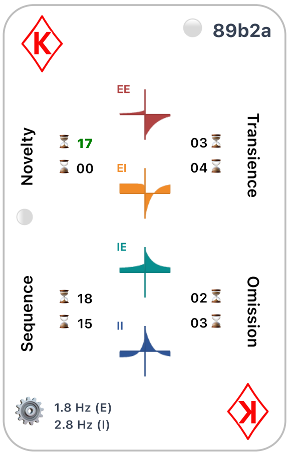
89b2a
EE
Novelty
17
00
03
04
Transience
EI
IE
Sequence
18
15
02
03
Omission
II
1.8 Hz (E)
2.8 Hz (I)

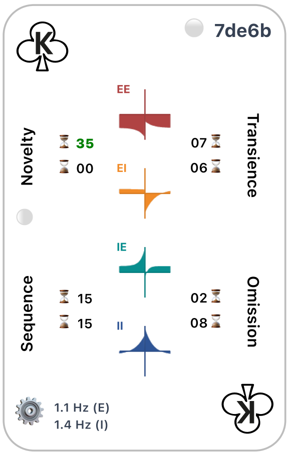
7de6b
EE
Novelty
35
00
07
06
Transience
EI
IE
Sequence
15
15
02
08
Omission
II
1.1 Hz (E)
1.4 Hz (I)

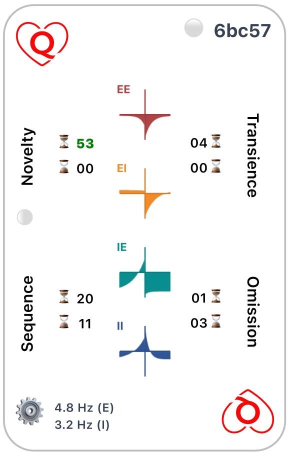
6bc57
EE
Novelty
53
00
04
00
Transience
EI
IE
Sequence
20
11
01
03
Omission
II
4.8 Hz (E)
3.2 Hz (I)

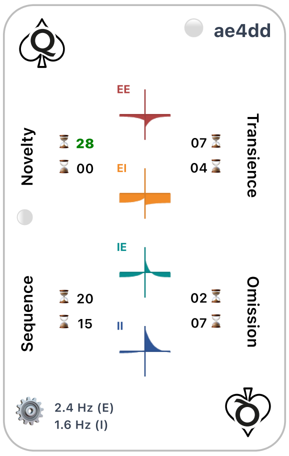
ae4dd
EE
Novelty
28
00
07
04
Transience
EI
IE
Sequence
20
15
02
07
Omission
II
2.4 Hz (E)
1.6 Hz (I)

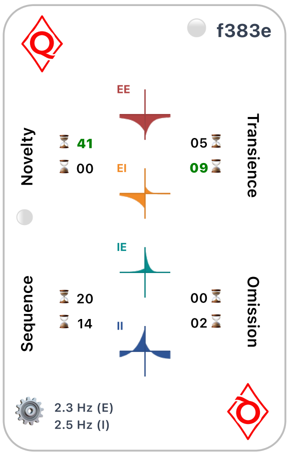
f383e
EE
Novelty
41
00
05
09
Transience
EI
IE
Sequence
20
14
00
02
Omission
II
2.3 Hz (E)
2.5 Hz (I)

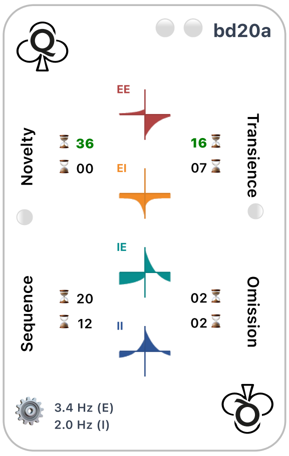
bd20a
EE
Novelty
36
00
16
07
Transience
EI
IE
Sequence
20
12
02
02
Omission
II
3.4 Hz (E)
2.0 Hz (I)

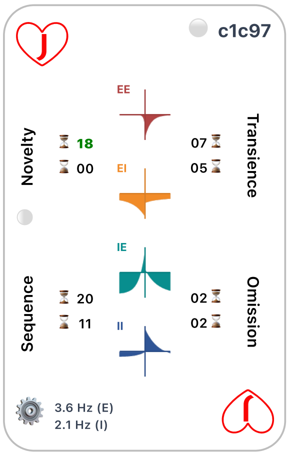
c1c97
EE
Novelty
18
00
07
05
Transience
EI
IE
Sequence
20
11
02
02
Omission
II
3.6 Hz (E)
2.1 Hz (I)

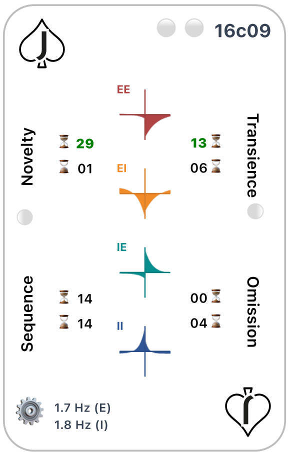
16c09
EE
Novelty
29
01
13
06
Transience
EI
IE
Sequence
14
14
00
04
Omission
II
1.7 Hz (E)
1.8 Hz (I)

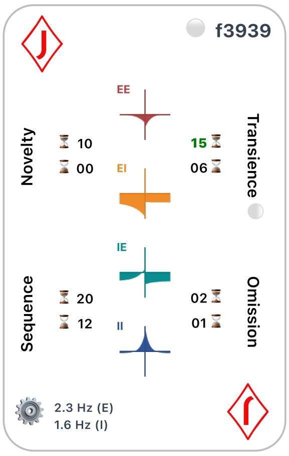
f3939
EE
Novelty
10
00
15
06
Transience
EI
IE
Sequence
20
12
02
01
Omission
II
2.3 Hz (E)
1.6 Hz (I)

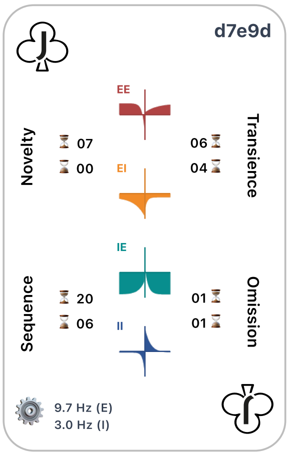
d7e9d
EE
Novelty
07
00
06
04
Transience
EI
IE
Sequence
20
06
01
01
Omission
II
9.7 Hz (E)
3.0 Hz (I)

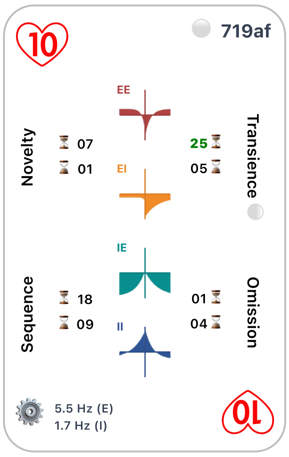
10
719af
EE
Novelty
07
01
EI
25
05
Transience
IE
Sequence
18
09
II
01
04
Omission
5.5 Hz (E)
1.7 Hz (I)

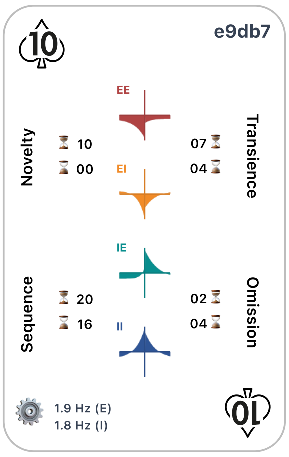
10
e9db7
EE
Novelty
10
00
EI
07
04
Transience
IE
Sequence
20
16
II
02
04
Omission
1.9 Hz (E)
1.8 Hz (I)

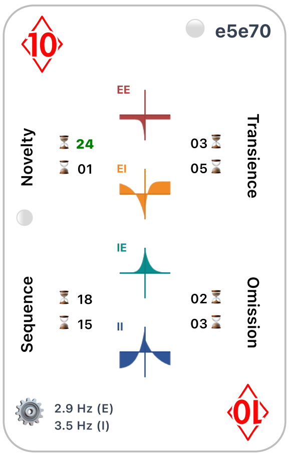
10
e5e70
EE
Novelty
24
01
EI
03
05
Transience
IE
Sequence
18
15
II
02
03
Omission
2.9 Hz (E)
3.5 Hz (I)

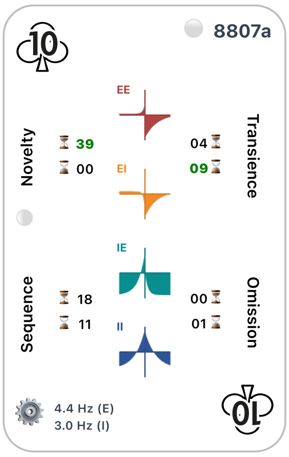
10
8807a
EE
Novelty
39
00
EI
04
09
Transience
IE
Sequence
18
11
II
00
01
Omission
4.4 Hz (E)
3.0 Hz (I)

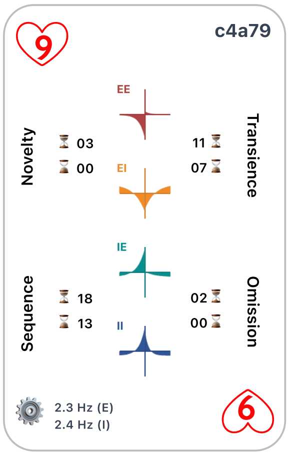
9
c4a79
EE
Novelty
03
00
EI
11
07
Transience
IE
Sequence
18
13
II
02
00
Omission
2.3 Hz (E)
2.4 Hz (I)

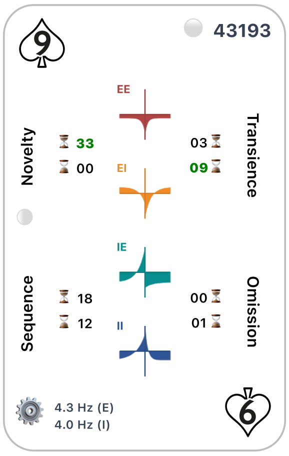
9
43193
EE
Novelty
33
00
EI
03
09
Transience
IE
Sequence
18
12
II
00
01
Omission
4.3 Hz (E)
4.0 Hz (I)

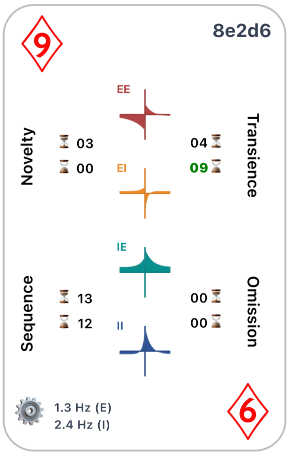
9
8e2d6
EE
Novelty
03
00
EI
04
09
Transience
IE
Sequence
13
12
II
00
00
Omission
1.3 Hz (E)
2.4 Hz (I)

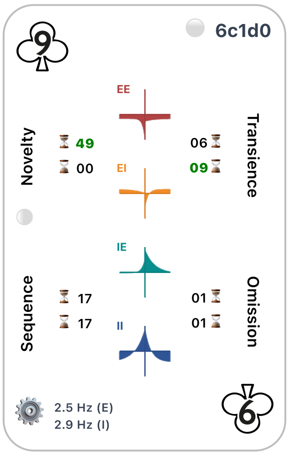
9
6c1d0
EE
Novelty
49
00
EI
06
09
Transience
IE
Sequence
17
17
II
01
01
Omission
2.5 Hz (E)
2.9 Hz (I)

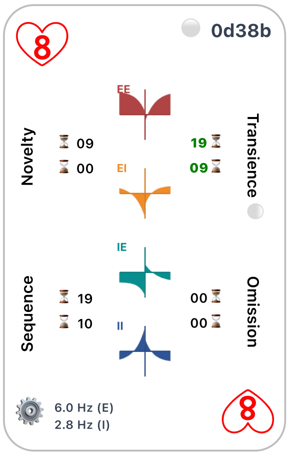
8
0d38b
EE
Novelty
09
00
EI
19
09
Transience
IE
Sequence
19
10
II
00
00
Omission
6.0 Hz (E)
2.8 Hz (I)

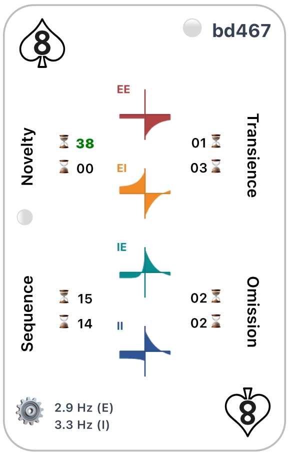
8
bd467
EE
Novelty
38
00
EI
01
03
Transience
IE
Sequence
15
14
II
02
02
Omission
2.9 Hz (E)
3.3 Hz (I)

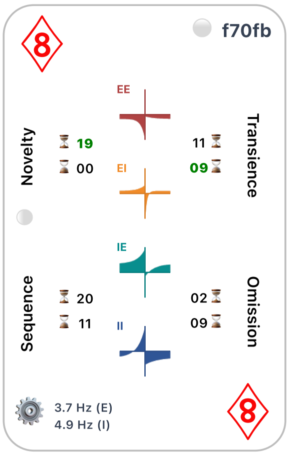
8
f70fb
EE
Novelty
19
00
EI
11
09
Transience
IE
Sequence
20
11
II
02
09
Omission
3.7 Hz (E)
4.9 Hz (I)

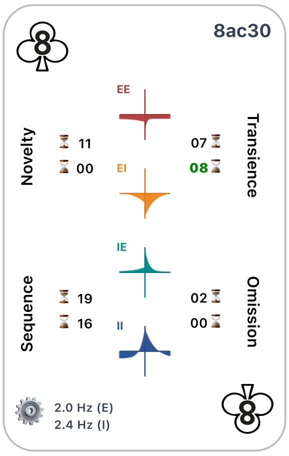
8
8ac30
EE
Novelty
11
00
EI
07
08
Transience
IE
Sequence
19
16
II
02
00
Omission
2.0 Hz (E)
2.4 Hz (I)

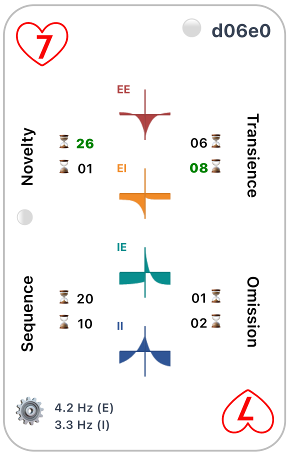
7
d06e0
EE
Novelty
26
01
EI
06
08
Transience
IE
Sequence
20
10
II
01
02
Omission
4.2 Hz (E)
3.3 Hz (I)

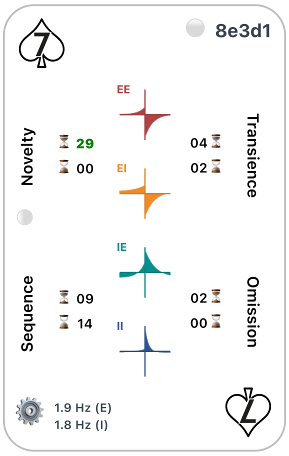
7
8e3d1
EE
Novelty
29
00
EI
04
02
Transience
IE
Sequence
09
14
II
02
00
Omission
1.9 Hz (E)
1.8 Hz (I)

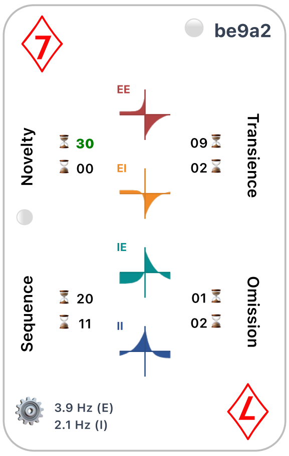
7
be9a2
EE
Novelty
30
00
EI
09
02
Transience
IE
Sequence
20
11
II
01
02
Omission
3.9 Hz (E)
2.1 Hz (I)

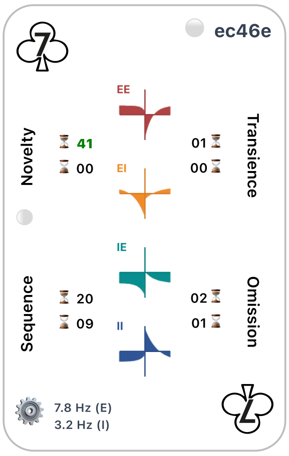
7
ec46e
EE
Novelty
41
00
EI
01
00
Transience
IE
Sequence
20
09
II
02
01
Omission
7.8 Hz (E)
3.2 Hz (I)

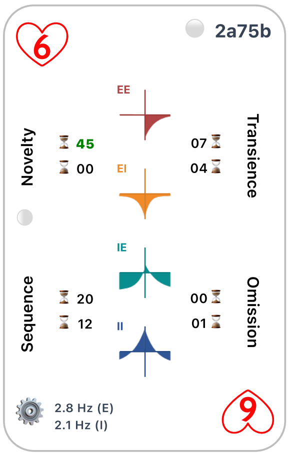
2a75b
EE
EI
IE
II
Novelty
45
00
07
04
Transience
Sequence
20
12
00
01
Omission
2.8 Hz (E)
2.1 Hz (I)

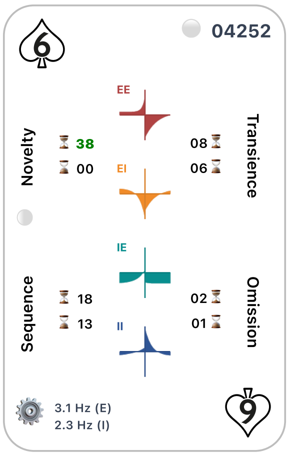
04252
EE
EI
IE
II
Novelty
38
00
08
06
Transience
Sequence
18
13
02
01
Omission
3.1 Hz (E)
2.3 Hz (I)

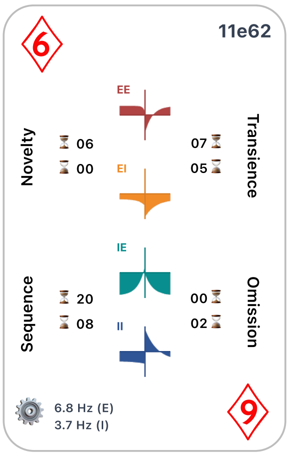
11e62
EE
EI
IE
II
Novelty
06
00
07
05
Transience
Sequence
20
08
00
02
Omission
6.8 Hz (E)
3.7 Hz (I)

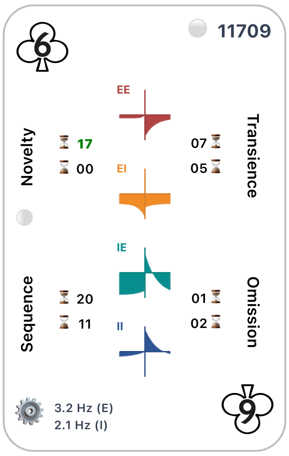
11709
EE
EI
IE
II
Novelty
17
00
07
05
Transience
Sequence
20
11
01
02
Omission
3.2 Hz (E)
2.1 Hz (I)

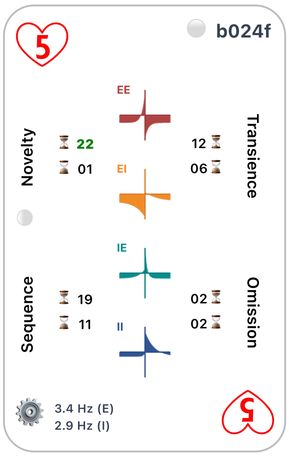
b024f
EE
EI
IE
II
Novelty
22
01
12
06
Transience
Sequence
19
11
02
02
Omission
3.4 Hz (E)
2.9 Hz (I)

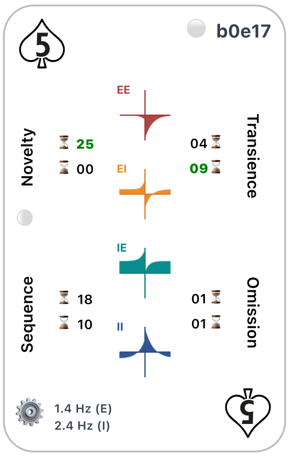
b0e17
EE
EI
IE
II
Novelty
25
00
04
09
Transience
Sequence
18
10
01
01
Omission
1.4 Hz (E)
2.4 Hz (I)

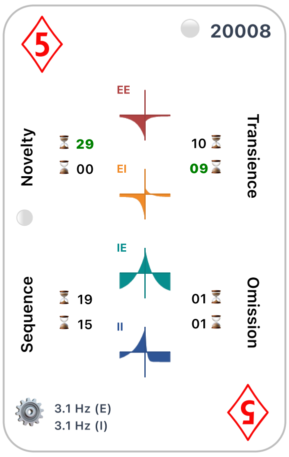
20008
EE
EI
IE
II
Novelty
29
00
10
09
Transience
Sequence
19
15
01
01
Omission
3.1 Hz (E)
3.1 Hz (I)

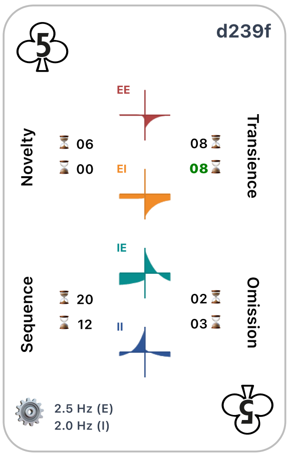
d239f
EE
EI
IE
II
Novelty
06
00
08
08
Transience
Sequence
20
12
02
03
Omission
2.5 Hz (E)
2.0 Hz (I)

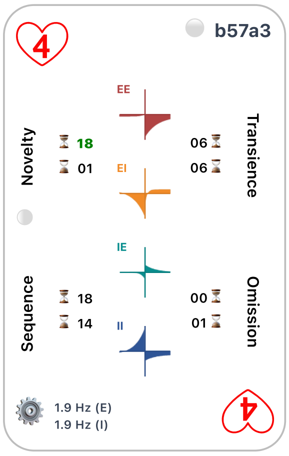
b57a3
EE
EI
IE
II
Novelty
18
01
06
06
Transience
Sequence
18
14
00
01
Omission
1.9 Hz (E)
1.9 Hz (I)

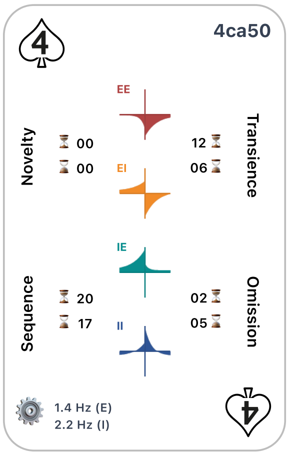
4ca50
EE
EI
IE
II
Novelty
00
00
12
06
Transience
Sequence
20
17
02
05
Omission
1.4 Hz (E)
2.2 Hz (I)

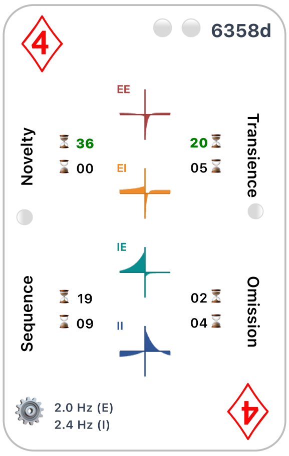
6358d
EE
EI
IE
II
Novelty
36
00
20
05
Transience
Sequence
19
09
02
04
Omission
2.0 Hz (E)
2.4 Hz (I)

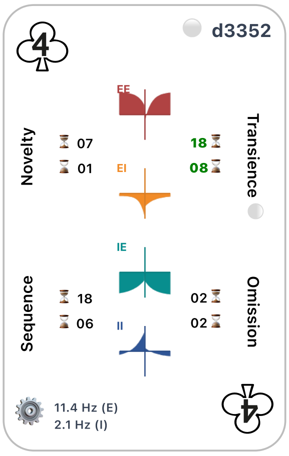
d3352
EE
EI
IE
II
Novelty
07
01
18
08
Transience
Sequence
18
06
02
02
Omission
11.4 Hz (E)
2.1 Hz (I)

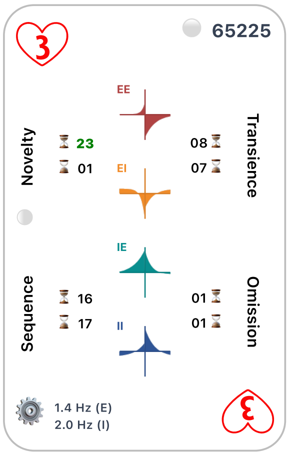
65225
EE
EI
IE
II
Novelty
23
01
08
07
Transience
Sequence
16
17
01
01
Omission
1.4 Hz (E)
2.0 Hz (I)

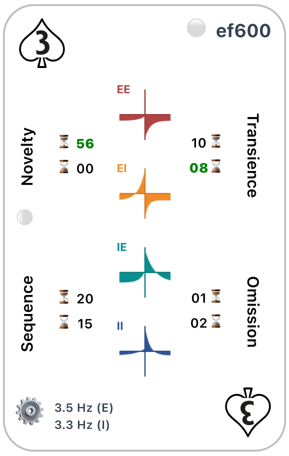
ef600
EE
EI
IE
II
Novelty
56
00
10
08
Transience
Sequence
20
15
01
02
Omission
3.5 Hz (E)
3.3 Hz (I)

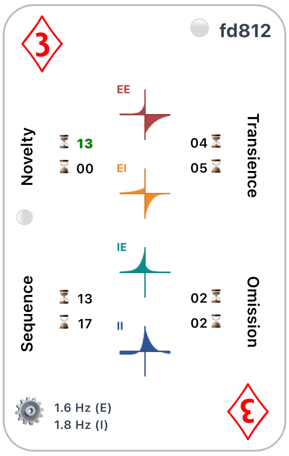
fd812
EE
EI
IE
II
Novelty
13
00
04
05
Transience
Sequence
13
17
02
02
Omission
1.6 Hz (E)
1.8 Hz (I)

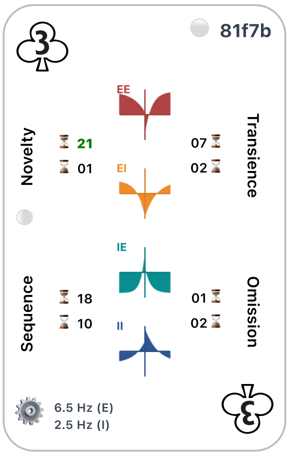
81f7b
EE
EI
IE
II
Novelty
21
01
07
02
Transience
Sequence
18
10
01
02
Omission
6.5 Hz (E)
2.5 Hz (I)

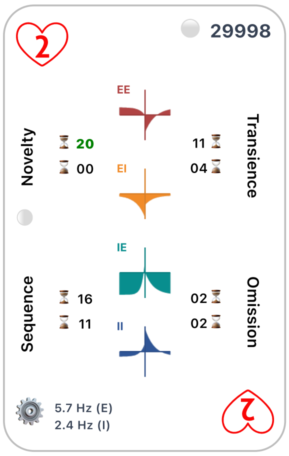
2
29998
EE
EI
IE
II
Novelty
20
00
Transience
11
04
Sequence
16
11
Omission
02
02
5.7 Hz (E)
2.4 Hz (I)

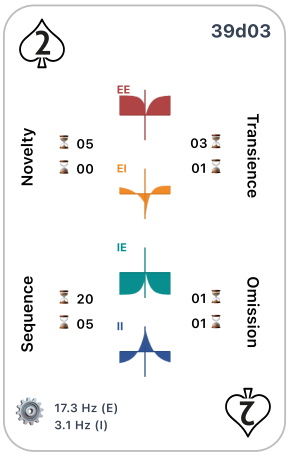
2
39d03
EE
EI
IE
II
Novelty
05
00
Transience
03
01
Sequence
20
05
Omission
01
01
17.3 Hz (E)
3.1 Hz (I)

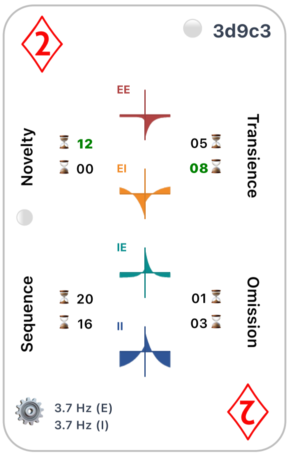
2
3d9c3
EE
EI
IE
II
Novelty
12
00
Transience
05
08
Sequence
20
16
Omission
01
03
3.7 Hz (E)
3.7 Hz (I)

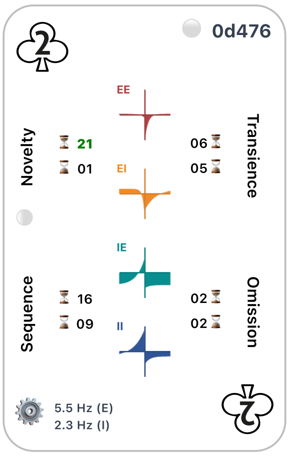
2
0d476
EE
EI
IE
II
Novelty
21
01
Transience
06
05
Sequence
16
09
Omission
02
02
5.5 Hz (E)
2.3 Hz (I)

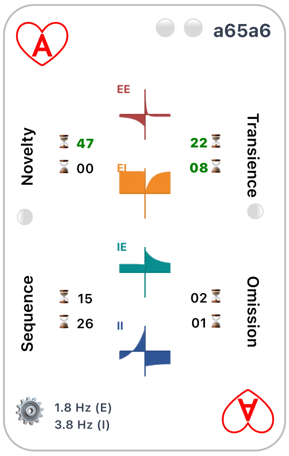
A
a65a6
EE
EI
IE
II
Novelty
47
00
Transience
22
08
Sequence
15
26
Omission
02
01
1.8 Hz (E)
3.8 Hz (I)

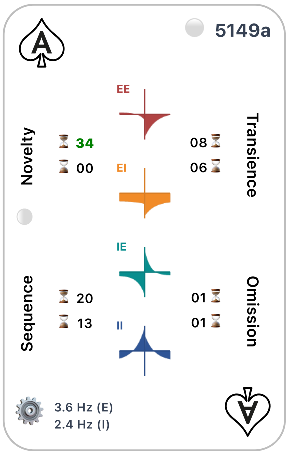
A
5149a
EE
EI
IE
II
Novelty
34
00
Transience
08
06
Sequence
20
13
Omission
01
01
3.6 Hz (E)
2.4 Hz (I)

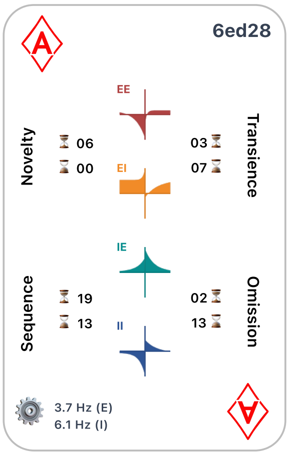
A
6ed28
EE
EI
IE
II
Novelty
06
00
Transience
03
07
Sequence
19
13
Omission
02
13
3.7 Hz (E)
6.1 Hz (I)

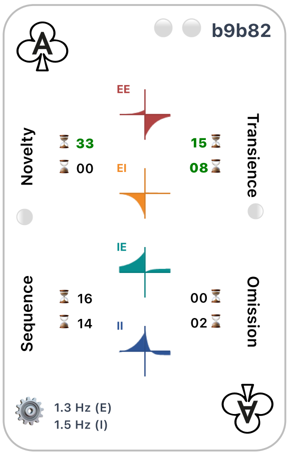
A
b9b82
EE
EI
IE
II
Novelty
33
00
Transience
15
08
Sequence
16
14
Omission
00
02
1.3 Hz (E)
1.5 Hz (I)

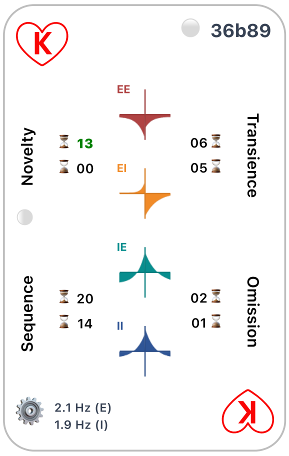
K
36b89
EE
EI
IE
II
Novelty
13
00
Transience
06
05
Sequence
20
14
Omission
02
01
2.1 Hz (E)
1.9 Hz (I)

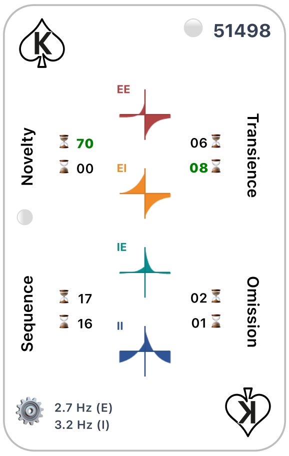
K
51498
EE
EI
IE
II
Novelty
70
00
Transience
06
08
Sequence
17
16
Omission
02
01
2.7 Hz (E)
3.2 Hz (I)

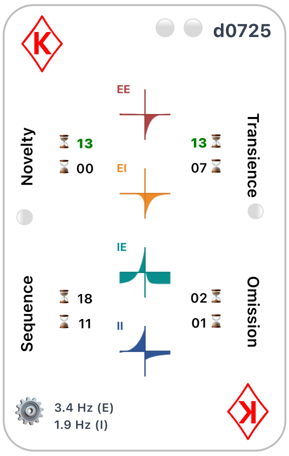
K
d0725
EE
EI
IE
II
Novelty
13
00
Transience
13
07
Sequence
18
11
Omission
02
01
3.4 Hz (E)
1.9 Hz (I)

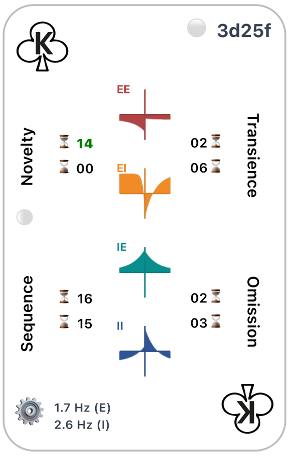
K
3d25f
EE
EI
IE
II
Novelty
14
00
Transience
02
06
Sequence
16
15
Omission
02
03
1.7 Hz (E)
2.6 Hz (I)

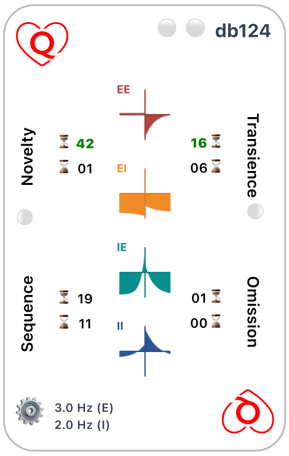
Q
db124
EE
EI
IE
II
Novelty
42
01
Transience
16
06
Sequence
19
11
Omission
01
00
3.0 Hz (E)
2.0 Hz (I)

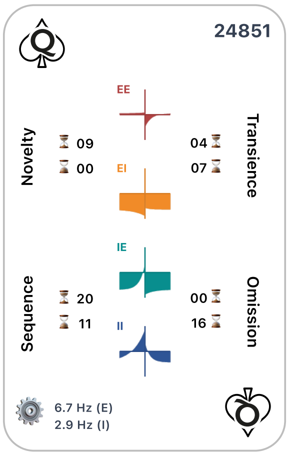
Q
24851
EE
EI
IE
II
Novelty
09
00
Transience
04
07
Sequence
20
11
Omission
00
16
6.7 Hz (E)
2.9 Hz (I)

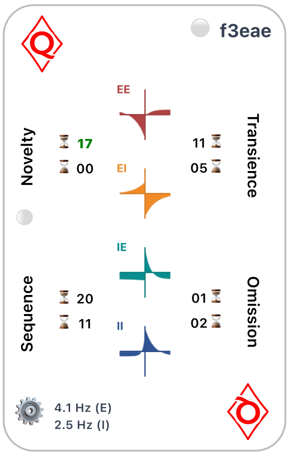
Q
f3eae
EE
EI
IE
II
Novelty
17
00
Transience
11
05
Sequence
20
11
Omission
01
02
4.1 Hz (E)
2.5 Hz (I)

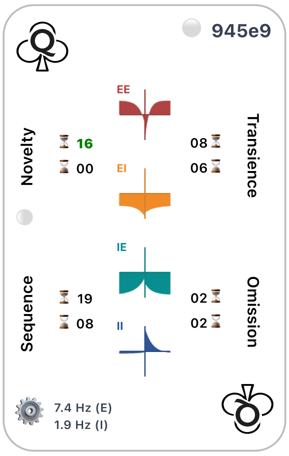
Q
945e9
EE
EI
IE
II
Novelty
16
00
Transience
08
06
Sequence
19
08
Omission
02
02
7.4 Hz (E)
1.9 Hz (I)

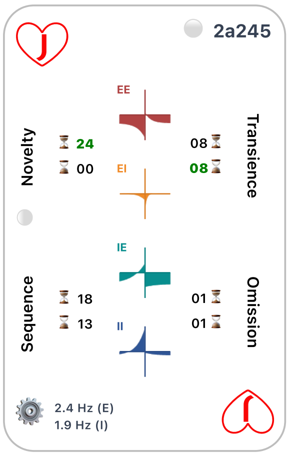
J
2a245
EE
EI
IE
II
Novelty
24
00
Transience
08
08
Sequence
18
13
Omission
01
01
2.4 Hz (E)
1.9 Hz (I)

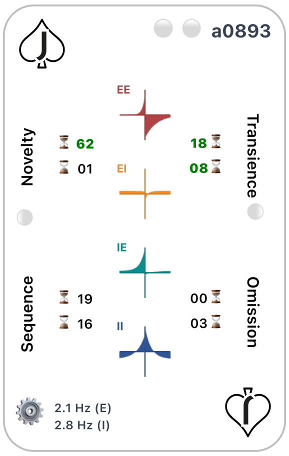
J
a0893
EE
EI
IE
II
Novelty
62
01
Transience
18
08
Sequence
19
16
Omission
00
03
2.1 Hz (E)
2.8 Hz (I)

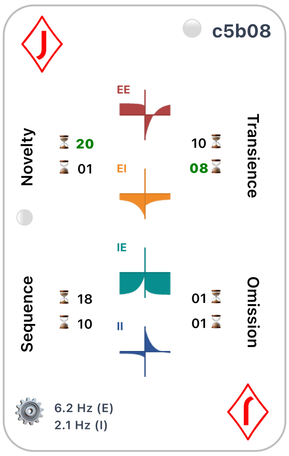
J
c5b08
EE
EI
IE
II
Novelty
20
01
Transience
10
08
Sequence
18
10
Omission
01
01
6.2 Hz (E)
2.1 Hz (I)

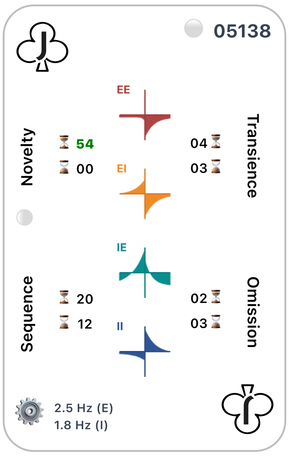
J
05138
EE
EI
IE
II
Novelty
54
00
Transience
04
03
Sequence
20
12
Omission
02
03
2.5 Hz (E)
1.8 Hz (I)

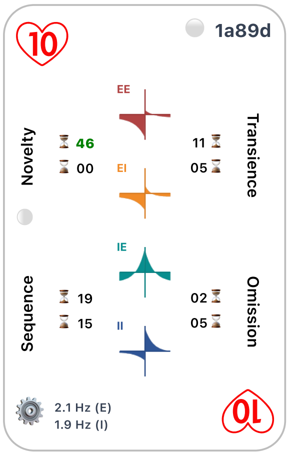
10
1a89d
EE
EI
IE
II
Novelty
46
00
Transience
11
05
Sequence
19
15
Omission
02
05
2.1 Hz (E)
1.9 Hz (I)

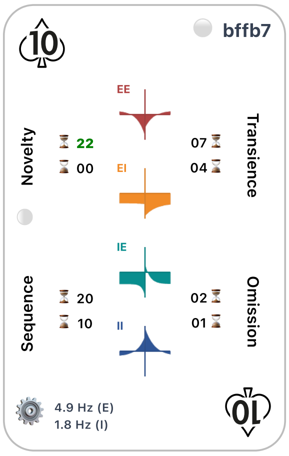
10
bffb7
EE
EI
IE
II
Novelty
22
00
Transience
07
04
Sequence
20
10
Omission
02
01
4.9 Hz (E)
1.8 Hz (I)

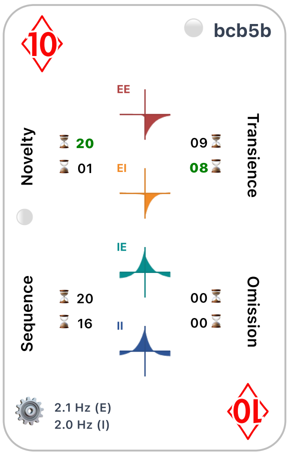
10
bcb5b
EE
EI
IE
II
Novelty
20
01
Transience
09
08
Sequence
20
16
Omission
00
00
2.1 Hz (E)
2.0 Hz (I)

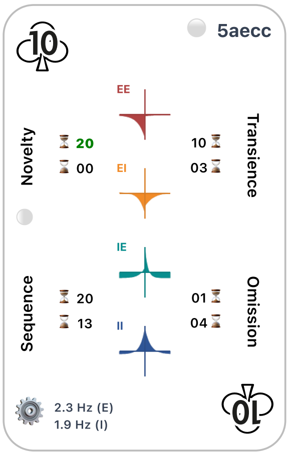
10
5aecc
EE
EI
IE
II
Novelty
20
00
Transience
10
03
Sequence
20
13
Omission
01
04
2.3 Hz (E)
1.9 Hz (I)

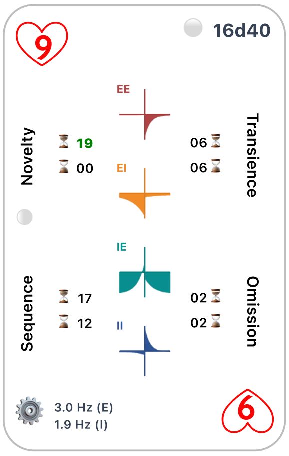
9
16d40
EE
EI
IE
II
Novelty
19
00
Transience
06
06
Sequence
17
12
Omission
02
02
3.0 Hz (E)
1.9 Hz (I)

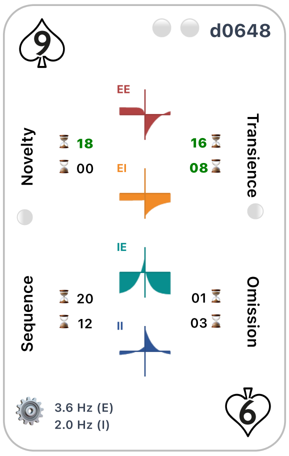
9
d0648
EE
EI
IE
II
Novelty
18
00
Transience
16
08
Sequence
20
12
Omission
01
03
3.6 Hz (E)
2.0 Hz (I)

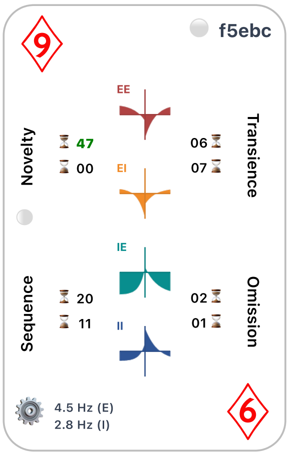
9
f5ebc
EE
EI
IE
II
Novelty
47
00
Transience
06
07
Sequence
20
11
Omission
02
01
4.5 Hz (E)
2.8 Hz (I)

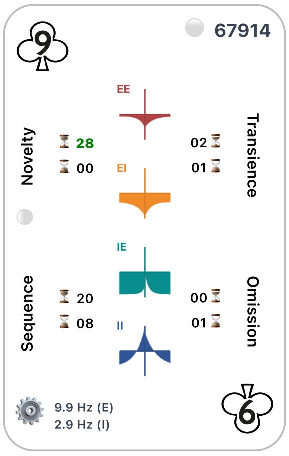
9
67914
EE
EI
IE
II
Novelty
28
00
Transience
02
01
Sequence
20
08
Omission
00
01
9.9 Hz (E)
2.9 Hz (I)

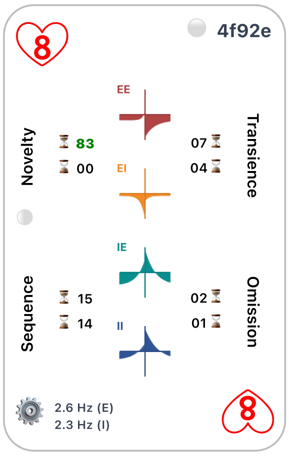
8
4f92e
EE
EI
IE
II
Novelty
83
00
Transience
07
04
Sequence
15
14
Omission
02
01
2.6 Hz (E)
2.3 Hz (I)

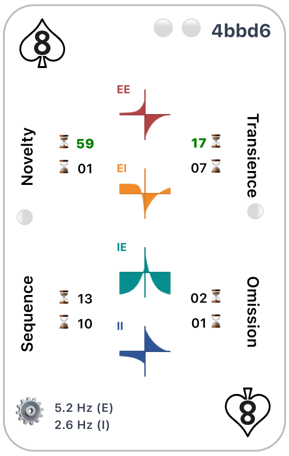
8
4bbd6
EE
EI
IE
II
Novelty
59
01
Transience
17
07
Sequence
13
10
Omission
02
01
5.2 Hz (E)
2.6 Hz (I)

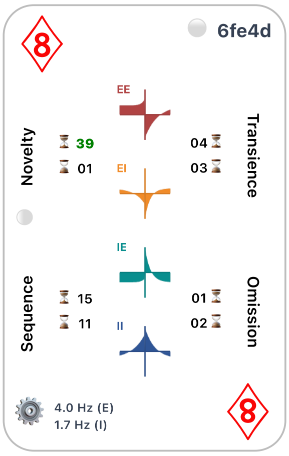
8
6fe4d
EE
EI
IE
II
Novelty
39
01
Transience
04
03
Sequence
15
11
Omission
01
02
4.0 Hz (E)
1.7 Hz (I)

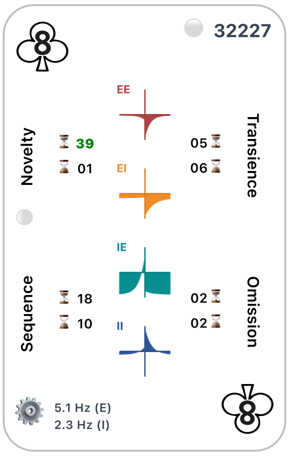
8
32227
EE
EI
IE
II
Novelty
39
01
Transience
05
06
Sequence
18
10
Omission
02
02
5.1 Hz (E)
2.3 Hz (I)

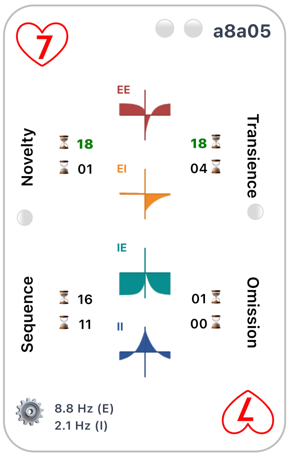
7
a8a05
EE
Novelty
18
01
18
04
Transience
EI
IE
Sequence
16
11
01
00
Omission
II
8.8 Hz (E)
2.1 Hz (I)

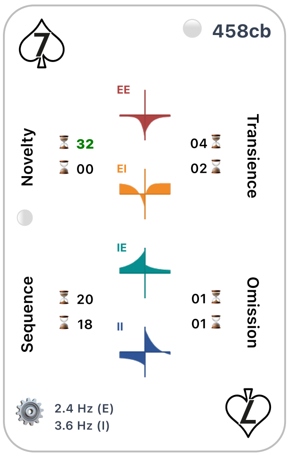
7
458cb
EE
Novelty
32
00
04
02
Transience
EI
IE
Sequence
20
18
01
01
Omission
II
2.4 Hz (E)
3.6 Hz (I)

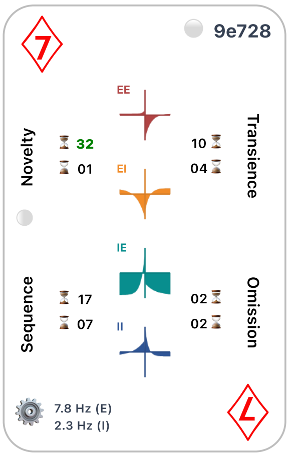
7
9e728
EE
Novelty
32
01
10
04
Transience
EI
IE
Sequence
17
07
02
02
Omission
II
7.8 Hz (E)
2.3 Hz (I)

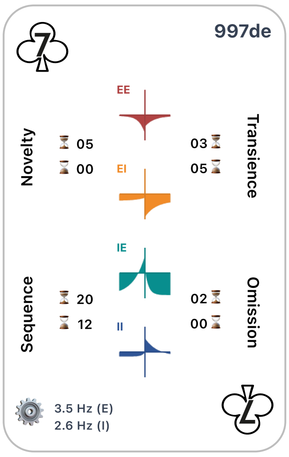
7
997de
EE
Novelty
05
00
03
05
Transience
EI
IE
Sequence
20
12
02
00
Omission
II
3.5 Hz (E)
2.6 Hz (I)

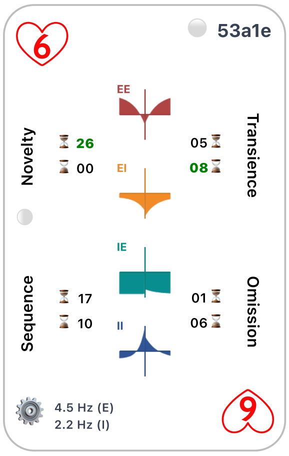
6
53a1e
EE
Novelty
26
00
05
08
Transience
EI
IE
Sequence
17
10
01
06
Omission
II
4.5 Hz (E)
2.2 Hz (I)

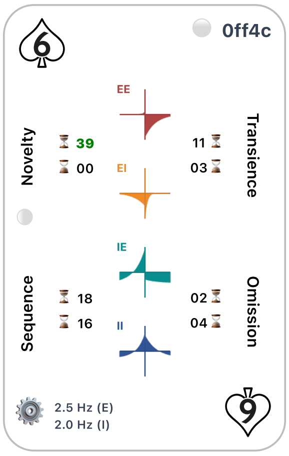
6
0ff4c
EE
Novelty
39
00
11
03
Transience
EI
IE
Sequence
18
16
02
04
Omission
II
2.5 Hz (E)
2.0 Hz (I)

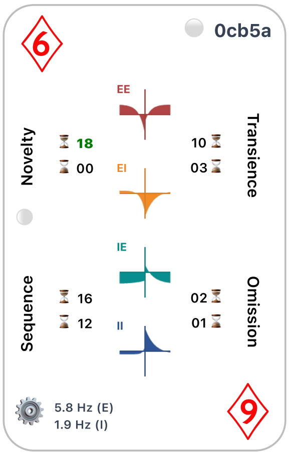
6
0cb5a
EE
Novelty
18
00
10
03
Transience
EI
IE
Sequence
16
12
02
01
Omission
II
5.8 Hz (E)
1.9 Hz (I)

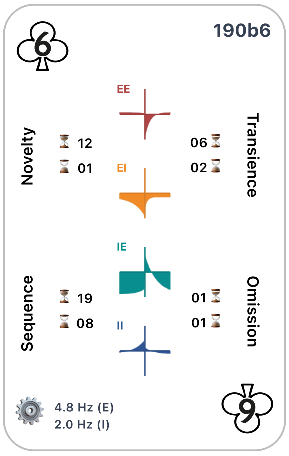
6
190b6
EE
Novelty
12
01
06
02
Transience
EI
IE
Sequence
19
08
01
01
Omission
II
4.8 Hz (E)
2.0 Hz (I)

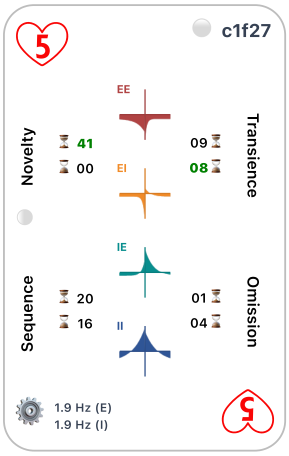
5
c1f27
EE
Novelty
41
00
09
08
Transience
EI
IE
Sequence
20
16
01
04
Omission
II
1.9 Hz (E)
1.9 Hz (I)

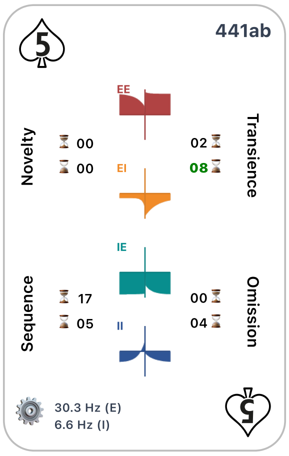
5
441ab
EE
Novelty
00
00
02
08
Transience
EI
IE
Sequence
17
05
00
04
Omission
II
30.3 Hz (E)
6.6 Hz (I)

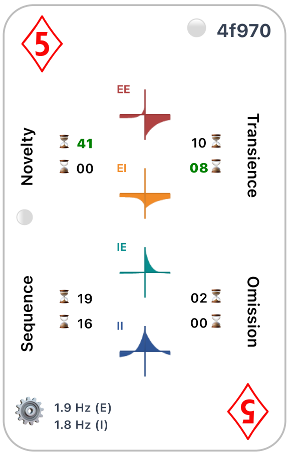
5
4f970
EE
Novelty
41
00
10
08
Transience
EI
IE
Sequence
19
16
02
00
Omission
II
1.9 Hz (E)
1.8 Hz (I)

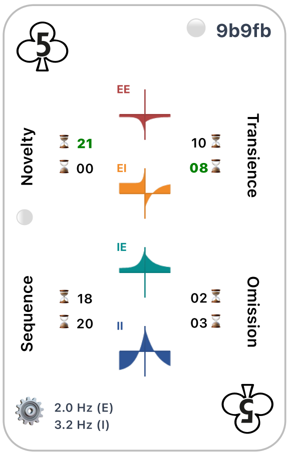
5
9b9fb
EE
Novelty
21
00
10
08
Transience
EI
IE
Sequence
18
20
02
03
Omission
II
2.0 Hz (E)
3.2 Hz (I)

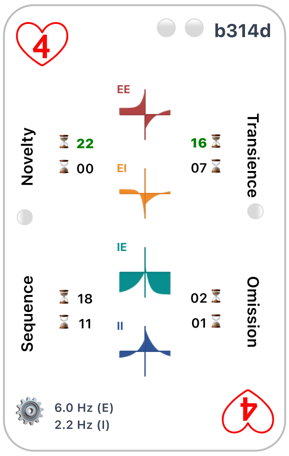
4
b314d
EE
Novelty
22
00
16
07
Transience
EI
IE
Sequence
18
11
02
01
Omission
II
6.0 Hz (E)
2.2 Hz (I)

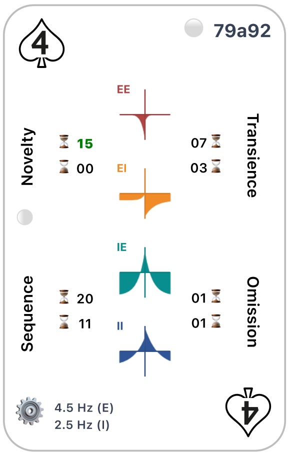
4
79a92
EE
Novelty
15
00
07
03
Transience
EI
IE
Sequence
20
11
01
01
Omission
II
4.5 Hz (E)
2.5 Hz (I)

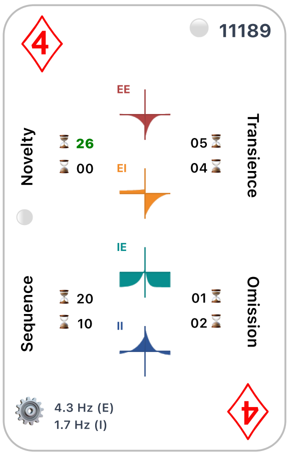
4
11189
EE
Novelty
26
00
05
04
Transience
EI
IE
Sequence
20
10
01
02
Omission
II
4.3 Hz (E)
1.7 Hz (I)

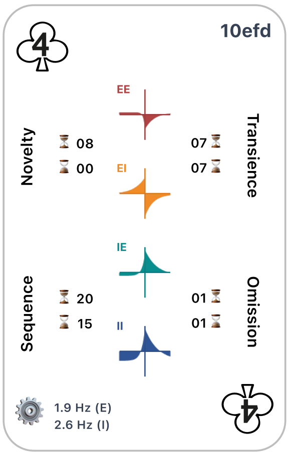
4
10efd
EE
Novelty
08
00
07
07
Transience
EI
IE
Sequence
20
15
01
01
Omission
II
1.9 Hz (E)
2.6 Hz (I)

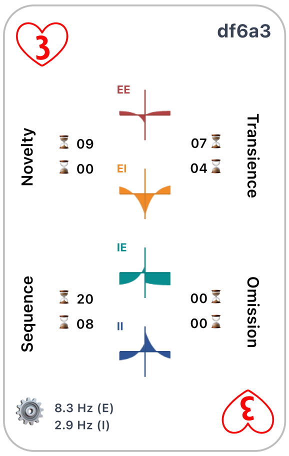
3
df6a3
EE
EI
IE
II
Novelty
09
00
07
04
Transience
Sequence
20
08
00
00
Omission
8.3 Hz (E)
2.9 Hz (I)

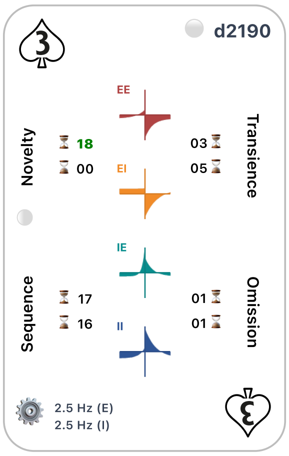
3
d2190
EE
EI
IE
II
Novelty
18
00
03
05
Transience
Sequence
17
16
01
01
Omission
2.5 Hz (E)
2.5 Hz (I)

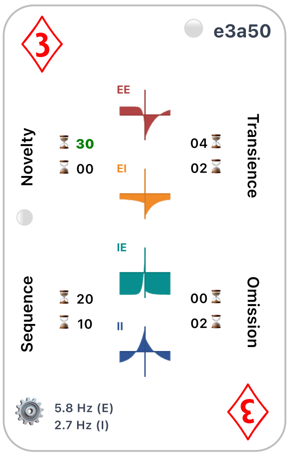
3
e3a50
EE
EI
IE
II
Novelty
30
00
04
02
Transience
Sequence
20
10
00
02
Omission
5.8 Hz (E)
2.7 Hz (I)

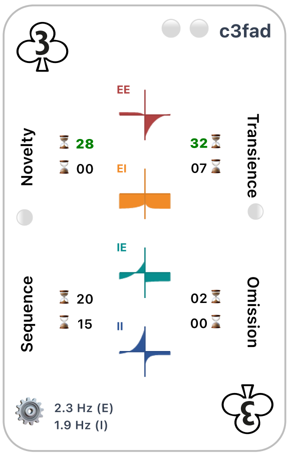
3
c3fad
EE
EI
IE
II
Novelty
28
00
32
07
Transience
Sequence
20
15
02
00
Omission
2.3 Hz (E)
1.9 Hz (I)

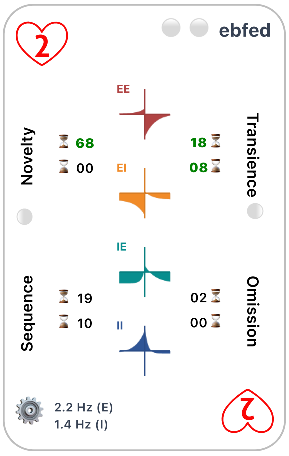
2
ebfed
EE
EI
IE
II
Novelty
68
00
18
08
Transience
Sequence
19
10
02
00
Omission
2.2 Hz (E)
1.4 Hz (I)

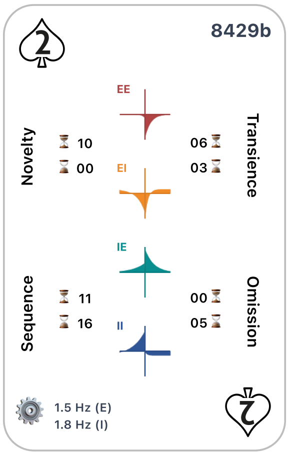
2
8429b
EE
EI
IE
II
Novelty
10
00
06
03
Transience
Sequence
11
16
00
05
Omission
1.5 Hz (E)
1.8 Hz (I)

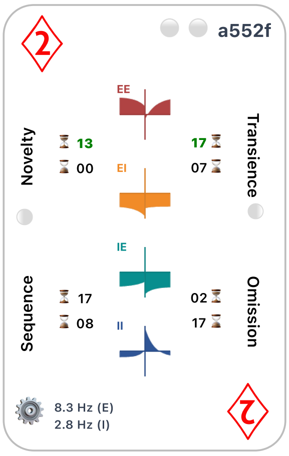
2
a552f
EE
EI
IE
II
Novelty
13
00
17
07
Transience
Sequence
17
08
02
17
Omission
8.3 Hz (E)
2.8 Hz (I)

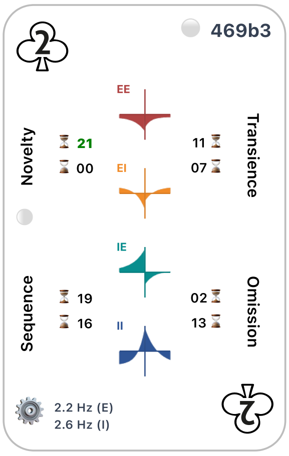
2
469b3
EE
EI
IE
II
Novelty
21
00
11
07
Transience
Sequence
19
16
02
13
Omission
2.2 Hz (E)
2.6 Hz (I)

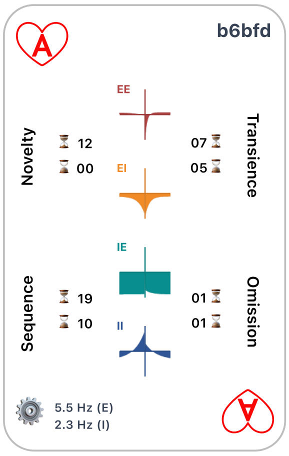
A
b6bfd
EE
EI
IE
II
Novelty
12
00
07
05
Transience
Sequence
19
10
01
01
Omission
5.5 Hz (E)
2.3 Hz (I)

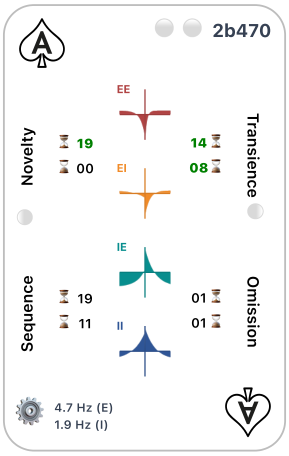
A
2b470
EE
EI
IE
II
Novelty
19
00
14
08
Transience
Sequence
19
11
01
01
Omission
4.7 Hz (E)
1.9 Hz (I)

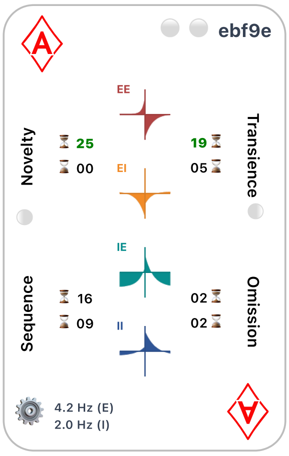
A
ebf9e
EE
EI
IE
II
Novelty
25
00
19
05
Transience
Sequence
16
09
02
02
Omission
4.2 Hz (E)
2.0 Hz (I)

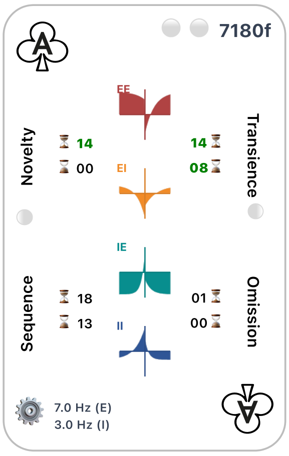
A
7180f
EE
EI
IE
II
Novelty
14
00
14
08
Transience
Sequence
18
13
01
00
Omission
7.0 Hz (E)
3.0 Hz (I)

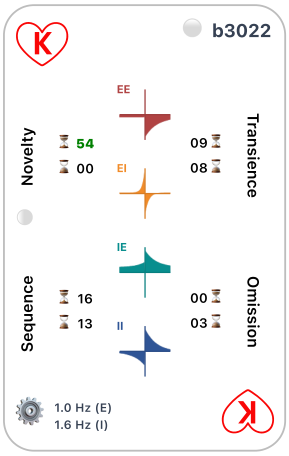
K
b3022
EE
EI
IE
II
Novelty
54
00
09
08
Transience
Sequence
16
13
00
03
Omission
1.0 Hz (E)
1.6 Hz (I)

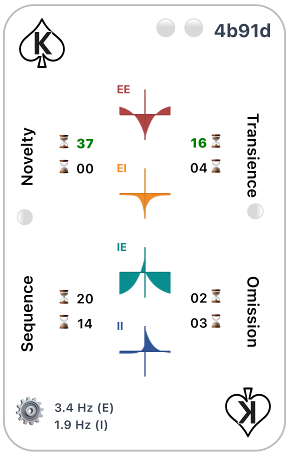
K
4b91d
EE
EI
IE
II
Novelty
37
00
16
04
Transience
Sequence
20
14
02
03
Omission
3.4 Hz (E)
1.9 Hz (I)

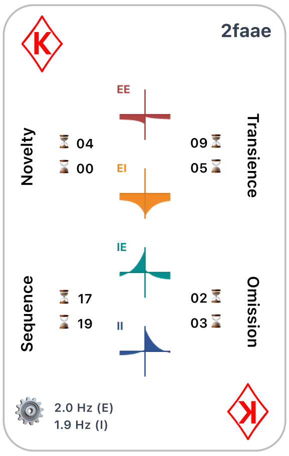
K
2faae
EE
EI
IE
II
Novelty
04
00
09
05
Transience
Sequence
17
19
02
03
Omission
2.0 Hz (E)
1.9 Hz (I)

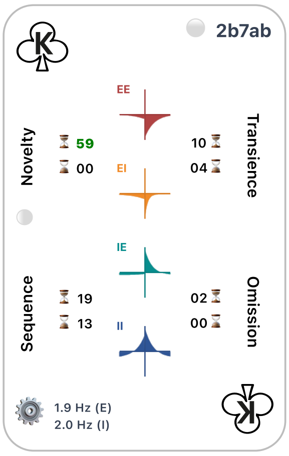
K
2b7ab
EE
EI
IE
II
Novelty
59
00
10
04
Transience
Sequence
19
13
02
00
Omission
1.9 Hz (E)
2.0 Hz (I)

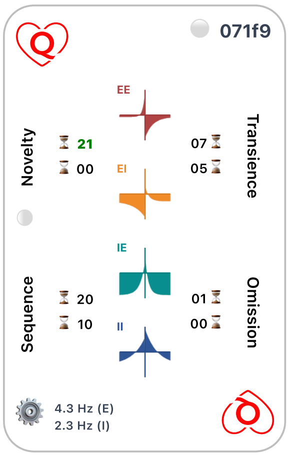
Q
071f9
EE
Novelty
21
00
07
05
Transience
EI
IE
Sequence
20
10
01
00
Omission
II
4.3 Hz (E)
2.3 Hz (I)

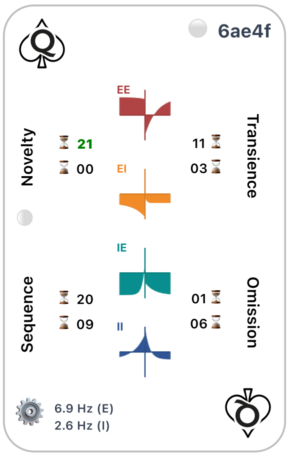
Q
6ae4f
EE
Novelty
21
00
11
03
Transience
EI
IE
Sequence
20
09
01
06
Omission
II
6.9 Hz (E)
2.6 Hz (I)

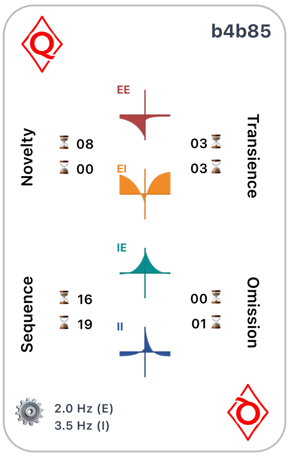
Q
b4b85
EE
Novelty
08
00
03
03
Transience
EI
IE
Sequence
16
19
00
01
Omission
II
2.0 Hz (E)
3.5 Hz (I)

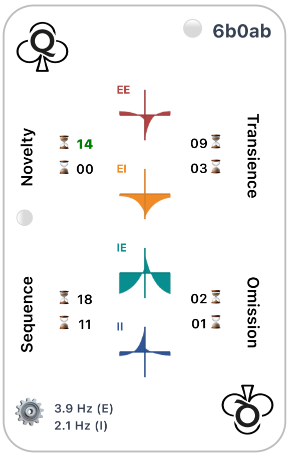
Q
6b0ab
EE
Novelty
14
00
09
03
Transience
EI
IE
Sequence
18
11
02
01
Omission
II
3.9 Hz (E)
2.1 Hz (I)

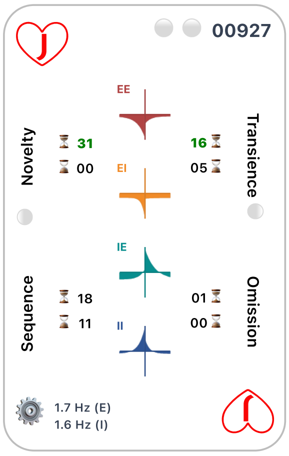
J
00927
EE
Novelty
31
00
16
05
Transience
EI
IE
Sequence
18
11
01
00
Omission
II
1.7 Hz (E)
1.6 Hz (I)

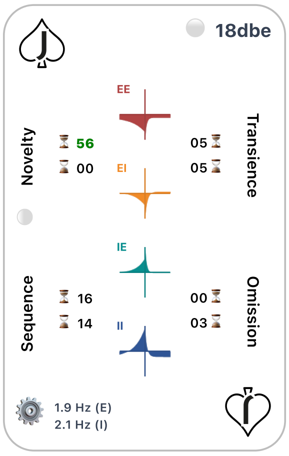
J
18dbe
EE
Novelty
56
00
05
05
Transience
EI
IE
Sequence
16
14
00
03
Omission
II
1.9 Hz (E)
2.1 Hz (I)

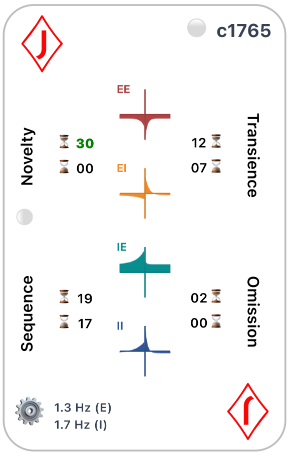
J
c1765
EE
Novelty
30
00
12
07
Transience
EI
IE
Sequence
19
17
02
00
Omission
II
1.3 Hz (E)
1.7 Hz (I)

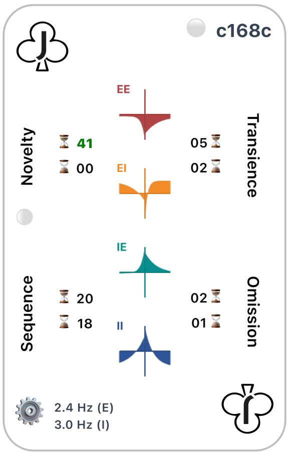
J
c168c
EE
Novelty
41
00
05
02
Transience
EI
IE
Sequence
20
18
02
01
Omission
II
2.4 Hz (E)
3.0 Hz (I)

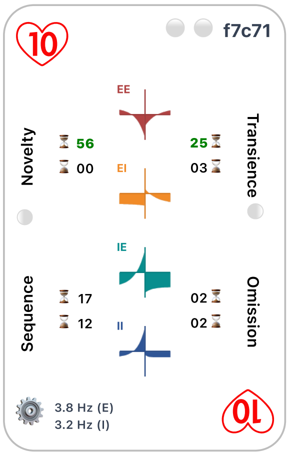
10
f7c71
EE
Novelty
56
00
25
03
Transience
EI
IE
Sequence
17
12
02
02
Omission
II
3.8 Hz (E)
3.2 Hz (I)

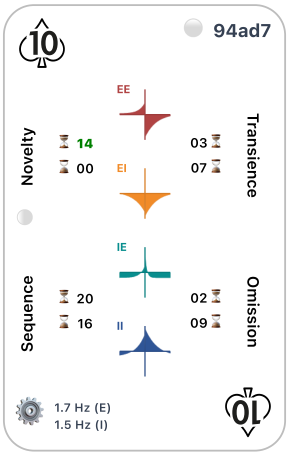
10
94ad7
EE
Novelty
14
00
03
07
Transience
EI
IE
Sequence
20
16
02
09
Omission
II
1.7 Hz (E)
1.5 Hz (I)

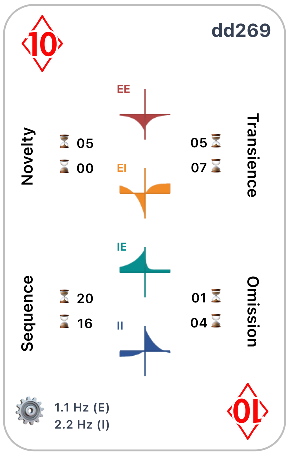
10
dd269
EE
Novelty
05
00
05
07
Transience
EI
IE
Sequence
20
16
01
04
Omission
II
1.1 Hz (E)
2.2 Hz (I)

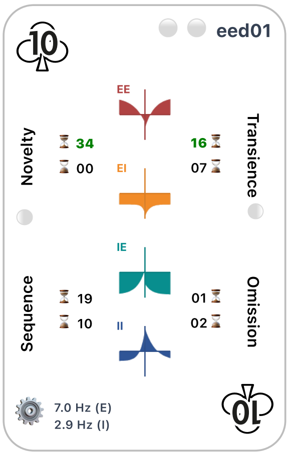
10
eed01
EE
Novelty
34
00
16
07
Transience
EI
IE
Sequence
19
10
01
02
Omission
II
7.0 Hz (E)
2.9 Hz (I)

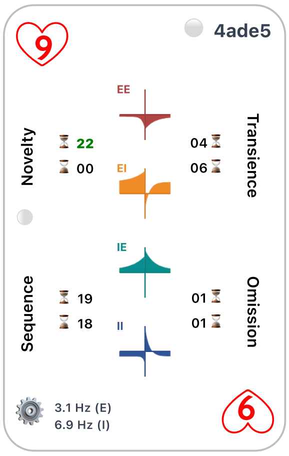
9
4ade5
EE
Novelty
22
00
04
06
Transience
EI
IE
Sequence
19
18
01
01
Omission
II
3.1 Hz (E)
6.9 Hz (I)

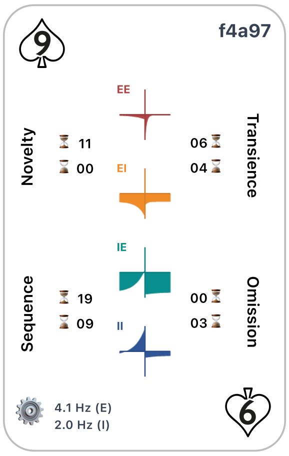
9
f4a97
EE
Novelty
11
00
06
04
Transience
EI
IE
Sequence
19
09
00
03
Omission
II
4.1 Hz (E)
2.0 Hz (I)

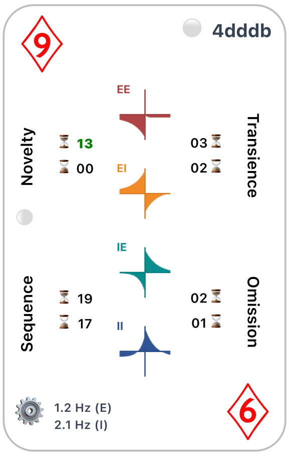
9
4dddb
EE
Novelty
13
00
03
02
Transience
EI
IE
Sequence
19
17
02
01
Omission
II
1.2 Hz (E)
2.1 Hz (I)

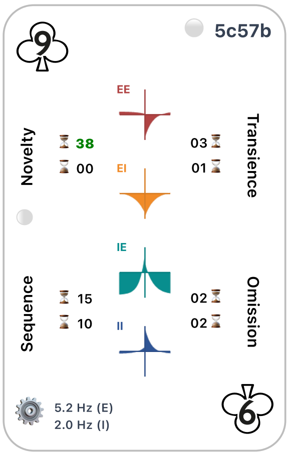
9
5c57b
EE
Novelty
38
00
03
01
Transience
EI
IE
Sequence
15
10
02
02
Omission
II
5.2 Hz (E)
2.0 Hz (I)

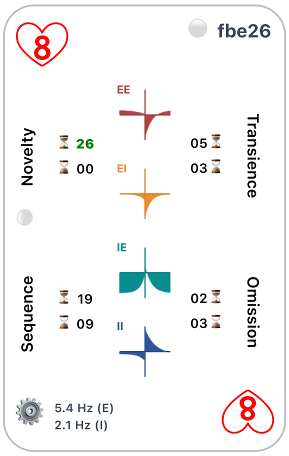
8
fbe26
Novelty
26
00
Transience
05
03
EE
EI
IE
II
Sequence
19
09
Omission
02
03
5.4 Hz (E)
2.1 Hz (I)

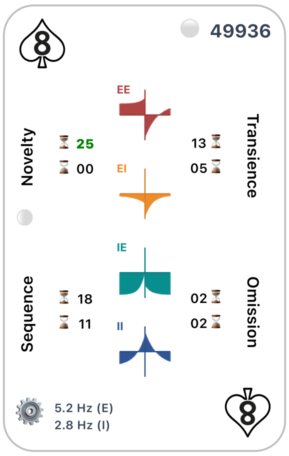
8
49936
Novelty
25
00
Transience
13
05
EE
EI
IE
II
Sequence
18
11
Omission
02
02
5.2 Hz (E)
2.8 Hz (I)

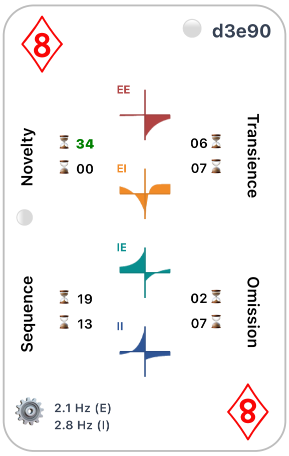
8
d3e90
Novelty
34
00
Transience
06
07
EE
EI
IE
II
Sequence
19
13
Omission
02
07
2.1 Hz (E)
2.8 Hz (I)

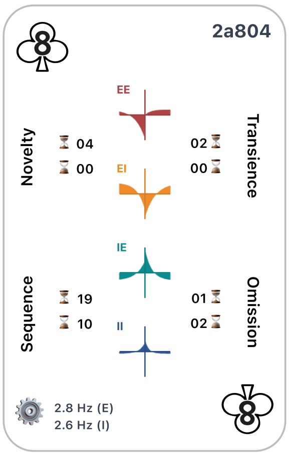
8
2a804
Novelty
04
00
Transience
02
00
EE
EI
IE
II
Sequence
19
10
Omission
01
02
2.8 Hz (E)
2.6 Hz (I)

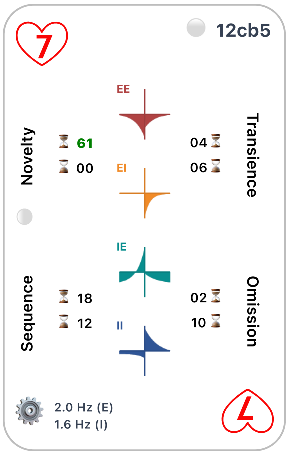
7
12cb5
Novelty
61
00
Transience
04
06
EE
EI
IE
II
Sequence
18
12
Omission
02
10
2.0 Hz (E)
1.6 Hz (I)

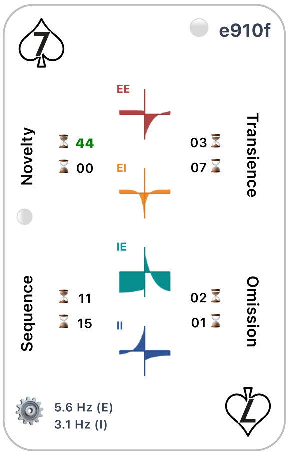
7
e910f
Novelty
44
00
Transience
03
07
EE
EI
IE
II
Sequence
11
15
Omission
02
01
5.6 Hz (E)
3.1 Hz (I)

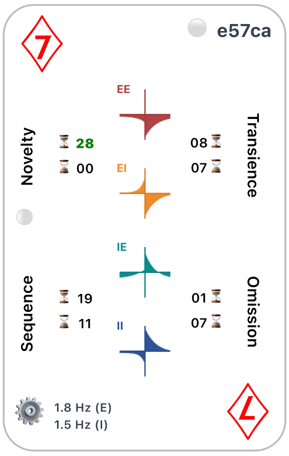
7
e57ca
Novelty
28
00
Transience
08
07
EE
EI
IE
II
Sequence
19
11
Omission
01
07
1.8 Hz (E)
1.5 Hz (I)

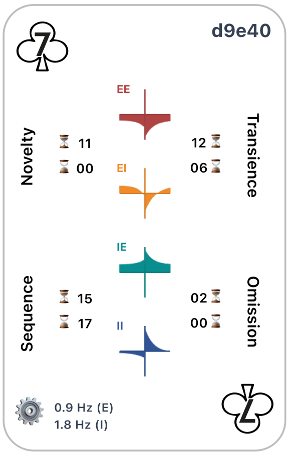
7
d9e40
Novelty
11
00
Transience
12
06
EE
EI
IE
II
Sequence
15
17
Omission
02
00
0.9 Hz (E)
1.8 Hz (I)

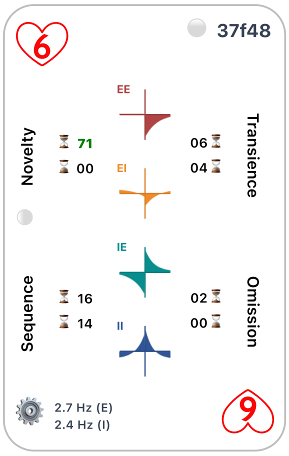
6
37f48
Novelty
71
00
Transience
06
04
EE
EI
IE
II
Sequence
16
14
Omission
02
00
2.7 Hz (E)
2.4 Hz (I)

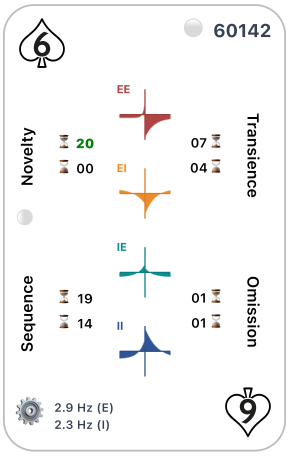
6
60142
Novelty
20
00
Transience
07
04
EE
EI
IE
II
Sequence
19
14
Omission
01
01
2.9 Hz (E)
2.3 Hz (I)

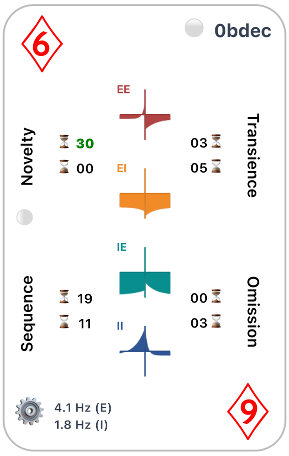
6
0bdec
Novelty
30
00
Transience
03
05
EE
EI
IE
II
Sequence
19
11
Omission
00
03
4.1 Hz (E)
1.8 Hz (I)

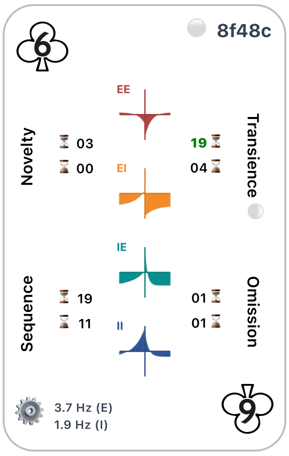
6
8f48c
Novelty
03
00
Transience
19
04
EE
EI
IE
II
Sequence
19
11
Omission
01
01
3.7 Hz (E)
1.9 Hz (I)

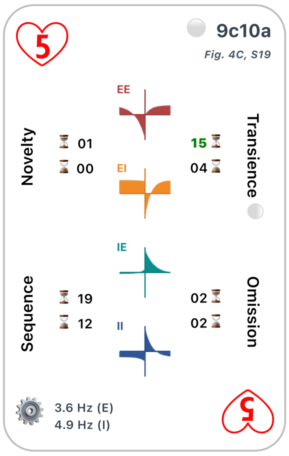
5
9c10a
Fig. 4C, S19
Novelty
01
00
Transience
15
04
EE
EI
IE
II
Sequence
19
12
Omission
02
02
3.6 Hz (E)
4.9 Hz (I)

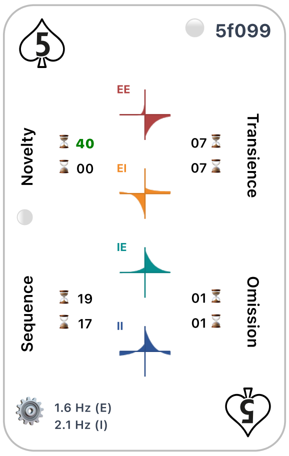
5
5f099
Novelty
40
00
Transience
07
07
EE
EI
IE
II
Sequence
19
17
Omission
01
01
1.6 Hz (E)
2.1 Hz (I)

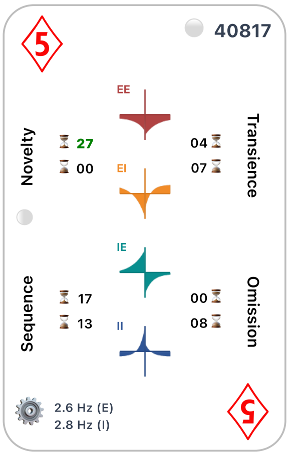
5
40817
Novelty
27
00
Transience
04
07
EE
EI
IE
II
Sequence
17
13
Omission
00
08
2.6 Hz (E)
2.8 Hz (I)

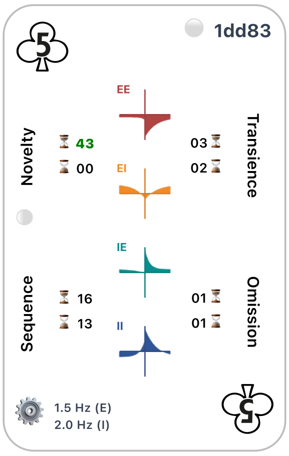
5
1dd83
Novelty
43
00
Transience
03
02
EE
EI
IE
II
Sequence
16
13
Omission
01
01
1.5 Hz (E)
2.0 Hz (I)

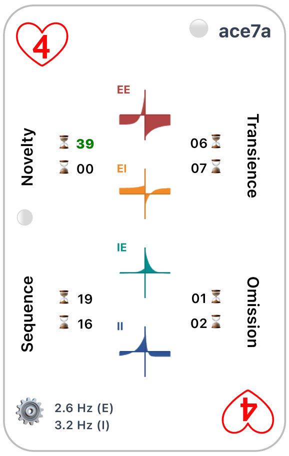
4
ace7a
EE
Novelty
39
00
06
07
Transience
EI
IE
Sequence
19
16
01
02
Omission
II
2.6 Hz (E)
3.2 Hz (I)

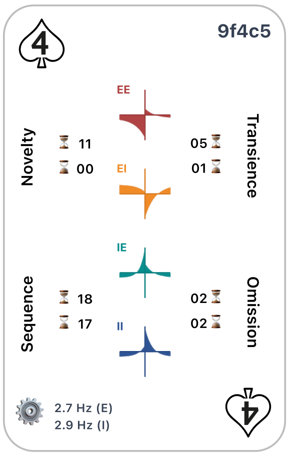
4
9f4c5
EE
Novelty
11
00
05
01
Transience
EI
IE
Sequence
18
17
02
02
Omission
II
2.7 Hz (E)
2.9 Hz (I)

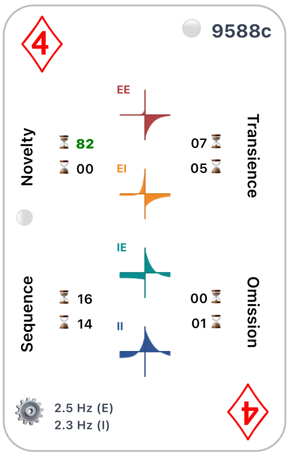
4
9588c
EE
Novelty
82
00
07
05
Transience
EI
IE
Sequence
16
14
00
01
Omission
II
2.5 Hz (E)
2.3 Hz (I)

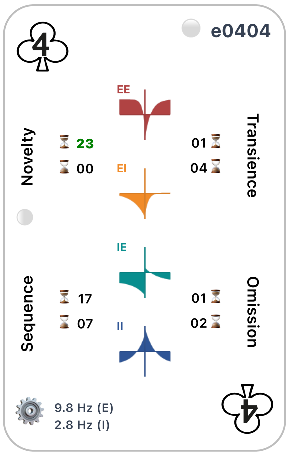
4
e0404
EE
Novelty
23
00
01
04
Transience
EI
IE
Sequence
17
07
01
02
Omission
II
9.8 Hz (E)
2.8 Hz (I)

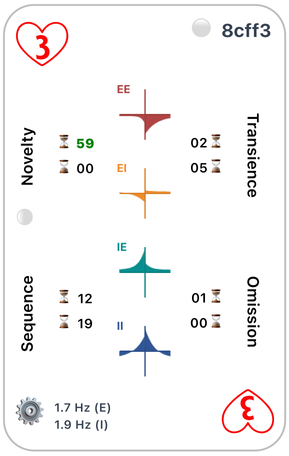
3
8cff3
EE
Novelty
59
00
02
05
Transience
EI
IE
Sequence
12
19
01
00
Omission
II
1.7 Hz (E)
1.9 Hz (I)

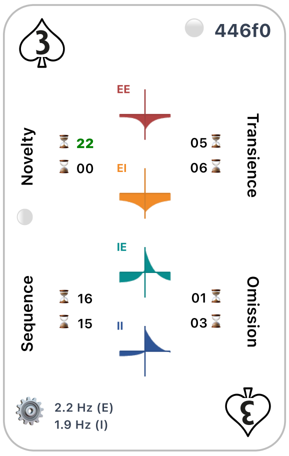
3
446f0
EE
Novelty
22
00
05
06
Transience
EI
IE
Sequence
16
15
01
03
Omission
II
2.2 Hz (E)
1.9 Hz (I)

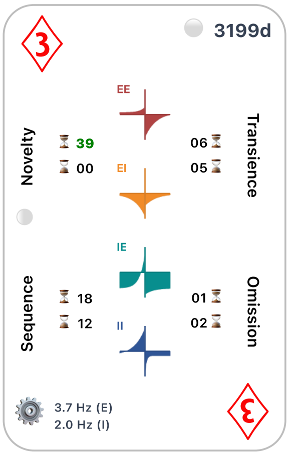
3
3199d
EE
Novelty
39
00
06
05
Transience
EI
IE
Sequence
18
12
01
02
Omission
II
3.7 Hz (E)
2.0 Hz (I)

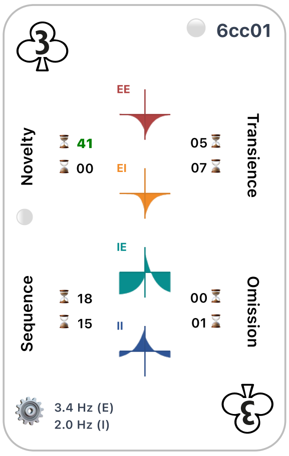
3
6cc01
EE
Novelty
41
00
05
07
Transience
EI
IE
Sequence
18
15
00
01
Omission
II
3.4 Hz (E)
2.0 Hz (I)

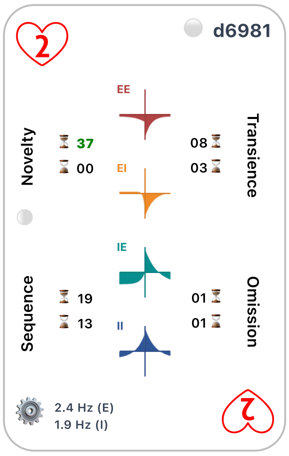
2
d6981
EE
Novelty
37
00
08
03
Transience
EI
IE
Sequence
19
13
01
01
Omission
II
2.4 Hz (E)
1.9 Hz (I)

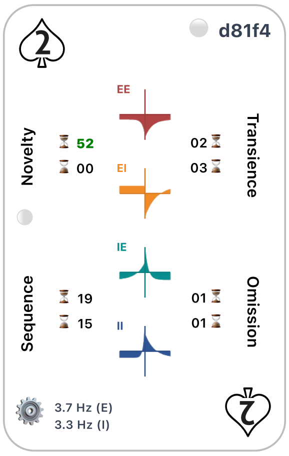
2
d81f4
EE
Novelty
52
00
02
03
Transience
EI
IE
Sequence
19
15
01
01
Omission
II
3.7 Hz (E)
3.3 Hz (I)

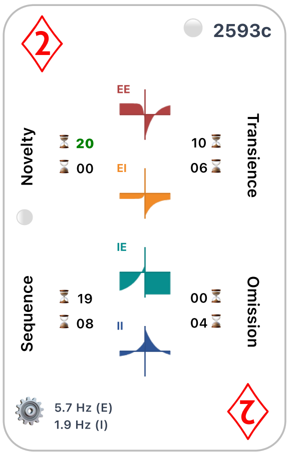
2
2593c
EE
Novelty
20
00
10
06
Transience
EI
IE
Sequence
19
08
00
04
Omission
II
5.7 Hz (E)
1.9 Hz (I)

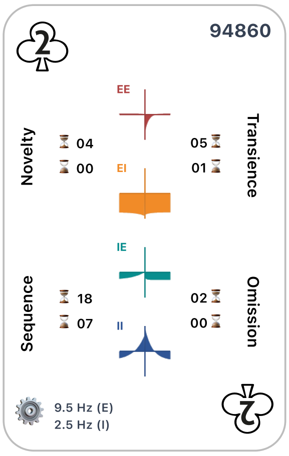
2
94860
EE
Novelty
04
00
05
01
Transience
EI
IE
Sequence
18
07
02
00
Omission
II
9.5 Hz (E)
2.5 Hz (I)

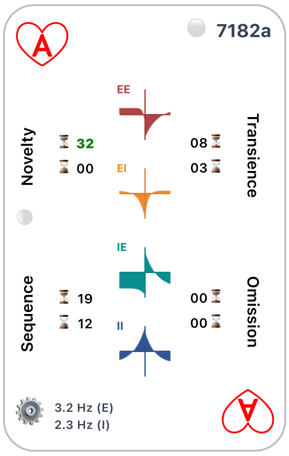
A
7182a
EE
Novelty
32
00
08
03
Transience
EI
IE
Sequence
19
12
00
00
Omission
II
3.2 Hz (E)
2.3 Hz (I)

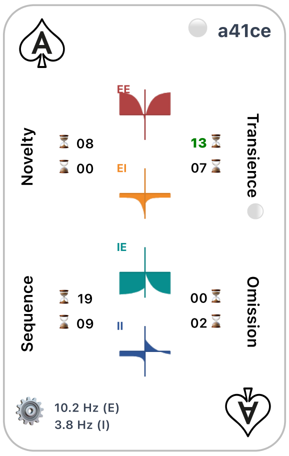
A
a41ce
EE
Novelty
08
00
13
07
Transience
EI
IE
Sequence
19
09
00
02
Omission
II
10.2 Hz (E)
3.8 Hz (I)

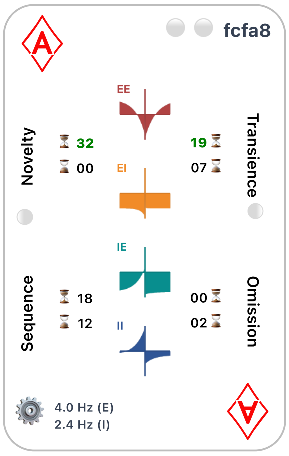
A
fcfa8
EE
Novelty
32
00
19
07
Transience
EI
IE
Sequence
18
12
00
02
Omission
II
4.0 Hz (E)
2.4 Hz (I)

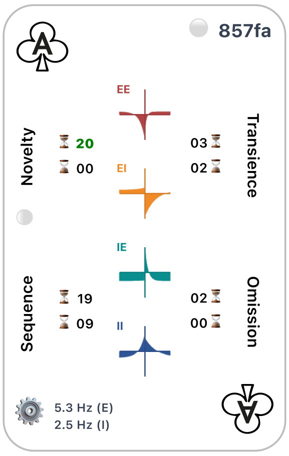
A
857fa
EE
Novelty
20
00
03
02
Transience
EI
IE
Sequence
19
09
02
00
Omission
II
5.3 Hz (E)
2.5 Hz (I)

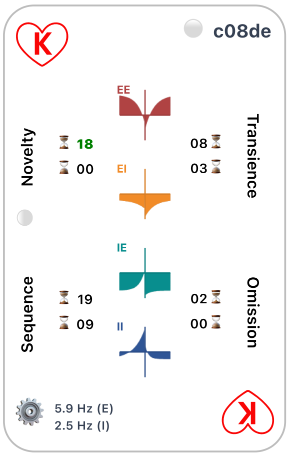
K
c08de
Novelty
18
00
Transience
08
03
EE
EI
IE
II
Sequence
19
09
Omission
02
00
5.9 Hz (E)
2.5 Hz (I)

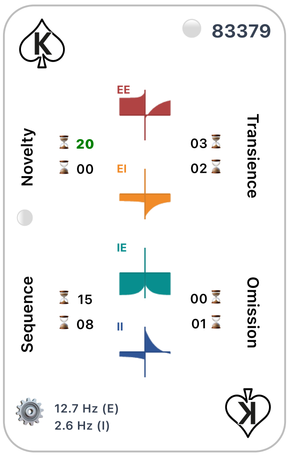
K
83379
Novelty
20
00
Transience
03
02
EE
EI
IE
II
Sequence
15
08
Omission
00
01
12.7 Hz (E)
2.6 Hz (I)

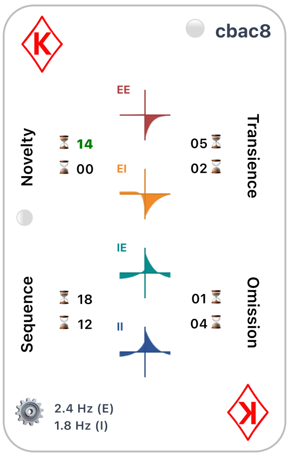
K
cbac8
Novelty
14
00
Transience
05
02
EE
EI
IE
II
Sequence
18
12
Omission
01
04
2.4 Hz (E)
1.8 Hz (I)

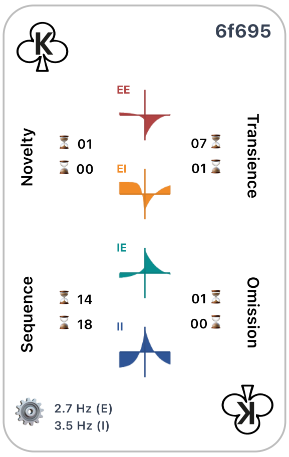
K
6f695
Novelty
01
00
Transience
07
01
EE
EI
IE
II
Sequence
14
18
Omission
01
00
2.7 Hz (E)
3.5 Hz (I)

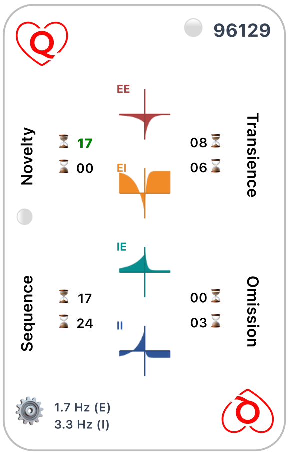
Q
96129
Novelty
17
00
Transience
08
06
EE
EI
IE
II
Sequence
17
24
Omission
00
03
1.7 Hz (E)
3.3 Hz (I)

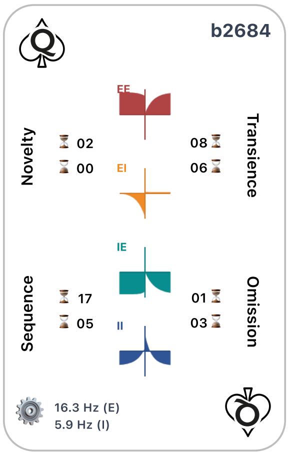
Q
b2684
Novelty
02
00
Transience
08
06
EE
EI
IE
II
Sequence
17
05
Omission
01
03
16.3 Hz (E)
5.9 Hz (I)

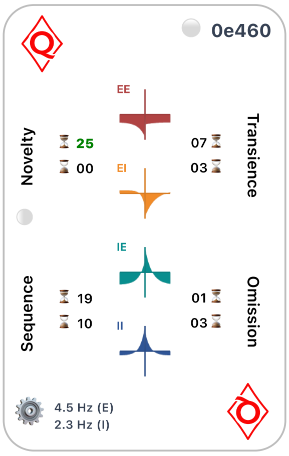
Q
0e460
Novelty
25
00
Transience
07
03
EE
EI
IE
II
Sequence
19
10
Omission
01
03
4.5 Hz (E)
2.3 Hz (I)

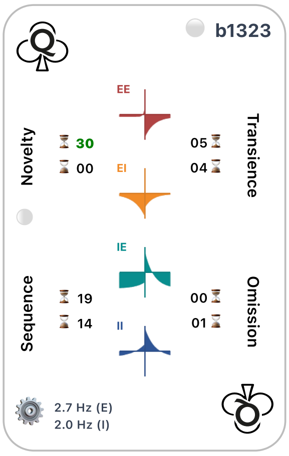
Q
b1323
Novelty
30
00
Transience
05
04
EE
EI
IE
II
Sequence
19
14
Omission
00
01
2.7 Hz (E)
2.0 Hz (I)

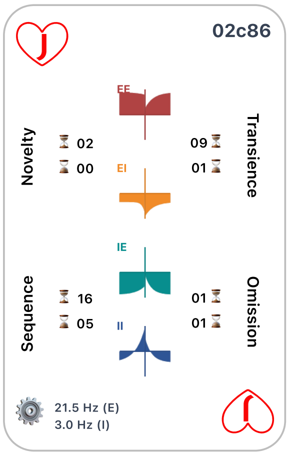
J
02c86
Novelty
02
00
Transience
09
01
EE
EI
IE
II
Sequence
16
05
Omission
01
01
21.5 Hz (E)
3.0 Hz (I)

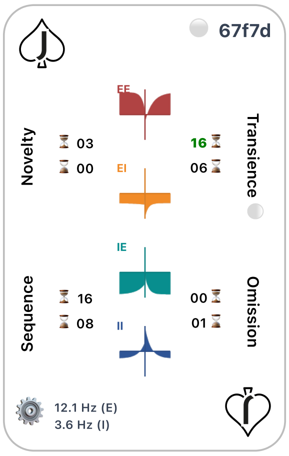
J
67f7d
Novelty
03
00
Transience
16
06
EE
EI
IE
II
Sequence
16
08
Omission
00
01
12.1 Hz (E)
3.6 Hz (I)

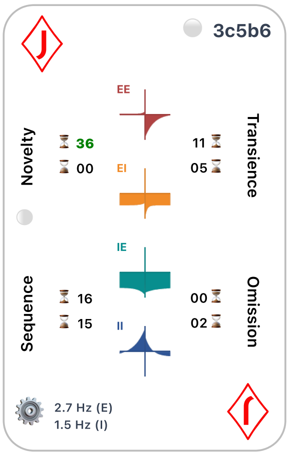
J
3c5b6
Novelty
36
00
Transience
11
05
EE
EI
IE
II
Sequence
16
15
Omission
00
02
2.7 Hz (E)
1.5 Hz (I)

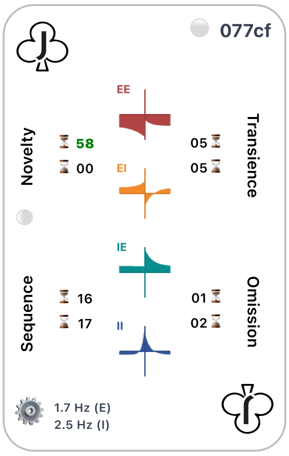
J
077cf
Novelty
58
00
Transience
05
05
EE
EI
IE
II
Sequence
16
17
Omission
01
02
1.7 Hz (E)
2.5 Hz (I)

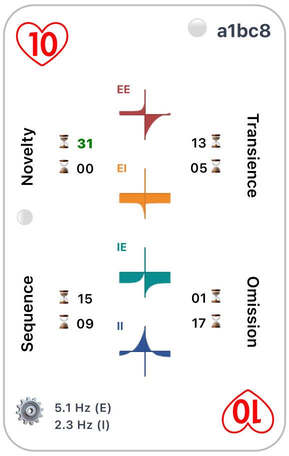
10
a1bc8
Novelty
31
00
Transience
13
05
EE
EI
IE
II
Sequence
15
09
Omission
01
17
5.1 Hz (E)
2.3 Hz (I)

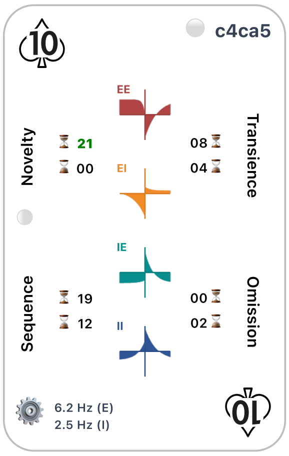
10
c4ca5
Novelty
21
00
Transience
08
04
EE
EI
IE
II
Sequence
19
12
Omission
00
02
6.2 Hz (E)
2.5 Hz (I)

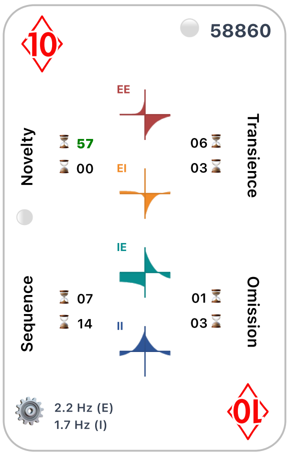
10
58860
Novelty
57
00
Transience
06
03
EE
EI
IE
II
Sequence
07
14
Omission
01
03
2.2 Hz (E)
1.7 Hz (I)

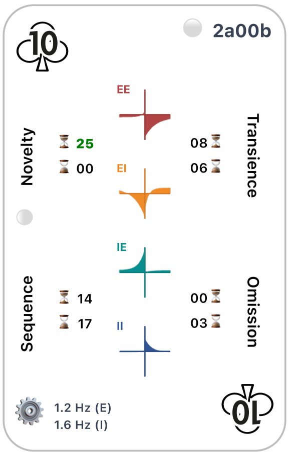
10
2a00b
Novelty
25
00
Transience
08
06
EE
EI
IE
II
Sequence
14
17
Omission
00
03
1.2 Hz (E)
1.6 Hz (I)

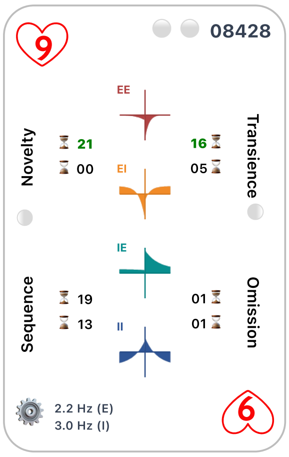
08428
Novelty
21
00
EE
EI
16
05
Transience
Sequence
19
13
IE
II
01
01
Omission
2.2 Hz (E)
3.0 Hz (I)

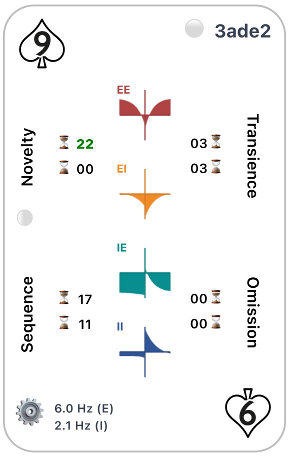
3ade2
Novelty
22
00
EE
EI
03
03
Transience
Sequence
17
11
IE
II
00
00
Omission
6.0 Hz (E)
2.1 Hz (I)

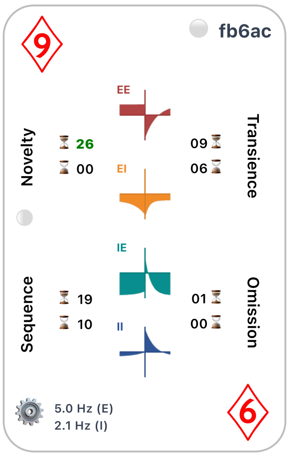
fb6ac
Novelty
26
00
EE
EI
09
06
Transience
Sequence
19
10
IE
II
01
00
Omission
5.0 Hz (E)
2.1 Hz (I)

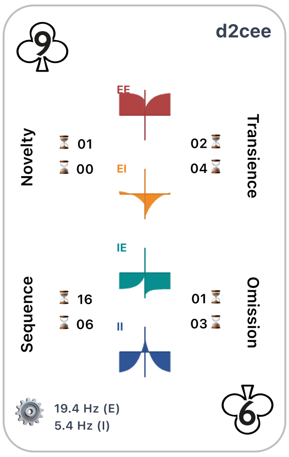
d2cee
Novelty
01
00
EE
EI
02
04
Transience
Sequence
16
06
IE
II
01
03
Omission
19.4 Hz (E)
5.4 Hz (I)

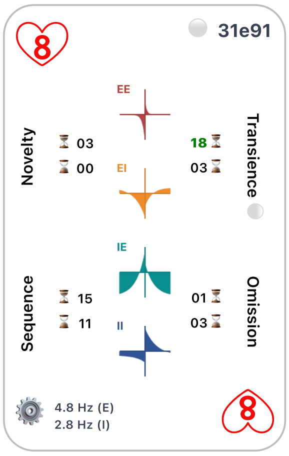
31e91
Novelty
03
00
EE
EI
18
03
Transience
Sequence
15
11
IE
II
01
03
Omission
4.8 Hz (E)
2.8 Hz (I)

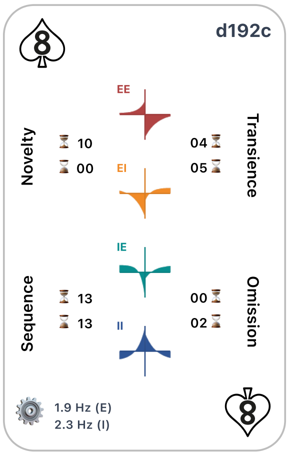
d192c
Novelty
10
00
EE
EI
04
05
Transience
Sequence
13
13
IE
II
00
02
Omission
1.9 Hz (E)
2.3 Hz (I)

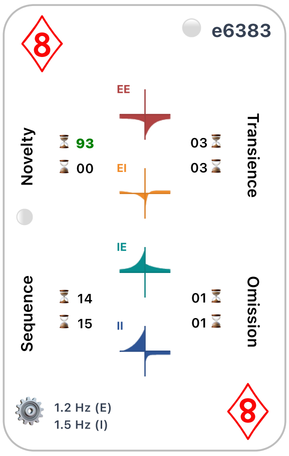
e6383
Novelty
93
00
EE
EI
03
03
Transience
Sequence
14
15
IE
II
01
01
Omission
1.2 Hz (E)
1.5 Hz (I)

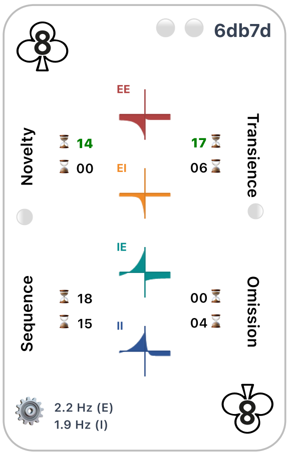
6db7d
Novelty
14
00
EE
EI
17
06
Transience
Sequence
18
15
IE
II
00
04
Omission
2.2 Hz (E)
1.9 Hz (I)

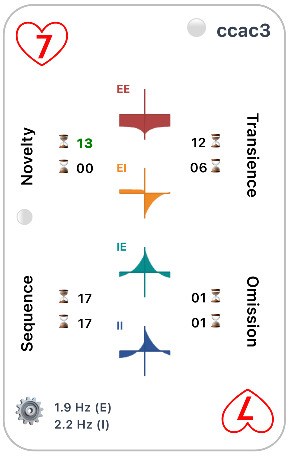
ccac3
Novelty
13
00
EE
EI
12
06
Transience
Sequence
17
17
IE
II
01
01
Omission
1.9 Hz (E)
2.2 Hz (I)

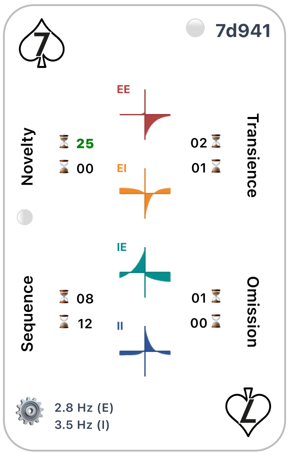
7d941
Novelty
25
00
EE
EI
02
01
Transience
Sequence
08
12
IE
II
01
00
Omission
2.8 Hz (E)
3.5 Hz (I)

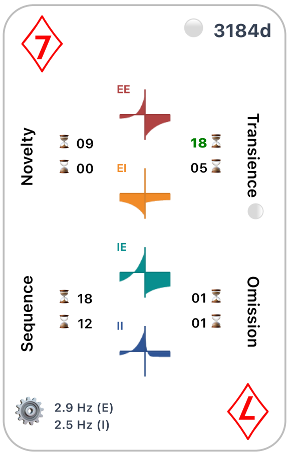
3184d
Novelty
09
00
EE
EI
18
05
Transience
Sequence
18
12
IE
II
01
01
Omission
2.9 Hz (E)
2.5 Hz (I)

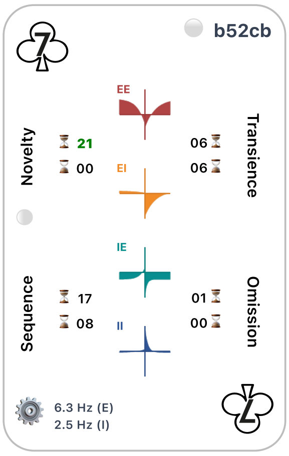
b52cb
Novelty
21
00
EE
EI
06
06
Transience
Sequence
17
08
IE
II
01
00
Omission
6.3 Hz (E)
2.5 Hz (I)

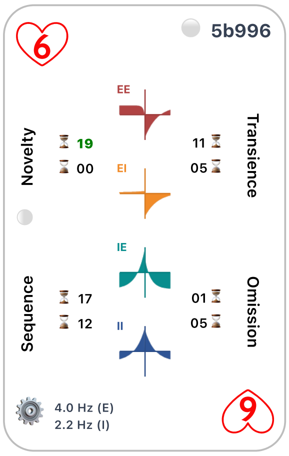
5b996
Novelty
19
00
EE
EI
11
05
Transience
Sequence
17
12
IE
II
01
05
Omission
4.0 Hz (E)
2.2 Hz (I)

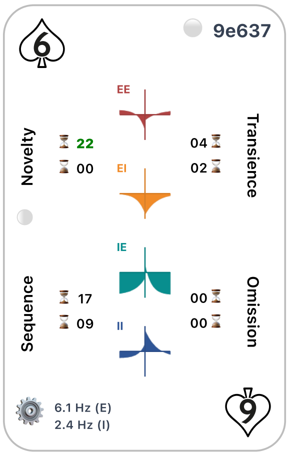
9e637
Novelty
22
00
EE
EI
04
02
Transience
Sequence
17
09
IE
II
00
00
Omission
6.1 Hz (E)
2.4 Hz (I)

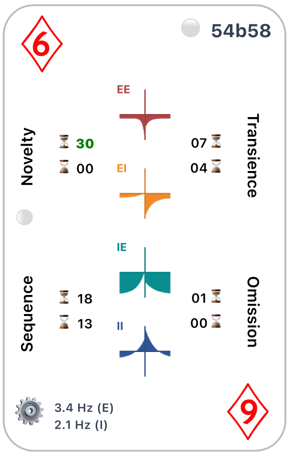
54b58
Novelty
30
00
EE
EI
07
04
Transience
Sequence
18
13
IE
II
01
00
Omission
3.4 Hz (E)
2.1 Hz (I)

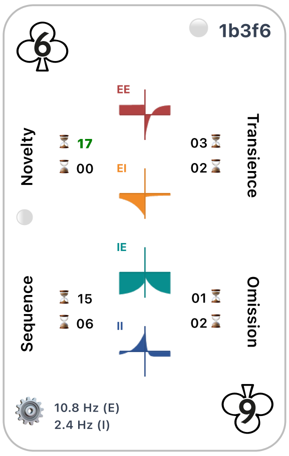
1b3f6
Novelty
17
00
EE
EI
03
02
Transience
Sequence
15
06
IE
II
01
02
Omission
10.8 Hz (E)
2.4 Hz (I)

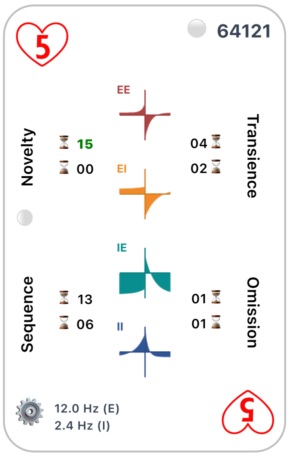
5
64121
EE
EI
IE
II
Novelty
15
00
04
02
Transience
Sequence
13
06
01
01
Omission
12.0 Hz (E)
2.4 Hz (I)

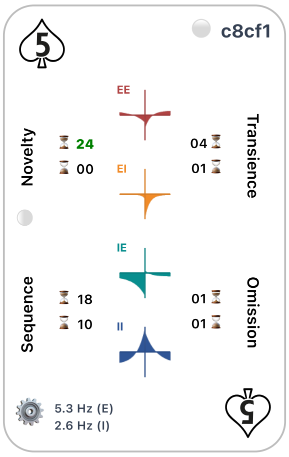
5
c8cf1
EE
EI
IE
II
Novelty
24
00
04
01
Transience
Sequence
18
10
01
01
Omission
5.3 Hz (E)
2.6 Hz (I)

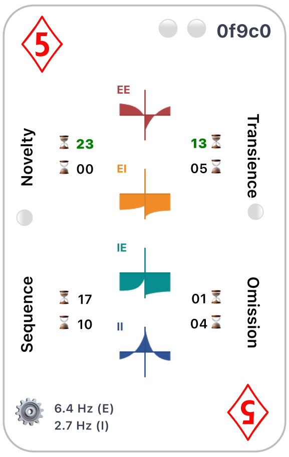
5
0f9c0
EE
EI
IE
II
Novelty
23
00
13
05
Transience
Sequence
17
10
01
04
Omission
6.4 Hz (E)
2.7 Hz (I)

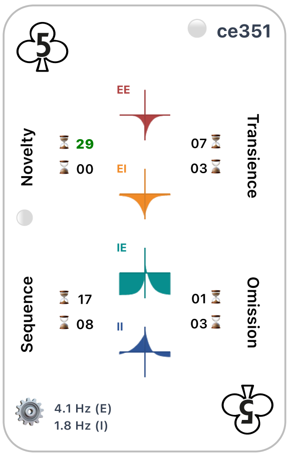
5
ce351
EE
EI
IE
II
Novelty
29
00
07
03
Transience
Sequence
17
08
01
03
Omission
4.1 Hz (E)
1.8 Hz (I)

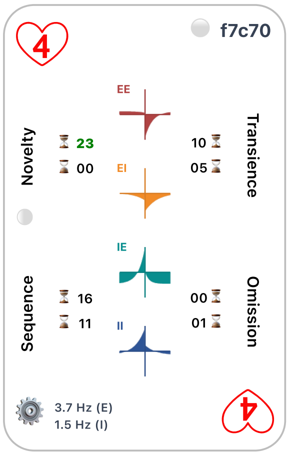
4
f7c70
EE
EI
IE
II
Novelty
23
00
10
05
Transience
Sequence
16
11
00
01
Omission
3.7 Hz (E)
1.5 Hz (I)

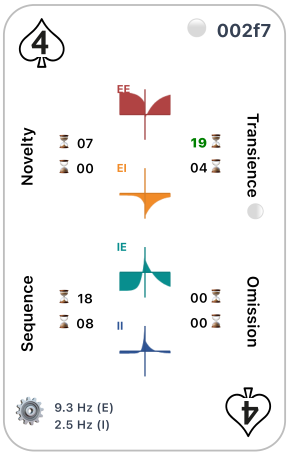
4
002f7
EE
EI
IE
II
Novelty
07
00
19
04
Transience
Sequence
18
08
00
00
Omission
9.3 Hz (E)
2.5 Hz (I)

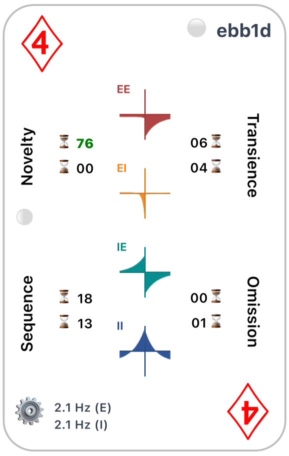
4
ebb1d
EE
EI
IE
II
Novelty
76
00
06
04
Transience
Sequence
18
13
00
01
Omission
2.1 Hz (E)
2.1 Hz (I)

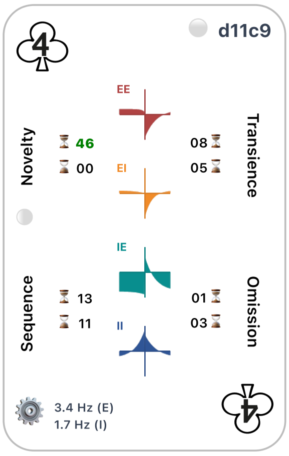
4
d11c9
EE
EI
IE
II
Novelty
46
00
08
05
Transience
Sequence
13
11
01
03
Omission
3.4 Hz (E)
1.7 Hz (I)

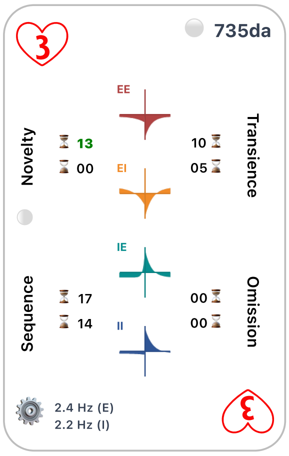
3
735da
EE
EI
IE
II
Novelty
13
00
10
05
Transience
Sequence
17
14
00
00
Omission
2.4 Hz (E)
2.2 Hz (I)

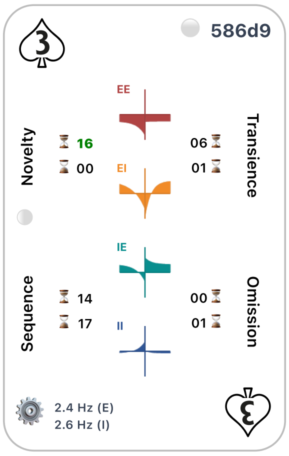
3
586d9
EE
EI
IE
II
Novelty
16
00
06
01
Transience
Sequence
14
17
00
01
Omission
2.4 Hz (E)
2.6 Hz (I)

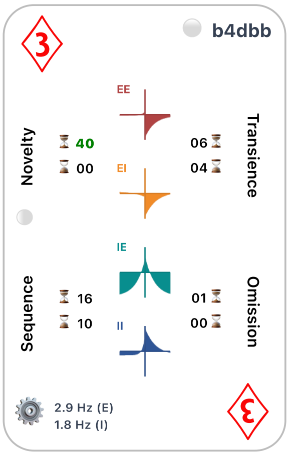
3
b4dbb
EE
EI
IE
II
Novelty
40
00
06
04
Transience
Sequence
16
10
01
00
Omission
2.9 Hz (E)
1.8 Hz (I)

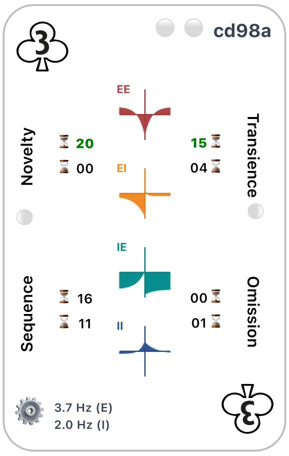
3
cd98a
EE
EI
IE
II
Novelty
20
00
15
04
Transience
Sequence
16
11
00
01
Omission
3.7 Hz (E)
2.0 Hz (I)

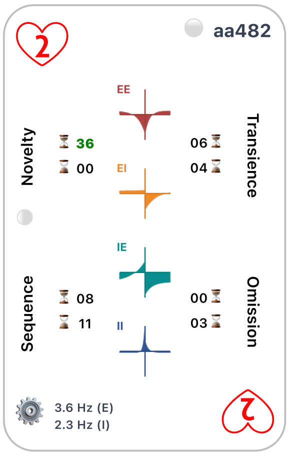
2
aa482
EE
EI
IE
II
Novelty
36
00
06
04
Transience
Sequence
08
11
00
03
Omission
3.6 Hz (E)
2.3 Hz (I)

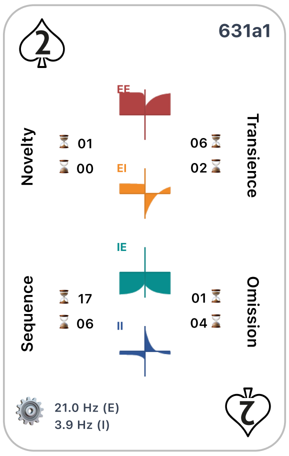
2
631a1
EE
EI
IE
II
Novelty
01
00
06
02
Transience
Sequence
17
06
01
04
Omission
21.0 Hz (E)
3.9 Hz (I)

### Table of rule quadruplet parameters

| Rule quadruplet ID | EE: $\tau_{pre}$, $\tau_{post}$, $\alpha$, $\beta$, $\gamma$, $\kappa$ | EI: $\tau_{pre}$, $\tau_{post}$, $\alpha$, $\beta$, $\gamma$, $\kappa$ | IE: $\tau_{pre}$, $\tau_{post}$, $\alpha$, $\beta$, $\gamma$, $\kappa$ | II: $\tau_{pre}$, $\tau_{post}$, $\alpha$, $\beta$, $\gamma$, $\kappa$ |
|---|---|---|---|---|
| **00028**ef499614197e2ce6a41689ef0f6 | 0.073,0.024,1.539,-0.746,-1.887,-0.443 | 0.090,0.018,0.501,-1.100,-0.434,-0.033 | 0.044,0.064,-1.111,0.138,1.791,0.759 | 0.069,0.012,-0.104,-0.156,1.724,0.961 |
| **002f7**614631abbf2369abc9f138c0a89 | 0.086,0.046,1.053,0.161,-1.424,-0.610 | 0.078,0.028,0.465,-0.617,-1.679,-0.840 | 0.097,0.025,-0.474,-0.246,1.253,1.579 | 0.050,0.014,-0.030,-0.090,0.745,1.701 |
| **00319**500f348db0ebd7af21051778bb8 | 0.052,0.063,1.501,-1.715,0.055,-1.102 | 0.058,0.074,0.481,-0.378,-1.584,-0.087 | 0.013,0.093,-0.967,1.058,-0.309,1.311 | 0.086,0.076,-0.862,0.666,0.963,0.989 |
| **00525**69281202c0d0528831365119893 | 0.096,0.022,1.201,-1.085,-1.704,0.091 | 0.038,0.060,0.526,-0.767,-1.276,0.568 | 0.014,0.055,-0.828,0.775,0.559,0.390 | 0.025,0.056,-0.754,0.694,1.327,0.600 |
| **006f9**224ac035f0821fc027fd7d03ac8 | 0.089,0.025,1.749,-1.257,-1.633,-0.713 | 0.020,0.087,1.137,-1.032,-0.833,-0.208 | 0.026,0.039,-1.652,1.594,-0.491,1.095 | 0.011,0.092,-1.562,0.822,1.879,1.822 |
| **00927**92d64c6713f67b36d6572c348e1 | 0.043,0.057,1.554,-1.670,-0.440,-1.232 | 0.091,0.023,0.814,-1.093,-0.054,-1.325 | 0.078,0.041,-1.636,1.292,1.512,0.518 | 0.035,0.047,-1.222,1.070,0.939,1.639 |
| **00ab6**b8a8c6dd0e0fe198217091f94d3 | 0.012,0.047,1.297,-1.389,-0.428,-0.571 | 0.014,0.053,1.662,-1.087,-0.738,-0.967 | 0.061,0.042,-0.969,1.280,1.078,0.187 | 0.095,0.065,-0.334,-0.426,1.988,1.431 |
| **00d85**be5a32b01f77971460d95438e6b | 0.078,0.039,1.530,-1.102,-1.674,-1.311 | 0.029,0.076,1.955,-1.478,-1.952,-1.203 | 0.046,0.062,-1.838,1.987,-1.838,1.131 | 0.030,0.053,-0.565,0.325,1.244,0.762 |
| **00dc4**23e09f87ddac9d0c5c8eb10403b | 0.074,0.059,1.804,-1.477,-0.982,-1.014 | 0.020,0.052,1.360,-1.933,-0.412,0.007 | 0.041,0.011,-1.510,1.160,-1.249,1.862 | 0.099,0.074,-1.067,0.376,0.893,1.840 |
| **0189c**7afe90a8808261ca3657b6869fd | 0.040,0.066,1.695,-1.177,-1.621,-1.034 | 0.062,0.047,1.222,-1.360,-0.892,-1.164 | 0.039,0.064,-1.843,1.210,0.075,0.963 | 0.022,0.031,-0.205,-0.072,1.215,1.837 |
| **01a3b**79859440f355308093007e403b0 | 0.029,0.059,1.505,-1.538,0.082,-1.049 | 0.060,0.021,0.614,-0.380,-1.058,-1.192 | 0.041,0.034,-0.522,0.291,0.591,1.313 | 0.062,0.036,-0.219,-0.151,1.242,1.116 |
| **01a6f**dbf6cf52691af3ffe18d345e835 | 0.087,0.026,0.922,-0.702,-0.680,0.924 | 0.042,0.069,1.225,-1.519,-0.539,-1.850 | 0.031,0.062,-1.895,0.754,1.916,0.453 | 0.052,0.085,-1.026,0.587,-0.039,1.393 |
| **01b62**12ddcbfc95bb7517c15759c35e2 | 0.083,0.055,1.123,-1.010,-0.816,0.187 | 0.080,0.051,0.993,-0.390,-1.014,-0.039 | 0.038,0.067,-0.627,1.020,1.474,0.488 | 0.016,0.075,-0.670,0.225,1.668,1.417 |
| **02004**ec498cbd879f2de36761f582ebf | 0.035,0.079,1.735,-1.630,-1.027,-1.368 | 0.059,0.093,1.355,-1.099,-0.862,-1.484 | 0.083,0.085,-1.812,1.943,1.565,0.093 | 0.020,0.062,-1.156,0.993,-0.861,1.292 |
| **021d2**b5861beffb2522ce976e3ffff93 | 0.014,0.073,0.615,-0.573,-0.352,-0.519 | 0.017,0.032,0.598,-0.456,1.871,-1.215 | 0.099,0.062,-0.459,0.292,1.286,0.315 | 0.079,0.019,-1.101,0.734,1.858,-0.679 |
| **023e3**2688212627ac12bd8e2a9cc2182 | 0.099,0.019,1.448,-0.965,-1.755,1.402 | 0.012,0.089,0.553,-0.748,1.564,-1.351 | 0.044,0.023,-1.888,0.878,0.429,1.993 | 0.053,0.066,-1.510,1.172,0.802,1.623 |
| **0286a**7f88886e576b302f39fd8ce9235 | 0.047,0.098,1.970,-1.400,-1.857,-1.896 | 0.031,0.027,0.978,-1.734,-1.893,-0.136 | 0.060,0.035,-1.841,0.790,1.958,1.605 | 0.096,0.018,-0.285,-0.110,1.985,1.650 |
| **02abe**c8a96f449978288b92f3447499d | 0.049,0.045,1.259,-1.202,-1.483,-0.934 | 0.067,0.074,0.776,-1.137,-0.486,1.247 | 0.073,0.043,-1.205,1.118,1.569,0.684 | 0.022,0.057,-1.525,1.248,0.348,1.692 |
| **02b11**252cadcaa8c35624023af10ad08 | 0.066,0.062,1.777,-1.915,0.598,-1.337 | 0.057,0.091,1.156,-1.676,-1.978,-0.463 | 0.083,0.035,-1.977,0.950,1.026,0.180 | 0.058,0.060,-0.980,0.753,1.768,0.502 |
| **02c52**4f5175884dcc09fc2d9fb609161 | 0.045,0.063,1.002,-0.954,-0.348,-1.101 | 0.041,0.072,1.605,-1.402,-0.518,-0.462 | 0.095,0.041,-1.883,1.913,1.850,1.586 | 0.065,0.056,-0.139,-0.156,0.830,1.426 |
| **02c86**2b73db41d48535eae4fd319f45e | 0.065,0.068,1.526,1.132,-1.912,-0.293 | 0.067,0.020,0.449,-1.337,-1.495,-1.186 | 0.049,0.067,-1.431,-0.101,1.378,0.591 | 0.052,0.073,-0.973,0.520,1.037,1.433 |
| **0350d**23a3598ac8d41fd839837fc3d2c | 0.074,0.050,1.847,-1.813,-1.922,0.144 | 0.094,0.085,1.882,-1.447,-0.554,-1.625 | 0.043,0.035,-1.216,1.654,1.797,-1.766 | 0.041,0.056,-0.427,0.021,1.662,1.846 |
| **03902**5116169fe7db40626e50d257cbf | 0.099,0.095,1.928,-1.508,-0.486,-1.692 | 0.080,0.084,1.333,-1.164,0.212,-1.760 | 0.084,0.033,-1.269,0.978,0.985,1.523 | 0.083,0.081,-0.746,0.098,1.431,1.823 |
| **03da0**9810bfb29c82f00c3bd7c9f83d2 | 0.054,0.042,1.539,-1.660,-1.361,0.198 | 0.070,0.077,0.922,-1.748,-0.194,-0.033 | 0.099,0.079,-0.960,0.181,1.581,0.376 | 0.079,0.034,-0.708,0.444,1.816,0.360 |
| **04041**c5e966ec0b9a6d091b865e56ea6 | 0.048,0.049,1.626,-1.958,-0.964,-0.262 | 0.091,0.012,0.538,-0.776,0.091,-0.463 | 0.094,0.040,-1.119,0.765,0.254,1.346 | 0.080,0.021,-1.055,0.701,1.929,1.217 |
| **04181**6cb956cae2153b1ae4759120f57 | 0.036,0.093,0.135,0.020,-0.408,-0.650 | 0.077,0.056,1.079,-1.210,-0.107,-1.435 | 0.084,0.031,-1.546,1.297,0.211,0.838 | 0.034,0.060,-0.483,0.336,1.935,0.710 |
| **04252**bc1c373428394e728a5fe6e8b84 | 0.096,0.020,1.639,-1.408,-1.909,1.149 | 0.087,0.045,0.405,-0.197,-0.891,-1.180 | 0.023,0.058,-1.294,0.872,0.044,0.523 | 0.053,0.093,-1.058,0.772,1.932,0.596 |
| **048d1**22f9c6f90bdf3c24d9b5ed43e71 | 0.051,0.031,1.672,-0.863,-1.640,-0.391 | 0.068,0.094,0.648,-0.112,-1.699,-1.530 | 0.048,0.068,-1.197,0.182,1.790,0.015 | 0.070,0.082,-0.725,0.108,1.945,1.447 |
| **04a9d**e0f96d4905e8c83c1339c6d6b16 | 0.013,0.061,1.034,-0.858,-0.279,-0.959 | 0.031,0.031,0.696,-0.572,-0.735,-0.718 | 0.086,0.049,-1.721,1.772,1.736,0.513 | 0.087,0.041,-0.401,0.293,-0.252,1.527 |
| **04e7d**16288e27aafb8049e9c39e5956b | 0.085,0.027,0.772,-0.673,-1.643,-0.146 | 0.090,0.081,1.031,-0.507,-1.958,0.729 | 0.070,0.098,-0.823,0.628,1.758,1.703 | 0.080,0.091,-1.545,1.469,1.642,-0.728 |
| **0501a**e322cc722a44eab0b599c9bdded | 0.097,0.021,1.547,-1.430,-1.920,1.325 | 0.038,0.060,0.282,-0.010,-1.880,-1.458 | 0.080,0.023,-0.882,0.915,-0.095,1.704 | 0.097,0.077,-1.364,0.850,1.846,1.192 |
| **05138**f4c8f6bf651fdee3265ae5bc6c6 | 0.067,0.013,1.497,-1.715,-1.697,-0.392 | 0.078,0.079,0.342,-0.389,-1.101,0.749 | 0.053,0.098,-1.046,0.395,1.343,1.828 | 0.079,0.072,-0.249,0.114,1.902,-0.535 |
| **0519b**622383297ca5b67b56f433984ad | 0.061,0.042,1.161,-1.329,-1.827,0.535 | 0.045,0.035,0.479,-0.646,1.473,-0.991 | 0.086,0.084,-1.515,1.170,1.925,1.909 | 0.097,0.072,-1.471,1.049,1.507,1.816 |
| **0519c**81917a4561e34d24e8cea8e1f99 | 0.098,0.010,1.548,-1.605,-0.473,0.120 | 0.016,0.078,0.738,-1.101,-0.848,-0.779 | 0.012,0.060,-1.881,0.896,1.949,1.883 | 0.085,0.062,-1.730,1.395,0.426,1.980 |
| **053c9**9cade86adff635794577b7f6dbc | 0.076,0.011,1.962,-1.539,-1.940,-0.361 | 0.047,0.034,0.929,-1.389,-1.897,-0.691 | 0.033,0.084,-1.860,0.773,1.005,0.820 | 0.088,0.071,-1.155,0.877,-0.525,1.948 |
| **0567a**38f4dc613d77604bc30ac6f058b | 0.088,0.021,1.161,-1.179,-1.196,-0.479 | 0.060,0.070,1.111,-0.667,1.039,-0.723 | 0.070,0.038,-0.499,0.715,1.339,0.729 | 0.095,0.037,-0.265,-0.215,1.309,1.610 |
| **058d7**5245b86f1399b67ae0e97d0dfa3 | 0.053,0.062,1.376,-1.191,-1.276,-0.145 | 0.021,0.019,1.001,-1.500,1.345,0.785 | 0.071,0.063,-1.696,1.183,1.852,0.776 | 0.075,0.097,-0.538,0.372,0.123,0.332 |
| **05a36**2bde6e01e9b0dccde0c705229b9 | 0.014,0.078,1.479,-1.761,0.161,-0.777 | 0.029,0.031,1.322,-0.883,-1.492,-1.023 | 0.053,0.047,-1.107,1.184,1.878,-0.119 | 0.068,0.047,-0.850,0.351,1.821,0.503 |
| **05ab3**149ae8582dd6b0d14835d770eb0 | 0.092,0.047,1.670,-1.723,-0.853,-0.543 | 0.092,0.056,0.596,-0.521,-0.445,-0.348 | 0.079,0.096,-1.758,1.954,1.487,1.389 | 0.053,0.077,-1.970,1.741,-0.308,1.742 |
| **05c65**5f66fe337f3808aad2cc2c14340 | 0.093,0.016,1.878,-0.496,-1.873,-0.594 | 0.100,0.014,0.647,-1.890,-0.913,1.098 | 0.030,0.089,-1.514,0.069,1.440,1.280 | 0.082,0.042,-0.611,0.160,0.961,1.929 |
| **05cae**c20f2319c608dceaa1d01c4f12f | 0.081,0.036,1.295,-1.259,-0.271,-0.177 | 0.039,0.065,0.628,-0.399,-0.903,-1.305 | 0.053,0.046,-0.577,0.506,1.314,0.601 | 0.037,0.027,-0.420,0.280,1.612,1.217 |
| **061ce**d3f3e0943e5c01e95a4bc71abb8 | 0.048,0.081,1.576,-1.188,-1.800,-0.835 | 0.019,0.084,1.577,-1.396,-1.414,0.146 | 0.090,0.035,-1.928,1.860,1.167,1.958 | 0.099,0.052,-0.242,-0.315,1.996,-0.223 |
| **064b6**8d972bd2d9d70527cc697d35e00 | 0.063,0.065,1.461,-1.209,-0.736,0.283 | 0.024,0.095,0.581,-1.332,-0.930,-1.124 | 0.044,0.016,-0.912,-0.026,1.268,1.269 | 0.069,0.081,-0.408,0.129,1.890,-0.226 |
| **0661e**da1630cada54f63344c56b4445e | 0.096,0.019,1.151,-1.136,-0.851,-0.847 | 0.090,0.079,1.365,-1.184,-1.703,0.477 | 0.061,0.030,-1.631,1.927,1.122,0.093 | 0.098,0.073,-1.909,1.367,1.045,1.767 |
| **0667e**02454848716739e01f8cc879a5f | 0.086,0.028,1.583,-1.476,-1.917,-0.133 | 0.071,0.073,0.486,-1.128,-0.166,0.271 | 0.077,0.032,-1.435,0.504,1.773,1.042 | 0.092,0.092,-0.438,0.018,1.531,1.050 |
| **066a3**2509cd925262bf53bee0a612913 | 0.039,0.092,1.095,-0.922,-0.277,-1.085 | 0.076,0.043,0.567,-0.700,-0.708,0.211 | 0.034,0.090,-0.778,0.313,-0.250,1.484 | 0.077,0.062,-0.615,0.427,0.718,0.576 |
| **069a1**04961d93e8dcd6185138dfe7678 | 0.045,0.032,1.507,-1.654,-0.522,-0.171 | 0.078,0.073,1.449,-1.459,-0.897,-1.698 | 0.044,0.036,-1.554,0.578,0.188,1.002 | 0.065,0.071,-0.739,0.337,1.935,-0.306 |
| **07149**993d0cf6d08a111835c0dc12471 | 0.043,0.064,0.901,-0.920,-1.255,-0.126 | 0.047,0.019,0.827,-0.700,-1.547,-1.205 | 0.071,0.066,-1.820,1.733,1.786,0.982 | 0.057,0.073,-1.813,1.666,1.779,0.495 |
| **071f9**e88c55e8b90bf462f37b0750776 | 0.060,0.017,1.462,-1.544,-0.884,0.981 | 0.029,0.085,0.846,-1.555,1.514,-1.511 | 0.026,0.038,-1.750,0.702,1.907,1.529 | 0.096,0.097,-1.169,0.545,1.747,1.319 |
| **07379**cb8b00a7d5175106723554f98de | 0.021,0.066,1.047,-1.074,0.570,-1.894 | 0.033,0.061,0.540,-0.190,-1.630,-0.072 | 0.048,0.062,-0.290,0.272,1.220,0.915 | 0.055,0.070,-0.184,0.094,1.290,-0.347 |
| **0774a**2880747ba5e2e4d149f8fc36170 | 0.073,0.013,1.672,-1.403,-1.533,-0.752 | 0.023,0.085,1.953,-1.806,-1.022,-0.254 | 0.074,0.030,-0.758,0.944,0.479,-1.249 | 0.023,0.076,-0.687,0.471,-0.328,0.754 |
| **07769**565c09a8ca7d55c3bd63299df57 | 0.020,0.077,0.825,-0.754,0.934,-1.579 | 0.081,0.032,0.563,-0.495,-0.429,0.009 | 0.063,0.086,-1.118,0.990,1.237,1.550 | 0.054,0.056,-0.690,0.465,-0.033,1.820 |
| **0779c**288f3d717d0f7b1986834d07713 | 0.058,0.047,1.801,-1.442,-1.600,-1.738 | 0.072,0.045,0.870,-1.160,-0.618,-0.277 | 0.053,0.086,-1.671,0.757,0.406,1.519 | 0.062,0.089,-0.361,0.082,1.928,-0.101 |
| **077cf**fb8cedf418b823aa567125d11d9 | 0.045,0.096,1.124,-1.570,0.244,-0.441 | 0.063,0.051,0.802,-0.574,-0.731,0.242 | 0.057,0.062,-1.435,1.978,1.373,-0.029 | 0.040,0.019,-1.476,1.345,0.937,1.317 |
| **07aa2**482fb1991c7e010eaf07995fb8d | 0.061,0.043,1.830,-1.661,-1.599,0.462 | 0.045,0.025,1.383,-1.659,-0.152,-0.945 | 0.077,0.090,-0.993,1.069,1.843,-1.305 | 0.085,0.038,-1.680,1.244,1.510,1.034 |
| **07c05**94517320c2cb3687909ea2a0c05 | 0.043,0.017,1.840,-1.839,-0.619,-0.654 | 0.075,0.017,1.909,-1.770,-1.853,-1.195 | 0.057,0.039,-1.941,1.956,-0.025,-0.708 | 0.039,0.043,-0.419,0.202,1.243,0.379 |
| **07d3d**4f8384b7b17c5b136a7be98edbd | 0.055,0.098,1.608,-1.124,-1.889,-1.841 | 0.097,0.054,1.662,-0.945,-1.697,-1.166 | 0.052,0.094,-1.786,1.899,1.666,0.479 | 0.087,0.067,-0.344,0.045,1.633,-0.285 |
| **08091**8fcb1886cb50d27259188e7a781 | 0.047,0.015,0.871,-0.786,-1.702,-0.087 | 0.050,0.066,0.799,-0.577,-1.226,-0.646 | 0.056,0.050,-1.035,1.092,0.075,1.746 | 0.075,0.046,-1.216,1.024,1.821,-0.197 |
| **080a3**82a47c43b9fcefe6a3a9bce61b9 | 0.078,0.026,1.957,-1.711,-1.553,0.458 | 0.066,0.080,1.456,-0.785,-1.359,-1.705 | 0.068,0.081,-1.910,1.936,-0.379,1.808 | 0.085,0.041,-1.048,0.708,1.348,0.145 |
| **0826e**e57ad8df99bed99d40aefd0c5ce | 0.026,0.062,1.892,-1.652,-0.569,-0.936 | 0.070,0.086,0.667,-1.072,-0.018,-0.733 | 0.013,0.087,-0.919,0.298,-0.580,1.197 | 0.064,0.020,-0.655,0.530,1.309,-0.317 |
| **08428**dfbdc2f4d2de65a32176305ac6a | 0.030,0.068,1.729,-1.729,-1.971,-0.430 | 0.039,0.042,1.810,-1.351,-0.864,-1.917 | 0.099,0.088,-1.720,1.974,1.172,-0.057 | 0.093,0.091,-0.876,0.344,1.114,0.981 |
| **08735**41608952b5c6c85a2954d17868a | 0.084,0.037,1.631,-1.862,-0.602,-0.502 | 0.029,0.042,0.815,-1.077,-0.501,1.018 | 0.068,0.086,-1.676,1.154,0.689,1.961 | 0.039,0.070,-1.604,1.384,1.645,0.972 |
| **088e2**0e542f0cc95217fd4af953c3354 | 0.048,0.078,1.589,-1.777,0.286,-1.427 | 0.015,0.091,0.489,-0.917,-0.987,0.890 | 0.098,0.068,-0.853,0.360,1.423,0.741 | 0.070,0.072,-0.483,-0.036,1.983,1.625 |
| **089ad**f0c07ade65d57fe1172446dc865 | 0.044,0.018,0.483,-0.486,-0.601,-0.274 | 0.077,0.072,1.781,-0.900,-1.260,-0.709 | 0.066,0.045,-1.206,1.952,-1.258,1.821 | 0.025,0.098,-0.680,0.477,-0.905,0.981 |
| **08ccf**5d571df5e50f0a36f558c12dc93 | 0.047,0.016,1.513,-1.533,-1.189,-0.445 | 0.045,0.040,0.361,-0.231,-1.737,-0.399 | 0.065,0.032,-0.960,0.762,-0.335,0.387 | 0.097,0.072,-0.346,0.161,0.638,0.184 |
| **09078**2d511dba60b31977029d77e042b | 0.073,0.086,0.290,-0.162,-0.644,-0.795 | 0.027,0.057,0.536,-0.451,-0.525,-0.584 | 0.074,0.028,-1.206,1.798,1.866,0.098 | 0.082,0.023,-0.410,0.207,1.697,0.826 |
| **091a7**fd6fee9900ddd98bc6977efa683 | 0.038,0.016,1.455,-1.728,-1.225,0.702 | 0.031,0.089,0.925,-1.031,-1.288,-0.673 | 0.014,0.052,-0.726,0.732,1.712,-1.513 | 0.040,0.060,-0.180,-0.284,1.541,1.632 |

| | | | | |
|---|---|---|---|---|
| **093e3**217e294182c85d6770960abe836 | 0.061,0.045,0.905,-0.573,-1.894,0.794 | 0.070,0.079,0.484,-1.967,0.235,1.244 | 0.049,0.089,-1.267,0.360,1.219,0.556 | 0.084,0.061,-1.024,0.660,1.867,0.187 |
| **093fd**8484f1824cb4d124dde65bb6017 | 0.022,0.086,0.523,-0.299,-0.623,-0.452 | 0.076,0.029,0.643,-1.401,-0.166,-0.542 | 0.062,0.015,-1.064,0.399,1.828,0.288 | 0.032,0.024,-0.812,0.707,0.615,0.996 |
| **09603**03486abd1c1850cfddca207dd01 | 0.084,0.022,1.808,-1.736,-1.817,0.667 | 0.014,0.043,1.502,-1.206,-1.082,-0.648 | 0.052,0.060,-1.607,1.103,1.522,1.782 | 0.052,0.059,-0.774,0.360,1.485,0.673 |
| **09681**46fa54ce4b0068537145011e08b | 0.092,0.017,1.188,-1.180,-1.822,0.390 | 0.014,0.051,0.398,-0.367,-0.023,0.212 | 0.068,0.064,-1.535,1.782,1.628,-0.326 | 0.093,0.079,-0.692,0.273,0.044,1.497 |
| **09796**678311c1ba5abf702e93eddb9a8 | 0.079,0.029,1.687,-1.414,-1.050,-1.074 | 0.074,0.097,0.586,-0.114,-0.999,-1.136 | 0.065,0.066,-0.272,0.294,-0.681,0.359 | 0.051,0.044,-0.740,0.482,1.683,0.689 |
| **097a6**5c189db784561b3e3ba3d5472d2 | 0.041,0.085,1.505,-0.792,-1.350,-0.980 | 0.094,0.027,0.097,0.056,-1.923,0.998 | 0.095,0.064,-0.670,0.199,0.534,-0.205 | 0.090,0.036,-1.366,1.095,1.779,0.605 |
| **0996f**4777cb663534b8759d5b36fe6fe | 0.025,0.043,1.284,-1.291,-0.820,-0.714 | 0.021,0.095,0.623,-1.521,-1.361,0.423 | 0.055,0.087,-0.706,0.261,0.722,1.035 | 0.041,0.079,-0.223,0.071,1.758,0.972 |
| **0a203**9abebaa7ffc6f065d74122a1e1a | 0.078,0.079,1.337,-1.127,-1.088,-1.453 | 0.041,0.049,0.762,-0.797,-0.927,-0.505 | 0.027,0.022,-1.409,1.060,0.387,1.770 | 0.068,0.023,-0.992,0.955,-0.129,1.286 |
| **0a578**e1b4ee104f6da5062070e341d59 | 0.065,0.097,1.621,-1.002,-0.944,-1.702 | 0.027,0.097,1.127,-1.026,-0.720,-1.816 | 0.055,0.047,-1.828,1.060,1.208,0.932 | 0.028,0.062,-0.076,-0.290,0.958,1.968 |
| **0a60a**4ecf140c609e1d3d8e5274595a4 | 0.013,0.085,1.734,-1.599,-0.165,-1.870 | 0.031,0.017,0.445,-0.386,-0.874,-1.103 | 0.096,0.092,-1.842,1.457,-0.210,1.263 | 0.092,0.089,-1.190,0.699,1.876,0.816 |
| **0a644**6687aa31047fa4d51200d6cefe3 | 0.071,0.062,1.342,-0.663,-0.748,-1.583 | 0.051,0.076,0.522,-0.024,-1.411,-0.082 | 0.085,0.069,-1.426,1.214,0.879,0.799 | 0.078,0.044,-0.286,-0.611,1.845,0.286 |
| **0a805**a818da00d6fe35231b4adc42b9f | 0.015,0.090,1.562,-1.611,-1.871,-0.216 | 0.069,0.088,0.542,-0.488,0.643,-0.437 | 0.036,0.065,-0.946,0.501,1.479,1.900 | 0.013,0.063,-1.226,1.046,1.951,0.387 |
| **0a866**37a1298da7fef702ed68f7d2abf | 0.098,0.046,0.568,-0.361,-1.150,0.544 | 0.042,0.060,1.389,-1.185,0.455,-1.059 | 0.044,0.052,-1.769,1.748,1.691,1.184 | 0.023,0.068,-0.634,0.305,-1.243,1.712 |
| **0a874**83af710cfe93fc7d5de0369654d | 0.054,0.061,0.094,0.144,-1.617,0.016 | 0.099,0.035,0.206,-0.455,-0.316,0.947 | 0.055,0.058,-1.554,0.836,0.404,1.567 | 0.074,0.053,-1.031,0.821,-0.394,1.686 |
| **0aab2**f102693aa016ea46b363415227f | 0.092,0.032,1.376,-0.852,-1.393,-0.287 | 0.051,0.077,0.495,-1.565,1.458,-0.927 | 0.086,0.097,-1.616,0.296,1.241,1.203 | 0.040,0.093,-1.298,0.992,0.478,1.504 |
| **0b314**cd987fac43e968deb9d97d4531d | 0.060,0.085,1.637,-1.308,-1.825,-0.737 | 0.028,0.072,0.384,-0.282,-0.710,-0.921 | 0.070,0.032,-0.267,0.021,1.483,0.564 | 0.053,0.092,-1.087,0.579,1.694,1.884 |
| **0b778**13f49731297fdbd32cf58fc57c2 | 0.049,0.050,0.396,-0.244,-1.611,-0.038 | 0.026,0.037,1.142,-1.059,1.095,-1.273 | 0.036,0.013,-1.286,1.468,0.765,1.674 | 0.051,0.080,-0.418,0.010,1.724,0.859 |
| **0ba1e**6bc3ea723e34d5b2a2204816293 | 0.082,0.029,1.325,-1.225,-1.622,0.055 | 0.015,0.078,0.528,-0.280,-0.442,-0.036 | 0.073,0.059,-0.522,0.976,-0.747,0.621 | 0.055,0.063,-0.670,0.484,1.816,-0.562 |
| **0bd33**6d0f7768d00fec7c2c6fdf329c2 | 0.054,0.084,1.002,-0.670,-1.285,-1.171 | 0.014,0.033,1.226,-1.747,-1.003,-1.532 | 0.038,0.067,-1.639,1.582,1.910,0.223 | 0.079,0.097,-0.338,-0.080,0.888,1.574 |
| **0bdec**4325330236ec718d3126f4229d8 | 0.087,0.019,0.933,-1.098,-0.431,0.667 | 0.081,0.056,0.706,-1.554,-0.545,-0.263 | 0.081,0.099,-1.980,0.716,0.718,0.265 | 0.019,0.077,-1.070,0.895,1.339,1.170 |
| **0bffb**b2ebf90780d2ec8f388bbad40c3 | 0.090,0.072,1.058,-0.821,-1.134,-0.842 | 0.037,0.053,1.636,-1.481,-1.667,0.137 | 0.078,0.060,-1.233,1.158,0.499,0.925 | 0.023,0.043,-0.429,0.224,0.962,1.268 |
| **0c00e**c51c5d4d9c403d9ac8fe6a9013c | 0.082,0.015,0.761,-1.081,-1.388,1.297 | 0.062,0.055,0.492,-0.512,0.963,-0.082 | 0.021,0.056,-1.221,1.402,-0.018,1.860 | 0.056,0.039,-0.617,0.335,1.716,1.068 |
| **0c1b6**a7db4217ab7f5e6944b880a9063 | 0.084,0.050,1.806,-1.643,-0.972,-0.404 | 0.085,0.071,0.947,-1.914,-1.708,0.166 | 0.063,0.098,-1.243,0.311,1.082,0.863 | 0.078,0.073,-1.511,1.392,1.577,-0.185 |
| **0c627**01b475ac1d7cd84d65c4a5a0e78 | 0.094,0.088,1.003,-0.622,-0.502,-1.880 | 0.066,0.044,1.001,-0.894,0.552,-1.474 | 0.096,0.096,-1.733,1.938,1.520,0.531 | 0.097,0.097,-0.666,0.156,0.761,1.671 |
| **0c62c**fc4e5a84ce728ce28b36a2060ae | 0.027,0.086,1.342,-1.245,-0.515,-0.725 | 0.034,0.028,0.395,-0.340,0.118,0.163 | 0.043,0.018,-0.534,0.637,0.913,0.356 | 0.042,0.043,-1.053,0.911,1.224,0.101 |
| **0c8bc**886b7fa84ce319b75691a4c7b99 | 0.051,0.068,1.057,-1.035,-1.437,-0.056 | 0.077,0.015,0.102,-0.125,-0.208,0.316 | 0.067,0.070,-0.448,0.057,1.091,1.859 | 0.039,0.048,0.028,-0.183,1.852,0.866 |
| **0cabf**0206859cb15d5062d8b8a22069e | 0.088,0.026,1.789,-1.872,-1.372,-0.176 | 0.056,0.019,0.370,-0.334,-0.997,-1.277 | 0.061,0.034,-0.976,0.494,1.222,0.563 | 0.066,0.085,-1.332,0.879,1.937,1.517 |
| **0cb5a**410f3db9f2efba46878c1b6a600 | 0.042,0.044,0.387,0.009,-0.675,-1.145 | 0.068,0.034,0.224,-0.076,-1.584,-1.293 | 0.065,0.080,-0.519,0.082,0.751,0.124 | 0.074,0.017,-0.097,-0.118,1.849,0.558 |
| **0cd01**45428c6f712397320c25356468a | 0.050,0.044,1.702,-1.866,-0.371,-1.023 | 0.014,0.053,0.984,-1.745,-0.346,0.308 | 0.014,0.037,-0.800,0.551,-0.249,0.700 | 0.013,0.057,-1.294,1.004,1.592,1.509 |
| **0cf28**91a6f8a0ef609b6943deaa2a644 | 0.089,0.049,1.608,-0.827,-1.718,-0.320 | 0.072,0.059,1.436,-1.719,-1.082,-1.628 | 0.089,0.045,-1.089,0.249,1.097,0.517 | 0.066,0.079,-0.464,-0.269,1.696,1.563 |
| **0cfae**1f131a0fe86a4a94d6a1b18010a | 0.076,0.018,0.650,-0.496,-1.344,0.063 | 0.018,0.063,0.661,-0.401,-1.492,-0.980 | 0.080,0.060,-1.795,1.929,1.672,1.895 | 0.022,0.089,-1.451,1.197,-0.093,1.150 |
| **0d0e4**593ed3c53e484983b7b97a4d7d3 | 0.090,0.072,1.872,-1.282,-1.903,-1.704 | 0.089,0.077,0.910,-1.364,0.801,-0.315 | 0.098,0.092,-1.907,1.049,1.394,0.961 | 0.048,0.046,-1.157,0.787,1.758,1.221 |
| **0d38b**11d86b514dc03afe2a4af6a859a | 0.080,0.067,1.776,-0.338,-1.893,-1.250 | 0.056,0.079,0.579,-0.138,-1.134,-1.842 | 0.095,0.047,-1.477,0.405,1.984,-1.422 | 0.064,0.076,-0.444,-0.116,1.216,1.782 |
| **0d476**f9d5adc48d7c449a80c7e272f9d | 0.024,0.028,1.961,-1.889,-1.882,-0.292 | 0.085,0.015,0.162,0.018,-0.814,-1.133 | 0.036,0.060,-0.612,0.129,-0.117,1.084 | 0.037,0.082,-1.381,1.194,-0.305,1.081 |
| **0d736**ce43236425c47be3197699569b8 | 0.095,0.011,1.696,-1.681,-0.962,-0.425 | 0.081,0.087,0.979,-0.761,0.237,-1.669 | 0.042,0.031,-1.047,1.448,1.673,-0.399 | 0.098,0.076,-0.504,0.166,1.216,0.616 |
| **0da30**062cde9c6ce6aa19570393fad79 | 0.060,0.023,1.623,-1.386,-1.935,0.130 | 0.035,0.073,1.013,-0.966,-1.992,0.187 | 0.049,0.033,-1.277,1.212,0.846,-0.710 | 0.061,0.095,-0.630,-0.043,1.928,1.751 |
| **0e1fb**cebb834b40aad3ff2de8a21c667 | 0.041,0.073,1.175,-1.294,0.121,-1.918 | 0.066,0.071,0.366,-0.333,0.147,-0.482 | 0.096,0.060,-0.217,0.228,-1.256,1.862 | 0.081,0.056,-0.587,0.334,1.636,1.095 |
| **0e460**5914c3d6c4abe923c370a9b0814 | 0.067,0.051,1.400,-1.709,-0.085,-0.564 | 0.078,0.026,0.409,-0.264,-1.337,-0.846 | 0.054,0.045,-0.618,0.026,1.770,1.145 | 0.039,0.037,-1.391,1.181,1.248,1.489 |
| **0e4c8**56e647b05a3e448c9f419dfee79 | 0.040,0.081,1.858,-1.615,-1.636,-1.378 | 0.076,0.036,0.412,-0.379,0.688,-0.030 | 0.097,0.092,-0.719,-0.032,1.924,1.030 | 0.017,0.011,-0.978,0.761,1.489,1.772 |
| **0e50d**6cac06a18010ee9b94a7e6dd90a | 0.091,0.065,1.921,-1.468,-1.812,-1.006 | 0.048,0.081,0.596,-0.706,-0.983,-0.554 | 0.067,0.071,-1.728,0.951,1.186,0.892 | 0.020,0.047,-1.747,1.583,0.066,1.706 |
| **0e5e1**15b4449a77f877a7150830efc9c | 0.050,0.039,0.770,-0.776,-0.532,-0.787 | 0.052,0.078,0.774,-1.567,-0.148,1.372 | 0.027,0.082,-1.829,1.170,1.755,1.592 | 0.047,0.073,-0.647,0.367,0.337,1.259 |
| **0e96b**343ec793b65374c1818756c25b9 | 0.098,0.012,1.743,-1.594,-1.879,1.189 | 0.074,0.081,1.031,-0.560,-1.163,-1.158 | 0.098,0.022,-1.835,1.236,1.537,1.477 | 0.050,0.053,-0.380,-0.025,1.880,1.219 |
| **0ef61**7da98f33dc2b1a442b3ea2341c9 | 0.047,0.092,1.956,-1.487,-1.803,0.087 | 0.018,0.030,0.453,-1.223,-0.854,-1.737 | 0.095,0.077,-1.672,0.128,1.962,-0.051 | 0.090,0.037,-1.188,0.850,1.151,1.262 |
| **0f103**79780eb7155e17bec5fe19efadb | 0.099,0.032,1.801,-1.253,-1.710,-1.245 | 0.028,0.098,0.843,-0.979,-1.311,-0.528 | 0.072,0.081,-0.999,0.645,0.148,0.302 | 0.013,0.087,-0.619,0.089,1.555,1.878 |
| **0f2de**1ff8bd1e297cd00a280f5be9073 | 0.095,0.074,1.772,-1.847,-0.473,-0.041 | 0.041,0.064,0.204,-0.372,-1.525,0.739 | 0.078,0.064,-0.968,0.896,1.367,-0.272 | 0.045,0.089,-1.223,0.978,-0.063,1.784 |
| **0f4d5**64cdd3096b128af0309ef4d9d3a | 0.093,0.053,0.780,-0.686,-0.939,-0.321 | 0.077,0.024,1.346,-0.679,-1.169,-1.122 | 0.068,0.023,-0.806,1.744,1.656,-0.379 | 0.096,0.080,-0.337,-0.063,1.646,0.692 |
| **0f6d3**6dae00b38315042d6b052f33a49 | 0.072,0.020,1.546,-1.478,-0.544,-0.828 | 0.097,0.035,0.946,-1.688,-0.571,0.378 | 0.099,0.027,-0.573,0.341,0.343,0.528 | 0.088,0.064,-1.041,0.873,1.551,-0.749 |
| **0f9c0**273016dc386161e405039071ddd | 0.094,0.064,1.731,-1.256,-1.098,-0.553 | 0.063,0.081,0.691,-1.907,-0.499,0.333 | 0.095,0.042,-1.510,0.605,-0.215,0.668 | 0.053,0.077,-1.062,0.547,1.862,1.547 |
| **0fdcf**1bfc8585c53057317250e86f46c | 0.017,0.087,1.395,-1.346,0.594,-1.963 | 0.087,0.091,1.566,-0.906,0.721,-1.960 | 0.091,0.080,-1.230,1.749,1.910,1.362 | 0.043,0.052,-0.666,0.403,0.825,1.696 |
| **0ff40**599284bf51cb34454c3e52d9779 | 0.045,0.024,0.465,-0.586,-0.811,0.252 | 0.074,0.089,0.388,-0.207,0.081,-0.293 | 0.048,0.079,-0.513,0.727,-1.098,1.086 | 0.081,0.076,-0.642,0.011,1.733,1.492 |
| **0ff4c**81db1aa26d1739be558f040bf74 | 0.085,0.014,0.893,-0.886,-1.052,0.133 | 0.019,0.090,0.519,-0.495,-1.636,-0.783 | 0.099,0.081,-1.159,0.574,0.188,1.863 | 0.092,0.063,-1.891,1.640,0.878,0.944 |
| **1002b**1628d06a8521faf240a2c021076 | 0.059,0.046,1.531,-1.554,-1.744,-0.347 | 0.055,0.025,1.840,-1.504,-1.048,-0.422 | 0.067,0.062,-1.990,1.933,1.402,1.967 | 0.058,0.049,-1.248,0.757,1.943,1.492 |
| **101ac**a73120483131e606b6fc2f62b0b | 0.097,0.084,1.851,-1.454,-1.268,-0.691 | 0.081,0.048,1.752,-1.729,-1.843,-1.111 | 0.094,0.028,-1.729,1.116,0.177,1.989 | 0.042,0.094,-0.441,0.018,1.489,1.190 |
| **105b0**7668a69401a67064383251345e2 | 0.046,0.061,1.867,-1.469,-1.681,-1.128 | 0.020,0.076,1.956,-1.397,0.169,-1.685 | 0.072,0.065,-1.843,1.912,1.854,1.336 | 0.076,0.045,-0.664,0.349,-0.117,1.937 |
| **10630**2620f38244fcf3182ae516a5de2 | 0.088,0.034,1.440,-1.197,-1.335,-0.660 | 0.063,0.068,0.967,-0.778,-1.939,-0.888 | 0.076,0.078,-1.364,1.137,-0.303,-0.050 | 0.080,0.075,-0.603,0.183,0.749,1.926 |
| **106c3**1c9b0f1cc27e2db62f434d2be10 | 0.099,0.032,1.869,-1.493,-1.187,-1.544 | 0.057,0.055,0.670,-0.695,-1.004,-0.569 | 0.070,0.049,-1.360,0.785,0.850,0.853 | 0.071,0.075,-0.161,-0.092,0.900,0.733 |
| **10823**830554c5b0bccaabbcc5f62223f | 0.073,0.029,1.786,-1.827,-0.783,-0.794 | 0.047,0.042,1.430,-1.836,-0.799,1.137 | 0.057,0.044,-1.595,1.334,0.059,1.422 | 0.020,0.061,-0.486,0.365,0.630,0.509 |
| **108c1**cbb616b14ece296ccefb3190676 | 0.068,0.032,1.580,-1.226,-1.878,-0.430 | 0.093,0.079,0.854,-1.319,-0.444,-1.033 | 0.095,0.061,-0.701,-0.036,0.982,1.350 | 0.033,0.052,-0.684,0.478,1.948,1.000 |
| **10a84**fd5091d1dc20ce3eac534a987f3 | 0.039,0.043,1.716,-1.747,-1.473,0.015 | 0.045,0.035,0.628,-1.851,-0.539,0.386 | 0.053,0.042,-1.932,0.804,1.421,0.515 | 0.084,0.011,-0.134,-0.174,1.525,1.078 |
| **10b53**b298304a104c5f80b1731232aa2 | 0.084,0.041,1.763,-1.320,-1.664,0.207 | 0.067,0.052,0.829,-0.806,-1.690,-0.241 | 0.051,0.051,-0.728,0.145,0.605,1.924 | 0.093,0.043,-1.066,0.769,0.949,0.322 |
| **10d09**bd5f860301d8af91e77780be74e | 0.048,0.023,1.450,-1.618,-1.742,0.094 | 0.094,0.047,1.358,-1.018,-1.931,-0.223 | 0.051,0.031,-0.795,0.442,0.823,1.435 | 0.082,0.044,-1.079,0.776,1.750,-0.510 |
| **10ece**43b5dd28ab04158005b71275e15 | 0.046,0.084,0.715,-0.688,-1.600,0.373 | 0.068,0.061,0.359,-0.637,1.616,-1.117 | 0.073,0.066,-1.162,0.921,0.599,0.773 | 0.042,0.079,-1.559,1.222,-0.994,1.531 |
| **10efa**ace1e9e08a2776090f546e60a0b | 0.038,0.064,1.846,-1.845,-1.684,-0.941 | 0.018,0.093,0.757,-0.806,-1.849,-0.440 | 0.088,0.047,-1.156,0.777,0.097,1.969 | 0.089,0.092,-0.976,0.332,1.404,1.790 |
| **10efd**fda217fe2f5f3746959e9e6415d | 0.073,0.012,1.224,-1.140,-0.660,-0.598 | 0.064,0.083,0.182,-0.235,-1.644,1.197 | 0.091,0.023,-0.664,0.471,1.620,1.302 | 0.098,0.027,-0.798,0.307,1.697,1.611 |
| **10fa2**51392ae4e849d950fb32f0643a2 | 0.074,0.081,1.007,-0.855,-0.954,-0.473 | 0.043,0.020,1.645,-1.467,-0.821,-0.239 | 0.066,0.082,-1.474,1.725,0.135,0.651 | 0.047,0.080,-0.302,-0.171,1.276,1.712 |
| **11189**1320f160d8f6fe46564d49cb95d | 0.043,0.060,1.649,-1.602,-1.107,-0.577 | 0.085,0.089,0.204,0.012,-1.989,0.103 | 0.019,0.044,-0.838,0.255,0.777,0.759 | 0.050,0.077,-1.369,1.120,1.780,1.266 |
| **1135b**8845799ab9d2aacbf3fab33c711 | 0.032,0.087,1.982,-1.681,-0.854,-1.339 | 0.088,0.083,0.662,-0.812,-1.969,1.149 | 0.023,0.075,-1.265,0.491,1.424,1.731 | 0.067,0.084,-0.493,-0.132,1.986,1.766 |
| **113cf**8a9c30dd27b527233dd12b43482 | 0.087,0.058,1.751,-1.750,-1.349,-0.854 | 0.044,0.080,1.318,-1.269,-0.413,-0.213 | 0.098,0.088,-1.886,1.906,1.893,1.107 | 0.042,0.091,-0.913,0.722,0.623,0.932 |

| | | | | |
|---|---|---|---|---|
| **113e4**4b0bf20e4c0c37b320a5015bd4c | 0.033,0.051,1.107,-1.168,0.961,-1.290 | 0.098,0.079,0.666,-0.452,-0.717,-0.896 | 0.046,0.016,-0.938,0.761,1.177,0.673 | 0.045,0.071,-0.300,0.120,1.477,0.663 |
| **11448**e53e952205d9def86241829dd9d | 0.012,0.060,1.535,-1.441,-0.643,-0.897 | 0.037,0.087,1.984,-1.200,0.871,-1.383 | 0.049,0.023,-1.070,1.998,-1.399,0.758 | 0.037,0.097,-0.605,0.053,1.512,1.456 |
| **11462**a75574e85a618c4abb0044f4dcc | 0.058,0.058,1.615,-1.605,-0.619,0.011 | 0.017,0.089,0.903,-0.856,0.151,-0.032 | 0.074,0.097,-1.486,1.607,-0.078,0.269 | 0.081,0.071,-1.369,0.973,0.921,0.711 |
| **115cb**2b3997839bedb80838e7dfb780a | 0.057,0.037,1.753,-1.753,-0.358,-0.679 | 0.089,0.081,0.816,-0.431,-0.933,-0.902 | 0.045,0.026,-0.341,0.443,1.148,0.454 | 0.040,0.077,-1.432,1.151,1.221,1.407 |
| **11638**f9f4e52a3bfbdc465840ea83da8 | 0.088,0.061,1.755,-1.351,-1.868,-1.031 | 0.075,0.016,0.929,-0.665,0.723,0.204 | 0.097,0.084,-1.394,1.310,1.940,1.576 | 0.016,0.060,-0.767,0.081,1.626,1.681 |
| **11709**fdeb5e670f67831b320758e3a73 | 0.076,0.020,1.616,-1.752,-0.731,0.130 | 0.065,0.048,0.963,-1.741,-0.296,-1.067 | 0.091,0.063,-1.942,0.927,1.941,0.200 | 0.091,0.097,-0.780,0.486,1.516,0.282 |
| **11885**9ad61671316d91a630773af412e | 0.034,0.076,0.768,-0.901,-0.896,-0.006 | 0.041,0.087,0.587,-0.385,0.242,-0.116 | 0.065,0.091,-0.991,1.782,-0.422,0.469 | 0.021,0.097,-1.363,0.999,0.875,1.495 |
| **11ce1**a34448f62d7c6054912d2da628c | 0.095,0.075,1.742,-1.806,-0.938,-1.039 | 0.022,0.060,1.364,-0.890,-0.631,1.192 | 0.096,0.099,-1.104,1.435,1.933,0.286 | 0.062,0.095,-0.444,-0.050,1.519,0.757 |
| **11e62**6a071b0d449f1b9c37c40a0929b | 0.058,0.015,1.995,-1.643,-1.049,-0.237 | 0.085,0.045,0.862,-1.133,-0.900,-0.329 | 0.060,0.054,-1.381,0.271,1.066,1.412 | 0.098,0.094,-0.299,-0.077,1.258,-0.096 |
| **11f57**7cce40b6bceea3426d88d29098c | 0.032,0.085,1.573,-1.456,0.191,-1.755 | 0.100,0.036,1.122,-1.855,0.161,0.048 | 0.035,0.058,-1.473,0.978,1.826,1.442 | 0.060,0.099,-0.425,-0.018,1.671,1.469 |
| **1208f**c7af1b1aa82e4407d11f7f0d652 | 0.078,0.033,1.169,-1.097,-0.789,-1.349 | 0.034,0.016,1.446,-0.889,0.209,-1.676 | 0.064,0.042,-1.218,1.916,-0.562,1.946 | 0.053,0.078,-0.871,0.577,0.640,1.433 |
| **12295**6599f65cd5ab8bf1a0ed4951739 | 0.047,0.045,1.970,-1.618,-1.945,-0.252 | 0.067,0.080,0.766,-1.342,-1.280,-0.436 | 0.086,0.042,-1.557,0.538,1.225,0.889 | 0.051,0.077,-0.610,0.367,0.122,1.483 |
| **1239f**9bcf931cc7f8b72f03de4c0051f | 0.079,0.067,1.694,-1.696,-1.420,-0.447 | 0.048,0.061,0.851,-0.733,-1.952,-0.368 | 0.093,0.041,-0.509,0.250,1.842,-0.177 | 0.089,0.050,-0.548,0.161,1.890,1.558 |
| **123a8**aad402c5a08a7079aa6f0a3a39c | 0.051,0.048,1.545,-1.582,-0.673,-0.584 | 0.018,0.038,0.828,-0.692,-1.943,-1.026 | 0.081,0.098,-1.492,1.845,1.745,-1.991 | 0.034,0.074,-1.079,0.945,0.854,0.421 |
| **1288c**a91c0f14d780a2c602b5910bd28 | 0.099,0.051,0.505,0.090,-1.391,0.052 | 0.054,0.033,0.045,-0.206,-1.184,1.383 | 0.059,0.074,-0.431,0.070,1.491,0.336 | 0.046,0.080,-0.601,0.380,0.570,0.864 |
| **128fe**90302072689dfa8c0c4eac11934 | 0.087,0.050,1.526,-1.597,-1.098,0.351 | 0.018,0.080,1.351,-1.355,-1.540,-0.947 | 0.041,0.050,-1.745,1.958,1.880,0.589 | 0.027,0.082,-1.374,1.110,-0.349,1.870 |
| **12a5d**c138e67de478290d2783c6870f1 | 0.079,0.079,1.539,-0.958,0.097,-1.175 | 0.056,0.085,0.753,-1.751,-0.340,-0.707 | 0.050,0.048,-1.753,0.301,1.779,1.232 | 0.018,0.080,-0.370,0.096,1.299,1.133 |
| **12afc**bd0bf054afaf4e3d7131305f781 | 0.044,0.017,1.865,-1.874,-0.278,-0.181 | 0.064,0.046,0.192,-0.560,-0.591,-0.955 | 0.022,0.062,-1.549,-0.018,1.012,1.908 | 0.064,0.085,-0.768,0.449,1.359,1.353 |
| **12b07**d1acb1788089fb801976a934118 | 0.084,0.039,0.672,-0.191,-1.910,-1.308 | 0.085,0.045,0.595,-0.403,-1.249,-0.646 | 0.073,0.056,-1.487,1.464,0.999,1.180 | 0.086,0.096,-1.600,1.281,0.362,1.281 |
| **12b8e**a4e7db49359da00fca20597741e | 0.098,0.013,1.894,-1.851,-1.267,1.078 | 0.075,0.012,0.911,-1.237,-1.936,-0.815 | 0.038,0.046,-0.327,0.122,-0.524,0.866 | 0.097,0.047,-0.087,-0.083,-0.061,1.445 |
| **12ba7**0c669cd2a3fddd9db03e8ecb57d | 0.055,0.025,1.785,-1.782,-0.843,-0.699 | 0.093,0.015,1.069,-0.663,-0.936,1.430 | 0.047,0.081,-1.188,0.843,1.747,1.474 | 0.055,0.083,-0.252,-0.194,1.978,0.108 |
| **12beb**121676e1c20419da156fe4f3e44 | 0.054,0.014,1.918,-1.866,-1.654,-0.383 | 0.065,0.045,1.023,-0.536,-1.379,-1.808 | 0.082,0.028,-1.241,0.510,1.773,-0.243 | 0.077,0.045,-0.720,0.384,1.083,0.445 |
| **12c5c**56c9a8575f5944d8396e0f22565 | 0.084,0.086,1.666,-1.307,-1.686,-0.258 | 0.097,0.019,1.930,-1.565,-1.477,-1.080 | 0.099,0.040,-1.869,1.902,0.216,0.595 | 0.098,0.032,-0.625,0.079,1.872,-0.270 |
| **12cb5**01929b14b3b2943d68e97d3a174 | 0.076,0.071,1.746,-1.730,-1.023,-1.663 | 0.040,0.066,0.544,-0.585,-1.576,0.009 | 0.095,0.062,-1.964,1.346,0.541,1.904 | 0.100,0.048,-0.421,0.197,1.960,-0.307 |
| **12cc9**017d7c88a987faeed2b95b7a1ad | 0.089,0.040,1.686,-1.411,-0.836,-0.641 | 0.047,0.083,0.539,-1.343,-1.828,0.425 | 0.093,0.019,-1.092,0.277,0.984,0.354 | 0.070,0.094,-0.361,0.161,-0.006,1.054 |
| **12f21**407ba6b8e0204e93761f2e7c497 | 0.079,0.062,0.505,-0.432,-0.391,-0.347 | 0.035,0.064,1.755,-1.489,-1.394,-0.046 | 0.064,0.032,-0.923,0.935,1.607,1.872 | 0.076,0.096,-1.516,0.854,1.630,1.374 |
| **12f90**f379fc435536cfa5319de0ade6f | 0.013,0.043,1.875,-1.901,-1.531,-1.168 | 0.072,0.077,1.321,-1.659,-1.346,-0.844 | 0.016,0.088,-1.008,0.566,1.364,0.181 | 0.049,0.030,-0.424,0.213,1.419,1.403 |
| **13041**1a62e251b211ecd9ac00b9e27d2 | 0.064,0.065,1.011,-0.735,-1.179,-0.316 | 0.042,0.078,0.901,-1.443,-0.975,-0.165 | 0.095,0.097,-0.884,0.447,-0.116,0.452 | 0.028,0.057,-0.273,-0.093,0.889,1.771 |
| **13655**0a57e488fc2118de4326b026e7b | 0.062,0.070,1.860,-1.784,-1.090,-0.150 | 0.040,0.016,0.449,-0.479,-1.920,-1.324 | 0.097,0.013,-1.013,0.502,0.856,-0.698 | 0.079,0.067,-1.479,1.142,1.785,0.969 |
| **13909**228554aa76a4726f3dde6f30cec | 0.064,0.020,0.396,-0.079,-1.796,-0.433 | 0.053,0.076,0.482,-0.233,-1.126,-1.096 | 0.092,0.038,-0.673,0.300,0.427,1.160 | 0.076,0.024,-0.254,-0.062,1.879,0.696 |
| **13947**6e2804dfd6c9204c9cbc896cdd8 | 0.081,0.046,1.621,-1.522,-1.751,-0.299 | 0.027,0.077,0.443,-0.660,-0.779,-0.161 | 0.073,0.034,-1.557,0.893,1.826,1.212 | 0.093,0.050,-1.177,0.868,1.426,1.005 |
| **13cbd**3112a46c6fa1f7086bb84cf7a28 | 0.030,0.042,0.479,-0.475,-0.048,-1.028 | 0.088,0.082,0.730,-0.358,-0.841,0.358 | 0.075,0.087,-1.180,1.949,-0.311,0.860 | 0.047,0.080,-1.832,1.572,-0.946,1.346 |
| **14012**b08c4bf0058c0e11b988de43153 | 0.013,0.075,1.608,-1.628,-1.605,-0.844 | 0.043,0.025,0.883,-1.899,-1.318,-1.702 | 0.096,0.092,-1.342,0.411,1.966,-0.488 | 0.078,0.063,-0.442,0.144,1.781,0.735 |
| **14091**72433ef332dbdf19f901cc4ff64 | 0.055,0.069,0.956,-0.778,-1.661,-0.003 | 0.026,0.074,1.233,-1.138,-1.596,-0.352 | 0.060,0.088,-1.916,1.244,1.912,1.861 | 0.012,0.091,-0.319,-0.128,1.753,1.701 |
| **1412d**6db93a7d3c9798d5942f56b1b1b | 0.096,0.021,1.567,-1.325,-1.847,-0.158 | 0.015,0.083,0.516,-0.466,-0.747,-0.345 | 0.033,0.097,-1.564,1.092,-0.168,1.565 | 0.016,0.076,-1.229,1.147,1.830,0.366 |
| **14217**dd2c9a6798894b629413227bc8f | 0.055,0.063,1.678,-1.691,-1.368,-0.986 | 0.093,0.032,0.858,-0.602,-0.938,-0.146 | 0.020,0.061,-1.699,1.930,1.604,0.312 | 0.087,0.075,-1.260,1.002,-0.373,1.966 |
| **143a7**0423b61420926e26e8a287ac79d | 0.085,0.041,1.828,-1.646,-1.701,0.692 | 0.062,0.067,1.276,-1.346,-1.985,-0.176 | 0.064,0.090,-1.101,0.768,1.497,1.098 | 0.041,0.068,-0.443,0.195,1.749,1.002 |
| **14424**12bdd5684d8922fb5e97531f2d9 | 0.068,0.035,1.723,-1.247,-1.309,-1.075 | 0.051,0.091,0.796,-1.526,-0.950,-0.480 | 0.072,0.097,-1.156,0.654,-0.060,-0.495 | 0.043,0.059,-0.249,0.061,0.804,0.976 |
| **144c9**0ec7e91156310c718f4eb0d4bdd | 0.053,0.056,1.698,-1.680,-1.013,-1.190 | 0.097,0.013,1.274,-0.921,-0.580,1.381 | 0.098,0.099,-0.816,0.380,1.931,1.329 | 0.039,0.020,-1.262,1.127,1.890,0.027 |
| **147f1**96b8ec9d3ec09823fc56ddb5b38 | 0.040,0.019,0.983,-0.498,-1.535,0.632 | 0.086,0.049,0.272,0.316,-1.487,-0.697 | 0.015,0.024,-1.937,0.415,1.645,0.474 | 0.060,0.026,-0.629,0.340,1.521,0.497 |
| **1481a**4a7d402c83bff2d33640475c14e | 0.012,0.082,1.707,-1.132,-0.964,-1.912 | 0.024,0.091,0.981,-1.503,0.402,0.761 | 0.088,0.030,-1.001,-0.080,1.872,1.740 | 0.044,0.051,-1.069,0.339,1.554,1.538 |
| **14890**3d40c7836ce4522c57b529f40e2 | 0.037,0.064,1.802,-1.196,-0.687,-1.248 | 0.084,0.081,0.528,-0.838,-1.300,-0.049 | 0.069,0.061,-1.933,0.178,1.370,1.916 | 0.098,0.037,-0.600,0.079,1.860,1.951 |
| **14bd3**99ca0f2e4b735a0da1f650e4bbe | 0.049,0.024,1.538,-1.384,-0.172,-1.328 | 0.021,0.052,0.951,-1.493,1.851,-1.872 | 0.090,0.040,-0.689,0.012,1.757,0.497 | 0.098,0.081,-1.007,0.822,0.545,0.190 |
| **14c14**4b94220f0f6551b615b047fdc33 | 0.044,0.061,1.617,-1.430,-1.036,-0.757 | 0.081,0.043,1.173,-0.886,0.282,-1.636 | 0.095,0.057,-1.040,0.825,1.950,0.166 | 0.075,0.040,-0.619,0.283,1.260,0.203 |
| **14eca**93cfe8ee43f05a5c1e2b06d84fa | 0.074,0.033,1.046,-0.521,-1.589,-0.571 | 0.095,0.045,0.694,-0.791,-0.152,-0.288 | 0.019,0.068,-1.779,1.254,0.206,1.451 | 0.087,0.017,-0.088,-0.021,0.719,-1.637 |
| **14fd3**f4e0bc15f4baf8bb2b551b1c7df | 0.030,0.081,1.419,-1.471,0.264,-1.001 | 0.018,0.014,0.716,-0.525,1.774,-0.934 | 0.065,0.031,-1.259,1.918,1.379,-0.047 | 0.075,0.041,-0.679,0.236,0.782,1.720 |
| **1506b**a19c0d0782e1dffa547742c39e3 | 0.034,0.024,1.397,-1.703,-1.064,-0.147 | 0.040,0.061,0.109,-0.467,-1.035,0.805 | 0.065,0.062,-0.456,-0.076,1.726,1.042 | 0.062,0.071,-0.654,0.446,0.928,1.056 |
| **15318**b1acaeed934c40271b6908cd2bc | 0.084,0.032,1.045,-1.010,-0.037,-0.677 | 0.019,0.066,1.725,-1.075,-1.864,-0.695 | 0.098,0.051,-0.864,1.031,1.935,0.287 | 0.054,0.095,-0.571,0.220,1.047,0.856 |
| **1531c**21e0937183b25160dcdd88d4a6e | 0.098,0.041,1.630,-1.450,-1.928,-0.459 | 0.074,0.037,1.402,-1.622,-0.495,1.301 | 0.080,0.059,-1.515,1.535,0.876,0.627 | 0.084,0.043,-0.724,0.188,1.918,1.696 |
| **15416**b7b813f4e895be92f699f41f325 | 0.013,0.066,1.266,-1.252,-0.689,-1.738 | 0.084,0.081,0.630,-0.774,0.134,-0.780 | 0.072,0.013,-1.721,1.077,1.893,1.853 | 0.071,0.099,-1.265,1.055,1.502,0.422 |
| **15799**15f169e27c9a9940055c25cfd12 | 0.047,0.068,1.662,-0.904,-1.892,-0.673 | 0.073,0.076,1.045,-1.028,-1.058,-0.964 | 0.038,0.082,-1.872,0.704,1.861,0.512 | 0.011,0.061,-0.522,0.225,0.593,1.347 |
| **157e5**63512026967d60d30e00de04c3f | 0.048,0.058,0.141,0.063,-0.927,-0.446 | 0.035,0.057,0.741,-0.709,-0.915,-1.753 | 0.063,0.059,-1.476,1.061,1.711,0.242 | 0.067,0.046,-0.486,0.344,1.009,0.124 |
| **15892**e894b627e7a9909417c542c4697 | 0.094,0.063,1.761,-1.486,-1.657,0.028 | 0.014,0.050,1.279,-1.158,-0.741,-0.139 | 0.070,0.077,-1.163,1.239,0.058,1.889 | 0.066,0.013,-0.261,-0.069,1.682,1.826 |
| **15d2a**7b346a037569915fd13868ab08b | 0.048,0.086,1.651,-1.384,-0.960,-1.353 | 0.042,0.018,0.771,-0.806,-0.380,-1.232 | 0.096,0.073,-1.689,1.109,1.941,1.091 | 0.062,0.088,-1.409,0.992,0.163,1.922 |
| **15db4**9df2013009341bb3824dd09b6f9 | 0.064,0.017,0.886,-0.598,-1.657,1.108 | 0.073,0.071,1.025,-1.646,-0.842,-1.176 | 0.036,0.048,-0.661,0.183,1.756,0.307 | 0.088,0.079,-0.855,0.471,1.121,0.927 |
| **15e85**4d7bc538a1c3241596abf6a14b6 | 0.047,0.049,1.712,-1.209,-1.763,-0.432 | 0.061,0.067,1.111,-1.582,-1.504,-0.599 | 0.041,0.020,-1.783,0.745,1.959,0.102 | 0.036,0.053,-1.092,0.782,1.901,0.723 |
| **15f7f**8385ba456f0a1a04d4a77fc7c93 | 0.090,0.080,0.949,-0.634,-0.968,0.099 | 0.012,0.015,0.278,-0.855,1.604,1.025 | 0.069,0.082,-1.698,0.659,1.910,1.079 | 0.093,0.047,-1.086,0.822,1.722,-0.492 |
| **15f8b**72cd7d1104c4b17cf50cce8f633 | 0.075,0.016,1.712,-1.580,-1.445,1.148 | 0.021,0.083,1.468,-0.670,-0.464,-0.809 | 0.052,0.093,-0.874,1.398,1.117,-0.776 | 0.095,0.041,-0.087,-1.116,0.943,1.704 |
| **15fae**7171f05a42a65421dfbda1937e8 | 0.063,0.011,1.558,-1.907,-1.162,1.325 | 0.097,0.053,0.976,-0.792,-0.869,-0.002 | 0.095,0.070,-0.648,0.650,1.574,0.336 | 0.053,0.062,-0.345,0.121,0.394,1.493 |
| **1604d**78ee47eb40081422413ab4dc695 | 0.091,0.055,1.909,-1.663,-0.754,-0.327 | 0.010,0.047,1.461,-1.768,-1.394,-0.361 | 0.098,0.075,-1.900,1.139,1.209,1.902 | 0.079,0.051,-1.424,1.197,1.688,-0.420 |
| **16287**295309d14b403455854b61777ee | 0.037,0.015,1.462,-1.492,-0.066,-1.308 | 0.013,0.084,0.518,-1.013,-0.933,-0.530 | 0.073,0.096,-1.106,0.150,-0.430,1.810 | 0.086,0.076,-0.876,0.574,0.588,1.190 |
| **16443**2ddef4323518de6b039cabf4ec5 | 0.038,0.096,1.803,-1.268,0.363,-1.189 | 0.092,0.062,0.900,-1.511,-1.027,0.059 | 0.037,0.061,-1.099,0.440,1.870,-0.576 | 0.061,0.064,-0.634,-0.068,1.723,1.955 |
| **16454**bb0edd715c7a7289ecccdf07adc | 0.016,0.028,1.375,-1.323,-1.276,-0.235 | 0.063,0.047,0.521,-0.627,-0.220,-1.155 | 0.048,0.047,-1.052,0.583,-0.119,0.351 | 0.068,0.069,-0.066,-0.389,1.553,0.803 |
| **1651c**86764e78c26bfa9b16303b072a0 | 0.055,0.023,1.662,-1.588,-0.576,-1.043 | 0.038,0.097,0.498,-0.650,1.316,0.328 | 0.044,0.094,-0.634,0.699,1.385,0.144 | 0.035,0.069,-1.058,0.441,1.457,1.761 |
| **1670d**b867bdebc34f2fbd0229a5a4fe4 | 0.089,0.036,0.441,-0.154,-0.807,-0.169 | 0.028,0.085,0.404,-0.568,-0.463,-0.044 | 0.067,0.096,-0.300,-0.041,1.552,0.015 | 0.027,0.033,-1.070,0.976,0.871,0.672 |
| **16c09**e4fd3cc68ccce349b95c1617f90 | 0.093,0.080,1.899,-1.801,-1.880,-0.387 | 0.093,0.058,0.716,-0.401,-1.220,-1.518 | 0.054,0.045,-1.683,1.624,1.846,-0.400 | 0.025,0.039,-1.735,1.611,0.832,1.493 |
| **16d40**f7eb4617ec568ecad2002319780 | 0.040,0.020,1.549,-1.626,-1.677,0.114 | 0.082,0.091,0.984,-1.160,-0.184,-1.868 | 0.069,0.056,-1.879,1.029,0.624,1.215 | 0.040,0.058,-1.356,1.301,1.936,-0.247 |
| **17098**b6705d49cc94d5ab40ee88712c2 | 0.092,0.048,1.500,-1.380,-1.634,1.296 | 0.070,0.090,0.494,-1.549,-0.533,1.304 | 0.061,0.050,-0.309,-0.194,0.007,1.861 | 0.045,0.026,-0.348,-0.018,1.821,0.701 |

| | | | | |
|---|---|---|---|---|
| **1716c**4e2ef28b65ccc9da7ec98f721c4 | 0.096,0.016,1.361,-1.114,-1.471,0.106 | 0.028,0.051,0.588,-0.611,-1.433,-0.647 | 0.035,0.078,-1.583,1.325,0.877,0.349 | 0.048,0.099,-0.479,0.101,1.094,0.991 |
| **17266**fe877ec3990588af1c910ecacd3 | 0.095,0.021,1.612,-0.958,-1.953,0.143 | 0.016,0.047,0.297,-0.345,-0.274,-0.394 | 0.058,0.092,-1.069,0.315,1.535,0.350 | 0.078,0.024,-0.286,-0.345,1.588,1.778 |
| **17378**0552eb11a3adaf3c676150f0306 | 0.088,0.023,0.882,-0.776,-1.439,-0.280 | 0.016,0.049,0.625,-0.275,-0.718,-1.475 | 0.067,0.017,-0.628,0.981,1.143,1.633 | 0.062,0.026,-0.110,-0.017,0.878,1.269 |
| **173a3**356ddd37325fe40ae1e8a9b68de | 0.093,0.022,1.892,-1.874,-1.588,-0.655 | 0.065,0.035,0.189,-0.162,-1.943,1.211 | 0.098,0.031,-0.640,0.448,0.238,0.824 | 0.044,0.046,-0.492,0.458,1.842,-0.495 |
| **173ee**edfa4bb39725414db30451f8c18 | 0.062,0.043,1.590,-1.503,-1.906,-0.099 | 0.073,0.034,1.423,-1.033,-0.397,0.038 | 0.067,0.013,-1.034,0.978,1.352,0.984 | 0.024,0.043,-0.141,-0.348,1.172,1.959 |
| **1746f**a273c5d7b0760fd4ca0921fa30d | 0.024,0.017,1.522,-1.609,-0.352,-0.850 | 0.025,0.032,0.311,-1.365,0.353,0.349 | 0.015,0.083,-0.863,0.172,1.095,0.811 | 0.074,0.052,-0.637,0.645,-0.355,0.368 |
| **174d9**e2fafb086f0a2d79735ea99926e | 0.073,0.087,1.620,-1.411,-0.666,-1.787 | 0.037,0.032,0.675,-0.936,1.923,-1.649 | 0.098,0.040,-1.256,0.929,1.923,0.374 | 0.072,0.037,-0.588,0.303,1.242,1.584 |
| **174fa**d231821dece229445417d83f23d | 0.048,0.023,0.752,-0.677,-1.042,-0.237 | 0.042,0.070,1.002,-1.088,-0.207,0.283 | 0.050,0.057,-1.734,1.706,1.423,1.055 | 0.055,0.042,-1.205,1.072,1.210,-0.242 |
| **177fd**f1dff05e16c05f34dea599ad02f | 0.075,0.022,0.473,-0.598,-0.438,1.250 | 0.098,0.087,0.051,-0.168,-0.765,1.099 | 0.033,0.036,-0.189,-0.030,1.449,1.724 | 0.038,0.024,-0.471,0.462,-1.217,1.226 |
| **17944**521f6e48fc6ea957bf23ac6937b | 0.087,0.048,1.722,-1.464,-1.909,-0.101 | 0.085,0.041,1.947,-1.829,0.199,-1.060 | 0.098,0.067,-1.742,1.816,1.894,1.045 | 0.049,0.098,-0.324,-0.189,1.845,1.351 |
| **17e88**c1c1ad0707e35b360bdce89f765 | 0.053,0.046,1.341,-1.292,0.204,-0.714 | 0.039,0.083,0.844,-0.634,0.562,-1.319 | 0.070,0.020,-1.787,1.885,1.278,-0.638 | 0.071,0.029,-0.899,0.559,1.041,1.790 |
| **17f5f**57695770abfe4ee8f6cea6bdd98 | 0.078,0.028,1.571,-1.304,-1.848,0.000 | 0.013,0.076,0.369,-0.433,-1.072,0.001 | 0.080,0.097,-1.037,0.213,1.700,0.495 | 0.072,0.071,-1.234,0.666,1.111,1.190 |
| **17fef**6db9e4958421da08499292b188c | 0.055,0.046,0.978,-0.825,-1.859,-0.410 | 0.087,0.052,0.237,-0.173,-0.273,-0.772 | 0.091,0.096,-1.390,0.592,-0.447,1.906 | 0.060,0.059,-0.455,0.138,1.802,0.978 |
| **180cf**04e538813973f290c1acf9ec9f5 | 0.085,0.018,1.828,-1.463,-1.872,0.152 | 0.013,0.063,0.765,-1.592,-1.940,-0.307 | 0.078,0.096,-1.868,0.553,1.778,1.270 | 0.050,0.043,-0.863,0.635,1.910,1.415 |
| **180ec**ab5c694332ce93549d5153a083d | 0.085,0.017,1.118,-1.175,-1.285,0.094 | 0.067,0.028,0.227,-0.181,-1.545,1.948 | 0.051,0.067,-0.252,0.364,-0.535,0.561 | 0.090,0.065,-0.411,0.107,1.126,1.573 |
| **1831d**caacf09fb1757002210cd48bdc2 | 0.073,0.035,1.051,-0.866,-1.762,-0.030 | 0.099,0.076,0.526,-0.432,-1.481,1.002 | 0.057,0.031,-0.535,0.294,1.616,0.927 | 0.049,0.088,-1.791,1.327,1.645,1.779 |
| **183de**11fc50bcc1e7f023001340491c8 | 0.042,0.070,1.223,-1.141,-0.400,-1.491 | 0.060,0.075,0.794,-0.438,-1.302,-0.474 | 0.093,0.050,-0.650,0.949,1.260,0.394 | 0.034,0.064,-0.501,0.411,0.969,0.333 |
| **188ac**fda5a4e7f155eda7080cca94754 | 0.081,0.059,1.459,-1.699,-0.601,-0.442 | 0.075,0.017,0.477,-0.575,-0.612,-0.335 | 0.022,0.081,-0.538,0.505,1.173,0.460 | 0.082,0.034,-1.437,1.276,1.139,1.284 |
| **18981**23a087fce4e763f181734204b2a | 0.039,0.087,1.617,-1.878,0.848,-0.812 | 0.023,0.016,1.624,-1.091,-0.355,1.452 | 0.045,0.052,-1.294,1.826,1.666,1.559 | 0.058,0.070,-0.615,0.048,1.870,1.329 |
| **18dbe**628764db7f7e59e9490f733f173 | 0.011,0.095,1.317,-1.543,-1.633,-0.921 | 0.018,0.081,0.938,-0.823,-1.970,-0.950 | 0.012,0.065,-1.965,1.940,1.927,0.918 | 0.015,0.080,-0.910,0.648,1.462,1.392 |
| **18e63**d4690c190f8ae14709aee6d0df7 | 0.039,0.052,0.876,-0.927,-0.809,-0.230 | 0.091,0.027,0.863,-0.420,-0.409,-1.534 | 0.040,0.090,-0.862,1.949,0.576,-1.818 | 0.082,0.086,-1.833,1.234,0.259,1.787 |
| **190b6**e9886d493a5a8c080ebcd01950d | 0.038,0.049,1.733,-1.643,-1.372,-0.068 | 0.019,0.086,0.819,-1.207,-1.169,-1.850 | 0.090,0.057,-1.335,0.248,1.935,0.261 | 0.070,0.089,-0.537,0.410,0.262,0.551 |
| **19326**bba2fbbb6dd506d8da41284e31d | 0.092,0.035,1.380,-1.365,-1.764,0.673 | 0.092,0.062,0.579,-0.473,-0.918,0.321 | 0.100,0.080,-1.328,1.157,1.755,-0.014 | 0.085,0.020,-1.074,0.919,1.628,-0.533 |
| **1934a**64a844ac3e8219d9ad44cd250c8 | 0.058,0.033,1.730,-1.654,-1.716,-1.955 | 0.065,0.012,0.453,-0.477,-1.770,-1.518 | 0.023,0.084,-0.833,0.557,-0.248,0.577 | 0.062,0.100,-0.864,0.641,1.530,0.813 |
| **19a0f**ed0ac7d02165a52ceba0f5cd987 | 0.029,0.024,1.455,-1.512,-1.175,-0.491 | 0.088,0.056,0.433,-0.408,-1.001,0.552 | 0.053,0.081,-0.923,0.548,-0.196,1.573 | 0.078,0.091,-1.519,1.239,0.845,1.135 |
| **19acc**fcaa8506c78afb9b093006cb2b8 | 0.066,0.088,0.575,-0.567,-0.727,0.348 | 0.028,0.084,1.103,-1.289,-1.240,-1.371 | 0.067,0.079,-1.486,1.156,0.636,0.337 | 0.041,0.076,-1.048,0.652,1.020,1.748 |
| **19b1a**b98ff227707ce54f6c527bbed41 | 0.094,0.034,1.667,-1.466,-1.141,1.215 | 0.066,0.080,0.774,-1.136,-0.008,0.128 | 0.089,0.089,-1.016,0.678,0.733,-0.321 | 0.081,0.071,-0.905,0.402,1.498,1.292 |
| **19b26**805458b2b48488b9fe767ac0684 | 0.058,0.066,1.275,-1.075,-0.405,-0.932 | 0.064,0.031,1.854,-0.851,-1.054,0.281 | 0.083,0.062,-1.041,1.831,1.058,-0.404 | 0.035,0.064,-0.461,0.160,-0.286,1.073 |
| **19cca**9cf3225b50f8ccfd58b3cfac685 | 0.016,0.064,1.435,-1.348,-1.755,-1.893 | 0.085,0.025,0.952,-0.584,-1.549,0.060 | 0.086,0.064,-1.429,1.913,-0.542,1.188 | 0.070,0.056,-1.356,0.981,1.221,1.635 |
| **19d78**03d2bf4bc45dcfb7ce141cd4037 | 0.082,0.036,1.627,-1.248,-1.754,0.318 | 0.025,0.038,0.938,-0.578,0.676,-1.742 | 0.086,0.064,-1.621,1.376,1.457,1.382 | 0.037,0.044,-0.790,0.306,1.701,1.119 |
| **1a2c6**772321f15eba4c09067a2b73179 | 0.092,0.054,0.018,-0.005,-1.056,0.786 | 0.060,0.084,0.265,-0.266,-1.282,1.333 | 0.068,0.059,-1.119,1.771,0.001,0.788 | 0.016,0.089,-1.537,1.182,-0.550,1.875 |
| **1a3f6**cfe51fe3000e3d40448f2deb88d | 0.038,0.046,1.611,-1.445,-0.426,-1.751 | 0.066,0.021,1.001,-0.853,0.550,-0.428 | 0.093,0.091,-1.298,1.595,-0.416,1.928 | 0.053,0.024,-1.489,1.349,0.646,-0.072 |
| **1a89d**f6c0cfd82a52f0b8b2b0c413ff4 | 0.073,0.076,1.642,-1.912,0.611,-1.599 | 0.042,0.071,0.772,-0.904,0.444,-0.530 | 0.068,0.062,-1.268,0.841,1.386,1.459 | 0.062,0.073,-0.948,0.892,1.668,-0.578 |
| **1a92d**ee6a4bf637c0e7841ecf52fc725 | 0.090,0.062,1.436,-0.724,-1.936,-1.444 | 0.027,0.046,1.904,-0.993,-1.443,1.133 | 0.093,0.061,-1.128,1.100,1.806,1.087 | 0.086,0.023,-1.176,0.289,1.958,0.613 |
| **1ab9e**88d034b9ec87c2f64c8b4459ed4 | 0.018,0.064,1.838,-1.586,-0.424,-1.125 | 0.063,0.019,0.892,-1.436,0.157,-0.965 | 0.033,0.040,-1.025,0.414,1.624,0.951 | 0.012,0.062,-1.495,1.102,0.713,1.416 |
| **1b095**7802d8eee0384d05f3e177787f5 | 0.053,0.057,1.709,-1.884,-1.078,-1.050 | 0.013,0.063,0.479,-0.253,-1.451,-0.340 | 0.028,0.060,-1.023,1.921,1.107,0.358 | 0.053,0.065,-1.417,1.226,0.101,1.718 |
| **1b147**63c69a1c634dd1129fbd0aa42ed | 0.060,0.021,0.531,-0.449,-1.337,0.430 | 0.019,0.044,0.984,-1.191,-1.712,-0.068 | 0.048,0.036,-1.290,1.320,1.135,0.104 | 0.030,0.084,-1.893,1.903,1.900,-0.401 |
| **1b25a**26cc690f5d2c93545c4ad97d3e0 | 0.036,0.099,1.605,-1.554,-0.859,-1.575 | 0.041,0.090,1.379,-1.172,-0.951,-1.283 | 0.096,0.070,-1.958,1.906,1.512,0.301 | 0.085,0.032,-0.307,0.047,1.471,0.858 |
| **1b3f6**4bbd7e26368539cb0cbbd5d2fd2 | 0.040,0.083,1.666,-1.141,-1.880,0.023 | 0.059,0.098,0.393,-0.522,-0.113,-1.458 | 0.078,0.074,-1.800,-0.048,1.495,1.059 | 0.012,0.078,-0.812,0.586,1.665,1.136 |
| **1bb7a**a8b8b60e49603668cf7c38ccc77 | 0.029,0.013,1.358,-1.486,-1.648,-0.220 | 0.064,0.019,0.076,-0.160,0.124,-1.627 | 0.024,0.065,-1.183,0.399,-0.240,1.930 | 0.071,0.063,-0.157,-0.274,1.989,1.335 |
| **1bbee**f7f15b80cb8f9d07204671d9b87 | 0.047,0.055,1.938,-1.722,-1.652,-0.510 | 0.043,0.052,1.760,-1.948,0.316,-1.686 | 0.064,0.062,-1.273,0.867,-0.252,1.549 | 0.040,0.055,-0.179,-0.141,1.224,1.122 |
| **1bea2**f3d9b5b67015faf220824122d0d | 0.067,0.025,1.603,-1.374,-1.894,0.766 | 0.054,0.015,0.954,-1.525,-0.072,-1.799 | 0.012,0.073,-1.335,0.640,0.469,1.212 | 0.022,0.067,-0.100,-0.117,1.149,1.094 |
| **1bedd**49ec94dc33ff4827d19a9d61bec | 0.092,0.044,1.772,-1.202,-1.580,-0.230 | 0.060,0.059,0.282,-0.072,-1.616,-0.949 | 0.051,0.085,-1.160,0.155,1.840,1.492 | 0.022,0.065,-0.387,0.216,0.731,1.316 |
| **1c106**b1ad98aafda8f2af547d457f38f | 0.055,0.054,1.702,-1.680,-1.155,-0.626 | 0.019,0.059,1.168,-1.906,-1.732,-0.861 | 0.076,0.034,-1.924,0.804,1.186,1.937 | 0.090,0.084,-0.505,-0.084,1.995,1.612 |
| **1c190**c09dc96a1a720a280a9e42275f3 | 0.049,0.011,1.279,-1.234,-1.692,0.129 | 0.026,0.098,0.690,-0.437,0.890,-0.526 | 0.084,0.083,-1.186,1.725,1.328,1.001 | 0.026,0.022,-1.365,1.241,0.262,1.510 |
| **1c2eb**19b151e108540d248c85f7a911a | 0.026,0.046,0.940,-0.864,-1.910,0.072 | 0.044,0.060,0.309,-0.594,0.952,-1.140 | 0.062,0.048,-1.742,1.041,-0.296,0.826 | 0.078,0.074,-1.494,1.306,-0.458,1.301 |
| **1c37e**781f596b35575764ef5417ff0e1 | 0.095,0.032,1.439,-0.836,-1.865,0.839 | 0.070,0.079,1.134,-1.594,-1.406,-1.633 | 0.068,0.098,-0.956,0.058,0.084,1.571 | 0.061,0.043,-0.384,0.193,1.788,0.202 |
| **1c440**e406d9cba9c04de48f257951a19 | 0.044,0.019,1.715,-1.903,-1.957,-0.095 | 0.038,0.022,0.230,-0.202,-1.534,-0.255 | 0.065,0.054,-0.562,0.315,0.017,0.624 | 0.092,0.065,-0.430,0.076,0.622,1.698 |
| **1c628**7bb5b5e31695b66b9346842f785 | 0.044,0.048,1.773,-1.503,-1.518,-1.050 | 0.068,0.070,1.275,-1.450,-1.285,-1.139 | 0.020,0.086,-1.239,0.928,1.029,-0.001 | 0.080,0.065,-1.220,0.797,0.988,1.613 |
| **1ca38**ed8004e29fe15b346fac44b69d1 | 0.017,0.078,1.449,-1.084,-0.173,-1.838 | 0.023,0.050,1.697,-1.647,1.605,-0.158 | 0.013,0.041,-1.743,1.922,1.353,1.158 | 0.013,0.052,-1.596,1.268,1.211,1.360 |
| **1cbf7**2257bae91ee998599cb388a5d44 | 0.049,0.058,1.799,-1.621,-1.080,-0.701 | 0.095,0.060,0.630,-0.474,-1.304,0.421 | 0.063,0.066,-0.595,0.451,1.931,-0.162 | 0.098,0.078,-1.334,0.835,1.457,1.768 |
| **1cce5**e07f035896cebb5afe7fde4891e | 0.045,0.053,1.465,-1.350,-0.950,-1.694 | 0.068,0.091,1.418,-1.478,-0.349,-0.759 | 0.036,0.049,-1.307,0.926,1.473,1.994 | 0.098,0.074,-0.286,-0.152,1.855,0.521 |
| **1ceb9**e9c8e9ef98ca1529546348bbff4 | 0.052,0.041,1.703,-1.531,-1.950,-0.053 | 0.089,0.032,0.566,-1.824,-1.748,1.633 | 0.097,0.037,-1.254,0.311,0.853,0.583 | 0.059,0.070,-0.716,0.551,1.294,0.663 |
| **1cf6a**274e0c40f95f51dbf73a36d8db3 | 0.088,0.039,1.974,-1.576,-1.916,-0.669 | 0.051,0.081,0.588,-0.361,-1.462,-0.584 | 0.071,0.090,-0.831,0.191,1.313,1.086 | 0.061,0.081,-1.668,1.168,1.907,1.123 |
| **1d29e**a5137115cb7e59b080742b887b1 | 0.077,0.023,1.689,-1.609,-1.907,0.212 | 0.058,0.081,1.314,-1.149,-0.556,-0.603 | 0.067,0.045,-1.642,1.587,1.762,0.697 | 0.096,0.079,-1.055,0.804,1.275,0.357 |
| **1d56c**7de71028f8af2cd4565928bc46c | 0.076,0.014,1.748,-1.704,-0.699,-0.714 | 0.047,0.097,0.597,-0.747,-0.745,-0.996 | 0.072,0.062,-1.847,0.876,0.891,0.620 | 0.088,0.092,-0.817,0.361,1.867,0.888 |
| **1d9f0**3d49977a57a696260c7f3bd8b66 | 0.096,0.037,1.179,-1.160,-0.818,0.696 | 0.024,0.087,0.941,-1.853,0.885,0.148 | 0.037,0.013,-1.729,1.328,-0.267,0.904 | 0.046,0.012,-1.240,0.919,1.900,1.585 |
| **1dc43**ab9e9f0136db8310a819a4a3b96 | 0.013,0.090,1.059,-1.016,-1.747,-0.346 | 0.017,0.087,0.080,-0.394,-0.336,1.304 | 0.029,0.066,-0.564,0.225,1.779,1.874 | 0.078,0.094,-1.138,0.579,0.751,1.363 |
| **1dd83**d72b1fa8fd514fff90fa9e19c6f | 0.086,0.019,0.890,-1.058,-1.142,-0.162 | 0.093,0.074,0.760,-0.397,-0.592,-0.548 | 0.033,0.077,-0.587,0.836,1.542,-0.296 | 0.084,0.036,-0.312,0.045,1.420,0.835 |
| **1dfe9**855fd28ed43ac2b65c1b47a2faf | 0.097,0.011,1.031,-0.567,-1.909,1.108 | 0.016,0.080,1.219,-0.873,0.434,-1.905 | 0.044,0.095,-1.955,1.486,1.615,1.877 | 0.091,0.092,-0.845,0.193,0.618,1.889 |
| **1e01e**2e1d74a8532fba5a91ad526c0a8 | 0.078,0.030,1.552,-1.295,-1.247,-0.228 | 0.059,0.022,0.669,-0.722,-1.344,-1.547 | 0.028,0.041,-1.102,0.668,0.705,1.574 | 0.054,0.083,-0.423,0.079,1.868,1.072 |
| **1e0d9**00067c126bf372a02a9f846d0df | 0.094,0.055,1.631,-1.336,-1.515,-1.180 | 0.019,0.082,1.929,-1.677,-0.618,-1.376 | 0.078,0.087,-1.962,1.680,1.057,1.730 | 0.012,0.089,-1.731,1.527,0.692,0.708 |
| **1e3c3**fa14be95b08a7e42871b3916fb3 | 0.086,0.028,1.901,-1.777,-0.188,-1.104 | 0.031,0.064,0.760,-0.578,-1.281,-0.265 | 0.013,0.090,-0.774,0.545,1.822,0.358 | 0.051,0.095,-0.658,0.313,0.565,0.445 |
| **1e5bd**0ac97d9273d6afe980b46c3ea7a | 0.080,0.056,0.458,-0.268,-0.808,-0.021 | 0.098,0.017,0.816,-0.447,-1.262,-0.900 | 0.034,0.090,-0.365,-0.100,1.757,0.653 | 0.056,0.038,-1.045,0.561,1.356,1.845 |
| **1e5e2**fabaf5dd9e1e4c52de5f2222154 | 0.021,0.019,1.205,-1.363,-1.532,-0.428 | 0.016,0.086,0.432,-0.274,-1.209,-0.096 | 0.023,0.069,-1.399,1.858,0.770,1.127 | 0.097,0.026,-1.774,1.424,1.828,1.160 |
| **1e8b9**c5189c3e28bbd201526fade96c5 | 0.016,0.060,1.799,-1.651,-1.652,-0.533 | 0.026,0.032,0.190,-0.905,1.027,0.848 | 0.020,0.026,-0.502,0.147,1.272,0.513 | 0.057,0.051,-0.254,-0.047,0.674,1.754 |
| **1ea09**a163ef99ebfb4fb72a7abd81326 | 0.046,0.066,0.955,-1.072,-0.083,-1.019 | 0.023,0.076,0.726,-0.631,0.747,-1.029 | 0.041,0.069,-1.583,1.886,1.935,0.548 | 0.096,0.098,-1.728,1.377,0.374,1.707 |
| **1eb84**7ec6516d8666757d848a65ff4e0 | 0.092,0.017,1.734,-1.819,-1.618,-0.366 | 0.093,0.077,0.666,-0.433,-0.996,1.086 | 0.061,0.060,-1.237,1.903,-1.170,1.951 | 0.027,0.062,-1.488,1.084,1.727,1.641 |

| | | | | |
|---|---|---|---|---|
| **1eb9f**35dc8af726f28e4fc830d468cb1 | 0.090,0.066,1.581,-1.513,-0.409,-1.418 | 0.025,0.027,0.315,-0.394,-0.517,0.597 | 0.038,0.091,-0.959,0.873,1.938,-0.815 | 0.020,0.017,-0.947,0.919,-0.287,1.174 |
| **1eed9**903fbdf1fd2f6cc143d289636f4 | 0.092,0.018,1.036,-0.273,-1.912,0.183 | 0.042,0.060,1.161,-1.454,-0.738,-1.268 | 0.084,0.019,-1.836,0.807,1.151,1.814 | 0.064,0.069,-0.405,-0.168,0.772,1.824 |
| **1ef0c**0e879adfb301d3995bed6079dd2 | 0.081,0.059,1.342,-0.886,-1.636,-0.826 | 0.074,0.043,0.827,-1.159,-0.760,-0.141 | 0.044,0.087,-0.927,0.224,1.037,1.613 | 0.043,0.075,-0.101,-0.243,1.866,0.973 |
| **1f08a**5d5224d42aa35dd1f8afeb42c05 | 0.076,0.022,1.780,-1.342,-1.595,-0.395 | 0.098,0.042,0.845,-0.742,-0.817,-0.517 | 0.051,0.033,-1.745,0.844,0.944,1.225 | 0.098,0.062,-0.060,-0.858,1.978,1.183 |
| **1f104**a4109d796fb809a5ba33c492424 | 0.080,0.022,1.969,-1.767,-1.813,-0.482 | 0.018,0.021,1.421,-1.064,-1.784,-1.693 | 0.097,0.057,-1.933,1.964,0.887,1.451 | 0.097,0.056,-0.587,0.289,1.747,-0.406 |
| **1f189**6def738ba7697cd47ce2fff8db3 | 0.051,0.031,1.936,-1.551,-1.865,-1.119 | 0.015,0.034,1.285,-1.012,-1.186,0.947 | 0.098,0.017,-1.927,1.957,1.660,-0.151 | 0.086,0.096,-1.209,0.621,-1.011,1.803 |
| **1f548**7406dfb1d57ef2c500561195019 | 0.036,0.089,1.803,-1.836,-1.045,-0.862 | 0.027,0.027,0.204,-0.097,-1.262,-0.444 | 0.060,0.063,-0.607,0.365,0.345,0.650 | 0.088,0.012,-0.895,0.545,1.727,0.000 |
| **1fc7f**c63d41c6a00381d44b91f8b33a2 | 0.073,0.062,1.026,-0.698,-1.178,-0.355 | 0.036,0.061,0.458,-0.606,-1.891,1.036 | 0.098,0.059,-0.915,0.563,-0.366,1.775 | 0.039,0.085,-0.953,0.530,0.468,1.469 |
| **1fccd**2c1bd9a16c9b805e29717b537bc | 0.060,0.025,0.902,-0.954,-0.080,-1.225 | 0.066,0.088,0.880,-0.653,1.940,-1.756 | 0.096,0.090,-1.279,1.822,1.898,0.095 | 0.090,0.087,-0.472,-0.189,1.575,1.543 |
| **1fdf8**ba63f9a477afd5f5afd4c7516cf | 0.079,0.049,1.533,-1.527,-0.051,-0.496 | 0.065,0.066,0.802,-0.604,-0.493,0.004 | 0.098,0.060,-1.816,1.949,-0.816,1.686 | 0.043,0.063,-0.465,0.105,1.625,0.286 |
| **1ffbb**fe50caad11f98b69557622a21a0 | 0.056,0.030,1.027,-0.893,-1.339,-0.687 | 0.024,0.059,0.732,-1.119,-0.849,0.087 | 0.024,0.025,-0.942,0.791,1.197,0.841 | 0.054,0.048,-0.735,0.608,1.073,0.230 |
| **20008**57c488b001c5fd7e10673e43438 | 0.037,0.061,1.771,-1.827,-0.257,-0.964 | 0.041,0.029,1.209,-1.446,0.771,-1.826 | 0.071,0.091,-1.914,1.066,1.858,1.630 | 0.011,0.089,-0.504,0.053,1.914,1.516 |
| **20674**4f72b65123587c20a9741ba82f2 | 0.094,0.042,1.739,-1.344,-1.812,-0.174 | 0.091,0.066,0.933,-0.816,-1.221,-0.884 | 0.046,0.027,-1.116,0.626,-0.328,1.326 | 0.032,0.047,-0.086,-0.254,1.638,1.794 |
| **206a7**6ae344ad5f765280692208340de | 0.084,0.039,1.798,-1.474,-1.082,-0.962 | 0.077,0.051,1.024,-0.681,-0.829,-0.074 | 0.099,0.076,-0.941,-0.039,1.798,1.844 | 0.067,0.071,-0.956,0.330,0.162,1.952 |
| **206d5**452357017f0de34f59f4af7e06d | 0.080,0.027,1.976,-1.934,-1.697,0.020 | 0.095,0.025,0.109,-0.207,0.037,-1.034 | 0.013,0.085,-0.318,0.111,1.185,0.249 | 0.089,0.064,-0.569,0.140,1.964,1.631 |
| **2087b**c93cdb84ee89580b2b6ff5e3f27 | 0.049,0.020,1.657,-1.801,-1.492,0.119 | 0.063,0.056,0.981,-1.184,-1.182,-0.144 | 0.059,0.097,-0.588,0.123,1.886,0.430 | 0.063,0.063,-1.148,0.753,1.402,1.563 |
| **20ac6**a94c29d766b9c605fed38aebe5a | 0.042,0.094,1.810,-1.327,0.308,-1.849 | 0.042,0.049,1.783,-1.452,-0.724,-0.857 | 0.067,0.034,-1.060,0.964,0.756,1.289 | 0.034,0.051,-0.204,-0.200,1.489,1.165 |
| **20bbe**6c3d41b4bceb58dde19894fc964 | 0.068,0.015,0.531,-0.273,-1.447,-0.307 | 0.072,0.051,1.226,-0.790,-0.883,-0.268 | 0.081,0.083,-1.708,1.659,1.941,0.269 | 0.012,0.083,-0.813,0.368,1.264,1.402 |
| **21237**2f423f96d49cee526fe3bed4694 | 0.046,0.019,1.519,-1.602,-0.796,-0.177 | 0.059,0.073,0.890,-0.772,-0.699,-1.198 | 0.064,0.012,-0.355,0.370,0.122,0.734 | 0.039,0.093,-0.767,0.285,1.747,1.867 |
| **214db**8c03a5caab24b440f9f8c7fe244 | 0.045,0.031,1.012,-1.080,-1.002,-0.102 | 0.019,0.054,1.232,-0.890,0.495,-0.254 | 0.069,0.075,-1.197,1.867,-0.657,0.903 | 0.011,0.041,-0.622,0.549,1.054,0.407 |
| **2161b**d31de96795f2f88f0824b531d2a | 0.049,0.016,1.487,-1.559,-1.899,-0.114 | 0.071,0.034,0.747,-0.533,-1.845,-1.945 | 0.039,0.064,-0.576,0.537,-0.731,0.199 | 0.079,0.052,-0.394,0.053,1.878,1.044 |
| **21807**6d2f9e4e75990584018109e4f46 | 0.082,0.019,0.944,-0.747,-1.102,-0.128 | 0.076,0.089,0.797,-1.179,-0.772,0.031 | 0.025,0.066,-1.721,1.034,-0.045,0.567 | 0.095,0.016,-1.082,0.728,1.926,-0.108 |
| **21a8b**9a5fc5171b4da06d89da43b8afb | 0.095,0.083,1.939,-1.331,-1.974,-0.580 | 0.035,0.047,1.577,-1.346,-0.750,-1.008 | 0.043,0.049,-1.793,1.593,1.843,1.725 | 0.083,0.076,-1.156,0.347,1.915,1.413 |
| **21cce**d26956d83164b1ee34306c3b266 | 0.013,0.085,1.752,-1.883,-1.915,-1.923 | 0.087,0.077,0.712,-0.576,-0.145,-1.003 | 0.098,0.016,-1.798,1.912,1.726,0.704 | 0.088,0.066,-1.567,1.158,1.854,1.838 |
| **21e9e**8dcb9af0a28f7323c17e315afc3 | 0.075,0.015,1.617,-1.887,-0.835,1.188 | 0.078,0.029,1.466,-1.158,-0.083,1.843 | 0.051,0.063,-1.113,1.413,1.624,0.659 | 0.085,0.016,-1.158,0.784,1.576,1.099 |
| **222d9**83e5fc1fff1e9a6db05c53b4cc6 | 0.031,0.022,1.582,-1.844,-1.646,-0.681 | 0.015,0.018,0.835,-0.828,-0.889,0.823 | 0.072,0.060,-1.406,1.768,1.381,1.358 | 0.096,0.090,-0.679,0.258,1.325,0.846 |
| **2247b**fc2e84ed800cbda9546385e75e3 | 0.048,0.037,1.720,-1.854,-1.647,0.481 | 0.039,0.022,0.512,-0.528,-0.881,-0.844 | 0.083,0.057,-1.117,0.947,1.870,-0.513 | 0.076,0.074,-0.548,0.079,1.960,1.253 |
| **22816**07196aa5d7ce08e6d36cbb213aa | 0.086,0.090,1.715,-1.559,-1.867,-0.591 | 0.095,0.012,1.349,-0.958,-0.684,-1.538 | 0.018,0.047,-0.689,0.973,1.258,0.751 | 0.061,0.021,-0.811,0.636,1.530,-0.400 |
| **22fb7**0562c4e9617b28fcab81ffb3e45 | 0.042,0.050,1.139,-0.945,-0.910,-1.227 | 0.089,0.012,0.925,-1.922,0.224,1.849 | 0.098,0.032,-0.856,0.348,1.084,0.454 | 0.089,0.072,-0.700,0.446,0.437,1.294 |
| **23130**a5daf3569282fe8317d7404cd58 | 0.023,0.080,1.743,-1.393,-0.891,-1.570 | 0.043,0.062,1.937,-1.663,-0.504,0.166 | 0.043,0.047,-1.860,1.947,1.834,1.318 | 0.056,0.092,-1.060,0.551,1.728,0.303 |
| **232f5**2e2d17ee2ddbd0a54507b920a5a | 0.099,0.036,1.048,-0.960,-1.271,-1.267 | 0.037,0.058,0.941,-0.657,-0.770,-1.083 | 0.062,0.034,-1.281,1.232,1.154,0.929 | 0.056,0.052,-0.269,-0.035,1.121,1.276 |
| **2344e**a9dbe3c4db3b6d713f76882b67e | 0.098,0.051,1.642,-1.668,-0.573,-0.774 | 0.052,0.089,1.530,-1.252,-0.596,0.096 | 0.050,0.054,-1.366,1.944,-0.067,0.496 | 0.051,0.031,-0.694,0.490,1.706,0.165 |
| **23841**f8b872d674b980a6a7f39a447a7 | 0.025,0.078,1.878,-1.864,0.252,-0.773 | 0.078,0.031,1.011,-0.325,-1.932,-1.113 | 0.034,0.064,-0.780,1.147,-0.411,1.646 | 0.063,0.048,-1.362,1.079,1.038,1.479 |
| **23da8**f528c22504143a8f990b14b85c7 | 0.062,0.056,1.458,-1.442,-1.819,-0.184 | 0.099,0.097,0.450,-1.291,1.296,-0.005 | 0.048,0.046,-1.130,0.277,1.903,1.894 | 0.026,0.029,-0.490,0.161,1.329,1.937 |
| **23e8c**57a76bc1d538b5876d196ad3e41 | 0.078,0.022,1.177,-1.061,-1.784,0.758 | 0.070,0.066,1.867,-1.133,-0.870,-1.788 | 0.073,0.084,-1.117,1.213,-0.430,1.253 | 0.054,0.056,-0.451,0.035,1.949,1.131 |
| **23eb9**890d9ddfa0c8837652d1e1e809a | 0.080,0.024,1.537,-1.414,-1.003,-0.511 | 0.094,0.081,0.319,-0.149,-1.549,0.200 | 0.044,0.057,-1.105,0.798,0.398,0.077 | 0.027,0.065,-1.315,1.225,0.046,0.596 |
| **246f1**4b1299502452bc6a79d2523d265 | 0.076,0.090,1.608,-1.396,-1.940,-0.798 | 0.055,0.054,1.359,-0.815,-0.657,-1.420 | 0.095,0.017,-1.000,1.388,1.077,1.252 | 0.078,0.057,-0.996,0.666,1.834,1.191 |
| **24851**92155d0401c4d6025ada8c5da4b | 0.048,0.085,1.155,-1.115,-0.535,0.035 | 0.037,0.089,0.628,-1.684,0.108,-0.428 | 0.062,0.041,-1.939,0.649,-0.370,1.896 | 0.037,0.098,-0.427,-0.007,0.784,1.289 |
| **2488a**45dd435e47b35bf4374abb10b20 | 0.044,0.091,1.770,-1.510,-0.962,-1.254 | 0.083,0.077,1.669,-1.022,-1.216,-0.679 | 0.056,0.065,-1.170,1.599,1.274,0.597 | 0.089,0.080,-0.244,-0.147,1.462,0.503 |
| **248d0**712e79a8af6d2491ee5a4768181 | 0.094,0.029,1.901,-1.629,-1.104,-0.298 | 0.014,0.073,1.424,-0.999,-1.946,-1.260 | 0.039,0.085,-1.694,1.721,-0.232,0.982 | 0.028,0.014,-0.632,0.620,-1.111,1.671 |
| **24c3c**1d02ab237aa51f366c10fe16c16 | 0.034,0.070,1.461,-1.296,0.257,-1.917 | 0.039,0.029,1.623,-1.095,-1.964,1.125 | 0.092,0.094,-1.173,1.549,1.353,1.146 | 0.098,0.073,-0.828,0.461,0.924,0.754 |
| **25249**63390982953c2d9aeabd1a299d3 | 0.063,0.047,1.141,-1.277,-1.678,-0.474 | 0.074,0.030,0.166,-0.108,-0.174,0.182 | 0.086,0.022,-0.610,0.846,0.895,0.518 | 0.083,0.062,-0.449,0.137,0.778,1.555 |
| **2538a**ce41752e06d932c4bc9851b4ae8 | 0.099,0.074,1.502,-1.234,-1.781,-0.045 | 0.045,0.018,0.429,-0.399,-0.975,-1.153 | 0.082,0.055,-0.929,0.485,-0.212,1.836 | 0.057,0.056,-0.910,0.806,0.276,0.628 |
| **25434**e2746e538fff97210217a05a9cc | 0.077,0.027,1.391,-0.744,-1.879,0.022 | 0.022,0.019,1.002,-1.151,-1.642,-1.174 | 0.097,0.055,-1.037,0.448,1.182,-0.717 | 0.040,0.065,-0.389,0.211,1.755,-0.198 |
| **25715**062600806c98cbd5628d3314237 | 0.047,0.018,1.963,-1.759,-1.699,-0.419 | 0.066,0.035,0.569,-1.995,1.518,-0.003 | 0.022,0.061,-1.843,0.830,-0.918,1.362 | 0.060,0.036,-0.274,-0.205,1.813,0.536 |
| **257b0**3946e35fb165d27ae037517dd31 | 0.080,0.067,1.927,-1.673,-1.829,0.428 | 0.096,0.045,1.966,-1.439,-1.762,-1.663 | 0.097,0.045,-1.990,1.628,1.332,-0.095 | 0.080,0.040,-0.613,0.349,1.597,0.109 |
| **25804**f4d1b74feeeb214d063c2593ca5 | 0.078,0.017,1.737,-1.907,-1.079,0.609 | 0.052,0.068,1.260,-1.790,1.965,-1.213 | 0.044,0.034,-1.166,0.872,0.913,1.109 | 0.062,0.058,-0.866,0.423,1.885,1.302 |
| **2593c**186bea098770726df8fd8c2484f | 0.068,0.013,1.942,-1.407,-1.696,-0.927 | 0.060,0.014,0.300,-0.710,-1.357,0.428 | 0.059,0.096,-1.665,0.210,0.000,1.921 | 0.085,0.080,-0.411,0.016,1.849,1.055 |
| **25a21**4ba4ba11e9a95bd3f943f39b152 | 0.082,0.046,1.809,-1.817,-0.941,-0.756 | 0.043,0.073,0.338,-1.099,-1.094,1.385 | 0.080,0.021,-1.117,0.527,0.160,1.275 | 0.088,0.081,-0.701,0.270,1.744,0.638 |
| **25cea**3c1879dac679e9fa661c276b684 | 0.089,0.011,1.594,-1.154,-1.784,-0.256 | 0.039,0.043,1.647,-1.129,-0.816,-1.141 | 0.092,0.032,-1.510,1.275,1.523,1.588 | 0.039,0.082,-1.471,0.685,1.822,1.816 |
| **25d88**fad48e369fb2e5c1b9279a5d422 | 0.060,0.056,1.444,-1.355,-0.816,-0.608 | 0.098,0.072,1.301,-0.395,-1.227,-1.464 | 0.078,0.090,-1.165,1.854,1.815,1.345 | 0.086,0.096,-1.096,0.679,0.648,1.241 |
| **263c3**79f34858354b25274f50ff301b5 | 0.079,0.017,1.752,-1.722,-1.894,0.433 | 0.053,0.050,0.531,-0.290,-1.552,-1.149 | 0.033,0.087,-1.102,0.850,0.364,0.356 | 0.055,0.100,-0.894,0.633,1.734,0.421 |
| **26404**496633bc1b30145a462aa142d85 | 0.022,0.086,1.812,-1.531,-1.607,-1.063 | 0.070,0.019,0.885,-1.346,-0.609,-0.030 | 0.065,0.019,-0.777,0.429,0.375,1.771 | 0.085,0.071,-0.473,0.207,1.929,-0.726 |
| **26547**d5e90c126e0040b380a4e9fcb75 | 0.046,0.029,1.542,-1.590,-1.473,0.580 | 0.068,0.088,1.307,-0.993,0.234,-0.248 | 0.060,0.032,-0.978,1.636,0.969,0.374 | 0.044,0.070,-0.590,0.275,1.769,0.638 |
| **2664c**4291cfee1f91fea6aa5e13172a9 | 0.083,0.035,1.040,-1.059,-0.370,-0.642 | 0.027,0.047,0.583,-0.505,-0.267,-0.172 | 0.015,0.035,-1.515,1.789,-0.962,0.677 | 0.015,0.028,-1.188,1.131,-0.101,0.879 |
| **2669a**a8de7806bfa08eef2487ed7eab3 | 0.047,0.082,1.205,-0.778,-1.680,-1.108 | 0.046,0.045,1.463,-1.324,0.295,-1.200 | 0.096,0.036,-1.682,1.200,1.933,1.997 | 0.096,0.015,-0.989,0.434,1.313,1.592 |
| **266ef**35ec932b21ed366c35bef5eae46 | 0.045,0.050,1.101,-1.303,0.101,-0.467 | 0.072,0.025,1.414,-1.319,-0.863,0.968 | 0.049,0.035,-1.756,1.929,1.408,1.377 | 0.085,0.041,-1.382,0.932,1.843,0.935 |
| **26825**76fcb2db3f7652b98ca1e591290 | 0.092,0.017,1.708,-1.811,-1.903,0.156 | 0.025,0.028,0.657,-0.798,0.409,-1.611 | 0.099,0.057,-1.429,1.030,0.310,1.151 | 0.083,0.038,-0.276,0.021,0.842,1.824 |
| **2686a**bb6d393cfa34d1dacf51e59284a | 0.066,0.048,1.453,-0.955,-1.840,-1.093 | 0.063,0.063,1.535,-1.036,-1.078,-1.176 | 0.055,0.032,-1.998,1.937,1.704,-0.085 | 0.065,0.071,-0.715,0.043,1.403,1.951 |
| **26bbe**4a419ae65091eea1ad26ae21e55 | 0.076,0.067,1.211,-0.717,-1.916,0.042 | 0.016,0.095,0.490,-0.341,-1.632,-0.995 | 0.044,0.059,-1.740,1.016,0.983,0.079 | 0.013,0.073,-0.509,0.356,1.267,0.677 |
| **26cee**49f3ec69024616768785a95230c | 0.098,0.024,1.832,-1.518,-1.874,-0.441 | 0.014,0.074,0.511,-0.730,0.234,-0.276 | 0.064,0.035,-0.507,0.449,-0.617,0.534 | 0.011,0.093,-0.962,0.723,1.474,1.502 |
| **26dd8**5619c2d8eeb8cf816c69b91239f | 0.045,0.066,0.476,-0.251,-1.699,-0.487 | 0.080,0.019,0.928,-0.512,-1.379,-0.249 | 0.093,0.067,-1.628,1.756,0.882,1.002 | 0.013,0.053,-0.466,0.197,0.521,1.426 |
| **2735c**861914ae0493db5e1e6667f54f6 | 0.081,0.095,1.851,-1.240,-0.641,-1.808 | 0.055,0.065,1.105,-1.086,-0.987,-0.159 | 0.035,0.059,-1.642,1.385,1.315,1.689 | 0.092,0.064,-1.101,0.947,0.114,0.704 |
| **276fd**561e3e349bcedb33d2b7c7d7113 | 0.093,0.038,1.884,-1.819,-1.587,-0.605 | 0.044,0.021,0.960,-0.919,-1.907,-0.186 | 0.034,0.049,-1.237,0.575,1.369,1.951 | 0.012,0.046,-0.330,0.049,1.208,1.774 |
| **279b8**2fd72af7a529ecbb1d91e520012 | 0.098,0.038,1.731,-1.622,-1.834,-0.368 | 0.081,0.017,0.888,-1.163,-1.030,-1.145 | 0.097,0.044,-1.566,0.907,1.399,0.542 | 0.014,0.041,-0.811,0.687,0.947,1.554 |
| **27ca0**35780fd3834d884a9d92c606bb1 | 0.052,0.029,1.063,-0.979,-1.521,-0.509 | 0.069,0.049,0.688,-0.876,-1.332,-0.520 | 0.095,0.059,-0.625,0.276,0.721,0.805 | 0.093,0.060,-0.259,0.011,1.802,0.257 |
| **28283**74e46af90d1a053eeee189d37f3 | 0.095,0.027,1.166,-1.379,-0.881,1.385 | 0.046,0.041,0.660,-0.879,-0.586,-0.028 | 0.052,0.039,-1.623,1.297,1.934,1.457 | 0.099,0.059,-1.501,1.368,0.622,0.583 |
| **28478**3cfd4c0ec85c01a558deb04158b | 0.011,0.086,1.939,-1.655,-0.782,-1.975 | 0.099,0.018,1.989,-1.809,-1.933,1.800 | 0.075,0.062,-1.783,1.914,1.945,1.972 | 0.064,0.069,-1.617,1.122,1.737,1.907 |

| | | | | |
|---|---|---|---|---|
| **28553**3a8a8df778906c4d22350499de8 | 0.054,0.045,0.560,-0.380,-0.440,-1.550 | 0.071,0.070,1.513,-1.218,-1.044,-0.584 | 0.098,0.088,-1.311,1.293,1.249,0.575 | 0.070,0.060,-1.156,0.874,0.927,1.105 |
| **2857b**474768dde564a68b9e47ee6fa42 | 0.086,0.049,1.076,-0.920,-1.858,-0.253 | 0.049,0.076,0.639,-0.714,-0.451,1.058 | 0.036,0.065,-1.143,1.149,0.124,1.573 | 0.019,0.077,-0.489,0.081,1.813,1.631 |
| **285ee**89818c9ef5a84d8cb6556f640ee | 0.098,0.040,0.895,-0.689,-0.928,0.483 | 0.044,0.085,0.871,-1.877,1.458,-0.256 | 0.069,0.062,-1.423,0.454,1.734,0.489 | 0.064,0.078,-1.851,1.304,1.256,0.979 |
| **28e9f**a1317b8171c53cb7d43733803fa | 0.066,0.029,1.225,-1.092,-1.585,-0.851 | 0.064,0.080,0.409,-1.098,0.726,0.186 | 0.031,0.066,-0.276,-0.034,1.811,0.692 | 0.091,0.055,-0.197,-0.085,0.739,0.896 |
| **28f11**ac311ca08b03a9c436e53ad42c4 | 0.054,0.022,1.320,-1.841,-0.471,1.291 | 0.095,0.051,0.407,-0.601,-0.627,0.851 | 0.083,0.030,-0.382,0.174,0.710,0.241 | 0.042,0.078,-1.451,1.285,1.662,0.617 |
| **292db**c51e6a51d22ce2a03261ac66a5c | 0.060,0.067,0.921,-0.627,-0.998,-0.887 | 0.025,0.077,1.122,-1.130,-0.996,-0.402 | 0.045,0.059,-1.140,0.931,0.825,0.515 | 0.071,0.062,-0.091,-0.426,1.682,0.971 |
| **29392**7ec77d8253ce46982511ea220ec | 0.092,0.064,1.577,-1.693,-1.509,-1.254 | 0.014,0.066,1.757,-1.572,-0.999,0.514 | 0.099,0.037,-1.368,1.990,-0.360,1.167 | 0.095,0.047,-0.991,0.694,1.923,0.917 |
| **2969d**f807495d134eac363db1b92ca86 | 0.052,0.032,1.053,-0.817,-1.017,-0.496 | 0.089,0.054,0.771,-1.904,-0.933,-0.996 | 0.057,0.062,-1.416,0.631,-0.628,0.015 | 0.074,0.057,-0.780,0.403,1.593,1.225 |
| **297cb**e12d44dafd8622763e5bd18ab7d | 0.043,0.034,1.401,-1.376,-0.575,-0.609 | 0.046,0.016,0.708,-1.732,-1.851,-0.062 | 0.061,0.020,-0.526,0.123,0.913,1.055 | 0.048,0.080,-0.176,-0.244,1.907,1.708 |
| **297db**e859bf8955438133f0bb516b887 | 0.046,0.060,1.771,-1.712,-0.486,-1.150 | 0.041,0.059,0.520,-0.663,-0.140,-1.638 | 0.072,0.080,-1.234,0.306,1.777,1.634 | 0.098,0.070,-1.167,0.888,1.282,0.617 |
| **298cd**a021898b3aa0cd5aed7115799e9 | 0.049,0.014,1.560,-1.459,-1.852,-0.172 | 0.071,0.087,1.269,-1.622,-0.050,-0.604 | 0.079,0.018,-1.558,1.016,1.598,0.501 | 0.094,0.064,-0.409,-0.018,1.613,0.523 |
| **29995**0efa92530ba1b8dc83f2207c22d | 0.077,0.052,0.468,-0.248,-0.394,-0.893 | 0.052,0.080,1.125,-0.963,-1.010,-0.129 | 0.044,0.056,-1.937,1.641,0.128,1.958 | 0.079,0.070,-0.762,0.150,1.627,1.082 |
| **29998**9f597582f4990155fbd58f4ee20 | 0.079,0.042,1.957,-1.660,-0.985,-0.306 | 0.063,0.067,0.408,-0.494,-0.580,-1.291 | 0.066,0.023,-1.420,0.362,1.108,1.603 | 0.072,0.034,-1.302,0.974,1.892,0.893 |
| **29c57**951109e2aeb0e667de8657bc79f | 0.070,0.040,1.414,-1.387,-1.653,0.428 | 0.022,0.082,1.071,-1.494,-0.296,-0.929 | 0.014,0.083,-1.114,0.632,-0.483,0.662 | 0.019,0.073,-0.122,-0.136,0.941,1.315 |
| **29d8d**ed3c3c8daff993bb6bac50422a5 | 0.071,0.065,1.905,-1.769,-0.975,-1.127 | 0.071,0.052,0.868,-1.169,1.954,-0.553 | 0.097,0.087,-0.733,0.364,1.880,0.539 | 0.055,0.062,-0.664,0.429,1.967,-0.477 |
| **29e70**0a8995f77d286ac9974f17a1751 | 0.097,0.039,1.907,-1.535,-1.786,0.121 | 0.069,0.032,0.623,-0.579,-0.686,-1.886 | 0.081,0.093,-1.845,0.460,1.694,1.957 | 0.091,0.067,-1.814,1.018,1.944,1.810 |
| **29eec**d9199c6298f24da08fbf546c8e4 | 0.054,0.037,1.646,-1.616,-1.505,0.435 | 0.030,0.025,0.521,-1.127,-0.578,-1.810 | 0.083,0.081,-0.865,0.121,1.964,0.473 | 0.055,0.070,-0.875,0.517,1.972,1.743 |
| **29fc0**eb1f55a6bfae908389ff2f0f57a | 0.078,0.098,1.359,-1.217,0.444,-1.734 | 0.016,0.053,0.287,-0.418,-0.961,-0.077 | 0.031,0.037,-1.247,0.867,-0.800,1.981 | 0.020,0.024,-0.319,0.185,1.416,1.767 |
| **2a00b**31145a5f664e731d9d7bfbbc989 | 0.084,0.027,1.389,-1.602,-1.757,0.496 | 0.028,0.083,0.566,-0.240,-1.290,-1.678 | 0.067,0.043,-0.558,0.705,-0.101,1.189 | 0.051,0.045,-0.371,0.349,0.525,0.051 |
| **2a026**1eba6e0080f46dc1286623d4a36 | 0.047,0.068,0.938,-0.646,-1.759,0.075 | 0.018,0.077,0.566,-0.924,-1.903,-1.018 | 0.080,0.084,-1.844,0.264,1.772,1.840 | 0.099,0.059,-1.025,0.865,-0.375,1.694 |
| **2a118**5b6939f03c3e25c13d84930f014 | 0.061,0.034,1.619,-1.169,-1.533,-0.393 | 0.019,0.080,0.663,-1.086,-0.503,-0.859 | 0.040,0.043,-1.619,0.516,0.979,1.054 | 0.095,0.060,-0.453,-0.048,1.055,1.100 |
| **2a218**7fab6aff45f48efa4f7aa2732a4 | 0.051,0.036,1.569,-1.498,-1.947,0.391 | 0.022,0.089,0.771,-0.808,-0.911,0.098 | 0.069,0.082,-1.500,1.184,1.084,0.581 | 0.076,0.072,-1.357,0.675,1.896,0.965 |
| **2a245**d3612f3fb86fc5b65a45de434e7 | 0.044,0.058,1.448,-1.792,0.437,-0.597 | 0.014,0.048,0.142,-0.141,-0.651,-0.253 | 0.089,0.097,-1.391,0.953,-0.114,0.798 | 0.087,0.065,-1.404,1.202,0.149,1.503 |
| **2a3ef**93330d704d92f690fc0b74258a5 | 0.040,0.064,0.856,-0.908,0.565,-0.701 | 0.062,0.063,0.563,-0.839,0.561,0.224 | 0.096,0.089,-1.000,0.536,-0.293,1.881 | 0.092,0.075,-1.290,0.978,-0.457,1.942 |
| **2a583**fb1c915076173a089e5bd2a5a2d | 0.014,0.062,1.629,-1.571,0.294,-1.459 | 0.019,0.037,0.783,-1.770,-0.119,0.027 | 0.033,0.083,-1.770,0.549,-0.546,1.899 | 0.040,0.051,-0.190,0.022,1.728,0.378 |
| **2a72a**a10c968221a1dce184fc9f83c67 | 0.067,0.023,1.572,-1.504,-1.699,0.556 | 0.044,0.025,1.064,-1.968,0.100,-0.561 | 0.047,0.086,-1.679,0.796,-0.079,1.185 | 0.046,0.064,-1.312,0.918,1.971,1.589 |
| **2a75b**47a61f99642d0faa3589cd54cc2 | 0.076,0.022,1.658,-1.642,-1.676,-0.079 | 0.034,0.045,1.344,-1.493,-1.645,-1.651 | 0.096,0.057,-1.664,0.986,1.021,0.847 | 0.088,0.098,-1.558,0.997,1.582,1.627 |
| **2a802**8c21266f41d7b194caf9b6d6d9b | 0.094,0.035,1.264,-1.341,-0.760,0.178 | 0.045,0.033,0.728,-1.484,-1.136,-1.237 | 0.045,0.026,-1.307,0.414,1.760,0.467 | 0.079,0.088,-0.268,-0.038,1.959,0.237 |
| **2a804**2649e8807dbab794b09a59556ca | 0.042,0.074,1.845,-1.537,-0.323,-1.648 | 0.081,0.058,0.762,-0.273,-1.537,-1.165 | 0.054,0.067,-1.083,0.788,1.221,0.746 | 0.029,0.042,-0.217,0.145,0.557,0.419 |
| **2a90f**0dd4d462fe2027002bcdfd2e85c | 0.087,0.064,1.484,-1.318,-1.909,0.208 | 0.073,0.065,0.805,-1.903,-0.316,-0.091 | 0.072,0.091,-1.855,0.839,-0.229,1.140 | 0.057,0.085,-1.665,1.458,1.898,0.695 |
| **2abdc**dc1a113e499829325ba99965362 | 0.044,0.072,0.753,-0.669,-1.445,0.098 | 0.042,0.022,1.009,-1.681,-1.648,-1.201 | 0.093,0.044,-1.556,0.685,1.267,1.125 | 0.095,0.060,-0.343,0.201,0.871,0.461 |
| **2acff**efc1345335302a4896286b4a6a7 | 0.060,0.039,1.700,-1.791,-1.550,0.073 | 0.089,0.091,0.422,-0.328,-1.543,0.526 | 0.060,0.067,-1.580,1.956,-1.455,-0.301 | 0.056,0.099,-0.578,0.286,1.025,0.683 |
| **2b470**b09dc2ccf94d08eda393b4c1d2a | 0.014,0.035,1.353,-1.197,-1.425,-1.104 | 0.036,0.080,0.077,0.024,-0.998,-0.288 | 0.076,0.064,-0.657,0.015,1.715,0.765 | 0.080,0.092,-0.726,0.259,1.767,1.510 |
| **2b5f6**f1d8a535698da2dd2f12bc93ce5 | 0.098,0.022,1.696,-1.131,-1.872,-0.816 | 0.082,0.040,0.961,-1.334,-1.482,-1.826 | 0.013,0.060,-1.387,0.535,-1.146,1.652 | 0.086,0.043,-1.117,0.800,1.950,0.268 |
| **2b7ab**12757ca7eca044b20040d06dd63 | 0.073,0.097,1.641,-1.622,-1.959,-0.416 | 0.021,0.090,1.116,-1.173,-1.970,-0.396 | 0.056,0.071,-1.146,1.016,1.498,0.062 | 0.066,0.069,-0.831,0.446,1.858,1.434 |
| **2b9bb**d9a350b02a473bf6f6c6dd07740 | 0.096,0.018,1.186,-0.798,-1.905,0.016 | 0.029,0.083,1.469,-0.907,-0.297,-0.540 | 0.024,0.062,-1.420,1.921,1.478,1.478 | 0.048,0.027,-0.578,-0.035,1.796,1.591 |
| **2ba5b**0bf95014964b4795cbb7ca50c61 | 0.044,0.034,1.689,-1.891,-0.105,-0.637 | 0.091,0.045,0.308,-0.719,0.224,0.033 | 0.067,0.076,-1.765,0.851,1.259,-0.199 | 0.065,0.019,-0.452,0.337,1.834,-1.338 |
| **2bb4d**e0f7254e4c17e27ca3dac6afc98 | 0.081,0.087,1.711,-1.515,-0.552,-0.600 | 0.022,0.068,1.299,-1.532,1.451,-1.826 | 0.088,0.087,-1.639,1.611,1.681,-1.561 | 0.076,0.038,-0.332,-0.198,1.874,1.217 |
| **2bc33**9c5846b7934a62f37ff4608f7e6 | 0.076,0.046,1.584,-1.433,-1.733,-1.095 | 0.060,0.072,0.208,-0.614,-0.452,1.478 | 0.082,0.031,-0.476,0.036,1.782,0.643 | 0.072,0.073,-1.150,0.808,0.519,1.339 |
| **2bc5f**b71f14e7541827ca7cc19897c8a | 0.087,0.022,1.348,-1.413,-1.690,0.840 | 0.082,0.041,1.066,-1.269,-0.088,-1.096 | 0.052,0.052,-1.722,1.330,1.397,0.151 | 0.092,0.085,-0.170,-0.348,0.953,1.721 |
| **2be09**674b7fe0f6adff227d880d63a8f | 0.020,0.041,0.844,-1.215,0.124,-0.963 | 0.016,0.017,0.533,-0.250,-1.448,0.341 | 0.100,0.096,-1.247,1.764,1.946,0.274 | 0.045,0.070,-0.936,0.555,1.213,0.651 |
| **2c026**f7bbc335879accd4c893c7496c8 | 0.076,0.058,1.361,-1.289,-1.231,-0.283 | 0.023,0.017,0.654,-0.583,-1.486,-1.969 | 0.035,0.036,-1.214,1.128,1.831,1.065 | 0.062,0.050,-0.825,0.626,1.281,0.897 |
| **2c3d4**6af807f603e0284d24f6841fdd7 | 0.017,0.018,1.726,-1.771,-1.914,-0.708 | 0.017,0.023,0.634,-0.857,0.197,0.062 | 0.017,0.066,-0.867,0.298,1.807,1.809 | 0.015,0.085,-0.605,0.127,1.976,1.447 |
| **2c842**0420e1c078571f34446926ec7ff | 0.050,0.040,1.407,-1.463,-1.607,-1.241 | 0.096,0.032,0.416,-0.704,0.502,-0.837 | 0.027,0.069,-0.663,0.227,0.849,1.958 | 0.096,0.062,-0.635,0.135,1.673,1.960 |
| **2c879**e0bfea8172edb68a74255378ca2 | 0.085,0.052,0.124,0.065,-1.455,-0.069 | 0.051,0.046,0.902,-0.811,-0.860,-0.782 | 0.097,0.100,-1.505,1.900,-0.406,0.169 | 0.012,0.094,-0.987,0.732,-0.792,1.562 |
| **2ca23**7e9435fe702ccdd71382d687b49 | 0.017,0.046,1.389,-1.414,-1.336,-0.408 | 0.025,0.047,1.100,-0.917,-0.822,-0.190 | 0.023,0.042,-0.947,1.495,1.226,-0.528 | 0.081,0.028,-1.348,0.875,1.958,1.522 |
| **2ca8e**ab82be440da2c74ff6cd91ba7bf | 0.072,0.056,0.842,-0.647,-1.782,-0.481 | 0.031,0.052,1.710,-1.088,-0.287,-0.491 | 0.069,0.011,-0.969,1.897,1.337,0.186 | 0.043,0.072,-0.255,0.152,0.181,0.444 |
| **2ceb2**f8da2e1284a4186ea18d55faf16 | 0.094,0.016,1.761,-1.665,-0.925,1.241 | 0.099,0.082,0.663,-0.211,-1.384,-0.197 | 0.028,0.076,-0.828,0.762,0.402,1.710 | 0.096,0.048,-0.457,-0.066,1.227,1.760 |
| **2cf5d**6134dd21e2e9572cba88023f29c | 0.082,0.056,1.254,-1.349,-1.259,-0.204 | 0.012,0.018,1.229,-0.545,0.840,0.755 | 0.098,0.098,-0.824,1.934,-0.207,0.774 | 0.019,0.094,-0.956,0.577,1.465,1.636 |
| **2d0ae**4cc94b43a5e5807cba46efc937c | 0.091,0.016,1.700,-1.581,-1.982,-0.004 | 0.064,0.064,1.116,-1.338,-0.903,-1.176 | 0.094,0.100,-1.531,1.082,1.376,-0.576 | 0.074,0.071,-0.194,-0.182,1.181,1.752 |
| **2d0ea**81e6b4d8db4b41a16c73bf025fa | 0.089,0.021,0.435,-0.350,-1.534,1.370 | 0.074,0.082,0.531,-0.216,-1.467,-0.294 | 0.066,0.062,-1.269,1.701,1.129,1.616 | 0.025,0.070,-0.991,0.880,-0.558,0.853 |
| **2d4e8**76f2f59310ac8b521ec267ec3b1 | 0.089,0.061,1.157,-0.908,-1.069,0.038 | 0.015,0.058,0.145,-0.197,-0.934,-0.148 | 0.098,0.078,-0.669,0.099,1.532,0.409 | 0.076,0.073,-0.979,0.624,1.540,1.275 |
| **2d68d**8db75dd573cb65dc26aa8fd1f3e | 0.092,0.083,1.675,-1.151,-1.857,-0.374 | 0.031,0.016,0.241,-0.137,-1.694,-0.120 | 0.090,0.028,-1.444,0.658,0.859,1.863 | 0.075,0.076,-1.270,1.178,1.761,-0.993 |
| **2d770**e10543bf7f19695caf24aab4035 | 0.087,0.026,0.987,-0.605,-1.026,-1.159 | 0.042,0.089,1.637,-1.558,-0.723,-1.186 | 0.039,0.050,-1.763,1.210,1.860,0.166 | 0.042,0.090,-1.000,0.413,1.321,1.466 |
| **2da68**1e21fa0a01d1a247297f989b669 | 0.044,0.059,1.309,-1.234,-1.425,-0.097 | 0.020,0.078,1.452,-1.345,0.908,-0.454 | 0.055,0.062,-1.730,1.925,1.538,1.248 | 0.061,0.088,-0.946,0.364,0.636,1.900 |
| **2dcfb**d05eefd2d9646acd0ede2a14820 | 0.046,0.037,1.299,-1.024,-1.537,-0.481 | 0.027,0.080,0.973,-1.426,-1.722,-0.173 | 0.082,0.093,-1.449,0.582,1.861,0.716 | 0.012,0.048,-0.169,-0.019,1.047,1.154 |
| **2ddf7**e12479d375a49ddfc41de196138 | 0.057,0.016,1.457,-1.724,-0.833,0.332 | 0.022,0.048,1.521,-1.130,-1.261,-0.333 | 0.071,0.060,-1.833,1.945,0.966,0.788 | 0.095,0.070,-1.274,0.740,0.734,1.660 |
| **2de43**59d52dabc4da63e7d7525479005 | 0.062,0.046,1.668,-1.688,-0.441,-1.039 | 0.041,0.098,1.102,-0.709,-1.230,-1.005 | 0.091,0.047,-0.314,0.264,0.568,0.510 | 0.058,0.067,-0.165,-0.101,1.953,0.729 |
| **2debf**69782aeba9d2862f4db79c4edf0 | 0.078,0.025,1.544,-1.597,-1.598,0.880 | 0.034,0.018,0.696,-0.617,-1.426,-0.086 | 0.055,0.079,-0.630,0.302,1.551,0.631 | 0.076,0.033,-0.057,-0.272,1.901,-0.565 |
| **2e004**42d115d64fff536a60d8a09d5c4 | 0.058,0.045,1.393,-1.659,0.436,-1.722 | 0.045,0.010,0.899,-1.194,-1.424,0.395 | 0.075,0.081,-1.296,0.685,1.722,1.310 | 0.098,0.041,-0.727,0.462,0.921,1.771 |
| **2e180**94467ce159863766fef4e582835 | 0.036,0.050,1.633,-1.746,-0.961,-0.556 | 0.039,0.042,1.325,-0.936,-0.806,-0.902 | 0.092,0.017,-1.404,1.859,1.804,-0.398 | 0.077,0.085,-0.820,0.068,1.699,1.862 |
| **2e2ec**a66edeeff26292d85a46a3ea2e7 | 0.012,0.062,1.531,-1.642,-0.173,-1.224 | 0.093,0.056,0.995,-1.163,-1.711,0.386 | 0.084,0.063,-1.367,0.806,1.320,1.744 | 0.054,0.098,-1.273,0.818,1.574,1.745 |
| **2e50c**138484fdf460d76685e5aa169f3 | 0.045,0.047,1.757,-1.106,-1.812,-0.539 | 0.092,0.071,0.733,-1.357,-1.194,0.678 | 0.044,0.084,-0.746,0.127,1.759,0.201 | 0.083,0.066,-1.811,1.319,0.554,1.653 |
| **2e6e6**8ea9934dc24b17bcab02be3aa5f | 0.087,0.035,0.610,-0.505,-0.650,-0.114 | 0.093,0.077,0.762,-0.523,0.796,-0.789 | 0.046,0.056,-0.950,1.125,1.544,1.871 | 0.068,0.095,-0.920,0.507,0.801,0.982 |
| **2ee6c**8c0ce93c2cf5835de69d212e86c | 0.020,0.088,1.511,-1.197,-0.679,-1.059 | 0.023,0.077,1.495,-1.425,-1.445,-1.334 | 0.031,0.071,-1.956,1.760,-0.163,-0.332 | 0.068,0.054,-0.661,0.241,0.947,1.299 |
| **2efb0**6104b408177ed138ca43d2f1b40 | 0.096,0.020,1.526,-1.404,-0.728,0.051 | 0.042,0.086,0.911,-0.493,-1.904,-0.286 | 0.071,0.055,-1.978,1.919,1.951,1.555 | 0.033,0.064,-1.461,1.315,1.629,0.082 |
| **2f2b2**8e84514a0fdb260154ec125e930 | 0.045,0.055,0.516,-0.468,0.025,-0.785 | 0.060,0.048,0.869,-0.375,-1.415,-0.522 | 0.076,0.036,-0.685,1.360,0.204,0.410 | 0.098,0.097,-1.450,1.136,0.994,0.750 |
| **2f43d**d6f2ad535d199ad5ad48822cb6b | 0.098,0.058,1.691,-1.590,-0.805,-0.981 | 0.058,0.049,1.241,-0.646,-0.382,-1.128 | 0.031,0.092,-1.082,1.752,1.312,0.218 | 0.094,0.051,-1.231,0.812,0.507,1.731 |

| | | | | |
|---|---|---|---|---|
| **2f45e**a285cbe166e36d95a5e1257f764 | 0.052,0.020,0.385,-0.344,-1.351,0.177 | 0.089,0.083,0.486,-0.297,-1.410,0.255 | 0.030,0.089,-0.271,0.754,1.310,-1.905 | 0.037,0.065,-1.124,1.073,-0.038,0.478 |
| **2faae**427a7345a6374c49a9ac6554249 | 0.082,0.052,0.890,-1.077,0.105,-0.179 | 0.071,0.053,1.087,-1.342,-0.687,-0.611 | 0.066,0.091,-1.694,1.256,0.666,1.881 | 0.086,0.070,-0.572,0.346,1.481,0.323 |
| **2fd1f**6ac03a7db0e8e921bbcb3ee8395 | 0.039,0.046,0.919,-1.145,-0.790,-0.523 | 0.021,0.084,0.823,-0.416,-1.146,-0.934 | 0.021,0.055,-1.271,1.889,-0.895,1.165 | 0.034,0.044,-0.665,0.544,1.308,0.453 |
| **2fdff**3e08ffe4e15d8fb3e1766016d19 | 0.059,0.050,1.975,-1.738,-1.743,-0.121 | 0.049,0.046,0.782,-0.654,-1.502,-1.760 | 0.025,0.051,-1.181,0.745,-0.745,-0.448 | 0.052,0.060,-0.646,0.414,0.231,1.108 |
| **30034**78cdec79f3ebf1398f9717b315d | 0.014,0.064,1.854,-1.619,-0.151,-1.600 | 0.026,0.015,1.140,-1.285,-0.794,-0.570 | 0.098,0.061,-1.867,1.144,1.167,1.358 | 0.063,0.083,-1.809,1.219,1.820,1.184 |
| **30097**a238ccfa477503c087ab5467b00 | 0.014,0.030,1.126,-1.386,0.705,-0.780 | 0.093,0.046,1.899,-1.593,-0.105,0.437 | 0.099,0.085,-1.338,1.193,1.841,0.000 | 0.045,0.057,-0.510,0.162,1.417,0.532 |
| **3010e**b477c5fca8ad3abb9cb5bf663d2 | 0.090,0.013,1.617,-1.465,-1.906,0.860 | 0.074,0.080,1.068,-1.523,-0.733,1.297 | 0.027,0.077,-1.853,1.569,1.737,1.496 | 0.091,0.055,-1.120,0.711,1.845,0.623 |
| **30132**941f4b7a6b756d8216a0e7759b3 | 0.062,0.062,0.805,-0.738,-0.169,-1.033 | 0.075,0.041,0.848,-0.966,-1.926,0.486 | 0.086,0.069,-0.530,0.451,-0.223,0.386 | 0.063,0.055,-0.559,0.447,0.675,0.761 |
| **302c2**5cea616a61ccf61aa661b51ea00 | 0.073,0.082,1.596,-1.370,-0.699,-0.246 | 0.053,0.031,1.041,-1.218,0.274,-0.345 | 0.036,0.036,-1.938,1.832,0.380,1.686 | 0.012,0.025,-0.375,0.345,1.778,-0.004 |
| **303fd**d966e11a20686d003a48d66c5af | 0.086,0.012,1.210,-1.059,-0.695,-0.731 | 0.056,0.014,1.599,-1.038,-0.734,0.035 | 0.046,0.038,-0.751,0.490,1.720,1.581 | 0.064,0.039,-1.273,0.568,1.863,1.074 |
| **30838**827ec3675e525adb9c1cbdd9453 | 0.040,0.088,-0.016,0.531,-1.411,-0.449 | 0.071,0.022,0.421,-0.014,-0.623,-0.820 | 0.060,0.057,-1.608,1.132,0.411,1.956 | 0.057,0.059,-0.691,-0.113,0.046,1.798 |
| **3088f**76a79182a85749b4daf3890f5f6 | 0.093,0.061,1.407,-1.227,-1.072,-0.672 | 0.066,0.050,0.424,-0.413,-0.457,1.094 | 0.085,0.090,-0.521,0.206,1.780,1.606 | 0.075,0.027,-0.236,0.154,1.228,-0.712 |
| **30cfc**fd87a85786190b0acaa54b53a2b | 0.045,0.070,1.608,-1.172,-1.914,-0.442 | 0.043,0.021,0.185,-0.416,-0.169,0.515 | 0.088,0.091,-1.637,0.297,1.635,0.723 | 0.060,0.066,-0.120,-0.015,0.848,0.052 |
| **30dfd**3aa6c64e90e86620899916618e4 | 0.015,0.018,1.609,-1.815,-1.238,-0.249 | 0.066,0.030,0.521,-0.427,-1.523,-0.642 | 0.094,0.063,-0.736,0.171,0.370,-0.043 | 0.051,0.071,-1.584,1.240,1.392,0.373 |
| **30ec9**417c66daef023357e22324ac4be | 0.096,0.023,1.645,-1.579,-1.302,0.577 | 0.022,0.055,0.477,-1.225,-0.344,-0.544 | 0.091,0.025,-0.943,0.520,-0.152,1.093 | 0.079,0.096,-1.025,0.536,1.904,1.835 |
| **3179a**2b1f4cf48a6cce3761a0e0f4083 | 0.057,0.013,1.563,-1.799,-1.869,-0.221 | 0.094,0.082,0.319,-0.284,0.207,0.024 | 0.030,0.068,-0.381,0.388,-0.287,0.929 | 0.055,0.064,-1.201,0.741,1.002,1.498 |
| **31806**413081a0497cba0eb8897f97685 | 0.079,0.048,1.319,-1.357,-0.321,-0.095 | 0.092,0.051,1.935,-1.730,-1.670,-1.255 | 0.092,0.084,-1.174,0.730,1.702,0.568 | 0.084,0.090,-0.825,0.441,1.626,0.615 |
| **3184d**1ef168504db0cc4e4935205ee06 | 0.098,0.046,1.599,-1.792,-0.826,1.069 | 0.093,0.088,0.987,-1.307,-0.176,-0.888 | 0.096,0.089,-1.898,1.367,-0.385,1.391 | 0.012,0.069,-1.584,1.381,0.974,1.080 |
| **3199d**8c38e389f851aa765ec3ef98771 | 0.087,0.039,0.969,-0.885,-0.999,0.209 | 0.089,0.080,0.421,-0.436,-0.509,-0.958 | 0.065,0.039,-1.658,0.781,-0.342,1.930 | 0.011,0.084,-0.543,0.315,-1.044,1.381 |
| **31a94**61d9457013aa0213211a1d09a94 | 0.087,0.022,1.002,-0.805,-1.204,0.989 | 0.014,0.014,1.170,-1.227,1.847,-0.434 | 0.013,0.099,-1.830,1.768,1.198,1.580 | 0.071,0.061,-1.517,1.327,1.516,-0.791 |
| **31c5a**dca06d802dbc8cb665108fbb061 | 0.071,0.022,0.620,-0.750,-0.130,0.612 | 0.025,0.057,0.838,-0.437,-0.488,-0.071 | 0.078,0.095,-0.876,1.945,-0.161,0.138 | 0.040,0.059,-0.594,0.385,0.931,0.657 |
| **31d37**1bd094c630c93808386ebc39aef | 0.087,0.027,1.708,-1.606,-1.757,0.148 | 0.059,0.018,0.639,-0.205,-1.949,-1.245 | 0.021,0.033,-0.535,0.897,0.685,-0.912 | 0.094,0.069,-1.534,1.197,1.177,0.719 |
| **31e45**b81a242d7b9446d0265e32505b5 | 0.066,0.066,1.723,-1.326,-1.861,-0.544 | 0.063,0.065,0.746,-1.113,0.145,-1.139 | 0.042,0.073,-1.915,1.204,0.990,1.401 | 0.027,0.090,-1.512,1.232,0.345,1.678 |
| **31e91**9caa0610bc69b0034ae045b3842 | 0.014,0.019,1.444,-1.457,1.043,-1.840 | 0.054,0.078,0.696,-0.354,-1.180,-1.821 | 0.069,0.067,-0.989,-0.017,1.401,1.869 | 0.087,0.073,-0.662,0.424,1.610,-0.408 |
| **31eda**9e4a65885a4d863a1d593f1345e | 0.096,0.053,1.509,-1.450,-1.129,0.401 | 0.089,0.055,1.698,-1.368,-0.389,-1.158 | 0.083,0.072,-1.078,1.310,-0.430,1.244 | 0.050,0.074,-1.529,1.164,1.671,1.403 |
| **32227**3622411f88062341c67aa0741ba | 0.040,0.037,1.114,-1.193,-1.619,-0.323 | 0.066,0.056,0.589,-0.941,-1.465,0.060 | 0.058,0.043,-1.587,0.513,0.174,1.878 | 0.049,0.045,-0.506,0.292,1.942,0.573 |
| **32362**0479659bed874e239248d5e4e3a | 0.041,0.053,1.058,-1.227,-1.883,-0.193 | 0.021,0.015,1.354,-1.236,1.774,-0.101 | 0.016,0.027,-1.480,1.935,1.907,1.908 | 0.021,0.065,-1.437,1.203,1.878,1.579 |
| **32539**d6630208156f7e111147e294991 | 0.048,0.082,1.051,-1.013,-0.747,0.089 | 0.071,0.070,0.639,-0.704,0.654,-1.701 | 0.078,0.096,-0.825,0.754,-0.544,0.380 | 0.080,0.060,-0.459,0.203,1.926,0.009 |
| **32562**f47d0a068c7e94646a45b387e88 | 0.077,0.061,1.975,-1.763,0.038,-1.251 | 0.024,0.026,0.736,-1.372,-1.378,-1.617 | 0.016,0.060,-1.052,0.370,-0.848,0.896 | 0.085,0.012,-0.514,0.080,1.849,-0.560 |
| **326bf**8328cc72ab8009964bfc73cdc73 | 0.045,0.059,1.097,-1.046,-1.704,-0.533 | 0.039,0.077,0.285,-0.092,-0.617,-1.040 | 0.095,0.043,-1.033,1.218,-0.697,1.901 | 0.098,0.066,-0.891,0.444,1.424,1.658 |
| **3275e**b96d444c56effde9205f563e4ea | 0.054,0.048,1.569,-1.234,-1.273,-0.306 | 0.016,0.098,0.543,-0.613,-0.316,-1.135 | 0.017,0.091,-1.934,0.598,1.293,1.559 | 0.011,0.034,-1.294,1.262,-1.396,0.539 |
| **32afd**39664999548009f63efb59786d3 | 0.042,0.031,1.673,-1.825,-0.799,-0.335 | 0.051,0.027,0.414,-0.711,-1.422,0.206 | 0.061,0.031,-0.627,0.288,-0.275,0.739 | 0.086,0.064,-0.455,-0.099,1.938,1.505 |
| **32fba**876711a6508c17fbb4634142fcd | 0.028,0.074,1.370,-1.129,-0.508,-1.166 | 0.041,0.021,1.724,-1.284,-0.583,-0.041 | 0.056,0.053,-1.771,1.900,1.538,1.734 | 0.042,0.063,-0.764,0.261,1.403,1.579 |
| **33347**23d603ee41cea44cd864fa9465e | 0.094,0.013,1.597,-1.590,-1.959,1.002 | 0.075,0.058,0.206,-0.540,1.769,-1.152 | 0.034,0.089,-0.415,0.253,1.665,0.408 | 0.080,0.068,-0.538,0.246,1.926,0.640 |
| **336f8**0a5f62728eaa9654a5e53f3727c | 0.053,0.040,1.360,-1.418,-0.378,-0.353 | 0.070,0.037,0.630,-0.446,-0.209,1.018 | 0.087,0.015,-1.503,1.921,1.725,1.180 | 0.023,0.032,-1.205,0.977,-0.757,1.392 |
| **338f5**a5b617b3d4a347275687bd3435c | 0.057,0.084,1.822,-1.488,-0.902,-1.196 | 0.040,0.095,1.122,-1.876,-0.922,-0.308 | 0.020,0.032,-1.370,0.725,-0.507,0.889 | 0.070,0.026,-1.448,1.181,1.910,-0.594 |
| **3395f**45c5916261980a75f825825b3c5 | 0.046,0.013,1.539,-1.952,-0.836,-0.195 | 0.083,0.045,0.587,-0.614,-0.962,0.436 | 0.037,0.026,-1.222,1.179,0.752,0.742 | 0.096,0.040,-0.585,0.362,0.929,1.101 |
| **33ac5**abe82735c3b52b6b96fc3534d52 | 0.045,0.027,1.410,-1.774,-1.822,-0.017 | 0.053,0.059,0.427,-0.282,-1.038,-0.621 | 0.046,0.048,-0.943,0.801,-0.450,1.072 | 0.086,0.073,-0.114,-0.469,1.477,1.801 |
| **33d94**43281028db613523baf35007c25 | 0.053,0.092,1.785,-1.835,-0.998,-1.316 | 0.086,0.040,0.498,-0.189,-1.529,-1.155 | 0.047,0.047,-0.837,0.913,-0.020,0.873 | 0.071,0.062,-0.200,-0.216,1.614,1.897 |
| **33e6d**719662eefc34e45d07cdd804ad1 | 0.081,0.064,1.743,-1.659,-1.669,-0.356 | 0.083,0.044,0.974,-1.419,1.579,-1.699 | 0.027,0.077,-1.941,1.632,-0.190,1.403 | 0.056,0.075,-0.949,0.501,1.668,1.346 |
| **34432**b5a787388aba2bcb36e64229ceb | 0.050,0.093,1.745,-1.149,-0.549,-1.956 | 0.091,0.017,1.846,-1.215,-1.075,-1.136 | 0.092,0.073,-1.820,1.851,1.864,0.139 | 0.078,0.060,-0.196,-0.107,0.793,0.232 |
| **34672**82297685bdbb70efbcfd43c44a4 | 0.033,0.069,1.572,-1.607,-0.500,-0.590 | 0.087,0.056,0.658,-0.884,-1.294,0.250 | 0.051,0.029,-0.852,0.601,-0.449,0.123 | 0.072,0.090,-0.226,-0.208,1.125,1.086 |
| **34871**5015464fd0efa96be86666fddd4 | 0.039,0.082,1.434,-1.240,-1.283,-1.360 | 0.045,0.045,0.681,-0.310,-0.418,-1.694 | 0.073,0.059,-0.738,1.342,-0.481,0.410 | 0.017,0.070,-0.431,0.134,1.882,1.755 |
| **349e5**f2c785568f7ecedcd626f752135 | 0.081,0.014,1.660,-1.877,-0.498,-0.059 | 0.076,0.062,1.059,-1.700,-0.974,-1.592 | 0.011,0.069,-0.438,0.240,1.498,0.505 | 0.086,0.068,-0.404,0.204,1.894,0.403 |
| **34a3e**86c5d58b4e3d983379201a98139 | 0.052,0.048,0.979,-0.973,0.086,-1.144 | 0.042,0.081,0.754,-1.120,1.266,-1.160 | 0.040,0.059,-1.382,1.096,-0.607,0.603 | 0.095,0.066,-0.335,0.245,0.798,-0.075 |
| **35024**266330f7af5c1e1b6d5430a49e9 | 0.022,0.046,1.942,-1.924,-1.890,-0.435 | 0.040,0.065,0.906,-1.500,-1.175,-1.069 | 0.095,0.099,-1.050,0.209,1.185,0.768 | 0.062,0.060,-0.915,0.613,1.554,1.515 |
| **3508d**ea86db64c35b4e61ddf41952427 | 0.047,0.030,1.559,-1.532,-1.290,0.409 | 0.080,0.023,0.975,-1.055,-0.875,-1.003 | 0.064,0.022,-0.452,0.151,1.407,0.397 | 0.058,0.075,-0.200,-0.262,1.987,1.229 |
| **35175**33a2316a6e7480a771bce22fbee | 0.059,0.082,1.324,-1.006,-1.344,-0.715 | 0.062,0.024,1.202,-1.017,-0.933,-1.645 | 0.098,0.038,-1.937,1.666,1.900,0.801 | 0.012,0.078,-0.595,0.210,-0.982,1.604 |
| **353f9**4a7156e3a5f482d63a00ff95299 | 0.082,0.069,1.526,-1.146,-1.745,0.079 | 0.037,0.036,1.007,-1.354,-0.440,-1.641 | 0.054,0.057,-1.307,0.685,1.804,0.427 | 0.017,0.091,-1.025,0.792,1.512,0.859 |
| **35dff**a7ca18bb129b1f4732d0bcaa09f | 0.012,0.076,0.945,-1.000,0.150,-1.272 | 0.018,0.087,0.518,-0.282,0.403,-0.064 | 0.077,0.089,-0.888,1.703,1.848,1.705 | 0.036,0.057,-0.561,0.345,0.682,1.399 |
| **35f3c**2c384f56b14ba9b965d15cfda9b | 0.053,0.038,1.117,-0.985,-1.635,-0.139 | 0.014,0.079,0.877,-1.306,-0.846,-1.009 | 0.100,0.012,-0.692,0.504,0.679,0.856 | 0.039,0.093,-1.608,1.364,-0.713,1.787 |
| **35fcf**3b3f8104b833e6932d36b7f416c | 0.071,0.042,1.727,-1.788,-0.546,-0.644 | 0.043,0.068,0.512,-0.630,-1.086,-0.460 | 0.046,0.073,-0.671,0.353,-0.200,0.373 | 0.073,0.035,-0.482,0.326,-0.025,1.568 |
| **36062**44940101eebb37f8fd5ff705a86 | 0.049,0.037,0.842,-0.835,-0.784,-0.195 | 0.029,0.074,0.324,-0.279,-0.551,0.123 | 0.086,0.038,-1.220,1.543,-0.534,1.026 | 0.040,0.087,-1.025,0.786,0.639,1.013 |
| **36569**fed5588c38329bb2924d54eb0aa | 0.020,0.052,0.944,-0.856,-0.295,-1.111 | 0.051,0.025,0.211,-0.146,-0.760,-0.832 | 0.060,0.094,-0.412,-0.146,1.899,1.867 | 0.092,0.022,-0.463,0.400,-0.138,1.776 |
| **36b89**5561aee8d137d2164f71adbae03 | 0.050,0.055,1.634,-1.744,-0.550,-0.988 | 0.092,0.026,0.822,-0.768,-1.793,0.843 | 0.095,0.059,-1.196,0.766,1.237,1.225 | 0.071,0.072,-0.917,0.512,1.847,1.484 |
| **370e0**b11f6ef21bf19ad31933dd8cc48 | 0.018,0.056,1.504,-1.300,-0.948,-0.659 | 0.020,0.020,0.136,-0.294,0.071,-1.741 | 0.012,0.062,-0.326,-0.161,0.871,1.857 | 0.092,0.040,-0.154,-0.114,1.714,-0.416 |
| **373da**29263e2e1a58bba9468fca425ac | 0.086,0.048,0.724,-0.521,-1.823,0.926 | 0.091,0.064,1.832,-1.248,-0.220,-0.590 | 0.057,0.040,-1.128,1.492,1.872,0.223 | 0.065,0.058,-0.343,-0.067,1.771,0.621 |
| **375f1**1ec6ef694335be759bdfffa94d3 | 0.044,0.042,1.292,-1.183,-1.289,-0.458 | 0.027,0.017,0.477,-0.434,-1.722,-1.000 | 0.019,0.080,-1.856,1.264,-1.114,1.891 | 0.086,0.091,-1.884,1.256,1.665,1.481 |
| **37972**6d061816af7904b356b5fc0aade | 0.048,0.050,1.456,-1.375,-1.562,0.269 | 0.023,0.061,1.654,-1.425,1.434,-0.355 | 0.073,0.085,-1.388,1.584,1.937,1.968 | 0.028,0.072,-0.796,0.473,1.920,1.704 |
| **37cda**13c13a9a53b77c30b6d4e9cbe56 | 0.078,0.043,1.410,-1.205,-0.758,-1.022 | 0.019,0.045,0.922,-1.471,0.686,-1.776 | 0.016,0.056,-1.141,0.773,1.859,1.938 | 0.086,0.066,-0.751,0.371,1.714,1.623 |
| **37efe**8e9a04d606b5fa7d5027a57b521 | 0.077,0.027,1.574,-1.189,-1.657,-0.863 | 0.052,0.033,0.937,-1.658,-1.833,-1.634 | 0.097,0.070,-1.814,0.903,-0.066,1.348 | 0.050,0.091,-0.631,0.141,1.902,1.622 |
| **37f48**88a465dc0918db5c5b427feba33 | 0.096,0.013,0.995,-0.929,-1.907,-0.049 | 0.093,0.092,0.434,-0.265,-0.672,-0.303 | 0.067,0.096,-0.712,0.753,1.210,-1.244 | 0.058,0.068,-0.742,0.425,1.361,1.106 |
| **37f77**0fab2dd84281a153965b223b8b1 | 0.077,0.093,0.836,-0.701,-0.167,-1.339 | 0.010,0.063,1.413,-1.377,0.914,0.015 | 0.016,0.089,-1.604,1.756,-1.939,1.926 | 0.044,0.055,-1.228,0.988,1.525,1.164 |
| **384d3**01376dab78088838ce6ac8dd2b5 | 0.071,0.048,1.514,-0.985,-1.762,-0.191 | 0.074,0.028,1.338,-1.068,-0.971,-0.866 | 0.063,0.038,-1.445,0.920,1.812,0.826 | 0.073,0.095,-0.163,-0.245,0.091,0.839 |
| **3872e**98f50de1d4c3d5e6b8d4fd376cf | 0.049,0.068,1.587,-1.618,0.671,-1.059 | 0.024,0.083,0.903,-0.466,0.693,-0.653 | 0.069,0.033,-1.151,1.674,1.493,0.161 | 0.074,0.072,-0.582,-0.269,1.712,1.516 |
| **387af**6b99190c1bbb8a0fc20d3a7a9f4 | 0.092,0.041,1.311,-1.294,-1.331,-0.860 | 0.075,0.079,0.745,-0.937,1.814,-1.135 | 0.041,0.094,-0.990,0.786,1.764,1.685 | 0.067,0.098,-0.543,0.084,1.291,1.821 |
| **38a14**aaec459fd3ab8189b1524b2e004 | 0.013,0.062,0.936,-1.071,-0.979,-0.936 | 0.093,0.081,0.553,-0.388,-1.569,0.186 | 0.079,0.090,-1.219,0.965,0.237,1.311 | 0.025,0.099,-0.317,-0.121,1.546,1.883 |
| **38a41**41e42f916544bca02bb9963cee9 | 0.035,0.075,1.587,-1.287,-0.911,-1.094 | 0.093,0.037,0.945,-0.440,-1.207,-1.835 | 0.062,0.083,-1.347,0.499,1.157,1.772 | 0.084,0.068,-0.545,-0.225,1.140,1.973 |

| | | | | |
|---|---|---|---|---|
| **38cfa**acc0b82aeb9a2106ffb232cde13 | 0.062,0.045,1.022,-0.965,-0.611,-0.316 | 0.074,0.056,1.165,-1.250,-1.471,0.078 | 0.076,0.053,-1.066,0.594,1.992,0.246 | 0.049,0.068,-0.002,-0.283,0.759,1.119 |
| **38f08**6749dc01303d0fcf1af9526ba95 | 0.079,0.031,1.568,-1.169,-1.819,-0.119 | 0.041,0.053,0.474,-0.421,0.033,-0.283 | 0.096,0.035,-1.083,1.232,-0.104,1.628 | 0.054,0.077,-0.564,0.412,1.065,0.198 |
| **38f4d**bb35b028cca98ff9052873fb74f | 0.037,0.067,0.658,-0.526,-1.538,-1.309 | 0.045,0.084,0.287,-0.301,1.845,-0.465 | 0.063,0.061,-0.834,1.211,1.912,1.627 | 0.047,0.065,-1.888,1.591,1.582,1.323 |
| **38fb1**37b0b73303764a5b50982bf62c3 | 0.100,0.048,1.546,-1.800,-0.887,0.141 | 0.057,0.072,0.583,-0.370,-0.259,-1.800 | 0.028,0.097,-1.086,0.736,-1.220,1.640 | 0.014,0.088,-1.235,0.939,1.751,1.370 |
| **39227**cd67d68bfdc9ddbb58d2ca826aa | 0.052,0.094,1.545,-1.517,-0.957,-1.477 | 0.028,0.042,0.956,-0.684,-0.234,-1.182 | 0.095,0.057,-1.395,1.912,1.277,0.900 | 0.094,0.087,-1.179,0.624,1.650,1.386 |
| **39304**48e5d8793dd6972ee96a768b6b8 | 0.046,0.048,1.383,-1.608,-1.768,-0.255 | 0.055,0.015,0.541,-0.810,0.861,-1.757 | 0.089,0.062,-1.660,1.294,-0.134,1.722 | 0.049,0.071,-0.792,0.515,0.748,1.405 |
| **397bc**d98c11d05d0b07717d899af1b59 | 0.089,0.052,1.845,-1.588,-1.765,-0.399 | 0.031,0.061,0.745,-1.019,-1.339,-1.872 | 0.042,0.030,-1.501,0.752,1.817,1.326 | 0.045,0.069,-1.266,1.143,1.708,0.528 |
| **39a67**2a5a5389bb3dd88bd8425d4f95b | 0.042,0.079,1.693,-1.464,0.091,-1.468 | 0.090,0.080,0.563,-1.202,0.111,-0.359 | 0.063,0.086,-1.490,0.443,1.658,0.667 | 0.087,0.089,-0.987,0.317,1.981,1.933 |
| **39d03**8b496c4f94b715ef043a0b28df3 | 0.042,0.031,1.873,-1.119,-0.874,-0.395 | 0.032,0.083,0.434,0.025,-1.896,-1.001 | 0.036,0.033,-1.455,-0.018,1.723,1.031 | 0.067,0.059,-0.698,0.148,1.608,1.444 |
| **3a280**2ea19de8f411ec3db49db546044 | 0.078,0.021,1.405,-1.329,-1.945,0.418 | 0.047,0.077,0.659,-1.383,-0.651,1.071 | 0.020,0.081,-0.596,0.159,1.411,1.454 | 0.035,0.091,-1.525,1.396,1.927,0.399 |
| **3a38c**a05994ded3e6557a7b217339c44 | 0.080,0.034,0.470,-0.385,-0.642,0.060 | 0.053,0.078,0.323,-0.869,-0.601,0.064 | 0.087,0.047,-1.548,0.488,0.620,1.756 | 0.068,0.092,-0.192,-0.045,1.246,0.900 |
| **3a56b**cc75131cd7748d67e9be4dfa343 | 0.080,0.086,1.876,-1.297,-1.969,-0.678 | 0.020,0.071,0.772,-1.130,-1.063,-0.168 | 0.043,0.025,-1.422,0.780,1.200,0.754 | 0.092,0.080,-1.399,0.830,1.818,1.256 |
| **3aa03**42ea703e33639bbdbc26f5a2cb2 | 0.085,0.056,0.513,-0.265,-0.717,-0.452 | 0.078,0.069,1.229,-1.814,-0.752,-0.025 | 0.042,0.034,-1.706,1.010,1.360,0.604 | 0.083,0.044,-0.146,-0.215,1.560,1.229 |
| **3aaed**f8e087f21f6f761680e28bd82e5 | 0.076,0.017,0.616,-0.489,-1.836,0.992 | 0.036,0.082,0.372,-0.423,-1.954,0.731 | 0.081,0.088,-1.544,1.240,-0.236,1.469 | 0.059,0.091,-1.957,1.590,1.970,1.536 |
| **3ac5f**02496e23d34f4da3ed6d32f1715 | 0.081,0.016,0.980,-0.804,-1.781,0.938 | 0.015,0.053,0.285,-0.311,-1.531,0.402 | 0.085,0.036,-1.031,1.004,0.554,1.638 | 0.047,0.059,-0.577,0.433,-0.651,1.428 |
| **3adab**4b056c8e887376676c41cfca72c | 0.088,0.071,1.027,-0.684,-1.177,-0.996 | 0.019,0.052,1.927,-1.226,0.379,-1.952 | 0.098,0.039,-1.231,1.940,-0.977,1.599 | 0.089,0.088,-0.391,-0.497,1.240,1.771 |
| **3ade2**67e8a202e0e6ad59529d68efed8 | 0.057,0.063,1.725,-1.060,-1.210,-1.050 | 0.071,0.078,0.461,-0.337,-1.756,-0.993 | 0.066,0.088,-1.137,0.296,1.073,-0.118 | 0.088,0.016,-0.543,0.270,1.850,-0.165 |
| **3ae11**e5d3fd0beab1a4bf72662a1723b | 0.054,0.017,0.905,-0.938,-1.335,-0.197 | 0.081,0.028,0.399,-0.570,0.160,-0.411 | 0.034,0.074,-0.660,0.400,1.766,0.737 | 0.038,0.022,-1.229,1.157,1.735,0.731 |
| **3ae17**57f9f5d5ebbac2e02041425afde | 0.079,0.080,1.717,-1.800,0.691,-1.053 | 0.093,0.044,0.645,-1.836,-0.127,-0.321 | 0.030,0.080,-1.472,0.359,1.078,1.908 | 0.084,0.081,-1.303,0.775,1.759,1.763 |
| **3ae38**b697ebda3a8b8179e6005b677f2 | 0.057,0.082,0.985,0.944,-1.695,-1.383 | 0.065,0.025,0.494,-0.071,-1.574,-0.835 | 0.098,0.088,-1.125,0.315,1.005,-0.483 | 0.092,0.018,0.028,-0.280,0.691,-0.115 |
| **3ae73**11c178361901f3d1fd0c2c1b19f | 0.081,0.023,1.751,-1.440,-1.530,-0.927 | 0.052,0.071,0.750,-1.801,-0.689,-0.323 | 0.065,0.070,-1.822,0.805,1.240,-0.498 | 0.094,0.047,-0.342,0.026,1.566,0.789 |
| **3b09f**89aefc8abc903213a471c961990 | 0.074,0.066,1.843,-1.921,-0.509,-1.566 | 0.054,0.045,1.129,-0.980,-1.913,-1.439 | 0.094,0.023,-1.436,1.245,1.793,0.693 | 0.088,0.055,-0.091,-0.303,1.731,1.141 |
| **3b394**4a60e3b9d67767470bd936bb083 | 0.097,0.057,0.508,0.093,-1.556,-1.137 | 0.024,0.073,1.418,-1.861,-1.529,-0.929 | 0.040,0.022,-1.947,1.432,1.191,1.931 | 0.084,0.077,-0.317,-0.264,1.707,1.569 |
| **3b8cd**f59702de41d2b8f613c4036ad9d | 0.060,0.012,1.637,-1.811,-1.587,0.610 | 0.021,0.063,0.780,-1.242,0.401,-0.483 | 0.031,0.029,-0.536,0.371,-0.270,1.689 | 0.078,0.086,-0.550,0.378,0.956,0.675 |
| **3baf8**94cf611a61b243ecd343cc58f6e | 0.078,0.053,1.385,-1.102,-1.786,0.450 | 0.063,0.082,0.764,-1.030,0.482,-1.398 | 0.047,0.071,-1.443,0.823,-0.015,0.532 | 0.085,0.031,-0.537,0.086,1.753,0.859 |
| **3bbc5**3b9ff4311df306534a99ba043a1 | 0.040,0.012,1.562,-1.791,-1.925,-0.037 | 0.070,0.020,0.263,-0.243,-0.902,-0.327 | 0.080,0.090,-1.641,0.888,0.010,0.693 | 0.097,0.052,-0.239,0.011,1.960,-1.390 |
| **3bcf1**b5480bf611125f4d09739cf1788 | 0.099,0.060,1.524,-1.459,-0.278,-0.632 | 0.072,0.091,1.144,-0.580,1.904,-1.510 | 0.075,0.065,-0.994,1.683,1.882,0.700 | 0.049,0.060,-1.419,0.779,1.775,0.869 |
| **3bd1b**41be8fc93a2cc45cffbe218aae3 | 0.074,0.025,1.961,-1.824,-1.828,-0.719 | 0.052,0.066,1.342,-1.895,-1.280,1.155 | 0.076,0.062,-1.914,1.407,-0.152,1.626 | 0.092,0.067,-0.925,0.643,-0.545,1.882 |
| **3bf54**fb1a347580cd99cf797486e181b | 0.089,0.012,1.627,-1.508,-1.899,-0.417 | 0.072,0.014,0.759,-0.638,-1.035,0.205 | 0.015,0.083,-1.027,1.486,-0.307,-0.903 | 0.082,0.054,-1.135,0.723,1.909,0.964 |
| **3c22d**9f512d7be1b0fd589d7f4949319 | 0.021,0.017,0.857,-0.694,-1.421,-0.234 | 0.098,0.015,1.308,-1.855,-0.466,1.545 | 0.067,0.041,-1.855,1.254,-0.508,1.877 | 0.054,0.060,-1.155,0.813,-0.362,1.420 |
| **3c2d6**68d6648ba37252be1c19b14f297 | 0.096,0.060,1.155,-0.704,-1.820,-0.633 | 0.090,0.065,1.540,-1.799,0.305,-1.213 | 0.089,0.082,-1.919,1.424,1.928,-0.642 | 0.050,0.065,-0.591,0.003,1.959,1.887 |
| **3c405**d30b5aff886f51118f3c72ff5f4 | 0.095,0.021,1.626,-1.598,-1.344,0.128 | 0.075,0.033,1.382,-1.557,-0.785,-0.865 | 0.093,0.051,-1.953,1.418,1.244,1.736 | 0.098,0.081,-1.237,0.546,1.963,1.632 |
| **3c5b6**a1835138b696e50ee40cd5c7393 | 0.072,0.017,1.361,-1.298,-1.726,0.176 | 0.015,0.036,0.564,-1.365,-0.871,0.165 | 0.088,0.047,-1.838,1.169,-0.207,-0.303 | 0.050,0.089,-1.259,1.068,0.799,1.086 |
| **3c777**0396ba0d59b4a2ea2cd2362347f | 0.064,0.039,0.677,-0.158,-1.016,0.082 | 0.074,0.078,1.180,-1.339,-0.367,-0.954 | 0.053,0.082,-1.343,0.162,1.894,0.393 | 0.046,0.061,-0.225,-0.151,1.904,0.087 |
| **3c7eb**7329dabe9ba33041ef273b037fa | 0.022,0.084,1.807,-1.653,-1.745,-0.563 | 0.083,0.020,0.702,-1.042,-1.574,-0.061 | 0.030,0.087,-1.914,1.011,1.606,1.989 | 0.055,0.046,-1.314,1.128,1.838,1.116 |
| **3ca10**f9d0ffd3805ac35d09ba852dbd7 | 0.094,0.011,1.644,-1.560,-1.728,-0.118 | 0.077,0.086,1.286,-0.805,0.304,-1.759 | 0.033,0.086,-1.274,1.615,-0.012,1.023 | 0.090,0.045,-1.130,0.671,1.519,1.456 |
| **3cb7e**a7a4a8fec4afae387338796856d | 0.089,0.023,1.763,-1.315,-1.345,-0.172 | 0.052,0.055,0.933,-1.427,-1.594,-0.935 | 0.071,0.057,-0.765,-0.048,1.793,1.016 | 0.050,0.041,-0.089,-0.088,1.810,0.337 |
| **3cc53**381c199e9f9c507a598980949ff | 0.086,0.098,1.931,-1.561,-0.233,-1.279 | 0.027,0.096,1.676,-1.051,-1.951,-1.757 | 0.040,0.039,-1.347,1.001,1.827,1.551 | 0.099,0.020,-0.354,-0.279,1.837,1.874 |
| **3d1d1**90c98cbe0544d1098937748713e | 0.085,0.015,0.772,-0.715,-1.848,0.167 | 0.026,0.061,0.288,-0.249,-1.166,-0.102 | 0.089,0.059,-0.322,0.461,-0.683,0.038 | 0.082,0.097,-0.637,0.289,0.685,1.269 |
| **3d25f**0178e19302f39c63200a284baa3 | 0.084,0.083,1.643,-1.848,0.046,-0.467 | 0.079,0.015,1.611,-0.775,-1.854,-1.582 | 0.092,0.096,-1.344,1.492,1.214,1.295 | 0.087,0.092,-0.304,-0.298,1.946,0.914 |
| **3d42f**420a281e3fadae0798f96db457f | 0.065,0.071,1.733,-1.395,-1.749,0.535 | 0.024,0.077,0.505,-0.397,-1.477,-0.138 | 0.089,0.092,-1.529,0.525,1.881,0.431 | 0.012,0.023,-0.708,0.632,1.683,0.426 |
| **3d9c3**8d394bd0be8617e8c1c2bee1fbc | 0.040,0.018,1.515,-1.619,-1.001,-0.189 | 0.083,0.076,1.703,-1.277,-0.885,-1.610 | 0.037,0.036,-1.416,1.230,1.223,0.381 | 0.077,0.100,-0.996,0.254,1.538,0.995 |
| **3dbb5**5a85e69a586108aad1bdc1a90d0 | 0.070,0.024,1.460,-1.749,-0.708,-0.145 | 0.052,0.017,0.583,-0.738,-0.538,0.425 | 0.081,0.087,-1.124,0.997,0.915,0.894 | 0.085,0.040,-0.551,0.428,1.843,-0.759 |
| **3dc84**15bea84b2a609c6661536263d76 | 0.095,0.082,1.772,-1.090,-1.840,-0.865 | 0.026,0.054,0.923,-1.370,-0.500,0.064 | 0.047,0.083,-0.850,0.296,1.853,1.033 | 0.056,0.078,-0.285,-0.179,0.711,1.520 |
| **3dd6e**8a18725a736637b395d14059d5e | 0.031,0.070,1.193,-1.110,-0.373,-0.490 | 0.045,0.079,0.416,-0.738,-0.293,0.231 | 0.078,0.085,-1.878,0.906,1.499,0.974 | 0.095,0.075,-1.810,1.271,1.754,0.401 |
| **3de4f**248de83c1eb8268d8d12d393c6e | 0.084,0.082,1.588,-1.452,-1.835,0.691 | 0.072,0.088,1.673,-1.185,-1.406,-1.300 | 0.034,0.037,-1.630,1.323,1.738,1.964 | 0.086,0.095,-0.721,-0.220,1.943,1.420 |
| **3e22e**f26dd71cb0b5089f1b93ce665e8 | 0.040,0.050,0.828,-0.878,-0.450,-0.435 | 0.075,0.056,0.928,-1.134,0.685,-0.857 | 0.077,0.086,-1.880,1.900,1.290,0.936 | 0.055,0.079,-0.586,0.458,-0.725,1.295 |
| **3ea21**bb0a19937daf3f9d581a9777a5b | 0.026,0.072,0.758,-0.735,-0.367,-0.519 | 0.023,0.038,0.183,-0.135,1.001,-0.238 | 0.096,0.062,-0.825,1.659,-0.686,0.774 | 0.079,0.046,-1.341,0.970,1.126,1.658 |
| **3ea5e**292f0f13bd0225205c6aa782a7f | 0.044,0.017,1.800,-1.607,-1.896,-0.054 | 0.013,0.016,1.306,-1.236,-0.993,1.000 | 0.050,0.092,-0.687,0.885,1.644,-0.329 | 0.067,0.058,-0.446,0.115,1.765,0.371 |
| **3eacb**05a43be53dcb230194acc1a0c3d | 0.096,0.058,1.561,-1.519,-1.629,0.789 | 0.087,0.079,0.537,-0.457,-0.609,-0.194 | 0.082,0.086,-1.307,1.014,1.827,1.050 | 0.065,0.092,-1.486,1.042,0.212,1.948 |
| **3eafb**8cfa859213e0316bcd587a30d89 | 0.077,0.027,1.875,-1.574,-1.168,-0.885 | 0.067,0.089,1.070,-1.792,-0.300,-1.282 | 0.076,0.036,-1.789,0.907,1.209,-0.021 | 0.097,0.040,-0.928,0.604,1.823,0.998 |
| **3eb18**3edc786c4b61f4ef73f1e6962db | 0.036,0.032,0.796,-0.731,-1.774,0.016 | 0.033,0.075,0.900,-0.709,0.590,0.294 | 0.084,0.064,-1.291,1.945,1.341,0.485 | 0.098,0.063,-1.356,0.725,1.360,1.454 |
| **3f5bc**d702c7d49094c5cfc4d4b420d0e | 0.078,0.024,1.372,-1.427,-0.749,-0.399 | 0.080,0.089,1.131,-1.248,-1.873,-1.199 | 0.043,0.047,-1.516,0.916,1.274,0.815 | 0.094,0.051,-0.233,-0.077,1.897,1.034 |
| **3f826**8cc5e87760b3c7843dee1dd2bf6 | 0.047,0.025,1.513,-1.665,-0.547,-0.529 | 0.097,0.015,1.364,-1.620,-1.319,1.132 | 0.099,0.088,-1.641,0.949,1.862,-0.061 | 0.086,0.071,-0.374,-0.265,1.957,1.554 |
| **3f841**75b881e552b20236f62fa287d1a | 0.078,0.048,1.141,-1.261,-0.953,-0.157 | 0.017,0.081,0.587,-0.393,-1.101,-0.722 | 0.076,0.098,-1.005,1.314,0.819,0.153 | 0.045,0.073,-1.626,1.361,0.481,1.699 |
| **3f86c**242db3bc4b1ddc019e2606231eb | 0.092,0.063,1.366,-1.169,-1.828,0.682 | 0.025,0.022,0.299,-0.166,-0.371,-0.118 | 0.098,0.059,-1.024,1.241,1.174,0.569 | 0.028,0.039,-0.602,0.209,1.930,1.852 |
| **3f8f6**326cfaaea1f3c78832f91ed6866 | 0.013,0.039,1.256,-1.148,-0.609,-1.670 | 0.014,0.089,0.996,-1.826,-0.482,-0.448 | 0.096,0.097,-1.916,0.686,0.582,1.923 | 0.089,0.074,-0.687,0.304,1.807,1.091 |
| **3fa51**47c8e6695773b2f0e9bf3f4a4bf | 0.098,0.042,1.661,-1.446,-1.942,-0.043 | 0.046,0.076,1.333,-1.941,-0.038,0.133 | 0.015,0.060,-1.076,0.475,1.826,1.810 | 0.088,0.063,-0.947,0.394,1.784,1.521 |
| **3fadf**08a9c3e902d31989032366bac35 | 0.064,0.051,0.760,-0.757,-0.121,-0.446 | 0.024,0.082,0.572,-0.659,0.768,-0.699 | 0.076,0.057,-1.652,1.564,1.298,1.972 | 0.092,0.058,-1.554,1.321,0.626,1.200 |
| **3fbaa**adf068c7a2e5c379027788ff19f | 0.045,0.036,1.105,-0.714,-1.187,-0.380 | 0.075,0.041,0.717,-1.708,-0.897,-1.164 | 0.071,0.094,-1.792,0.308,-0.132,1.197 | 0.072,0.046,-1.790,1.358,1.822,1.136 |
| **3fcc3**8e75c4c64e8d7ac509dc11e2db6 | 0.075,0.078,0.858,-0.739,-0.838,-0.351 | 0.041,0.053,0.310,-0.078,-0.800,-1.109 | 0.057,0.015,-1.589,1.779,1.814,1.398 | 0.089,0.049,-0.041,-0.346,1.960,0.179 |
| **3fd54**e1e4e346939f37fc75f3574f5b3 | 0.043,0.062,1.314,-1.296,-1.072,-0.729 | 0.086,0.057,1.980,-1.052,-1.776,1.753 | 0.097,0.088,-1.160,1.912,1.953,-0.260 | 0.055,0.050,-1.040,0.673,1.821,-0.063 |
| **4021f**905cfedb90102e58f335cb5f4a5 | 0.078,0.047,1.435,-1.520,-0.176,-0.569 | 0.024,0.052,1.526,-1.257,-0.730,-1.792 | 0.067,0.064,-1.917,1.966,0.839,1.510 | 0.098,0.054,-0.591,0.041,1.881,0.704 |
| **40230**be990d2b7d57eee3eec12bb925c | 0.063,0.043,0.773,-0.440,-1.863,-0.899 | 0.013,0.079,0.607,-0.544,0.766,-0.249 | 0.073,0.065,-1.276,1.355,1.875,0.121 | 0.045,0.060,-1.066,0.715,1.085,1.728 |
| **40305**608ab46b35dcf8c2fd79d04c1ce | 0.076,0.048,1.055,-1.245,-1.563,1.155 | 0.054,0.024,0.866,-0.758,-1.379,-1.056 | 0.082,0.045,-0.784,0.469,1.646,1.303 | 0.078,0.041,-0.040,-0.202,1.351,0.427 |
| **403ab**3cb43e68f75ff95751a64d2e90d | 0.064,0.080,0.342,-0.341,-1.799,1.142 | 0.078,0.063,0.417,-0.313,1.506,-1.140 | 0.042,0.084,-0.608,1.613,-0.277,0.545 | 0.047,0.086,-1.084,0.694,1.291,1.280 |
| **404cd**0ea0ef354af42cb2ac51024f66b | 0.057,0.040,1.103,-1.087,-0.510,-0.912 | 0.061,0.040,1.843,-1.091,-1.717,1.604 | 0.040,0.053,-1.142,1.481,1.277,1.739 | 0.082,0.066,-1.092,0.380,1.994,0.120 |
| **405ff**020ba7caa28b1b73b1e7c7e5dd0 | 0.023,0.086,1.698,-1.643,-1.701,-1.117 | 0.092,0.052,1.980,-1.592,-0.078,-0.000 | 0.079,0.019,-1.353,1.908,-0.102,1.984 | 0.045,0.043,-0.640,0.310,1.876,0.835 |

| | | | | |
|---|---|---|---|---|
| **407932**c8086ba74b94eb39a892e55832 | 0.048,0.035,1.600,-1.863,-0.317,-0.413 | 0.095,0.021,0.429,-0.895,0.574,0.146 | 0.022,0.088,-1.149,0.482,-0.233,1.489 | 0.051,0.045,-0.897,0.563,1.908,1.445 |
| **408173**a69f1c841578608e452deaccc1 | 0.090,0.066,1.737,-1.873,-0.664,-0.504 | 0.043,0.082,1.480,-0.986,-1.373,-1.809 | 0.057,0.099,-1.711,1.612,-1.551,1.892 | 0.046,0.041,-0.557,0.308,0.691,1.533 |
| **4088ed**a54e78ef6cb21dbc340b8a6109 | 0.085,0.020,1.533,-1.318,-1.876,-0.098 | 0.100,0.012,1.363,-1.425,-1.512,-0.298 | 0.099,0.060,-1.401,0.830,-0.711,1.675 | 0.059,0.081,-0.711,0.090,1.183,1.825 |
| **40d39b**f48fa87e93331726bddd278430 | 0.093,0.016,1.614,-1.361,-0.889,-0.865 | 0.088,0.062,0.971,-1.857,-1.894,-0.226 | 0.072,0.066,-1.845,0.861,1.694,-0.496 | 0.013,0.072,-1.200,0.958,1.494,1.425 |
| **40ece**211863d94aa45133db43bdb7f88 | 0.099,0.069,1.794,-1.661,-0.716,-1.076 | 0.070,0.033,1.296,-1.038,-1.216,-1.489 | 0.068,0.063,-1.653,1.982,1.346,0.940 | 0.027,0.079,-1.495,1.232,1.786,1.030 |
| **41376**ba3cebae7dec979c75b64ccd053 | 0.013,0.063,1.539,-1.808,0.519,-0.846 | 0.093,0.022,0.506,-0.204,-0.660,-1.152 | 0.065,0.097,-1.126,1.924,0.304,-0.956 | 0.056,0.038,-1.422,1.380,0.873,-0.357 |
| **4155f**b3a3628c732f22c22d9f9a8d04f | 0.042,0.066,1.037,-0.866,-0.785,-1.200 | 0.017,0.097,0.809,-0.693,0.515,-0.517 | 0.090,0.037,-0.922,0.991,-0.181,0.993 | 0.091,0.072,-0.505,-0.173,1.521,1.669 |
| **416168**fa2ae09b13c060eb2236a1fc9a | 0.046,0.046,1.922,-1.383,-1.603,-1.691 | 0.026,0.087,1.087,-1.905,0.734,0.102 | 0.015,0.087,-0.474,-0.153,1.154,1.864 | 0.018,0.093,-1.393,0.876,1.686,1.767 |
| **417be**898f814fa6ce54ccecfd9a53f1e | 0.078,0.071,1.611,-1.132,-1.679,-0.516 | 0.025,0.064,0.756,-1.108,-1.078,-0.973 | 0.021,0.035,-1.421,0.630,-0.727,1.678 | 0.018,0.092,-0.109,-0.208,0.488,1.425 |
| **417e0**32a2acd5dbc01507ffe470dcf89 | 0.061,0.056,1.691,-1.740,-0.636,-0.411 | 0.079,0.016,0.997,-1.398,-1.476,0.038 | 0.048,0.077,-1.109,0.750,-0.763,0.946 | 0.022,0.069,-1.383,1.119,1.345,1.433 |
| **419d6**a4e56ad8f00f7da26f5e34f18be | 0.064,0.041,1.815,-1.518,-1.975,-0.208 | 0.049,0.022,0.278,-0.338,0.022,-1.263 | 0.019,0.031,-1.152,0.976,1.916,0.713 | 0.046,0.028,-0.477,0.319,0.849,1.486 |
| **41bcd**2e82fba1775576e9d53d3f89c76 | 0.086,0.035,0.733,-0.628,-1.903,1.212 | 0.090,0.062,0.654,-0.464,-1.146,0.779 | 0.045,0.086,-0.830,1.131,-0.957,0.440 | 0.093,0.042,-1.159,0.649,1.664,1.787 |
| **41e41**a937d4087618b16cc6dfc2d816d | 0.096,0.019,1.514,-1.162,-1.938,0.463 | 0.077,0.063,0.588,-0.702,-0.707,-0.532 | 0.038,0.086,-1.794,1.222,-0.289,-0.124 | 0.078,0.075,-0.310,-0.030,1.287,0.695 |
| **41f836**c0224ffc8ddc58dfe9ee5e4835 | 0.073,0.012,0.676,-0.636,-0.654,0.884 | 0.025,0.084,1.905,-0.829,-1.078,-0.386 | 0.099,0.082,-1.043,1.926,1.754,-0.423 | 0.067,0.021,-0.219,0.079,0.708,-0.512 |
| **4204**289bb21ef0102b1a9a08eb4a40da | 0.080,0.036,1.601,-0.968,-1.261,-1.559 | 0.059,0.046,1.217,-1.830,-0.905,-0.905 | 0.065,0.016,-1.775,0.910,0.796,0.800 | 0.023,0.052,-0.509,0.351,1.681,0.678 |
| **4216a**802eb0001263544a8140182c028 | 0.076,0.047,0.935,-0.736,-0.996,-0.396 | 0.042,0.060,0.471,-0.925,-0.201,1.014 | 0.016,0.085,-0.809,0.512,-1.441,1.674 | 0.032,0.035,-0.309,0.136,1.899,1.140 |
| **4230e**c182e72e5ab9bf595581f5e6763 | 0.070,0.012,1.974,-1.323,-1.987,-0.989 | 0.027,0.041,0.575,-0.856,-0.807,-1.095 | 0.031,0.030,-1.901,0.673,1.910,1.426 | 0.098,0.088,-0.744,0.023,1.964,0.473 |
| **42398**c83d7c45c7e283541746b59c0ae | 0.062,0.060,1.589,-1.871,-1.873,1.486 | 0.053,0.091,0.305,-0.254,-0.790,-0.377 | 0.028,0.098,-0.220,-0.084,1.650,0.770 | 0.023,0.090,-1.270,1.053,1.913,0.892 |
| **42942**d0ed32b455586b6c3887c6fa4ec | 0.098,0.086,0.428,-0.125,-1.145,0.173 | 0.051,0.084,0.506,-1.002,-0.610,0.944 | 0.078,0.040,-0.727,0.434,0.040,0.572 | 0.042,0.099,-1.420,0.928,1.902,1.266 |
| **42b29**c83c9195ed794e8660ced5c402f | 0.072,0.058,1.664,-1.839,-0.952,-1.078 | 0.097,0.046,1.165,-1.278,0.590,-1.416 | 0.097,0.040,-1.856,1.760,1.813,1.258 | 0.054,0.028,-1.431,1.279,1.741,0.398 |
| **42bbc**ac265dc9c9595931cad536ac3d1 | 0.094,0.064,1.916,-1.692,-1.107,0.054 | 0.064,0.043,0.157,-0.964,0.906,0.325 | 0.062,0.043,-1.857,0.347,1.910,1.734 | 0.095,0.071,-0.470,0.136,1.936,0.143 |
| **42f4d**c000c6af207defbee9a9f66cd2a | 0.074,0.020,0.751,-0.577,-1.767,-0.938 | 0.045,0.078,1.109,-1.529,-0.563,-0.783 | 0.091,0.083,-1.626,1.213,-0.135,0.724 | 0.070,0.014,-0.674,0.553,1.012,1.030 |
| **43193**fc1ae288901ee15d4a3dfd67272 | 0.017,0.021,1.323,-1.475,-0.716,-0.558 | 0.046,0.044,0.941,-0.616,-1.591,-0.851 | 0.058,0.054,-1.750,1.353,-0.213,1.338 | 0.020,0.068,-0.981,0.609,1.535,1.084 |
| **431ea**5aab7bf1d7c28a1a22008a242ff | 0.047,0.048,1.437,-1.311,-0.592,-0.999 | 0.073,0.088,0.478,-0.160,-1.036,-1.031 | 0.078,0.089,-0.822,0.504,-0.607,0.508 | 0.057,0.088,-0.336,0.287,-0.394,0.414 |
| **434fa**3d9586b7078a8d3abcc419d9a6f | 0.080,0.028,1.248,-1.103,-1.743,1.115 | 0.079,0.019,0.710,-0.616,-0.799,-0.769 | 0.046,0.093,-1.000,0.753,-0.481,0.781 | 0.021,0.057,-1.415,1.108,1.627,1.244 |
| **4351a**776c73904b251e9b08900848b89 | 0.087,0.084,1.899,-1.854,-0.337,-0.968 | 0.048,0.085,1.113,-0.606,0.217,-1.518 | 0.030,0.090,-0.842,0.843,0.696,0.929 | 0.052,0.043,-0.398,-0.064,1.565,1.388 |
| **4359b**e6e6313c8f8486ff8bf7d855a2f | 0.088,0.075,0.804,-0.890,-0.121,-0.350 | 0.089,0.061,0.931,-0.762,-0.759,-0.199 | 0.065,0.043,-1.445,1.632,1.865,0.555 | 0.063,0.074,-1.399,1.177,-0.168,1.519 |
| **43935**e45a67cd6cf5290657295017217 | 0.046,0.010,1.616,-1.625,-1.738,-0.050 | 0.039,0.084,0.547,-0.367,-0.551,-1.772 | 0.074,0.018,-1.524,0.721,1.243,0.778 | 0.085,0.084,-0.431,-0.180,1.621,1.676 |
| **43ac9**1b4c897f23f7283a3dc21af5644 | 0.042,0.076,0.718,-0.572,-0.826,-0.563 | 0.091,0.050,0.910,-1.000,1.453,-0.130 | 0.082,0.082,-0.469,0.000,1.634,1.529 | 0.041,0.059,-0.054,-0.047,1.580,-0.435 |
| **43aca**be46cabeb8f0687ba697527c68e | 0.093,0.028,0.965,-0.791,-1.302,-0.362 | 0.025,0.084,1.894,-1.060,-1.698,0.937 | 0.097,0.096,-1.444,1.931,1.859,0.228 | 0.075,0.045,-1.541,1.020,0.483,0.783 |
| **43c8b**f3bf45dc40f4acf03baa3e78c2b | 0.045,0.049,1.720,-1.320,-1.809,-0.637 | 0.098,0.073,0.407,-0.621,-0.623,0.099 | 0.059,0.027,-1.895,0.459,1.775,1.789 | 0.087,0.040,-1.534,1.188,0.494,1.818 |
| **43d2f**7a7dea966af0058db529bb910f4 | 0.045,0.061,1.873,-1.800,-1.673,-0.123 | 0.019,0.055,0.590,-0.777,0.989,-1.583 | 0.070,0.064,-1.069,0.609,1.001,0.347 | 0.091,0.098,-0.379,-0.284,1.913,1.245 |
| **43d4e**b0f0eb1ce5a3b4a0959d3c6ee20 | 0.012,0.082,1.463,-1.280,-0.999,-0.854 | 0.023,0.018,0.009,-0.125,0.184,0.781 | 0.067,0.081,-0.167,-0.326,1.588,1.202 | 0.062,0.065,-0.703,0.527,-0.098,1.151 |
| **43dde**1bd2cc58265de834830c8bfd84f | 0.040,0.083,1.911,-1.604,-1.090,-1.303 | 0.085,0.058,1.771,-1.513,-1.727,-1.158 | 0.093,0.037,-1.043,0.636,1.830,1.800 | 0.063,0.097,-0.323,-0.214,1.399,1.867 |
| **441ab**af4388fe0a5c66a55b95e32b59e | 0.030,0.064,1.617,0.261,0.134,-1.170 | 0.094,0.019,0.769,-1.041,-1.077,-0.579 | 0.051,0.053,-1.609,0.125,0.655,0.053 | 0.068,0.048,-1.369,0.946,0.359,0.852 |
| **445302**b3bbdbafb17a5ceea000c4c912 | 0.038,0.063,0.996,-0.837,-0.615,-1.604 | 0.060,0.084,0.399,-0.881,-0.984,0.649 | 0.077,0.089,-0.868,0.231,1.331,1.231 | 0.092,0.094,-0.141,-0.394,1.984,1.869 |
| **446f0**082cd34cad03d098e2d97584eff | 0.055,0.041,0.835,-0.940,-0.529,-0.438 | 0.081,0.082,0.714,-0.945,-0.536,-0.471 | 0.072,0.051,-1.618,1.122,1.644,0.656 | 0.100,0.059,-0.970,0.630,1.895,0.305 |
| **447d1**9d76b8116c2d8bbe0653deb00e5 | 0.015,0.086,1.765,-1.585,-0.325,-1.341 | 0.062,0.056,0.676,-0.478,-0.471,0.052 | 0.083,0.061,-1.082,1.171,1.089,0.339 | 0.080,0.060,-0.758,0.384,0.946,0.980 |
| **44b45**d5184625a728e7382512444941d | 0.079,0.033,1.549,-1.649,-1.504,-0.505 | 0.021,0.088,0.395,-0.734,-1.898,1.043 | 0.020,0.047,-0.680,0.591,1.569,1.590 | 0.078,0.084,-1.597,1.331,1.934,0.343 |
| **44b4a**47d21729709eb7c36acffd03a32 | 0.099,0.065,0.679,-0.529,-1.734,1.244 | 0.033,0.066,0.735,-0.702,-0.853,-1.030 | 0.040,0.024,-1.133,0.788,-0.126,1.805 | 0.055,0.064,-1.049,0.542,1.673,1.374 |
| **44bc4**dd0b14abba68ac3010cfe51d9e5 | 0.043,0.036,0.808,-0.905,-1.415,0.594 | 0.038,0.024,0.536,-0.355,1.917,-1.893 | 0.089,0.027,-0.251,0.154,1.720,1.844 | 0.096,0.052,-0.358,-0.014,1.855,-0.134 |
| **44c735**a372ecb0fdbc1a8c0dab6b4fb3 | 0.051,0.030,0.423,-0.327,-0.975,-0.846 | 0.046,0.072,0.378,-0.352,0.097,0.292 | 0.028,0.043,-0.706,0.980,-0.486,0.854 | 0.084,0.053,-0.610,0.147,1.107,1.272 |
| **44d11**41d278bd6514b41b580ca4016bb | 0.017,0.026,1.758,-1.854,-1.382,-0.902 | 0.064,0.077,0.650,-0.645,-0.219,-0.241 | 0.021,0.082,-1.799,1.827,0.385,0.553 | 0.037,0.081,-1.593,1.001,1.958,1.919 |
| **44de0**481e538ed2bbcf31bcce0f98f60 | 0.076,0.020,1.511,-1.604,-1.132,0.726 | 0.064,0.054,1.419,-1.619,-0.304,-1.408 | 0.055,0.065,-0.984,0.687,1.545,0.608 | 0.093,0.071,-0.653,0.009,1.875,1.531 |
| **44e246**f80ef310891159d2c015efd530 | 0.051,0.040,1.951,-1.578,-1.911,-0.853 | 0.020,0.071,0.466,-0.214,-0.968,-0.929 | 0.065,0.085,-0.823,0.447,1.153,0.129 | 0.075,0.089,-0.246,-0.353,1.758,0.668 |
| **44e65**f7263b01630f81bd6a7305881ae | 0.096,0.045,1.672,-0.970,-1.055,-1.334 | 0.068,0.076,1.248,-1.329,-0.941,-1.507 | 0.025,0.092,-1.608,0.549,0.213,1.913 | 0.088,0.073,-1.291,0.939,0.373,1.223 |
| **44f4e**b69eed234a655eff0058fb242c1 | 0.068,0.022,1.735,-1.596,-0.958,-0.521 | 0.047,0.065,0.855,-1.075,-0.074,0.796 | 0.057,0.068,-1.876,1.441,0.528,1.902 | 0.023,0.075,-1.387,1.066,1.669,0.857 |
| **450a2**81ed1ea8c89625e826ddafb9549 | 0.051,0.014,1.694,-1.658,-1.523,-0.006 | 0.016,0.082,0.294,-0.214,-0.464,-0.793 | 0.043,0.032,-0.420,-0.002,1.038,1.237 | 0.033,0.043,-0.701,0.584,-0.182,1.018 |
| **452435**e1912bc1e55f6e5651d3bf8798 | 0.097,0.023,1.897,-1.670,-1.577,0.239 | 0.023,0.081,0.646,-1.186,-0.575,0.759 | 0.099,0.090,-1.078,0.533,1.351,-0.436 | 0.068,0.077,-1.445,0.953,0.659,1.575 |
| **452a7**c24beccac4c9b70e15502568fb7 | 0.056,0.083,1.638,-1.419,-0.448,-1.127 | 0.026,0.030,1.610,-1.613,-1.700,-0.737 | 0.033,0.074,-1.225,1.268,1.602,-0.020 | 0.045,0.054,-0.305,-0.066,1.133,1.347 |
| **453524**b91ac980f0eff274f1f74554c2 | 0.067,0.023,0.814,-0.846,-0.545,-0.373 | 0.094,0.036,0.603,-0.648,0.152,0.138 | 0.076,0.054,-1.199,1.387,1.188,-0.664 | 0.071,0.018,-0.423,0.064,1.648,0.964 |
| **455388**dd19658e951fde4b280ee90f72 | 0.078,0.057,1.519,-1.607,-1.668,0.598 | 0.014,0.070,1.093,-1.491,-1.539,-0.365 | 0.099,0.026,-1.491,0.977,1.141,1.568 | 0.052,0.023,-0.161,-0.011,1.593,1.480 |
| **456de**e5d4fcf1782c237157dfe4cebf0 | 0.094,0.061,1.549,-1.433,-0.652,-0.311 | 0.084,0.095,1.180,-0.901,0.860,-0.410 | 0.065,0.096,-1.009,1.546,0.795,0.739 | 0.070,0.071,-0.403,0.166,1.365,0.192 |
| **4576c**2e172df564167729e0c4de72183 | 0.044,0.055,1.619,-1.614,-1.290,0.040 | 0.073,0.075,0.795,-0.322,0.721,-1.683 | 0.095,0.012,-0.735,0.778,1.763,0.675 | 0.099,0.097,-1.553,0.702,0.919,1.367 |
| **457a1**f05235563dff0345e55dcac87b4 | 0.032,0.037,1.462,-0.893,-1.824,-0.144 | 0.045,0.038,0.231,-0.403,0.249,-1.840 | 0.031,0.071,-1.726,0.372,1.222,0.273 | 0.088,0.024,-1.513,1.037,1.890,0.903 |
| **4584d**41a988fb4ae861582b2f2d7df33 | 0.094,0.028,1.932,-1.599,-1.298,-0.884 | 0.045,0.095,1.264,-1.303,0.850,-1.835 | 0.043,0.082,-1.241,0.793,1.192,0.725 | 0.026,0.055,-0.060,-0.302,1.371,1.662 |
| **458cb**098b7747ee913ab5c74ed210e59 | 0.047,0.026,1.605,-1.700,-1.007,-1.378 | 0.025,0.050,1.021,-0.597,-0.522,-0.996 | 0.070,0.096,-1.742,1.959,0.424,1.448 | 0.086,0.074,-1.140,0.854,1.496,-0.431 |
| **45a0e**d646b78dccb8617b6922d1a5b96 | 0.077,0.014,1.931,-1.823,-1.226,-0.199 | 0.042,0.077,0.854,-0.410,-1.165,-1.781 | 0.046,0.070,-1.308,1.680,-0.245,1.053 | 0.059,0.045,-0.559,0.408,0.849,0.877 |
| **45db0**a3b10dff20ff6eef9bf576367af | 0.052,0.077,1.934,-1.720,-1.166,-0.539 | 0.061,0.023,0.798,-0.843,-0.104,-1.360 | 0.091,0.036,-0.590,0.300,1.161,1.180 | 0.090,0.093,-0.491,-0.150,1.787,1.286 |
| **45ef6**b6445abeed84f28fe15a0c0779f | 0.055,0.020,0.494,-0.405,-0.992,-0.404 | 0.026,0.053,1.306,-1.333,-1.937,-1.903 | 0.077,0.061,-1.820,1.441,1.627,-0.635 | 0.065,0.091,-1.225,0.899,0.331,1.459 |
| **45fac**e51450e9858a591ad18503218ee | 0.046,0.088,1.338,-1.247,-0.579,-0.819 | 0.020,0.045,1.683,-1.220,-1.058,1.033 | 0.084,0.038,-0.775,0.584,1.881,1.489 | 0.070,0.077,-0.310,-0.237,1.104,1.368 |
| **462881**269594ac1665344c7683787cc0 | 0.054,0.061,0.554,-0.482,-1.451,-0.065 | 0.098,0.085,1.342,-0.814,0.106,-0.869 | 0.067,0.083,-0.869,1.279,1.823,0.916 | 0.092,0.098,-0.954,0.620,1.798,-0.124 |
| **466262**386801c8e0228effb6ee250355 | 0.081,0.082,1.504,-1.343,-0.668,-0.655 | 0.094,0.048,1.627,-1.190,-1.627,-0.085 | 0.014,0.033,-1.196,1.900,1.186,0.630 | 0.044,0.042,-0.308,0.288,0.780,-0.345 |
| **4698d**ee9ab1255d9ff462230ee0830e7 | 0.040,0.064,1.812,-1.658,-0.198,-1.266 | 0.098,0.069,1.024,-1.466,-0.985,-0.166 | 0.089,0.098,-1.730,0.737,0.236,1.186 | 0.046,0.074,-0.268,-0.139,1.884,1.513 |
| **469b3**823e3f369400230819e5e944007 | 0.045,0.075,1.789,-1.897,-0.411,-0.547 | 0.072,0.065,1.631,-1.379,-0.553,-0.874 | 0.071,0.095,-1.791,1.920,-1.257,1.975 | 0.099,0.052,-1.067,0.396,1.837,1.844 |
| **46cdb**a64f3bca796dc60bbc712804a4c | 0.089,0.088,0.666,-0.481,-1.660,-0.032 | 0.011,0.075,0.823,-0.741,1.803,0.874 | 0.038,0.098,-1.153,1.878,1.328,0.668 | 0.011,0.023,-0.900,0.701,1.692,1.400 |
| **46dfa**9280e954167229a29ab61dc7332 | 0.075,0.048,1.678,-1.418,-1.503,-1.169 | 0.099,0.014,1.494,-0.686,-0.646,-1.499 | 0.064,0.013,-0.198,0.101,1.945,1.342 | 0.048,0.068,-1.513,1.004,1.370,1.948 |
| **46e94**42671fffce769342c3851652706 | 0.061,0.037,1.565,-1.446,-0.643,-1.096 | 0.092,0.052,1.021,-1.354,-1.987,0.932 | 0.021,0.053,-0.131,0.018,1.497,0.251 | 0.095,0.064,-0.357,-0.192,1.847,1.481 |

| | | | | |
|---|---|---|---|---|
| **47311**ca62283076bdd6bcb132e4d1fda | 0.080,0.030,1.538,-1.187,-1.403,-1.021 | 0.013,0.055,1.380,-1.460,-1.949,0.029 | 0.079,0.063,-1.874,1.977,-0.723,1.257 | 0.029,0.064,-1.168,0.839,-0.251,1.560 |
| **475079**f4353e141baff594cd8eddf92d | 0.024,0.050,1.224,-1.086,-1.431,-0.530 | 0.038,0.056,1.683,-1.370,-0.722,-1.701 | 0.052,0.038,-1.904,1.950,1.156,1.329 | 0.070,0.086,-0.831,0.224,1.430,1.094 |
| **475a4**dec52b156d0a41ad862bf51c89b | 0.091,0.045,1.809,-0.569,-0.929,-0.234 | 0.099,0.014,0.342,-0.033,-1.858,-1.340 | 0.020,0.045,-1.820,0.237,1.275,0.793 | 0.085,0.056,-1.320,1.166,0.183,0.679 |
| **476c6**11fca92855dd530bec53ad8881f | 0.075,0.081,1.389,-0.986,-1.734,-0.113 | 0.035,0.050,0.787,-1.096,0.237,-0.859 | 0.093,0.090,-1.627,0.745,1.333,1.340 | 0.058,0.094,-0.261,-0.235,1.302,0.956 |
| **476db**4374ffd7470f624219fa894b75a | 0.082,0.027,1.457,-1.320,-1.543,-0.332 | 0.052,0.031,0.727,-0.550,-0.707,-0.654 | 0.091,0.083,-1.137,0.569,1.374,1.857 | 0.031,0.073,-0.961,0.494,1.271,1.626 |
| **477c3**dea65065107f2e308bf804d0a50 | 0.087,0.048,0.959,-0.837,-0.607,-1.287 | 0.025,0.070,0.507,-0.381,-0.995,-0.899 | 0.032,0.088,-1.329,0.965,-0.029,0.610 | 0.093,0.051,-0.047,-0.154,0.846,0.333 |
| **4782d**aee2663324d7e782efe071c2d59 | 0.080,0.012,1.851,-1.484,-1.946,0.503 | 0.064,0.020,0.403,-0.586,-0.505,-1.575 | 0.069,0.053,-1.840,0.623,1.873,0.945 | 0.097,0.032,-1.489,1.312,0.140,1.903 |
| **486963**b797b4be2ea13c283845210f43 | 0.016,0.086,1.536,-1.644,-1.052,-0.960 | 0.018,0.061,0.525,-0.640,-0.882,-1.433 | 0.084,0.023,-0.440,0.168,1.796,0.191 | 0.061,0.098,-0.804,0.492,1.933,1.550 |
| **487377**b57613c0981199ffdd0e71ef99 | 0.052,0.066,1.223,-1.146,-0.378,-1.212 | 0.078,0.075,0.297,-0.498,0.171,0.137 | 0.045,0.038,-0.630,0.573,0.389,1.082 | 0.093,0.068,-0.803,0.430,1.834,0.878 |
| **4880c**84687dcbee89302058c6b6750d9 | 0.089,0.017,0.392,-0.071,-1.242,0.255 | 0.024,0.063,0.788,-0.980,-1.606,-0.805 | 0.073,0.087,-1.623,0.571,0.731,1.703 | 0.078,0.075,-1.945,1.485,1.034,1.552 |
| **488fa**5daef6ab05b8c588277c35e6e9c | 0.057,0.014,1.112,-1.082,-1.160,-0.721 | 0.077,0.038,0.813,-1.319,-1.214,1.772 | 0.072,0.080,-0.568,0.160,1.168,0.386 | 0.040,0.086,-0.651,0.196,0.079,1.912 |
| **490c9**48f5ba849a26ae4051d0d8e10da | 0.057,0.096,1.055,-1.079,0.019,-0.738 | 0.079,0.080,1.191,-0.856,1.884,-1.902 | 0.055,0.015,-0.616,0.849,1.545,-0.193 | 0.045,0.091,-0.908,0.409,1.571,1.132 |
| **4936e**7477ee111008474bbdfae653b3d | 0.030,0.045,1.788,-1.888,-1.163,0.014 | 0.085,0.079,0.974,-1.543,0.208,-1.243 | 0.097,0.079,-0.825,0.259,1.014,1.691 | 0.082,0.044,-0.486,0.158,1.890,0.916 |
| **4978b**d447aa02622eff7931ca327e892 | 0.062,0.079,1.109,-1.047,-0.133,-1.158 | 0.072,0.059,1.914,-1.186,-1.653,0.803 | 0.045,0.070,-1.374,1.821,0.903,0.490 | 0.012,0.064,-0.921,0.682,-0.513,1.292 |
| **49936**b5f21c195ae35b44f6e961a6099 | 0.098,0.046,0.565,-0.032,-1.625,0.264 | 0.056,0.087,1.116,-1.259,-1.967,-0.702 | 0.046,0.038,-1.812,0.868,0.408,0.952 | 0.094,0.071,-0.678,0.128,1.338,1.191 |
| **49dce**514c947b281c3d41ea1d7647b02 | 0.044,0.049,0.762,-0.981,-0.692,0.606 | 0.025,0.027,0.264,-0.188,0.203,-0.098 | 0.056,0.062,-1.200,1.835,1.353,0.060 | 0.018,0.042,-0.175,-0.017,1.879,1.327 |
| **4a20b**cca14a988fa37b8f2c60783fcf1 | 0.068,0.030,1.662,-1.625,-1.835,0.432 | 0.013,0.059,0.230,-0.634,1.803,-0.199 | 0.099,0.041,-0.283,-0.194,1.721,1.392 | 0.056,0.068,-0.156,-0.124,1.260,1.782 |
| **4a748**39b8e8611de0369939463c8a8c4 | 0.063,0.075,1.538,-1.111,-1.844,-0.456 | 0.043,0.025,0.743,-0.815,-1.720,0.510 | 0.062,0.020,-1.486,0.658,1.892,1.330 | 0.026,0.058,-0.321,0.014,1.633,1.238 |
| **4a8a0**5c99da32e3a3ab18c0e4620b334 | 0.043,0.054,1.669,-1.751,0.092,-1.371 | 0.058,0.058,1.187,-0.982,-1.933,-1.126 | 0.079,0.041,-0.932,1.081,-0.106,-1.151 | 0.076,0.096,-0.260,-0.111,1.395,0.785 |
| **4a91b**e7bdc332616f6424194721c8d97 | 0.069,0.051,0.848,-0.627,-0.423,-0.478 | 0.069,0.090,1.360,-1.976,-0.776,-1.176 | 0.031,0.074,-1.984,0.323,1.458,1.919 | 0.044,0.062,-0.793,0.214,1.965,1.496 |
| **4aaf2**d1e423e5a92292ba8640b4c4f57 | 0.020,0.042,0.702,-0.644,-1.403,-1.160 | 0.037,0.076,0.549,-0.640,0.316,-0.629 | 0.067,0.048,-1.275,0.974,1.607,0.841 | 0.083,0.093,-0.231,-0.112,0.929,1.235 |
| **4abe9**36a01e9f1c57b28e67e7817c011 | 0.075,0.059,1.259,-0.793,-1.504,-1.074 | 0.021,0.078,1.362,-1.880,0.129,1.369 | 0.062,0.085,-1.980,1.309,1.682,1.717 | 0.099,0.088,-0.376,-0.682,1.869,1.307 |
| **4ade5**7bae4891b4ca5257d9fc32494bb | 0.076,0.021,1.502,-1.586,-0.496,-0.476 | 0.034,0.097,1.284,-0.637,-1.531,0.614 | 0.074,0.099,-1.690,1.961,1.879,1.294 | 0.033,0.013,-0.176,-0.144,1.888,-0.471 |
| **4adea**50d7375189ce248d689ca7f4b2c | 0.090,0.046,1.673,-0.717,-1.135,-0.692 | 0.075,0.064,0.254,-0.106,-1.159,-0.066 | 0.032,0.064,-1.575,0.249,1.251,1.433 | 0.016,0.063,-0.840,0.626,1.587,0.842 |
| **4aeca**a0c5875732aa93fc5dfa7107cce | 0.059,0.021,1.537,-1.555,-1.440,1.048 | 0.010,0.060,0.693,-0.456,1.405,-1.322 | 0.043,0.062,-0.593,0.738,1.814,1.050 | 0.078,0.075,-0.601,0.278,1.512,0.344 |
| **4aed1**5d9b16af7d47b85cbc675736bd0 | 0.033,0.029,1.554,-1.582,-1.124,-0.093 | 0.094,0.049,0.485,-0.703,-0.664,-0.602 | 0.096,0.043,-1.762,0.689,1.687,-0.688 | 0.081,0.089,-0.417,-0.062,1.347,1.528 |
| **4aff2**dd8c98ef6b16ec560acc09aa246 | 0.043,0.058,1.717,-1.498,-1.678,-0.870 | 0.048,0.045,1.536,-1.155,0.003,-0.646 | 0.099,0.073,-1.749,1.400,1.823,1.946 | 0.062,0.075,-1.059,0.385,1.214,0.956 |
| **4b549**70ac348afd7ff6ea4547a07c48d | 0.048,0.021,1.841,-1.585,-1.981,-0.565 | 0.071,0.070,0.436,-0.581,-1.035,-0.331 | 0.020,0.063,-0.551,0.346,1.383,-0.266 | 0.082,0.094,-1.407,1.034,0.699,1.778 |
| **4b91d**e39b7cf149764a3828eca1bec52 | 0.070,0.077,1.968,-1.545,-1.433,-1.343 | 0.037,0.023,0.308,-0.390,-0.820,-1.298 | 0.098,0.072,-1.697,0.491,1.235,1.979 | 0.055,0.025,-0.752,0.638,1.336,0.491 |
| **4b98a**673179a3a67eaf73a7b5f71c3c5 | 0.021,0.079,1.911,-1.656,-1.471,-0.886 | 0.062,0.054,1.601,-1.544,-1.208,-0.444 | 0.049,0.047,-1.490,1.111,1.269,0.122 | 0.029,0.044,-0.903,0.511,0.758,1.497 |
| **4bbd6**c29bd9817b177cf213b03518f00 | 0.070,0.033,1.643,-1.507,-1.889,1.032 | 0.098,0.018,0.263,0.231,-1.511,-1.977 | 0.054,0.059,-1.080,0.164,1.720,1.174 | 0.096,0.069,-0.099,-0.182,1.935,-0.619 |
| **4bf27**4f87b3ca549b0f4129574d6108c | 0.049,0.024,1.607,-1.846,-0.459,-0.311 | 0.026,0.081,1.400,-1.173,-0.510,-0.593 | 0.049,0.062,-1.653,1.918,1.651,1.019 | 0.083,0.078,-1.193,0.813,1.416,0.895 |
| **4c097**156d427f170646c83179b05eeb0 | 0.070,0.059,1.571,-1.422,-0.402,-1.050 | 0.023,0.062,1.054,-1.242,-0.855,-0.836 | 0.038,0.077,-1.501,1.383,1.337,0.456 | 0.029,0.044,-1.112,1.093,-0.405,0.392 |
| **4c4aa**d8f484ef6cfc1b8b39163e1c31b | 0.072,0.059,0.339,0.302,-1.394,-0.755 | 0.047,0.082,0.730,-1.830,-1.314,0.567 | 0.052,0.066,-0.717,-0.057,1.778,1.364 | 0.025,0.091,-0.401,-0.096,1.387,1.829 |
| **4c6a6**b023078f17e36f95ffc669df3b2 | 0.059,0.030,1.162,-0.632,-0.804,-1.832 | 0.098,0.040,0.443,-1.412,-1.446,0.569 | 0.018,0.015,-0.810,0.238,0.865,1.114 | 0.082,0.055,-0.994,0.675,1.542,1.335 |
| **4c91d**e6de047f7f2b74c6865eef0f748 | 0.056,0.062,1.786,-1.774,-0.430,-0.597 | 0.013,0.064,1.770,-1.564,1.117,-1.651 | 0.094,0.052,-1.721,1.974,1.527,1.240 | 0.033,0.062,-0.500,0.111,1.912,1.491 |
| **4c9c3**559e968d145586a4ed913d297a9 | 0.089,0.054,1.228,-1.114,-1.351,0.713 | 0.076,0.030,1.386,-1.051,-1.519,0.681 | 0.042,0.071,-1.382,1.881,1.409,1.051 | 0.098,0.042,-0.560,0.008,1.739,1.548 |
| **4ca50**301cc32c60f7cc9055e94d0239f | 0.089,0.030,1.683,-1.717,-0.821,-0.525 | 0.093,0.090,0.688,-0.506,-1.870,0.720 | 0.024,0.093,-1.041,1.216,0.477,1.454 | 0.052,0.074,-1.591,1.376,1.433,0.685 |
| **4cc27**2d0273535298b86f0ad7b7e462c | 0.054,0.023,1.351,-1.592,-1.840,0.808 | 0.026,0.087,0.490,-0.937,-0.145,-0.109 | 0.014,0.021,-1.783,0.935,1.413,1.178 | 0.083,0.062,-0.633,0.299,1.818,1.010 |
| **4cc4a**acdd7e5d65475dc8093565cc1921 | 0.065,0.042,1.410,-1.685,-1.877,1.627 | 0.018,0.056,0.090,-0.312,-0.040,0.165 | 0.054,0.031,-0.660,0.042,0.828,1.985 | 0.061,0.064,-1.371,1.232,1.048,0.156 |
| **4cdb2**a76971fdfda2ae11ef150335794 | 0.048,0.041,1.463,-1.469,-1.049,-0.773 | 0.041,0.069,0.867,-1.066,0.321,-1.725 | 0.025,0.050,-1.507,0.979,1.208,1.865 | 0.056,0.071,-0.974,0.642,1.286,1.988 |
| **4cdb4**b3c351e060bdb10eca787de5056 | 0.026,0.081,1.632,-1.882,-0.496,-0.458 | 0.056,0.051,0.520,-0.428,-0.775,-0.905 | 0.090,0.056,-0.602,0.288,1.028,0.713 | 0.082,0.087,-0.434,0.373,1.943,-1.033 |
| **4d066**595cb27a758e19bca9bc4cb71d5 | 0.059,0.015,1.630,-1.966,-1.175,1.170 | 0.062,0.088,0.855,-0.664,-0.035,0.406 | 0.056,0.021,-0.725,0.941,1.932,0.158 | 0.098,0.094,-1.234,0.914,0.942,0.287 |
| **4d1bd**c8fdaef60efaea5fe1b13d16e65 | 0.077,0.035,1.438,-1.585,-0.720,0.604 | 0.059,0.086,0.825,-0.339,-0.198,0.160 | 0.068,0.041,-0.432,0.627,1.211,0.625 | 0.083,0.065,0.048,-1.251,1.934,1.692 |
| **4d68b**bd62f00bc26348ee8271e003e32 | 0.047,0.072,1.910,-1.291,-1.696,-1.329 | 0.098,0.047,1.012,-1.955,0.337,-1.741 | 0.041,0.070,-1.332,0.405,1.156,0.878 | 0.071,0.034,-0.875,0.376,1.952,1.956 |
| **4d915**306e0eb5122e33797002e25d913 | 0.090,0.023,0.432,-0.092,-1.616,-1.065 | 0.046,0.037,1.083,-0.691,-0.665,-1.052 | 0.055,0.086,-1.778,1.616,1.857,1.855 | 0.093,0.094,-1.123,0.197,1.246,1.533 |
| **4d9b2**fc3af097ef7378f6fe59f513ce8 | 0.026,0.047,1.068,-1.152,0.504,-0.453 | 0.019,0.060,0.584,-0.299,-0.217,-0.881 | 0.081,0.071,-0.374,0.227,0.807,1.920 | 0.082,0.067,-0.496,-0.079,1.876,1.261 |
| **4d9bb**55225683ce1dc046f3ec0267ac3 | 0.086,0.031,0.607,-0.451,-1.157,0.799 | 0.020,0.065,1.612,-1.813,0.015,-0.829 | 0.045,0.059,-1.589,1.317,-0.070,0.603 | 0.056,0.019,-0.442,0.034,1.654,1.619 |
| **4dd13**9608aac71db5108a6af829cd48d | 0.068,0.077,1.779,-1.269,-1.732,-1.115 | 0.015,0.060,1.057,-1.003,-1.058,-1.616 | 0.088,0.095,-1.929,1.067,1.963,1.385 | 0.088,0.055,-1.349,0.796,1.918,1.036 |
| **4dddb**e62dc20632cb04e1014788cf92c | 0.012,0.092,1.056,-1.234,0.198,-1.154 | 0.086,0.086,0.397,-0.232,-1.426,1.355 | 0.092,0.044,-0.130,0.107,1.641,-1.351 | 0.078,0.042,-1.031,0.731,1.635,1.221 |
| **4dec8**9123f74ff95ccc5f2956443c620 | 0.030,0.047,0.829,-0.390,-1.659,-1.006 | 0.062,0.024,0.808,-0.675,-1.837,-0.758 | 0.093,0.027,-1.293,0.589,0.460,1.903 | 0.026,0.068,-1.514,1.172,-0.226,1.604 |
| **4e729**9a6b4af98dbde1c3ec3b02f7ec5 | 0.079,0.042,1.861,-1.802,-1.945,-1.720 | 0.043,0.013,0.767,-0.695,-0.772,0.397 | 0.097,0.061,-1.155,0.952,1.883,1.134 | 0.085,0.061,-0.485,-0.091,1.879,1.576 |
| **4e78e**da0dd9d2690203cf332ab3b49dd | 0.077,0.054,1.968,-1.671,-1.640,-0.383 | 0.035,0.079,0.929,-0.659,-1.094,-1.457 | 0.044,0.022,-1.051,0.852,0.032,1.283 | 0.075,0.075,-1.272,1.090,1.925,-0.814 |
| **4e82d**8cd35bfea33e28bf7be21c02a0d | 0.072,0.039,1.532,-1.258,-1.018,-0.725 | 0.018,0.052,1.603,-1.114,0.696,-0.242 | 0.063,0.091,-0.641,0.815,1.731,-0.141 | 0.063,0.074,-0.596,0.436,1.321,-0.400 |
| **4ed07**0fc417670f8708e630d46f5d633 | 0.061,0.064,1.220,-1.450,0.293,-1.764 | 0.068,0.058,0.401,-0.230,-1.004,0.394 | 0.074,0.057,-1.208,1.910,-0.747,1.476 | 0.096,0.027,-1.143,1.039,0.857,-0.255 |
| **4ed47**39f5d0fd11f16ef24ebb48c10ee | 0.067,0.049,1.972,-1.796,-1.817,-0.318 | 0.076,0.014,0.665,-0.777,-0.796,-0.453 | 0.081,0.097,-0.348,-0.062,1.104,1.793 | 0.063,0.068,-1.001,0.638,1.826,1.898 |
| **4ed80**2bb87367a0e3cff51c09a769ced | 0.031,0.053,1.837,-1.819,0.711,-1.132 | 0.014,0.098,1.573,-0.998,0.828,-0.409 | 0.066,0.054,-1.662,1.945,1.219,1.660 | 0.050,0.040,-1.254,0.717,1.130,1.965 |
| **4ee12**d69fa0a8a808fdca370ffc85ef5 | 0.047,0.054,1.909,-1.636,-0.449,-1.691 | 0.017,0.054,1.192,-1.883,-0.942,1.020 | 0.037,0.095,-1.596,0.727,1.682,1.974 | 0.011,0.073,-0.378,0.167,0.152,1.241 |
| **4ee9d**610979de2eeb536a0a9fefbd055 | 0.075,0.091,1.657,-1.254,-0.506,-0.751 | 0.020,0.019,0.013,0.020,-1.531,0.124 | 0.016,0.079,-1.211,0.133,1.761,1.086 | 0.091,0.015,-0.115,0.003,1.218,0.606 |
| **4eee2**c906bcd36d08b0a73a07f25da23 | 0.081,0.022,1.435,-1.313,-1.536,1.191 | 0.021,0.067,0.687,-0.753,-1.002,0.044 | 0.062,0.036,-0.791,0.549,0.224,1.408 | 0.057,0.046,-1.468,1.111,1.580,0.673 |
| **4efb8**329898a68bcfbe060221c491812 | 0.075,0.057,1.535,-0.777,-0.736,-0.213 | 0.026,0.043,0.512,-1.014,-0.375,-1.882 | 0.068,0.041,-0.705,0.067,1.157,-0.878 | 0.017,0.093,-0.633,0.526,1.860,0.103 |
| **4f052**902fbda2d00b8cf1da0c51c7317 | 0.045,0.066,0.326,0.205,-1.320,-1.359 | 0.066,0.088,0.713,-1.311,0.850,-0.487 | 0.016,0.044,-1.966,1.102,1.955,1.211 | 0.058,0.020,-0.489,0.315,1.844,-0.737 |
| **4f2fe**872568399ee85bad7795af92dd3 | 0.088,0.089,1.660,-1.758,-0.647,-0.391 | 0.039,0.087,0.811,-0.698,-0.412,-1.604 | 0.012,0.038,-1.877,1.545,1.885,1.310 | 0.072,0.070,-0.648,0.342,1.923,0.949 |
| **4f303**bea4199892f105bb05be3478894 | 0.043,0.086,1.965,-1.848,-1.720,-0.584 | 0.093,0.082,0.843,-0.501,-1.378,-0.217 | 0.080,0.044,-1.705,1.360,0.354,1.262 | 0.056,0.053,-1.040,0.751,1.044,1.001 |
| **4f5c8**317cd99c7d74964bf881ce64434 | 0.051,0.082,1.490,-1.029,-1.448,-1.279 | 0.098,0.030,0.386,-0.388,-1.731,1.257 | 0.095,0.087,-0.654,0.121,1.211,0.226 | 0.082,0.073,-1.377,0.938,1.945,1.373 |
| **4f92e**1dcfcd8c11f07952741ae865fb5 | 0.075,0.019,0.863,-1.136,-0.809,0.419 | 0.027,0.031,0.854,-0.982,-0.321,-1.264 | 0.078,0.061,-1.461,0.817,1.839,1.308 | 0.095,0.076,-0.289,-0.237,1.969,1.037 |
| **4f970**fec7626594c8104913c350f4d09 | 0.084,0.026,1.332,-1.520,-1.070,0.664 | 0.095,0.032,0.754,-0.890,-0.452,-0.349 | 0.040,0.055,-0.472,0.387,1.620,0.081 | 0.087,0.053,-1.440,1.096,1.521,1.687 |
| **4fa0b**13d3a3963aee792343b74bc625f | 0.020,0.096,1.909,-1.766,0.346,-1.707 | 0.018,0.016,1.061,-0.891,-0.557,0.929 | 0.069,0.063,-1.870,1.948,1.381,1.955 | 0.042,0.043,-0.337,-0.191,1.932,1.820 |

| | | | | |
|---|---|---|---|---|
| **4fa18**90045891fd0d1bc98d89e0da450 | 0.039,0.018,0.670,-0.587,-1.264,-0.807 | 0.022,0.075,0.194,-0.181,-1.091,-0.129 | 0.075,0.016,-1.632,1.221,-0.265,0.900 | 0.016,0.082,-1.533,1.324,-0.923,1.380 |
| **4fac6**f16a182732bfb84f66ed0ef2ea6 | 0.039,0.079,1.857,-1.510,-0.491,-0.831 | 0.052,0.027,1.036,-1.373,-1.599,0.099 | 0.092,0.039,-1.237,0.088,1.699,1.775 | 0.042,0.018,-0.490,0.381,0.607,0.473 |
| **4faef**96d6d19fe269c017e3cdb3a0cc3 | 0.048,0.041,1.855,-1.577,-0.616,-1.367 | 0.053,0.053,1.944,-1.353,-0.463,-1.208 | 0.052,0.066,-1.342,1.335,1.511,1.110 | 0.069,0.052,-1.406,1.222,-0.319,1.254 |
| **4fcce**068e256ed7bf7142a5878fb53a7 | 0.096,0.043,1.612,-1.668,-1.922,0.390 | 0.018,0.051,0.181,-0.291,-0.650,1.541 | 0.053,0.034,-0.626,0.790,1.222,-0.194 | 0.031,0.057,-0.817,0.443,0.274,1.658 |
| **4fe2d**96f4c781c8676d4d7818ab634ad | 0.097,0.016,1.729,-1.300,-1.586,-0.574 | 0.039,0.080,0.686,-1.047,1.736,-1.698 | 0.046,0.034,-1.242,0.665,0.727,1.202 | 0.053,0.024,-0.823,0.579,1.780,1.363 |
| **4ff34**62ea8f98b3aabd82104817c6cbb | 0.096,0.029,1.546,-1.485,-1.553,0.365 | 0.094,0.082,0.583,-1.385,-0.582,1.070 | 0.076,0.030,-1.838,1.243,0.269,1.782 | 0.096,0.074,-0.452,0.186,1.778,0.283 |
| **4fff1**6a96788b333afb2673d548d92b5 | 0.046,0.050,1.457,-1.450,-1.696,-0.513 | 0.064,0.036,1.164,-0.548,-1.968,-1.040 | 0.094,0.053,-0.675,1.043,-0.075,-0.237 | 0.095,0.044,-0.357,-0.266,1.827,1.647 |
| **500e3**1599af8cd9f5a8b5d0df05d48bc | 0.084,0.040,1.203,-1.244,-0.591,-0.868 | 0.013,0.071,0.740,-0.674,-0.113,-0.396 | 0.093,0.086,-0.670,0.763,0.163,1.109 | 0.026,0.035,-0.245,0.132,1.776,1.261 |
| **5030e**e4d8da9563a646530eb5189272f | 0.099,0.021,1.870,-1.690,-1.466,0.344 | 0.023,0.074,0.898,-1.386,-0.858,-1.258 | 0.014,0.056,-0.696,0.248,1.501,0.672 | 0.068,0.067,-0.439,-0.043,1.738,1.683 |
| **503bc**8e4e7b1fd27e80976681653a082 | 0.095,0.062,0.727,-0.584,-1.533,-0.825 | 0.048,0.077,0.888,-0.486,-1.362,-1.061 | 0.076,0.096,-1.869,1.945,-0.578,1.821 | 0.073,0.089,-0.825,0.651,0.526,0.421 |
| **504e7**a9b71eb2868a69e6c3bf001fa3a | 0.029,0.014,1.510,-1.523,-1.865,-0.667 | 0.032,0.038,0.463,-0.819,-0.200,0.128 | 0.021,0.037,-1.401,0.743,0.507,1.078 | 0.067,0.059,-1.361,1.042,0.792,1.176 |
| **50592**d102be43e09d9fa00c974e056fc | 0.087,0.026,1.476,-1.000,-1.110,0.036 | 0.017,0.083,0.387,-0.353,1.650,-1.196 | 0.098,0.039,-1.034,-0.130,1.837,1.882 | 0.049,0.065,-1.257,0.746,1.314,1.453 |
| **507f0**dbf7a35dc8408bd764bcb57080b | 0.092,0.018,0.594,-0.429,-1.797,1.245 | 0.018,0.051,1.047,-0.571,-1.731,1.229 | 0.025,0.053,-1.005,1.875,1.056,1.943 | 0.054,0.083,-1.909,1.326,1.971,1.618 |
| **508b0**482656e4c80f3c4e6f1cc3c1307 | 0.049,0.043,1.583,-1.168,-1.024,-0.925 | 0.068,0.084,0.522,-0.823,-0.633,-0.634 | 0.092,0.052,-1.929,0.407,1.724,0.495 | 0.058,0.020,-1.261,0.991,1.772,1.408 |
| **50b53**aa95dfd2c458f1c252c4f2f5641 | 0.051,0.011,1.907,-1.919,-0.879,-0.582 | 0.047,0.040,1.473,-1.262,-0.441,-0.542 | 0.031,0.058,-1.573,1.548,1.371,1.306 | 0.097,0.067,-0.497,-0.620,1.994,1.401 |
| **50f81**2e19e6ae353ac26b62b87540468 | 0.036,0.026,1.229,-1.645,-0.216,0.126 | 0.032,0.021,0.171,-0.244,1.466,-0.336 | 0.087,0.028,-0.930,1.171,1.201,1.921 | 0.088,0.029,-0.511,0.279,1.862,-0.046 |
| **5105a**5ef02f2d983acacc3586a4f7819 | 0.092,0.058,1.457,-1.579,0.262,-1.850 | 0.015,0.025,0.309,-0.484,-1.598,0.244 | 0.099,0.090,-0.540,0.122,0.574,0.351 | 0.088,0.032,-0.787,0.587,1.134,0.263 |
| **51498**aa6212398beb134f26a16a3e4e6 | 0.060,0.030,0.501,-0.609,-0.807,0.637 | 0.071,0.082,0.524,-0.622,-0.916,1.038 | 0.031,0.065,-1.021,0.939,1.377,0.403 | 0.075,0.097,-0.181,-0.391,1.435,0.973 |
| **5149a**7040b6724ca8d46f262e60a3113 | 0.085,0.048,1.861,-1.783,-1.279,-0.389 | 0.079,0.079,1.125,-1.594,-0.882,-0.169 | 0.043,0.053,-0.819,0.581,1.695,-1.301 | 0.077,0.078,-0.284,-0.256,1.415,1.820 |
| **514ab**b5ca72cf775da8b4f5216f4e723 | 0.052,0.023,1.454,-1.977,-0.435,1.106 | 0.057,0.052,0.442,-0.147,-0.699,-1.196 | 0.096,0.082,-0.242,0.592,1.667,-1.722 | 0.046,0.070,-0.481,0.386,-0.347,0.984 |
| **514b1**f39ece508f0cc471082431c67e1 | 0.028,0.028,1.063,-1.117,-1.233,-0.053 | 0.033,0.091,0.404,-0.167,0.797,-1.102 | 0.078,0.077,-1.099,1.587,0.172,1.053 | 0.016,0.044,-1.223,1.091,-0.135,1.023 |
| **51555**e9683710150efe6cc2bc019439d | 0.057,0.016,1.715,-1.630,-1.586,0.230 | 0.044,0.051,1.134,-0.711,-0.928,-0.638 | 0.070,0.028,-0.497,0.386,0.976,1.590 | 0.034,0.050,-0.745,0.454,1.395,1.139 |
| **51598**f09e2693d7ca7da57654b39aae7 | 0.012,0.063,0.821,-0.828,-0.030,-0.640 | 0.072,0.028,1.330,-0.969,-1.449,-0.165 | 0.065,0.088,-0.753,0.619,1.169,1.938 | 0.084,0.097,-1.656,1.373,-0.614,1.593 |
| **51600**90178e3c514eea1910201cd34aa | 0.050,0.050,1.776,-1.807,-0.991,0.006 | 0.068,0.061,0.876,-0.675,-0.342,-1.820 | 0.064,0.095,-1.833,1.166,-0.496,1.016 | 0.091,0.022,-1.036,0.740,0.966,0.155 |
| **5192d**2b1aa80495af609f3170b709315 | 0.066,0.022,1.463,-1.336,-0.644,-0.157 | 0.028,0.067,1.229,-0.826,-1.090,-0.797 | 0.099,0.084,-0.765,0.278,1.860,0.228 | 0.090,0.089,-0.289,-0.720,1.734,1.493 |
| **5194e**055608b3e256af5d85b0dd33529 | 0.085,0.020,1.793,-1.785,-1.641,0.420 | 0.069,0.084,1.457,-1.430,0.921,-0.666 | 0.069,0.033,-1.828,1.933,1.875,1.691 | 0.079,0.073,-1.446,0.814,1.858,1.479 |
| **52057**4ab9499b5dd56f812e7854ed0de | 0.033,0.027,0.843,-0.833,-0.972,-1.028 | 0.050,0.039,0.692,-0.822,-1.746,-0.995 | 0.063,0.044,-1.874,1.246,0.819,0.987 | 0.064,0.097,-0.769,0.413,1.596,1.086 |
| **520cf**c2d5d43493ef8730b37eed3bffc | 0.079,0.064,1.275,-1.257,-0.516,-1.234 | 0.087,0.039,0.761,-0.392,-1.521,0.447 | 0.017,0.055,-0.655,0.849,1.359,0.386 | 0.096,0.037,-0.771,0.562,1.261,0.391 |
| **52800**6b8afa90f3bddf1d5907fbdd8fa | 0.099,0.078,1.695,-1.580,0.418,-1.090 | 0.083,0.077,1.546,-1.411,-1.022,1.492 | 0.079,0.026,-0.856,0.741,0.830,1.647 | 0.045,0.098,-0.704,0.267,1.395,0.639 |
| **5288c**9722a0d914d024e185bfb204293 | 0.013,0.086,1.345,-1.359,0.698,-1.118 | 0.098,0.080,1.473,-1.222,-1.914,-1.463 | 0.077,0.082,-1.025,0.895,1.071,1.246 | 0.095,0.073,-0.704,0.415,0.819,1.845 |
| **52bb5**169cdbfa41a2f26865ade5c68b0 | 0.067,0.063,0.810,-0.385,-1.935,-0.925 | 0.019,0.082,0.838,-1.068,-0.820,0.250 | 0.061,0.088,-1.382,0.741,0.768,1.140 | 0.047,0.047,-1.157,0.888,0.827,1.142 |
| **53162**c5811d3e7e025601fe9b3fb7cd1 | 0.074,0.027,1.844,-1.456,-0.997,-1.906 | 0.018,0.066,1.263,-1.872,-1.948,1.212 | 0.061,0.074,-1.255,0.486,1.400,1.917 | 0.073,0.071,-0.463,-0.003,1.740,1.112 |
| **531ff**26d0db73902c57bc86231b0da42 | 0.024,0.073,1.913,-1.845,0.322,-1.362 | 0.013,0.018,1.157,-0.876,-1.443,1.053 | 0.100,0.055,-1.640,1.941,1.815,0.886 | 0.062,0.018,-1.018,0.722,1.806,0.338 |
| **5340f**1452ded225dfb959b0e2df956d5 | 0.097,0.042,1.590,-1.330,-1.125,0.440 | 0.027,0.074,1.481,-1.357,1.319,-1.835 | 0.066,0.025,-0.978,0.884,0.468,0.465 | 0.087,0.072,-1.126,0.449,1.436,1.520 |
| **535c7**387c544bd4ecf90d615ca05fc7b | 0.099,0.012,0.984,-0.932,-1.269,1.327 | 0.027,0.021,1.094,-0.885,1.988,1.854 | 0.014,0.088,-0.591,0.910,1.825,1.021 | 0.067,0.093,-1.086,0.527,1.711,1.916 |
| **539b7**2396147abc17bad511814085482 | 0.098,0.014,0.838,-0.723,-1.617,0.960 | 0.014,0.014,0.971,-0.877,-0.011,-1.082 | 0.099,0.066,-1.786,1.598,1.318,1.880 | 0.098,0.051,-0.341,-0.042,1.344,0.807 |
| **53a1e**b618f89db005b75106a5d111c9b | 0.095,0.090,1.715,-0.924,-1.132,-1.175 | 0.091,0.069,1.320,-1.655,-1.891,-1.348 | 0.097,0.051,-1.617,0.761,0.186,0.005 | 0.099,0.045,-0.438,0.130,0.926,1.323 |
| **53b42**4510bad040047c6430bea8ee9d5 | 0.088,0.025,1.158,-0.841,-1.077,-0.665 | 0.091,0.011,1.400,-1.155,-1.604,-0.201 | 0.033,0.061,-1.816,1.078,0.571,1.895 | 0.078,0.019,-1.339,0.854,1.716,1.476 |
| **53cc0**1bb78376e7b8b198cfe4a1a562e | 0.083,0.018,1.153,-0.675,-1.942,-0.044 | 0.099,0.079,1.398,-1.069,-1.790,-1.205 | 0.040,0.042,-1.750,1.068,-0.114,1.245 | 0.054,0.091,-0.110,-0.197,0.825,0.847 |
| **54610**f16b5dd5ac8d0cc217d93a2a81d | 0.044,0.087,1.409,-1.165,-0.723,-0.986 | 0.076,0.051,0.775,-0.984,-0.460,-0.836 | 0.089,0.067,-1.886,0.668,1.883,1.945 | 0.092,0.094,-0.731,0.133,1.777,1.420 |
| **5472a**e44eb080ef529a0b5ea47fcd637 | 0.078,0.040,1.882,-1.431,-1.894,-0.658 | 0.097,0.012,0.990,-0.503,-1.022,-1.878 | 0.026,0.057,-1.968,1.932,0.713,1.920 | 0.088,0.031,-0.571,0.032,1.081,1.458 |
| **54ad7**f1cf93c09953845d9616720f0c3 | 0.058,0.051,1.961,-1.285,-1.982,-1.499 | 0.025,0.063,1.093,-1.391,-0.422,-1.405 | 0.055,0.069,-1.951,0.923,1.712,0.663 | 0.091,0.051,-0.122,-0.360,1.614,0.754 |
| **54b58**59075e47e9a1d268e4811a2a913 | 0.028,0.016,1.442,-1.640,-0.516,-1.128 | 0.053,0.062,0.903,-1.278,-1.599,0.436 | 0.075,0.056,-1.951,1.096,0.499,0.922 | 0.076,0.069,-1.353,1.024,1.405,0.948 |
| **55026**eb38fd5a1aff16c5174d28b08bc | 0.073,0.061,1.074,-0.900,-1.262,-0.412 | 0.096,0.070,1.946,-1.378,-1.426,-0.673 | 0.090,0.087,-1.492,1.772,1.675,-0.480 | 0.095,0.067,-0.476,0.029,0.988,1.353 |
| **55137**bd765ccdedac9289ee7567d93fb | 0.067,0.088,1.713,-1.492,-1.709,-0.400 | 0.081,0.093,0.751,-0.556,-1.826,1.446 | 0.066,0.036,-0.424,0.971,-0.303,-1.387 | 0.082,0.027,-0.235,-0.180,1.830,-0.687 |
| **55300**37cc61273dca1dc99d1deb07009 | 0.088,0.026,1.854,-1.618,-1.407,0.240 | 0.021,0.055,0.868,-1.507,-0.589,-1.581 | 0.093,0.095,-1.486,0.521,0.431,0.306 | 0.097,0.066,-0.910,0.350,1.731,1.412 |
| **5535e**39b1708af0ee2503a62c3c9738f | 0.065,0.076,1.984,-1.389,-0.938,-1.263 | 0.096,0.071,1.455,-1.534,-1.112,-1.143 | 0.040,0.068,-1.696,0.790,1.803,0.398 | 0.040,0.059,-0.267,-0.213,0.971,1.839 |
| **5566b**7cc96a0c53c0af775b3e95ec1f3 | 0.098,0.016,1.814,-1.282,-1.882,0.788 | 0.090,0.068,1.047,-0.670,-1.774,1.047 | 0.012,0.012,-0.958,1.033,0.235,-0.823 | 0.074,0.062,-1.749,1.376,1.303,0.135 |
| **55934**e47186bafd8202ed2cf3450e3eb | 0.065,0.047,1.673,-1.921,0.002,0.032 | 0.050,0.016,0.476,-0.648,0.488,-1.199 | 0.075,0.070,-0.709,0.219,0.234,1.952 | 0.087,0.095,-0.908,0.016,1.970,1.182 |
| **55afb**2d36207a1f9abe26ae148d0c492 | 0.049,0.062,1.520,-1.427,-0.817,-0.580 | 0.076,0.017,0.472,-0.724,-0.585,0.077 | 0.071,0.070,-1.197,0.459,1.765,1.631 | 0.096,0.054,-0.929,0.609,1.833,0.405 |
| **55b60**13ca644d1ad8dbd3d901d2b8eb8 | 0.089,0.030,0.767,-0.460,-1.705,0.018 | 0.043,0.087,0.708,-0.763,-0.662,-0.770 | 0.012,0.076,-0.852,0.603,0.911,0.520 | 0.051,0.094,-0.960,0.514,1.878,1.297 |
| **55b7a**c2fbd33495f19d132da4e63385c | 0.073,0.024,1.743,-1.459,-1.845,0.146 | 0.052,0.082,0.749,-0.533,-0.986,0.152 | 0.064,0.089,-1.861,1.559,1.936,1.541 | 0.033,0.054,-1.258,0.902,-0.385,1.813 |
| **55cb1**39c420ef46c807e99ba7b7bc60d | 0.078,0.018,1.025,-1.111,-0.925,-0.267 | 0.022,0.027,0.647,-0.548,-0.950,-1.216 | 0.018,0.056,-0.939,0.859,-1.406,1.393 | 0.075,0.084,-0.131,-0.371,0.829,1.764 |
| **55fa7**b7bba2751ec75c46f65329b4dc5 | 0.060,0.015,1.624,-1.703,-0.950,-0.969 | 0.043,0.052,0.753,-0.710,-0.556,-1.947 | 0.084,0.044,-1.216,1.005,-0.019,1.294 | 0.089,0.097,-0.535,0.359,1.509,0.252 |
| **55fb0**40839908e096bab6365f6bbe041 | 0.068,0.056,0.886,-0.814,-1.565,-0.398 | 0.092,0.083,1.345,-0.745,-1.574,0.504 | 0.050,0.085,-1.186,1.524,-0.017,0.838 | 0.078,0.075,-0.615,-0.083,0.999,1.735 |
| **566b4**62c3cc5bd8d7f7d6b1331ac8781 | 0.088,0.083,1.515,-1.192,-1.875,0.678 | 0.069,0.059,1.619,-1.063,-1.273,-1.410 | 0.041,0.077,-1.348,0.819,1.742,0.550 | 0.059,0.059,-0.806,0.310,1.654,0.628 |
| **566ec**f6ed2aa7e60166b9508477e166c | 0.091,0.024,1.562,-1.612,-1.353,0.522 | 0.027,0.055,0.469,-0.319,-0.221,-1.175 | 0.045,0.032,-0.768,0.847,-0.889,1.149 | 0.015,0.089,-0.644,0.315,1.477,1.544 |
| **5676b**a1eb917c4f35b82079039d6ffc6 | 0.079,0.012,1.666,-1.940,-1.468,0.642 | 0.040,0.047,1.033,-0.984,-1.548,-1.098 | 0.022,0.032,-0.637,0.396,1.087,1.463 | 0.054,0.067,-0.769,0.314,1.996,1.711 |
| **5699a**6818c19131f1c6e4664154489b8 | 0.068,0.017,1.201,-1.290,-1.819,1.184 | 0.074,0.052,1.103,-0.642,-0.633,0.198 | 0.096,0.051,-1.715,1.962,1.570,0.267 | 0.027,0.060,-1.010,0.484,0.684,1.858 |
| **56b1b**f99fe9b616a4ac8448b1426d1e6 | 0.090,0.059,1.933,-1.342,-1.909,-0.900 | 0.036,0.017,1.944,-1.858,-1.006,-1.106 | 0.061,0.056,-1.963,1.621,1.643,1.804 | 0.086,0.035,-0.643,0.201,1.834,0.214 |
| **56b79**0ba1e60e081dc261631d402d072 | 0.051,0.078,1.651,-1.425,-1.436,-0.377 | 0.021,0.091,0.677,-0.325,-0.468,-1.304 | 0.069,0.026,-0.782,0.855,-0.576,0.980 | 0.063,0.075,-0.629,0.084,1.157,1.215 |
| **56dbd**64eda75929464c086ef9f954376 | 0.092,0.037,1.646,-1.505,-1.281,-0.613 | 0.077,0.016,1.265,-1.288,-1.919,1.392 | 0.089,0.060,-1.616,1.246,0.293,1.957 | 0.086,0.058,-0.722,0.360,1.943,0.504 |
| **56dfa**7fb5e7376701974ede35bae33e1 | 0.041,0.042,0.743,-0.788,-0.422,-0.362 | 0.013,0.016,0.728,-0.417,-1.429,-0.505 | 0.071,0.059,-1.231,1.879,-1.039,1.952 | 0.046,0.075,-1.379,1.120,1.931,0.156 |
| **56e8f**8d4fdc641284cd8b879d00bc3a4 | 0.049,0.043,0.986,-0.449,-1.850,-0.186 | 0.030,0.082,0.654,-0.763,-1.898,-0.666 | 0.089,0.061,-0.755,0.135,-0.051,0.997 | 0.095,0.092,-0.637,0.371,1.484,-0.546 |
| **56fd7**d38d1534fc0f976c69ae21acf27 | 0.016,0.060,1.344,-1.219,-0.153,-1.710 | 0.031,0.062,1.379,-0.720,1.777,-1.889 | 0.042,0.067,-1.152,1.978,1.752,-0.584 | 0.091,0.057,-1.402,1.027,0.268,0.959 |
| **572d5**42f3175559e0d7166cbd81f22c3 | 0.095,0.060,0.767,-0.733,-0.519,-0.670 | 0.016,0.045,0.100,-0.214,-1.591,1.237 | 0.064,0.096,-1.666,1.051,0.039,1.922 | 0.086,0.068,-0.985,0.773,1.470,0.310 |
| **5758b**7941cb6a8bceba354fbc59242a3 | 0.079,0.042,1.740,-1.228,-1.337,-0.941 | 0.023,0.086,0.319,-0.302,-0.074,-0.486 | 0.067,0.056,-0.758,0.400,-0.339,0.894 | 0.052,0.029,-1.229,1.199,-0.170,0.559 |
| **576aa**241beac99c5a2c8fbe660ad4112 | 0.045,0.017,1.740,-1.878,-1.613,-0.276 | 0.045,0.075,0.581,-0.250,-0.879,0.488 | 0.081,0.067,-0.632,0.901,1.659,0.729 | 0.012,0.065,-1.053,0.452,1.715,1.922 |

| | | | | |
|---|---|---|---|---|
| **57bb9**3932e0e300a25f3f472c90dd8a1 | 0.046,0.080,1.089,-0.800,0.256,-1.892 | 0.091,0.072,1.071,-1.370,-1.847,1.234 | 0.059,0.058,-1.885,1.107,1.770,1.378 | 0.029,0.089,-1.787,1.358,1.931,1.378 |
| **57f17**8a32edd08884b094f9bcc74eade | 0.051,0.071,0.470,-0.515,-1.205,0.323 | 0.030,0.063,0.822,-0.534,0.438,-0.254 | 0.021,0.025,-0.991,1.852,1.891,-0.602 | 0.028,0.023,-1.279,1.093,1.848,0.973 |
| **57f78**d2ada5194da25f2a729684f50d2 | 0.044,0.058,1.114,-1.262,0.294,-1.564 | 0.097,0.081,0.456,-0.558,-1.497,1.237 | 0.039,0.033,-1.199,0.796,1.259,0.801 | 0.087,0.029,-0.209,-0.099,1.644,0.486 |
| **581eb**8bba6103386e4dca71518505fa5 | 0.064,0.026,1.945,-1.809,-1.951,0.391 | 0.029,0.057,0.563,-0.552,0.120,-0.161 | 0.099,0.014,-0.669,0.557,0.710,0.750 | 0.085,0.084,-0.527,-0.180,1.687,1.455 |
| **583ee**34d0b3181d309f62fe0c8e99a91 | 0.063,0.049,1.509,-1.660,-1.743,0.008 | 0.026,0.049,0.735,-0.813,1.357,-0.335 | 0.053,0.064,-1.799,1.704,1.663,0.525 | 0.047,0.081,-0.619,0.460,1.980,-0.081 |
| **585dd**71e9b59ff1afa7f5eb15e009584 | 0.096,0.028,1.331,-1.282,-1.653,1.656 | 0.078,0.033,0.471,-0.500,-1.041,0.191 | 0.099,0.070,-1.693,1.019,1.497,0.481 | 0.092,0.039,-0.354,0.177,0.421,1.964 |
| **586d9**a44428e77b843ce1d3c79597572 | 0.040,0.061,1.122,-1.398,-0.019,-0.553 | 0.046,0.092,0.915,-0.259,-1.949,-1.487 | 0.050,0.065,-1.444,1.762,0.267,-0.697 | 0.036,0.058,-1.106,1.033,0.103,0.467 |
| **58860**b72eb1a437516a92b850a401bde | 0.085,0.021,0.661,-0.615,-1.706,1.160 | 0.053,0.032,0.199,-0.080,-1.451,-0.426 | 0.082,0.079,-1.408,0.932,1.764,-0.321 | 0.082,0.071,-0.550,0.309,0.929,1.108 |
| **58919**0d9bc04c47a3a97de9787be8f2e | 0.022,0.053,1.662,-1.455,-0.177,-1.571 | 0.095,0.051,1.273,-1.463,1.103,-0.071 | 0.094,0.091,-1.937,1.762,1.135,1.413 | 0.061,0.047,-0.931,0.735,1.602,-0.361 |
| **58a4e**493068e1d93fd9bd7d990b91bd4 | 0.099,0.019,1.911,-1.668,-1.788,0.610 | 0.093,0.078,1.453,-1.915,0.256,0.116 | 0.068,0.093,-1.951,0.800,1.900,1.352 | 0.011,0.098,-1.181,0.437,1.741,1.953 |
| **58acf**f646a5d26059caac060f5cae7f6 | 0.071,0.049,1.872,-1.906,-0.983,-0.391 | 0.051,0.028,0.487,-0.324,-1.607,-1.061 | 0.077,0.014,-1.380,0.757,1.494,0.812 | 0.091,0.074,-1.083,0.501,1.983,0.951 |
| **58c2f**550ddddd38fd482f21885a03a71 | 0.062,0.078,1.755,-1.253,-1.490,-0.920 | 0.038,0.070,0.355,-0.578,0.053,-0.916 | 0.091,0.082,-1.716,0.463,1.862,1.024 | 0.047,0.068,-0.893,0.815,0.812,0.372 |
| **58e56**170e19c6729da28e31609b9d08f | 0.076,0.019,1.538,-1.287,-1.720,0.494 | 0.095,0.017,1.292,-1.697,-0.306,-0.540 | 0.061,0.052,-1.758,0.954,1.805,1.968 | 0.097,0.075,-0.628,0.152,1.627,0.218 |
| **58f66**226df253338a147a170202ac97b | 0.037,0.021,1.308,-1.210,-1.090,-0.435 | 0.027,0.044,1.603,-1.068,-0.773,-0.169 | 0.099,0.100,-0.720,0.149,1.910,0.127 | 0.026,0.091,-1.090,0.458,1.257,1.171 |
| **59126**f1abc157546345ed540eef6ebd3 | 0.081,0.057,1.272,-1.419,-1.128,0.128 | 0.038,0.063,0.273,-0.733,-0.022,0.705 | 0.075,0.065,-1.112,0.369,-0.007,1.874 | 0.098,0.069,-0.637,0.208,1.579,1.231 |
| **592cd**b1975dc67679106d0b4207c293d | 0.043,0.082,1.808,-1.601,-0.338,-1.462 | 0.079,0.085,1.075,-1.800,0.174,-1.301 | 0.062,0.087,-1.588,0.821,0.877,1.356 | 0.085,0.074,-0.194,-0.281,1.998,1.013 |
| **593bd**2b0a111fea3500f17e3818dac54 | 0.091,0.043,0.853,-0.597,-1.883,0.841 | 0.028,0.080,0.706,-0.122,-1.077,-1.907 | 0.045,0.039,-1.560,1.866,1.817,0.375 | 0.086,0.089,-1.021,0.064,1.797,1.774 |
| **59557**9171d40be5da1256980afb5a66c | 0.084,0.053,1.721,-1.622,-0.991,-1.161 | 0.093,0.047,1.667,-1.465,-0.866,-0.091 | 0.036,0.030,-1.303,1.400,1.076,1.568 | 0.044,0.068,-0.403,-0.087,1.125,1.676 |
| **595c6**0d6c4bf316d60cefe8646572ed9 | 0.028,0.068,1.637,-1.421,-0.407,-1.088 | 0.055,0.018,0.712,-1.874,0.118,-0.159 | 0.059,0.022,-1.695,0.547,1.888,0.089 | 0.088,0.070,-1.592,1.161,1.948,-0.126 |
| **59607**37466321e15415d45c2c003eab9 | 0.049,0.082,1.885,-1.864,-0.806,-0.441 | 0.069,0.045,0.955,-0.682,-0.850,-1.344 | 0.036,0.083,-1.398,1.043,-0.258,1.319 | 0.059,0.038,-0.403,-0.148,1.842,1.360 |
| **598e0**976c4401a4b0ac846166c375f65 | 0.080,0.099,1.948,-1.180,-1.451,-1.189 | 0.015,0.047,1.501,-0.969,-0.878,-1.281 | 0.070,0.014,-1.957,1.714,1.848,1.966 | 0.056,0.060,-1.719,0.973,1.844,0.723 |
| **59abf**3b050d1b82fd86504da5cebd822 | 0.030,0.044,1.738,-1.668,-1.773,-0.174 | 0.015,0.063,1.627,-1.344,-1.011,-0.307 | 0.083,0.060,-1.549,1.487,0.865,1.512 | 0.058,0.086,-0.291,-0.693,1.853,1.665 |
| **59d7a**123f64e7874a0802ce4cdabbdbb | 0.053,0.040,1.071,-1.221,-1.906,0.956 | 0.025,0.048,0.105,-0.206,0.301,1.102 | 0.084,0.062,-0.740,1.044,1.161,0.201 | 0.071,0.036,-0.719,0.343,1.902,1.594 |
| **59dc0**e34973a85718d311166fc88de38 | 0.086,0.030,0.874,-0.788,-1.929,-0.450 | 0.074,0.084,0.394,-0.173,-1.191,-0.389 | 0.050,0.054,-1.225,1.420,0.533,-0.275 | 0.091,0.056,-0.479,0.226,1.955,0.281 |
| **59e67**cf26c8cf1c46865b489a0ccc9c0 | 0.083,0.066,1.687,-1.570,-0.789,0.087 | 0.047,0.054,1.895,-1.906,0.337,-0.792 | 0.047,0.029,-1.531,1.354,1.836,1.856 | 0.058,0.098,-0.618,0.359,0.838,0.348 |
| **59ed8**3687e45e74bad211f9210153e77 | 0.056,0.042,1.507,-1.543,0.035,-1.125 | 0.089,0.075,0.880,-0.875,-0.359,-0.393 | 0.056,0.058,-0.760,0.156,1.932,1.963 | 0.021,0.020,-0.761,0.694,-1.391,1.880 |
| **5a480**cef828b2b3c2b5657be211da5e2 | 0.044,0.030,1.247,-1.608,-1.138,0.169 | 0.041,0.075,0.368,-0.336,-0.859,-0.829 | 0.070,0.084,-0.626,0.295,1.243,0.065 | 0.090,0.075,-0.185,-0.224,1.454,1.663 |
| **5a4a5**04bd219c93578fc70740d9fc9ae | 0.041,0.031,1.331,-1.443,-1.377,0.157 | 0.010,0.079,0.520,-0.551,-0.606,-0.182 | 0.083,0.013,-1.958,1.916,1.607,0.960 | 0.080,0.065,-0.596,0.382,1.891,-0.054 |
| **5a4db**20d6aaeeae0be09b1e307e31b3c | 0.085,0.044,1.097,-0.931,-1.901,0.867 | 0.022,0.074,0.648,-0.557,-0.444,0.987 | 0.099,0.067,-0.983,0.741,1.131,1.854 | 0.056,0.076,-1.132,0.948,-0.285,0.598 |
| **5a59c**a0b567d5de14a9050a3f3530491 | 0.016,0.068,0.770,-0.571,-0.151,-1.233 | 0.020,0.079,1.606,-1.461,-0.327,-1.523 | 0.073,0.056,-1.833,1.921,1.285,1.191 | 0.060,0.072,-1.001,0.504,1.649,1.505 |
| **5a6d0**2874552afd8fae92d76a4bc77ac | 0.081,0.054,1.624,-1.533,-1.774,0.253 | 0.021,0.086,0.372,-0.360,0.474,0.179 | 0.039,0.066,-0.865,0.893,0.650,1.138 | 0.014,0.046,-0.195,-0.220,0.919,1.562 |
| **5a9a6**f6a9dc685f82f6ac3a5be72c354 | 0.032,0.052,1.961,-1.847,-1.730,-0.380 | 0.051,0.080,0.567,-0.617,-1.142,-1.685 | 0.051,0.065,-1.932,0.343,1.179,1.476 | 0.077,0.034,-0.880,0.695,0.889,0.778 |
| **5aa05**82ab7bafab285fdeeefa00bb3a4 | 0.099,0.048,1.962,-1.616,-1.593,-0.058 | 0.052,0.046,0.492,-0.336,-0.434,-1.797 | 0.056,0.068,-0.924,0.639,-0.406,0.729 | 0.057,0.100,-0.603,0.413,0.939,0.214 |
| **5aec4**5a6806450c5aa2adc9e32c646ee | 0.038,0.075,1.480,-1.068,-1.390,-1.807 | 0.027,0.076,0.965,-1.041,-1.960,-1.552 | 0.045,0.086,-1.761,1.626,1.647,-1.508 | 0.087,0.075,-1.264,1.068,1.753,-0.125 |
| **5aecc**67be29755b847a23074ef9e5e5b | 0.046,0.053,1.807,-1.861,-0.060,-1.283 | 0.068,0.044,0.644,-0.609,-0.559,-0.815 | 0.022,0.039,-1.552,1.250,1.727,0.566 | 0.044,0.035,-1.406,1.288,1.047,1.036 |
| **5af3d**6ada296e1e028df27ae470e4879 | 0.055,0.042,0.974,-0.875,-1.743,0.781 | 0.012,0.042,1.385,-1.645,-0.217,0.011 | 0.071,0.097,-1.612,1.487,0.811,1.285 | 0.013,0.082,-1.081,0.812,-1.032,1.736 |
| **5af4a**61a73f39f625b649e876822dcc3 | 0.085,0.021,0.920,-0.662,-1.249,0.105 | 0.040,0.028,0.288,-0.705,-0.339,1.102 | 0.078,0.033,-0.532,0.167,0.407,1.121 | 0.023,0.089,-0.959,0.632,1.109,1.683 |
| **5b05f**91322aef2e870a3972874c331dd | 0.048,0.071,0.479,-0.411,-0.341,-0.695 | 0.015,0.071,0.356,-0.171,-0.793,-1.378 | 0.094,0.061,-1.261,1.283,0.209,0.341 | 0.043,0.084,-0.677,0.639,1.872,-0.404 |
| **5b3d1**6545370bb34502be97bceb707d5 | 0.059,0.019,1.669,-1.567,-0.678,-0.598 | 0.027,0.044,1.425,-1.598,-0.440,-0.479 | 0.057,0.030,-1.724,1.391,-0.080,1.521 | 0.059,0.072,-0.537,0.215,1.948,0.048 |
| **5b6d8**dd5bbb702de2ade758c7efddb4a | 0.049,0.055,1.076,-1.158,-0.241,-0.262 | 0.029,0.085,0.767,-0.558,-1.383,-1.254 | 0.079,0.070,-1.039,1.219,1.545,-0.269 | 0.096,0.093,-0.403,-0.081,0.701,1.713 |
| **5b7ec**07c57425df6fbcee265f7aedd9a | 0.083,0.025,1.931,-1.642,-1.682,0.505 | 0.028,0.080,1.362,-0.906,-1.257,-1.257 | 0.026,0.085,-1.414,1.911,-0.823,-0.420 | 0.016,0.069,-0.382,0.337,1.298,0.062 |
| **5b996**0aff23d2b73b5418f211bcf39a2 | 0.098,0.013,1.798,-1.417,-1.357,-0.433 | 0.100,0.083,0.459,-0.389,-1.547,0.069 | 0.045,0.063,-0.803,0.119,1.277,1.707 | 0.077,0.076,-0.500,0.054,1.401,1.485 |
| **5ba49**144eb933d40ce206ba242f4bd5e | 0.045,0.050,1.722,-1.800,-0.053,-1.074 | 0.097,0.089,1.932,-1.534,-1.747,0.322 | 0.032,0.051,-0.262,0.312,0.914,-0.553 | 0.089,0.041,-0.166,-0.365,1.598,1.219 |
| **5be17**f699d6431b9245ba5ada95e6ffb | 0.041,0.085,1.837,-1.635,-1.682,-1.241 | 0.085,0.064,0.698,-1.205,-0.746,-0.311 | 0.039,0.083,-1.911,1.204,1.255,0.631 | 0.062,0.026,-0.680,0.487,1.681,1.884 |
| **5be36**4e2fedcacf9586c6605a88c783e | 0.069,0.065,1.015,-0.906,-1.894,0.348 | 0.021,0.065,1.683,-1.863,-1.157,0.889 | 0.076,0.089,-1.769,1.872,-0.324,0.358 | 0.046,0.095,-0.542,0.110,0.864,1.190 |
| **5bf80**7c56f88a467bb6b3245cdf25cfc | 0.077,0.059,1.126,-0.931,-1.674,-0.130 | 0.027,0.051,1.738,-1.333,-1.588,-0.247 | 0.088,0.055,-1.211,1.096,-0.384,1.879 | 0.076,0.062,-0.559,0.008,0.418,1.526 |
| **5c029**994397838af52a2764698eff778 | 0.060,0.019,1.827,-1.704,-1.925,0.121 | 0.025,0.078,0.729,-0.625,-1.522,-1.088 | 0.094,0.039,-1.376,0.931,-0.493,0.527 | 0.094,0.050,-0.463,0.150,1.082,0.200 |
| **5c419**1c8537c6252488bcf80f251f957 | 0.038,0.059,1.491,-1.513,-0.114,-1.526 | 0.036,0.052,1.937,-1.377,0.380,-0.199 | 0.084,0.099,-1.624,1.837,1.857,1.344 | 0.093,0.068,-1.814,1.225,1.791,1.627 |
| **5c57b**63255666ff731711efa3a1e6e1d | 0.060,0.070,1.557,-1.353,-1.760,-0.153 | 0.070,0.075,0.344,-0.274,-1.011,-0.850 | 0.063,0.040,-1.221,0.160,1.554,1.917 | 0.048,0.051,-0.412,0.259,1.771,0.460 |
| **5c67c**4fcebecba411f5607b1ed0fb675 | 0.088,0.034,1.543,-1.240,-1.386,0.216 | 0.031,0.083,1.210,-1.050,-0.708,-1.234 | 0.085,0.059,-1.925,1.300,1.214,1.094 | 0.064,0.069,-1.675,0.986,1.966,1.629 |
| **5ca92**4bf416215eddce1f44ab498144b | 0.015,0.096,1.100,-0.810,-0.465,-1.183 | 0.042,0.086,0.797,-0.455,-1.080,-0.135 | 0.063,0.059,-0.827,1.127,0.315,0.623 | 0.020,0.094,-0.991,0.435,0.888,1.784 |
| **5cbb9**d4611a087a8edb1c6cdcd0201e7 | 0.090,0.045,1.038,-0.852,-1.446,0.028 | 0.022,0.054,1.954,-0.980,-1.260,-0.221 | 0.070,0.090,-1.202,1.914,0.455,0.519 | 0.086,0.064,-0.204,-0.585,1.870,0.078 |
| **5cd89**c96893232324d18bd5937b28fa7 | 0.082,0.023,1.649,-1.768,-1.936,1.314 | 0.062,0.015,0.345,-0.231,-1.053,-1.197 | 0.059,0.060,-0.646,0.624,1.875,-0.823 | 0.081,0.059,-0.090,-0.365,1.985,1.603 |
| **5d09e**4bdde96655270ae4871c42deedc | 0.087,0.064,1.530,-1.129,-1.845,-1.726 | 0.014,0.046,1.945,-1.308,-1.941,0.939 | 0.095,0.090,-1.684,1.892,1.554,0.542 | 0.054,0.055,-0.691,0.172,1.952,0.827 |
| **5d23a**39ac63594433e4f4e040aadd6a2 | 0.043,0.088,1.333,-1.213,-0.171,-1.489 | 0.083,0.090,0.949,-0.550,-1.023,-0.203 | 0.055,0.075,-1.038,1.884,-1.202,0.627 | 0.076,0.071,-1.154,1.014,1.633,-0.070 |
| **5d359**39bb4e8f624082fbee47538cb77 | 0.086,0.017,0.802,-0.864,-0.508,-0.190 | 0.078,0.060,0.794,-0.868,-1.574,0.547 | 0.044,0.056,-0.941,0.819,-1.052,0.608 | 0.061,0.075,-0.474,0.150,1.276,1.252 |
| **5d3a5**c9704504a83c782468ab0900682 | 0.089,0.044,1.956,-1.913,-1.117,-0.117 | 0.016,0.034,0.693,-0.755,1.672,0.042 | 0.065,0.061,-1.068,1.075,1.034,1.020 | 0.027,0.078,-0.673,0.265,1.504,1.573 |
| **5d3c6**51b3134886a56b78b5ad6f83c4f | 0.079,0.070,0.470,-0.379,-0.605,0.076 | 0.090,0.069,0.707,-0.602,-1.713,0.114 | 0.074,0.059,-0.568,0.492,-0.176,0.585 | 0.051,0.065,-1.275,1.129,-0.003,0.951 |
| **5d7dc**dbcddb662c1c359abe1f7c7a8ab | 0.092,0.087,1.062,-0.833,-1.906,0.653 | 0.062,0.044,0.716,-0.933,-0.897,-1.104 | 0.051,0.055,-0.799,0.389,1.785,-0.034 | 0.054,0.083,-0.898,0.589,1.366,1.440 |
| **5da4b**7798736d5cb9e8b0c84442def2c | 0.077,0.073,1.907,-1.875,-0.289,-0.611 | 0.021,0.065,0.294,-0.254,-1.372,-0.973 | 0.025,0.064,-0.936,0.183,0.833,1.925 | 0.037,0.058,-0.274,0.156,1.837,0.276 |
| **5dd9f**9e5ca6e561b96baf2a98fd76eb8 | 0.084,0.051,0.960,-0.937,-0.653,-0.904 | 0.065,0.048,0.531,-0.540,-0.974,1.600 | 0.061,0.088,-0.709,1.032,1.192,-0.932 | 0.088,0.060,-0.584,0.076,1.975,0.895 |
| **5dddd**e7ebb39d2ab353954a46e0833de | 0.099,0.022,1.819,-1.424,-1.585,-0.443 | 0.079,0.093,1.964,-1.897,-0.531,-0.041 | 0.041,0.018,-1.687,1.936,1.680,-0.526 | 0.045,0.035,-1.319,1.164,1.911,-0.200 |
| **5de4e**a6449ca4d2f60187abde0cca5b8 | 0.048,0.049,1.229,-1.197,-1.646,0.102 | 0.064,0.075,0.574,-0.805,-0.230,0.079 | 0.078,0.067,-0.985,0.850,-0.046,1.137 | 0.083,0.029,-0.234,0.203,-0.346,1.313 |
| **5deee**87c5c7844a7b7965348157d94ae | 0.014,0.022,1.327,-1.252,-1.084,-1.787 | 0.042,0.047,0.866,-0.869,-0.804,-0.701 | 0.016,0.086,-1.095,1.140,0.197,-0.881 | 0.091,0.093,-0.317,-0.360,1.957,0.631 |
| **5e1ad**213c89f31e490eccedf5392336e | 0.091,0.088,1.132,-1.163,-0.334,-0.852 | 0.098,0.012,1.571,-1.569,0.818,-1.728 | 0.099,0.051,-1.912,1.936,1.891,1.365 | 0.093,0.040,-1.309,0.743,1.662,1.704 |
| **5e256**7c17608a65bd437a8e6b862d821 | 0.081,0.059,1.964,-1.179,-1.820,-0.123 | 0.093,0.012,1.707,-1.184,-0.922,-1.167 | 0.047,0.015,-1.083,0.631,0.954,1.402 | 0.015,0.067,-0.037,-0.548,-0.674,1.298 |
| **5e258**ddb99a01d107a84439b91b1aa9b | 0.014,0.050,1.701,-1.798,-1.111,-0.511 | 0.065,0.061,0.973,-1.184,-0.594,-0.948 | 0.033,0.037,-1.957,1.310,-0.231,1.303 | 0.045,0.078,-1.623,1.301,1.739,1.368 |
| **5e287**d2181e93802c7d85ac8e0ec6661 | 0.088,0.064,0.559,-0.474,0.002,-1.000 | 0.021,0.050,1.437,-1.179,-0.373,-1.453 | 0.071,0.066,-1.712,1.947,0.618,1.264 | 0.067,0.097,-1.079,0.664,-0.110,1.892 |

| | | | | |
|---|---|---|---|---|
| **5e3fc**bd3e97c462644b428aa3c50ca3b | 0.039,0.081,0.892,-0.648,-0.998,-1.285 | 0.073,0.019,1.329,-0.982,0.292,-1.677 | 0.066,0.089,-1.839,1.933,1.868,1.960 | 0.096,0.086,-1.323,0.888,1.520,0.403 |
| **5e46d**f25ab7bb4403d02fb5d4e0141db | 0.071,0.014,1.752,-1.402,-1.405,-0.389 | 0.024,0.074,0.906,-0.896,-0.578,-1.056 | 0.030,0.063,-1.004,0.654,-0.940,0.931 | 0.066,0.031,-0.947,0.560,1.559,1.483 |
| **5e5cd**015ef920418c0839f1a232253d5 | 0.076,0.055,1.366,-1.634,-0.029,-0.395 | 0.070,0.088,0.526,-1.480,-0.802,-0.173 | 0.037,0.055,-1.612,0.487,1.408,0.073 | 0.022,0.066,-0.556,0.433,1.953,0.735 |
| **5e924**b061fc7a36fb10cb07f892e98a2 | 0.023,0.033,1.928,-1.903,-0.841,-0.490 | 0.099,0.039,1.121,-1.125,-0.776,-0.801 | 0.036,0.037,-0.673,0.316,1.792,0.123 | 0.036,0.068,-0.996,0.634,0.489,1.201 |
| **5eb19**ee0436c7cd21e7256c7b469f6ec | 0.089,0.018,1.490,-1.650,-1.204,0.279 | 0.019,0.050,0.487,-0.257,-0.792,-0.792 | 0.084,0.067,-1.281,1.706,1.854,-0.292 | 0.091,0.084,-0.955,0.148,1.969,1.661 |
| **5ec47**a3987dc462364e48f052259f80f | 0.064,0.028,1.566,-1.837,-0.956,0.049 | 0.016,0.027,0.448,-0.992,0.691,0.040 | 0.017,0.083,-1.032,0.476,1.511,1.124 | 0.080,0.056,-1.338,1.137,0.922,1.428 |
| **5ed6a**f076365b2049e1f22a2967e632d | 0.083,0.024,1.700,-1.338,-1.894,-0.123 | 0.016,0.055,1.188,-0.677,-0.637,0.033 | 0.097,0.071,-1.075,1.210,1.068,1.191 | 0.056,0.060,-0.181,-0.616,1.848,1.866 |
| **5f045**888c46095141b25ddcd413b79a2 | 0.099,0.077,1.435,-1.891,-1.136,-0.431 | 0.022,0.014,1.441,-1.394,0.464,-0.557 | 0.050,0.065,-1.632,1.920,1.922,0.635 | 0.088,0.074,-0.845,0.569,1.467,1.338 |
| **5f099**d40485109f266a98d7a45796ccd | 0.068,0.013,1.510,-1.579,-1.928,1.281 | 0.047,0.050,0.645,-0.558,0.151,-1.015 | 0.068,0.072,-0.962,0.916,1.870,0.469 | 0.067,0.071,-0.512,0.216,1.694,0.779 |
| **5f3da**49f65a4c7bf77c1f4b7d09e09c8 | 0.067,0.080,1.489,-1.520,-0.502,-0.978 | 0.015,0.054,0.812,-1.038,-0.579,1.313 | 0.063,0.031,-1.012,0.900,1.167,1.755 | 0.096,0.069,-0.177,-0.034,1.506,-0.416 |
| **5f3f0**4a0f1ba52de95177e75a5460b91 | 0.043,0.065,0.489,-0.359,-1.722,-0.441 | 0.017,0.032,0.698,-0.394,-0.579,-0.798 | 0.034,0.047,-1.288,1.947,1.320,1.634 | 0.089,0.048,-1.152,0.578,1.750,0.446 |
| **5f559**aba820ff5f112e17357bd1a8e01 | 0.098,0.029,1.642,-1.951,-0.867,0.971 | 0.058,0.016,1.116,-1.413,-1.894,-1.167 | 0.073,0.033,-1.254,0.813,1.935,0.309 | 0.072,0.047,-1.125,0.928,1.303,1.397 |
| **5f55d**d4e18996afb6aa40e763b97b512 | 0.045,0.080,0.838,-0.636,-0.971,-0.970 | 0.068,0.068,1.500,-1.021,0.659,-1.167 | 0.097,0.033,-1.569,1.928,1.497,1.407 | 0.087,0.059,-0.521,0.125,1.348,0.667 |
| **5fb24**cf50ba20855eb949f52a3fe3f14 | 0.076,0.012,1.561,-1.877,-0.298,0.424 | 0.096,0.049,0.317,-0.534,1.943,0.465 | 0.077,0.026,-1.067,1.913,1.211,0.224 | 0.016,0.098,-1.345,0.941,1.366,1.905 |
| **5fc0d**ca3d3587ef3d7fb85e0c0a62a93 | 0.076,0.037,1.863,-1.781,-0.087,-1.140 | 0.068,0.074,1.335,-0.664,-1.227,-0.542 | 0.079,0.056,-1.555,1.692,1.870,1.459 | 0.066,0.044,-0.217,-0.205,1.771,0.274 |
| **5fde9**f0c8673680dcf24e37b8210abda | 0.065,0.056,0.232,0.000,-0.334,-1.695 | 0.022,0.032,1.085,-0.664,0.014,-1.868 | 0.098,0.053,-1.383,1.770,1.726,0.499 | 0.071,0.090,-0.208,-0.059,1.342,-0.113 |
| **5ff66**e06cc5bb705c9a28ce145f6fc37 | 0.099,0.052,1.601,-1.656,-1.163,-0.080 | 0.036,0.017,1.050,-0.796,-0.497,-1.396 | 0.078,0.068,-1.591,1.715,1.134,1.369 | 0.023,0.085,-0.134,-0.257,1.540,1.157 |
| **5ffe9**26f2d3220499655872bc9b0e031 | 0.083,0.087,1.610,-1.272,-0.539,-1.913 | 0.057,0.075,1.686,-1.791,1.317,-1.153 | 0.095,0.035,-1.773,1.466,1.897,1.921 | 0.100,0.094,-1.440,0.619,1.567,1.746 |
| **60142**ab4f099919fd6bba70ce7c36104 | 0.071,0.014,1.334,-1.507,-1.102,0.930 | 0.076,0.084,0.422,-0.230,-1.021,-0.602 | 0.055,0.052,-0.337,0.172,0.345,0.501 | 0.094,0.049,-0.958,0.647,1.265,0.801 |
| **6023e**9a5eb24b0b3cc05868cf1a1ad8c | 0.080,0.038,1.734,-1.606,-1.565,-0.468 | 0.016,0.096,0.789,-1.385,-1.317,0.897 | 0.039,0.066,-1.652,1.009,0.551,1.386 | 0.033,0.096,-0.389,0.015,1.187,1.488 |
| **60495**bfccba62573678692f8fb7fa3dd | 0.041,0.039,1.587,-1.339,-1.658,-1.190 | 0.075,0.028,1.545,-1.597,-0.634,-0.799 | 0.096,0.068,-1.244,1.140,-0.073,-0.379 | 0.088,0.064,0.307,-1.079,1.292,1.416 |
| **60787**6b46f5a4ff5df6fb1c6ba0a71f6 | 0.048,0.015,1.851,-1.314,-1.366,-1.533 | 0.085,0.079,1.145,-1.317,-1.888,-0.936 | 0.098,0.065,-1.528,0.203,1.704,-0.465 | 0.025,0.095,-0.709,0.077,1.578,1.893 |
| **60790**f7edf3ba02ad8c9b94c96607ae3 | 0.036,0.070,1.748,-1.497,0.827,-1.447 | 0.030,0.094,0.373,-0.308,-0.473,0.101 | 0.017,0.076,-0.763,0.043,-0.204,1.854 | 0.047,0.047,-0.428,0.261,-0.859,1.170 |
| **60d83**1d25772e02bc9e0ad07929dafc5 | 0.097,0.035,1.665,-1.480,-1.960,0.424 | 0.058,0.033,0.881,-0.681,-1.210,1.022 | 0.077,0.029,-0.640,0.910,1.756,1.394 | 0.098,0.070,-0.124,-0.043,1.952,-1.057 |
| **60e85**efaa76de4ff0c123bd6dbd5423f | 0.076,0.046,1.955,-1.654,-1.473,-0.909 | 0.052,0.081,0.700,-0.365,-1.951,-0.170 | 0.033,0.084,-1.631,1.025,1.328,1.866 | 0.047,0.081,-1.845,1.390,1.870,0.658 |
| **60f03**b86a4d9e6bdfa5a5d0bb333bd62 | 0.038,0.030,1.547,-1.540,-1.790,-0.098 | 0.095,0.022,1.009,-1.567,-0.666,1.811 | 0.074,0.086,-1.248,0.630,0.809,1.664 | 0.090,0.043,-0.553,0.336,1.163,0.608 |
| **60f4d**a94df42a069de587dd016385556 | 0.050,0.015,1.830,-1.842,-1.839,1.088 | 0.013,0.083,1.148,-1.688,-1.333,-1.611 | 0.096,0.014,-0.619,0.119,1.406,-0.435 | 0.075,0.076,0.057,-0.279,1.909,-0.589 |
| **610e4**17e9c0354418a7a5823872fb21b | 0.096,0.037,1.502,-1.600,-0.740,-0.431 | 0.036,0.024,0.304,-0.194,-0.753,-0.726 | 0.077,0.013,-1.328,1.088,1.036,0.740 | 0.050,0.097,-1.114,0.644,1.727,0.779 |
| **61173**da60c15f85c084612eab041eb60 | 0.079,0.068,1.936,-1.705,-1.812,-0.220 | 0.094,0.064,1.447,-1.228,-1.365,-0.171 | 0.041,0.031,-1.831,1.971,0.346,0.912 | 0.048,0.053,-0.727,0.572,1.948,0.424 |
| **6121d**7b195337e28d0cc9ed31f512be5 | 0.054,0.028,1.944,-1.934,-0.735,-0.182 | 0.065,0.057,1.492,-1.459,-1.424,-0.810 | 0.066,0.014,-1.494,1.077,1.775,1.943 | 0.075,0.049,-1.212,0.744,1.951,1.297 |
| **61579**e5539e549c19288200338419841 | 0.093,0.076,1.661,-1.895,-0.043,-0.665 | 0.074,0.047,1.048,-1.340,-0.566,-1.898 | 0.095,0.093,-1.869,0.949,-0.578,1.413 | 0.087,0.090,-0.463,0.014,0.353,1.520 |
| **6168b**0c1d9c72a73dd59a6309bdfa7c4 | 0.089,0.024,1.629,-1.689,-1.104,-0.714 | 0.085,0.049,0.950,-1.932,0.433,1.321 | 0.020,0.011,-1.865,1.518,1.606,1.713 | 0.016,0.061,-1.591,1.359,0.773,1.442 |
| **61c30**94ea22b1dcf044ac79834f004d9 | 0.092,0.012,1.110,-0.973,-1.282,0.865 | 0.038,0.090,0.754,-0.815,-0.854,-0.704 | 0.028,0.093,-1.885,1.135,-1.105,1.399 | 0.053,0.088,-1.205,0.740,1.854,1.862 |
| **61cca**e86c55711ba82291e0ed0c1ae82 | 0.098,0.078,1.703,-1.251,-1.542,-0.936 | 0.062,0.077,0.392,-0.490,0.323,-0.086 | 0.050,0.082,-0.619,0.326,1.385,0.334 | 0.022,0.067,-1.223,0.929,1.930,1.006 |
| **61ce4**e8d11972ee0ffd5e2b781f696ad | 0.100,0.070,1.196,-0.799,-1.769,-0.027 | 0.014,0.024,1.052,-1.344,-0.454,-1.786 | 0.074,0.064,-0.818,0.451,0.447,1.146 | 0.082,0.096,-0.489,-0.124,1.943,1.170 |
| **61d27**a524416160c4c35fa3a5bc34c90 | 0.046,0.042,1.091,-1.112,-0.407,-0.604 | 0.100,0.043,0.496,-0.797,0.005,-1.409 | 0.013,0.061,-0.842,0.455,-0.598,0.520 | 0.050,0.078,-1.442,1.081,0.895,1.440 |
| **61e6a**d737ab5da62e4002ced0485600d | 0.048,0.058,1.577,-1.890,-0.876,-0.145 | 0.096,0.055,0.635,-0.804,-1.215,0.404 | 0.083,0.084,-0.503,0.467,-0.271,-0.397 | 0.098,0.039,-0.372,0.084,1.198,1.257 |
| **62165**c20f3af1d933eb1b79f8c7a896b | 0.056,0.026,1.164,-1.135,-1.706,-0.994 | 0.047,0.036,0.235,-0.142,-0.707,0.250 | 0.065,0.052,-0.475,0.545,-0.292,1.443 | 0.034,0.068,-0.988,0.825,0.852,0.671 |
| **6233d**885468b675fccbe0661017f6560 | 0.053,0.062,0.638,-0.469,-0.969,-0.531 | 0.073,0.088,1.377,-1.649,1.057,-0.841 | 0.031,0.097,-1.858,1.750,1.928,1.909 | 0.058,0.070,-1.490,1.123,1.553,1.206 |
| **627bb**e33ae37a905cebaff26f92e5201 | 0.063,0.046,1.656,-1.680,-0.499,-0.960 | 0.017,0.044,0.491,-0.799,-0.063,-1.303 | 0.031,0.099,-1.951,0.730,1.053,1.808 | 0.063,0.066,-0.661,0.490,1.384,0.647 |
| **628b3**0edcc22e9cf8ab95b413c1437e0 | 0.096,0.086,1.135,-1.232,-0.442,-0.698 | 0.062,0.018,0.213,-0.088,-0.541,-0.038 | 0.087,0.057,-1.111,1.971,1.850,-1.274 | 0.090,0.083,-1.508,1.136,1.874,1.465 |
| **62bc7**2f0ca3baf28306fb978f3b1aea6 | 0.068,0.022,0.414,-0.429,-0.654,-0.136 | 0.028,0.089,0.594,-1.057,-1.058,0.451 | 0.037,0.035,-1.423,0.964,1.464,-0.156 | 0.056,0.059,-0.731,0.427,1.903,1.457 |
| **62f85**d564f3ed354d5f825f31be4123d | 0.014,0.082,0.906,-0.596,-1.726,-1.080 | 0.030,0.020,0.639,-0.226,-1.384,-0.183 | 0.093,0.086,-1.574,1.833,1.859,0.000 | 0.026,0.061,-0.422,0.109,1.840,0.365 |
| **62fb5**8734af6055d6e0ba7da5f87f2ab | 0.044,0.087,1.701,-1.680,-0.356,-0.591 | 0.064,0.040,0.518,-1.539,0.204,-0.222 | 0.098,0.056,-0.756,0.039,1.941,0.135 | 0.044,0.023,-1.339,1.262,1.624,0.677 |
| **6303e**664bdd46b6ae1d0e8eb5bed1b61 | 0.057,0.051,1.153,-1.162,-1.061,-1.077 | 0.019,0.021,0.729,-0.558,-0.410,-1.624 | 0.031,0.062,-0.945,0.956,1.718,1.488 | 0.050,0.070,-1.356,1.087,-0.157,1.830 |
| **63092**0db86e11dc627478e6e06a4f8a2 | 0.060,0.066,1.614,-1.251,-1.658,0.366 | 0.032,0.097,0.595,-0.769,1.200,-1.253 | 0.083,0.049,-1.864,0.265,1.755,1.928 | 0.093,0.094,-0.237,-0.877,1.925,1.688 |
| **631a1**450e9706be44b2e3c29d3359eec | 0.071,0.011,1.703,-0.216,-0.955,-0.338 | 0.086,0.082,0.456,0.044,-1.684,-0.035 | 0.063,0.055,-1.310,-0.041,0.592,0.701 | 0.044,0.030,-0.248,0.044,1.763,-0.335 |
| **6358d**a791b55b6325072333a91bcc4b7 | 0.017,0.054,0.629,-0.546,-1.631,-0.219 | 0.012,0.089,0.951,-0.824,-1.367,0.164 | 0.019,0.085,-1.032,1.020,-0.192,0.887 | 0.078,0.096,-0.796,0.434,1.761,0.406 |
| **63a11**3cdfdeb5f2828f6624d8b58cdfc | 0.055,0.016,1.335,-1.124,-1.731,0.664 | 0.097,0.058,1.374,-1.322,-1.382,-1.817 | 0.094,0.053,-1.813,0.874,0.724,0.839 | 0.060,0.024,-0.732,0.409,1.922,0.470 |
| **63ba6**8ef011d59b8eccbdeff549d57d0 | 0.047,0.093,1.900,-1.311,-0.675,-1.653 | 0.015,0.039,1.761,-1.249,0.245,-1.864 | 0.093,0.080,-1.916,1.499,1.887,0.963 | 0.073,0.065,-1.205,0.393,1.977,0.401 |
| **64121**85c64fd10f2d44833d0c9d4e25a | 0.041,0.023,1.484,-1.345,-1.641,0.581 | 0.089,0.085,0.129,0.211,-1.943,0.123 | 0.070,0.064,-0.694,-0.130,1.551,0.206 | 0.023,0.072,-0.904,0.657,1.577,1.148 |
| **6413f**cdc6721ae9d648984882f89f74a | 0.058,0.076,0.848,-0.811,-1.294,0.030 | 0.085,0.041,0.201,-0.281,0.157,-0.524 | 0.059,0.036,-0.826,0.532,0.724,0.771 | 0.076,0.069,-0.341,-0.238,1.867,1.113 |
| **6415e**946cf43219506c674dcbaeb27cc | 0.039,0.025,1.144,-1.358,-1.194,-0.170 | 0.015,0.015,0.834,-0.927,-1.115,0.881 | 0.078,0.045,-1.178,1.267,1.307,-0.409 | 0.097,0.073,-0.371,0.106,1.636,0.377 |
| **6452e**e22ab945394db7c0dfa90ae08bc | 0.081,0.066,0.902,-0.741,-1.145,-0.252 | 0.026,0.033,0.477,-0.170,-1.852,-1.438 | 0.094,0.055,-1.701,1.928,1.499,-0.221 | 0.086,0.052,-0.957,0.482,0.726,1.600 |
| **645d3**34f039be55f7b4c38542feb76b9 | 0.050,0.092,1.278,-0.959,-1.739,-1.340 | 0.017,0.040,1.039,-1.041,-1.626,1.833 | 0.084,0.088,-1.382,1.943,1.570,1.334 | 0.099,0.072,-0.197,-0.497,1.968,1.499 |
| **6474d**a10b1ceac2745d2e466b7d18ca5 | 0.073,0.029,1.206,-0.694,-1.678,-0.123 | 0.072,0.050,1.444,-1.623,-1.120,-1.286 | 0.066,0.063,-0.995,0.550,0.257,0.108 | 0.048,0.081,-0.074,-0.378,1.926,0.610 |
| **64867**f84e4063e0ace30b828e78b9b26 | 0.056,0.083,0.541,-0.477,0.190,-0.921 | 0.065,0.077,1.307,-1.029,1.444,-0.098 | 0.079,0.087,-0.985,0.864,1.834,1.636 | 0.052,0.053,-0.481,0.066,1.618,0.340 |
| **64920**a44b8022531b93155fead430933 | 0.070,0.046,1.092,-0.947,-0.863,-0.423 | 0.091,0.084,1.779,-1.417,-1.264,-0.695 | 0.051,0.024,-1.736,1.826,-0.380,1.778 | 0.098,0.083,-0.501,0.092,1.722,-0.083 |
| **64e3b**5395c529f5832032a5c572a167a | 0.086,0.095,1.858,-1.373,-0.376,-1.611 | 0.073,0.090,0.968,-0.717,-0.767,-1.735 | 0.065,0.035,-0.748,0.450,1.306,0.451 | 0.048,0.038,-1.357,1.155,0.946,1.476 |
| **64f1d**13574fbc7cb79f6df03c488746e | 0.047,0.059,1.805,-1.660,-1.865,-1.141 | 0.089,0.086,0.688,-0.563,-0.614,-1.052 | 0.079,0.093,-1.260,0.972,1.876,0.617 | 0.053,0.044,-0.462,0.261,1.191,1.705 |
| **65156**7a1264aa24c5a48aabfb71fdee6 | 0.068,0.073,1.916,-1.712,-1.597,-1.581 | 0.069,0.053,1.364,-1.559,-1.623,0.267 | 0.015,0.062,-1.415,1.097,1.492,1.411 | 0.024,0.084,-1.307,0.998,1.544,1.597 |
| **65225**2b70946dee17e3015eb5c76c1c7 | 0.081,0.023,1.169,-1.161,-1.525,0.780 | 0.065,0.028,0.473,-0.293,-0.765,-1.198 | 0.048,0.089,-1.050,0.912,1.801,1.843 | 0.074,0.076,-1.684,1.334,1.888,0.925 |
| **65235**5069f8225367b218966a0c0671c | 0.012,0.043,1.209,-1.418,-1.130,-1.262 | 0.038,0.063,0.713,-0.655,-0.577,0.681 | 0.020,0.037,-0.668,0.965,1.319,-0.250 | 0.058,0.040,-0.371,0.124,1.141,0.153 |
| **6528b**b052f4936587665edb880c1f96a | 0.092,0.030,1.249,-1.208,-0.974,-0.500 | 0.039,0.066,1.097,-0.721,-1.319,-1.136 | 0.044,0.030,-1.433,1.674,1.063,0.096 | 0.054,0.079,-0.983,0.761,-0.578,1.438 |
| **6541b**448e55d6aed3f3afed0c6930a12 | 0.050,0.020,1.972,-1.799,-0.327,-1.465 | 0.080,0.023,0.703,-1.297,-1.138,0.305 | 0.099,0.051,-1.283,-0.001,1.455,1.307 | 0.026,0.085,-0.626,0.340,0.993,0.879 |
| **654d5**0f20fc00d4cc4e5106163e855bf | 0.014,0.022,1.590,-1.811,-0.923,-0.322 | 0.030,0.019,0.259,-0.322,-1.939,0.959 | 0.017,0.016,-0.273,0.322,1.142,-0.494 | 0.046,0.063,-0.596,0.296,1.202,1.783 |
| **65526**32b49ae53eb68039c20b84371e3 | 0.086,0.046,1.743,-1.422,-1.879,-0.089 | 0.018,0.082,1.854,-1.358,1.404,-1.740 | 0.092,0.068,-1.542,1.889,1.295,-0.408 | 0.033,0.066,-1.152,0.806,0.643,1.889 |
| **65693**9752b200e4fadd4637f646f9f41 | 0.045,0.043,1.537,-1.823,-0.809,-0.554 | 0.060,0.054,0.141,-0.178,-0.725,1.157 | 0.056,0.062,-0.262,0.287,0.928,-0.089 | 0.062,0.064,-0.300,-0.040,1.131,1.166 |

| | | | | |
|---|---|---|---|---|
| **658e5**3a9a8701c28e0296c74354a7c84 | 0.046,0.098,1.757,-1.399,-0.390,-1.141 | 0.075,0.035,1.204,-0.427,-1.949,0.053 | 0.054,0.033,-1.577,1.915,1.092,1.738 | 0.058,0.084,-1.163,0.330,1.282,1.372 |
| **65fc3**c3eccc41a1d4abab069af3cdebf | 0.091,0.049,1.656,-1.509,-1.564,0.801 | 0.086,0.080,1.686,-1.836,0.083,-0.538 | 0.052,0.069,-1.621,1.133,-0.036,1.760 | 0.050,0.090,-1.346,0.612,1.915,1.317 |
| **660ac**c32b1a670e1ff1494729d27b4c9 | 0.079,0.012,1.856,-1.953,-1.260,-0.078 | 0.023,0.019,0.608,-0.558,-0.938,0.683 | 0.082,0.031,-0.958,1.237,0.266,0.146 | 0.011,0.047,-0.479,0.172,1.902,1.403 |
| **66733**af00ac71af904b0fb8ec4a08470 | 0.097,0.083,1.651,-1.043,-1.743,-1.675 | 0.042,0.086,1.952,-1.294,-1.221,-1.538 | 0.022,0.012,-1.104,1.162,1.460,1.210 | 0.048,0.094,-0.920,0.307,1.622,1.637 |
| **6675b**5dc75c228b9830ba3f8cd64362a | 0.087,0.081,1.656,-1.777,-1.281,0.277 | 0.032,0.076,0.768,-0.458,-0.094,-0.557 | 0.095,0.049,-0.670,0.801,1.550,0.234 | 0.037,0.074,-0.654,0.300,1.305,1.622 |
| **66765**d9ad1abcd53b4e1740183224135 | 0.066,0.074,1.664,-1.044,-0.158,-1.769 | 0.049,0.092,0.430,-0.685,-0.804,0.489 | 0.077,0.042,-1.902,1.173,0.144,1.470 | 0.049,0.083,-0.607,0.171,1.181,0.458 |
| **66788**8020c50740aee0b8a2a1fc2fc0d | 0.070,0.034,1.743,-1.991,-0.509,-0.138 | 0.017,0.084,1.031,-1.123,-1.057,-1.632 | 0.051,0.065,-1.580,1.103,1.106,0.511 | 0.012,0.071,-0.573,0.314,-0.626,1.818 |
| **6694b**bd0f2c930161f444191f39e7a69 | 0.025,0.037,1.585,-1.846,-0.842,-1.379 | 0.016,0.031,0.692,-0.644,0.788,-0.001 | 0.043,0.066,-1.432,1.887,0.765,1.878 | 0.051,0.068,-0.493,0.290,1.344,0.989 |
| **66c3a**d3f36cc67a11a291392eee6cc52 | 0.084,0.049,0.373,-0.302,-0.706,0.012 | 0.029,0.086,0.794,-0.318,-1.493,0.009 | 0.077,0.042,-0.935,1.891,1.412,1.197 | 0.095,0.077,-0.271,-0.104,0.121,1.891 |
| **67504**67a39f43c76b3acb1ef42178a24 | 0.078,0.035,1.502,-1.296,-1.913,0.648 | 0.095,0.039,1.976,-1.296,-0.318,1.213 | 0.094,0.030,-1.110,1.624,1.922,0.676 | 0.076,0.029,-0.671,0.298,0.807,1.725 |
| **6759b**1e7e970a9c610a83efb1be695ce | 0.064,0.078,0.136,-0.057,-1.343,0.191 | 0.078,0.017,0.925,-0.535,-1.942,-0.875 | 0.023,0.099,-1.865,1.925,1.117,1.682 | 0.091,0.058,-0.544,0.005,1.914,1.634 |
| **675f7**1e65faf71ece6d1685431950e51 | 0.011,0.073,0.665,-0.546,-0.567,-1.040 | 0.017,0.093,0.495,-0.468,0.117,-0.236 | 0.031,0.087,-0.811,0.450,1.207,1.896 | 0.031,0.052,-1.145,0.921,0.935,1.595 |
| **67607**50d8a034d6cc5c0fda9e9031094 | 0.048,0.031,1.501,-1.815,-0.700,-0.632 | 0.044,0.097,0.535,-0.327,-0.557,-1.199 | 0.072,0.065,-0.877,1.220,0.360,-1.717 | 0.092,0.076,-1.418,1.006,1.967,1.451 |
| **676a8**f3d39e14cb7543ef9f4cbfa7ade | 0.049,0.064,1.933,-1.437,-0.733,-0.898 | 0.045,0.014,1.678,-1.522,-1.716,-1.406 | 0.060,0.017,-1.933,1.126,1.645,1.306 | 0.023,0.077,0.012,-0.308,1.622,0.536 |
| **67914**cf2c9ef283b14410761b40419a8 | 0.059,0.040,1.742,-1.826,-0.437,-0.428 | 0.047,0.064,0.597,-1.255,-1.124,-1.050 | 0.028,0.023,-1.634,0.371,1.243,0.409 | 0.091,0.070,-1.094,0.476,1.360,1.550 |
| **67a0d**a5ead754980ea3dd85bdf68173f | 0.071,0.056,0.797,-0.561,-1.122,-1.137 | 0.079,0.055,1.879,-1.787,-0.016,-0.191 | 0.050,0.066,-0.759,0.808,0.557,0.733 | 0.064,0.023,-0.876,0.581,1.802,0.325 |
| **67d6e**56fa698b27d2a21ef50e2837eb3 | 0.042,0.053,1.643,-1.748,-0.570,-1.086 | 0.071,0.090,0.469,-1.287,-0.789,1.188 | 0.063,0.030,-1.826,0.920,0.458,1.796 | 0.098,0.084,-1.211,0.772,1.740,1.029 |
| **67f7d**24f4727e13474ec3399782b4b24 | 0.059,0.025,0.900,0.429,-1.777,-1.094 | 0.024,0.092,0.600,-1.821,-1.829,-0.186 | 0.027,0.059,-1.721,0.276,1.226,0.685 | 0.045,0.048,-0.450,0.130,1.425,0.739 |
| **68062**4539798c00ca5175460696e0914 | 0.029,0.071,1.087,-1.138,-1.133,-0.400 | 0.015,0.025,1.125,-1.435,0.046,-0.200 | 0.044,0.062,-1.343,1.261,1.234,0.963 | 0.066,0.017,-0.415,0.177,1.808,1.413 |
| **68316**cb0dab10fca93cd7a541ce63a7d | 0.051,0.030,1.715,-1.499,-1.659,-0.662 | 0.079,0.082,0.452,-0.201,-0.694,-1.523 | 0.091,0.049,-1.255,0.292,1.017,1.777 | 0.075,0.068,-0.126,-0.310,1.886,1.198 |
| **6839c**83fdc870ae8431ea273cd936668 | 0.081,0.027,1.825,-1.167,-1.523,-1.083 | 0.079,0.026,1.739,-1.439,-1.345,-1.616 | 0.098,0.027,-1.875,1.079,1.111,1.194 | 0.081,0.039,-0.691,0.635,-0.895,1.805 |
| **68549**0ea5f7714b1a19e01840546e894 | 0.022,0.056,1.147,-1.328,1.112,-0.933 | 0.020,0.097,1.135,-0.728,1.112,-1.803 | 0.065,0.047,-1.011,1.148,-0.471,1.505 | 0.097,0.078,-0.595,0.204,1.945,-0.299 |
| **68682**3a43358c8156ce492e2fd5f0caf | 0.089,0.012,1.767,-1.821,-1.233,1.118 | 0.042,0.070,1.459,-1.379,-1.886,-1.689 | 0.042,0.045,-1.522,1.131,-0.000,0.598 | 0.081,0.056,-0.698,0.238,1.661,1.384 |
| **68790**08629f4aac39be7c59615de8d48 | 0.073,0.019,0.966,-0.710,-1.305,-1.091 | 0.038,0.085,1.887,-1.262,-1.366,-0.710 | 0.036,0.044,-1.470,1.875,1.801,0.776 | 0.037,0.044,-0.832,0.746,-0.615,1.130 |
| **68875**cb9212d077665fd3b378aa1ff72 | 0.055,0.044,1.092,-1.000,-0.422,-1.525 | 0.023,0.062,1.440,-1.313,1.770,0.215 | 0.037,0.065,-1.284,1.928,-0.350,0.672 | 0.063,0.037,-0.436,0.068,1.523,1.090 |
| **689d0**52919c26378e3fe1353c2bb6286 | 0.076,0.024,1.820,-1.774,-1.486,-0.381 | 0.062,0.018,0.691,-0.668,-1.212,-1.510 | 0.038,0.064,-0.481,0.277,-0.018,0.593 | 0.064,0.044,-0.520,0.215,1.800,0.402 |
| **68fc7**c2bcb6ce8af5bd1502d6d4854ee | 0.068,0.013,1.993,-1.705,-1.601,-1.090 | 0.046,0.052,0.795,-1.457,-1.964,-0.738 | 0.094,0.035,-1.789,0.348,1.801,1.490 | 0.085,0.043,-1.175,0.881,1.256,1.445 |
| **6900e**8526d6181c9d8f20238488314d2 | 0.070,0.026,1.525,-1.589,-1.940,1.031 | 0.038,0.082,0.212,-0.136,-0.878,-0.240 | 0.026,0.022,-0.658,0.381,-0.040,0.314 | 0.053,0.091,-0.717,0.529,1.217,0.233 |
| **690ea**80034d936df705c008a088ada29 | 0.023,0.043,1.532,-1.380,-0.235,-1.320 | 0.040,0.082,1.423,-0.921,-0.369,-0.869 | 0.097,0.031,-1.739,1.875,1.813,0.181 | 0.058,0.041,-1.222,0.760,1.927,1.001 |
| **6910e**cd3451f9ec3c3603a36c40cf603 | 0.043,0.079,1.835,-1.868,0.212,-1.334 | 0.084,0.092,1.154,-1.673,1.556,-0.990 | 0.040,0.081,-1.839,1.307,1.352,0.848 | 0.033,0.078,-1.398,0.933,0.830,1.822 |
| **694eb**501d874d17ddd2c123edf808f37 | 0.013,0.046,0.965,-0.626,-1.119,-1.766 | 0.024,0.078,1.391,-0.903,-1.171,-0.552 | 0.096,0.072,-1.670,1.505,1.087,0.645 | 0.055,0.048,-0.398,-0.056,0.507,1.601 |
| **6964f**188dadbf7367fc313e84c8b3d28 | 0.088,0.027,1.604,-1.722,-0.615,-1.074 | 0.013,0.042,1.137,-1.904,-1.294,0.312 | 0.041,0.021,-1.726,1.118,-0.797,1.914 | 0.076,0.092,-1.327,1.062,-0.122,1.250 |
| **696dc**f90c35e519757a392a35108037b | 0.095,0.035,1.895,-1.445,-1.840,-0.348 | 0.043,0.062,1.286,-1.484,-1.149,-1.701 | 0.080,0.083,-1.950,1.008,-0.155,1.579 | 0.013,0.067,-0.756,0.671,-0.930,0.506 |
| **69761**e612b03e1dc77801baaee975828 | 0.045,0.025,0.870,-0.843,-0.544,-0.157 | 0.011,0.072,1.211,-0.877,1.155,-1.903 | 0.026,0.067,-0.898,0.647,0.829,1.911 | 0.072,0.061,-0.099,-0.388,1.949,0.564 |
| **6980c**0b7fac5ac32d48834664c33db2c | 0.030,0.039,1.887,-1.432,-1.779,0.197 | 0.086,0.019,1.019,-1.967,-1.891,0.079 | 0.068,0.083,-0.929,-0.073,1.571,-0.038 | 0.084,0.087,0.124,-0.294,1.168,-0.622 |
| **69e31**a6e314d4f5a411b72267aa723af | 0.020,0.035,1.359,-1.392,-0.549,-1.082 | 0.018,0.089,0.512,-0.359,-0.092,-0.219 | 0.044,0.080,-1.474,1.683,1.346,0.295 | 0.053,0.050,-1.924,1.423,1.701,1.820 |
| **6a0ef**03835c7e018227df0177e4c1c75 | 0.030,0.082,1.646,-1.269,-1.250,-1.453 | 0.065,0.054,1.506,-1.148,0.661,-0.187 | 0.095,0.084,-1.187,0.341,1.881,1.976 | 0.057,0.043,-0.284,-0.329,1.749,1.176 |
| **6a41f**0088ea7dfb26e01ac6194341332 | 0.059,0.081,1.154,-1.028,-1.317,-0.913 | 0.025,0.061,0.695,-0.636,-0.288,-1.098 | 0.081,0.095,-1.958,1.961,1.927,-0.965 | 0.026,0.081,-1.655,1.391,0.069,1.570 |
| **6a607**a1f6cdc1a700dc2bb70fa9eedfe | 0.088,0.087,0.251,-0.298,0.127,-0.138 | 0.068,0.089,0.160,-0.314,0.845,0.213 | 0.057,0.066,-1.296,1.473,1.643,0.496 | 0.047,0.048,-0.667,0.301,1.905,1.693 |
| **6ab76**1393a8c6be172ec084a9cf590fe | 0.044,0.045,1.409,-1.697,-1.741,0.018 | 0.062,0.063,0.618,-0.291,-1.591,-0.894 | 0.070,0.027,-0.361,0.551,1.547,0.889 | 0.015,0.034,-0.462,0.389,-1.812,1.536 |
| **6ae4f**cffc7fe7760bbc89dd263f71c28 | 0.092,0.076,1.954,-1.257,-1.480,-0.072 | 0.013,0.093,0.946,-1.886,0.659,-1.460 | 0.071,0.021,-1.628,0.408,0.691,1.764 | 0.035,0.086,-0.427,-0.020,1.944,1.490 |
| **6af08**7352ef5ad5870659591d9faa2de | 0.074,0.019,1.335,-1.269,-1.090,-1.164 | 0.015,0.021,0.745,-1.277,1.003,-0.494 | 0.097,0.063,-1.741,1.362,-0.754,0.952 | 0.067,0.076,-0.747,0.381,-0.257,1.815 |
| **6b0ab**1cfe223bb821489e80176af512c | 0.044,0.061,1.614,-1.393,-1.619,-0.543 | 0.084,0.057,0.757,-0.847,-1.568,-1.281 | 0.080,0.096,-1.076,0.118,1.707,1.162 | 0.079,0.048,-0.248,0.018,0.484,1.586 |
| **6b190**7a71308cd1dd718cfdfa3dcf32f | 0.040,0.083,1.508,-1.218,-0.728,-1.449 | 0.019,0.078,1.065,-0.912,0.791,-1.829 | 0.095,0.044,-0.807,0.657,1.393,0.192 | 0.091,0.076,-0.732,0.119,1.660,1.750 |
| **6b250**e8f139e858458d9b1aa437df533 | 0.041,0.069,1.074,-0.895,-1.796,-1.425 | 0.037,0.032,1.019,-0.741,1.877,-1.080 | 0.095,0.064,-1.062,1.529,1.599,1.296 | 0.082,0.097,-0.447,0.113,-1.021,1.685 |
| **6b34b**d5751465e73cb4b2c4596c707a7 | 0.078,0.071,1.867,-1.593,-1.785,-0.988 | 0.036,0.017,1.197,-1.153,-1.588,-1.333 | 0.098,0.031,-1.882,1.394,1.698,1.459 | 0.097,0.090,-0.780,0.212,1.481,1.306 |
| **6b699**32c89a0a504fbbbc01ea9d851d3 | 0.019,0.097,1.651,-1.302,-1.209,-0.560 | 0.071,0.031,0.730,-1.915,-1.331,0.670 | 0.094,0.086,-1.162,0.427,-0.710,0.649 | 0.028,0.078,-1.119,0.763,0.641,1.393 |
| **6b7e8**8ff4e10f70b87be583b6bc3bbc7 | 0.058,0.029,1.513,-1.613,-1.165,-0.157 | 0.045,0.017,0.574,-0.504,-0.524,-1.432 | 0.017,0.047,-1.702,1.903,1.581,1.376 | 0.048,0.059,-0.438,0.087,1.902,0.832 |
| **6b871**3b7ac66e048e64f3f8a5801a923 | 0.087,0.046,1.154,-1.015,-1.770,1.205 | 0.056,0.057,1.506,-0.746,-0.952,-1.909 | 0.055,0.035,-0.888,1.169,1.574,-1.027 | 0.095,0.027,-0.023,-0.474,1.315,1.829 |
| **6ba3b**805e9aafd373a639f9278f797c5 | 0.012,0.021,1.839,-1.897,-0.570,-1.199 | 0.084,0.025,1.931,-0.826,-1.861,-1.195 | 0.041,0.097,-0.618,1.146,0.874,-0.248 | 0.042,0.052,-0.903,0.440,1.811,1.500 |
| **6bbb2**5580d5d76f20bd9f7c7315085a3 | 0.042,0.061,0.962,-0.986,-0.822,-0.685 | 0.065,0.040,0.390,-0.454,-0.935,-0.193 | 0.082,0.059,-0.856,0.680,-0.338,0.383 | 0.055,0.043,-0.320,0.075,1.821,1.778 |
| **6bc57**6f82eb33e6b485126864b34ac44 | 0.047,0.033,1.758,-1.853,-1.550,-0.727 | 0.078,0.078,0.515,-0.387,-1.051,-0.026 | 0.024,0.091,-1.678,0.537,-0.205,1.971 | 0.029,0.086,-1.315,0.871,1.921,1.263 |
| **6c089**b082193a4fbe1c56d6a117a5931 | 0.066,0.076,1.554,-1.197,-1.201,-0.849 | 0.019,0.036,0.248,-0.321,-1.069,-0.093 | 0.098,0.034,-0.360,-0.120,1.264,1.202 | 0.045,0.089,-0.218,0.287,1.537,-0.801 |
| **6c1a1**78c5e2abf0a78dc5b6e6233fc22 | 0.082,0.028,1.694,-1.798,-0.609,-0.934 | 0.034,0.018,0.438,-0.570,-1.917,0.011 | 0.052,0.014,-1.286,0.601,0.039,1.887 | 0.060,0.059,-0.134,-0.158,1.831,0.203 |
| **6c1d0**1fbe6f5d770ffa45a727118fddb | 0.069,0.017,1.225,-1.464,-0.136,-1.127 | 0.024,0.073,0.843,-0.672,-0.790,-0.155 | 0.074,0.025,-1.974,1.925,1.874,1.113 | 0.045,0.074,-1.716,1.245,1.553,1.445 |
| **6c1d1**e830c876c0496baa1d245fc4a72 | 0.023,0.049,1.750,-1.921,-1.177,-0.781 | 0.070,0.078,1.609,-1.287,-1.962,-0.640 | 0.046,0.020,-1.559,1.804,-1.199,0.254 | 0.060,0.059,-0.704,0.536,1.944,-0.459 |
| **6c3eb**fa316cd5be315d658e6d681973b | 0.017,0.085,1.810,-1.838,0.076,-0.722 | 0.069,0.048,0.638,-1.109,-0.754,1.212 | 0.016,0.027,-1.823,1.198,1.262,0.510 | 0.044,0.014,-0.633,0.384,1.961,1.188 |
| **6c6b1**55bc7721d5b4beb461d90c5475c | 0.013,0.044,1.969,-1.779,-0.202,-1.239 | 0.096,0.023,0.629,-1.974,-1.512,-0.323 | 0.012,0.028,-1.407,0.419,1.726,0.071 | 0.013,0.051,-1.623,1.408,-0.671,1.765 |
| **6c7be**e7ba7707b2efaeb6ddf7c828b28 | 0.082,0.061,1.571,-1.448,-1.355,-1.690 | 0.083,0.072,0.864,-0.329,-0.343,-0.982 | 0.094,0.061,-0.848,1.267,1.392,0.390 | 0.066,0.041,-0.921,0.565,1.525,1.306 |
| **6c874**6a21d155cdc6269a573aa6723a1 | 0.087,0.043,1.331,-1.259,-0.287,-0.598 | 0.069,0.085,1.413,-0.699,-1.906,-1.647 | 0.012,0.086,-1.043,0.743,1.003,1.565 | 0.089,0.046,-0.939,0.349,1.402,1.904 |
| **6cc01**3e6ec21e95e54a7b39ba8886ba7 | 0.061,0.049,1.479,-1.445,-1.203,-0.860 | 0.049,0.026,0.399,-0.417,-1.547,-1.220 | 0.074,0.062,-1.656,0.635,1.944,0.833 | 0.074,0.076,-0.199,-0.272,1.407,1.904 |
| **6cd88**e6bd8dbe8cc7b1c7aa6dba3e8f8 | 0.086,0.061,1.726,-1.732,-1.471,-0.151 | 0.063,0.016,0.736,-0.632,-1.648,1.151 | 0.096,0.097,-0.734,0.410,1.730,-0.083 | 0.088,0.053,-0.486,0.142,1.530,0.890 |
| **6ce21**786cdb875f1d6d79ab913e16d75 | 0.013,0.045,1.848,0.216,-1.282,-1.026 | 0.029,0.079,0.579,-0.500,-0.617,-1.073 | 0.039,0.034,-1.406,0.104,0.154,0.533 | 0.090,0.061,-0.719,0.570,0.664,-0.529 |
| **6cfd4**42de3785120a416829ab5fd9310 | 0.032,0.029,1.451,-1.428,-0.436,-1.334 | 0.015,0.098,0.781,-0.494,1.670,-0.383 | 0.066,0.083,-0.606,0.668,1.424,0.727 | 0.085,0.055,-0.617,0.023,1.919,0.768 |
| **6d0fe**20def39ab48e7c37235dbb8e0b1 | 0.045,0.055,1.417,-1.382,-0.959,0.186 | 0.045,0.061,1.177,-0.336,-1.250,-1.191 | 0.029,0.060,-0.709,1.229,-0.288,1.144 | 0.056,0.060,-1.113,0.453,1.834,1.263 |
| **6d10d**fa1c6874a450f0fa46cbf3fa165 | 0.080,0.071,1.154,-1.025,-1.684,0.393 | 0.082,0.029,1.345,-0.896,-0.877,-0.756 | 0.095,0.030,-1.545,1.932,-0.254,0.661 | 0.033,0.034,-1.195,0.871,1.636,1.317 |
| **6d4a9**6e4b9ec5dbcb5f0d22e1534227d | 0.043,0.027,1.883,-1.839,-1.777,-0.679 | 0.016,0.027,0.875,-1.657,0.047,-1.167 | 0.082,0.034,-1.935,0.755,1.964,1.376 | 0.051,0.075,-0.716,0.409,1.373,1.276 |
| **6d4fe**d3afc86bffeba2083ea273815ba | 0.060,0.028,0.742,-0.598,-1.860,-0.545 | 0.072,0.039,0.610,-1.385,1.867,-0.277 | 0.067,0.053,-0.735,0.189,1.543,1.352 | 0.076,0.012,-1.009,0.734,1.889,-0.539 |

| | | | | |
|---|---|---|---|---|
| **6d543**f46d439e1fd306b6111302b9e5d | 0.044,0.019,1.958,-1.518,-1.680,-0.525 | 0.028,0.030,1.521,-1.858,-1.058,-0.550 | 0.018,0.063,-0.815,0.197,0.475,1.513 | 0.096,0.072,-0.012,-0.210,1.696,-1.607 |
| **6d67a**044dbfdf6800d540263c655241a | 0.053,0.051,1.716,-1.885,-1.604,-0.532 | 0.045,0.068,0.438,-1.010,-0.763,1.393 | 0.079,0.088,-1.231,0.454,0.848,1.892 | 0.099,0.034,-0.159,-0.169,1.811,-0.106 |
| **6d7ce**44843f387593cfb7865ff3e20f2 | 0.053,0.022,1.912,-1.762,-1.755,0.731 | 0.043,0.018,0.557,-0.829,-0.088,-0.860 | 0.031,0.089,-1.095,0.512,1.750,0.889 | 0.019,0.073,-0.639,0.444,1.422,0.596 |
| **6d8a5**7a2ed662d5f09d30f163bc6a46d | 0.078,0.068,1.809,-1.531,-0.726,-1.341 | 0.084,0.025,1.022,-1.722,-1.924,-0.026 | 0.068,0.037,-0.463,0.076,1.863,0.292 | 0.096,0.049,-0.961,0.630,1.891,1.196 |
| **6da46**a67a5e888e81820d746258a48d9 | 0.039,0.018,1.653,-1.838,-1.677,-0.303 | 0.016,0.039,1.374,-1.833,-0.817,-0.956 | 0.041,0.086,-1.114,0.700,-0.181,1.135 | 0.067,0.076,-1.326,0.857,1.826,1.021 |
| **6db7d**585e26d141a18297afca7effe8d | 0.015,0.042,1.528,-1.797,0.093,-0.848 | 0.067,0.018,0.405,-0.636,0.005,-1.760 | 0.056,0.079,-0.822,0.418,-0.187,1.964 | 0.012,0.093,-0.696,0.477,1.677,1.152 |
| **6dcb6**7567ce565d2ad042bf14f1a489f | 0.052,0.040,1.588,-1.391,-1.872,-0.299 | 0.088,0.049,1.831,-0.549,-1.856,-1.301 | 0.078,0.093,-0.803,0.677,1.337,0.996 | 0.024,0.049,-0.718,0.474,-0.064,1.238 |
| **6de7e**2a544f0e53f66112d4d6567ee7a | 0.095,0.031,1.448,-1.136,-1.128,-0.827 | 0.053,0.065,1.262,-0.753,-1.528,-1.024 | 0.076,0.034,-1.477,1.510,0.066,0.427 | 0.079,0.092,-0.648,0.254,0.213,1.119 |
| **6e40b**ce0964e8fb31d890c94dd9c8193 | 0.099,0.043,1.748,-0.991,-1.567,-0.088 | 0.054,0.070,0.661,-1.065,-1.020,-0.525 | 0.038,0.062,-0.677,0.111,1.764,0.177 | 0.050,0.080,-1.227,0.844,1.980,0.868 |
| **6e4ad**148f9393cd0341c0a7dfdfc7679 | 0.052,0.050,1.921,-1.797,-0.752,-1.030 | 0.019,0.093,0.871,-1.529,-1.099,-0.973 | 0.023,0.055,-1.133,0.484,1.507,1.018 | 0.086,0.044,-0.619,0.625,-0.250,0.344 |
| **6e5c7**54b25d55da2e87b8176d684256c | 0.049,0.078,1.886,-1.588,-1.887,-0.683 | 0.044,0.062,0.576,-0.843,-0.678,-0.970 | 0.051,0.029,-1.444,0.633,1.847,1.871 | 0.077,0.071,-1.934,1.541,1.958,1.263 |
| **6e781**e9a3df31a2cfbd36718ef5053b0 | 0.042,0.087,0.654,-0.504,-1.302,-0.532 | 0.093,0.087,0.761,-0.650,-1.279,0.646 | 0.062,0.055,-0.641,0.704,1.501,-0.540 | 0.019,0.092,-0.294,0.040,1.600,0.842 |
| **6e9a1**568b2e70186a1c51877cb8b8ab3 | 0.041,0.038,1.588,-1.438,-1.730,-0.923 | 0.017,0.085,1.124,-1.136,0.799,-0.016 | 0.044,0.091,-1.831,1.965,0.910,0.516 | 0.036,0.080,-1.279,1.065,1.556,0.106 |
| **6ea91**5b85f48304e8f40f06c9d0b5a90 | 0.088,0.078,1.302,-1.031,-0.300,-1.674 | 0.038,0.092,1.304,-0.764,-0.755,-1.081 | 0.099,0.094,-1.012,1.067,0.433,0.523 | 0.092,0.049,-0.460,-0.085,1.122,1.685 |
| **6eba0**5d6140c580af0d3447fbd9255c7 | 0.020,0.081,1.843,-1.833,-1.564,-1.176 | 0.097,0.050,0.509,-0.778,0.356,-0.585 | 0.014,0.026,-1.275,1.112,1.886,1.465 | 0.061,0.084,-1.284,0.876,1.879,1.370 |
| **6ecc9**9b43954f87435517374169f95c4 | 0.045,0.058,1.356,-0.820,-1.568,-1.758 | 0.046,0.015,0.810,-1.074,1.194,-1.908 | 0.089,0.029,-1.874,1.093,1.883,1.976 | 0.088,0.076,-1.870,1.084,1.807,1.694 |
| **6ed28**cdb0fc3f58a45f9716b0b54d5db | 0.013,0.075,1.845,-1.659,-0.662,-1.299 | 0.098,0.016,1.980,-0.888,-1.412,0.859 | 0.070,0.062,-1.211,1.272,1.225,0.739 | 0.091,0.044,-0.454,0.382,0.587,-0.994 |
| **6edd6**9bb4ddb2242b3efbbe3a535c19b | 0.043,0.031,1.772,-1.644,-1.763,-0.089 | 0.033,0.045,1.561,-1.517,0.825,-1.456 | 0.064,0.060,-1.974,1.879,1.330,-0.088 | 0.049,0.028,-0.368,0.187,-0.030,1.171 |
| **6ee19**71895267ac9bc5396d5d47af332 | 0.010,0.057,0.789,-0.755,-0.489,-1.215 | 0.042,0.077,0.577,-0.332,0.194,-1.734 | 0.069,0.079,-1.570,1.826,-0.873,1.779 | 0.020,0.067,-1.054,0.811,0.727,1.470 |
| **6ef2b**91d72d5f231c07151f6e34809b3 | 0.095,0.014,1.233,-1.497,-1.184,1.271 | 0.020,0.030,0.954,-1.055,-1.233,-1.568 | 0.076,0.084,-1.444,1.069,1.881,0.217 | 0.078,0.097,-1.252,1.010,-0.210,1.417 |
| **6f104**6893f5138e1991d2c8a937aadb1 | 0.063,0.060,1.223,1.107,-1.512,-0.528 | 0.069,0.017,0.190,-1.618,-1.512,-0.013 | 0.059,0.037,-1.455,0.114,0.008,0.663 | 0.060,0.099,-0.243,-0.225,1.186,1.396 |
| **6f695**b3fa862f31d17336df9f5770c61 | 0.078,0.054,1.962,-1.844,-1.215,-0.359 | 0.094,0.019,0.561,-0.096,-1.093,-1.343 | 0.100,0.069,-0.954,0.711,1.322,0.526 | 0.083,0.048,-0.317,-0.470,1.954,1.791 |
| **6f819**40444300ad7f10473e49c995444 | 0.042,0.084,1.476,-1.411,-0.754,-1.189 | 0.039,0.026,1.264,-0.737,0.928,-1.368 | 0.060,0.083,-0.795,1.197,1.890,0.485 | 0.093,0.085,-0.926,0.347,0.632,1.919 |
| **6f9d6**61a5394eb263293b49ac7e7e88d | 0.052,0.089,0.843,-0.614,0.414,-1.655 | 0.071,0.068,0.509,-0.660,0.065,-0.801 | 0.069,0.087,-1.501,0.693,1.823,1.898 | 0.053,0.048,-0.482,0.218,1.646,1.276 |
| **6fb2e**bd5ed61c1e385ab06cda40d9c53 | 0.091,0.056,1.764,-1.495,-1.002,-0.085 | 0.096,0.053,0.564,-1.739,0.602,-0.982 | 0.098,0.098,-0.728,-0.161,1.930,0.154 | 0.066,0.099,-0.878,0.260,1.703,1.495 |
| **6fe4d**3b375b511c5fc9724184477ab2f | 0.098,0.024,1.105,-0.568,-1.925,0.421 | 0.041,0.079,0.150,0.116,-1.577,-0.844 | 0.043,0.026,-0.683,0.187,1.763,0.205 | 0.079,0.099,-0.761,0.310,1.942,1.559 |
| **6fe7f**60af4d79d3eee04b17e1bc7c70e | 0.049,0.054,1.046,-0.829,-1.935,-0.502 | 0.016,0.061,1.592,-1.473,0.717,1.105 | 0.027,0.079,-0.944,1.117,1.669,1.512 | 0.072,0.051,-0.367,-0.146,1.939,0.660 |
| **700cb**f90fc373e69c57a5a2ffebf6b75 | 0.047,0.021,0.481,-0.497,-0.812,0.169 | 0.018,0.064,0.607,-0.399,-1.300,-0.628 | 0.025,0.043,-1.185,1.536,1.614,0.274 | 0.081,0.097,-0.636,0.329,-0.015,1.749 |
| **70129**c1687d8f60fc8bbeaa99271fa9a | 0.076,0.019,1.638,-1.570,-1.171,0.376 | 0.077,0.019,1.249,-0.797,1.161,-1.078 | 0.032,0.084,-0.413,0.527,-0.194,1.077 | 0.019,0.090,-0.374,-0.150,1.444,1.263 |
| **70511**31563274d5f6c3cd7181fe67d7f | 0.043,0.059,0.741,-0.665,0.351,-1.082 | 0.080,0.020,1.023,-1.688,-0.668,0.490 | 0.086,0.029,-0.760,0.250,0.501,1.509 | 0.074,0.030,-0.118,-0.211,1.988,0.702 |
| **71038**d41d424213de739cb84dcb1a815 | 0.096,0.013,1.937,-1.997,-1.858,0.812 | 0.066,0.072,0.554,-0.390,-1.247,-0.023 | 0.023,0.069,-0.499,0.576,1.533,0.457 | 0.019,0.039,-1.342,1.261,-0.965,1.541 |
| **71178**453437c3dd9fabede1b8d1551a8 | 0.040,0.065,1.750,-1.397,-0.287,-1.908 | 0.030,0.019,1.521,-1.040,-1.921,-1.029 | 0.096,0.042,-1.915,1.909,1.780,1.370 | 0.098,0.058,-1.197,0.627,1.004,1.406 |
| **7161a**599fc4b9b8a4faf08436281cd8a | 0.049,0.050,1.416,-1.423,-0.644,-0.442 | 0.075,0.058,0.634,-0.499,-0.975,1.005 | 0.077,0.058,-0.511,0.347,1.441,0.087 | 0.064,0.079,-0.220,-0.332,1.581,0.517 |
| **71738**b0ae53a30ee2b38cad4a4383363 | 0.088,0.018,1.438,-1.763,-1.677,-0.102 | 0.091,0.078,1.290,-0.936,-1.217,-0.595 | 0.076,0.059,-1.785,1.193,1.611,0.213 | 0.026,0.044,-0.469,0.198,1.610,1.394 |
| **717ac**4d9a5d3c850f26b672dfa2ff3a0 | 0.099,0.039,1.478,-1.850,-0.748,-0.002 | 0.052,0.050,0.357,-0.169,-1.308,-0.630 | 0.030,0.034,-0.607,0.613,0.096,0.551 | 0.063,0.047,-0.012,-0.225,1.468,1.076 |
| **717b7**9bf6f50b0aa36dafb9476e8e310 | 0.014,0.072,1.136,-1.035,0.827,-1.926 | 0.032,0.018,0.340,-0.622,-1.001,0.263 | 0.023,0.048,-1.373,0.807,1.878,1.175 | 0.092,0.086,-1.325,0.944,1.900,1.544 |
| **7180f**7fbb21919803ea4dd8374151fdf | 0.072,0.074,1.903,-0.965,-1.758,-0.328 | 0.092,0.061,0.473,-0.148,-0.855,-1.319 | 0.045,0.026,-1.503,0.411,0.771,1.971 | 0.027,0.074,-0.953,0.600,0.942,1.356 |
| **7182a**48c34886c39a86e7f7a72b9f15f | 0.098,0.013,1.979,-1.585,-1.829,-0.945 | 0.023,0.046,0.432,-0.265,-1.774,-1.709 | 0.093,0.031,-1.718,0.879,1.938,-0.365 | 0.077,0.088,-1.211,0.617,1.571,1.750 |
| **71943**36c74429c8d36f19b4217912d1d | 0.066,0.063,1.486,-1.103,-1.875,0.016 | 0.011,0.020,0.948,-1.000,-1.013,-1.382 | 0.058,0.066,-1.748,1.070,1.680,1.690 | 0.065,0.020,-1.450,1.048,1.894,1.445 |
| **719af**4dcf135b1e3dfbb7d3e2dd73224 | 0.040,0.023,1.698,-1.470,-1.095,-0.773 | 0.099,0.013,0.344,-0.571,-1.361,-0.259 | 0.098,0.068,-1.922,0.103,1.948,1.939 | 0.059,0.099,-1.380,1.027,1.618,1.446 |
| **71c53**947cfeba14a21e97340a876530e | 0.092,0.065,1.867,-1.623,-1.888,0.062 | 0.097,0.083,0.746,-0.964,1.805,-1.852 | 0.074,0.042,-0.716,0.519,1.069,1.133 | 0.039,0.042,-0.258,-0.161,1.836,1.705 |
| **71c72**bb2cb0fede2d3ddcb159d0c6819 | 0.097,0.079,0.878,-0.611,-1.747,0.053 | 0.065,0.090,0.375,-0.333,0.994,-0.604 | 0.075,0.098,-1.103,1.319,0.148,0.625 | 0.041,0.087,-0.418,-0.173,1.434,1.714 |
| **71cc5**f5206f68e7a5e013689438b0486 | 0.047,0.063,1.494,-1.469,-0.212,-0.722 | 0.080,0.075,1.482,-1.315,-0.625,-0.811 | 0.066,0.088,-0.989,0.996,0.908,0.531 | 0.083,0.093,-0.282,-0.148,1.854,0.382 |
| **71ce4**3d3ff5fbe29ea1e05aa98abfcc9 | 0.062,0.021,1.653,-1.918,-1.185,0.442 | 0.026,0.073,0.935,-0.453,-0.409,0.209 | 0.062,0.020,-0.693,1.925,1.580,0.329 | 0.071,0.072,-1.524,1.306,0.750,0.424 |
| **71dfb**a8e0720b78bca1ed6f5082d5325 | 0.086,0.029,1.736,-1.813,0.263,-0.550 | 0.025,0.090,0.840,-1.051,0.878,-0.097 | 0.033,0.061,-1.802,1.921,1.162,1.913 | 0.021,0.025,-1.204,1.042,1.800,1.055 |
| **71e8a**922f6aa6cf51d62968c1e17ccb5 | 0.065,0.058,1.332,-1.315,-0.266,-0.918 | 0.014,0.079,1.853,-1.787,0.202,-1.213 | 0.035,0.062,-1.903,1.796,-0.099,1.925 | 0.080,0.012,-0.503,0.101,1.775,0.874 |
| **7234f**97244002811a303162f248603d3 | 0.099,0.082,1.739,-1.511,-1.871,-0.531 | 0.076,0.098,1.131,-0.787,-0.051,-1.241 | 0.095,0.035,-0.697,0.583,1.474,-0.468 | 0.022,0.048,-0.719,0.552,1.353,1.156 |
| **7247b**057e8ae31d27ac7c81da7e52648 | 0.060,0.057,1.122,-0.999,-0.347,-1.930 | 0.097,0.075,1.207,-0.888,-1.568,-0.505 | 0.070,0.055,-0.439,0.410,1.251,-0.911 | 0.085,0.062,-0.743,0.331,1.395,1.123 |
| **7248b**dd367f76a8cd312e8455fbb3760 | 0.079,0.016,1.190,-1.003,-1.119,-0.480 | 0.028,0.042,1.298,-0.993,-1.308,-0.532 | 0.078,0.058,-1.624,1.910,-0.346,0.491 | 0.055,0.087,-1.479,0.791,0.719,1.501 |
| **72644**c17207a1ca56c0e4addc7668b78 | 0.098,0.056,1.874,-1.427,-1.785,-0.441 | 0.066,0.033,1.558,-1.011,-1.512,-1.288 | 0.067,0.025,-1.514,1.610,1.014,1.086 | 0.026,0.061,-0.329,-0.073,0.310,1.482 |
| **7281a**47b9942c9d52d821a9fa89d4c23 | 0.056,0.086,1.422,-1.303,-1.067,-0.214 | 0.093,0.012,0.719,-1.685,-0.979,0.290 | 0.077,0.073,-1.876,0.838,1.609,-1.388 | 0.054,0.045,-1.298,1.128,0.427,1.914 |
| **72868**6e6404924f72e0954d2db2bde15 | 0.085,0.022,1.772,-1.322,-1.239,-0.912 | 0.072,0.015,1.211,-1.267,-1.911,0.245 | 0.032,0.089,-1.233,0.957,1.149,-0.353 | 0.057,0.077,-0.529,0.338,0.985,0.155 |
| **72b13**201aeffc045149647922b165adc | 0.059,0.086,1.921,-1.411,-0.419,-1.970 | 0.023,0.015,1.699,-1.401,0.199,-1.468 | 0.060,0.098,-1.334,0.993,1.959,1.024 | 0.096,0.055,-1.180,0.385,1.270,1.527 |
| **72b4e**ef39e165d811eea87bbcfba0f76 | 0.047,0.075,0.673,-0.649,0.003,-1.029 | 0.015,0.086,0.628,-0.348,0.787,-0.570 | 0.099,0.064,-1.109,1.954,1.195,1.925 | 0.086,0.065,-0.781,0.422,1.834,1.114 |
| **732a4**77def55a3d9225d15fa0e482085 | 0.049,0.028,1.474,-1.562,-1.211,-0.127 | 0.050,0.080,0.172,-0.179,-1.887,0.079 | 0.033,0.034,-1.356,0.626,0.875,1.604 | 0.011,0.100,-0.314,0.154,-0.390,1.142 |
| **732ae**516a88704de47ac4eb063045731 | 0.030,0.030,1.301,-1.221,-0.866,-0.748 | 0.080,0.015,0.299,-0.552,0.555,-1.334 | 0.075,0.066,-1.318,0.519,1.816,1.739 | 0.060,0.047,-0.551,0.236,1.624,1.437 |
| **7340c**8e68bd13d769f32c668bdbe6b67 | 0.068,0.053,1.750,-1.879,-0.775,-0.660 | 0.041,0.047,0.609,-0.262,-1.798,-1.141 | 0.024,0.020,-0.921,0.715,1.154,1.359 | 0.019,0.050,-0.897,0.919,1.488,-0.447 |
| **735da**e483fcfcaf9545380e5ea315b8e | 0.048,0.073,1.865,-1.969,-1.054,-0.292 | 0.058,0.074,0.640,-0.246,-1.920,-1.170 | 0.048,0.014,-0.944,0.755,1.108,1.002 | 0.057,0.072,-0.505,0.412,1.254,-0.054 |
| **73be3**461589cc0b38d8a0ee39600ece8 | 0.093,0.020,1.029,-0.764,-1.417,1.105 | 0.093,0.087,1.050,-0.734,-1.961,-0.037 | 0.019,0.056,-1.613,0.695,1.798,0.779 | 0.086,0.019,-0.986,0.430,1.681,1.814 |
| **73cc3**fd2953a11ae383a618ad779718b | 0.048,0.073,1.781,-1.934,-0.197,-0.999 | 0.077,0.016,0.673,-0.688,0.123,-0.023 | 0.054,0.056,-1.230,1.295,1.495,0.442 | 0.056,0.065,-1.379,0.907,1.076,1.844 |
| **740e6**2ff89f0642c1eb319ae76fb9b5f | 0.043,0.080,1.465,-1.274,-1.192,-1.331 | 0.070,0.079,1.581,-1.094,0.331,-1.777 | 0.062,0.062,-1.584,1.879,1.647,0.677 | 0.063,0.047,-0.665,0.330,1.695,0.808 |
| **741c1**3f2cc332aa2e2146cfcbcba06c2 | 0.056,0.060,1.385,-1.559,-1.863,-0.589 | 0.081,0.067,0.536,-0.197,-1.224,-0.285 | 0.094,0.057,-0.580,1.049,-0.344,-1.107 | 0.097,0.060,-0.481,0.117,1.466,0.785 |
| **7427f**3a9a4d3a222b6cb994c7ad06bcd | 0.097,0.021,0.553,-0.543,-1.629,0.729 | 0.018,0.052,0.538,-0.526,-1.329,1.389 | 0.095,0.039,-0.959,0.869,1.389,0.766 | 0.086,0.059,-1.126,0.516,1.955,1.486 |
| **7437e**c82c2c908b4b70267cba1f56bfc | 0.040,0.030,1.738,-1.692,-1.838,0.152 | 0.092,0.085,1.157,-1.178,-0.715,-0.528 | 0.081,0.083,-0.554,-0.111,1.867,1.350 | 0.079,0.057,-0.676,0.577,-0.258,0.983 |
| **745bc**049c31a49d1d28bfd8202edb2f0 | 0.098,0.019,1.444,-1.747,-1.005,0.894 | 0.094,0.082,1.609,-1.007,-1.811,-0.053 | 0.099,0.073,-1.288,1.687,1.047,0.316 | 0.090,0.075,-0.900,0.627,1.716,0.271 |
| **74804**1e9ff4d95466128b7b1391f0deb | 0.062,0.027,1.688,-1.280,-1.936,-0.473 | 0.093,0.086,0.237,-0.304,-1.134,0.519 | 0.037,0.048,-1.190,0.269,1.327,1.974 | 0.059,0.098,-0.110,-0.406,1.873,1.436 |
| **75025**6e28c37b342ae475720a1699f6a | 0.049,0.051,1.629,-1.223,-0.419,-1.201 | 0.098,0.084,0.572,-1.076,0.743,-0.804 | 0.035,0.093,-1.566,0.388,1.558,1.577 | 0.089,0.044,-0.078,-0.183,0.830,-0.247 |
| **75210**79898952e3855d10a34e99cb7ed | 0.092,0.040,0.615,-0.586,-0.070,-1.080 | 0.089,0.090,0.615,-0.940,-0.138,1.147 | 0.073,0.077,-0.570,0.374,1.856,-0.012 | 0.040,0.060,-0.133,0.038,0.766,0.132 |

| | | | | |
|---|---|---|---|---|
| **75285**b0e2f9d513411f3f6487684ecdc | 0.091,0.045,1.551,-1.271,-0.936,-0.591 | 0.052,0.088,0.986,-1.942,-1.754,-0.420 | 0.056,0.096,-1.872,0.824,-0.315,0.634 | 0.087,0.047,-0.541,0.155,1.048,1.591 |
| **753dd**5eaa6be059ddb36d54ce04ce970 | 0.069,0.017,1.116,-1.065,-0.819,-1.067 | 0.020,0.090,0.821,-0.907,1.304,-0.335 | 0.087,0.099,-0.998,0.932,1.594,-0.552 | 0.010,0.072,-0.643,0.297,-0.561,1.783 |
| **75451**a466a2cc830baea912065e7e37c | 0.045,0.014,1.703,-1.895,-0.280,-0.332 | 0.087,0.017,1.537,-1.019,-0.598,0.140 | 0.075,0.071,-1.171,1.893,1.844,-1.281 | 0.022,0.039,-0.850,0.475,1.938,1.689 |
| **75e50**4a5118310088af8f0ab3fc77234 | 0.080,0.046,1.328,-1.321,-1.829,0.739 | 0.092,0.017,0.521,-0.168,-1.593,0.927 | 0.083,0.014,-0.529,1.345,1.101,-0.428 | 0.039,0.046,-0.737,0.624,1.519,0.321 |
| **767d5**2a15c34ee705d799a3143232629 | 0.027,0.022,1.157,-1.526,-0.861,-0.265 | 0.018,0.019,0.671,-0.639,-1.841,-0.998 | 0.013,0.098,-1.027,1.072,1.671,-0.325 | 0.029,0.065,-1.366,1.169,0.147,1.095 |
| **767de**3ecc0dec007efff3b69797e27e9 | 0.054,0.080,1.586,-1.265,-1.419,-0.269 | 0.088,0.079,1.451,-1.186,-1.007,-0.955 | 0.065,0.096,-1.256,0.605,1.164,0.800 | 0.035,0.059,-0.950,0.638,-0.224,1.544 |
| **76830**2bb57c6c16609e53b0e3a59e598 | 0.068,0.023,1.061,-1.067,-1.271,0.908 | 0.062,0.054,0.854,-0.781,-0.942,0.552 | 0.063,0.067,-0.558,0.529,1.684,0.222 | 0.026,0.076,-0.938,0.693,-0.524,1.333 |
| **76d27**7f808bdfdc5ca47b78c090eb3b8 | 0.054,0.069,1.458,-1.862,0.890,-0.288 | 0.032,0.072,0.621,-0.490,0.808,-0.058 | 0.041,0.069,-1.444,1.950,1.084,1.218 | 0.091,0.086,-0.650,0.490,0.754,-0.149 |
| **7700d**a4f4db8847e3a8d49fe385084a3 | 0.011,0.077,1.463,-1.196,-0.008,-1.952 | 0.095,0.030,1.404,-1.853,-1.313,-1.129 | 0.088,0.098,-1.141,0.589,0.256,1.128 | 0.087,0.053,-0.875,0.743,1.700,-0.594 |
| **774b7**e085d7bd4be0e529690c1484711 | 0.083,0.079,1.463,-1.346,-0.636,-0.772 | 0.047,0.032,1.033,-1.137,1.458,-0.814 | 0.036,0.036,-1.743,1.936,1.894,1.247 | 0.076,0.084,-0.362,0.148,-0.777,1.246 |
| **77c75**60af169bd8c16a5b0cf8f9c40b3 | 0.091,0.021,1.261,-1.192,-0.713,-0.102 | 0.017,0.078,1.890,-1.053,-1.962,-1.785 | 0.048,0.076,-1.459,1.355,1.456,1.117 | 0.098,0.012,-1.787,1.179,1.782,1.157 |
| **77dd5**61975f3a1efa0254c73c0c95b8e | 0.047,0.082,0.924,-0.713,-0.453,-0.430 | 0.013,0.042,1.044,-1.084,0.525,0.447 | 0.095,0.090,-1.430,0.444,1.190,1.909 | 0.037,0.068,-0.782,0.566,0.921,0.333 |
| **7806a**254afb142b94090278e6be445c2 | 0.078,0.081,1.773,-1.436,-1.657,-1.887 | 0.069,0.086,0.700,-0.842,-1.872,0.760 | 0.072,0.042,-1.964,1.876,-0.351,0.856 | 0.053,0.069,-1.156,0.913,1.708,0.832 |
| **7820f**05a896caa1b722f2fe7746629ba | 0.096,0.049,1.905,-1.581,-1.090,-0.458 | 0.047,0.056,0.516,-1.284,1.048,-1.850 | 0.066,0.018,-1.542,0.455,1.097,1.436 | 0.070,0.094,-1.771,1.339,1.391,1.319 |
| **78253**042bbf4a8acf1e18a4e5fb8c0b1 | 0.036,0.021,1.960,-1.722,-1.871,-0.535 | 0.077,0.017,0.987,-0.505,-1.997,-1.240 | 0.027,0.037,-0.460,0.357,-0.217,0.343 | 0.065,0.030,-0.045,-0.302,0.571,1.511 |
| **783e5**9c2b2bd192dd37eba29bb39ad84 | 0.043,0.031,0.754,-0.874,0.445,-0.827 | 0.021,0.013,0.742,-0.579,-0.975,-0.631 | 0.029,0.091,-1.141,1.218,0.267,1.554 | 0.045,0.030,-0.859,0.688,1.712,0.153 |
| **78446**e8f3cc64740947c30940fe47a55 | 0.019,0.067,1.931,-1.568,-0.238,-1.912 | 0.040,0.048,1.184,-0.833,-1.832,-1.676 | 0.050,0.059,-1.880,1.888,-0.267,1.121 | 0.068,0.026,-1.018,0.666,1.869,-0.041 |
| **7858f**8a1f172cf8ba64a64f6888a1663 | 0.037,0.039,1.755,-1.248,-1.589,-0.818 | 0.011,0.032,0.767,-1.880,-0.005,-1.760 | 0.086,0.027,-1.274,0.202,1.288,1.487 | 0.049,0.083,-1.415,0.949,1.986,1.424 |
| **78b9d**f6e3de8fe759d8159fca18a65e9 | 0.090,0.021,1.671,-1.556,-1.697,-0.166 | 0.047,0.063,0.568,-0.580,-0.635,0.132 | 0.071,0.052,-1.031,0.824,-0.003,0.886 | 0.020,0.069,-0.672,0.307,1.785,1.836 |
| **78e7b**455f36a5e8cf5d578b17204b407 | 0.065,0.066,1.833,-1.443,-1.347,-1.812 | 0.083,0.048,1.508,-0.882,-1.057,-0.954 | 0.068,0.059,-0.493,0.483,0.366,0.995 | 0.064,0.092,-1.049,0.693,1.572,0.346 |
| **7925c**79deb9e1bd1901aa3a6524876ae | 0.074,0.077,1.800,-1.326,-1.732,-0.421 | 0.084,0.070,0.206,0.111,-0.155,-1.978 | 0.040,0.047,-0.928,0.263,1.142,1.684 | 0.013,0.023,-0.624,0.577,0.943,0.593 |
| **79546**53c7e750aeef79b7b2d80927aeb | 0.051,0.022,1.457,-1.823,-0.872,0.118 | 0.072,0.053,0.171,-0.153,-0.809,1.308 | 0.073,0.058,-0.630,0.652,1.738,0.027 | 0.076,0.042,-0.557,0.295,-0.027,1.842 |
| **798fa**9a1e4bbc78202bc5abc53268b4e | 0.084,0.060,1.214,-1.034,-1.898,-0.018 | 0.045,0.078,1.219,-1.202,-1.242,-0.945 | 0.082,0.041,-1.831,1.360,-0.494,1.660 | 0.073,0.093,-1.064,0.445,1.714,1.534 |
| **79a92**ca7ac5464ff15a498e92a712514 | 0.021,0.028,1.853,-1.833,-0.438,-1.526 | 0.075,0.026,0.908,-1.400,-1.195,0.475 | 0.057,0.053,-1.059,0.137,1.761,1.800 | 0.099,0.090,-0.682,-0.079,1.959,1.480 |
| **79c6d**190080170af308c4a3af9dee09d | 0.072,0.052,1.215,-1.063,-1.359,-0.209 | 0.058,0.077,1.523,-1.463,0.298,-0.231 | 0.072,0.023,-1.645,1.921,0.086,1.666 | 0.031,0.073,-1.073,0.514,-0.331,1.935 |
| **79d0d**bb3c06e0027e367ea0c2eb0ccbc | 0.099,0.083,1.910,-1.277,-1.792,-0.988 | 0.068,0.011,1.508,-1.016,-1.398,-1.670 | 0.089,0.035,-1.908,1.914,1.314,1.074 | 0.079,0.096,-0.581,0.456,0.553,0.052 |
| **7a410**43b497d5724f0c279b5e69c9633 | 0.073,0.048,1.553,-1.393,-1.050,-0.848 | 0.020,0.058,1.123,-0.844,0.472,-0.912 | 0.067,0.039,-1.128,1.234,0.402,1.131 | 0.089,0.080,-0.757,0.042,1.383,1.595 |
| **7a898**131665bebbe034e7a328a0d35e0 | 0.026,0.090,1.680,-1.358,-0.614,-1.575 | 0.024,0.080,0.539,-0.586,0.824,-0.551 | 0.094,0.059,-1.898,1.398,1.789,-0.116 | 0.087,0.039,-0.331,0.108,0.137,1.864 |
| **7a919**68e4468dfd7adec01fe2c35bb9b | 0.099,0.068,0.672,-0.573,-1.174,-0.065 | 0.074,0.063,1.147,-0.408,-1.063,-1.454 | 0.053,0.015,-0.764,1.470,-0.301,1.642 | 0.040,0.083,-0.886,0.510,1.435,1.384 |
| **7aabc**882b4da713712faa01d18269a41 | 0.097,0.058,1.296,-1.046,-1.814,-0.048 | 0.046,0.080,0.831,-1.071,-0.136,0.018 | 0.037,0.034,-0.892,0.272,1.647,1.928 | 0.027,0.066,-0.304,0.019,0.413,1.029 |
| **7aaf8**408a1fbdc717424437ea38fa802 | 0.079,0.098,1.981,-1.701,-1.383,-1.288 | 0.026,0.042,0.645,-0.612,-0.534,-1.734 | 0.018,0.077,-1.848,1.742,1.363,0.850 | 0.017,0.092,-1.346,1.032,1.817,1.544 |
| **7ac5e**19b7bc704193ccef8766a4cacf5 | 0.094,0.019,1.888,-1.628,-1.182,-0.328 | 0.013,0.069,1.830,-1.446,-0.692,0.250 | 0.056,0.037,-1.702,1.959,1.627,0.471 | 0.040,0.048,-0.602,0.344,1.558,0.985 |
| **7ad89**1983d0d16fd7110371d0e3406e8 | 0.062,0.058,1.934,-1.593,-1.855,-1.050 | 0.085,0.022,0.551,-0.687,-0.500,-0.592 | 0.055,0.067,-1.384,1.033,1.204,-1.094 | 0.028,0.080,-1.319,1.096,0.775,1.348 |
| **7b126**62c669cdaeeedd6cd1268f3ebfb | 0.090,0.019,1.499,-1.632,-0.908,-0.048 | 0.014,0.075,0.789,-0.854,0.186,-1.151 | 0.096,0.084,-1.819,1.403,-0.593,1.696 | 0.047,0.088,-1.873,1.637,1.889,0.956 |
| **7b249**dae3ae467e8f1d837927e6c63d0 | 0.042,0.096,1.685,-1.422,-0.105,-1.257 | 0.061,0.030,0.502,-0.454,0.108,0.042 | 0.074,0.084,-1.712,1.736,0.348,1.811 | 0.052,0.067,-1.054,0.794,0.868,0.919 |
| **7b34b**25ac29a17c1188ee85ddd27c001 | 0.084,0.053,0.933,-0.755,-1.054,0.002 | 0.075,0.072,0.546,-0.721,1.098,-1.037 | 0.068,0.050,-1.045,0.896,1.256,-0.125 | 0.020,0.068,-1.110,0.824,1.212,1.513 |
| **7b5d0**482be94d1d2732c44300235d090 | 0.092,0.057,1.751,-1.366,-1.033,-0.156 | 0.036,0.070,1.943,-1.089,-1.038,-1.206 | 0.049,0.075,-1.821,1.813,1.935,1.434 | 0.080,0.060,-0.705,0.411,-0.402,1.496 |
| **7b5f6**cfb546f0482ead8059808c52de4 | 0.054,0.090,0.365,-0.210,-1.762,-0.610 | 0.097,0.074,0.968,-0.437,-0.781,-1.743 | 0.051,0.042,-0.746,0.967,0.922,0.799 | 0.088,0.012,0.004,-0.293,1.871,0.942 |
| **7b673**5d0e3a1c5058ea121e4511c5e19 | 0.098,0.068,1.704,-1.124,-1.797,0.739 | 0.037,0.053,0.207,-0.004,-0.707,-1.753 | 0.036,0.049,-0.593,0.035,1.653,0.672 | 0.029,0.062,-1.129,1.030,-1.094,1.018 |
| **7b74f**29eac50bad289e0ae96963b0e20 | 0.086,0.079,0.956,-0.991,-0.843,-0.166 | 0.067,0.033,0.218,-0.110,-0.939,0.013 | 0.015,0.037,-0.990,1.942,1.493,1.361 | 0.086,0.046,-0.603,0.365,1.351,1.494 |
| **7b899**ef93b1479e1a34f25e5ee4293ef | 0.052,0.034,1.812,-1.523,-1.821,-0.581 | 0.097,0.078,0.638,-1.055,-0.053,-1.213 | 0.011,0.090,-0.432,0.089,1.866,0.682 | 0.013,0.017,-0.012,-0.048,1.526,1.243 |
| **7bb6c**6ada7d5b1746e52d2ad33983266 | 0.014,0.037,1.647,-1.755,-0.384,-1.486 | 0.045,0.089,0.873,-1.485,-1.871,-0.202 | 0.054,0.086,-1.850,0.988,-0.318,-0.271 | 0.044,0.039,-0.577,0.382,1.360,1.798 |
| **7be91**316b76380e9a1ea4ebd4e4579fa | 0.048,0.040,1.040,-0.814,-1.867,-0.435 | 0.033,0.080,0.805,-0.659,-0.606,-0.211 | 0.036,0.034,-1.664,1.790,1.920,0.834 | 0.029,0.053,-1.607,1.296,-1.023,1.817 |
| **7bf79**4e98f00ee22471436eda4b84515 | 0.095,0.037,1.571,-1.422,-1.242,1.080 | 0.074,0.076,1.042,-1.832,0.135,-1.186 | 0.039,0.069,-1.432,0.644,-0.051,0.654 | 0.050,0.074,-1.334,0.914,1.437,1.397 |
| **7c028**999c4f9e80567694e710d7cf8e2 | 0.044,0.031,0.850,-0.554,-0.896,-0.909 | 0.100,0.052,0.688,-0.687,-1.382,-1.915 | 0.096,0.073,-1.869,0.570,-0.123,1.705 | 0.072,0.071,-1.754,1.466,-0.113,1.942 |
| **7c0bd**f47f83b54d9c7e5517a108163f2 | 0.090,0.016,1.457,-1.456,-0.920,0.145 | 0.011,0.041,1.670,-1.426,-0.683,-1.273 | 0.041,0.073,-1.553,1.923,1.303,0.713 | 0.089,0.096,-1.364,0.989,1.859,0.022 |
| **7c0c8**f448b8f7ab81a1a175c80a50933 | 0.033,0.043,0.632,-0.580,-1.178,-0.306 | 0.081,0.085,1.819,-1.347,-1.674,-0.930 | 0.093,0.057,-1.308,1.671,-1.030,0.505 | 0.035,0.069,-1.203,1.173,1.693,-0.375 |
| **7c1c8**e00a98c6338fdf82b6b4ed2ba6d | 0.030,0.066,1.209,-1.165,-0.448,-0.546 | 0.087,0.052,1.562,-1.002,0.377,-1.861 | 0.097,0.095,-1.597,1.883,1.221,0.622 | 0.035,0.065,-0.585,0.082,1.886,0.891 |
| **7c822**02298908c296f32f144fc3a0213 | 0.095,0.012,1.983,-1.984,-1.869,1.049 | 0.017,0.020,0.230,-0.236,-1.396,-0.547 | 0.063,0.030,-0.440,0.292,0.816,0.935 | 0.094,0.052,-0.971,0.851,1.905,-0.813 |
| **7d102**ebd603ceddfd7045719ade98b85 | 0.076,0.043,1.185,-0.956,-1.352,0.127 | 0.055,0.045,1.205,-1.232,-1.021,-1.235 | 0.060,0.055,-1.502,1.062,1.822,0.575 | 0.029,0.096,-0.603,0.312,-0.918,1.623 |
| **7d154**e6af5752e22519d5adaf17ba88f | 0.077,0.040,1.917,-0.729,-1.785,-1.887 | 0.037,0.072,0.417,-0.169,-1.477,-1.751 | 0.022,0.063,-1.539,0.362,1.243,1.303 | 0.062,0.062,-0.893,0.584,0.861,1.577 |
| **7d16e**2104d88ae0baee8c34ec4e49af8 | 0.061,0.026,1.495,-1.466,-1.852,-0.015 | 0.072,0.021,1.624,-1.889,-1.288,-0.453 | 0.013,0.059,-1.595,1.483,1.874,0.486 | 0.099,0.085,-0.603,0.096,1.731,0.821 |
| **7d451**530583effb1b51a4a939dc50311 | 0.082,0.098,1.940,-1.454,-1.312,-1.825 | 0.094,0.014,1.932,-1.863,-1.446,-1.492 | 0.043,0.013,-1.822,1.254,1.889,1.694 | 0.043,0.022,-0.319,0.092,1.787,1.447 |
| **7d5bb**bb2fd2f72f35f6d9745aff1ba14 | 0.048,0.058,1.559,-1.815,-0.906,0.466 | 0.050,0.017,1.001,-1.897,1.446,-1.249 | 0.028,0.085,-1.869,1.115,1.699,1.028 | 0.076,0.080,-1.576,0.939,1.920,1.644 |
| **7d5d7**c16dc2d12a0bc78db101f9da09b | 0.058,0.050,1.755,-1.779,-0.107,-0.993 | 0.050,0.077,1.234,-0.723,-1.665,-0.453 | 0.048,0.091,-1.206,1.135,0.640,1.932 | 0.096,0.039,-0.591,0.161,1.797,0.362 |
| **7d63d**00a8840188f7e3bf5ba566d074b | 0.062,0.056,1.674,-1.088,-1.677,-0.834 | 0.044,0.021,0.549,-0.742,-0.781,-0.700 | 0.098,0.090,-1.610,0.312,0.819,1.858 | 0.058,0.083,-0.566,0.067,1.127,1.558 |
| **7d941**82797f8914f519f3b6b0aa772ca | 0.063,0.020,0.993,-1.015,-1.806,0.264 | 0.042,0.055,0.397,-0.034,-1.949,-0.357 | 0.086,0.098,-0.800,0.247,0.397,1.851 | 0.046,0.027,-1.080,0.927,1.101,0.244 |
| **7d9f3**08c33f16b2dc35a229c6b6cde51 | 0.086,0.046,1.299,-0.639,-1.841,-0.296 | 0.073,0.045,1.059,-1.375,-0.379,-1.634 | 0.059,0.078,-1.979,0.931,1.660,0.746 | 0.097,0.055,-0.156,-0.727,1.939,1.839 |
| **7da5b**e63fbfdfc43562a49c6708ae5d6 | 0.063,0.055,0.872,-0.827,-1.806,-0.875 | 0.014,0.052,0.299,-0.265,1.859,-0.728 | 0.027,0.042,-1.297,1.921,-0.039,1.401 | 0.056,0.043,-1.316,1.256,0.756,0.422 |
| **7da99**07f4c53fd56e2bba76b0f04ec63 | 0.098,0.032,1.647,-1.851,-0.793,1.113 | 0.011,0.050,0.885,-0.818,-1.911,-1.664 | 0.084,0.037,-1.858,1.977,-1.227,0.922 | 0.059,0.065,-1.495,1.371,1.788,-0.055 |
| **7dc9b**18852b7a83ed95f150a64ec1276 | 0.051,0.071,1.761,-1.766,-1.158,-0.504 | 0.045,0.037,0.303,-0.594,-0.427,-0.141 | 0.023,0.035,-1.652,1.066,1.826,0.629 | 0.073,0.050,-1.043,0.690,1.974,1.810 |
| **7de6b**02e831924c3f769fd210b40657f | 0.058,0.048,1.239,-1.975,0.373,-0.511 | 0.080,0.094,0.430,-0.254,-0.955,-0.024 | 0.022,0.033,-1.534,1.981,-0.837,1.238 | 0.063,0.075,-1.438,1.214,1.721,1.272 |
| **7e070**5cca7cc20e5dc6d1db98e2eb4af | 0.048,0.055,0.928,-0.673,-1.768,-1.188 | 0.087,0.020,1.096,-0.801,-1.989,0.732 | 0.022,0.049,-1.509,1.305,1.100,0.937 | 0.094,0.095,-0.684,0.426,1.720,-0.184 |
| **7e0df**f5888b37c881304127c3a3a081e | 0.064,0.046,1.798,-1.600,-1.839,0.186 | 0.036,0.035,1.532,-1.478,-0.689,-1.890 | 0.091,0.012,-1.938,1.933,1.882,-0.661 | 0.012,0.095,-0.580,0.273,-1.055,1.637 |
| **7e5c8**32ae3ccb8cca53abb562399b5ae | 0.078,0.019,1.793,-1.756,-0.429,-0.333 | 0.026,0.076,1.762,-1.208,1.115,-1.439 | 0.012,0.085,-1.102,1.353,-0.294,1.174 | 0.087,0.026,-0.343,-0.185,1.879,0.402 |
| **7e7d7**24f32291f9f06565c6857ee08fd | 0.078,0.047,1.109,-0.692,-1.764,0.884 | 0.040,0.034,0.140,0.020,-1.054,-0.762 | 0.044,0.055,-0.702,0.088,1.409,0.855 | 0.021,0.063,-0.413,0.397,1.523,-0.135 |
| **7e87d**30fca5d3916f7751fb9a206ce71 | 0.029,0.049,1.912,-1.570,-0.859,-1.265 | 0.094,0.081,0.954,-1.589,-0.649,-0.947 | 0.054,0.078,-1.951,0.637,1.155,1.267 | 0.048,0.088,-0.498,0.361,-0.899,0.981 |
| **7ef9d**a0976803e0b4d34d0e9a2c50e1e | 0.089,0.095,1.544,-1.010,-1.608,-0.976 | 0.081,0.075,0.759,-0.953,-0.896,-0.156 | 0.093,0.048,-1.815,0.959,-0.125,1.383 | 0.098,0.026,-0.934,0.505,1.805,1.555 |

| | | | | |
|---|---|---|---|---|
| **7f3b4**2830e50de4f1c704e2b9f370b59 | 0.086,0.061,1.639,-1.669,-1.722,0.179 | 0.017,0.074,0.978,-0.708,-1.557,-1.147 | 0.065,0.032,-1.188,1.212,-0.257,1.633 | 0.051,0.084,-0.595,0.221,1.642,0.900 |
| **7f708**d88ea53ff20efdcd6608771bcc8 | 0.038,0.016,0.844,-0.791,-1.001,-0.512 | 0.030,0.054,1.388,-0.896,-0.885,0.562 | 0.049,0.052,-0.972,1.692,1.181,1.562 | 0.098,0.063,-0.096,-0.665,1.702,1.918 |
| **7f85e**1767b248b4b8fa99f4b6e046308 | 0.032,0.010,1.219,-1.070,-1.914,-0.427 | 0.072,0.052,0.903,-1.386,-0.508,-1.760 | 0.097,0.059,-1.875,0.913,0.022,0.647 | 0.059,0.071,-0.599,0.423,-0.762,1.511 |
| **7fd22**d3b249ebc7ee043ed64a9c4383e | 0.094,0.021,1.458,-1.405,-1.922,0.535 | 0.062,0.045,0.602,-0.528,-0.813,-0.354 | 0.070,0.054,-0.889,0.887,-1.218,0.838 | 0.027,0.096,-0.846,0.504,0.791,1.432 |
| **7fd25**2be3f4ecdefbf470a11465c3551 | 0.016,0.062,1.982,-1.861,0.170,-1.412 | 0.064,0.033,0.717,-1.563,1.199,-1.163 | 0.012,0.027,-1.496,0.913,1.081,0.744 | 0.058,0.047,-0.447,0.209,1.882,-0.213 |
| **7fe8a**8554daf398fc7003e4ccae8609a | 0.073,0.025,1.066,-1.099,-0.229,-1.617 | 0.088,0.050,0.808,-1.147,-0.901,-1.076 | 0.086,0.087,-1.522,0.741,-0.274,1.268 | 0.033,0.078,-0.646,0.384,0.476,1.629 |
| **7ffa8**4ad4644da57c25f9682e9acaec6 | 0.064,0.057,1.758,-1.407,-1.624,-0.480 | 0.085,0.079,0.664,-1.465,-0.483,0.089 | 0.013,0.093,-1.271,0.516,-1.610,1.027 | 0.042,0.053,-0.756,0.564,0.468,1.080 |
| **8060c**d06ec34982a7f50bd833f065b62 | 0.039,0.025,0.729,-0.598,-1.231,-0.548 | 0.098,0.070,0.709,-0.791,-0.208,0.664 | 0.092,0.092,-1.866,1.934,1.356,0.533 | 0.062,0.081,-0.345,-0.022,0.776,1.552 |
| **8062b**5f7dd1702c2cd8da082d8ad8ba5 | 0.047,0.026,1.475,-1.675,-1.127,-0.181 | 0.086,0.021,0.187,-0.408,0.624,-0.559 | 0.055,0.047,-1.221,0.892,1.289,0.854 | 0.091,0.060,-0.769,0.490,1.642,0.399 |
| **80aa6**bc775b13376f0c35872b52043a8 | 0.077,0.030,0.661,-0.806,-0.583,-0.045 | 0.076,0.027,0.261,-0.167,-0.402,-0.978 | 0.029,0.037,-0.622,0.675,1.186,1.923 | 0.088,0.026,-0.242,-0.027,1.784,1.431 |
| **80baf**033476bac4dc3578e691db1c1f4 | 0.093,0.015,0.208,-0.013,-1.795,1.139 | 0.051,0.084,1.914,-1.521,-1.708,-0.481 | 0.042,0.096,-1.856,1.516,1.482,1.487 | 0.082,0.093,-0.610,0.351,0.610,0.740 |
| **80e17**fcc74d45e7b763d677e281b2914 | 0.040,0.022,1.152,-1.132,-1.257,-0.681 | 0.089,0.012,1.344,-1.899,1.961,0.903 | 0.100,0.066,-1.779,1.969,1.925,-1.432 | 0.095,0.049,-1.483,0.986,1.664,0.383 |
| **811d1**042f90509fce8322f835206bb61 | 0.058,0.061,1.452,-1.527,-1.797,-0.392 | 0.016,0.093,0.382,-0.624,-0.679,0.808 | 0.018,0.062,-0.665,0.483,1.596,1.804 | 0.040,0.012,-1.205,1.119,0.577,1.891 |
| **813fa**c3ae2b5856ce2cde4e4eb23be5d | 0.045,0.095,1.969,-1.866,-0.482,-1.145 | 0.068,0.061,0.656,-0.463,-1.487,-0.841 | 0.085,0.075,-1.710,1.022,0.637,1.071 | 0.089,0.058,-0.261,-0.062,1.186,1.142 |
| **8151e**42a06999b03b9a82d9a9c375f16 | 0.081,0.044,1.895,-1.591,-1.859,-0.004 | 0.039,0.068,1.649,-0.623,-1.275,-1.809 | 0.051,0.071,-0.988,1.285,1.410,0.289 | 0.058,0.035,-0.051,-0.324,0.579,1.418 |
| **817db**3d1094fed9bb5cfc99655dfcee3 | 0.011,0.085,1.616,-1.801,-0.435,-0.589 | 0.080,0.053,0.636,-0.358,-0.703,-0.319 | 0.095,0.067,-0.678,1.076,1.865,-0.045 | 0.056,0.050,-1.254,1.189,1.793,-0.751 |
| **817f2**bb15b0aacab477e20dd119f65b7 | 0.039,0.059,1.703,-1.623,-1.026,-0.511 | 0.069,0.023,0.691,-1.483,0.011,-0.509 | 0.046,0.037,-0.546,0.131,1.161,0.686 | 0.029,0.041,-0.003,-0.203,1.713,1.375 |
| **819c4**c03f9a1d360358ac4615f292d1e | 0.054,0.043,1.187,-1.107,-1.894,0.250 | 0.027,0.079,0.937,-0.722,0.957,-1.913 | 0.034,0.071,-1.025,1.088,1.791,1.334 | 0.085,0.085,-1.609,1.386,-0.034,1.720 |
| **81b70**241d221441f8a2864bdad332526 | 0.014,0.066,1.733,-1.809,0.032,-1.842 | 0.043,0.057,1.049,-0.804,0.043,-0.949 | 0.095,0.062,-1.425,1.718,-0.273,1.148 | 0.087,0.073,-1.308,0.801,1.498,1.509 |
| **81ed2**6c0cecd1287bf21d8dca0babb9f | 0.052,0.049,0.755,-0.683,-1.306,0.514 | 0.058,0.028,0.612,-0.393,0.094,-1.804 | 0.051,0.089,-0.549,0.747,1.262,0.233 | 0.078,0.065,-1.169,0.566,1.801,1.619 |
| **81f7b**79ddaf0688fbe6bc103ef0e5741 | 0.053,0.093,1.723,-0.664,-1.967,-0.944 | 0.091,0.058,0.593,0.074,-1.904,-1.861 | 0.041,0.031,-0.972,0.178,1.038,1.557 | 0.098,0.082,-0.175,-0.329,1.533,0.945 |
| **8213c**f7dcf27886e8240007761c92a2e | 0.054,0.062,0.938,-0.823,-1.103,-0.870 | 0.087,0.016,1.276,-0.733,-0.914,-0.900 | 0.077,0.091,-0.692,0.836,0.481,1.675 | 0.042,0.092,-0.352,-0.186,1.206,1.453 |
| **82190**25df690200dbf8fe82a6d149c66 | 0.034,0.014,1.929,-1.841,-0.940,-0.641 | 0.044,0.057,1.932,-1.006,-1.915,-1.457 | 0.074,0.090,-1.912,1.959,1.435,1.720 | 0.043,0.078,-0.897,0.406,0.918,0.950 |
| **823b5**ade4e31662021bc1c2392b3c68c | 0.099,0.051,0.785,-0.637,-0.446,-0.364 | 0.089,0.038,1.534,-1.226,-1.455,0.893 | 0.053,0.096,-1.384,0.993,1.934,1.686 | 0.026,0.040,-0.656,0.517,0.941,0.856 |
| **82515**17986ea6af3471a483ea7f95a1e | 0.064,0.033,0.424,-0.141,-1.680,-0.450 | 0.026,0.047,1.381,-0.981,-0.040,-1.631 | 0.055,0.070,-1.800,1.778,1.938,1.641 | 0.084,0.094,-0.160,-0.824,1.359,1.899 |
| **82ddd**d8aab8025ae2a4171a846c12b67 | 0.092,0.017,1.590,-1.046,-1.750,-0.326 | 0.017,0.096,1.533,-1.423,-1.642,-1.630 | 0.091,0.063,-1.934,1.176,1.643,0.213 | 0.058,0.074,-0.184,-0.278,1.457,0.914 |
| **82e55**2e454cf354d18291424c755736e | 0.098,0.066,1.441,-1.514,-1.865,0.175 | 0.063,0.057,0.654,-0.363,-1.948,-0.381 | 0.078,0.082,-1.250,1.694,1.341,-1.122 | 0.081,0.077,-1.041,0.665,1.860,0.832 |
| **8314c**8d2492fb17c589fbea4ce4e2b77 | 0.012,0.086,1.860,-1.612,-0.707,-1.557 | 0.075,0.050,1.969,-1.750,-1.228,-0.692 | 0.016,0.052,-1.483,1.949,1.171,0.589 | 0.045,0.085,-0.658,0.174,1.467,1.689 |
| **8319c**298b48abdfdd07141cf390f7d1f | 0.066,0.016,1.645,-1.411,-1.840,-0.042 | 0.070,0.094,0.677,-1.494,-0.560,-0.671 | 0.013,0.035,-1.808,0.665,-1.097,1.938 | 0.095,0.088,-0.432,-0.257,1.838,1.820 |
| **83379**6f31bd7c6a7da38cb21ec86e7f8 | 0.094,0.023,1.564,-0.633,-1.146,0.319 | 0.072,0.081,0.481,-1.221,-1.779,-0.176 | 0.043,0.038,-1.974,0.242,0.866,0.792 | 0.082,0.084,-1.777,1.487,1.863,-0.222 |
| **839df**093bab3561cae723b022b8c85d7 | 0.094,0.032,1.887,-1.804,-0.943,-0.210 | 0.075,0.077,0.903,-0.706,-1.837,-1.916 | 0.031,0.076,-1.322,0.495,-0.192,1.091 | 0.091,0.056,-0.417,0.080,1.242,0.770 |
| **839e3**39dd1470fe4bc65ce5d0dbec015 | 0.015,0.063,1.442,-1.509,-1.032,-1.282 | 0.061,0.053,1.421,-1.541,1.688,1.623 | 0.085,0.091,-1.630,1.951,1.935,0.404 | 0.087,0.093,-1.183,0.420,1.000,1.561 |
| **83ae5**167b2a2e3b040ba3cefc06a813b | 0.050,0.065,1.431,-1.155,-1.861,0.039 | 0.018,0.085,1.332,-1.216,0.503,-1.894 | 0.058,0.019,-1.084,0.819,0.481,1.715 | 0.046,0.013,-0.658,0.442,1.822,1.188 |
| **83ded**cc54d1233f9485eb117c96c91d2 | 0.068,0.025,1.439,-1.257,-0.598,-0.409 | 0.070,0.030,0.181,-0.345,-1.732,1.776 | 0.015,0.022,-0.706,0.136,-0.310,1.933 | 0.076,0.049,-1.647,1.069,1.964,0.667 |
| **83eb4**cb008a52068a4d244bb14eee35b | 0.042,0.047,1.740,-1.522,-0.280,-0.737 | 0.093,0.013,0.700,-1.217,-1.956,-1.180 | 0.021,0.063,-1.627,0.282,1.654,1.945 | 0.069,0.019,-0.162,-0.141,1.747,1.079 |
| **83f17**354920262080c8de0323a745861 | 0.042,0.076,1.824,-1.351,-1.952,-1.586 | 0.043,0.060,1.180,-1.002,0.521,-1.266 | 0.054,0.088,-1.353,1.042,0.255,1.978 | 0.098,0.046,-0.983,0.482,1.826,-0.277 |
| **83ff0**b1493d8a8500a5bbc73c5ce4360 | 0.086,0.022,1.024,-0.772,-1.767,0.687 | 0.059,0.053,0.493,-0.537,-0.174,1.709 | 0.047,0.056,-0.969,1.192,1.529,1.356 | 0.049,0.038,-0.482,0.293,-0.797,1.711 |
| **8429b**6fafe2ce55cac185586784e08b9 | 0.044,0.023,0.822,-0.817,-1.783,-0.295 | 0.022,0.062,0.326,-0.135,-1.394,-1.050 | 0.068,0.096,-0.998,1.009,1.715,0.694 | 0.016,0.097,-1.431,1.263,0.143,0.962 |
| **84390**f146b85141dc2bec6f68dd5aea5 | 0.070,0.055,1.835,-0.357,-1.379,-0.841 | 0.092,0.030,0.463,-1.346,-1.094,-0.029 | 0.066,0.091,-0.597,0.026,1.784,-0.645 | 0.044,0.080,-1.148,0.895,0.374,1.007 |
| **843b9**223d93929f5cadd23cec15ccec6 | 0.075,0.073,1.580,-1.595,-0.489,-0.328 | 0.077,0.085,1.014,-1.488,0.121,-1.152 | 0.025,0.043,-1.731,1.113,1.618,1.520 | 0.048,0.029,-0.888,0.819,1.269,0.290 |
| **844da**2aec2c82d780280a13b52dd2501 | 0.077,0.050,1.637,-1.804,0.129,-0.599 | 0.056,0.049,1.825,-1.381,1.661,0.782 | 0.073,0.082,-1.127,1.611,1.808,0.591 | 0.012,0.061,-0.505,0.388,0.643,0.245 |
| **8462c**ebd9d1c2f34376e25af47ed1813 | 0.038,0.065,1.110,-0.801,-1.949,-0.635 | 0.079,0.088,1.273,-1.443,0.137,-1.386 | 0.040,0.053,-1.156,0.556,1.948,1.396 | 0.015,0.087,-1.054,0.764,-0.207,1.253 |
| **8472d**19aa36e3256d2494665b4ed8cfe | 0.091,0.065,1.653,-1.216,-1.670,0.626 | 0.073,0.077,1.399,-1.312,0.040,0.345 | 0.076,0.067,-1.791,1.847,1.028,0.559 | 0.055,0.073,-1.235,0.802,0.666,0.687 |
| **84862**52e13c6865c311ed3b4888cf76b | 0.062,0.086,1.723,-1.433,-1.067,-0.697 | 0.028,0.045,1.499,-1.075,-1.827,-1.348 | 0.056,0.032,-1.418,1.908,0.606,1.841 | 0.084,0.075,-1.378,0.825,0.903,1.726 |
| **84a70**061bb573157533134526cdd5d9e | 0.051,0.043,1.531,-1.414,-1.094,-0.126 | 0.071,0.055,0.563,-0.845,-0.053,-0.642 | 0.078,0.055,-1.254,0.623,1.714,0.342 | 0.063,0.072,-0.528,0.250,0.612,1.107 |
| **84b5b**54b85838a48623ccc6d4bc3db98 | 0.070,0.023,1.782,-1.672,-1.608,-1.405 | 0.083,0.068,1.700,-1.363,-1.339,-0.905 | 0.095,0.081,-0.998,0.745,0.043,1.261 | 0.094,0.068,-1.271,0.860,1.469,0.546 |
| **84dbd**f6a00f05f84b66a00609efc2bdc | 0.070,0.066,1.326,-1.055,-1.556,-0.458 | 0.093,0.021,1.023,-1.148,-0.132,-1.154 | 0.094,0.079,-0.692,0.407,-0.411,1.297 | 0.069,0.100,-0.344,0.079,-0.253,0.918 |
| **85080**e204e3c56c9b93a5b70684f3249 | 0.012,0.077,1.838,-1.753,-1.145,-1.420 | 0.034,0.071,0.931,-0.526,-1.596,-0.775 | 0.034,0.082,-1.311,1.864,1.920,0.091 | 0.050,0.064,-0.435,0.030,1.869,0.918 |
| **850b1**2f9c6626c48dde741dabe7dcfa0 | 0.074,0.032,1.996,-1.983,-1.224,-0.360 | 0.043,0.026,0.455,-0.489,-1.461,0.385 | 0.015,0.037,-1.332,0.800,1.389,1.942 | 0.049,0.022,-0.077,-0.218,1.460,1.389 |
| **85355**101440942681c323d468e51471a | 0.074,0.047,1.441,-1.094,-1.464,-0.405 | 0.068,0.015,0.631,-1.825,1.896,-0.355 | 0.049,0.081,-1.541,0.516,1.703,1.094 | 0.075,0.040,-1.217,0.869,1.410,1.146 |
| **85545**1fccad42a395f3c3c752c7cf77d | 0.096,0.025,1.669,-1.560,-1.427,0.536 | 0.075,0.060,0.115,-0.063,-1.919,1.174 | 0.016,0.098,-0.954,0.491,0.519,0.481 | 0.012,0.032,-0.574,0.513,1.503,1.028 |
| **8557e**9222218d90a1ba7aa38e68eac8b | 0.076,0.029,0.601,-0.556,-1.321,1.298 | 0.072,0.048,1.473,-0.823,-0.766,-1.898 | 0.059,0.078,-1.062,1.478,1.849,-1.402 | 0.058,0.074,-0.473,0.032,0.493,1.706 |
| **8572e**4f192f8b39fbcdb5501740b4044 | 0.077,0.056,1.835,-1.695,-1.028,-1.892 | 0.026,0.055,0.302,-0.421,-0.867,0.856 | 0.093,0.065,-0.650,0.251,1.480,0.718 | 0.016,0.086,-1.550,1.083,1.585,1.920 |
| **857fa**a3ba42d30d20cfd872cd34c6362 | 0.014,0.038,1.609,-1.446,-1.074,-1.481 | 0.082,0.053,0.253,-0.035,-1.619,0.132 | 0.023,0.053,-0.709,0.254,1.887,0.029 | 0.090,0.042,-0.951,0.704,0.783,0.942 |
| **85af1**8bda2265b875e9e56b7a490e344 | 0.077,0.015,1.214,-1.003,-1.626,-0.171 | 0.059,0.057,0.721,-0.549,-1.087,-1.354 | 0.098,0.068,-1.754,1.195,1.256,0.335 | 0.033,0.015,-0.098,-0.080,1.919,1.507 |
| **85d03**141d7dd617352d202650fb646f8 | 0.036,0.036,1.935,-1.942,-1.930,-0.982 | 0.093,0.047,0.178,-0.197,-0.747,0.611 | 0.022,0.020,-0.685,0.430,1.436,1.110 | 0.066,0.043,-0.402,0.217,0.454,1.902 |
| **8602b**66aa20188999c7cd857cdc4a1fc | 0.075,0.025,0.999,-0.880,-1.556,-0.226 | 0.065,0.028,0.324,-0.398,0.261,-1.373 | 0.047,0.091,-0.928,0.544,0.908,0.748 | 0.066,0.072,-0.389,-0.080,1.706,1.524 |
| **86165**9eddfdc40e4ca3fb13420effadc | 0.059,0.024,1.563,-1.733,-1.869,-0.320 | 0.028,0.049,0.139,-0.136,-0.930,0.036 | 0.098,0.037,-0.210,0.078,1.100,0.122 | 0.014,0.072,-1.150,1.070,1.916,0.587 |
| **86799**1ad11230a6a719aee5eb4ab8b2e | 0.084,0.093,1.714,-1.469,-0.787,-1.574 | 0.078,0.048,1.061,-0.839,-1.373,-0.908 | 0.042,0.020,-0.333,0.247,1.229,0.470 | 0.087,0.094,-0.455,0.151,1.261,0.532 |
| **86d16**0e21aa6e6ff2982aa1335723fe5 | 0.080,0.070,0.903,-0.894,-1.715,0.819 | 0.048,0.061,0.797,-0.520,0.281,1.123 | 0.092,0.015,-0.702,1.297,1.381,0.650 | 0.073,0.030,-1.113,0.818,1.985,-0.460 |
| **86e2d**c103b010eaef45a4d579327c791 | 0.074,0.073,1.362,-1.126,-0.198,-1.500 | 0.067,0.056,0.592,-0.413,-1.673,-1.765 | 0.098,0.080,-1.971,1.268,0.772,0.386 | 0.097,0.064,-0.190,-0.291,1.798,1.348 |
| **86f48**b6a8ead8a3f929bfc99acf4585b | 0.020,0.091,1.586,-1.423,-0.714,-0.601 | 0.047,0.076,1.304,-1.783,-1.317,0.340 | 0.049,0.045,-1.688,1.091,1.446,0.833 | 0.094,0.062,-1.967,1.390,1.907,0.840 |
| **871fc**6691ed0ca2897f1da8117b7da8a | 0.060,0.022,1.830,-1.546,-1.931,-1.681 | 0.077,0.089,0.897,-1.252,0.465,-1.145 | 0.016,0.067,-1.681,0.749,1.495,1.360 | 0.029,0.034,-1.060,0.885,1.885,0.813 |
| **87229**55f5260570177b5283bf2b714c8 | 0.076,0.020,1.632,-1.098,-1.819,-1.636 | 0.063,0.088,0.863,-1.379,-0.883,-0.486 | 0.034,0.059,-1.738,0.732,0.788,0.823 | 0.015,0.081,-0.295,0.142,0.144,0.900 |
| **874ec**031bc9bcea67a4fdcfc96c9ad5f | 0.096,0.066,1.404,-1.443,-1.435,-0.104 | 0.074,0.060,1.335,-1.226,-1.675,1.591 | 0.012,0.027,-1.397,1.831,1.811,1.212 | 0.077,0.042,-1.475,1.307,1.700,-0.487 |
| **879f5**6a479d9ea4025c122e1cc782c06 | 0.074,0.074,1.993,-1.784,-1.774,-1.190 | 0.020,0.018,1.099,-1.340,-1.921,-0.854 | 0.048,0.058,-1.686,1.009,1.288,1.709 | 0.059,0.078,-0.586,0.258,-0.122,1.627 |
| **87aa3**b39658fa337d9678df7f3975e20 | 0.054,0.014,1.815,-0.318,-1.950,-0.486 | 0.073,0.097,0.684,-0.686,-1.333,-1.176 | 0.037,0.034,-1.912,0.142,0.747,1.872 | 0.034,0.084,-1.087,0.695,1.522,1.051 |
| **87b54**265a72626a9797c83287ca44bb0 | 0.037,0.044,1.185,-1.135,-1.905,-1.018 | 0.089,0.016,0.620,-0.759,0.551,0.571 | 0.077,0.026,-0.659,0.562,1.255,0.829 | 0.025,0.062,-1.001,0.685,1.099,1.364 |

| | | | | |
|---|---|---|---|---|
| 8807aafa12440822b4ae7ac6f0dc0b8d | 0.098,0.015,1.970,-1.906,-1.247,0.497 | 0.087,0.011,0.763,-0.606,-1.752,-1.207 | 0.021,0.051,-1.852,0.934,1.047,1.728 | 0.086,0.076,-0.445,-0.229,1.506,1.558 |
| 88161719330768 15f76fd05b9cd4c931 | 0.046,0.011,1.523,-1.461,-1.108,-0.304 | 0.070,0.073,1.375,-1.541,-1.460,-1.691 | 0.064,0.036,-1.927,1.145,0.096,-0.081 | 0.089,0.038,-0.359,-0.137,1.735,1.543 |
| 88349cc63b69a5286beb733b726047ce | 0.094,0.051,1.591,-1.238,-1.840,-0.348 | 0.071,0.046,1.977,-1.927,-0.887,0.536 | 0.032,0.098,-1.729,1.735,0.963,1.940 | 0.088,0.028,-0.556,0.078,1.817,0.776 |
| 884cb1999af0c2d029d447940b8266b1 | 0.043,0.073,1.650,-1.528,-1.751,-0.946 | 0.021,0.062,0.500,-0.523,-1.005,-0.909 | 0.066,0.054,-1.297,0.506,1.881,0.701 | 0.087,0.041,-0.794,0.360,1.887,0.945 |
| 88a4f947a65a413868c8f94c8591f986 | 0.075,0.063,1.445,-1.285,-1.586,-1.068 | 0.037,0.048,1.241,-1.443,-0.533,-1.197 | 0.059,0.088,-1.912,1.368,1.888,1.048 | 0.047,0.049,-1.491,1.433,1.136,-0.113 |
| 88aa687c9854c4ff9180e0febbaa4cbd | 0.081,0.041,1.915,-1.594,-0.923,-1.279 | 0.040,0.077,1.301,-1.756,-1.813,-1.804 | 0.071,0.025,-1.975,0.871,1.546,-0.374 | 0.097,0.030,-0.741,0.327,1.479,1.945 |
| 88c9d075819c162123bc0b58b5a06931 | 0.022,0.085,1.799,-1.402,-1.881,-1.249 | 0.085,0.081,1.964,-1.991,-0.402,0.458 | 0.051,0.096,-1.625,1.203,-0.197,1.833 | 0.099,0.056,-1.588,1.270,1.034,0.318 |
| 88dcec3ea5f7cc7186f8f809a8971d39 | 0.085,0.048,1.150,-1.231,-1.946,0.902 | 0.040,0.077,0.633,-0.425,-0.822,-0.194 | 0.078,0.044,-0.739,0.946,1.122,0.244 | 0.053,0.032,-0.230,0.105,1.103,1.035 |
| 89335ec5ad5b9765000951efe5da11c9 | 0.013,0.090,1.542,-1.326,0.973,-1.952 | 0.018,0.063,1.894,-1.274,-0.462,0.806 | 0.091,0.056,-1.340,1.941,1.727,0.650 | 0.092,0.041,-0.293,-0.119,1.762,-1.296 |
| 8964fc4a952c58c309a8d652f2a7aa45 | 0.089,0.023,1.099,-0.806,-1.888,-0.604 | 0.094,0.012,0.609,-0.340,-1.655,0.957 | 0.088,0.080,-0.904,0.771,1.394,1.489 | 0.015,0.029,-0.373,0.206,1.952,1.894 |
| 8973658085ed18bd864cc04ed2a804e6 | 0.081,0.099,1.478,-1.226,-0.153,-1.884 | 0.083,0.047,1.245,-0.985,-1.725,-0.150 | 0.099,0.092,-1.929,1.752,1.171,1.027 | 0.093,0.057,-0.110,-0.201,1.815,-0.507 |
| 897e9205a727acd2c0095e5000479352 | 0.062,0.047,1.442,-1.015,-1.826,-0.277 | 0.094,0.068,1.493,-1.380,-0.215,-1.438 | 0.071,0.066,-1.717,0.926,1.958,1.377 | 0.037,0.051,-0.565,0.015,1.991,1.657 |
| 89886a43ac8816056cc3df55d8ec7291 | 0.050,0.088,1.432,-1.428,-1.605,0.349 | 0.099,0.033,0.909,-0.796,-0.834,-1.001 | 0.094,0.035,-0.446,0.488,0.264,0.985 | 0.090,0.095,-0.641,0.428,0.979,0.413 |
| 89b2ad0a52e155e2b7efcb83568ab380 | 0.094,0.065,1.332,-1.284,-1.340,0.184 | 0.056,0.015,0.769,-0.356,-1.459,-0.112 | 0.058,0.074,-1.441,1.892,1.852,0.440 | 0.058,0.036,-1.569,1.192,1.936,1.317 |
| 8a936774a71877c1299ab38534741049 | 0.082,0.021,0.351,-0.408,-0.591,1.474 | 0.013,0.015,0.655,-0.591,0.564,-1.086 | 0.097,0.096,-1.261,1.258,0.761,0.708 | 0.060,0.076,-0.698,0.650,-0.270,0.499 |
| 8ac301ac1138f9d7fb54dd2346f6bac8 | 0.011,0.079,1.719,-1.884,-0.612,-0.121 | 0.074,0.088,1.938,-1.830,-1.920,-0.848 | 0.029,0.085,-1.875,1.938,1.484,0.280 | 0.077,0.034,-1.173,0.747,1.848,1.824 |
| 8ac817c5b194dc3f216e555954aca6b3 | 0.047,0.060,1.951,-1.828,-0.517,-0.818 | 0.096,0.091,1.461,-1.197,-0.976,-1.003 | 0.097,0.095,-1.295,0.660,0.230,0.691 | 0.052,0.073,-1.168,0.748,1.027,0.840 |
| 8ad101959efd3688d82b45284ab6c121 | 0.026,0.071,0.816,-0.548,-1.664,-0.577 | 0.084,0.040,0.681,-0.830,-1.708,-1.327 | 0.053,0.063,-1.736,0.836,0.586,1.008 | 0.075,0.077,-0.119,-0.328,1.411,1.588 |
| 8ae6702e1333366777d59c0cc9c0b3cc | 0.064,0.020,1.652,-1.531,-1.928,-0.118 | 0.012,0.071,0.884,-0.472,-1.970,-0.666 | 0.062,0.084,-0.324,0.580,1.292,-0.884 | 0.072,0.073,-1.729,1.316,1.610,1.064 |
| 8b107579d9b2ed374e5178565e27f8ab | 0.046,0.086,1.518,-1.838,-1.043,-1.038 | 0.084,0.026,0.529,-0.554,0.686,-0.693 | 0.098,0.037,-1.749,1.887,1.832,0.822 | 0.050,0.090,-1.881,1.790,1.819,-0.501 |
| 8b22376d11d4239d7110399afe3df867 | 0.049,0.036,0.564,-0.477,-1.938,-0.175 | 0.079,0.075,0.338,-0.731,0.742,1.143 | 0.076,0.067,-1.504,1.519,1.698,1.162 | 0.069,0.064,-0.926,0.736,1.009,0.183 |
| 8b518d15e6c936fc5dc7fa86bbd45a13 | 0.092,0.036,1.334,-1.589,-1.152,0.018 | 0.098,0.041,0.707,-0.964,-0.539,0.934 | 0.058,0.087,-1.694,0.915,-0.254,1.641 | 0.029,0.066,-1.472,1.253,0.384,1.249 |
| 8b78f25eaa8de963ebd8d9b8db95e435 | 0.090,0.044,0.620,-0.520,-0.877,0.076 | 0.042,0.034,0.632,-0.542,-0.428,-0.348 | 0.072,0.038,-0.597,0.407,1.780,1.692 | 0.083,0.066,-0.422,0.093,0.797,1.388 |
| 8bdd683739c4c11ed0ac7f0cf90dfa3d | 0.085,0.026,0.364,-0.260,-1.658,1.368 | 0.069,0.059,0.738,-0.588,-0.678,-0.206 | 0.097,0.033,-0.640,1.001,-0.040,0.597 | 0.066,0.034,-0.337,0.228,0.205,1.465 |
| 8bf72902b6f647cba2110aa98a63be05 | 0.071,0.060,1.901,-1.668,0.007,-1.193 | 0.043,0.088,0.831,-1.535,-0.396,-1.047 | 0.088,0.064,-1.051,0.322,1.059,0.482 | 0.097,0.064,-0.511,0.278,1.920,-0.683 |
| 8c781cc2c9f57cf0bc1e708663beadc3 | 0.034,0.044,1.331,-1.380,-1.663,0.195 | 0.061,0.051,0.138,-0.127,-1.663,0.162 | 0.038,0.016,-0.698,0.383,0.389,0.787 | 0.046,0.049,-0.594,0.462,1.970,0.621 |
| 8c7c1c871fbaf5eaa6e06d47924387d7 | 0.078,0.011,1.570,-1.485,-1.316,0.181 | 0.096,0.049,0.592,-0.268,-1.008,-1.546 | 0.063,0.030,-0.618,0.525,1.345,0.482 | 0.094,0.034,-0.967,0.686,1.929,-0.493 |
| 8c88f858831a99e5a591bf4251fcbecb | 0.093,0.049,1.879,-1.895,-0.257,-1.263 | 0.086,0.091,1.322,-1.559,-1.110,-1.423 | 0.097,0.088,-1.158,0.788,1.322,-1.276 | 0.071,0.052,-0.331,0.166,-0.059,1.708 |
| 8c918199011d0d3b24749f78fee40b86 | 0.083,0.065,1.235,-1.238,-1.108,0.357 | 0.014,0.038,0.323,-0.255,-0.466,-0.081 | 0.012,0.029,-0.822,1.295,0.949,0.679 | 0.037,0.073,-1.674,1.704,1.345,-0.609 |
| 8c99187b5eb36068a2c9f7213a718af1 | 0.012,0.086,1.705,-1.411,1.092,-1.841 | 0.066,0.050,1.000,-1.870,1.555,-1.633 | 0.095,0.074,-1.119,0.466,0.589,1.136 | 0.054,0.079,-0.289,-0.035,0.815,1.376 |
| 8ca35030ce58e495933bd0692017cb3c | 0.052,0.023,1.623,-1.392,-1.192,-0.145 | 0.086,0.092,0.192,-0.147,-1.776,0.583 | 0.077,0.098,-0.591,-0.001,1.554,-0.405 | 0.036,0.011,-0.647,0.554,1.798,-0.193 |
| 8ca3a54e326412c3303c6467252f8cf0 | 0.027,0.016,1.384,-1.676,-1.857,-0.075 | 0.054,0.053,0.473,-0.299,-1.664,-0.097 | 0.029,0.012,-0.161,0.116,1.450,-0.170 | 0.064,0.055,-0.537,0.192,1.843,1.275 |
| 8cd195d0240ba3b4124cf452d1661371 | 0.096,0.063,1.462,-1.066,-1.675,-0.861 | 0.045,0.062,1.948,-1.600,-0.718,-0.580 | 0.062,0.061,-1.581,1.643,1.828,0.722 | 0.084,0.024,-1.001,0.496,1.741,1.636 |
| 8cdf107b69a738dc437aef601c726734 | 0.042,0.021,1.490,-1.618,-1.164,-0.122 | 0.078,0.077,1.864,-1.280,-1.680,-1.075 | 0.048,0.029,-1.199,1.184,1.393,0.217 | 0.069,0.024,-0.025,-0.234,1.828,0.390 |
| 8cff32f1b19c29167a78a572cb7533d2 | 0.090,0.024,0.613,-0.678,-0.718,-0.308 | 0.055,0.080,0.380,-0.314,0.048,-0.413 | 0.040,0.047,-1.679,1.879,0.487,1.398 | 0.072,0.083,-1.082,0.686,1.763,1.319 |
| 8d001284abdd914d6731d3c2158013d0 | 0.095,0.085,0.988,-0.711,-0.737,-1.810 | 0.027,0.079,0.800,-0.885,-1.097,0.715 | 0.090,0.036,-1.294,1.413,-0.358,1.961 | 0.042,0.091,-1.388,0.983,1.910,1.472 |
| 8d2309ba4482cb5ec49ab85e4a57f092 | 0.048,0.014,1.324,-1.271,-1.469,-0.030 | 0.012,0.033,0.577,-0.645,-1.468,-1.246 | 0.097,0.019,-1.843,1.038,1.358,0.874 | 0.012,0.067,-1.160,0.845,-0.520,1.742 |
| 8dcfc216bbf500c3b560a95fd2afa0d9 | 0.074,0.027,1.620,-1.411,-1.927,0.502 | 0.087,0.076,1.412,-0.974,-1.835,-1.219 | 0.056,0.059,-1.765,1.211,-1.092,1.526 | 0.087,0.023,-0.478,-0.061,1.858,1.514 |
| 8dd684085f37b8c4a3704acb5ac676aa | 0.091,0.049,0.577,-0.657,0.698,-1.782 | 0.058,0.015,0.213,-0.438,0.673,-0.921 | 0.029,0.097,-1.537,0.846,1.169,1.177 | 0.020,0.057,-1.032,0.885,0.193,1.300 |
| 8e03887745814e853c4fa1c244d4d742 | 0.048,0.081,1.933,-1.322,-1.653,-1.784 | 0.074,0.096,1.139,-0.728,-1.154,0.015 | 0.094,0.071,-0.921,0.250,1.212,1.708 | 0.069,0.014,-0.518,0.313,0.766,0.647 |
| 8e231190f857539590f0b2d07e4dfbb3 | 0.094,0.047,1.948,-1.792,-1.292,-1.076 | 0.048,0.010,0.220,-0.577,0.799,-0.547 | 0.070,0.042,-0.710,0.368,-0.598,1.361 | 0.093,0.076,-1.368,1.003,1.896,0.823 |
| 8e2d669589e47b3927b3d4f4e0d9c606 | 0.054,0.095,0.907,-0.964,0.485,-0.554 | 0.018,0.031,0.292,-0.204,-0.882,0.166 | 0.055,0.038,-1.040,1.544,1.682,0.964 | 0.043,0.019,-1.162,0.966,1.809,1.534 |
| 8e3d15f3878663f427cc94a840234f12 | 0.061,0.050,0.488,-0.462,-1.731,0.552 | 0.072,0.081,0.414,-0.243,-1.961,0.468 | 0.081,0.049,-0.923,0.676,1.388,0.793 | 0.026,0.079,-0.495,0.430,1.706,0.233 |
| 8e79e4d810c0457efea28315567f4dc7 | 0.090,0.039,1.683,-0.747,-1.791,-0.522 | 0.069,0.042,0.657,-1.719,-0.855,0.272 | 0.032,0.030,-1.435,0.508,1.255,0.598 | 0.060,0.033,-0.569,0.456,0.996,0.278 |
| 8e909cf2fc00d89494ea63b8f158b247 | 0.037,0.083,1.113,-0.884,-1.793,-1.297 | 0.086,0.026,1.972,-1.870,-1.972,-0.506 | 0.058,0.085,-1.876,1.965,1.889,1.652 | 0.094,0.056,-1.391,1.154,1.672,0.275 |
| 8ec2af806f26c6ef1f60d4ec315ad213 | 0.044,0.091,1.950,-1.886,0.739,-1.265 | 0.055,0.036,0.426,-0.318,-1.614,-1.262 | 0.032,0.073,-0.909,0.202,1.813,1.039 | 0.083,0.057,-0.266,0.066,1.641,-0.541 |
| 8ed9cfa5efe4ffcadaf88414b91335d0 | 0.074,0.062,0.706,-0.240,-0.749,-1.942 | 0.020,0.089,1.819,-0.914,1.490,-1.783 | 0.057,0.095,-1.245,1.774,-0.478,0.755 | 0.034,0.065,-0.628,0.519,-0.153,0.436 |
| 8f0b2529640bf8a490fbdfdf8fe880c4 | 0.055,0.013,1.209,-1.344,-1.346,0.185 | 0.054,0.018,1.342,-0.925,-1.084,-1.792 | 0.095,0.071,-0.756,0.914,1.976,-0.829 | 0.071,0.049,-0.480,0.342,0.253,0.979 |
| 8f3bb6610f350875d9dac9fa911ec71f | 0.024,0.077,1.618,-1.878,-0.802,-1.173 | 0.046,0.023,0.285,-0.091,-1.949,-0.222 | 0.074,0.060,-1.205,1.340,1.579,1.421 | 0.018,0.097,-1.403,1.259,-0.232,0.895 |
| 8f48c5bc4503a71371011e392d8addc8 | 0.045,0.047,1.816,-1.742,-0.974,-0.438 | 0.073,0.079,0.452,-1.942,-1.183,1.800 | 0.019,0.096,-0.852,0.333,1.495,0.567 | 0.027,0.080,-1.587,1.249,1.686,1.872 |
| 8f631878bc261f200c71b390950c1fd6 | 0.041,0.038,0.739,-0.507,-1.651,-0.466 | 0.078,0.042,0.629,-0.443,-1.423,-0.859 | 0.026,0.011,-1.054,0.653,0.855,-0.449 | 0.080,0.046,-0.273,-0.318,1.923,1.914 |
| 8f7a9acb4bd630cf0afe6a8ac3afed3e | 0.085,0.068,1.038,-1.051,-0.732,-0.737 | 0.027,0.024,0.708,-0.729,-0.603,-0.258 | 0.033,0.098,-0.359,0.312,-0.466,0.425 | 0.086,0.046,-0.420,0.213,0.693,0.903 |
| 8fb25346310eb0c3489da43780148821 | 0.034,0.026,1.814,-1.529,-0.805,-1.343 | 0.057,0.068,0.721,-1.694,-0.881,0.050 | 0.065,0.078,-0.865,0.352,0.381,-0.080 | 0.091,0.052,-0.189,-0.324,1.938,0.188 |
| 8fff0327e6ce66bb99226a4ca8d89edf | 0.054,0.010,1.104,-1.322,-0.867,-0.005 | 0.074,0.059,0.711,-0.631,-1.550,-0.903 | 0.070,0.038,-0.758,0.456,0.178,0.754 | 0.097,0.067,-0.476,0.180,1.770,-0.241 |
| 9026773f8ccf1c4f54a022cdcc67f2cf | 0.050,0.015,1.359,-1.405,-1.137,-0.641 | 0.018,0.026,1.324,-1.147,-1.344,-1.106 | 0.095,0.049,-1.750,1.899,1.391,1.483 | 0.087,0.035,-0.555,0.337,0.403,1.921 |
| 904e846a6b6c36010cd2c6c043cd106d | 0.041,0.073,1.577,-1.827,-0.156,-0.665 | 0.066,0.073,0.943,-1.056,-1.889,-0.543 | 0.044,0.089,-0.729,0.337,-0.360,1.393 | 0.081,0.072,-1.138,0.775,1.135,1.412 |
| 908d97682d4a8044a1c3ff218cc19ee1 | 0.098,0.044,1.720,-1.529,-0.793,-1.324 | 0.086,0.054,0.717,-0.441,-1.607,-1.425 | 0.070,0.031,-1.199,0.658,1.362,0.508 | 0.081,0.021,-1.587,1.232,1.932,1.436 |
| 9115edbd9379ecd65067145c6cb98e2a | 0.038,0.050,1.057,-0.541,-1.818,-1.070 | 0.046,0.029,0.393,-0.793,-0.782,-1.732 | 0.079,0.049,-1.919,0.749,1.891,-1.502 | 0.043,0.054,-0.266,-0.009,1.295,1.610 |
| 9117f681fa7319de03e57463b90ac28d | 0.099,0.016,1.704,-1.212,-1.736,-0.554 | 0.043,0.072,0.814,-1.757,-1.285,1.238 | 0.084,0.014,-1.196,0.518,1.605,0.547 | 0.054,0.075,-0.824,0.405,1.641,1.915 |
| 91580a1078377b40237c0153cc81f386 | 0.050,0.067,1.747,-1.633,0.256,-1.253 | 0.031,0.046,1.510,-1.089,0.742,-0.229 | 0.081,0.033,-1.860,1.952,1.919,1.911 | 0.067,0.061,-1.743,1.088,0.759,1.656 |
| 9174981d6db32e419d3f4f46c42556fd | 0.060,0.034,1.736,-1.523,-1.958,-0.407 | 0.067,0.069,1.024,-1.201,-1.160,0.489 | 0.039,0.019,-0.810,0.661,1.518,0.792 | 0.067,0.025,-0.792,0.642,1.612,0.557 |
| 91a4df0cc6098f05a8446d78bd22315c | 0.093,0.066,1.701,-1.773,-0.834,-1.030 | 0.098,0.052,1.032,-1.163,-1.895,1.536 | 0.073,0.080,-1.533,1.090,0.340,0.790 | 0.082,0.069,-0.428,-0.016,1.587,1.694 |
| 91a647f06d7f4564cda34f784db8a29f | 0.070,0.045,1.545,-1.349,-1.478,-0.344 | 0.023,0.042,0.253,-0.274,-0.142,-1.554 | 0.064,0.055,-1.788,0.848,1.401,1.039 | 0.038,0.074,-0.226,0.028,1.466,0.909 |
| 91bde65183a7457b1ff89f05d9e26eb0 | 0.086,0.023,1.598,-1.556,-0.982,-0.269 | 0.090,0.087,0.778,-1.397,-0.394,-0.919 | 0.063,0.079,-0.434,0.108,1.597,-0.327 | 0.011,0.077,-0.611,0.535,0.384,0.569 |
| 9201358ce4953c77a4e5a85007426e98 | 0.095,0.026,1.395,-1.332,-1.363,0.657 | 0.085,0.088,1.117,-1.211,-0.267,-1.513 | 0.034,0.090,-0.371,0.127,1.127,0.592 | 0.094,0.073,-0.442,0.111,0.585,1.593 |
| 92068334f12a82280725ae8cfa472fcc | 0.041,0.082,1.665,-0.890,-1.815,-1.491 | 0.059,0.026,1.881,-1.149,-0.307,0.773 | 0.094,0.062,-1.688,1.889,1.248,0.503 | 0.074,0.033,-0.474,-0.781,1.926,1.876 |
| 920719a208b90f677870700e8e94c46b | 0.051,0.023,0.488,-0.427,-0.530,-0.131 | 0.014,0.078,0.682,-0.742,1.858,0.314 | 0.031,0.090,-1.539,1.813,0.765,1.653 | 0.060,0.076,-1.110,0.850,-0.670,1.563 |
| 921ecc93d7c7ea1ca19794001caaf616 | 0.041,0.023,1.424,-1.658,-0.991,0.621 | 0.017,0.065,1.145,-1.485,-1.446,0.655 | 0.014,0.063,-1.073,0.936,1.518,1.726 | 0.090,0.047,-0.777,0.607,0.995,0.728 |

| | | | | |
|---|---|---|---|---|
| **9244e**de2e0c5ba0f3e85557757a67ace | 0.028,0.035,1.081,-1.118,-1.332,-0.167 | 0.013,0.088,1.307,-0.902,-0.612,-0.943 | 0.036,0.081,-1.926,1.814,1.617,1.820 | 0.085,0.089,-1.129,0.402,1.519,0.718 |
| **9278b**8c5bfbfe8a4debcebd99edbecf0 | 0.046,0.059,1.576,-1.025,-1.937,-1.854 | 0.076,0.065,1.037,-1.224,0.537,-1.075 | 0.054,0.064,-1.375,1.096,1.563,-0.368 | 0.068,0.052,-0.170,-0.303,1.496,1.718 |
| **92a23**06dd8f97e9337eacfef61b4794a | 0.023,0.015,1.828,-1.740,-1.494,-0.744 | 0.016,0.090,0.594,-0.489,-1.110,-0.489 | 0.074,0.013,-0.775,0.620,1.722,0.600 | 0.098,0.071,-0.429,-0.146,1.694,1.350 |
| **92b44**c4b312414f65b8b9a347878bdb9 | 0.080,0.020,1.473,-1.500,-1.785,-0.043 | 0.022,0.073,1.792,-1.821,-0.470,0.334 | 0.097,0.066,-1.568,1.904,-0.541,0.688 | 0.043,0.080,-1.031,1.004,1.557,-0.338 |
| **92d02**605c6d8d3cbc421719268f88bb2 | 0.049,0.054,0.947,-0.623,-1.897,-0.293 | 0.038,0.074,1.272,-1.784,0.709,-1.743 | 0.057,0.038,-1.086,0.420,1.830,1.097 | 0.085,0.040,-0.586,0.092,1.952,1.768 |
| **9300e**330a89d5ed3e8bf4830784235f4 | 0.094,0.063,1.375,-1.366,-0.758,-0.868 | 0.039,0.066,0.672,-1.902,1.531,1.070 | 0.072,0.047,-1.668,0.856,1.567,1.817 | 0.094,0.091,-0.699,0.270,1.691,1.470 |
| **930d2**a86db1366f3852ceefc14e9b5b9 | 0.070,0.034,1.916,-1.760,-0.793,-1.203 | 0.023,0.074,0.622,-1.869,-1.501,-0.174 | 0.073,0.026,-1.423,0.232,1.803,1.935 | 0.089,0.085,-1.207,0.878,1.972,0.877 |
| **9317d**5d3a7e97d28a5888fa06371762f | 0.092,0.018,1.747,-1.478,-1.509,1.100 | 0.011,0.055,1.396,-1.438,-1.415,-0.918 | 0.026,0.061,-1.977,1.843,0.680,1.242 | 0.020,0.097,-1.415,1.246,1.594,0.674 |
| **931bb**348ae2d0368bce95e4d30726d33 | 0.098,0.021,1.183,-0.816,-1.876,1.555 | 0.029,0.082,0.749,-0.714,-1.333,0.451 | 0.087,0.069,-1.546,1.135,1.337,1.875 | 0.014,0.081,-1.451,0.990,1.935,1.589 |
| **935ca**47a42dcd87a768f3c8196dde6b7 | 0.078,0.061,1.308,-1.766,0.686,-0.628 | 0.098,0.052,1.966,-1.447,-1.697,-0.860 | 0.045,0.064,-1.945,1.963,0.992,1.936 | 0.087,0.077,-0.592,0.079,1.888,1.293 |
| **93613**799846ba56d1e3157c828af7b33 | 0.040,0.061,1.881,-1.829,-1.707,-0.295 | 0.074,0.045,0.614,-1.406,1.694,-0.633 | 0.061,0.056,-0.685,0.042,1.757,1.333 | 0.028,0.070,-1.155,0.769,1.516,1.313 |
| **93719**9034a68e796cf1e1299fe434d8b | 0.092,0.087,1.112,-0.884,-1.010,-0.826 | 0.020,0.080,1.010,-0.496,-1.155,-0.042 | 0.087,0.032,-0.865,1.517,0.590,1.618 | 0.048,0.057,-0.146,-0.182,1.782,0.275 |
| **93aff**42659f03a11a612c8f31b3676f8 | 0.098,0.026,0.880,-0.745,-0.922,-0.739 | 0.075,0.050,1.052,-1.200,-0.566,-1.649 | 0.098,0.098,-0.924,0.821,-0.391,0.423 | 0.023,0.093,-0.596,0.461,0.447,0.745 |
| **93baf**c9ad9fa230b50654653f49d5877 | 0.081,0.060,0.575,-0.259,-1.562,-0.277 | 0.059,0.029,1.349,-1.255,-0.408,-1.181 | 0.065,0.067,-1.698,1.870,0.521,1.560 | 0.094,0.036,-0.455,-0.046,1.398,1.310 |
| **93cf7**77083329858eb2ef6d4e160177d | 0.030,0.084,1.771,-1.727,0.243,-0.952 | 0.094,0.033,0.458,-0.236,-0.439,-0.977 | 0.060,0.034,-0.930,0.813,1.899,1.199 | 0.083,0.100,-0.044,-0.401,1.677,0.032 |
| **93e4c**cce437934c21624d8d365685ad5 | 0.070,0.019,1.929,-1.596,-1.902,-0.960 | 0.070,0.081,1.578,-1.312,-1.523,-0.627 | 0.077,0.034,-1.824,1.729,-1.432,0.842 | 0.071,0.074,-1.053,0.484,1.311,1.238 |
| **93e59**ab0150560a21c73983b35e1aed1 | 0.073,0.072,1.909,-1.552,-1.154,-0.675 | 0.067,0.059,0.760,-0.616,-0.298,-0.281 | 0.089,0.072,-1.130,0.605,1.872,1.995 | 0.042,0.062,-0.284,0.160,1.065,0.178 |
| **944dd**b38685bc9a6ddf19dc3db1723b8 | 0.094,0.017,1.083,-0.796,-1.215,0.337 | 0.064,0.088,1.937,-1.342,-1.575,-1.341 | 0.073,0.090,-1.628,1.286,1.889,0.013 | 0.041,0.064,-0.168,-0.094,0.122,1.043 |
| **945e9**4cd111226df6857efaf75156946 | 0.030,0.043,1.662,-0.990,-1.941,-1.099 | 0.078,0.052,0.546,-1.717,-0.704,-1.091 | 0.052,0.068,-1.607,0.212,0.724,1.057 | 0.058,0.052,-0.709,0.601,1.317,0.066 |
| **946e9**11a74877df3b8ae221d00788017 | 0.077,0.056,1.050,-0.657,-1.894,-1.355 | 0.048,0.057,1.381,-0.794,-1.913,-1.666 | 0.100,0.053,-1.901,1.936,1.522,0.631 | 0.082,0.093,-0.769,0.036,1.926,1.329 |
| **94850**7cb0135fd7d02a0546997948f38 | 0.074,0.018,1.963,-1.881,-1.864,1.350 | 0.096,0.082,1.377,-1.140,-1.564,-0.411 | 0.029,0.029,-0.934,0.677,-0.202,0.479 | 0.062,0.012,-0.518,0.060,1.953,0.976 |
| **94860**a15a286993b1845f8f22c877036 | 0.029,0.021,1.546,-1.503,-0.758,-0.049 | 0.042,0.064,0.469,-1.919,-0.183,-0.270 | 0.012,0.097,-0.292,0.003,0.084,0.302 | 0.059,0.080,-0.544,0.154,1.443,1.122 |
| **94977**3d5087fa5a2c786a1ccbdb22b42 | 0.023,0.022,1.639,-1.803,-1.802,-0.116 | 0.022,0.066,0.846,-0.951,1.147,-1.398 | 0.081,0.040,-1.901,1.951,1.794,1.063 | 0.077,0.056,-1.928,1.702,1.795,0.881 |
| **94ad7**04c0df59292b4506acbdd23114f | 0.090,0.044,0.957,-0.906,-1.192,0.546 | 0.092,0.087,0.782,-0.675,-1.395,-1.168 | 0.013,0.036,-1.229,1.018,1.069,0.479 | 0.094,0.071,-0.907,0.573,1.746,1.390 |
| **94d98**2bfe7188637413560688cac9f99 | 0.077,0.038,1.166,-1.326,-1.931,1.118 | 0.035,0.096,1.019,-0.839,-0.793,-1.521 | 0.015,0.050,-1.909,1.808,0.084,-1.326 | 0.053,0.060,-1.010,0.756,1.782,1.030 |
| **94de2**1fe3394e7efaf55ab22b9a7bc7c | 0.077,0.078,0.951,-0.721,-0.888,-0.568 | 0.093,0.051,1.703,-1.161,-0.488,-1.102 | 0.078,0.074,-1.563,1.941,1.941,1.201 | 0.072,0.041,-0.916,0.620,1.902,-0.501 |
| **94f0c**4598735ea2288bf5164b7696ea7 | 0.079,0.074,0.817,-0.735,0.345,-1.163 | 0.099,0.092,1.022,-0.982,-1.437,-0.896 | 0.085,0.059,-1.750,0.905,0.740,1.890 | 0.099,0.041,-0.633,0.217,1.639,1.690 |
| **94f8f**72db03843d7949abe5ecd5b1318 | 0.066,0.055,0.389,-0.240,-1.828,-0.057 | 0.092,0.079,0.870,-0.423,-0.979,-1.340 | 0.095,0.098,-0.572,0.369,-0.326,1.593 | 0.090,0.091,-0.840,0.434,0.913,1.236 |
| **955e9**25f4ae9c5ce4b6e46538a82af7c | 0.050,0.029,1.631,-0.979,-1.919,-0.662 | 0.067,0.058,1.018,-0.975,-1.275,-0.955 | 0.067,0.099,-1.290,0.262,-0.213,1.444 | 0.080,0.098,-1.312,0.438,0.941,1.675 |
| **95638**e9d80e9fbd01addac087b7caef8 | 0.092,0.062,1.164,-1.217,-1.000,-0.227 | 0.073,0.021,1.088,-0.863,-0.539,1.536 | 0.031,0.077,-1.459,1.811,1.203,1.534 | 0.038,0.097,-1.415,1.103,-0.134,1.269 |
| **95690**a44d18736264bfbeadfdf2d460b | 0.051,0.015,1.909,-1.808,-1.749,0.412 | 0.069,0.078,0.935,-1.916,-0.719,-0.489 | 0.053,0.029,-0.839,0.268,0.866,1.387 | 0.093,0.075,-0.922,0.680,0.188,1.459 |
| **95842**b875d2e836468e9b520afb635aa | 0.076,0.026,1.029,-0.934,-1.020,0.031 | 0.095,0.081,0.713,-0.832,-0.382,-0.439 | 0.038,0.067,-1.091,0.825,1.072,1.635 | 0.068,0.074,-0.702,0.547,0.936,0.474 |
| **9588c**593d6494e6d2181a1360692fee6 | 0.055,0.016,0.871,-0.891,-1.909,0.500 | 0.073,0.020,0.529,-0.609,-0.536,1.311 | 0.055,0.067,-1.596,1.272,1.907,-0.108 | 0.078,0.039,-1.839,1.438,1.910,1.456 |
| **9589d**4f6ce53b50c96b18eb5a4306071 | 0.043,0.017,1.345,-1.340,-1.607,-0.153 | 0.041,0.017,1.932,-1.555,-0.851,-1.463 | 0.043,0.097,-1.736,1.798,1.523,1.145 | 0.083,0.065,-0.656,0.055,1.272,1.618 |
| **95c6a**4bec277d1d34dfde39eb531dbe9 | 0.098,0.026,1.695,-1.776,-1.913,-0.174 | 0.022,0.078,0.852,-0.700,0.313,-1.098 | 0.094,0.053,-1.245,1.064,0.970,1.994 | 0.059,0.092,-1.008,0.680,1.635,0.962 |
| **95e95**266ca2f6e0e485b81bb3a1b4723 | 0.021,0.078,1.824,-1.698,-1.863,-1.455 | 0.075,0.046,1.985,-1.336,-0.610,-1.229 | 0.027,0.065,-1.841,1.971,1.507,1.469 | 0.087,0.071,-1.084,0.119,1.654,1.811 |
| **96054**482c0a55dde439f4a5ff66be3a2 | 0.063,0.022,0.786,-0.650,-1.689,1.304 | 0.030,0.016,1.347,-0.963,-0.826,-0.241 | 0.051,0.014,-1.580,1.960,1.939,0.192 | 0.086,0.086,-1.423,0.648,1.010,1.106 |
| **96129**7ae5d087645beba16166e007ffc | 0.031,0.070,1.289,-1.322,-0.984,-0.328 | 0.014,0.062,1.826,-0.805,-1.693,-1.882 | 0.013,0.093,-0.517,0.650,1.638,0.988 | 0.012,0.069,-0.155,-0.111,0.406,1.077 |
| **96407**906fc94d1daf7c6013d903f365e | 0.049,0.014,1.050,-1.238,-1.939,-0.625 | 0.094,0.079,0.587,-0.469,-1.040,0.480 | 0.061,0.065,-0.611,0.592,1.479,0.796 | 0.095,0.096,-1.170,0.702,1.769,1.369 |
| **96616**673f74d6e9822a134c052885d07 | 0.043,0.050,0.061,-0.012,-1.219,0.013 | 0.087,0.069,0.094,-0.020,-1.003,0.293 | 0.078,0.022,-0.475,0.753,1.188,-0.529 | 0.052,0.064,-0.528,0.288,1.805,0.825 |
| **968ca**310c03c3f47392e6a31a6415fd7 | 0.100,0.036,1.456,-1.123,-1.759,-0.459 | 0.015,0.058,0.355,-0.859,0.870,0.457 | 0.048,0.021,-1.799,0.964,1.909,1.126 | 0.056,0.097,-0.648,0.086,1.438,1.761 |
| **968d3**a53e170676045ba036c89deee1f | 0.091,0.088,1.823,-0.978,-1.547,-1.878 | 0.046,0.098,1.783,-1.330,-0.347,-1.078 | 0.072,0.040,-1.920,1.886,1.569,0.851 | 0.059,0.053,-1.007,0.640,0.605,1.380 |
| **96b6f**2ec0541ad906ff497d88f472bca | 0.063,0.094,1.777,-1.540,0.385,-1.933 | 0.039,0.047,1.168,-1.172,-1.111,0.542 | 0.032,0.033,-0.436,0.337,1.271,-0.312 | 0.081,0.064,-0.416,-0.022,0.805,0.920 |
| **96c23**f6ef96f7a18bf87f5d40700d9a6 | 0.048,0.058,1.777,-1.798,-1.746,-0.039 | 0.018,0.073,0.812,-0.521,-0.746,-0.721 | 0.039,0.063,-1.299,1.741,0.956,-0.308 | 0.050,0.041,-1.171,0.751,1.286,1.778 |
| **96ce2**7961d85c179044672648a134ab5 | 0.087,0.030,1.580,-1.939,-0.369,-0.643 | 0.086,0.061,0.429,-0.344,-0.798,-0.253 | 0.072,0.038,-0.540,0.286,1.085,0.857 | 0.069,0.062,-0.590,0.488,1.682,-0.546 |
| **96d51**7f36decd74b28dbdf5e22222f15 | 0.097,0.019,1.598,-1.877,-0.703,-0.227 | 0.074,0.078,1.660,-1.199,-1.610,1.046 | 0.028,0.087,-0.813,1.014,-1.317,0.623 | 0.054,0.073,-0.644,0.298,1.891,1.259 |
| **970ea**837b29d890b4795eb18062ab59d | 0.069,0.020,1.079,-1.154,-0.763,-0.162 | 0.029,0.070,0.858,-0.930,-0.727,-1.073 | 0.043,0.025,-1.580,1.278,1.521,1.168 | 0.096,0.052,-0.710,0.508,0.916,0.921 |
| **972c6**0702f414d06895b70a8f37e9874 | 0.075,0.047,1.948,-1.605,-1.639,-0.344 | 0.021,0.082,0.558,-0.439,1.316,-1.751 | 0.071,0.031,-0.500,0.008,1.723,0.919 | 0.035,0.077,-0.341,-0.012,1.050,1.290 |
| **97380**847eef8529dc7671373f2be5d75 | 0.067,0.089,1.533,-1.574,0.239,-0.674 | 0.094,0.087,0.832,-1.422,1.381,0.170 | 0.081,0.088,-1.334,1.176,1.887,-0.430 | 0.078,0.070,-0.149,-0.572,1.931,1.408 |
| **97387**3185b55bc81606fb240708d07a7 | 0.030,0.057,0.574,-0.489,0.214,-1.922 | 0.042,0.047,0.906,-0.414,0.153,-0.942 | 0.048,0.060,-0.296,0.440,0.112,1.151 | 0.054,0.068,-0.902,0.269,1.862,1.710 |
| **97584**14e6f575dee7ec31365a8b6a533 | 0.098,0.016,1.551,-1.254,-0.982,-0.353 | 0.023,0.080,0.482,-1.013,0.008,-1.179 | 0.068,0.045,-1.670,0.555,0.276,1.788 | 0.069,0.049,-1.524,1.346,1.615,0.924 |
| **97914**3d8cdf3318bdf26587e82ed9e0c | 0.074,0.052,1.914,-1.789,-0.549,-1.193 | 0.093,0.018,0.449,-0.322,0.380,0.358 | 0.090,0.093,-0.844,1.510,0.127,0.684 | 0.037,0.076,-0.806,0.491,0.158,1.893 |
| **97993**715a77395fdd6ae73c2adc63813 | 0.098,0.084,1.770,-1.416,-0.577,-1.726 | 0.021,0.074,1.921,-1.185,0.597,-1.113 | 0.021,0.099,-1.401,1.968,1.178,0.618 | 0.093,0.059,-0.381,-0.672,1.831,1.787 |
| **97a52**89a6e3e1ef70ad17742fbfa5752 | 0.085,0.020,1.711,-1.836,-1.112,-0.108 | 0.070,0.054,0.677,-0.220,-0.780,-1.609 | 0.090,0.013,-1.784,1.871,1.897,1.620 | 0.098,0.015,-0.732,0.204,1.824,1.512 |
| **97b3a**23df08236d397c39e71f2814048 | 0.060,0.058,1.928,-1.010,-1.620,-1.919 | 0.062,0.057,0.433,-0.407,-0.758,-1.727 | 0.078,0.049,-1.657,0.274,1.209,1.601 | 0.096,0.044,-0.350,0.067,1.874,-0.616 |
| **97c88**10d2a5442a6011458b8ba051219 | 0.028,0.052,1.482,-1.822,-1.454,-0.492 | 0.076,0.024,0.629,-0.822,-0.719,0.343 | 0.082,0.029,-0.194,-0.007,1.397,-0.139 | 0.073,0.041,-1.054,0.908,1.773,-0.432 |
| **97e65**b6a895ba79c0b561cdda5185c34 | 0.090,0.019,1.861,-1.573,-1.950,0.475 | 0.012,0.048,0.921,-1.226,-1.507,1.880 | 0.097,0.097,-1.144,1.294,0.656,0.669 | 0.094,0.092,-0.166,-0.169,1.244,0.354 |
| **97f57**d7f8092fd559913454f1911532e | 0.065,0.063,1.003,-0.977,-1.082,-0.046 | 0.024,0.043,1.211,-0.777,-0.767,-0.083 | 0.046,0.087,-1.280,1.876,1.965,1.410 | 0.057,0.079,-0.649,0.495,1.971,-0.599 |
| **98038**c614466e98ef04d504571385f7d | 0.033,0.061,1.619,-1.548,-1.953,-0.368 | 0.091,0.039,0.712,-0.628,-1.099,-1.419 | 0.041,0.033,-0.646,0.338,1.428,1.000 | 0.096,0.089,-0.444,-0.161,1.881,1.056 |
| **982d4**5f172607284b94ba12fffb6c19a | 0.057,0.042,0.381,-0.350,-1.566,-0.082 | 0.062,0.024,0.297,-0.225,0.744,-1.570 | 0.013,0.031,-0.327,0.602,-1.234,-0.536 | 0.082,0.058,-1.793,1.327,1.828,1.114 |
| **984cc**2d01ad8ab8c735809bf4859a2b5 | 0.070,0.020,1.794,-1.789,-1.299,0.046 | 0.019,0.087,1.918,-1.281,-1.512,0.048 | 0.069,0.065,-1.591,1.947,1.873,0.249 | 0.028,0.027,-0.675,0.414,0.655,1.375 |
| **986fa**e494fb87c1d760d9fdda3e4779d | 0.075,0.078,1.512,-1.086,-1.594,-0.557 | 0.069,0.020,0.663,-0.956,-0.955,-0.329 | 0.084,0.047,-1.754,0.776,1.305,1.128 | 0.051,0.067,-0.199,-0.034,1.401,0.478 |
| **98a47**3d25f66636e2e889df8b7351cf5 | 0.067,0.058,1.357,-1.141,-0.298,-1.015 | 0.086,0.036,0.632,-1.419,-0.209,-0.931 | 0.036,0.050,-0.739,0.145,1.414,0.307 | 0.098,0.083,-0.757,0.135,1.904,1.019 |
| **99049**f585a051a84e0997a6a4d712b41 | 0.087,0.015,1.636,-1.800,-1.719,0.558 | 0.028,0.086,0.400,-0.240,0.291,-0.410 | 0.034,0.048,-1.286,1.912,-1.055,0.958 | 0.046,0.023,-0.584,0.338,1.004,1.988 |
| **991a5**170f21c5979361ab477c6eab972 | 0.079,0.083,1.943,-1.537,-0.954,-1.183 | 0.076,0.025,0.608,-0.928,-1.348,-0.432 | 0.057,0.035,-0.995,0.397,-0.253,1.976 | 0.074,0.075,-0.747,0.502,1.422,0.891 |
| **991b6**0342de77a495c488c7c123338bf | 0.048,0.054,1.565,-1.467,-0.360,-1.081 | 0.099,0.033,0.871,-1.340,-1.118,-0.894 | 0.029,0.060,-1.918,0.644,0.762,1.736 | 0.087,0.099,-1.800,1.291,1.796,0.964 |
| **991cb**24b5a80e41bee97a73146e51615 | 0.073,0.064,1.460,-1.478,-1.877,-0.940 | 0.097,0.080,0.774,-0.726,-0.180,0.052 | 0.096,0.045,-1.216,1.452,0.522,0.731 | 0.093,0.071,-0.743,0.528,1.780,-0.321 |
| **9923a**38eff129c1c26b678258ee57c2b | 0.087,0.027,1.838,-1.564,-1.558,0.769 | 0.026,0.050,1.841,-1.852,-1.675,1.100 | 0.035,0.024,-1.693,1.876,1.599,1.066 | 0.085,0.026,-0.691,0.208,1.625,1.626 |

| | | | | |
|---|---|---|---|---|
| **9926e**a3f0a80c8ae4d96523510a0231b | 0.074,0.026,0.460,-0.183,-1.555,-1.331 | 0.088,0.048,1.322,-1.112,-1.416,-0.183 | 0.045,0.096,-1.945,1.436,1.114,1.141 | 0.094,0.091,-0.948,0.831,1.326,-0.718 |
| **996d0**e29a7833197eeedd44758edb0b7 | 0.098,0.047,1.507,-1.871,-0.862,1.420 | 0.095,0.034,1.059,-1.873,-0.109,1.627 | 0.075,0.036,-1.636,1.003,1.933,1.700 | 0.064,0.082,-1.062,0.865,0.544,0.880 |
| **997de**c3253729cc112d7213d65b718af | 0.072,0.044,1.744,-1.784,-0.415,-0.672 | 0.088,0.071,1.260,-1.897,-1.574,0.282 | 0.027,0.089,-1.938,1.044,1.095,1.568 | 0.098,0.023,-0.674,0.506,0.712,0.153 |
| **998ee**32de40e51a4b051cd2f974396e1 | 0.044,0.045,0.548,-0.561,-0.594,0.481 | 0.098,0.079,1.973,-0.726,-1.511,-1.931 | 0.042,0.062,-0.920,1.434,0.642,0.545 | 0.097,0.064,-0.191,-0.375,1.529,1.176 |
| **99991**a103d48b6592237fbfc1c4542e8 | 0.041,0.046,1.727,-1.563,-1.939,0.101 | 0.097,0.054,0.928,-1.242,-0.853,-0.383 | 0.058,0.042,-1.598,0.774,0.664,0.584 | 0.087,0.055,-0.196,-0.501,1.817,1.689 |
| **99994**0cfabeffcff80b5a8b0bfd69fa8 | 0.020,0.050,1.891,-1.806,-1.485,-0.439 | 0.031,0.060,0.591,-1.131,-0.346,-0.652 | 0.080,0.025,-1.872,0.785,-0.263,1.353 | 0.016,0.072,-1.401,1.054,0.790,1.626 |
| **999ce**a72f37541517ce3b1a56b88cec5 | 0.030,0.063,1.962,-1.825,-0.985,-0.472 | 0.097,0.026,0.700,-1.954,1.818,-1.118 | 0.031,0.087,-1.045,0.307,1.551,1.458 | 0.099,0.070,-1.403,0.513,1.877,1.370 |
| **99b68**74fc81ca3323a5f7f1699666785 | 0.088,0.086,1.301,-0.978,-0.803,-1.728 | 0.036,0.016,1.473,-1.243,-1.937,0.026 | 0.011,0.020,-1.642,1.928,0.184,1.566 | 0.092,0.058,-1.478,1.043,0.860,1.319 |
| **9a0a8**b67785a53471a8c3902f9447d4d | 0.098,0.098,1.768,-0.935,-1.794,-1.380 | 0.073,0.067,1.946,-1.332,-1.561,-1.595 | 0.035,0.096,-1.855,1.958,-0.319,0.360 | 0.083,0.016,-1.018,0.636,1.886,-1.113 |
| **9a4e2**0a4204d3f49d1ee9c3ac8b9cc1c | 0.044,0.030,1.629,-1.744,-1.649,0.992 | 0.012,0.064,0.863,-0.669,0.842,-1.021 | 0.094,0.064,-1.617,1.938,1.732,1.236 | 0.092,0.074,-0.626,0.211,1.557,1.151 |
| **9a526**8553bd03d7f74d065f8598c371e | 0.060,0.032,1.610,-1.287,-0.809,-0.289 | 0.019,0.051,0.456,-0.635,-0.441,-1.824 | 0.076,0.053,-1.930,0.370,1.908,0.421 | 0.099,0.033,-0.140,-0.351,1.952,0.696 |
| **9a8e9**2722009587afac5549796a90e16 | 0.018,0.026,1.382,-1.738,-1.674,-0.190 | 0.026,0.058,0.391,-0.247,-0.095,-1.268 | 0.056,0.041,-0.611,0.463,1.785,0.601 | 0.047,0.054,-0.163,0.014,0.572,0.723 |
| **9a9af**2c8d21856c4da80fceb0242e77a | 0.066,0.029,1.805,-1.839,-1.757,0.558 | 0.044,0.066,0.661,-0.497,0.328,-1.463 | 0.096,0.097,-0.894,0.616,1.806,1.785 | 0.045,0.073,-1.442,1.178,1.497,1.500 |
| **9aeeb**976ff9c695fcd5728016c5dccaa | 0.074,0.051,1.353,-1.671,-0.166,-0.211 | 0.048,0.085,0.718,-0.672,-1.051,-0.161 | 0.095,0.014,-0.981,0.896,0.312,1.229 | 0.066,0.094,-1.385,0.859,1.769,0.981 |
| **9afa3**5e2e99d22717ca84d3832118e6a | 0.055,0.086,1.210,-1.248,0.830,-1.568 | 0.017,0.054,0.393,-0.307,0.923,0.148 | 0.088,0.094,-1.487,1.932,1.884,0.407 | 0.039,0.082,-0.919,-0.079,1.945,1.771 |
| **9b0f4**4bfdf67371b340d082af9deb999 | 0.031,0.015,0.981,-1.097,-0.716,-1.058 | 0.027,0.029,0.162,-0.220,-0.968,0.805 | 0.080,0.055,-0.504,0.344,1.212,0.769 | 0.048,0.069,-0.905,0.724,1.577,0.824 |
| **9b20f**510514e3da2270e33b66915c98e | 0.086,0.024,0.415,-0.229,-1.355,0.412 | 0.023,0.083,0.299,0.013,-0.768,-1.345 | 0.035,0.014,-0.734,0.849,1.196,0.581 | 0.052,0.032,-0.235,0.083,0.866,0.406 |
| **9b2ff**8eeca967713a9053727cb6767ab | 0.074,0.022,1.752,-1.897,-0.842,-0.624 | 0.033,0.089,0.959,-0.798,-1.544,-1.879 | 0.092,0.086,-1.279,0.831,1.880,-0.709 | 0.093,0.073,-1.333,1.019,1.329,0.697 |
| **9b4c7**b1f67c4c354307fe451c52fa10f | 0.042,0.015,1.737,-1.840,-1.875,0.089 | 0.023,0.072,0.193,-0.514,-0.982,0.846 | 0.053,0.073,-0.338,0.226,1.471,-0.423 | 0.076,0.082,-0.232,-0.231,1.479,1.457 |
| **9b767**a24996cc66c7bde3756ec87e136 | 0.081,0.053,1.576,-1.210,-1.886,-0.250 | 0.012,0.076,1.376,-1.825,-0.997,0.361 | 0.036,0.042,-1.964,1.289,0.808,1.888 | 0.098,0.062,-0.076,-0.315,1.762,-0.047 |
| **9b86d**d5301d4da0ab79c6b0cf956221b | 0.044,0.027,1.698,-1.633,-1.824,0.468 | 0.082,0.084,0.264,-0.216,-1.275,0.362 | 0.075,0.016,-1.288,0.427,1.473,0.715 | 0.048,0.073,-1.379,0.849,1.761,1.800 |
| **9b9fb**6c971cd396beef4151cb72386b8 | 0.050,0.025,1.246,-1.347,-0.221,-0.652 | 0.068,0.013,1.420,-0.787,-1.532,0.927 | 0.069,0.034,-1.248,1.627,1.207,1.602 | 0.096,0.061,-1.378,0.524,1.845,1.746 |
| **9ba4b**761ae4a5680a91b59ee78a9f365 | 0.073,0.047,1.506,-1.342,-0.555,-0.512 | 0.075,0.083,0.685,-1.310,-0.929,-0.025 | 0.030,0.046,-1.631,0.867,1.069,0.917 | 0.010,0.068,-0.464,0.255,-0.443,1.423 |
| **9c10a**23b883eba91730eac6bb94a9e01 | 0.047,0.070,1.790,-1.350,-0.474,-1.561 | 0.044,0.060,1.389,-0.728,-1.900,-0.043 | 0.072,0.038,-1.341,1.250,1.779,0.350 | 0.054,0.040,-0.440,0.073,1.979,-0.278 |
| **9c1b0**05085e7b9c195df81be8a47b1aa | 0.041,0.043,0.635,-0.595,-1.151,0.319 | 0.017,0.095,0.595,-0.949,0.272,-0.873 | 0.013,0.093,-1.470,0.208,1.409,1.891 | 0.047,0.082,-0.620,0.284,0.983,1.124 |
| **9c223**2c305c327ba960112f4d6f6d76f | 0.097,0.011,1.958,-1.477,-1.941,1.193 | 0.044,0.043,0.406,-0.514,-1.633,-0.070 | 0.042,0.062,-1.873,0.753,0.730,0.879 | 0.047,0.068,-1.928,1.565,0.864,1.790 |
| **9c369**ea4ce0d2a6be91642d8aa8fe1eb | 0.045,0.069,1.410,-1.180,-1.932,-0.326 | 0.044,0.088,1.060,-1.586,-0.436,-0.503 | 0.036,0.091,-0.641,0.265,-0.124,0.983 | 0.088,0.054,-1.124,0.646,1.928,0.753 |
| **9c75e**1efdecbfe91969ad82a4c6067c5 | 0.039,0.093,0.848,-0.522,-1.258,-1.465 | 0.016,0.073,1.809,-1.296,-0.804,-0.090 | 0.079,0.069,-1.575,1.960,1.738,1.050 | 0.082,0.054,-0.569,0.428,1.384,-1.015 |
| **9cc3f**fbcd3c2e077ebc08124704668a0 | 0.049,0.076,1.862,-1.571,-0.433,-1.767 | 0.022,0.054,1.872,-1.383,-1.024,-1.265 | 0.085,0.045,-0.933,1.309,1.180,-0.168 | 0.086,0.099,-0.840,0.283,1.022,1.592 |
| **9d004**c0d5bbfb5933f78f787029ca381 | 0.027,0.022,1.855,-1.734,-1.025,-0.720 | 0.035,0.051,1.066,-0.602,-1.624,-0.990 | 0.031,0.077,-1.964,1.913,1.859,1.213 | 0.085,0.050,-0.641,0.027,0.770,1.503 |
| **9d26e**0c1e3e307e4d93e7c60c4070fab | 0.038,0.024,0.883,-1.014,-1.283,-0.030 | 0.062,0.030,1.038,-1.523,1.878,-0.827 | 0.090,0.022,-1.937,1.678,1.872,1.817 | 0.041,0.082,-0.976,0.631,0.260,1.437 |
| **9d65a**91cbf997fdd2e22eafcf819d17f | 0.063,0.082,1.663,-1.437,-1.386,-0.544 | 0.026,0.058,1.250,-1.231,0.179,-0.681 | 0.092,0.016,-1.031,0.675,1.839,0.660 | 0.072,0.058,-0.508,0.111,1.765,0.501 |
| **9da8e**9f681df5a44b7dba777acde4ed0 | 0.050,0.070,0.163,-0.006,-1.009,-0.812 | 0.013,0.078,0.647,-0.340,-1.842,0.567 | 0.071,0.096,-0.548,0.680,0.997,1.848 | 0.080,0.053,-0.393,0.009,1.971,0.343 |
| **9dc89**d8d17ca6560b1cdbfd4eb6b7b3b | 0.036,0.044,1.754,-1.466,-1.924,-0.839 | 0.092,0.046,0.963,-0.659,-1.447,-1.348 | 0.097,0.098,-1.790,1.101,1.800,0.672 | 0.089,0.045,-0.291,0.033,1.617,-0.589 |
| **9e637**55701c1377bd7e45ba7e2230c0e | 0.099,0.049,1.913,-1.729,-0.413,-0.797 | 0.089,0.057,0.344,-0.343,-0.853,-0.790 | 0.049,0.078,-1.098,0.195,1.240,0.671 | 0.095,0.046,-0.632,0.366,1.120,0.580 |
| **9e63a**a19ac13950a249e55736dc3f9e8 | 0.088,0.038,1.729,-1.771,-1.127,-0.038 | 0.086,0.062,1.338,-0.930,-1.859,-0.797 | 0.050,0.079,-1.156,0.981,0.057,0.965 | 0.090,0.086,-0.660,0.164,0.558,1.922 |
| **9e728**9cb1235d4ae5326bf04b3634e0f | 0.045,0.017,1.926,-1.865,-1.346,0.211 | 0.034,0.063,0.282,0.075,-1.452,-1.834 | 0.075,0.029,-1.118,0.125,0.540,1.946 | 0.024,0.087,-0.796,0.564,1.705,0.911 |
| **9e83d**c516ea0a57c129169abf7a017d5 | 0.040,0.048,0.757,-0.690,-1.042,-0.108 | 0.079,0.022,0.675,-0.890,-0.121,-0.573 | 0.059,0.062,-0.986,0.707,1.807,0.744 | 0.089,0.071,-1.124,0.546,1.967,1.512 |
| **9e89e**8143a2f1ad0eef2f4cbf3155e49 | 0.071,0.057,1.692,-1.162,-1.000,-1.629 | 0.068,0.026,0.283,-0.288,0.218,0.112 | 0.048,0.096,-1.101,0.518,1.281,1.720 | 0.050,0.053,-0.097,-0.258,0.803,1.008 |
| **9ed25**d43f1d5bcbf1bdd0501c177503c | 0.095,0.046,1.629,-1.478,-1.815,0.388 | 0.089,0.037,0.440,-0.172,-1.896,-1.277 | 0.082,0.027,-1.171,0.504,1.401,0.806 | 0.089,0.075,-1.139,0.694,1.926,1.439 |
| **9ee39**81f4cc15b0fb56934a729da1551 | 0.012,0.073,1.399,-1.394,0.426,-0.786 | 0.064,0.044,1.492,-1.917,-1.670,-0.050 | 0.015,0.062,-1.643,1.042,1.831,0.639 | 0.087,0.052,-1.251,1.132,0.294,0.527 |
| **9f348**91e52d2d1017f06cc45d0ee9c76 | 0.066,0.032,0.955,-0.938,-1.592,1.147 | 0.012,0.024,0.710,-0.837,1.952,-1.873 | 0.047,0.039,-1.740,1.905,0.477,0.808 | 0.027,0.093,-0.548,0.144,0.323,1.871 |
| **9f3da**b05c7a6760fe70c2ddda2fcb413 | 0.061,0.017,0.962,-1.062,-0.951,0.987 | 0.049,0.052,1.136,-1.555,-1.733,0.817 | 0.068,0.092,-1.660,1.076,1.902,1.753 | 0.094,0.049,-0.734,0.570,1.008,0.696 |
| **9f489**f9dbdca67ad959c2e60d3b2593f | 0.046,0.040,1.449,-1.129,-1.640,-0.637 | 0.039,0.020,0.739,-0.930,-1.771,-0.589 | 0.059,0.036,-1.565,0.691,1.857,0.585 | 0.092,0.065,-0.956,0.765,0.805,0.192 |
| **9f4c5**e0d4e22d9594868535d65db34c4 | 0.075,0.079,1.766,-1.996,0.641,-1.291 | 0.080,0.063,0.664,-0.236,-1.522,-0.334 | 0.065,0.037,-0.936,0.641,0.908,1.913 | 0.054,0.049,-1.263,0.914,1.972,0.791 |
| **9f654**89867e975145e2f4706dad45095 | 0.038,0.047,1.541,-1.482,-1.366,-1.004 | 0.014,0.088,0.650,-0.745,0.366,-1.511 | 0.097,0.061,-0.812,0.188,1.915,0.494 | 0.081,0.036,-0.618,0.258,1.927,1.554 |
| **9f747**b7b388509e6942012c01b7d90b9 | 0.013,0.068,1.651,-1.696,-0.776,-1.248 | 0.091,0.044,1.075,-0.954,-0.464,-1.457 | 0.057,0.027,-1.815,1.814,1.430,1.862 | 0.098,0.084,-1.891,1.242,1.959,1.713 |
| **9f9c2**9200897b36ffb0fec855c6e2f8d | 0.047,0.022,1.786,-1.676,-1.425,-0.709 | 0.019,0.016,0.888,-1.022,-1.250,-1.267 | 0.070,0.021,-1.818,1.485,-0.328,1.951 | 0.087,0.069,-1.616,1.118,1.157,0.946 |
| **9fa64**3d19fbd612777f1943452217ad0 | 0.085,0.054,1.681,-1.492,-1.239,-0.108 | 0.095,0.043,1.008,-0.801,-1.919,-0.339 | 0.043,0.031,-1.115,1.126,0.497,-0.648 | 0.020,0.099,-0.626,0.279,0.616,1.203 |
| **9fcc5**b4dfc73d1d0875a57b09c2f8509 | 0.085,0.030,1.605,-1.510,-0.732,-0.431 | 0.022,0.053,1.262,-1.128,-1.843,-1.741 | 0.065,0.036,-0.416,0.324,1.463,0.227 | 0.066,0.069,-0.128,-0.391,1.787,1.192 |
| **9fea4**272dc6220f63e4a7d72cd7b7376 | 0.088,0.061,0.684,-0.804,-1.455,1.992 | 0.036,0.037,0.831,-1.230,1.611,1.543 | 0.083,0.087,-1.814,1.579,1.857,1.086 | 0.068,0.022,-0.789,0.297,1.819,1.822 |
| **9ffa9**ca48dfe678d2c6545045ff803c4 | 0.085,0.062,0.444,-0.458,-1.866,1.134 | 0.027,0.067,0.872,-0.644,-0.168,-0.794 | 0.070,0.061,-1.298,1.721,1.816,-0.943 | 0.095,0.068,-0.201,-0.102,1.117,0.364 |
| **a00b9**ff2b8f523a1535b7bda099d23fa | 0.086,0.051,1.469,-0.981,-1.336,-0.359 | 0.026,0.073,0.830,-1.494,-0.657,-0.517 | 0.066,0.061,-0.757,0.224,1.275,0.318 | 0.036,0.097,-0.890,0.478,0.597,1.613 |
| **a06ec**e4dd656a94d7468ad15452912dd | 0.071,0.069,1.266,-1.464,-0.861,0.924 | 0.085,0.051,0.562,-0.473,1.941,-0.737 | 0.029,0.097,-0.213,-0.030,1.065,1.559 | 0.029,0.053,-1.659,1.584,0.379,0.226 |
| **a0893**1766cad4ef3b594cbcb93ef3b00 | 0.090,0.020,1.223,-1.211,-1.728,1.501 | 0.019,0.025,0.730,-0.627,-0.379,-0.025 | 0.055,0.037,-1.023,1.153,-0.277,1.698 | 0.048,0.061,-1.566,1.147,1.818,1.392 |
| **a0967**3a05824c1a5385f235f884fc785 | 0.020,0.095,1.113,-0.935,-0.638,-1.184 | 0.065,0.017,1.512,-1.190,-1.908,0.937 | 0.050,0.036,-0.662,0.586,0.466,1.333 | 0.086,0.074,-0.711,0.252,1.605,1.332 |
| **a0a8b**614f08043a548552b7a6c135efd | 0.052,0.037,1.491,-1.681,-1.784,0.599 | 0.040,0.077,0.548,-0.450,-0.575,-0.236 | 0.073,0.019,-1.326,1.356,1.691,1.600 | 0.014,0.088,-0.448,0.127,1.223,1.494 |
| **a0e15**d4e20387b78968df42998760fad | 0.028,0.066,1.123,-1.097,0.351,-1.202 | 0.078,0.064,1.224,-0.533,0.225,-0.927 | 0.055,0.075,-0.549,0.948,1.442,1.038 | 0.080,0.037,-1.487,1.119,1.743,-0.442 |
| **a0f7c**acdb6b47d05215675a88c424b21 | 0.053,0.041,1.778,-1.807,-0.060,-1.206 | 0.083,0.052,0.753,-0.889,-0.814,-0.120 | 0.090,0.037,-0.729,0.113,1.666,1.751 | 0.064,0.080,-0.366,-0.007,1.729,0.731 |
| **a1143**4eef15d574231b419d7d7f691a7 | 0.013,0.086,1.436,-1.785,0.388,-1.033 | 0.026,0.019,0.489,-0.274,-0.621,-0.512 | 0.044,0.049,-0.791,1.016,1.177,0.206 | 0.028,0.069,-1.483,1.203,0.755,0.902 |
| **a1383**91a4132ce4422646a1a1615108f | 0.095,0.032,1.817,-1.570,-1.258,0.279 | 0.070,0.023,0.629,-1.415,-0.449,-1.267 | 0.100,0.083,-1.708,0.233,1.300,1.564 | 0.089,0.082,-0.887,0.386,1.751,1.695 |
| **a14af**ad5795a131d8facfdc66e4fc30e | 0.066,0.076,1.929,-1.356,-1.805,-0.880 | 0.055,0.021,0.480,-0.673,-0.143,0.472 | 0.096,0.098,-1.620,0.630,1.899,0.686 | 0.028,0.062,-0.391,-0.061,1.953,1.854 |
| **a1741**2435bd338d87f0969a28ffdb5a1 | 0.079,0.063,0.682,-0.004,-1.478,-0.640 | 0.069,0.028,0.311,-0.369,-1.006,-0.287 | 0.044,0.034,-0.420,0.038,1.418,0.589 | 0.049,0.061,-1.304,0.903,1.244,1.813 |
| **a1bc8**a88e8870084561317e43a4b0537 | 0.077,0.014,1.823,-1.575,-1.840,0.690 | 0.049,0.015,0.687,-1.664,-0.093,-1.656 | 0.036,0.026,-0.997,0.439,-0.638,0.612 | 0.038,0.063,-0.463,0.145,1.370,1.697 |
| **a1cc7**857c2565ecb38ccbdc6bc21d70d | 0.061,0.062,1.934,-1.778,-0.739,-1.141 | 0.032,0.048,0.293,-0.294,-0.824,-0.339 | 0.091,0.052,-1.076,0.774,-0.161,-0.359 | 0.067,0.081,-0.969,0.532,1.674,0.778 |
| **a1f4f**3fd920a63970176790263b31f34 | 0.037,0.060,0.741,-0.611,0.079,-1.200 | 0.027,0.044,0.909,-0.567,-1.521,-0.192 | 0.016,0.061,-1.565,1.800,1.592,0.668 | 0.070,0.068,-1.043,0.447,0.375,1.321 |
| **a1f61**702d3b12651777466b05d7f204c | 0.094,0.071,1.131,-0.818,-1.313,-0.209 | 0.016,0.085,0.506,-0.486,-0.087,0.351 | 0.090,0.021,-0.607,0.572,1.104,-1.020 | 0.062,0.067,-1.127,0.684,0.637,1.957 |
| **a21ef**9edbe1c9fdc6bbd060f7d2010b2 | 0.041,0.059,1.972,-1.492,-1.369,-1.911 | 0.076,0.035,1.556,-1.470,-1.916,-1.385 | 0.093,0.013,-1.884,1.138,1.536,0.947 | 0.049,0.074,-0.610,0.169,1.804,1.075 |

| | | | | |
|---|---|---|---|---|
| **a22e4**5087894c21c8da237825d0ef2e2 | 0.035,0.022,1.402,-1.124,-0.792,-1.215 | 0.066,0.053,0.599,-1.229,0.509,-1.467 | 0.011,0.073,-0.630,0.131,1.349,0.562 | 0.063,0.059,-0.652,0.126,1.215,1.680 |
| **a2480**82ca5967e3cc549095979f85efc | 0.082,0.017,1.762,-1.552,-1.866,-0.824 | 0.016,0.025,0.375,-0.373,-0.732,-1.067 | 0.099,0.077,-0.655,0.385,0.203,0.391 | 0.091,0.074,-0.875,0.461,0.982,0.937 |
| **a275d**09d1475444a2012a39971984c78 | 0.077,0.069,1.568,-1.419,-0.767,-0.728 | 0.069,0.085,1.040,-0.588,-0.612,-1.562 | 0.080,0.037,-0.738,0.690,1.465,1.005 | 0.063,0.044,-0.860,0.640,1.338,0.748 |
| **a279a**0f665a26cae3e889f1776cdb6c9 | 0.070,0.063,0.659,-0.615,-0.184,-0.645 | 0.040,0.096,1.953,-1.057,-0.904,-1.046 | 0.099,0.037,-1.365,1.798,1.942,1.610 | 0.037,0.070,-0.370,-0.143,1.546,1.680 |
| **a28aa**bea5ab350e9677d895b9ae23641 | 0.067,0.083,1.705,-1.528,-1.147,-0.198 | 0.057,0.089,0.493,-0.398,-1.005,1.221 | 0.096,0.090,-0.881,1.187,1.155,0.520 | 0.067,0.065,-1.533,1.011,1.696,1.153 |
| **a2905**1b718678962604525aced58f434 | 0.087,0.073,1.270,-0.916,-0.998,-0.866 | 0.075,0.072,1.352,-1.032,-0.473,-0.538 | 0.074,0.094,-1.837,1.823,1.368,1.730 | 0.019,0.075,-0.981,0.751,0.347,1.029 |
| **a2aae**3a21149bb30acc05d2f0e3f03e1 | 0.011,0.050,1.597,-1.820,-0.741,-0.387 | 0.024,0.092,0.360,-0.254,1.907,-0.111 | 0.097,0.058,-1.026,1.811,1.914,0.655 | 0.062,0.068,-1.526,0.859,1.815,1.836 |
| **a2c0d**d81031714d994873a74ff11d60c | 0.096,0.057,1.129,-1.219,0.009,-0.502 | 0.037,0.073,0.856,-0.758,-0.699,-0.136 | 0.082,0.077,-1.850,1.913,1.440,1.183 | 0.077,0.076,-0.796,0.393,0.335,1.306 |
| **a2ce3**6ccd07ba560ad9e853c530485dd | 0.083,0.024,0.982,-0.901,-1.534,-0.185 | 0.048,0.085,0.794,-0.802,-1.005,0.838 | 0.079,0.023,-1.335,1.725,1.939,-0.884 | 0.044,0.073,-1.919,1.569,1.122,1.641 |
| **a2cfe**ac225912d8f4146e376a7dfc86e | 0.079,0.082,1.349,-1.319,-0.928,0.420 | 0.038,0.023,0.535,-0.695,1.873,-0.220 | 0.079,0.024,-1.685,1.927,1.957,1.005 | 0.032,0.082,-0.944,0.523,0.058,1.536 |
| **a2d1a**6e87dee2cb236f2a8d27249f481 | 0.057,0.064,1.382,-1.028,-1.685,-0.583 | 0.035,0.087,0.843,-0.897,1.054,-0.201 | 0.073,0.012,-1.767,1.699,1.898,0.617 | 0.095,0.037,-1.293,0.872,1.337,0.556 |
| **a2dd4**d565885a97698279ad8081d54a0 | 0.092,0.034,1.774,-1.765,-0.933,-1.206 | 0.094,0.073,0.916,-1.377,-1.132,1.418 | 0.053,0.055,-0.738,0.458,1.306,0.378 | 0.074,0.019,-0.655,0.364,1.648,1.630 |
| **a3300**7ed7105230b5018ccdcebf20186 | 0.047,0.018,1.939,-1.926,-0.427,-1.565 | 0.093,0.031,1.311,-1.926,-0.946,-0.911 | 0.095,0.071,-1.820,0.795,1.275,0.813 | 0.066,0.071,-0.932,0.806,1.295,-0.017 |
| **a33ae**cc50feedf528b4415de594c253f | 0.086,0.019,1.414,-1.729,-0.605,1.021 | 0.033,0.079,1.238,-0.669,-0.425,0.109 | 0.064,0.042,-0.999,1.747,1.839,1.361 | 0.087,0.059,-0.439,0.314,1.887,-0.997 |
| **a33c3**5e356e289b3330b61c6bbed5169 | 0.015,0.044,0.957,-0.849,-0.268,-0.957 | 0.070,0.061,1.368,-0.941,-0.435,-0.780 | 0.070,0.073,-1.328,1.623,-0.896,1.383 | 0.028,0.017,-1.128,1.073,0.849,-0.161 |
| **a37b8**7c81ef980308cc883f84b926c77 | 0.019,0.088,1.894,-1.638,-0.788,-1.306 | 0.023,0.076,1.185,-1.678,1.697,0.687 | 0.051,0.085,-1.713,1.330,1.717,1.014 | 0.086,0.073,-0.135,-0.123,0.909,0.217 |
| **a3a01**fa8aef864b7b0e1e5190e78a286 | 0.070,0.079,1.799,-0.464,-1.434,-1.613 | 0.099,0.087,0.806,-1.803,-1.815,0.950 | 0.021,0.044,-1.364,0.385,1.080,1.247 | 0.064,0.012,-0.150,0.102,0.314,0.305 |
| **a3a9d**4ddad4e375be90866eb67100efa | 0.088,0.026,1.000,-0.558,-1.905,0.524 | 0.030,0.047,0.450,-0.672,1.432,-0.743 | 0.066,0.095,-1.487,0.816,0.511,1.094 | 0.047,0.017,-0.206,0.201,0.819,-1.784 |
| **a3aae**d98050b45bf77ceb7644b384147 | 0.046,0.025,0.562,-0.432,-0.858,-0.685 | 0.052,0.016,0.744,-0.845,-1.843,-0.331 | 0.019,0.085,-1.677,0.965,1.786,1.451 | 0.077,0.065,-0.398,-0.145,1.763,1.527 |
| **a3ac2**86fbe7a96961ff001cc2cc1b3d0 | 0.072,0.060,1.481,-1.641,-1.499,0.061 | 0.033,0.033,0.834,-0.714,-0.537,-0.527 | 0.071,0.036,-1.798,1.813,1.891,1.956 | 0.075,0.070,-1.451,1.098,0.599,1.310 |
| **a3c2f**97b18373dd89cc76998547df0ec | 0.067,0.037,1.755,-1.834,-1.853,-0.036 | 0.088,0.061,0.537,-1.237,0.741,1.429 | 0.033,0.024,-1.378,1.309,1.484,1.125 | 0.039,0.012,-0.813,0.598,1.629,1.495 |
| **a3f62**19057029d5decba9de7d1c840d2 | 0.022,0.060,1.571,-1.523,-1.018,-0.445 | 0.034,0.057,1.935,-1.267,1.860,-1.814 | 0.096,0.089,-1.888,1.941,1.950,1.995 | 0.081,0.096,-1.812,0.996,1.747,1.375 |
| **a404f**b9cfaaf020ff746d13a93b0e458 | 0.027,0.057,1.086,-0.879,-1.781,-0.448 | 0.062,0.066,0.345,0.003,-1.461,0.278 | 0.099,0.099,-0.863,1.171,-0.001,0.378 | 0.067,0.074,-0.322,0.093,0.699,0.129 |
| **a4178**9620543081da6cc0a07f51966ee | 0.084,0.066,1.826,-1.308,-1.142,-0.372 | 0.057,0.091,1.230,-0.571,-0.565,-1.836 | 0.034,0.097,-1.813,1.081,-0.229,1.327 | 0.067,0.076,-1.423,0.471,1.642,1.601 |
| **a41ce**d23fc2c6d8674c88c224fd751d2 | 0.049,0.046,0.852,0.248,-1.239,-1.470 | 0.026,0.027,0.457,-0.835,-0.624,-1.930 | 0.058,0.069,-1.866,0.240,1.663,0.529 | 0.090,0.021,-0.455,0.264,0.676,-0.263 |
| **a4596**6877d5bdbf446cebecf4ad8b97a | 0.056,0.022,1.724,-1.416,-0.925,-0.885 | 0.081,0.087,0.588,-1.912,-1.466,1.875 | 0.072,0.026,-1.703,0.483,1.895,0.712 | 0.049,0.060,-1.165,0.954,0.149,1.754 |
| **a471a**f48dfe72da448976c1589bfd13b | 0.098,0.028,1.289,-0.967,-1.549,0.172 | 0.018,0.042,1.294,-0.990,1.527,-0.976 | 0.042,0.065,-1.650,1.874,1.845,1.713 | 0.083,0.097,-1.458,0.568,1.756,1.907 |
| **a49db**a52e0a9731176ad26d919951651 | 0.079,0.050,0.505,-0.429,-0.813,0.525 | 0.013,0.080,1.955,-1.364,-1.015,-1.762 | 0.089,0.099,-1.357,1.680,1.325,0.676 | 0.066,0.097,-0.481,-0.023,0.220,1.753 |
| **a4a2a**ef300e4b0c2e5198ce123178415 | 0.043,0.030,1.768,-1.833,-0.881,-0.652 | 0.052,0.093,1.308,-0.579,-0.698,-1.013 | 0.060,0.062,-1.364,1.943,0.974,0.255 | 0.095,0.082,-1.126,0.832,1.119,-0.218 |
| **a51cd**24f6373cbce2e35c929efe0df5d | 0.078,0.014,0.997,-0.764,-1.860,0.765 | 0.098,0.072,1.260,-1.463,-0.624,-1.158 | 0.096,0.068,-1.729,1.151,1.381,0.140 | 0.048,0.042,-0.286,-0.017,1.549,1.573 |
| **a52a9**24e990d30fcc89cd93937353cd7 | 0.043,0.013,1.975,-1.944,-1.895,0.894 | 0.025,0.075,0.477,-0.277,-0.886,-0.698 | 0.071,0.058,-0.419,0.264,1.929,0.787 | 0.056,0.080,-0.414,0.148,1.138,0.898 |
| **a547a**6d80081e76809ce745b350fbb59 | 0.075,0.061,1.504,-1.184,-0.637,-0.905 | 0.024,0.048,0.281,-0.467,0.355,-1.346 | 0.059,0.025,-0.939,0.275,1.438,0.879 | 0.077,0.042,-1.114,0.740,1.891,1.234 |
| **a552f**a5be532b490db6d338647a069ad | 0.087,0.069,1.497,-0.809,-0.871,-0.365 | 0.055,0.073,0.602,-1.544,-0.129,-0.700 | 0.062,0.088,-0.868,0.284,-0.274,0.208 | 0.072,0.041,-0.557,0.327,1.180,0.307 |
| **a5650**57b06b625acf4ffcf5a7b119609 | 0.039,0.021,1.087,-1.495,-1.923,0.503 | 0.045,0.027,0.694,-0.809,-0.422,0.474 | 0.064,0.031,-1.659,1.460,0.925,1.205 | 0.083,0.049,-0.631,0.401,1.119,1.033 |
| **a576d**b49d43bc9a1033544b78c0fc7a4 | 0.092,0.058,1.027,-1.148,-0.061,-0.258 | 0.044,0.095,0.885,-0.782,-0.928,-1.185 | 0.085,0.100,-1.874,1.870,-0.115,0.383 | 0.082,0.073,-0.646,0.490,-0.695,1.732 |
| **a59c5**3d7341e683cd70f36b2fddcc249 | 0.033,0.071,0.977,-0.426,-0.649,-1.339 | 0.028,0.039,1.319,-1.028,-0.486,-0.934 | 0.086,0.060,-1.891,1.184,1.414,1.471 | 0.093,0.028,-0.245,-0.831,1.852,1.122 |
| **a5b82**3f21300be2a76bf7bd8edc701d7 | 0.043,0.012,1.590,-1.849,-1.198,-0.340 | 0.026,0.097,0.563,-0.281,-0.537,-1.033 | 0.094,0.083,-0.895,0.718,-0.282,1.956 | 0.058,0.029,-1.334,1.027,1.916,1.582 |
| **a5b84**37c53d8123ede210b0c3b183171 | 0.098,0.027,1.348,-1.290,-0.819,0.427 | 0.099,0.018,1.189,-1.148,-1.601,-0.172 | 0.065,0.099,-0.862,0.617,-0.430,0.453 | 0.083,0.099,-0.703,0.539,0.567,0.192 |
| **a6040**35f58f0c427069fb7a2ad24c1e8 | 0.077,0.029,0.401,0.543,-1.806,-0.601 | 0.033,0.075,0.497,-0.471,-0.386,-1.681 | 0.092,0.028,-1.123,0.351,0.546,1.297 | 0.026,0.082,-0.732,0.367,0.799,1.500 |
| **a6061**8299dba1182536ed91e2ab72c04 | 0.042,0.054,1.134,-0.997,-0.417,-1.168 | 0.065,0.015,1.519,-1.481,-1.009,-1.084 | 0.052,0.040,-0.995,1.363,1.817,-1.700 | 0.078,0.095,-0.657,0.260,0.146,1.751 |
| **a6113**7def9e17f978707a589b04cab43 | 0.049,0.041,1.796,-1.560,-1.766,0.253 | 0.020,0.096,0.573,-1.470,-0.842,-0.859 | 0.059,0.064,-1.396,0.096,1.351,0.531 | 0.067,0.093,-1.085,0.712,1.405,0.869 |
| **a616d**8624560342da94b792170012966 | 0.051,0.049,1.283,1.792,-1.797,-1.945 | 0.079,0.047,1.159,-1.644,-1.926,-1.522 | 0.044,0.042,-1.834,0.306,0.280,0.423 | 0.066,0.067,-0.225,-0.591,1.708,1.307 |
| **a659a**22120647015dd8ed03105fa5543 | 0.060,0.050,0.665,-0.561,-1.645,0.284 | 0.026,0.061,1.476,-0.828,-0.224,-1.097 | 0.019,0.050,-1.094,1.926,-0.147,1.426 | 0.062,0.084,-0.176,-0.296,0.913,1.092 |
| **a65a6**9c50e6281298f5949009b48ddfd | 0.011,0.064,1.077,-1.157,1.099,-0.419 | 0.049,0.076,1.793,-0.832,-0.920,-0.006 | 0.074,0.031,-1.092,1.929,1.429,-0.056 | 0.090,0.094,-0.655,0.167,-0.103,1.380 |
| **a6686**d1e63cd977b388e773eb9b22ff0 | 0.036,0.050,1.654,-1.804,-1.147,-0.623 | 0.033,0.034,0.159,-0.167,-0.888,-0.005 | 0.027,0.047,-0.515,0.267,1.214,0.270 | 0.091,0.043,-0.346,-0.065,1.690,1.689 |
| **a6701**630d53b65b4a155bc7b87ab8849 | 0.096,0.085,1.764,-1.511,-1.789,-0.830 | 0.020,0.018,1.416,-1.862,-1.881,1.038 | 0.019,0.071,-1.722,1.449,-0.648,0.885 | 0.085,0.085,-1.470,0.859,1.840,1.477 |
| **a6824**e8da1fb0d782a5b2f58ec1e32fd | 0.062,0.046,1.461,-1.225,-1.126,-0.461 | 0.097,0.051,1.385,-1.478,-0.222,0.344 | 0.081,0.083,-0.880,0.449,1.449,0.470 | 0.057,0.066,-0.321,-0.177,1.945,1.438 |
| **a6da6**91c4ae9230f4058894843c87801 | 0.034,0.035,0.806,-0.804,-0.707,-0.032 | 0.034,0.080,1.511,-1.535,-0.783,-1.697 | 0.026,0.090,-1.711,1.812,1.744,-1.427 | 0.042,0.068,-0.446,0.216,-0.734,1.336 |
| **a6dc5**30b8241ac695a44c3e9fcc87f9c | 0.084,0.084,1.109,-0.990,-1.806,0.201 | 0.034,0.071,1.436,-0.630,-0.684,-0.668 | 0.100,0.040,-0.846,1.617,1.052,0.366 | 0.058,0.091,-0.737,0.021,1.904,1.667 |
| **a7294**cda9fc19ec612823fe61f67d860 | 0.087,0.063,0.493,-0.500,-0.549,0.165 | 0.024,0.033,0.652,-0.374,-1.204,-1.834 | 0.099,0.092,-1.277,1.929,1.091,1.852 | 0.052,0.081,-1.688,1.476,-0.396,1.542 |
| **a743c**0bc44870808f4fc8a7289a4f236 | 0.038,0.035,1.619,-1.840,-1.739,0.454 | 0.024,0.078,0.889,-0.649,-0.647,-0.564 | 0.070,0.100,-1.615,1.816,1.905,1.536 | 0.023,0.086,-0.498,0.220,1.241,1.218 |
| **a76da**3c003e4327cf179bdb8c67fa2f9 | 0.068,0.058,1.542,-1.098,-1.760,-0.588 | 0.061,0.035,0.668,-1.916,0.056,-0.618 | 0.096,0.061,-1.724,0.513,1.912,0.605 | 0.038,0.052,-0.824,0.784,-0.228,0.490 |
| **a77f8**4d55c993f55c0f13d6d4b3f9459 | 0.023,0.075,1.757,-1.500,-1.309,-1.998 | 0.072,0.022,1.349,-0.886,0.166,-1.925 | 0.099,0.064,-0.778,1.313,0.926,-0.910 | 0.073,0.057,-0.190,-0.137,1.737,0.489 |
| **a7df4**99350cc29d8621847e5649a5186 | 0.051,0.016,1.570,-1.758,-0.581,-0.002 | 0.076,0.053,0.814,-1.407,-0.736,-0.994 | 0.062,0.053,-0.602,0.136,0.802,0.736 | 0.057,0.059,-0.136,0.047,1.815,-0.568 |
| **a8003**4f8fb1429e62aef3519356648f8 | 0.089,0.015,1.652,-1.010,-1.573,-0.542 | 0.027,0.069,1.239,-1.128,-0.502,-1.909 | 0.026,0.062,-1.991,1.373,1.588,-0.440 | 0.076,0.070,-0.517,-0.174,1.530,1.278 |
| **a875a**71fac5c8954f372ceeba5debad8 | 0.043,0.031,1.645,-1.815,-1.792,0.178 | 0.055,0.017,0.533,-0.517,-1.748,-1.276 | 0.077,0.081,-1.862,1.148,-0.146,0.690 | 0.049,0.064,-1.208,1.151,1.514,-0.319 |
| **a876a**e005e1868665b17e976b90ce0bf | 0.078,0.014,1.719,-0.910,-1.933,-0.054 | 0.097,0.084,0.388,-0.227,-0.766,-1.394 | 0.073,0.081,-1.264,-0.051,0.975,1.653 | 0.071,0.046,-0.742,0.426,1.931,1.028 |
| **a87d6**713bfb0d6777e309ff5eac396dc | 0.077,0.018,1.659,-1.821,-1.767,1.192 | 0.042,0.079,1.273,-1.903,-0.859,1.443 | 0.038,0.083,-1.879,1.708,-0.050,0.686 | 0.092,0.069,-1.168,0.873,0.702,1.703 |
| **a884c**45b986a7ad2d2c7415574e43f6f | 0.013,0.038,1.655,-1.783,-1.697,-1.056 | 0.022,0.037,0.627,-1.108,0.299,1.148 | 0.099,0.065,-0.470,-0.057,1.778,0.624 | 0.019,0.079,-0.566,0.438,0.098,0.781 |
| **a891f**5eb33710b42f12932d89c55c55f | 0.058,0.020,0.420,-0.390,-1.005,-0.828 | 0.021,0.019,0.962,-0.696,0.806,0.636 | 0.095,0.065,-1.335,1.951,1.086,1.256 | 0.065,0.058,-0.305,-0.162,1.146,1.720 |
| **a8a05**716e02a32c42cbd41bbcb331808 | 0.045,0.045,1.254,-0.789,-1.298,-0.269 | 0.098,0.046,0.080,-0.034,-0.746,-0.024 | 0.043,0.029,-1.283,0.261,1.186,0.589 | 0.081,0.068,-0.718,0.250,1.625,1.569 |
| **a8c3e**e56dfed1858a2ecdb096436b960 | 0.035,0.064,1.820,-1.809,-1.106,-0.487 | 0.077,0.060,0.451,-0.136,-1.947,-0.222 | 0.040,0.032,-0.674,0.360,-0.714,1.329 | 0.095,0.083,-0.666,0.321,1.972,-0.696 |
| **a8e80**f52f1354cf10daabd06883204d0 | 0.041,0.019,1.569,-1.826,-1.820,0.963 | 0.016,0.019,0.298,-0.890,1.602,-0.133 | 0.056,0.050,-1.252,0.480,0.870,1.154 | 0.090,0.066,-0.785,0.598,1.838,-1.099 |
| **a8e85**d97bfb3cd801676da7a5355d953 | 0.040,0.057,1.567,-1.677,0.314,-1.904 | 0.015,0.046,0.679,-0.806,-1.729,0.112 | 0.029,0.018,-1.226,1.076,1.039,1.480 | 0.055,0.062,-1.109,1.053,1.896,-0.805 |
| **a95d0**ef1b11b799eedc37ce171c0d438 | 0.031,0.078,1.775,-1.491,0.618,-1.535 | 0.074,0.063,1.209,-1.804,-1.716,0.381 | 0.050,0.072,-1.142,0.571,0.881,0.917 | 0.052,0.055,-1.250,0.907,0.997,1.475 |
| **a99c0**61818bf869d3f65d9208ecc1f1c | 0.078,0.025,1.803,-1.896,-0.888,-0.391 | 0.036,0.064,0.385,-0.219,-1.158,-1.140 | 0.094,0.060,-0.587,0.530,0.076,0.393 | 0.038,0.100,-0.218,-0.062,1.235,1.245 |
| **a9b28**3d9394d4dd66e0bc1a6863ba93f | 0.059,0.085,1.117,-0.999,-0.670,-1.153 | 0.018,0.079,1.011,-0.910,0.921,-1.216 | 0.075,0.079,-0.962,1.059,1.942,-0.706 | 0.094,0.066,-0.478,-0.049,1.821,1.613 |
| **a9c0f**9a717786c1bf0e2173cd51fc469 | 0.062,0.067,1.716,-1.616,-1.343,-0.685 | 0.058,0.075,0.282,-0.321,0.888,-1.484 | 0.066,0.052,-1.063,0.669,1.361,1.388 | 0.090,0.059,-1.573,1.238,1.753,1.295 |

| | | | | |
|---|---|---|---|---|
| **a9f39**4605db65cccbad8188471017f0c | 0.063,0.061,1.618,-1.674,0.254,-1.053 | 0.094,0.013,1.651,-0.947,1.372,-0.145 | 0.092,0.071,-1.217,1.970,1.951,1.424 | 0.027,0.065,-1.143,0.451,1.952,1.444 |
| **a9f6a**9987bde192427c2a2b4d50765e8 | 0.098,0.013,1.805,-1.498,-1.793,0.897 | 0.012,0.037,1.308,-1.420,-0.942,-1.609 | 0.071,0.071,-1.029,0.783,1.003,0.539 | 0.094,0.083,-1.165,0.777,1.487,0.404 |
| **aa01c**15dd8763134bc739567112d6a95 | 0.095,0.061,1.052,-0.738,-1.901,0.044 | 0.026,0.038,1.137,-1.427,0.630,1.374 | 0.019,0.043,-1.499,1.947,1.580,-0.432 | 0.028,0.048,-1.025,0.721,1.576,1.606 |
| **aa02c**c008a7fc94aedabd0e4232daed4 | 0.049,0.046,1.028,-1.024,-1.119,-0.067 | 0.089,0.062,0.479,-0.701,-0.253,0.151 | 0.053,0.041,-1.437,0.737,1.936,1.649 | 0.043,0.068,-0.102,-0.059,1.134,0.444 |
| **aa0de**9b43151b65701aeff19c1dc13de | 0.033,0.024,0.917,-0.910,-1.665,-0.482 | 0.081,0.020,1.223,-0.889,-1.942,1.152 | 0.024,0.050,-1.477,1.887,0.974,0.680 | 0.039,0.075,-1.520,1.141,1.768,1.148 |
| **aa19d**8a7b9e788d0d43d51f9e9bc155f | 0.013,0.059,1.701,-1.858,-0.468,-1.127 | 0.084,0.047,0.550,-0.302,-1.150,-1.362 | 0.034,0.031,-0.926,0.906,1.301,1.131 | 0.080,0.069,-0.480,0.208,1.210,0.940 |
| **aa26c**7c6f738cc87a1548d65a791953f | 0.023,0.065,1.759,-1.809,-0.966,-0.621 | 0.015,0.061,1.133,-0.929,0.108,-1.887 | 0.088,0.044,-0.809,1.076,1.754,1.936 | 0.097,0.096,-0.441,-0.012,1.597,1.877 |
| **aa28b**20c790814068891dff2bfe2df39 | 0.087,0.019,1.614,-1.964,-0.806,0.300 | 0.099,0.023,0.701,-0.590,-0.727,-1.174 | 0.069,0.028,-1.383,1.392,1.452,-0.131 | 0.080,0.017,-0.702,0.381,1.807,1.525 |
| **aa311**c8e41bada48ddfef41603045586 | 0.099,0.027,0.912,-0.787,-1.674,0.508 | 0.030,0.069,0.741,-0.465,-0.356,-1.277 | 0.063,0.025,-1.230,1.724,0.941,0.640 | 0.080,0.040,-1.278,0.949,1.849,0.464 |
| **aa356**9f38e8f9c3aacd7bb32c663eeb8 | 0.045,0.027,1.692,-1.718,-0.602,-0.298 | 0.017,0.049,1.926,-0.785,-0.841,-1.112 | 0.082,0.060,-0.922,1.164,1.582,1.228 | 0.035,0.068,-1.191,0.867,0.750,0.667 |
| **aa451**eed3de093f86100d572f803c0f6 | 0.062,0.021,1.375,-1.758,-0.996,1.104 | 0.066,0.080,0.216,-1.061,-0.513,1.617 | 0.088,0.067,-1.841,0.588,1.823,0.426 | 0.090,0.073,-0.417,0.231,0.895,0.679 |
| **aa482**4a2d44362e2e06ac4a54d2a069a | 0.033,0.054,0.773,-0.636,-0.968,-0.889 | 0.084,0.095,0.118,-0.013,-0.775,0.022 | 0.055,0.097,-1.055,0.458,-0.823,1.298 | 0.025,0.012,-0.802,0.740,0.649,1.519 |
| **aa58d**ed646dc27999504a135bacf17b3 | 0.082,0.062,1.842,-1.385,-1.831,-0.202 | 0.017,0.087,0.328,-0.529,-0.632,0.060 | 0.094,0.041,-1.041,0.816,-0.792,0.626 | 0.051,0.067,-0.427,-0.165,0.580,1.800 |
| **aa86b**5cefeefbf9e7d75d5e2822a6605 | 0.043,0.065,1.932,-1.785,-0.474,-1.204 | 0.014,0.016,1.321,-1.258,-1.908,-0.497 | 0.020,0.062,-1.844,1.957,0.293,0.850 | 0.011,0.050,-1.289,1.173,0.749,0.583 |
| **aa89d**7e761b57a7c6c3adb7cb7371a40 | 0.060,0.020,1.087,-1.003,-1.622,-1.195 | 0.072,0.058,0.737,-0.873,-0.520,1.081 | 0.021,0.069,-1.155,1.085,1.461,1.095 | 0.085,0.021,-1.026,0.765,1.572,-0.630 |
| **aa987**d8f586ed9a000a540b1649558d5 | 0.013,0.084,1.631,-1.788,-1.899,-0.552 | 0.046,0.043,1.419,-1.049,0.161,-1.221 | 0.095,0.042,-1.636,1.862,1.887,1.002 | 0.092,0.080,-1.413,0.785,1.917,1.428 |
| **aab28**1ab90fdf3c670222f575d9f759c | 0.048,0.071,1.851,-1.751,-1.661,-0.142 | 0.037,0.061,0.605,-1.384,0.302,-1.098 | 0.081,0.065,-1.018,0.250,-0.060,1.237 | 0.080,0.057,-0.754,0.499,1.139,1.030 |
| **aad22**054ca636a9b3ac70b8ae2dba83a | 0.031,0.083,0.096,0.027,0.192,-1.541 | 0.070,0.039,1.049,-1.245,-1.270,1.122 | 0.055,0.058,-1.750,1.520,1.733,1.712 | 0.059,0.066,-0.799,0.531,1.982,1.398 |
| **aae63**e55f8b6feaadf19d18d72fe3004 | 0.038,0.047,0.672,-0.587,-0.412,-1.102 | 0.049,0.058,0.532,-1.400,1.361,0.277 | 0.044,0.042,-1.946,1.143,1.307,1.414 | 0.098,0.054,-1.129,0.658,1.301,1.498 |
| **ab221**d1770a01c0e9605a0c0706a4202 | 0.086,0.068,1.727,-1.931,-0.711,0.276 | 0.092,0.091,0.547,-0.283,-1.115,0.053 | 0.066,0.088,-0.240,0.191,0.796,0.028 | 0.098,0.071,-0.714,0.545,1.069,-0.063 |
| **ab42c**a6f06443971c356133ec80f1e89 | 0.067,0.054,1.460,-1.620,-0.542,-0.598 | 0.042,0.088,0.153,-0.096,0.169,-0.852 | 0.052,0.099,-1.178,1.293,-0.226,-1.430 | 0.011,0.096,-0.909,0.757,1.680,1.018 |
| **ab598**b47f78bbbc5078ed0bfb397a2b8 | 0.035,0.086,1.527,-1.353,-0.622,-1.518 | 0.016,0.017,0.400,-0.318,-1.364,-0.973 | 0.028,0.068,-0.646,0.093,1.906,1.900 | 0.074,0.096,-0.955,0.415,1.978,0.580 |
| **ab5ef**0eddb71abde4ff3e7f3df939cc8 | 0.069,0.062,1.610,-1.177,-0.757,-1.701 | 0.044,0.021,1.914,-1.288,-1.458,0.495 | 0.033,0.057,-1.074,0.989,1.098,1.862 | 0.057,0.070,-0.425,-0.037,1.210,0.782 |
| **ab62f**d608ac3cd7a8665a05dc241336f | 0.048,0.066,1.932,-1.683,-1.174,-0.836 | 0.065,0.016,0.719,-1.525,-1.521,-0.420 | 0.032,0.028,-0.708,0.231,0.151,0.752 | 0.092,0.063,-0.409,0.245,1.178,0.171 |
| **ab8f3**f6cedcf6f4656fff1c3bbad368f | 0.023,0.024,1.236,-1.395,-1.201,-1.487 | 0.058,0.040,0.898,-0.467,-0.680,-0.914 | 0.091,0.013,-1.341,1.964,1.493,1.655 | 0.097,0.093,-1.360,0.559,1.739,1.735 |
| **ab918**985f63ed08dee728ad0c6a8c5b5 | 0.035,0.086,1.648,-1.564,-0.932,-1.245 | 0.011,0.046,0.465,-0.365,-0.461,0.344 | 0.084,0.087,-1.284,1.396,-0.244,1.608 | 0.097,0.060,-1.488,1.274,-0.050,1.118 |
| **abf94**575438e3eecebc59ae7d27cbecf | 0.052,0.065,1.924,-1.644,-1.799,-1.037 | 0.037,0.022,0.771,-0.337,-0.879,-1.030 | 0.039,0.058,-0.757,1.727,1.259,-0.135 | 0.080,0.076,-0.258,-0.420,1.578,1.548 |
| **ac0f0**d23e0b6c7b292d6ddf6c3b2c268 | 0.060,0.057,1.674,-1.791,-0.861,0.664 | 0.018,0.047,1.988,-1.199,-1.717,1.771 | 0.025,0.087,-1.198,1.740,0.896,1.238 | 0.038,0.097,-0.295,-0.017,1.644,-0.213 |
| **ac691**d7d0025f726717bcd63759dc4a7 | 0.043,0.068,1.049,-0.835,-1.774,0.017 | 0.022,0.040,0.322,-1.205,-0.259,-0.405 | 0.034,0.089,-1.118,0.203,1.935,0.474 | 0.097,0.038,-0.836,0.761,0.727,-0.261 |
| **ac6a7**7e0573e7a9a5b3654f9fff2dc8e | 0.085,0.074,1.920,-1.537,-1.397,-0.489 | 0.019,0.051,1.607,-1.565,-0.100,-0.950 | 0.099,0.059,-1.865,1.260,1.948,0.830 | 0.077,0.043,-0.458,0.099,1.610,0.703 |
| **ac7c2**0be9c62c8f288e2c22561b05499 | 0.089,0.060,1.922,-1.791,-1.634,1.383 | 0.033,0.081,0.808,-0.946,1.493,-1.299 | 0.064,0.066,-1.317,0.819,1.831,0.963 | 0.079,0.075,-0.278,-0.330,1.799,1.669 |
| **ac81c**6a528e3c57ea4769424c5f51b2f | 0.096,0.013,1.332,-0.908,-1.160,-0.514 | 0.070,0.060,0.537,-0.646,-1.800,0.493 | 0.016,0.084,-1.464,0.603,-0.179,1.097 | 0.050,0.055,-0.114,-0.141,1.113,1.156 |
| **ac8b3**ed26807d5365d3bb13ad75b4c4d | 0.050,0.051,1.583,-1.785,-0.599,0.435 | 0.013,0.019,0.863,-0.593,-0.657,-0.938 | 0.017,0.046,-1.059,1.750,1.315,0.284 | 0.071,0.090,-0.323,-0.205,1.942,0.775 |
| **ac96f**bb3597f54768b01028084279b9b | 0.036,0.083,0.976,-0.549,-1.614,-1.664 | 0.029,0.024,0.999,-0.995,1.938,-1.789 | 0.099,0.048,-1.652,1.892,-0.527,1.541 | 0.085,0.087,-1.085,0.710,1.569,-0.200 |
| **ace7a**7e3fe513a5a28ecd040447e12b7 | 0.068,0.028,1.178,-1.575,-0.539,1.330 | 0.037,0.052,0.884,-0.660,-0.733,0.105 | 0.035,0.048,-0.486,0.446,1.548,0.338 | 0.012,0.061,-0.146,-0.097,1.783,1.070 |
| **acf26**06d9f4bec4641e7f46c23d18e85 | 0.074,0.019,1.380,-1.790,-0.764,0.355 | 0.060,0.042,1.014,-0.948,0.656,-1.342 | 0.060,0.062,-1.730,1.706,1.398,0.425 | 0.068,0.084,-1.314,0.894,0.481,1.203 |
| **acfc5**66b9f406473c9a128aa8b82d9ec | 0.098,0.034,1.074,-0.627,-1.444,0.691 | 0.035,0.053,0.417,-0.424,-0.905,-1.940 | 0.096,0.082,-0.782,0.370,1.094,-1.121 | 0.048,0.046,-0.543,0.205,1.856,1.716 |
| **ad137**d50d23422f05006416065e22587 | 0.041,0.020,1.413,-1.514,-0.700,-0.674 | 0.095,0.048,1.382,-1.927,-1.546,0.304 | 0.075,0.086,-1.751,1.190,1.440,-0.271 | 0.077,0.096,-1.900,1.435,1.822,1.308 |
| **ad161**81c69f6da4c347f1ffcc2b15ddd | 0.061,0.019,1.423,-1.516,-1.133,-0.089 | 0.063,0.039,0.886,-0.543,-1.147,0.018 | 0.053,0.021,-0.876,1.700,-0.162,1.896 | 0.098,0.057,-1.048,0.861,1.119,-0.262 |
| **ad362**7ea6be0c2f8f78db634021747d8 | 0.052,0.027,1.629,-1.276,-1.845,-0.019 | 0.077,0.011,0.255,-0.130,-0.999,-1.704 | 0.080,0.034,-1.176,0.374,-0.086,1.675 | 0.082,0.076,-1.016,0.440,1.943,1.317 |
| **ad50a**b86acb3fc4c229df97f3059cc0d | 0.090,0.065,1.398,-1.459,-0.580,-0.526 | 0.042,0.060,0.665,-0.825,-1.837,0.574 | 0.047,0.018,-0.641,0.510,1.775,0.739 | 0.097,0.095,-1.331,1.107,1.026,0.508 |
| **ad78b**f12befcbee9a24eb7955a04ae98 | 0.071,0.048,0.569,-0.189,-1.504,-1.211 | 0.020,0.063,0.577,-0.492,0.310,-1.009 | 0.062,0.045,-0.602,0.090,1.778,1.886 | 0.066,0.097,-0.936,0.431,-0.209,1.827 |
| **ad8c4**72a1f30dde1d3193206eac5d9ec | 0.089,0.036,1.894,-1.922,-1.371,-0.217 | 0.016,0.083,0.481,-0.282,-0.915,-1.852 | 0.050,0.041,-0.927,0.734,-0.195,0.954 | 0.087,0.055,-0.436,0.132,0.966,1.665 |
| **ad965**0725de7174271300732fb92097a | 0.099,0.069,1.819,-1.061,-0.732,-1.669 | 0.013,0.078,1.199,-1.924,0.103,-1.024 | 0.015,0.075,-1.957,0.835,1.342,1.261 | 0.010,0.070,-0.608,0.310,1.404,1.214 |
| **ada10**3c2f9a7ba4c8c3903df92d0ecc9 | 0.028,0.094,1.365,-1.475,0.309,-1.825 | 0.065,0.022,0.585,-0.373,-0.516,-1.654 | 0.084,0.091,-1.584,1.972,-0.968,0.457 | 0.070,0.071,-1.145,0.592,1.952,1.456 |
| **adc34**3e154b3a3ae50e729eb650784b7 | 0.043,0.032,1.721,-1.592,-1.500,-0.541 | 0.057,0.054,1.077,-0.400,-0.862,-1.585 | 0.053,0.061,-0.530,0.774,1.629,0.474 | 0.074,0.065,-0.116,-0.367,1.569,1.078 |
| **adcc3**d23e1dd41a2240a97bac0acebc1 | 0.076,0.066,1.829,-1.262,-1.866,-1.372 | 0.077,0.082,0.327,0.024,-1.344,-0.942 | 0.069,0.086,-1.208,0.554,-0.642,1.505 | 0.088,0.040,-0.674,0.359,0.971,1.906 |
| **adeff**901737094267183fb25039d4fd5 | 0.078,0.020,1.727,-1.282,-1.095,-1.168 | 0.080,0.045,1.026,-0.406,-1.929,-0.944 | 0.073,0.059,-1.466,0.491,0.211,1.906 | 0.037,0.071,-0.621,-0.101,1.312,1.635 |
| **adf98**8595a75ccf980c948fffecd8337 | 0.046,0.087,1.549,-1.883,1.218,-0.959 | 0.039,0.048,1.024,-1.166,0.949,0.415 | 0.057,0.013,-1.808,1.830,1.401,1.159 | 0.056,0.063,-0.462,0.159,1.956,1.476 |
| **ae273**b7745a0c065c1b8578712693c48 | 0.044,0.098,1.939,-0.431,0.145,-1.614 | 0.066,0.048,0.422,0.177,-1.461,-1.275 | 0.058,0.090,-1.375,0.278,0.602,0.102 | 0.092,0.060,-1.439,1.103,0.133,1.508 |
| **ae33c**63078cdb03aa8f831c1d04635a7 | 0.062,0.090,1.686,-1.624,-0.464,-0.394 | 0.034,0.048,0.660,-0.403,-1.149,-1.628 | 0.095,0.075,-0.961,0.737,0.554,0.721 | 0.059,0.091,-0.530,-0.015,1.783,1.189 |
| **ae4dd**4f73ef9f30fdd8f02669e6997ce | 0.045,0.056,0.830,-0.908,-0.460,-0.171 | 0.069,0.051,0.366,-0.731,-0.091,0.203 | 0.035,0.047,-0.448,0.267,0.750,0.226 | 0.058,0.059,-0.319,0.182,1.938,0.221 |
| **ae60a**53c60f90c455b709c4e262e0218 | 0.075,0.041,0.909,-0.926,-1.289,1.347 | 0.095,0.042,0.488,-0.743,0.456,-0.312 | 0.085,0.085,-1.091,0.707,1.961,-0.403 | 0.025,0.040,-0.307,0.208,0.970,0.366 |
| **aeb84**4a5d2f336cf49fb59fa7077d048 | 0.078,0.022,1.565,-1.774,-1.108,0.504 | 0.097,0.051,1.028,-0.628,0.682,-1.019 | 0.076,0.097,-0.846,1.466,1.525,0.509 | 0.051,0.091,-0.751,0.465,-0.037,1.375 |
| **aecea**42518deb5424abb2c3f993036aa | 0.083,0.050,1.071,-0.842,-0.536,-1.437 | 0.058,0.020,0.467,-0.475,-1.427,0.064 | 0.060,0.049,-1.274,0.717,0.898,0.730 | 0.041,0.068,-0.040,-0.198,0.969,1.460 |
| **aed03**63d14c047494f07fbcba87472c1 | 0.079,0.058,1.819,-1.300,-1.900,-1.194 | 0.031,0.087,0.714,-0.880,1.586,-0.400 | 0.069,0.081,-0.457,-0.002,1.728,1.188 | 0.062,0.024,-0.036,-0.212,1.817,0.801 |
| **aed7e**fedc1414163d1e57f4443801f3f | 0.019,0.063,1.906,-1.729,-1.458,-0.549 | 0.071,0.066,1.141,-1.877,-0.807,-0.043 | 0.052,0.047,-1.037,0.427,0.937,0.745 | 0.054,0.071,-0.302,0.035,1.685,0.489 |
| **aed83**980e2a4e44df99747156cf60a8b | 0.017,0.015,1.621,-1.589,-0.433,-1.289 | 0.090,0.045,0.545,-1.005,-0.459,-1.350 | 0.060,0.069,-0.999,0.401,1.005,1.368 | 0.083,0.059,-0.345,0.053,1.086,1.288 |
| **aedf1**5c0197fb18ccebcd394ebd3826b | 0.057,0.033,1.541,-1.385,-1.435,-0.095 | 0.017,0.042,0.649,-1.391,-1.541,-0.245 | 0.066,0.080,-0.836,0.110,1.198,1.229 | 0.054,0.076,-0.178,-0.105,0.905,1.093 |
| **aeefd**85be155e2c84c5ececbd689fe57 | 0.059,0.057,1.600,-1.869,-0.880,0.647 | 0.094,0.029,0.708,-1.141,-0.341,0.736 | 0.048,0.086,-0.692,0.270,1.232,0.432 | 0.083,0.051,-0.403,-0.085,1.074,1.383 |
| **af0a6**d2222662a1c17ae2e8733328c08 | 0.092,0.050,1.853,-1.606,-1.573,0.179 | 0.027,0.082,1.644,-1.240,-0.428,-1.861 | 0.069,0.047,-1.781,1.840,1.019,0.548 | 0.066,0.082,-1.096,0.594,1.116,1.404 |
| **af0d9**8e9ad13e602d8a3eb78417159a9 | 0.059,0.054,1.605,-1.595,-1.567,-0.544 | 0.068,0.047,0.548,-0.461,-0.316,-0.189 | 0.080,0.063,-0.754,0.812,1.842,1.048 | 0.081,0.094,-1.160,0.719,1.364,1.336 |
| **af107**2e9965159f213562db2e0da47de | 0.045,0.019,1.606,-1.632,-1.922,-0.449 | 0.093,0.026,0.477,-1.070,-0.503,-0.204 | 0.011,0.090,-1.641,0.936,-0.151,0.631 | 0.073,0.094,-0.737,0.489,1.906,0.418 |
| **af2aa**22ce4608d535142b73e542b963f | 0.076,0.052,1.780,-1.272,-0.634,-1.321 | 0.022,0.014,1.928,-1.113,-0.278,-0.051 | 0.100,0.033,-1.630,1.275,1.898,1.772 | 0.060,0.050,0.156,-1.366,1.558,1.292 |
| **af49b**67d56eeab0099bdc3376ec20b66 | 0.034,0.071,0.752,-0.798,-0.800,-0.120 | 0.014,0.058,0.289,-0.301,0.326,-0.816 | 0.020,0.012,-0.314,0.250,0.859,0.947 | 0.042,0.089,-0.358,0.083,0.962,1.163 |
| **af652**c048e67216d27b5e1ab5be2ad5c | 0.099,0.028,1.793,-1.729,-1.784,0.590 | 0.020,0.020,0.580,-0.830,0.662,-0.939 | 0.098,0.060,-1.719,1.384,-0.229,0.902 | 0.027,0.090,-0.838,0.247,1.979,1.649 |
| **af77f**88ff8828dddeaa9925ba096ecd3 | 0.042,0.054,1.661,-1.355,-0.511,-1.223 | 0.016,0.096,0.360,-1.322,-0.507,1.202 | 0.098,0.051,-1.221,0.198,1.925,0.307 | 0.096,0.093,-1.014,0.183,1.966,1.392 |
| **af882**a5eb1c230f9c7913206a371d7af | 0.058,0.055,0.575,-0.474,-1.309,-0.321 | 0.066,0.093,0.934,-1.276,-0.644,1.574 | 0.036,0.056,-1.313,1.232,1.646,1.095 | 0.019,0.088,-0.646,0.318,1.681,0.891 |

| | | | | |
|---|---|---|---|---|
| **afc8b**236ad0d3e2ae1df64a744d0a803 | 0.097,0.027,1.358,-1.465,-0.753,0.224 | 0.031,0.012,0.497,-0.839,-0.853,-0.774 | 0.098,0.068,-0.476,0.061,1.457,-0.572 | 0.065,0.064,-0.550,0.133,1.922,1.028 |
| **afca0**2880f942375e3ac036ce67a559b | 0.034,0.062,1.696,-1.860,-1.812,-0.703 | 0.015,0.020,0.541,-0.578,-1.239,-1.101 | 0.069,0.022,-1.249,1.045,1.231,1.468 | 0.097,0.045,-1.530,1.192,1.544,1.489 |
| **b024f**9a5bd35e7878650263b3677f285 | 0.024,0.022,1.546,-1.849,-0.672,1.237 | 0.098,0.073,1.275,-1.651,0.487,-0.813 | 0.015,0.046,-0.851,0.646,1.335,0.376 | 0.072,0.014,-1.204,1.032,0.986,0.125 |
| **b02b1**0421fa527e4bd26306777616a85 | 0.053,0.021,1.385,-1.383,-1.172,-0.842 | 0.021,0.084,0.709,-0.813,-1.907,-1.303 | 0.069,0.058,-0.824,0.660,1.260,-1.093 | 0.097,0.088,-0.790,0.410,1.654,0.862 |
| **b02f0**d470dd4987200635023637f818b | 0.047,0.043,1.556,-1.796,-1.339,-0.082 | 0.014,0.022,1.017,-0.920,-1.968,0.938 | 0.098,0.070,-1.642,1.880,1.362,0.070 | 0.051,0.068,-1.394,1.309,0.322,0.219 |
| **b0539**130e9b2e27e8f94983ec4121a73 | 0.081,0.086,0.652,-0.679,-1.110,0.464 | 0.049,0.049,1.030,-0.530,-0.803,-1.422 | 0.097,0.035,-1.716,1.973,1.694,0.609 | 0.029,0.096,-0.370,0.030,1.635,0.639 |
| **b07ae**83c9161fe35d03b33eba33fa393 | 0.089,0.021,1.729,-1.504,-1.880,0.893 | 0.059,0.085,1.069,-1.054,0.108,-1.533 | 0.074,0.031,-1.142,0.734,0.039,1.850 | 0.066,0.077,-1.212,1.001,1.234,0.105 |
| **b0851**d4b0204f066f5f782e80a7591be | 0.093,0.058,1.751,-1.279,-1.658,-1.227 | 0.075,0.061,0.420,-0.217,-1.741,0.601 | 0.075,0.035,-1.618,1.421,0.045,0.561 | 0.092,0.076,-1.304,0.416,1.916,1.459 |
| **b0903**e3359e62c453fa7826b1a4a9c03 | 0.066,0.042,1.714,-1.721,-1.041,-1.567 | 0.083,0.051,0.656,-0.419,-1.256,-1.456 | 0.080,0.068,-1.113,0.916,1.728,0.380 | 0.064,0.087,-0.907,0.625,0.879,1.340 |
| **b0e17**0ca1043c10b57e1d121eebf83ae | 0.056,0.039,1.241,-1.230,-1.924,-0.368 | 0.044,0.019,0.489,-0.302,-0.736,0.414 | 0.030,0.031,-1.093,1.862,-0.694,0.718 | 0.065,0.051,-1.042,0.709,1.595,1.023 |
| **b1323**eed8ad0996c456f5a662c812ab6 | 0.071,0.011,1.562,-1.705,-1.266,0.583 | 0.062,0.082,0.987,-1.028,-0.888,-1.457 | 0.069,0.071,-1.572,0.949,1.463,0.244 | 0.078,0.095,-1.163,0.781,1.839,0.830 |
| **b14cf**078d8577c5d6d66038ba265b7d1 | 0.095,0.036,1.971,-1.508,-1.841,-1.068 | 0.093,0.075,1.292,-1.782,-1.030,0.235 | 0.073,0.059,-1.711,0.995,-0.224,1.382 | 0.067,0.068,-1.057,0.638,1.874,0.754 |
| **b1552**afc08ebed9677e34c868253446f | 0.074,0.048,1.792,-1.541,-1.161,-1.197 | 0.056,0.083,0.395,-0.956,-0.049,-0.198 | 0.026,0.046,-0.277,-0.072,1.431,1.721 | 0.062,0.067,-1.133,0.885,0.106,1.911 |
| **b160a**ce52f568fad0069dc74c3860a1d | 0.072,0.050,1.558,-1.225,-1.894,-1.127 | 0.011,0.049,1.679,-1.632,1.667,-1.837 | 0.094,0.035,-1.528,1.586,1.866,1.386 | 0.085,0.075,-0.685,0.343,0.597,1.695 |
| **b175c**9eca090dd7083c6fbda298a46a9 | 0.023,0.056,1.899,-1.500,-1.807,-1.903 | 0.021,0.023,1.602,-1.295,-1.229,-0.171 | 0.016,0.064,-1.405,1.346,0.619,1.348 | 0.040,0.063,-0.103,-0.244,1.648,0.931 |
| **b19af**20b8517d89ee9c7e1efcb318129 | 0.099,0.064,1.724,-1.258,-1.879,-0.449 | 0.023,0.065,1.114,-1.577,1.694,-1.398 | 0.064,0.053,-1.036,0.442,1.097,1.655 | 0.049,0.065,-0.413,-0.095,1.809,1.867 |
| **b1d97**3925d1404d4ebf5a42e3c23c3f4 | 0.049,0.048,1.607,-1.722,-1.518,-0.029 | 0.099,0.014,1.200,-1.935,-0.445,0.314 | 0.042,0.050,-1.754,1.158,1.109,0.484 | 0.034,0.066,-0.763,0.713,-0.165,0.393 |
| **b1dc9**735961d80ca477a8f7e316da8bc | 0.043,0.065,0.812,-0.597,-0.884,-0.447 | 0.079,0.071,1.494,-1.307,-0.685,-0.312 | 0.049,0.059,-1.390,0.991,1.924,1.079 | 0.066,0.049,-0.997,0.580,1.419,0.968 |
| **b1e28**2d9b4ead3bb0bec4495306c18f4 | 0.094,0.064,1.812,-1.652,-1.973,-0.322 | 0.041,0.075,1.653,-1.750,-1.072,-1.939 | 0.067,0.031,-1.412,1.214,1.153,-1.731 | 0.081,0.059,-0.346,-0.089,1.979,1.424 |
| **b1fb2**01010005cfa2576b6d1b517a0e0 | 0.088,0.023,1.768,-1.921,-1.703,1.081 | 0.053,0.014,0.685,-1.041,-1.364,1.181 | 0.068,0.054,-1.639,0.968,1.931,0.455 | 0.023,0.078,-1.814,1.581,0.134,1.857 |
| **b2103**7e91336d432fa322b5dc74a1c54 | 0.089,0.060,1.155,-0.691,-1.959,0.119 | 0.029,0.074,0.352,-0.790,1.084,-1.179 | 0.057,0.040,-1.647,0.542,1.893,0.782 | 0.016,0.077,-1.503,1.235,1.564,1.820 |
| **b224f**93327d8ba51ab3eacfa89638e78 | 0.093,0.012,1.466,-0.893,-1.618,0.115 | 0.057,0.085,0.929,-1.712,0.095,0.201 | 0.011,0.085,-1.825,0.647,1.576,1.943 | 0.037,0.069,-0.200,-0.170,1.774,0.790 |
| **b2295**0245fdd78bc0e251c65c975ae79 | 0.092,0.014,1.623,-0.958,-1.800,0.688 | 0.056,0.088,1.323,-0.952,-1.126,-1.878 | 0.089,0.055,-1.953,1.095,-0.226,1.785 | 0.061,0.083,-1.012,0.223,1.521,1.648 |
| **b243d**c2fd3b9cda962fa70c5aac89994 | 0.046,0.064,0.647,-0.721,-0.010,-1.133 | 0.077,0.090,0.373,-0.125,-1.399,-0.301 | 0.076,0.041,-0.294,0.181,1.770,0.358 | 0.073,0.072,-0.531,0.138,1.942,1.594 |
| **b24bc**29ecd3ba0b13bcb5d2a99419d58 | 0.012,0.061,1.584,-1.628,-0.172,-1.216 | 0.064,0.060,0.362,-0.202,-0.221,-0.175 | 0.016,0.086,-1.005,1.052,-0.182,1.925 | 0.051,0.073,-0.971,0.541,0.217,1.509 |
| **b24c5**4ba2260d12f3ecef8c2f9be0985 | 0.087,0.050,1.276,-1.107,-1.523,-0.960 | 0.074,0.048,0.901,-0.854,-0.174,0.360 | 0.011,0.053,-1.352,1.918,1.089,-0.198 | 0.066,0.068,-0.658,0.214,0.967,1.423 |
| **b24e8**da5be216d744bfbb33a7bbbc7ce | 0.099,0.056,1.536,-1.239,-1.707,-0.631 | 0.031,0.050,1.376,-0.806,0.062,-1.030 | 0.085,0.025,-0.987,1.333,1.752,0.539 | 0.042,0.075,-0.207,-0.100,-0.719,1.844 |
| **b2684**2c4a5377e256ba2a4974432b611 | 0.064,0.046,1.904,0.191,-1.970,-0.352 | 0.070,0.061,0.581,-0.515,0.015,-1.517 | 0.073,0.028,-1.367,-0.176,1.566,0.190 | 0.051,0.061,-0.339,-0.225,1.227,0.687 |
| **b2731**1725c70eadd9a41f0eb0e14a957 | 0.085,0.065,1.050,-1.278,-1.784,0.476 | 0.029,0.063,0.312,-0.274,-0.081,-0.166 | 0.037,0.091,-0.666,0.568,-0.815,0.922 | 0.085,0.060,-0.332,-0.166,1.893,1.204 |
| **b2735**dfafdde0efe713b5b7715f364d6 | 0.052,0.043,0.998,-1.243,0.126,-0.496 | 0.018,0.085,0.283,-0.348,0.878,0.076 | 0.095,0.022,-1.702,1.857,1.170,1.475 | 0.062,0.045,-1.190,0.823,0.802,1.259 |
| **b2beb**76d654d9c2c4ccc6a53597db044 | 0.054,0.022,0.418,-0.427,-1.179,0.119 | 0.083,0.058,0.625,-0.391,-0.871,0.188 | 0.097,0.078,-1.491,1.824,1.438,1.888 | 0.060,0.082,-0.648,0.408,1.242,1.381 |
| **b2d41**be16e702acde34fed6fad64af15 | 0.043,0.078,0.863,-0.670,0.869,-1.988 | 0.094,0.080,1.482,-0.698,0.775,-1.371 | 0.071,0.072,-1.177,1.917,1.488,0.825 | 0.042,0.019,-1.146,0.864,0.911,1.630 |
| **b2da0**7a5d231ff7ac1f0c5998ce13533 | 0.094,0.047,1.884,-1.904,-1.847,0.688 | 0.041,0.091,0.884,-0.593,-1.680,-1.352 | 0.087,0.031,-1.967,1.782,1.838,0.974 | 0.017,0.028,-1.544,1.427,1.640,1.611 |
| **b2e23**c6ee0d6671245fb469cf0d2c806 | 0.075,0.053,1.531,-1.339,-0.553,-0.559 | 0.023,0.058,0.608,-1.207,-0.853,-1.120 | 0.060,0.074,-0.722,0.263,1.706,-0.889 | 0.093,0.011,-0.654,0.299,1.816,0.362 |
| **b3022**9de87d919bcfbe34064a071720b | 0.093,0.021,1.513,-1.776,-1.828,0.008 | 0.028,0.018,0.373,-0.306,-1.571,1.739 | 0.083,0.066,-1.143,1.594,1.344,0.405 | 0.088,0.087,-1.239,1.209,1.628,-0.975 |
| **b3038**b7c4c991e796833278f738679e5 | 0.013,0.065,1.682,-1.631,-0.312,-1.505 | 0.019,0.022,1.530,-0.918,-0.145,-1.607 | 0.022,0.060,-1.073,1.626,1.963,0.881 | 0.052,0.061,-1.392,1.030,-0.141,1.703 |
| **b3061**805c4a47845556f3ac6981d343c | 0.042,0.047,1.560,-1.884,-1.834,0.617 | 0.047,0.063,0.604,-0.572,1.220,-1.506 | 0.068,0.061,-0.898,0.672,1.828,1.933 | 0.030,0.071,-0.695,0.410,1.952,1.565 |
| **b30ec**783b56585e888ce3816313b7a5e | 0.081,0.042,1.818,-1.362,-1.752,-0.352 | 0.066,0.026,1.092,-1.317,-1.050,-0.837 | 0.026,0.071,-1.788,1.264,-0.522,0.696 | 0.089,0.033,-0.297,-0.349,1.902,0.761 |
| **b311b**8e870731bb7e6b6b380c6dc25ba | 0.096,0.036,1.495,-1.401,-1.892,0.864 | 0.021,0.070,0.594,-0.369,-1.060,-0.825 | 0.050,0.035,-0.580,0.683,1.604,-1.252 | 0.040,0.054,-0.070,-0.306,1.637,1.806 |
| **b314d**6ded76bafa0a985f0090dd1a63b | 0.082,0.027,1.476,-0.897,-1.936,1.577 | 0.099,0.022,0.219,-0.005,-0.783,-1.547 | 0.030,0.083,-1.100,0.142,1.417,0.985 | 0.080,0.065,-0.573,0.231,1.420,0.916 |
| **b31bf**769e5d10f860612bcc1622d5216 | 0.047,0.086,1.891,-1.852,-1.745,0.345 | 0.065,0.035,1.426,-1.119,1.192,0.789 | 0.098,0.060,-0.808,1.488,1.417,0.689 | 0.030,0.080,-0.715,0.380,1.576,1.720 |
| **b31f2**f35c8ada757e6b17787b334ec1a | 0.051,0.062,0.986,-0.789,-0.482,-1.109 | 0.028,0.020,1.172,-1.817,-1.794,0.616 | 0.053,0.088,-1.642,0.760,0.370,1.399 | 0.066,0.038,-1.127,0.860,1.111,1.115 |
| **b32be**db11d582cf5311432fdb8ea31dd | 0.089,0.071,1.122,-1.513,-1.329,1.196 | 0.083,0.083,0.759,-0.452,0.629,-0.914 | 0.098,0.017,-1.131,1.947,1.589,0.718 | 0.016,0.068,-0.512,0.294,0.583,1.302 |
| **b38ea**974a4fec80b6bb2e713be031604 | 0.045,0.029,1.405,-1.525,-0.272,-0.266 | 0.042,0.079,1.436,-1.745,-1.660,-1.112 | 0.069,0.064,-1.223,0.917,0.488,0.154 | 0.037,0.013,-0.214,0.077,1.217,0.806 |
| **b3a59**235cf35d7bd1fc719b8ece4c4a1 | 0.021,0.085,1.714,-1.716,0.296,-1.733 | 0.012,0.054,1.181,-0.811,-0.326,-0.850 | 0.085,0.072,-1.316,1.847,-0.684,1.340 | 0.092,0.075,-0.624,-0.223,1.784,1.757 |
| **b3d12**58f4a9e80fabf8db071ecc1046a | 0.051,0.042,0.724,-0.574,-1.284,-0.777 | 0.068,0.080,1.374,-1.341,0.058,-1.929 | 0.018,0.068,-0.842,0.610,-0.685,1.950 | 0.092,0.055,-0.565,0.169,1.866,1.145 |
| **b3e17**d000479153e66a989d9526dbb5e | 0.072,0.028,0.970,-0.870,-1.460,0.603 | 0.092,0.083,1.191,-0.853,-0.396,-1.032 | 0.092,0.098,-1.858,1.942,1.904,0.698 | 0.024,0.047,-1.695,1.524,-1.122,1.818 |
| **b4032**3cfeed146826d35baadf9311a5c | 0.035,0.022,1.653,-1.733,-0.984,-0.644 | 0.073,0.043,1.258,-1.143,-1.425,-0.592 | 0.079,0.020,-1.323,1.155,0.007,1.159 | 0.097,0.057,-0.657,0.353,1.973,-0.477 |
| **b40ed**ca1d0832fb247b288abba02acd4 | 0.078,0.067,1.050,-0.678,-1.064,-1.030 | 0.095,0.053,0.999,-0.964,-0.858,0.317 | 0.057,0.072,-1.508,1.029,-0.126,1.977 | 0.046,0.072,-1.003,0.651,1.615,0.869 |
| **b418b**2338be2eb4a72118069ae076674 | 0.042,0.017,0.559,-0.543,-1.747,0.854 | 0.088,0.057,0.503,-0.285,-0.979,0.055 | 0.084,0.039,-0.564,0.534,1.482,0.065 | 0.051,0.031,-1.281,1.140,0.106,1.641 |
| **b460e**57c7ceaaf5626e7a54658d3f901 | 0.058,0.053,1.903,-1.559,-1.465,0.088 | 0.019,0.028,1.005,-1.204,-0.398,-0.216 | 0.048,0.068,-1.429,0.782,1.911,0.808 | 0.071,0.062,-0.501,0.155,-0.079,1.521 |
| **b475d**f7ff33aae80cc69162b21061e7b | 0.045,0.088,1.507,-1.063,-1.720,-1.052 | 0.097,0.062,1.383,-1.850,-1.162,-0.414 | 0.095,0.050,-1.969,1.017,1.769,1.287 | 0.064,0.022,-1.162,0.975,1.250,0.597 |
| **b4877**394073a3c30a64a27fa42a586bb | 0.069,0.018,0.732,-0.633,-1.275,1.127 | 0.050,0.075,1.687,-1.851,-1.537,-1.907 | 0.026,0.062,-1.697,1.815,-0.853,0.434 | 0.095,0.083,-1.033,0.667,1.393,1.089 |
| **b4b85**56d3673679cee40e18e2b635f93 | 0.040,0.084,1.549,-1.612,-0.163,-0.862 | 0.057,0.087,1.635,-0.614,-1.282,-1.460 | 0.061,0.058,-0.667,0.513,1.859,1.586 | 0.037,0.053,-0.301,0.076,1.531,0.454 |
| **b4cc1**f249a02890b124e2e9cf11602a8 | 0.051,0.062,0.706,-0.259,-1.854,-1.877 | 0.023,0.033,0.719,-0.922,-1.657,1.032 | 0.093,0.077,-0.923,0.629,0.542,1.106 | 0.098,0.075,-0.222,-0.377,1.848,1.828 |
| **b4dbb**00ecf4418da1c79dadf63b9b4ed | 0.075,0.025,1.772,-1.853,-1.856,0.342 | 0.090,0.091,0.479,-0.373,-1.440,-0.307 | 0.074,0.070,-1.769,0.767,1.683,1.543 | 0.083,0.020,-0.507,0.331,1.229,0.876 |
| **b4df4**7cebd1dbce5494fda170c3b7acd | 0.087,0.046,1.724,-1.684,-0.695,-0.412 | 0.095,0.047,0.513,-0.420,0.384,-0.962 | 0.084,0.089,-1.289,0.817,1.100,1.677 | 0.044,0.068,-0.218,-0.266,1.855,0.386 |
| **b4e7d**dc8903caf7a692da636c90a058d | 0.046,0.084,1.558,-1.567,0.901,-1.154 | 0.023,0.037,0.945,-0.924,1.879,0.917 | 0.039,0.063,-1.422,1.981,1.811,0.209 | 0.028,0.064,-1.452,1.253,0.545,0.401 |
| **b4f7a**17000939ffdd9c5e68b3a1fe3f9 | 0.091,0.079,1.037,-0.790,-1.082,-0.432 | 0.080,0.048,1.953,-1.155,-1.798,1.342 | 0.080,0.096,-1.662,1.870,1.264,1.638 | 0.048,0.081,-1.782,1.088,1.712,1.371 |
| **b5090**181f9e1e688e702d3cc7bbd07f2 | 0.089,0.044,1.878,-1.668,-1.874,-1.333 | 0.043,0.061,1.142,-0.834,-0.824,-0.130 | 0.073,0.032,-1.499,1.939,-0.043,-1.328 | 0.053,0.075,-1.471,0.903,1.330,0.906 |
| **b52cb**0b61931b059e82a26cb67fee1ed | 0.081,0.057,1.904,-1.291,-1.148,-0.995 | 0.082,0.091,0.416,-0.265,-1.601,-0.126 | 0.015,0.038,-0.610,0.183,-1.063,0.686 | 0.022,0.059,-0.853,0.803,1.163,0.154 |
| **b5414**a9d7a2c62ed9245a568333fe653 | 0.019,0.023,1.222,-1.480,-0.001,-0.981 | 0.040,0.038,0.562,-0.376,-0.085,-0.050 | 0.029,0.060,-1.291,1.978,-0.639,1.866 | 0.095,0.063,-1.664,1.370,1.036,1.118 |
| **b547d**a32c788a56216cc4d95e6752312 | 0.069,0.051,1.714,-1.393,-1.795,-1.076 | 0.073,0.074,1.474,-1.892,-0.692,-1.539 | 0.099,0.096,-1.917,1.040,1.843,0.454 | 0.083,0.073,-0.105,-0.390,1.730,1.267 |
| **b54da**8eecfeb3fdb7c738eae98879c61 | 0.069,0.011,0.168,-0.098,-1.075,1.321 | 0.014,0.057,1.575,-0.960,-0.225,-1.079 | 0.096,0.027,-1.179,1.938,1.883,1.308 | 0.060,0.029,-0.830,0.628,1.185,0.695 |
| **b55b0**dbc828894c5b8dfcc0df950f712 | 0.047,0.028,1.132,-1.121,-1.054,-0.984 | 0.053,0.030,0.818,-0.590,-1.341,-0.290 | 0.090,0.077,-1.641,1.941,1.538,0.872 | 0.078,0.057,-1.514,1.112,1.728,1.563 |
| **b5749**7d41ae926b18c69f0ee930766fa | 0.078,0.021,1.648,-1.713,-0.602,-0.432 | 0.045,0.050,1.068,-1.140,-0.884,-1.482 | 0.086,0.093,-1.712,1.266,-0.025,0.664 | 0.038,0.075,-0.604,0.147,0.761,1.969 |
| **b57a3**6707f31a6f0b141a0d90d9e20a9 | 0.098,0.023,1.666,-1.765,-1.055,0.132 | 0.016,0.088,0.767,-0.520,-0.958,-1.665 | 0.085,0.029,-0.787,0.790,0.290,-0.274 | 0.083,0.077,-0.596,0.442,-0.287,1.238 |
| **b5946**05a889d1fa282017141a97c2653 | 0.052,0.065,1.579,-1.538,-0.960,-0.205 | 0.031,0.054,1.960,-1.600,-0.522,0.047 | 0.095,0.065,-1.347,1.876,-0.915,1.302 | 0.064,0.055,-1.597,0.924,1.973,1.620 |

| | | | | |
|---|---|---|---|---|
| **b5dbd**2c53539e89450b1fecc45e2e5b5 | 0.052,0.013,1.537,-1.551,-1.475,0.679 | 0.084,0.049,1.463,-1.003,-1.071,-0.160 | 0.045,0.062,-1.054,1.061,1.802,0.875 | 0.092,0.054,-1.133,0.522,1.184,1.678 |
| **b5ecb**a118ee57bffc91049be2edbde1b | 0.076,0.027,1.612,-1.822,-0.580,-0.181 | 0.045,0.090,0.194,-0.517,1.682,-0.150 | 0.064,0.081,-1.017,0.763,1.529,-0.040 | 0.074,0.075,-0.700,0.146,1.801,1.569 |
| **b6046**df01143641fd9b89a40b88b6730 | 0.091,0.083,0.637,-0.332,-0.573,-0.601 | 0.056,0.038,0.608,-0.694,-0.346,-0.042 | 0.071,0.025,-1.780,1.318,1.794,0.611 | 0.035,0.051,-1.421,1.156,1.592,1.000 |
| **b6188**60565788bd60c0ab9eb25c802a3 | 0.040,0.065,0.702,-0.639,-0.338,-0.393 | 0.042,0.043,0.781,-0.960,1.153,-0.454 | 0.050,0.037,-0.715,0.944,0.468,-0.470 | 0.082,0.062,-0.389,0.273,0.655,0.308 |
| **b6198**c9f08bdc4e786e77150c0f6ae04 | 0.086,0.043,1.881,-1.873,-0.513,-1.030 | 0.089,0.026,1.195,-1.405,-0.530,-1.583 | 0.097,0.056,-1.298,0.864,1.339,0.833 | 0.075,0.052,-0.268,-0.091,1.716,1.441 |
| **b62e8**275bc77f1fb5722351852e794af | 0.084,0.094,1.552,-1.699,-1.533,0.425 | 0.013,0.078,0.370,-0.265,1.253,-0.853 | 0.083,0.066,-1.235,1.938,1.966,-0.765 | 0.037,0.050,-1.070,0.932,-0.627,1.719 |
| **b6514**1d286600f07f857cc791c64c8e3 | 0.077,0.026,1.721,-1.752,-0.973,-0.699 | 0.062,0.030,1.346,-1.007,0.104,-1.244 | 0.098,0.088,-1.893,1.653,1.302,1.873 | 0.022,0.035,-1.474,1.261,1.712,0.999 |
| **b6580**a75c61dc778f35b5082924f6a82 | 0.052,0.080,1.797,-1.472,-1.665,-1.975 | 0.047,0.060,0.999,-0.750,-1.235,-1.329 | 0.026,0.038,-1.361,1.206,1.661,0.801 | 0.098,0.051,-1.571,1.090,1.257,1.753 |
| **b65bc**ea3bd97f7aefe634276edab858d | 0.020,0.054,0.433,-0.312,-0.031,-1.421 | 0.089,0.050,0.928,-1.409,-1.300,1.771 | 0.085,0.063,-1.763,1.307,1.911,0.786 | 0.053,0.097,-1.282,1.031,1.341,1.283 |
| **b6aeb**c5e58ad157c2aa72e3ad637102b | 0.027,0.067,1.066,-1.188,-0.232,-1.295 | 0.077,0.082,0.447,-0.184,-0.897,-0.477 | 0.015,0.058,-0.662,0.926,1.410,-0.078 | 0.034,0.012,-0.212,0.146,1.815,-0.362 |
| **b6b56**56a0f4f46c47bc2e249fab361de | 0.073,0.064,0.657,-0.602,-0.745,-0.183 | 0.092,0.074,0.546,-0.274,0.324,-1.409 | 0.070,0.069,-0.387,0.274,1.830,0.493 | 0.068,0.030,-0.117,-0.286,1.779,1.642 |
| **b6bfd**d46ac2bdce4a0dfe50c6776a52d | 0.016,0.078,1.952,-1.829,-1.977,-0.209 | 0.031,0.046,0.578,-0.854,-1.945,-1.668 | 0.048,0.046,-1.807,0.721,0.219,0.018 | 0.089,0.083,-1.160,0.822,0.671,1.139 |
| **b705c**dbb8149cf48fbeb72a5a731a7f3 | 0.065,0.028,1.175,-1.426,-0.389,-0.728 | 0.012,0.046,0.231,-0.291,-1.350,1.558 | 0.073,0.100,-0.772,0.875,-0.487,1.532 | 0.073,0.033,-1.486,1.321,1.876,-0.651 |
| **b7666**42bd5c6309dd2711850bd97f61f | 0.081,0.074,1.058,-1.090,-0.413,-0.952 | 0.045,0.077,0.151,-0.236,0.580,0.082 | 0.055,0.077,-1.208,1.278,-0.313,1.003 | 0.065,0.079,-1.671,1.258,1.158,1.218 |
| **b76a0**f68a748a2cecc300e26563091a5 | 0.016,0.055,1.887,-1.783,-1.925,-0.215 | 0.085,0.045,1.728,-1.277,-1.549,-1.185 | 0.036,0.094,-1.792,1.487,1.849,1.570 | 0.088,0.063,-0.131,-0.404,1.880,-0.278 |
| **b76de**bff751a19277796e5accee115a5 | 0.051,0.022,1.321,-1.433,-0.817,-0.322 | 0.092,0.060,0.222,-0.114,-1.697,1.133 | 0.053,0.058,-0.261,0.198,-0.422,1.048 | 0.090,0.034,-1.126,0.829,1.940,-0.337 |
| **b7826**8bbbc7afeb1c401cd74f1b0c7ea | 0.046,0.037,1.830,-1.733,-0.652,-0.518 | 0.096,0.022,1.849,-1.221,-1.173,-1.075 | 0.099,0.098,-1.430,1.627,0.577,1.746 | 0.089,0.037,-0.474,0.199,0.974,0.309 |
| **b7994**221404588dcfba456f9d8078bed | 0.076,0.016,1.692,-1.230,-1.448,-0.566 | 0.053,0.045,1.952,-1.179,-1.526,-1.572 | 0.093,0.029,-1.751,1.326,1.339,1.309 | 0.047,0.065,-1.308,1.139,1.061,-0.064 |
| **b7bbd**be86435499ddaf9ea697dafbff7 | 0.029,0.074,0.366,-0.372,-1.358,0.541 | 0.052,0.089,0.095,-0.191,-0.897,1.145 | 0.054,0.056,-0.340,0.311,1.050,0.707 | 0.082,0.094,-0.585,-0.184,1.900,1.856 |
| **b7bef**59feb31f42184d49eb93f412df5 | 0.084,0.039,1.401,-0.927,-1.904,-0.898 | 0.080,0.069,1.128,-1.480,-0.641,-1.589 | 0.027,0.044,-1.619,1.122,-0.846,0.572 | 0.036,0.086,-0.542,0.364,0.376,0.788 |
| **b7d0f**cc204152f54eaaac8cf61f69c6a | 0.086,0.018,0.768,-0.615,-1.291,-0.668 | 0.013,0.056,0.126,-0.294,-1.146,0.910 | 0.054,0.090,-1.630,0.998,1.424,1.660 | 0.076,0.036,-1.397,1.228,0.558,0.993 |
| **b7d40**1b13b155556ff1fdafcdcfb1646 | 0.099,0.022,0.971,-0.696,-1.621,0.350 | 0.033,0.018,0.346,-0.301,-0.079,-0.243 | 0.077,0.044,-0.424,0.245,0.668,1.051 | 0.039,0.082,-0.483,0.308,0.847,0.436 |
| **b846c**23a4f3abf1bd8e44a30be8d2d88 | 0.091,0.069,0.698,-0.723,-1.106,-0.000 | 0.016,0.054,0.722,-0.581,-1.852,-0.725 | 0.030,0.082,-0.440,0.347,1.205,0.478 | 0.041,0.095,-0.271,-0.124,1.206,1.440 |
| **b86cc**fe1d1a83c4cccb92a8a6a0d6c29 | 0.081,0.016,0.685,-0.598,-1.923,-0.241 | 0.032,0.021,0.557,-0.637,-0.247,0.860 | 0.100,0.054,-1.085,0.569,1.912,0.961 | 0.023,0.098,-0.647,0.451,-0.656,0.927 |
| **b8863**e241d9cea64fc1770636d6d4d1f | 0.039,0.049,1.902,-1.512,-1.574,-1.133 | 0.023,0.052,1.541,-1.511,-1.006,0.708 | 0.054,0.063,-1.067,0.571,1.169,1.980 | 0.032,0.053,-0.893,0.576,1.716,0.360 |
| **b88fa**640a14b620f100b6feb7d857fc0 | 0.044,0.075,1.591,-1.182,-1.935,-0.643 | 0.017,0.016,0.033,0.005,-1.851,0.003 | 0.060,0.057,-0.842,0.330,1.181,0.455 | 0.037,0.053,-0.589,0.391,0.302,1.788 |
| **b8b2c**2d224794705d8c88174a83b2c5b | 0.047,0.027,1.290,-1.232,-1.637,0.024 | 0.012,0.058,1.108,-0.857,1.921,0.161 | 0.057,0.011,-1.118,1.587,1.419,0.969 | 0.026,0.037,-0.566,0.164,1.977,1.968 |
| **b8c06**3a346c4651c02d95699a02a1352 | 0.075,0.014,1.667,-1.831,-1.901,-0.513 | 0.091,0.084,0.865,-0.488,-1.050,-0.836 | 0.031,0.097,-0.872,0.759,0.551,0.721 | 0.072,0.060,-0.924,0.663,1.946,-0.073 |
| **b901c**556b0274f56dffe321845248d6f | 0.064,0.029,0.533,-0.611,-0.056,-0.512 | 0.091,0.066,0.282,-0.091,-1.041,-0.319 | 0.079,0.055,-0.123,0.166,1.460,-0.743 | 0.078,0.045,-0.232,0.102,1.849,-0.662 |
| **b9254**161deb96f4a64fa3b8dbaaabbf8 | 0.070,0.020,0.055,0.038,-0.625,-0.598 | 0.085,0.097,1.021,-0.705,-1.412,-0.427 | 0.086,0.053,-0.933,0.481,1.785,1.315 | 0.055,0.042,0.000,-0.417,1.959,1.854 |
| **b95d3**311f03561ceddd283dcd024778e | 0.099,0.044,1.919,-1.409,-1.894,-0.933 | 0.030,0.088,1.635,-1.484,0.859,-1.233 | 0.095,0.056,-1.600,1.103,1.898,1.929 | 0.090,0.043,-0.077,-0.131,-0.045,1.983 |
| **b96ef**7c62516791839c9bd816e094690 | 0.088,0.024,1.578,-1.558,-1.324,-0.408 | 0.068,0.068,0.635,-0.319,-1.500,-0.308 | 0.030,0.073,-0.756,0.864,-1.268,1.110 | 0.050,0.033,-1.201,1.002,1.730,0.843 |
| **b96fc**0713bf2edaf274b079c408aad26 | 0.027,0.026,1.769,-1.817,-0.726,-1.160 | 0.066,0.060,0.221,-0.093,-1.146,-0.342 | 0.061,0.038,-0.902,0.410,0.007,0.794 | 0.048,0.084,-0.704,0.322,0.953,1.948 |
| **b96fc**488f1cb6be3e39e8d12ec95fc19 | 0.056,0.075,1.533,-1.476,-1.114,-0.058 | 0.053,0.032,0.606,-1.836,0.049,0.482 | 0.095,0.041,-1.589,0.728,0.068,1.346 | 0.073,0.073,-1.348,0.890,1.479,1.531 |
| **b97d2**98176c969a661a9a9e16b321b55 | 0.060,0.032,1.841,-1.785,-0.623,-1.101 | 0.030,0.041,0.504,-0.818,-1.347,-1.008 | 0.059,0.026,-1.430,0.633,1.795,0.118 | 0.086,0.033,-0.455,0.287,1.129,0.547 |
| **b98a1**cafd75a3ade0645a9f582ab6da0 | 0.044,0.062,1.611,-1.460,-1.168,-0.947 | 0.086,0.099,1.375,-1.115,-0.092,-1.590 | 0.030,0.096,-1.380,1.606,1.891,0.085 | 0.051,0.081,-1.132,0.950,-0.090,1.298 |
| **b9b82**fb27eb0da5a66fd96e45453ac45 | 0.078,0.064,0.635,-0.646,-0.941,0.209 | 0.050,0.059,0.997,-0.977,0.022,-1.301 | 0.038,0.095,-1.357,1.508,-0.147,0.678 | 0.024,0.093,-1.027,0.802,1.883,1.459 |
| **ba05c**24987ad3f1e69c310a4da8d2210 | 0.069,0.087,1.837,-1.698,0.229,-1.190 | 0.090,0.015,1.357,-1.699,0.814,1.318 | 0.075,0.079,-1.920,1.646,1.969,1.935 | 0.086,0.068,-0.662,0.325,1.837,0.344 |
| **ba330**c5aa1d387829557996e7cc97f00 | 0.055,0.025,1.767,-1.579,-1.792,-0.139 | 0.020,0.038,1.887,-1.259,-0.305,1.001 | 0.078,0.098,-1.482,1.807,1.365,1.268 | 0.082,0.081,-1.159,0.917,-0.558,0.968 |
| **ba5a3**65543dc3b228db6fb8a7a17239c | 0.083,0.015,1.174,-0.778,-0.941,-0.497 | 0.056,0.048,0.995,-1.489,-1.474,-0.350 | 0.070,0.083,-1.810,0.569,-0.302,1.849 | 0.068,0.080,-1.860,1.255,1.879,1.649 |
| **ba74f**f6d316345cd3faa0e5832f43a83 | 0.036,0.074,1.004,-0.493,-0.317,-0.957 | 0.059,0.049,0.486,-1.179,-1.050,-0.853 | 0.061,0.055,-1.541,0.284,-0.391,1.816 | 0.020,0.049,-0.470,0.411,1.251,0.290 |
| **ba7fb**c82f5511f047f6b8c7d230d8c6c | 0.092,0.065,1.638,-1.704,-0.641,-0.034 | 0.073,0.058,0.244,-0.232,-1.589,0.991 | 0.034,0.064,-0.413,0.229,0.669,0.091 | 0.057,0.063,-0.630,0.221,1.067,1.561 |
| **ba8fb**4b8d32db5bb14cfc9b4e072809e | 0.085,0.050,1.037,-0.320,-0.842,-0.012 | 0.033,0.079,0.619,-0.610,-1.915,-1.191 | 0.025,0.084,-1.498,0.365,1.215,0.525 | 0.017,0.073,-0.821,0.677,1.792,0.378 |
| **bac3d**eeef80bfcbc1790bcff6efc8864 | 0.091,0.072,1.126,-0.734,-1.415,-0.331 | 0.082,0.069,1.427,-1.651,-1.916,-1.897 | 0.036,0.058,-1.886,0.927,1.413,1.940 | 0.081,0.061,-0.462,0.046,1.810,1.455 |
| **bae8d**a7a22ea732ccfbb4abad51cc21a | 0.061,0.047,1.957,-1.460,-1.542,-0.544 | 0.046,0.053,0.612,-1.938,-0.342,-0.429 | 0.098,0.041,-1.779,0.416,1.898,-0.660 | 0.074,0.057,-1.267,1.014,1.910,1.051 |
| **bb2b2**ecf23558522c82274079f602f4b | 0.086,0.054,1.699,-1.998,-1.106,-0.139 | 0.075,0.085,0.877,-0.847,0.354,-1.220 | 0.058,0.068,-1.895,1.964,1.664,-1.838 | 0.092,0.073,-1.319,0.959,0.905,1.637 |
| **bb5c4**f8f1399f9688ae5a3a13004f8af | 0.058,0.061,0.721,-0.673,-0.674,-0.742 | 0.048,0.056,1.036,-0.660,0.306,-0.133 | 0.075,0.086,-1.185,1.849,-0.239,1.591 | 0.026,0.098,-1.099,0.730,1.909,0.735 |
| **bb66f**835bb0b63af60cfd3ba769a904c | 0.059,0.057,0.663,-0.472,-1.137,-0.917 | 0.066,0.075,0.698,-0.757,0.207,-0.340 | 0.038,0.073,-1.225,1.186,0.598,1.742 | 0.087,0.058,-1.390,1.206,0.661,0.822 |
| **bb6c5**1bdce4da4d30477641aa5a02031 | 0.046,0.049,1.760,-1.750,-1.646,-0.346 | 0.029,0.037,0.430,-0.645,-0.959,-0.196 | 0.089,0.056,-1.116,0.590,1.067,0.412 | 0.042,0.012,-0.031,-0.064,1.012,1.403 |
| **bb85f**3762019f39bebddd5226e02a3f9 | 0.016,0.066,1.906,-1.719,-1.793,-1.061 | 0.099,0.027,0.982,-1.689,0.071,-0.237 | 0.061,0.089,-1.108,0.339,1.770,1.123 | 0.029,0.041,-0.355,0.188,1.077,1.257 |
| **bb9ca**1f722853ddb4c7109f8a73b6e10 | 0.013,0.042,0.726,-0.725,1.594,-1.501 | 0.078,0.094,0.574,-0.467,-0.794,0.017 | 0.075,0.084,-1.935,1.757,0.672,1.928 | 0.011,0.077,-1.491,1.175,-1.780,1.922 |
| **bbb4f**b65fe642248949c18d37b3191d3 | 0.017,0.035,0.704,-0.773,-1.477,-0.240 | 0.025,0.021,0.315,-0.172,-1.943,-1.001 | 0.034,0.062,-0.681,0.839,1.354,0.796 | 0.091,0.071,-0.447,0.022,1.884,0.875 |
| **bbc16**d1014f87c9d894601d66a53910c | 0.075,0.069,0.378,-0.180,-1.755,0.097 | 0.021,0.017,0.593,-0.441,1.247,-1.078 | 0.098,0.025,-0.546,0.453,1.237,1.513 | 0.050,0.090,-1.016,0.692,0.176,0.997 |
| **bbcf4**94bf50343ab168c61f4748acc00 | 0.088,0.032,1.816,-1.392,-1.687,-0.917 | 0.065,0.096,1.033,-1.113,-0.177,-1.868 | 0.094,0.058,-1.822,0.930,0.042,1.949 | 0.015,0.063,-0.692,0.562,1.795,0.876 |
| **bc2d1**48629508e43b3c8147099708771 | 0.081,0.088,1.706,-1.622,-0.946,-0.187 | 0.016,0.090,1.645,-0.731,-0.115,-1.830 | 0.056,0.048,-1.246,1.879,1.761,0.602 | 0.094,0.070,-0.559,-0.213,1.657,1.645 |
| **bc542**ef4de0d27ec0090694ac926c809 | 0.078,0.060,0.620,-0.273,-1.621,0.041 | 0.026,0.044,0.149,-0.317,-1.216,0.011 | 0.067,0.070,-1.339,0.260,0.071,1.913 | 0.042,0.026,-0.458,0.290,1.728,1.280 |
| **bc597**c8a26c51553ab70ac692ab6b8ef | 0.048,0.035,0.470,-0.385,-0.860,-0.148 | 0.012,0.081,0.396,-0.470,-0.592,-1.127 | 0.037,0.057,-1.396,0.656,1.668,1.703 | 0.064,0.086,-1.446,1.001,1.755,1.845 |
| **bcad6**723b78dc96eb6c2a9c5311269f0 | 0.045,0.040,1.674,-1.856,-1.843,0.152 | 0.039,0.025,0.833,-0.518,-1.124,-0.367 | 0.039,0.057,-0.979,1.361,-0.318,0.146 | 0.055,0.051,-1.078,1.090,1.668,-1.546 |
| **bcb5b**fdfdcd7a7243bfe374cb017b1d4 | 0.071,0.020,1.811,-1.792,-1.518,-0.235 | 0.047,0.097,0.736,-0.657,-1.874,-0.216 | 0.072,0.050,-1.708,1.402,1.146,1.486 | 0.042,0.061,-0.578,0.260,1.694,1.802 |
| **bcc72**362eb2fdb7e231db74ceb27c2b1 | 0.093,0.030,1.849,-1.365,-1.200,-0.002 | 0.013,0.064,0.617,-1.508,-1.066,-0.194 | 0.100,0.016,-0.917,0.010,1.031,1.874 | 0.040,0.087,-0.163,-0.181,1.684,0.852 |
| **bcfc2**2f46c66bec4e2e18932e4027c19 | 0.086,0.052,1.537,-1.132,-1.654,-0.388 | 0.059,0.045,1.468,-1.467,-0.338,-1.462 | 0.062,0.053,-1.528,1.162,1.312,-0.621 | 0.085,0.091,-0.099,-0.505,1.919,0.046 |
| **bd20a**bca308d4e7264122ace09edb692 | 0.095,0.069,1.709,-1.630,-1.695,0.313 | 0.028,0.030,0.906,-1.358,-0.856,-1.857 | 0.085,0.082,-0.821,0.191,1.784,0.395 | 0.087,0.067,-1.080,0.636,1.876,1.270 |
| **bd409**f8fdb6c21ce774b4ad4004a1b86 | 0.041,0.049,0.687,-0.555,-1.688,-0.331 | 0.014,0.037,0.126,-0.378,0.039,0.244 | 0.072,0.058,-1.732,0.785,-0.528,1.571 | 0.039,0.064,-0.712,0.530,0.381,1.172 |
| **bd40d**c4109719212471c0b74cfba7123 | 0.072,0.056,1.719,-1.522,-0.785,-0.954 | 0.096,0.068,1.339,-1.228,-0.756,-1.532 | 0.094,0.066,-1.272,0.919,1.140,-0.040 | 0.051,0.088,-0.808,0.492,1.513,0.815 |
| **bd467**90781f6121a23e899c79fa245ac | 0.077,0.020,1.589,-1.837,-1.137,0.008 | 0.094,0.053,0.703,-0.363,-1.538,0.821 | 0.080,0.019,-1.223,0.848,1.945,1.503 | 0.097,0.043,-0.535,0.248,1.116,-0.098 |
| **bd4db**d926025e38ef44a017147058b93 | 0.040,0.058,1.226,-1.277,-0.322,-0.976 | 0.046,0.035,0.853,-0.638,-0.678,-1.826 | 0.099,0.097,-1.419,1.426,1.163,1.117 | 0.091,0.026,-0.517,0.188,1.831,0.695 |
| **bd539**a8181e45d28148f84f69f184f9b | 0.069,0.030,1.985,-1.945,-1.902,0.050 | 0.019,0.065,0.489,-0.380,0.586,-1.702 | 0.095,0.071,-0.899,0.691,0.561,0.829 | 0.030,0.063,-0.205,-0.062,1.777,1.646 |
| **bd6b3**9559c6af594aee41462291e55c4 | 0.093,0.029,1.726,-1.684,-1.892,-0.087 | 0.039,0.014,0.682,-0.670,-1.034,-0.668 | 0.099,0.097,-0.648,0.403,1.685,-0.369 | 0.026,0.095,-0.621,0.266,1.318,1.481 |

| | | | | |
|---|---|---|---|---|
| **bd8214**d9b40977af07fcd406a7a5294f | 0.095,0.016,1.534,-1.313,-1.722,0.271 | 0.044,0.078,0.774,-0.868,-0.732,-0.402 | 0.098,0.060,-1.548,1.104,-0.438,1.514 | 0.054,0.065,-0.495,-0.008,1.857,1.500 |
| **bdab1**b5504c22b4dd15784951a754bac | 0.094,0.026,0.639,-0.319,-1.615,-0.348 | 0.055,0.027,0.344,-0.250,-0.942,-1.483 | 0.082,0.060,-1.153,0.920,0.883,0.377 | 0.020,0.092,-1.222,0.925,1.337,1.388 |
| **bdb45**cd94057ddf69714ac637d15bc70 | 0.014,0.086,1.960,-1.675,0.308,-1.885 | 0.040,0.049,1.215,-1.375,-0.833,-1.202 | 0.015,0.088,-1.998,1.939,1.958,-0.244 | 0.010,0.059,-1.505,1.261,1.240,1.994 |
| **bdb90**b7e1f82f3f4afffc371c5d8fcd0 | 0.046,0.019,1.535,-1.871,-1.385,0.692 | 0.075,0.018,0.536,-0.546,-1.212,1.435 | 0.073,0.049,-1.618,0.662,1.824,1.721 | 0.046,0.023,-0.448,0.284,0.633,1.304 |
| **be0ec**8ce0fce6747cf80f476e04f5955 | 0.039,0.047,1.868,-1.820,-0.672,-0.569 | 0.080,0.046,0.459,-0.415,-1.466,0.029 | 0.033,0.076,-1.739,0.888,1.755,1.683 | 0.071,0.098,-0.175,-0.251,1.937,0.691 |
| **be330**e188acc433e4075e911594fb6f5 | 0.046,0.017,1.076,-0.831,-1.846,-0.406 | 0.084,0.065,0.179,-0.251,-0.892,0.909 | 0.013,0.027,-0.869,0.383,0.852,0.831 | 0.027,0.031,-1.098,1.072,1.317,-0.427 |
| **be4eb**c117b3a6578d94f202b1f7b06d6 | 0.095,0.027,1.822,-1.674,-1.612,-0.917 | 0.012,0.048,1.504,-1.068,-0.581,-1.439 | 0.092,0.096,-1.870,1.888,0.615,1.960 | 0.079,0.079,-0.597,-0.100,1.820,1.499 |
| **be4f5**e46078daff4f57013a4c4fbf445 | 0.096,0.057,0.619,-0.483,-1.097,0.046 | 0.041,0.075,0.527,-0.833,-0.737,1.806 | 0.078,0.043,-1.196,0.794,1.795,1.883 | 0.024,0.064,-0.954,0.735,0.810,0.897 |
| **be775**18054031dc82b6f4b5715a72e15 | 0.083,0.021,1.404,-1.254,-1.221,-0.015 | 0.081,0.058,1.872,-1.867,0.324,-1.464 | 0.038,0.042,-1.774,1.730,0.040,1.452 | 0.029,0.041,-1.016,0.679,1.970,1.850 |
| **be7e4**137f9c707f7500cd7dac3b4da93 | 0.058,0.086,1.267,-0.903,-1.419,-1.299 | 0.021,0.031,1.213,-0.727,1.868,-0.313 | 0.094,0.091,-1.235,1.738,1.899,0.581 | 0.072,0.039,-0.534,0.201,0.155,1.007 |
| **be9a2**16bd4a783b9ce9fbad1701b5176 | 0.092,0.021,1.622,-1.395,-1.850,1.146 | 0.071,0.017,0.359,-0.122,-1.940,-1.006 | 0.097,0.038,-0.436,-0.140,1.801,0.701 | 0.037,0.083,-0.378,-0.029,1.874,1.668 |
| **bea35**186719525a975de3e0d21cf398b | 0.030,0.033,1.849,-1.722,-1.928,-0.878 | 0.042,0.074,0.608,-1.950,-1.949,0.190 | 0.041,0.066,-0.858,0.081,0.609,1.502 | 0.078,0.043,-0.388,0.012,1.903,1.453 |
| **bea54**87c2c806d65e42c0e2967c57626 | 0.088,0.062,1.191,-1.172,-1.389,0.129 | 0.018,0.053,0.867,-0.813,-0.980,0.275 | 0.090,0.026,-1.826,1.864,1.781,1.667 | 0.085,0.074,-1.280,0.781,0.370,1.732 |
| **beaa9**2c27d5586b139e77af62ff2f250 | 0.039,0.050,1.534,-1.762,-1.283,-0.000 | 0.030,0.015,0.581,-0.448,-1.397,0.473 | 0.026,0.079,-0.853,1.688,0.828,-0.846 | 0.087,0.036,-0.749,0.583,0.846,0.553 |
| **beb00**1af3af4cc32e6ce4e5568f4922f | 0.056,0.027,1.069,-0.891,-1.056,-0.235 | 0.063,0.059,0.504,-1.025,-0.240,-0.033 | 0.076,0.040,-0.700,0.190,0.756,1.084 | 0.026,0.058,-0.425,0.278,1.519,0.755 |
| **bec6a**5b8627cd740149ac331b8653e5f | 0.052,0.022,1.844,-1.655,-1.226,-0.443 | 0.073,0.071,1.811,-1.346,-0.509,-0.142 | 0.051,0.041,-1.967,1.882,1.893,1.339 | 0.090,0.038,-0.039,-0.998,1.906,1.073 |
| **bee0e**1b0a7abd3887e30471a0958986e | 0.084,0.043,1.568,-1.191,-1.175,0.289 | 0.039,0.042,0.785,-1.244,-1.497,0.340 | 0.056,0.057,-1.333,0.631,1.909,0.203 | 0.029,0.095,-1.526,1.013,0.918,1.841 |
| **bf281**a9a8d6b28e6d89197cb0942e45d | 0.016,0.016,1.821,-1.568,-1.528,-0.998 | 0.060,0.067,0.920,-1.519,-1.816,-1.871 | 0.066,0.056,-1.991,0.433,1.916,-0.812 | 0.063,0.053,-0.003,-0.553,1.984,1.964 |
| **bf811**fbc59faee6e6fc43ebba6889bd5 | 0.046,0.044,1.967,-1.738,-1.631,-0.161 | 0.072,0.096,1.018,-1.914,-0.831,-1.004 | 0.084,0.045,-1.740,0.646,0.294,0.926 | 0.099,0.029,-0.209,-0.285,1.819,0.405 |
| **bf8b3**55ab9e1d18190d67166814fa578 | 0.093,0.014,1.881,-1.930,-1.482,0.127 | 0.098,0.051,1.264,-1.903,0.007,-0.268 | 0.088,0.096,-1.028,0.657,1.483,-0.407 | 0.083,0.040,-1.290,0.982,1.206,1.299 |
| **bfab3**fd0bf6de3ae6948fbc2e5702ee1 | 0.058,0.020,0.973,-1.078,-0.437,-0.566 | 0.014,0.086,0.468,-0.721,0.913,-0.171 | 0.064,0.036,-1.658,1.082,-1.077,1.882 | 0.050,0.032,-0.924,0.762,1.787,-0.848 |
| **bfb1a**ce73d4550c3d45f03c5aa3537d4 | 0.096,0.021,0.259,-0.235,0.029,-1.401 | 0.014,0.071,1.277,-0.639,0.322,-0.725 | 0.072,0.082,-0.620,1.453,1.764,-0.522 | 0.096,0.066,-0.273,-0.155,1.520,1.005 |
| **bfbb0**56ecb763411f8f6459042219ea8 | 0.034,0.062,1.466,-1.654,0.024,-1.305 | 0.053,0.050,0.994,-0.947,1.256,-1.012 | 0.098,0.017,-1.328,1.845,-0.195,1.876 | 0.082,0.061,-1.329,0.970,1.250,0.779 |
| **bfbf0**4bd4657303ef5d56c5314235566 | 0.065,0.022,1.243,-1.193,-0.185,-0.978 | 0.074,0.072,0.912,-1.279,-1.200,-1.737 | 0.096,0.095,-1.250,0.345,-0.139,0.488 | 0.082,0.084,-1.481,0.875,1.833,1.129 |
| **bfd8a**4f002f5d76cac95b17afa1c0767 | 0.011,0.087,1.693,-1.913,0.009,-1.217 | 0.061,0.088,0.707,-0.766,-0.585,0.034 | 0.045,0.046,-0.937,0.868,1.022,1.325 | 0.042,0.036,-1.216,1.056,1.517,1.262 |
| **bfe0d**5f131cd5888b7c00593ed6e2d1e | 0.057,0.060,0.998,-1.017,-1.029,0.688 | 0.036,0.081,0.543,-1.290,-0.140,0.765 | 0.060,0.076,-0.610,0.189,1.960,-0.295 | 0.086,0.039,-1.756,1.419,1.730,0.721 |
| **bffb7**082b23d9a48addae3db39d55091 | 0.067,0.061,1.923,-1.766,-0.681,-0.983 | 0.085,0.050,0.467,-1.103,-0.826,-0.047 | 0.039,0.023,-0.996,0.332,1.131,-0.662 | 0.075,0.061,-0.818,0.549,1.533,1.064 |
| **c0112**c796b90b1eb299e73c758cf8b6d | 0.043,0.046,1.974,-1.424,-1.742,-1.040 | 0.017,0.024,1.397,-1.604,-0.085,-1.449 | 0.099,0.047,-1.896,1.169,1.700,0.046 | 0.067,0.014,-0.166,-0.447,1.851,1.721 |
| **c023b**772f406005284b1b80af086f6c4 | 0.045,0.088,1.585,-1.774,-1.547,0.056 | 0.041,0.057,0.622,-0.659,1.688,-1.436 | 0.064,0.070,-0.578,0.398,1.163,0.644 | 0.022,0.045,-0.698,0.498,1.867,1.141 |
| **c02f4**6ed72af33ae2d3f183a893b8189 | 0.054,0.022,1.871,-1.758,-0.373,-1.673 | 0.071,0.047,1.593,-1.135,-0.939,-1.094 | 0.032,0.081,-1.680,1.628,-0.454,1.110 | 0.057,0.066,-1.049,0.543,1.843,1.049 |
| **c052f**03a2e986588a850cf5d5e29560c | 0.076,0.047,1.921,-1.777,-1.394,-1.098 | 0.060,0.056,0.417,-0.262,-0.644,-0.454 | 0.051,0.060,-1.210,1.091,1.180,0.774 | 0.067,0.057,-1.329,1.057,1.445,0.907 |
| **c0548**b0255c9feb3f378840b2dbdc654 | 0.083,0.019,1.794,-1.793,-1.827,1.093 | 0.094,0.014,0.758,-1.218,1.634,-1.112 | 0.078,0.048,-1.174,0.841,1.924,1.689 | 0.040,0.075,-0.192,-0.039,1.897,0.485 |
| **c0825**971dfd228851158c962ece4f258 | 0.093,0.038,1.084,-0.928,-1.827,0.963 | 0.057,0.088,0.546,-0.571,-1.472,-0.189 | 0.053,0.063,-0.423,0.210,1.041,0.018 | 0.065,0.090,-0.642,0.280,0.595,1.563 |
| **c08de**5e231f63220a9d88ed657eb8489 | 0.069,0.057,1.682,-0.894,-1.375,-1.118 | 0.071,0.082,1.047,-1.608,-1.735,-0.884 | 0.042,0.042,-1.548,0.560,-0.263,1.143 | 0.025,0.084,-1.005,0.681,0.694,1.404 |
| **c0aac**f416420598f3b6efe68c3a0f796 | 0.013,0.079,1.411,-1.670,0.797,-1.123 | 0.019,0.026,0.160,-0.190,0.368,-0.452 | 0.068,0.061,-1.300,0.779,1.855,1.859 | 0.038,0.093,-1.881,1.558,0.097,1.822 |
| **c0d47**d8b8ea5476430f023a25967623d | 0.091,0.041,1.087,-0.863,-1.826,0.593 | 0.039,0.055,0.226,-0.336,-0.237,-0.187 | 0.055,0.017,-1.869,1.238,1.299,1.922 | 0.098,0.077,-0.729,0.462,1.691,0.153 |
| **c0e59**f9095ee554a8ec4c9fdd100ae1d | 0.077,0.044,1.660,-1.769,-1.181,0.221 | 0.042,0.030,0.561,-0.331,-0.786,-1.117 | 0.058,0.042,-0.746,0.769,0.958,1.054 | 0.055,0.087,-1.433,1.137,1.913,0.357 |
| **c0e7f**98dec884d3199e6f6ddcb1c3595 | 0.015,0.080,1.128,-1.015,0.260,-1.689 | 0.091,0.018,1.125,-0.500,-1.200,-1.687 | 0.077,0.066,-0.678,0.717,1.300,0.723 | 0.075,0.081,-1.012,0.334,1.502,1.540 |
| **c0e97**a12bb87fceb164e6341398481af | 0.011,0.097,1.671,-1.486,0.833,-1.839 | 0.058,0.080,1.546,-1.556,-0.524,-0.034 | 0.082,0.055,-0.613,0.362,1.748,0.732 | 0.089,0.028,-0.778,0.388,1.738,1.151 |
| **c1368**9bb3febd8e346022e0654dbbab5 | 0.043,0.034,1.508,-1.400,-1.014,-0.453 | 0.067,0.074,0.892,-1.395,-0.842,0.064 | 0.066,0.034,-1.284,0.257,1.730,1.252 | 0.045,0.037,-0.527,0.223,1.658,1.004 |
| **c1411**5e119707521ec475a9fcf60e4ab | 0.048,0.014,1.166,-1.288,-1.342,-0.389 | 0.020,0.082,0.654,-0.461,-1.780,-0.871 | 0.066,0.055,-1.347,1.244,0.012,1.176 | 0.082,0.094,-1.206,0.562,1.689,1.523 |
| **c14f1**2ada4f5774225178c559b852daa | 0.077,0.017,1.193,-1.088,-0.990,-0.556 | 0.072,0.035,0.836,-0.926,-0.672,-1.309 | 0.022,0.039,-1.231,0.756,-0.156,1.288 | 0.048,0.037,-0.820,0.486,1.839,1.148 |
| **c1520**ff7d7742a91d9751c11756f1741 | 0.093,0.063,0.743,-0.174,-0.958,-0.624 | 0.014,0.047,1.009,-1.831,-0.060,1.510 | 0.060,0.080,-1.490,0.475,1.929,1.103 | 0.060,0.039,-0.742,0.356,1.224,0.809 |
| **c168c**4a75c020e41c2d89e88cf775347 | 0.099,0.024,1.742,-1.826,-0.777,-1.018 | 0.020,0.097,1.444,-0.881,-1.839,-0.912 | 0.089,0.029,-1.844,1.772,1.564,1.935 | 0.065,0.099,-1.155,0.387,1.945,1.523 |
| **c16ac**b01a49e882b101098facd55a19a | 0.047,0.025,1.421,-1.112,-1.263,-0.931 | 0.054,0.080,0.645,-0.403,-1.899,-0.519 | 0.028,0.029,-0.833,0.308,-0.180,0.986 | 0.094,0.079,-0.747,0.531,1.267,-0.659 |
| **c1765**57178fa9473b61025c1b4651bf6 | 0.017,0.027,1.394,-1.581,-1.002,-0.451 | 0.028,0.078,0.788,-0.844,0.807,-0.167 | 0.010,0.061,-1.356,1.741,0.214,0.704 | 0.017,0.077,-1.226,1.146,1.352,0.550 |
| **c17ec**2744acf589dcea54726d96a3474 | 0.097,0.043,1.512,-1.865,-0.300,-0.260 | 0.063,0.075,0.977,-0.650,-1.505,-1.149 | 0.055,0.030,-0.967,0.789,-0.218,1.747 | 0.068,0.042,-0.529,0.031,1.781,1.787 |
| **c19b3**7da5f9bac77646ef63de4f06b07 | 0.052,0.031,1.891,-1.925,-1.219,-0.642 | 0.070,0.080,1.383,-1.761,-0.950,-0.738 | 0.013,0.056,-1.829,1.356,0.271,1.243 | 0.060,0.086,-1.123,0.794,0.307,1.338 |
| **c1c97**373a5f6130958a3ee641a505f39 | 0.044,0.010,1.814,-1.848,-1.431,-0.664 | 0.098,0.089,0.849,-1.146,-0.300,-1.164 | 0.097,0.045,-1.233,0.395,0.972,1.674 | 0.098,0.062,-0.695,0.373,1.759,0.299 |
| **c1f27**7eb7c8a13e7186c5438ca5d3936 | 0.013,0.058,1.568,-1.815,-1.241,-0.728 | 0.042,0.014,0.529,-0.683,0.406,-1.124 | 0.063,0.024,-1.118,0.908,1.804,1.093 | 0.098,0.096,-1.919,1.332,1.670,1.839 |
| **c2074**d33ca9eb4d9b190630721b729f2 | 0.024,0.057,1.372,-1.626,0.682,-0.886 | 0.027,0.046,1.402,-0.941,0.339,-1.028 | 0.090,0.081,-1.195,0.987,1.645,1.726 | 0.081,0.067,-0.499,-0.012,0.535,1.636 |
| **c26dd**7a13ee80a655fb17afdced9baa6 | 0.040,0.019,1.913,-1.692,-1.109,-0.428 | 0.045,0.066,0.561,-0.728,-0.721,-1.898 | 0.070,0.088,-1.091,0.239,0.724,0.300 | 0.076,0.056,-0.303,-0.228,1.945,1.409 |
| **c2811**219bfae7af6bd91c5a162d43a66 | 0.047,0.039,1.001,-0.801,-1.907,0.346 | 0.014,0.072,0.393,-0.729,-1.490,0.528 | 0.052,0.048,-1.501,0.999,1.118,1.637 | 0.044,0.031,-0.345,0.195,1.125,0.849 |
| **c28df**1e614846f2897c8c7ad36caf723 | 0.057,0.014,1.674,-1.745,-0.760,-0.484 | 0.087,0.015,0.753,-0.537,-1.528,1.676 | 0.085,0.019,-0.621,0.304,1.664,1.469 | 0.031,0.090,-0.245,-0.056,0.950,1.371 |
| **c2b8f**d5676a17bbcc722a04728273eca | 0.037,0.048,1.793,-1.915,0.464,-1.214 | 0.049,0.021,0.806,-1.907,-1.767,-0.079 | 0.047,0.018,-1.415,0.411,1.900,0.754 | 0.085,0.096,-0.456,0.240,1.316,0.248 |
| **c2c1e**f12e37b663b4853f1486d89221e | 0.062,0.057,1.905,-1.474,-1.010,-0.987 | 0.091,0.017,1.289,-1.835,-0.341,-1.155 | 0.054,0.096,-1.872,0.907,1.423,1.399 | 0.075,0.052,-0.317,-0.004,1.835,-0.182 |
| **c2cf4**4ed2e945a88dd07df577086d095 | 0.014,0.078,1.930,-1.315,-1.843,-1.726 | 0.086,0.012,1.166,-1.865,-1.931,0.751 | 0.096,0.018,-1.478,0.261,1.780,1.991 | 0.042,0.041,-1.403,1.286,1.781,-0.191 |
| **c2da5**3a9b0a3b0d3d679b30b697b1cf9 | 0.091,0.019,1.058,-0.615,-1.841,0.867 | 0.053,0.094,0.609,-0.792,-0.161,0.148 | 0.038,0.034,-1.418,1.117,-0.734,1.476 | 0.033,0.059,-1.074,0.870,1.187,0.320 |
| **c2dd6**f1f40c7ff4ac30f3d58a5bca460 | 0.014,0.046,1.704,-1.771,-1.404,-1.094 | 0.025,0.095,1.338,-0.944,-1.567,-0.081 | 0.060,0.064,-1.843,1.929,1.496,0.285 | 0.090,0.090,-1.553,1.390,1.195,-0.565 |
| **c2fc0**e31cd66369dc3e5a780d0ab07ab | 0.087,0.099,0.720,-0.559,-0.939,-0.451 | 0.044,0.035,0.521,-0.552,-0.878,-0.870 | 0.033,0.020,-1.815,1.847,1.094,-0.071 | 0.058,0.030,-0.739,0.571,1.819,1.317 |
| **c3446**fd789069475bf715540fd9c9583 | 0.014,0.070,1.725,-1.943,0.183,-1.316 | 0.066,0.079,0.532,-0.598,0.309,-1.011 | 0.012,0.042,-1.693,1.316,0.927,1.539 | 0.041,0.061,-1.251,1.019,1.536,1.557 |
| **c34a8**ec1d592a57279a6cd3c145e5e59 | 0.084,0.051,0.576,-0.427,-1.771,0.772 | 0.032,0.089,0.795,-0.751,-1.474,-0.187 | 0.072,0.027,-0.497,0.319,1.290,0.669 | 0.022,0.031,-0.629,0.487,1.918,1.255 |
| **c34f4**2b742a95f4c05f2000cb9509c60 | 0.083,0.019,1.806,-1.693,-1.840,-0.086 | 0.042,0.034,1.411,-1.643,-0.427,-0.367 | 0.036,0.058,-1.500,1.806,1.087,-1.563 | 0.049,0.068,-0.739,0.562,0.172,0.887 |
| **c3610**0f56052eb8cd4845b1de37cd24f | 0.022,0.059,0.807,-0.777,-0.271,-0.506 | 0.020,0.043,0.429,-0.397,-0.936,0.040 | 0.055,0.064,-1.322,1.379,0.305,0.815 | 0.039,0.036,-1.687,1.473,1.677,0.829 |
| **c367f**b0975dd3377059229099c6ee43e | 0.085,0.027,1.579,-1.860,-0.947,0.734 | 0.016,0.054,1.200,-1.282,1.064,0.066 | 0.067,0.067,-1.815,1.958,1.195,0.393 | 0.097,0.042,-1.534,1.323,1.728,0.186 |
| **c3785**06b4f9235b5f20bd7a9bf43eb51 | 0.039,0.078,1.382,-1.345,-0.442,-0.003 | 0.074,0.030,1.889,-0.665,-0.956,-0.584 | 0.061,0.062,-1.205,1.901,1.915,1.458 | 0.048,0.083,-0.946,0.436,0.573,0.980 |
| **c3b3f**6b8881b3d7a9f9b37352f0508c8 | 0.063,0.043,1.579,-1.708,-0.600,-0.720 | 0.015,0.072,0.405,-0.775,0.031,-0.285 | 0.021,0.073,-1.694,0.851,0.121,1.933 | 0.086,0.054,-0.540,0.284,1.918,0.270 |
| **c3bd4**a0e8d3b90591b8e0f448e191b02 | 0.025,0.051,0.381,-0.352,-0.916,0.129 | 0.055,0.080,0.453,-0.347,1.714,-1.879 | 0.045,0.037,-1.507,1.934,1.052,0.447 | 0.060,0.056,-1.111,0.919,0.444,0.737 |

| | | | | |
|---|---|---|---|---|
| **c3c29**f025d6c53761ced7ec0a71a6352 | 0.026,0.085,1.756,-1.843,0.266,-0.660 | 0.087,0.028,0.562,-1.510,1.131,-1.232 | 0.089,0.094,-1.556,0.188,1.736,1.752 | 0.084,0.063,-1.480,1.084,1.601,1.057 |
| **c3f3a**58c14ddd3804009d80f38b79476 | 0.089,0.053,1.463,-0.643,-1.932,-0.438 | 0.092,0.097,1.947,-1.597,-1.923,-0.011 | 0.035,0.071,-1.939,1.136,1.143,1.424 | 0.072,0.063,-0.877,0.397,1.909,0.240 |
| **c3fad**58627f20ee9c85dcd67853ff77d | 0.073,0.025,1.487,-1.289,-1.938,-0.249 | 0.021,0.057,0.722,-1.292,0.248,0.178 | 0.073,0.082,-1.187,0.868,-0.043,0.800 | 0.023,0.060,-1.663,1.510,-0.534,1.332 |
| **c3fb9**14d0ea7caca58d4c254bfd919e3 | 0.064,0.036,0.945,-0.755,-1.077,-0.241 | 0.083,0.086,1.929,-1.584,-1.304,-0.333 | 0.076,0.090,-1.590,1.722,0.989,-1.177 | 0.057,0.034,-1.013,0.834,1.435,0.335 |
| **c42f8**178eba2efe5c1b8edbf5b6a6f72 | 0.095,0.023,1.083,-0.679,-1.939,0.561 | 0.037,0.072,1.162,-1.232,0.269,-0.331 | 0.068,0.082,-0.976,0.945,0.187,0.402 | 0.078,0.044,-0.150,-0.189,0.857,1.648 |
| **c4338**9bf83ef0cc46eefff4766febf7b | 0.048,0.018,1.822,-1.865,-1.692,0.012 | 0.091,0.023,0.385,-0.925,-0.004,0.539 | 0.012,0.025,-0.792,0.408,1.548,1.403 | 0.058,0.049,-1.443,1.226,1.982,0.835 |
| **c4454**1c21fbd57e1b71ea180eaa28122 | 0.046,0.013,1.640,-1.487,-1.905,-0.342 | 0.073,0.083,0.620,-1.241,0.496,-1.823 | 0.095,0.033,-1.592,0.349,1.670,0.225 | 0.017,0.041,-1.234,1.104,1.885,1.163 |
| **c4a79**d432a4d60a6e1e236d76615e96a | 0.049,0.042,1.777,-1.809,0.268,-1.847 | 0.078,0.081,0.549,-0.193,-0.974,-1.081 | 0.073,0.058,-0.528,0.265,0.325,1.746 | 0.029,0.044,-1.467,1.207,1.500,1.756 |
| **c4a90**a9c0c38e8d8450ed5b71c7ef859 | 0.043,0.083,1.384,-1.058,-1.949,-0.521 | 0.027,0.087,1.530,-0.864,0.717,-1.509 | 0.093,0.091,-1.752,1.971,1.661,0.766 | 0.025,0.075,-1.138,0.551,0.237,1.798 |
| **c4ca5**5ca5119e51d0509d401438a4a03 | 0.093,0.017,1.849,-1.126,-1.860,-0.693 | 0.035,0.086,0.439,-0.228,0.298,-1.957 | 0.061,0.043,-0.415,-0.002,1.256,0.111 | 0.085,0.057,-0.721,0.259,1.726,1.044 |
| **c4dc1**ff29a9ac062a5a2b79ed77fce6b | 0.091,0.056,1.740,-1.660,-1.647,-0.788 | 0.075,0.055,1.317,-1.489,-0.695,0.081 | 0.074,0.039,-1.938,1.946,1.814,1.380 | 0.055,0.075,-1.467,1.173,1.843,1.324 |
| **c4e57**a729e9023099fa81540a92d106e | 0.012,0.078,1.539,-1.239,-0.409,-1.903 | 0.063,0.076,0.965,-0.321,0.081,-1.462 | 0.084,0.046,-1.158,1.384,0.721,1.320 | 0.073,0.075,-0.126,-0.884,1.057,1.513 |
| **c4ea8**ccca5e1a6f067b2d33548450863 | 0.061,0.096,0.538,-0.472,-1.139,0.026 | 0.039,0.083,0.670,-0.477,-1.166,-1.403 | 0.093,0.043,-1.789,1.816,1.426,1.132 | 0.088,0.033,-0.594,0.240,1.946,1.097 |
| **c4f0e**610938ac3987c2d648f352d26f7 | 0.011,0.050,1.734,-1.749,-1.126,-1.235 | 0.041,0.055,1.966,-1.297,-1.005,1.109 | 0.083,0.062,-1.229,1.967,-0.348,1.212 | 0.053,0.061,-1.382,0.671,0.425,1.915 |
| **c4f6a**43186dd98aac9440a344ac52d34 | 0.077,0.040,0.544,-0.399,-1.189,0.097 | 0.068,0.040,0.357,-0.208,-0.484,-1.260 | 0.028,0.052,-0.718,0.684,1.735,0.156 | 0.091,0.050,-1.674,1.344,1.798,0.721 |
| **c4fc4**92353224792e0f5e21be73ba821 | 0.081,0.025,1.024,-1.083,-0.772,-0.909 | 0.058,0.086,0.565,-0.656,0.044,-0.942 | 0.059,0.086,-1.664,1.029,0.231,0.626 | 0.082,0.067,-0.537,0.117,1.203,1.724 |
| **c50a8**e5852d5881d60dddeaf44788e9b | 0.099,0.083,1.579,-1.118,-1.927,-0.350 | 0.089,0.040,1.928,-1.016,-1.197,-1.057 | 0.067,0.032,-0.783,1.059,1.678,0.583 | 0.079,0.078,-0.101,-0.753,1.471,1.460 |
| **c50da**2a52c1370884a7e92264b266473 | 0.082,0.026,0.651,-0.450,-1.047,0.385 | 0.024,0.018,1.406,-0.961,-0.553,-1.140 | 0.042,0.095,-1.711,1.108,1.072,1.858 | 0.056,0.063,-1.269,0.753,1.085,0.705 |
| **c5103**4ce155898351e24643d718dd3a4 | 0.047,0.062,1.749,-1.567,-1.748,0.469 | 0.015,0.097,0.511,-0.240,-1.132,-1.511 | 0.072,0.062,-1.899,0.448,-0.206,1.187 | 0.025,0.041,-0.130,0.068,1.927,-0.191 |
| **c5198**f596da30f5c0c2b1cae9467e0ea | 0.057,0.027,0.454,0.102,-1.594,-0.499 | 0.047,0.036,0.358,-0.097,-1.393,-1.075 | 0.030,0.074,-1.983,1.414,0.346,0.471 | 0.055,0.026,-1.390,0.963,1.686,1.470 |
| **c522e**d4040a7a2b5aa5888b4265be823 | 0.099,0.044,1.886,-1.663,-1.919,-0.824 | 0.049,0.047,0.632,-0.594,-0.876,0.113 | 0.054,0.019,-1.019,0.908,1.803,0.495 | 0.068,0.071,-1.154,0.743,1.065,1.901 |
| **c5744**0efe1c7890ae34f1d358a367d2c | 0.084,0.043,0.975,-0.937,-0.077,-1.024 | 0.076,0.014,0.470,-0.813,-1.625,1.112 | 0.015,0.088,-1.174,0.348,1.295,1.027 | 0.027,0.080,-1.356,1.105,-0.984,1.738 |
| **c585a**ad7a694d308bf10770a12fe8cd5 | 0.013,0.047,1.765,-1.751,-1.262,-0.665 | 0.096,0.079,0.727,-0.923,0.180,-0.485 | 0.024,0.028,-0.740,0.344,1.675,1.973 | 0.093,0.063,-1.571,1.000,0.714,1.784 |
| **c5a86**6bd1a989982f14a108dbd5499dd | 0.046,0.088,1.592,-1.775,-1.783,-0.439 | 0.031,0.033,0.536,-0.937,1.205,-0.710 | 0.057,0.013,-0.585,0.485,1.219,0.751 | 0.096,0.086,-0.281,0.002,0.900,1.380 |
| **c5b08**6ad7d381fa66ea3c192513cde43 | 0.061,0.036,1.191,-0.576,-1.911,-0.125 | 0.067,0.061,0.579,-1.136,-0.701,-1.864 | 0.039,0.050,-1.708,0.455,0.344,1.110 | 0.052,0.041,-0.214,0.103,1.752,-0.288 |
| **c5dd0**69c739f4f741f79c3df85ade163 | 0.036,0.075,0.514,-0.498,-0.162,-0.346 | 0.025,0.074,0.521,-0.425,1.430,-1.030 | 0.032,0.061,-0.941,1.210,1.269,0.415 | 0.057,0.025,-0.599,0.332,1.501,1.100 |
| **c5e42**7038184c5b0952d2ff14b15a6c0 | 0.071,0.042,1.919,-1.744,-1.594,0.045 | 0.024,0.097,1.265,-1.832,0.425,-1.701 | 0.096,0.084,-1.914,1.146,-0.811,0.616 | 0.075,0.083,-1.450,1.135,0.098,1.444 |
| **c6012**2723eea3017fca5d46b890c0246 | 0.048,0.080,0.796,-0.701,-1.088,-0.385 | 0.021,0.067,0.612,-0.778,1.618,-0.431 | 0.047,0.083,-1.235,1.026,-0.347,1.559 | 0.071,0.044,-0.710,0.378,1.826,0.515 |
| **c602e**6b3b181917caa362e094a197211 | 0.097,0.010,1.124,-1.478,-0.789,0.226 | 0.099,0.072,1.071,-0.796,0.784,-1.378 | 0.048,0.093,-0.500,0.531,1.114,0.499 | 0.081,0.075,-0.397,-0.102,1.762,1.146 |
| **c627e**2920c28790405c6d3a8517b63f3 | 0.062,0.056,1.702,-1.545,-1.027,-0.670 | 0.092,0.028,0.629,-0.379,-1.176,-1.077 | 0.086,0.049,-0.502,0.287,1.906,-0.394 | 0.053,0.046,-1.268,1.041,1.344,0.960 |
| **c6339**4bd554d535827125eeec41ced3b | 0.089,0.060,1.486,-1.204,-0.712,-0.456 | 0.014,0.090,1.453,-1.918,0.331,-0.908 | 0.028,0.038,-1.865,1.174,-0.231,1.457 | 0.040,0.049,-0.802,0.295,1.933,1.089 |
| **c652e**1bbfbfcc29d6e48289095c8feeb | 0.094,0.042,1.789,-1.672,-0.405,-0.903 | 0.040,0.083,0.955,-0.783,-1.947,-1.176 | 0.043,0.029,-0.687,0.607,-0.514,1.406 | 0.046,0.050,-1.216,1.165,0.888,-0.111 |
| **c6590**84e3b3942f96c57bc6f366efd1b | 0.030,0.049,1.301,-1.029,-1.090,-1.335 | 0.070,0.079,1.621,-0.999,-1.955,-0.073 | 0.055,0.019,-1.759,1.924,1.735,0.012 | 0.054,0.046,-0.922,0.676,-0.122,1.416 |
| **c6719**f3d11a1b5a58b811d6373879a37 | 0.012,0.028,0.612,-0.588,-1.276,-0.955 | 0.013,0.030,0.199,-0.182,-0.711,1.280 | 0.050,0.033,-0.860,1.872,1.769,1.069 | 0.043,0.093,-1.448,1.190,-0.569,1.518 |
| **c6884**1e2064478e9dda6b20156abcd22 | 0.072,0.069,0.707,-0.712,-1.700,1.148 | 0.081,0.089,0.800,-0.772,1.389,-1.541 | 0.083,0.033,-1.710,1.886,1.872,1.692 | 0.048,0.084,-0.683,0.382,-0.380,1.910 |
| **c6c48**72e9a34fbee178fbf7ef0097a8f | 0.020,0.077,0.838,-0.622,-1.224,-0.998 | 0.039,0.081,1.012,-0.823,-1.154,-0.615 | 0.086,0.027,-1.540,1.412,-0.097,1.325 | 0.095,0.035,-0.456,0.038,1.494,0.773 |
| **c6dd3**a81c47621acba416fcd5e58a82d | 0.098,0.023,1.830,-1.393,-1.942,0.944 | 0.086,0.015,0.652,-0.609,-1.325,-1.001 | 0.097,0.024,-0.385,-0.012,1.006,0.458 | 0.026,0.028,-1.328,1.211,1.924,0.679 |
| **c6f3c**91c51e889b122d2d34d7879c6d1 | 0.082,0.017,1.637,-1.634,-1.845,-0.546 | 0.054,0.063,0.585,-0.578,-1.280,-1.255 | 0.048,0.064,-1.771,0.952,-0.393,1.445 | 0.061,0.021,-0.860,0.756,0.982,0.846 |
| **c7004**0d192304dd4ef0175070813922b | 0.057,0.063,1.557,-1.979,0.826,-0.558 | 0.096,0.096,0.540,-1.246,-1.842,1.833 | 0.016,0.049,-1.194,0.580,0.841,1.383 | 0.080,0.060,-1.000,0.613,1.699,1.455 |
| **c7046**c8a6d722eb43e2ee5f6cd68c93c | 0.044,0.082,1.915,-1.536,-1.615,-0.856 | 0.087,0.067,0.509,-1.605,-0.808,1.244 | 0.045,0.042,-0.511,0.130,1.209,0.521 | 0.098,0.057,-0.947,0.636,0.538,1.670 |
| **c7821**f9492abb6aaceef40072f5a8201 | 0.049,0.048,1.759,-1.477,-1.067,-0.742 | 0.077,0.049,0.839,-1.900,0.054,0.075 | 0.062,0.088,-1.429,0.396,1.750,0.449 | 0.050,0.066,-0.811,0.438,1.785,1.295 |
| **c7def**9ea46e538b794ba86a5fddc140c | 0.053,0.053,1.748,-1.439,-1.471,-0.870 | 0.023,0.085,0.702,-1.374,0.049,-0.805 | 0.082,0.062,-0.883,0.249,0.518,0.957 | 0.029,0.085,-0.375,-0.012,1.971,1.953 |
| **c80e1**8d8cd1f692f6529d57c80f8cdb2 | 0.023,0.066,1.556,-1.207,-0.742,-1.768 | 0.099,0.013,0.730,-0.733,-0.304,-0.469 | 0.014,0.060,-0.909,0.535,0.499,1.763 | 0.054,0.091,-1.233,0.864,-0.267,1.438 |
| **c8420**68d2ec26384a6c209de6058f570 | 0.048,0.098,1.638,-1.201,-0.501,-1.829 | 0.040,0.075,1.218,-1.816,0.748,-0.707 | 0.091,0.091,-1.934,1.866,0.367,0.335 | 0.060,0.098,-0.277,-0.233,1.518,1.493 |
| **c8669**f84c13cf99d49e3f70061262ad0 | 0.083,0.073,0.332,-0.219,0.139,-1.495 | 0.014,0.088,0.739,-0.650,-1.347,0.528 | 0.095,0.075,-1.291,1.689,0.361,1.886 | 0.099,0.045,-0.318,0.169,-0.050,1.477 |
| **c8892**d018668a203c783c93e4ce7eaf7 | 0.032,0.059,1.745,-1.718,-1.148,-0.254 | 0.042,0.050,1.384,-1.022,-0.151,0.031 | 0.052,0.093,-0.792,0.517,0.666,1.235 | 0.056,0.042,-0.735,0.191,1.385,1.781 |
| **c8ab2**b4c56790b0d3958946b540695d4 | 0.017,0.055,1.884,-1.077,-1.965,-1.802 | 0.090,0.090,0.473,-0.670,-1.892,0.423 | 0.018,0.073,-1.210,0.357,1.484,0.383 | 0.083,0.042,-0.618,0.117,1.958,1.213 |
| **c8cac**c359d7da6eedb8bac2abe12ed6a | 0.095,0.016,1.806,-1.582,-1.625,-0.351 | 0.071,0.060,1.945,-1.250,-0.818,-1.746 | 0.089,0.035,-1.971,1.901,1.794,-0.257 | 0.033,0.073,-1.706,1.209,1.970,1.345 |
| **c8cf1**73a53c83587ac5fa36e8bd62035 | 0.045,0.096,1.165,-1.004,-0.674,-0.446 | 0.034,0.018,0.366,-0.438,-1.972,-0.367 | 0.046,0.083,-0.926,0.679,0.403,-1.293 | 0.091,0.061,-1.142,0.577,1.524,1.707 |
| **c8e6f**a8dd4ea1853c8df6233f144a4c1 | 0.058,0.027,1.133,-1.049,-1.702,-0.291 | 0.058,0.096,0.638,-1.518,1.183,0.092 | 0.086,0.089,-1.215,0.228,1.212,1.793 | 0.017,0.073,-1.160,0.903,0.673,1.453 |
| **c8fb7**35f1b20f8f849cf8d9ff8611cb8 | 0.024,0.021,1.445,-1.429,-1.154,-0.818 | 0.043,0.067,1.005,-1.720,-0.803,-0.330 | 0.022,0.044,-1.882,0.971,1.117,1.533 | 0.098,0.077,-0.486,-0.149,1.722,0.851 |
| **c9616**b4031bb4e683ae1416b2791abb5 | 0.066,0.056,1.365,-1.254,-1.093,0.487 | 0.074,0.063,1.527,-0.767,-1.616,-1.163 | 0.022,0.092,-1.082,1.374,1.222,-0.205 | 0.052,0.061,0.015,-0.020,0.920,-0.554 |
| **c976d**a6c71adc360ec89295a4788d79b | 0.035,0.034,1.260,-1.101,-1.801,-1.081 | 0.048,0.023,0.986,-1.084,-1.307,0.540 | 0.053,0.091,-1.663,1.705,-0.408,0.852 | 0.052,0.035,-0.697,0.542,0.793,0.844 |
| **c979e**6a40242946d84ec3cc1d22cf570 | 0.095,0.021,1.774,-1.837,-1.155,0.089 | 0.011,0.086,0.618,-0.497,0.283,-1.738 | 0.093,0.021,-1.845,1.748,1.768,1.641 | 0.049,0.096,-1.821,1.554,-0.406,1.883 |
| **c97f7**496e0ad7aa031d161d5055882f8 | 0.089,0.048,1.726,-0.837,-1.969,-1.310 | 0.037,0.089,0.340,-1.398,0.299,-0.062 | 0.046,0.038,-0.816,0.177,1.465,0.366 | 0.058,0.080,-0.327,0.110,1.917,1.015 |
| **c9bc0**0d50ca71d101ca745c6c4da4fb0 | 0.035,0.025,1.708,-1.201,-1.902,-1.153 | 0.018,0.024,0.762,-1.858,-1.357,-1.357 | 0.035,0.065,-1.458,0.526,0.307,0.645 | 0.082,0.040,-0.681,0.443,0.984,0.388 |
| **c9cad**1f99f34238bce9fb4987d9f4c7f | 0.040,0.018,1.587,-1.532,-1.034,-0.207 | 0.019,0.039,0.246,-1.760,-1.899,1.467 | 0.086,0.036,-1.689,0.213,1.135,0.832 | 0.065,0.087,-1.624,1.420,1.694,0.507 |
| **c9f44**8265b96fbb82bd8382f7c0f94f0 | 0.080,0.058,1.509,-1.523,-0.149,-1.222 | 0.016,0.033,0.390,-0.663,1.154,-1.883 | 0.093,0.097,-1.317,0.307,1.442,1.890 | 0.059,0.073,-1.026,0.734,1.283,1.682 |
| **ca0a7**83f485e23e4445e0978eb84a050 | 0.040,0.081,0.798,-0.762,-1.352,-1.183 | 0.037,0.045,0.587,-0.410,-0.363,-0.806 | 0.071,0.091,-0.778,0.847,1.868,1.658 | 0.083,0.072,-0.765,0.459,1.765,1.463 |
| **ca0dc**7377a23eb898af3d57a0c724cbd | 0.092,0.081,1.177,-0.982,-0.870,-0.382 | 0.036,0.020,1.575,-1.615,0.592,1.325 | 0.092,0.068,-1.545,1.909,1.947,-0.594 | 0.063,0.075,-0.560,0.050,1.437,1.850 |
| **ca1c4**b7b24629bc2d0bc4c0010f79004 | 0.098,0.069,1.075,-1.342,-1.887,1.048 | 0.090,0.040,0.550,-0.483,-0.916,1.022 | 0.024,0.038,-1.216,1.489,1.918,-0.851 | 0.087,0.029,-0.286,-0.093,1.560,1.819 |
| **ca1d5**80efe1d6cdad5f6a3ad17cff4fc | 0.098,0.046,0.533,-0.396,-1.708,-0.385 | 0.091,0.075,1.838,-0.892,-0.917,-1.429 | 0.091,0.025,-1.518,1.945,1.922,0.566 | 0.066,0.083,-0.432,-0.141,1.658,1.550 |
| **ca1ed**bde96a52a1a389f22a9386ba3d2 | 0.036,0.080,1.598,-1.447,-0.308,-1.277 | 0.078,0.062,1.756,-1.979,-1.937,0.002 | 0.035,0.038,-1.418,1.066,1.839,1.122 | 0.095,0.035,-0.652,0.148,1.908,1.409 |
| **ca664**79370beb9205d698466c1d7cae2 | 0.079,0.056,1.407,-1.110,-0.764,-1.638 | 0.045,0.077,1.485,-1.450,0.125,-0.127 | 0.096,0.094,-1.850,1.866,-0.115,0.506 | 0.096,0.084,-0.671,-0.048,1.885,1.123 |
| **ca87c**08e9e54767025a99382475a50a8 | 0.043,0.074,1.752,-1.608,0.191,-1.905 | 0.020,0.065,1.383,-1.742,1.698,-1.124 | 0.053,0.050,-1.776,1.726,-0.397,1.798 | 0.051,0.087,-1.065,0.472,1.772,1.866 |
| **ca95c**55cefcbc7077050298fd259e099 | 0.054,0.076,1.409,-1.046,-0.408,-1.283 | 0.043,0.070,0.593,-1.900,0.035,1.050 | 0.077,0.063,-0.901,-0.025,1.812,1.967 | 0.081,0.074,-1.722,1.344,1.901,1.096 |
| **ca9a2**29e5c20f603e2546aca250e21b3 | 0.036,0.018,1.076,-1.084,-0.743,-0.432 | 0.063,0.087,0.991,-0.634,-0.487,-1.195 | 0.045,0.016,-1.577,1.949,1.739,0.911 | 0.096,0.070,-0.225,-0.404,1.122,1.616 |
| **caaeb**2ed510a6acba24d9e5b34ff18d4 | 0.070,0.058,0.418,-0.339,0.093,-0.983 | 0.072,0.033,0.634,-0.235,-1.979,0.813 | 0.081,0.064,-0.590,0.395,0.881,1.802 | 0.060,0.066,-0.626,0.281,0.799,1.433 |

| | | | | |
|---|---|---|---|---|
| **cacbe**49103005ee1bd2329b22273c718 | 0.020,0.061,1.576,-1.716,-0.575,-0.743 | 0.044,0.042,0.817,-0.646,1.951,-0.470 | 0.084,0.057,-1.137,1.563,1.823,1.309 | 0.068,0.053,-0.710,0.181,1.157,1.443 |
| **caea4**57518b6eaa72454a170eb374628 | 0.041,0.056,0.768,-0.608,-0.923,-0.762 | 0.013,0.046,0.859,-1.027,1.798,-0.185 | 0.077,0.019,-1.138,1.003,1.816,1.945 | 0.092,0.066,-1.517,1.106,1.523,0.433 |
| **caf21**6ff5aaf777fb3ebc2b42de3cb2b | 0.013,0.049,1.688,-1.385,-0.802,-1.940 | 0.031,0.081,0.603,-0.778,-1.036,-0.623 | 0.074,0.030,-1.860,0.912,1.368,-1.779 | 0.054,0.056,-1.215,1.036,-0.518,1.556 |
| **cb17d**6fce54dcade9d294e431d882526 | 0.067,0.052,1.996,-1.684,-1.893,-0.571 | 0.028,0.077,1.391,-0.954,-1.137,-0.246 | 0.037,0.060,-0.764,0.768,-0.780,1.647 | 0.092,0.091,-1.796,1.180,1.939,0.614 |
| **cb5b6**3b0f026bb99303f802e8ef6441b | 0.065,0.052,1.577,-1.653,-0.188,-0.436 | 0.068,0.084,1.181,-1.953,-0.370,-0.129 | 0.082,0.062,-1.438,0.775,-0.385,1.846 | 0.081,0.091,-0.532,-0.189,1.830,1.907 |
| **cb7d9**6c2976e72d126488af75ae71423 | 0.069,0.044,0.666,-0.616,-1.914,1.370 | 0.012,0.059,0.418,-0.203,-0.643,-1.156 | 0.095,0.012,-0.993,1.835,0.857,0.545 | 0.050,0.079,-0.502,0.196,1.913,1.702 |
| **cb8bf**3ab23d15533ccfb11cd0762b2e3 | 0.045,0.083,1.339,-1.240,-0.337,-0.475 | 0.018,0.026,0.823,-0.564,-0.618,-0.958 | 0.062,0.083,-1.720,1.803,1.648,1.031 | 0.059,0.021,-0.196,-0.012,1.412,-0.460 |
| **cba09**91adfbdfe1429e29b5c2ce7d9d1 | 0.015,0.083,0.880,-1.058,-1.753,-0.427 | 0.077,0.075,0.760,-0.542,-0.449,-0.224 | 0.098,0.044,-1.547,1.955,1.901,0.663 | 0.070,0.065,-0.747,0.671,0.136,0.820 |
| **cbac8**33dc83377fb13e7eb379a99f319 | 0.046,0.072,0.741,-0.739,-1.590,0.062 | 0.089,0.017,0.472,-0.346,-1.506,-1.153 | 0.076,0.080,-0.404,0.141,1.086,0.399 | 0.082,0.065,-0.726,0.416,1.640,0.935 |
| **cbb58**9d00a53b2194c619511c4eddfc5 | 0.098,0.038,1.416,-1.194,-0.625,-0.102 | 0.032,0.074,0.824,-1.913,0.138,-1.661 | 0.044,0.096,-1.480,0.335,-0.387,0.927 | 0.060,0.080,-0.931,0.620,1.365,1.059 |
| **cbbaa**18178ef860b5fb905888e5c6e5b | 0.082,0.050,0.448,-0.392,-1.257,0.160 | 0.056,0.071,0.964,-0.486,-0.882,-1.730 | 0.048,0.033,-1.241,1.978,-0.864,-0.803 | 0.041,0.076,-0.558,0.414,1.122,0.593 |
| **cbe5c**c5dbce3ea040fd374143148953c | 0.022,0.085,1.019,-1.167,-1.814,-0.487 | 0.024,0.068,0.214,-0.059,-1.349,-0.215 | 0.025,0.059,-0.843,1.958,0.887,0.862 | 0.092,0.073,-0.457,0.123,1.633,1.310 |
| **cc04a**3b0f29eb13f702b2ec10892f26b | 0.098,0.021,1.487,-0.946,-1.888,-1.177 | 0.045,0.073,1.182,-1.923,-0.878,-0.731 | 0.095,0.078,-1.969,1.071,0.584,1.359 | 0.079,0.093,-1.145,0.859,-0.589,1.647 |
| **cc284**94777b9a02299687f205e5109be | 0.075,0.042,1.517,-1.240,-1.331,-0.110 | 0.011,0.066,0.804,-1.581,-1.736,-1.047 | 0.045,0.100,-1.924,0.826,0.074,1.427 | 0.060,0.023,-0.771,0.545,1.925,0.619 |
| **cc340**75caaa13dec5f0965142f3b18df | 0.062,0.027,1.283,-1.186,-0.664,-1.034 | 0.035,0.086,1.138,-1.201,-1.321,-1.317 | 0.095,0.034,-1.318,0.894,1.620,0.293 | 0.094,0.042,-0.107,-0.101,1.179,-0.141 |
| **cc4b5**efd04933e44db86b533bcb5576b | 0.038,0.037,1.134,-1.109,-1.287,-0.441 | 0.013,0.081,0.578,-0.505,-1.813,0.179 | 0.083,0.085,-0.562,0.532,0.337,0.268 | 0.055,0.049,-0.890,0.551,1.322,0.846 |
| **cc55b**75afd25ea65d169d0f5b82a352b | 0.090,0.033,1.756,-1.396,-1.710,0.411 | 0.035,0.078,1.320,-1.078,1.369,-1.851 | 0.066,0.036,-1.780,1.355,1.912,1.416 | 0.083,0.065,-1.589,0.746,1.806,1.424 |
| **cc565**176800a7db17f6a4705466f636d | 0.094,0.018,1.686,-1.365,-1.904,1.656 | 0.021,0.087,0.530,-0.725,0.301,-0.608 | 0.081,0.023,-1.758,1.085,0.247,1.169 | 0.048,0.090,-1.218,0.935,-0.135,1.524 |
| **cc795**9b6b6cea33c14538dfd3efb5f41 | 0.036,0.049,1.453,-1.089,-1.868,-1.066 | 0.018,0.031,0.267,-0.442,-1.399,-0.727 | 0.030,0.094,-0.878,0.219,-0.130,1.174 | 0.053,0.031,-1.236,1.028,1.953,0.795 |
| **cc7eb**2e88aa67c94b82d0720de78796a | 0.093,0.020,1.728,-1.647,-1.028,-1.083 | 0.090,0.032,0.609,-1.067,0.185,-0.954 | 0.083,0.039,-1.178,0.354,1.568,0.459 | 0.066,0.069,-0.229,-0.046,1.244,0.620 |
| **cc9b2**fadbca7fb16bf5ff412235da206 | 0.092,0.020,1.545,-1.040,-1.840,0.705 | 0.024,0.047,0.370,-0.747,-0.649,-0.798 | 0.081,0.094,-1.918,0.396,1.520,1.500 | 0.015,0.025,-0.312,0.243,-0.998,1.841 |
| **cc9cd**5427043ef39f71c1aaa82262de8 | 0.084,0.018,1.661,-1.610,-1.866,-0.458 | 0.087,0.083,0.848,-1.590,0.156,-0.985 | 0.087,0.064,-1.817,0.950,0.665,0.356 | 0.091,0.038,-1.395,1.299,1.684,-1.189 |
| **cca37**72262bc786f8a65a0b31222fd4a | 0.061,0.022,1.781,-1.647,-0.715,-0.166 | 0.097,0.067,0.188,-0.136,-1.872,0.075 | 0.011,0.041,-1.254,0.386,0.572,0.457 | 0.097,0.045,-1.062,0.828,1.267,0.821 |
| **ccac3**22fc7e1655211e2d8357e86bdc6 | 0.049,0.044,1.470,-1.949,-0.322,-0.307 | 0.086,0.014,0.544,-0.291,-1.826,-0.134 | 0.072,0.091,-0.804,0.340,1.885,1.573 | 0.094,0.065,-1.424,1.009,1.628,0.937 |
| **ccb2c**eb8722cdbd31f81de810626bcc4 | 0.088,0.034,1.184,-1.294,-1.032,0.553 | 0.075,0.064,0.825,-1.981,0.057,1.664 | 0.023,0.061,-1.692,0.904,1.687,1.280 | 0.041,0.065,-0.094,-0.217,1.806,1.458 |
| **ccb50**db8ecc5bcc155d7574a8e1f08f9 | 0.089,0.063,1.543,-1.313,-0.959,-1.150 | 0.056,0.064,1.304,-1.236,-0.153,0.401 | 0.095,0.073,-1.081,1.619,-0.544,0.527 | 0.095,0.016,-0.719,0.412,1.643,0.365 |
| **ccfc2**d374e4002b52de451b0558a3747 | 0.079,0.025,1.677,-1.771,-1.112,0.062 | 0.058,0.044,0.870,-0.986,0.383,-1.417 | 0.098,0.051,-1.466,1.202,0.478,1.243 | 0.097,0.096,-0.561,-0.106,1.899,1.303 |
| **cd246**0b61b8628858adde8b4a32ae16e | 0.090,0.077,1.697,-1.911,-1.780,0.748 | 0.028,0.079,1.534,-1.594,-1.648,-0.675 | 0.017,0.059,-1.909,1.934,1.628,0.497 | 0.046,0.058,-0.354,0.032,1.885,1.469 |
| **cd448**da04daa186831cc0c053602d32d | 0.097,0.012,1.948,-1.436,-1.922,0.209 | 0.015,0.078,1.757,-1.265,-0.422,-1.533 | 0.099,0.062,-1.738,1.694,1.317,0.148 | 0.100,0.020,-1.111,0.707,0.927,1.438 |
| **cd4a8**7afcc6b822156b400378f23a959 | 0.073,0.058,1.593,-1.919,-0.186,-0.107 | 0.019,0.099,0.685,-0.476,-0.932,-0.359 | 0.089,0.085,-1.160,1.778,1.109,1.813 | 0.021,0.083,-0.467,0.251,1.748,1.525 |
| **cd658**710577da6b765139fb9e258e8f3 | 0.031,0.065,0.640,-0.567,-0.675,-0.344 | 0.069,0.055,0.985,-1.117,-0.265,-1.119 | 0.048,0.078,-1.568,1.160,-0.522,1.297 | 0.072,0.053,-0.898,0.450,1.739,1.034 |
| **cd980**248420b89829691c76bb0a13eff | 0.059,0.013,0.807,-0.728,-1.411,0.284 | 0.013,0.025,0.447,-0.552,-0.369,-1.818 | 0.063,0.049,-1.265,1.022,-0.342,1.266 | 0.057,0.099,-0.617,0.247,0.468,1.445 |
| **cd98a**a0e2835875332a5bd26346e24ca | 0.050,0.049,1.849,-1.549,-1.126,-1.137 | 0.015,0.075,0.389,-0.568,0.464,-1.341 | 0.070,0.064,-1.943,0.849,-0.327,1.143 | 0.090,0.036,-1.627,1.527,0.466,0.518 |
| **cdc6f**41dd7d3acdaaa3b4c1643f60724 | 0.054,0.029,1.955,-1.892,-1.588,0.481 | 0.085,0.067,0.971,-1.241,-1.786,-1.102 | 0.057,0.042,-1.880,0.817,1.227,0.692 | 0.084,0.077,-0.139,-0.318,1.979,0.646 |
| **cdd23**1f0b52b260e62f8bd6997d222f6 | 0.041,0.059,1.551,-1.578,-0.593,-0.369 | 0.074,0.075,1.471,-0.954,1.319,-1.494 | 0.090,0.060,-1.386,1.929,1.133,0.597 | 0.073,0.070,-0.817,0.625,1.675,-0.631 |
| **cddb7**5f3022f2b581fdb37bdc43b2582 | 0.092,0.015,1.157,-1.249,-1.206,0.136 | 0.099,0.086,0.842,-0.316,-1.442,-0.383 | 0.075,0.060,-0.605,0.399,0.760,1.492 | 0.093,0.099,-0.285,-0.438,1.941,1.253 |
| **ce167**b8809aa66ecd9b3d7e20f228655 | 0.045,0.029,1.546,-1.432,-1.785,-0.639 | 0.060,0.061,0.664,-1.762,0.134,-0.695 | 0.050,0.053,-1.719,0.400,1.773,1.939 | 0.012,0.069,-0.646,0.402,0.048,1.850 |
| **ce1cf**b72d2da6663b063243a0812b1d8 | 0.027,0.042,1.665,-1.367,-1.922,0.072 | 0.092,0.013,1.176,-1.126,-1.216,-1.057 | 0.072,0.046,-1.537,0.120,1.204,1.160 | 0.018,0.037,-1.735,1.299,1.353,1.501 |
| **ce23b**7b6ee4dcc71ca205a3e8b4a7a2e | 0.096,0.016,1.577,-1.202,-1.690,-0.122 | 0.058,0.068,0.655,-1.212,0.001,-0.322 | 0.040,0.028,-1.319,0.755,0.677,0.548 | 0.056,0.062,-0.201,-0.020,1.706,0.342 |
| **ce351**5366cc761f3f284aa5fad35dc3f | 0.040,0.033,1.897,-1.890,-1.883,-0.989 | 0.038,0.051,0.415,-0.489,-1.367,-1.515 | 0.045,0.033,-1.418,0.534,1.322,0.678 | 0.038,0.092,-1.718,1.389,1.916,1.524 |
| **ceab5**5f68ca46bd54ec8e6edb0ac96d6 | 0.096,0.078,1.329,-1.331,-0.316,-1.343 | 0.021,0.020,0.767,-0.932,-0.298,0.312 | 0.094,0.096,-0.636,0.169,1.048,0.589 | 0.051,0.076,-1.123,0.608,1.620,1.778 |
| **ceb0e**16b7f90e4603d69a428cf7aaaf6 | 0.089,0.072,1.555,-1.304,-0.988,-0.203 | 0.023,0.069,0.800,-0.716,-1.269,-0.844 | 0.096,0.059,-0.823,0.467,-0.085,1.592 | 0.018,0.066,-0.563,0.306,1.175,1.055 |
| **ceb2c**6f3d0b8c85c9ac3adfba9756db0 | 0.062,0.052,1.669,-1.209,-1.689,-0.855 | 0.037,0.037,1.514,-1.511,-1.804,-1.548 | 0.037,0.043,-1.709,1.175,1.788,0.070 | 0.074,0.080,-0.159,-0.042,1.311,-0.120 |
| **cec13**f038999b2db8a449d22d440d1d9 | 0.090,0.017,0.614,-0.538,-1.426,1.059 | 0.023,0.084,1.136,-1.220,-0.625,-1.831 | 0.083,0.063,-1.418,1.269,0.964,-0.855 | 0.059,0.076,-0.256,0.090,0.934,0.804 |
| **cecab**ae608f334cf55b9f24c92ea450a | 0.081,0.033,1.652,-1.814,0.092,-1.241 | 0.030,0.051,1.541,-1.116,-0.686,-1.213 | 0.086,0.082,-1.586,1.935,-0.240,0.916 | 0.049,0.029,-0.439,0.305,1.708,0.842 |
| **cecc5**e4f4388dcd92eee4741a0382ca7 | 0.096,0.088,1.870,-1.308,-1.608,-0.933 | 0.032,0.075,1.366,-1.141,-1.363,0.112 | 0.038,0.032,-1.700,1.873,-0.657,1.361 | 0.083,0.076,-1.406,0.443,1.953,1.347 |
| **ceff9**aed5d8ceacb3d871116c5209591 | 0.100,0.050,1.609,-1.742,-0.598,-0.888 | 0.091,0.074,1.143,-0.683,-0.514,-1.892 | 0.058,0.028,-0.800,1.063,-0.089,0.317 | 0.050,0.070,-0.245,-0.031,1.014,1.256 |
| **cf0b4**315fd9b16026ba602a95ee34729 | 0.039,0.016,1.109,-1.180,-1.180,-1.083 | 0.012,0.088,0.578,-0.311,-1.033,-1.521 | 0.050,0.019,-1.661,1.797,-1.104,1.225 | 0.085,0.073,-0.866,0.405,1.977,0.931 |
| **cf221**ef81999d70415790858dbfe726f | 0.057,0.048,1.514,-1.416,-1.538,0.768 | 0.028,0.079,1.181,-1.465,-1.621,-0.033 | 0.023,0.058,-1.518,0.791,0.146,1.397 | 0.040,0.045,-0.369,0.237,1.747,-0.728 |
| **cf29f**c3330140577b2abc222e32a579d | 0.096,0.025,1.595,-1.687,-1.863,0.877 | 0.046,0.030,0.439,-0.326,-0.681,-0.198 | 0.047,0.060,-0.795,1.105,1.930,0.672 | 0.026,0.099,-0.896,0.621,1.952,1.254 |
| **cf49b**0f6b31bc73fd507882f43a31fb0 | 0.055,0.075,1.087,-0.549,-1.175,-1.806 | 0.073,0.071,1.893,-0.867,-0.474,0.139 | 0.030,0.090,-0.900,0.786,1.281,1.586 | 0.033,0.099,-1.082,0.756,0.974,0.275 |
| **cf5ec**7f3dcdfc447c7d3de22bdce8080 | 0.099,0.018,1.743,-1.418,-1.786,0.042 | 0.057,0.086,0.976,-0.796,-0.381,-1.345 | 0.078,0.044,-0.353,0.119,0.017,1.761 | 0.078,0.049,-0.157,-0.380,1.670,1.606 |
| **cf621**d24ed64e7797d7fb5f82d405ec2 | 0.073,0.036,0.588,-0.579,-0.783,0.713 | 0.089,0.082,1.208,-1.839,-0.837,0.898 | 0.037,0.092,-1.022,0.391,-0.417,0.865 | 0.021,0.020,-1.041,0.933,-0.548,1.482 |
| **cf71b**56e5610e952c1942d1a29622f89 | 0.065,0.042,1.708,-1.874,-0.604,-0.864 | 0.045,0.083,1.943,-1.346,-1.688,1.589 | 0.025,0.087,-0.789,0.991,1.657,1.550 | 0.045,0.075,-0.888,0.561,1.686,0.669 |
| **cf877**c51bb485e70a8f817290de078df | 0.056,0.087,1.709,-1.671,-0.497,-0.917 | 0.040,0.048,0.673,-1.032,-1.190,0.913 | 0.084,0.094,-0.612,0.187,0.684,0.574 | 0.075,0.072,-0.918,0.547,1.392,0.937 |
| **cf966**1edea3daca6eae35af65b611684 | 0.060,0.057,1.347,-1.265,-0.311,-0.935 | 0.043,0.082,0.605,-0.843,-0.751,-1.038 | 0.079,0.034,-1.815,0.662,0.382,1.664 | 0.089,0.089,-0.415,-0.202,1.596,1.463 |
| **cfac9**11f2e99387338912f81ed45a345 | 0.077,0.026,1.847,-1.854,-1.088,-1.081 | 0.023,0.025,0.344,-0.869,-1.881,0.386 | 0.080,0.058,-1.691,0.439,1.781,1.919 | 0.012,0.085,-0.628,0.375,-1.046,1.913 |
| **cfd96**95c0709ed3b092f00d87387975a | 0.087,0.028,1.746,-1.258,-1.753,-0.408 | 0.014,0.083,0.972,-1.714,-0.164,-0.855 | 0.054,0.088,-0.894,0.220,0.260,0.991 | 0.038,0.097,-0.504,-0.035,1.214,1.825 |
| **cfe83**5a0d84a1ed004a8d509299f6a68 | 0.049,0.055,1.489,-1.177,-1.873,-0.424 | 0.079,0.043,1.395,-1.495,-1.168,0.141 | 0.047,0.031,-1.813,1.623,0.037,0.906 | 0.022,0.042,-0.190,-0.071,1.927,1.031 |
| **cffd3**3c5f41c2cb78a1a03433998f6df | 0.052,0.065,1.460,-1.046,-1.593,-0.839 | 0.084,0.044,1.498,-0.598,-0.025,-1.573 | 0.074,0.068,-0.437,-0.089,1.880,0.470 | 0.037,0.014,-1.804,1.317,1.478,1.893 |
| **d040a**3e647b7a883a1fb397f43d8c68b | 0.050,0.051,1.763,-1.363,-1.243,-1.214 | 0.054,0.089,1.166,-1.588,-1.218,-0.663 | 0.036,0.089,-1.383,0.410,1.678,1.711 | 0.084,0.090,-0.739,0.013,1.865,1.341 |
| **d059a**20aa7d95e0154856c815bbf5d26 | 0.063,0.021,1.503,-1.849,0.099,-0.397 | 0.095,0.080,0.294,-0.558,-0.679,0.967 | 0.073,0.042,-0.868,0.369,0.411,1.692 | 0.082,0.070,-0.856,0.489,1.979,0.681 |
| **d0648**1e51440d055b5f513f00a0d575d | 0.096,0.016,1.890,-1.540,-1.489,-0.341 | 0.090,0.074,0.712,-1.374,-1.482,0.031 | 0.045,0.061,-1.553,0.614,1.042,1.598 | 0.065,0.080,-0.622,0.370,1.685,0.569 |
| **d06e0**c21a914e9afe470b14092d71e73 | 0.048,0.039,1.562,-1.457,-1.160,-1.006 | 0.022,0.038,1.465,-1.919,0.212,-1.383 | 0.036,0.087,-1.940,1.314,1.922,0.168 | 0.060,0.096,-0.500,-0.128,1.297,1.261 |
| **d0725**5c627cfc052b4fce87fb4e7f94a | 0.040,0.049,1.903,-1.832,-1.717,-0.142 | 0.041,0.019,0.208,-0.238,-1.399,-0.657 | 0.016,0.047,-1.059,0.449,0.926,1.953 | 0.082,0.065,-0.554,0.382,-0.436,1.848 |
| **d0c80**0c226c6a99f9349d74ad0a08290 | 0.051,0.047,1.078,-1.107,-0.793,-0.201 | 0.087,0.033,1.346,-1.399,-0.457,-1.793 | 0.096,0.070,-1.797,1.022,1.310,1.284 | 0.094,0.098,-0.623,0.275,1.379,0.392 |
| **d0f18**03a080ada448eac0ddd1fe753a2 | 0.027,0.037,1.142,-1.361,-1.094,-0.701 | 0.061,0.015,0.339,-0.263,-0.403,-0.579 | 0.016,0.099,-1.227,1.239,-0.420,0.540 | 0.085,0.074,-1.034,0.524,1.561,1.293 |
| **d11c9**4110d44131ba81432c8ceb366dd | 0.090,0.025,1.441,-1.069,-1.945,-0.194 | 0.054,0.081,0.185,0.004,-1.949,-0.199 | 0.079,0.063,-1.702,0.718,1.804,-0.158 | 0.094,0.075,-1.536,1.265,0.922,1.119 |

| | | | | |
|---|---|---|---|---|
| **d11fd**a0ea9f0f44d4355412fdefce152 | 0.020,0.014,0.840,-0.797,-1.431,-0.054 | 0.058,0.051,1.035,-0.549,-0.843,-0.992 | 0.072,0.072,-1.236,1.464,-0.258,1.138 | 0.019,0.069,-0.106,0.016,0.921,0.255 |
| **d131c**5bcd670622b741e72899e9fc30c | 0.076,0.028,1.449,-0.873,-1.663,-1.043 | 0.051,0.065,0.927,-1.352,-1.279,-0.858 | 0.091,0.049,-1.232,0.587,-0.587,1.235 | 0.079,0.095,-0.166,-0.477,1.186,1.463 |
| **d1428**9a25cb01c74943d26304330b566 | 0.056,0.090,1.510,-1.533,-0.789,-1.116 | 0.019,0.082,0.710,-0.735,0.257,0.030 | 0.061,0.049,-1.692,1.746,1.253,1.925 | 0.051,0.029,-0.686,0.556,0.588,1.324 |
| **d157c**1c5c33bb781cb34436a13a28565 | 0.058,0.093,1.775,-1.505,-0.128,-0.952 | 0.041,0.035,1.255,-1.647,-1.086,-1.641 | 0.053,0.098,-1.970,1.193,1.812,0.454 | 0.091,0.035,-1.150,0.789,1.891,0.269 |
| **d1705**51701c5ea983524bf0153287ae7 | 0.041,0.033,0.837,-1.058,-0.979,0.303 | 0.065,0.045,0.750,-0.723,0.321,0.295 | 0.069,0.089,-1.368,1.841,1.249,-0.394 | 0.086,0.034,-0.486,0.436,-0.179,1.483 |
| **d17c1**12c2c194665d3f8f58671788a9e | 0.055,0.050,1.605,-1.567,-1.616,0.596 | 0.089,0.044,0.707,-0.676,0.279,-1.530 | 0.046,0.087,-1.088,1.128,-0.386,0.433 | 0.072,0.093,-1.288,1.009,1.192,-0.066 |
| **d192c**500a27bc492e248e050aa6cfcb8 | 0.098,0.048,0.756,-0.705,-1.062,0.540 | 0.043,0.085,0.669,-0.340,-0.799,-1.748 | 0.071,0.033,-1.144,1.447,-0.591,-1.368 | 0.086,0.062,-1.404,1.032,1.111,1.346 |
| **d1db5**b7d39b9a515359e06345c7ef044 | 0.058,0.064,1.412,-1.071,-1.338,-1.236 | 0.066,0.035,0.309,-0.611,1.242,-1.465 | 0.081,0.051,-1.310,0.907,1.023,0.525 | 0.062,0.019,-0.477,0.260,1.110,0.102 |
| **d1e8b**03d5afd1c14b149c882a991067c | 0.056,0.030,1.770,-1.680,-0.578,-1.175 | 0.096,0.048,1.832,-1.832,-1.890,-1.324 | 0.069,0.044,-1.854,1.271,1.929,0.832 | 0.085,0.097,-0.473,-0.049,1.982,0.626 |
| **d2190**35b44684adf90c6ba6540bd5779 | 0.057,0.068,0.692,-0.911,-1.175,0.530 | 0.097,0.014,0.839,-0.547,-1.633,0.029 | 0.060,0.021,-0.787,0.568,1.149,1.861 | 0.084,0.043,-1.099,0.676,1.955,0.724 |
| **d2270**0eb2a54978b795901c7116e4890 | 0.020,0.049,0.705,-0.680,0.707,-1.166 | 0.049,0.093,0.591,-0.552,-0.470,-0.216 | 0.056,0.050,-1.333,1.911,-1.423,1.908 | 0.091,0.032,-1.397,1.176,1.589,-0.502 |
| **d239f**27fecd995434286ef37a36fec95 | 0.062,0.014,1.192,-1.165,-0.573,-0.244 | 0.080,0.013,1.018,-1.351,-1.395,0.051 | 0.096,0.098,-0.962,0.459,1.465,0.244 | 0.084,0.070,-1.007,0.761,0.387,1.280 |
| **d26f4**67c9ca9b0c99cfddc11b5ade763 | 0.046,0.020,1.839,-1.870,-1.360,-0.456 | 0.069,0.075,1.632,-1.904,-1.839,-1.326 | 0.068,0.079,-1.866,1.303,0.162,1.855 | 0.099,0.076,-0.743,0.185,1.937,1.676 |
| **d2a6d**c2be5d0471e0f5a114f46a1ea61 | 0.060,0.029,1.559,-1.101,-1.651,-1.181 | 0.076,0.074,0.695,-0.359,-1.955,-0.439 | 0.070,0.100,-1.950,0.948,0.706,1.544 | 0.080,0.076,-0.583,0.244,1.369,0.475 |
| **d2a7f**b9266ed0a6d85a34452d403d447 | 0.044,0.042,1.193,-1.351,-0.314,-0.589 | 0.080,0.032,0.644,-1.070,-0.412,0.156 | 0.018,0.080,-0.310,0.115,1.448,0.524 | 0.074,0.051,-1.408,1.214,0.716,1.239 |
| **d2aa8**6a6c5d57d1bac84d9d0e152ab1b | 0.053,0.037,1.451,-1.293,-1.935,0.699 | 0.014,0.053,0.928,-0.540,-1.594,-1.516 | 0.063,0.032,-1.856,1.843,1.803,1.243 | 0.043,0.027,-1.194,0.857,1.803,0.996 |
| **d2af5**9e85486b30319f2eb1e881da716 | 0.037,0.012,1.215,-1.140,-1.849,-0.120 | 0.085,0.012,0.514,-0.202,-1.411,-1.700 | 0.012,0.078,-0.390,0.188,-0.612,0.522 | 0.082,0.014,-0.138,-0.291,1.863,1.647 |
| **d2c99**2f05be2f28e62adf72393da39af | 0.041,0.078,1.907,-1.863,-0.377,-0.555 | 0.091,0.028,0.434,-0.590,-1.551,-0.938 | 0.054,0.046,-0.340,0.023,0.546,0.943 | 0.097,0.071,-1.637,1.364,0.212,1.841 |
| **d2cee**6e4124bb71ba5384e2b5c77d001 | 0.058,0.047,1.894,0.383,-1.596,-0.745 | 0.088,0.087,1.633,-1.357,-1.897,-1.027 | 0.029,0.050,-1.061,0.137,-0.363,0.733 | 0.062,0.059,-0.481,-0.362,1.422,1.319 |
| **d2d94**0911a3d8cf37625746dd6ef0415 | 0.074,0.049,1.776,-1.448,-0.617,-1.915 | 0.045,0.070,1.945,-1.587,-0.974,-1.162 | 0.061,0.025,-1.246,1.167,1.846,-0.758 | 0.047,0.097,-1.378,0.954,1.522,0.718 |
| **d2e1a**9957e4082d8f3f150489d509c14 | 0.077,0.046,1.977,-1.732,-0.971,-1.108 | 0.078,0.053,1.077,-1.244,-1.981,-1.884 | 0.023,0.089,-1.917,0.921,1.880,1.187 | 0.095,0.044,-0.319,-0.084,1.421,1.502 |
| **d2ecf**4fcc0ae90625a32d582d2013c78 | 0.039,0.017,1.903,-1.897,-1.765,0.341 | 0.085,0.017,0.634,-1.152,-0.676,-1.198 | 0.083,0.084,-1.300,0.545,0.819,0.130 | 0.030,0.099,-0.566,0.312,1.841,0.929 |
| **d304a**f6450ca455316f6fe1afe08947f | 0.094,0.098,1.188,-0.930,-1.534,-0.184 | 0.023,0.073,0.168,-0.180,-0.604,-0.571 | 0.072,0.073,-1.439,0.837,1.217,-1.249 | 0.065,0.027,-1.067,0.993,0.816,0.854 |
| **d320a**1dfb03498a1884404fa653327c9 | 0.053,0.018,1.375,-1.407,-1.929,1.055 | 0.047,0.052,0.601,-1.295,-0.960,0.964 | 0.099,0.099,-0.501,0.135,0.140,0.578 | 0.079,0.082,-0.573,0.330,-0.384,1.742 |
| **d3213**7c892443383e6c58c611b71439c | 0.097,0.083,0.733,-0.498,-0.964,-0.234 | 0.097,0.029,1.930,-1.267,-1.663,-1.578 | 0.063,0.036,-1.910,1.982,-0.152,0.950 | 0.073,0.028,-0.168,0.096,1.174,-1.660 |
| **d3310**ab3a3d8176ed9eebaa690e34aea | 0.025,0.080,0.970,-1.324,-0.219,-0.606 | 0.021,0.017,0.255,-0.153,0.046,-1.239 | 0.094,0.099,-1.276,1.644,1.883,0.907 | 0.014,0.020,-0.147,0.120,0.940,1.090 |
| **d3352**41faf51e28e6742f08fabdd51ef | 0.052,0.052,1.614,-0.092,-1.943,-0.933 | 0.075,0.035,0.336,-0.550,-0.966,-1.950 | 0.072,0.097,-1.815,-0.079,1.187,1.268 | 0.091,0.084,-0.605,0.370,0.371,1.018 |
| **d343c**0b796021ecb82af96d79e808538 | 0.078,0.026,1.303,-1.199,-1.434,-0.383 | 0.081,0.066,0.550,-0.461,-1.367,0.472 | 0.012,0.082,-0.604,0.223,-0.830,1.935 | 0.056,0.069,-0.229,-0.117,1.929,1.561 |
| **d35e6**9afa7cf678ea8b9c5b269bb8b8c | 0.048,0.073,0.722,-0.653,0.297,-1.668 | 0.042,0.023,1.436,-1.440,0.335,-0.946 | 0.065,0.064,-1.850,1.947,1.491,0.901 | 0.014,0.035,-0.654,0.557,-1.724,1.902 |
| **d3a37**c887d9ea97e79a8eb984018068e | 0.042,0.072,1.902,-1.379,-1.868,-1.695 | 0.090,0.096,1.967,-1.711,-1.440,0.932 | 0.066,0.061,-1.871,1.926,0.778,0.999 | 0.043,0.043,-1.682,1.365,1.058,0.994 |
| **d3a5e**6c3fe1dfe4c59b0162e65aa8524 | 0.045,0.066,1.683,-1.385,-0.726,-1.644 | 0.059,0.046,0.610,-0.893,-0.343,-1.207 | 0.014,0.039,-0.318,0.161,-0.118,0.166 | 0.096,0.031,-1.166,0.950,1.592,-0.584 |
| **d3e90**fa276f988efada642baf6f8524f | 0.096,0.040,1.877,-1.948,-1.570,0.134 | 0.021,0.079,1.095,-0.663,-1.698,-1.293 | 0.088,0.059,-1.525,1.948,-1.192,1.508 | 0.081,0.054,-1.004,0.804,-0.284,1.630 |
| **d3ec8**59b6ea2e16b81555715cc526466 | 0.090,0.017,1.956,-1.848,-1.880,-0.918 | 0.067,0.073,0.388,-0.879,-0.365,0.759 | 0.059,0.068,-1.398,0.677,0.773,1.057 | 0.049,0.070,-0.668,0.284,1.601,1.835 |
| **d40dd**5c3ee9c93865a05d4beddcd75dc | 0.091,0.020,1.748,-1.902,-1.715,0.158 | 0.095,0.080,0.679,-0.556,-0.676,0.004 | 0.069,0.086,-0.792,0.860,-0.107,0.162 | 0.098,0.066,-0.516,-0.052,1.946,1.411 |
| **d44ab**1e0d4076a96a22d40cedfbb0d68 | 0.056,0.054,1.606,-1.657,-1.794,-0.016 | 0.073,0.022,0.907,-1.015,0.204,-0.449 | 0.084,0.035,-0.760,0.550,1.654,0.873 | 0.076,0.069,-0.361,0.040,1.754,0.143 |
| **d456c**4f38cf3f9bfc3acab60daf0b98f | 0.049,0.039,1.780,-1.550,-0.927,-1.793 | 0.014,0.094,0.429,-0.225,-1.133,-1.094 | 0.067,0.077,-1.257,1.019,1.375,0.373 | 0.091,0.080,-0.686,0.228,1.771,1.035 |
| **d5234**4332ee0fe3cfac833161c9be780 | 0.011,0.080,1.490,-1.185,0.341,-1.919 | 0.039,0.078,0.557,-0.299,-1.608,-0.659 | 0.070,0.047,-1.285,0.472,1.897,1.912 | 0.065,0.067,-0.432,0.442,1.349,-1.134 |
| **d52c6**7e22b20055318945df429c4a54f | 0.046,0.056,1.713,-1.803,-0.184,-1.059 | 0.013,0.086,0.657,-0.617,-1.813,-0.763 | 0.073,0.096,-1.149,0.279,1.888,1.858 | 0.083,0.096,-0.463,-0.127,1.443,1.381 |
| **d52d5**87d16480a9ecb76332e9567dbf1 | 0.089,0.027,1.052,-0.346,-1.634,-0.438 | 0.019,0.090,0.891,-1.173,-0.740,-1.865 | 0.078,0.091,-1.863,0.698,0.019,0.520 | 0.036,0.031,-0.278,0.065,1.517,1.562 |
| **d5373**ae7093ea265747d42b94f242fa9 | 0.066,0.044,1.786,-1.707,-1.181,-0.651 | 0.026,0.079,1.818,-1.126,-0.467,-1.884 | 0.064,0.047,-1.725,1.908,1.493,-0.534 | 0.083,0.037,-1.134,0.638,1.928,0.288 |
| **d5435**ea13997ceb82b36ea362971cb2d | 0.078,0.052,1.071,-0.684,-1.318,-0.606 | 0.058,0.073,0.521,-1.054,-0.688,0.112 | 0.019,0.059,-1.880,0.562,1.578,1.942 | 0.061,0.095,-1.442,1.043,0.672,1.458 |
| **d544f**95c6bcfb3843d829d9154e27908 | 0.044,0.022,1.234,-1.260,-1.672,0.073 | 0.078,0.014,1.928,-1.063,-0.889,-0.678 | 0.037,0.069,-1.110,1.954,1.562,0.376 | 0.097,0.041,-0.315,-0.311,1.931,1.593 |
| **d556a**57fe15f4b4628f6fa0bde2fb439 | 0.054,0.076,0.925,-0.725,-1.143,-0.703 | 0.019,0.062,0.467,-0.406,-0.942,0.094 | 0.017,0.021,-0.984,1.116,1.090,0.787 | 0.061,0.071,-0.406,-0.144,1.539,1.324 |
| **d564a**762d2e5e7484bfa15f3b30d72d6 | 0.084,0.071,1.955,-1.534,-0.552,-1.218 | 0.043,0.069,0.377,-0.067,-0.804,-1.338 | 0.098,0.024,-1.922,1.130,0.525,1.284 | 0.078,0.068,-0.473,-0.021,1.985,0.285 |
| **d5652**5c4961ea506adbc7842bfdcbbb9 | 0.070,0.037,1.172,-1.147,-0.855,0.316 | 0.077,0.046,0.760,-0.629,0.271,-0.723 | 0.047,0.076,-0.674,1.257,-0.614,-0.376 | 0.033,0.064,-0.330,0.141,-0.117,0.894 |
| **d566a**1a0d5da2548390ca9739449654a | 0.066,0.089,0.458,-0.263,-1.645,-0.119 | 0.041,0.096,1.030,-0.841,-1.046,-0.717 | 0.048,0.049,-1.918,1.969,-1.832,1.407 | 0.088,0.083,-1.406,0.776,1.441,1.395 |
| **d567e**0676e13738c6059ee53a2cab17b | 0.058,0.079,1.279,-1.521,0.294,-0.196 | 0.024,0.070,0.943,-0.584,0.881,-1.820 | 0.053,0.062,-1.337,1.743,-0.236,0.885 | 0.094,0.057,-0.674,0.085,1.723,1.746 |
| **d57a3**eb8863e3e69a33b9cf6b78840eb | 0.055,0.011,0.382,-0.319,-1.262,1.192 | 0.027,0.046,0.618,-0.790,-1.058,-0.021 | 0.081,0.034,-1.924,1.290,1.898,1.636 | 0.077,0.053,-0.577,0.373,1.765,0.891 |
| **d58ff**d69e51d1422d2873f022fa97549 | 0.048,0.012,1.806,-1.806,-1.826,-0.019 | 0.095,0.050,1.693,-0.815,-0.654,-0.554 | 0.082,0.062,-0.846,1.883,0.828,-0.037 | 0.072,0.079,-0.734,0.619,-0.539,1.164 |
| **d6314**7a936c84bb717e0d035682f25a7 | 0.055,0.054,1.714,-1.858,0.314,-0.743 | 0.070,0.013,1.870,-1.805,-0.888,0.806 | 0.016,0.030,-1.578,1.789,1.948,0.955 | 0.021,0.069,-0.459,0.097,1.259,1.215 |
| **d6535**882de73a438285929f82c1579e4 | 0.062,0.038,1.615,-1.164,-1.784,-0.749 | 0.029,0.089,1.083,-1.548,0.330,-0.753 | 0.099,0.069,-1.489,0.614,1.314,1.751 | 0.092,0.062,-0.404,0.068,1.527,0.152 |
| **d65ed**d49b7902caec51b8d558c5c0fe3 | 0.048,0.025,1.461,-1.682,-0.263,-0.206 | 0.078,0.056,0.484,-0.471,-1.582,0.025 | 0.099,0.097,-1.127,0.753,1.795,1.289 | 0.082,0.060,-0.583,0.448,0.582,0.986 |
| **d6981**c0bc5d3f65ad7d0517306374c83 | 0.045,0.067,1.842,-1.994,-1.806,-0.483 | 0.065,0.012,0.476,-0.390,-1.267,-0.925 | 0.099,0.033,-1.386,0.795,1.769,0.601 | 0.083,0.070,-1.667,1.207,1.993,1.631 |
| **d6c4a**ead2693fd8cd6d5f4f4f77a33e1 | 0.076,0.057,1.916,-1.725,-0.988,-1.048 | 0.076,0.059,0.691,-0.596,-1.706,1.352 | 0.075,0.079,-1.284,1.036,1.265,0.650 | 0.060,0.069,-1.341,0.613,1.530,1.979 |
| **d6dc4**f4524d6d9117a679d28efcaafac | 0.090,0.080,0.767,-0.753,-0.210,-0.674 | 0.034,0.050,0.637,-0.171,-0.329,-1.060 | 0.097,0.077,-0.784,1.554,1.850,0.257 | 0.033,0.097,-1.251,0.677,-0.303,1.763 |
| **d7039**f16e2e1ea0c6d8f6c4373f99fa3 | 0.045,0.072,1.520,-1.740,-0.678,0.105 | 0.039,0.091,1.479,-0.934,-0.705,-0.304 | 0.099,0.066,-1.552,1.951,1.728,0.785 | 0.084,0.098,-1.478,0.838,-0.356,1.908 |
| **d73d5**68ebc5a0e97b09bbe42f61a3399 | 0.080,0.076,1.899,-1.935,-1.303,-1.918 | 0.038,0.053,1.340,-1.147,-1.437,-0.081 | 0.040,0.055,-1.175,1.336,-0.310,1.506 | 0.013,0.042,-1.223,1.102,-0.862,1.517 |
| **d79f7**b546a8260cf23937448f941f90f | 0.095,0.056,1.190,-0.867,-1.082,-0.938 | 0.069,0.015,1.261,-1.799,-1.511,-0.602 | 0.086,0.081,-1.309,0.936,1.639,0.458 | 0.056,0.099,-0.253,-0.093,1.547,1.198 |
| **d7a29**276524f9e423ebaedd06e0d3eeb | 0.077,0.086,1.885,-1.528,-1.852,0.140 | 0.037,0.013,1.915,-1.615,-0.324,-0.851 | 0.057,0.051,-1.670,1.967,1.311,-0.435 | 0.029,0.017,-1.675,1.588,1.211,0.374 |
| **d7c2e**cd8bf68b9004d034571370b5183 | 0.052,0.055,1.048,-1.097,-0.220,-0.943 | 0.040,0.017,0.805,-0.434,-1.369,-0.136 | 0.052,0.060,-0.432,0.598,-0.251,0.599 | 0.089,0.052,-1.049,0.631,0.656,1.478 |
| **d7cfd**f50dfbc7ad3b731b3bf0bece956 | 0.091,0.037,1.420,-1.314,-1.793,0.462 | 0.014,0.086,0.590,-0.442,0.099,-0.372 | 0.049,0.034,-0.832,1.144,-0.730,1.361 | 0.042,0.046,-0.613,0.390,1.729,0.648 |
| **d7d2e**7a56590659b7291cab8bb894427 | 0.099,0.053,1.761,-1.614,-1.187,-0.798 | 0.030,0.087,1.426,-0.969,-1.296,-1.081 | 0.050,0.057,-1.600,1.991,1.686,-1.492 | 0.033,0.068,-1.094,0.659,0.550,1.845 |
| **d7e9d**e3d59a1ac4bdebc95e4d8f19934 | 0.096,0.019,1.933,-1.493,-0.350,-1.150 | 0.016,0.077,0.448,-0.670,-0.812,-1.142 | 0.024,0.043,-1.371,0.267,1.413,0.705 | 0.042,0.015,-0.237,0.177,1.172,-1.022 |
| **d7f85**ad92d8551a1f21becb24f37ae7d | 0.069,0.054,1.047,-0.924,-1.442,0.804 | 0.031,0.081,1.952,-1.013,-0.238,-1.838 | 0.046,0.012,-1.332,1.985,1.472,-1.571 | 0.026,0.093,-1.240,0.766,1.376,1.127 |
| **d81f4**4626b79f0e7d7b30f70eb06f20b | 0.054,0.020,1.495,-1.845,-1.233,-1.086 | 0.087,0.058,1.215,-0.765,-1.890,0.002 | 0.024,0.065,-1.068,0.661,1.878,0.974 | 0.072,0.013,-1.218,0.958,0.997,1.524 |
| **d8552**81e5ec2790673beb8604670b3e3 | 0.072,0.033,1.614,-1.690,-0.564,-0.354 | 0.027,0.033,0.649,-0.460,-0.402,-0.399 | 0.075,0.088,-0.923,0.553,1.620,1.020 | 0.092,0.035,-0.504,0.218,0.275,1.882 |
| **d863f**1a63e300c6b8780256150cb7a32 | 0.030,0.016,1.372,-1.606,-0.626,-0.731 | 0.014,0.051,1.130,-1.127,-1.935,0.719 | 0.064,0.016,-1.475,1.618,1.398,1.476 | 0.048,0.080,-1.033,0.773,1.809,0.493 |
| **d8750**50eae9b8c206bec8488eb6b0e40 | 0.048,0.026,1.954,-1.876,-1.969,-0.281 | 0.091,0.087,1.205,-0.993,-1.020,-1.018 | 0.054,0.036,-1.143,0.844,0.818,-0.081 | 0.084,0.066,-1.316,1.013,1.914,-0.039 |

| | | | | |
|---|---|---|---|---|
| **d8759**511d857723b9aea8f70bbc8410a | 0.078,0.043,0.725,-0.634,-0.928,0.074 | 0.036,0.088,0.797,-0.823,0.234,-0.811 | 0.091,0.042,-1.779,1.240,0.820,1.154 | 0.066,0.080,-1.139,0.575,1.629,1.522 |
| **d8e39**3421472f483c65705931e67f7e8 | 0.096,0.019,1.938,-1.824,-1.811,0.266 | 0.019,0.049,0.880,-0.944,-1.129,-0.612 | 0.069,0.065,-1.548,1.420,0.818,0.213 | 0.097,0.045,-0.953,0.529,1.804,0.736 |
| **d8e3d**4b4c56e5eaae61542ef9ec4b39b | 0.090,0.041,1.533,-1.659,-1.413,-0.223 | 0.078,0.044,0.689,-0.635,-0.044,1.075 | 0.083,0.087,-1.284,1.870,1.669,-0.650 | 0.047,0.067,-0.474,0.004,1.981,1.774 |
| **d8fe8**b96c4ad5741bd50f1a43a5a15fe | 0.050,0.039,1.937,-1.811,-1.006,-0.403 | 0.072,0.091,0.808,-1.931,0.195,-0.142 | 0.047,0.088,-0.891,0.212,1.235,0.494 | 0.013,0.078,-1.126,0.715,1.667,1.871 |
| **d9027**79032b4e0fbe33f49b7eea52adc | 0.049,0.030,1.075,-1.092,-1.879,0.518 | 0.075,0.022,0.977,-0.943,1.109,0.824 | 0.088,0.041,-0.813,0.964,0.551,1.072 | 0.047,0.039,-0.942,0.678,0.025,1.378 |
| **d92a6**1e9c141859859562eb2934734d9 | 0.050,0.057,1.943,-1.872,-0.346,-0.561 | 0.065,0.042,0.919,-1.509,0.456,-1.180 | 0.034,0.098,-1.115,0.898,1.277,0.922 | 0.049,0.039,-0.274,0.142,1.760,0.546 |
| **d996a**9d068b244ad4aecad9f9cf47daa | 0.097,0.028,1.488,-1.275,-1.900,-0.494 | 0.021,0.077,0.241,-0.283,-1.567,-0.130 | 0.042,0.053,-1.166,0.804,-0.368,1.589 | 0.098,0.078,-1.865,1.731,-0.495,1.697 |
| **d9e40**10b4f4d1f9b7a5046adbbdcd81e | 0.066,0.052,1.382,-1.664,-0.810,-0.347 | 0.080,0.049,0.643,-0.343,-1.024,-0.418 | 0.058,0.053,-0.986,1.946,1.804,0.623 | 0.064,0.082,-1.399,1.308,1.948,-0.433 |
| **da15b**6c4cf8c4681f131c9aa7c533657 | 0.064,0.065,1.712,-1.796,-0.757,-0.912 | 0.078,0.078,0.308,-0.660,0.887,0.152 | 0.065,0.027,-1.101,0.724,1.697,1.792 | 0.026,0.060,-0.889,0.690,0.920,1.442 |
| **da3ce**df2df7e4b172cd764faf39e97e5 | 0.041,0.058,1.347,-0.901,-1.150,-0.275 | 0.053,0.032,0.370,-0.373,-1.875,-0.075 | 0.047,0.032,-0.354,0.101,-0.237,0.611 | 0.053,0.029,-0.140,-0.317,1.923,1.877 |
| **da4b6**03348700bbba0faffb1fab6dd80 | 0.056,0.038,1.083,-0.916,-1.584,-0.893 | 0.090,0.045,0.991,-0.686,-1.225,0.148 | 0.031,0.038,-1.549,1.766,0.839,0.729 | 0.087,0.098,-1.310,0.892,1.353,0.820 |
| **da7f1**03bccffa3f9de51597d9cf49050 | 0.052,0.099,1.147,-0.937,-0.241,-1.950 | 0.033,0.081,1.205,-0.988,0.246,-1.550 | 0.046,0.059,-1.830,1.961,0.975,1.526 | 0.058,0.060,-0.698,0.435,1.496,1.197 |
| **da87d**03f10572e043a2b694fae49d859 | 0.086,0.048,0.798,-0.746,-0.888,0.600 | 0.047,0.047,0.681,-0.596,-1.292,0.032 | 0.036,0.078,-1.169,1.232,-0.287,1.041 | 0.054,0.058,-0.544,0.282,0.224,1.356 |
| **da9c8**b398c201531a5df7e1588422840 | 0.092,0.082,1.675,-0.862,-1.703,-0.587 | 0.074,0.072,1.127,-1.703,-0.908,-0.903 | 0.038,0.036,-1.963,0.790,1.819,1.406 | 0.062,0.087,-0.164,-0.184,1.925,0.284 |
| **daca9**fde1f4ca2968e3f779da26d96c8 | 0.015,0.046,0.656,-0.740,-0.804,0.047 | 0.052,0.021,0.265,-0.287,0.072,-1.883 | 0.100,0.053,-1.833,1.256,0.595,1.234 | 0.092,0.083,-1.770,1.585,1.511,-0.396 |
| **db042**96546bd64322e0d509393fefb89 | 0.090,0.039,1.060,-0.527,-1.285,0.003 | 0.024,0.071,0.573,-1.085,-1.468,-0.536 | 0.056,0.019,-1.890,0.808,1.096,0.766 | 0.018,0.094,-1.208,0.843,1.504,1.183 |
| **db124**9ada2a3d8d41f8207ef4e24d5ba | 0.074,0.024,1.086,-1.002,-1.610,0.019 | 0.095,0.089,0.837,-1.829,0.371,-0.184 | 0.065,0.033,-1.908,0.945,1.359,1.881 | 0.096,0.070,-0.911,0.550,1.719,0.817 |
| **db206**94f58c76913b6223c6d5b60c9d3 | 0.019,0.036,1.609,-1.668,0.320,-1.808 | 0.018,0.046,1.498,-1.023,-0.997,-0.132 | 0.095,0.022,-1.818,1.918,1.867,1.953 | 0.066,0.024,-0.352,-0.092,1.873,-0.122 |
| **db3b2**e498ec83fdc22ff2e2b1edbdd13 | 0.081,0.056,1.670,-1.585,-0.049,-0.984 | 0.039,0.053,1.544,-0.719,-1.704,-0.490 | 0.031,0.094,-1.033,1.954,-0.181,0.480 | 0.011,0.091,-0.642,0.595,0.129,0.224 |
| **db5cf**15f29464a70449d632c78971ee5 | 0.070,0.029,1.392,-1.274,-1.625,-0.069 | 0.014,0.016,1.030,-0.587,-0.155,0.936 | 0.088,0.093,-0.457,0.479,1.430,1.498 | 0.088,0.072,-0.797,0.422,0.965,1.603 |
| **dba74**d2c4d00cd01a4ee14a823dc7d74 | 0.052,0.044,1.676,-1.587,-0.690,-0.970 | 0.098,0.057,0.947,-0.603,-1.778,-0.936 | 0.047,0.098,-1.961,0.940,1.314,0.869 | 0.090,0.086,-0.847,0.370,1.883,0.081 |
| **dba9d**e6cff6c8e9ab4f882fbb477f781 | 0.090,0.021,0.879,-0.632,-1.549,1.061 | 0.030,0.081,1.432,-1.198,-1.930,-1.043 | 0.035,0.089,-1.849,1.908,1.789,0.228 | 0.097,0.036,-1.024,0.624,0.911,1.868 |
| **dbc9f**c037f7294f071c84ed8068f21ee | 0.054,0.014,1.674,-1.892,-0.773,-0.136 | 0.063,0.043,0.753,-1.490,1.610,-0.965 | 0.070,0.061,-1.725,1.047,1.076,1.171 | 0.037,0.087,-0.581,0.191,1.497,1.784 |
| **dbd81**64caa0a94a38b28b2ef34f31ae1 | 0.075,0.029,1.339,-1.146,-1.498,0.239 | 0.060,0.053,0.857,-1.470,-1.889,0.073 | 0.030,0.058,-1.803,0.897,1.579,-1.014 | 0.038,0.082,-0.773,0.538,1.755,0.736 |
| **dbdbb**8e03cc36f84847afc921e350d2f | 0.048,0.044,1.541,-1.560,-0.265,-0.982 | 0.013,0.077,0.662,-1.476,1.793,-0.114 | 0.082,0.097,-1.751,0.647,1.141,0.929 | 0.088,0.070,-1.009,0.977,1.374,-1.161 |
| **dc317**ba6e6730a9a28d9207c7f1a8c1a | 0.025,0.073,0.690,-0.668,-0.849,0.139 | 0.019,0.029,1.257,-1.693,-0.522,-1.784 | 0.014,0.056,-1.770,1.675,1.444,-1.453 | 0.041,0.063,-1.048,0.777,1.636,1.485 |
| **dc3fd**d266592b856673684707b47a16b | 0.042,0.066,0.735,-0.775,-1.663,-0.263 | 0.026,0.033,0.235,-0.271,-0.511,-0.677 | 0.054,0.059,-1.059,0.306,1.610,1.993 | 0.051,0.051,-0.044,-0.186,1.932,0.948 |
| **dc48b**3cbf361171445eacc71607460ea | 0.047,0.081,1.882,-1.620,-1.420,-1.379 | 0.079,0.086,1.433,-1.165,0.109,-1.608 | 0.099,0.070,-1.803,1.669,1.871,1.454 | 0.083,0.073,-0.209,-0.058,1.744,0.276 |
| **dc4d2**afeb73c80becfac21af8d36c3a9 | 0.074,0.073,1.658,-1.683,-1.668,-0.835 | 0.029,0.057,0.374,-0.330,-1.951,0.002 | 0.013,0.028,-0.700,0.688,1.242,-0.722 | 0.063,0.093,-0.536,0.204,1.003,1.073 |
| **dc573**f2a094b08ee97b8ee4b1e7a4602 | 0.065,0.035,1.912,-1.485,-1.934,-0.311 | 0.076,0.081,0.997,-1.478,-0.923,-0.950 | 0.067,0.045,-0.554,0.161,1.979,-1.607 | 0.043,0.065,-1.195,0.742,1.871,1.893 |
| **dce62**13ee214382fa0b5e987f758f7e8 | 0.091,0.098,1.859,-1.620,-0.711,-1.814 | 0.084,0.035,0.804,-0.407,-1.572,-1.112 | 0.098,0.057,-0.543,0.281,1.838,0.162 | 0.028,0.027,-0.633,0.590,1.586,-0.027 |
| **dd269**571c0bd9d8145ae1e58bc3379f7 | 0.054,0.063,1.460,-1.483,-0.498,-0.660 | 0.041,0.073,1.061,-0.617,-0.520,-1.524 | 0.013,0.097,-0.797,1.009,1.098,1.536 | 0.097,0.055,-1.336,0.990,1.781,0.441 |
| **dd338**ce1d18f230fb0f128d7f4871d0b | 0.098,0.044,1.455,-1.040,-1.851,-0.264 | 0.080,0.051,0.505,-0.621,-0.353,1.058 | 0.092,0.097,-0.500,-0.004,1.152,1.234 | 0.024,0.081,-0.426,0.021,1.407,1.922 |
| **dd68d**ef53316c2d6dc61d060dd4e491c | 0.086,0.041,1.378,-1.392,-0.790,-0.769 | 0.014,0.035,0.481,-0.677,-0.840,-0.453 | 0.095,0.089,-0.712,0.127,1.746,0.129 | 0.077,0.076,-0.773,0.310,0.727,1.709 |
| **dd850**9509b32775013e9b158896e37d3 | 0.056,0.066,1.794,-1.036,-1.922,-0.412 | 0.072,0.030,0.300,-0.336,-1.015,-0.959 | 0.085,0.013,-1.842,0.388,1.487,1.973 | 0.096,0.082,-0.490,0.027,1.892,0.873 |
| **dd8eb**6c10257725cae4563626f3466ac | 0.097,0.061,0.594,-0.705,-0.375,-0.859 | 0.022,0.051,0.340,-0.177,-1.280,0.790 | 0.084,0.073,-0.599,0.668,1.877,1.852 | 0.020,0.027,-1.387,1.256,-0.399,1.969 |
| **dd974**65eafef1cd3f18bc2e93b275856 | 0.081,0.045,1.671,-1.254,-1.851,-0.967 | 0.028,0.050,1.825,-1.347,1.253,-1.857 | 0.070,0.076,-1.820,1.981,1.028,1.508 | 0.072,0.099,-0.696,-0.083,0.991,1.623 |
| **ddd7e**c5c2a39e9a0a99394b16fa2bbbc | 0.076,0.030,1.540,-1.345,-1.522,-0.376 | 0.033,0.023,1.099,-1.903,-0.920,0.816 | 0.089,0.041,-0.640,0.329,-0.002,0.528 | 0.015,0.071,-0.343,-0.116,1.532,1.831 |
| **dde53**3a4f734b53127d8f60ecbf3ca9a | 0.022,0.024,1.167,-0.935,-0.936,-0.880 | 0.033,0.068,0.625,-0.988,-1.311,-1.804 | 0.084,0.028,-1.995,0.432,1.942,-1.118 | 0.085,0.054,-1.540,1.042,1.999,1.485 |
| **de070**69686bc9b39ae1222f20ff051a8 | 0.077,0.076,1.650,-1.372,-0.824,-1.186 | 0.061,0.088,0.907,-0.607,-0.561,-1.865 | 0.061,0.096,-1.855,1.095,1.889,1.735 | 0.096,0.100,-0.128,-0.069,1.842,-0.615 |
| **de08d**a12d866a8d9d5d814f94fac03d7 | 0.077,0.027,0.571,-0.057,-1.162,-0.936 | 0.019,0.083,0.112,-0.096,-0.495,-0.227 | 0.097,0.062,-1.521,0.823,0.459,-0.325 | 0.055,0.093,-0.982,0.462,1.007,1.440 |
| **de0d6**5cd194cf78d1b153961233c7cfb | 0.078,0.020,1.915,-1.949,-1.578,0.590 | 0.039,0.075,0.527,-0.568,-0.974,-0.089 | 0.058,0.088,-0.698,0.350,1.907,0.212 | 0.098,0.072,-0.386,-0.008,1.417,0.724 |
| **de469**f3328856ad5cf19a9f95f9e874b | 0.053,0.048,1.511,-1.594,-0.945,-0.583 | 0.094,0.024,0.739,-1.234,-0.479,-0.737 | 0.014,0.012,-1.507,0.884,-0.315,1.552 | 0.013,0.017,-0.169,0.146,-1.161,0.788 |
| **de781**573b0a3a4114b9abaa3865a486c | 0.065,0.095,1.651,-1.202,-1.692,-1.068 | 0.018,0.089,1.424,-0.964,0.398,-1.543 | 0.053,0.062,-1.125,1.026,1.633,1.153 | 0.091,0.097,-1.016,0.646,0.728,0.726 |
| **de7c3**17aec74d595c96cc6c868f7277e | 0.089,0.050,1.854,-1.839,0.489,-1.214 | 0.037,0.063,0.503,-0.345,-0.713,-0.923 | 0.059,0.032,-0.994,0.649,1.475,0.360 | 0.049,0.054,-0.618,0.340,1.940,0.790 |
| **deb03**11e38ec4f963d727eed107d4204 | 0.081,0.042,0.456,-0.054,-1.141,-1.771 | 0.095,0.087,0.276,-0.272,-1.057,1.246 | 0.095,0.090,-1.870,1.910,1.122,1.593 | 0.017,0.023,-0.518,0.443,1.488,0.320 |
| **deb5b**6d156ee9f9435eda9b2db966318 | 0.063,0.077,1.207,-1.213,-0.164,-0.170 | 0.059,0.045,0.419,-0.562,-1.056,-1.070 | 0.076,0.015,-1.678,0.450,1.868,1.972 | 0.091,0.079,-1.440,1.040,1.909,0.351 |
| **deef5**8e83eb571bf87e364a25b7f4bc4 | 0.083,0.088,1.067,-1.088,0.372,-1.072 | 0.085,0.092,0.673,-0.475,1.281,-0.019 | 0.097,0.059,-0.921,1.854,1.151,0.937 | 0.046,0.082,-0.817,0.374,1.947,1.218 |
| **df263**ffc7895ae05454cf424f322cb69 | 0.053,0.022,1.267,-1.327,-0.405,-0.143 | 0.019,0.082,0.412,-0.222,-1.288,-1.317 | 0.059,0.064,-1.344,0.522,1.405,0.542 | 0.076,0.048,-1.059,0.800,1.904,-0.265 |
| **df6a3**fa15977081cbc5cfe00c2582d55 | 0.051,0.051,1.750,-1.596,-0.176,-0.501 | 0.068,0.081,0.522,-0.127,-1.271,-1.210 | 0.072,0.062,-0.525,0.027,0.114,0.766 | 0.081,0.067,-0.290,-0.080,1.217,0.729 |
| **df6da**9ddcb224b56cec65326885505ba | 0.075,0.014,1.687,-1.711,-1.930,-0.867 | 0.018,0.065,0.342,-0.326,-1.489,-0.009 | 0.068,0.091,-1.134,0.759,1.444,0.991 | 0.067,0.024,-0.481,0.258,1.541,0.746 |
| **df783**2efec28217983eac92e9d14b462 | 0.098,0.056,1.695,-1.412,-1.162,-0.094 | 0.012,0.089,1.138,-0.670,-0.819,-1.267 | 0.098,0.058,-1.959,1.922,1.528,1.310 | 0.097,0.080,-0.526,0.081,1.370,0.199 |
| **df8c5**2a84cb7438a3a91c184309c2484 | 0.038,0.053,1.532,-1.428,-0.198,-0.571 | 0.096,0.032,0.627,-1.386,-0.499,-1.269 | 0.097,0.036,-1.900,0.279,1.604,1.963 | 0.098,0.075,-1.218,1.055,0.750,0.321 |
| **df8fd**04924eadf37085e82bfb712e50b | 0.076,0.050,1.819,-0.886,-0.830,-1.941 | 0.037,0.045,0.669,-0.778,-1.642,-1.229 | 0.089,0.086,-1.706,0.425,1.169,0.407 | 0.054,0.086,-0.421,-0.079,1.933,0.970 |
| **df9e3**c2d55fe8b912260d560650bf906 | 0.075,0.048,1.751,-1.656,-1.796,-1.031 | 0.050,0.035,1.026,-0.597,-1.254,0.702 | 0.012,0.039,-1.261,1.972,0.166,0.554 | 0.016,0.075,-1.196,1.009,1.364,0.359 |
| **dfd55**cfec930d91c8272ccf44d93ef45 | 0.067,0.026,1.460,-1.751,-1.031,0.302 | 0.070,0.040,0.600,-0.620,-0.914,-0.794 | 0.073,0.031,-1.602,0.736,1.320,0.752 | 0.097,0.060,-0.422,-0.064,1.894,0.673 |
| **dfd8b**a4b72e7252002140f95208b9f0b | 0.075,0.046,1.099,-0.918,-0.737,-0.300 | 0.086,0.035,0.858,-1.372,-1.564,-0.539 | 0.082,0.035,-1.378,0.179,1.867,1.903 | 0.087,0.045,-0.578,0.221,1.787,1.227 |
| **dff94**6e1cec758ae1e02c825d17bc52b | 0.084,0.030,1.794,-1.634,-0.904,-1.092 | 0.076,0.017,1.255,-1.283,-1.896,-0.463 | 0.079,0.034,-0.811,0.309,1.833,0.035 | 0.044,0.063,-0.576,0.312,1.151,1.098 |
| **e00dc**2897ecca8b8ea8b9bb7d28907c9 | 0.091,0.068,1.534,-1.760,-0.810,-0.783 | 0.043,0.037,0.157,-0.138,-0.753,0.178 | 0.058,0.096,-0.630,0.773,-0.332,-1.030 | 0.011,0.027,-1.084,1.002,-0.713,1.794 |
| **e018d**f5a1072cdb1e50b8c7c764cb911 | 0.057,0.027,1.986,-1.742,-1.730,-0.600 | 0.062,0.072,1.113,-1.504,-0.562,-1.149 | 0.013,0.070,-1.978,0.914,1.869,1.724 | 0.099,0.044,-1.187,0.683,1.366,1.845 |
| **e0404**cbee2208c268f6a49e36b162e42 | 0.046,0.020,1.909,-1.261,-1.718,-1.159 | 0.027,0.093,0.457,-0.429,-1.205,-1.237 | 0.097,0.092,-1.746,0.495,1.916,-1.912 | 0.082,0.077,-0.148,-0.523,1.867,1.496 |
| **e046a**f3ac8bd2eae95185912323870b7 | 0.048,0.040,1.813,-1.417,-1.406,-1.156 | 0.094,0.060,1.263,-1.447,-1.864,-0.631 | 0.081,0.071,-1.617,0.745,0.547,1.000 | 0.055,0.077,-0.379,0.330,1.962,-0.752 |
| **e058b**52b7d5c74c51dbc248f2171a4c5 | 0.098,0.055,1.979,-1.648,-1.976,0.810 | 0.021,0.023,0.970,-1.894,-1.355,-0.364 | 0.012,0.038,-1.321,0.677,1.487,0.387 | 0.038,0.059,-0.259,0.013,1.705,1.041 |
| **e05fa**6d9fda7b5799079be172064fb98 | 0.097,0.016,1.827,-1.494,-1.116,0.438 | 0.013,0.014,1.956,-1.363,-0.282,1.104 | 0.024,0.083,-1.404,1.106,1.283,1.968 | 0.051,0.078,-1.256,0.484,1.852,0.701 |
| **e0723**48186975ea8a2bebfb0a39c1cb1 | 0.073,0.039,1.121,-0.982,-1.595,-0.113 | 0.061,0.042,0.903,-0.958,1.020,-1.610 | 0.093,0.096,-1.689,1.693,-1.207,1.750 | 0.068,0.037,-1.197,0.774,1.676,1.458 |
| **e074b**60a51fde61f6203c059b3cd0302 | 0.047,0.075,1.608,-1.471,-1.824,0.141 | 0.034,0.052,1.843,-1.306,-0.909,-0.891 | 0.029,0.037,-1.476,1.582,1.177,0.494 | 0.087,0.026,-0.554,0.410,-0.295,1.784 |
| **e090f**2ab44c287ad48d94da38bf47559 | 0.057,0.047,1.027,-1.001,-1.813,1.340 | 0.026,0.032,0.179,-0.649,1.804,-0.959 | 0.026,0.061,-1.806,0.785,0.984,-0.039 | 0.083,0.093,-1.815,1.310,1.085,0.947 |
| **e0b7b**005191baf527bf42ecbd6adce84 | 0.049,0.047,1.074,-1.158,-1.660,-0.379 | 0.043,0.077,0.649,-0.536,-0.984,-0.939 | 0.099,0.060,-0.726,0.578,1.613,0.188 | 0.085,0.061,-1.456,1.204,0.947,1.519 |

| | | | | |
|---|---|---|---|---|
| **e0b7b**b643a1bdde958b369fa33d24e8e | 0.049,0.018,1.582,-1.845,-1.612,0.938 | 0.094,0.090,0.646,-0.376,-0.507,-1.133 | 0.084,0.063,-1.693,1.875,1.942,0.704 | 0.015,0.078,-1.565,1.359,1.543,1.363 |
| **e0b89**d33009c5fa2901befa8133355f1 | 0.048,0.042,1.785,-1.799,-1.917,-0.225 | 0.047,0.097,1.054,-1.069,-1.194,-0.224 | 0.041,0.027,-1.082,0.680,1.709,0.256 | 0.051,0.075,-1.049,0.447,1.969,1.625 |
| **e0f2c**17c45c42653b9aef4c3b02140f9 | 0.035,0.087,1.349,-0.901,-1.816,-0.670 | 0.067,0.042,0.904,-1.288,0.867,-1.211 | 0.099,0.053,-1.498,0.371,1.576,1.813 | 0.041,0.014,-0.824,0.537,1.814,1.595 |
| **e106f**860bb5923c78733de3f1a4dfb5e | 0.034,0.055,0.644,-0.692,0.046,-0.545 | 0.092,0.072,1.907,-1.043,-1.909,1.202 | 0.031,0.048,-0.874,0.871,1.124,1.979 | 0.027,0.044,-0.762,0.398,1.797,1.453 |
| **e13f6**d06258c21b63ac1a002fd89fc90 | 0.078,0.028,1.809,-1.514,-1.797,0.485 | 0.098,0.020,0.513,-0.621,-0.682,-1.388 | 0.014,0.089,-0.875,0.368,0.160,1.155 | 0.036,0.030,-0.173,0.014,1.500,1.226 |
| **e1c64**a23096a676bf912809a3eef67c8 | 0.088,0.061,1.796,-1.873,-1.549,1.020 | 0.050,0.088,0.490,-0.601,-0.639,-0.433 | 0.037,0.099,-0.983,0.465,-0.477,0.914 | 0.030,0.069,-1.608,1.245,0.604,1.876 |
| **e1cac**aa33cdff29d06bd73f085ff8174 | 0.013,0.056,1.319,-1.181,-0.887,-1.323 | 0.031,0.035,0.370,-0.566,-1.231,-0.189 | 0.057,0.063,-1.721,0.870,1.265,0.768 | 0.071,0.057,-0.298,-0.141,1.478,1.539 |
| **e1f29**2ede30b4c7d4f82e134985d1948 | 0.052,0.076,1.523,-1.360,-0.122,-1.612 | 0.066,0.062,0.224,-0.169,-1.860,1.051 | 0.090,0.054,-0.625,0.269,1.320,-0.242 | 0.069,0.086,-0.704,0.317,1.763,0.739 |
| **e2123**ba7dc227ebf5adfd5ab2c30fbec | 0.092,0.063,1.846,-1.606,-1.567,0.430 | 0.080,0.017,1.961,-1.241,-0.545,0.512 | 0.029,0.030,-1.245,1.877,1.675,1.941 | 0.018,0.068,-0.779,0.406,1.919,1.313 |
| **e231b**c3c31a9d9f2816bec19468d8357 | 0.042,0.064,1.716,-1.544,-1.405,-0.966 | 0.072,0.062,1.445,-1.883,-1.958,-1.159 | 0.057,0.061,-1.821,1.132,-0.592,0.546 | 0.017,0.063,-0.804,0.621,-0.425,1.524 |
| **e2457**b8dddd5dc8b2b78a304c747389f | 0.043,0.077,1.456,-1.485,-0.425,-0.887 | 0.019,0.098,0.672,-0.528,0.231,-1.874 | 0.062,0.063,-1.244,0.830,0.097,1.529 | 0.071,0.043,-0.214,0.113,1.947,-1.117 |
| **e248c**2c8ad10f2a21b97011ec460ee80 | 0.046,0.048,1.044,-1.003,-1.859,0.238 | 0.092,0.071,0.945,-0.420,-1.043,-1.767 | 0.076,0.073,-1.244,0.972,-0.366,1.111 | 0.050,0.025,-0.891,0.813,0.072,0.927 |
| **e2723**95423e59fda45021f9739e568be | 0.076,0.047,1.780,-1.519,-0.559,-1.157 | 0.098,0.011,1.336,-1.743,-1.890,-1.418 | 0.017,0.090,-1.750,0.739,-0.030,1.812 | 0.080,0.012,-0.453,0.333,0.683,-0.545 |
| **e374c**013a2385a2ae10f28a5588ed37f | 0.085,0.020,1.755,-1.365,-1.728,-0.751 | 0.057,0.091,1.008,-1.955,1.869,-1.477 | 0.074,0.062,-1.950,1.137,0.970,1.013 | 0.088,0.074,-1.793,1.382,1.866,1.366 |
| **e3a50**10e8cb9152318da2b21bf8216d7 | 0.087,0.014,1.675,-1.345,-1.284,-0.921 | 0.063,0.079,0.939,-1.428,-1.804,-0.458 | 0.040,0.014,-1.600,0.630,0.182,1.912 | 0.096,0.066,-1.311,0.676,1.370,1.862 |
| **e402e**3672e665522d95ebbf838f62262 | 0.087,0.051,1.171,-0.950,-1.488,-0.020 | 0.031,0.087,0.790,-1.304,0.007,-0.198 | 0.056,0.080,-1.829,1.423,1.896,0.944 | 0.058,0.097,-1.320,0.761,1.746,1.871 |
| **e40a9**6c6d1f70502ac182fa58278b279 | 0.031,0.021,1.592,-1.620,-1.834,-0.457 | 0.093,0.082,1.008,-1.763,-0.871,-0.234 | 0.094,0.018,-1.826,0.626,1.731,1.926 | 0.098,0.087,-0.244,0.012,1.802,-0.417 |
| **e4154**5cbb23e206edbffacf7fc6fd428 | 0.047,0.023,1.654,-1.572,-1.759,-0.295 | 0.091,0.054,1.302,-1.161,-0.230,-1.152 | 0.060,0.081,-1.134,0.934,1.496,0.198 | 0.083,0.043,-0.329,0.069,1.879,-0.422 |
| **e44f0**920c9fe7156283cd031a3d89439 | 0.052,0.065,1.884,-1.641,-1.084,-0.214 | 0.040,0.066,1.911,-1.840,-0.605,-0.920 | 0.021,0.049,-1.812,1.846,1.255,0.126 | 0.098,0.073,-0.310,-0.295,1.833,-0.369 |
| **e4835**f6f5efeef45dec95a8c64f7dd5f | 0.070,0.057,1.965,-1.039,-1.786,-1.317 | 0.053,0.085,0.327,-0.150,-1.159,-0.989 | 0.072,0.055,-1.877,0.400,0.840,1.874 | 0.064,0.047,-0.128,-0.281,1.674,1.924 |
| **e484c**c1f3fdbb97114fa9532ae4f5c01 | 0.073,0.019,1.950,-1.825,-1.712,-0.018 | 0.024,0.082,0.884,-0.927,-0.132,-0.856 | 0.019,0.090,-1.885,1.953,-0.475,0.752 | 0.053,0.068,-1.909,1.642,1.555,1.192 |
| **e4a28**a909f22743043368910c21a8974 | 0.089,0.019,1.413,-1.634,-0.860,-0.180 | 0.076,0.052,0.966,-0.678,-0.783,-1.419 | 0.035,0.085,-1.055,1.277,-0.572,-0.473 | 0.083,0.041,-0.744,0.241,1.925,1.694 |
| **e4aac**aab192946176de1480a917d9211 | 0.028,0.078,1.142,-0.857,-0.725,-1.720 | 0.045,0.063,0.326,-0.674,1.315,0.296 | 0.040,0.086,-1.811,1.565,-0.292,1.454 | 0.076,0.041,-0.942,0.785,0.085,1.137 |
| **e4f9b**e67da27e2d3fc8f489c0542aa1e | 0.041,0.061,1.430,-1.574,-1.806,0.312 | 0.057,0.088,0.462,-0.957,0.169,0.608 | 0.059,0.045,-1.427,0.964,1.592,-0.463 | 0.050,0.034,-0.720,0.484,1.774,1.526 |
| **e5073**e47d101b028aaf394fdc057e73c | 0.056,0.078,1.040,-1.196,-0.036,-0.258 | 0.032,0.097,0.767,-0.819,-0.648,-1.154 | 0.055,0.038,-1.433,1.075,1.708,1.010 | 0.069,0.032,-0.152,-0.065,1.567,1.199 |
| **e51ac**f4b457e6a1ae15b2ccedfb7a740 | 0.044,0.025,1.652,-1.769,-0.597,-1.093 | 0.030,0.079,1.507,-1.017,-1.907,0.210 | 0.055,0.043,-1.283,1.263,1.733,1.975 | 0.079,0.041,0.003,-0.353,1.117,0.178 |
| **e5246**81f66f54f8483f6d240bcfac099 | 0.040,0.072,1.320,-1.218,0.154,-1.153 | 0.069,0.069,1.150,-1.735,-0.936,-0.002 | 0.025,0.083,-0.601,0.010,-0.262,1.831 | 0.048,0.072,-1.060,0.863,0.694,0.643 |
| **e5433**250c9071a0509653a84461827cd | 0.048,0.079,1.518,-1.332,-0.637,-1.403 | 0.041,0.031,0.812,-0.540,0.238,-0.651 | 0.059,0.060,-1.187,1.494,1.846,1.335 | 0.095,0.064,-0.363,-0.211,0.978,1.454 |
| **e5486**ee77d5e0072a085df04f488353d | 0.088,0.017,0.977,-0.920,-1.683,1.197 | 0.081,0.086,0.764,-0.800,-0.081,-0.930 | 0.060,0.051,-1.480,1.181,0.574,0.685 | 0.054,0.035,-0.574,0.487,0.945,0.374 |
| **e5531**0606baf24e316907c504d8d958e | 0.052,0.087,1.501,-1.255,-0.127,-1.107 | 0.064,0.057,0.933,-1.469,0.711,-1.312 | 0.067,0.028,-1.440,0.912,1.867,1.071 | 0.092,0.073,-0.416,-0.021,1.693,1.197 |
| **e56ee**7b13188a0847ae931e600f6ee76 | 0.052,0.029,1.153,-1.266,-1.211,0.025 | 0.083,0.041,0.360,-0.465,0.024,0.866 | 0.100,0.059,-1.249,1.231,0.535,0.593 | 0.093,0.035,-0.730,0.586,0.410,0.489 |
| **e57b7**d8757db88f34128ce882fd0fdda | 0.068,0.055,1.233,-1.105,-1.075,0.086 | 0.016,0.018,1.967,-1.429,-1.572,0.471 | 0.093,0.086,-1.705,1.956,1.624,1.634 | 0.019,0.053,-0.598,0.335,-1.937,1.723 |
| **e57ca**a2eacad7dc1c10eb9dd8535ccef | 0.089,0.058,1.741,-1.884,-1.384,-0.386 | 0.085,0.035,0.323,-0.281,-1.484,0.977 | 0.076,0.098,-0.568,0.316,1.039,0.370 | 0.086,0.032,-0.384,0.246,1.929,-0.867 |
| **e5e70**2a88f558169f5d987dfedb587ea | 0.017,0.014,1.570,-1.786,-0.006,-1.062 | 0.023,0.070,1.879,-1.267,-1.471,-1.774 | 0.038,0.026,-1.918,1.946,1.745,1.738 | 0.099,0.053,-1.335,0.652,1.193,1.639 |
| **e5eca**7f5d6608baabe45d9c61e5e7b4a | 0.050,0.052,1.671,-1.829,-0.164,-0.518 | 0.038,0.095,0.127,-0.085,-0.904,0.180 | 0.056,0.100,-0.203,0.243,-0.332,0.540 | 0.079,0.097,-1.576,1.151,1.848,0.980 |
| **e5efd**4c574514896a0b98459a9222411 | 0.073,0.066,1.246,-1.173,-0.979,0.247 | 0.033,0.023,0.763,-1.069,1.566,0.108 | 0.100,0.062,-1.098,0.406,1.633,1.589 | 0.047,0.057,-1.115,0.662,1.046,1.474 |
| **e5f6a**a13086a811b33745897a0c3de11 | 0.074,0.071,1.667,-1.305,-1.314,-1.446 | 0.067,0.035,0.881,-0.748,0.560,-1.644 | 0.038,0.030,-1.426,1.654,-0.235,1.755 | 0.076,0.062,-1.196,0.862,0.863,0.511 |
| **e634f**dcf8dad7c7cc7e4d817c84b792f | 0.066,0.048,1.980,-0.932,-1.890,-1.576 | 0.097,0.074,0.468,-0.589,-0.429,-0.674 | 0.081,0.047,-0.645,-0.042,1.293,0.962 | 0.051,0.068,-0.235,-0.162,1.865,1.231 |
| **e6383**cba1be47501a6b31169693b723e | 0.054,0.048,0.795,-1.115,-1.474,-0.328 | 0.015,0.069,0.628,-0.526,-0.838,-0.385 | 0.090,0.053,-1.776,1.981,0.474,1.564 | 0.013,0.091,-1.430,1.228,-0.566,1.552 |
| **e63be**882f70f59aab8cb7081f66b5e41 | 0.090,0.024,0.646,-0.497,-1.119,-0.892 | 0.057,0.061,0.431,-0.264,1.189,-0.286 | 0.098,0.099,-0.634,1.019,1.366,0.801 | 0.052,0.065,-1.367,1.040,0.398,1.709 |
| **e6440**26f1b250cb0a319e7237023af5c | 0.096,0.040,1.781,-1.500,-1.620,-0.079 | 0.070,0.024,0.456,-1.172,-0.065,-0.284 | 0.074,0.055,-1.053,0.401,1.941,0.195 | 0.050,0.094,-1.737,1.405,1.308,1.857 |
| **e66d0**cae2b9e851604d978475d6fcc96 | 0.081,0.026,1.880,-1.919,-0.464,-1.054 | 0.046,0.018,0.223,-0.112,-1.876,-1.391 | 0.080,0.097,-0.560,0.203,-0.097,0.399 | 0.017,0.085,-1.268,1.081,0.910,1.192 |
| **e6f08**70ed0b5ac62f7be803948e72d34 | 0.078,0.082,1.720,-1.285,-0.892,-0.561 | 0.066,0.061,1.384,-1.625,1.006,-1.098 | 0.014,0.087,-1.922,1.314,1.787,1.635 | 0.088,0.016,-0.283,-0.153,1.087,1.046 |
| **e6f1c**f2c1080a94309dfaefccccf675a | 0.076,0.028,1.507,-1.825,-0.745,-0.065 | 0.023,0.043,0.212,-0.187,-0.410,-0.102 | 0.089,0.036,-1.644,1.793,0.279,0.531 | 0.098,0.082,-0.703,0.552,1.776,-0.612 |
| **e70b4**4fab7445f640e38930547e1e3c4 | 0.061,0.081,1.121,-0.807,-1.562,-0.945 | 0.080,0.045,1.551,-1.423,-0.317,0.885 | 0.057,0.043,-1.278,1.328,1.394,1.843 | 0.097,0.090,-1.200,0.577,1.879,1.373 |
| **e7173**3f7a0c75a7ced5ae159ba129f58 | 0.054,0.040,0.340,-0.290,-1.393,0.136 | 0.030,0.086,0.292,-0.322,0.613,0.401 | 0.076,0.030,-1.582,1.913,0.993,1.968 | 0.024,0.035,-0.735,0.576,1.531,1.328 |
| **e744c**3ed212f09d82ffa9b0962b5ca37 | 0.049,0.031,1.665,-1.726,-1.531,-1.309 | 0.080,0.043,1.350,-1.307,0.255,-1.280 | 0.054,0.082,-1.810,1.939,-0.735,1.509 | 0.086,0.049,-1.328,1.234,1.407,-1.292 |
| **e74bd**494ccdd9fcd7ebc7e1e805ddc4f | 0.020,0.063,1.639,-1.585,0.163,-1.895 | 0.064,0.058,0.949,-0.611,-0.940,-0.421 | 0.087,0.062,-1.437,1.307,0.004,1.527 | 0.032,0.025,-0.500,0.359,0.742,0.750 |
| **e7514**d7fe70616344c6494d048fff872 | 0.040,0.084,1.796,-1.930,-0.931,-0.728 | 0.086,0.049,0.568,-0.802,-0.070,-0.144 | 0.091,0.073,-1.586,0.896,1.940,1.342 | 0.024,0.057,-1.462,1.274,0.948,1.433 |
| **e765c**78fff916ba636fa577f41a5a74d | 0.047,0.053,1.408,-1.435,-0.298,-0.235 | 0.081,0.088,1.168,-1.170,1.880,-1.758 | 0.033,0.085,-0.785,0.511,1.544,1.778 | 0.079,0.035,-0.120,-0.086,1.090,0.372 |
| **e778c**2e98937b9364435c419f7b20fa7 | 0.033,0.020,1.700,-1.623,-1.053,-1.595 | 0.031,0.011,0.722,-1.572,-0.162,-0.344 | 0.060,0.043,-1.863,0.857,1.349,1.203 | 0.039,0.051,-1.310,1.053,1.742,0.899 |
| **e7be6**578e4ccc5b3d676bea94cd39255 | 0.040,0.070,1.580,-1.736,-0.213,-1.375 | 0.014,0.024,0.477,-0.521,-0.602,-0.150 | 0.068,0.100,-1.485,1.211,1.103,1.141 | 0.072,0.055,-1.282,1.186,-0.715,1.480 |
| **e7d78**203951a77bceec22589420191c8 | 0.083,0.061,0.711,-0.636,-0.841,-0.185 | 0.018,0.026,1.005,-1.035,-0.557,1.163 | 0.092,0.082,-1.040,1.217,-0.387,1.838 | 0.057,0.073,-0.360,0.023,1.629,1.757 |
| **e7e5e**88dc6ffbf4e7bd69b81ca35741a | 0.019,0.065,0.659,-0.436,-0.809,-1.855 | 0.059,0.027,1.836,-1.505,-1.159,1.625 | 0.043,0.039,-1.308,1.758,0.342,1.730 | 0.085,0.070,-0.269,-0.309,1.749,0.805 |
| **e7ed0**76bf2d7b32408396dddb6797ca1 | 0.021,0.035,1.889,-1.772,-1.792,-0.455 | 0.086,0.071,0.610,-0.447,-0.204,-1.286 | 0.030,0.065,-0.865,0.608,-0.275,1.258 | 0.097,0.040,-0.141,-0.451,1.877,1.243 |
| **e868c**d2eb48127e9bc2ae70521c984e7 | 0.066,0.067,1.960,-1.568,-1.017,-0.239 | 0.062,0.021,0.427,-1.201,-0.346,-0.907 | 0.036,0.085,-1.907,0.657,1.595,0.312 | 0.072,0.071,-0.659,0.347,0.290,1.677 |
| **e8725**59edf49aa885d73ef7687eb5d37 | 0.050,0.068,1.050,-0.834,-1.768,-0.970 | 0.013,0.088,1.616,-1.248,-1.006,-0.627 | 0.094,0.060,-1.810,1.879,1.847,0.967 | 0.098,0.067,-1.446,1.153,0.302,1.096 |
| **e888c**7a8601a02ca5855b14a118466bb | 0.047,0.020,1.351,-0.956,-1.851,-0.580 | 0.022,0.030,0.156,-0.031,-0.871,-1.587 | 0.072,0.049,-1.115,0.388,0.175,1.627 | 0.087,0.043,-0.544,0.033,1.449,1.029 |
| **e89f2**cf9bb1d922915ddafc1279b0c48 | 0.058,0.028,0.934,-0.698,-1.762,-0.474 | 0.013,0.015,0.604,-0.586,-0.484,-0.788 | 0.075,0.060,-1.765,1.168,0.877,1.922 | 0.026,0.099,-1.246,0.465,1.861,1.717 |
| **e8ccb**ac048413df60f90e9ad4950c81c | 0.025,0.028,1.153,-1.194,-1.389,-0.308 | 0.099,0.025,0.893,-0.854,-1.489,0.426 | 0.054,0.080,-0.939,0.649,-0.027,0.297 | 0.097,0.071,-1.249,0.739,1.816,0.797 |
| **e9046**fc04210fb9f6e63da8347a2536f | 0.018,0.022,1.745,-1.736,-1.369,-1.024 | 0.063,0.071,0.389,-0.497,-0.045,-1.644 | 0.051,0.014,-0.272,-0.060,1.185,1.594 | 0.062,0.048,-0.132,-0.162,1.200,1.492 |
| **e910f**0b0b553c22631ac46a2c625f644 | 0.061,0.019,1.845,-1.596,-1.832,-0.072 | 0.024,0.024,0.134,0.070,-1.134,-1.877 | 0.062,0.088,-1.088,0.275,1.612,0.156 | 0.061,0.057,-0.530,0.301,-0.277,1.466 |
| **e9625**f671dfee8f0e4856ed874f53e45 | 0.069,0.060,1.813,-1.853,-0.615,-0.839 | 0.041,0.065,1.369,-0.884,-1.962,0.035 | 0.056,0.026,-1.224,1.990,-0.682,1.038 | 0.054,0.049,-0.722,0.392,1.961,0.321 |
| **e9876**f00aded933393d2174f23ef883b | 0.079,0.072,1.820,-1.786,-1.287,-0.321 | 0.047,0.077,1.533,-0.980,-1.776,-0.289 | 0.094,0.023,-1.559,1.939,1.480,0.077 | 0.052,0.080,-0.858,0.358,0.585,1.713 |
| **e987b**609e3e8323ab4e67cb1472b070f | 0.078,0.064,0.783,-0.645,-0.857,-0.425 | 0.088,0.078,1.371,-1.418,-1.210,0.551 | 0.060,0.088,-1.887,1.860,-0.353,0.950 | 0.087,0.083,-0.547,0.314,-0.639,1.525 |
| **e9887**d4ecf8d9319638d7cd80c1c37c5 | 0.044,0.029,1.478,-1.390,-1.122,-0.501 | 0.025,0.044,0.771,-0.479,-0.792,-0.754 | 0.073,0.064,-0.669,0.862,1.864,0.704 | 0.070,0.060,-0.878,0.776,1.263,-0.503 |
| **e98e5**08c674d79a92fadb21218e79a21 | 0.056,0.085,1.558,-1.489,-0.336,-0.921 | 0.086,0.082,1.407,-0.997,0.512,-1.111 | 0.062,0.086,-1.733,1.884,1.821,1.629 | 0.032,0.055,-0.444,0.024,1.705,1.333 |
| **e99dd**cba0ad1ff3a50938d5b1de0ddbc | 0.093,0.020,1.257,-1.350,-1.113,1.009 | 0.098,0.022,1.122,-1.021,-0.832,-0.376 | 0.027,0.038,-1.050,0.998,0.690,0.468 | 0.082,0.036,-0.499,0.148,1.424,0.229 |

| | | | | |
|---|---|---|---|---|
| **e9c05**b695345f9c9d0c2d382c1c681b0 | 0.082,0.036,1.675,-1.462,-1.173,0.957 | 0.092,0.029,0.494,-0.569,-0.172,-0.743 | 0.083,0.088,-0.947,0.418,0.021,0.502 | 0.032,0.029,-0.598,0.345,1.459,0.501 |
| **e9db7**21fcccfd1263da61e81958a3451 | 0.048,0.079,1.664,-1.816,-0.185,-0.799 | 0.084,0.052,0.524,-0.401,-0.715,-1.125 | 0.099,0.028,-0.790,0.436,1.828,0.701 | 0.083,0.071,-0.738,0.391,1.395,1.551 |
| **e9deb**598aef246d0c522d09de6d70795 | 0.078,0.088,1.051,-0.848,-0.248,-1.390 | 0.052,0.064,0.920,-1.217,-0.296,-0.867 | 0.094,0.088,-1.851,1.803,-0.132,-0.596 | 0.084,0.049,-1.196,0.837,1.870,1.052 |
| **e9dfa**405e24f882c5a236bc6cf51b949 | 0.100,0.045,1.604,-1.293,-1.955,-0.273 | 0.028,0.071,1.199,-1.415,0.044,-1.776 | 0.071,0.064,-1.654,0.876,1.952,0.775 | 0.063,0.058,-1.615,1.438,1.809,-0.116 |
| **e9f37**6892fca7f779bb543bbe86864a3 | 0.099,0.080,1.738,-1.090,-1.740,-0.119 | 0.013,0.023,0.470,-0.597,-0.822,-0.250 | 0.071,0.094,-1.887,0.617,1.901,0.523 | 0.062,0.046,-0.452,-0.107,1.613,1.151 |
| **ea404**b6ff85f9b7a5b67a71e739ca12f | 0.078,0.086,1.692,-1.526,-1.776,0.195 | 0.080,0.020,0.998,-1.392,-0.422,-1.180 | 0.098,0.097,-1.584,0.937,1.311,-0.132 | 0.085,0.078,-0.072,-0.406,1.410,1.187 |
| **ea50f**a36aeafbdd40242ebd7dd336a1c | 0.083,0.049,0.150,-0.069,-1.646,1.441 | 0.039,0.079,1.435,-1.259,-0.950,-1.244 | 0.050,0.016,-1.430,1.706,0.632,1.700 | 0.082,0.097,-0.391,-0.207,1.605,1.675 |
| **ea830**604f240dcc55f962a4734d3208a | 0.045,0.042,1.878,-1.740,-0.615,-0.807 | 0.043,0.020,0.603,-1.347,-1.016,-0.807 | 0.054,0.077,-1.081,0.418,1.775,0.215 | 0.084,0.012,-0.179,-0.151,1.876,0.211 |
| **ea976**ce4928c37d2334c4257fe329565 | 0.043,0.050,1.815,-1.713,-1.054,-0.907 | 0.076,0.049,0.554,-1.042,-1.046,0.055 | 0.065,0.060,-1.852,0.939,-0.213,0.276 | 0.023,0.081,-0.940,0.849,1.766,0.471 |
| **eaa3b**39cd4197bdcb9aa2625a945b3ed | 0.080,0.078,1.666,-1.279,-1.837,-0.256 | 0.091,0.053,0.682,-0.871,-1.000,-0.228 | 0.071,0.038,-1.298,0.789,-0.456,0.262 | 0.054,0.060,-0.309,-0.003,1.868,0.881 |
| **eaacd**6e37614b749d4278db830a17e29 | 0.058,0.091,1.544,-1.383,-0.958,-0.073 | 0.072,0.020,1.191,-0.900,-0.787,-1.887 | 0.066,0.079,-0.828,0.616,1.873,0.352 | 0.094,0.022,-0.593,0.396,0.584,0.936 |
| **eaeb9**74fc944ee5d819a50789215374d | 0.047,0.055,1.479,-1.668,-0.668,-0.629 | 0.027,0.021,0.161,-0.137,0.616,-0.533 | 0.049,0.040,-0.689,0.799,1.897,0.303 | 0.023,0.042,-1.174,0.975,0.808,1.279 |
| **eb057**02c23c0fd7e08926f4c4933c922 | 0.054,0.061,1.443,-1.501,-0.814,-0.880 | 0.032,0.036,0.501,-0.532,-0.998,0.068 | 0.097,0.069,-0.619,0.493,0.617,0.374 | 0.040,0.071,-0.741,0.678,1.203,0.164 |
| **eb06b**e64bca48b821774fe64632fa1ab | 0.052,0.028,1.955,-1.879,-1.668,-0.314 | 0.040,0.061,0.889,-1.292,-0.702,-0.491 | 0.065,0.020,-0.714,0.337,1.540,0.439 | 0.040,0.051,-0.447,0.151,1.263,1.813 |
| **eb725**e4e5272304945d8888ca876c0ae | 0.093,0.036,0.991,-0.816,-1.170,-0.960 | 0.044,0.072,0.395,-1.225,0.349,1.238 | 0.077,0.090,-1.446,0.338,1.558,1.753 | 0.093,0.012,-1.914,1.583,1.948,1.102 |
| **eb736**8a4cd758cd87b92898dc10d886b | 0.098,0.069,0.132,0.204,-1.676,0.217 | 0.094,0.024,1.242,-0.760,-1.891,-1.880 | 0.068,0.042,-1.223,1.119,1.824,-1.924 | 0.073,0.072,-0.576,0.230,1.590,0.404 |
| **eb88d**2ee552034cb169c62be572af497 | 0.062,0.059,1.526,-1.035,-1.120,-1.868 | 0.018,0.066,1.626,-1.212,-0.255,-1.775 | 0.052,0.025,-1.854,1.875,1.015,1.596 | 0.084,0.091,-0.219,-0.087,1.076,-0.015 |
| **eb897**47082a3c5c1d5a35b8261b0390c | 0.081,0.053,0.858,-0.664,-0.664,-1.087 | 0.019,0.086,0.362,-0.287,0.631,-0.611 | 0.029,0.061,-1.398,1.271,0.847,1.942 | 0.017,0.071,-0.779,0.542,1.480,1.238 |
| **eb9ed**e0425cbfd667dc2a89ecb477164 | 0.058,0.020,1.425,-1.265,-1.933,-0.399 | 0.099,0.088,0.262,-0.031,-1.381,-0.052 | 0.031,0.046,-0.226,0.113,1.088,0.237 | 0.084,0.064,-0.142,-0.000,0.848,0.348 |
| **ebabd**629a73783d48fcb557564e62427 | 0.090,0.011,1.763,-1.703,-1.883,-0.508 | 0.040,0.030,1.104,-0.950,-0.668,-1.585 | 0.065,0.024,-1.925,1.908,1.922,0.184 | 0.047,0.060,-0.161,-0.423,1.883,1.975 |
| **ebb18**177a6398ba24cebfb873afc0c75 | 0.048,0.019,1.284,-1.687,-0.450,-0.377 | 0.094,0.050,0.359,-0.585,0.032,-0.179 | 0.068,0.072,-0.574,0.253,-0.138,1.691 | 0.072,0.029,-0.218,-0.025,1.930,1.815 |
| **ebb1d**e77d43b89db68b4135caa4a11c0 | 0.087,0.018,1.539,-1.777,-1.877,-0.162 | 0.100,0.017,0.355,-0.319,-0.110,-1.584 | 0.096,0.093,-1.478,1.456,-0.822,0.691 | 0.081,0.093,-1.212,0.626,1.757,1.879 |
| **ebb35**24d79add1fd7c3346c96cbc528f | 0.043,0.048,1.989,-1.869,-0.641,-0.600 | 0.013,0.043,0.608,-0.565,-1.276,-1.235 | 0.020,0.069,-1.972,1.880,1.864,1.396 | 0.033,0.013,-1.303,1.244,0.952,0.456 |
| **ebd62**a2f4f86a2af9dd07d65b800b9fe | 0.042,0.097,1.772,-1.436,-1.114,-0.817 | 0.078,0.055,1.886,-1.373,-1.093,-0.914 | 0.017,0.055,-1.834,1.917,1.027,1.407 | 0.046,0.086,-1.363,0.794,1.817,1.383 |
| **ebd80**abfea06e007316927bb9c71ac7b | 0.044,0.063,0.991,-0.992,-1.154,-0.022 | 0.072,0.041,1.943,-1.615,-1.426,0.914 | 0.041,0.074,-1.510,1.684,0.286,0.566 | 0.062,0.077,-0.681,0.360,1.273,0.129 |
| **ebed3**5bc88ef9f89b0db08f378ef88a6 | 0.089,0.012,1.744,-1.846,-1.347,0.604 | 0.091,0.020,0.724,-0.854,-1.902,-0.624 | 0.071,0.020,-0.473,0.193,0.777,1.812 | 0.095,0.071,-0.386,0.077,1.456,1.171 |
| **ebf9e**f25a7c8ce8756cce91477643401 | 0.059,0.021,1.771,-1.731,-1.854,1.292 | 0.052,0.059,0.215,-0.120,-0.620,-1.227 | 0.064,0.050,-0.712,0.138,1.273,0.768 | 0.083,0.068,-1.516,1.352,0.951,0.356 |
| **ebfed**70556a4d1fb9d60cf05750b9e1d | 0.080,0.048,1.126,-1.075,-1.693,0.358 | 0.062,0.070,0.357,-0.616,0.334,-0.607 | 0.083,0.021,-1.014,0.582,0.691,0.568 | 0.022,0.072,-1.893,1.716,1.972,1.551 |
| **ec255**2f162c69a4ea14bbea12d7434aa | 0.046,0.088,1.569,-1.203,-0.696,-1.787 | 0.086,0.054,1.751,-1.331,-1.030,-1.716 | 0.058,0.070,-1.853,1.804,0.898,0.261 | 0.070,0.071,-1.847,1.452,1.036,1.404 |
| **ec42e**91b3fa8d858117b4ca0041621db | 0.042,0.016,1.885,-1.669,-1.927,-0.595 | 0.058,0.059,1.538,-1.505,1.070,-1.598 | 0.078,0.045,-1.672,1.955,-0.148,0.663 | 0.100,0.083,-0.981,0.575,-0.692,1.879 |
| **ec46e**ab357cbc92b7d460882079df4e1 | 0.055,0.027,1.362,-0.911,-1.680,-0.287 | 0.093,0.078,0.801,-0.398,-0.797,-1.953 | 0.074,0.043,-1.825,0.674,1.240,-1.285 | 0.098,0.040,-0.670,0.182,1.619,0.314 |
| **ec4f1**08639d68b575fd3ed9d0a8a9960 | 0.080,0.078,1.428,-1.321,0.183,-1.048 | 0.069,0.035,0.833,-1.081,-0.179,-0.137 | 0.048,0.060,-1.877,1.760,1.426,1.038 | 0.058,0.052,-0.483,0.222,0.814,1.149 |
| **ec4fc**acfd5208afee3de69a9f54b555c | 0.047,0.092,1.098,-1.209,0.770,-0.717 | 0.011,0.070,0.582,-1.524,1.985,-0.266 | 0.068,0.040,-1.062,0.211,1.354,1.268 | 0.074,0.032,-0.800,0.641,1.161,0.649 |
| **ec5dd**bf4eb2fb28148b6b0218ea6c4c6 | 0.056,0.015,1.118,-0.422,-1.891,0.890 | 0.094,0.025,0.644,-0.536,-1.256,-0.019 | 0.097,0.066,-1.937,0.523,0.958,0.167 | 0.053,0.078,-1.265,0.726,1.326,0.639 |
| **ec877**b25f8d20e00055400ec991b4867 | 0.090,0.072,1.674,-1.565,-0.425,-1.807 | 0.025,0.024,1.534,-1.955,-0.488,-0.774 | 0.061,0.086,-1.756,1.375,1.829,0.220 | 0.049,0.075,-0.747,0.580,1.948,0.300 |
| **ecaf8**c77444632a298b835ebd7f49760 | 0.015,0.080,1.612,-1.631,-1.658,-0.579 | 0.054,0.051,1.428,-1.078,-1.398,-0.035 | 0.094,0.026,-1.484,1.845,1.081,1.384 | 0.048,0.094,-0.916,0.169,1.854,1.651 |
| **ecb18**a1c795f2f52bdc0baed382a6d32 | 0.070,0.060,1.963,-1.617,-0.840,-1.421 | 0.013,0.038,0.621,-0.469,-0.941,-1.107 | 0.044,0.036,-1.401,1.294,-0.543,1.430 | 0.015,0.045,-0.894,0.695,0.092,1.037 |
| **ed121**95e761256231789efceb471d9ac | 0.034,0.026,1.097,-0.967,-0.795,-0.426 | 0.046,0.062,0.358,-0.408,-0.822,0.395 | 0.054,0.026,-1.401,1.031,1.353,1.345 | 0.026,0.058,-1.195,0.759,1.296,1.798 |
| **ed1f2**cfda06b9ca3d018e7d7637ca5f0 | 0.096,0.013,1.009,-0.823,-1.879,0.300 | 0.080,0.071,0.405,-0.296,-0.401,0.120 | 0.073,0.029,-1.510,1.707,1.305,0.297 | 0.075,0.062,-0.373,-0.136,1.828,1.885 |
| **ed327**ead2fa723b6b186abb548af4250 | 0.051,0.074,1.764,-1.604,-1.892,-1.358 | 0.043,0.074,1.540,-1.995,0.752,-1.053 | 0.026,0.087,-1.521,1.398,-0.614,0.638 | 0.071,0.091,-0.373,0.226,1.751,-0.244 |
| **eda2a**3ef20bcbb85f38d934b4bc3ff51 | 0.080,0.021,1.713,-1.813,-0.925,-0.368 | 0.024,0.059,1.134,-1.531,-0.819,-1.151 | 0.030,0.045,-1.847,1.151,1.994,0.256 | 0.081,0.030,-0.148,-0.202,1.847,1.146 |
| **eda99**2ca0cde183f9334f65599bbf6a3 | 0.048,0.084,1.952,-1.689,-1.384,-0.363 | 0.035,0.066,1.950,-1.618,-0.541,-1.500 | 0.094,0.040,-1.975,1.945,1.731,0.173 | 0.091,0.086,-0.384,0.038,1.055,0.230 |
| **edb06**def4d2260fb0048f2f3d7d43c16 | 0.061,0.053,0.674,-0.655,-1.301,0.930 | 0.022,0.051,0.491,-0.425,1.371,-0.949 | 0.098,0.061,-1.219,1.575,1.406,0.834 | 0.039,0.054,-1.205,0.977,1.373,1.082 |
| **edd8d**80e2e50ac4d44a2276b3e5931ab | 0.079,0.024,1.200,-1.024,-1.824,1.285 | 0.041,0.050,1.307,-0.959,-0.601,-1.254 | 0.065,0.063,-1.367,1.338,1.981,1.285 | 0.055,0.059,-0.689,0.436,0.524,1.805 |
| **edf94**a34544ca24abae74d9ec6fbc96b | 0.057,0.040,1.735,-1.546,-1.735,-1.833 | 0.015,0.085,0.973,-0.691,-1.153,-0.235 | 0.063,0.033,-1.110,1.089,1.467,1.179 | 0.098,0.074,-0.076,-0.845,1.597,1.814 |
| **ee152**d9c5c169a6f734229c41f0f5cc4 | 0.054,0.039,1.217,-1.036,-1.727,-0.768 | 0.021,0.082,1.448,-1.800,0.464,-0.555 | 0.077,0.029,-1.402,1.315,0.485,1.299 | 0.081,0.065,-0.792,0.558,0.382,0.856 |
| **ee5c9**3c269a0ce310ca27748e2cc89cf | 0.052,0.044,0.895,-0.867,-0.470,-0.438 | 0.026,0.091,1.732,-1.296,-0.411,-0.931 | 0.026,0.038,-1.804,1.920,0.585,1.024 | 0.086,0.066,-1.295,1.183,0.945,-0.872 |
| **ee645**0c2ffdb0bce2ab1c6eccc50b093 | 0.044,0.072,1.226,-0.859,-1.542,-1.214 | 0.088,0.080,1.079,-1.460,0.010,-0.517 | 0.060,0.090,-1.996,1.104,1.430,0.498 | 0.090,0.061,-0.606,0.284,0.173,1.744 |
| **ee6d8**ac45f0843fc5d6c8cf69dfba2c9 | 0.050,0.024,1.174,-1.456,-1.318,1.054 | 0.093,0.087,0.751,-0.802,0.275,0.547 | 0.088,0.081,-1.552,1.722,1.490,1.400 | 0.061,0.044,-0.583,0.377,1.939,0.128 |
| **eeb79**65e5440d91efa1967421b09e273 | 0.050,0.024,1.040,-1.143,-0.929,-0.213 | 0.065,0.079,0.477,-1.712,0.191,-0.008 | 0.025,0.088,-1.686,0.355,1.400,1.802 | 0.055,0.092,-1.440,1.208,1.465,1.168 |
| **eecbd**e74812db219e285582fa74f2112 | 0.051,0.049,0.547,-0.463,-0.822,-0.062 | 0.042,0.074,0.396,-0.523,-0.764,-0.123 | 0.044,0.057,-1.840,1.209,-0.779,0.794 | 0.093,0.018,-0.468,0.060,1.882,1.022 |
| **eed01**fcf90b59f33711435eda8056407 | 0.074,0.088,1.920,-1.241,-1.225,-0.986 | 0.033,0.019,0.569,-1.234,-0.756,-0.530 | 0.049,0.059,-1.581,0.347,0.614,1.346 | 0.082,0.036,-0.329,-0.192,1.764,1.588 |
| **eed30**78978290d2f9918a8238799b49b | 0.081,0.015,1.902,-1.693,-1.529,0.162 | 0.090,0.023,1.208,-1.911,-1.680,-0.604 | 0.022,0.083,-1.232,0.607,-0.533,0.626 | 0.098,0.034,-1.007,0.706,0.574,1.791 |
| **eee3a**1091e897ab030d4182ba731eeda | 0.043,0.029,0.575,-0.478,-1.078,-0.128 | 0.071,0.024,0.783,-0.355,-1.382,0.430 | 0.078,0.036,-0.363,0.340,0.989,0.694 | 0.076,0.081,0.118,-0.731,1.948,0.344 |
| **ef292**470e3d052c6a33b29f3364b37ee | 0.042,0.070,0.790,-0.742,-0.801,-0.211 | 0.050,0.078,1.267,-1.164,-1.664,0.135 | 0.046,0.099,-0.767,1.069,1.556,0.277 | 0.087,0.071,-1.131,0.978,-0.274,1.481 |
| **ef600**8fad0213814c71573f5bdf51e08 | 0.039,0.045,1.474,-1.883,-1.154,0.305 | 0.023,0.050,0.393,-0.671,-0.739,1.250 | 0.075,0.038,-1.091,0.699,1.281,0.460 | 0.037,0.073,-0.818,0.610,1.486,0.353 |
| **ef932**02fffcf30827a46765d0151b2f7 | 0.049,0.084,1.694,-1.427,-1.677,-1.047 | 0.070,0.051,1.022,-1.298,1.206,-1.838 | 0.082,0.097,-1.124,0.782,-0.054,1.460 | 0.095,0.090,-0.513,-0.308,1.790,1.842 |
| **ef9ac**ab4932c9de248413fdc79fcad8d | 0.054,0.022,0.977,-1.185,-0.814,0.115 | 0.047,0.096,0.542,-0.160,-1.747,-0.789 | 0.028,0.097,-0.913,0.920,-1.148,1.913 | 0.077,0.049,-0.426,0.248,0.657,0.829 |
| **efbfd**ad2ede1118145995dc53042f267 | 0.097,0.022,0.363,-0.064,-1.851,0.633 | 0.039,0.089,0.679,-0.405,-1.546,-1.067 | 0.096,0.038,-1.319,1.153,-0.851,1.229 | 0.089,0.021,-0.974,0.568,1.830,1.212 |
| **f0673**3e138012e0dd847816322f687dc | 0.097,0.032,1.011,-0.955,-0.625,-1.071 | 0.025,0.047,0.156,-0.354,-1.455,0.687 | 0.046,0.086,-0.973,0.714,1.982,-0.656 | 0.086,0.049,-0.184,-0.238,1.938,1.989 |
| **f0676**43720bfcee3c8c7733556900772 | 0.040,0.040,0.925,-1.322,0.095,0.062 | 0.028,0.061,1.139,-0.812,-1.688,-1.308 | 0.093,0.038,-1.399,1.786,-0.375,0.859 | 0.033,0.046,-0.941,0.807,0.507,1.411 |
| **f09da**59261dd7f79802312c34b965135 | 0.067,0.055,0.907,-0.725,-1.667,0.511 | 0.048,0.090,1.673,-1.211,-1.025,-1.072 | 0.092,0.076,-1.741,1.913,-0.487,1.513 | 0.019,0.092,-0.341,-0.113,1.964,1.181 |
| **f0f1c**7e8fab2ce535fba953058479e8f | 0.079,0.021,1.818,-1.775,-1.113,0.138 | 0.023,0.019,0.355,-0.428,0.006,0.476 | 0.057,0.058,-0.413,0.155,-0.215,1.374 | 0.097,0.046,-0.723,0.205,1.816,0.330 |
| **f0f45**93d4730caca3b9d04a199dba131 | 0.098,0.020,1.970,-1.732,-1.962,-0.236 | 0.022,0.021,0.337,-0.284,-0.555,-1.567 | 0.099,0.047,-0.847,0.621,1.870,-0.906 | 0.097,0.077,-0.633,0.361,0.375,1.502 |
| **f0f6a**6e1ef4e07ec121935d1bcb9154c | 0.097,0.019,1.008,-0.993,-1.894,1.129 | 0.043,0.051,1.253,-0.711,-1.902,-0.401 | 0.064,0.065,-0.898,0.854,1.796,0.469 | 0.073,0.091,-0.505,-0.247,1.357,1.868 |
| **f0fb3**d056d77db87f1c06df62ae67441 | 0.087,0.020,1.819,-1.251,-1.775,-0.648 | 0.099,0.076,0.995,-1.699,-0.703,-1.511 | 0.013,0.032,-1.855,0.805,1.636,0.384 | 0.098,0.033,-0.179,-0.184,1.462,1.404 |
| **f107b**17d72ec1daaa0e19219943ff322 | 0.056,0.051,1.097,-1.285,-0.673,-0.534 | 0.091,0.042,0.280,-0.465,0.556,-0.360 | 0.080,0.070,-1.231,0.797,1.386,0.653 | 0.096,0.089,-0.693,0.475,0.957,0.827 |
| **f10a6**49edeedbf1da3c6499dc4dd4463 | 0.079,0.031,1.578,-1.270,-1.372,-1.137 | 0.033,0.083,0.771,-0.302,-0.963,-1.609 | 0.079,0.046,-1.706,1.262,1.608,1.492 | 0.098,0.042,-0.170,-0.556,1.665,1.954 |

| | | | | |
|---|---|---|---|---|
| f1278d0643eb95524189b0496d897da3 | 0.018,0.038,1.803,-1.525,-1.839,-0.889 | 0.068,0.022,0.438,-0.371,-0.868,-1.813 | 0.047,0.043,-1.617,0.636,-0.022,1.059 | 0.040,0.035,-0.274,-0.147,1.944,1.430 |
| f15f6de1027ed515fc34d61818891a81 | 0.050,0.041,1.926,-1.647,-1.983,-0.199 | 0.061,0.072,1.949,-1.436,-0.956,-1.014 | 0.028,0.063,-1.820,1.948,1.658,-0.364 | 0.095,0.093,-0.716,-0.012,1.975,0.741 |
| f186aac43c67456c9e54d4d3a247e10a | 0.095,0.020,1.589,-1.720,-1.898,0.939 | 0.017,0.056,1.240,-0.826,1.665,0.709 | 0.076,0.084,-1.207,1.881,1.954,0.301 | 0.054,0.021,-1.068,0.826,1.401,-0.656 |
| f19bec85c2d4104a767b4b821891852d | 0.039,0.083,1.562,-1.836,0.378,-1.105 | 0.078,0.092,0.605,-0.820,1.119,-0.925 | 0.095,0.046,-1.277,1.180,1.042,-1.392 | 0.027,0.077,-1.154,0.754,0.754,1.923 |
| f1b5babd998fc56014d701f18c30bea9 | 0.044,0.037,1.602,-1.311,-1.568,-1.009 | 0.052,0.080,0.914,-1.913,0.157,-0.896 | 0.077,0.089,-1.934,0.454,1.967,1.678 | 0.034,0.080,-0.737,0.437,1.519,1.100 |
| f1e7d9a3d5e3fd540be07c1575f3618b | 0.096,0.022,1.345,-1.557,-0.448,-0.458 | 0.030,0.090,0.627,-0.645,-0.272,-0.111 | 0.080,0.041,-1.697,1.492,0.646,0.448 | 0.018,0.061,-1.109,0.920,-0.574,1.220 |
| f255c7227478c0264b20bfe9f91a5c51 | 0.061,0.026,1.735,-1.864,-1.634,-0.593 | 0.074,0.039,0.148,-0.134,-1.061,1.520 | 0.097,0.061,-1.040,1.135,1.873,-0.234 | 0.087,0.050,-0.376,-0.175,1.826,1.008 |
| f29e2f37c97102acbd3added101c3755 | 0.049,0.024,0.712,-0.437,-1.693,0.344 | 0.082,0.060,0.835,-0.822,0.011,-1.084 | 0.089,0.037,-1.413,0.465,1.391,1.674 | 0.036,0.060,-1.016,0.472,1.791,1.407 |
| f2ec15565ab42521b2a8dfe09d7c745d | 0.085,0.067,0.788,-0.445,-1.211,-1.325 | 0.012,0.083,0.896,-0.582,-0.944,-0.677 | 0.026,0.040,-1.485,1.884,-0.682,1.963 | 0.088,0.074,-1.823,1.141,1.858,1.350 |
| f2ec80a3529fd7537c5baac6ff13eb7c | 0.087,0.086,1.599,-1.444,-1.633,-0.436 | 0.056,0.061,1.371,-0.944,0.602,-1.920 | 0.084,0.087,-0.706,0.777,1.774,-0.022 | 0.084,0.019,-0.913,0.493,1.983,0.354 |
| f343085c91597d12f3ccb4931c81df5a | 0.023,0.045,1.911,-1.657,-1.476,-1.441 | 0.066,0.089,1.399,-1.081,0.378,-1.902 | 0.081,0.063,-1.823,1.930,0.878,1.222 | 0.029,0.094,-0.434,-0.142,1.433,1.691 |
| f382add1a47d1d3eba639d1a1462c3ff | 0.091,0.051,1.632,-1.335,-1.035,-0.648 | 0.083,0.016,1.373,-1.849,1.339,-0.675 | 0.094,0.089,-1.911,1.729,1.333,0.794 | 0.035,0.041,-0.568,0.327,-0.319,1.841 |
| f383edb06284e4a302cd9c6f2a575723 | 0.040,0.067,1.755,-1.867,-0.809,-0.573 | 0.035,0.082,1.377,-1.412,0.329,-0.556 | 0.025,0.063,-1.974,1.972,1.512,0.262 | 0.058,0.075,-0.870,0.389,1.050,1.874 |
| f38945958d0819286d5aad86e37908e7 | 0.089,0.027,1.941,-1.798,-1.738,-0.022 | 0.029,0.034,0.542,-0.307,-0.769,-0.663 | 0.098,0.035,-0.624,0.540,1.805,0.751 | 0.097,0.063,-0.783,0.188,1.962,1.263 |
| f38aa79750b137a7ce22b60759e568ce | 0.049,0.017,1.923,-1.860,-0.394,-1.934 | 0.093,0.081,0.712,-1.403,-0.646,-1.717 | 0.047,0.090,-0.966,0.152,0.098,0.609 | 0.097,0.040,-0.952,0.544,1.763,1.462 |
| f3939e49a8981e22a267b9ddc56e0c05 | 0.042,0.061,0.657,-0.635,-0.537,-0.338 | 0.031,0.090,1.262,-1.899,0.035,-1.014 | 0.071,0.063,-0.992,0.659,-0.126,0.468 | 0.033,0.035,-0.731,0.607,1.900,1.426 |
| f39ec1bac63bfd924fd2d132b9bba5e4 | 0.059,0.031,1.281,-1.526,-0.426,0.348 | 0.019,0.035,0.420,-0.498,-0.328,-1.317 | 0.061,0.060,-0.536,0.246,0.959,0.877 | 0.026,0.075,-1.401,1.239,-0.516,1.376 |
| f3a11735cb8ece38aada46ae9eeabb64 | 0.050,0.057,1.511,-1.716,-1.890,-0.193 | 0.017,0.092,1.085,-1.632,-0.940,0.589 | 0.073,0.065,-0.735,0.262,1.921,0.638 | 0.047,0.041,-0.279,0.102,1.843,0.282 |
| f3c5cdc7986da88c9a66fc1ef25319a1 | 0.094,0.060,1.148,-1.190,-0.284,-0.350 | 0.043,0.097,0.545,-1.502,0.363,0.972 | 0.085,0.061,-0.498,-0.026,1.455,1.508 | 0.097,0.052,-1.431,0.953,1.826,1.937 |
| f3c7b268837cea99aa191814a966e36f | 0.069,0.020,1.111,-0.868,-1.863,0.520 | 0.021,0.090,1.256,-1.057,-0.984,-1.156 | 0.072,0.050,-1.919,1.452,1.626,1.203 | 0.045,0.091,-0.621,-0.001,1.967,1.872 |
| f3eae4daf237a1045aeb55eadbe59ab6 | 0.044,0.081,1.927,-1.633,-0.410,-1.659 | 0.085,0.089,0.312,-0.348,-1.923,0.991 | 0.040,0.093,-0.705,0.329,1.783,-0.071 | 0.067,0.018,-1.120,0.867,1.584,1.134 |
| f429f6ba7529e498e62a18e1f8aac60a | 0.045,0.035,1.458,-0.698,-1.679,-1.055 | 0.051,0.024,0.611,-0.532,-1.194,-0.365 | 0.042,0.083,-0.148,-0.124,1.908,-0.290 | 0.059,0.070,-0.479,0.164,0.199,0.767 |
| f47141e75692089ed0b8b51459072611 | 0.089,0.048,1.454,-0.915,-0.938,-1.026 | 0.065,0.051,0.706,-0.754,-1.851,0.029 | 0.029,0.052,-1.617,0.822,1.410,0.985 | 0.066,0.082,-1.145,0.558,1.960,1.001 |
| f48696ba204f330b8068538142ece40a | 0.062,0.046,1.981,-1.391,-1.947,-1.336 | 0.070,0.070,1.552,-1.850,-1.005,-0.705 | 0.089,0.030,-0.938,0.073,1.695,1.594 | 0.091,0.069,-0.519,0.269,1.508,-0.524 |
| f48f33d60d39bfe05c2d780deb87fb19 | 0.034,0.051,1.611,-1.900,-0.182,-0.475 | 0.053,0.087,0.208,-0.174,-0.646,0.178 | 0.015,0.096,-0.684,0.490,1.246,0.587 | 0.013,0.059,-0.529,0.479,1.901,0.045 |
| f4a970d6c8da02e0682cc7539e2953e3 | 0.012,0.084,1.585,-1.621,-1.305,-0.194 | 0.019,0.057,1.030,-1.929,-0.408,-1.656 | 0.093,0.074,-1.874,0.733,-0.120,1.279 | 0.099,0.080,-1.923,1.647,-0.137,1.823 |
| f4d0843b28a9166889dc17fdff8774d9 | 0.088,0.046,1.454,-1.664,-0.055,-1.918 | 0.099,0.067,1.312,-1.110,-0.945,-1.036 | 0.033,0.033,-1.167,1.275,1.823,-1.358 | 0.088,0.035,-1.008,0.750,1.318,1.319 |
| f4e653bcf90e890a9101ab33adce18be | 0.053,0.068,1.705,-1.391,-0.820,-0.969 | 0.023,0.090,0.754,-1.353,0.275,-0.675 | 0.088,0.099,-1.178,0.518,-0.278,1.212 | 0.043,0.082,-0.995,0.608,1.127,1.536 |
| f4e9496de76f63c0007def4ad6316957 | 0.054,0.040,1.006,-0.669,-1.717,-0.918 | 0.048,0.082,1.192,-1.061,0.527,-1.027 | 0.078,0.081,-1.794,1.900,1.249,-0.683 | 0.032,0.046,-1.273,1.097,1.052,0.545 |
| f4ecda30a434707ac3d2efa37bda2079 | 0.050,0.063,1.693,-1.420,-1.813,-0.456 | 0.090,0.042,1.643,-1.575,-1.533,-0.635 | 0.050,0.065,-1.532,0.989,1.258,1.680 | 0.060,0.071,-0.344,-0.110,1.631,1.127 |
| f4ed1338f48c95083061f88c7d4b628a | 0.023,0.078,1.394,-1.408,0.327,-1.872 | 0.081,0.083,0.803,-0.449,-1.146,0.151 | 0.044,0.015,-0.852,1.473,1.651,-0.197 | 0.097,0.041,-0.558,0.078,1.958,0.883 |
| f538735eecabbef1a4f7a1e78b2a79a5 | 0.047,0.022,1.758,-1.285,-1.330,-1.038 | 0.069,0.044,0.441,-1.325,-1.313,1.094 | 0.046,0.098,-1.738,0.099,1.764,1.580 | 0.036,0.056,-1.439,1.072,1.877,1.698 |
| f551ca44b045d75d3b2532ca93ecef31 | 0.060,0.079,0.900,-0.764,-0.580,-0.602 | 0.041,0.094,1.052,-0.725,0.170,-1.294 | 0.093,0.045,-1.142,1.627,0.046,1.649 | 0.094,0.076,-1.270,0.876,1.499,0.922 |
| f5608eab86bb75d336ed803957cecfa2 | 0.093,0.070,1.645,-1.478,-1.773,0.670 | 0.029,0.071,0.361,-1.005,0.362,0.046 | 0.099,0.060,-1.795,0.417,1.488,0.308 | 0.092,0.075,-1.325,0.614,1.962,1.897 |
| f594adab4488922a2e658d201b73ac39 | 0.056,0.058,1.249,-1.284,-1.087,-1.191 | 0.065,0.089,0.392,-0.284,0.026,-0.477 | 0.029,0.087,-1.136,1.245,-1.780,1.947 | 0.094,0.031,-1.685,1.375,1.827,-0.104 |
| f595e4f068f121c165ed275540457195 | 0.061,0.051,1.440,-0.993,-1.899,-0.742 | 0.025,0.045,0.813,-1.749,1.059,-0.388 | 0.045,0.056,-1.316,0.821,1.407,0.297 | 0.051,0.032,-0.410,0.226,1.822,0.839 |
| f5a11fb5d8afd807c2727420adc27263 | 0.055,0.024,1.372,-1.295,-0.080,-0.967 | 0.081,0.060,1.003,-0.818,-1.889,0.103 | 0.060,0.067,-0.792,0.673,-0.051,0.335 | 0.073,0.097,-1.211,0.912,0.486,0.873 |
| f5ebcf83421698e422e4c784447fd076 | 0.082,0.087,1.979,-1.446,-1.746,-0.857 | 0.082,0.058,0.629,-0.329,-0.906,-1.652 | 0.098,0.043,-1.767,0.522,1.537,1.473 | 0.098,0.041,-1.451,0.897,1.879,1.011 |
| f616200e449fc0b589f4fea448f1a2ab | 0.049,0.082,0.834,-0.683,-1.940,-0.326 | 0.042,0.097,0.597,-0.330,-0.935,-0.343 | 0.032,0.092,-1.822,1.904,-0.101,1.800 | 0.011,0.071,-1.409,1.013,0.396,1.725 |
| f64e6cac525a868581ac3a84ec453706 | 0.085,0.060,0.791,-0.409,-1.312,-0.543 | 0.039,0.018,0.224,-0.324,-1.439,-0.144 | 0.097,0.047,-1.221,0.487,0.788,1.201 | 0.068,0.082,-1.791,1.432,1.942,1.519 |
| f68939fc0aac0721b2963e0f3452ded5 | 0.063,0.042,1.044,-0.687,-1.217,-0.655 | 0.042,0.029,0.281,-0.205,-1.177,-0.190 | 0.039,0.062,-1.496,0.649,1.802,-0.437 | 0.064,0.091,-1.299,0.922,1.470,0.431 |
| f68db3f0e1682ad9293d5533df4317dd | 0.074,0.086,1.891,-1.476,-1.936,-0.148 | 0.070,0.087,0.616,-0.737,-0.765,0.167 | 0.064,0.065,-1.787,1.148,0.985,1.884 | 0.069,0.080,-0.605,0.101,1.589,1.557 |
| f70f6a825a7104fa5963ab0ce2936245 | 0.045,0.080,1.796,-1.854,0.128,-0.982 | 0.044,0.038,0.530,-0.985,0.253,-1.052 | 0.088,0.081,-1.451,0.228,1.791,1.650 | 0.098,0.077,-0.896,0.362,1.366,1.558 |
| f70fbd17b74ea79ce94a014fafc3fcac | 0.071,0.030,1.671,-1.859,0.106,-0.818 | 0.014,0.049,0.647,-0.493,-1.913,0.351 | 0.068,0.059,-1.573,1.956,-0.577,0.652 | 0.098,0.058,-1.533,1.183,-0.417,1.718 |
| f73452bdfc859f21df092b1c1681131c | 0.097,0.067,1.855,-1.522,-1.047,-0.741 | 0.037,0.014,1.138,-1.236,-0.822,0.110 | 0.025,0.040,-1.469,1.588,1.713,0.528 | 0.084,0.062,-0.548,0.373,1.563,-0.687 |
| f73e8d5f0a2d2fc4f33c8b7d7f71ea28 | 0.018,0.050,1.173,-1.207,-0.802,-0.387 | 0.038,0.076,0.745,-1.413,-1.088,0.228 | 0.069,0.083,-1.130,0.607,-0.213,0.177 | 0.045,0.055,-0.291,-0.088,1.107,1.341 |
| f75a032c3d3792c6fcf2442476a5cfd3 | 0.047,0.022,1.596,-1.307,-1.685,0.467 | 0.053,0.076,1.185,-1.456,-1.054,-1.177 | 0.046,0.031,-0.511,-0.003,1.442,1.267 | 0.081,0.061,-0.039,-0.388,1.481,0.412 |
| f7c1a29cbae540afacbb6919bbc69d0b | 0.036,0.025,1.343,-1.421,-1.262,-0.135 | 0.019,0.096,0.905,-0.968,-1.930,-0.356 | 0.043,0.014,-1.576,1.446,1.307,0.869 | 0.097,0.051,-0.493,0.142,1.378,1.256 |
| f7c399a4526c595a4618b38f49914866 | 0.049,0.023,1.658,-1.410,-1.789,-0.034 | 0.074,0.078,0.582,-0.886,-0.697,-0.163 | 0.023,0.060,-0.963,0.284,1.915,1.181 | 0.028,0.072,-1.643,1.215,1.998,1.483 |
| f7c70e2a08facd650f0fa0dedf859285 | 0.058,0.043,1.642,-1.457,-1.585,-0.091 | 0.096,0.088,0.181,-0.153,-0.732,-0.298 | 0.031,0.050,-0.707,0.109,0.658,1.904 | 0.082,0.076,-0.380,0.227,0.397,1.051 |
| f7c7175f99d4426a0058a6ff80a4a511 | 0.094,0.068,1.313,-1.067,-0.695,-1.133 | 0.096,0.088,0.748,-1.156,0.595,-0.068 | 0.054,0.074,-1.479,0.910,-0.220,1.434 | 0.020,0.057,-1.717,1.379,0.155,1.734 |
| f7cd1d8fa900ae330ab752c7a76208a0 | 0.044,0.011,1.626,-1.923,-1.377,-0.037 | 0.092,0.073,0.787,-1.905,1.159,-0.880 | 0.014,0.047,-1.644,0.877,-0.277,0.729 | 0.043,0.042,-1.352,1.206,1.970,0.917 |
| f7ece5b83029e5011fb290bf83ff11fd | 0.065,0.067,1.346,-1.317,-1.780,-0.708 | 0.064,0.051,0.513,-0.402,-1.879,0.985 | 0.086,0.024,-1.035,1.631,-0.291,-1.610 | 0.090,0.083,-1.111,0.996,1.473,-0.694 |
| f856601336881dc5f501e52120f25ced | 0.040,0.098,0.911,-0.622,-0.153,-1.049 | 0.054,0.088,0.990,-0.931,-0.929,-0.804 | 0.079,0.060,-1.106,0.830,0.104,0.556 | 0.063,0.097,-1.530,1.222,0.084,1.037 |
| f870e658d5a842a17efd0cf725794b39 | 0.097,0.034,1.407,-1.036,-1.335,0.638 | 0.092,0.086,0.964,-1.135,-1.652,-0.981 | 0.092,0.063,-1.658,0.648,0.941,0.798 | 0.097,0.040,-0.271,-0.027,1.299,0.383 |
| f87842dbe91ba1379dcab17203073445 | 0.015,0.050,1.757,-1.899,-0.373,-0.879 | 0.073,0.046,1.268,-0.868,-0.704,-1.341 | 0.064,0.042,-1.012,1.238,1.686,-0.003 | 0.069,0.071,-0.260,-0.222,1.834,0.979 |
| f879225d9e79356c166c8006ae72634c | 0.098,0.019,1.724,-1.581,-1.897,-0.512 | 0.018,0.012,0.946,-1.826,0.331,1.113 | 0.093,0.046,-1.566,0.796,1.791,1.388 | 0.065,0.076,-0.954,0.734,1.887,0.488 |
| f896e5f12a44d9aef74e5ba168550ee5 | 0.042,0.081,1.917,-1.743,-1.783,-1.812 | 0.078,0.087,0.504,-0.945,-0.820,1.491 | 0.098,0.049,-1.242,0.666,1.997,0.156 | 0.082,0.013,-0.340,0.132,1.705,0.305 |
| f8a072798377a86a94968720e12c03ae | 0.043,0.036,1.960,-1.762,-1.100,-0.127 | 0.025,0.089,0.389,-0.835,-1.968,0.030 | 0.098,0.034,-1.757,0.615,-0.117,1.138 | 0.099,0.060,-0.398,0.060,1.833,-0.815 |
| f901290292fa133127d7a37d46f673aa | 0.089,0.067,1.440,-1.386,-0.426,-0.545 | 0.026,0.086,0.574,-0.748,-1.733,-0.572 | 0.029,0.093,-0.535,0.121,-0.282,1.155 | 0.036,0.093,-1.086,1.083,1.553,-0.219 |
| f967771e3ab3c89fdc3d1bda0a2afb85 | 0.052,0.045,1.514,-0.889,-1.545,-0.910 | 0.076,0.046,1.083,-1.876,-1.480,1.084 | 0.029,0.038,-1.050,0.518,1.022,-0.164 | 0.097,0.053,-0.069,-0.245,1.705,-1.062 |
| f9859f46bc326222ce17ba3e339f0fa1 | 0.077,0.057,1.951,-1.469,-1.049,-0.501 | 0.028,0.076,0.674,-1.912,-0.407,-0.514 | 0.088,0.014,-1.616,0.323,1.309,0.415 | 0.039,0.045,-0.780,0.548,1.951,0.640 |
| f9cce6f0e2d10302f3801ce7cb2d3f5c | 0.080,0.022,1.170,-0.836,-1.780,-0.515 | 0.055,0.016,0.872,-0.546,-1.989,0.091 | 0.088,0.053,-0.463,0.133,1.342,0.386 | 0.078,0.098,-1.761,1.220,-0.203,1.793 |
| fa1c85d5aa9d55c67d9ae314981d2e40 | 0.092,0.024,0.815,-0.643,-1.488,-0.254 | 0.075,0.086,0.790,-1.276,0.778,0.869 | 0.028,0.098,-1.750,1.285,0.543,1.891 | 0.051,0.025,-1.588,1.401,1.130,1.225 |
| fa231b34663397bb0e487b321c08ff74 | 0.088,0.084,1.519,-1.910,0.096,-0.965 | 0.022,0.056,0.852,-0.794,-1.911,-0.301 | 0.040,0.033,-1.642,1.915,1.809,0.163 | 0.098,0.078,-0.576,-0.083,1.889,1.665 |
| fa40100d08ec71720704233c862b38bf | 0.074,0.061,1.128,-0.732,-0.375,-1.537 | 0.061,0.093,0.865,-1.331,-0.362,-0.480 | 0.063,0.068,-1.901,0.612,1.611,1.593 | 0.076,0.057,-0.685,0.300,1.370,1.345 |
| fa4d79168021194712078571 6f54d465 | 0.037,0.063,1.532,-1.418,-0.917,-1.601 | 0.061,0.041,0.393,-0.419,-0.556,-1.213 | 0.097,0.050,-1.744,0.686,1.965,1.355 | 0.041,0.092,-0.997,0.703,1.956,1.109 |
| fab4190330ef7d069296dd097b24ed5e | 0.088,0.022,1.191,-0.993,-1.653,-0.017 | 0.098,0.057,1.421,-1.647,-0.530,-1.093 | 0.064,0.036,-1.985,1.334,1.815,1.973 | 0.099,0.075,-0.474,0.018,1.642,1.104 |

| | | | | |
|---|---|---|---|---|
| fb0f8c9af3084954bdbae4c41ed626db | 0.078,0.030,1.904,-1.565,-1.819,-0.785 | 0.029,0.076,0.660,-1.434,-0.980,-0.366 | 0.034,0.044,-0.918,0.391,-0.265,1.414 | 0.091,0.034,-0.477,0.056,1.963,1.438 |
| fb10e58e7b6a2ef4e06ab74df0268ee7 | 0.058,0.060,1.348,-1.715,-0.093,-0.493 | 0.060,0.085,0.937,-0.585,-1.600,-1.458 | 0.072,0.084,-1.277,1.147,0.663,1.173 | 0.089,0.098,-1.173,0.715,1.587,1.605 |
| fb25d6307b13f765db4669d8f160e172 | 0.063,0.034,1.363,-1.737,-0.362,-0.572 | 0.033,0.089,0.482,-0.268,-0.999,-0.691 | 0.022,0.038,-0.828,0.979,-0.193,1.232 | 0.053,0.045,-0.730,0.488,1.577,1.456 |
| fb39e04ba4d797378b1a790fc68c78f2 | 0.023,0.055,1.586,-1.300,-1.435,-1.063 | 0.021,0.095,0.590,-0.629,0.231,-1.472 | 0.089,0.063,-0.733,0.129,1.197,0.276 | 0.075,0.065,-0.095,-0.308,1.293,1.350 |
| fb4b23d1c0c419dd2d82357b7cb48f02 | 0.087,0.085,1.749,-1.499,-1.724,0.029 | 0.062,0.074,0.779,-0.747,0.628,-1.036 | 0.098,0.028,-1.291,0.989,0.994,1.501 | 0.066,0.076,-0.723,0.282,1.005,1.373 |
| fb6acfae7e28617612a52bbf3e4f060e | 0.093,0.023,1.741,-1.175,-1.846,0.016 | 0.030,0.083,0.738,-1.286,-1.583,-1.271 | 0.044,0.072,-1.555,0.511,1.845,0.188 | 0.076,0.090,-0.646,0.357,1.861,0.299 |
| fb7b6d5ceed2ecfd0c5b040c0b3e4c8f | 0.046,0.078,1.620,-1.446,-0.814,-0.841 | 0.012,0.092,1.530,-1.949,-1.163,1.509 | 0.076,0.100,-1.458,1.129,-0.313,1.400 | 0.082,0.073,-0.477,-0.164,1.765,1.642 |
| fba405a15e8f97e548370d72c175ddea | 0.068,0.040,1.979,-1.561,-1.905,-1.383 | 0.040,0.023,0.300,-0.376,-0.401,0.026 | 0.014,0.053,-0.647,0.131,1.912,1.521 | 0.048,0.057,-0.129,0.060,0.928,-0.274 |
| fbbf326afdce526fc84b987387919699 | 0.051,0.068,1.831,-1.452,-1.655,-0.466 | 0.050,0.049,1.713,-1.738,-0.266,-1.314 | 0.069,0.071,-1.278,0.557,1.183,1.926 | 0.037,0.069,-0.102,-0.292,1.877,1.079 |
| fbcc216d7eb6546a27699fad4ba1a59c | 0.045,0.022,0.855,-0.845,-0.700,-0.376 | 0.054,0.076,0.136,-0.122,0.338,-0.691 | 0.026,0.053,-1.449,1.661,1.454,-1.483 | 0.058,0.041,-0.433,0.326,0.848,0.722 |
| fbe263f69912cda73dab66bbb1d4bc52 | 0.054,0.077,1.705,-1.475,-1.398,-0.203 | 0.042,0.041,0.251,-0.270,-1.956,-0.510 | 0.049,0.038,-1.051,0.259,1.095,0.700 | 0.077,0.043,-0.423,0.242,1.889,-0.515 |
| fc0ac698038990a79e5d64369a2bfb85 | 0.083,0.057,0.640,-0.451,-0.953,-0.261 | 0.018,0.095,0.098,-0.134,0.254,-0.162 | 0.073,0.069,-1.542,0.634,1.140,1.912 | 0.035,0.045,-1.097,0.969,0.720,0.948 |
| fc59c49488ee81b8da91dc241c2b1b0e | 0.090,0.027,0.633,-0.808,-0.619,0.079 | 0.028,0.097,0.821,-0.628,-1.030,-0.654 | 0.076,0.078,-1.911,1.939,1.436,1.031 | 0.078,0.097,-0.733,0.399,1.663,1.187 |
| fc66b3ee22f9a564fb0d7a6e8b9c14ea | 0.095,0.075,1.935,-1.299,-0.389,-1.859 | 0.066,0.098,1.759,-0.274,-1.911,-1.848 | 0.057,0.081,-1.203,0.812,-0.022,1.490 | 0.060,0.056,-1.180,0.447,1.818,0.902 |
| fc71730b5f8e6dc0f709e036afd5e574 | 0.093,0.022,1.089,-0.696,-1.572,0.095 | 0.081,0.074,0.835,-1.335,-0.897,0.117 | 0.085,0.097,-1.735,0.786,1.622,0.527 | 0.069,0.023,-1.478,1.130,1.706,1.482 |
| fc9b3fb66a3e65b1c6860b6485d3d373 | 0.067,0.087,0.754,-0.692,-0.449,-0.313 | 0.042,0.052,0.482,-0.289,-1.524,-0.559 | 0.068,0.047,-0.673,0.641,0.420,0.312 | 0.068,0.017,-1.334,1.164,1.852,-1.238 |
| fcbd4caf1ace07248fb3f51ca2e41622 | 0.049,0.072,1.681,-1.491,-1.680,-0.716 | 0.096,0.082,1.774,-1.418,-0.536,-0.880 | 0.060,0.028,-1.343,1.712,0.106,0.865 | 0.058,0.062,-1.162,0.977,1.860,-0.592 |
| fcdc33b5127dc3762e0199e71cf2a268 | 0.024,0.023,1.366,-1.368,-1.284,-1.011 | 0.079,0.027,1.214,-1.448,-1.604,1.573 | 0.082,0.091,-1.697,1.442,1.778,-0.051 | 0.083,0.080,-1.729,1.324,1.737,0.922 |
| fce0be1298d6170480d601fc0c1b6cf8 | 0.090,0.012,1.908,-1.751,-1.931,0.523 | 0.092,0.083,0.538,-0.310,-1.635,-0.621 | 0.013,0.056,-1.529,0.642,-0.848,1.295 | 0.042,0.099,-1.177,0.876,1.960,1.004 |
| fcfa8b3ee7811a9a51e88bb8ef77f3cf | 0.088,0.074,1.751,-1.081,-1.741,-1.595 | 0.014,0.033,1.020,-1.918,0.065,-0.472 | 0.063,0.072,-1.903,0.925,-0.115,1.186 | 0.073,0.077,-0.989,0.749,-0.546,1.695 |
| fcfc64ba0886245b49a275b42d52a2fe | 0.071,0.061,1.389,-1.263,-1.094,-0.216 | 0.066,0.066,0.942,-1.891,-1.321,0.969 | 0.059,0.077,-1.679,1.006,0.438,0.612 | 0.037,0.083,-1.645,1.444,1.865,0.772 |
| fd26f5ec45d510fab422f8fa271faaa2 | 0.080,0.030,1.942,-1.912,-1.757,0.640 | 0.041,0.053,1.145,-0.728,-1.812,-0.902 | 0.069,0.044,-1.717,1.539,1.682,-0.009 | 0.041,0.064,-0.373,-0.040,-0.293,1.853 |
| fd2feda5cc58cde04c0e6b3f2bf4f3a2 | 0.046,0.085,1.546,-1.264,-1.161,-1.140 | 0.073,0.080,0.947,-0.937,-1.223,-1.011 | 0.070,0.098,-1.649,1.306,-0.515,0.021 | 0.020,0.074,-1.697,1.403,-0.785,1.774 |
| fd7191ce46963c4df491fac9bf60a3f8 | 0.084,0.056,1.377,-1.494,-1.381,0.037 | 0.093,0.049,0.192,-0.251,-1.955,1.918 | 0.065,0.064,-1.695,1.190,-0.815,0.524 | 0.086,0.076,-1.341,0.982,1.601,1.564 |
| fd8124d08a12641f4b2b0c5c8e3475e7 | 0.087,0.023,0.616,-0.581,-0.803,0.452 | 0.044,0.082,0.208,-0.180,-1.585,0.370 | 0.025,0.036,-1.624,1.723,0.621,1.881 | 0.056,0.021,-1.436,1.239,1.753,1.878 |
| fdacb0925b24f105faf859cf31c2698a | 0.044,0.062,1.250,-1.180,-1.723,0.176 | 0.073,0.022,1.120,-1.365,0.656,-1.781 | 0.085,0.055,-1.661,1.293,-0.133,1.771 | 0.076,0.060,-0.969,0.594,1.839,0.231 |
| fdd7aa077285160dce8d048cd99fcceb | 0.035,0.019,1.223,-0.722,-1.839,-0.431 | 0.064,0.088,0.653,-0.750,-1.711,-0.408 | 0.097,0.051,-1.414,0.078,1.886,-0.061 | 0.079,0.091,-0.205,-0.011,1.521,-0.257 |
| fdef8e195405132b58e5003082c9636d | 0.059,0.073,1.641,-1.376,-1.902,-1.487 | 0.016,0.080,0.743,-0.534,-0.720,-1.251 | 0.069,0.035,-0.587,0.350,1.351,0.475 | 0.014,0.056,-1.074,0.842,1.593,1.436 |
| fe1ff196adeebe12d39ccfcf2467096d | 0.038,0.074,1.864,-1.325,-1.532,-1.413 | 0.045,0.056,0.896,-0.868,-0.723,0.084 | 0.066,0.041,-1.534,1.141,-0.510,1.888 | 0.078,0.035,-0.394,-0.009,0.758,0.556 |
| fe2f9e8d280694b5f291c688a8e77024 | 0.069,0.075,1.728,-1.398,-1.667,-0.444 | 0.093,0.040,1.698,-0.747,-1.204,1.290 | 0.082,0.043,-0.653,0.927,1.584,0.142 | 0.072,0.090,-0.929,0.810,1.166,-0.540 |
| fe3c62fccc9ea7461665b715c141e839 | 0.073,0.019,1.553,-1.364,-1.428,-0.373 | 0.017,0.077,0.669,-0.640,-0.930,-1.218 | 0.099,0.026,-0.584,0.190,0.631,1.788 | 0.098,0.066,-0.488,-0.101,1.990,0.851 |
| fe4c196aacee75e6aaef15de0b149cc4 | 0.099,0.085,1.465,-0.727,-1.864,-0.349 | 0.027,0.050,0.480,-0.550,0.039,-1.351 | 0.099,0.096,-1.869,0.583,1.637,1.464 | 0.079,0.086,-0.125,-0.226,0.994,0.815 |
| feb39fa36337cc5a4c51d1fab0ea411b | 0.096,0.059,1.613,-1.684,0.182,-0.648 | 0.069,0.089,1.192,-1.125,0.805,-1.625 | 0.057,0.089,-1.793,1.770,1.357,0.661 | 0.052,0.072,-0.609,0.257,1.294,1.215 |
| fee2ceda251de93e7e7ec7a2e56c16af | 0.098,0.017,1.681,-1.276,-1.777,0.142 | 0.064,0.097,0.801,-1.857,-0.374,-0.365 | 0.053,0.030,-1.145,0.518,0.688,-0.556 | 0.062,0.018,-0.690,0.521,1.868,0.374 |
| ff04278a89858c6ea354534fe667fb4a | 0.080,0.024,1.934,-1.718,-0.722,-0.957 | 0.031,0.079,0.949,-1.867,0.922,-0.206 | 0.013,0.023,-1.674,0.900,0.778,0.958 | 0.067,0.049,-0.719,0.246,1.877,1.156 |
| ff4c3e4d20bdab7924d800052e4d5b6e | 0.092,0.017,0.632,-0.644,-0.826,1.180 | 0.065,0.046,1.807,-1.154,-1.783,1.679 | 0.036,0.032,-1.324,1.878,1.755,1.935 | 0.088,0.031,-1.033,0.349,1.711,1.439 |
| 0b83a1513cf3498b4b330e365c568264 | 0.085,0.064,0.787,-0.486,-1.174,-0.871 | 0.048,0.058,0.974,-0.593,-1.754,-0.564 | 0.074,0.045,-1.053,1.302,-0.038,1.330 | 0.026,0.094,-1.603,1.186,0.726,1.430 |
| 0cc6b0f6ec02ff851b69149d3b780daf | 0.065,0.059,0.102,0.037,-1.351,0.401 | 0.055,0.089,0.465,-1.115,0.477,0.665 | 0.068,0.065,-1.418,1.131,-0.479,0.796 | 0.097,0.070,-0.245,-0.397,1.748,0.870 |
| 0e255ef2138d2d50b00799d58f0c7c82 | 0.045,0.051,1.048,-0.704,-1.643,-1.105 | 0.052,0.047,1.800,-1.394,0.237,-0.928 | 0.031,0.037,-1.566,1.834,1.729,0.357 | 0.060,0.046,-0.625,0.221,1.127,1.330 |
| 0fbc9c464a6911c80873c26b26bd2f7c | 0.078,0.074,0.529,-0.349,-1.356,-0.301 | 0.082,0.090,0.787,-0.506,0.775,-0.100 | 0.033,0.037,-0.647,1.317,1.144,1.606 | 0.041,0.054,-0.722,0.516,0.833,0.366 |
| 1296812b7fb3cfacbe8f3a851eab4278 | 0.066,0.085,0.453,-0.041,-1.150,-1.771 | 0.091,0.068,1.565,-0.738,-0.496,-1.099 | 0.057,0.092,-1.473,1.755,1.847,1.951 | 0.020,0.097,-0.091,-0.006,0.527,0.214 |
| 14700dd78f9628a3634b5b72fb39e6c0 | 0.095,0.064,0.419,-0.229,-1.713,-0.152 | 0.091,0.070,1.601,-0.816,1.202,0.097 | 0.092,0.098,-0.370,0.551,1.378,0.558 | 0.085,0.060,-0.752,0.100,1.965,1.394 |
| 17d52bc9af5f4ca20053d8f40aadada3 | 0.081,0.080,0.618,-0.238,-0.982,-1.557 | 0.030,0.079,1.683,-1.281,1.758,-1.546 | 0.099,0.062,-1.829,1.922,1.850,1.541 | 0.016,0.056,-0.507,0.281,0.651,1.520 |
| 1e5b5f9fa6946d86c5a317563d5e92ec | 0.091,0.091,1.048,-0.601,-1.106,-0.641 | 0.038,0.079,1.418,-1.197,-1.465,0.170 | 0.081,0.033,-1.072,0.751,1.884,0.945 | 0.099,0.052,-1.090,0.403,1.907,0.743 |
| 2af3c5d54c6a35498d47d8312825b3c6 | 0.054,0.074,1.187,-1.060,-1.468,-0.486 | 0.034,0.059,0.685,-0.930,1.580,1.331 | 0.034,0.032,-1.170,1.703,1.557,-1.313 | 0.032,0.093,-0.061,-0.223,0.488,0.750 |
| 2ea91ec79399d1f296a002dea9200e4d | 0.070,0.062,0.498,-0.345,-0.808,-0.657 | 0.037,0.072,1.352,-0.790,-1.884,0.702 | 0.062,0.013,-1.276,1.921,1.281,1.933 | 0.096,0.048,-1.224,0.594,1.811,1.754 |
| 389710744cd5c8ccfc4cb241e8c74dbc | 0.081,0.090,0.521,-0.113,-1.911,-0.643 | 0.025,0.042,1.062,-0.985,0.078,-0.232 | 0.085,0.085,-1.344,1.216,1.347,0.833 | 0.092,0.040,-0.478,0.191,1.171,0.012 |
| 3c5935916004b6490dd8b0c1f2dd4192 | 0.082,0.059,1.756,-1.243,-1.838,-0.195 | 0.022,0.058,1.938,-1.242,-0.179,-1.342 | 0.080,0.028,-1.525,1.407,1.527,1.657 | 0.042,0.042,-0.763,0.351,0.928,1.464 |
| 429ad189a04df4ecf490d9daff95f5d7 | 0.048,0.086,0.295,0.052,-1.577,-1.286 | 0.068,0.037,0.728,-0.400,-0.787,-0.899 | 0.081,0.066,-1.020,1.134,0.590,1.857 | 0.072,0.043,-0.257,0.027,0.746,1.010 |
| 4377f15f8354760797ae1cb52eb59534 | 0.097,0.034,1.046,-0.643,-1.484,-1.039 | 0.056,0.052,1.139,-0.831,-1.295,0.436 | 0.099,0.033,-1.438,1.659,1.415,0.402 | 0.040,0.063,-0.581,0.402,1.459,0.088 |
| 52fb10539e48e2065965717d12898798 | 0.086,0.059,0.651,-0.340,-0.961,-0.817 | 0.087,0.081,1.389,-1.064,-0.248,-1.431 | 0.060,0.024,-1.898,1.926,1.054,1.895 | 0.062,0.071,-0.323,-0.129,1.797,0.995 |
| 5621445bc672195de82fed807ec75fd0 | 0.094,0.087,0.476,-0.068,-1.070,-1.205 | 0.045,0.085,0.595,-0.335,0.103,-0.941 | 0.097,0.073,-1.384,1.827,1.101,-0.831 | 0.098,0.070,-0.242,-0.093,0.358,1.498 |
| 668b8536615f21f942f65ead3f6124da | 0.091,0.061,0.673,-0.268,-1.276,-1.405 | 0.093,0.071,1.443,-1.430,-0.502,0.478 | 0.084,0.012,-1.447,1.527,1.925,0.929 | 0.053,0.058,-0.452,0.181,0.325,1.683 |
| 68f5eb3b72ec57dbfb19e8f31c823bb9 | 0.084,0.083,0.819,-0.471,-0.771,-0.699 | 0.097,0.095,1.068,-1.274,0.680,-1.450 | 0.070,0.044,-1.537,1.066,0.937,1.852 | 0.076,0.077,-1.231,0.727,1.378,1.511 |
| 79627d8e2ecfc3ede61e727ee71bf788 | 0.062,0.081,0.590,-0.314,-1.516,-0.733 | 0.052,0.054,1.188,-0.674,-1.957,-0.363 | 0.041,0.094,-1.594,1.932,-0.369,1.874 | 0.069,0.039,-0.642,0.323,0.922,1.415 |
| 7b1bff41083a49c53111fd84c7a1bd16 | 0.058,0.064,0.786,-0.510,-1.496,-1.186 | 0.016,0.071,1.468,-1.329,0.752,0.166 | 0.090,0.090,-1.237,1.295,0.801,0.701 | 0.087,0.092,-0.614,-0.305,1.953,1.743 |
| 7c90cb0730398dadd468dae01f3a0c51 | 0.065,0.058,0.656,-0.405,-1.257,-0.966 | 0.027,0.068,1.540,-1.119,-1.892,-0.813 | 0.054,0.069,-1.845,1.961,1.142,1.971 | 0.095,0.057,-0.607,-0.028,1.665,1.803 |
| 8183a302e8a1b440cf56a95a85abb671 | 0.073,0.081,0.866,-0.617,-1.261,-0.444 | 0.019,0.045,1.600,-1.167,-1.488,0.650 | 0.020,0.064,-1.540,1.862,0.469,1.954 | 0.080,0.013,-0.524,0.240,0.904,1.290 |
| 8e35d045c015cef7ff1a6f85452dd688 | 0.067,0.066,0.793,-0.338,-1.080,-0.970 | 0.016,0.032,0.718,-0.844,-0.045,-0.367 | 0.065,0.038,-1.097,0.531,1.644,1.536 | 0.045,0.090,-1.084,0.396,1.926,1.437 |
| 91ab19a5765cbcfb371b649096f85d5a | 0.045,0.077,0.256,0.118,-0.255,-1.344 | 0.091,0.062,1.386,-1.465,0.528,0.369 | 0.057,0.044,-0.637,0.659,-0.016,0.765 | 0.041,0.055,-1.209,0.980,0.712,0.484 |
| 952cb774657763e45e82e343424ed432 | 0.074,0.081,0.600,-0.232,-1.427,-1.072 | 0.013,0.080,1.335,-1.048,0.046,-1.640 | 0.056,0.029,-1.404,1.477,1.206,1.235 | 0.075,0.095,-1.223,0.603,1.734,1.633 |
| 96fd7088a10f95bded8c053066e116ba | 0.052,0.074,0.394,-0.144,-1.135,-1.137 | 0.039,0.013,1.314,-0.866,-1.065,1.146 | 0.097,0.063,-0.770,0.697,1.935,0.308 | 0.034,0.058,-0.137,-0.195,1.741,1.474 |
| a184fd42eecdac76f98e3abb6e64ecd8 | 0.078,0.064,0.869,-0.588,-1.814,-0.795 | 0.011,0.098,1.611,-0.747,-0.743,-1.540 | 0.054,0.087,-1.006,1.821,-0.877,0.335 | 0.011,0.056,-0.596,0.338,1.961,1.184 |
| a5091b8a2efd1e54c139c7026923b9cc | 0.055,0.063,0.296,-0.023,-1.730,-0.908 | 0.045,0.076,0.930,-0.704,-0.591,-0.595 | 0.066,0.093,-1.737,1.690,1.883,1.951 | 0.095,0.061,-0.622,0.256,1.069,1.260 |
| ae4b33237da307003964558ad4c9677f | 0.081,0.076,1.180,-0.814,-1.407,-0.385 | 0.012,0.033,1.617,-1.364,-0.249,-1.313 | 0.047,0.051,-1.552,1.465,1.728,1.383 | 0.087,0.045,-1.060,0.687,1.351,0.227 |
| b7a48b0fa4e4dd520f4b6e1bfaea7b55 | 0.057,0.039,0.922,-0.615,-1.199,-1.850 | 0.031,0.062,1.581,-0.850,0.277,-1.754 | 0.096,0.036,-1.371,1.758,1.867,0.385 | 0.038,0.063,-0.505,-0.049,1.270,1.948 |
| b7c67a89ab21ab4edfad0f4738da8321 | 0.049,0.080,0.293,-0.100,-1.051,-1.245 | 0.023,0.080,1.581,-0.859,-1.881,0.763 | 0.025,0.075,-1.029,1.905,0.802,0.489 | 0.023,0.055,-0.724,0.533,1.646,0.831 |
| c9a6ebe5adbe809ddcbed8a35018c0ea | 0.082,0.047,0.835,-0.486,-1.543,-1.120 | 0.054,0.054,1.807,-1.198,-1.405,-1.279 | 0.028,0.064,-0.998,1.120,1.946,1.390 | 0.050,0.055,-0.433,0.034,1.667,1.581 |

| | | | | |
|---|---|---|---|---|
| **d2023**5da66e018baee0730474166e637 | 0.069,0.021,1.147,-0.849,-1.600,0.011 | 0.011,0.014,1.452,-1.645,1.524,-0.701 | 0.037,0.075,-1.132,1.314,1.847,-1.334 | 0.042,0.036,-1.137,1.080,-0.603,1.062 |
| **dbf94**392f0fc4d87e0551a398dbb55e1 | 0.055,0.076,0.503,-0.264,-1.257,-0.977 | 0.087,0.083,0.888,-0.610,-0.520,0.943 | 0.046,0.084,-0.765,1.267,1.170,0.808 | 0.049,0.071,-1.216,0.779,1.148,1.033 |
| **e5a4b**efbe2bb54afe82cf041cdc6396d | 0.042,0.057,0.263,-0.056,-1.210,-0.821 | 0.020,0.071,1.280,-1.036,1.215,-1.496 | 0.053,0.095,-1.907,1.626,0.665,1.914 | 0.070,0.065,-0.552,0.178,1.799,0.499 |
| **e6450**6ea08f1c78c1e77f166aa1fb0d8 | 0.082,0.094,1.033,-0.614,-1.876,-1.817 | 0.022,0.020,1.341,-0.747,-0.884,-0.606 | 0.032,0.086,-1.025,1.741,1.896,0.521 | 0.080,0.075,-0.315,-0.374,1.208,1.616 |
| **f4a63**9c160f53042e301c3e1090a99df | 0.064,0.023,0.758,-0.444,-1.644,-0.181 | 0.014,0.081,1.463,-1.072,0.536,-0.160 | 0.080,0.062,-1.920,1.854,1.244,1.818 | 0.025,0.088,-0.746,0.049,1.918,1.563 |
| **fab97**7d6038914f8c0e89ea301e8d3a0 | 0.066,0.098,0.069,0.077,0.363,-1.697 | 0.076,0.049,0.944,-0.690,-0.776,0.820 | 0.086,0.060,-1.183,1.674,-0.660,1.580 | 0.071,0.098,-0.518,0.124,1.937,0.709 |
| **fe6cc**5f5ff4ad282119a1e9a83bb59ed | 0.064,0.022,0.296,-0.155,-0.612,-0.148 | 0.044,0.057,1.012,-0.944,0.406,-1.114 | 0.099,0.020,-0.978,0.445,1.638,1.892 | 0.052,0.090,-0.169,-0.335,1.370,0.973 |
| **0cbec**b3b8e7cf8d16f502803f73f9100 | 0.046,0.084,0.672,-0.596,-1.238,-0.230 | 0.028,0.027,0.930,-0.793,1.884,-0.801 | 0.070,0.064,-1.288,1.943,1.527,-1.305 | 0.081,0.020,-0.534,0.262,1.658,-0.748 |
| **19281**4670e527cfd763c46552cbc6d3f | 0.075,0.092,0.668,-0.304,-0.834,-1.901 | 0.087,0.049,1.845,-1.114,1.415,-1.948 | 0.096,0.092,-1.346,1.943,1.460,1.107 | 0.084,0.032,-0.474,-0.136,1.902,1.051 |
| **2199d**5d32b1a1e2f05a6700c44139e5e | 0.033,0.019,0.392,-0.358,-1.145,0.109 | 0.047,0.025,0.415,-0.504,1.128,-0.163 | 0.087,0.036,-0.911,1.097,0.957,-0.675 | 0.039,0.071,-0.424,0.077,1.758,1.582 |
| **229cf**7495fb723d8d2c376c8aada50d3 | 0.088,0.057,0.809,-0.520,-1.103,-1.405 | 0.026,0.025,0.791,-0.744,1.739,-0.710 | 0.080,0.070,-1.104,1.381,0.277,1.454 | 0.078,0.094,-0.735,0.283,1.238,1.236 |
| **4baeb**b0d3dcf4ffd2a1e0d452002c3c9 | 0.080,0.054,0.774,-0.634,-1.669,0.525 | 0.077,0.060,1.652,-1.134,1.722,-1.104 | 0.075,0.051,-0.733,1.522,0.551,-0.936 | 0.052,0.083,-0.318,-0.251,1.065,1.227 |
| **8141a**8c617d5ae6d38d5d73dd3a1ade5 | 0.081,0.080,0.375,-0.039,-1.297,-1.334 | 0.073,0.022,1.224,-1.167,1.423,-1.759 | 0.093,0.067,-1.417,1.940,-0.343,0.455 | 0.055,0.048,-0.331,0.130,0.918,0.433 |
| **98ba2**074c5379538863425088111caac | 0.063,0.030,1.636,-1.497,-1.841,-0.149 | 0.026,0.094,0.355,-0.929,1.861,-0.172 | 0.097,0.069,-0.755,0.564,0.476,-0.902 | 0.070,0.070,-0.542,0.073,1.871,1.423 |
| **b81f2**38e7944dc4eb91d1ebeb1dae165 | 0.058,0.064,0.427,-0.176,-0.720,-1.080 | 0.083,0.082,0.335,-0.409,1.209,-1.007 | 0.079,0.061,-0.979,0.647,1.923,0.862 | 0.029,0.091,-1.282,0.730,1.775,1.784 |
| **b9adf**b6dfddafbd42ec0ca4b68b471fd | 0.091,0.062,1.374,-0.979,-0.781,-1.675 | 0.026,0.077,1.425,-1.244,0.979,-0.427 | 0.033,0.040,-1.339,1.420,1.503,1.909 | 0.027,0.066,-0.081,-0.183,-0.269,1.127 |
| **f183c**0cf4f534b5bdf5f028325fa4ec4 | 0.066,0.031,1.622,-1.607,-1.882,-0.138 | 0.073,0.043,0.904,-1.833,1.871,0.246 | 0.096,0.071,-1.568,1.481,1.524,-1.110 | 0.090,0.065,-0.907,0.634,1.655,0.262 |
| **2426f**f02eaf3fee3e24de816474a8c74 | 0.040,0.070,0.892,-0.704,-1.594,-1.423 | 0.026,0.052,1.012,-0.923,1.821,-0.755 | 0.082,0.066,-1.048,1.601,1.495,-1.814 | 0.088,0.047,-0.632,0.459,1.246,-0.005 |
| **3d73e**09370ebe4a62bb208fb9ce36372 | 0.053,0.071,0.600,-0.321,-1.835,-1.145 | 0.013,0.070,1.668,-0.883,0.228,-1.690 | 0.050,0.089,-1.497,1.955,1.268,1.382 | 0.028,0.094,-0.562,0.013,1.750,1.530 |
| **9f2fa**46b0c44546bf090eb021d3db4c3 | 0.068,0.083,0.310,-0.009,-1.678,-0.768 | 0.042,0.077,1.127,-0.855,1.869,-0.531 | 0.097,0.020,-1.294,1.544,1.945,1.717 | 0.073,0.060,-1.675,1.286,1.708,0.083 |
| **f0bf0**37480ae805afee50c0039124da0 | 0.068,0.064,1.409,-1.195,-0.532,-0.773 | 0.073,0.045,1.049,-1.252,0.372,-0.343 | 0.056,0.072,-1.086,0.971,1.696,-0.922 | 0.041,0.021,-0.773,0.546,1.048,1.336 |
| **3924c**df59da9afffd2f82c75ca7553d1 | 0.051,0.050,1.757,-1.622,0.062,-1.364 | 0.085,0.049,1.110,-1.302,1.071,-1.482 | 0.076,0.086,-1.893,1.947,1.060,-0.818 | 0.088,0.019,-1.813,1.317,1.736,0.911 |
| **49adf**5ee46e40175f642c8ff80992f6c | 0.059,0.083,1.700,-1.403,-0.461,-1.165 | 0.035,0.045,1.435,-1.320,0.929,-0.801 | 0.082,0.054,-0.737,0.889,0.973,-1.471 | 0.074,0.037,-0.140,-0.452,1.439,1.612 |
| **5c87a**5bcfdc6ef74d61a1f7d22d216e8 | 0.093,0.072,1.918,-1.831,-1.883,-1.158 | 0.073,0.087,0.975,-0.664,1.533,-0.355 | 0.091,0.078,-0.951,1.949,1.149,-0.804 | 0.084,0.026,-0.917,0.561,1.097,1.785 |
| **6b434**3005acd1ffb7fe68072a2e42923 | 0.094,0.078,0.409,-0.150,-0.724,-1.085 | 0.034,0.070,1.429,-0.732,0.623,-1.110 | 0.045,0.062,-1.095,1.878,1.148,0.657 | 0.078,0.020,-1.869,1.532,1.071,1.017 |
| **7c810**8c97b85df2c87f10baf93d4301e | 0.087,0.095,0.271,0.050,-0.227,-1.615 | 0.045,0.083,0.764,-0.828,-0.672,1.469 | 0.066,0.074,-1.224,1.006,0.432,1.899 | 0.035,0.061,-1.179,0.862,0.910,0.609 |
| **7fbd6**d82a5994bec3fc5d7dbd1ee9ace | 0.085,0.082,1.111,-0.726,-0.490,-1.601 | 0.014,0.087,1.857,-1.298,-1.831,-1.785 | 0.080,0.098,-1.080,1.130,1.920,-0.762 | 0.081,0.053,-1.055,0.616,1.103,1.328 |
| **a00fe**eeb909e4320c023a537f7baf3c1 | 0.093,0.071,1.207,-1.298,0.398,-0.750 | 0.066,0.037,1.037,-1.205,1.589,0.408 | 0.096,0.062,-1.152,1.639,1.790,-1.666 | 0.013,0.078,-1.336,1.057,-1.173,1.557 |
| **ac424**9370b4dc433f2de466c51aec0f6 | 0.070,0.076,0.934,-0.650,-1.133,-1.064 | 0.065,0.066,1.953,-1.841,1.502,-0.400 | 0.081,0.050,-1.465,1.799,1.914,-1.123 | 0.020,0.045,-0.626,0.453,0.042,1.318 |
| **ec15f**9dc19d79f6dea4217b531fb16eb | 0.051,0.086,1.278,-0.949,-1.943,-1.165 | 0.022,0.023,1.500,-1.023,1.520,-0.866 | 0.080,0.081,-0.627,1.046,1.903,-1.751 | 0.074,0.037,-1.418,1.088,1.326,-0.446 |
| **0cffd**475a88b8677f47cafddaef7644f | 0.058,0.081,0.339,-0.247,0.941,-1.900 | 0.048,0.074,0.767,-0.608,-1.742,-0.518 | 0.077,0.088,-1.297,1.361,-0.480,0.563 | 0.076,0.080,-1.169,0.757,1.832,0.864 |
| **1d869**e6519477647c089b51c5f14da87 | 0.056,0.085,0.290,-0.239,0.976,-1.349 | 0.046,0.074,0.938,-1.323,-0.705,0.662 | 0.080,0.082,-1.503,1.095,-0.631,1.786 | 0.094,0.098,-1.283,0.950,1.642,0.313 |
| **39827**20dddf50e7cf60c636b10b80172 | 0.054,0.064,0.267,-0.164,0.393,-1.337 | 0.095,0.045,1.209,-0.810,-0.618,0.070 | 0.067,0.014,-1.413,1.888,-0.102,-1.444 | 0.072,0.095,-1.122,0.236,1.557,1.285 |
| **3b292**08fae3080dd268b58611abfff00 | 0.036,0.064,0.003,0.071,0.764,-1.708 | 0.034,0.095,0.707,-0.436,-0.282,-0.659 | 0.083,0.041,-1.610,1.952,-1.528,1.818 | 0.030,0.050,-1.116,0.770,1.876,1.267 |
| **3cb44**4efa6a61807ffc2633186b5ab18 | 0.088,0.095,0.332,-0.109,0.643,-1.926 | 0.012,0.083,1.009,-1.160,-0.335,0.177 | 0.062,0.030,-1.495,1.654,-0.640,1.929 | 0.080,0.098,-1.582,1.086,1.488,0.790 |
| **48d7e**61bab2ce51765c645e8273db59a | 0.059,0.092,0.417,-0.301,1.215,-1.712 | 0.081,0.085,0.597,-0.506,0.303,0.146 | 0.058,0.090,-1.151,1.634,-0.674,1.212 | 0.063,0.026,-0.772,0.439,0.798,0.978 |
| **52729**76c65e5dde478f61d47f902bdc2 | 0.042,0.099,0.412,-0.169,0.237,-1.219 | 0.061,0.039,0.402,-0.410,0.210,-0.105 | 0.058,0.065,-1.750,1.832,-1.270,1.324 | 0.059,0.056,-1.106,0.840,-0.238,1.410 |
| **5823d**0a425f1c7482eb89617d5c736d9 | 0.036,0.068,0.234,-0.116,1.168,-1.795 | 0.063,0.075,0.746,-0.788,-1.426,0.235 | 0.090,0.036,-1.924,1.836,0.386,0.792 | 0.061,0.068,-1.303,1.149,1.294,0.449 |
| **69a5b**bc44ee765e7212f3f431b3e59ec | 0.051,0.085,0.657,-0.504,1.223,-1.947 | 0.095,0.084,1.705,-1.440,-0.354,1.192 | 0.057,0.041,-1.304,1.843,-0.947,1.552 | 0.081,0.098,-1.702,1.114,1.779,0.467 |
| **6bbc6**0b0d5cda39ba906a65314102aa5 | 0.036,0.083,0.652,-0.453,1.166,-1.780 | 0.028,0.046,1.527,-1.082,0.311,-0.755 | 0.029,0.039,-1.370,1.646,-0.995,1.456 | 0.094,0.099,-1.548,1.025,0.676,0.318 |
| **7f793**ca55228ece75f7f39c2ddc7fa9b | 0.048,0.092,0.629,-0.412,1.151,-1.903 | 0.074,0.055,0.924,-0.775,-0.481,1.401 | 0.076,0.063,-0.878,1.538,-0.668,0.372 | 0.094,0.091,-1.438,1.056,-0.019,1.000 |
| **c4cba**9503e2245308b4384d709f84b19 | 0.015,0.090,0.177,0.008,0.821,-1.581 | 0.017,0.052,0.877,-0.618,-0.067,-0.182 | 0.027,0.068,-1.697,1.836,-0.728,1.917 | 0.063,0.088,-1.253,0.673,1.873,0.456 |
| **daeaa**4cc76b58bc993e33e2dfac3ba23 | 0.016,0.048,0.560,-0.389,0.178,-1.515 | 0.063,0.055,0.576,-0.211,-0.815,-0.040 | 0.098,0.062,-1.678,1.944,-0.407,1.344 | 0.078,0.021,-1.611,0.917,1.340,0.708 |
| **f14de**eee9176bb58637cbb51aaa89d36 | 0.076,0.092,0.339,-0.246,0.857,-1.539 | 0.016,0.084,1.729,-1.032,-0.589,0.770 | 0.096,0.068,-1.298,1.861,1.252,0.496 | 0.070,0.092,-1.406,0.971,-1.025,1.567 |
